# A surrogate workflow for virtual population development: designing an amenable proxy for a hypothesis-test inspired nonlinear goodness-of-fit

**DOI:** 10.1101/2025.04.26.649406

**Authors:** Lu Huang, Yinbo Chen, Satyendra Suryawanshi, Amir Molavi, Eric Sison, Brian J. Schmidt

**Affiliations:** Quantitative Pharmacology, Data, and Analytics, Bristol Myers Squibb, Princeton, NJ, USA; Clinical Pharmacology, Oncology, Bristol Myers Squibb, Princeton, NJ, USA; Quantitative Pharmacology, Data, and Analytics, Bristol Myers Squibb, Cambridge, MA, USA; Research Compute, Bristol Myers Squibb, Princeton, NJ, USA; Madrigal Pharmaceuticals, PA 19428, USA

**Keywords:** quantitative systems pharmacology, virtual population, prevalence weights, convex optimization, immuno-oncology

## Abstract

QSP models are increasingly being applied to drug development. Calibrating virtual populations (VPops) to clinical data helps explore patient variability, study populations of interest, and predict clinical outcomes for new therapies.

In previous work, we have developed the QSP Toolbox (1, 2), a library of tools and workflows for developing VPops. The aim is to create a VPop that fits clinical data from a variety of sources and with a variety of datatypes, allowing for differences in the quality of the data.

However, our VPop workflow has been hampered by the need to perform repeated optimization of “prevalence weights”. In this article, we explain how we sped up our VPop development workflow by using convex quadratic objective function as a sort of proxy for our highly nonlinear goodness-of-fit function.

We illustrate with two case studies the efficacy of our “surrogate” workflow for generating VPops with good *p*-fits. We find that the surrogate workflow can calibrate VPops much faster than the original workflow because it saves so much time on the prevalence-weighting optimizations.

**Graphic abstract:** 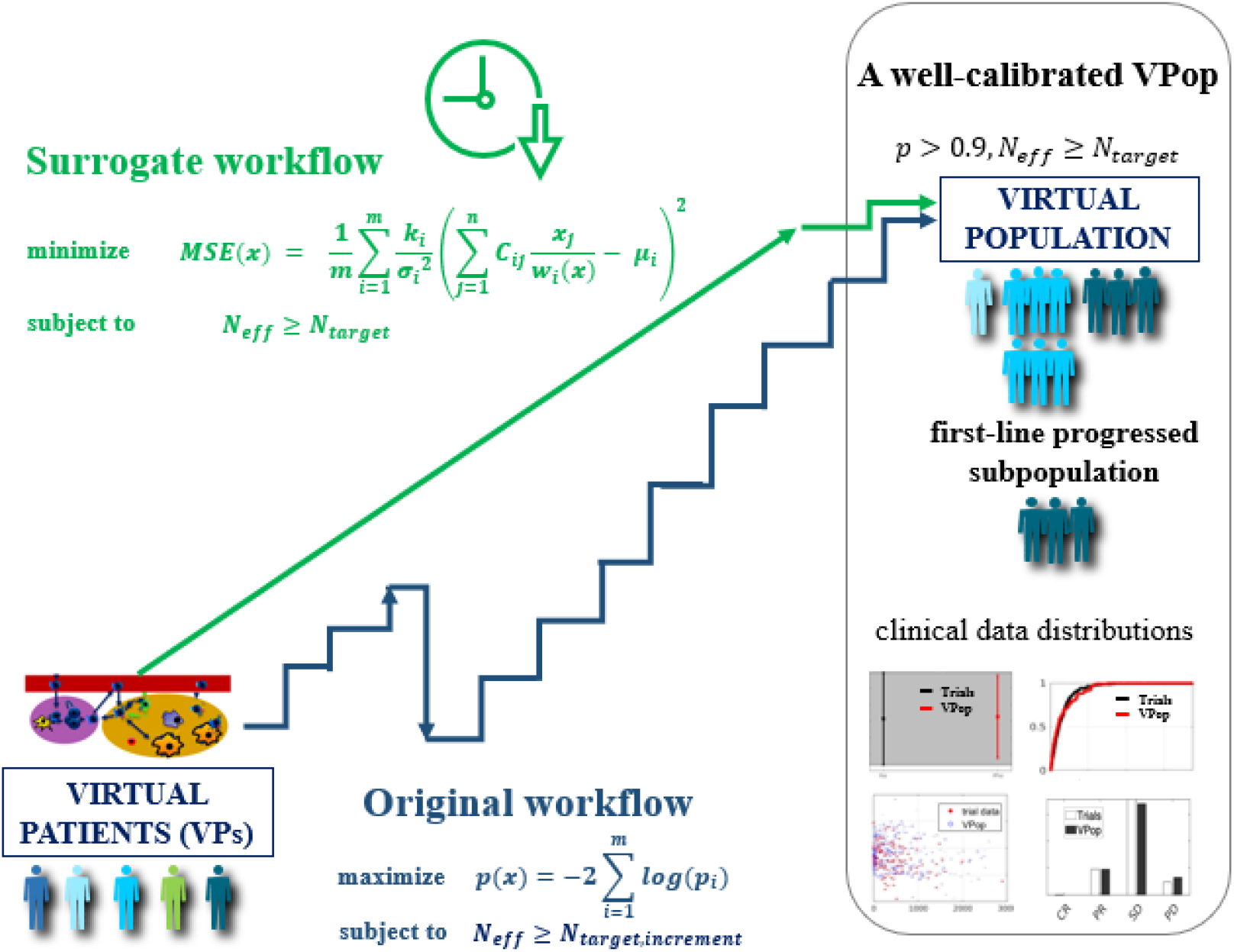

## I. Introduction

Quantitative Systems Pharmacology (QSP) modeling integrates biology and pathology to study system-level responses to drug treatment (3–9). A QSP model is typically a system of ordinary differential equations (ODEs). Each complete set of parameter values determines a unique dynamical trajectory, thereby defining a virtual patient (VP). Ensembles of VPs are used to explore clinical variability and uncertainty (10–14). We focus on VPs with physiologically plausible behavior, identified with the parameter space of the QSP model. Generally speaking, a virtual population (VPop) is any probability distribution on this parameter space, representing a clinical population with diverse characteristics.

As reviewed in (2), various methods for generating VPops have been utilized in literature, differing in goodness-of-fit (GOF) criteria, simulation strategies, sampling techniques, weighting and resampling approaches (13–23). These methods vary in suitability for different calibration data, data distribution assumptions, and computational demands. While a uniform VPop approach is challenging due to varying needs, continued innovation is anticipated to advance VPop applications for more complex cases.

In the theoretical framework implicit in our VPop generation workflow (1, 2, 14), a VPop is modeled as a weighted discrete distribution on parameter space. These weights, normalized, are called “prevalence weights”. The VPs themselves, as well as their prevalence weights, evolve during the VPop generation process. A “VP cohort” is essentially an unstructured finite set of plausible VPs; a prevalence-weighted VP cohort is a kind of VPop.

A “well-calibrated” VPop must fit the available clinical data, which is usually heterogeneous and includes various data types (e.g., different biomarkers or clinical outcomes) from distinct experiments (e.g., multiple clinical trials or interventions). For GOF we use a metric *p* inspired by Fisher’s combined probability test that can accommodate the number and variety of relevant clinical observations (1, 2, 14). The function *p* is defined in more detail in **Supplementary File 1**. *The p-value of Fisher’s combined test as GOF metric*; larger values of *p* connote better fits. Note that, while *p*(*x*) depends explicitly on the prevalence weights *x*, it also depends, implicitly, on the VPs in the VP cohort underlying the VPop.

Our VPop development workflow (1, 2, 14) iteratively adjusts the number and locations of VPs in the working VP cohort as it evolves towards one with higher GOF. This process resembles a random walk in population space, allowing the original VP cohort to evolve into a quite different one with completely different VPs. Each random step to the next VP cohort is guided by the working VPop, which is determined by assigning optimal prevalence weights to the working VP cohort. The optimal prevalence weights are *argmax*{*p*(*x*)}, the maximizer of *p*. The goal is to produce a VPop that has high *p* (e.g., *p* > 0.9) and moderately large “effective sample size”. To quantify effective sample size *N*_*eff*_ we have used the reciprocal of the Simpson index of diversity (24): the *N*_*eff*_ of a population of *n* VPs with normalized prevalence weights *x* = (*x*_1_, *x*_2_, …, *x*_*n*_)^*T*^ is given by the function

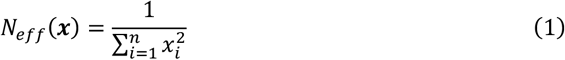

Prevalence-weighting a fixed VP cohort is done by maximizing *p* (or other GOF function) subject to the linear constraints that the weights are non-negative and sum to 1 (see **Methods**: *Optimization of prevalence weights*). Note that *N*_*eff*_(*x*) *< n* unless all prevalence weights equal 1/*n*, which is the equiprobable weighting of the cohort.

In the present work, we explore the use of a “proxy” GOF function of computationally amenable form to help expedite the search for VPops with high *p* and moderately large *N*_*eff*_. The heuristic idea is that a very good fit should have high GOF no matter what GOF function you use to measure it; and we venture that our alternative workflow, which runs along optimizing an amenable GOF function instead of *p*, will nonetheless lead to high-*p* VPops with large enough *N*_*eff*_.

Our proxy GOF function has the form exp(−*MSE*(*x*|*C*)), where

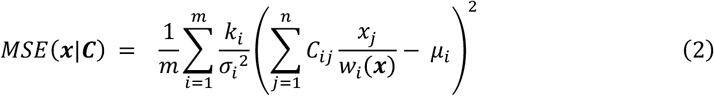

Note that maximizing the proxy GOF function is equivalent to minimizing *MSE*. We think in terms of minimizing *MSE* rather than maximizing exp(−*MSE*(*x*|*C*)).

In Formula (2), *m* denotes the number of biomarkers or other observables. The clinical data for biomarker *i* consists of *k*_*i*_ observations with sample mean *μ*_*i*_ and sample standard deviation *σ*_*i*_. There are *n* VPs in the cohort, and *C* is an *m × n* matrix. The entry *C*_*ij*_ is the simulation value of biomarker *i* for the *j*th VP unless that is undefined or not applicable, in which case *C*_*ij*_ = 0. The quantity *w*_*i*_(*x*) is the “subpopulation weight” of biomarker *i*, that is, the sum of the prevalence weights of the VPs subject to that biomarker. (See **Methods**: *The mean-squared-error* for more information.)

What makes *MSE*(*x*|*C*) technically amenable to our purposes is that it is, conditionally, a convex quadratic function of *x*. If we somehow knew the subpopulation weights *W*_*i*_ we could then minimize

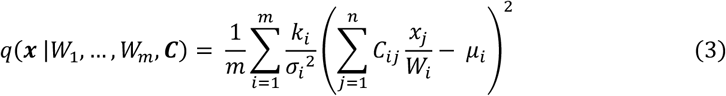

over prevalence weights that satisfy the *n* linear constraints that *w*_*i*_(*x*) = *W*_*i*_ in addition to *x ≥***0** and ∑ *x*_*i*_ = 1. We have named this function *q* as a reminder that it is quadratic; and we will write *q*(*x*) instead of *q*(*x* |*W*_1_, …, *W*_*m*_, *C*) when the conditioning values are not important.

Since *q*(*x*) and 1/*N*_*eff*_(*x*) are both convex functions of *x*, the minimizer of *MSE* subject to a lower bound constraint on *N*_*eff*_ along with the linear constraints *x ≥***0** and ∑ *x*_*i*_ = 1 is a minimizer of

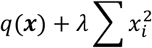

(for some *λ ≥*0) subject only to the linear constraints (25). Since *q*(*x*) and 1/*N*_*eff*_(*x*) are both *quadratic* convex functions of *x*, the problem has now been reduced to a linearly constrained least-squares problem. Such optimization problems are exceptionally tame computationally and can be solved very quickly (26). (See **Methods**: *Optimization of prevalence weights*.)

When we do this in practice (that is, when we find prevalence weights all throughout VPop development workflow by minimizing *q* + *λ*/*N*_*eff*_ subject to linear constraints instead of maximizing *p* subject to a constraint on *N*_*eff*_) we often end up with a VPop that appears, qualitatively, to fit the data distributions fairly well. When the prevalence weights underlying the VPop are adjusted to maximize *p*, the resulting VPop often has high enough *p* and usually appears to fit the data distribution even better. This way of producing a high-*p* VPop by driving down *MSE* is our “surrogate workflow”.

Our surrogate workflow largely follows the logic of the original workflow; it is just the iterated step at the heart of both algorithms that differs. The original workflow increments *N*_*eff*_ while maintaining high *p*; whereas the surrogate workflow maintains the desired target *N*_*eff*_ as it tries to decrease *MSE*. (See **Methods**: *The surrogate workflow*.)

A considerable amount of time can be saved on optimization using the surrogate workflow. Previously, a typical VPop workflow, fitting to 150 observables, would require hundreds of iterations of sampling and weighting, and could take days to run even with hundreds of parallel processors. The surrogate workflow cuts that time in half, or better. For example, in *Case Study 2* on VPop calibration with a large QSP model for non-small cell lung cancer (NSCLC), it reduced the time by 80-90%!

Unfortunately, the surrogate workflow does not *always* succeed in practice (though, for that matter, neither does the original workflow). We have had to put considerable thought into the design of *MSE* to make *e*^−*MSE*^ a good proxy for *p*, and have had to meet a couple of technical challenges:

The form (2) of *MSE* makes sense when all observables are simple quantitative measurements. However, as *p* involves observables of other types (e.g., multivariate, categorical, distributional) and we would like *MSE* to emulate *p*, we must attempt to encode the p-values for a single statistical comparison into a number of rows of the matrix *C*. We discuss this challenge and our *ad hoc* techniques for handling it in **Supplementary File 1**: *Constructing matrices for non-scalar data-type comparisons*.

Another challenge is what we call the “subpopulation problem”, by which we refer to certain complications imposed upon our surrogate workflow due to the unfortunate fact that *MSE*(*x*|*C*) of Formula (2) is not convex in *x*. We define a “subpopulation” as a subset of VPs sharing a common characteristic, such as nonresponse to therapy or mechanistic dropouts in clinical trial simulations due to progressive disease. For example, in immune-oncology therapies, a subpopulation could be anti-Programmed Death 1 (anti-PD1) progressed patients undergoing second-line treatment; or patients excluding those who have dropped out at a given scan time, based on Response Evaluation Criteria in Solid Tumors (RECIST) criteria (27).

The formula for *p* (see **Supplementary File 1**: *The p-value of Fisher’s combined test as GOF metric*) supports integration of subpopulation data straightforwardly; but the way such data is accommodated in Formula (2) for *MSE* unfortunately complicates our surrogate workflow. To deal with this difficulty, we devise ways to condition on the subpopulation weights so that we only ever have to minimize functions of *x* like *q*(*x* |*W*_1_, …, *W*_*m*_, *C*) of Formula (3). See **Methods**: *The mean-squared-error* and **Supplementary File 1**: *The subpopulation problem in the surrogate workflow* for details.

## II. Methods

### Optimization of prevalence weights

Suppose one has a VP cohort with good coverage. How can one assign prevalence weights to the VPs to create a VPop whose statistical predictions agree well with the observed clinical distributions? One way to accomplish this is to posit a scalar GOF function *f* and, fixing the VPs in the cohort, to search for an assignment of weights that maximizes *f*. That is, one seeks prevalence weights that

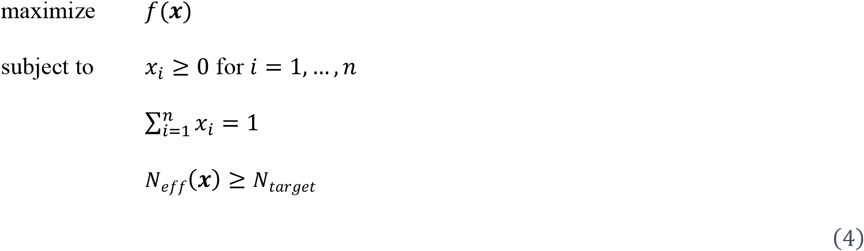

Here, *x* = (*x*_1_, *x*_2_, …, *x*_*n*_)^*T*^ is the vector of prevalence weights, *n* denotes the number of VPs; and 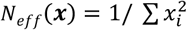, the effective sample size defined in Formula (1), is constrained to be larger than *N*_*target*_.

We refer to the optimization (4) of the prevalence weights of VPs in a fixed cohort as “prevalence-weighting”. If we change the GOF function from *f* to *g* and recompute prevalence weights with respect to *g*, we refer to this operation as “rebalancing” the prevalence weights.

These optimization (4) would be much easier to perform if *f* were given by (the negative of) a sum-of-squares, rather than *f* = *p* (25, 28, 29). If the GOF function had the form −*q* or *e*^−*q*^, where *q* is a convex quadratic function, then maximizing GOF subject to a hard lower-bound constraint on *N*_*eff*_(*x*) = 1/ ∑ *x*^2^ would be equivalent to solving the *L*^2^-penalized minimization problem

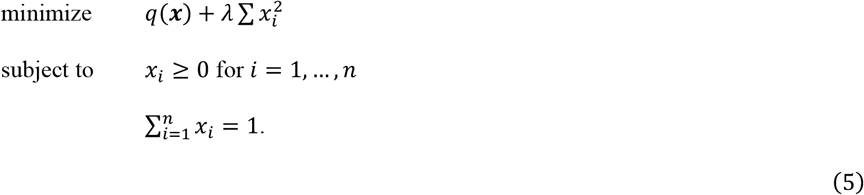

with suitably chosen penalization parameter *λ ≥*0. With its convex quadratic objective function and linear constraints, Problem (5) can be solved much, much faster than Problem (4). This technical advantage is at the heart of the algorithm for our surrogate workflow.

### The mean-squared-error

Building on earlier work (28, 29), we have further developed the *MSE* metric to more closely emulate p-value GOF components, incorporates subpopulation calibrations including mechanistic dropouts, as well as multivariate considerations (e.g. correlations and joint distributions).

Let *x* = (*x*_1_, *x*_2_, …, *x*_*n*_)^*T*^ be a (column) vector of prevalence weights for *n* VPs, defining a VPop on a fixed VP cohort. Suppose there are *m* generalized biomarker measurements on the VPs, or some subset of the VPs. The *i*’th entry of the vector *Cx*, where *C* is the *m × n* matrix *C* defined just below, will represent the average value of the *i*’th generalized biomarker as predicted by the VPop with prevalence weights *x*.

Let

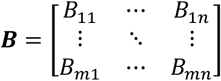

denote the *m × n* matrix of simulated values: *B*_*ij*_ is the simulated value of biomarker *i* on the *j*’th VP. Missing values (NaNs) in *B* correspond to missing measurements (e.g. mechanistic dropouts due to progressive disease) or VPs missing from a subpopulation (e.g., second-line treated patient subsets). Let *C* denote the matrix obtained from *B* by replacing all NaNs with 0 and let *K* be the *m × n* “mask” matrix whose entry *K*_*ij*_ = 0 if the corresponding entry of *B* is NaN and *K*_*ij*_ = 1 if it is not.

For convenience of this narrative, we first consider the ideal situation with no subpopulations and where all biomarkers are simple quantitative measurements. The clinical data for biomarker *i* is a sample of *k*_*i*_ observations with sample mean *μ*_*i*_ and sample standard deviation *σ*_*i*_. In this context, we may think of prevalence-weighting as a weighted least-squares problem: we solve for parameters *x* = (*x*_1_, *x*_2_, …, *x*_*n*_)^*T*^ subject to linear non-negativity and sum-to-one constraints, that minimize the weighted sum-of-squared-errors:

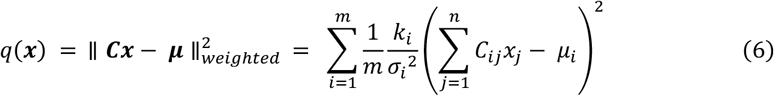

In this formulation, *C* is the model matrix of simulated values, while ***μ*** = (*μ*_1_, *μ*_2_, …, *μ*_*m*_)^*T*^ represents the data.

We weight the squared residuals in these formulas by 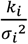 so that each term in the sum has variance approximately equal to 1, under the assumption that the observations are independently sampled from a population with mean 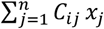 as predicted by the prevalence weighting *x*. For clinical data of the simplest form, i.e., large sample averages of (scalar) biomarker measurements, this scaling makes *q* compatible with the corresponding set of t-statistics that contribute to *p* (see **Supplementary File 1**: *The p-value of Fisher’s combined test as GOF metric*). Furthermore, in certain cases where we only had to fit VPop predictions to mean values, weighting the squared residuals in this way appears to have improved the performance of the proxy function.

Formula (2) for *MSE* is a straightforward generalization of Formula (6) that also accommodates subpopulation data. Let *S*_*i*_ denote the set of VPs who subject to measurement of the generalized biomarker corresponding to row *i* of ***C*** (though the data value may be NaN). If ***x*** is a system of normalized prevalence weights, let

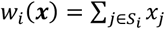

denote the total prevalence weight of all the VPs in the set *S*_*i*_, which we refer to as the “subpopulation weight” corresponding to row *i* of ***C***.

Formula (2) says that

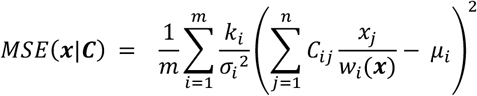

The expression 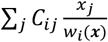 in the preceding formula indeed represents the renormalized subpopulation average, thanks to the fact that *C*_*ij*_ = 0 if *j ∉S*_*i*_. Note that

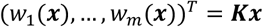

where ***K*** denotes the *m × n* mask matrix defined above.

The problem is that *MSE*(*x*|*C*) is no longer a simple quadratic function in ***x*** for fixed ***C*** because of its nonlinear dependence on the linear forms *w*_*i*_. Nonetheless, *MSE*(***x***|***C***) retains its simple quadratic structure when restricted to any non-empty domain of feasible solutions ***x*** that satisfy linear constraints of the form (*w*_1_(***x***), …, *w*_*m*_(***x***)) = ***W***, where ***W*** is a fixed vector of subpopulation weights. This means that the problem of minimizing *MSE*(*x*|*C*) subject to linear constraints ***Kx*** = ***W*** reduces to the problem of minimizing the convex quadratic form *q*(***x*** |***W, C***) given by Formula (3) in the Introduction.

We discuss this further in **Supplementary File 1**: *The subpopulation problem in the surrogate workflow*.

### The surrogate workflow

**Figure 1** provides an overview of the VPop development workflow in QSP Toolbox. Our surrogate VPop calibration workflow modifies the original workflow only in the iterated step of the VPop development. The first stage of the surrogate workflow is the same. The goal of this stage is to produce a VP cohort with good coverage. A large number of initial VPs are generated by dispersed sampling of parameter values from physiologically reasonable ranges. Each VP is then simulated, and those with implausible trajectories are screened out.

**Figure 1.**
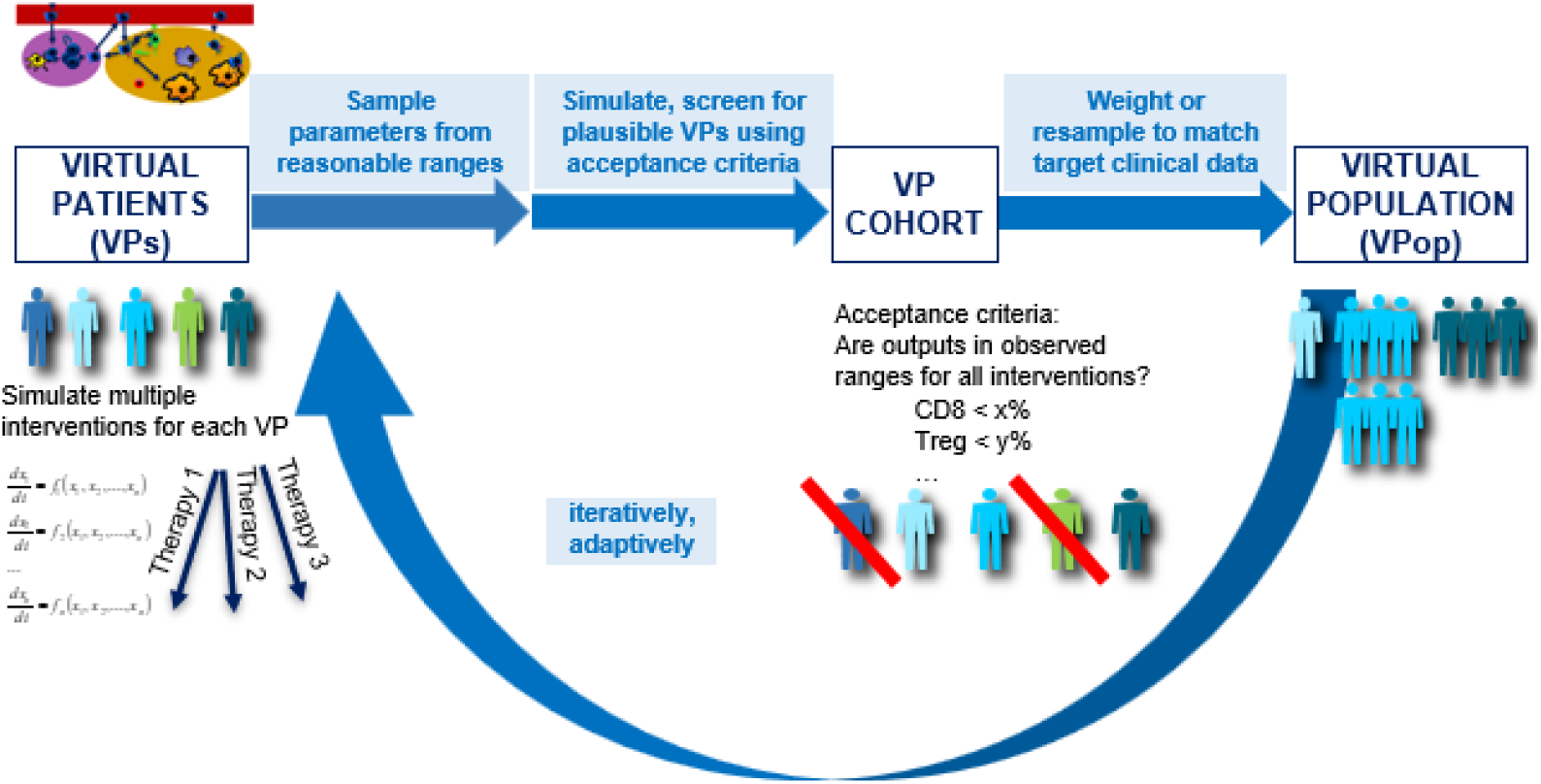
High-level overview of the VPop development workflow in QSP Toolbox. The workflow of developing a VPop from a QSP model in QSP Toolbox involves two stages: First, we generate a VP cohort by sampling parameter values from physiologically reasonable ranges and screening them for plausibility. In the second stage, we iteratively resample and reweight VPs to produce a well-calibrated VPop, measured by a high *p* GOF and a large *N*_*eff*_.

The second stage of the *original* workflow takes this screened VP cohort as its input and subjects it to random perturbations in an iterative process that involves further sampling and screening to generate new plausible VPs. The process is designed to cause the working VP cohort to evolve in a manner that maintains *p* above a threshold (we want *p ≥*0.9) while gradually increasing *N*_*eff*_. When the working VP cohort has issues finding a good fit with *p ≥*0.9 after increasing *N*_*eff*_, the algorithm will reduce the number of VPs through clustering and restart from a low *N*_*eff*_ again (2). The goal is to develop a VPop with *p* as large as possible and with large enough *N*_*eff*_ (e.g., *N*_*eff*_ > 150). Such a VPop is the output of the workflow.

Our *surrogate* workflow uses the same QSP Toolbox functions for the sampling and screening, which still requires time to simulate and screen new VPs for plausibility. However, it saves time on optimizing prevalence weights by solving linearly constrained least-squares problems, saving almost all the prevalence-weighting effort demanded of the original workflow.

We first describe an *idealized* surrogate workflow for VPop development; the looser variant thereof that we implement in practice is discussed at the end of this section. The idealized surrogate workflow is designed so that the basic step, operating on a VPop with effective sample size *N*_*eff*_, transforms it into a different VPop with the same *N*_*eff*_ but smaller or equal *MSE*.

Furthermore, the idealized surrogate workflow described here is limited to cases without “subpopulation problems”. Assume for now that every VP is subject to every generalized biomarker. That way, given simulated data *C*, we only have to minimize the simple sum-of-squares *q*(*x*) of Formula (6). We explain how we handle subpopulations in **Supplementary File 1**: *The subpopulation problem in the surrogate workflow*.

To initialize the VPop development stage of the idealized surrogate workflow, set the “working VP cohort” to be the one produced by the first stage of the workflow, as described above. Assign prevalence weights to the cohort by minimizing *MSE* subject to a constraint on *N*_*eff*_, e.g., that *N*_*eff*_ *≥*150. This can be accomplished by a bisection search for the value of *λ* such that *N*_*eff*_(*x*_*\**_(*λ*)) = 150, where *x*_*\**_(*λ*) denotes the constrained minimizer of *MSE* + *λ*/*N*_*eff*_. This requires constrained minimization of *L*^2^-penalized *MSE* for several different values of *λ* (and it is this extra work that we neglect to do in practice).

Now, repeat the following basic step:

> [Input a VPop]
>
> First, **delete** from the VP cohort underlying this VPop all those VPs whose prevalence weights equal zero (there can be many of these).
>
> Second, **add** new plausible VPs to the VP cohort by pseudo-random sampling around the current VPs. The sampling rate near a VP is an increasing function of its prevalence weight (e.g. proportional). Screen the new VPs and keep only the plausible ones. Ideally, the newly sampled VPs just replace the lately discarded VPs; at worst, the cohort size increases by a limited amount.
>
> Third, **prevalence-weight** the working VP cohort by minimizing *MSE* subject to the constraint on *N*_*eff*_ along with the non-negativity and sum-to-one constraints.
>
> [Output this VPop]

As an operation on VPops, the basic step keeps *N*_*eff*_ constant while *MSE* decreases (or stays the same). If we only ever eliminate VPs with prevalence weights equal to zero, then the *MSE* of the VPop can only decrease (or stay the same) throughout VPop development.

The basic step is iterated until certain (admittedly arbitrary) end-criteria are met, or until the working cohort becomes too large to work with. For example, we may decide to stop after 100 iterations of the basic step, or when working cohort gets too big, whichever comes first. Then we rebalance the final working VPop according to *p*, and that VPop is the output of our surrogate workflow. With any luck it will prove to have *p* close enough to 1, and the surrogate workflow will have succeeded.

If the VP cohort gets too big, but doesn’t achieve high enough *p*, then regular operation of the surrogate workflow ceases and we intervene to reduce the size of the VP cohort through automated clustering (30). After clustering, we may continue by rebooting the surrogate workflow with the clustered VPop as initial VPop, or we may revert with it to the original workflow. For example, one could use the final VPop of the surrogate workflow as a seed for the particle swarm optimization (PSO) algorithm we use to maximize *p*. The surrogate workflow, if it fails to calibrate a VPop, can serve at least as a sort of “burn-in” phase to this end.

The case studies presented below did not use the idealized algorithm described above. We cut a couple of corners to save time. In our QSP Toolbox, the surrogate workflow does not bother to find the value of the penalty hyperparameter *λ* such that *N*_*eff*_(*x*_*\**_(*λ*)) = *N*_*tarwwet*_ (= 150, for example). Instead, *λ* is adjusted adaptively: if the *N*_*eff*_ is currently less than *N*_*tarwwet*_ we try a larger *λ* in the next round, and if it is greater than *N*_*tarwwet*_ we use a smaller *λ*. We cut a corner, too, when we delete some VPs with tiny but non-zero prevalence weights. The effect of these shortcuts is that the *MSE* of the developing VPop can bounce up and down, though the eventual trend, visible in **Figure 5**, is for *MSE* to decrease overall once *N*_*eff*_ has more or less stabilized.

### Algorithm

The algorithm runs on a cluster using MATLAB Parallel Server (MPS) on MATLAB R2022a. The QSP Toolbox is available open-source at https://github.com/BMSQSP/QSPToolbox.

The algorithm is included as a ‘linearCalibration’ module, and the surrogate VPop development workflow is wrapped in the ‘expandVPopEffN’ function within the QSP Toolbox. To run the surrogate workflow, expandVPopEffNOptions.linearExpandFlag needs to be set true. This setup was used for both case studies below. For *Case Study 1*, all the model and data are public, and the code to reproduce the work is provided in **Supplementary File 3**.

## III. Results

We incorporated the surrogate workflow into the automated VPop development process in the QSP Toolbox. Here we report two case studies using this surrogate workflow to calibrate VPops for two QSP models. The first is a published Antibody Drug Conjugate (ADC) QSP model from the QSP Toolbox examples. The second is a large-scale in-house immuno-oncology (I-O) QSP model for NSCLC.

The first case study demonstrates that the surrogate workflow is able to generate high-quality VPops comparable to those produced by the original workflow. The second case study highlights the speed-up afforded by the surrogate workflow, which is especially advantageous for large-scale models.

### Case Study 1: ADC QSP model VPop calibration

We first applied this workflow to a previously published ADC QSP model (1) to demonstrate its effectiveness, referring to the VPs as virtual xenografts (VXs). This case study is presented as updated example 06 in the QSP Toolbox. It shows how to set up the surrogate workflow to develop VPops from an existing VX cohort. The model, data and MATLAB script for this case study are included in **Supplementary File 3**, easily modifiable for application to other models.

The surrogate workflow successfully calibrated a VPop that matches the summary statistics (28 means, 28 standard deviations, and 28 bin frequencies) across several experimental datasets. Both were run on 20 parallel workers. The surrogate workflow required the same number of iterations as the original workflow and generated a VPop with *p* = 0.95 and *N*_*eff*_ = 40. However, since the convex optimization is much faster than nonlinear particle swarm optimization (PSO), the entire VPop development process is significantly faster. The 20 iterations of surrogate workflow followed by one round of rebalancing the weights to maximize *p* took 77 minutes, as compared to 182 minutes.

This well-calibrated VPop presents visually similar fits to those produced using the original workflow. Biomarker dynamics (**Figure 2a-l**) and parameter distributions (**Figure 2m**) are well captured by this VPop. Parallel results from original workflow shown in **Supplementary Figure 2**.

**Figure 2.**
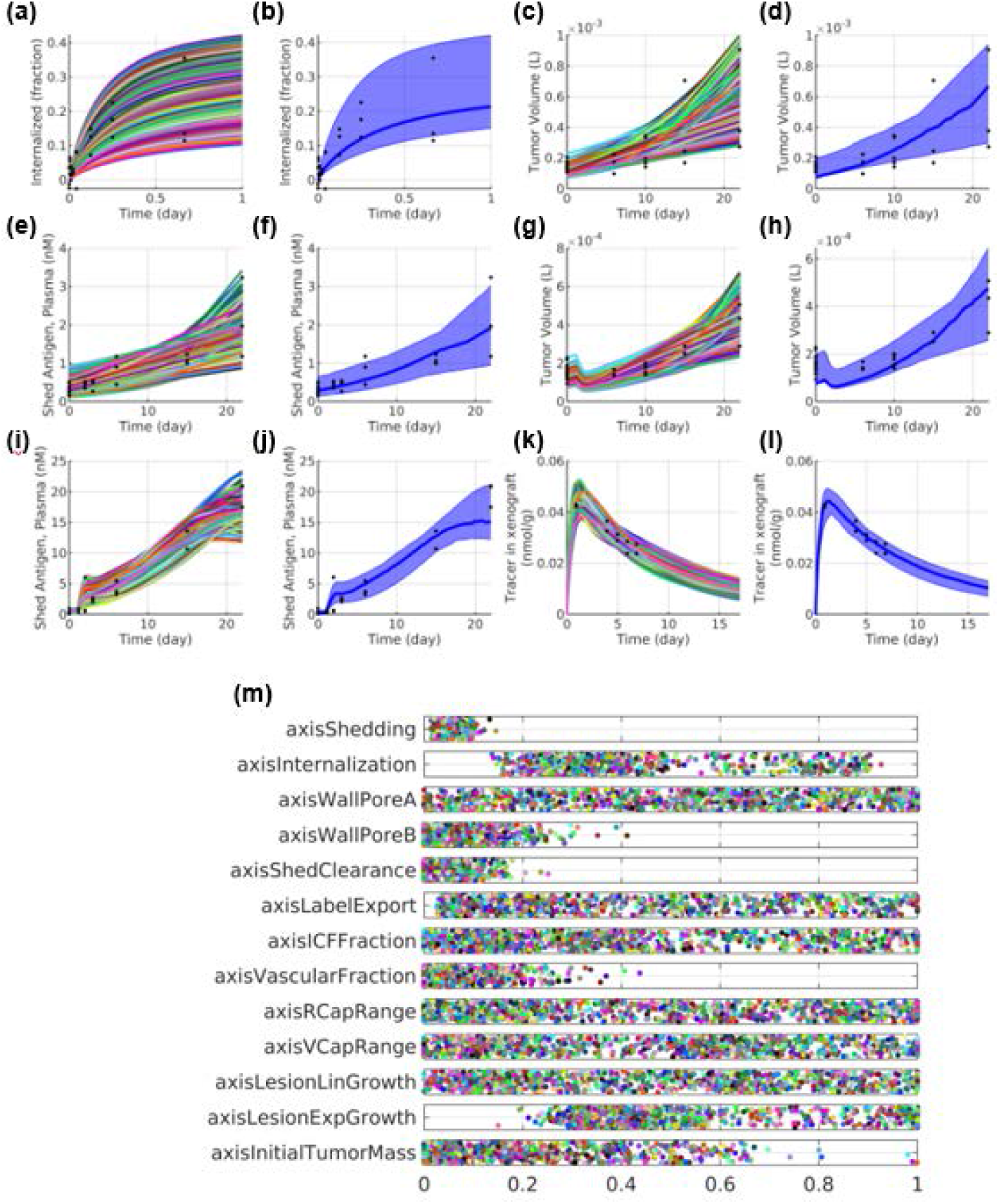
Demonstration of ADC model VPop generated from the surrogate workflow. Using the initial VX cohort from the example 06 in QSP Toolbox, we developed a VPop using our surrogate workflow. The plots show simulated outcomes for each VX in the cohort for different conditions: **(a)** cell culture; **(c)** and **(e)** buffer injection; **(g)** and **(i)** naked antibody injection; (**k)** 89Zr-labeled antibody injection. Plots **(b)**, (**d)**, (**f)**, (**h)**, (**j)**, and (**l)** display the 5th, 50th, and 95th percentiles of the prevalence-weighted VPop. Plot (**m)** shows scaled mechanistic axis coefficients of 1000 VXs, where each axis corresponds to a model parameter, each dot represent a VX, and each parameter is scaled within its axis range.

### Case Study 2: I-O QSP NSCLC VPop calibration

We applied the surrogate workflow to a large-scale in-house I-O QSP platform model for NSCLC to demonstrate its improvement in efficiency and scalability. This model was chosen for its sizable scope and complexity in modeling patient variability and disease progression across multiple clinical trials and data types. Application of our workflow to the NSCLC model demonstrates its effectiveness in generating high-quality VPops with significantly reduced computational time compared to the original workflow.

The I-O QSP NSCLC platform model is a therapeutic response model that includes key pathways like cytotoxic T-lymphocyte antigen 4 (CTLA-4), PD-1, a novel immune checkpoint, and innate immune pathways. It captures major immune cell life cycles, cytokine-mediated feedbacks, and important clinical biomarkers such as lesion responses and immune cell infiltrates (**Figure 3**). Developed in SimBiology, the model consists of 375 ODEs, 1037 reactions, 1031 rules, and over 550 references. The model structure and parameters were determined using separate pathway-model fits for model components like cell-cell contact, confinement, two-dimensional molecular interactions, immune cell life cycle, recruitment, cytokine-mediated feedback loops, and cancer killing. It has been developed in a stepwise manner, adding new pathways and calibrating with new clinical data to support clinical development.

**Figure 3.**
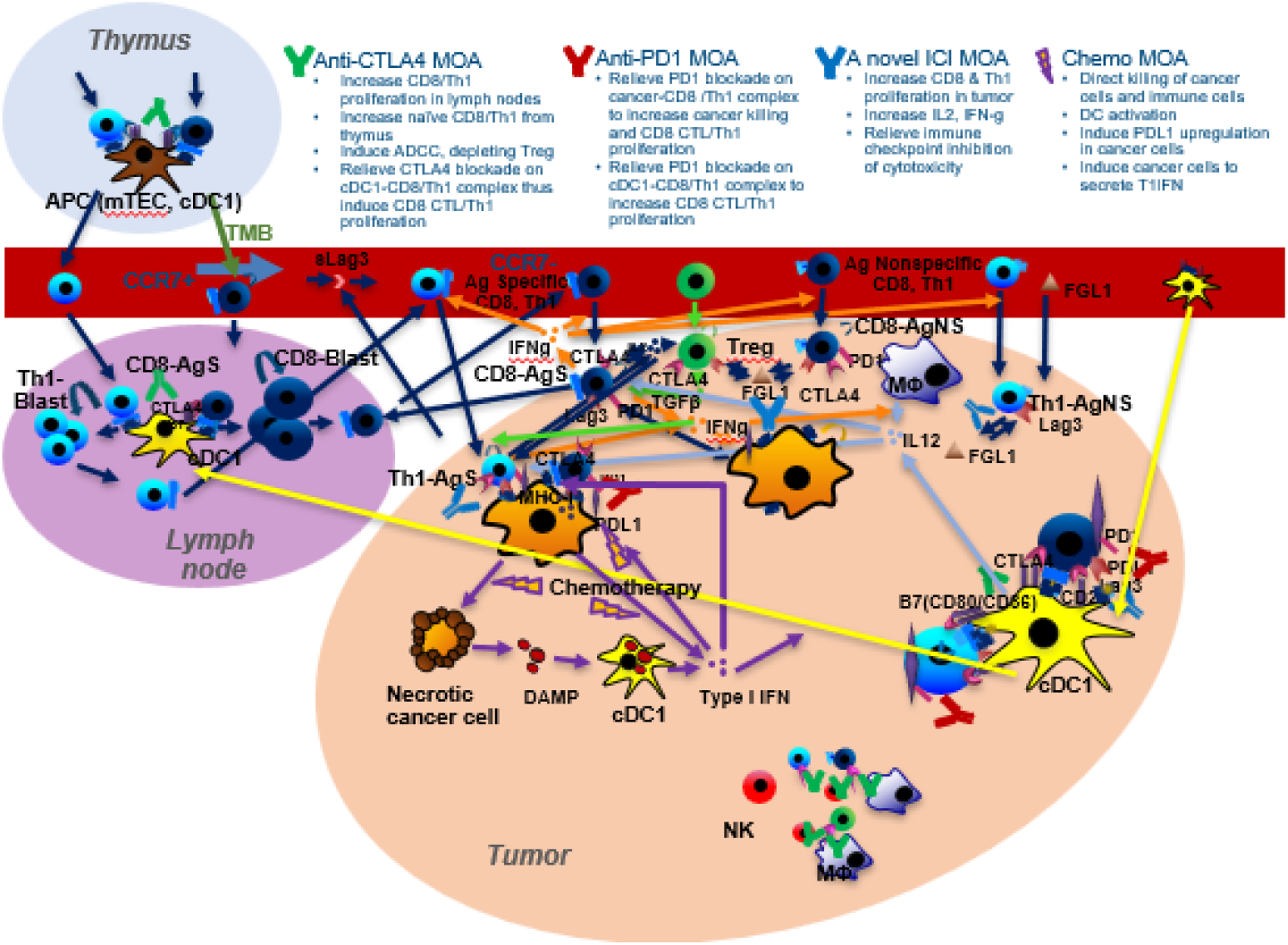
Overview of the in-house I-O QSP platform model.

In this case study, using the surrogate workflow, we developed a VPop for non-squamous NSCLC that captures the population variability of clinical biomarkers, including immune cell infiltrates, PDL1, absolute lymphocyte count (ALC) and tumor mutational burden (TMB), as well as lesion response at multiple time points across multiple therapies and lines of intervention. The fit was performed across 11 therapies, including combinations of chemotherapy, anti-PD1, anti-CTLA-4, and a novel immune checkpoint inhibitor (ICI), and involved PDL1 stratified subpopulations. The GOF functions (both *p* and *MSE*) include comparisons of VPop predictions with the clinical data for 130 means, 130 variances, 121 distributions, 212 binned distributions, 15 two-dimensional distributions, and 20 correlations. For the evaluation of *p*, these comparisons were effected, respectively, by the t-test, F-test, Kolmogorov-Smirnov test, Fisher’s exact test, Peacock test, and modification of Fisher’s r-to-z transformation (31, 32).

We incorporated the surrogate VPop development algorithm into the QSP Toolbox. VPop development began with a first stage of computationally demanding initial dispersed sampling, here involving over one million simulations, and screening for plausible VPs; while the computational demands of the second stage of iterative simulation and optimization are made much more practical with this surrogate VPop development workflow (**Figure 4**). As we decreased *MSE* while maintaining *N*_*eff*_ around *N*_*tarwwet*_ during the VPop development process, *p* generally increased and eventually achieved a target value higher than 0.9 after one final round of rebalancing of prevalence weights according to *p* (**Figure 5**). At the end of the evolution phase of 100 iterations of the surrogate VPop development algorithm, the final VPop achieved *p* = 0.9995 with *N*_*eff*_ = 150 in a single “rebalancing” run.

**Figure 4.**
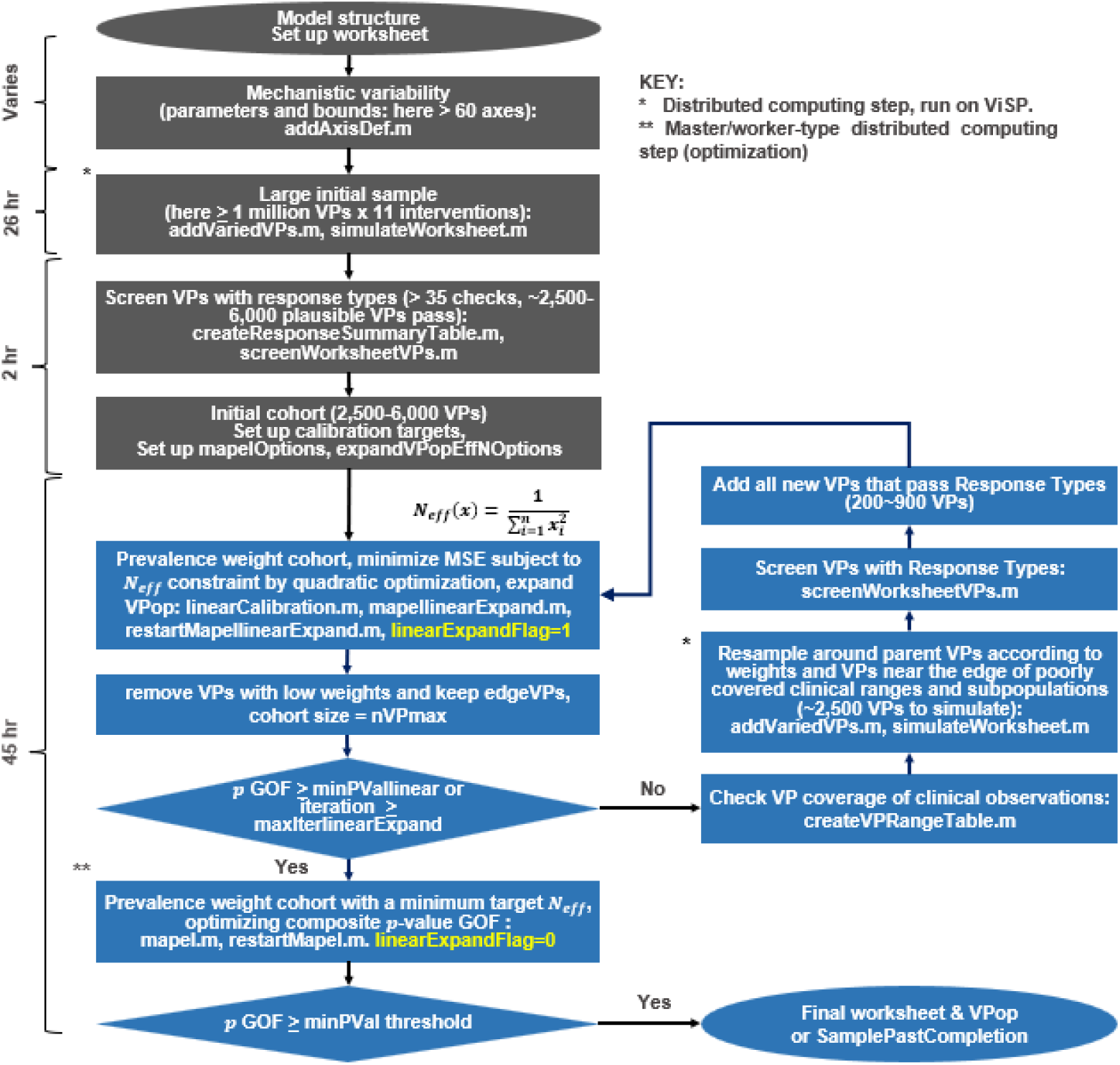
The algorithmic surrogate workflow of I-O QSP NSCLC VPop development. The first stage of the surrogate workflow is the same as the original: generating a VP cohort by sampling and screening for plausibility. The second stage differs, with highlighted in blue, involving the iterative steps for the VPop development. The diagram also lists the QSP Toolbox functions used for each step. The runtime and VP numbers shown are specific to the I-O QSP NSCLC VPop development in *Case Study 2*. The distributed computing step runs on ViSP (33).

**Figure 5.**
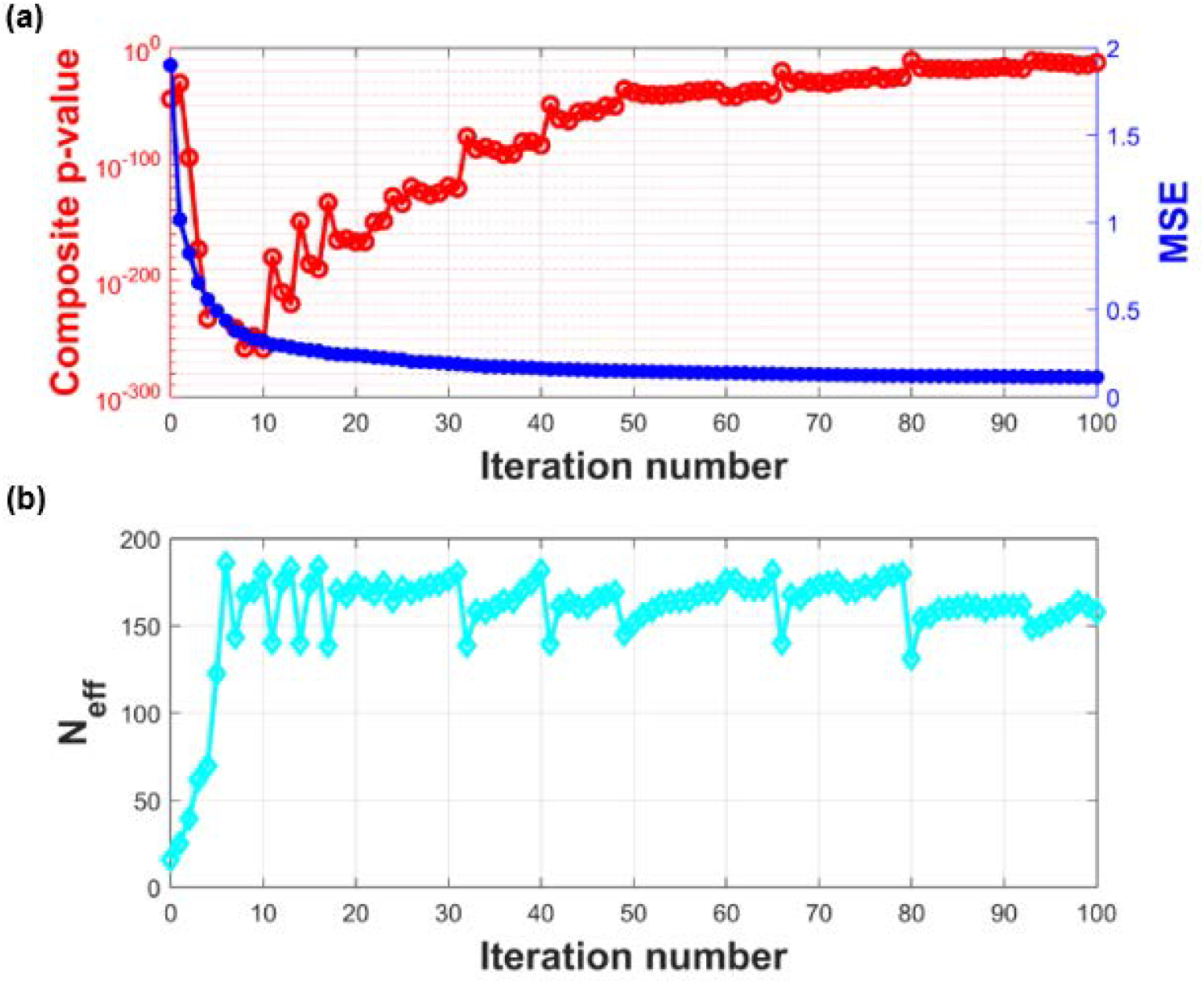
The algorithm performance of I-O QSP NSCLC VPop development. As we decrease *MSE* while maintaining the effective sample size *N*_*eff*_ around a target value of 150 during VPop development, the GOF *p* tended to increase and eventually achieves a high *p* > 0.9 after “rebalancing”. The plots show **(a)** the two GOFs and **(b)** the effective sample size throughout the VPop development iterations.

The VPop developed from the surrogate workflow implies predicted biomarker distributions that agree well with their corresponding clinical distributions (**Figures 6–8** and **Supplementary Figures 3–4**).

**Figure 6.**
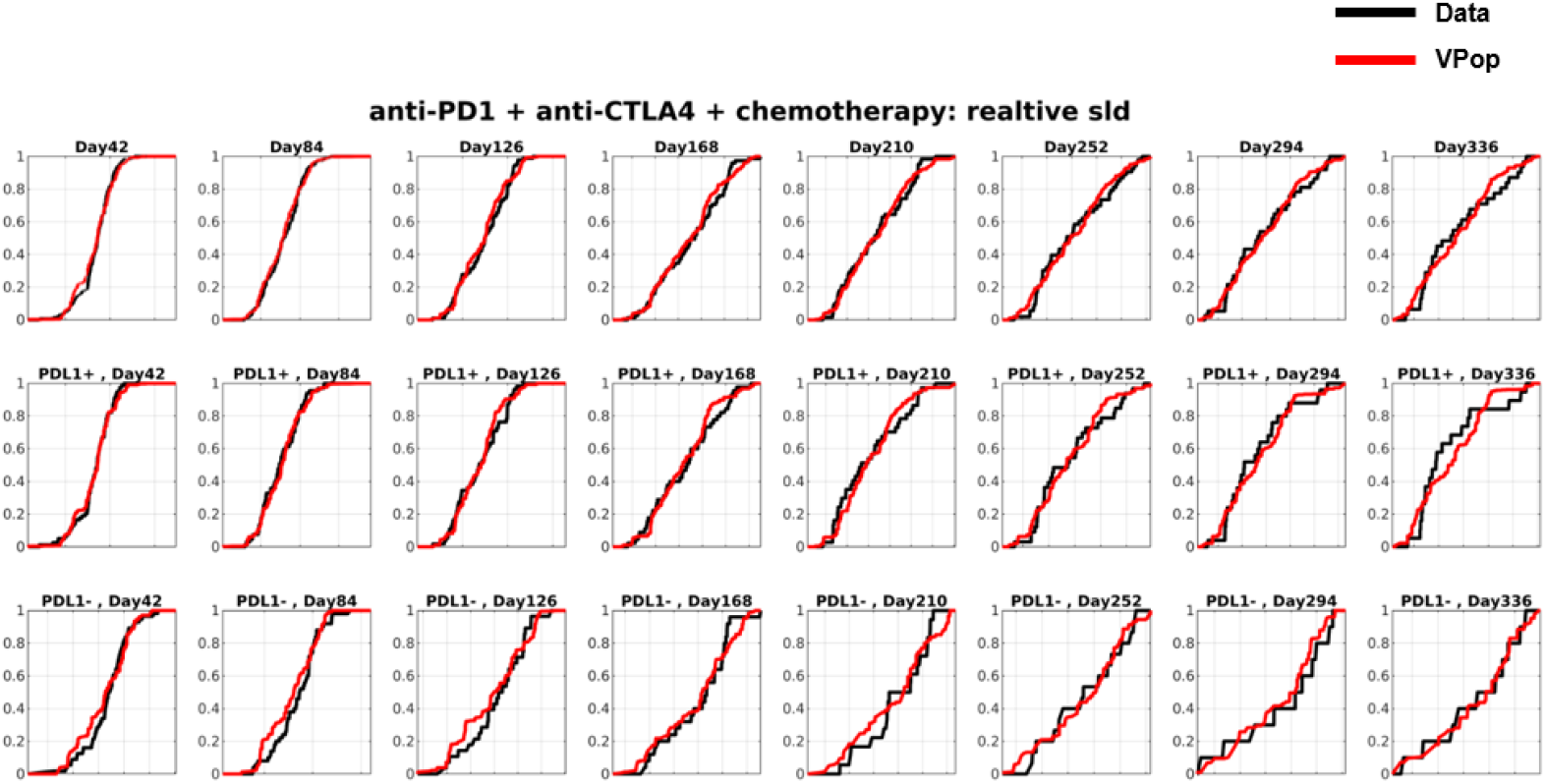
The I-O QSP NSCLC VPop simultaneously captures 1D clinical distributions. Shown are a subset of the clinical distributions - lesion size change in response to a first-line combination of anti-PD1, anti-CTLA4 and chemotherapy, stratified by PDL1 expression levels. The black curve represents the cumulative distribution function (cdf) extracted from internal clinical data, while the red curve shows the cdf of the calibrated VPop. The VPop closely matches the observed marginal distributions, with no assumptions needed for the distributions due to the empirical comparison strategy.

**Figure 7.**
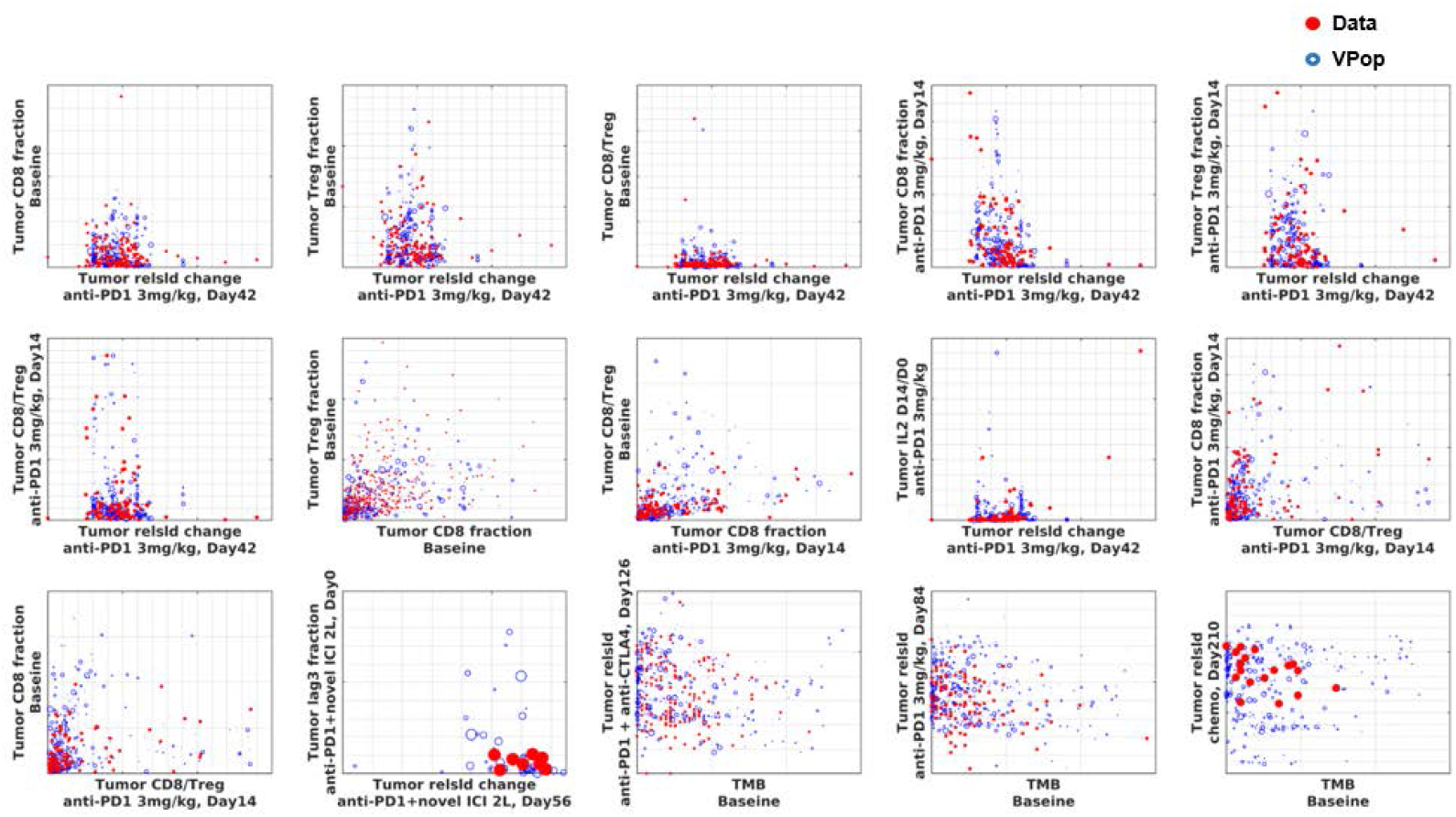
The I-O QSP NSCLC VPop simultaneously captures 2D (joint) clinical distributions. Each plot shows the joint distribution between clinical biomarkers or endpoints. Red dots represent data extracted from internal clinical trials, and blue circles represent VPs, with larger circles indicating higher prevalence. The plots show good agreement between the VPop and clinical data, with no assumptions needed for the joint distribution due to the empirical comparison strategy.

**Figure 8.**
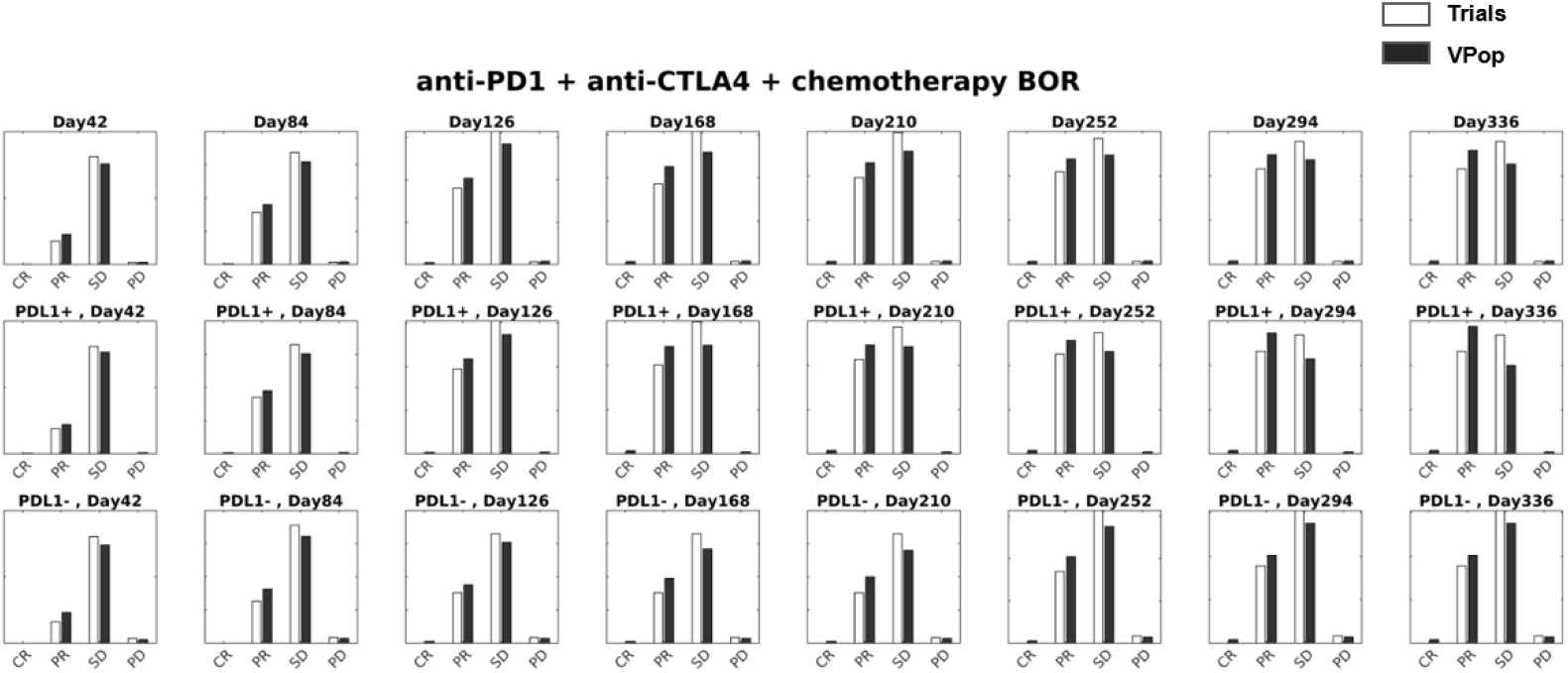
The I-O QSP NSCLC VPop captures best index lesion responses. The I-O QSP NSCLC VPop captures best index lesion response at multiple time points and for multiple interventions. Shown here are a subset of the response rates - best overall response (BOR) to first-line combination of anti-PD1, anti-CTLA4 and chemotherapy, stratified by PDL1 expression levels. The white bars represent the clinical data and the black bars represent the VPop response.

The NSCLC VPop development took approximately 45 hours on 1,500 workers (with MPS) using the surrogate workflow; while using the original workflow, it would take over 116 iterations and might take more than 300 hours to complete. In fact, we stopped the original workflow after the first clustering and never completed it due to lack of computational resources, so its runtime is only a lower bound estimate. The surrogate workflow has already shown a runtime reduction of over 85% compared to the original workflow.

To assess model performance and predictability, we withheld data on combination therapy of chemotherapy and anti-PD1 in PDL1-negative patients. We simulated patient recruitment differences by resampling VPs with replacement according to prevalence weight. The observed data fell within the modeled 95% confidence intervals for index lesion response at multiple time points (**Figure 9**). In-house clinical trial data were used to calculate the index lesion RECIST classification for the combination therapy.

**Figure 9.**
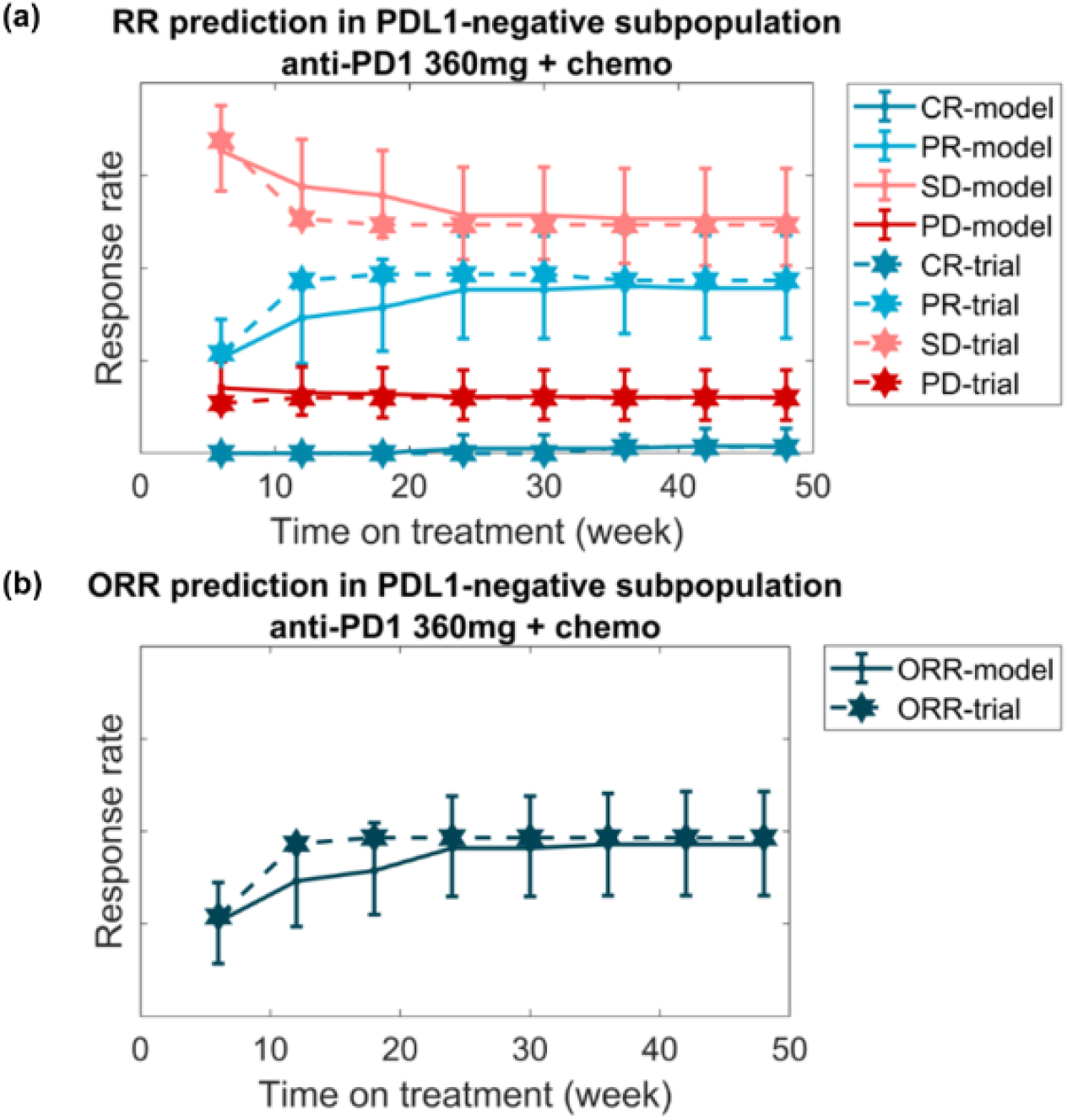
Validation of I-O QSP NSCLC VPop on independent dataset. The calibrated VPop accurately predicted the withheld data. Prediction of **(a)** response rates and **(b)** overall response rates to the combination of anti-PD1 and chemotherapy in the PDL1-negative subpopulation match the hold-out trial data. Solid lines show the predicted response rates from the VPop, while dashed lines represent the observed trial response rates. Error bars indicate the 95th percentile range from 200 alternate in silico trials, each with 68 VPs, as in the clinical trial. Response rates are based on RECIST criteria from index lesions, scaled to account for unequivocal progression.

## IV. Discussion

Our surrogate workflow reduces computational burden and accelerates VPop generation, which is crucial for timely decision-making in the pharmaceutical industry. Quickly generating accurate and robust VPops enhances the applicability of QSP models.

However, the simulation time for large QSP models with low plausibility rates remains a problem. Our surrogate workflow still requires time to simulate new VPs for plausibility screening. AI/ML models have been explored to accelerate plausibility detection (21, 22), but accurately representing the complex dynamical model and identifying small plausible regions remain challenging. Integrating our surrogate workflow with model reduction techniques and AI/ML algorithms could further enhance VPop development efficiency.

As an emerging and innovative tool, there are still many open questions about VPop development. One key question is how to select a minimal set of variable parameters to define a VP and avoid overfitting. Our surrogate workflow offers an efficient method for generating new VPops, accelerating the exploration of parameter selection strategies.

It is also important to recognize that multiple VPops may fit observed data equally well but yield different predictions. To better assess prediction uncertainty, one strategy involves identifying other VPops that fit the data adequately (14). While exhaustively exploring alternative VPop space requires further research and is beyond the scope of this paper, our surrogate workflow could potentially advance uncertainty quantification and improve QSP model credibility by reducing the time needed to generate alternative VPops.

## V. Conclusion

We have discussed a modification of our VPop calibration workflow that significantly speeds up the VPop development process while still achieving high-quality fits. This “surrogate workflow” uses a sum-of-squares objective function along with *L*^2^-regularization to maintain well-distributed prevalence weights. Like *p*, the GOF function *MSE* of the surrogate workflow is also designed to handle multiple data types and subpopulations, albeit less elegantly. Our heuristic approach has proven robust across several case studies, including multiple in-house programs.

Our surrogate workflow provides a scalable and widely applicable method to develop VPops more efficiently, which could ultimately enhance the predictive power and utility of QSP models in clinical settings.

## Acknowledgments

We sincerely thank our former colleague Abed E. Alnaif for his initial work on linear calibration and bagging exploration while at Bristol Myers Squibb.

We would like to thank our QPDA colleague Huy Vo for technical support; our former colleague Ronny Straube and current QPDA colleagues Laura Woo, Yili Qian, Kiyoto Tanemura, Alexander Ratushny and Pegy Foteinou for their insightful discussions and feedback as we develop and apply VPop methods; and our IT colleagues Amir Ghasemi, Deepika Battula, Peter Foster, and former BMS colleagues Sudhindra Rao and Kai Zhao for their IT support. We gratefully acknowledge the suggestions from Ricardo Paxson and Arthur Goldsipe of MathWorks® for improving aspects of the QSP Toolbox.

Additionally, we greatly appreciate Gaohua Lu and Anna Kondic for providing feedback on the manuscript.

## Author contributions

LH: Conceptualization, Algorithm design and implementation, Model development, Data analysis, Simulations, Writing - Original draft, Visualization

YC: Model development, Writing - Review and Editing

SS: Data acquisition and visualization, Writing - Review and Editing

AM: Writing - Review and Editing

ES: Cloud computing support, Writing - Review and Editing

BJS: Conceptualization, Algorithm design, Model development, Data analysis

## Supplementary Materials

**Supplementary File 1**. Additional methodological context

### The p-value of Fisher’s combined test as GOF metric

The choice of *p* as GOF metric offers a direct quantitative method to combine different data type comparisons while accounting for sample size, and allows for straightforward integration of data comparisons from disjoint datasets.

The logic of Fisher’s combined test is as follows: Suppose we know the p-values (*p*_*i*_) of *m* hypothesis tests and we wish to test the combined null hypothesis that all *m* of those null hypotheses are true. Provided that the p-values (which may be thought of as random variables) are statistically independent, the statistic 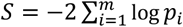 has a chi-squared distribution with 2*m* degrees of freedom and can serve as the test statistic for the combined null hypothesis. The p-value *p*_*S*_ associated with this test statistic is the probability that a 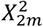 random variable is greater than *S*.

To build our GOF functions *p*, we combine the p-values from separate comparisons for each biomarker measurement, clinical endpoint, or other observable type using the formulas for Fisher’s combined probability test, defining *p* = *p*_*S*_ as above, so that 0 ≤ *p* ≤ 1, with *p* = 1 being a perfect fit.

Though *p* is a p-value, we do not use it as such. For hypothesis tests, it is *small* values of the p-value that are of significance; but we want *p* to measure goodness-of-fit, as a tool of meta-analysis, and hope to attain fairly *large* values of it (e.g., *p* ≥ 0.9). We stress that the GOF function *p* is not to be interpreted as a p-value in the statistical sense. This is just as well, because many of the observables that we combine are not statistically independent, anyway: for example, when patients in a contingent “subpopulation” of a larger population (e.g., those who undergo second-line treatment) are subject to further measurements. On account of these dependencies, *p* is an inaccurate p-value (it won’t have a uniform distribution, as a p-value is supposed to). But *p* needs not be a reliable p-value for our purposes: we just want it to serve as a sensible GOF function.

### Constructing matrices for non-scalar data-type comparisons

As described in (1, 2, 14), frequentist statistical methods such as t-tests, F-tests, contingency tables, K-S tests, Fisher’s r-to-z transformation, and multidimensional K-S tests such as the Peacock test and related comparisons (31, 32) were used in our QSP Toolbox for comparisons between VPop simulations and clinical data. We computed the p-value from Fisher’s combined test as our GOF function *p*, as explained in **Supplementary File 1**: *The p-value of Fisher’s combined test as GOF metric*.

The *MSE* measure of GOF is designed to capture the components of Fisher’s combined p-value GOF (*p*), including multivariate considerations such as correlations and joint distributions. This is essential for capturing the complex relationships between different clinical variables and ensuring that the VPops accurately represent the observed clinical distributions. We have found that our surrogate workflow achieves better fits to the observed clinical data when we make the terms of *MSE* more like the test statistics in the respective hypothesis-test argument.

The methods we used to construct matrices for different data type comparisons are illustrated in **Supplementary Figure 1**.

First, for fitting to the variance of various biomarkers, we reformulated the variance formula, and instead fit the second moment of the mean (**Supplementary Figure 1a**). By fitting both the first and second moments of the mean simultaneously, the variance is effectively captured as well. For fitting to the correlation, we proceeded similarly and fit the mean of the product of the two relevant biomarkers (**Supplementary Figure 1b**).

Second, for fitting to binned data, marginal, and joint distribution data, we convert them all into categorical binned data distributions (**Supplementary Figure 1c-e**): The marginal cumulative distribution function (cdf) extracted from data is divided into 20 distinct bins along the probability function (**Supplementary Figure 1d)**. Similarly, the joint cdf of two biomarkers is divided into a 5×5 grid (**Supplementary Figure 1e**). Each bin was represented by a different row in the matrices. In the *C* matrix, a VP is assigned 1 if it belongs in the corresponding bin, and 0 otherwise. In the *μ* vector, the experimental data point is the observed frequency of that bin.

Each data type is further normalized by the number of rows in that observation to maintain balanced weighting among data types. Additionally, we implemented the flexibility to independently weight each observation (matrix row), allowing users to assign different weights based on various factors, such as data quality considerations.

### Reformulation of the mean-squared-error

We note here an interesting formal property of *q*(***x***|*W*_1_, …, *W*_*m*_, ***C***).

Recalling the definitions of the data-cleaned model matrix ***C*** and the mask matrix ***K*** from **Methods:** *The mean-squared-error*, we can easily verify that ***C*** = ***C*** *∘* ***K***, i.e., *C*_*ij*_ = (***C*** *∘* ***K***)_*ij*_ = *C*_*ij*_*K*_*ij*_

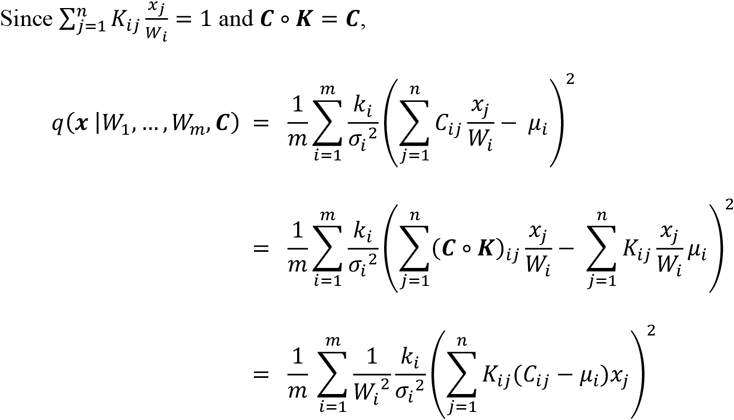

If we let ***D*** denote the *m* × *n* matrix with entries *D*_*ij*_ = *K*_*ij*_(*C*_*ij*_ *− μ*_*i*_), we can write

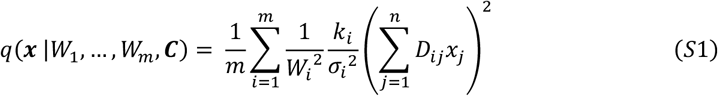

In this formula, the weight on the *i*’th squared residual is proportional to 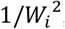,where *W*_*i*_ is the *i*’th subpopulation weight. For this insight alone, Formula (S1) deserves to be a numbered equation in the present document!

We have effectively used Formula (S1) in our code, working with entities such as ***D***, but we have not yet used the essential insight it confers about the relationship between the subpopulation weights *W*_*i*_ and the factors 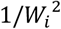 in the weighted sum-of-squares.

### The subpopulation problem in the surrogate workflow

If there is no real subpopulation data at all, i.e., if every *W*_*i*_ = 1, then there is no subpopulation problem, and prevalence-weighting a fixed cohort is a matter of minimizing the simple sum-of-squares in Formula (6). If all subpopulation weights *W*_*i*_ were known *a priori*, then we could use the convex quadratic function *q*(*x* |*W*_1_, …, *W*_*m*_, *C*) of Formula (3), minimizing it over the polytope of prevalence weightings that satisfy the linear constraints

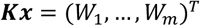

along with the usual constraints that ***x* ≥ 0** and that **∑** *x*_*ij*_ = **1**.

Now, if there were only one or two proper subpopulations, i.e., if only one or two *W*_*i*_ are strictly less than 1, one could try to minimize *MSE*(*x*|*C*) by minimizing, over all feasible choices of the subpopulation weights *W*_*i*_, the constrained minima over *x* of *q*(*x* |*W*_1_, …, *W*_*m*_, *C*). However, as we are dealing with tens or hundreds of proper subpopulation observables, this is not a practical approach for our purposes, and instead we do the following.

When there is subpopulation data, the surrogate workflow is basically the same, except for a tweak to the basic step. Instead of *q*(***x***) as in Formula (6), we are going to minimize a function of the form *q*(***x*** |*W*_1_, …, *W*_*m*_, ***C***) as in Formula (3), subject to constraints on the subpopulation weights *w*_*i*_(*x*) = *W*_*i*_. A natural suggestion is constrain the subpopulation weights *W*_1_, …, *W*_*m*_ of the current VPop to equal those of the input VPop, and to minimize *q*(***x*** |*W*_1_, …, *W*_*m*_, ***C***) subject to the linear constrains *w*_*i*_(*x*) = *W*_*i*_, *i* = 1,2, …, *m*, together with other constraints that ***x*** ≥ **0**, ∑ *x*_*i*_ = 1, and *N*_*eff*_(*x*) ≥ *N*_*target*_ (the latter constraint can be enforced by *L*^2^-penalization). If we do this, the output VPop cannot have greater *MSE* than the input VPop. This is because the input VPop corresponds to a specific system of prevalence weights the VP cohort of the output VPop: the prevalence weights of VPs that were in the input VPop are unchanged, and *new* VPs all have weight 0, and the VPs that were removed from the VP cohort of the input VPop all had weight 0 to begin with. As the minimum over a larger set of possible VPops, the *MSE* of the output VPop can be no larger than that of the input VPop.

The only problem now is that, though the basic step is designed to make *MSE* smaller, it keeps the subpopulation weights exactly the same. Our solution is to propose alternative estimates 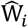 of the subpopulation weights and, in parallel with the first proposal, also minimize 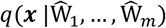 subject to the constraints 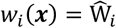 in addition to the other constraints. One way to obtain estimates of the *W*_*i*_ is to use the full-population data, i.e., biomarkers for which *W*_*i*_ = 1, of which we have plenty, in practice. We provisionally optimize prevalence weights 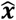 using the full-population data and estimate the subpopulation weights as 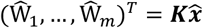.We end up two alternative prevalence weightings of the current VP cohort, and we choose the one that has lower *MSE* as the output VPop.

**Supplementary File 2**. Supplementary figures

**Supplementary Figure 1.**
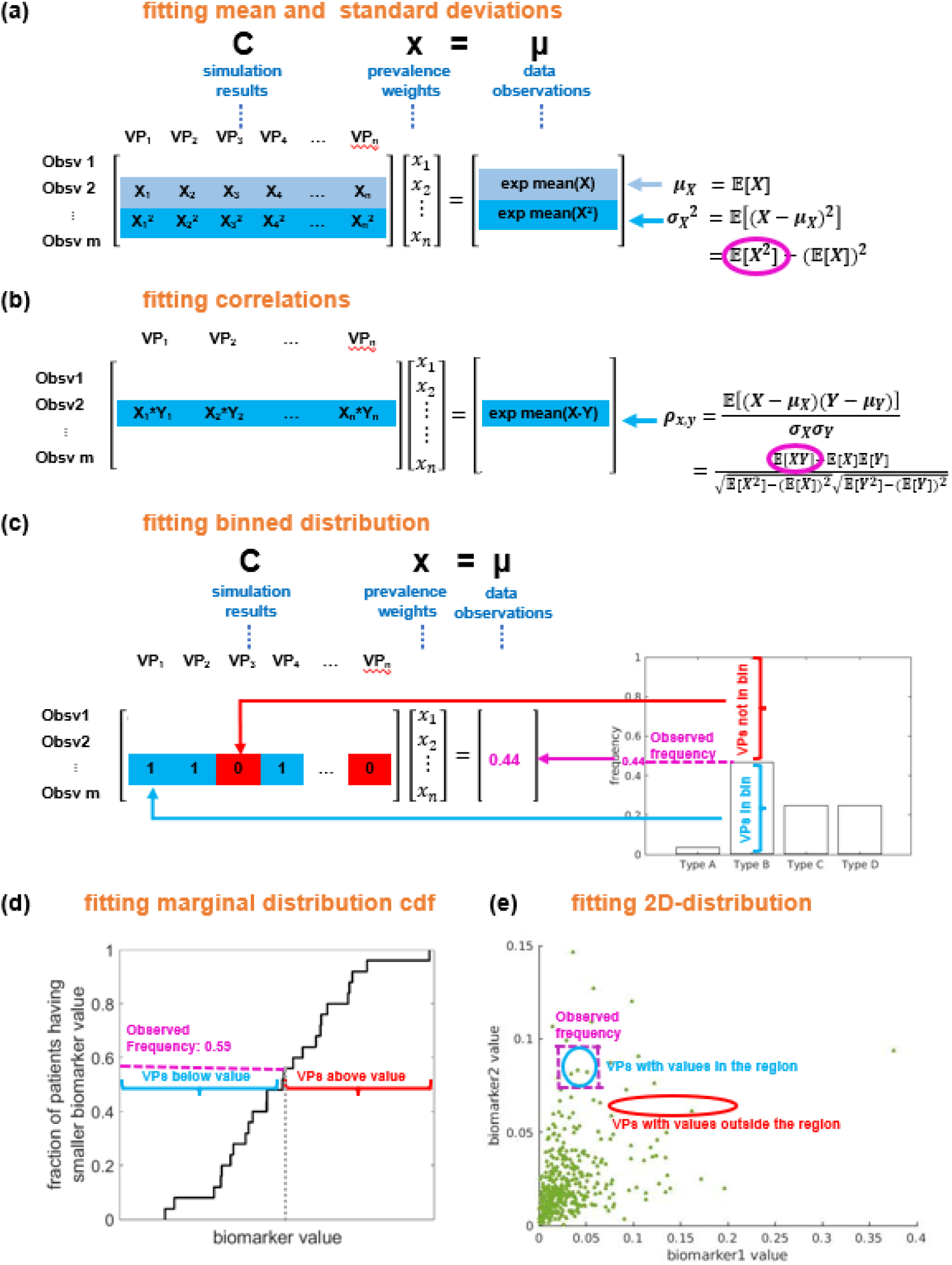
Linear matrices construction for different data-type comparisons. **(a)** Variance fitting: we fit the second moment of the mean by reformulating the variance formula. **(b)** Correlation fitting: we fit the mean of the product of the two biomarkers by reformulating the correlation formula. **(c-e)** Binned, marginal, and joint distribution data: we converted them into categorical binned data distributions. Each bin is a row in the matrices, with the *C* matrix assigning 1 if a VP is in the bin, and 0 otherwise. The *μ* vector shows the observed frequency of each bin.

**Supplementary Figure 2.**
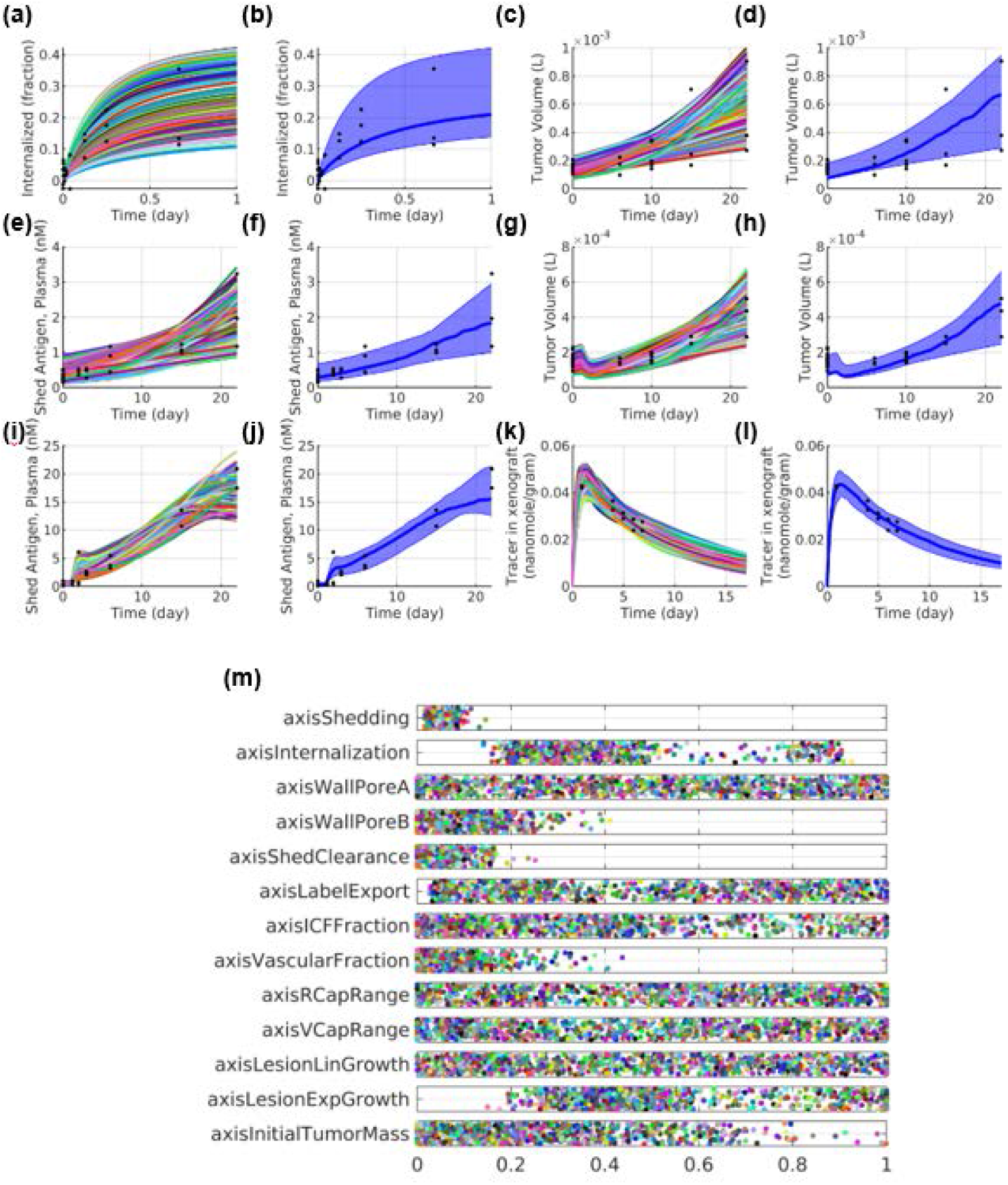
Demonstration of ADC model VPop generated from the original workflow. Shown is the comparison to experimental data from VPop developed using the original workflow after 20 iterations. This VPop also has *p* = 0.95 and *N*_*eff*_ = 40. The plots show simulated outcomes for each VX in the cohort for different conditions: **(a)** cell culture; **(c)** and **(e)** buffer injection; **(g)** and **(i)** naked antibody injection; (**k)** 89Zr-labeled antibody injection. Plots **(b)**, (**d)**, (**f)**, (**h)**, (**j)**, and (**l)** display the 5th, 50th, and 95th percentiles of the prevalence-weighted VPop. Plot (**m)** shows scaled mechanistic axis coefficients of 1000 VXs, where each axis corresponds to a model parameter, each dot represent a VX, and each parameter is scaled within its axis range.

**Supplementary Figure 3.**
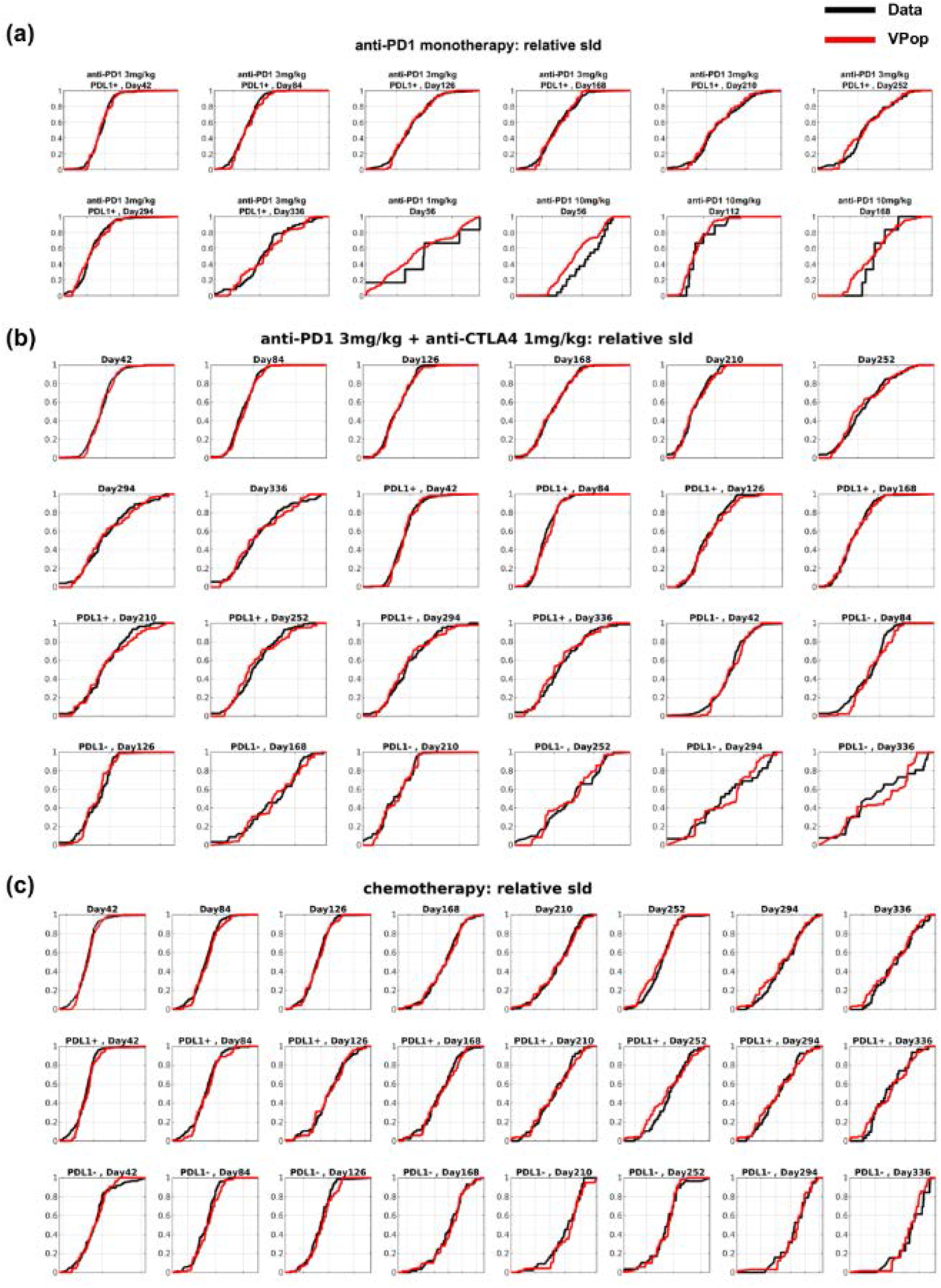
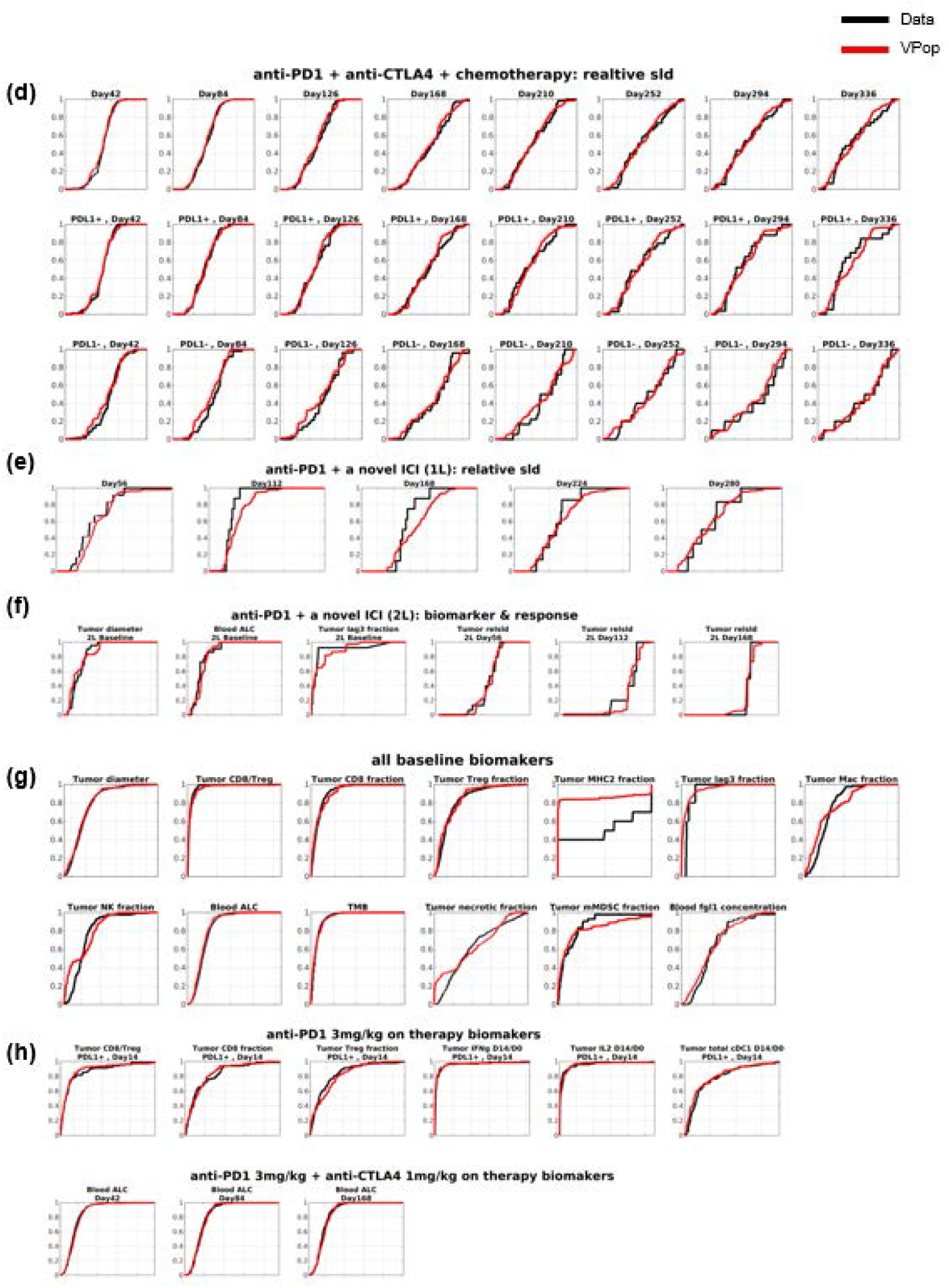
The I-O QSP NSCLC VPop simultaneously captures 1D clinical distributions across multiple trials. Shown are all the clinical distributions used in the calibration of I-O QSP NSCLC VPop. The VPop captures lesion size changes for: **(a)** anti-PD1 monotherapy, **(b)** anti-PD1 and anti-CTLA4 combination therapy, **(c)** chemotherapy; **(d)** combination therapy of anti-PD1, anti-CTLA4 and chemotherapy, **(e)** first-line combination therapy of a novel ICI and anti-PD1, and **(f)** second-line combination therapy of a novel ICI and anti-PD1. Some clinical trials are stratified by PDL1 expression levels. Additionally, the VPop captures clinical distributions of biomarker at **(g)** baseline and **(h)** on therapy across several trials. The black curve represents the cumulative distribution function (cdf) extracted from internal clinical data, while the red curve shows the cdf of the calibrated VPop. The VPop closely matches the observed marginal distributions, with no assumptions needed for the distributions due to the empirical comparison strategy.

**Supplementary Figure 4.**
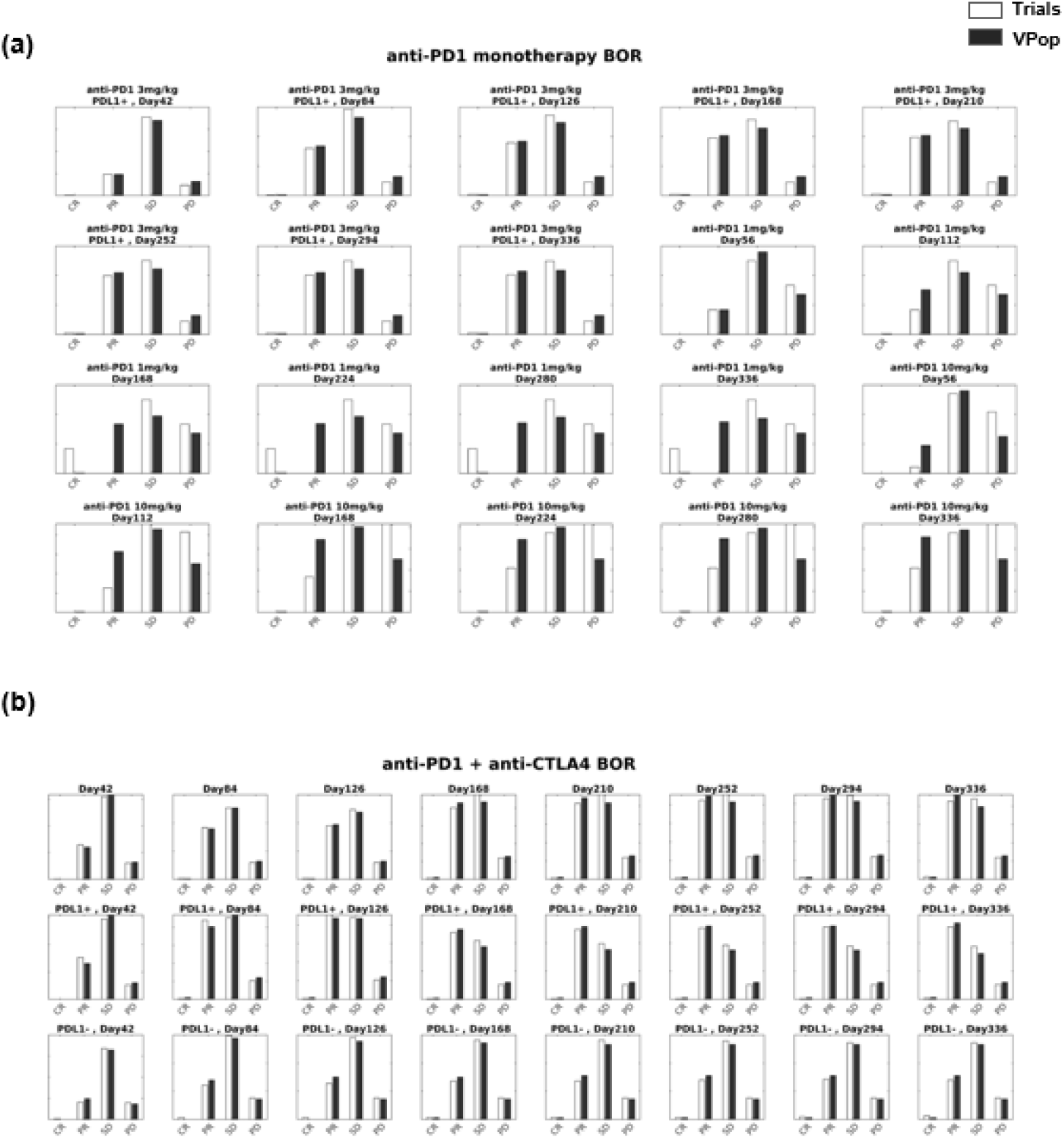
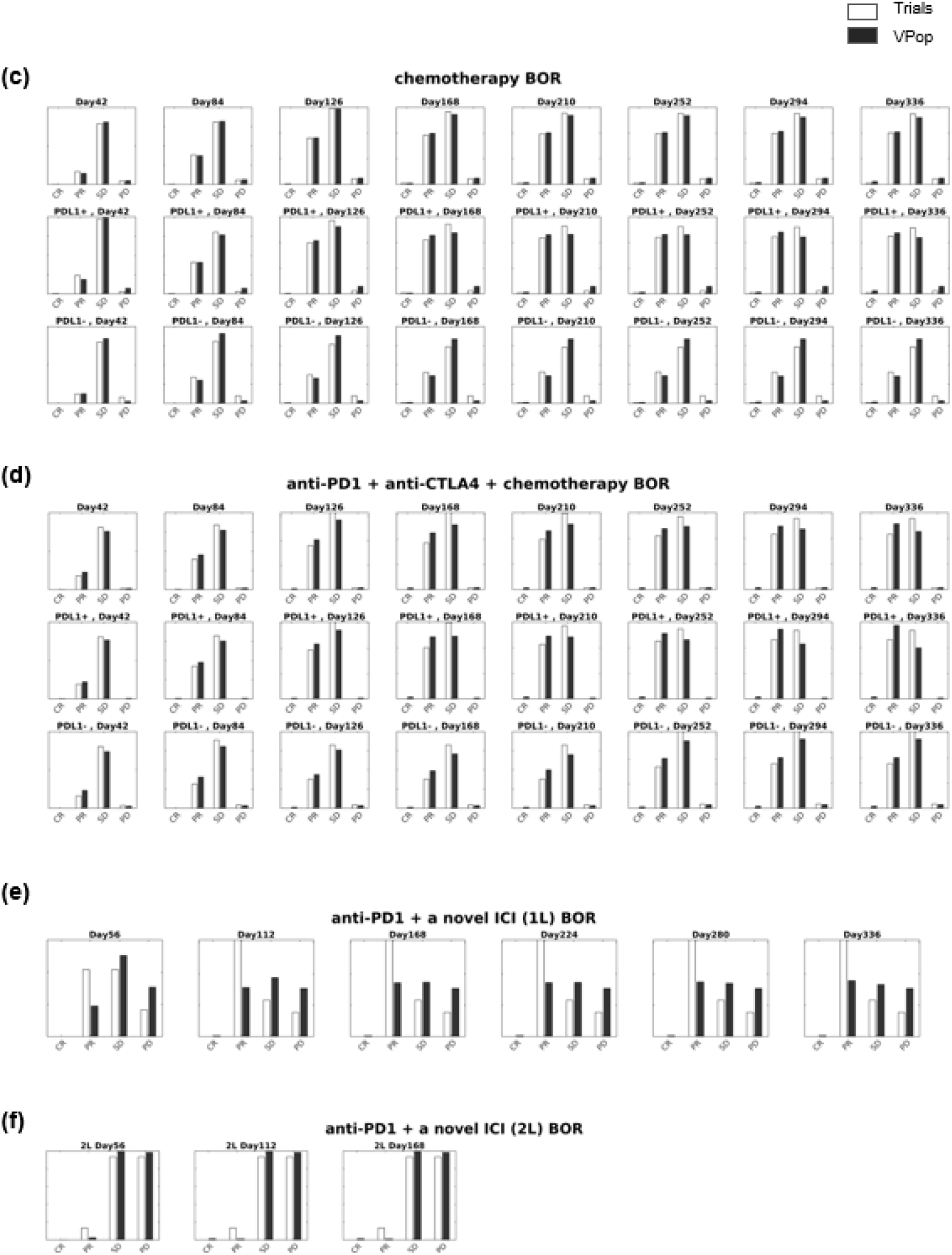
The I-O QSP NSCLC VPop simultaneously captures clinical BOR across multiple trials. Shown are all the clinical response used in the calibration of I-O QSP NSCLC VPop. The VPop captures the best overall response (BOR) to: **(a)** anti-PD1 monotherapy, **(b)** anti-PD1 and anti-CTLA4 combination therapy, **(c)** chemotherapy, **(d)** combination therapy of anti-PD1, anti-CTLA4 and chemotherapy, **(e)** first-line combination therapy of a novel ICI and anti-PD1, and **(f)** second-line combination therapy of a novel ICI and anti-PD1. The white bars represent the clinical data and the black bars represent the VPop response.

**Supplementary File 3**. QSP Toolbox and scripts for *Case Study 1*

