## Supplementary File 3 for "A surrogate workflow for virtual population development: designing an amenable proxy for a hypothesis-test inspired nonlinear goodness-of-fit": step1_preloading_initialVPCohort.m

```
%% This script prepares the VP cohort as a initial VPop for calibration
% worksheet - unweighted VP cohort, preloaded from example folders
% mapelOptions - calibration targets

clear all;
clc;

% Toolbox initialization
% toolboxPath = 'DRIVE:/DIRs/QSPToolbox';
% addpath(toolboxPath);
initQSPToolbox;

%% load in the worksheet (with adc model) and regenerate results.
myWorksheetFileName = 'example_cohort_worksheet1'; % check this RT?
myWorksheet = loadWorksheet(myWorksheetFileName);
mySimulateOptions = simulateOptions();
mySimulateOptions.rerunExisting = true; 

% add in target time point to ensure we store the simulation results at the correct time points
myMapelOptions = mapelOptions;
myMapelOptions.expData = convertResponseTypeToExpDataTable(myWorksheet, 'N87_agx', 'USUBJID');
myWorksheet.simProps.sampleTimes = sort(unique([0:.025:35,myMapelOptions.expData.time'])','ascend');

% adjust the relativeTolerance 
% we had separate simuations confirm that this relativeTolerance setting speeds up the solver, 
% and gives numerically similar results as the worksheet original setting 1e-12
myWorksheet.simProps.relativeTolerance = 1E-6;

% record only target biomarkers to save storage space
myWorksheet.simProps.saveElementResultIDs={'parrule_payload_fraction_of_initial_bound_intracellular'
    'parrule_tumor_volume'
    'parrule_shed_total_plasma_concentration'
    'parrule_payload_total_apparent_pet_tumor_nanomole_per_gram'};

% simulate and save the worksheet with results
myWorksheet = simulateWorksheet(myWorksheet, mySimulateOptions);

saveWorksheetAutoSplit(myWorksheet,'example_cohort_worksheet_SIM');


%% load in the mapelOptions from the example VPop
myVPop = loadVPop('example_vpop');

% update the VPop, add expDataID to check for the calibration target, 
% for use in linearCalibration and expandVPopEffN
myVPop=getSimData(myVPop,myWorksheet)
myVPop=myVPop.addTableSimVals
myVPop=myVPop.addPredTableVals

myVPop.pwStrategy = 'direct';
myVPop.optimizeType = 'gapso';
myMapelOptions = initializeVPopPropertiesToOption(myVPop);
myMapelOptions.minEffN = 0;

% We had identified several time/experiment datapoints that would be 
% difficult to match and set the influence of these on the optimization to 0.
myMapelOptions.mnSDTable{9,'weightSD'}=0;
myMapelOptions.mnSDTable{9,'weightMean'}=0;
myMapelOptions.binTable{9,'weight'}=0;

myMapelOptions.mnSDTable{20,'weightSD'}=0;
myMapelOptions.mnSDTable{20,'weightMean'}=0;
myMapelOptions.binTable{20,'weight'}=0;

saveMapelOptions(myMapelOptions,'example_MapelOptions');
```
