## Supplementary File 3 for "A surrogate workflow for virtual population development: designing an amenable proxy for a hypothesis-test inspired nonlinear goodness-of-fit": step2_CS1_surrogateVPopWorkflow.m

```
clear all;
clc;

% Toolbox initialization
% toolboxPath = 'DRIVE:/DIRs/QSPToolbox';
% addpath(toolboxPath);
initQSPToolbox;

%% We demonstrate our new surrogate VPop developement workflow with expandVPopEffN with linearCalibration
% A automatic VPop resampling and reweighting workflow to generate 
% a well calibrated VPop (p-value>0.9) using wrapper function expandVPopEffN

%% load the initial VP cohort: worksheet and MapelOptions prepared from step1_preprocessing.m
myWorksheet = loadWorksheet('example_cohort_worksheet_SIM');
myMapelOptions = loadMapelOptions('example_MapelOptions');

%% load the initial VPop
myVPop = loadVPop('example_vpop');

% update the VPop, add expDataID to check for the calibration target, for use in linearCalibration and expandVPopEffN
myVPop=getSimData(myVPop,myWorksheet)
myVPop=myVPop.addTableSimVals
myVPop=myVPop.addPredTableVals

myVPop.pwStrategy = 'direct';
myVPop.optimizeType = 'gapso';

myExpandVPopEffNOptions = expandVPopEffNOptions;
% An identifying suffix
myExpandVPopEffNOptions.suffix = 'example06SurrogateWf';
myExpandVPopEffNOptions.wsIterCounter = 0;
% Normally target 150 and often pause at 50
% in initial run.  We'll stop here too in order to
% adjust expansion sized
myExpandVPopEffNOptions.targetEffN = 40;
% myMapelOptions.minEffN = 0;
myExpandVPopEffNOptions.maxNewPerIter= 10;
myExpandVPopEffNOptions.expandCohortSize = 5000;
myExpandVPopEffNOptions.effNDelta = 2;
myExpandVPopEffNOptions.minPVal = 0.9;
myExpandVPopEffNOptions.nTries = 1;
myExpandVPopEffNOptions.nRetries = 1;
myExpandVPopEffNOptions.verbose = true;
myExpandVPopEffNOptions.restartPVal = 1E-4;
myExpandVPopEffNOptions.expandRandomStart = 0;
myExpandVPopEffNOptions.varyMethod = 'gaussian';
myExpandVPopEffNOptions.resampleStd = 0.05;
myExpandVPopEffNOptions.expandEdgeVPs=true;
myExpandVPopEffNOptions.maxNewPerOld = 3;
% Here, we enforce any checks on sampled VPs,
% before simulation and adjust VPs if needed.
myExpandVPopEffNOptions.screenFunctionName = '';
myExpandVPopEffNOptions.nCluster = 400;

% all above seetings are the same as the original workflow 
% Different settings to run the surrogate workflow: make sure set linearExpandFlag=true
myExpandVPopEffNOptions.linearExpandFlag = true;
myExpandVPopEffNOptions.minEffNlinearflag = true;
% also set cohortSize and maxIter if running this workflow
myExpandVPopEffNOptions.linearExpandCohortSize = 5000;
myExpandVPopEffNOptions.maxIterlinearExpand = 20;
myExpandVPopEffNOptions.nVPMax = 1000;
myExpandVPopEffNOptions.minPVallinear = 0.1;


% This takes a while to run.  Every iteration will
% write to file when a better VPop is found
% and a new worksheet will write to file
% when new VPs are added.
% You can delete intermediate iterations between your start file and
% stopping point.  They are included to allow you to more easily
% track progress.
[newWorksheet, newVPop] = expandVPopEffN(myWorksheet,myExpandVPopEffNOptions,myMapelOptions,myVPop);
```
