## Supplementary File 3 for "A surrogate workflow for virtual population development: designing an amenable proxy for a hypothesis-test inspired nonlinear goodness-of-fit": convertResultTableToStruct.m

```
function myWorksheet = convertResultTableToStruct(myWorksheet, myResultTable)
% This function takes a worksheet and a resultTable
% and overwrites the results in the worksheet with the
% results in the table
%
% ARGUMENTS
%  myWorksheet:            A worksheet; the results field
%                          will be overwritten.     
%  fileName:               (optional) If provided, results will be written
%                           to file        
%
% RETURNS
%  myResultTable:          A matlab table with the results.  
%
continueFlag = true;
if nargin > 2
    warning(['Too many input arguments to ',mfilename,'. Require: myWorksheet, myResultTable.  Exiting.'])
    continueFlag = false;
elseif nargin > 1
    continueFlag = true; 
elseif nargin >= 1
	fileName = '';
    continueFlag = true; 	
elseif nargin < 1 
    warning(['Insufficient input arguments to ',mfilename,'. Require: myWorksheet, myResultTable.  Exiting.'])
    continueFlag = false;   
end

if continueFlag
	interventionIDs = getInterventionIDs(myWorksheet);
	vpIDs = getVPIDs(myWorksheet);
	tableInterventionIDs = unique(myResultTable.('interventionID'),'stable');
	tableVPIDs = myResultTable.Properties.VariableDescriptions(4:end);
	myOutputTimes = unique(myResultTable.('time'),'stable');
	mySaveElementResultIDs = unique(myResultTable.('elementResultID'),'stable')';
	[nInterventions, nVPs] = size(myWorksheet.results);
	if (length(interventionIDs) ~= sum(ismember(tableInterventionIDs,interventionIDs)))
		warning(['Interventions in myResultTable should match internventions in myWorksheet call to ',mfilename,'. Exiting.'])
		continueFlag = false;		
	end
	if (length(vpIDs) ~= sum(ismember(tableVPIDs,vpIDs)))
		warning(['VPs in myResultTable should match internventions in myWorksheet call to ',mfilename,'. Exiting.'])
		continueFlag = false;		
	end	
	% We do not check the times or saveElementResultIDs here.
end	


if continueFlag
    nOutputTimes = length(myOutputTimes);
    nSaveElementResultIDs = length(mySaveElementResultIDs);
    resultArray = cell(nInterventions, nVPs);
    for vpCounter = 1 : nVPs
        vpID = vpIDs(vpCounter);
        tableCol = find(ismember(tableVPIDs,vpID))+3;
        for interventionCounter = 1 : nInterventions
            interventionID = interventionIDs{interventionCounter};
            simStruct.Data = nan(nOutputTimes,nSaveElementResultIDs+1);
            simStruct.Names = ['time',mySaveElementResultIDs];
            simStruct.Data(:,1) = myOutputTimes;
            myWorksheet.results{interventionCounter,vpCounter};
            for resultCounter = 1 : nSaveElementResultIDs
                curSaveElementResultID = mySaveElementResultIDs{resultCounter};			
                rowIndices = find(ismember(myResultTable{:,'elementResultID'},{curSaveElementResultID}) & ismember(myResultTable.('interventionID'),interventionID));
                simStruct.Data(:,resultCounter+1) = myResultTable{rowIndices,tableCol};
            end
            resultArray{interventionCounter,vpCounter} = simStruct;
        end
    end	
end
    
if continueFlag
	myWorksheet.results = resultArray;
end
    
    
```
