## Supplementary File 3 for "A surrogate workflow for virtual population development: designing an amenable proxy for a hypothesis-test inspired nonlinear goodness-of-fit": createSimResultTable.m

```
function myResultTable = createSimResultTable(myWorksheet, fileName)
% This function takes a worksheet results structure
% and creates a simulation result table from it.
% Note that you need to include the file extension in
% myFileName.
%
% ARGUMENTS
%  myWorksheet:            A worksheet; the results field
%                          will be overwritten.     
%  fileName:               (optional) If provided, results will be written
%                           to file        
%
% RETURNS
%  myResultTable:          A matlab table with the results.  
%
continueFlag = true;
addFlag = false;
if nargin > 2
    warning(['Too many input arguments to ',mfilename,'. Require: myWorksheet, and optionally fileName.  Exiting.'])
    continueFlag = false;
elseif nargin > 1
    continueFlag = true; 
elseif nargin >= 1
	fileName = '';
    continueFlag = true; 	
elseif nargin < 1 
    warning(['Insufficient input arguments to ',mfilename,'. Require: myWorksheet, and optionally fileName.  Exiting.'])
    continueFlag = false;   
end

if continueFlag
	interventionIDs = getInterventionIDs(myWorksheet);
	vpIDs = getVPIDs(myWorksheet);
	[nInterventions, nVPs] = size(myWorksheet.results);
	if ((nInterventions ~= length(interventionIDs)) || (nVPs ~= length(vpIDs)))
		warning(['Results should be populated before call to ',mfilename,'. Exiting.'])
		continueFlag = false;
	else
		addFlag = true;
	end
end	

myOutputTimes = myWorksheet.simProps.sampleTimes;
mySaveElementResultIDs = myWorksheet.simProps.saveElementResultIDs;
mySaveElementResultIDs = unique(mySaveElementResultIDs,'stable');
nOutputTimes = length(myOutputTimes);
nSaveElementResultIDs = length(mySaveElementResultIDs);

resultMatrix = nan(nOutputTimes*nSaveElementResultIDs*nInterventions,nVPs+1);
resultMatrix(:,1) = repmat(myOutputTimes,nSaveElementResultIDs*nInterventions,1);
variableStrings = cell(nSaveElementResultIDs*nInterventions*nOutputTimes,1);
interventionStrings = cell(nSaveElementResultIDs*nInterventions*nOutputTimes,1);

vpCounter = 1;
while addFlag
	interventionCounter = 1;
	while interventionCounter <= nInterventions
		curResults = myWorksheet.results{interventionCounter,vpCounter};
		for resultCounter = 1 : nSaveElementResultIDs
			startRow = ((resultCounter-1)*nOutputTimes+1)+(interventionCounter-1)*nOutputTimes*nSaveElementResultIDs;
			endRow = ((resultCounter)*nOutputTimes)+(interventionCounter-1)*nOutputTimes*nSaveElementResultIDs;
			variableStrings(startRow:endRow)=mySaveElementResultIDs(resultCounter);
			interventionStrings(startRow:endRow)=interventionIDs(interventionCounter);
			if strcmp(class(curResults),'struct')
                curIndex = find(ismember(curResults.Names,mySaveElementResultIDs{resultCounter}));
				resultMatrix(startRow:endRow,vpCounter+1) = curResults.Data(:,curIndex);
            end
            
		end	
		interventionCounter = interventionCounter+1;
	end
	if vpCounter < nVPs 
		vpCounter = vpCounter+1;
	else
		addFlag = false;
	end
end	

if continueFlag
	myResultTable = array2table(resultMatrix,'VariableNames',genvarname(['time',vpIDs]));
	T2 = cell2table([variableStrings,interventionStrings],'VariableNames',{'elementResultID','interventionID'});
	myResultTable = [T2,myResultTable];
    myResultTable.Properties.VariableDescriptions = ['elementResultID','interventionID','time',vpIDs];
	if length(fileName) > 0
		writetable(myResultTable, fileName);
	end
end

end
    
    
```
