## Supplementary File 3 for "A surrogate workflow for virtual population development: designing an amenable proxy for a hypothesis-test inspired nonlinear goodness-of-fit": plotVarsAcrossVPIntervention.m

```
function plotHandleVector = plotVarsAcrossVPIntervention(myWorksheet, myPlotVarsOptions)
% This function plots multiple variables separately for VP-Intervention simulations.
%
% ARGUMENTS
% myWorksheet:          (required) Worksheet with simulation results
% myPlotVarsOptions:    (required) A plotVarsOptions structure to adjust the plotting
%
% RETURNS
% plotHandle
%
% Check number of inputs.
% We define fail behavior as warn rather than error here
% This just seems a more reasonable way to fail if we can generate plots
% rather than interrupting higher level execution with an error.
plotHandleVector = [];

if nargin > 2
    flagContinue = false;
    warning(['Too many input arguments for ',mfilename,', require: myWorksheet, myPlotVarsOptions. Exiting.'])
elseif nargin > 1
    flagContinue = true;
else
    flagContinue = false;
    warning(['Insufficient input arguments for ',mfilename,', require: myWorksheet, myPlotVarsOptions. Exiting.']) 
end

if flagContinue
    if ~strcmp(class(myPlotVarsOptions),'plotVarsOptions')
        flagContinue = false;
        warning(['Invalid plotVarsOptions object for ',mfilename,', exiting.'])
    end           
end

if flagContinue
    if length(myPlotVarsOptions.vpIDs) < 1
        myPlotVarsOptions.vpIDs = getVPIDs(myWorksheet);
    end
end

if flagContinue
    flagContinue = myPlotVarsOptions.verify(myWorksheet);
    if ~flagContinue
        warning(['The plotVarsOptions failed verification for ',mfilename,', exiting.'])
    end
end


if flagContinue
    vpIDs = myPlotVarsOptions.vpIDs;
    % Check whether the desired VPs exists
    allVPIDs = getVPIDs(myWorksheet);
    keepIndices = find(ismember(allVPIDs,vpIDs));
	vpIDs = allVPIDs(keepIndices);
	allInterventionIDs = getInterventionIDs(myWorksheet);
    interventionIDs = myPlotVarsOptions.interventionIDs; 
	keepInterventionIndices = find(ismember(allInterventionIDs,interventionIDs));	
	interventionIDs = allInterventionIDs(keepInterventionIndices);
    varNames = myPlotVarsOptions.varNames;
end

% Experiment data not included here

% TO CONSIDER:
% May want to scan VPs for the simulation lag parameter and
% update the plot options
xVarName = 'time';   
legendEntries = cell(1,0);
if flagContinue
    % Now we can finally generate the plots!
    figure;
    nVP = length(keepIndices);
	nIntervention = length(keepInterventionIndices);
    % Reseed the random number generator to get consistent if randomly sequenced plot colors
    rng('default');
    randColors = rand(nVP,3);
	for interventionCounter = 1 : nIntervention
		interventionIndex = keepInterventionIndices(interventionCounter);
		for vpCounter = 1 : nVP
			vpIndex = keepIndices(vpCounter);
			legendEntries{1,vpCounter} = myWorksheet.vpDef{vpIndex}.('ID');       
			wshVarNames = myWorksheet.results{interventionIndex,vpIndex}.Names;
			timeIndex = find(ismember(wshVarNames,xVarName));
			varIndices = find(ismember(wshVarNames,varNames));
			timeVal = myWorksheet.results{interventionIndex,vpIndex}.Data(:,timeIndex);
			varVals = myWorksheet.results{interventionIndex,vpIndex}.Data(:,varIndices);
			figure;
			hold on
			for plotCounter = 1 : length(varIndices)
				plotHandleVector(plotCounter) = plot(timeVal-myPlotVarsOptions.xShiftSim, varVals(:,plotCounter), myPlotVarsOptions.lineSpecs{plotCounter}, 'LineWidth',4); 
			end
			if (strcmp(myPlotVarsOptions.scale,'xlogylin') || strcmp(myPlotVarsOptions.scale,'xlogylog'))
				set(gca,'xscale','log'); % gca
			end
			if (strcmp(myPlotVarsOptions.scale,'xlogylog') || strcmp(myPlotVarsOptions.scale,'xlinylog'))
				set(gca,'yscale','log'); % gca
			end        

			if length(myPlotVarsOptions.yLim) > 0
				ylim(myPlotVarsOptions.yLim);
			end 
			if length(myPlotVarsOptions.xLabelPretty) > 0
				xlabel(myPlotVarsOptions.xLabelPretty);
			else
				xlabel(xVarName,'interpreter','none');
			end
			if length(myPlotVarsOptions.yLabelPretty) > 0
				ylabel(myPlotVarsOptions.yLabelPretty);
			else
				ylabel(varName,'interpreter','none');
			end    
			if length(myPlotVarsOptions.xLim) > 0
				xlim(myPlotVarsOptions.xLim);
			end    
			if length(myPlotVarsOptions.yLim) > 0
				ylim(myPlotVarsOptions.yLim);
			end     
			grid on
			set(gca,'fontsize', 18);
			if myPlotVarsOptions.flagLegend
				legend(legendEntries,'interpreter','none');
			end
			if myPlotVarsOptions.flagSave
				if length(myPlotVarsOptions.fileName) > 0
					print([myPlotVarsOptions.fileName,'_',vpIDs{vpCounter},'_',interventionIDs{interventionCounter},'.tif'],'-dtiff','-r300');
				else
					theDate = date;
					formatOut = 'yymmdd';
					theDate = datestr(theDate,formatOut);            
					print([vpIDs{vpCounter},'_',interventionIDs{interventionCounter},'_',theDate,'.tif'],'-dtiff','-r300');
				end
			end
		end
	end
end

end
```
