## Supplementary File 3 for "A surrogate workflow for virtual population development: designing an amenable proxy for a hypothesis-test inspired nonlinear goodness-of-fit": plotVarsOptions.m

```
classdef plotVarsOptions
% Here, we define the plotOptions class to help quickly make acceptable
% plots of results
%
%  Properties
%      flagSave:        Boolean (true/false) indicating whether to save the
%                       figure.  Default is false.
%      fileName:        String indicating the save filename, '.tif' will be
%                       appended.  Leave as '' for a default name if saving.
%      flagLegend:      Boolean (true/false) indicating whether to add a
%                       legend. Default is false.
%      scale:           String indicating X/Y scale, valid strings are:
%                       'xlinylin' (default)
%                       'xlinylog'
%                       'xlogylog'
%                       'xlogylin'
%      xLim:            1x2 numeric matrix of X plotting limits.  Leave as
%                       [] for MATLAB's default.
%      yLim:            1x2 numeric matrix of Y plotting limits.  Leave as
%                       [] for MATLAB's default.
%      xShiftSim:       Double, x-value (time) to shift 
%                       simulation results by. (Optional), leave as 0
%                       so as not to shift.
%      xLabelPretty:    String, will be applied to xlabel. Optional,
%                       if left as '' a default label will be used.
%      yLabelPretty:    String, will be applied to ylabel. Optional,
%                       if left as '' a default label will be used.
%      vpIDs:           Cell array of strings, VP IDs from the worksheet to
%                       include.  (Optional) If {} or plotting a population
%                       all VPs are included.
%      varNames:        Cell array, variable names from the results to plot.  
%      interventionIDs: Cell array, intervention IDs from the worksheet to plot.
%      lineSpecs
%
   properties
      flagSave
      fileName
      flagLegend
      scale
      xLim
      yLim
      xShiftSim
      xLabelPretty
      yLabelPretty
      vpIDs
      varNames
      interventionIDs
	  lineSpecs
   end
   methods
      function obj = set.flagSave(obj,flagvalue)
          if (flagvalue == true) || (flagvalue == false)
              obj.flagSave = flagvalue;
          else
            error('Invalid flagSave value, expecting true/false')
          end
      end
      function obj = set.fileName(obj,myFileName)
          if (ischar(myFileName) == true) 
              obj.fileName = myFileName;
          else
              error('Invalid fileName value, expecting a string')
          end
      end           
      function obj = set.flagLegend(obj,flagvalue)
          if (flagvalue == true) || (flagvalue == false)
              obj.flagLegend = flagvalue;
          else
            error('Invalid flagLegend value, expecting true/false')
          end
      end         
      function obj = set.scale(obj,scalevalue)
          if (strcmpi(scalevalue,'xlinylin') ||...
               strcmpi(scalevalue,'xlinylog') ||...
               strcmpi(scalevalue,'xlogylog') || ...
               strcmpi(scalevalue,'xlogylin'))
            obj.scale = scalevalue;
          else
            error('Invalid scale setting, allowed setting: "xlinylin," "xlinylog," "xlogylog," "xlogylin."')
          end
      end       
      function obj = set.xLim(obj,xLimVal)
          if ((isnumeric(xLimVal) == true) &&...
                  ((length(xLimVal) == 0) || (length(xLimVal) == 2)))
              obj.xLim = xLimVal;
          else
            error('Invalid xLim value, expecting a 1x2 numeric matrix')
          end
      end 
      function obj = set.yLim(obj,ylim)
          if ((isnumeric(ylim) == true) &&...
                  ((length(ylim) == 0) || (length(ylim) == 2)))
              obj.yLim = ylim;
          else
            error('Invalid yLim value, expecting a 1x2 numeric matrix')
          end
      end 
      function obj = set.xShiftSim(obj,xShiftSimVal)
          if (isnumeric(xShiftSimVal) == true) 
              obj.xShiftSim = xShiftSimVal;
          else
            error('Invalid xShiftSim value, expecting a numeric value')
          end
      end 
      function obj = set.xLabelPretty(obj,xLabelPrettyVal)
          if (ischar(xLabelPrettyVal) == true) 
              obj.xLabelPretty = xLabelPrettyVal;
          else
            error('Invalid xLabelPretty value, expecting a string')
          end
      end       
      function obj = set.yLabelPretty(obj,yLabelPrettyVal)
          if (ischar(yLabelPrettyVal) == true) || (iscell(yLabelPrettyVal) == true)
              obj.yLabelPretty = yLabelPrettyVal;
          else
            error('Invalid yLabelPretty value, expecting a string (or cell array of strings for multiple lines).')
          end
      end      
      function obj = set.vpIDs(obj,myVPIDs)
          if (iscell(myVPIDs) == true) 
              obj.vpIDs = myVPIDs;
          else
            error('Invalid myVPIDs value, expecting a cell array')
          end
      end     
      function obj = set.varNames(obj,myVarNames)
          if (iscell(myVarNames) == true) 
              obj.varNames = myVarNames;
          else
            error('Invalid myVarName value, expecting a cell array')
          end
      end 
      function obj = set.interventionIDs(obj,myInterventionNames)
          if (iscell(myInterventionNames) == true) 
              obj.interventionIDs = myInterventionNames;
          else
            error('Invalid myInterventionNames value, expecting a cell array')
          end
      end    
      function obj = set.lineSpecs(obj,myLineSpecs)
          if (iscell(myLineSpecs) == true) 
              obj.lineSpecs = myLineSpecs;
          else
            error('Invalid myLineSpecs value, expecting a cell array')
          end
      end   	  
      
      function passCheck = verify(obj,myWorksheet)
          passCheck = true;
          passCheck = obj.verifyVPIDs(myWorksheet);
          passInterventions = obj.verifyInterventionIDs(myWorksheet);
          passCheck = passCheck && passInterventions;
          [nInterventionResults, nVPResults] = size(myWorksheet.results);
          myInterventionIDs = getInterventionIDs(myWorksheet);
          if passInterventions
              if nInterventionResults == length(myInterventionIDs)
                  myIndices = find(ismember(myInterventionIDs,obj.interventionIDs));
                  for interventionCounter = 1 : length(myIndices)
                      myResultClasses = cellfun(@class,myWorksheet.results(myIndices(interventionCounter),:), 'UniformOutput', false);
                      nSimResults = sum(strcmp(myResultClasses,'struct'));
                      if nSimResults == nVPResults;
                          passCheck = passCheck && true;
                      else
                          warning(['Simulation results not fully populated for specified intervention in worksheet provided to ',mfilename,'.'])
                          passCheck = false;
                      end
                  end
              else
                  warning(['Simulation results not fully populated for specified intervention in worksheet provided to ',mfilename,'.'])
                  passCheck = false;
              end
          end
      end
          
      function passCheck = verifyVPIDs(obj,myWorksheet)  
          passCheck = true;
          vpIDs = obj.vpIDs;
          % Check whether the desired VPs exists
          allVPIDs = getVPIDs(myWorksheet);
          keepIndices = find(ismember(allVPIDs,vpIDs));
          vpIDs = allVPIDs(keepIndices);
          if length(keepIndices) < 1
              warning(['Specified vpIDs not in worksheet provided to ',mfilename,'.'])
              passCheck = false;
          end
      end
          
      function passCheck = verifyInterventionIDs(obj,myWorksheet)  
          passCheck = true;      
          interventionIDsSpec = obj.interventionIDs;
          interventionIDs = getInterventionIDs(myWorksheet);
          if sum(ismember(interventionIDs,interventionIDsSpec)) < length(interventionIDsSpec)
              passCheck = false;
              warning(['Specified interventionIDs all not in worksheet provided to ',mfilename,'.'])
          end
      end
          
      
      function value = get(obj,propName)
          switch propName
              case 'flagSave'
                  value = obj.flagSave;
              case 'fileName'
                  value = obj.fileName;                  
              case 'flagLegend'
                  value = obj.flagLegend;
              case 'scale'
                  value = obj.scale;                  
              %case 'axesCor'
              %    value = obj.axesCor;
              case 'xLim'
                  value = obj.xLim;
              case 'yLim'
                  value = obj.yLim;
              case 'xShiftSim'
                  value = obj.xShiftSim;
              case 'xLabelPretty'
                  value = obj.xLabelPretty;
              case 'yLabelPretty'
                  value = obj.yLabelPretty;
              case 'vpIDs'
                  value = obj.vpIDs;
              case 'varNames'
                  value = obj.varNames;  
              case 'interventionIDs'
                  value = obj.interventionIDs;       
              case 'lineSpecs'
                  value = obj.lineSpecs;  				  
              otherwise
                  error(['Error: ',propName ,' is not a valid ',mfilename,' property.'])
          end
      end      

      % The constructor method must have the same name as the class
      function obj = plotVarsOptions()
            obj.flagSave = false;
            obj.fileName = '';            
            obj.flagLegend = false;
            obj.scale = 'xlinylin';
            obj.xLim = [];
            obj.yLim = [];
            obj.xShiftSim = 0;
            obj.xLabelPretty = '';
            obj.yLabelPretty = '';   
            obj.vpIDs = {};
            obj.varNames = {};
            obj.interventionIDs = {};
            obj.lineSpecs = {};
			
      end
   end
end
```
