## Supplementary File 3 for "A surrogate workflow for virtual population development: designing an amenable proxy for a hypothesis-test inspired nonlinear goodness-of-fit": prccSensitivityOptions.m

```
classdef prccSensitivityOptions
    % Options object for running PRCC sensitivity analysis.
    % Note that the sensitivity analysis must be run with the returned
    % worksheet from a sampling step
    % PROPERTIES
    % analyzeElementResultIDs: Worksheet elementIDs to run analysis
    %                           on.
    % analyzeInterventionIDs:  Worksheet interventions to run analysis
    %                           on.
    % analyzeTimes:            (Required) Simulation time points to run analysis
    %                           at.  If not specified, the analysis will be run
    %                           at all sampled timepoints.  Also,
    %                           this should correstpond to an exact sampled
    %                           simulation timepoint as interpolation is not
    %                           implemented
    % poolClose:                Whether to close the pool at the end of
    %                            calculations
    % poolRestart:              Whether to restart the pool at the beginning
    %                            of calculations
    % clusterID:                Which cluster to use, parallel.defaultClusterProfile is default
    % nWorkers:                 How many workers in the pool to use.  Set to nan to
    %                            use the default (default).
    %
    %  TODO: add in subsampling to assess if enough VPs are run
    %        for consistent results
    %
    %  confidenceInterval:      '1' computes confidence intervals for PRCCs
    %                           by sampling VPs according to prevalence weights
    %                           '0' computes PRCCs from all VPs in worksheet
    %  sampleSize:              number of samples ( used to compute CI)
    %  cohortSize:              number of VPs per sample
    %
    %  figstoPlot:              a cell array indicating the type of PRCC to plot
    %                           options: prcc , prccw, prccabs , prccwabs, prccregr, prccwregr
    %
    %  doseresponsetoPlot:      a cell array indicating in which order
    %                           dose-reseponse curves are to be plotted,
    %                  options: prccorder, prccabsorder, prccregrorder, prccworder, prccwabsorder, prccwregrorder
    %
    %  whiskerPlot:            '1' plots PRCCs as whisker plots instead of bar plots
    %
    %  vpWeights:               weight vector of dimension nVP x 1
    %
    %  numVP:                  number of worksheet VPs (computed automatically)
    
    properties
        
        
        analyzeElementResultIDs
        analyzeInterventionIDs
        analyzeTimes
        poolClose
        poolRestart
        clusterID
        nWorkers
        confidenceInterval
        sampleSize
        cohortSize
        figstoPlot
        doseresponsetoPlot
        whiskerPlot
        vpWeights
        numVP
        totalbin
        
    end
    
    methods(Static)
        
        function [corre_xy_weight]= weightedCorr(myParamsOrderedRes, myDataOrderedRes, w, absCCflag)
            
            x = myParamsOrderedRes;
            y = myDataOrderedRes;
            n= length(x);
            mean_x_weight = sum(w.*x);
            mean_y_weight = sum(w.*y);
            
            %if absCCflag == false -> don't take absolute values of residuals
            if absCCflag==false
                
                residual_x_weight = (x-mean_x_weight);
                residual_y_weight = (y-mean_y_weight);
                
                %elseif absCCflag == true -> take absolute values of residuals
            elseif absCCflag==true
                
                residual_x_weight = abs(x-mean_x_weight);
                residual_y_weight = abs(y-mean_y_weight);
                
                
                %elseif same as absCCflag == false
            else
                
                residual_x_weight = (x-mean_x_weight);
                residual_y_weight = (y-mean_y_weight);
                
            end
            
            variance_x_weight= (n/(n-1))*sum(w.*residual_x_weight.^2);
            variance_y_weight= (n/(n-1))*sum(w.*residual_y_weight.^2);
            std_x_weight =  sqrt(variance_x_weight);
            std_y_weight= sqrt(variance_y_weight);
            corre_xy_weight= (n/(n-1))*((sum(w.*residual_x_weight.*residual_y_weight))/(std_x_weight*std_y_weight));
            % if there is one biomarker data, this might cause numerical issues 
            % so we put a logic here, to force corr within (-1,1)
            if corre_xy_weight>0
                corre_xy_weight = min(corre_xy_weight,1);
            elseif corre_xy_weight<0
                corre_xy_weight=max(corre_xy_weight,-1);
            end      
        end
        
        
        
        function [PickedWeights, PickedVP]= samplefromWeights(newWorksheet_input, myPRCCSensitivityOptions )
            
            CohortSize= myPRCCSensitivityOptions.cohortSize;
            nSample=myPRCCSensitivityOptions.sampleSize;
            allVPIDs = getVPIDs(newWorksheet_input);
            PickedVP=NaN(nSample,CohortSize);
            PickedWeights = NaN(nSample,CohortSize);
            
            myWeights= myPRCCSensitivityOptions.vpWeights;
            
            [pwSort, pwInd]=sort(myWeights);
            nVP=length(pwSort);
            pwSort_cumsum=NaN(nVP,1);
            for i=1:nVP
                pwSort_cumsum(i)=sum(pwSort(1:i)); % cumulative weight function
            end
            
            
            for k=1:nSample
                for j=1:CohortSize
                    vp_rand=rand(1);
                    ind_rand=find(vp_rand<pwSort_cumsum); % select VP based on weight
                    PickedVP(k,j)=pwInd(ind_rand(1)); % assign original VP index
                    PickedWeights(k,j) = myWeights(pwInd(ind_rand(1)));
                end
                PickedWeights(k,:) = PickedWeights(k,:)./sum(PickedWeights(k,:));
            end
            
            
            
        end
        
        function [doseresponses]= doseresponseGenerator(myWorksheet, myPRCCSensitivityOptions)
            w=myPRCCSensitivityOptions.vpWeights';
            allElementResultIDs = myWorksheet.simProps.saveElementResultIDs;
            allSampleTimes = myWorksheet.simProps.sampleTimes;
            allInterventionIDs = getInterventionIDs(myWorksheet); % Myintervations PRCC handles this.
            % find indices for myOutputVariable and mySampleTimes
            myOutputVariables = myPRCCSensitivityOptions.analyzeElementResultIDs;
            mySampleTimes = myPRCCSensitivityOptions.analyzeTimes;
            myInterventionIDs = myPRCCSensitivityOptions.analyzeInterventionIDs;
            [~,myOutputVariableIndex] = ismember(myOutputVariables,allElementResultIDs);
            
            [~,mySampleTimesIndices] = ismember(mySampleTimes,allSampleTimes);
            
            myPRCCSensitivityOptions.analyzeInterventionIDs = getInterventionIDs(myWorksheet);
            % get names of all mechanistic axes (Parameters)
            myAxisDefs = getAxisDefIDs(myWorksheet)';
            % define number of Interventions, SampleTimes and OutputVariables
            nInterventionIDs = length(myInterventionIDs);
            nTimes = length(mySampleTimes);
            nOutputs = length(myOutputVariables);
            mysampletimes= myPRCCSensitivityOptions.analyzeTimes;
            totalbin= myPRCCSensitivityOptions.totalbin;
            
            for n=1:nOutputs
                for i=1:nInterventionIDs
                    % curResult.Data (nTimes x nVPs) contains times series for all VPs
                    % for specified Intervention and OutputVariable
                    curResult = getResultOutputforIntervention(myWorksheet,myInterventionIDs{i},allElementResultIDs{myOutputVariableIndex(n)});
                    curData = curResult.Data(mySampleTimesIndices,2:end);
                    myData = curData'; % nVPs x nTimes
                    
                    myVPCoeffs = getVPCoeffs(myWorksheet); % nParams x nVPs
                    myParams = myVPCoeffs'; % nVPs x nParams
                    
                    % % find indices for myOutputVariable and mySampleTimes
                    [~,myOutputVariableIndex] = ismember(myOutputVariables,allElementResultIDs);
                    [~,mySampleTimesIndices] = ismember(mySampleTimes,allSampleTimes);
                    
                    %elementsids_int=myPRCCSensitivityOptions.analyzeElementResultIDs
                    intervation= i;
                    
                    [doseresponses]= prccSensitivityOptions.binGenerator(myData,myParams ,intervation,  myWorksheet, totalbin);
                    
                end
            end
        end
        
        
        %[~,doseresponses]= runPRCCSensitivity(curWorksheet, myPRCCSensitivityOptions, w, 1);
        
        
        
        function [myPRCC] = runPRCCsample(newWorksheet_input, myPRCCSensitivityOptions, curElementResultIDstring)
            confidenceinterval= myPRCCSensitivityOptions.confidenceInterval;
            [PickedWeights, PickedVP]= prccSensitivityOptions.samplefromWeights(newWorksheet_input, myPRCCSensitivityOptions);
            if myPRCCSensitivityOptions.poolRestart
                if ~isempty(gcp('nocreate'))
%                     delete(gcp); % commented this to not restart everytime
                else
                    % First check the default number of workers, if needed
                    myPRCCSensitivityOptions = checkNWorkers(myPRCCSensitivityOptions);
                    myPool = parpool(myPRCCSensitivityOptions.clusterID,myPRCCSensitivityOptions.nWorkers,'SpmdEnabled',false);
                    
                end
            end
            if confidenceinterval==1
                disp('computing PRCC samples')
                nSample= myPRCCSensitivityOptions.sampleSize;
                myPRCCSample = cell(nSample,1);
                if myPRCCSensitivityOptions.poolRestart==false
                    for j=1:nSample
                        curWorksheet = keepVPs(newWorksheet_input, PickedVP(j,:));
                        [myPRCCSample{j}]= runPRCCSensitivity(curWorksheet, myPRCCSensitivityOptions, PickedWeights(j,:)');
                        disp(['sample: ',num2str(j),'/',num2str(nSample),' | ElementResultID: ',curElementResultIDstring,newline])
                        %disp(['sample: ',num2str(j),'/',num2str(nSample),' | ElementResultID: '])
                    end
                else
                    parfor j=1:nSample
                        curWorksheet = keepVPs(newWorksheet_input, PickedVP(j,:));
                        [myPRCCSample{j}]= runPRCCSensitivity(curWorksheet, myPRCCSensitivityOptions, PickedWeights(j,:)');
                        disp(['sample: ',num2str(j),'/',num2str(nSample),' | ElementResultID: ',curElementResultIDstring,newline])
                        %disp(['sample: ',num2str(j),'/',num2str(nSample),' | ElementResultID: '])
                    end
                    
                end
                
                
                w= myPRCCSensitivityOptions.vpWeights';
                [myPRCCWorksheet]= runPRCCSensitivity(newWorksheet_input, myPRCCSensitivityOptions, w);
                myPRCC.myPRCCWorksheet= myPRCCWorksheet;
                myPRCC.myPRCCSample= myPRCCSample;
                [myPRCC]= prccSensitivityOptions.computePRCCci(newWorksheet_input,myPRCCSensitivityOptions,myPRCC);
                
                
            else
                
                w= myPRCCSensitivityOptions.vpWeights';
                [myPRCCWorksheet]= runPRCCSensitivity(newWorksheet_input, myPRCCSensitivityOptions, w);
                myPRCC.myPRCCWorksheet= myPRCCWorksheet;
                myPRCC.myPRCCSample= {};
            end
            
            doseresponses= prccSensitivityOptions.doseresponseGenerator(newWorksheet_input , myPRCCSensitivityOptions);
            %alldoseresponses{i}= doseresponses;
            myPRCC.doseresponses=doseresponses;
        end
        
        function [myPRCC]= computePRCCci(newWorksheet_input,myPRCCSensitivityOptions , myPRCC)
            nSample= myPRCCSensitivityOptions.sampleSize;
            nAxes = length(getAxisDefIDs(newWorksheet_input)');
            PRCCwabsMatrix = zeros(nSample, nAxes);
            PRCCabsMatrix = zeros(nSample, nAxes);
            PRCCwMatrix = zeros(nSample, nAxes);
            PRCCMatrix = zeros(nSample, nAxes);
            PRCCwregrMatrix = zeros(nSample, nAxes);
            PRCCregrMatrix = zeros(nSample, nAxes);
            for i=1:nSample
                PRCCwabsMatrix(i,:)=myPRCC.myPRCCSample{i}{1}.PRCCwabs;
                PRCCabsMatrix(i,:)=myPRCC.myPRCCSample{i}{1}.PRCCabs;
                PRCCwMatrix(i,:)=myPRCC.myPRCCSample{i}{1}.PRCCw;
                PRCCMatrix(i,:)=myPRCC.myPRCCSample{i}{1}.PRCC;
                PRCCwregrMatrix(i,:)=myPRCC.myPRCCSample{i}{1}.PRCCwregr;
                PRCCregrMatrix(i,:)=myPRCC.myPRCCSample{i}{1}.PRCCregr;
            end
            
            PRCCwabsMean = mean(PRCCwabsMatrix);
            PRCCabsMean = mean(PRCCabsMatrix);
            PRCCwMean = mean(PRCCwMatrix);
            PRCCMean = mean(PRCCMatrix);
            PRCCwregrMean = mean(PRCCwregrMatrix);
            PRCCregrMean = mean(PRCCregrMatrix);
            
            PRCCwabsMedian = median(PRCCwabsMatrix);
            PRCCabsMedian = median(PRCCabsMatrix);
            PRCCwMedian = median(PRCCwMatrix);
            PRCCMedian = median(PRCCMatrix);
            PRCCwregrMedian = median(PRCCwregrMatrix);
            PRCCregrMedian = median(PRCCregrMatrix);
            
            PRCCwabsPrct = prctile(PRCCwabsMatrix,[2.5 97.5]);
            PRCCabsPrct = prctile(PRCCabsMatrix,[2.5 97.5]);
            PRCCwPrct = prctile(PRCCwMatrix,[2.5 97.5]);
            PRCCPrct = prctile(PRCCMatrix,[2.5 97.5]);
            PRCCwregrPrct = prctile(PRCCwregrMatrix,[2.5 97.5]);
            PRCCregrPrct = prctile(PRCCregrMatrix,[2.5 97.5]);
            
            
            myPRCC.myPRCCavg.PRCCMean = PRCCMean;
            myPRCC.myPRCCavg.PRCCwMean =PRCCwMean;
            myPRCC.myPRCCavg.PRCCabsMean = PRCCabsMean ;
            myPRCC.myPRCCavg.PRCCwabsMean = PRCCwabsMean ;
            myPRCC.myPRCCavg.PRCCregrMean = PRCCregrMean ;
            myPRCC.myPRCCavg.PRCCwregrMean = PRCCwregrMean ;
            
            myPRCC.myPRCCavg.PRCCMedian = PRCCMedian;
            myPRCC.myPRCCavg.PRCCwMedian = PRCCwMedian;
            myPRCC.myPRCCavg.PRCCabsMedian = PRCCabsMedian ;
            myPRCC.myPRCCavg.PRCCwabsMedian = PRCCwabsMedian ;
            myPRCC.myPRCCavg.PRCCregrMedian = PRCCregrMedian ;
            myPRCC.myPRCCavg.PRCCwregrMedian = PRCCwregrMedian ;
            
            myPRCC.myPRCCavg.cf = PRCCPrct;
            myPRCC.myPRCCavg.cfw = PRCCwPrct;
            myPRCC.myPRCCavg.cfabs = PRCCabsPrct;
            myPRCC.myPRCCavg.cfwabs = PRCCwabsPrct;
            myPRCC.myPRCCavg.cfregr = PRCCregrPrct;
            myPRCC.myPRCCavg.cfwregr = PRCCwregrPrct;
            
            myPRCC.myPRCCavg.PRCCMatrix = PRCCMatrix;
            myPRCC.myPRCCavg.PRCCwMatrix = PRCCwMatrix;
            myPRCC.myPRCCavg.PRCCabsMatrix = PRCCabsMatrix ;
            myPRCC.myPRCCavg.PRCCwabsMatrix = PRCCwabsMatrix ;
            myPRCC.myPRCCavg.PRCCregrMatrix = PRCCregrMatrix ;
            myPRCC.myPRCCavg.PRCCwregrMatrix = PRCCwregrMatrix ;
            
            
            %myPRCC.myPRCCavg.Names= myPRCC.myPRCCSample{i}{1}.Names;
            
        end
        
        function [final]= binGenerator(myData,myParams ,intervation,  myWorksheet, totalbin)
            axisnumber= length(myWorksheet.axisProps.axisDef);
            
            for ix= 1:axisnumber
                name= myWorksheet.axisProps.axisDef{ix}.id;
                param= myParams(:,ix);
                sizebin= totalbin;
                [N, edges] = histcounts(param, sizebin);
                
                data_attime= myData;
                
%                 for outputix=1:size(myParams, 2) % this for-loop over outputix can be completely removed
                    final_per_axis =[];
                    std_per_axis = [];
                    cihalfa =[];
                    data=data_attime;
                    %edges_data=struct();
                    for edgesix= 1:length(edges)-1
                        %edges_name=['edge',num2str(edgesix)];
                        %edges_output=['edgeoutput',num2str(edgesix)];
                        
                        location_boolean= (param >= edges(edgesix)  & param < edges(edgesix+1));
                        location_ix= find(location_boolean ==1);
                        data_to = data(location_ix);
                        param_edges=param(location_ix);
                        %edges_data.(edges_output)=data_to;
                        %edges_data.(edges_name)= param_edges;
                        avg_data= mean(data_to);
                        std_data= std(data_to);
                        final_per_axis=[final_per_axis; avg_data];
                        std_per_axis = [std_per_axis; std_data];
                        cihalf= 1.96*std_data/sqrt(length(data_to));
                        cihalfa =[cihalfa ; cihalf];
                    end
                    outputf= final_per_axis;
                    
                    
                    outputfnew= outputf(isnan(outputf)==0);
                    cv= std(outputfnew)/mean(outputfnew);
                    
                    
                    
                    confidenceinterval=  cihalfa;
                    final(intervation, ix).output=outputf;
                    final(intervation, ix).param=param;
                    final(intervation, ix).edges=edges;
                    %final(intervation, ix).edgesdata=edges_data;
                    final(intervation, ix).N=N;
                    final(intervation, ix).confidenceinterval=confidenceinterval;
                    final(intervation, ix).cv= cv;
                    final(intervation, ix).sizebin= sizebin;
                    output= final(intervation, ix).output;
                    
%                 end
            end
        end
        
        function [pvalue]= pvaluePRCC(N,dparams, ccvalue, absoption)
            
            
            
            
            Tcc = ccvalue*sqrt((N-2-dparams)/(1-ccvalue^2));
            
            if absoption==false
                pvalue= 2*tcdf(Tcc,N-dparams-2); % if cc can be positive or negative
            else
                pvalue= tcdf(Tcc,N-dparams-2);% if cc can only be positive
                
            end
        end
        
        function [doseresponseFigs,PRCCfigs] = plotPRCC(newWorksheet_input,  myPRCC,myPRCCSensitivityOptions,nCC, ntoPlot)
            %doseresponseFig,PRCCfig, absPRCCfig , weightPRCCfig,  absweightPRCCfig
            CCType= 'PRCC';
            myElementResultIDs = myPRCCSensitivityOptions.analyzeElementResultIDs;
            myInterventions=myPRCCSensitivityOptions.analyzeInterventionIDs;
            myTimes=myPRCCSensitivityOptions.analyzeTimes;
            doseresponsetoPlot=myPRCCSensitivityOptions.doseresponsetoPlot;
            figstoPlot=myPRCCSensitivityOptions.figstoPlot;
            confidenceinterval=myPRCCSensitivityOptions.confidenceInterval;
            whisker= myPRCCSensitivityOptions.whiskerPlot;
            if whisker ==1 && confidenceinterval==0
                disp(['To generate WhiskerPlots recompute PRCC with confidenceInterval=1',newline, '... generating barplot instead'])
                
                
            end
            if ~isempty(figstoPlot)
                nFigsToPlot = length(figstoPlot);
                PRCCfigs = gobjects(nFigsToPlot,1);
            end
            plotcounter= 0;
            curERIdx= eval(ntoPlot);
            if ismember('prcc',figstoPlot) % section of PRCC
                if confidenceinterval==1 && whisker==0
                    Y = myPRCC{curERIdx}.myPRCCavg.PRCCMean; %% Caution: this is actually weighted mean?
                    Name='PRCC';
                elseif confidenceinterval==0
                    Y = myPRCC{curERIdx}.myPRCCWorksheet{1}.PRCC;
                    Name='PRCC';
                elseif whisker==1 && confidenceinterval==1
                    Y = myPRCC{curERIdx}.myPRCCavg.PRCCMedian; %% Caution: this is actually weighted median?
                    Name='PRCC (whisker)';
                    
                end
                %plotcounter= plotcounter +1;
                %PRCCfigs(plotcounter)= figure('Name',Name);
                [~, PRCC_Ind] = sort(abs(Y),'descend');
                Y_sorted = Y(PRCC_Ind);
                X_sorted=categorical(myPRCC{curERIdx}.myPRCCWorksheet{1}.Names(PRCC_Ind));
                loopb=1;
                if nCC > 20
                    loopb= ceil(nCC/20);
                end
                for ll= 1:loopb
                    plotcounter= plotcounter +1;
                    PRCCfigs(plotcounter)= figure('Name',Name);
                    axa=gca;
                    n_start = 1+(ll-1)*20;
                    n_end = min(20+(ll-1)*20,nCC);
                    Yfinal= Y_sorted(n_start:n_end);
                    Xfinal= X_sorted(n_start:n_end);
                    nCCt= length(Yfinal);
                    if whisker == 0 || (whisker == 1 && confidenceinterval ==0)
                        b=barh(Yfinal');
                        
                        Xfinalt= Xfinal';
                        hold on
                        set(axa, 'Xtick', -1:0.5:1,'YTick',1:nCCt,  'YTickLabels',Xfinalt);
                        set(axa,'TickLabelInterpreter','none')
                        axis([-1 1 0.5 (nCCt+0.5)]);
                        title([{myInterventions{1}, myElementResultIDs{1},['day ', num2str(myTimes)]}],'Interpreter','none')
                        %title([{myInterventions, myElementResultIDs,['day ', num2str(myTimes)]}],'Interpreter','none')
                        xlabel('PRCC')
                        set(gcf,'Position',[1300 300 600 500])
                        confidenceinterval= myPRCCSensitivityOptions.confidenceInterval;
                        if confidenceinterval==1
                            cf= myPRCC{curERIdx}.myPRCCavg.cf(:,PRCC_Ind);
                            %cf = cf(:,1:nCC);
                            cf= cf(:, n_start:n_end);
                            errorbar(Yfinal',1:nCCt,Yfinal-cf(1,:),cf(2,:)-Yfinal,'.', 'horizontal','LineWidth',1.5);
                            %errorbar(Y,1:2,[0.02 0.02],'.', 'horizontal');
                        end
                        hold off
                    elseif whisker==1 && confidenceinterval==1 % generate boxplot
                        PRCCMatrix= myPRCC{curERIdx}.myPRCCavg.PRCCMatrix(:,PRCC_Ind);
                        fitaxis=PRCCMatrix(:, n_start:n_end);
                        boxplot(fitaxis, 'orientation', 'horizontal')
                        hold on
                        set(gca, 'Xtick', -1:0.5:1, 'YTickLabels',Xfinal);
                        set(gca,'TickLabelInterpreter','none')
                        
                    end
                end
            end
            %Type of PRCC
            if ismember('prccabs',figstoPlot)
                %Towards Absolute
                if confidenceinterval==1 && whisker==0
                    Y = myPRCC{curERIdx}.myPRCCavg.PRCCMean;
                    Yabs = myPRCC{curERIdx}.myPRCCavg.PRCCabsMean;
                    Name='Absolute PRCC';
                elseif confidenceinterval==0
                    Y = myPRCC{curERIdx}.myPRCCWorksheet{1}.PRCC;
                    Yabs = myPRCC{curERIdx}.myPRCCWorksheet{1}.PRCCabs;
                    Name='Absolute PRCC';
                elseif whisker==1 && confidenceinterval==1
                    Name='Absolute PRCC (whisker)';
                    Y = myPRCC{curERIdx}.myPRCCavg.PRCCMedian;
                    Yabs = myPRCC{curERIdx}.myPRCCavg.PRCCabsMedian;
                    
                end
                X=categorical(myPRCC{curERIdx}.myPRCCWorksheet{1}.Names);
                isneg= find(Y<0);
                isnegnames= X(isneg);
                loopb=1;
                if nCC > 20
                    loopb= ceil(nCC/20);
                end
                for ll= 1:loopb
                    plotcounter=plotcounter +1;
                    PRCCfigs(plotcounter)= figure('Name',Name);
                    [~, PRCC_ABSInd] = sort(Yabs,'descend');
                    Yabs_sorted = Yabs(PRCC_ABSInd);
                    Xsorted=categorical(myPRCC{curERIdx}.myPRCCWorksheet{1}.Names(PRCC_ABSInd));
                    axa=gca;
                    n_start = 1+(ll-1)*20;
                    n_end = min(20+(ll-1)*20,nCC);
                    Yabs_final = Yabs_sorted(n_start:n_end);
                    Xabs_final = Xsorted(n_start:n_end);
                    %pvaluencc= myPRCC{i}{1}.pvalueabs(PRCC_ABSIndabs);
                    %pvaluencc=pvaluencc(1:nCC);
                    
                    nCCt= length(Yabs_final');
                    if whisker == 0 || (whisker == 1 && confidenceinterval ==0)
                        babs=barh(Yabs_final');
                        babs.FaceColor = 'flat';
                        
                        hold on
                        if confidenceinterval==1
                            cf =myPRCC{curERIdx}.myPRCCavg.cfabs(:,PRCC_ABSInd);
                            cf= cf(:,n_start:n_end);
                            errorbar(Yabs_final',1:nCCt,Yabs_final-cf(1,:),cf(2,:)-Yabs_final,'.', 'horizontal','LineWidth',1.5);
                        end
                        % Setting Colors
                        blue= [0 0 1];
                        red= [1 .0 0];
                        index= ismember(Xabs_final,isnegnames );
                        for icolor=1:nCCt
                            redorblue= index(icolor);
                            if redorblue ==1
                                babs.CData(icolor,:)= red;
                            else
                                babs.CData(icolor,:)= blue;
                                
                            end
                        end
                        %Probably needs to be min aspects _ focus
                        set(gca, 'Xtick', 0:0.2:1, 'YTick',1:nCCt,  'YTickLabels',Xabs_final);
                        set(gca,'TickLabelInterpreter','none')
                        axis([0 1 0.5 (nCCt+0.5)]);
                        title([{myInterventions{1}, ...
                            myElementResultIDs{1}, ...
                            ['day ', num2str(myTimes)]}],'Interpreter','none')
                        xlabel('PRCCabs')
                        set(gcf,'Position',[1300 300 600 500])
                        hold off
                    elseif whisker==1 && confidenceinterval==1
                        PRCCabsMatrix= myPRCC{curERIdx}.myPRCCavg.PRCCabsMatrix(:,PRCC_ABSInd);
                        fitaxis=PRCCabsMatrix(:, n_start:n_end);
                        boxplot(fitaxis, 'orientation', 'horizontal')
                        hold on
                        set(gca, 'YTickLabels',Xabs_final);
                        set(gca,'TickLabelInterpreter','none')
                    end
                    
                end
            end
            
            %Type of PRCC
            if ismember('prccregr',figstoPlot)
                if confidenceinterval==1 && whisker==0
                    Y = myPRCC{curERIdx}.myPRCCavg.PRCCMean;
                    Yregr = myPRCC{curERIdx}.myPRCCavg.PRCCregrMean;
                    Name='regr PRCC';
                elseif confidenceinterval==0
                    Y = myPRCC{curERIdx}.myPRCCWorksheet{1}.PRCC;
                    Yregr = myPRCC{curERIdx}.myPRCCWorksheet{1}.PRCCregr;
                    Name='regr PRCC';
                elseif whisker==1 && confidenceinterval==1
                    Name='regr PRCC (whisker)';
                    Y = myPRCC{curERIdx}.myPRCCavg.PRCCMedian;
                    Yregr = myPRCC{curERIdx}.myPRCCavg.PRCCregrMedian;
                    
                end
                X=categorical(myPRCC{curERIdx}.myPRCCWorksheet{1}.Names);
                isneg= find(Y<0);
                isnegnames= X(isneg);
                loopb=1;
                if nCC > 20
                    loopb= ceil(nCC/20);
                end
                for ll= 1:loopb
                    plotcounter=plotcounter +1;
                    PRCCfigs(plotcounter)= figure('Name',Name);
                    [~, PRCC_regrInd] = sort(Yregr,'descend');
                    Yregr_sorted = Yregr(PRCC_regrInd);
                    Xsorted=categorical(myPRCC{curERIdx}.myPRCCWorksheet{1}.Names(PRCC_regrInd));
                    axa=gca;
                    n_start = 1+(ll-1)*20;
                    n_end = min(20+(ll-1)*20,nCC);
                    Yregr_final = Yregr_sorted(n_start:n_end);
                    Xregr_final = Xsorted(n_start:n_end);
                    %pvaluencc= myPRCC{i}{1}.pvalueregr(PRCC_regrIndregr);
                    %pvaluencc=pvaluencc(1:nCC);
                    
                    nCCt= length(Yregr_final');
                    if whisker == 0 || (whisker == 1 && confidenceinterval ==0)
                        bregr=barh(Yregr_final');
                        bregr.FaceColor = 'flat';
                        
                        hold on
                        if confidenceinterval==1
                            cf =myPRCC{curERIdx}.myPRCCavg.cfregr(:,PRCC_regrInd);
                            cf= cf(:,n_start:n_end);
                            errorbar(Yregr_final',1:nCCt,Yregr_final-cf(1,:),cf(2,:)-Yregr_final,'.', 'horizontal','LineWidth',1.5);
                        end
                        % Setting Colors
                        blue= [0 0 1];
                        red= [1 .0 0];
                        index= ismember(Xregr_final,isnegnames );
                        for icolor=1:nCCt
                            redorblue= index(icolor);
                            if redorblue ==1
                                bregr.CData(icolor,:)= red;
                            else
                                bregr.CData(icolor,:)= blue;
                                
                            end
                        end
                        %Probably needs to be min aspects _ focus
                        set(gca, 'Xtick', 0:0.2:1, 'YTick',1:nCCt,  'YTickLabels',Xregr_final);
                        set(gca,'TickLabelInterpreter','none')
                        axis([0 1 0.5 (nCCt+0.5)]);
                        title([{myInterventions{1}, ...
                            myElementResultIDs{1}, ...
                            ['day ', num2str(myTimes)]}],'Interpreter','none')
                        xlabel('PRCCregr')
                        set(gcf,'Position',[1300 300 600 500])
                        hold off
                    elseif whisker==1 && confidenceinterval==1
                        PRCCregrMatrix= myPRCC{curERIdx}.myPRCCavg.PRCCregrMatrix(:,PRCC_regrInd);
                        fitaxis=PRCCregrMatrix(:, n_start:n_end);
                        boxplot(fitaxis, 'orientation', 'horizontal')
                        hold on
                        set(gca, 'YTickLabels',Xregr_final);
                        set(gca,'TickLabelInterpreter','none')
                    end
                    
                end
            end
            
            if ismember('prccw',figstoPlot)
                if confidenceinterval==1 && whisker==0
                    Yw = myPRCC{curERIdx}.myPRCCavg.PRCCwMean; % Lu: incorrect! weighted PRCC mean should be just myPRCCavg.PRCCMean
                    Name='Weighted PRCC';
                    
                elseif confidenceinterval==0
                    Yw = myPRCC{curERIdx}.myPRCCWorksheet{1}.PRCCw;
                    Name='Weighted PRCC';
                elseif whisker==1 && confidenceinterval==1
                    Yw = myPRCC{curERIdx}.myPRCCavg.PRCCwMedian;
                    Name='Weighted PRCC (whisker)';
                    
                end
                [~, PRCCw_Ind] = sort(abs(Yw),'descend');
                Y_sorted = Yw(PRCCw_Ind);
                X_sorted=categorical(myPRCC{curERIdx}.myPRCCWorksheet{1}.Names(PRCCw_Ind));
                
                %pvalue
                %pvalue= myPRCC{i}{1}.pvalue(PRCCw_Ind);
                %pvaluencc= pvalue(1:nCC);
                loopb=1;
                if nCC > 20
                    loopb= ceil(nCC/20);
                end
                for ll= 1:loopb
                    n_start = 1+(ll-1)*20;
                    n_end = min(20+(ll-1)*20,nCC);
                    Yfinalw= Y_sorted(n_start:n_end);
                    Xfinalw= X_sorted(n_start:n_end);
                    plotcounter=plotcounter +1;
                    PRCCfigs(plotcounter)= figure('Name',Name);
                    nCCt=length(Yfinalw');
                    if whisker ==0 || (whisker == 1 && confidenceinterval ==0)
                        
                        bw=barh(Yfinalw');
                        hold on
                        if confidenceinterval==1
                            cf= myPRCC{curERIdx}.myPRCCavg.cfw(:,PRCCw_Ind);
                            cf = cf(:,n_start:n_end);
                            errorbar(Yfinalw',1:nCCt,Yfinalw-cf(1,:),cf(2,:)-Yfinalw,'.', 'horizontal','LineWidth',1.5);
                        end
                        set(gca, 'Xtick', -1:0.5:1, 'YTick',1:nCCt, 'YTickLabels',Xfinalw);
                        set(gca,'TickLabelInterpreter','none')
                        axis([-1 1 0.5 (nCCt+0.5)]);
                        title([{myInterventions{1}, ...
                            myElementResultIDs{1}, ...
                            ['day ', num2str(myTimes)]}],'Interpreter','none')
                        xlabel('wPRCC')
                        set(gcf,'Position',[1300 300 600 500])
                        hold off
                    elseif whisker==1 && confidenceinterval==1
                        
                        PRCCwMatrix= myPRCC{curERIdx}.myPRCCavg.PRCCwMatrix(:,PRCCw_Ind);
                        fitaxis=PRCCwMatrix(:, n_start:n_end);
                        boxplot(fitaxis, 'orientation', 'horizontal')
                        hold on
                        set(gca, 'YTickLabels',Xfinalw);
                        set(gca,'TickLabelInterpreter','none')
                    end
                end
            end
            if ismember('prccwabs',figstoPlot)
                if confidenceinterval==1 && whisker==0
                    Yw = myPRCC{curERIdx}.myPRCCavg.PRCCwMean; % Lu: incorrect! weighted PRCC mean should be just myPRCCavg.PRCCMean
                    Ywabs = myPRCC{curERIdx}.myPRCCavg.PRCCwabsMean; %PRCC w values % Lu: incorrect! weighted PRCCabs mean should be just myPRCCavg.PRCCabsMean
                    Name='Absolute Weighted PRCC';
                    
                elseif confidenceinterval==0
                    Yw = myPRCC{curERIdx}.myPRCCWorksheet{1}.PRCCw;
                    Ywabs = myPRCC{curERIdx}.myPRCCWorksheet{1}.PRCCwabs; %PRCC w values
                    Name='Absolute Weighted PRCC';
                elseif whisker==1 && confidenceinterval==1
                    Name='Absolute Weighted PRCC (whisker)';
                    Yw = myPRCC{curERIdx}.myPRCCavg.PRCCwMedian;
                    Ywabs = myPRCC{curERIdx}.myPRCCavg.PRCCwabsMedian; %PRCC w values
                    
                end
                % Weighted PRCC
                X=categorical(myPRCC{curERIdx}.myPRCCWorksheet{1}.Names); %Names
                isneg= find(Yw<0); %Record which ones are negative
                isnegnames = X(isneg); %Of the one's that are negative save name.
                loopb=1;
                if nCC > 20
                    loopb= ceil(nCC/20);
                end
                for ll= 1:loopb
                    n_start = 1+(ll-1)*20;
                    n_end = min(20+(ll-1)*20,nCC);
                    plotcounter=plotcounter +1;
                    PRCCfigs(plotcounter)= figure('Name',Name); %title
                    [~, PRCC_ABSIndw] = sort(Ywabs,'descend');
                    Ywabs_sorted = Ywabs(PRCC_ABSIndw);
                    Xsorted=categorical(myPRCC{curERIdx}.myPRCCWorksheet{1}.Names(PRCC_ABSIndw));% Names Ywabs sorted
                    Ywabs_final= Ywabs_sorted(n_start:n_end); %values of PRCCwabs sorted
                    Xwabs_final= Xsorted(n_start:n_end); %Most important one's
                    nCCt=length(Ywabs_final);
                    if whisker ==0 || (whisker == 1 && confidenceinterval ==0)
                        bwabs=barh(Ywabs_final');
                        bwabs.FaceColor = 'flat';
                        hold on
                        if confidenceinterval==1
                            cf= myPRCC{curERIdx}.myPRCCavg.cfwabs(:,PRCC_ABSIndw);
                            cf = cf(:,n_start:n_end);
                            errorbar(Ywabs_final',1:nCCt,Ywabs_final-cf(1,:),cf(2,:)-Ywabs_final,'.', 'horizontal','LineWidth',1.5);
                            
                        end
                        % Setting Colors
                        blue= [0 0 1];
                        red= [1 .0 0];
                        index= ismember(Xwabs_final,isnegnames );
                        for icolor=1:nCCt
                            redorblue= index(icolor);
                            if redorblue ==1
                                bwabs.CData(icolor,:)= red;
                            else
                                bwabs.CData(icolor,:)= blue;
                                
                            end
                        end
                        
                        set(gca, 'Xtick', 0:0.2:1,'YTick',1:nCCt,  'YTickLabels',Xwabs_final );
                        set(gca,'TickLabelInterpreter','none')
                        %pwabs p value
                        %labels=strcat('p:',num2str(pvaluewncc, '%0.3f'));
                        %text((Yfinalw +0.05), bwabs.XData',labels,'VerticalAlignment','middle')
                        axis([0 1 0.5 (nCCt+0.5)]);
                        title([{myInterventions{1}, ...
                            myElementResultIDs{1}, ...
                            ['day ', num2str(myTimes)]}],'Interpreter','none')
                        xlabel('PRCCabsweighted')
                        hold off
                        
                        
                    elseif whisker==1 && confidenceinterval==1
                        PRCCwabsMatrix= myPRCC{curERIdx}.myPRCCavg.PRCCwabsMatrix(:,PRCC_ABSIndw);
                        fitaxis=PRCCwabsMatrix(:, n_start:n_end);
                        boxplot(fitaxis, 'orientation', 'horizontal')
                        hold on
                        set(gca, 'YTickLabels',Xwabs_final);
                        set(gca,'TickLabelInterpreter','none')
                        
                    end
                    
                end
            end
            if ismember('prccwregr',figstoPlot)
                if confidenceinterval==1 && whisker==0
                    Yw = myPRCC{curERIdx}.myPRCCavg.PRCCwMean; % Lu: incorrect! weighted PRCC mean should be just myPRCCavg.PRCCMean
                    Ywregr = myPRCC{curERIdx}.myPRCCavg.PRCCwregrMean; %PRCC w values % Lu: incorrect! weighted PRCCregr mean should be just myPRCCavg.PRCCregrMean
                    Name='regr Weighted PRCC';
                    
                elseif confidenceinterval==0
                    Yw = myPRCC{curERIdx}.myPRCCWorksheet{1}.PRCCw;
                    Ywregr = myPRCC{curERIdx}.myPRCCWorksheet{1}.PRCCwregr; %PRCC w values
                    Name='regr Weighted PRCC';
                elseif whisker==1 && confidenceinterval==1
                    Name='regr Weighted PRCC (whisker)';
                    Yw = myPRCC{curERIdx}.myPRCCavg.PRCCwMedian;
                    Ywregr = myPRCC{curERIdx}.myPRCCavg.PRCCwregrMedian; %PRCC w values
                    
                end
                % Weighted PRCC
                X=categorical(myPRCC{curERIdx}.myPRCCWorksheet{1}.Names); %Names
                isneg= find(Yw<0); %Record which ones are negative
                isnegnames = X(isneg); %Of the one's that are negative save name.
                loopb=1;
                if nCC > 20
                    loopb= ceil(nCC/20);
                end
                for ll= 1:loopb
                    n_start = 1+(ll-1)*20;
                    n_end = min(20+(ll-1)*20,nCC);
                    plotcounter=plotcounter +1;
                    PRCCfigs(plotcounter)= figure('Name',Name); %title
                    [~, PRCC_regrIndw] = sort(Ywregr,'descend');
                    Ywregr_sorted = Ywregr(PRCC_regrIndw);
                    Xsorted=categorical(myPRCC{curERIdx}.myPRCCWorksheet{1}.Names(PRCC_regrIndw));% Names Ywregr sorted
                    Ywregr_final= Ywregr_sorted(n_start:n_end); %values of PRCCwregr sorted
                    Xwregr_final= Xsorted(n_start:n_end); %Most important one's
                    nCCt=length(Ywregr_final);
                    if whisker ==0 || (whisker == 1 && confidenceinterval ==0)
                        bwregr=barh(Ywregr_final');
                        bwregr.FaceColor = 'flat';
                        hold on
                        if confidenceinterval==1
                            cf= myPRCC{curERIdx}.myPRCCavg.cfwregr(:,PRCC_regrIndw);
                            cf = cf(:,n_start:n_end);
                            errorbar(Ywregr_final',1:nCCt,Ywregr_final-cf(1,:),cf(2,:)-Ywregr_final,'.', 'horizontal','LineWidth',1.5);
                            
                        end
                        % Setting Colors
                        blue= [0 0 1];
                        red= [1 .0 0];
                        index= ismember(Xwregr_final,isnegnames );
                        for icolor=1:nCCt
                            redorblue= index(icolor);
                            if redorblue ==1
                                bwregr.CData(icolor,:)= red;
                            else
                                bwregr.CData(icolor,:)= blue;
                                
                            end
                        end
                        
                        set(gca, 'Xtick', 0:0.2:1,'YTick',1:nCCt,  'YTickLabels',Xwregr_final );
                        set(gca,'TickLabelInterpreter','none')
                        %pwregr p value
                        %labels=strcat('p:',num2str(pvaluewncc, '%0.3f'));
                        %text((Yfinalw +0.05), bwregr.XData',labels,'VerticalAlignment','middle')
                        axis([0 1 0.5 (nCCt+0.5)]);
                        title([{myInterventions{1}, ...
                            myElementResultIDs{1}, ...
                            ['day ', num2str(myTimes)]}],'Interpreter','none')
                        xlabel('PRCCregrweighted')
                        hold off
                        
                        
                    elseif whisker==1 && confidenceinterval==1
                        PRCCwregrMatrix= myPRCC{curERIdx}.myPRCCavg.PRCCwregrMatrix(:,PRCC_regrIndw);
                        fitaxis=PRCCwregrMatrix(:, n_start:n_end);
                        boxplot(fitaxis, 'orientation', 'horizontal')
                        hold on
                        set(gca, 'YTickLabels',Xwregr_final);
                        set(gca,'TickLabelInterpreter','none')
                        
                    end
                    
                end
            end            
            %Dose Function
            if confidenceinterval==1
                Y = myPRCC{curERIdx}.myPRCCavg.PRCCMean;
                Yw = myPRCC{curERIdx}.myPRCCavg.PRCCwMean;
                Yregr = myPRCC{curERIdx}.myPRCCavg.PRCCregrMean;
                Ywregr = myPRCC{curERIdx}.myPRCCavg.PRCCwregrMean;
                [~, PRCC_Ind] = sort(abs(Y),'descend');
                [~, PRCC_Indw] = sort(abs(Yw),'descend');
                [~, PRCC_Indregr] = sort(abs(Yregr),'descend');
                [~, PRCC_Indwregr] = sort(abs(Ywregr),'descend');
            else
                Y = myPRCC{curERIdx}.myPRCCWorksheet{1}.PRCC;
                Yw = myPRCC{curERIdx}.myPRCCWorksheet{1}.PRCCw;
                Yregr = myPRCC{curERIdx}.myPRCCWorksheet{1}.PRCCregr;
                Ywregr = myPRCC{curERIdx}.myPRCCWorksheet{1}.PRCCwregr;
                
                [~, PRCC_Ind] = sort(abs(Y),'descend');
                
                [~, PRCC_Indw] = sort(abs(Yw),'descend');
                
                [~, PRCC_Indregr] = sort(abs(Yregr),'descend');
                
                [~, PRCC_Indwregr] = sort(abs(Ywregr),'descend');
                
                
            end
            
            if nCC >0 && nCC<=4
                dimensionsncclist = {[4,1]};
                loop=1;
                %doser=4;
                %dosec = 1;
                
            elseif nCC >4 && nCC<=6
                dimensionsncclist  ={[3,2]};
                loop=1;
                %doser=3;
                %dosec = 2;
                
            elseif nCC > 6 && nCC <=10
                dimensionsncclist={[5,2]};
                loop=1;
                %doser=5;
                %dosec = 1;
                
            elseif nCC > 10 && nCC <=15
                dimensionsncclist={[5 , 3]};
                loop=1;
                
            elseif nCC >15 && nCC <= 20
                dimensionsncclist={[5,4]};
                loop=1;
                
            elseif nCC >20
               % dimensionsncclist= {[5,4] , [5,4] , [5,4]};
                loop= ceil(nCC/20);
                dimensionsncclist= repmat({[5,4]},1,loop);
            end
            
            for nccl= 1:loop
                dimensionsncc= dimensionsncclist{nccl};
                doser= dimensionsncc(1);
                dosec= dimensionsncc(2);
                ncc_start = 1+(nccl-1)*20;
                ncc_end = min(20+(nccl-1)*20,nCC);
                net= ncc_end -ncc_start +1;
                %if ~isempty(doseresponsetoPlot)
                nDoseResponsePlots = length(doseresponsetoPlot);
                doseresponseFigs = gobjects(nDoseResponsePlots,1);
                for ndose=1:nDoseResponsePlots
                    dosestoplot_i= doseresponsetoPlot(ndose);
                    if ismember('prccorder', dosestoplot_i)
                        order= PRCC_Ind;
                        order= PRCC_Ind(ncc_start:ncc_end);
                        FigName = 'PRCC';
                    elseif ismember('prccabsorder', dosestoplot_i)
                        order=PRCC_Indabs(ncc_start:ncc_end);
                        FigName = 'Absolute PRCC';
                    elseif ismember('prccregrorder', dosestoplot_i)
                        order=PRCC_Indregr(ncc_start:ncc_end);
                        FigName = 'regr PRCC';
                    elseif ismember('prccwabsorder', dosestoplot_i)
                        order= PRCC_Indwabs(ncc_start:ncc_end);
                        FigName = 'Absolute Weighted PRCC';
                    elseif ismember('prccworder', dosestoplot_i)
                        order= PRCC_Indw(ncc_start:ncc_end);
                        FigName = 'Weighted PRCC';
                    elseif ismember('prccwregrorder', dosestoplot_i)
                        order=PRCC_Indwregr(ncc_start:ncc_end);
                        FigName = 'regr Weighted PRCC';    
                    end
                    
                    %weighted end , starting dose response.
                    finalnew= myPRCC{curERIdx}.doseresponses;
                    outputname= 'Output';
                    Name= ['dose-response curves for: ', FigName];
                    doseresponseFigs(ndose)=figure('Name',Name);
                    
                    for ix2=1:net
                        ix= order(ix2);
                        %subplot(doser,dosec,ix2)
                        subplot(doser,dosec,ix2)
                        output= finalnew(1,ix).output;
                        index=1:length(output);
                        if any(isnan(output))
                            
                            disp(['Some bins of the dose-response curve contain no data points.',newline, ...
                                 'Consider increasing the number of VPs']);
                            M=output;
                            index=find(~isnan(M));
                            %idx=find(diff(index)~=1);
                            %A=[idx(1);diff(idx);numel(index)-idx(end)];
                            %outputchunk=mat2cell(M(~isnan(M)),A,1);
                        end
                        
                        
                        
                        %output(isnan(output))=0;
                        cv= finalnew(1,ix).cv;
                        if isnan(cv)
                            cv='NaN';
                        end
                        %Confidence interval
                        cf=finalnew(1,ix).confidenceinterval;
                        %cv(isnan(output))=0;
                        %Axis name
                        axisname= {newWorksheet_input.axisProps.axisDef{ix}.id, strcat('cv:', string(cv))};
                        %titletemp= {strcat('monotonic:',  string(finalnew{i}(1,ix).monotonic))};
                        %title(titletemp , 'FontSize',8);
                        %xlabel, ylabel , title
                        xlabel(axisname, 'FontSize', 10);
                        ylabel(outputname, 'FontSize', 10);
                        titlep = {myInterventions{1}, myElementResultIDs{1},['day',num2str(myTimes)] } ;
                        %titlep = {myInterventions{1}, myElementResultIDs{1},['day',num2str(myTimes)], ['order:',dosestoplot_i{1}] } ;
                        
                        sgtitle(titlep ,'fontweight','bold','FontSize', 11, 'Interpreter','none');
                        set(gcf,'Position',[1 500 600 800])
                        if any(isnan(output))
                            %
                            hold on
                            edges_filtered=finalnew(1,ix).edges(1,2:end);
                            plot(edges_filtered(index), output(index));
                            errorbar(edges_filtered(index),output(index),cf(index));
                        else
                            hold on
                            plot(finalnew(1,ix).edges(1,2:end), output);
                            errorbar(finalnew(1,ix).edges(1,2:end),output,cf);
                            
                            
                        end
                        hold off
                        
                        
                    end
                end
            end
            
            if ismember('CV',figstoPlot)
                
                Name= 'Cv per parameter';
                Cvf=[];
                for ix= 1:length(newWorksheet_input.axisProps.axisDef)
                    cvix= myPRCC{1}.doseresponses(1,ix).cv;
                    Cvf=[Cvf; cvix];
                end
                Y= Cvf;
                [~, PRCC_Ind] = sort(abs(Y),'descend');
                Y_sorted = Y(PRCC_Ind);
                X_sorted=categorical(myPRCC{curERIdx}.myPRCCWorksheet{1}.Names(PRCC_Ind));
                loopb=1;
                if nCC > 20
                    loopb= ceil(nCC/20);
                end
                for ll= 1:loopb
                    plotcounter= plotcounter +1;
                    Cvfigs(plotcounter)= figure('Name',Name);
                    axa=gca;
                    n_start = 1+(ll-1)*20;
                    n_end = min(20+(ll-1)*20,nCC);
                    Yfinal= Y_sorted(n_start:n_end);
                    Xfinal= X_sorted(n_start:n_end);
                    nCCt= length(Yfinal);
                    b=barh(Yfinal');
                    Xfinalt= Xfinal';
                    
                    xplotmax = ceil(max(abs(Yfinal')));
                    hold on
                    if xplotmax <=1
                        set(axa, 'Xtick', -1:0.5:1,'YTick',1:nCCt,  'YTickLabels',Xfinalt);
                        axis([-1 1 0.5 (nCCt+0.5)]);
                    else
                        set(axa, 'Xtick', -xplotmax:xplotmax/2:xplotmax,'YTick',1:nCCt,  'YTickLabels',Xfinalt);
                        axis([-xplotmax xplotmax 0.5 (nCCt+0.5)]);
                    end
                    set(axa,'TickLabelInterpreter','none')
                    title([{myInterventions{1}, myElementResultIDs{1},['day ', num2str(myTimes)]}],'Interpreter','none')
                    %title([{myInterventions, myElementResultIDs,['day ', num2str(myTimes)]}],'Interpreter','none')
                    xlabel('Cv')
                    set(gcf,'Position',[1300 300 600 500])
                    
                end
            end
            
            
            
        end
        
        
        
        function [tablePRCCs, tablesample] = tableGenerator(newWorksheet_input, myPRCC,myPRCCSensitivityOptions, ntoPlot)
            %clear table
            counter=1;
            
            i=eval(ntoPlot);
            
            myPRCCnew=myPRCC{i};
            
            axisf= myPRCCnew.myPRCCWorksheet{1}.Names;
            nAxis= length(axisf);
            
            analyzeTimes=myPRCCSensitivityOptions.analyzeTimes;
            
            
            
            interventions= myPRCCSensitivityOptions.analyzeInterventionIDs{1};
            sizebin= myPRCCnew.doseresponses(1,1).sizebin;
            
            output= myPRCCnew.myPRCCWorksheet{1}.outputVariable;
            
            outputtemp = [output];
            
            sizebintemp= [sizebin];
            
            Outputc = repmat({outputtemp},length(axisf),1);
            interventionstemp= [interventions];
            interventionsc = repmat({interventionstemp},length(axisf),1);
            analyzeTimestemp= [analyzeTimes];
            analyzeTimesc=  repmat({analyzeTimestemp},length(axisf),1);
            
            sizebinc=  repmat({sizebintemp},length(axisf),1);
            
            
            
            if myPRCCSensitivityOptions.confidenceInterval==1
                %PRCC Weighted abs
                PRCCwabsMean= myPRCCnew.myPRCCavg.PRCCwabsMean;
                cfwabs= myPRCCnew.myPRCCavg.cfwabs;
                %PRCC  abs
                PRCCabsMean= myPRCCnew.myPRCCavg.PRCCabsMean;
                cfabs= myPRCCnew.myPRCCavg.cfabs;
                %PRCC Weighted regr
                PRCCwregrMean= myPRCCnew.myPRCCavg.PRCCwregrMean;
                cfwregr= myPRCCnew.myPRCCavg.cfwregr;
                %PRCC  regr
                PRCCregrMean= myPRCCnew.myPRCCavg.PRCCregrMean;
                cfregr= myPRCCnew.myPRCCavg.cfregr;
                
                %PRCC Weighted
                PRCCwMean = myPRCCnew.myPRCCavg. PRCCwMean;
                cfw= myPRCCnew.myPRCCavg.cfw;
                
                %PRCC
                PRCCMean = myPRCCnew.myPRCCavg.PRCCMean;
                cf= myPRCCnew.myPRCCavg.cf;
                
                
                cfcombined= string(cf(1,:)) + ' ' + string(cf(2, :));
                cfS= cellstr(cfcombined');
                
                
                cfwcombined= string(cfw(1,:)) + ' ' + string(cfw(2, :));
                cfwS= cellstr(cfwcombined');
                
                
                
                cfwabscombined= string(cfwabs(1,:)) + ' ' + string(cfwabs(2, :));
                cfwabsS= cellstr(cfwabscombined');
                
                cfabscombined= string(cfabs(1,:)) + ' ' + string(cfabs(2, :));
                cfabsS= cellstr(cfabscombined');
                
                cfwregrcombined= string(cfwregr(1,:)) + ' ' + string(cfwregr(2, :));
                cfwregrS= cellstr(cfwregrcombined');
                
                cfregrcombined= string(cfregr(1,:)) + ' ' + string(cfregr(2, :));
                cfregrS= cellstr(cfregrcombined');               
                
                
                PRCCwabsMedian= myPRCCnew.myPRCCavg.PRCCwabsMedian;
                PRCCabsMedian= myPRCCnew.myPRCCavg.PRCCabsMedian;
                PRCCwregrMedian= myPRCCnew.myPRCCavg.PRCCwregrMedian;
                PRCCregrMedian= myPRCCnew.myPRCCavg.PRCCregrMedian;
                
                PRCCwMedian = myPRCCnew.myPRCCavg.PRCCwMedian;
                PRCCMedian = myPRCCnew.myPRCCavg.PRCCMedian;
                
                
                PRCCwabsMatrix= myPRCCnew.myPRCCavg.PRCCwabsMatrix;
                PRCCabsMatrix= myPRCCnew.myPRCCavg.PRCCabsMatrix;
                PRCCwregrMatrix= myPRCCnew.myPRCCavg.PRCCwregrMatrix;
                PRCCregrMatrix= myPRCCnew.myPRCCavg.PRCCregrMatrix;
                
                PRCCwMatrix = myPRCCnew.myPRCCavg.PRCCwMatrix;
                PRCCMatrix = myPRCCnew.myPRCCavg.PRCCMatrix;
                nsample = length(PRCCMatrix);
                
                axis_sample= myPRCCnew.myPRCCWorksheet{1}.Names;
                output_sample= myPRCCnew.myPRCCWorksheet{1}.outputVariable;
                output_sampletemp = [output_sample];
                interventions_sampletemp= [ interventions];
                analyzeTime_sampletemp= [analyzeTimes];
                sizebin_sampletemp= [sizebin];
                Outputc_sample = repmat({output_sampletemp},nAxis*nsample,1);
                
                
                interventionS = repmat({interventions_sampletemp},nAxis*nsample,1);
                analyzeTimeS= repmat({analyzeTime_sampletemp},nAxis*nsample,1);
                
                sizebinS= repmat({sizebin_sampletemp},nAxis*nsample,1);
                
                
                PRCCwabssample= {};
                PRCCabssample= {};
                PRCCwregrsample= {};
                PRCCregrsample= {};
                PRCCwsample= {};
                PRCCsample= {};
                axisS={};
                samplenumberall={};
                PRCCMeanSf=      { };
                PRCCwMeanSf=      { };
                PRCCabsMeanSf=      { };
                PRCCwabsMeanSf=      { };
                PRCCregrMeanSf=      { };
                PRCCwregrMeanSf=      { };
                
                PRCCMedianSf=      { };
                PRCCwMedianSf=      { };
                PRCCabsMedianSf=      { };
                PRCCwabsMedianSf=      { };
                PRCCregrMedianSf=      { };
                PRCCwregrMedianSf=      { };
                
                for ix =1:nAxis
                    
                    PRCCwabssample= [PRCCwabssample; num2cell(PRCCwabsMatrix(:,ix))] ;
                    PRCCabssample= [PRCCabssample; num2cell(PRCCabsMatrix(:,ix))] ;
                    PRCCwregrsample= [PRCCwregrsample; num2cell(PRCCwregrMatrix(:,ix))] ;
                    PRCCregrsample= [PRCCregrsample; num2cell(PRCCregrMatrix(:,ix))] ;
                    PRCCwsample= [PRCCwsample; num2cell(PRCCwMatrix(:,ix))] ;
                    PRCCsample= [PRCCsample; num2cell(PRCCMatrix(:,ix))] ;
                    axis_sampletemp = [axis_sample{ix}];
                    axis_samplec = repmat({axis_sampletemp},nsample,1);
                    
                    samplenumber=num2cell(1:nsample)';
                    
                    axisS= [axisS; axis_samplec] ;
                    
                    samplenumberall= [samplenumberall; samplenumber] ;
                    
                    
                    PRCCMeantemp  = [PRCCMean(ix)];
                    PRCCwMeantemp  = [PRCCwMean(ix)];
                    PRCCabsMeantemp = [PRCCabsMean(ix)];
                    PRCCwabsMeantemp = [PRCCwabsMean(ix)];
                    PRCCregrMeantemp = [PRCCregrMean(ix)];
                    PRCCwregrMeantemp = [PRCCwregrMean(ix)];
                    
                    PRCCMediantemp = [PRCCMedian(ix)];
                    PRCCwMediantemp = [PRCCwMedian(ix)];
                    PRCCabsMediantemp = [PRCCabsMedian(ix)];
                    PRCCwabsMediantemp = [PRCCwabsMedian(ix)];
                    PRCCregrMediantemp = [PRCCregrMedian(ix)];
                    PRCCwregrMediantemp = [PRCCwregrMedian(ix)];
                    
                    PRCCMeanS     = repmat({PRCCMeantemp},nsample,1);
                    PRCCwMeanS= repmat({PRCCwMeantemp},nsample,1);
                    PRCCabsMeanS= repmat({PRCCabsMeantemp},nsample,1);
                    PRCCwabsMeanS= repmat({PRCCwabsMeantemp},nsample,1);
                    PRCCregrMeanS= repmat({PRCCregrMeantemp},nsample,1);
                    PRCCwregrMeanS= repmat({PRCCwregrMeantemp},nsample,1);
                    
                    PRCCMedianS= repmat({PRCCMediantemp},nsample,1);
                    PRCCwMedianS= repmat({PRCCwMediantemp},nsample,1);
                    PRCCabsMedianS= repmat({PRCCabsMediantemp},nsample,1);
                    PRCCwabsMedianS= repmat({PRCCwabsMediantemp},nsample,1);
                    PRCCregrMedianS= repmat({PRCCregrMediantemp},nsample,1);
                    PRCCwregrMedianS= repmat({PRCCwregrMediantemp},nsample,1);
                    PRCCMeanSf=  [ PRCCMeanSf ; PRCCMeanS ];
                    PRCCwMeanSf=  [ PRCCwMeanSf ; PRCCwMeanS ];
                    PRCCabsMeanSf=  [ PRCCabsMeanSf ; PRCCabsMeanS ];
                    PRCCwabsMeanSf=  [PRCCwabsMeanSf  ; PRCCwabsMeanS ];
                    PRCCregrMeanSf=  [ PRCCregrMeanSf ; PRCCregrMeanS ];
                    PRCCwregrMeanSf=  [PRCCwregrMeanSf  ; PRCCwregrMeanS ];
                    
                    PRCCMedianSf=  [ PRCCMedianSf ; PRCCMedianS ];
                    PRCCwMedianSf=  [ PRCCwMedianSf ; PRCCwMedianS ];
                    PRCCabsMedianSf=  [ PRCCabsMedianSf ; PRCCabsMedianS ];
                    PRCCwabsMedianSf=  [ PRCCwabsMedianSf ; PRCCwabsMedianS ];
                    PRCCregrMedianSf=  [ PRCCregrMedianSf ; PRCCregrMedianS ];
                    PRCCwregrMedianSf=  [ PRCCwregrMedianSf ; PRCCwregrMedianS ];
                    
                    
                end
                
                
                tablesample= table(interventionS, Outputc_sample, analyzeTimeS, axisS, samplenumberall, PRCCsample,PRCCMeanSf, PRCCMedianSf, PRCCwsample, PRCCwMeanSf, ...
                    PRCCwMedianSf,  PRCCabssample, PRCCabsMeanSf, PRCCabsMedianSf,  PRCCwabssample, PRCCwabsMeanSf, PRCCwabsMedianSf, ...
                    PRCCregrsample, PRCCregrMeanSf, PRCCregrMedianSf,  PRCCwregrsample, PRCCwregrMeanSf, PRCCwregrMedianSf, sizebinS);
                
                PRCCwabsMeanc= num2cell(PRCCwabsMean');
                PRCCabsMeanc= num2cell(PRCCabsMean');
                PRCCwregrMeanc= num2cell(PRCCwregrMean');
                PRCCregrMeanc= num2cell(PRCCregrMean');
                
                PRCCwMeanc = num2cell(PRCCwMean');
                PRCCMeanc = num2cell(PRCCMean');
                
                
                PRCCwabsMedianc= num2cell(PRCCwabsMedian');
                PRCCabsMedianc= num2cell(PRCCabsMedian');
                PRCCwregrMedianc= num2cell(PRCCwregrMedian');
                PRCCregrMedianc= num2cell(PRCCregrMedian');
                
                PRCCwMedianc = num2cell(PRCCwMedian');
                PRCCMedianc = num2cell(PRCCMedian');
                
                
                tablePRCCs= table(interventionsc, Outputc, analyzeTimesc,  axisf, PRCCMeanc,cfS,  PRCCwMeanc, cfwS,  PRCCabsMeanc, cfabsS,  PRCCwabsMeanc, cfwabsS, ...
                    PRCCregrMeanc, cfregrS,  PRCCwregrMeanc, cfwregrS, ...
                    PRCCMedianc, PRCCwMedianc,  PRCCabsMedianc, PRCCwabsMedianc, PRCCregrMedianc, PRCCwregrMedianc, sizebinc);
                
            else
                
                PRCCwabs= myPRCCnew.myPRCCWorksheet{1}.PRCCwabs;
                PRCCabs= myPRCCnew.myPRCCWorksheet{1}.PRCCabs;
                PRCCwregr= myPRCCnew.myPRCCWorksheet{1}.PRCCwregr;
                PRCCregr= myPRCCnew.myPRCCWorksheet{1}.PRCCregr;
                PRCCw = myPRCCnew.myPRCCWorksheet{1}.PRCCw;
                PRCC = myPRCCnew.myPRCCWorksheet{1}.PRCC;
                CV = [myPRCCnew.doseresponses.cv];
                
                
                PRCCwabsf= num2cell(PRCCwabs');
                PRCCabsf= num2cell(PRCCabs');
                PRCCwregrf= num2cell(PRCCwregr');
                PRCCregrf= num2cell(PRCCregr');
                PRCCwf = num2cell(PRCCw');
                PRCCf = num2cell(PRCC');
                CVf = num2cell(CV');
                
                tablePRCCs= table(interventionsc, Outputc, analyzeTimesc, ...
                    axisf, PRCCf, PRCCwf,  PRCCabsf, PRCCwabsf, PRCCregrf, PRCCwregrf, CVf, sizebinc);
                
                tablesample= {};
                
                
            end
            
            
            
        end
        
        
        
%         function [S, rT ,sign]=fcorr(R)
%             % Routine for the calculation of top-down correlation coefficient
%             % R: matrix of ranks by columns.   S: Savage scores
%             % based on Iman, R.L. and Conover W. J., A Measure of Top-Down
%             % Correlation. Technometrics, August 1987, vol. 29 (3)
%             [n k]=size(R);
%             S=zeros(n,k);
%             for s=1:k
%                 [R_temp,x]=sortrows(R,s);
%                 for i=1:n
%                     %        i
%                     for j=i:n
%                         S(i,s);
%                         S(i,s)=S(i,s)+1/R_temp(j,s); % Savage Scores
%                     end
%                 end
%                 S(x,s)=S(:,s);
%             end
%             if k<3
%                 [rT]=corr(S);
%                 rT=rT(2)
%                 p=normpdf(sqrt(n-1)*rT);
%                 sign=p
%             else % for k>2
%                 a1=sum(S,2);
%                 S1=mean(max(S));
%                 a2=sum(a1.*a1);
%                 b=k;
%                 CT=((a2-n*b^2)/(b^2*(n-S1)))
%                 p=chi2pdf(b*(n-1)*CT,n-1);
%                 sign=p
%                 rT=CT;
%             end
%             
%             
%         end
    end
    
    
    methods
        
        function obj = set.numVP(obj,numVP)
            if isnumeric(numVP)
                obj.numVP= (numVP);
            else
                error(['NumVP should be numeric.'])
            end
        end
        
        
        
        
        function obj = set.vpWeights(obj,vpWeights)
            if isnumeric(vpWeights) && length(vpWeights) == get(obj, 'numVP')
                %if isnumeric(vpWeights)
                obj.vpWeights= (vpWeights);
            else
                error(['Length of vpWeights should be equal to numVP.'])
            end
        end
        
        
        
        
        
        function obj = set.whiskerPlot(obj,whiskerPlot)
            if isnumeric(whiskerPlot)
                obj.whiskerPlot= (whiskerPlot);
            else
                error(['Invalid whiskerPlotspecified for ',mfilename,', a number should be specified.'])
            end
        end
        
        
        function obj = set.confidenceInterval(obj,confidenceInterval)
            if isnumeric(confidenceInterval)
                obj.confidenceInterval = (confidenceInterval);
            else
                error(['Invalid confidence interval specified for ',mfilename,', a number 0 or 1, should be specified.'])
            end
        end
        function obj = set.sampleSize(obj,sampleSize)
            if isnumeric(sampleSize)
                obj.sampleSize = (sampleSize);
            else
                error(['Invalid nsample specified for ',mfilename,', should be > 0 integer, should be specified.'])
            end
        end
        function obj = set.cohortSize(obj,cohortSize)
            if isnumeric(cohortSize)
                obj.cohortSize = (cohortSize);
            else
                error(['Invalid cohort size specified for ',mfilename,', > 0 integer, should be specified.'])
            end
        end
        
        
        function obj = set.doseresponsetoPlot(obj,doseresponsetoPlot)
            if iscell(doseresponsetoPlot)
                obj.doseresponsetoPlot = (doseresponsetoPlot);
            else
                error(['Invalid dosestoplot specified for ',mfilename,', a cell array of variable ID strings should be specified.'])
            end
        end
        
        
        function obj = set.figstoPlot(obj,figstoPlot)
            if iscell(figstoPlot)
                obj.figstoPlot = (figstoPlot);
            else
                error(['Invalid figstoplot specified for ',mfilename,', a cell array of variable ID strings should be specified.'])
            end
        end
        
        
        function obj = set.analyzeElementResultIDs(obj,myAnalyzeElementResultIDs)
            if iscell(myAnalyzeElementResultIDs)
                obj.analyzeElementResultIDs = (myAnalyzeElementResultIDs);
            else
                error(['Invalid analyzeElementResultIDs specified for ',mfilename,', a cell array of variable ID strings should be specified.'])
            end
        end
        
        function obj = set.analyzeInterventionIDs(obj,myAnalyzeInterventionIDs)
            if iscell(myAnalyzeInterventionIDs)
                obj.analyzeInterventionIDs = (myAnalyzeInterventionIDs);
            else
                error(['Invalid analyzeInterventionIDs specified for ',mfilename,', a cell array of intervention ID strings should be specified.'])
            end
        end
        
        function obj = set.analyzeTimes(obj,myAnalyzeTime)
            failFlag = false;
            if isnumeric(myAnalyzeTime)
                obj.analyzeTimes = (myAnalyzeTime);
            else
                failFlag = true;
            end
            if failFlag
                error(['Invalid analyzeTimes specified for ',mfilename,'.'])
            end
        end
        
        function obj = set.poolClose(obj,myInput)
            if islogical(myInput)
                obj.poolClose = myInput;
            else
                error(['Property poolClose in ',mfilename,' must be logical.'])
            end
        end
        
        function obj = set.poolRestart(obj,myInput)
            if islogical(myInput)
                obj.poolRestart = myInput;
            else
                error(['Property poolRestart in ',mfilename,' must be logical.'])
            end
        end
        
        function obj = set.nWorkers(obj,myNWorkers)
            if (isnumeric(myNWorkers) == true)
                obj.nWorkers = myNWorkers;
            else
                error(['Invalid nWorkers value in ',mfilename,'.'])
            end
        end
        
        function obj = set.clusterID(obj,myClusterID)
            if ischar(myClusterID)
                obj.clusterID = myClusterID;
            else
                error(['Invalid clusterID in ',mfilename,'.'])
            end
        end
        
        function value = get(obj,propName)
            switch propName
                case 'analyzeElementResultIDs'
                    value = obj.analyzeElementResultIDs;
                case 'analyzeInterventionIDs'
                    value = obj.analyzeInterventionIDs;
                case 'analyzeTimes'
                    value = obj.analyzeTimes;
                case 'poolRestart'
                    value = obj.poolRestart;
                case 'poolClose'
                    value = obj.poolClose;
                case 'nWorkers'
                    value = obj.nWorkers;
                case 'clusterID'
                    value = obj.clusterID;
                case 'confidenceinterval'
                    value=obj.clusterID;
                    
                case 'nsample'
                    value=obj.clusterID;
                    
                case 'cohortsize'
                    value=obj.clusterID;
                    
                case 'dosestoplot'
                    value=obj.doseresponsetoPlot;
                    
                case 'figstoplot'
                    value=obj.figstoPlot;
                    
                case 'whiskerPlot'
                    value=obj.whiskerPlot;
                    
                case 'numVP'
                    value=obj.numVP;
                    
                case 'vpWeights'
                    value=obj.vpWeights;
                    
                    
                    
                    
                    
                    
                    
            end
        end
        
        function passCheck = verify(obj,myWorksheet)
            passCheck = true;
            myOutputVariables = obj.analyzeElementResultIDs;
            mySampleTimes = obj.analyzeTimes;
            myInterventionIDs = obj.analyzeInterventionIDs;
            
            if ~ischar(myOutputVariables) && ~iscellstr(myOutputVariables)
                warning(['OutputVariable is not a single string or a list of strings in ',mfilename,'.  Exiting.'])
                passCheck = false;
            end
            
            allElementResultIDs = myWorksheet.simProps.saveElementResultIDs;
            if sum(~ismember(myOutputVariables,allElementResultIDs)) > 0
                warning(['myWorkSheet is missing the desired OutputVariables in ',mfilename,'.  Exiting.'])
                continueFlag = false;
            end
            
            if sum(mySampleTimes >= 0) < length(mySampleTimes)
                warning(['mySampleTimes must be a vector of nonnegative values in ',mfilename,'.  Exiting.'])
                passCheck = false;
            end
            
            allSampleTimes = myWorksheet.simProps.sampleTimes;
            if sum(ismember(mySampleTimes,allSampleTimes)) < length(mySampleTimes)
                warning(['mySampleTimes is not a subset of SampleTimes in myWorkSheet in ',mfilename,'.  Exiting.'])
                passCheck = false;
            end
            
            allInterventionIDs = getInterventionIDs(myWorksheet);
            if sum(ismember(myInterventionIDs,allInterventionIDs)) < length(myInterventionIDs)
                warning(['myInterventionIDs is not a subset of the intervention IDs in myWorkSheet in ',mfilename,'.  Exiting.'])
                passCheck = false;
            end
            
            allInterventionIDs = getInterventionIDs(myWorksheet);
            allVPIDs = getVPIDs(myWorksheet);
            [dummy, nVPResults]= size(myWorksheet.results);
            if nVPResults ~= length(allVPIDs)
                warning(['Results for all VPs not available from worksheet in ',mfilename,'.  Exiting.'])
                passCheck = false;
            end
            
            %           if (obj.nBootstraps < 1) && ~(strcomp(subSampleSplitType,'none'))
            %               warning(['If splitting into subsamples in ',mfilename,', nBootstraps must be > 0.'])
            %               passCheck = false;
            %           end
            
            if passCheck
                for interventionCounter = 1 : length(myInterventionIDs)
                    myInterventionID = myInterventionIDs{interventionCounter};
                    
                    interventionIndex = find(ismember(allInterventionIDs, myInterventionID));
                    checkResults = myWorksheet.results(interventionIndex,:);
                    if sum(sum(arrayfun(@(i) isstruct(checkResults{i}),1:nVPResults)) ) < length(allVPIDs)
                        %failCheckVPs = ~(arrayfun(@(i) isstruct(checkResults{i}),1:nVPResults));
                        %failCheckVPIndices = find(failCheckVPs);
                        warning(['Results for all VPs not available from worksheet in ',mfilename,'.'])
                        passCheck = false;
                    else
                        for vpCounter = 1 : nVPResults
                            if sum(ismember(checkResults{1, vpCounter}.Names,obj.analyzeElementResultIDs)) < length(obj.analyzeElementResultIDs)
                                passCheck = false;
                            end
                        end
                        if ~passCheck
                            warning(['Not all analyzeElementResultIDs identified in the worksheet results provided to ',mfilename,'.'])
                        end
                    end
                end
            end
            if passCheck
                ntimefail = 0;
                for vpCounter = 1 : nVPResults
                    timeIndex = find(ismember(checkResults{1, vpCounter}.Names,'time'));
                    timeVals = checkResults{1, vpCounter}.Data(:,timeIndex);
                    % Might want to add a tolerance here... but
                    % MATLAB has been generally robust for small
                    % numerical issues so far
                    if sum(timeVals == obj.analyzeTimes) < 1
                        ntimefail = ntimefail+1;
                    end
                end
                if ntimefail > 0
                    passCheck = false;
                    warning(['Not all sample times align with the desired analyzeTime in the results provided to ',mfilename,'.'])
                    passCheck = false;
                end
            end
        end
        
        
        % The constructor method must have the same name as the class
        function obj = prccSensitivityOptions(myWorksheet)
            
            obj.analyzeElementResultIDs={};
            obj.analyzeInterventionIDs={};
            obj.analyzeTimes=0;
            obj.poolClose = true;
            obj.poolRestart = true;
            obj.clusterID = parallel.defaultClusterProfile;
            obj.nWorkers = nan;
            obj.confidenceInterval=0;
            obj.sampleSize=200;
            obj.cohortSize=200;
            obj.doseresponsetoPlot={};
            obj.figstoPlot={};
            obj.whiskerPlot=1;
            obj.numVP= length(myWorksheet.vpDef);
            obj.vpWeights= ones(1, obj.numVP)./sum(ones(1,obj.numVP));
            
            obj.totalbin=20;
            
            
        end
    end
    
    
end
```
