## Supplementary File 3 for "A surrogate workflow for virtual population development: designing an amenable proxy for a hypothesis-test inspired nonlinear goodness-of-fit": readAllResultStruct.m

```
function myWorksheet = readAllResultStruct(myWorksheet, myFileName, myVPColName, myInterventionColName, myTimeColName)
% This function takes a text file and reads results
% and over-writes the worksheet results structure.
% Note that you need to include the file extension in
% myFileName.
%
% ARGUMENTS
%  myWorksheet:            A worksheet; the results field
%                          will be overwritten.                    
%  myFileName:             The name of the file to read.
%                          It is assumed .csv implies a comma-separated
%                          value file.  Otherwise, tab-delimited
%                          is assumed
%  myInterventionColname:  Column header for the interventions
%  myVPColName:            Column header for the VPs
%  myTimeColName:          Column header for the time
%
% RETURNS
%  myWorksheet:            A worksheet with the overwritten result
%                          structure.  
%
continueFlag = true;

if nargin > 5
    warning(['Too many input arguments to ',mfilename,'. Require: myWorksheet, myFileName; optional: myInterventionColname, myVPColName, myTimeColName.  Exiting.'])
    continueFlag = false;
elseif nargin > 4
    continueFlag = true;    
elseif nargin > 3
    continueFlag = true;
    myVPColName = 'vpID';    
elseif nargin > 2
    continueFlag = true;
    myVPColName = 'vpID';
elseif nargin > 1
    continueFlag = true;
    myVPColName = 'vpID';    
    myInterventionColname = 'interventionID';        
elseif nargin < 2 
    warning(['Insufficient input arguments to ',mfilename,'. Require: myWorksheet, myFileName; optional: myInterventionColname, myVPColName, myTimeColName.  Exiting.'])
    continueFlag = false;   
end

if continueFlag
    if ~strcmp(class(myFileName),'char')
        warning(['Expecting a string for a filename in call to ',mfilename,'.  Exiting.'])
        continueFlag = false; 
    end    
end


if continueFlag
    tokens = regexp(myFileName,'(.*)\.(\w*)','tokens');
     if length(tokens) > 0
            noPathName = strsplit(tokens{1}{1},filesep);
            filePath = strjoin(noPathName(1:length(noPathName)-1),filesep);
            noPathName = noPathName{length(noPathName)};
            fileTypeExt = tokens{1}{2};
     else
            warning(['Cannot load file in ',mfilename,', file should have an extension.  Exiting.'])
            continueFlag = false;
     end
end

if continueFlag
    if length(filePath) < 1
        filePath=which([noPathName,'.',fileTypeExt]);
        filePath=filePath(1:end-(length([noPathName,'.',fileTypeExt])+1));
    end
end

if continueFlag
    if exist([filePath,filesep,noPathName,'.',fileTypeExt], 'file') ~= 2
        warning(['Cannot locate file in ',mfilename,'.  Exiting.'])
        continueFlag = false;
    end
end

if continueFlag
    % Assume 1 header line
    addpath(filePath);
    if strcmp(fileTypeExt,'csv') 
        delimiterIn = ',';
    else
        delimiterIn = '\t';
    end
    fID = fopen([noPathName,'.',fileTypeExt]);
    unparsedHeaders=fgetl(fID);
    allHeaders = strsplit(unparsedHeaders,delimiterIn);    
    fclose(fID);
    
    % Assume headers are in A.textdata{1,1}
    myVPIDIndex = find(ismember(allHeaders,myVPColName));
    if length(myVPIDIndex) ~= 1
        warning(['Unable to identify VP header in ',noPathName,'.',fileTypeExt,'.  Exiting.'])
        continueFlag = false;
    end
    myInterventionIndex = find(ismember(allHeaders,myInterventionColName));
    if length(myInterventionIndex) ~= 1
        warning(['Unable to identify intervention header in ',noPathName,'.',fileTypeExt,'.  Exiting.'])
        continueFlag = false;
    end   
    myTimeIndex = find(ismember(allHeaders,myTimeColName));
    if length(myTimeIndex) ~= 1
        warning(['Unable to identify time header in ',noPathName,'.',fileTypeExt,'.  Exiting.'])
        continueFlag = false;
    end      
    
    if continueFlag
        opts = detectImportOptions([noPathName,'.',fileTypeExt]);
        if ~(strcmp(opts.VariableNames{myVPIDIndex},myVPColName) & strcmp(opts.VariableNames{myInterventionIndex},myInterventionColName))
            warning(['Unable to identify VP and intervention header in ',noPathName,'.',fileTypeExt,'.  Exiting.'])
            continueFlag = false;            
        end
    end
    if continueFlag
        opts.VariableTypes{myVPIDIndex} = 'char';
        opts.VariableTypes{myInterventionIndex} = 'char';
        T = readtable([noPathName,'.',fileTypeExt],opts);
        % We will convert to a cell array of matrices that we
        % can read into 
        tVarNames = T.Properties.VariableNames;
        if (length(tVarNames) ~= length(allHeaders))
            warning(['Unable to read ',noPathName,'.',fileTypeExt,' correctly.  Exiting.'])
            continueFlag = false;
        end
    end
    
    if continueFlag        
        allVPIDs = T{:,myVPIDIndex};
        allVPIDs = unique(allVPIDs,'stable');
        allInterventionIDs = T{:,myInterventionIndex};
        allInterventionIDs = unique(allInterventionIDs);    
        nNewVPs = length(allVPIDs);
        nNewInterventions = length(allInterventionIDs);
        newResults = cell(nNewInterventions,nNewVPs);
        dataColIndices = [myVPIDIndex,myInterventionIndex,myTimeIndex];
        dataColIndices = [setdiff(1:length(allHeaders), dataColIndices)];
        dataNames = ['time', allHeaders(dataColIndices)];
        dataColIndices = [myTimeIndex,dataColIndices];
        testData = T{:,dataColIndices};
        if ~strcmp(class(testData),'double')
            warning(['Results in ',noPathName,'.',fileTypeExt,' not all numeric.  Exiting.'])
            continueFlag = false;
        end
        
    end
    
    if continueFlag  
        allWshVPIDs = getVPIDs(myWorksheet);
        allWshInterventionIDs = getInterventionIDs(myWorksheet);
        if ((sum(ismember(allVPIDs,allWshVPIDs)) ~= sum(ismember(allInterventionIDs,allWshInterventionIDs))) | (sum(ismember(allVPIDs,allWshVPIDs)) == 0))
            warning(['VPIDs in ',noPathName,'.',fileTypeExt,' do not all match supplied worksheet.  Overwriting worksheet VPs to facilitate analysis of data in results.'])
            if length(allWshVPIDs) >= 1
                newVariants = repmat(myWorksheet.vpDef{1}.variants,length(allVPIDs));
                allCoefficients = getVPCoeffs(myWorksheet);
                allCoefficients = allCoefficients(:,1);
                newCoefficients = repmat(allCoefficients,length(allVPIDs));
                myWorksheet = removeVPs(myWorksheet, allWshVPIDs);
                myWorksheet = createVPs(myWorksheet, allVPIDs, newVariants, newCoefficients);
                allWshVPIDs = allVPIDs;
            else
                allWshVPIDs = allVPIDs;
                myWorksheet = createVPs(myWorksheet, allWshVPIDs, repmat(cell(1,1),1,length(allWshVPIDs)));
            end
        end
        if ((sum(ismember(allInterventionIDs,allWshInterventionIDs)) ~= sum(ismember(allInterventionIDs,allWshInterventionIDs))) | (sum(ismember(allInterventionIDs,allWshInterventionIDs)) == 0))
            warning(['InterventionIDs in ',noPathName,'.',fileTypeExt,' do not all match supplied worksheet.  Overwriting worksheet interventions to facilitate analysis of data in results.'])
            myWorksheet = removeInterventions(myWorksheet, allWshInterventionIDs);
            for interventionCounter = 1 : length(allInterventionIDs)
                myWorksheet = createIntervention(myWorksheet, allInterventionIDs{interventionCounter}, cell(0,2));
            end
            allWshInterventionIDs = allInterventionIDs;
        end
    end
    
    for interventionCounter = 1 : nNewInterventions
        interventionSelect(:,interventionCounter) = ismember(T{:,myInterventionIndex},allWshInterventionIDs{interventionCounter});
    end
    
    if continueFlag
        nSimulations = nNewVPs*nNewInterventions;
        newResults = cell(1,nSimulations);
        if ~isempty(gcp('nocreate'))
            delete(gcp);
        end
        parpool;        
        parfor simulationCounter = 1 : nSimulations
            simStruct = struct();
            vpCounter = ceil(simulationCounter/nNewInterventions);
            interventionCounter = simulationCounter-(vpCounter-1)*nNewInterventions;
            vpSelect = ismember(T{:,myVPIDIndex},allWshVPIDs{vpCounter});
            curRows = find( vpSelect &  interventionSelect(:,interventionCounter));
            simStruct.Names = dataNames;
            simStruct.Data = T{curRows,dataColIndices};                
            newResults{simulationCounter} = simStruct;
        end
        newResults = reshape(newResults,[nNewInterventions,nNewVPs]);
        myWorksheet.results = newResults;
        delete(gcp)
    end
end
end
    
    
```
