## Supplementary File 3 for "A surrogate workflow for virtual population development: designing an amenable proxy for a hypothesis-test inspired nonlinear goodness-of-fit": removeVPopInterventions.m

```
function myVPop = removeVPopInterventions(myVPop, myInterventionIDs)
% This function removes interventions from a VPop or VPopRECIST
% object.  It removes associated rows from the tables.
% Then it removes associated simData as well.
%
% ARGUMENTS
%  myVPop
%  myInterventionIDs: A list of interventions.
%
% RETURNS
%  newVPop:         The new VPop with the interventions removed.
%
% TODO: ADD MORE CHECKING OF INPUTS
% 

% Drop unneeded rows from simData
if ~isempty(myVPop.simData)
    interventionIndex = ismember(myVPop.simData.rowInfoNames,'interventionID');
	keepRows = ~ismember(myVPop.simData.rowInfo(:,interventionIndex),myInterventionIDs);
	myVPop.simData.Data = myVPop.simData.Data(keepRows,:);
    myVPop.simData.rowInfo = myVPop.simData.rowInfo(keepRows,:)
end
mnsdDataFlag = false;
binDataFlag = false;
distDataFlag = false;
distData2DFlag = false;	
corDataFlag = false;
brDataFlag = false;	
rDataFlag = false;	

subpopInterventions = cell(1,0);
for cellCounter = 1:numel(myVPop.subpopTable{:,'interventionID'})
	subpopInterventions = [subpopInterventions, myVPop.subpopTable{cellCounter,'interventionID'}{1}];
end

if sum(ismember(myInterventionIDs, subpopInterventions)) > 0
    myInterventionIDs = setdiff(myInterventionIDs,subpopInterventions); % make sure interventions used to specify subpopulations are not removed.
% 	warning(['Unable to remove specified interventions in ',mfilename,'.  Removing interventions used to specify subpopulations is not currently supported.  Exiting...'])
end

% Next scan through the VPop 
myMnSDTable = myVPop.mnSDTable;
myBinTable = myVPop.binTable;
myDistTable = myVPop.distTable;
myDistTable2D = myVPop.distTable2D;
myCorTable = myVPop.corTable;
if isa(myVPop,'VPopRECIST')
	myBRTableRECIST = myVPop.brTableRECIST;
	myRTableRECIST = myVPop.rTableRECIST;
else
	myBRTableRECIST = [];
	myRTableRECIST = [];
end

if ~isempty(myMnSDTable)
	[nRows, nCols] = size(myMnSDTable);
	if nRows > 0
		keepRows = ~ismember(myMnSDTable.('interventionID'),myInterventionIDs);
        if sum(keepRows) > 0
            myMnSDTable = myMnSDTable(keepRows,:);
            mnsdDataFlag = true;
        else
            myMnSDTable = [];
        end 
        myVPop.mnSDTable = myMnSDTable;        
	end
end

if ~isempty(myBinTable)
	[nRows, nCols] = size(myBinTable);
	if nRows > 0
		keepRows = ~ismember(myBinTable.('interventionID'),myInterventionIDs);
        if sum(keepRows) > 0
            myBinTable = myBinTable(keepRows,:);
            binDataFlag = true;
        else
            myBinTable = [];
        end
        myVPop.binTable = myBinTable;
	end
end

if ~isempty(myDistTable)
	[nRows, nCols] = size(myDistTable);
	if nRows > 0
		keepRows = ~ismember(myDistTable.('interventionID'),myInterventionIDs);
        if sum(keepRows) > 0
            myDistTable = myDistTable(keepRows,:);
            distDataFlag = true;
        else
            myDistTable = [];
        end
        myVPop.distTable = myDistTable;
	end
end

if ~isempty(myDistTable2D)
	[nRows, nCols] = size(myDistTable2D);
	if nRows > 0
		keepRows = ~(ismember(myDistTable2D.('interventionID1'),myInterventionIDs) | ismember(myDistTable2D.('interventionID2'),myInterventionIDs));
        if sum(keepRows) > 0
            myDistTable2D = myDistTable2D(keepRows,:);
            distData2DFlag = true;
        else
            myDistTable2D = [];
        end
        myVPop.distTable2D = myDistTable2D;
	end
end

if ~isempty(myCorTable)
	[nRows, nCols] = size(myCorTable);
	if nRows > 0
		keepRows = ~(ismember(myCorTable.('interventionID1'),myInterventionIDs) | ismember(myCorTable.('interventionID2'),myInterventionIDs));
		if sum(keepRows) > 0
            myCorTable = myCorTable(keepRows,:);
            corDataFlag = true;
        else
            myCorTable = [];
        end
        myVPop.corTable = myCorTable;
	end
end

if ~isempty(myBRTableRECIST)
	[nRows, nCols] = size(myBRTableRECIST);
	if nRows > 0
		keepRows = ~ismember(myBRTableRECIST.('interventionID'),myInterventionIDs);
        if sum(keepRows) > 0
            myBRTableRECIST = myBRTableRECIST(keepRows,:);
            brDataFlag = true;		
        else
            myBRTableRECIST = [];
        end
        myVPop.brTableRECIST = myBRTableRECIST;  
        myVPop.simData.brData = myVPop.simData.brData(keepRows,:);	
        myVPop.simData.brRowInfo = myVPop.simData.brRowInfo(keepRows,:);        
	end
end

if ~isempty(myRTableRECIST)
	[nRows, nCols] = size(myRTableRECIST);
	if nRows > 0
		keepRows = ~ismember(myRTableRECIST.('interventionID'),myInterventionIDs);
        if sum(keepRows) > 0
            myRTableRECIST = myRTableRECIST(keepRows,:);
            rDataFlag = true;
        else
            myRTableRECIST = [];
        end
        myVPop.rTableRECIST = myRTableRECIST;
		myVPop.simData.rData = myVPop.simData.rData(keepRows,:);
		myVPop.simData.rRowInfo = myVPop.simData.rRowInfo(keepRows,:);
	end
end

% It would be more efficient to do
% this as we remove the rows.  But this
% checking is conceptually faster to write without
% much performance penalty.
if ~isempty(myVPop.simData)
	             
	[nEntries, ~] = size(myVPop.simData.Data);
    mnSDRows = nan(nEntries,1);
    binRows = nan(nEntries,1);
    distRows = nan(nEntries,1);
	distRows2D = cell(nEntries,2);
	corRows = cell(nEntries,2);
    rRows = nan(nEntries,1);
    brRows = nan(nEntries,1);    
	
	expVarIndex = ismember(myVPop.simData.rowInfoNames,'expVarID');
	elementIDIndex = ismember(myVPop.simData.rowInfoNames,'elementID');
	elementTypeIndex = ismember(myVPop.simData.rowInfoNames,'elementType');
	expTimeIndex = ismember(myVPop.simData.rowInfoNames,'time');
	
	for rowCounter = 1 : nEntries
			
		interventionID = myVPop.simData.rowInfo{rowCounter,interventionIndex};
		elementID = myVPop.simData.rowInfo{rowCounter,elementIDIndex};
		elementType = myVPop.simData.rowInfo{rowCounter,elementTypeIndex};
		expTime = myVPop.simData.rowInfo{rowCounter,expTimeIndex};
		
		% To avoid re-searching for the right rows on every
		% iteration mapel, we provide the indices here 
		if mnsdDataFlag
			temp = find((ismember(myMnSDTable{:,'interventionID'},interventionID)) & ((myMnSDTable{:,'time'})==expTime) & (ismember(myMnSDTable{:,'elementID'},elementID)) & (ismember(myMnSDTable{:,'elementType'},elementType)));
			if ~isempty(temp)
				mnSDRows(rowCounter) = temp;
			end
		end
		if binDataFlag
			temp = find((ismember(myBinTable{:,'interventionID'},interventionID)) & ((myBinTable{:,'time'})==expTime) & (ismember(myBinTable{:,'elementID'},elementID)) & (ismember(myBinTable{:,'elementType'},elementType)));
			if ~isempty(temp)
				binRows(rowCounter) = temp;
			end
		end
		if distDataFlag
			temp = find((ismember(myDistTable{:,'interventionID'},interventionID)) & ((myDistTable{:,'time'})==expTime) & (ismember(myDistTable{:,'elementID'},elementID)) & (ismember(myDistTable{:,'elementType'},elementType)));
			if ~isempty(temp)
				distRows(rowCounter) = temp;
			end
		end                    
		if distData2DFlag
			% One row of source data may be involved in
			% multiple 2D distributions
			temp = find((ismember(myDistTable2D{:,'interventionID1'},interventionID)) & ((myDistTable2D{:,'time1'})==expTime) & (ismember(myDistTable2D{:,'elementID1'},elementID)) & (ismember(myDistTable2D{:,'elementType1'},elementType)));
			if ~isempty(temp)
				distRows2D{rowCounter,1} = temp;
			end
			temp = find((ismember(myDistTable2D{:,'interventionID2'},interventionID)) & ((myDistTable2D{:,'time2'})==expTime) & (ismember(myDistTable2D{:,'elementID2'},elementID)) & (ismember(myDistTable2D{:,'elementType2'},elementType)));
			if ~isempty(temp)
				distRows2D{rowCounter,2} = temp;
			end						
		end
		if corDataFlag
			% One row of source data may be involved in
			% multiple 2D distributions
			temp = find((ismember(myCorTable{:,'interventionID1'},interventionID)) & ((myCorTable{:,'time1'})==expTime) & (ismember(myCorTable{:,'elementID1'},elementID)) & (ismember(myCorTable{:,'elementType1'},elementType)));
			if ~isempty(temp)
				corRows{rowCounter,1} = temp;
			end
			temp = find((ismember(myCorTable{:,'interventionID2'},interventionID)) & ((myCorTable{:,'time2'})==expTime) & (ismember(myCorTable{:,'elementID2'},elementID)) & (ismember(myCorTable{:,'elementType2'},elementType)));
			if ~isempty(temp)
				corRows{rowCounter,2} = temp;
			end						
		end 
		if brDataFlag
			% One row of source data may be involved in
			% multiple 2D distributions
            temp = find((ismember(myBRTableRECIST{:,'interventionID'},interventionID)) & ((myBRTableRECIST{:,'time'})==expTime) );
			if ~isempty(temp)
				brRows(rowCounter) = temp;		
			end				
		end
 		if rDataFlag
			% One row of source data may be involved in
			% multiple 2D distributions
            temp = find((ismember(myRTableRECIST{:,'interventionID'},interventionID)) & ((myRTableRECIST{:,'time'})==expTime) );
			if ~isempty(temp)
				rRows(rowCounter) = temp;		
			end				
		end 
    end
	myVPop.simData.binRows = binRows;
	myVPop.simData.mnSDRows = mnSDRows;
	myVPop.simData.distRows = distRows;
	myVPop.simData.distRows2D = distRows2D;
	myVPop.simData.corRows = corRows;
	if brDataFlag
		myVPop.simData.brRows = brRows;
    elseif isa(myVPop, 'VPopRECIST')
        myVPop.simData.brRows = [];
    end
	if rDataFlag
		myVPop.simData.rRows = rRows;
    elseif isa(myVPop, 'VPopRECIST')
        myVPop.simData.rRows = [];
    end
	
	% Next update the individual GOF statistics
    myVPop = myVPop.addPredTableVals();
	myVPop = evaluateGOF(myVPop);

end
end
```
