## Supplementary File 3 for "A surrogate workflow for virtual population development: designing an amenable proxy for a hypothesis-test inspired nonlinear goodness-of-fit": runPRCCSensitivity.m

```
function [myCorrCoefs] = runPRCCSensitivity(myWorksheet, myPRCCSensitivityOptions, w)

%clear finalf

% Here, calculate Partial Rank Correlation Coefficients
%
% ARGUMENTS:
%  myWorksheet              A worksheet data structure with VPs and
%                            coefficients.
%  myPRCCSensitivityOptions A prccSensitivityOptions object.
%
% RETURNS:
%  myCorrCoeffs             cell array (nIntervention x nOutputVariables)
%                           cell.Names: names of axis definitions (Params)
%                           cell.Times: provided Sample Times (Times)
%                           cell.Pearson: matrix of Pearson correlation coefficients (nTimes x nParams)
%                           cell.Spearman: matrix of Spearman correlation coefficients (nTimes x nParams)
%                           cell.PRCC: matrix of partial rank correlation coefficients (nTimes x nParams)
%                           cell.interventionID: string with current interventionID
%                           cell.outputVariable: string with output variable
%
continueFlag = true;
if nargin > 5 % Come back here modify
    warning(['Too many input arguments provided to ',mfilename,'.  Expecting myWorksheet, myOutputVariables, mySampleTimes.  Exiting.'])
    continueFlag = false;
elseif nargin > 1
    continueFlag = true;
elseif nargin > 0
    myPRCCSensitivityOptions = prccSensitivityOptions;
    myPRCCSensitivityOptions.analyzeElementResultIDs = myWorksheet.simProps.saveElementResultIDs;
    myPRCCSensitivityOptions.analyzeInterventionIDs = getInterventionIDs(myWorksheet);
    myPRCCSensitivityOptions.analyzeTimes = myWorksheet.simProps.sampleTimes;
    continueFlag = true;
else
    warning(['Too few input arguments provided to ',mfilename,'.  Expecting myWorksheet, and optionally myPRCCSensitivityOptions.  Exiting.'])
    continueFlag = false;
end

if continueFlag
    continueFlag = myPRCCSensitivityOptions.verify(myWorksheet);
    if ~(continueFlag)
        warning(['Please correct the options provided to ',mfilename, '.'])
    else
        myOutputVariables = myPRCCSensitivityOptions.analyzeElementResultIDs;
        mySampleTimes = myPRCCSensitivityOptions.analyzeTimes;
        myInterventionIDs = myPRCCSensitivityOptions.analyzeInterventionIDs;
    end
end

if continueFlag
    % get parameter samples from Sobol for all VPs and all axes
    myVPCoeffs = getVPCoeffs(myWorksheet); % nParams x nVPs
    myParams = myVPCoeffs'; % nVPs x nParams
    [nVPs,nParams] = size(myParams);
    if nVPs < (nParams + 1)
        warning(['Number of Virtual Patients is smaller than the number of parameter axes in ',mfilename,' ... not a good idea!'])
    end
end

myCorrCoefs = cell(1,0);

if continueFlag
    allElementResultIDs = myWorksheet.simProps.saveElementResultIDs;
    allSampleTimes = myWorksheet.simProps.sampleTimes;
    allInterventionIDs = getInterventionIDs(myWorksheet); % Myintervations PRCC handles this.
    % find indices for myOutputVariable and mySampleTimes
    [~,myOutputVariableIndex] = ismember(myOutputVariables,allElementResultIDs);
    [~,mySampleTimesIndices] = ismember(mySampleTimes,allSampleTimes);
    
    %mySampleTimes= myPRCCSensitivityOptions.analyzeElementResultIDs = myWorksheet.simProps.saveElementResultIDs;
    mySampleTimes= myPRCCSensitivityOptions.analyzeTimes;
    myPRCCSensitivityOptions.analyzeInterventionIDs = getInterventionIDs(myWorksheet);
    
    %myOutputVariables= myPRCCSensitivityOptions.analyzeTimes = myWorksheet.simProps.sampleTimes;
    
    
    % get names of all mechanistic axes (Parameters)
    myAxisDefs = getAxisDefIDs(myWorksheet)';
    % define number of Interventions, SampleTimes and OutputVariables
    nInterventionIDs = length(myInterventionIDs);
    nTimes = length(mySampleTimes);
    nOutputs = length(myOutputVariables);
    
    % define output cell array
    myCorrCoefs = cell(nInterventionIDs, nOutputs);
    
    % As a precaution, restart any existing parallel
    % pools
    %     if myPRCCSensitivityOptions.poolRestart
    %         if ~isempty(gcp('nocreate'))
    %             delete(gcp);
    %         end
    %     end
    
    %     if isempty(gcp('nocreate'))
    %         % First check the default number of workers, if needed
    %         myPRCCSensitivityOptions = checkNWorkers(myPRCCSensitivityOptions);
    %         myPool = parpool(myPRCCSensitivityOptions.clusterID,myPRCCSensitivityOptions.nWorkers,'SpmdEnabled',false);
    %     end
    mySampleTimes= myPRCCSensitivityOptions.analyzeTimes;
    
    for n=1:nOutputs
        
        for i=1:nInterventionIDs
            
            myCorrCoefs{i,n}.Names = myAxisDefs;
            myCorrCoefs{i,n}.Times = mySampleTimes;
            myCorrCoefs{i,n}.interventionID = myInterventionIDs{i};
            myCorrCoefs{i,n}.outputVariable = allElementResultIDs{myOutputVariableIndex(n)};
            
            
            % curResult.Data (nTimes x nVPs) contains times series for all VPs
            % for specified Intervention and OutputVariable
           
            curResult = getResultOutputforIntervention(myWorksheet,myInterventionIDs{i},allElementResultIDs{myOutputVariableIndex(n)});
            
           
            curData = curResult.Data(mySampleTimesIndices,2:end);
            myData = curData'; % nVPs x nTimes
            
            myVPCoeffs = getVPCoeffs(myWorksheet); % nParams x nVPs
            myParams = myVPCoeffs'; % nVPs x nParams
            %curResultm = getResultOutputforIntervention(myWorksheetm,myInterventionIDs{i},allElementResultIDs{myOutputVariableIndex(n)});
            %curDatam = curResultm.Data(mySampleTimesIndices,2:end);
            %myDatam = curDatam'; % nVPs x nTimes
            
            %myVPCoeffsm = getVPCoeffs(myWorksheetm); % nParams x nVPs
            %myParamsm = myVPCoeffsm'; % nVPs x nParams
            
            
            
            
            
            % % find indices for myOutputVariable and mySampleTimes
            [~,myOutputVariableIndex] = ismember(myOutputVariables,allElementResultIDs);
            [~,mySampleTimesIndices] = ismember(mySampleTimes,allSampleTimes);
            
            %             %elementsids_int=myPRCCSensitivityOptions.analyzeElementResultIDs
            %             intervation= i;
            %             if doseresponse==1
            %                 [finalf]= runprccsensitivityc.bingen(myData,myParams ,intervation,  myWorksheet);
            %             else
            %                 finalf=[];
            %             end
            
            
            % PearsonCorrMatrix (nParams x nTimes) contains Pearson correlation
            % coefficient between each axis and the OutputVariable at specified time
            
            PearsonCorrMatrix = corr(myParams,myData);
            myCorrCoefs{i,n}.Pearson = PearsonCorrMatrix';
            
            % compute Spearman correlationusing rank-ordered
            % parameters and outputs
            
            
            myDataOrdered = tiedrank(myData);
            myParamsOrdered = tiedrank(myParams);
        
            
            
            % SpearmanCorrMatrix (nParams x nTimes) contains Spearman correlation
            % coefficient between each axis and the OutputVariable at specified time
            
            SpearmanCorrMatrix = corr(myParamsOrdered,myDataOrdered);
            myCorrCoefs{i,n}.Spearman = SpearmanCorrMatrix';
            
            % compute partial rank correlation matrix using rank-ordered data
            % and parameters in myDataOrdered and myParamsOrdered
            % PRCCMatrix: nParams x nTimes
            PRCCMatrix = zeros(size(myParamsOrdered,2),size(myDataOrdered,2));
           
            
            
            PRCCMatrixabs = zeros(size(myParamsOrdered,2),size(myDataOrdered,2));
            PRCCMatrixw = zeros(size(myParamsOrdered,2),size(myDataOrdered,2));
            PRCCMatrixabsw= zeros(size(myParamsOrdered,2),size(myDataOrdered,2));
            pvalue = zeros(size(myParamsOrdered,2),size(myDataOrdered,2));
            pvalueabs = zeros(size(myParamsOrdered,2),size(myDataOrdered,2));
            pvaluew = zeros(size(myParamsOrdered,2),size(myDataOrdered,2));
            pvaluewabs= zeros(size(myParamsOrdered,2),size(myDataOrdered,2));
            PRCCMatrixregr = PRCCMatrixabs;
            pvalueregr = pvalueabs;
            PRCCMatrixwregr = PRCCMatrixabs;
            pvaluewregr = pvalueabs;

            
            for j=1:nTimes % for all time points
                curDataOrderedCol = myDataOrdered(:,j); % nVPs x 1
                
                for k=1:nParams % for all parameter axes
                    curParamsOrderedCol = myParamsOrdered(:,k); % nVPs x 1
                    curParamsOrderedReduced = myParamsOrdered;
                    % eliminate kth VP's parameter ...
                    curParamsOrderedReduced(:,k) = [];
                    % ... and add constant vector to account for constant
                    % offset in the regression: p_k = beta0 + beta_i p_i (i~=k)
                    curParamsOrderedReduced = [ones(size(curParamsOrderedReduced,1),1), ...
                        curParamsOrderedReduced]; % nVPs x nParams
                    
                    % find linear regression coefficients for curDataOrderedCol
                    % and curParamsOrderedCol with respect to curParamsOrderedReduced
                    
                    betaParams = curParamsOrderedReduced\curParamsOrderedCol; % nParams x 1
                    betaData = curParamsOrderedReduced\curDataOrderedCol; % nParams x 1
                    
                    curParamsOrderedReg = curParamsOrderedReduced*betaParams; % nVPs x 1
                    curDataOrderedReg = curParamsOrderedReduced*betaData; % nVPs x 1
                    
                    % compute residuals
                    myParamsOrderedRes = curParamsOrderedCol - curParamsOrderedReg;
                    myDataOrderedRes = curDataOrderedCol - curDataOrderedReg;
                    
                    % compute correlation coefficients between rank-ordered
                    % residuals

                    wequal = (ones(1,length(myDataOrderedRes))./length(myDataOrderedRes))';
                    dparams=length(myWorksheet.axisProps.axisDef)-1;
                    nSample=length(myDataOrderedRes);
                    
                    %PRCC matrix abs
                    PRCCMatrixabs(k,j)= myPRCCSensitivityOptions.weightedCorr(abs(myParamsOrderedRes),abs(myDataOrderedRes), wequal, false);
                    %PRCCMatrixabs(k,j)= myPRCCSensitivityOptions.weightedCorr(abs(myParamsOrderedRes),abs(myDataOrderedRes), wequal, false);
                    ccvalue= PRCCMatrixabs(k,j);
                    pvalueabs(k,j)= myPRCCSensitivityOptions.pvaluePRCC(nSample,dparams, ccvalue, true);
                    %pvalueabs(k,j)= myPRCCSensitivityOptions.pvaluePRCC(nSample,dparams, ccvalue, false);
                    
                    %PRCC matrix abs weighted
                    % weighted linear regression + weighted correlation
                    W = diag(w);
                    betaParams = (W*curParamsOrderedReduced)\(w.*curParamsOrderedCol); % nParams x 1
                    betaData = (W*curParamsOrderedReduced)\(w.*curDataOrderedCol); % nParams x 1
                    curParamsOrderedRegw = curParamsOrderedReduced*betaParams; % nVPs x 1
                    curDataOrderedRegw = curParamsOrderedReduced*betaData; % nVPs x 1
                    % compute residuals
                    myParamsOrderedResw = curParamsOrderedCol - curParamsOrderedRegw;
                    myDataOrderedResw = curDataOrderedCol - curDataOrderedRegw;
                    PRCCMatrixabsw(k,j)= myPRCCSensitivityOptions.weightedCorr(abs(myParamsOrderedResw), abs(myDataOrderedResw), w, false);
                    %PRCCMatrixabsw(k,j)= myPRCCSensitivityOptions.weightedCorr(abs(myParamsOrderedRes), abs(myDataOrderedRes), w, false);
                    ccvalue=   PRCCMatrixabsw(k,j);
                    pvaluewabs(k,j)= myPRCCSensitivityOptions.pvaluePRCC(nSample,dparams, ccvalue, true);
                    %pvaluewabs(k,j)= myPRCCSensitivityOptions.pvaluePRCC(nSample,dparams, ccvalue, false);
                    
                    %PRCC matrix
                    PRCCMatrix(k,j) = corr(myParamsOrderedRes,myDataOrderedRes);
                    ccvalue=   PRCCMatrix(k,j);
                    pvalue(k,j)= myPRCCSensitivityOptions.pvaluePRCC(nSample,dparams, ccvalue, false);

                    %PRCC matrix w
                    % weighted linear regression + weighted correlation
                    PRCCMatrixw(k,j)= myPRCCSensitivityOptions.weightedCorr(myParamsOrderedResw, myDataOrderedResw, w, false);
                    ccvalue=   PRCCMatrixw(k,j);
                    pvaluew(k,j)= myPRCCSensitivityOptions.pvaluePRCC(nSample,dparams, ccvalue, false);
                    
                    
                    % nonlinear regression-based PRCC 
                    x = myParamsOrderedRes;
                    y = myDataOrderedRes;
                    modelfun = @(b,x)b(1) + b(2)*x + b(3)*x.^2; % not sure if we need to go for higher degree, so far a simple quadratic polynomial seems enough
                    beta0 = [0 1 1];
                    mdl = fitnlm(x,y,modelfun,beta0);                    
                    PRCCMatrixregr(k,j)= sqrt(mdl.Rsquared.Ordinary);
                    ccvalue= PRCCMatrixregr(k,j);
                    pvalueregr(k,j)= myPRCCSensitivityOptions.pvaluePRCC(nSample,dparams, ccvalue, true);
                    
                    
                    % nonlinear regression-based weighted PRCC 
                    x = myParamsOrderedResw;
                    y = myDataOrderedResw;
                    modelfun = @(b,x)b(1) + b(2)*x + b(3)*x.^2;
                    beta0 = [0 1 1];
                    mdl = fitnlm(x,y,modelfun,beta0,'Weights',w);                    
                    PRCCMatrixwregr(k,j)= sqrt(mdl.Rsquared.Ordinary);
                    ccvalue= PRCCMatrixwregr(k,j);
                    pvaluewregr(k,j)= myPRCCSensitivityOptions.pvaluePRCC(nSample,dparams, ccvalue, true);
                end
            end

            myCorrCoefs{i,n}.PRCC = PRCCMatrix';
            myCorrCoefs{i,n}.pvalue=pvalue;
            myCorrCoefs{i,n}.PRCCabs = PRCCMatrixabs';
            myCorrCoefs{i,n}.pvalueabs= pvalueabs;
            myCorrCoefs{i,n}.PRCCw = PRCCMatrixw';
            myCorrCoefs{i,n}.pvaluew=pvaluew;
            myCorrCoefs{i,n}.PRCCwabs=    PRCCMatrixabsw';
            myCorrCoefs{i,n}.pvaluewabs= pvaluewabs;

            myCorrCoefs{i,n}.PRCCregr = PRCCMatrixregr';
            myCorrCoefs{i,n}.pvalueregr= pvalueregr;
            myCorrCoefs{i,n}.PRCCwregr = PRCCMatrixwregr';
            myCorrCoefs{i,n}.pvaluewregr= pvaluewregr;

        end
    end

    
    %     if myPRCCSensitivityOptions.poolClose
    %         if ~isempty(gcp('nocreate'))
    %             delete(gcp);
    %         end
    %     end
    
else
    warning(['Exiting ',mfilename,'.'])
end
end
        
```
