## Supplementary File 3 for "A surrogate workflow for virtual population development: designing an amenable proxy for a hypothesis-test inspired nonlinear goodness-of-fit": simulateWorksheetIndDoses.m

```
function myWorksheet = simulateWorksheetIndDoses(myWorksheet, mySimulateOptions)
% This function "crosses" the VPs in a worksheet plus alterations from 
% applied axes with the interventions
% defined int the worksheet and the runs each simulation
% Simulations are run in parallel, if PCT is available.
%
% This is a special variation on simulateWorksheet to allow for the
% specification of individual doses for each VP.  As such, it
% assumes specific modifications have been to the worksheet that
% aren't explicitly supported by the toolbox functions.
%
% ARGUMENTS
% myWorksheet: A worksheet to simulate, with special modifications.
%              This function will look for the field:
%              'vpDoseModifications'
%              Within the each intervention.
%              The vpDoseModification field should contain a 1xnVP cell
%              array, and each entry in the cell array should contain
%              a nDoseObject x 1 cell array of structured arrays,
%              each with fields: 'Name', 'Amount', 'Interval', 'Rate',
%              and 'RepeatCount'.
%              so to reference 1 dose modification that is:
%              doseStructuredArray = myWorksheet.interventions{interventionCounter}.('vpDoseModifications'){1,vpCounter}{doseStructCounter}
%              Also note this function therefore ignores doses specified in
%              myWorksheet.interventions{interventionCounter}.('definition')
%              
% mySimulateOptions: an instance of a simulateOptions object
% 
% RETURNS
% myWorksheet: the original worksheet plus
%              the simulation results

flagContinue = true;

% First check input arguments
if nargin > 2
    warning(['Too many input arguments to',mfilename,'. Require: myWorksheet; Optional: simulateOptions.'])
    flagContinue = false;
elseif nargin > 1
    flagContinue = true;     
elseif nargin > 0
    mySimulateOptions = simulateOptions;   
    flagContinue = true; 
else
    warning(['Insufficient input arguments to ',mfilename,'. Requires at least: myWorksheet; Optional: simulateOptions.'])
    flagContinue = false;    
end

% We don't support optimization in this version of simulate
if flagContinue
    if sum(ismember({'none'},mySimulateOptions.optimizeType)) < 1
        warning(['Optimization methods are not currently supported in ',mfilename,', only simulation.'])
        flagContinue = false;
    end
end

if flagContinue
    passCheck = mySimulateOptions.verify(myWorksheet);
    if ~passCheck
        warning(['Specified simulation options for ',mfilename,' not valid.'])
        flagContinue = false;        
    end
end

if flagContinue
    % We will assume compiled models are still valid.
    if ~(isequal(class(myWorksheet.compiled.model),'SimBiology.export.Model'))
        warning(['No exported model associated with myWorksheet prior to ',mfilename,', exporting and accelerating.'])
        myWorksheet = compileModel(myWorksheet,true);
        if ~(isequal(class(myWorksheet.compiled.model),'SimBiology.export.Model'))
             warning(['Unable to compile (export & accelerate) model associated with myWorksheet in ',mfilename,'.'])
             flagContinue = false;
        end
    elseif ~myWorksheet.compiled.model.isAccelerated
        warning(['No accelerated model associated with myWorksheet prior to ',mfilename,', accelerating.'])        
        myWorksheet = compileModel(myWorksheet,true);
        if ~(isequal(class(myWorksheet.compiled.model),'SimBiology.export.Model'))
             warning(['Unable to compile (export & accelerate) model associated with myWorksheet in ',mfilename,'.'])
             flagContinue = false;
        end        
    end
end


% Now verify input worksheet
if flagContinue
    vpIDs = getVPIDs(myWorksheet);
    interventionIDs = getInterventionIDs(myWorksheet);
    nVPs = length(vpIDs);
    nInterventions = length(interventionIDs);
    if ((nVPs < 1) || (nInterventions < 1))
        warning(['Insufficient VPs and interventions for ',mfilename,'.'])
        flagContinue = false;
    end
    % At least we can check the specified variables are
    % recongized as elements.  We won't make 'time' requires as specified 
    % but we will add this at the end in regardless.
    if(sum(ismember(myWorksheet.simProps.saveElementResultIDs,myWorksheet.compiled.elements(:,1))) < length(myWorksheet.simProps.saveElementResultIDs))
        warning(['Unable to identify all indicated saveElementResultIDs as elements in myWorksheet in call to ',mfilename,'.'])
        flagContinue = false;
    end    
    
    % May want to add a check for indDoses:
    % sum(ismember(fields(curIntervention),'vpDoseModifications')) > 0    
    
end

if flagContinue
    
    % We reset the random number generator if specified
    % this will be relevant if we are generating initial guesses
    % for optimization
    if mySimulateOptions.intSeed > -1
        rng(mySimulateOptions.intSeed, 'twister');
    end        
    
    filterFailedRunVPs = mySimulateOptions.filterFailedRunVPs;
    rerunExisting = mySimulateOptions.rerunExisting; 
    optimizeAxisIDs = mySimulateOptions.optimizeAxisIDs;
    optimizeType = mySimulateOptions.optimizeType;

    % We could force re-run for optimization.
    %if sum(ismember({'fmincon'},optimizeType)) > 0
    %    rerunExisting = true;
    %else
    %    rerunExisting = mySimulateOptions.rerunExisting;
    %end
    responseTypeID = mySimulateOptions.responseTypeID;
    % Can only write slices with the parfor loop counter,
    % so simResults needs to be 1-D rather than nInterventions X nVPs
    simResults = cell(1, nVPs * nInterventions);
    % We also need a vector to ultimately track which simulations we will
    % run. We include an option to avoid rerunning all, which is useful
    % for some calling functions.
    nSimulations = nInterventions * nVPs;
    flagRunSim = ones(1, nSimulations);
    flagRunVP = ones(1, nVPs);
    [nInterventionResults, nVPResults] = size(myWorksheet.results);
    if ((nInterventionResults == nInterventions) && (nVPResults == nVPs))
        if ~(rerunExisting)
            myResultClasses = cellfun(@class,myWorksheet.results, 'UniformOutput', false);
            % Results should be stored in a structure, we assume 
            % if a structure is provided then it is a valid result
        	flagRunVP = sum(strcmp(myResultClasses,'struct'),1);
            flagRunVP = ~(flagRunVP == nInterventions);
            flagRunSim = zeros(1, nSimulations);
            for vpCounter = 1 : nVPs
                % increment intervention first, then VP
                if flagRunVP(vpCounter)
                    simulationStartIndex = 1 + (vpCounter - 1) * nInterventions;
                    simulationEndIndex = nInterventions + (vpCounter - 1) * nInterventions;                    
                    flagRunSim(simulationStartIndex:simulationEndIndex) = ones(1,nInterventions);
                %else
                %    flagRunSim = cat(2, flagRunSim, zeros(1, nInterventions));
                end
            end
        end
        % This may consume some nontrivial memory with larger
        % runs, so clear it out.
        clear myResultClasses
    else
        % Otherwise, we don't know which result is paired with which
        % VP & intervention.  We should clear everything out and force
        % a re-run
        myWorksheet.results = cell(nInterventions, nVPs);
    end

    % We need to get the parameters ready in serial before 
    % calling parfor
    %elementNamesDefaultValues = myWorksheet.compiled.elements;
    [updateValues, updateDoses] = getSimulateValuesDoses(myWorksheet, flagRunVP, flagRunSim);
    
    % Specify simulation length and sample times
    exportedModel = myWorksheet.compiled.model;
    exportedModel.SimulationOptions.OutputTimes = myWorksheet.simProps.sampleTimes;
    exportedModel.SimulationOptions.StopTime = max(myWorksheet.simProps.sampleTimes);
    exportedModel.SimulationOptions.AbsoluteTolerance = myWorksheet.simProps.absoluteTolerance;
    exportedModel.SimulationOptions.RelativeTolerance = myWorksheet.simProps.relativeTolerance;    
    exportedModel.SimulationOptions.AbsoluteToleranceScaling = myWorksheet.simProps.absoluteToleranceScaling;    
    exportedModel.SimulationOptions.AbsoluteToleranceStepSize = myWorksheet.simProps.absoluteToleranceStepSize;    
    exportedModel.SimulationOptions.MaximumWallClock = myWorksheet.simProps.maximumWallClock;
    exportedModel.SimulationOptions.SolverType = myWorksheet.simProps.solverType;
    % It doesn't appear the exportedModel has a property similar to states 
    % to log in the configset object, so this will be done manually.
    mySaveElementResultIDs = myWorksheet.simProps.saveElementResultIDs;    
    % As a precaution, restart any existing parallel 
    % pools 
    if ~isempty(gcp('nocreate'))
        delete(gcp);
    end
    parpool;
	% in 2017a this message can cause issues, especially when
	% trying some of the global optimization methods
	% You may want this on for some applications,
	% but it is disabled here.
	parfevalOnAll(gcp(), @warning, 0, 'off', 'SimBiology:Simulation:NonFiniteRHS');    

    simResults = cell(1,nSimulations);
    % Only 'none' is supported here
    %if sum(ismember({'none'},optimizeType)) > 0
    parfor simulationCounter = 1 : nSimulations
        %for simulationCounter = 1 : nSimulations
        if flagRunSim(simulationCounter) > 0
            % increment intervention first, then VP
            % so we can complete simulations for a given VP
            vpCounter = floor((simulationCounter-1)/(nInterventions)) + 1;
            interventionCounter = simulationCounter - (vpCounter - 1) * nInterventions;
            % Retrieve the pre-specified parameter alterations & doses
            % for the current run
            currentModelValues = updateValues(interventionCounter, vpCounter,:);
            currentModelValues = reshape(currentModelValues,[],1);
            
            curIntervention = myWorksheet.interventions{interventionCounter};
            % We assume the ID's are pre-ordered to match
            % TODO: May want to error-proof this with some
            % sort of lookup, but wasn't needed for initial runs
            curDoseUpdateArray = curIntervention.('vpDoseModifications'){1,vpCounter};
            curDoses = {};
            for doseTypeCounter = 1 : length(curDoseUpdateArray)
                curUpdateStruct = curDoseUpdateArray{doseTypeCounter};
                theDoseID = curUpdateStruct.('Name');
                curDose = getdose(exportedModel,theDoseID);
                curDose.Amount = curUpdateStruct.('Amount');
                curDose.Interval = curUpdateStruct.('Interval');
                curDose.Rate = curUpdateStruct.('Rate');
                curDose.RepeatCount = curUpdateStruct.('RepeatCount');
                curDoses = [curDoses,curDose];
            end
            
            % Simulate with specified sampling interval
            try
                simData = simulate(exportedModel, currentModelValues, curDoses);
                simData = convertSimData(simData);
                % Reformat results into data table format
                if length(mySaveElementResultIDs) > 0
                    [~, keepIndices] = ismember(['time',mySaveElementResultIDs], simData.Names);
                    simData.Names = simData.Names(keepIndices);
                    simData.Data = simData.Data(:,keepIndices);
                end
                simResults{simulationCounter} = simData;
            catch
                % If the simulation fails, we need to move on
                simResults{simulationCounter} = [];
            end
        else
            simResults{simulationCounter} = [];
        end
    end
    simResults = reshape(simResults, [nInterventions, nVPs]);
    if sum(flagRunVP) < nVPs
        myWorksheet.results(:,find(flagRunVP)) = simResults(:,find(flagRunVP));
    else
        myWorksheet.results = simResults;
    end
    
    %end
    
    simResults = reshape(simResults, [nInterventions, nVPs]);
    % Clean up the worker pool
    delete(gcp)
    if sum(flagRunVP) < nVPs
        myWorksheet.results(:,find(flagRunVP)) = simResults(:,find(flagRunVP));
    else
        myWorksheet.results = simResults;
    end
    
    
 
    % Filter VPs with failed results?  These are potentially useful as a
    % diagnostic
    if filterFailedRunVPs
        filterVPids = cell(1,0);
        allVPIDs = getVPIDs(myWorksheet);
        for vpCounter = 1 : nVPs
            if sum(cellfun('isempty',myWorksheet.results(:,vpCounter))) > 0
                filterVPids{length(filterVPids)+1} = allVPIDs{vpCounter};
            end
        end
        myWorksheet = removeVPs(myWorksheet, filterVPids);
    end

else
    warning(['Could not complete ',mfilename,'. Returning original worksheet.'])
end
end
```
