## Supplementary File 3 for "A surrogate workflow for virtual population development: designing an amenable proxy for a hypothesis-test inspired nonlinear goodness-of-fit": addVariedVPs.m

```
function myWorksheet = addVariedVPs(myWorksheet, myVaryAxesOptions)
% Take an existing worksheet and add VPs varied along mechanistic axes to it.
%
% ARGUMENTS
% myWorksheet:          a worksheet
% myVaryAxesOptions:    an instance of a varyAxesOptions object.
%
% RETURNS
% myWorksheet
%
% First check the number of input arguments and apply defaults as needed
continueFlag = false;
if nargin > 2
    warning(['Too many input arguments to ',mfilename, '. Arguments should be: myWorksheet; and optionally: myVaryAxesOptions.'])
    continueFlag = false;
elseif nargin > 1
    continueFlag = true;
elseif nargin > 0
    myVaryAxesOptions = varyAxesOptions();
    myVaryAxesOptions.baseVPIDs = getVPIDs(myWorksheet);
    myVaryAxesOptions.varyAxisIDs = getAxisDefIDs(myWorksheet);
    continueFlag = true;
else
    warning(['Insufficient input arguments to ',mfilename, '. Arguments should be: myWorksheet; and optionally: myVaryAxesOptions.'])
end

% Check whether the input arguments make sense
if continueFlag
    passTestFlag = myVaryAxesOptions.verify(myWorksheet);
    if ~passTestFlag
        continueFlag = false;
    end
end

if continueFlag
    nBaseVPs = length(myVaryAxesOptions.baseVPIDs);
    if nBaseVPs > 1
        % If the selected VPs are not consistent WRT their variants, we
        % will at least warn the user to make sure they are aware.
        VPID1 = myVaryAxesOptions.baseVPIDs{1};
        myVP1 = getVP(myWorksheet,VPID1);
        myVP1Variants = myVP1.variants;
        variantComparePassFlag = true;
        if ((strcmp(myVaryAxesOptions.varyMethod,'sobol')) || (strcmp(myVaryAxesOptions.varyMethod,'saltelli')) || (strcmp(myVaryAxesOptions.varyMethod,'lh')))
            warning('A sobol or saltelli sequence, or latin hypercube sample, for sensitivity analysis doesnt make sense with multiple base VPs specified. Pick a single base VP.')
            continueFlag = false;
        else
            for vpCounter = 2 : nBaseVPs
                curVPID = myVaryAxesOptions.baseVPIDs{vpCounter};
                myVP = getVP(myWorksheet,curVPID);
                myVPVariants = myVP.variants;
                if length(setxor(myVP1Variants,myVPVariants)) > 0
                    variantComparePassFlag = false;
                end
            end
            if ~(variantComparePassFlag)
                warning(['Note that the variants for all VPs being varied are not consistent in ',mfilename, '.  Proceeding with creating the VPs anyways.'])
            end
        end
    end
end
    
if continueFlag    
    if ~strcmp(myVaryAxesOptions.varyMethod, 'none')    
        if length(myVaryAxesOptions.varyAxisIDs) < 1
            continueFlag = false;
            warning(['Must vary at least one axis in ',mfilename, '.'])
        end 
    end        
end

additionalIDString = myVaryAxesOptions.additionalIDString;
% Apply checks to the input argument values
testVPIDs = getVPIDs(myWorksheet);
if continueFlag
    if ~(ischar(additionalIDString)) == 1
        warning(['The argument additionalIDString should be a character.'])
        continueFlag = false;    
    end
    baseVPPassFlag = true;
    for baseVPCounter = 1 : nBaseVPs
        if ((sum(ismember(testVPIDs, myVaryAxesOptions.baseVPIDs{baseVPCounter})) > 1)) 
            baseVPPassFlag = false;
        end
    end
    if length(unique(myVaryAxesOptions.baseVPIDs)) < length(myVaryAxesOptions.baseVPIDs)
        baseVPPassFlag = false;
    end
    if ~baseVPPassFlag
        warning(['Degeneracy detected in specified worksheet and/or base VP IDs.'])
        continueFlag = false;
    end
end

if continueFlag
	existingVPCoefficients = getVPCoeffs(myWorksheet);
    
	allAxisIDs = getAxisDefIDs(myWorksheet);    
    variedAxisIndices = find(ismember(allAxisIDs, myVaryAxesOptions.varyAxisIDs));
	[nAxis,nVP] = size(existingVPCoefficients);
	if strcmp(myVaryAxesOptions.varyMethod,'localpca')
		if nAxis > (nVP+1)
			warning(['Not enough VPs for localpca option in ',mfilename,'.  Number of VPs should be at least 1 greater than the number of axes.  Exiting.'])
			continueFlag = false;
		end	
		if length(variedAxisIndices) < nAxis
			warning(['All axes should be varied for localpca option in ',mfilename,'.  Exiting.'])
			continueFlag = false;
		end	
	end
end

% Now for actually creating the varied VPs
if continueFlag
    
    % First we reset the random number generator if specified
    if myVaryAxesOptions.intSeed > -1
        rng(myVaryAxesOptions.intSeed, 'twister');
    end    
    
    if length(additionalIDString) > 0
        additionalIDString = ['_', additionalIDString, '_'];
    else
        additionalIDString = '_';
    end
    
    nAxis = length(allAxisIDs);
    nRAxis = length(myVaryAxesOptions.varyAxisIDs);
    % Even though a little slower, we do this with a for loop to maintain
    % the order in baseVPIDs
    baseVPIndices = nan(1,nBaseVPs);
    for baseVPCounter = 1 : nBaseVPs
        baseVPIndices(baseVPCounter) = find(ismember(testVPIDs, myVaryAxesOptions.baseVPIDs{baseVPCounter}));
    end
    baseVPCoefficients = existingVPCoefficients(:,baseVPIndices);
    if strcmp(myVaryAxesOptions.varyMethod,'uniform')
        newVPIDs = cell(1,myVaryAxesOptions.newPerOld * nBaseVPs);
        randomCoefficients = rand(nRAxis, myVaryAxesOptions.newPerOld * nBaseVPs);
        for baseVPCounter = 1 : nBaseVPs
            for newVPCounter = 1 : myVaryAxesOptions.newPerOld
                newVPIDs{1, newVPCounter + (baseVPCounter-1) * myVaryAxesOptions.newPerOld} = [myVaryAxesOptions.baseVPIDs{baseVPCounter},additionalIDString,num2str(newVPCounter)];
            end
        end   
    elseif strcmp(myVaryAxesOptions.varyMethod,'gaussian')
        newVPIDs = cell(1,myVaryAxesOptions.newPerOld * nBaseVPs);
        randomCoefficients = nan(nRAxis,myVaryAxesOptions.newPerOld * nBaseVPs);
        for baseVPCounter = 1 : nBaseVPs
            % * 0.05 rather than * 0.1 is less aggressive for the gaussian
            % but also finds more solutions if the output is constrained            
            randomCoefficients(:, ((baseVPCounter-1) * myVaryAxesOptions.newPerOld + 1) : (baseVPCounter*myVaryAxesOptions.newPerOld)) = (randn(nRAxis,myVaryAxesOptions.newPerOld) * myVaryAxesOptions.gaussianStd) + repmat(baseVPCoefficients(variedAxisIndices, baseVPCounter), 1, myVaryAxesOptions.newPerOld);
            for newVPCounter = 1 : myVaryAxesOptions.newPerOld
                newVPIDs{1, newVPCounter + (baseVPCounter-1) * myVaryAxesOptions.newPerOld} = [myVaryAxesOptions.baseVPIDs{baseVPCounter},additionalIDString,num2str(newVPCounter)];
            end
        end
        indicesNeg = find(randomCoefficients < 0);
        randomCoefficients(indicesNeg) = 0;
        indicesBig = find(randomCoefficients > 1);
        randomCoefficients(indicesBig) = 1;    
    elseif strcmp(myVaryAxesOptions.varyMethod,'localpca')
        newVPIDs = cell(1,myVaryAxesOptions.newPerOld * nBaseVPs);
        randomCoefficients = nan(nRAxis,myVaryAxesOptions.newPerOld * nBaseVPs);
		% Transpose VPs from columns to rows for pdist2
		vpDistance = pdist2(existingVPCoefficients',existingVPCoefficients');
        for baseVPCounter = 1 : nBaseVPs
			% Find the VPs closest to the current base
            curDistance = vpDistance(baseVPIndices(baseVPCounter),:);
			% Get just enough for a square matrix so we can compute principal components
			[~, sortI] = sort(curDistance,'ascend');
            % Exclude VPs that are too close.
            % This is a bit arbitrary.
            excludeI = find(curDistance<=1E-8);
            sortI = setdiff(sortI, excludeI);
            sortI = [baseVPIndices(baseVPCounter), sortI];
			% Check for linear independence
            % This will iterate until a set of
            % basis vectors of sufficient number are found,
            % Starting with the nearest VPs and expanding.
            existingVPCoefficients = existingVPCoefficients(:,sortI);
            % Check independence of the non-parent VPs
            [addBasisVPCoefficients, VPind] = getLinearIndependentVectors(existingVPCoefficients(:,2:end));
            addBasisVPCoefficients = [existingVPCoefficients(:,1),addBasisVPCoefficients];
            VPind = [1, (VPind+1)];
            
            if length(VPind) >= nAxis + 1
                % Bounds will be adjusted in the function
                randomCoefficients(:, ((baseVPCounter-1) * myVaryAxesOptions.newPerOld + 1) : (baseVPCounter*myVaryAxesOptions.newPerOld)) = resamplePCASpace(addBasisVPCoefficients,1,myVaryAxesOptions.gaussianStd,myVaryAxesOptions.newPerOld,[zeros(nAxis,1),ones(nAxis,1)]);
                for newVPCounter = 1 : myVaryAxesOptions.newPerOld
                    newVPIDs{1, newVPCounter + (baseVPCounter-1) * myVaryAxesOptions.newPerOld} = [myVaryAxesOptions.baseVPIDs{baseVPCounter},additionalIDString,num2str(newVPCounter)];
                end                
            else
                % * 0.05 rather than * 0.1 is less aggressive for the gaussian
                % but also finds more solutions if the output is constrained            
                randomCoefficients(:, ((baseVPCounter-1) * myVaryAxesOptions.newPerOld + 1) : (baseVPCounter*myVaryAxesOptions.newPerOld)) = (randn(nRAxis,myVaryAxesOptions.newPerOld) * 0.05* myVaryAxesOptions.gaussianStd) + repmat(baseVPCoefficients(variedAxisIndices, baseVPCounter), 1, myVaryAxesOptions.newPerOld);
                for newVPCounter = 1 : myVaryAxesOptions.newPerOld
                    newVPIDs{1, newVPCounter + (baseVPCounter-1) * myVaryAxesOptions.newPerOld} = [myVaryAxesOptions.baseVPIDs{baseVPCounter},additionalIDString,num2str(newVPCounter)];
                end
                indicesNeg = find(randomCoefficients < 0);
                randomCoefficients(indicesNeg) = 0;
                indicesBig = find(randomCoefficients > 1);
                randomCoefficients(indicesBig) = 1;                     
            end
        end 		
    elseif strcmp(myVaryAxesOptions.varyMethod,'lh')
        newVPIDs = cell(1,myVaryAxesOptions.newPerOld * nBaseVPs);
        randomCoefficients = lhsdesign(myVaryAxesOptions.newPerOld * nBaseVPs,nRAxis);
        randomCoefficients = randomCoefficients';
        for baseVPCounter = 1 : nBaseVPs          
            for newVPCounter = 1 : myVaryAxesOptions.newPerOld
                newVPIDs{1, newVPCounter + (baseVPCounter-1) * myVaryAxesOptions.newPerOld} = [myVaryAxesOptions.baseVPIDs{baseVPCounter},additionalIDString,num2str(newVPCounter)];
            end
        end      
    elseif strcmp(myVaryAxesOptions.varyMethod,'sobol')
        newVPIDs = cell(1,myVaryAxesOptions.newPerOld * nBaseVPs);
        mySobolSet = sobolset(nRAxis);
        if myVaryAxesOptions.intSeed == 0
            disp(['Note: current behavior of Sobol sampling when intSeed is 0 is not to scramble in ',mfilename,'.  This forces the RNG to reseed with 0.'])
        else
            disp(['Note: current behavior of Sobol sampling when intSeed is not 0 is to scramble in ',mfilename,'.'])
            mySobolSet = scramble(mySobolSet,'MatousekAffineOwen');
        end
        randomCoefficients = net(mySobolSet,myVaryAxesOptions.newPerOld);
        randomCoefficients = transpose(randomCoefficients);
        for baseVPCounter = 1 : nBaseVPs
            for newVPCounter = 1 : myVaryAxesOptions.newPerOld
                newVPIDs{1, newVPCounter + (baseVPCounter-1) * myVaryAxesOptions.newPerOld} = [myVaryAxesOptions.baseVPIDs{baseVPCounter},additionalIDString,num2str(newVPCounter)];
            end
        end
    elseif strcmp(myVaryAxesOptions.varyMethod,'none')
        newVPIDs = cell(1,myVaryAxesOptions.newPerOld * nBaseVPs);
        randomCoefficients = nan(nRAxis,myVaryAxesOptions.newPerOld * nBaseVPs);
        for baseVPCounter = 1 : nBaseVPs
            for newVPCounter = 1 : myVaryAxesOptions.newPerOld
                newVPIDs{1, newVPCounter + (baseVPCounter-1) * myVaryAxesOptions.newPerOld} = [myVaryAxesOptions.baseVPIDs{baseVPCounter},additionalIDString,num2str(newVPCounter)];
                randomCoefficients(:,(baseVPCounter-1)* myVaryAxesOptions.newPerOld + newVPCounter) = baseVPCoefficients(:,baseVPCounter);
            end
        end
    else        
        % For Saltelli, we structure the sampling
        % to maximize the efficiency of estimating the
        % Sobol sensitivity indices.  For a desciption, see:
        % Saltelli, A., et al, Global Sensitivity Analysis:
        % The Primer. 2009. Pages 164-167.
        % I have left this as the unscrambled sequence
        % until I can investigate the best way to scramble
        mySobolSet = sobolset(nRAxis*2);
        if myVaryAxesOptions.intSeed == 0
            disp(['Note: current behavior of Sobol/Saltelli sampling when intSeed is 0 is not to scramble in ',mfilename,'.'])
        else
            disp(['Note: current behavior of Sobol/Saltelli sampling when intSeed is not 0 is to scramble in ',mfilename,'.'])
            mySobolSet = scramble(mySobolSet,'MatousekAffineOwen');
        end        
        randomCoefficientsA = net(mySobolSet,myVaryAxesOptions.newPerOld);      
        randomCoefficientsB = randomCoefficientsA(:,(nRAxis+1):nRAxis*2);
        randomCoefficientsA = randomCoefficientsA(:,1:nRAxis);
        randomCoefficientsC = nan(nRAxis*myVaryAxesOptions.newPerOld,nRAxis);
        for aCounter = 1 : nRAxis
            cRowStart = (aCounter-1) * myVaryAxesOptions.newPerOld + 1;
            randomCoefficientsC(cRowStart:(cRowStart+myVaryAxesOptions.newPerOld-1), :) = randomCoefficientsB;
            randomCoefficientsC(cRowStart:(cRowStart+myVaryAxesOptions.newPerOld-1), aCounter) = randomCoefficientsA(:,aCounter);
        end 
        randomCoefficients = [randomCoefficientsA; randomCoefficientsB; randomCoefficientsC];
        clear randomCoefficientsA randomCoefficientsB randomCoefficientsC
        randomCoefficients = transpose(randomCoefficients);
        % We construct the VPIDs so we can search on _C_ _B_ and _A_ in the
        % name.  Beware when of this naming base VPs and
        % running the Saltelli option 
        % We run with N * (k + 2) samples. 
        newVPIDs = cell(1, nBaseVPs * myVaryAxesOptions.newPerOld * (nRAxis + 2));
        for newVPCounter = 1 : myVaryAxesOptions.newPerOld
            newVPIDs{1,newVPCounter}=[myVaryAxesOptions.baseVPIDs{1},additionalIDString,'A_',num2str(newVPCounter)];
        end        
        for newVPCounter = 1 : myVaryAxesOptions.newPerOld
            newVPIDs{1,myVaryAxesOptions.newPerOld + newVPCounter}=[myVaryAxesOptions.baseVPIDs{1},additionalIDString,'B_',num2str(newVPCounter)];
        end       
        for newVPACounter = 1 : nRAxis
            for newVPBCounter = 1 : myVaryAxesOptions.newPerOld
                newVPIDs{1, 2*myVaryAxesOptions.newPerOld + (newVPACounter - 1) * myVaryAxesOptions.newPerOld + newVPBCounter} = [myVaryAxesOptions.baseVPIDs{1},additionalIDString,'C__B_',num2str(newVPBCounter),'_A_',num2str(newVPACounter)];
            end
        end
    end
    
    nNewVPs = length(newVPIDs);
    nWshVPs = length(testVPIDs);
    [nInterventions, ~] = size(myWorksheet.interventions);
    fieldNames = fields(myWorksheet);
    newVPCoefficients = nan(nAxis, nNewVPs);
    if ~strcmp(myVaryAxesOptions.varyMethod,'saltelli')
        extensionPerBaseVP = myVaryAxesOptions.newPerOld;
    else
        extensionPerBaseVP = myVaryAxesOptions.newPerOld * (nRAxis + 2);
    end
    for fieldCounter = 1 : length(fieldNames)
        curField = fieldNames{fieldCounter};
        % We need to modify VP-related fields in the worksheet
        if strcmp(curField, 'vpDef')
            myWorksheet.(curField) = [myWorksheet.(curField), cell(1, nNewVPs)];
            for baseVPCounter = 1 : nBaseVPs
                baseVPIndex = baseVPIndices(baseVPCounter);
                startIndex = (baseVPCounter - 1) * extensionPerBaseVP + 1;
                endIndex = (baseVPCounter) * extensionPerBaseVP;
                myWorksheet.(curField)(nWshVPs+startIndex : nWshVPs+endIndex) = {myWorksheet.(curField){1,baseVPIndex}};
                % update the ID's appropriately
                for childVPCounter = 1 : extensionPerBaseVP
                    newVPIndex = startIndex + (childVPCounter-1);
                    myWorksheet.(curField){1,nWshVPs+newVPIndex}.ID = newVPIDs{newVPIndex};
                end
            end
        elseif strcmp(curField, 'axisProps')
            for baseVPCounter = 1 : nBaseVPs
                startIndex = (baseVPCounter - 1) * extensionPerBaseVP + 1;
                endIndex = (baseVPCounter) * extensionPerBaseVP;               
                newVPCoefficients(:,startIndex:endIndex) = repmat(baseVPCoefficients(:,baseVPCounter),1, extensionPerBaseVP);
                if length(variedAxisIndices) > 0
                    newVPCoefficients(variedAxisIndices,startIndex:endIndex) = randomCoefficients(:,startIndex:endIndex);
                end
            end
            newVPCoefficients = cat(2, existingVPCoefficients, newVPCoefficients);
            
            myWorksheet.(curField).axisVP.coefficients = newVPCoefficients;
            
        % We will not enforce zeroing out old results,
        % to avoid the potential extra execution time
        % associated with re-running old results.
        elseif strcmp(curField, 'results')
            previousResults = myWorksheet.(curField);
            [nInterventionResults, nVPResults] = size(previousResults);
            if nInterventionResults == nInterventions;
                myWorksheet.(curField)(:,(nWshVPs+1):(nWshVPs+nNewVPs)) = cell(nInterventions, nNewVPs);
            else
                myWorksheet.(curField) = cell(0,0);
            end
        end
    end
else
    warning(['Unable to run ',mfilename,', returning input worksheet.'])
end
end
```
