## Supplementary File 3 for "A surrogate workflow for virtual population development: designing an amenable proxy for a hypothesis-test inspired nonlinear goodness-of-fit": bestNofM.m

```
function myWorksheet = bestNofM(myWorksheet, myBestNofMOptions)
% Try M random VPs based on an initial VP, and then keep the N best in
% a new worksheet. 
% Note that axes, where variability will be introduced, should also be
% set up before calling this worksheet.
%
% ARGUMENTS
% myWorksheet:       Starting worksheet.  Note the mechanistic axes must be
%                    defined first before calling this function.
% myBestNofMOptions: Options for testing new VPs with the varied axes
%
% RETURNS
% myWorksheet:       An updated worksheet that incudes the best VPs
%


% First check whether sufficient input arguments are provided
continueFlag = true;
if nargin > 2
    warning(['Too many input arguments to ',mfilename, '. Arguments should be: myWorksheet and myBestNofMOptions.'])
    continueFlag = false;
elseif nargin < 2
    warning(['Too few input arguments to ',mfilename, '. Arguments should be: myWorksheet and myBestNofMOptions.'])
    continueFlag = false;    
end

% Check whether the input arguments make sense
if continueFlag
    passTestFlag = myBestNofMOptions.verify(myWorksheet);
    if ~passTestFlag
        continueFlag = false;
    end
end

if continueFlag
    % For now, if we save to file, enforce incrementing the save file name.
    % This may help troubleshooting, and keeping track of longer runs.
    % Might want to make this into another input argument.
    incrementSaveName = false;
end

% ADD IN ENFORCE SOME AXES TO VARY HERE, LOAD ALL WSH AXES IF NONE
% SPECIFIED
if continueFlag
    if length(myBestNofMOptions.varyAxisIDs) < 1
    	myBestNofMOptions.varyAxisIDs = getAxisDefIDs(myWorksheet);
    end
end

if continueFlag
    if myBestNofMOptions.intSeed > -1
        rng(myBestNofMOptions.intSeed, 'twister');
    end
    % First pull out the base VP into a new worksheet
    myWorksheet = copyWorksheet(myWorksheet, {myBestNofMOptions.baseVPID});
    mySimulateOptions = simulateOptions;
    mySimulateOptions.rerunExisting = false;
    mySimulateOptions.filterFailedRunVPs = true;   
    baseVPid = myBestNofMOptions.baseVPID;
    % Add the varied VPs
    tic
    saveTime = 0;
    for k = 1:myBestNofMOptions.repeatK 
        myVaryOptions = varyAxesOptions;
        allVPIDs = getVPIDs(myWorksheet);
        if sum(ismember(allVPIDs,baseVPid)) < 1
            baseVPid = allVPIDs{1};
        end
        myVaryOptions.baseVPIDs = {baseVPid};
        myVaryOptions.newPerOld = myBestNofMOptions.tryM;
        myVaryOptions.varyMethod = myBestNofMOptions.varyMethod;
        myVaryOptions.additionalIDString = num2str(k);
        myVaryOptions.varyAxisIDs = myBestNofMOptions.varyAxisIDs;
        myWorksheet = addVariedVPs(myWorksheet, myVaryOptions);
        % Run the simulation
        myWorksheet = simulateWorksheet(myWorksheet,mySimulateOptions);
        % Evaluate the response type and pick the best
        myResponseTypeResult = evaluateResponseType(myWorksheet, myBestNofMOptions.responseTypeID);
        myVPvalues = myResponseTypeResult.vpValues;
        myVPvalueCutoff = sort(myVPvalues, 'ascend');
        % It is possible if we have a lot of ties with an identical
        % objective value that we may keep extra VPs.        
        bestVPvalueCutoff = myVPvalueCutoff(1);
        bestVPindex = find(myResponseTypeResult.vpValues <= bestVPvalueCutoff);
        if length(bestVPindex) > 1
            bestVPindex = bestVPindex(1);
        end
        % Best case, we have at least NtoKeep, but this many may not have
        % complete successfully
        myVPvalueCutoff = myVPvalueCutoff(min(myBestNofMOptions.keepN,sum(~isnan(myVPvalueCutoff))));
        myVPindices = find(myResponseTypeResult.vpValues <= myVPvalueCutoff);
        allVPIDs = getVPIDs(myWorksheet);
        % If this is too long, we will just keep up to the max N
        if length(myVPindices) > myBestNofMOptions.keepN
            round1values = myResponseTypeResult.vpValues(myVPindices);
            [round1valuesSort, I] = sort(round1values, 'ascend');
            myVPindices = myVPindices(I);
            myVPindices = myVPindices(1:myBestNofMOptions.keepN);
        end
        vpToKeep = allVPIDs(myVPindices);
        
        baseVPid = allVPIDs{bestVPindex};
        % Now just pull the best VPs into a new worksheet.
        myWorksheet = copyWorksheet(myWorksheet, vpToKeep, true);
        if length(myBestNofMOptions.saveFile) > 0
            % Limit the file saves to every 60 minutes, since it
            % may take time to write all the data if the files
            % get big.
            if (((toc/60-saveTime) > 60) || (k == KrepeatM)) 
                saveTime = toc/60;
                if incrementSaveName
                    saveWorksheet(myWorksheet, [myBestNofMOptions.saveFile,'_',num2str(k)]);
                else
                    saveWorksheet(myWorksheet, myBestNofMOptions.saveFile);
                end
            end
        end
        if myBestNofMOptions.verbose
            disp(['Completed iteration ',num2str(k),' of ',num2str(myBestNofMOptions.repeatK),' in ',mfilename,'.']);
            disp(['Elapsed time in worksheet iterations is ',num2str(toc/60),' min.']);
        end        
    end
else
    warning(['Unable to run ',mfilename,', returning input worksheet.'])
    myWorksheet = myWorksheet;
end
end
```
