## Supplementary File 3 for "A surrogate workflow for virtual population development: designing an amenable proxy for a hypothesis-test inspired nonlinear goodness-of-fit": calculateControlCoefficients.m

```
function myControlCoefficientsResults = calculateControlCoefficients(myWorksheet,myCalculateControlCoefficientsOptions)
% This function takes a worksheet from a local SA 
% (runControlCoefficientSimulation) and gets the simulation results of interest
% This is based on the local SA method in:
% Chen X, Hickling TP, Vicini P. A mechanistic, multiscale mathematical 
% model of immunogenicity for therapeutic proteins: part 1-theoretical
% model. CPT: pharmacometrics & systems pharmacology. 2014;3:e133.
%
% ARGUMENTS
% myWorksheet:                            an instance of a worksheet, 
%                                         output from runUnivariateSweep()
% myCalculateControlCoefficientsOptions: an instance of a calculateControlCoefficientsOptions
%                                         object
%
% RETURNS
% myControlCoefficientsResults:                 A structured array with the desired results.
%                                               The fields are:
%                               (varname):      the (varname) field contains the
%                                               results from the analysis
%                                               Each is reported as a (nAxis + 1)
%                                               x 2 matrix
%                               axisDefIDs:     a 1 x nAxis + 1 cell array to record
%                                               the name of axes that were
%                                               varied.
%                               interventionID: a record of the intervention
%                                         
%
myControlCoefficientsResults = struct();
continueFlag = false;
if nargin > 2
    warning(['Too many input arguments to ',mfilename, '. Arguments should be: myWorksheet and myCalculateControlCoefficientsOptions.'])
    continueFlag = false;
elseif nargin > 1
    continueFlag = true;
else
    warning(['Insufficient input arguments to ',mfilename, '. Arguments should be: myWorksheet and myCalculateControlCoefficientsOptions.'])
    continueFlag = false;
end

if continueFlag
    continueFlag = myCalculateControlCoefficientsOptions.verify(myWorksheet);
    if ~(continueFlag)
       warning(['Please correct the options provided to ',mfilename, '.']) 
    end
    myResultIDs = myCalculateControlCoefficientsOptions.analyzeElementResultIDs;
    nResultIDs = length(myResultIDs);
    if nResultIDs < 1
        warning(['At least one valid model output must be specified in analyzeElementResultIDs in options provided to ',mfilename,'.'])
        continueFlag = false;
    end    
end

if continueFlag
    allVPIDs = getVPIDs(myWorksheet);
    nVPs = length(allVPIDs);
    allInterventionIDs = getInterventionIDs(myWorksheet);
    myInterventionIndex = find(ismember(allInterventionIDs, myCalculateControlCoefficientsOptions.interventionID));
    Ymat = nan(nVPs-1,nResultIDs);
    allAxisIDs=getAxisDefIDs(myWorksheet);
    for resultCounter = 1 : nResultIDs
        vpCounter = 1;
        curResults = myWorksheet.results{myInterventionIndex,vpCounter};
        timeIndex = find(ismember(curResults.Names,'time'));
        timeVals = curResults.Data(:,timeIndex);
        %timeIndex = find(timeVals == myCalculateControlCoefficientsOptions.analyzeTime);
        resultIndex = find(ismember(curResults.Names,myResultIDs{resultCounter}));
        % Chen references the max deviation
        refResults = curResults.Data(:,resultIndex);
        for vpCounter = 2 : nVPs
            % We take for granted non-degeneracy of referenced result
            % element ID's.
            curResults = myWorksheet.results{myInterventionIndex,vpCounter};
            resultIndex = find(ismember(curResults.Names,myResultIDs{resultCounter}));
            curAxisID = myCalculateControlCoefficientsOptions.varyAxisIDs{vpCounter-1};
            curAxisIndex = find(ismember(allAxisIDs, curAxisID));
            refElementValue = myWorksheet.axisProps.axisVP.calculateElementValues([curAxisIndex,1],myWorksheet);
            endElementValue = myWorksheet.axisProps.axisVP.calculateElementValues([curAxisIndex,vpCounter],myWorksheet);
            nElements = length(refElementValue);
            if nElements > 1
                warning(['Axis ',curAxisID,' is problematic in ',mfilename,' with ',num2str(nElements),' elements. Each axis should have 1 element for the analysis. Assigning nan.'])
                ccp = nan;
            else
                ccp = ((curResults.Data(:,resultIndex)-refResults)/(endElementValue-refElementValue)*refElementValue)./refResults;
                ccp = abs(ccp);
            end
            Ymat(vpCounter-1, resultCounter) = max(ccp);
        end
    end
    for resultCounter = 1 : nResultIDs
        myControlCoefficientsResults.(myResultIDs{resultCounter}) = Ymat(:, resultCounter);
    end
    %myControlCoefficientsResults.('time') = myCalculateControlCoefficientsOptions.analyzeTime;
    myControlCoefficientsResults.('axisDefIDs') = myCalculateControlCoefficientsOptions.varyAxisIDs;
    myControlCoefficientsResults.('interventionID') = myCalculateControlCoefficientsOptions.interventionID;    
else
    warning(['Unable to run ',mfilename, '.'])
end
end
```
