## Supplementary File 3 for "A surrogate workflow for virtual population development: designing an amenable proxy for a hypothesis-test inspired nonlinear goodness-of-fit": bestNofMOptions.m

```
classdef bestNofMOptions
% Options object for randomly creating, testing, and slecting VPs in 
% a worksheet with the bestNofM function
%
% PROPERTIES
% baseVPid:       (required to update) ID string of the baseline VP to
%                 base the variations on
% responseTypeID: (required to update) ID string of the response type to 
%                 implement as the objective
% keepN:          The keepN best VPs will be kept
% tryM:           The each run will randomly generate tryM VPs
% repeatK:        Repeat the tryM tries repeatK times.  Note 1
%                 here means to run once rather than run once and repeat
%                 again.
%                 This is available to keep Worksheet sizes smaller 
%                 (memory) while still using the available cores.
% verbose:        If true, write some information to screen so you can 
%                 verify process is executing
% saveFile:       (optional) if provided, each run of the algorithm will be saved to 
%                 this file in .mat format.  Default is an empty string, 
%                 '', which results in no iterative save
% varyAxisIDs:    Axes IDs to vary, provided preferably as a 1XL  
%                 cell array of strings. If provided, only the specified 
%                 axes will be varied.  If empty, all will be varied.
% intSeed:        (optional) an integer, used to reseed the random number generator, 
%                 if given. Otherwise, -1 to ignore.
% varyMethod:      (optional) When generating new VPs, this is the
%                 distribution to draw from.
%                        'uniform'  a uniform distribution for each variable
%                        'sobol'    a sobol sequence is used
%                        'lh'      latin hypercube

   properties
        baseVPID;
        responseTypeID;
        keepN;
        tryM;
        repeatK;
        verbose;
        saveFile;
        varyAxisIDs;
        intSeed;
        varyMethod; 
   end
   methods
     
       
      function obj = set.baseVPID(obj,myBaseVPID)
          if ischar(myBaseVPID)
              obj.baseVPID = (myBaseVPID);
          else
              error(['Invalid baseVPID specified for ',mfilename,', a VP ID string should be specified.'])
          end
      end
      
      function obj = set.responseTypeID(obj,myResponseTypeID)
          if ischar(myResponseTypeID)
              obj.responseTypeID = (myResponseTypeID);
          else
              error(['Invalid responseTypeID specified for ',mfilename,', a response type ID string should be specified.'])
          end
      end
      
      function obj = set.keepN(obj,myKeepN)
          failFlag = false;
          if isnumeric(myKeepN) 
              if isequal(size(myKeepN),[1 1])
                  if round(myKeepN) > 0
                      obj.keepN = round(myKeepN);
                  else
                      failFlag = true;
                  end
              else
                  failFlag = true;
              end
          else
              failFlag = true;
          end
          if failFlag
              error(['Invalid keepN specified for ',mfilename,'. A positive number should be specified.'])
          end
      end      
      
      function obj = set.tryM(obj,myTryM)
          failFlag = false;
          if isnumeric(myTryM) 
              if isequal(size(myTryM),[1 1])
                  if round(myTryM) > 0
                      obj.tryM = round(myTryM);
                  else
                      failFlag = true;
                  end
              else
                  failFlag = true;
              end
          else
              failFlag = true;
          end
          if failFlag
              error(['Invalid tryM specified for ',mfilename,'. A positive number should be specified.'])
          end
      end     
  
      function obj = set.repeatK(obj,myRepeatK)
          failFlag = false;
          if isnumeric(myRepeatK) 
              if isequal(size(myRepeatK),[1 1])
                  if round(myRepeatK) > 0
                      obj.repeatK = round(myRepeatK);
                  else
                      failFlag = true;
                  end
              else
                  failFlag = true;
              end
          else
              failFlag = true;
          end
          if failFlag
              error(['Invalid repeatK specified for ',mfilename,'. A positive number should be specified.'])
          end
      end             
      
      function obj = set.verbose(obj,myVerbose)
          if islogical(myVerbose)
              obj.verbose = myVerbose;
          else
              error(['Invalid verbose specified for ',mfilename,', a boolean (true/false) should be specified.'])
          end
      end
      
      function obj = set.saveFile(obj,mySaveFile)
          if ischar(mySaveFile)
              obj.saveFile = (mySaveFile);
          else
              error(['Invalid saveFile specified for ',mfilename,', a saveFile string should be specified.'])
          end
      end      
      
      function obj = set.varyAxisIDs(obj,myVaryAxisIDs)
          if iscell(myVaryAxisIDs)
              obj.varyAxisIDs = (myVaryAxisIDs);
          else
              error(['Invalid varyAxisIDs specified for ',mfilename,', a cell array of axis ID strings should be specified.'])
          end
      end         
      
      function obj = set.intSeed(obj,myIntSeed)
          if ((mod(myIntSeed,1) == 0) && (myIntSeed>=-1))
              obj.intSeed = (myIntSeed);
          else
              error(['Invalid intSeed specified for ',mfilename,', a non-negative integer should be specified, or -1 to ignore.'])
          end
      end              

      function obj = set.varyMethod(obj,myVaryMethod)
          allowedSettings = {'uniform','sobol','lh'};
          if sum(ismember(allowedSettings, lower(myVaryMethod)))>0
              obj.varyMethod = lower(myVaryMethod);
          else
              error('Invalid varyMethod specified for ',mfilename,', should specify one of: ',strjoin(allowedSettings,', '),'.')
          end
      end 
      
      function value = get(obj,propName)
          switch propName
              case 'baseVPID'
                  value = obj.baseVPID;
              case 'responseTypeID'
                  value = obj.responseTypeID;
              case 'keepN'
                  value = obj.keepN; 
              case 'tryM'
                  value = obj.tryM;                   
              case 'repeatK'
                  value = obj.repeatK;  
              case 'verbose'
                  value = obj.verbose;  
              case 'saveFile'
                  value = obj.saveFile;
              case 'varyAxisIDs'
                  value = obj.varyAxisIDs;                    
              case 'intSeed'
                  value = obj.intSeed; 
              case 'varyMethod'
                  value = obj.varyMethod;                   
          end
      end 
      
      function passCheck = verify(obj,myWorksheet)
          % Verify the specified bestNofMOptions are acceptable for a 
          % given worksheet.
          %
          % ARGUMENTS
          %  (self)
          %  myWorksheet:  a worksheet data structure
          %
          % RETURNS
          %  passCheck:    a boolean (true/false) indicating whether
          %                the specified varyAxesOptions pass the check
          %
          passCheck = true;
          allVPIDs = getVPIDs(myWorksheet);
          if sum(ismember(allVPIDs, obj.baseVPID)) < 1
              warning(['Specified baseVPID should all be selected from VPIDs the worksheet in ',mfilename,'.'])
              passCheck = false;
          end
          allResponseTypeIDs = getResponseTypeIDs(myWorksheet);
          if sum(ismember(allResponseTypeIDs, obj.responseTypeID)) < 1
              warning(['Specified responseTypeID should all be selected from responseTypeID the worksheet in ',mfilename,'.'])
              passCheck = false;
          end          
          allAxisDefIDs = getAxisDefIDs(myWorksheet);
          if sum(ismember(allAxisDefIDs, obj.varyAxisIDs)) < length(obj.varyAxisIDs)
              warning(['Specified varyAxisIDs should all be specified from axis IDs the worksheet in ',mfilename,'.'])
              passCheck = false;
          end     
          if sum(length(obj.varyAxisIDs)< length(unique(obj.varyAxisIDs)))
              warning(['Specified varyAxisIDs should all be unique in ',mfilename,'.'])
              passCheck = false;
          end              
          if (obj.keepN > obj.tryM)
              warning(['Specified tryM not be less than keepN in ',mfilename,'.'])
              passCheck = false;
          end                 
      end
      
      % The constructor method must have the same name as the class
      function obj = bestNofMOptions()
          obj.baseVPID = '';
          obj.responseTypeID = '';
          obj.keepN = 10;
          obj.tryM = 100;
          obj.repeatK = 1;
          obj.verbose = true;
          obj.saveFile = '';
          obj.varyAxisIDs = cell(1,0);
          obj.intSeed = -1;
          obj.varyMethod = 'uniform';
      end
   end
    
end
```
