## Supplementary File 3 for "A surrogate workflow for virtual population development: designing an amenable proxy for a hypothesis-test inspired nonlinear goodness-of-fit": calculateControlCoefficientsOptions.m

```
classdef calculateControlCoefficientsOptions
% Options object setting up the the control coefficient analysis.
% Note that the analysis must be run with the returned
% worksheet from the simulate step (runControlCoefficientSimulation).
%
%
% PROPERTIES
% interventionID:          (Required) Worksheet intervention to run
%                          analysis on.
% analyzeElementResultIDs: (Required) Worksheet outputs to run analysis
%                          on.
% varyAxisIDs:             (Required) IDs for the varied axes
    
   properties
      interventionID
      analyzeElementResultIDs
      varyAxisIDs
   end
   
   methods

      function obj = set.interventionID(obj,myInterventionID)
          if ischar(myInterventionID)
              obj.interventionID = myInterventionID;
          else
              error(['Invalid interventionID specified for ',mfilename,', an intervention ID string should be specified.'])
          end
      end     
      
      function obj = set.analyzeElementResultIDs(obj,myAnalyzeElementResultIDs)
          if iscell(myAnalyzeElementResultIDs)
              obj.analyzeElementResultIDs = (myAnalyzeElementResultIDs);
          else
              error(['Invalid analyzeElementResultIDs specified for ',mfilename,', a cell array of variable ID strings should be specified.'])
          end
      end     
      
      function obj = set.varyAxisIDs(obj,myVaryAxisIDs)
          if iscell(myVaryAxisIDs)
              obj.varyAxisIDs = (myVaryAxisIDs);
          else
              error(['Invalid varyAxisIDs specified for ',mfilename,', a cell array of axis ID strings should be specified.'])
          end
      end         

      
      function value = get(obj,propName)
          switch propName                 
              case 'interventionID'
                  value = obj.interventionID;   
              case 'analyzeElementResultIDs'
                  value = obj.analyzeElementResultIDs;          
              case 'varyAxisIDs'
                  value = obj.varyAxisIDs;                    
          end
      end     
      
      function passCheck = verify(obj,myWorksheet)
          passCheck = true;
          allInterventionIDs = getInterventionIDs(myWorksheet);
          allVPIDs = getVPIDs(myWorksheet);
          [~, nVPResults]= size(myWorksheet.results);
          if sum(ismember(allInterventionIDs,  obj.interventionID)) < 1
              warning(['Specified interventionID should be selected from interventionIDs the worksheet in ',mfilename,'.'])
              passCheck = false;
          end     
          if length(allVPIDs) ~= (length(obj.varyAxisIDs)+1)
              warning(['Number of VPs in the worksheet in ',mfilename,' not expected given the number of varyAxisIDs.'])
              passCheck = false;
          end
          if passCheck
              interventionIndex = find(ismember(allInterventionIDs, obj.interventionID));
              checkResults = myWorksheet.results(interventionIndex,:);
              if sum(sum(arrayfun(@(i) isstruct(checkResults{i}),1:nVPResults)) ) < length(allVPIDs)
                  warning(['Results for all VPs not available from worksheet in ',mfilename,'.'])
                  passCheck = false;
              else
                  for vpCounter = 1 : nVPResults
                      if sum(ismember(checkResults{1, vpCounter}.Names,obj.analyzeElementResultIDs)) < length(obj.analyzeElementResultIDs)
                          passCheck = false;
                      end
                  end
                  if ~passCheck
                      warning(['Not all analyzeElementResultIDs identified in the worksheet results provided to ',mfilename,'.'])
                  end
              end
          end
      end
      
      % The constructor method must have the same name as the class
      function obj = calculateControlCoefficientsOptions() 
          obj.interventionID='';
          obj.analyzeElementResultIDs={};
          obj.varyAxisIDs={};           
      end
   end
end
```
