## Supplementary File 3 for "A surrogate workflow for virtual population development: designing an amenable proxy for a hypothesis-test inspired nonlinear goodness-of-fit": clusterPickOptions.m

```
classdef clusterPickOptions
% This class defines options for pickClusterVPs
%
% PROPERTIES:
%  algorithm:       kmedoid algorithm to use.  These are taken from MATLAB
%                    and valid options include:
%                       'pam':   classical Partition Around Medoids
%                       'clara': a good alternative if clustering more than
%                                a couple thousand VPs.  Scales better than
%                                PAM.
%                       'small': uses an algorithm similar to k-means to
%                                find medoids
%                       'large': similar to 'small' but uses a random
%                                sample
%                       'auto' (default):  automatically pick the algorithm
%                                based on the size of the problem.  A
%                                similar strategy to MATLAB's kmedoids
%                                algorithm is used.
%  nClusters:       number of clusters to include
%  clusterAxisIDs:  a cell array of IDs of the axes to cluster  
%  clusterElement:  an N element x 4 cell array of model outputs
%                   in the results to use as a basis.  Each row contains:
%                   elementID, elementType (i.e. parameter, compartment, ....), interventionID, time;
%  clusterID:       which cluster to use, parallel.defaultClusterProfile is default
%  distance:        distance metric to use. 
%                       'correlation': default
%                       'seuclidean':  squared euclidean
%                       'euclidean'
%  normalizeType:   type of method to center and scale the data.  
%                   Valid options: min-max and z-score
%  intSeed:         Integer for seeding the random number generator
%  edgeVPFlag:      Boolean flag.  If true, VPs from the the extremes
%                   of the cluster elements will be used to supplement
%                   the clustering results, to make sure
%                   the range of coverage for all of the axes
%                   and elements is maintained.
%  pointsDefaultLastOnly:   When defining clusters by default method, if true,
%                           for responseTypeElementPoints only the last time
%                           point will be used (boolean)
%  poolClose:       whether to close the pool at the end of the optimization  
%  poolRestart:     whether to restart the pool at the beginning of the optimization 
%  maxIter:         Maximum number of iterations
%  nWorkers:        how many workers in the pool to use.  Set to nan to
%                    use the default (default).
%  replicates:      Number of replicates
%  verbose:         Boolean flag.  Whether to return more information to
%                   screen

   properties
	  algorithm       
      nClusters
      clusterAxisIDs
      clusterElement
      clusterID	  
      distance
      normalizeType
      intSeed
      edgeVPFlag
      pointsDefaultLastOnly
	  poolClose
	  poolRestart		  
	  maxIter
      nWorkers	  
	  replicates
      verbose
   end
   
   methods
      function obj = set.algorithm(obj,myAlgorithm)
          allowedSettings = {'pam','clara','small','large','auto'};
          if sum(ismember(allowedSettings, lower(myAlgorithm))>0)
              obj.algorithm = lower(myAlgorithm);
          else
              error(['Invalid algorithm specified for ',mfilename,', should specify one of: ',strjoin(allowedSettings,', '),'.'])
          end
      end 	         
       
      function obj = set.nClusters(obj,myNClusters)
          if ismatrix(myNClusters)
              if isequal(size(myNClusters),[1 1])
                  if myNClusters > 0
                    % Enforce integer
                    obj.nClusters = round(myNClusters);
                  else
                    error(['Invalid nClusters specified for ',mfilename,', a positive integer must be specified.'])
                  end
              else
                  error(['Invalid nClusters specified for ',mfilename,', a positive integer must be specified.'])
              end
          else
              error(['Invalid nClusters specified for ',mfilename,', a positive integer must be specified.'])
          end
      end 
      
      function obj = set.clusterAxisIDs(obj,myClusterAxisIDs)
          if iscell(myClusterAxisIDs)
              obj.clusterAxisIDs = (myClusterAxisIDs);
          else
              error(['Invalid clusterAxisIDs specified for ',mfilename,', a cell array of axis ID strings should be specified.'])
          end
      end

      function obj = set.clusterElement(obj,myClusterElement)
          if iscell(myClusterElement)
              % Cluster 
              [~, nCols] = size(myClusterElement);
              if nCols == 4
                  obj.clusterElement = myClusterElement;
              else
                  error(['Invalid clusterElement specified for ',mfilename,', an Nx4 cell array of form {elementID, elementType, interventionID, time; ...} should be specified.'])
              end
          else
              error(['Invalid clusterElement specified for ',mfilename,', an Nx4 cell array of form {elementID, elementType, interventionID, time; ...} should be specified.'])
          end
      end
	  
      function obj = set.clusterID(obj,myClusterID)
          if ischar(myClusterID)
              obj.clusterID = myClusterID;
          else
            error(['Invalid clusterID in ',mfilename,'.'])
          end
      end  

      function obj = set.normalizeType(obj,myNormalizeType)
          allowedSettings = {'min-max','z-score'};
          if sum(ismember(allowedSettings, lower(myNormalizeType))>0)
              obj.normalizeType = lower(myNormalizeType);
          else
              error(['Invalid normalizeType specified for ',mfilename,', should specify one of: ',strjoin(allowedSettings,', '),'.'])
          end
      end 

      function obj = set.intSeed(obj,myIntSeed)
          if ((mod(myIntSeed,1) == 0) && (myIntSeed>=-1))
              obj.intSeed = (myIntSeed);
          else
              error(['Invalid intSeed specified for ',mfilename,', a non-negative integer should be specified, or -1 to ignore.'])
          end
      end
      
      function obj = set.edgeVPFlag(obj,myEdgeVPFlag)
          if islogical(myEdgeVPFlag)
              obj.edgeVPFlag = myEdgeVPFlag;
          else
              error(['Invalid edgeVPFlag specified for ',mfilename,', true/false should be specified.'])
          end
      end
      
      function obj = set.pointsDefaultLastOnly(obj,myPointsDefaultLastOnly)
          if islogical(myPointsDefaultLastOnly)
              obj.pointsDefaultLastOnly = myPointsDefaultLastOnly;
          else
              error(['Invalid pointsDefaultLastOnly specified for ',mfilename,', a boolean (true/false) should be specified.'])
          end
      end    

      function obj = set.poolClose(obj,myInput)
          if islogical(myInput)
               obj.poolClose = myInput;
          else
               error(['Property poolClose in ',mfilename,' must be logical.'])
          end
      end  	  
	  
      function obj = set.poolRestart(obj,myInput)
          if islogical(myInput)
               obj.poolRestart = myInput;
          else
               error(['Property poolRestart in ',mfilename,' must be logical.'])
          end
      end 	  
	  
      function obj = set.maxIter(obj,myMaxIter)
          if ismatrix(myMaxIter)
              if isequal(size(myMaxIter),[1 1])
                  if myMaxIter > 0
                    % Enforce integer
                    obj.maxIter = round(myMaxIter);
                  else
                    error(['Invalid maxIter specified for ',mfilename,', a positive integer must be specified.'])
                  end
              else
                  error(['Invalid maxIter specified for ',mfilename,', a positive integer must be specified.'])
              end
          else
              error(['Invalid maxIter specified for ',mfilename,', a positive integer must be specified.'])
          end
      end 	  
	  
      function obj = set.nWorkers(obj,myNWorkers)
          if (isnumeric(myNWorkers) == true)
              obj.nWorkers = myNWorkers;
          else
            error(['Invalid nWorkers value in ',mfilename,'.'])
          end
      end
	  
      function obj = set.replicates(obj,myReplicates)
          if ismatrix(myReplicates)
              if isequal(size(myReplicates),[1 1])
                  if myReplicates > 0
                    % Enforce integer
                    obj.replicates = round(myReplicates);
                  else
                    error(['Invalid replicates specified for ',mfilename,', a positive integer must be specified.'])
                  end
              else
                  error(['Invalid replicates specified for ',mfilename,', a positive integer must be specified.'])
              end
          else
              error(['Invalid replicates specified for ',mfilename,', a positive integer must be specified.'])
          end
      end 	  
      
      function obj = set.distance(obj,myInput)
          allowedSettings = {'correlation','euclidean','seuclidean'};
          if sum(ismember(allowedSettings, lower(myInput))>0)
              obj.distance = lower(myInput);
          else
              error(['Invalid distance metric specified for ',mfilename,', should specify one of: ',strjoin(allowedSettings,', '),'.'])
          end
      end      
      
      function obj = set.verbose(obj,myInput)
          if islogical(myInput)
               obj.verbose = myInput;
          else
               error(['Property verbose in ',mfilename,' must be logical.'])
          end
      end        
     
      function value = get(obj,propName)
          switch propName
              case 'algorithm'
                  value = obj.algorithm;                
              case 'nClusters'
                  value = obj.nClusters;
              case 'clusterAxisIDs'
                  value = obj.clusterAxisIDs;
              case 'clusterElement'
                  value = obj.clusterElement;
              case 'clusterID'
                  value = obj.clusterID; 
              case 'distance'
                  value = obj.distance;                    
              case 'normalizeType'
                  value = obj.normalizeType;                  
              case 'intSeed'
                  value = obj.intSeed;
              case 'edgeVPFlag'
                  value = obj.edgeVPFlag;               
              case 'pointsDefaultLastOnly'
                  value = obj.pointsDefaultLastOnly;   
              case 'poolRestart'
                  value = obj.poolRestart;
              case 'poolClose'
                  value = obj.poolClose;				  
              case 'maxIter'
                  value = obj.maxIter;  
              case 'nWorkers'
                  value = obj.nWorkers;  				  
              case 'replicates'
                  value = obj.replicates;  	
              case 'verbose'
                  value = obj.verbose;                   
              otherwise
                  error(['Error: ',propName ,' is not a valid ',mfilename,' property.'])
          end
      end
      
      function passCheck = verify(obj,myWorksheet)
          passCheck = true;
          allVPIDs = getVPIDs(myWorksheet);
          nWshVPs = length(allVPIDs);
          myNClusters = obj.nClusters;
          if myNClusters > nWshVPs
              warning(['Specified nClusters must be <=  in the number of worksheet VPs in ',mfilename,'.'])
              passCheck = false;
          end
          allAxisIDs = getAxisDefIDs(myWorksheet);
          if sum(ismember(allAxisIDs, obj.clusterAxisIDs)) < length(obj.clusterAxisIDs)
              warning(['Specified clusterAxisIDs should all be specified from axisIDs the worksheet ',mfilename,'.'])
              passCheck = false;
          end
          if ~(isequal(class(myWorksheet.compiled.model),'SimBiology.export.Model'))
              myWorksheet = compileModel(myWorksheet);
              if ~(isequal(class(myWorksheet.compiled.model),'SimBiology.export.Model'))
                  warning(['Unable to compile model associated with myWorksheet in ',mfilename,'.'])
                  passCheck = false;
              end
          end
          if passCheck
              elementInterventionsToCheck = (obj.clusterElement(:,3));
              interventionsToCheck = unique(elementInterventionsToCheck);
              timesToCheck = obj.clusterElement(:,4);
              modelElements = myWorksheet.compiled.elements;
              modelElements = modelElements(:,1:2);   
              modelElements = cell2table(modelElements,'VariableNames',{'ONE','TWO'});
              myElements = obj.clusterElement(:,1:2);
              myElements = cell2table(myElements,'VariableNames',{'ONE','TWO'});
              % This is simplified, we simply check that all unique
              % elementName, elementType pairs in the 
              % cluster elements are also model.
              % If required variables are not recorded in the results,
              % this could create problems.
              elementList = union(modelElements,myElements);
              flagPresent = ismember(modelElements,myElements,'rows');
              [nUnique,~] = size(unique(myElements,'rows')); 
              if sum(flagPresent) ~= nUnique
                  warning(['Specified clusterElement ID, type pairs not all present in myWorksheet in ',mfilename,'.'])
                  passCheck = false;
              end
              modelInterventionIDs = getInterventionIDs(myWorksheet);
              if sum(ismember(modelInterventionIDs,interventionsToCheck)) ~= length(interventionsToCheck)
                  warning(['Specified clusterElement intervention IDs not all present in myWorksheet in ',mfilename,'.'])
                  passCheck = false;
              end           
              if sum(~((cell2mat(timesToCheck))>=0)) > 0
                  % We could do more to actually check if these times are in
                  % the results...                  
                  warning(['Specified clusterElement times are not all non-negative in ',mfilename,'.'])
                  passCheck = false;
              end
              
			  [nClusterElements, ~] = size(obj.clusterElement);
              nClusterAxis = length(obj.clusterAxisIDs);
              
              if obj.edgeVPFlag == true
                  % I should write a better algorithm for getting
                  % the edge VPs and call that here, then check
                  % how big the edge VP set is against these
                  % values.
                  if nWshVPs < 2 * (nClusterElements + nClusterAxis)
                    warning(['There are insufficient VPs in available to ensure enforcement of edgeVPFlag == true in ',mfilename,'.'])
                    passCheck = false;                      
                  end
                  if myNClusters < 2 * (nClusterElements + nClusterAxis)
                    warning(['When edgeVPFlag == true in ',mfilename,', the specified number of clusters includes the number of VPs from edge elements. At least 2x the number of elements and axis should be specified for nClusters.'])
                    passCheck = false;                      
                  end                  
              end     

			  % Also check that all of the outputs are populated
			  keepIndices = ones(nClusterAxis+nClusterElements,1);
			  allInterventionIDs = getInterventionIDs(myWorksheet);
			  for theElementCounter = 1 : nClusterElements
				  theElementName = obj.clusterElement{theElementCounter, 1};
				  theElementType = obj.clusterElement{theElementCounter, 2};
				  theInterventionID = obj.clusterElement{theElementCounter, 3};
				  theSampleTime = obj.clusterElement{theElementCounter, 4};
				  theInterventionIndex = find(ismember(allInterventionIDs, theInterventionID));
				  % All VPs should be run with the same model, with the same result
				  % elements
				  theVPCounter = 1;
				  curResultStruct = myWorksheet.results{theInterventionIndex,theVPCounter};
				  timeIndex = find(ismember(curResultStruct.Names, 'time'));
				  % We assume element IDs are unique, even without type information.
				  elementIndex = find(ismember(curResultStruct.Names, theElementName));
				  for theVPCounter = 1 : nWshVPs
					  curResultStruct = myWorksheet.results{theInterventionIndex,theVPCounter};
					  curTime = curResultStruct.Data(:,timeIndex);
					  curElement = curResultStruct.Data(:,elementIndex);
					  y_new = interp1(curTime,curElement,theSampleTime,'linear');
					  theMatrixToCluster(nClusterAxis+theElementCounter,theVPCounter)=y_new;
				  end
				  curNans = isnan(theMatrixToCluster(nClusterAxis+theElementCounter,:));
				  if sum(curNans) > 0
					  warning(['Unable to cluster element ',theElementName,' of type ',theElementType,' for intervention ',theInterventionID,' at time ',num2str(theSampleTime),' due to NaNs.'])
					  keepIndices(nClusterAxis+theElementCounter) = nan;
					  passCheck = false; 
				  end
			  end					  	  
          end
      end

      function obj = setDefaultFromWorksheet(obj,myWorksheet)
          % This method sets default axes and clustering elements 
          % based on the axes and responseTypes in the worksheet
          % 
          %
          % Arguments:
          %  myWorksheet:        a worksheet that will be used for
          %                      clustering
          %
          % Returns:
          %  An object with updated cluster settings
          %          
          allVPIDs = getVPIDs(myWorksheet);
          nWshVPs = length(allVPIDs);
          obj.normalizeType = 'min-max';
          obj.clusterAxisIDs = getAxisDefIDs(myWorksheet);
          myClusterElement = cell(0, 4);
          for responseTypeCounter = 1 : length(myWorksheet.responseTypes)
              curResponseType = myWorksheet.responseTypes{responseTypeCounter};
              
              for responseTypeElementCounter = 1 :length(curResponseType.elements)
                      curElement = curResponseType.elements{responseTypeElementCounter};
                      if strcmp(class(curElement),'responseTypeElementPoints')
                          expData = (getExpData(curElement.expDataID, myWorksheet));
                          if obj.pointsDefaultLastOnly
                            clusterTimes = max(expData.(curElement.expDataTimeVar));
                          else
                            clusterTimes = unique(expData.(curElement.expDataTimeVar));
                            clusterTimes = sort(clusterTimes(find(~isnan(clusterTimes))),'ascend');
                          end
                          for timeCounter = 1 : length(clusterTimes)
                            clusterTime = clusterTimes(timeCounter);
                            newClusterElement = {curElement.modelYVar, curElement.modelYVarType, curElement.interventionID, clusterTime};
                            [nPrevious, ~] = size(myClusterElement);
                            if nPrevious > 0
                                columnSum = ismember(myClusterElement(:,1),newClusterElement{1}) + ismember(myClusterElement(:,2),newClusterElement{2}) + ismember(myClusterElement(:,3),newClusterElement{3}) + (cell2mat(myClusterElement(:,4))==newClusterElement{4});
                                if max(columnSum) < 4
                                    myClusterElement = cat(1, myClusterElement,newClusterElement);
                                end
                            else
                              myClusterElement = cat(1, myClusterElement,newClusterElement);
                            end
                          end
                      elseif strcmp(class(curElement),'responseTypeElementAxis')
                          % This is a pass-through, we already cluster
                          % based on axis by default
                          myClusterElement = myClusterElement;
                      elseif strcmp(class(curElement),'responseTypeElementBounds')
                          time01 = curElement.referenceTime(1);
                          time02 = curElement.referenceTime(2);
                          if isinf(time01)
                              time01 = 0;
                          end
                          if isinf(time02)
                              time02 = max(myWorksheet.simProps.sampleTimes);
                          elseif time02 > max(myWorksheet.simProps.sampleTimes)
                              time02 = max(myWorksheet.simProps.sampleTimes);
                              warning(['Response type ',curResponseType.id,' responseTypeElementBounds element ',curElement.id,' set to record past the last sample timepoint.  Using the last for clustering.'])
                          end
                          clusterTime = time01;
                          myClusterElement = cat(1, myClusterElement,{curElement.modelYVar, curElement.modelYVarType, curElement.interventionID, clusterTime});
                          clusterTime = time02;
                          myClusterElement = cat(1, myClusterElement,{curElement.modelYVar, curElement.modelYVarType, curElement.interventionID, clusterTime});
                      else
                          error(['Currently, only responseTypeElementPoints, responseTypeElementBounds supported in ',mfilename,'.'])
                      end
              end
          end
          obj.clusterElement = myClusterElement;
      
      
          % We'll default to 4x the number of clusterAxes and Elements
          [nClusterElements, ~] = size(obj.clusterElement);
          nClusterAxis = length(obj.clusterAxisIDs);
          obj.nClusters = min(4 * (nClusterElements + nClusterAxis), nWshVPs);

          % We'll only set edgeVPFlag if it looks like there
          % are enough VPs to support.
          if nWshVPs >= 2 * (nClusterElements + nClusterAxis)
              obj.edgeVPFlag = true;
          else
              obj.edgeVPFlag = false;
          end
      end
      
      
      function obj = setClusterElementFromOptions(obj, myObj)
          % This method sets default clustering elements based on the 
          % VPop or mapelOptions object.  Rather than using
          % response types, the targets 
          % will be used to set the "element" clustering.
          % You can call "setDefaultFromWorksheet" first
          % to set the axes to cluseter on, and this will overwrite the
          % elements from the call to "setDefaultFromWorksheet."
          %
          % Arguments:
          %  myObj:        a mapelOptions or VPop object
          %
          % Returns:
          %  An object with updated cluster settings
          %
          
          obj.normalizeType = 'min-max';
          myClusterElement = cell(0, 4);
          % We won't consider weight, we will just use all timepoint,
          % variables, interventions of potential interest
          if ~isempty(myObj.mnSDTable)
              myClusterElement = [myClusterElement; table2cell(myObj.mnSDTable(:,{'elementID','elementType','interventionID','time'}))];
          end
          if ~isempty(myObj.distTable)
              myClusterElement = [myClusterElement; table2cell(myObj.distTable(:,{'elementID','elementType','interventionID','time'}))];
          end
          if ~isempty(myObj.binTable)
              myClusterElement = [myClusterElement; table2cell(myObj.binTable(:,{'elementID','elementType','interventionID','time'}))];
          end
          if ~isempty(myObj.distTable2D)
              myClusterElement = [myClusterElement; table2cell(myObj.distTable2D(:,{'elementID1','elementType1','interventionID1','time1'}))];
              myClusterElement = [myClusterElement; table2cell(myObj.distTable2D(:,{'elementID2','elementType2','interventionID2','time2'}))];
          end
          if ~isempty(myObj.corTable)
              myClusterElement = [myClusterElement; table2cell(myObj.corTable(:,{'elementID1','elementType1','interventionID1','time1'}))];
              myClusterElement = [myClusterElement; table2cell(myObj.corTable(:,{'elementID2','elementType2','interventionID2','time2'}))];
          end
          [~,idx]=unique(cell2table(myClusterElement),'rows');
          myClusterElement = myClusterElement(idx,:);
          obj.clusterElement = myClusterElement;
      
          % We'll only set edgeVPFlag if it looks like there
          % are enough VPs to support.
          [nSubpops, ~] = size(myObj.subpopTable);
          if nSubpops > 0
              nWshVPs = length(myObj.subpopTable{1,'vpIDs'}{1});
              % We'll default to 4x the number of clusterAxes and Elements
              [nClusterElements, ~] = size(obj.clusterElement);
              nClusterAxis = length(obj.clusterAxisIDs);
              obj.nClusters = min(4 * (nClusterElements + nClusterAxis), nWshVPs);              
              if nWshVPs >= 2 * (nClusterElements + nClusterAxis)
                  obj.edgeVPFlag = true;
              else
                  obj.edgeVPFlag = false;
              end
          else
              disp(['Subpopulation table not yet defined in mapelOptions or VPop object provided in setClusterElementFromOptions.  Not updating edgeVPFlag or nClusters for ',mfilename,'.'])
          end
      end      
      
      
      
      % The constructor method must have the same name as the class
      function obj = clusterPickOptions()
	      obj.algorithm = 'auto';          
          obj.nClusters = 1;
          obj.clusterAxisIDs = {};
          obj.clusterElement = cell(0,4);
          obj.clusterID = parallel.defaultClusterProfile;		  
	      obj.distance = 'correlation';          
          obj.normalizeType = 'min-max';
          obj.intSeed = -1;
          obj.edgeVPFlag = false;
          obj.pointsDefaultLastOnly = true;
          obj.poolClose = true;  
		  obj.poolRestart = true;		  
		  obj.maxIter = 1000;
          obj.nWorkers = nan;		  
	      obj.replicates = 10;
          obj.verbose = true;
      end
    end
end
```
