## Supplementary File 3 for "A surrogate workflow for virtual population development: designing an amenable proxy for a hypothesis-test inspired nonlinear goodness-of-fit": clusterTestOptions.m

```
classdef clusterTestOptions
% This class defines options for testClusterSizes
%
% PROPERTIES:
%  rangeNClusters:   set the range of cluster sizes to test.  A 1x2 matrix of integers.
%  clusterAxisIDs:   specify any axes that will be used as a basis for clustering.
%  clusterElement:   an N element x 4 cell array of model outputs
%                    in the results to use as a basis.  Each row contains:
%                    elementID, elementType (i.e. parameter, compartment, ....), interventionID, time;
%  distance:        distance metric to use. 
%                       'correlation': default
%                       'seuclidean':  squared euclidean
%  normalizeType:    type of method to center and scale the data.  
%                    Valid options: min-max and z-score
%  intSeed:          Integer for seeding the random number generator
%  clusterCutoff:    A measure between 0 and 1 of the amount of variability in
%                    the original cohort to maintain.  The nearest number of
%                    clusters will be included in the output.
%  pointsDefaultLastOnly:   When defining clusters by default method, if true,
%                           for responseTypeElementPoints only the last time
%                           point will be used (boolean)

   properties
      rangeNClusters
      clusterAxisIDs
      clusterElement
      distance
      normalizeType
      intSeed
      clusterCutoff
      pointsDefaultLastOnly
   end
   
   methods
      function obj = set.rangeNClusters(obj,myRangeNClusters)
          if ismatrix(myRangeNClusters)
              if isequal(size(myRangeNClusters),[1 2])
                  % Enforce integer
                  obj.rangeNClusters = round(myRangeNClusters);
              else
                  error(['Invalid rangeNClusters specified for ',mfilename,', a 1 x 2 numeric matrix must be specified.'])
              end
          else
              error(['Invalid rangeNClusters specified for ',mfilename,', a 1 x 2 numeric matrix must be specified.'])
          end
      end
      
      function obj = set.clusterCutoff(obj,myClusterCutoff)
        if (isequal(size(myClusterCutoff), [1 1]) && (myClusterCutoff > 0) && (myClusterCutoff < 1))
            obj.clusterCutoff = myClusterCutoff;
        else
            error(['Invalid clusterCutoff specified for ',mfilename,', a number > 0 and < 1 should be specified.'])
        end
      end
      
      function obj = set.clusterAxisIDs(obj,myClusterAxisIDs)
          if iscell(myClusterAxisIDs)
              obj.clusterAxisIDs = (myClusterAxisIDs);
          else
              error(['Invalid clusterAxisIDs specified for ',mfilename,', a cell array of axis ID strings should be specified.'])
          end
      end

      function obj = set.clusterElement(obj,myclusterElement)
          if iscell(myclusterElement)
              % Cluster 
              [~, nCols] = size(myclusterElement);
              if nCols == 4
                  obj.clusterElement = myclusterElement;
              else
                  error(['Invalid clusterElement specified for ',mfilename,', an Nx3 cell array of form {elementID, elementType, interventionID; ...} should be specified.'])
              end
          else
              error(['Invalid clusterElement specified for ',mfilename,', an Nx3 cell array of form {elementID, elementType, interventionID; ...} should be specified.'])
          end
      end
      
      function obj = set.distance(obj,myInput)
          allowedSettings = {'correlation','seuclidean'};
          if sum(ismember(allowedSettings, lower(myInput))>0)
              obj.distance = lower(myInput);
          else
              error(['Invalid distance metric specified for ',mfilename,', should specify one of: ',strjoin(allowedSettings,', '),'.'])
          end
      end      

      function obj = set.normalizeType(obj,myNormalizeType)
          allowedSettings = {'min-max','z-score'};
          if sum(ismember(allowedSettings, lower(myNormalizeType))>0)
              obj.normalizeType = lower(myNormalizeType);
          else
              error(['Invalid normalizeType specified for ',mfilename,', should specify one of: ',strjoin(allowedSettings,', '),'.'])
          end
      end 

      function obj = set.intSeed(obj,myIntSeed)
          if ((mod(myIntSeed,1) == 0) && (myIntSeed>=-1))
              obj.intSeed = (myIntSeed);
          else
              error(['Invalid intSeed specified for ',mfilename,', a non-negative integer should be specified, or -1 to ignore.'])
          end
      end
      
      function obj = set.pointsDefaultLastOnly(obj,myPointsDefaultLastOnly)
          if islogical(myPointsDefaultLastOnly)
              obj.pointsDefaultLastOnly = myPointsDefaultLastOnly;
          else
              error(['Invalid pointsDefaultLastOnly specified for ',mfilename,', a boolean (true/false) should be specified.'])
          end
      end      
     
      function value = get(obj,propName)
          switch propName
              case 'rangeNClusters'
                  value = obj.rangeNClusters;
              case 'clusterAxisIDs'
                  value = obj.clusterAxisIDs;
              case 'clusterElement'
                  value = obj.clusterElement;  
              case 'distance'
                  value = obj.distance;                    
              case 'normalizeType'
                  value = obj.normalizeType;                  
              case 'intSeed'
                  value = obj.intSeed;
              case 'clusterCutoff'
                  value = obj.clusterCutoff;
              case 'pointsDefaultLastOnly'
                  value = obj.pointsDefaultLastOnly;
              otherwise
                  error(['Error: ',propName ,' is not a valid ',mfilename,' property.'])
          end
      end
      
      function passCheck = verify(obj,myWorksheet)
          passCheck = true;
          allVPIDs = getVPIDs(myWorksheet);
          myRangeNClusters = obj.rangeNClusters;
          if myRangeNClusters(2) < myRangeNClusters(1)
              warning(['Specified rangeNClusters(2) should be >= rangeNClusters(2) in ',mfilename,'.'])
              passCheck = false;
          end
          if myRangeNClusters(1) < 1
              warning(['Specified rangeNClusters(1) should be > 1 in ',mfilename,'.'])
              passCheck = false;
          end
          if myRangeNClusters(2) > length(allVPIDs)
              warning(['Specified rangeNClusters(2) should be <=  in the number of worksheet VPs in ',mfilename,'.'])
              passCheck = false;
          end
          if myRangeNClusters(2) < 2 
              warning(['Specified rangeNClusters(2) should be >= 2 in ',mfilename,'.'])
              passCheck = false;
          end
          allAxisIDs = getAxisDefIDs(myWorksheet);
          if sum(ismember(allAxisIDs, obj.clusterAxisIDs)) < length(obj.clusterAxisIDs)
              warning(['Specified clusterAxisIDs should all be specified from axisIDs the worksheet ',mfilename,'.'])
              passCheck = false;
          end
          if ~(isequal(class(myWorksheet.compiled.model),'SimBiology.export.Model'))
              myWorksheet = compileModel(myWorksheet);
              if ~(isequal(class(myWorksheet.compiled.model),'SimBiology.export.Model'))
                  warning(['Unable to compile model associated with myWorksheet in ',mfilename,'.'])
                  passCheck = false;
              end
          end
          if passCheck
              elementInterventionsToCheck = (obj.clusterElement(:,3));
              interventionsToCheck = unique(elementInterventionsToCheck);
              timesToCheck = obj.clusterElement(:,4);
              modelElements = myWorksheet.compiled.elements;
              modelElements = modelElements(:,1:2);   
              modelElements = cell2table(modelElements,'VariableNames',{'ONE','TWO'});
              myElements = obj.clusterElement(:,1:2);
              myElements = cell2table(myElements,'VariableNames',{'ONE','TWO'});
              % This is simplified, we simply check that all unique
              % elementName, elementType pairs in the 
              % cluster elements are also model.
              % If required variables are not recorded in the results,
              % this could create problems.
              elementList = union(modelElements,myElements);
              flagPresent = ismember(modelElements,myElements,'rows');
              [nUnique,~] = size(unique(myElements,'rows')); 
              if sum(flagPresent) ~= nUnique
                  warning(['Specified clusterElement ID, type pairs not all present in myWorksheet in ',mfilename,'.'])
                  passCheck = false;
              end
              modelInterventionIDs = getInterventionIDs(myWorksheet);
              if sum(ismember(modelInterventionIDs,interventionsToCheck)) ~= length(interventionsToCheck)
                  warning(['Specified clusterElement intervention IDs not all present in myWorksheet in ',mfilename,'.'])
                  passCheck = false;
              end           
              if sum(~((cell2mat(timesToCheck))>=0)) > 0
                  % We could do more to actually check if these times are in
                  % the results...                  
                  warning(['Specified clusterElement times are not all non-negative in ',mfilename,'.'])
                  passCheck = false;
              end
          end
      end

      function obj = setDefaultFromWorksheet(obj,myWorksheet)
          allVPIDs = getVPIDs(myWorksheet);
          nWshVPs = length(allVPIDs);
          obj.normalizeType = 'min-max';
          obj.clusterAxisIDs = getAxisDefIDs(myWorksheet);
          myClusterElement = cell(0, 4);
          for responseTypeCounter = 1 : length(myWorksheet.responseTypes)
              curResponseType = myWorksheet.responseTypes{responseTypeCounter};
              
              for responseTypeElementCounter = 1 :length(curResponseType.elements)
                      curElement = curResponseType.elements{responseTypeElementCounter};
                      if strcmp(class(curElement),'responseTypeElementPoints')
                          expData = (getExpData(curElement.expDataID, myWorksheet));
                          if obj.pointsDefaultLastOnly
                            clusterTimes = max(expData.(curElement.expDataTimeVar));
                          else
                            clusterTimes = unique(expData.(curElement.expDataTimeVar));
                            clusterTimes = sort(clusterTimes(find(~isnan(clusterTimes))),'ascend');
                          end
                          for timeCounter = 1 : length(clusterTimes)
                            clusterTime = clusterTimes(timeCounter);
                            newClusterElement = {curElement.modelYVar, curElement.modelYVarType, curElement.interventionID, clusterTime};
                            [nPrevious, ~] = size(myClusterElement);
                            if nPrevious > 0
                                columnSum = ismember(myClusterElement(:,1),newClusterElement{1}) + ismember(myClusterElement(:,2),newClusterElement{2}) + ismember(myClusterElement(:,3),newClusterElement{3}) + (cell2mat(myClusterElement(:,4))==newClusterElement{4});
                                if max(columnSum) < 4
                                    myClusterElement = cat(1, myClusterElement,newClusterElement);
                                end
                            else
                              myClusterElement = cat(1, myClusterElement,newClusterElement);
                            end
                          end
                      elseif strcmp(class(curElement),'responseTypeElementAxis')
                          % This is a pass-through, we already cluster
                          % based on axis by default
                          myClusterElement = myClusterElement;
                      elseif strcmp(class(curElement),'responseTypeElementBounds')
                          time01 = curElement.referenceTime(1);
                          time02 = curElement.referenceTime(2);
                          if isinf(time01)
                              time01 = 0;
                          end
                          if isinf(time02)
                              time02 = max(myWorksheet.simProps.sampleTimes);
                          elseif time02 > max(myWorksheet.simProps.sampleTimes)
                              time02 = max(myWorksheet.simProps.sampleTimes);
                              warning(['Response type ',curResponseType.id,' responseTypeElementBounds element ',curElement.id,' set to record past the last sample timepoint.  Using the last for clustering.'])
                          end
                          clusterTime = time01;
                          myClusterElement = cat(1, myClusterElement,{curElement.modelYVar, curElement.modelYVarType, curElement.interventionID, clusterTime});
                          clusterTime = time02;
                          myClusterElement = cat(1, myClusterElement,{curElement.modelYVar, curElement.modelYVarType, curElement.interventionID, clusterTime});
                      else
                          error(['Currently, only responseTypeElementPoints, responseTypeElementBounds supported in ',mfilename,'.'])
                      end
              end
          end
          obj.clusterElement = myClusterElement;
      end
      
      % The constructor method must have the same name as the class
      function obj = clusterTestOptions()
          obj.rangeNClusters = [2 2];
          obj.clusterAxisIDs = {};
          obj.clusterElement = cell(0,4);
	      obj.distance = 'correlation';           
%           obj.centerType = 'mean';
%           obj.scaleType = 'std';
          obj.normalizeType = 'min-max';
          obj.intSeed = -1;
          obj.clusterCutoff = 0.85;
          obj.pointsDefaultLastOnly = true;
      end
    end
end
```
