## Supplementary File 3 for "A surrogate workflow for virtual population development: designing an amenable proxy for a hypothesis-test inspired nonlinear goodness-of-fit": runControlCoefficientsSimulationsOptions.m

```
classdef runControlCoefficientsSimulationsOptions
% Options object for running a local control coefficient analysis
%
% PROPERTIES
%  baseVPID:                 (Required) A VP ID from the worksheet must be
%                            a starting parameterization of the model.
%                            Relevant variants will be loaded and the
%                            sensitivity analysis will be applied
%                            across this VP's axes.
%  varyAxisIDs:              (Required) Axes that will be included in the 
%                            sensitivity analysis.
%  interventionID:           (Required) The sensitivity analysis will only 
%                            be performed for one worksheet intervention.
%  saveElementResultIDs:     (Required) Just the selected simulation 
%                            outputs will be
%                            written to the resulting worksheet.  This
%                            helps reduce the size of the output for large 
%                            simulation runs.
%  saveFileName:             (Optional) file to save results to during run.
%                            If set to '', the run will not be saved while
%                            in progress.  Note it may generally be better
%                            not to save to file if I/O times are too
%                            lengthy, which may happen with large
%                            worksheets.  Default is ''.
%  simulateWorksheet:        (Optional) Boolean (true/false), whether to 
%                            run the simulations (or just set up the 
%                            worksheet).  Default is true.
%  verbose:                  (Optional) Boolean (true/false), whether to 
%                            write the run status to screen.  Default is
%                            true.
%
   properties
      baseVPID
      varyAxisIDs
      interventionID
      saveElementResultIDs
      saveFileName
      simulateWorksheet
      verbose
   end
   
   methods

      function obj = set.baseVPID(obj,myBaseVPID)
          if ischar(myBaseVPID)
              obj.baseVPID = myBaseVPID;
          else
              error(['Invalid baseVPID specified for ',mfilename,', a VP ID string should be specified.'])
          end
      end
      
      function obj = set.varyAxisIDs(obj,myVaryAxisIDs)
          if iscell(myVaryAxisIDs)
              obj.varyAxisIDs = (myVaryAxisIDs);
          else
              error(['Invalid varyAxisIDs specified for ',mfilename,', a cell array of axis ID strings should be specified.'])
          end
      end   

      function obj = set.interventionID(obj,myInterventionID)
          if ischar(myInterventionID)
              obj.interventionID = myInterventionID;
          else
              error(['Invalid interventionID specified for ',mfilename,', an intervention ID string should be specified.'])
          end
      end     
      
      function obj = set.saveElementResultIDs(obj,mySaveElementResultIDs)
          if iscell(mySaveElementResultIDs)
              obj.saveElementResultIDs = (mySaveElementResultIDs);
          else
              error(['Invalid saveElementResultIDs specified for ',mfilename,', a cell array of variable ID strings should be specified.'])
          end
      end     
      
      function obj = set.saveFileName(obj,mySaveFileName)
          if ischar(mySaveFileName)
              obj.saveFileName = mySaveFileName;
          else
              error(['Invalid saveFileName specified for ',mfilename,', a file name string should be specified.'])
          end
      end   
      
      function obj = set.simulateWorksheet(obj, mySimulateWorksheet)
          if ~islogical(mySimulateWorksheet)
              warning(strcat('Setting for simulateWorksheet must be logical (true, false) in ',mfilename,'.'))
          else
              obj.simulateWorksheet = mySimulateWorksheet;
          end
      end      
      
      function obj = set.verbose(obj, myVerbose)
          if ~islogical(myVerbose)
              warning(strcat('Setting for verbose must be logical (true, false) in ',mfilename,'.'))
          else
              obj.verbose = myVerbose;
          end
      end
     
      function value = get(obj,propName)
          switch propName
              case 'baseVPID'
                  value = obj.baseVPID;
              case 'nRandomizationsPerSample'
                  value = obj.nRandomizationsPerSample;
              case 'varyAxisIDs'
                  value = obj.varyAxisIDs;                   
              case 'interventionID'
                  value = obj.interventionID;   
              case 'saveElementResultIDs'
                  value = obj.saveElementResultIDs;  
              case 'saveFileName'
                  value = obj.saveFileName; 
              case 'simulateWorksheet'
                  value = obj.simulateWorksheet;                    
              case 'verbose'
                  value = obj.verbose;                     
          end
      end     
      
      function passCheck = verify(obj,myWorksheet)
          passCheck = true;
          allVPIDs = getVPIDs(myWorksheet);
          if sum(ismember(allVPIDs, obj.baseVPID)) < 1
              warning(['Specified baseVPID should all be specified from VPIDs the worksheet in ',mfilename,'.']);
              passCheck = false;
          end
          allAxisDefIDs = getAxisDefIDs(myWorksheet);
          if sum(ismember(allAxisDefIDs, obj.varyAxisIDs)) < length(obj.varyAxisIDs)
              warning(['Specified varyAxisIDs should all be specified from axis IDs the worksheet in ',mfilename,'.']);
              passCheck = false;
          end 
          % All axes should also have just 1 parameter for this
          % method, otherwise we can't gaurantee scaling all of the
          % parameters by a fixed relative percent
          nAxisFail = 0;
          for axisCounter = 1 : length(obj.varyAxisIDs)
              axisIndex = find(ismember(allAxisDefIDs,obj.varyAxisIDs{axisCounter}));
              if length(myWorksheet.axisProps.axisDef{axisIndex}.elementNames) > 1
                  nAxisFail = nAxisFail+1;
              end
              if nAxisFail > 0
                  warning(['Specified varyAxisIDs should all have just 1 element/parameter in ',mfilename,'.'])
                  passCheck = false;
              end
          end
          
          allInterventionIDs = getInterventionIDs(myWorksheet);
          if sum(ismember(allInterventionIDs,  obj.interventionID)) < 1
              warning(['Specified interventionID should be selected from interventionIDs the worksheet in ',mfilename,'.']);
              passCheck = false;
          end       
          
          if ~(isequal(class(myWorksheet.compiled.model),'SimBiology.export.Model'))
              myWorksheet = compileModel(myWorksheet, false);
              if ~(isequal(class(myWorksheet.compiled.model),'SimBiology.export.Model'))
                  warning(['Unable to compile model associated with myWorksheet in ',mfilename,'.'])
                  passCheck = false;
              end
          end
          
          if passCheck
              % Ideally, we only want to select saveElementResultIDs from
              % variables that are written to results, but this wasn't
              % immediately clear to me where these would be stored in the
              % SimBiology model object prior to obtaining results.
              % At least we can check the specified variables are
              % recognized as elements.
              if(sum(ismember(obj.saveElementResultIDs,myWorksheet.compiled.elements(:,1))) < length(obj.saveElementResultIDs))
                  warning(['Unable to identify all indicated saveElementResultIDs as elements in ',mfilename,'.'])
                  passCheck = false;
              end
          end
          
      end
      
      % The constructor method must have the same name as the class
      function obj = runControlCoefficientsSimulationsOptions() 
          obj.baseVPID = '';
          obj.varyAxisIDs={};
          obj.interventionID='';
          obj.saveElementResultIDs={};
          obj.saveFileName='';
          obj.simulateWorksheet=true;
          obj.verbose = true;
      end
   end
    
end
```
