## Supplementary File 3 for "A surrogate workflow for virtual population development: designing an amenable proxy for a hypothesis-test inspired nonlinear goodness-of-fit": sobolSampleOptions.m

```
classdef sobolSampleOptions
% Options object for running Sobol sampling.  Note that the sensitivity
% analysis must subsequently be run on the returned worksheet. 
%
% PROPERTIES
%  baseVPID:                 (Required) A VP ID from the worksheet must be
%                            a starting parameterization of the model.
%                            Relevant variants will be loaded and the
%                            sensitivity analysis will be applied
%                            across this VP's axes.
%  nRandomizationsPerSample: (Optional) This determines how many sampling 
%                            points will will be used.  Note that after 
%                            sampling we can check the confidence intervals 
%                            to verify a suitable sampling size has been 
%                            selected. In Saltelli's method for first and 
%                            total sensitivity indices, we run N * (k + 2) 
%                            simulations total, where N is 
%                            nRandomizationsPerSample and k is the
%                            number of axes to vary.  Default: 1000,
%                            more likely needed.
%  varyAxisIDs:              (Required) Axes that will be included in the 
%                            sensitivity analysis.
%  interventionID:           (Required) The sensitivity analysis will only 
%                            be performed for one worksheet intervention.
%  saveElementResultIDs:     (Required) Just the selected simulation 
%                            outputs will be
%                            written to the resulting worksheet.  This
%                            helps reduce the size of the output for large 
%                            simulation runs. 
%  saveFileName:             (Optional) file to save results to during run.
%                            If set to '', the run will not be saved while
%                            in progress.  Note it may generally be better
%                            not to save to file if I/O times are too
%                            lengthy, which may happen with large
%                            worksheets.  Default is ''.
%  maxBatchSimulateN:        (Optional) The sampling simulations will be 
%                            sent to simulateWorksheet in sizes not exceeding
%                            maxBatchSimulateN.  Reducing maxBatchSimulateN
%                            may help reduce maximum memory utilization
%                            due to memory management and distributing
%                            over available cores at the expense of 
%                            longer computation times due the additional
%                            data manipulation.  Default is 250, you
%                            will likely want to try running at
%                            >= N * (k + 2) to try to complete the sampling
%                            in one run.
%  intSeed:                  (Optional) A non-negative integer seed to  
%                            initialize the random number generator.  Set 
%                            to -1 to avoid changing the state of the 
%                            random number generator.  Default is -1.
%  simulateWorksheet:           (Optional) Boolean (true/false), whether to 
%                            run the simulations (or just set up the 
%                            worksheet).  Default is true.
%  verbose:                  (Optional) Boolean (true/false), whether to 
%                            write the run status to screen.  Default is
%                            true.
%
   properties
      baseVPID;
      nRandomizationsPerSample;
      varyAxisIDs;
      interventionID;
      saveElementResultIDs;
      saveFileName;
      maxBatchSimulateN;
      intSeed;
      simulateWorksheet;
      verbose;
   end
   
   methods

      function obj = set.baseVPID(obj,myBaseVPID)
          if ischar(myBaseVPID)
              obj.baseVPID = myBaseVPID;
          else
              error(['Invalid baseVPID specified for ',mfilename,', a VP ID string should be specified.'])
          end
      end

      function obj = set.nRandomizationsPerSample(obj,myNRandomizationsPerSample)
          failFlag = false;
          if isnumeric(myNRandomizationsPerSample)
              if isequal(size(myNRandomizationsPerSample),[1 1])
                  if myNRandomizationsPerSample > 0
                      obj.nRandomizationsPerSample = round(myNRandomizationsPerSample);
                  else
                      failFlag = true;
                  end
              else
                  failFlag = true;
              end
          else
              failFlag = true;
          end
          if failFlag
              error(['Invalid myNRandomizationsPerSample specified for ',mfilename,'. A positive number should be specified.'])
          end
      end
      
      function obj = set.varyAxisIDs(obj,myVaryAxisIDs)
          if iscell(myVaryAxisIDs)
              obj.varyAxisIDs = (myVaryAxisIDs);
          else
              error(['Invalid varyAxisIDs specified for ',mfilename,', a cell array of axis ID strings should be specified.'])
          end
      end   

      function obj = set.interventionID(obj,myInterventionID)
          if ischar(myInterventionID)
              obj.interventionID = myInterventionID;
          else
              error(['Invalid interventionID specified for ',mfilename,', an intervention ID string should be specified.'])
          end
      end     
      
      function obj = set.saveElementResultIDs(obj,mySaveElementResultIDs)
          if iscell(mySaveElementResultIDs)
              obj.saveElementResultIDs = (mySaveElementResultIDs);
          else
              error(['Invalid saveElementResultIDs specified for ',mfilename,', a cell array of variable ID strings should be specified.'])
          end
      end     
      
      function obj = set.saveFileName(obj,mySaveFileName)
          if ischar(mySaveFileName)
              obj.saveFileName = mySaveFileName;
          else
              error(['Invalid saveFileName specified for ',mfilename,', a file name string should be specified.'])
          end
      end   
      
      function obj = set.maxBatchSimulateN(obj,myMaxBatchSimulateN)
          failFlag = false;
          if isnumeric(myMaxBatchSimulateN)
              if isequal(size(myMaxBatchSimulateN),[1 1])
                  if myMaxBatchSimulateN > 0
                      obj.maxBatchSimulateN = round(myMaxBatchSimulateN);
                  else
                      failFlag = true;
                  end
              else
                  failFlag = true;
              end
          else
              failFlag = true;
          end
          if failFlag
              error(['Invalid myMaxBatchSimulateN specified for ',mfilename,'. A positive number should be specified.'])
          end
      end  
      
      function obj = set.intSeed(obj, myRandomSeed)
          if ((myRandomSeed >= -1) && (mod(myRandomSeed,1) == 0))
              obj.intSeed = myRandomSeed;
          else
              warning(['Setting for intSeed for random number generator must be a nonegative integer in ',mfilename,', or -1 to ignore. Refusing to update.']);
          end
      end
      
      function obj = set.simulateWorksheet(obj, mySimulateWorksheet)
          if ~islogical(mySimulateWorksheet)
              warning(strcat('Setting for simulateWorksheet must be logical (true, false) in ',mfilename,'.'))
          else
              obj.simulateWorksheet = mySimulateWorksheet;
          end
      end      
      
      function obj = set.verbose(obj, myVerbose)
          if ~islogical(myVerbose)
              warning(strcat('Setting for verbose must be logical (true, false) in ',mfilename,'.'))
          else
              obj.verbose = myVerbose;
          end
      end
     
      function value = get(obj,propName)
          switch propName
              case 'baseVPID'
                  value = obj.baseVPID;
              case 'nRandomizationsPerSample'
                  value = obj.nRandomizationsPerSample;
              case 'varyAxisIDs'
                  value = obj.varyAxisIDs;                   
              case 'interventionID'
                  value = obj.interventionID;   
              case 'saveElementResultIDs'
                  value = obj.saveElementResultIDs;  
              case 'saveFileName'
                  value = obj.saveFileName; 
              case 'maxBatchSimulateN'
                  value = obj.maxBatchSimulateN; 
              case 'intSeed'
                  value = obj.intSeed;  
              case 'simulateWorksheet'
                  value = obj.simulateWorksheet;                    
              case 'verbose'
                  value = obj.verbose;                     
          end
      end     
      
      function passCheck = verify(obj,myWorksheet)
          passCheck = true;
          allVPIDs = getVPIDs(myWorksheet);
          if sum(ismember(allVPIDs, obj.baseVPID)) < 1
              warning(['Specified baseVPID should all be specified from VPIDs the worksheet in ',mfilename,'.']);
              passCheck = false;
          end
          allAxisDefIDs = getAxisDefIDs(myWorksheet);
          if sum(ismember(allAxisDefIDs, obj.varyAxisIDs)) < length(obj.varyAxisIDs)
              warning(['Specified varyAxisIDs should all be specified from axis IDs the worksheet in ',mfilename,'.']);
              passCheck = false;
          end 
          allInterventionIDs = getInterventionIDs(myWorksheet);
          if sum(ismember(allInterventionIDs,  obj.interventionID)) < 1
              warning(['Specified interventionID should be selected from interventionIDs the worksheet in ',mfilename,'.']);
              passCheck = false;
          end       
          
          if ~(isequal(class(myWorksheet.compiled.model),'SimBiology.export.Model'))
              myWorksheet = compileModel(myWorksheet, false);
              if ~(isequal(class(myWorksheet.compiled.model),'SimBiology.export.Model'))
                  warning(['Unable to compile model associated with myWorksheet in ',mfilename,'.'])
                  passCheck = false;
              end
          end
          
          if passCheck
              % Ideally, we only want to select saveElementResultIDs from
              % variables that are written to results, but this wasn't
              % immediately clear to me where these would be stored in the
              % SimBiology model object prior to obtaining results.
              % At least we can check the specified variables are
              % recongized as elements.
              if(sum(ismember(obj.saveElementResultIDs,myWorksheet.compiled.elements(:,1))) < length(obj.saveElementResultIDs))
                  warning(['Unable to identify all indicated saveElementResultIDs as elements in ',mfilename,'.'])
                  passCheck = false;
              end
          end
          
      end
      
      % The constructor method must have the same name as the class
      function obj = sobolSampleOptions() 
          obj.baseVPID = '';
          obj.nRandomizationsPerSample=1000;
          obj.varyAxisIDs={};
          obj.interventionID='';
          obj.saveElementResultIDs={};
          obj.saveFileName='';
          obj.maxBatchSimulateN = 250;
          obj.intSeed = 0;
          obj.simulateWorksheet=true;
          obj.verbose = true;
      end
   end
    
end
```
