## Supplementary File 3 for "A surrogate workflow for virtual population development: designing an amenable proxy for a hypothesis-test inspired nonlinear goodness-of-fit": sobolSensitivityOptions.m

```
classdef sobolSensitivityOptions
% Options object for running Sobol sensitivity analysis.
% Note that the sensitivity analysis must be run with the returned
% worksheet from the sampling step.
%
% PROPERTIES
% interventionID:          (Required) Worksheet intervention to run
%                          analysis on.
% analyzeElementResultIDs: (Required) Worksheet outputs to run analysis
%                          on.
% analyzeTime:             (Required) Simulation time point to run analysis
%                          at.  If not specified, the analysis will be run
%                          at t=0, which is not very informative.  Also,
%                          this should correstpond to an exact sampled
%                          simulation timepoint as itnerpolation is not
%                          implemented
% subSampleSplitType:      (Optional) If specified, the sensitivity indices
%                          will be recomputed using a subsamples of 
%                          smaller nRandomizationsPerSample.  This is
%                          intended to give a sense whether the sensitivity
%                          indices and confidence intervals have stabilized
%                          Default is 'none'.
% nBootstraps:             (Optional) The sampling results will be
%                          bootstrapped to develop confidence intervals.
%                          The default is 1,000
% intSeed               A non-negative integer seed to initialize the 
%                          random number generator.  Set to -1 to avoid
%                          changing the state of the random number
%                          generator.  Default is -1.
    
   properties
      interventionID;
      analyzeElementResultIDs;
      analyzeTime;
      subSampleSplitType;
      nBootstraps;
      intSeed;      
   end
   
   methods

      function obj = set.interventionID(obj,myInterventionID)
          if ischar(myInterventionID)
              obj.interventionID = myInterventionID;
          else
              error(['Invalid interventionID specified for ',mfilename,', an intervention ID string should be specified.'])
          end
      end     
      
      function obj = set.analyzeElementResultIDs(obj,myAnalyzeElementResultIDs)
          if iscell(myAnalyzeElementResultIDs)
              obj.analyzeElementResultIDs = (myAnalyzeElementResultIDs);
          else
              error(['Invalid analyzeElementResultIDs specified for ',mfilename,', a cell array of variable ID strings should be specified.'])
          end
      end     
      
      function obj = set.analyzeTime(obj,myAnalyzeTime)
          failFlag = false;
          if isnumeric(myAnalyzeTime)
              if isequal(size(myAnalyzeTime),[1 1])
                  obj.analyzeTime = (myAnalyzeTime);
              else
                  failFlag = true;
              end
          else
              failFlag = true;
          end
          if failFlag
              error(['Invalid myAnalyzeTime specified for ',mfilename,'. A nonnegative number should be specified.'])
          end
      end      
      
      function obj = set.subSampleSplitType(obj,mySubSampleSplitType)
          allowedValues = {'none','base2'};
          if sum(ismember(allowedValues,lower(mySubSampleSplitType))) > 0
              obj.subSampleSplitType = lower(mySubSampleSplitType);
          else
              error(['Invalid subSampleSplitType specified for ',mfilename,'. A "none" or "base2" should be specified.'])
          end
      end     
     
      function obj = set.nBootstraps(obj,myNBootstraps)
          failFlag = false;
          if isnumeric(myNBootstraps)
              if isequal(size(myNBootstraps),[1 1])
                  obj.nBootstraps = (myNBootstraps);
              else
                  failFlag = true;
              end
          else
              failFlag = true;
          end
          if failFlag
              error(['Invalid nBootstraps specified for ',mfilename,'. A nonnegative number should be specified.'])
          end
      end
      
      function obj = set.intSeed(obj, myRandomSeed)
          if ((myRandomSeed >= -1) && (mod(myRandomSeed,1) == 0))
              obj.intSeed = myRandomSeed;
          else
              warning(['Setting for intSeed for random number generator must be a nonegative integer in ',mfilename,', or -1 to ignore. Refusing to update.']);
          end
      end      
      
      function value = get(obj,propName)
          switch propName                 
              case 'interventionID'
                  value = obj.interventionID;   
              case 'analyzeElementResultIDs'
                  value = obj.analyzeElementResultIDs;  
              case 'analyzeTime'
                  value = obj.analyzeTime;         
              case 'subSampleSplitType'
                  value = obj.subSampleSplitType;  
              case 'nBootstraps'
                  value = obj.nBootstraps;  
              case 'intSeed'
                  value = obj.intSeed;                    
          end
      end     
      
      function passCheck = verify(obj,myWorksheet)
          passCheck = true;
          allInterventionIDs = getInterventionIDs(myWorksheet);
          allVPIDs = getVPIDs(myWorksheet);
          [dummy, nVPResults]= size(myWorksheet.results);
          if sum(ismember(allInterventionIDs,  obj.interventionID)) < 1
              warning(['Specified interventionID should be selected from interventionIDs the worksheet in ',mfilename,'.'])
              passCheck = false;
          end
          if nVPResults ~= length(allVPIDs)
              warning(['Results for all VPs not available from worksheet in ',mfilename,'.'])
              passCheck = false;
          end
          if (obj.nBootstraps < 1) && ~(strcomp(subSampleSplitType,'none'))
              warning(['If splitting into subsamples in ',mfilename,', nBootstraps must be > 0.'])
              passCheck = false;
          end          
          if passCheck
              interventionIndex = find(ismember(allInterventionIDs, obj.interventionID));
              checkResults = myWorksheet.results(interventionIndex,:);
              if sum(sum(arrayfun(@(i) isstruct(checkResults{i}),1:nVPResults)) ) < length(allVPIDs)
                  %failCheckVPs = ~(arrayfun(@(i) isstruct(checkResults{i}),1:nVPResults));
                  %failCheckVPIndices = find(failCheckVPs);
                  warning(['Results for all VPs not available from worksheet in ',mfilename,'.'])
                  passCheck = false;
              else
                  for vpCounter = 1 : nVPResults
                      if sum(ismember(checkResults{1, vpCounter}.Names,obj.analyzeElementResultIDs)) < length(obj.analyzeElementResultIDs)
                          passCheck = false;
                      end
                  end
                  if ~passCheck
                      warning(['Not all analyzeElementResultIDs identified in the worksheet results provided to ',mfilename,'.'])
                  end
                  ntimefail = 0;
                  for vpCounter = 1 : nVPResults
                      timeIndex = find(ismember(checkResults{1, vpCounter}.Names,'time'));
                      timeVals = checkResults{1, vpCounter}.Data(:,timeIndex);
                      % Might want to add a tolerance here... but
                      % MATLAB has been generally robust for small
                      % numerical issues so far
                      if sum(timeVals == obj.analyzeTime) < 1
                          ntimefail = ntimefail+1;
                      end
                  end
                  if ntimefail > 0
                      passCheck = false;
                      warning(['Not all sample times align with the desired analyzeTime in the results provided to ',mfilename,'.'])
                      passCheck = false;
                  end
              end
          end
      end
      
      % The constructor method must have the same name as the class
      function obj = sobolSensitivityOptions() 
          obj.interventionID='';
          obj.analyzeElementResultIDs={};
          obj.analyzeTime=0;
          obj.subSampleSplitType='none';
          obj.nBootstraps=1E3; 
          obj.intSeed=-1;           
      end
   end
end
```
