## Supplementary File 3 for "A surrogate workflow for virtual population development: designing an amenable proxy for a hypothesis-test inspired nonlinear goodness-of-fit": varyAxesOptions.m

```
classdef varyAxesOptions
% Options object for introducing random VPs into a worksheet with the
% varyAxes function
%
% PROPERTIES
%  baseVPIDs:           a cell array of strings consisting of VP IDs
%                       in the worksheet that will base used as a base -
%                       their parameter sets will be inherited
%  newPerOld:           how many random VPs to create per baseVPID 
%  varyMethod:          method for varying the axes.  Allowed values are:
%                        'uniform' a uniform distribution for each variable
%                        'sobol' a sobol sequence is used
%                        'saltelli' one of the variations on Saltelli's
%                                   method for making the best use of
%                                   samples for calculating the sensitivity
%                                   indices
%                        'gaussian' a normal distribution for each axis
%                        'lh'      latin hypercube
%                        'none'     no randomization is applied to the
%                                   children
%                        'localpca' sample along PCAs for the nearest VPs
%  additionalIDString:  an additional string to add to the VP IDs
%                       generated.  This can be helpful for ensuring
%                       each VP ID is unique, even with multiple runs
%                       and filtering VPs
%  varyAxisIDs:         a cell array of strings, each being an axis
%                       in the worksheet
%  gaussianStd:         Only used if 'gaussian' or 'localPCA' is the varyMethod.
%                       For gaussian, this is a normalized standard deviation, 
%                       since the axes are scaled between 0 and 1.
%                       For localPCA, this is a relative  re-scaling of the 
%                       standard deviation along all principal components when sampling
%  intSeed:             A non-negative integer seed to initialize the 
%                       random number generator.  Set to -1 to avoid
%                       changing the state of the random number generator.
%

   properties
      baseVPIDs;
      newPerOld;
      varyMethod;
      additionalIDString;
      varyAxisIDs;
      gaussianStd;
      intSeed;
   end
   methods
       
      function obj = set.baseVPIDs(obj,myBaseVPIDs)
          if iscell(myBaseVPIDs)
              obj.baseVPIDs = (myBaseVPIDs);
          else
              error(['Invalid baseVPIDs specified for ',mfilename,', a cell array of VPs ID strings should be specified.'])
          end
      end

      function obj = set.newPerOld(obj,myNewPerOld)
          failFlag = false;
          if isnumeric(myNewPerOld) 
              if isequal(size(myNewPerOld),[1 1])
                  if round(myNewPerOld) > 0
                      obj.newPerOld = round(myNewPerOld);
                  else
                      failFlag = true;
                  end
              else
                  failFlag = true;
              end
          else
              failFlag = true;
          end
          if failFlag
              error(['Invalid newPerOld specified for ',mfilename,'. A positive number should be specified.'])
          end
      end

      function obj = set.varyMethod(obj,myVaryMethod)
          allowedSettings = {'uniform','sobol','saltelli','gaussian','lh','none','localpca'};
          if sum(ismember(allowedSettings, lower(myVaryMethod)))>0
              obj.varyMethod = lower(myVaryMethod);
          else
              error('Invalid varyMethod specified for ',mfilename,', should specify one of: ',strjoin(allowedSettings,', '),'.')
          end
      end 
      
      function obj = set.additionalIDString(obj,myAdditionalIDString)
          if ischar(myAdditionalIDString)
              obj.additionalIDString = (myAdditionalIDString);
          else
              error(['Invalid additionalIDString specified in ',mfilename,', a string should be specified.'])
          end
      end      
      
      function obj = set.varyAxisIDs(obj,myVaryAxisIDs)
          if iscell(myVaryAxisIDs)
              obj.varyAxisIDs = (myVaryAxisIDs);
          else
              error(['Invalid varyAxisIDs specified for ',mfilename,', a cell array of axis ID strings should be specified.'])
          end
      end          

      function obj = set.gaussianStd(obj,myGaussianStd)
          failFlag = false;
          if isnumeric(myGaussianStd) 
              if isequal(size(myGaussianStd),[1 1])
                  if myGaussianStd >= 0
                      obj.gaussianStd = (myGaussianStd);
                  else
                      failFlag = true;
                  end
              else
                  failFlag = true;
              end
          else
              failFlag = true;
          end
          if failFlag
              error(['Invalid gaussianStd specified for ',mfilename,'. A nonegative integer should be specified.'])
          end
      end
    
      function obj = set.intSeed(obj,myIntSeed)
          if ((mod(myIntSeed,1) == 0) && (myIntSeed>=-1))
              obj.intSeed = (myIntSeed);
          else
              error(['Invalid intSeed specified for ',mfilename,', a non-negative integer should be specified, or -1 to ignore.'])
          end
      end            
      
      function value = get(obj,propName)
          switch propName
              case 'baseVPIDs'
                  value = obj.baseVPIDs;
              case 'newPerOld'
                  value = obj.newPerOld;
              case 'varyMethod'
                  value = obj.varyMethod; 
              case 'additionalIDString'
                  value = obj.additionalIDString;                   
              case 'varyAxisIDs'
                  value = obj.varyAxisIDs;  
              case 'gaussianStd'
                  value = obj.gaussianStd;  
              case 'intSeed'
                  value = obj.intSeed;     
              otherwise
                  error(['Error: ',propName ,' is not a valid ',mfilename,' property.'])                  
          end
      end
      
      function passCheck = verify(obj,myWorksheet)
          % Verify the specified varyAxesOptions are acceptable for a 
          % given worksheet.
          %
          % ARGUMENTS
          %  (self)
          %  myWorksheet:  a worksheet data structure
          %
          % RETURNS
          %  passCheck:    a boolean (true/false) indicating whether
          %                the specified varyAxesOptions pass the check
          %
          passCheck = true;
          allVPIDs = getVPIDs(myWorksheet);
          if sum(ismember(obj.baseVPIDs, allVPIDs)) < length(obj.baseVPIDs)
              warning(['Specified baseVPIDs should all be selected from VPIDs the worksheet in ',mfilename,'.'])
              passCheck = false;
          end
          allAxisDefIDs = getAxisDefIDs(myWorksheet);
          if sum(ismember(allAxisDefIDs, obj.varyAxisIDs)) < length(obj.varyAxisIDs)
              warning(['Specified varyAxisIDs should all be specified from axis IDs the worksheet in ',mfilename,'.'])
              passCheck = false;
          end          
      end
      
      % The constructor method must have the same name as the class
      function obj = varyAxesOptions()
          obj.baseVPIDs = cell(1,0);
          obj.newPerOld = 100;
          obj.varyMethod = 'uniform';
          obj.additionalIDString='';
          obj.varyAxisIDs = {};  
          obj.gaussianStd = 0.05;
          obj.intSeed = -1;
      end
   end
    
end
```
