## Supplementary File 3 for "A surrogate workflow for virtual population development: designing an amenable proxy for a hypothesis-test inspired nonlinear goodness-of-fit": createResponseSummaryTable.m

```
function myResponseSummaryTable = createResponseSummaryTable(myWorksheet, myResponseTypeID, flagParameters)
% Make a copy of a worksheet; if VPs are specified, they will be selected.
%
% ARGUMENTS
% myWorksheet: a worksheet
% myResponseTypeID: a cell array, 1XnVP, of string identifiers
% flagParameters: whether to break parameter axes down to parameters or just
%                 to use coefficients.
%
% RETURNS
% myResultTable: data from cohort, which should be more suitable
%              for subsequent analysis
%

% Perform initial checks on the provided arguments
flagContinue = true;
myResponseSummaryTable = resultTable({});
if nargin > 3
    warning([mfilename,' requires input arguments: myWorksheet, responseTypeID, and optionally flagParameters.  Too many arguments provided.'])
    flagContinue = false;
elseif nargin > 2
    if flagParameters
        warning([mfilename,' currently only supports flagParameters == false; reassigning.'])
        flagParameters = false;
    end     
elseif nargin > 1
    flagParameters = false;
else
    warning([mfilename,' requires input arguments: myWorksheet, responseTypeID, and optionally flagParameters.  Insufficient arguments provided.'])
    flagContinue = false;
end

if flagContinue
    myResponseTypeIDs = getResponseTypeIDs(myWorksheet);
    if sum(ismember(myResponseTypeIDs, myResponseTypeID)) < 1
        flagContinue = false;
        warning([mfilename,' could not find the indicated response type ID, ',myResponseTypeID,', in the provided worksheet.'])
    end
end

if flagContinue
    [nInterventionResult, nVPResult] = size(myWorksheet.results);
    nInterventions = length(getInterventionIDs(myWorksheet));
    nVPs = length(getVPIDs(myWorksheet));
    if (nInterventionResult == 0) || (nVPResult == 0)
        warning(['Results not provided in worksheet provided to ',mfilename,'.'])
        flagContinue = false;
    elseif (nInterventionResult < nInterventions) || (nVPResult < nVPs)
        warning(['Result dimensions (nInterventions x nVPs) not consistent with worksheet size in call to ',mfilename,'.'])
        flagContinue = false;        
    end
end

if flagContinue
    myResponseType = getResponseType(myWorksheet, myResponseTypeID);
    [nResponseTypeElements, dummyVar] = size(myResponseType.elements);
    myAxisDefIDs = getAxisDefIDs(myWorksheet);
        
    if ~(flagParameters)
        
        % First set up the names for the rows/columns
        myColNames = getVPIDs(myWorksheet);
        myRowNames = getResponseTypeElementIDs(myWorksheet, myResponseTypeID);
        myRowNames = [myRowNames, 'vpValues'];
        myRowNames = [myAxisDefIDs, myRowNames];
        
        % Now get the values
        myCoeffValues = getVPCoeffs(myWorksheet);
        myResponseTypeResult = evaluateResponseType(myWorksheet, myResponseTypeID);
        myData = [myCoeffValues; myResponseTypeResult.values; myResponseTypeResult.vpValues];
        myResponseSummaryTable = resultTable({myResponseTypeID,myData,myColNames,myRowNames});
        
    else
        error([mfilename,' currently only supports flagParameters == false.'])
    end
else
    warning(['Unable to generate a resultTable structure in call to ',mfilename,', returning an empty resultTable structure.'])
end
```
