## Supplementary File 3 for "A surrogate workflow for virtual population development: designing an amenable proxy for a hypothesis-test inspired nonlinear goodness-of-fit": createScreenTable.m

```
function myScreenTable = createScreenTable(myWorksheet, myResponseTypeIDs, flagSetBoundsFromResults)
% Create a screening table.  This can be used as an input argument
% to a screening function.  It shows the response type IDs, response type 
% element IDs, and additional information on the response types and their 
% evaluations.
%
% ARGUMENTS
%  myWorksheet:              A worksheet structure, required.  If the results
%                             are populated, they will be used to set allowed
%                             upper values for screening unless flagSetBoundsFromResults
%                             is false.
%  myResponseTypeID:         (optional) A cell array of responseTypeID strings.
%                             They will be used to pick which to use for the
%                             screenTable.  If none are given, all
%                             responseTypes will be used.
%  flagSetBoundsFromResults: (optional) A boolean, whether to use worksheet results
%                             to set nonzero bounds on the response types.  Default
%                             is true, if worksheet results are populated.
%
% RETURNS
%  screenTable:              A table with critical information for
%                             screeningVPs.  Can be used as an input
%                             argument for screenWorksheetVPs.
%

continueFlag = false;
if nargin > 3
    warning(['Too many input arguments to ',mfilename, '. Arguments should be: myWorksheet; and optionally: myResponseTypeIDs, flagSetBoundsFromResults.'])
    continueFlag = false;
elseif nargin > 2
    continueFlag = true;
elseif nargin > 1
    flagSetBoundsFromResults = true;
    continueFlag = true;
	myResponseTypeIDs = getResponseTypeIDs(myWorksheet);
elseif nargin > 0
    continueFlag = true;
    flagSetBoundsFromResults = true;	
	myResponseTypeIDs = getResponseTypeIDs(myWorksheet);
else
	continueFlag = false;
    warning(['Insufficient input arguments to ',mfilename, '. Arguments should be: myWorksheet; and optionally: myResponseTypeIDs, flagSetBoundsFromResults.'])
end

if continueFlag
	allResponseTypeIDs = getResponseTypeIDs(myWorksheet);
	if sum(ismember(myResponseTypeIDs,allResponseTypeIDs)) < length(myResponseTypeIDs)
		continueFlag = false;
		warning(['Provided myResponseTypeIDs in ',mfilename, ' not all found in myWorksheet.'])
	end
end		

if continueFlag
	% Check to see if we will populate the bounds from the
	% worksheet results.
    vpIDs = getVPIDs(myWorksheet);
    interventionIDs = getInterventionIDs(myWorksheet);
    nVPs = length(vpIDs);
    nInterventions = length(interventionIDs);
    [nInterventionResults, nVPResults] = size(myWorksheet.results);
    myResultClasses = cellfun(@class,myWorksheet.results, 'UniformOutput', false);
    % Results should be stored in a structure, we assume 
    % if a structure is provided then it is a valid result
    flagVPComplete = sum(strcmp(myResultClasses,'struct'),1) == nInterventions;
    
    if ((sum(flagVPComplete) == nVPs) && flagSetBoundsFromResults)
		getValues = true;
	elseif flagSetBoundsFromResults
		disp(['Unable to get complete results for worksheet provided to ',mfilename,'.  Setting responseType limits to 0.'])
		getValues = false;	
	else
		getValues = false;
	end	
end		


if continueFlag

	tableResponseTypeIDs = cell(0,1);
	tableResponseTypeElementIDs = cell(0,1);	
	tableResponseTypeElementTypes = cell(0,1);
	tableInterventionIDs = cell(0,1);		
	tableModelYVars = cell(0,1);
	tableModelYVarsTypes = cell(0,1);
	tableWeights = zeros(0,1);
	tableValues = zeros(0,1);	
	myAxisIDs = getAxisDefIDs(myWorksheet);
	nAxis = length(myAxisIDs);
	for responseTypeCounter = 1: length(myResponseTypeIDs)
        curTableResponseTypeElementIDs = (getResponseTypeElementIDs(myWorksheet,myResponseTypeIDs{responseTypeCounter}))';	
		nResponseTypeElementIDs = length(curTableResponseTypeElementIDs);
		curTableResponseTypeIDs = cell(nResponseTypeElementIDs+1,1);		
		curTableResponseTypeElementTypes = cell(nResponseTypeElementIDs+1,1);
		curTableInterventionIDs = cell(nResponseTypeElementIDs+1,1);		
		curTableModelYVars = cell(nResponseTypeElementIDs+1,1);
		curTableModelYVarsTypes = cell(nResponseTypeElementIDs+1,1);		
		curTableValues = zeros(nResponseTypeElementIDs+1,1);
		curTableWeights = zeros(nResponseTypeElementIDs+1,1);
		for responseTypeElementCounter = 1: nResponseTypeElementIDs
			curTableResponseTypeIDs{responseTypeElementCounter} = myResponseTypeIDs{responseTypeCounter};
			curTableModelYVars{responseTypeElementCounter} = myWorksheet.responseTypes{responseTypeCounter}.elements{responseTypeElementCounter}.modelYVar;
			curTableModelYVarsTypes{responseTypeElementCounter} = myWorksheet.responseTypes{responseTypeCounter}.elements{responseTypeElementCounter}.modelYVarType;
			curTableInterventionIDs{responseTypeElementCounter} = myWorksheet.responseTypes{responseTypeCounter}.elements{responseTypeElementCounter}.interventionID;
			curTableValues(responseTypeElementCounter) = 0;
			curTableResponseTypeElementTypes{responseTypeElementCounter} = class(myWorksheet.responseTypes{responseTypeCounter}.elements{responseTypeElementCounter});
			curTableWeights(responseTypeElementCounter) = myWorksheet.responseTypes{responseTypeCounter}.elements{responseTypeElementCounter}.weight;
		end
		% Append an element for the sum.
		curTableResponseTypeElementIDs{nResponseTypeElementIDs+1,1} = 'vpValues';
		curTableResponseTypeElementTypes{nResponseTypeElementIDs+1} = '';
		curTableResponseTypeIDs{nResponseTypeElementIDs+1} = myResponseTypeIDs{responseTypeCounter};
		curTableInterventionIDs{nResponseTypeElementIDs+1} = '';		
		curTableModelYVars{nResponseTypeElementIDs+1} = '';
		curTableModelYVarsTypes{nResponseTypeElementIDs+1} = '';		
		curTableValues(nResponseTypeElementIDs+1) = 0;
		curTableWeights(nResponseTypeElementIDs+1) = 1;		
		
		% Get screening limits from the current worksheet, if needed.
		if getValues
			myResponseSummaryTable = createResponseSummaryTable(myWorksheet, allResponseTypeIDs{responseTypeCounter});
			curTableValues = max(myResponseSummaryTable.values((nAxis+1):end,:),[],2);
		end
		
		% Append the current set from the current response type.
		tableResponseTypeIDs = [tableResponseTypeIDs;curTableResponseTypeIDs];
		tableResponseTypeElementIDs = [tableResponseTypeElementIDs;curTableResponseTypeElementIDs];
		tableResponseTypeElementTypes = [tableResponseTypeElementTypes;curTableResponseTypeElementTypes];
		tableInterventionIDs = [tableInterventionIDs;curTableInterventionIDs];
		tableModelYVars = [tableModelYVars;curTableModelYVars];
		tableModelYVarsTypes = [tableModelYVarsTypes;curTableModelYVarsTypes];
		tableValues = [tableValues;curTableValues];
		tableWeights = [tableWeights;curTableWeights];
	end
	myScreenTable = table(tableResponseTypeIDs,tableResponseTypeElementIDs,tableResponseTypeElementTypes,tableInterventionIDs,tableModelYVars,tableModelYVarsTypes,tableWeights,tableValues);
	myScreenTable.Properties.VariableNames = {'responseTypeID','responseTypeElementID','responseTypeElementType','interventionID','modelYVar','modelYVarType','weight','value'};
else
	myScreenTable = [];
    warning(['Unable to complete ',mfilename,'.  Exiting.'])
end

end
```
