## Supplementary File 3 for "A surrogate workflow for virtual population development: designing an amenable proxy for a hypothesis-test inspired nonlinear goodness-of-fit": pickClusterVPs.m

```
function medoidData = pickClusterVPs(myWorksheet, myClusterPickOptions)
% Run a clustering algorithm and propose VPs to keep.
% This is done in two parts:
% first "edge" VPs are picked, 
% then remaining VPs are clustered.
%
% ARGUMENTS
%  myWorksheet:           starting worksheet
%  myClusterPickOptions:  optional, a cluster options object
%                         if not provided, the function will attempt
%                         to construct one based on the provided worksheet.
%
% RETURNS
% medoidData:             a structure with the clustering results,
%                          including the IDs of VPs to keep.
%
%
medoidData = struct();
continueFlag = false;
% Verify the input arguments.
if nargin > 2
    warning(['Too many input arguments to ',mfilename, '. Arguments should be: myWorksheet; optionally: myClusterPickOptions.'])
    continueFlag = false; 
elseif nargin > 1
    allVPIDs = getVPIDs(myWorksheet);
    continueFlag = true;      
elseif nargin > 0
    allVPIDs = getVPIDs(myWorksheet);
    myClusterPickOptions = clusterPickOptions();
    myClusterPickOptions = myClusterPickOptions.setDefaultFromWorksheet(myWorksheet);
    continueFlag = true; 
    mySimulateOptions = simulateOptions;     
else
    warning(['Insufficient input arguments to ',mfilename, '. Arguments should be: myWorksheet; optionally: myClusterPickOptions.'])
    continueFlag = false; 
end

% Additional proofing that the input arguments make sense.
if continueFlag
    continueFlag = myClusterPickOptions.verify(myWorksheet);
end

if continueFlag
    
    if myClusterPickOptions.poolRestart
        if ~isempty(gcp('nocreate'))
            delete(gcp);
        end
    end    
    if isempty(gcp('nocreate'))
        % First check the default number of workers, if needed
        myClusterPickOptions = checkNWorkers(myClusterPickOptions);
        myPool = parpool(myClusterPickOptions.clusterID, myClusterPickOptions.nWorkers,'SpmdEnabled',false);        
    end    
   
    if myClusterPickOptions.intSeed > -1
        rng(myClusterPickOptions.intSeed, 'twister');
    end

    % We need to create a matrix of data values that will then be
    % normalized and clustered.  It will be (nClusterAxis + nClusterOutput)
    % x nVP in size.
    nClusterAxis = length(myClusterPickOptions.clusterAxisIDs);
    [nClusterOutput, ~] = size(myClusterPickOptions.clusterElement); 
    nVP = length(allVPIDs);
    theMatrixToCluster = nan((nClusterAxis + nClusterOutput), nVP);
    theRowNames = cell(nClusterAxis + nClusterOutput,1);
    theRowNames(1:nClusterAxis) = myClusterPickOptions.clusterAxisIDs;
    for theElementCounter = 1:nClusterOutput
        theName = [myClusterPickOptions.clusterElement{theElementCounter, 1},'___',myClusterPickOptions.clusterElement{theElementCounter, 2},'___',myClusterPickOptions.clusterElement{theElementCounter, 3},'___',num2str(myClusterPickOptions.clusterElement{theElementCounter, 4})];
        theRowNames{nClusterAxis+theElementCounter} = theName;
    end
    % Now we need to add the data
    worksheetAxisIndices = nan(nClusterAxis,1);
    allAxisIDs = getAxisDefIDs(myWorksheet);
    allInterventionIDs = getInterventionIDs(myWorksheet);
    for axisCounter = 1 : nClusterAxis
        worksheetAxisIndices(axisCounter) = find(ismember(allAxisIDs,myClusterPickOptions.clusterAxisIDs{axisCounter}));
    end
    allAxisCoefs = getVPCoeffs(myWorksheet);
    theMatrixToCluster(1:nClusterAxis,:) = allAxisCoefs(worksheetAxisIndices,:);
    % The outputs need to be retrieved for each VP separately...
    % TODO: might want to add some check here for whether results exist and
    % if not to simulateWorksheet
    for theElementCounter = 1 : nClusterOutput
        theElementName = myClusterPickOptions.clusterElement{theElementCounter, 1};
        theElementType = myClusterPickOptions.clusterElement{theElementCounter, 2};
        theInterventionID = myClusterPickOptions.clusterElement{theElementCounter, 3};
        theSampleTime = myClusterPickOptions.clusterElement{theElementCounter, 4};
        theInterventionIndex = find(ismember(allInterventionIDs, theInterventionID));
        % All VPs should be run with the same model, with the same result
        % elements
        theVPCounter = 1;
        curResultStruct = myWorksheet.results{theInterventionIndex,theVPCounter};
        timeIndex = find(ismember(curResultStruct.Names, 'time'));
        % We assume element IDs are unique, even without type information.
        elementIndex = find(ismember(curResultStruct.Names, theElementName));
        for theVPCounter = 1 : nVP
            curResultStruct = myWorksheet.results{theInterventionIndex,theVPCounter};
            curTime = curResultStruct.Data(:,timeIndex);
            curElement = curResultStruct.Data(:,elementIndex);
            y_new = interp1(curTime,curElement,theSampleTime,'linear');
            theMatrixToCluster(nClusterAxis+theElementCounter,theVPCounter)=y_new;
        end
    end

    originalMatrix = theMatrixToCluster;
        
    if strcmp(myClusterPickOptions.normalizeType,'min-max') 
        denominatorTerm = max(theMatrixToCluster,[],2)-min(theMatrixToCluster,[],2) + ((max(theMatrixToCluster,[],2)-min(theMatrixToCluster,[],2))==0);
        theMatrixToCluster = (theMatrixToCluster - min(theMatrixToCluster,[],2)*ones(1, size(theMatrixToCluster,2)))./(denominatorTerm*ones(1, size(theMatrixToCluster,2)));
        theMatrixToCluster = theMatrixToCluster ;
    elseif strcmp(myClusterPickOptions.normalizeType,'z-score')
        denominatorTerm = std(theMatrixToCluster,[],2) + ((max(theMatrixToCluster,[],2)-min(theMatrixToCluster,[],2))==0);
        theMatrixToCluster = theMatrixToCluster./(denominatorTerm*ones(1, size(theMatrixToCluster,2)));
        theMatrixToCluster = theMatrixToCluster - mean(theMatrixToCluster,2)*ones(1, size(theMatrixToCluster,2));
    end
    
    myNClusters = myClusterPickOptions.nClusters;
    myIndices = nan(1,myNClusters);
    % We may remove entries so we need to map back to the original
    % matrix
    myIndicesMap = [1 : nVP]';
    % If we have stated that we want to include edge VPs, first we pull
    % 2 VPs from each axis or output, one for the min and one for the max,
    % then we will cluster the remaining VPs    
    % We could improve this by scanning for a minimal set that covers all
    % the edges in case there are a few VPs that cover the same edges.
    % For now, just pick the edge VPs randomly; this will also influence
    % the other dimensions.
    % I should write a better algorithm for getting
    % the edge VPs and call that here, and also in clusterPickOptions
    nVPFromEdges = 0;
    edgeSets = cell(2*(nClusterAxis + nClusterOutput),1);
    if myClusterPickOptions.edgeVPFlag
        for curClusterAxisOutputCounter = 1 : (nClusterAxis + nClusterOutput)
            curMaxVal = max(theMatrixToCluster(curClusterAxisOutputCounter,:));
            curMaxIndex = find(theMatrixToCluster(curClusterAxisOutputCounter,:) >= curMaxVal);
            curMinVal = min(theMatrixToCluster(curClusterAxisOutputCounter,:));
            curMinIndex = find(theMatrixToCluster(curClusterAxisOutputCounter,:) <= curMinVal);
            edgeSets{(curClusterAxisOutputCounter-1)*2+1,1} = allVPIDs(curMinIndex);
            edgeSets{(curClusterAxisOutputCounter-1)*2+2,1} = allVPIDs(curMaxIndex);
        end
        edgeSets = getMinimalEdgeSet(edgeSets);
        edgeIndices = find(ismember(allVPIDs,edgeSets));
        myIndices(1:length(edgeIndices)) = edgeIndices;
        if myClusterPickOptions.verbose
            disp(['Found ',num2str(length(edgeIndices)),' edge VPs, total requested number of VPs post clustering is ',num2str(myNClusters),'.  Proceeding.'])
        end
		if length(edgeIndices) >= myNClusters
			warning(['Edge VPs are greater than number of requested clustered VPs in ',mfilename,'.  Returning edge VPs but not clustering.'])
		end
        nVPFromEdges = length(edgeIndices);
        myIndicesMap(edgeIndices) = [];
        theMatrixToCluster(:,edgeIndices) = [];
    else
        edgeIndices = [];
    end
    
    % VPs should be rows for clustering
    theMatrixToCluster = transpose(theMatrixToCluster);    
    
    medoidIndices = nan(myNClusters-length(edgeIndices),1);
    if sum(isnan(myIndices)) > 0
       % opts = statset('MaxIter',myClusterPickOptions.maxIter,'UseParallel',1);
           if myClusterPickOptions.intSeed > -1
                s = RandStream('mlfg6331_64','Seed',myClusterPickOptions.intSeed);  % Random number stream
                opts = statset('MaxIter',myClusterPickOptions.maxIter,'UseParallel',1,'UseSubstreams',1,'Streams',s);
           else
                s = RandStream('mlfg6331_64','Seed',0);  % used in example04
                opts = statset('MaxIter',myClusterPickOptions.maxIter,'UseParallel',1,'UseSubstreams',1,'Streams',s);
           end
        if sum(ismember(myClusterPickOptions.algorithm,'auto')) > 0
            if nVP <= 3000
                 myAlgorithm = 'pam';
            elseif nVP <= 10000
                 myAlgorithm = 'small';
            else
                 myAlgorithm = 'large';
            end 
        else
            myAlgorithm = myClusterPickOptions.algorithm;
        end                
        

        [idx,theMedoids,sumd,D,midx,info] = kmedoids(theMatrixToCluster, (myNClusters - nVPFromEdges), 'Algorithm', myAlgorithm, 'Distance', myClusterPickOptions.distance, 'Start','plus', 'Replicates', myClusterPickOptions.replicates, 'Options',opts);     
        
        for medoidCounter = 1 : (myNClusters - nVPFromEdges)
            curIndices = find(ismember(theMatrixToCluster,theMedoids(medoidCounter,:),'rows'));
            % We may need to filter duplicate solutions
            % For example, in case the GA carried duplicate VPs forward.
            % So we make sure to pick out one VP per medoid.        
            medoidIndices(medoidCounter) = myIndicesMap(curIndices(1));
            myIndices(nVPFromEdges + medoidCounter) = myIndicesMap(curIndices(1));
        end
    else
        idx = myIndices;
        theMedoids = nan(0,nClusterAxis + nClusterOutput);
		sumd = nan;
		D = nan;
		midx = nan;
		info = '';
    end
    %myIndices = find(ismember(theMatrixToCluster,theMedoids,'rows'));
    medoidData.('medioidVPIDs') = allVPIDs(medoidIndices);
    medoidData.('edgeVPIDs') = allVPIDs(edgeIndices);
    medoidData.('pickedVPIDs') = allVPIDs(myIndices);
    medoidData.('allVPIDs') = allVPIDs;
    medoidData.('clusteredMatrix') = transpose(theMatrixToCluster);
    medoidData.('untransformedMatrix') = originalMatrix;
    medoidData.('clusterIndices') = transpose(idx);
    medoidData.('clusterVPIDs') = setdiff(allVPIDs,allVPIDs(edgeIndices));
    medoidData.('theMedoids') = transpose(theMedoids);
    medoidData.('sumd') = transpose(sumd);
	medoidData.('D') = transpose(D);
	medoidData.('midx') = transpose(midx);
	medoidData.('info') = info;
    
    % Clean up the pool, if needed
    if myClusterPickOptions.poolClose
        if ~isempty(gcp('nocreate'))
            delete(gcp);
        end
    end    
    
else
    warning(['Unable to run ',mfilename,'.'])
end
```
