## Supplementary File 3 for "A surrogate workflow for virtual population development: designing an amenable proxy for a hypothesis-test inspired nonlinear goodness-of-fit": runControlCoefficientsSimulations.m

```
function myFinalWorksheet = runControlCoefficientsSimulation(myWorksheet,myRunControlCoefficientSimulationsOptions)
% This function runs a llocal control coefficient analysis
% This is based on the local SA method in:
% Chen X, Hickling TP, Vicini P. A mechanistic, multiscale mathematical 
% model of immunogenicity for therapeutic proteins: part 1-theoretical
% model. CPT: pharmacometrics & systems pharmacology. 2014;3:e133.
%
% ARGUMENTS
% myWorksheet:              An instance of a worksheet
% myRunControlCoefficientSimulationsOptions: An instance of a runControlCoefficientSimulationsOptions
%                           object. This must be provided.
%
% RETURNS
% myWorksheet:              Final worksheet with sweep VPs, and with 
%                           simulation results if specified
%
continueFlag = false;
if nargin > 2
    warning(['Too many input arguments to ',mfilename, '. Arguments should be: myWorksheet and myRunControlCoefficientSimulationsOptions.'])
    continueFlag = false;
elseif nargin > 1
    continueFlag = true;
else
    warning(['Insufficient input arguments to ',mfilename, '. Arguments should be: myWorksheet and myRunControlCoefficientSimulationsOptions.'])
    continueFlag = false;
end

if continueFlag
    continueFlag = myRunControlCoefficientSimulationsOptions.verify(myWorksheet);
    if ~(continueFlag)
       warning(['Please correct the options provided to ',mfilename, '.']) 
    end
end

tic;
saveTime = 0;

if continueFlag
    
    % Prepare the worksheet
    removeInterventionIDs = getInterventionIDs(myWorksheet);
    removeInterventionIDs = setdiff(removeInterventionIDs, {myRunControlCoefficientSimulationsOptions.interventionID});
    myWorksheet=copyWorksheet(myWorksheet,{myRunControlCoefficientSimulationsOptions.baseVPID});
    myWorksheet = removeInterventions(myWorksheet, removeInterventionIDs);
    if myRunControlCoefficientSimulationsOptions.verbose
        disp(['Creating VPs in ',mfilename,'.']);
    end
    % Since this is a univariate sweep of axis values and we currently
    % only support the high/low point (we may make this variable in the
    % future), we just create the VPs here 
    allAxisIDs = getAxisDefIDs(myWorksheet);    
    nAxis = length(allAxisIDs);
    defaultCoefficients = getVPCoeffs(myWorksheet);
    nVaryAxis = length(myRunControlCoefficientSimulationsOptions.varyAxisIDs);
    testVPIDs = getVPIDs(myWorksheet);
    newVPIDs = cell(1,nVaryAxis);
    newVPVariants = cell(1,nVaryAxis);
    newVPAxis = cell(1,nVaryAxis);    
    baseVP = getVP(myWorksheet,myRunControlCoefficientSimulationsOptions.baseVPID);
    baseVPIndex = find(ismember(testVPIDs,myRunControlCoefficientSimulationsOptions.baseVPID));
    newVPCoeffs = getVPCoeffs(myWorksheet);
    newVPCoeffs = newVPCoeffs(:,baseVPIndex);
    newVPCoeffs = newVPCoeffs * ones(1,nVaryAxis);
    for axisCounter = 1 : nVaryAxis
        newVPvariants{1,axisCounter} = baseVP.variants;        
        newVPIDString = [myRunControlCoefficientSimulationsOptions.baseVPID,'_axis_',num2str(axisCounter),'_alt'];
        newVPIDs{1,axisCounter} = newVPIDString;
    end
    myWorksheet = createVPs(myWorksheet,newVPIDs,newVPvariants,newVPCoeffs);
    for axisCounter = 1 : nVaryAxis
        newVPIndex = axisCounter + 1;
        curAxisID = myRunControlCoefficientSimulationsOptions.varyAxisIDs{axisCounter};
        axisIndex = find(ismember(allAxisIDs,curAxisID));
        elementValue = myWorksheet.axisProps.axisVP.calculateElementValues([axisIndex,baseVPIndex],myWorksheet);
        % Chen increases by 1%
        elementValue = elementValue*1.01;
        myWorksheet.axisProps.axisVP = myWorksheet.axisProps.axisVP.calculateCoefficientFromValues([axisIndex,newVPIndex],elementValue,myWorksheet);
    end
    
    if myRunControlCoefficientSimulationsOptions.verbose
        disp(['Done.'])
    end      
    allVPIDs = getVPIDs(myWorksheet);
    if myRunControlCoefficientSimulationsOptions.simulateWorksheet
        % Prepare for simulations
        mySimulateOptions = simulateOptions();
        % We need to know if runs fail   
        mySimulateOptions.filterFailedRunVPs=false;
        myWorksheet.simProps.saveElementResultIDs = myRunControlCoefficientSimulationsOptions.saveElementResultIDs;        
        if myRunControlCoefficientSimulationsOptions.verbose
            disp(['Beginning simulations in ',mfilename,'.'])
        end
        myFinalWorksheet = simulateWorksheet(myWorksheet, mySimulateOptions);
        if myRunControlCoefficientSimulationsOptions.verbose
            disp(['Completed simulations in ',mfilename,'.'])
            disp(['Elapsed time is ',num2str(toc/60),' min.'])
        end
    else
        myFinalWorksheet = myWorksheet;
    end
    if length(myRunControlCoefficientSimulationsOptions.saveFileName) > 0
        saveWorksheet(myFinalWorksheet, myRunControlCoefficientSimulationsOptions.saveFileName);
    end
else
    warning(['Unable to run ',mfilename, '. Returning input worksheet.'])
    myFinalWorksheet = myWorksheet;
end
end
```
