## Supplementary File 3 for "A surrogate workflow for virtual population development: designing an amenable proxy for a hypothesis-test inspired nonlinear goodness-of-fit": runSobolSample.m

```
function myFinalWorksheet = runSobolSample(myWorksheet,mySobolSampleOptions)
% This function runs a scan of the parameter space using a Sobol sequence
% based sampling scheme that has been optimized for subsequent sensitivity
% analysis. 
% For a desciption, see:
% Saltelli, A., et al, Global Sensitivity Analysis:
% The Primer. 2009. Pages 164-167.
% This function runs the associated simulations, so run time may be
% lengthy.
%
% ARGUMENTS
% myWorksheet:          An instance of a worksheet
% mySobolSampleOptions: An instance of a sobolSampleOptions object.  This
%                       must be provided.
%
% RETURNS
% myWorksheet: Final worksheet with sampling results
%
continueFlag = false;
if nargin > 2
    warning(['Too many input arguments to ',mfilename, '. Arguments should be: myWorksheet and mySobolSampleOptions.'])
    continueFlag = false;
elseif nargin > 1
    continueFlag = true;
else
    warning(['Insufficient input arguments to ',mfilename, '. Arguments should be: myWorksheet and mySobolSampleOptions.'])
    continueFlag = false;
end

if continueFlag
    continueFlag = mySobolSampleOptions.verify(myWorksheet);
    if ~(continueFlag)
       warning(['Please correct the options provided to ',mfilename, '.']) 
    end
end

tic;
saveTime = 0;

if continueFlag
    % Reset the rng if specified
    if mySobolSampleOptions.intSeed > -1
        rng(mySobolSampleOptions.intSeed, 'twister');
    end
    % Update varyOptions
    myVaryAxesOptions = varyAxesOptions;
    myVaryAxesOptions.baseVPIDs = {mySobolSampleOptions.baseVPID};
    myVaryAxesOptions.newPerOld = mySobolSampleOptions.nRandomizationsPerSample;
    myVaryAxesOptions.varyMethod = 'saltelli';
    myVaryAxesOptions.varyAxisIDs = mySobolSampleOptions.varyAxisIDs;  
    myVaryAxesOptions.intSeed = mySobolSampleOptions.intSeed;
    % Prepare the worksheet
    removeInterventionIDs = getInterventionIDs(myWorksheet);
    removeInterventionIDs = setdiff(removeInterventionIDs, {mySobolSampleOptions.interventionID});
    myWorksheet = removeInterventions(myWorksheet, removeInterventionIDs);
    allVPIDs = getVPIDs(myWorksheet);
    if mySobolSampleOptions.verbose
        disp(['Creating VPs in ',mfilename,'.']);
    end    
    myWorksheet = addVariedVPs(myWorksheet, myVaryAxesOptions);
    if mySobolSampleOptions.verbose
        disp(['Done.']);
    end      
    % Remove the original VPs in the worksheet
    myWorksheet = removeVPs(myWorksheet, allVPIDs);
    allVPIDs = getVPIDs(myWorksheet);
    % To avoid overloading the memory in case there are a lot of VPs,
    % (memory overhead may expand in PARFOR)
    % we have an option to run the SSA in batch mode.
    batchSize = mySobolSampleOptions.maxBatchSimulateN;
    nBatches = ceil(length(allVPIDs)/batchSize);
    mySimulateOptions = simulateOptions();
    % We need to know if runs fail, due to the way the Saltelli 
    % method structures the A,B,Ci's    
    mySimulateOptions.filterFailedRunVPs=false;
    myWorksheet.simProps.saveElementResultIDs = mySobolSampleOptions.saveElementResultIDs;
    if mySobolSampleOptions.simulateWorksheet
        if nBatches > 1
            for batchCounter = 1 : nBatches
                startIndex = (batchCounter-1) * batchSize + 1;
                if batchCounter < nBatches
                    endIndex = (batchCounter) * batchSize;
                else
                    endIndex = length(allVPIDs);
                end
                simulateVPIDs = allVPIDs(startIndex:endIndex);            
                mySimulateWorksheet = copyWorksheet(myWorksheet, simulateVPIDs, false);
                myWorksheet = removeVPs(myWorksheet,simulateVPIDs);

                mySimulateWorksheet = simulateWorksheet(mySimulateWorksheet, mySimulateOptions);
                %mySimulateWorksheet = filterResults(mySimulateWorksheet, mySobolSampleOptions.saveElementResultIDs);
                if batchCounter == 1
                    myFinalWorksheet = mySimulateWorksheet;
                else
                    myFinalWorksheet = mergeWorksheets(myFinalWorksheet, mySimulateWorksheet);
                end
                % Limit the file saves to every 3 hr, since saves
                % may take time to write all the data if the files
                % get big.
                if length(mySobolSampleOptions.saveFileName) > 0
                    if (toc/60-saveTime) > 3*60
                        saveTime = toc/60;
                        saveWorksheet(myFinalWorksheet, mySobolSampleOptions.saveFileName);
                    end  
                end

                if mySobolSampleOptions.verbose
                    disp(['Completed simulating batch ',num2str(batchCounter),' of ',num2str(nBatches),' in ',mfilename,'.']);
                    disp(['Elapsed time in batch iterations is ',num2str(toc/60),' min.']);
                end            

            end

        else
            if mySobolSampleOptions.verbose
                disp(['Beginning simulations in ',mfilename,'.']);
            end
            myFinalWorksheet = simulateWorksheet(myWorksheet, mySimulateOptions);
            if mySobolSampleOptions.verbose
                disp(['Completed simulations in ',mfilename,'.']);
                disp(['Elapsed time in batch iterations is ',num2str(toc/60),' min.']);
            end
        end
    else
        myFinalWorksheet = myWorksheet;
    end
    if length(mySobolSampleOptions.saveFileName) > 0
        saveWorksheet(myFinalWorksheet, mySobolSampleOptions.saveFileName);
    end
else
    warning(['Unable to run ',mfilename, '. Returning input worksheet.'])
    myFinalWorksheet = myWorksheet;
end
end
```
