## Supplementary File 3 for "A surrogate workflow for virtual population development: designing an amenable proxy for a hypothesis-test inspired nonlinear goodness-of-fit": runSobolSensitivity.m

```
function mySobolResults = runSobolSensitivity(myWorksheet,mySobolSensitivityOptions)
% This function takes a worksheet from Sobol sampling (Saltelli method) and
% calculates the sensitivity indices
%
% ARGUMENTS
% myWorksheet:               an instance of a worksheet, output from
%                            runSobolSample()
% mySobolSensitivityOptions: an instance of a sobolSensitivityOptions
%                            object
%
% RETURNS
% mySobolResults:            A data structure with the desired results.
%                            The fields are:
%                             (varname):      the (varname) field contains the
%                                             results from the bootstrapped
%                                             sensitivity analysis.  Lower
%                                             CI is 2.5 percentile, upper
%                                             CI is 97.5 percentile.
%                                              firstOrderMedian
%                                              firstOrderUpperCI
%                                              firstOrderLowerCI
%                                              totalMedian
%                                              totalUpperCI
%                                              totalLowerCI
%                                              varianceEst1Median
%                                              varianceEst1UpperCI
%                                              varianceEst1LowerCI
%                                              varianceEst2Median
%                                              varianceEst2UpperCI
%                                              varianceEst2LowerCI
%                                             Each is reported as a (nAxis + 1)
%                                             x nSplitLevels matrix.
%                             time:           simulation time the sensitivity
%                                             indices were computer for
%                             axisValues:     an nAxis x (N * (k+2)) matrix 
%                                             that keeps the sampled parameter
%                                             coefficient values.
%                             axisDefIDs:     a 1 x nAxis cell array to record
%                                             the name of axes that were
%                                             varied.
%                             interventionID: a record of the intervention
%                             colNames:       column names for the matrices
%                                             in the (varname) result
%                                             matrices
%                             sampleSize:     a record of the sample size
%                                             at each of the nSplitLevels
%                                             for each column in the 
%                                             (varname) result matrices.
%                                         
%
mySobolResults = struct();
continueFlag = false;
if nargin > 2
    warning(['Too many input arguments to ',mfilename, '. Arguments should be: myWorksheet and mySobolSensitivityOptions.'])
    continueFlag = false;
elseif nargin > 1
    continueFlag = true;
else
    warning(['Insufficient input arguments to ',mfilename, '. Arguments should be: myWorksheet and mySobolSensitivityOptions.'])
    continueFlag = false;
end

if continueFlag
    continueFlag = mySobolSensitivityOptions.verify(myWorksheet);
    if ~(continueFlag)
       warning(['Please correct the options provided to ',mfilename, '.']) 
    end
    myResultIDs = mySobolSensitivityOptions.analyzeElementResultIDs;
    nResultIDs = length(myResultIDs);
    if nResultIDs < 1
        warning(['At least one valid model output must be specified in analyzeElementResultIDs in options provided to ',mfilename,'.'])
        continueFlag = false;
    end    
end

if continueFlag
    % Reset the rng if specified
    if mySobolSensitivityOptions.intSeed > -1
        rng(mySobolSensitivityOptions.intSeed, 'twister');
    end    
    % Sobol/Saltelli
    % number of samples: N(k+2)
    % we can get k and then back-calculate
    % N, rather than requiring these be specified.
    % In theory we could add an extra check at that point to make sure
    % the A, B, and Ci matrices from the Saltelli method agree
    % with what is expected.
    allCoefficients = getVPCoeffs(myWorksheet);
    maxVals = max(allCoefficients,[],2);
    minVals = min(allCoefficients,[],2);
    % The sampling should be performed with the same base VP, so
    % we will ignore axes that don't appear to change.
    theTolerance = 1E-9;
    alteredAxisIndices = find((maxVals - minVals) > theTolerance);
    alteredCoefficients = allCoefficients(alteredAxisIndices, :);
    allAxisIDs = getAxisDefIDs(myWorksheet);
    alteredAxisIDs = allAxisIDs(alteredAxisIndices);
    nAlteredAxisIDs = length(alteredAxisIDs);
    if nAlteredAxisIDs < 1
        warning(['All axes in the worksheet provided to ',mfilename,' do not vary.'])
        continueFlag = false;
    end
end

if continueFlag
    nBootstraps = mySobolSensitivityOptions.nBootstraps;
    [nAllAxis, nVPs] = size(allCoefficients);
    nRandomizationsPerSample = nVPs/(nAlteredAxisIDs+2);
    % We could in theory test this against VP IDs and A,B,C
    % to verify.
    allInterventionIDs = getInterventionIDs(myWorksheet);
    myInterventionIndex = find(ismember(allInterventionIDs, mySobolSensitivityOptions.interventionID));
    % Rather than reconstructing A, B, C parameter matrices
    % We create the y(A), y(B), yi(Ci)
    YA = nan(nRandomizationsPerSample, 1, nResultIDs);
    YB = nan(nRandomizationsPerSample, 1, nResultIDs);
    YC = nan(nRandomizationsPerSample, nAlteredAxisIDs, nResultIDs);
    for vpCounter = 1 : nVPs
        curResults = myWorksheet.results{myInterventionIndex,vpCounter};
        timeIndex = find(ismember(curResults.Names,'time'));
        timeVals = curResults.Data(:,timeIndex);
        timeIndex = find(timeVals == mySobolSensitivityOptions.analyzeTime);
        for resultCounter = 1 : nResultIDs
            % We take for granted non-dengeracy of referenced result
            % element ID's.
            resultIndex = find(ismember(curResults.Names,myResultIDs{resultCounter}));
            if vpCounter <= nRandomizationsPerSample
                YA(vpCounter, 1, resultCounter) = curResults.Data(timeIndex,resultIndex);
            elseif vpCounter <= 2*nRandomizationsPerSample
                YB(vpCounter-nRandomizationsPerSample, 1, resultCounter) = curResults.Data(timeIndex,resultIndex);
            else
                % The sampling should be set up so we increment through
                % each Ci VP, then each Ci.
                YCiCounter = ceil((vpCounter - 2*nRandomizationsPerSample) / nRandomizationsPerSample);
                YCiVPcounter = vpCounter - 2*nRandomizationsPerSample - (YCiCounter-1)*nRandomizationsPerSample;
                YC(YCiVPcounter,YCiCounter,resultCounter) = curResults.Data(timeIndex,resultIndex);
            end
        end
    end
    % Now we split into subsample fractions, e.g. in case we want to see
    % the stability of the solutions with increasing sample size
    % A better way to do this would be to actually generate the Sobol
    % sequences and re-run the model, but this option is
    % provided as a quicker method
    if strcmp(mySobolSensitivityOptions.subSampleSplitType,'base2')
        nSplitLevels = max(floor(log2(nRandomizationsPerSample)),1);
    end
    
    if sum(ismember({'base2','none'},mySobolSensitivityOptions.subSampleSplitType)) > 0
        if (strcmp(mySobolSensitivityOptions.subSampleSplitType,'none') || (nSplitLevels<2))
            nSplitLevels = 1;
        elseif (mod(log2(nRandomizationsPerSample),1)>0) && strcmp('base2',mySobolSensitivityOptions.subSampleSplitType)
            nSplitLevels = nSplitLevels + 1;
        end
        sampleSize = nan(nSplitLevels,1);
        resampledIndices = cell(max(1,nBootstraps),nSplitLevels);
        theIndices = (1 : nRandomizationsPerSample)';
        for splitCounter = 1 : nSplitLevels
            if splitCounter == 1
                curSplitSize = nRandomizationsPerSample;
            elseif strcmp('base2',mySobolSensitivityOptions.subSampleSplitType)
                curSplitPower = (nSplitLevels + 1 - splitCounter);
                curSplitSize = 2^curSplitPower;
            end
            sampleSize(splitCounter,1) = curSplitSize;
            for bootstrapCounter = 1 : max(nBootstraps,1)
                if nBootstraps > 0
                    resampledIndices{bootstrapCounter,splitCounter} = datasample(theIndices,curSplitSize);
                else
                    resampledIndices{1,splitCounter} = theIndices;
                end
            end
        end
        for resultCounter = 1 : nResultIDs
            firstOrderMedian=nan(nSplitLevels,nAlteredAxisIDs+1);
            firstOrderUpperCI=nan(nSplitLevels,nAlteredAxisIDs+1);
            firstOrderLowerCI=nan(nSplitLevels,nAlteredAxisIDs+1);
            totalMedian=nan(nSplitLevels,nAlteredAxisIDs+1);
            totalUpperCI=nan(nSplitLevels,nAlteredAxisIDs+1);
            totalLowerCI=nan(nSplitLevels,nAlteredAxisIDs+1);  
            % We also add in the total variance in case we want to
            % back out the magnitude of the sensitivity indices later
            varianceMedian=nan(nSplitLevels,nAlteredAxisIDs+1);
            varianceUpperCI=nan(nSplitLevels,nAlteredAxisIDs+1);
            varianceLowerCI=nan(nSplitLevels,nAlteredAxisIDs+1);            
            colNames=cell(1,nAlteredAxisIDs+1);
            curResultID = myResultIDs{resultCounter};
            mySobolResults.(curResultID) = struct();
            % There are a few references for different ways to implement
            % the A, B, and Ci matrices to calculate the sensitivity indices.
            % Note that several options are presented in:
            % Saltelli, A., et al, Global Sensitivity Analysis:
            % The Primer. 2009. Pages 165-166.
            % Nossent and Bauwens compare
            % alternate implementations in:
            % Nossent, J. and W. Bauwens, Optimising the convergence of
            % a Sobol sensitivity analysis for an environmental model:
            % application of an appropriate estimate for the square of
            % the expectation value and the total variance, in 2012
            % International Congress on Environmental Modelling and Software,
            % R. Seppelt, et al., Editors. 2012: Leipzig, Germany.               
            for splitCounter = 1 : nSplitLevels
                curSplitSize=sampleSize(splitCounter,1);
                SplitSiValues = nan(nAlteredAxisIDs + 1, max(nBootstraps,1));
                SplitSTiValues = nan(nAlteredAxisIDs + 1, max(nBootstraps,1));
                SplitVarianceValues1 = nan(1,max(nBootstraps,1));
                SplitVarianceValues2 = nan(1,max(nBootstraps,1));
                for bootstrapCounter = 1 : max(nBootstraps,1) 
                    YBCur = YB(resampledIndices{bootstrapCounter,splitCounter},1,resultCounter);
                    YACur = YA(resampledIndices{bootstrapCounter,splitCounter},1,resultCounter);    
                    fosq_cross = (sum(YACur(:, 1, 1).*YBCur(:, 1, 1)))/curSplitSize;
                    sqfo_cross = (sum((YBCur(:, 1, 1)).^2)+sum((YACur(:, 1, 1)).^2))/(2*(curSplitSize-1)); 
                    % Note Saltelli 2009 p. 166 makes the argument the
                    % estimates for fo and the variance are improved by using
                    % both A and B points. 
                    % parfor didn't improve performance here
                    fosq_b = ((sum(YBCur(:, 1, 1)))/curSplitSize)^2;
                    sqfo_b = (sum((YBCur(:, 1, 1)).^2))/(curSplitSize - 1);
                    %fosq_a = ((sum(YACur(:, 1, 1)))/curSplitSize)^2;
                    %sqfo_a = (sum((YACur(:, 1, 1)).^2))/(curSplitSize - 1);
                     for axisCounter = 1 : nAlteredAxisIDs                          
                        YCCur = YC(resampledIndices{bootstrapCounter,splitCounter},axisCounter,resultCounter);
                        uhat_a = sum(YACur(:, 1, 1) .* YCCur(:,1,1))/(curSplitSize - 1);
                        uhat_b = sum(YBCur(:, 1, 1) .* YCCur(:,1,1))/(curSplitSize - 1);
                        % Both Nossent and Saltelli discuss the advantages
                        % of using A & B points, espcially in the Si's.
                        SplitSiValues(axisCounter,bootstrapCounter) = (uhat_a - fosq_cross)/(sqfo_cross - fosq_cross);
                        % Also tried this one, but it did not impact
                        % results
                        % SplitSiValues(axisCounter,bootstrapCounter) = (uhat_a - fosq_a)/(sqfo_a - fosq_a);
                        % With the test function, this version of the
                        % equation produced large confidence intervals
                        % for the STi, especially the noninfluential ones
                        % SplitSTiValues(axisCounter,bootstrapCounter) = 1-(uhat_b - fosq_cross)/(sqfo_cross - fosq_cross);
                        % Instead, this form gave small CI for the
                        % noninfluential terms.
                        SplitSTiValues(axisCounter,bootstrapCounter) = 1-(uhat_b - fosq_b)/(sqfo_b - fosq_b);
                     end
                     % We'll include an estimate for the total variance
                     % to return as well.  There are a few options to
                     % choose, although I didn't benchmark these
                     % directly.  I'll provide two estimates for now
                     % so they can be compared.                    
                     SplitVarianceValues1(bootstrapCounter) = (sqfo_cross - fosq_cross);
                     SplitVarianceValues2(bootstrapCounter) = (sqfo_b - fosq_b);
                end
                SplitSiValues(nAlteredAxisIDs + 1,:) = sum(SplitSiValues(1:nAlteredAxisIDs,:),1);
                SplitSTiValues(nAlteredAxisIDs + 1,:) = sum(SplitSTiValues(1:nAlteredAxisIDs,:),1);
                % hist(SplitSTiValues(6,:))
                if nBootstraps < 2
                    firstOrderMedian(splitcounter,:)=SplitSiValues;
                    firstOrderUpperCI(splitcounter,:)=nan(1,nAlteredAxisIDs + 1);
                    firstOrderLowerCI(splitcounter,:)=nan(1,nAlteredAxisIDs + 1);
                    totalMedian(splitcounter,:)=SplitSTiValues;
                    totalUpperCI(splitCounter,:)=nan(1,nAlteredAxisIDs + 1);
                    totalLowerCI(splitCounter,:)=nan(1,nAlteredAxisIDs + 1);
                    varianceEst1Median(splitCounter,:)=SplitVarianceValues1;
                    varianceEst1UpperCI(splitCounter,:)=nan;
                    varianceEst1LowerCI(splitCounter,:)=nan;
                    varianceEst2Median(splitCounter,:)=SplitVarianceValues2;
                    varianceEst2UpperCI(splitCounter,:)=nan;
                    varianceEst2LowerCI(splitCounter,:)=nan;
                else
                    firstOrderMedian(splitCounter,:)=(prctile(SplitSiValues,50,2))';
                    firstOrderUpperCI(splitCounter,:)=(prctile(SplitSiValues,97.5,2))';
                    firstOrderLowerCI(splitCounter,:)=(prctile(SplitSiValues,2.5,2))';
                    totalMedian(splitCounter,:)=(prctile(SplitSTiValues,50,2))';
                    totalUpperCI(splitCounter,:)=(prctile(SplitSTiValues,97.5,2))';
                    totalLowerCI(splitCounter,:)=(prctile(SplitSTiValues,2.5,2))';
                    varianceEst1Median(splitCounter,:)=(prctile(SplitVarianceValues1,50,2))';
                    varianceEst1UpperCI(splitCounter,:)=(prctile(SplitVarianceValues1,97.5,2))';
                    varianceEst1LowerCI(splitCounter,:)=(prctile(SplitVarianceValues1,2.5,2))';
                    varianceEst2Median(splitCounter,:)=(prctile(SplitVarianceValues2,50,2))';
                    varianceEst2UpperCI(splitCounter,:)=(prctile(SplitVarianceValues1,97.5,2))';
                    varianceEst2LowerCI(splitCounter,:)=(prctile(SplitVarianceValues1,2.5,2))';                    
                end   
            end
            mySobolResults.(curResultID).('firstOrderMedian') = flipud(firstOrderMedian);
            mySobolResults.(curResultID).('firstOrderUpperCI') = flipud(firstOrderUpperCI);
            mySobolResults.(curResultID).('firstOrderLowerCI') = flipud(firstOrderLowerCI);
            mySobolResults.(curResultID).('totalMedian') = flipud(totalMedian);
            mySobolResults.(curResultID).('totalUpperCI') = flipud(totalUpperCI);
            mySobolResults.(curResultID).('totalLowerCI') = flipud(totalLowerCI);
            mySobolResults.(curResultID).('varianceEst1Median') = flipud(varianceEst1Median);
            mySobolResults.(curResultID).('varianceEst1UpperCI') = flipud(varianceEst1UpperCI);
            mySobolResults.(curResultID).('varianceEst1LowerCI') = flipud(varianceEst1LowerCI);
            mySobolResults.(curResultID).('varianceEst2Median') = flipud(varianceEst2Median);
            mySobolResults.(curResultID).('varianceEst2UpperCI') = flipud(varianceEst2UpperCI);
            mySobolResults.(curResultID).('varianceEst2LowerCI') = flipud(varianceEst2LowerCI);            
        end
        mySobolResults.('time') = mySobolSensitivityOptions.analyzeTime;
        mySobolResults.('axisValues') = alteredCoefficients;
        mySobolResults.('axisDefIDs') = alteredAxisIDs;
        mySobolResults.('interventionID') = mySobolSensitivityOptions.interventionID;
        mySobolResults.('colNames') = cat(2,alteredAxisIDs,'total');
        mySobolResults.('sampleSize') = flipud(sampleSize);        
    else
        warning('Specified split type not yet supported. Exiting.')
    end
  
else
    warning(['Unable to run ',mfilename, '.'])
end
end
```
