## Supplementary File 3 for "A surrogate workflow for virtual population development: designing an amenable proxy for a hypothesis-test inspired nonlinear goodness-of-fit": runUnivariateSweep.m

```
function myFinalWorksheet = runUnivariateSweep(myWorksheet,myUnivariateSweepOptions)
% This function runs a local sweep of the parameter space around a single
% parameterization / VP. This function implements a simple
% approach whereby a base point (VP) is specified and we simply vary
% each parameter one at a time between the high and low bounds.
% This is meant to be run in series, prior to
% runUnivariateSweepOutputAnalysis
%
% ARGUMENTS
% myWorksheet:              An instance of a worksheet
% myUnivariateSweepOptions: An instance of a univariateSampleOptions
%                           object. This must be provided.
%
% RETURNS
% myWorksheet:              Final worksheet with sweep VPs, and with 
%                           simulation results if specified
%
continueFlag = false;
if nargin > 2
    warning(['Too many input arguments to ',mfilename, '. Arguments should be: myWorksheet and myUnivariateSweepOptions.'])
    continueFlag = false;
elseif nargin > 1
    continueFlag = true;
else
    warning(['Insufficient input arguments to ',mfilename, '. Arguments should be: myWorksheet and myUnivariateSweepOptions.'])
    continueFlag = false;
end

if continueFlag
    continueFlag = myUnivariateSweepOptions.verify(myWorksheet);
    if ~(continueFlag)
       warning(['Please correct the options provided to ',mfilename, '.']) 
    end
end

tic;
saveTime = 0;

if continueFlag
    
    % Prepare the worksheet
    removeInterventionIDs = getInterventionIDs(myWorksheet);
    removeInterventionIDs = setdiff(removeInterventionIDs, {myUnivariateSweepOptions.interventionID});
    myWorksheet=copyWorksheet(myWorksheet,{myUnivariateSweepOptions.baseVPID});
    myWorksheet = removeInterventions(myWorksheet, removeInterventionIDs);
    if myUnivariateSweepOptions.verbose
        disp(['Creating VPs in ',mfilename,'.']);
    end
    % Since this is a univariate sweep of axis values and we currently
    % only support the high/low point (we may make this variable in the
    % future), we just create the VPs here 
    allAxisIDs = getAxisDefIDs(myWorksheet);    
    nAxis = length(allAxisIDs);
    defaultCoefficients = getVPCoeffs(myWorksheet);
    nVaryAxis = length(myUnivariateSweepOptions.varyAxisIDs);
    testVPIDs = getVPIDs(myWorksheet);
    newVPIDs = cell(1,2*nVaryAxis);
    newVPVariants = cell(1,2*nVaryAxis);
    baseVP = getVP(myWorksheet,myUnivariateSweepOptions.baseVPID);
    baseVPIndex = find(ismember(testVPIDs,myUnivariateSweepOptions.baseVPID));
    newVPCounter = 0;
    newVPCoeffs = getVPCoeffs(myWorksheet);
    newVPCoeffs = newVPCoeffs(:,baseVPIndex);
    newVPCoeffs = newVPCoeffs * ones(1,nVaryAxis*2);
    for axisCounter = 1 : nVaryAxis
        curAxisID = myUnivariateSweepOptions.varyAxisIDs{axisCounter};
        newVPCounter = newVPCounter + 1;
        newVPIDString = [myUnivariateSweepOptions.baseVPID,'_axis_',num2str(axisCounter),'_low'];
        newVPIDs{1,newVPCounter} = newVPIDString;
        newVPvariants{1,newVPCounter} = baseVP.variants;
        newVPaxis{1,newVPCounter} = myWorksheet.axisProps.axisVP(:,baseVPIndex);
        axisIndex = find(ismember(allAxisIDs,curAxisID));
        newVPCoeffs(axisIndex,newVPCounter) = 0;
        newVPCounter = newVPCounter + 1;
        newVPIDString = [myUnivariateSweepOptions.baseVPID,'_axis_',num2str(axisCounter),'_high'];
        newVPIDs{1,newVPCounter} = newVPIDString;
        newVPvariants{1,newVPCounter} = baseVP.variants;
        newVPCoeffs(axisIndex,newVPCounter) = 1;
    end
    myWorksheet = createVPs(myWorksheet,newVPIDs,newVPvariants,newVPCoeffs);
    if myUnivariateSweepOptions.verbose
        disp(['Done.']);
    end      
    allVPIDs = getVPIDs(myWorksheet);

    myWorksheet.simProps.saveElementResultIDs = myUnivariateSweepOptions.saveElementResultIDs; 
    if myUnivariateSweepOptions.simulateWorksheet
        % Prepare for simulations
        mySimulateOptions = simulateOptions();
        % We need to know if runs fail   
        mySimulateOptions.filterFailedRunVPs=false;
        if myUnivariateSweepOptions.verbose
            disp(['Beginning simulations in ',mfilename,'.']);
        end
        myFinalWorksheet = simulateWorksheet(myWorksheet, mySimulateOptions);
        if myUnivariateSweepOptions.verbose
            disp(['Completed simulations in ',mfilename,'.']);
            disp(['Elapsed time is ',num2str(toc/60),' min.']);
        end
    else
        myFinalWorksheet = myWorksheet;
    end
    if length(myUnivariateSweepOptions.saveFileName) > 0
        saveWorksheet(myFinalWorksheet, myUnivariateSweepOptions.saveFileName);
    end
else
    warning(['Unable to run ',mfilename, '. Returning input worksheet.'])
    myFinalWorksheet = myWorksheet;
end
end
```
