## Supplementary File 3 for "A surrogate workflow for virtual population development: designing an amenable proxy for a hypothesis-test inspired nonlinear goodness-of-fit": runUnivariateSweepOutputAnalysis.m

```
function mySweepResults = runUnivariateSweepOutputAnalysis(myWorksheet,myUnivariateSweepOutputAnalysisOptions)
% This function takes a worksheet from a univariate sweep 
% (runUnivariateSweep) and gets the simulation results of interest
%
% ARGUMENTS
% myWorksheet:                            an instance of a worksheet, 
%                                         output from runUnivariateSweep()
% myUnivariateSweepOutputAnalysisOptions: an instance of a univariateSweepOutputAnalysisOptions
%                                         object
%
% RETURNS
% mySweepResults:            A data structure with the desired results.
%                            The fields are:
%                             (varname):      the (varname) field contains the
%                                             results from the analysis
%                                             Each is reported as a (nAxis + 1)
%                                             x 2 matrix
%                             time:           simulation time for which the
%                                             outputs are reported for
%                             axisDefIDs:     a 1 x nAxis + 1 cell array to record
%                                             the name of axes that were
%                                             varied.
%                             interventionID: a record of the intervention
%                                         
%
mySweepResults = struct();
continueFlag = false;
if nargin > 2
    warning(['Too many input arguments to ',mfilename, '. Arguments should be: myWorksheet and myUnivariateSweepOutputAnalysisOptions.'])
    continueFlag = false;
elseif nargin > 1
    continueFlag = true;
else
    warning(['Insufficient input arguments to ',mfilename, '. Arguments should be: myWorksheet and myUnivariateSweepOutputAnalysisOptions.'])
    continueFlag = false;
end

if continueFlag
    continueFlag = myUnivariateSweepOutputAnalysisOptions.verify(myWorksheet);
    if ~(continueFlag)
       warning(['Please correct the options provided to ',mfilename, '.']) 
    end
    myResultIDs = myUnivariateSweepOutputAnalysisOptions.analyzeElementResultIDs;
    nResultIDs = length(myResultIDs);
    if nResultIDs < 1
        warning(['At least one valid model output must be specified in analyzeElementResultIDs in options provided to ',mfilename,'.'])
        continueFlag = false;
    end    
end

if continueFlag
    allVPIDs = getVPIDs(myWorksheet);
    nVPs = length(allVPIDs);
    allInterventionIDs = getInterventionIDs(myWorksheet);
    myInterventionIndex = find(ismember(allInterventionIDs, myUnivariateSweepOutputAnalysisOptions.interventionID));
    Ymat = nan(nVPs,nResultIDs);
    for vpCounter = 1 : nVPs
        curResults = myWorksheet.results{myInterventionIndex,vpCounter};
        timeIndex = find(ismember(curResults.Names,'time'));
        timeVals = curResults.Data(:,timeIndex);
        timeIndex = find(timeVals == myUnivariateSweepOutputAnalysisOptions.analyzeTime);
        for resultCounter = 1 : nResultIDs
            % We take for granted non-degeneracy of referenced result
            % element ID's.
            resultIndex = find(ismember(curResults.Names,myResultIDs{resultCounter}));
            Ymat(vpCounter, resultCounter) = curResults.Data(timeIndex,resultIndex);
            
        end
    end
    for resultCounter = 1 : nResultIDs
        lowerIndices = [1,2:2:nVPs]';
        upperIndices = [1,3:2:nVPs]';
        mySweepResults.(myResultIDs{resultCounter}) = [Ymat(lowerIndices, resultCounter),Ymat(upperIndices, resultCounter)];
    end
    mySweepResults.('time') = myUnivariateSweepOutputAnalysisOptions.analyzeTime;
    mySweepResults.('axisDefIDs') = ['baseline',myUnivariateSweepOutputAnalysisOptions.varyAxisIDs];
    mySweepResults.('interventionID') = myUnivariateSweepOutputAnalysisOptions.interventionID;    
else
    warning(['Unable to run ',mfilename, '.'])
end
end
```
