## Supplementary File 3 for "A surrogate workflow for virtual population development: designing an amenable proxy for a hypothesis-test inspired nonlinear goodness-of-fit": screenWorksheetVPs.m

```
function myWorksheet = screenWorksheetVPs(myWorksheet, myScreenTable, verboseFlag, myScreenVPIDs)
% Screen VPs in a worksheet with a screen table.  This function will
% take a worksheet, use the responseTypes on that worksheet to compare
% to a screen table.
%
% ARGUMENTS
%  myWorksheet:              A worksheet structure, required.  The results
%                             all need to be populated.
%  myScreenTable:            (optional) A screen table resulting from
%                             createScreenTable.  If none is provided,
%                             all of the responseTypes on the worksheet
%                             will be used to create one, with the
%                             requirement that all passing VPs are
%                             evaluated with a value of zero for each
%                             weighted responseTypeElement.  Note that
%                             cutoff values other than 0 for each
%                             responseTypeElement can be used if you
%                             provide your own screenTable.
%  verboseFlag:             (optional) A boolean, whether to report summary
%                            results to the screen.  Default is true.
%  myScreenVPIDs            (optional) A subset of VPIDs to screen,
%                            Default is all wokrsheet VPs.  VPs
%                            not included in the screen list are allowed to
%                            pass.
%
% RETURNS
%  myWorksheet:             A screened worksheet.
%

continueFlag = false;
if nargin > 4
    warning(['Too many input arguments to ',mfilename, '. Arguments should be: myWorksheet; and optionally: myScreenTable, verboseFlag, myScreenVPIDs.'])
    continueFlag = false;
elseif nargin > 3
    continueFlag = true;    
elseif nargin > 2
    myScreenVPIDs = getVPIDs(myWorksheet);
    continueFlag = true;
elseif nargin > 1
    myScreenVPIDs = getVPIDs(myWorksheet);    
    verboseFlag = true;
    continueFlag = true;
elseif nargin > 0
    myScreenVPIDs = getVPIDs(myWorksheet);    
    continueFlag = true;
    verboseFlag = true;	
    flagSetBoundsFromResults = false;
    allResponseTypeIDs = getResponseTypeIDs(myWorksheet);
	myScreenTable = createScreenTable(myWorksheet, allResponseTypeIDs, flagSetBoundsFromResults);
else
	continueFlag = false;
    warning(['Insufficient input arguments to ',mfilename, '. Arguments should be: myWorksheet; and optionally: myScreenTable, verboseFlag, myScreenVPIDs.'])
end


if continueFlag
	% Check to see if we will populate the bounds from the
	% worksheet results.
    allVPIDs = getVPIDs(myWorksheet);
    interventionIDs = getInterventionIDs(myWorksheet);
    nVPs = length(allVPIDs);
    nInterventions = length(interventionIDs);
    [nInterventionResults, nVPResults] = size(myWorksheet.results);
    myResultClasses = cellfun(@class,myWorksheet.results, 'UniformOutput', false);
    % Results should be stored in a structure, we assume 
    % if a structure is provided then it is a valid result
    flagVPComplete = sum(strcmp(myResultClasses,'struct'),1) == nInterventions;
    
    if ~(sum(flagVPComplete) == nVPs)
		continueFlag = false;
        warning(['Unable to get complete results for worksheet provided to ',mfilename,'.'])
    end	
    
end		


if continueFlag
	% Check to see if all responseTypes from the screenTable
    % are in the worksheet.
    myResponseTypeIDs = unique(myScreenTable.responseTypeID);
    allResponseTypeIDs = getResponseTypeIDs(myWorksheet);
    if sum(~ismember(myResponseTypeIDs,allResponseTypeIDs)) > 0
        missingResponseTypeIDs = myResponseTypeIDs(~ismember(myResponseTypeIDs,allResponseTypeIDs));
        warning(['Unable to find responseTypeIDs:',strjoin(missingResponseTypeIDs,', '),' from the screenTable for the worksheet provided to ',mfilename,'.'])
        continueFlag = false;
    end
end


if continueFlag
	% Check to see if all responseTypeElements from the screenTable
    % are in the worksheet.
    nResponseTypes = length(myResponseTypeIDs);
    for responseTypeCounter = 1 : nResponseTypes
        checkRows = ismember(myScreenTable.responseTypeID,myResponseTypeIDs{responseTypeCounter});
        myResponseTypeElementIDs = unique(myScreenTable{checkRows,'responseTypeElementID'});
        allResponseTypeElementIDs = getResponseTypeElementIDs(myWorksheet,myResponseTypeIDs{responseTypeCounter});
        % Pre-VP summary values are also added to the calculations
        allResponseTypeElementIDs = [allResponseTypeElementIDs,'vpValues'];
        if sum(~ismember(myResponseTypeElementIDs,allResponseTypeElementIDs)) > 0
            missingResponseTypeElementIDs = myResponseTypeElementIDs(~ismember(myResponseTypeElementIDs,allResponseTypeElementIDs));
            warning(['Unable to find responseTypeElementIDs:',strjoin(missingResponseTypeElementIDs,', '),' from the screenTable for the worksheet provided to ',mfilename,'.'])
            continueFlag = false;
        end       
    end
end


if continueFlag
    % Split into screening and non-screening worksheets.
    myVPIDsNoScreen = setdiff(allVPIDs,myScreenVPIDs,'stable');
    myWorksheetNoScreen = copyWorksheet(myWorksheet, myVPIDsNoScreen);
    myWorksheet = copyWorksheet(myWorksheet, myScreenVPIDs);
    
    myCoeffs = getVPCoeffs(myWorksheet);
    [nAxis, nVPs] = size(myCoeffs);
    newInvalidIndices = nan(1,0);
    for responseTypeCounter = 1 : nResponseTypes
        curTable = createResponseSummaryTable(myWorksheet, allResponseTypeIDs{responseTypeCounter});
        checkRows = ismember(myScreenTable.responseTypeID,myResponseTypeIDs{responseTypeCounter});
        checkResponseTypeElementIDs = myScreenTable.responseTypeElementID(checkRows);
        curTableRowIndices = cellfun(@(c)find(strcmp(c,curTable.rowNames)),checkResponseTypeElementIDs,'UniformOutput',false);
        curTableRowIndices = cell2mat(curTableRowIndices);
        curFailChecks = curTable.values(curTableRowIndices,:)>myScreenTable.value(checkRows);
        curInvalidIndices = find(sum(curFailChecks,1)>0);
        curFailN = length(curInvalidIndices);
        if verboseFlag
            if length(curInvalidIndices) > 0
                newInvalidIndices = [newInvalidIndices, curInvalidIndices];
                curInvalidRTEN = sum(curFailChecks,2);
                curInvalidRTEIndices = find(curInvalidRTEN>0);
                [~,curInvalidRTEIndicesSort] = sort(curInvalidRTEN(curInvalidRTEIndices),'descend');
                curInvalidRTEIndices = curInvalidRTEIndices(curInvalidRTEIndicesSort);
                for invalidRTECounter = 1 : length(curInvalidRTEIndices)
                    disp(['Screen for responseTypeElement ',checkResponseTypeElementIDs{curInvalidRTEIndices(invalidRTECounter)},' in responseType ',myResponseTypeIDs{responseTypeCounter},' failed ',num2str(curInvalidRTEN(curInvalidRTEIndices(invalidRTECounter))),' of ',num2str(nVPs),' VPs, ',num2str(100*(curInvalidRTEN(curInvalidRTEIndices(invalidRTECounter)))/nVPs),'%.'])
                end
            end
        end
        
    end
    newInvalidIndices = sort(unique(newInvalidIndices),'ascend');
    splitVPIDs = getVPIDs(myWorksheet);	
    validIndices = find(~ismember([1:nVPs],newInvalidIndices));
    validVPIDs = splitVPIDs(validIndices);
    nPass = length(validIndices);    
    if verboseFlag
        disp(['Screening passed ',num2str(nPass),' of ',num2str(nVPs),' VPs in ',mfilename,', ',num2str(100*nPass/nVPs),'%.'])
    end
    myWorksheet = copyWorksheet(myWorksheet,validVPIDs); 
    myWorksheet = mergeWorksheets(myWorksheetNoScreen,myWorksheet);
    
else
    warning(['Unable to complete ',mfilename,'.  Exiting.'])
    myWorksheet = [];
end

end
```
