## Supplementary File 3 for "A surrogate workflow for virtual population development: designing an amenable proxy for a hypothesis-test inspired nonlinear goodness-of-fit": testClusterSizes.m

```
function clusterData = testClusterSizes(myWorksheet, myClusterTestOptions)
% Evaluate WCSS and TSS metrics using kmeans
%
% ARGUMENTS
% myWorksheet:        starting worksheet
% myClusterTestOptions:   optional, a cluster options object
%
% RETURNS
% clusterData:        a structure with the clustering results
%
%
clusterData = struct();
continueFlag = false;
% Verify the input arguments.
if nargin > 2
    warning(['Too many input arguments to ',mfilename, '. Arguments should be: myWorksheet, myClusterTestOptions.'])
    continueFlag = false; 
elseif nargin > 1
    allVPIDs = getVPIDs(myWorksheet);
    continueFlag = true;
elseif nargin > 0
    allVPIDs = getVPIDs(myWorksheet);
    myClusterTestOptions = clusterTestOptions();
    myClusterTestOptions = myClusterTestOptions.setDefaultFromWorksheet(myWorksheet);
    continueFlag = true; 
else
    warning(['Insufficient input arguments to ',mfilename, '. Arguments should be: myWorksheet, myClusterTestOptions.'])
    continueFlag = false; 
end

% Additional proofing that the input arguments make sense.
if continueFlag
    continueFlag = myClusterTestOptions.verify(myWorksheet);
end

if continueFlag
    if myClusterTestOptions.intSeed > -1
        rng(myClusterTestOptions.intSeed, 'twister');
    end

    % We need to create a matrix of data values that will then be
    % normalized and clustered.  It will be (nClusterAxis + nClusterOutput)
    % x nVP in size.
    nClusterAxis = length(myClusterTestOptions.clusterAxisIDs);
    [nClusterOutput, dummy] = size(myClusterTestOptions.clusterElement); 
    nVP = length(allVPIDs);
    theMatrixToCluster = nan((nClusterAxis + nClusterOutput), nVP);
    theRowNames = cell(nClusterAxis + nClusterOutput,1);
    theRowNames(1:nClusterAxis) = myClusterTestOptions.clusterAxisIDs;
    for theElementCounter = 1:nClusterOutput
        theName = [myClusterTestOptions.clusterElement{theElementCounter, 1},'___',myClusterTestOptions.clusterElement{theElementCounter, 2},'___',myClusterTestOptions.clusterElement{theElementCounter, 3},'___',num2str(myClusterTestOptions.clusterElement{theElementCounter, 4})];
        theRowNames{nClusterAxis+theElementCounter} = theName;
    end
    % Now we need to add the data
    worksheetAxisIndices = nan(nClusterAxis,1);
    allAxisIDs = getAxisDefIDs(myWorksheet);
    allInterventionIDs = getInterventionIDs(myWorksheet);
    for axisCounter = 1 : nClusterAxis
        worksheetAxisIndices(axisCounter) = find(ismember(allAxisIDs,myClusterTestOptions.clusterAxisIDs{axisCounter}));
    end
    allAxisCoefs = getVPCoeffs(myWorksheet);
    theMatrixToCluster(1:nClusterAxis,:) = allAxisCoefs(worksheetAxisIndices,:);
    % The outputs need to be retrieved for each VP separately...
    % TODO: might want to add some check here for whether results exist and
    % if not to simulateWorksheet
    for theElementCounter = 1 : nClusterOutput
        theElementName = myClusterTestOptions.clusterElement{theElementCounter, 1};
        theElementType = myClusterTestOptions.clusterElement{theElementCounter, 2};
        theInterventionID = myClusterTestOptions.clusterElement{theElementCounter, 3};
        theSampleTime = myClusterTestOptions.clusterElement{theElementCounter, 4};
        theInterventionIndex = find(ismember(allInterventionIDs, theInterventionID));
        % All VPs should be run with the same model, with the same result
        % elements
        theVPCounter = 1;
        curResultStruct = myWorksheet.results{theInterventionIndex,theVPCounter};
        timeIndex = find(ismember(curResultStruct.Names, 'time'));
        % We assume element IDs are unique, even without type information.
        elementIndex = find(ismember(curResultStruct.Names, theElementName));
        for theVPCounter = 1 : nVP
            curResultStruct = myWorksheet.results{theInterventionIndex,theVPCounter};
            curTime = curResultStruct.Data(:,timeIndex);
            curElement = curResultStruct.Data(:,elementIndex);
            y_new = interp1(curTime,curElement,theSampleTime,'linear');
            theMatrixToCluster(nClusterAxis+theElementCounter,theVPCounter)=y_new;
        end
    end

    originalMatrix = theMatrixToCluster;
    if sum(ismember(myClusterTestOptions.distance,'seuclidean')) > 0
        myDistance = 'sqeuclidean';
    elseif sum(ismember(myClusterTestOptions.distance,'correlation')) > 0
        myDistance = 'correlation';
    end
    % any variables with zero variation will be a constant, 0, following
    % normalization and there
    % will be zero distance among these points in the dimension with the
    % shared value
    if strcmp(myClusterTestOptions.normalizeType,'min-max') 
        denominatorTerm = max(theMatrixToCluster,[],2)-min(theMatrixToCluster,[],2) + ((max(theMatrixToCluster,[],2)-min(theMatrixToCluster,[],2))==0);
        theMatrixToCluster = (theMatrixToCluster - min(theMatrixToCluster,[],2)*ones(1, size(theMatrixToCluster,2)))./(denominatorTerm*ones(1, size(theMatrixToCluster,2)));
        theMatrixToCluster = theMatrixToCluster ;
    elseif strcmp(myClusterTestOptions.normalizeType,'z-score')
        denominatorTerm = std(theMatrixToCluster,[],2) + ((max(theMatrixToCluster,[],2)-min(theMatrixToCluster,[],2))==0);
        theMatrixToCluster = theMatrixToCluster./(denominatorTerm*ones(1, size(theMatrixToCluster,2)));
        theMatrixToCluster = theMatrixToCluster - mean(theMatrixToCluster,2)*ones(1, size(theMatrixToCluster,2));
    end
    
    % VPs should be rows for clustering
    theMatrixToCluster = transpose(theMatrixToCluster);
    
    nClusterMin = myClusterTestOptions.rangeNClusters(1);
    nClusterMax = myClusterTestOptions.rangeNClusters(2);    
    kMeanResult = cell(1, nClusterMax);
    
    if isempty(gcp('nocreate'))
        parpool;
    else
        delete(gcp)
        parpool;
    end      
    
    
    % computing the Total sum of squares matrix
    meanPoint = mean(theMatrixToCluster,1);
    meanPoint = theMatrixToCluster - ones(nVP,1)*meanPoint; 
    tss = sum(sum(meanPoint.*meanPoint));
    
    % First we need to assess the number of required clusters
    parfor nCluster = nClusterMin : nClusterMax
        curResult = struct();
        [idx,theMeans,sumd,D] = kmeans(theMatrixToCluster, nCluster, 'Distance', myDistance, 'Replicates',5,'MaxIter',100);
        % sumd within-cluster sums of point-to-medoid distances in the k-by-1 vector sumd
        % D distances from each point to every medoid in the n-by-k matrix D
        % We want theMeans, within-cluster SS, and total SS
        % We will calculate them.
        wcss = nan(nCluster,1);
        for theGroupCounter = 1:nCluster
            curIndices = find(idx==theGroupCounter);
            curPoints = theMatrixToCluster(curIndices,:);
            wcss(theGroupCounter) = sum(sum((curPoints - ones(length(curIndices),1)*theMeans(theGroupCounter,:)).^2,2));
        end
        curResult.('nCluster') = nCluster;
        curResult.('clusterMeans') = theMeans;
        curResult.('clusterMember') = idx;
        curResult.('wcsstss') = sum(wcss)/tss;
        kMeanResult{nCluster} = curResult;
    end

    kMeanResult = kMeanResult(nClusterMin : nClusterMax); 
    nClustersTested = length(kMeanResult);
    
    % For now, let's just return this result
    clusterMetric = nan(length(kMeanResult),2);
    for counter=1:length(kMeanResult)
        curResult = kMeanResult{counter};
        clusterMetric(counter, 1) = curResult.('nCluster');
        clusterMetric(counter, 2) = 1-curResult.('wcsstss');
    end
    if length(kMeanResult) > 5
        kernel = ones(5, 1) / 5; % 5x1 mean kernel
        % Convolve keeping size of clusterMetric
        newClusterMetric(:, 2) = conv(clusterMetric(:, 2), kernel, 'same');
        % Reset the edges that had the zero-convolution edges
        newClusterMetric(1, 2) = clusterMetric(1, 2);
        newClusterMetric(2, 2) = [1/3, 1/3, 1/3] * clusterMetric(1:3, 2);
        newClusterMetric(nClustersTested, 2) = clusterMetric(nClustersTested, 2);
        newClusterMetric(nClustersTested-1, 2) = [1/3, 1/3, 1/3] * clusterMetric(nClustersTested-2:nClustersTested, 2);
        clusterMetric(:, 2) = newClusterMetric(:, 2);
    end
    myCutoff = myClusterTestOptions.clusterCutoff;
    myIndices = clusterMetric(:, 2) >= myCutoff;
    if sum(myIndices) > 0
        myIndices = find(myIndices);
        myIndex = myIndices(1);
        myNClusters = clusterMetric(myIndex, 1);
    else
        myIndex = length(myIndices);
        myNClusters = clusterMetric(myIndex, 1);
    end
    
    clusterData.('kmeanTest') = clusterMetric;
    clusterData.('allVPIDs') = allVPIDs;
    clusterData.('clusteredMatrix') = transpose(theMatrixToCluster);
    clusterData.('untransformedMatrix') = originalMatrix;
    clusterData.('varNames') = theRowNames;
    clusterData.('nClusters') = myNClusters;
    
else
    warning(['Unable to run ',mfilename,'.'])
end
```
