## Supplementary File 3 for "A surrogate workflow for virtual population development: designing an amenable proxy for a hypothesis-test inspired nonlinear goodness-of-fit": getLinearIndependentVectors.m

```
function [VPCoeffsLI, VPind] = getLinearIndependentVectors(VPCoeffs)
% This function takes an input matrix where columns are
% VPs and tested for linear independence. 
% It will preferentially pick out columns in the order provided. 
%
% ARGUMENTS
%  VPCoeffs:	A matrix of VP coefficients
%
% RETURNS
%  VPCoeffsLI:	A matrix of VP coefficients
%				 that are linearly independent.
%  VPind:		Indices of the VPs in the returned
%				 matrix from the input matrix
%

% This will force to select the first vector of 
% collinar vector pairs, if any
%VPCoeffs = fliplr(VPCoeffs);
[nRows, nCols] = size(VPCoeffs);
VPCoeffsT = VPCoeffs';
R1=1;
% We will force what is probably a larger tolerance value to
% be more conservative
tolerance = 1E-8;
excludeInd = nan(1,0);
for I=1:size(VPCoeffsT,1)
    R2=rank(VPCoeffsT(1:I,:),tolerance);
    if R2~=R1
        excludeInd = [excludeInd,I];
    end
    R1=R2+1;
end
VPind = [1:nCols];
VPind(excludeInd) = [];
VPCoeffsLI = VPCoeffs(:,VPind);
end
```
