## Supplementary File 3 for "A surrogate workflow for virtual population development: designing an amenable proxy for a hypothesis-test inspired nonlinear goodness-of-fit": getMinimalEdgeSet.m

```
function edgeVPIDs = getMinimalEdgeSet(parentVPs)
% This function takes a cell array of cell arrays of identification strings
% as an imput and returns a cell of identification strings
% including at least one indentificaiton string from each set in the 
% original cell array of cell arrays.
%
% This function was written to help with the selection of a minimum set of
% edge VPs.  i.e. where each inner array 
% {{VPID1, VPID2}; {VPID2, VPID3, VPID4}; ... }
% Could have multiple VPs that are a tied for some characteristic, i.e.
% being and edge, and we want to provide a cell array of VPIDs that 
% includes one VP from all the edges, preferably the smallest set possible.
%
% Arguments:
%  parentVPs   A cell array of cell array of strings.  i.e.:
%               {{VPID1, VPID2}; {VPID2, VPID3, VPID4}; {VPID5}; ... }
%
% Returns:
%  edgeVPIDs   A cell array of strings containing at least one entry from
%               from every set in parentVPs.  i.e.:
%               {VPID2, VPID5, ... }
%

nParentCandidates = length(parentVPs);
parentVPsPass = cell(1,length(parentVPs));
parentsPassCounter = 1;
for checkCounter =1 :  nParentCandidates
    curCheckParents = parentVPs{checkCounter};
    % We need to compare to other parents in the list to
    % 1. pick the best parent that might cover multiple edges and
    % 2. minimize double counting for edges
    [~, curCheckSize] = size(curCheckParents);
    
    if checkCounter < nParentCandidates
        curCheckscore = zeros(nParentCandidates-(checkCounter),curCheckSize);
        % First check the current set of parents against the ones
        % further down the list in case one of the parents
        % in the current set appears there.
        for otherParentCounter = (checkCounter + 1) : nParentCandidates
            testParents = parentVPs{otherParentCounter};
            curCheckscore(otherParentCounter - checkCounter,:) = ismember(curCheckParents,testParents);
        end
        otherParentIndices = (checkCounter + 1) : nParentCandidates;
        % After we have the comparison for the current parents,
        % we pick the one with highest score as compared to
        % other rows
        sumParentScores = sum(curCheckscore,1);
        bestIndex = find(sumParentScores == max(sumParentScores));
        if length(bestIndex) > 1
            % In case of a tie
            bestIndex = bestIndex(1);
        end
        % Now we note the other VP sets that contained the best
        % parent
        curCheckscore = curCheckscore(:,bestIndex);
        % These are the rows where we found a match and can
        % substitute in the current best parent
        substituteOtherParentIndex = otherParentIndices(find(curCheckscore));
        if length(substituteOtherParentIndex) > 0
            parentVPs(substituteOtherParentIndex) = {{curCheckParents{bestIndex}}};
        end
        parentVPsPass{checkCounter} = curCheckParents{bestIndex};
        
    else
        % For the last entry, we do no comparison
        parentVPsPass{checkCounter} = curCheckParents{1};
    end
    
end
edgeVPIDs = unique(parentVPsPass,'stable');
end
```
