## Supplementary File 3 for "A surrogate workflow for virtual population development: designing an amenable proxy for a hypothesis-test inspired nonlinear goodness-of-fit": resamplePCASpace.m

```
function myVPs = resamplePCASpace(myNeighborsAndSelf,myVPIndex,relStd,nChildren,outputBounds)
% This function transform VPs into PCA space, resamples around the parent VP
% according to their variance, and transforms the new VPs back.
%
% ARGUMENTS
%  myNeighborsAndSelf: matrix with VP of interest and neightbording VPs.
%                     VPs are assumed to be column vectors.
%  myVPIndex:         column index of "parent" VP to sample around
%  relStd:            Relative scaling of standard deviation for sampling.  The value
%                     will be squared to scale the relative variance of resampling in 
%                     PCA space, which will also take into account the variance
%                     of each principal component
%  nChildren:         number of children VPs
%  outputBounds:      an nAxis x 2 matrix of bounds on the output
%                     values above or below will be truncated to this
%                     value.
%                     i.e. a matrix with rows [0 1] if the input is
%                     normalized between 0 and 1.
%
% RETURNS
%  myVPs:             VPs that have been resampled
%
[nDim,nParents] = size(myNeighborsAndSelf);
myNeighborsAndSelf=myNeighborsAndSelf';
myMean = mean(myNeighborsAndSelf,1);
myNeighborsAndSelf=myNeighborsAndSelf-mean(myNeighborsAndSelf,1);

% Use MATLAB's function
[eigenV,transData,varV,tsd,varN] = pca(myNeighborsAndSelf,'Algorithm','eig');

% Now that we have eigenVectors, transform just the VPs of interest
transData = myNeighborsAndSelf(myVPIndex,:)*eigenV;

% Sample in PCA space taking the eigenvalues and
% relative scaling into consideration
% reScale = 1/sqrt((2*pi())^nDim*sum(varV));
reScale = 1/sqrt(sum(varV));
transData=transData+reScale*mvnrnd(zeros(1,nDim),(relStd^2)*(varV.*eye(nDim)),nChildren);

% Transform back
myVPs = transData * (eigenV') + myMean;
myVPs = myVPs';
[replaceRow,replaceCol] = find(myVPs < outputBounds(:,1));
replaceIndices = find(myVPs < outputBounds(:,1));
if ~isempty(replaceRow)
    myVPs(replaceIndices) = outputBounds(replaceRow,1);
end
[replaceRow,replaceCol] = find(myVPs > outputBounds(:,2));
replaceIndices = find(myVPs > outputBounds(:,2));
if ~isempty(replaceIndices)
    myVPs(replaceIndices) = outputBounds(replaceRow,2);
end

end
```
