## Supplementary File 3 for "A surrogate workflow for virtual population development: designing an amenable proxy for a hypothesis-test inspired nonlinear goodness-of-fit": selectNbest.m

```
function newWorksheet = selectNbest(oldWorksheet, responseTypeID, NtoKeep)
% Select the N best VPs in a worksheet
%
% ARGUMENTS
% oldWorksheet: Starting worksheet
% baseVPid: ID string of the baseline VP to base the variations on
% responseTypeID: ID string of the response type to implement as the
%                 objective
% NtoKeep: The N best VPs will be kept
% MtoTry: The monte-carlo run will generate the M best VPs
% KrepeatM: Repeat the M tries K times, keeping the overall N best
%           This is available to keep Worksheet sizes smaller while still
%           using the available cores.
% verbose:  If true, write some information to screen so you can verify
%           process is executing
%
% RETURNS
% newWorksheet: A worksheet with the best VPs.
%

continueFlag = false;
% Verify the input arguments.
if nargin > 7
    warning(['Too many input arguments to ',mfilename, '. Arguments should be: oldWorksheet, baseVPid, responseTypeID; and optionally: NtoKeep, MtoTry, KrepeatM.'])
elseif nargin > 6
    continueFlag = true;
elseif nargin > 5
    verbose = true;    
    continueFlag = true;
elseif nargin > 4
    verbose = true;    
    KrepeatM = 1;
    continueFlag = true;
elseif nargin > 3
    verbose = true;    
    continueFlag = true;
    KrepeatM = 1;    
    MtoTry = 100;
    warning(['MtoTry missing in call to ',mfilename, '. Using ',num2str(MtoTry),'.']);
elseif nargin > 2
    verbose = true;    
    continueFlag = true;
    KrepeatM = 1;    
    MtoTry = 100;
    NtoKeep = 10;
    warning(['MtoTry and NtoKeep missing in call to ',mfilename, '. Using ',num2str(MtoTry),' and ',num2str(NtoKeep),'.']);
else
    warning(['Insufficient input arguments to ',mfilename, '. Arguments should be: oldWorksheet, baseVPid, responseTypeID; and optionally: NtoKeep, MtoTry, verbose.']);
end
    
% Additional proofing that the input arguments make sense.
if continueFlag
    % We only support monte-carlo right now
    varyMethod = 'montecarlo';
    testVPIDs = getVPIDs(oldWorksheet);
    if ~islogical(verbose)
        warning(strcat('Setting for verbose must be logical (true, false) in ',mfilename,'.'))
        continueFlag = false;  
    end
    if sum(ismember(testVPIDs, baseVPid)) < 1
        warning(strcat('VP ID not in worksheet: ',baseVPid,'.'))
        continueFlag = false;
    end
    testRTids = getResponseTypeIDs(oldWorksheet);
    if sum(ismember(testRTids, responseTypeID)) < 1
        warning(strcat('Response type ID not in worksheet: ',responseTypeID,' in ',mfilename,'.'))
        continueFlag = false;
    end    
    if sum(ismember({'montecarlo'},varyMethod)) < 1
        warning(['Selected varyMethod not supported  in ',mfilename,'.'])
        continueFlag = false;
    end
    if isnumeric(NtoKeep) == false
        warning(['Specify a number for NtoKeep in ',mfilename,'.'])
        continueFlag = false;
    end
    if isnumeric(MtoTry) == false
        warning(['Specify a number for MtoTry  in ',mfilename,'.'])
        continueFlag = false;
    end    
    if isnumeric(KrepeatM) == false
        warning(['Specify a number for KrepeatM in ',mfilename,'.'])
        continueFlag = false;
    end        
end

if continueFlag
    if NtoKeep > MtoTry
        warning(['NtoKeep should be larger than MtoTry in ',mfilename,'.'])
        continueFlag = false;
    end
end

if continueFlag
    % First pull out the base VP into a new worksheet
    newWorksheet = copyWorksheet(oldWorksheet, {baseVPid}, false);
    % Add the varied VPs
    tic
    for k = 1:KrepeatM 
        newWorksheet = addVariedVPs(newWorksheet, baseVPid, MtoTry, varyMethod, num2str(k));
        % Run the simulation
        newWorksheet = simulateWorksheet(newWorksheet);
        % Evaluate the response type and pick the best
        myResponseTypeResult = evaluateResponseType(newWorksheet, responseTypeID);
        myVPvalues = myResponseTypeResult.vpValues;
        myVPvalueCutoff = sort(myVPvalues, 'ascend');
        % It is possible if we have a lot of ties with an identical
        % objective value that we may keep extra VPs.        
        bestVPvalueCutoff = myVPvalueCutoff(1);
        bestVPindex = find(myResponseTypeResult.vpValues <= bestVPvalueCutoff);
        % Best case, we have at least NtoKeep, but this many may not have
        % complete successfully
        myVPvalueCutoff = myVPvalueCutoff(min(NtoKeep,sum(~isnan(myVPvalueCutoff))));
        myVPindices = find(myResponseTypeResult.vpValues <= myVPvalueCutoff);
        allVPIDs = getVPIDs(newWorksheet);
        vpToKeep = allVPIDs(myVPindices);
        baseVPid = allVPIDs{bestVPindex};
        % Now just pull the best VPs into a new worksheet.
        newWorksheet = copyWorksheet(newWorksheet, vpToKeep, true);
        if verbose
            disp(['Completed iteration ',num2str(k),' of ',num2str(KrepeatM),' in ',mfilename,'.']);
            disp(['Elapsed time in worksheet iterations is ',num2str(toc/60),' min.']);
        end
    end
end
        
if continueFlag == false
    warning(['Unable to run ',mfilename,', returning input worksheet.'])
    newWorksheet = oldWorksheet;
end
end
```
