## Supplementary File 3 for "A surrogate workflow for virtual population development: designing an amenable proxy for a hypothesis-test inspired nonlinear goodness-of-fit": varyAxes.m

```
function newWorksheet = varyAxes(oldWorksheet, myVaryAxesOptions)
% Create a worksheet by varying selected VPs randomly along their
% mechanistic axes.
%
% ARGUMENTS
% oldWorksheet:      Starting worksheet.  Note the mechanistic axes must be
%                    defined first before calling this function.
% myVaryAxesOptions: Options for creating new VPs with the varied axes
%
% RETURNS
% newWorksheet:      Worksheet that incudes the base VPs and the new VPs
%

% First check whether sufficient input arguments are provided
continueFlag = false;
if nargin > 2
    warning(['Too many input arguments to ',mfilename, '. Arguments should be: oldWorksheet; and optionally: myVaryAxesOptions.'])
    continueFlag = false;
elseif nargin > 1
    continueFlag = true;
elseif nargin > 0
    myVaryAxesOptions = varyAxesOptions();
    myVaryAxesOptions.baseVPIDs = getVPIDs(oldWorksheet);
    myVaryAxesOptions.varyAxisIDs = getAxisDefIDs(oldWorksheet);
    continueFlag = true;
else
    warning(['Insufficient input arguments to ',mfilename, '. Arguments should be: oldWorksheet; and optionally: myVaryAxesOptions.'])
end

% Check whether the input arguments make sense
if continueFlag
    passTestFlag = myVaryAxesOptions.verify(oldWorksheet);
    if ~passTestFlag
        continueFlag = false;
    end
    if length(myVaryAxesOptions.varyAxisIDs) < 1
        continueFlag = false;
        warning(['Must vary at least one axis in ',mfilename, '.'])
    end
    if length(myVaryAxesOptions.baseVPIDs) < 1
        continueFlag = false;
        warning(['Must select at least one base VP in ',mfilename, '.'])
    end
end

if continueFlag
    myBaseVPIDs = myVaryAxesOptions.baseVPIDs;
    % If we pass the checks, we can create the new worksheet and add the
    % VPs.
    newWorksheet = copyWorksheet(oldWorksheet, myBaseVPIDs, false);
    for vpCounter = 1 : length(myBaseVPIDs)
        subCallVaryOptions = myVaryAxesOptions;
        subCallVaryOptions.baseVPIDs = {myBaseVPIDs{vpCounter}};
        newWorksheet = addVariedVPs(newWorksheet, subCallVaryOptions);
    end
else
    warning(['Unable to run ',mfilename,', returning input worksheet.'])
    newWorksheet = oldWorksheet;
end
```
