## Supplementary File 3 for "A surrogate workflow for virtual population development: designing an amenable proxy for a hypothesis-test inspired nonlinear goodness-of-fit": checkUpdateObjectVersion.m

```
function myUpdatedObject = checkUpdateObjectVersion(myInputObject)
% This sample code checks objects against
% known characteristics of prior versions to make sure 
% the most up-to-date ones are being used
%
% ARGUMENTS: 
%  myInputObject:     A VPop, VPopRECIST, VPopRECISTnoBin, 
%                      mapelOptions, mapelOptionsRECIST, or 
%                      mapelOptionsRECISTnoBin object to convert.
%
% RETURNS:
%  myUpdatedObject    A VPop, VPopRECIST, VPopRECISTnoBin, 
%                      mapelOptions, mapelOptionsRECIST, or 
%                      mapelOptionsRECISTnoBin object with
%                      the new binTable.


continueFlag = true;

if nargin > 1
    continueFlag = false;
    warning(['Too many input arguments for ',mfilename,'. Should provide: a VPop, VPopRECIST, VPopRECISTnoBin, mapelOptions, mapelOptionsRECIST, or mapelOptionsRECISTnoBin.'])
    continueFlag = false;	
	
elseif nargin > 0
    continueFlag = true;
else
    warning(['Insufficient input arguments for ',mfilename,'. Should provide: a VPop, VPopRECIST, VPopRECISTnoBin, mapelOptions, mapelOptionsRECIST, or mapelOptionsRECISTnoBin.'])
    continueFlag = false;
end

if ~(isa(myInputObject,'VPop') || isa(myInputObject,'VPopRECIST') || isa(myInputObject,'VPopRECISTnoBin') || isa(myInputObject,'mapelOptions') || isa(myInputObject,'mapelOptionsRECIST') || isa(myInputObject,'mapelOptionsRECISTnoBin'))
	warning(['Wrong input arguments for ',mfilename,'. Should provide: a VPop, VPopRECIST, VPopRECISTnoBin, mapelOptions, mapelOptionsRECIST, or mapelOptionsRECISTnoBin.'])
	continueFlag = false;
end


if continueFlag
	if ~isempty(myInputObject.binTable)
		% This is a portion to update the bin tables from an old
		% format to the new one with allowed variable bin numbers.
		% The transition was made around in-house rev1043.	
		if sum(ismember(myInputObject.binTable.Properties.VariableNames,'binEdge1')) > 0
			myInputObject = updateBinTableFormat(myInputObject)
			disp(['Deprecated binTable format automatically patched in ',mfilename,'.  You may want to manually verify the columns are correct.'])
		end
    end
    myUpdatedObject = myInputObject;
	if isa(myInputObject, 'VPopRECISTnoBin') 
		% VPopRECISTnoBin objects are deprecated.
		disp(['Deprecated VPopRECISTnoBin format detected in ',mfilename,'.  Attempting to patch to the new combined VPopRECIST version automatically.'])
		myUpdatedObject = VPopRECIST;
        myUpdatedObject.coeffsTable=myInputObject.coeffsTable;	
		myUpdatedObject.coeffsDist=myInputObject.coeffsDist;
		myUpdatedObject.pwStrategy = 'direct';							  
        myUpdatedObject.indexTable = [];
        myUpdatedObject.binEdges = [];
        myUpdatedObject.binMidPoints = [];
        myUpdatedObject.binProbs = [];
        myUpdatedObject.pws = myInputObject.pws;
        myUpdatedObject.mnSDTable = myInputObject.mnSDTable;
        myUpdatedObject.binTable = myInputObject.binTable;
        myUpdatedObject.distTable = myInputObject.distTable;          
        myUpdatedObject.distTable2D = myInputObject.distTable2D;	
		myUpdatedObject.corTable = myInputObject.corTable;
		myUpdatedObject.brTableRECIST = myInputObject.brTableRECIST;
		myUpdatedObject.rTableRECIST = myInputObject.rTableRECIST;
        myUpdatedObject.expData = myInputObject.expData;          
        myUpdatedObject.simData = myInputObject.simData;
        myUpdatedObject.gofMn = myInputObject.gofMn;
        myUpdatedObject.gofSD = myInputObject.gofSD;
        myUpdatedObject.gofBin = myInputObject.gofBin;
        myUpdatedObject.gofDist = myInputObject.gofDist;          
		myUpdatedObject.gofDist2D = myInputObject.gofDist2D;  
		myUpdatedObject.gofCor = myInputObject.gofCor;
		myUpdatedObject.gofBR = myInputObject.gofBR;
		myUpdatedObject.gofR = myInputObject.gofR;
        myUpdatedObject.gof = myInputObject.gof;          
        myUpdatedObject.spreadOut = myInputObject.spreadOut;
        myUpdatedObject.minIndPVal = myInputObject.minIndPVal;			  
        myUpdatedObject.useEffN = myInputObject.useEffN;
        myUpdatedObject.exactFlag = myInputObject.exactFlag; 
        myUpdatedObject.optimizeTimeLimit = myInputObject.optimizeTimeLimit;
        myUpdatedObject.optimizeType = myInputObject.optimizeType;    
        myUpdatedObject.optimizePopSize = myInputObject.optimizePopSize;    
		myUpdatedObject.objectiveLimit = myInputObject.objectiveLimit;
        myUpdatedObject.intSeed = myInputObject.intSeed;
        myUpdatedObject.tol = myInputObject.tol;
        myUpdatedObject.nIters = myInputObject.nIters;
        myUpdatedObject.minEffN = myInputObject.minEffN;
        myUpdatedObject.relSLDvar = myInputObject.relSLDvar;
        myUpdatedObject.absALDVar = myInputObject.absALDVar;
        myUpdatedObject.crCutoff = myInputObject.crCutoff;            
        myUpdatedObject.recistSimFilter = myInputObject.recistSimFilter;		
		disp(['VPopRECISTnoBin automatically updated.  You may want to manually verify it looks OK.'])		
	elseif isa(myInputObject, 'mapelOptionsRECISTnoBin')
		% mapelOptionsRECISTnoBin objects are deprecated.
		disp(['Deprecated mapelOptionsRECISTnoBin format detected in ',mfilename,'.  Attempting to patch to the new combined mapelOptionsRECIST version automatically.'])
		myUpdatedObject = mapelOptionsRECIST;
        myUpdatedObject.expData = myInputObject.expData;
        myUpdatedObject.mnSDTable = myInputObject.mnSDTable;
        myUpdatedObject.binTable = myInputObject.binTable;
        myUpdatedObject.distTable = myInputObject.distTable;    
		myUpdatedObject.distTable2D = myInputObject.distTable2D; 
		myUpdatedObject.corTable = myInputObject.corTable; 
        myUpdatedObject.brTableRECIST = myInputObject.brTableRECIST;
        myUpdatedObject.rTableRECIST = myInputObject.rTableRECIST; 
		myUpdatedObject.pwStrategy = 'direct';		  
        myUpdatedObject.nBins = 2;
        myUpdatedObject.initialProbs = [];
        myUpdatedObject.equalBinBreaks = false;		
        myUpdatedObject.initialPWs = myInputObject.initialPWs;		  
        myUpdatedObject.randomStart = myInputObject.randomStart;
        myUpdatedObject.nIters = myInputObject.nIters;
        myUpdatedObject.tol = myInputObject.tol;
        myUpdatedObject.spreadOut = myInputObject.spreadOut;
        % Older mapelOptions class definitions may have
        % a missing minIndPVal
        if ~isempty(myInputObject.minIndPVal)
            myUpdatedObject.minIndPVal = myInputObject.minIndPVal;		
        else
            myUpdatedObject.minIndPVal = 0;
        end
        myUpdatedObject.useEffN = myInputObject.useEffN;
        myUpdatedObject.exactFlag = myInputObject.exactFlag;
        myUpdatedObject.optimizeTimeLimit = myInputObject.optimizeTimeLimit;
        myUpdatedObject.optimizeType = myInputObject.optimizeType;
        myUpdatedObject.optimizePopSize = myInputObject.optimizePopSize;
        % Older mapelOptions class definitions may have
        % a missing minIndPVal
        if ~isempty(myInputObject.objectiveLimit)
            myUpdatedObject.objectiveLimit = myInputObject.objectiveLimit;		
        else
            myUpdatedObject.objectiveLimit = -Inf;
        end		  
        myUpdatedObject.intSeed = myInputObject.intSeed;   
        myUpdatedObject.minEffN = myInputObject.minEffN;     
        myUpdatedObject.relSLDvar = myInputObject.relSLDvar;
        myUpdatedObject.absALDVar = myInputObject.absALDVar;
        myUpdatedObject.crCutoff = myInputObject.crCutoff;  
		disp(['mapelOptionsRECISTnoBin automatically updated in ',mfilename,'.  You may want to manually verify it looks OK.'])	
	end
	if isa(myUpdatedObject, 'VPopRECIST') || isa(myUpdatedObject, 'VPop')
        [nRows, nCols] = size(myUpdatedObject.distTable2D);
        if nRows > 0
            if sum(ismember(myUpdatedObject.distTable2D.Properties.VariableNames,{'combinedQuadrants'})) > 0
                myUpdatedObject.distTable2D.('combinedQuadrants') = [];
                disp(['VPop/VPopRECIST distTable2D automatically updated to remove combinedQuadrants in ',mfilename,'.  You may want to manually verify it looks OK.'])
            end
        end
    end
    % Also add the subpopulation number column, if needed.
	myUpdatedObject = updateTablesWithSubpopReference(myUpdatedObject);
    % Convert the subpopulation into cell table format, if needed.
	myUpdatedObject = updateTablesWithSubpopformat(myUpdatedObject);
    % Also check if the new predIndices columns are added to the mnSDTable and binTable
    commonNames = loadCommonNames();
	if isa(myUpdatedObject, 'VPopRECIST') || isa(myUpdatedObject, 'VPop') || isa(myUpdatedObject, 'mapelOptionsRECIST') || isa(myUpdatedObject, 'mapelOptions')   
        myMnSDTable = myUpdatedObject.mnSDTable;
        [nRows, ~] = size(myMnSDTable);
        if nRows > 0
            if sum(ismember(myMnSDTable.Properties.VariableNames,'predIndices')) < 1
                predIndices = repmat({{nan}},nRows,1);
                if isa(myUpdatedObject,'VPop') || isa(myUpdatedObject,'mapelOptions')
                    tableVariableNames = commonNames.VPOPTABLEVARNAMESFIXED;
                    nDataHeaderCols = length(commonNames.VPOPTABLEVARNAMESFIXED);
                else
                    tableVariableNames = commonNames.VPOPRECISTTABLEVARNAMESFIXED;
                    nDataHeaderCols = length(commonNames.VPOPRECISTTABLEVARNAMESFIXED);
                end
                tableVariableNames = [tableVariableNames,{'weightMean', 'weightSD', 'expN', 'expMean', 'expSD', 'predN', 'predIndices', 'predMean', 'predSD'}];
                myMnSDTable = [myMnSDTable, cell2table(predIndices)];
                myMnSDTable = myMnSDTable(:,tableVariableNames);
                myUpdatedObject.mnSDTable = myMnSDTable;
                disp(['mnSDTable table automatically updated to include predIndices column in ',mfilename,'.'])
            end
			if sum(ismember(myMnSDTable.Properties.VariableNames,'logN')) < 1
				logN = false(nRows,1);
                if isa(myUpdatedObject,'VPop') || isa(myUpdatedObject,'mapelOptions')
                    tableVariableNames = commonNames.VPOPTABLEVARNAMESFIXED;
                    nDataHeaderCols = length(commonNames.VPOPTABLEVARNAMESFIXED);
                else
                    tableVariableNames = commonNames.VPOPRECISTTABLEVARNAMESFIXED;
                    nDataHeaderCols = length(commonNames.VPOPRECISTTABLEVARNAMESFIXED);
                end
                tableVariableNames = [tableVariableNames,{'logN','weightMean', 'weightSD', 'expN', 'expMean', 'expSD', 'predN', 'predIndices', 'predMean', 'predSD'}];
                myMnSDTable = [myMnSDTable, array2table(logN)];
                myMnSDTable = myMnSDTable(:,tableVariableNames);
                myUpdatedObject.mnSDTable = myMnSDTable;
                disp(['mnSDTable table automatically updated to include lognormal column in ',mfilename,'.'])
			end
			
        end
    end
	if isa(myUpdatedObject, 'VPopRECIST') || isa(myUpdatedObject, 'VPop') || isa(myUpdatedObject, 'mapelOptionsRECIST') || isa(myUpdatedObject, 'mapelOptions')   
        myBinTable = myUpdatedObject.binTable;
        [nRows, ~] = size(myBinTable);
        if nRows > 0
            if sum(ismember(myBinTable.Properties.VariableNames,'predIndices')) < 1
                predIndices = repmat({{nan}},nRows,1);
                if isa(myUpdatedObject,'VPop') || isa(myUpdatedObject,'mapelOptions')
                    tableVariableNames = commonNames.VPOPTABLEVARNAMESFIXED;
                    nDataHeaderCols = length(commonNames.VPOPTABLEVARNAMESFIXED);
                else
                    tableVariableNames = commonNames.VPOPRECISTTABLEVARNAMESFIXED;
                    nDataHeaderCols = length(commonNames.VPOPRECISTTABLEVARNAMESFIXED);
                end
                tableVariableNames = [tableVariableNames,{'weight','binEdges','expN','expBins','predN','predIndices','predBins'}];
                myBinTable = [myBinTable, cell2table(predIndices)];
                myBinTable = myBinTable(:,tableVariableNames);
                myUpdatedObject.binTable = myBinTable;
                disp(['binTable automatically updated to include predIndices column in ',mfilename,'.'])
            end
        end
    end
    if isa(myUpdatedObject, 'VPopRECIST') || isa(myUpdatedObject, 'VPop')
        % Also make sure these are up-to-date, in case an earlier version
        % is loaded
        if ~isempty(myUpdatedObject.subpopTable)
            myUpdatedObject=myUpdatedObject.addTableSimVals;
        else
            disp(['No subpopTable available in the current ',class(myUpdatedObject),' object in ',mfilename,'.  Be sure to run createSubpopTable() with the relevant worksheet, and then run the addTableSimVals to make sure the data is up-to-date.'])
        end
    else
        if isempty(myUpdatedObject.subpopTable)
            disp(['No subpopTable available in the current ',class(myUpdatedObject),' object in ',mfilename,'.  Be sure to run createSubpopTable() with the relevant worksheet.'])
        end
    end
else
	myUpdatedObject = myInputObject;
	warning(['Unable to run ',mfilename,'. Returning input object.'])
end
```
