## Supplementary File 3 for "A surrogate workflow for virtual population development: designing an amenable proxy for a hypothesis-test inspired nonlinear goodness-of-fit": expandVPopEffNAutoSplitSave.m

```
function [myWorksheet, newVPop] = expandVPopEffNAutoSplitSave(myWorksheet,myExpandVPopEffNOptions,myMapelOptions, hotStartVPop)
% This function takes a worksheet, options object
% mapelOptions object, as well as a reference 
% worksheet for response type bounds,
% tries to fit a VPop. Once successful it adds
% VPs to the worksheet, then tries to fit a VPop with higher effN,
% expanding the effective N in MAPEL in through consecutive iterations.  
% Essentially, it will run MAPEL iteratively, generating VPops until it 
% finds one with an acceptable GOF, generate new VPs based on the most 
% weighted VPs, and select VPs from each most weighted "seed" VP to add.
% Worksheets and VPops are written to file as the algorithm progresses.
% After each iteration, acceptable VPop results and the new
% worksheet are written to file, before the minEffN is increased in MAPEL.
% The algorithm will keep running until it finds a VPop that satisfies the 
% target effN.
% 
% ARGUMENTS:
%  myWorksheet:             A worksheet to expand
%  myExpandVPopEffNOptions: An expandVPopEffNOptions object
%  myMapelOptions:          A mapelOptions object
%  hotStartVPop:            A VPop object, will try to hot
%                            start from this solution if provided.
%                            Leave empty or '' if no hot start desired.
%
% RETURNS:
%  myWorksheet
%  newVPop
%
%  NOTES: 
%   The algorithm
%   may encounter problems if an old suffix and
%   wsIterCounter combination are re-used, if existing 
%   VPs in the worksheet have names identical to those 
%   selected by the algorithm.
%

continueFlag = false;
if nargin > 4
    warning(['Too many input arguments to ',mfilename, '. Arguments should be: myWorksheet, myExpandVPopEffNOptions, and myMapelOptions. Optionally: hotStartVPop.'])
    continueFlag = false;
    hotStartVPop = '';    
elseif nargin > 3
    continueFlag = true;
else 
    warning(['Insufficient input arguments to ',mfilename, '. Arguments should be: myWorksheet, myExpandVPopEffNOptions, and myMapelOptions. Optionally: hotStartVPop.'])
    continueFlag = false;
end

if continueFlag
    if ~strcmp(class(myExpandVPopEffNOptions),'expandVPopEffNOptions')
        warning(['Argument myExpandVPopEffNOptions to ',mfilename, ' should be of class expandVPopEffNOptions.'])
        continueFlag = false;
    end
    if ~ismember(class(myMapelOptions),{'mapelOptions','mapelOptionsRECIST','mapelOptionsRECISTnoBin'})
        warning(['Argument myMapelOptions to ',mfilename, ' should be of class mapelOptions or mapelOptionsRECIST or mapelOptionsRECISTnobin.'])
        continueFlag = false;
    end      
end 
    
if continueFlag
    allResponseTypeIDs = getResponseTypeIDs(myWorksheet);
    if length(myExpandVPopEffNOptions.plausibleResponseTypeIDs) == 0
        myExpandVPopEffNOptions.plausibleResponseTypeIDs = allResponseTypeIDs;
        disp(['No plausibleResponseTypeIDs defined by myExpandVPopEffNOptions in call to ',mfilename, '.  Using all worksheet responseTypeIDs.'])
        continueFlag = true;
    elseif sum(ismember(myExpandVPopEffNOptions.plausibleResponseTypeIDs,allResponseTypeIDs)) < length(myExpandVPopEffNOptions.plausibleResponseTypeIDs)
        warning(['The response types for myExpandVPopEffNOptions were not all found in the worksheet provided to ',mfilename, '.'])
        continueFlag = false;
    end   
end

if continueFlag    
    
    % INITIALIZATION STARTS
    suffix = myExpandVPopEffNOptions.suffix;
    wsIterCounter = myExpandVPopEffNOptions.wsIterCounter;
    targetEffN = myExpandVPopEffNOptions.targetEffN;
    maxNewPerIter = myExpandVPopEffNOptions.maxNewPerIter;
    effNDelta = myExpandVPopEffNOptions.effNDelta;
    minPVal = myExpandVPopEffNOptions.minPVal;
    nTries = myExpandVPopEffNOptions.nTries;
    nRetries = myExpandVPopEffNOptions.nRetries;
    verbose = myExpandVPopEffNOptions.verbose;
    expandCohortSize = myExpandVPopEffNOptions.expandCohortSize;
    restartPVal = myExpandVPopEffNOptions.restartPVal;

    % Maximal optimizePopSize.  We'll try to use smaller to save time.
    maxOptimizePopSize = myMapelOptions.optimizePopSize;

    % Set these for fast optimization
    myMapelOptions.exactFlag = false;
    % Enforce using effN, since
    % we enforce using it for the VPops
    myMapelOptions.useEffN = true;
    % Also minimize pool restarts
    myMapelOptions.poolRestart=false;
    myMapelOptions.poolClose=false;
    
    % We will manually refresh the pool once at the start.
    if ~isempty(gcp('nocreate'))
        delete(gcp);
    end
    if isempty(gcp('nocreate'))
        % We will use default pool settings
        mySimulateOptions = simulateOptions;
        mySimulateOptions = checkNWorkers(mySimulateOptions);		
        myPool = parpool(mySimulateOptions.clusterID,mySimulateOptions.nWorkers,'SpmdEnabled',false);
    end	    

    % Make sure results are present.  Here, we don't re-simulate VPs if 
    % results are present.
    mySimulateOptions.rerunExisting=false;
	mySimulateOptions.poolRestart=false;
	mySimulateOptions.poolClose=false; 
    myWorksheet = simulateWorksheet(myWorksheet,mySimulateOptions);

	% Set up a reference worksheet size and effN
	% to help with clustering decisions
    startNVPs = length(getVPIDs(myWorksheet));
    startEffN = myMapelOptions.minEffN;	
    % Basis and nVPs for clustering
	basisEffN = 0;    
	nSteps = floor(((startEffN - basisEffN) / effNDelta)); 
	basisNVPs = floor(startNVPs - nSteps * maxNewPerIter);
	
	curEffN = startEffN;

    % Create a screen table
    % so we can enforce new VPs will need pass within bounds
    % set by the starting worksheet.
    myScreenTable = createScreenTable(myWorksheet, myExpandVPopEffNOptions.plausibleResponseTypeIDs, true);
    
    % Note intSeed will be incremented +1 soon after.
    if (~ismember(class(hotStartVPop),{'VPop','VPopRECIST'}))
        nVPopsFound = 0;
        if (myExpandVPopEffNOptions.useMapelIntSeed) && (myMapelOptions.intseed > -1)
            intSeedTarget = myMapelOptions.intseed-1;
        else
            intSeedTarget = -1;
        end
    else
        nVPopsFound = 1;
        oldVPop = hotStartVPop;
        oldVPop.useEffN = true;
        oldVPop.exactFlag = true;
        oldVPop = evaluateGOF(oldVPop);	
        if (myExpandVPopEffNOptions.useMapelIntSeed) && (myMapelOptions.intseed > -1)    
            intSeedTarget = myMapelOptions.intseed-1;
        else
            intSeedTarget = oldVPop.intSeed-1;
        end
    end
    
    
    
    % INITIALIZATION ENDS
	

    % START ITERATIONS
    % OUTER LOOP
    while curEffN < (targetEffN + effNDelta)
        % Set the initial best p value to a negative
		% number to force the algorithm to record
		% and write out the first VPop calibration
		% attempt, especially to aid troubleshooting
		% if calibrations fail.
        bestPVal = -1;
        myMapelOptions.minEffN = curEffN;
        myTestCounter = 0;		
		
        % NOW FOR A GIVEN EFFN ITERATE UNTIL WE FIND A GOOD VPOP
        % AND EXPAND THE WORKSHEET WHEN WE DO
        while bestPVal < minPVal
            myTestCounter = myTestCounter + 1;
            % We check whether to reseed with myTestCounter
            % for repeatability of the sequence
            intSeedTarget = intSeedTarget+1;
            myMapelOptions.intSeed=intSeedTarget;
            
            % We will try to restart with initial probabilities, if
            % available, and refresh worksheet data if this is the the first
            % iteration.  We'll also enforce the MAPEL options in
            % case they are tweaked.  Note we don't just directly restart
            % since there may be new simData in the worksheet.
            
            if (myTestCounter == 1) && (nVPopsFound > 0)
                myExtraPWs = [];                
                % This is the first iteration with the current effN
                % and we have already found VPops before
                newVPop = initializeOptionPropertiesToVPop(myMapelOptions);
                if isa(myMapelOptions,'mapelOptionsRECIST')                   
                    newVPop.recistSimFilter = createRECISTSimFilter(myWorksheet, newVPop, false);                    
                end		
                newVPop = newVPop.getSimData(myWorksheet);
                newVPop.subpopTable = updateSubpopTableVPs(newVPop.subpopTable,myWorksheet);
                if ismember(newVPop.pwStrategy,'bin')
                    newVPop = newVPop.assignIndices(myWorksheet, myMapelOptions);
                    % We assume the old bin probs are compatible
                    % with the new indices assignments.  That is,
                    % there are an equal number of bins and axes.
                    newVPop.binProbs = oldVPop.binProbs;
                else
                    % Otherwise we are weighting on VPs directly.
                    % We need to assign pws for added vps, also
                    % the CoeffsTable
                    newVPop=newVPop.assignCoeffs(myWorksheet);
                    [~,nVP_nobin]=size(newVPop.coeffsTable);
                    nVP_diff=nVP_nobin-length(oldVPop.pws);
                    if nVP_diff > 0
                        % If we have more VPs to calibrate in this
                        % VPop than the last, we can provide an
                        % intelligent guess on spreading the PWs
                        % to the new VPs.
                        newVPop.pws = [oldVPop.pws,zeros(1,nVP_diff)];
                        newVPop.pws = transferPWtoChildren(newVPop,nVP_diff,0);
                        myExtraPWs = transferPWtoChildren(newVPop,nVP_diff,2);
                    else
                        % If there are the same number of VPs
                        % just use the old PWs
                        % nVP_diff should not be < 0
                        % pws_addition=sum(ones(1,nVP_diff)*1/nVP_nobin);
                        % newVPop.pws=[oldVPop.pws*(1-pws_addition),
                        % ones(1,nVP_diff)*1/nVP_nobin]; 
                        newVPop.pws = oldVPop.pws;
                    end
                end
                
                
                % TEST: IF WE ARE LOADING IN A FULLY SUCCESSFUL
                % VPOP SO WE CAN SKIP MAPEL               
                newVPop.useEffN = true;
                newVPop.exactFlag = true;
                newVPop=addTableSimVals(newVPop);                
                newVPop = evaluateGOF(newVPop);
				newVPop.exactFlag = myMapelOptions.exactFlag;
                curVPopEffN = 1/sum(newVPop.pws.^2);
                if ~(~(newVPop.gof < minPVal) && (curVPopEffN >= curEffN))
                    % We randomize here with magnitude 
                    % myExpandVPopEffNOptions.expandRandomStart
                    % If the magnitude is set to infinity, we just
                    % fully randomize before "restarting"
                    % If set to zero, no randomization will be applied
                    if isinf(myExpandVPopEffNOptions.expandRandomStart)
                        if ismember(newVPop.pwStrategy,'direct')
                            newVPop = newVPop.startPWs(myWorksheet,true);
                        else
                            newVPop = newVPop.startProbs(true);
                        end	
                        newVPop = restartMapel(newVPop, 0, myExtraPWs);
                    else
                        newVPop = restartMapel(newVPop, myExpandVPopEffNOptions.expandRandomStart, myExtraPWs);
                    end
                end
            else
                % For consistency, on the first iteration
                % if a VPop isn't provided we will use the
                % myExpandVPopEffNOptions.expandRandomStart
                % value to decide randomization status.
                curMapelOptions = myMapelOptions;
                if (myTestCounter == 1) 
                    curMapelOptions.randomStart = myExpandVPopEffNOptions.expandRandomStart;
                end
                newVPop = mapel(myWorksheet, curMapelOptions);
            end
            
            % Now check the results for the current fresh start
            % iteration
            % Force the exact flag for this check.
            newVPop.useEffN = true;
            newVPop.exactFlag = true;
            newVPop = evaluateGOF(newVPop);
            curVPopEffN = 1/sum(newVPop.pws.^2);
            if verbose
                disp(['Previous best pvalue ',num2str(max(0,bestPVal)),' with effN ',num2str(curEffN),'.'])
                disp(['Current solution effN ', num2str(curVPopEffN), ' with pvalue ',num2str(newVPop.gof),' in ',mfilename,'.'])
                disp(['Current iteration ', num2str(myTestCounter),'.'])
            end
            
            % If the results didn't look too bad, we will restart
            if ((newVPop.gof < minPVal) && (newVPop.gof >= restartPVal) && (curVPopEffN >= curEffN))
                disp(['This pvalue not quite at acceptance, restarting MAPEL with this as initial guess.  Using myMapelOptions.randomStart value to set the magnitude of randomization.'])
                myRandomStart = myMapelOptions.randomStart;
                restartCounter = 1;
                if (newVPop.gof > bestPVal)
                    % Save here in case this is the best
                    mySaveName = ['vpop_',suffix,'_iter',num2str(wsIterCounter),'_effN',num2str(curEffN),'_curBest'];
                    saveVPop(newVPop, mySaveName);
                    bestPVal = newVPop.gof;   
                end								
                while (((restartCounter <= nRetries) && ~(newVPop.gof ...
                                                          >= minPVal)) || ((newVPop.gof > (0.96 * minPVal)) && ~(newVPop.gof >= minPVal)))
                    % We will keep trying a restart until we get an
                    % OK value or we run out of retries.  If we're
                    % really close we will let the algorithm
                    % keep iterating.
                    restartVPop = newVPop;
                    restartVPop.useEffN = myMapelOptions.useEffN;
                    restartVPop.exactFlag = myMapelOptions.exactFlag;
                    intSeedTarget = intSeedTarget+1;
                    restartVPop.intSeed = intSeedTarget;
                    % Decrease the tolerance stringency for restarting
                    restartVPop.tol = newVPop.tol*0.1;
                    restartVPop = restartMapel(restartVPop, myRandomStart);
                    restartVPop.tol = newVPop.tol;
                    restartVPop.useEffN = true;
                    restartVPop.exactFlag = true;
                    restartVPop = evaluateGOF(restartVPop);				
                    restartVPopEffN = 1/sum(restartVPop.pws.^2);
                    if verbose
                        disp(['Previous best pvalue ',num2str(max(0,bestPVal)),' with effN ',num2str(curEffN),'.'])
                        disp(['Current restart solution effN ', num2str(restartVPopEffN), ' with pvalue ',num2str(restartVPop.gof),' in ',mfilename,'.'])
                        disp(['Current restart iteration ', num2str(restartCounter),'.'])
                    end
                    restartCounter = restartCounter + 1;
                    if ((restartVPop.gof >= newVPop.gof) && (restartVPopEffN >= curEffN))
                        newVPop = restartVPop;
                        curVPopEffN = restartVPopEffN;
                        if (newVPop.gof > bestPVal)
                            % We also do this during the restart loop so we can
                            % track progress
                            mySaveName = ['vpop_',suffix,'_iter',num2str(wsIterCounter),'_effN',num2str(curEffN),'_curBest'];
                            saveVPop(newVPop, mySaveName);
                            bestPVal = newVPop.gof;   
                        end
                    end
                end
            end
            if ((newVPop.gof > bestPVal) && (curVPopEffN >= ...
                                             curEffN))
                % This handles the case where we found a better
                % VPop than we had but it didn't trigger a restart
                mySaveName = ['vpop_',suffix,'_iter',num2str(wsIterCounter),'_effN',num2str(curEffN),'_curBest'];
                saveVPop(newVPop, mySaveName);
                bestPVal = newVPop.gof;
            end
            % If we have surpassed nTries VPop iterations without success 
            % we will try to adjust the VPs
            
            if (mod(myTestCounter,nTries) == 0) && (nVPopsFound > 0) && ~((newVPop.gof > minPVal) && (1/sum(newVPop.pws.^2) >= curEffN))
                % We will try adding VPs based on the last, best
                % VPop if the effN is not much bigger than the basis
                if curEffN <= (basisEffN + 2*effNDelta)
                    if ((expandCohortSize > 0) && (maxNewPerIter > 0))
                        % There should be no -1 intseeds.
                        rng(newVPop.intSeed, 'twister');
                        [myWorksheet, newPassNames] = expandWorksheetVPsFromVPop(myWorksheet,oldVPop, myMapelOptions,suffix,wsIterCounter, maxNewPerIter, myScreenTable, expandCohortSize, myExpandVPopEffNOptions.varyMethod, myExpandVPopEffNOptions.resampleStd, myExpandVPopEffNOptions.maxNewPerOld, myExpandVPopEffNOptions.expandEdgeVPs, myExpandVPopEffNOptions.selectByParent, myExpandVPopEffNOptions.screenFunctionName);
                        % We force myTestCounter to 0
                        % so we can generate a new
                        % initial VPop spreading
                        % PWs from parents to children.
                        myTestCounter = 0;
                        bestPVal = -1;
                        wsIterCounter = wsIterCounter + 1;                       
                        if verbose
                            disp(['Unable to find an acceptable VPop with initial worksheet in ',num2str(nTries),' VPop fit restarts, added ', num2str(length(newPassNames)), ' VPs to the worksheet in ',mfilename,' to start worksheet iteration ',num2str(wsIterCounter),'.'])
                        end       
                        saveWorksheetAutoSplit(myWorksheet,['myWorksheet_',suffix,'_iter',num2str(wsIterCounter)]);                         
                    end    
                else
                    % Otherwise, if we've increased the effN a bit,
                    % we will try clustering the worksheet
                    % and continuing from a lower effN
                    if ((expandCohortSize > 0) && (maxNewPerIter > 0))
                        % Decide now far to "reduce" the worksheet
                        reduceFactor = 0;
                        nSteps = floor(((curEffN - basisEffN) / effNDelta)*reduceFactor);
                        curEffN = basisEffN + nSteps * effNDelta; 
						nVPTarget = basisNVPs + nSteps * maxNewPerIter;
						if nVPTarget <= 0
							nVPTarget = startNVPs;
							curEffN = startEffN;
						end
                        % Cluster on worksheet axes, data being calibrated
                        % and keeping the edge VPs
                        myClusterPickOptions = clusterPickOptions;       
                        myClusterPickOptions = myClusterPickOptions.setDefaultFromWorksheet(myWorksheet);
                        myClusterPickOptions = myClusterPickOptions.setClusterElementFromOptions(myMapelOptions);  
                        myClusterPickOptions.intSeed = intSeedTarget;
                        myClusterPickOptions.edgeVPFlag = true;
                        myClusterPickOptions.nClusters = nVPTarget;
                        myClusterPickOptions.algorithm = 'auto';
                        myClusterPickOptions.distance = 'correlation';
                        myMedoidResult = pickClusterVPs(myWorksheet,myClusterPickOptions);
                        myVPIDs = myMedoidResult.('pickedVPIDs');
                        [myWorksheet, oldVPop] = ...
                            subsetWorksheetVPop(myWorksheet, newVPop, myVPIDs, false);
                        % We also need to update the  current
                        % effN, pValue
                        bestPVal = -1;
                        myMapelOptions.minEffN = curEffN;
                        wsIterCounter = wsIterCounter + 1;
                        % We reset the testCounter so
                        % we can force trying to use the current
                        % VPop PW guess from the subset
                        myTestCounter = 0;
                        if verbose
                            disp(['Unable to find an acceptable VPop with initial worksheet in ',num2str(nTries),' VPop fit restarts, clustered to ', num2str(nVPTarget), ' VPs to the worksheet in ',mfilename,' to start worksheet iteration ',num2str(wsIterCounter),' with effN ',num2str(curEffN),'.'])
                        end                            
                        saveWorksheetAutoSplit(myWorksheet,['myWorksheet_',suffix,'_iter',num2str(wsIterCounter)]);                         
                    end
                end
            end
        end
        nVPopsFound = nVPopsFound+1;
        oldVPop = newVPop;
        % Increment target
        curEffN = curEffN + effNDelta;
        myMapelOptions.minEffN = curEffN;

        if curEffN <= targetEffN
            if expandCohortSize > 0
                % Get ready to expand VPs
                wsIterCounter = wsIterCounter+1;
                % There should be no -1 intseeds.
                rng(newVPop.intSeed, 'twister');
                [myWorksheet, newPassNames] = expandWorksheetVPsFromVPop(myWorksheet, oldVPop, myMapelOptions, suffix,wsIterCounter, maxNewPerIter, myScreenTable, expandCohortSize, myExpandVPopEffNOptions.varyMethod, myExpandVPopEffNOptions.resampleStd, myExpandVPopEffNOptions.maxNewPerOld, myExpandVPopEffNOptions.expandEdgeVPs, myExpandVPopEffNOptions.selectByParent, myExpandVPopEffNOptions.screenFunctionName);
                % Save the new worksheet that will be used.
                saveWorksheetAutoSplit(myWorksheet,['myWorksheet_',suffix,'_iter',num2str(wsIterCounter)]);   
                if verbose
                    disp(['Added ', num2str(length(newPassNames)), ' VPs to the worksheet in ',mfilename,' to start worksheet iteration ',num2str(wsIterCounter),'.'])
                end
            end
        end
    end
	% Close the pool before exiting.
	if ~isempty(gcp('nocreate'))
		delete(gcp);
	end
else
    warning(['Exiting ',mfilename, '. Returning input worksheet and hot start VPop.'])    
    newVPop = hotStartVPop;
end
end
```
