## Supplementary File 3 for "A surrogate workflow for virtual population development: designing an amenable proxy for a hypothesis-test inspired nonlinear goodness-of-fit": mapelOptionsRECISTnoBin.m

```
classdef mapelOptionsRECISTnoBin
% Options and data required to to run MAPEL
%
% expData:           (Required) Contains experimental data.  Must be
%                    populated before calling MAPEL.
% mnSDTable:         (Required) Contains mean/SD data to match.  Must be
%                    populated with experimental data before calling MAPEL,
%                    or leave unassigned if no mean/SD targets.
% binTable:          (Required) containing data on a binned pdf function to
%                    match.  Must be
%                    populated with experimental data before calling MAPEL,
%                    or leave unassigned if no bin targets.
% distTable:         (Required) containing data on a cdf function to
%                    match.  Must be
%                    populated with experimental data before calling MAPEL,
%                    or leave unassigned if no distribution targets.
% distTable2D:       (Required) containing data on a 2D pdf function to
%                    match.  Must be
%                    populated with experimental data before calling MAPEL,
%                    or leave unassigned if no 2D distribution targets.
% corTable:
% brTableRECIST:     (Required) A table with best RECIST responses.
% rTableRECIST:      (Required) A table with RECIST responses.
% initialPWs:        (Optional) 
%                     - A 1 nVP vector of initial pws. OR
%                     - (default) If set to be [], probabilities are
%                       initialized to be uniform.  The default is []. OR
%                     - if set to -1, we will try to find a near optimum starting
%                       point based on a linearized problem formulation.
% randomStart:       (Optional) If 0, the initial 
%                    parameters are not perturbed in anyway.  Otherwise the 
%                    transformed initial bin probabilities are perturbed  
%                    with random (normally distributed) noise that has
%                    normalized standard deviation of the specified value.  
%                    If no initialPWs are specified, a value of zero 
%                    results in uniform bin probabilities and a value >0 
%                    results in uniform normal initial bin probabilities  
%                    (renormalized so the margin is 1).
% nIters:            (Optional) number of iterations for the simplex 
%                    solver, ignored for other optimizeType options.  The
%                    default is 10,000.
% tol:               (Optional) Stopping tolerance for the solvers.
%                    Default is 1E-3.
% spreadOut:         (Optional) Extra argument for "findFit" function that 
%                    tries to force more VPs to be used in the solution.
%                    This is also addressed with minEffN.  Default is 0.
% minIndPVal:        Used for optimization, if nonzero, will  
%                    apply a large penalty if any of the individual pvalues.
%                    fall below the target.
% useEffN:           (Optional) Boolean (true/false). Whether to use the  
%                    total number of PWs above or effective N measure for 
%                    optimization and calculating GOF statistics.  It is 
%                    strongly recommended to set this to false during
%                    optimization.  It is set to false by
%                    default.
% exactFlag:         Whether to check to apply Fisher's exact test instead
%                    of the chi-square approximation for binned distribution
%                    test.  This may be strictly correct but can slow
%                    calculations, often with small impacts on GOF
%                    calculations.						  
% optimizeTimeLimit: (Optional) Time limit to stop the solver, does not 
%                    apply to the simplex method.  Default is 10*60 s,
%                    which will not be sufficient in many cases.
% optimizeType:      (Optional) "pso," "ga," or "simplex".  Default is
%                    "pso"
% optimizePopSize:   (Optional) only applies to "pso" and "ga".  This is
%                    the population size to use in the optimization runs.
%                    Default is 1000.
% objectiveLimit:    stopping condition for optimization
% intSeed:           (Optional) Random seed, set to -1 to avoid re-seeding
%                    the random number generator.  Default is -1.
% minEffN:           (Optional) Distinct from useEffN, this is an additional
%                    constraint placed on optimization.  Solutions that
%                    do not meet this EffN criteria are heavily penalized.
%                    The default is 0.
% relSLDvar:         a variable name that will be used for RECIST classification
% absALDVar:         a variable to indicate lesion size, used for CR cutoff
% crCutoff:          numeric value for diameter to use for CR cutoff

   properties
      expData
      mnSDTable
      binTable
      distTable
      distTable2D	  
	  corTable
      brTableRECIST
      rTableRECIST      
      initialPWs
      randomStart
      nIters
      tol
      spreadOut
	  minIndPVal	  
      useEffN    
      exactFlag			   
      optimizeTimeLimit
      optimizeType
      optimizePopSize 
	  objectiveLimit	  	  
      intSeed
      minEffN
      relSLDvar 
      absALDVar
      crCutoff
      
   end

   methods     

      function obj = set.expData(obj,myExpData)
          obj.expData = myExpData;
      end       
       
      function obj = set.mnSDTable(obj,myExpMnSdData)
          obj.mnSDTable = myExpMnSdData;
      end

      function obj = set.binTable(obj,myExpBinData)
          obj.binTable = myExpBinData;
      end
      
      function obj = set.brTableRECIST(obj,myBRTableRECIST)
          obj.brTableRECIST = myBRTableRECIST;
      end 
      
      function obj = set.rTableRECIST(obj,myRTableRECIST)
          obj.rTableRECIST = myRTableRECIST;
      end 
	  
      function obj = set.distTable(obj,myExpDistData)
          obj.distTable = myExpDistData;
      end   

      function obj = set.distTable2D(obj,myExpDistData2D)
          obj.distTable2D = myExpDistData2D;
      end   	  
      function obj = set.corTable(obj,myCorTable)
          obj.corTable = myCorTable;
      end 	 	 

      function obj = set.initialPWs(obj,myInitialPWs)
          obj.initialPWs = myInitialPWs;
      end      
      
      function obj = set.randomStart(obj,myRandomStart)
          if (myRandomStart >= 0)
               obj.randomStart = myRandomStart;
          else
               error(['Property randomStart in ',mfilename,' must be > 0.'])
          end
      end  
      
      function obj = set.nIters(obj,myNIters)
          if (myNIters > 0) && (mod(myNIters,1) == 0)
               obj.nIters = myNIters;
          else
               error(['Property nIters in ',mfilename,' must be a positive integer.'])
          end
      end            
      
      function obj = set.tol(obj,myTol)
          if (myTol > 0)
               obj.tol = myTol;
          else
               error(['Property tol in ',mfilename,' must be > 0.'])
          end
      end      
       
      function obj = set.spreadOut(obj,mySpreadOut)
          if (mySpreadOut >= 0)
               obj.spreadOut = mySpreadOut;
          else
               error(['Property spreadOut in ',mfilename,' must be >= 0.'])
          end
      end  
	  
      function obj = set.minIndPVal(obj,myMinIndPVal)
          if (myMinIndPVal >= 0) & (myMinIndPVal <= 1)
               obj.minIndPVal = myMinIndPVal;
          else
               error(['Property minIndPVal in ',mfilename,' must be between 0 and 1.'])
          end
      end 		  

      function obj = set.useEffN(obj,myUseEffN)
          if islogical(myUseEffN)
               obj.useEffN = myUseEffN;
          else
               error(['Property useEffN in ',mfilename,' must be logical.'])
          end
      end  

      function obj = set.exactFlag(obj,myFlag)
          if islogical(myFlag)
               obj.exactFlag = myFlag;
          else
               error(['Property exactFlag in ',mfilename,' must be logical.'])
          end
      end   
      
      function obj = set.optimizeTimeLimit(obj,myOptimizeTimeLimit)
          obj.optimizeTimeLimit = myOptimizeTimeLimit;
      end  
      
      function obj = set.optimizeType(obj,myOptimizeType)
          if sum(ismember({'pso','ga','simplex'},lower(myOptimizeType))) == 1
              obj.optimizeType = lower(myOptimizeType);
          else
              error(['Property optimizeType in ',mfilename,' must be "ga," "pso," or "simplex."'])
          end
      end        
      
      function obj = set.optimizePopSize(obj,myPopulationSize)
          if ((isnumeric(myPopulationSize) == true) && (myPopulationSize >= 0))
              obj.optimizePopSize = myPopulationSize;
          else
            error('Invalid optimizePopSize value')
          end
      end     
      
      function obj = set.objectiveLimit(obj,myObjectiveLimit)
          if ((isnumeric(myObjectiveLimit) == true))
              obj.objectiveLimit = myObjectiveLimit;
          else
            error('Invalid objectiveLimit value')
          end
      end 	  	  
	  
      function obj = set.intSeed(obj,myIntSeed)
          if ((mod(myIntSeed,1) == 0) && (myIntSeed>=-1))
              obj.intSeed = (myIntSeed);
          else
              error(['Invalid intSeed specified for ',mfilename,', a non-negative integer should be specified, or -1 to ignore.'])
          end
      end
      
      function obj = set.minEffN(obj,myMinEffN)
          if ((isnumeric(myMinEffN) == true) && (myMinEffN >= 0))
              obj.minEffN = myMinEffN;
          else
              error('Invalid minEffN value')
          end
      end       
      
      function obj = set.relSLDvar(obj,myRelSLDvar)
          obj.relSLDvar = myRelSLDvar;
      end       
      
      function obj = set.absALDVar(obj,myAbsALDVar)
          obj.absALDVar = myAbsALDVar;
      end     
      
      function obj = set.crCutoff(obj,myCRCutoff)
          obj.crCutoff = myCRCutoff;
      end           
      
      
      function value = get(obj,propName)
          switch propName
              case 'brTableRECIST'
                  value = obj.brTableRECIST;            
              case 'expData'
                  value = obj.expData;
              case 'mnSDTable'
                  value = obj.mnSDTable;
              case 'binTable'
                  value = obj.binTable;                  
              case 'distTable'
                  value = obj.distTable; 	
              case 'distTable2D'
                  value = obj.distTable2D; 				  
              case 'corTable'
                  value = obj.corTable;					 
              case 'brTableRECIST'
                  value = obj.brTableRECIST; 
              case 'rTableRECIST'
                  value = obj.rTableRECIST;                   
              case 'initialPWs'
                  value = obj.initialPWs;
              case 'randomStart'
                  value = obj.randomStart;
              case 'nIters'
                  value = obj.nIters;
              case 'tol'
                  value = obj.tol;
              case 'spreadOut'
                  value = obj.spreadOut;
              case 'minIndPVal'
                  value = obj.minIndPVal;				  
              case 'useEffN'
                  value = obj.useEffN;
              case 'exactFlag'
                  value = obj.exactFlag;   	  
              case 'optimizeTimeLimit'
                  value = obj.optimizeTimeLimit;
              case 'optimizeType'
                  value = obj.optimizeType;  
              case 'optimizePopSize'
                  value = obj.optimizePopSize;
              case 'objectiveLimit'
                  value = obj.objectiveLimit;					  
              case 'intSeed'
                  value = obj.intSeed;   
              case 'minEffN'
                  value = obj.minEffN; 
              case 'relSLDvar'
                  value = obj.relSLDvar; 
              case 'absALDVar'
                  value = obj.absALDVar; 
              case 'crCutoff'
                  value = obj.crCutoff;                   
              otherwise
                  error(['Error: ',propName ,' is not a valid ',mfilename,' property.'])
          end
      end

      % TODO: ADD METHODS TO VERIFY AGAINST WORKSHEET
            
      function obj = mapelOptionsRECISTnoBin()
          % This is the constructor method for instances of mapelOptions
          % objects.
          %
          obj.expData = [];
          obj.mnSDTable = [];
          obj.binTable = [];
          obj.distTable = [];   
		  obj.distTable2D = []; 
		  obj.corTable = []; 					   
          obj.brTableRECIST = [];
          obj.rTableRECIST = [];          
          obj.initialPWs = [];
          obj.randomStart = 0;
          obj.nIters = 10000;
          obj.tol = 1E-3;
          obj.spreadOut = 0;
          obj.minIndPVal = 0;			  		  		  
          obj.useEffN = false;
          obj.exactFlag = true;
          obj.optimizeTimeLimit = 10*60;
          obj.optimizeType = 'pso';
          obj.optimizePopSize = 1000;
		  obj.objectiveLimit = -Inf;		  
          obj.intSeed = -1;   
          obj.minEffN = 0;     
          obj.relSLDvar = [];
          obj.absALDVar = [];
          obj.crCutoff = nan;          
      end
   end
end
```
