## Supplementary File 3 for "A surrogate workflow for virtual population development: designing an amenable proxy for a hypothesis-test inspired nonlinear goodness-of-fit": VPopRECISTnoBin.m

```
classdef VPopRECISTnoBin
% This is a virtual population class.  This contains the results if a call
% to MAPEL.  There should be too much here to that needs to
% be changed directly, with the biggest possible exception being adjusting 
% useEffN before re-evaluating the GOF.  However, there is a substantial
% amount of customization possible by resuming fits with an existing
% VPop so you may find reasons to modify the properties.
%
% This object uses RECIST-based classifications for VP responses.  This
% impacts population observations.  For example, once a VP has observed "PD"
% biomarkers for later time points will not be included in calculations since
% this VP would generally be off trial.
%
% Note that the "noBin" designation means prevalence weights are applied directly
% to VPs and the VPs are not grouped into bins based on their parameter axes.
%
% PROPERTIES:
%  coeffsTable:  (Don't manipulate) A nAxis X nVP matrix with VP the
%                coefficients
%  coeffsDist:   (Don't manipulate) A nVP X vVP matrix with VP the
%                distance for the coefficients
%  pws:          (Don't manipulate) A 1xnVP vector of prevalence weights,
%                which are calculated based on the assignPWs.
%  expData:      (Don't manipulate) A table of experimental data used to
%                guide optimization.  It is usually taken from the
%                mapelOptions.
%  simData:      (Don't manipulate) A table of simulation results that 
%                pairs to the experimental data.  Usually assigned in
%                MAPEL.
%  mnSDTable:    (Don't manipulate) Summary table of experimental and 
%                simulated (weighted) mean and standard deviation.  Usually
%                copied from the mapelOptions and updated with
%                the weighted simulation results from the VPop fitting.
%                If you want to adjust a fit and restart, you may want to 
%                adjust some of the weights here.
%  binTable:     (Don't manipulate) Summary table of experimental and  
%                simulated binned (weighted) data for a virtual population
%                Usually copied from the mapelOptions and updated with
%                the weighted simulation results from the VPop fitting.
%                If you want to adjust a fit and restart, you may want to 
%                adjust some of the weights here.
%  distTable:    (Don't manipulate) Summary table of experimental and  
%                simulated distribution data for a virtual population
%                Usually copied from the mapelOptions and updated with
%                the weighted simulation results from the VPop fitting.
%                If you want to adjust a fit and restart, you may want to 
%                adjust some of the weights here. 			
%  distTable2D:	 (Don't manipulate) A table to enable calibrating 2D distributions
%  corTable:	 (Don't manipulate) A table to enable calibrating pairwise correlations																					
%  brTableRECIST: (Don't manipulate) A table with best RECIST responses
%  rTableRECIST: (Don't manipulate) A table with RECIST responses to better 
%                calibrate the distributions at each time step
%  gofMn:        (Don't manipulate) Goodness of fit statistics for 
%                individual endpoint/time mean comparisons between data
%                and virtual population.  Usually calcualated by the
%                evaluateGOF function.
%  gofSD:        (Don't manipulate) Goodness of fit statistics for 
%                individual endpoint/time standard deviation comparisons  
%                between data and virtual population.  Usually calculated 
%                by the evaluateGOF function.
%  gofBin:       (Don't manipulate) Goodness of fit statistics for 
%                individual endpoint/time bin distribution comparisons 
%                between data and virtual population.  Usually calculated 
%                by the evaluateGOF function.
%  gofDist:      (Don't manipulate) Goodness of fit statistics for 
%                empirical distribution comparisons 
%                between data and virtual population.  Usually calculated 
%                by the evaluateGOF function.	
%  gofCor:			
%  gofBR:
%  gofR:
%  gof:          (Don't manipulate) Composite goodness-of-fit result.
%                Usually updated by a call to the evaluateGOF.
%  useEffN:      Whether or not to use the effN for optimization and
%                evaluation of the GOF.  It is generally recommended to
%                useEffN for evaluation of the final fit but not
%                during optimization.
%  exactFlag:    Whether to check to apply Fisher's exact test instead
%                of the chi-square approximation for binned distribution
%                test.  This may be strictly correct but can slow
%                calculations, often with small impacts on GOF
%                calculations.																	  	  
%  spreadOut:    Used for optimization, if nonzero, will  
%                apply a penalty proportional to spreadOut to increase how
%                the prevalence weight is distributed.
%  minIndPVal:   Used for optimization, if nonzero, will  
%                apply a large penalty if any of the individual pvalues.
%                fall below the target.
%  optimizeTimeLimit:   Time limit for optimizing the VPop in s.
%  optimizeType:        Type of optimization algorithm to employ: "pso,"
%                       "ga," or "simplex".  Default is
%                       "pso".
%  optimizePopSize:     Number of solutions to try in each optimization
%                       generation.  Only applies to "pso" and "ga".  This 
%                       is the population size to use in the optimization 
%                       runs. Default is 1000.
%  objectiveLimit:		stopping condition for optimization
%  intSeed:             A non-negative integer seed to initialize the 
%                       random number generator.  Set to -1 to avoid
%                       changing the state of the random number generator.
%  tol:                 Numerical tolerance for optimization.
%  nIters:              Maximum number of iterations for fminsearch.
%  minEffN:             Minimum effective N.  A large penalty is applied
%                       if the effN drops below this N during optimization,
%                       better ensuring solutions that weight multiple VPs.
%                       The minEffN setting operates independently of 
%                       useEffN, and often works better than modifying 
%                       spreadOut.  The default is 0.
% relSLDvar:         a variable name that will be used for RECIST classification
% absALDVar:         a variable to indicate lesion size, used for CR cutoff
% crCutoff:          numeric value for diameter to use for CR cutoff
% recistSimFilter:   A structure with data on which simulated patients are
%                    on therapy

properties
    coeffsTable
	coeffsDist
    pws
    expData
    simData      
    % We combine simulation and experimental results
    % into the same table.
    mnSDTable
    binTable
    distTable
    distTable2D  
    corTable
    brTableRECIST
    rTableRECIST    
    gofMn
    gofSD
    gofBin
    gofDist	
    gofDist2D    
	gofCor	   
    gofBR    
    gofR    
    gof
    useEffN  
    exactFlag			 
    spreadOut   
	minIndPVal
    optimizeTimeLimit
    optimizeType
    optimizePopSize
	objectiveLimit	
    intSeed  
    tol
    nIters
    minEffN
    relSLDvar 
    absALDVar
    crCutoff    
    recistSimFilter    
end

methods
      function obj = set.coeffsTable(obj,myCoeffsTable)
          obj.coeffsTable = myCoeffsTable;
      end
      function obj = set.coeffsDist(obj,myCoeffsDist)
          obj.coeffsDist = myCoeffsDist;
      end	  
      function obj = set.pws(obj,myPWs)
          obj.pws = myPWs;
      end

      function obj = set.mnSDTable(obj,myMnSDTable)
          obj.mnSDTable = myMnSDTable;
      end

      function obj = set.binTable(obj,myBinTable)
          obj.binTable = myBinTable;
      end

      function obj = set.distTable(obj,myDistTable)
          obj.distTable = myDistTable;
      end    
      
      function obj = set.distTable2D(obj,myDistTable)
          obj.distTable2D = myDistTable;
      end          
      function obj = set.corTable(obj,myCorTable)
          obj.corTable = myCorTable;
      end			
      function obj = set.brTableRECIST(obj,myBRTableRECIST)
          obj.brTableRECIST = myBRTableRECIST;
      end 
      
      function obj = set.rTableRECIST(obj,myRTableRECIST)
          obj.rTableRECIST = myRTableRECIST;
      end 
      
      function obj = set.expData(obj,myExpData)
          obj.expData = myExpData;
      end

      function obj = set.simData(obj,mySimData)
          obj.simData = mySimData;
      end

      function obj = set.gofMn(obj,myGOF)
          obj.gofMn = myGOF;
      end

      function obj = set.gofSD(obj,myGOF)
          obj.gofSD = myGOF;
      end
      
      function obj = set.gofBin(obj,myGOF)
          obj.gofBin = myGOF;
      end     

      function obj = set.gofDist(obj,myGOF)
          obj.gofDist = myGOF;
      end   
      
      function obj = set.gofDist2D(obj,myGOF)
          obj.gofDist2D = myGOF;
      end               

      function obj = set.gofCor(obj,myGOF)
          obj.gofCor = myGOF;
      end 
      function obj = set.gofBR(obj,myGOF)
          obj.gofBR = myGOF;
      end      
      
      function obj = set.gofR(obj,myGOF)
          obj.gofR = myGOF;
      end           
      
      function obj = set.gof(obj,myGOF)
          obj.gof = myGOF;
      end          
      
      function obj = set.useEffN(obj,myUseEffN)
          if islogical(myUseEffN)
               obj.useEffN = myUseEffN;
          else
               error(['Property useEffN in ',mfilename,' must be logical.'])
          end
      end  
      
      function obj = set.exactFlag(obj,myFlag)
          obj.exactFlag = myFlag;
      end         
      
      function obj = set.spreadOut(obj,mySpreadOut)
          if (mySpreadOut >= 0)
               obj.spreadOut = mySpreadOut;
          else
               error(['Property spreadOut in ',mfilename,' must be >= 0.'])
          end
      end    
	  
      function obj = set.minIndPVal(obj,myMinIndPVal)
          if (myMinIndPVal >= 0) & (myMinIndPVal <= 1)
               obj.minIndPVal = myMinIndPVal;
          else
               error(['Property minIndPVal in ',mfilename,' must be between 0 and 1.'])
          end
      end 	 	  
      
      function obj = set.optimizeTimeLimit(obj,myOptimizeTimeLimit)
          obj.optimizeTimeLimit = myOptimizeTimeLimit;
      end  
      
      function obj = set.optimizeType(obj,myOptimizeType)
          if sum(ismember({'pso','ga','simplex'},lower(myOptimizeType))) == 1
              obj.optimizeType = lower(myOptimizeType);
          else
              error(['Property optimizeType in ',mfilename,' must be "ga," "pso," or "simplex"'])
          end
      end          
      
      function obj = set.optimizePopSize(obj,myPopulationSize)
          if ((isnumeric(myPopulationSize) == true) && (myPopulationSize >= 0))
              obj.optimizePopSize = myPopulationSize;
          else
            error('Invalid optimizePopSize value')
          end
      end          

      function obj = set.objectiveLimit(obj,myObjectiveLimit)
          if ((isnumeric(myObjectiveLimit) == true))
              obj.objectiveLimit = myObjectiveLimit;
          else
            error('Invalid objectiveLimit value')
          end
      end 	 	  
      
      function obj = set.intSeed(obj,myIntSeed)
          if ((mod(myIntSeed,1) == 0) && (myIntSeed>=-1))
              obj.intSeed = (myIntSeed);
          else
              error(['Invalid intSeed specified for ',mfilename,', a non-negative integer should be specified, or -1 to ignore.'])
          end
      end      
      
      function obj = set.tol(obj,myTol)
          if (myTol > 0)
               obj.tol = myTol;
          else
               error(['Property tol in ',mfilename,' must be > 0.'])
          end
      end       
      
      function obj = set.nIters(obj,myNIters)
          if (myNIters > 0) && (mod(myNIters,1) == 0)
               obj.nIters = myNIters;
          else
               error(['Property nIters in ',mfilename,' must be a positive integer.'])
          end
      end         
      
      function obj = set.minEffN(obj,myMinEffN)
          if ((isnumeric(myMinEffN) == true) && (myMinEffN >= 0))
              obj.minEffN = myMinEffN;
          else
            error('Invalid minEffN value')
          end
      end       
      
      function obj = set.relSLDvar(obj,myRelSLDvar)
          obj.relSLDvar = myRelSLDvar;
      end       
      
      function obj = set.absALDVar(obj,myAbsALDVar)
          obj.absALDVar = myAbsALDVar;
      end     
      
      function obj = set.crCutoff(obj,myCRCutoff)
          obj.crCutoff = myCRCutoff;
      end       
      
      function obj = set.recistSimFilter(obj,myRECISTSimFilter)
          obj.recistSimFilter = myRECISTSimFilter;
      end
      
      function value = get(obj,propName)
          switch propName
              case 'coeffsTable'
                  value = obj.coeffsTable;
              case 'coeffsDist'
                  value = obj.coeffsDist;				  
              case 'pws'
                  value = obj.pws; 
              case 'mnSDTable'
                  value = obj.mnSDTable; 
              case 'binTable'
                  value = obj.binTable; 
              case 'distTable'
                  value = obj.distTable; 
              case 'distTable2D'
                  value = obj.distTable2D; 				  
              case 'corTable'
                  value = obj.corTable;								 											 
              case 'brTableRECIST'
                  value = obj.brTableRECIST;   
              case 'rTableRECIST'
                  value = obj.rTableRECIST;   
              case 'expData'
                  value = obj.expData;                  
              case 'simData'
                  value = obj.simData;            
              case 'gofMn'
                  value = obj.gofMn;      
              case 'gofSD'
                  value = obj.gofSD;   
              case 'gofBin'
                  value = obj.gofBin;   
              case 'gofDist'
                  value = obj.gofDist;                				  
              case 'gofDist2D'
                  value = obj.gofDist2D;   
              case 'gofCor'
                  value = obj.gofCor; 
              case 'gofBR'
                  value = obj.gofBR;   
              case 'gofR'
                  value = obj.gofR;                     
              case 'gof'
                  value = obj.gof;                     
              case 'spreadOut'
                  value = obj.spreadOut;
              case 'minIndPVal'
                  value = obj.minIndPVal;					  
              case 'useEffN'
                  value = obj.useEffN;
              case 'exactFlag'
                  value = obj.exactFlag;     
              case 'optimizeTimeLimit'
                  value = obj.optimizeTimeLimit;
              case 'optimizeType'
                  value = obj.optimizeType;  
              case 'optimizePopSize'
                  value = obj.optimizePopSize;
              case 'objectiveLimit'
                  value = obj.objectiveLimit;				  
              case 'intSeed'
                  value = obj.intSeed;             
              case 'tol'
                  value = obj.tol;
              case 'nIters'
                  value = obj.nIters;       
              case 'minEffN'
                  value = obj.minEffN;  
              case 'relSLDvar'
                  value = obj.relSLDvar; 
              case 'absALDVar'
                  value = obj.absALDVar; 
              case 'crCutoff'
                  value = obj.crCutoff; 
              case 'recistSimFilter'
                  value = obj.recistSimFilter;                  
              otherwise
                  error(['Error: ',propName ,' is not a valid ',mfilename,' property.'])
          end
      end

      function obj = assignCoeffs(obj,myWorksheet)
          mycoeffsTable = getVPCoeffs(myWorksheet);
          obj.coeffsTable = mycoeffsTable;
      end
	  
      function obj = startPWs(obj, myWorksheet, myRandomStart)
          % This method initializes pws, and is used
          % prior to the optization.
          %
          % ARGUMENTS
          %  (self)
          %  myRandomStart:  A boolean variable (true/false), if true
          %                  prevalence weights will be set
          %                  randomly and if false they will be uniform
           if nargin < 1
                myRandomStart = false;
           end
          mycoeffsTable = getVPCoeffs(myWorksheet);
          [nAxis, nVP] = size(mycoeffsTable);
           if ~myRandomStart
                myUniformStartProbs = ones(1,nVP) ./ nVP;
           else
                myUniformStartProbs = rand([1, nVP]);
				myUniformStartProbsSum=sum(myUniformStartProbs);
                for axisCounter = 1 : nVP
                    myUniformStartProbs(1,axisCounter) = myUniformStartProbs(1,axisCounter)/myUniformStartProbsSum;
                end
           end
           obj.pws = myUniformStartProbs;
      end

     function obj = getSimData(obj, myWorksheet)
           % Get all of the simulation datapoints for the VPs that will
           % be needed for calculating population statistics
           %
           % ARGUMENTS
           %  (self):      Note that the following properties should be
           %               initialized (experimental data) before calling this
           %               method:
           %                mnSDTable
           %                binTable
           %                distTable
		   %                distTable2D
		   %				corTable				  
		   %                brTableRECIST
		   %                rTableRECIST           
		   %                recistSimFilter;
           %  myWorksheet: A worksheet with the simulation results.
           %
           % RETURNS
           %  (self):      The VPop object is returned, but with an updated
           %               simData property.
           %
           %
           continueFlag = true;
           mnsdDataFlag = false;
           binDataFlag = false;
           distDataFlag = false;
           distData2DFlag = false;
		   corDataFlag = false;						  
           brDataFlag = false;	
           rDataFlag = false;		              
           myMnSdData = obj.mnSDTable;
           myBinTable = obj.binTable;
		   myDistTable = obj.distTable;
		   myDistTable2D = obj.distTable2D;		   
		   myCorTable = obj.corTable;							   
           myBRTableRECIST = obj.brTableRECIST;
           myRTableRECIST = obj.rTableRECIST;           
           myRECISTFilter = obj.recistSimFilter;
           
           % We want the {expVarID, element, elementType, time} sets
           % from the experimental summaries that will be paired
           % with real data.
           rowInfoNames = {'expVarID','interventionID','elementID','elementType','time'};
		   rowInfoNames2D = {'expVarID1','expVarID2','interventionID1','interventionID2','elementID1','elementID2','elementType1','elementType2','time1','time2'};
           brInfoNames = {'expVarID','interventionID','elementID','elementType','time'};
           rInfoNames = brInfoNames;
           myDataSource = cell2table(cell(0,5), 'VariableNames', rowInfoNames);
           if ~isempty(myMnSdData)
               myDataSource = [myDataSource; myMnSdData(:,rowInfoNames)];
               mnsdDataFlag = true;
           end
           if ~isempty(myBinTable)
               myDataSource = [myDataSource; myBinTable(:,rowInfoNames)];
               binDataFlag = true;
           end           
           if ~isempty(myDistTable)
               myDataSource = [myDataSource; myDistTable(:,rowInfoNames)];
               distDataFlag = true;
           end     
           if ~isempty(myDistTable2D)
               distData2DFlag = true;
		   end
           if ~isempty(myCorTable)
               corDataFlag = true;			   
           end
           if ~isempty(myBRTableRECIST)
               brDataFlag = true;
           end     
           if ~isempty(myRTableRECIST)
               rDataFlag = true;
           end                
		   
           [nrows,ncols] = size(myDataSource);
           if nrows < 1
               warning(['No mnSDTable, binTable, or distTable assigned before calling getSimData in ',mfilename,'. The data to get is not known. Exiting.'])
               continueFlag = false;
           end
           
           % TODO: verify check for RECIST data "Source"
           
           if continueFlag
               % Eliminate duplicate rows
               myDataSource = unique(myDataSource, 'rows');
               myDataSource = sortrows(myDataSource,{'interventionID','elementID','time','elementType'},{'ascend','ascend','ascend','ascend'});

               [nEntries, ~] = size(myDataSource);

               % Info on each row of simvalues
               rowInfo = table2cell(myDataSource);
               % We will need to use a structure, like we did for results,
               % since some of the VP names will certainly be longer
               % than what is permitted by variable name lengths
               % Also, use the values
               % Row indices for the data are also added to speed subsequent calculations.

               vpIDs = getVPIDs(myWorksheet);
               nVPs = length(vpIDs);
               % Simulation results for each VP for each row
               dataValues = nan(nEntries, length(vpIDs));
               interventionIDIndex = find(ismember(rowInfoNames,'interventionID'));
               elementIDIndex = find(ismember(rowInfoNames,'elementID'));
               elementTypeIndex = find(ismember(rowInfoNames,'elementType'));
               expTimeIndex = find(ismember(rowInfoNames,'time'));
               simData.Data = dataValues;
               simData.rowInfoNames = rowInfoNames;
               simData.rowInfo = rowInfo;
               simData.vpIDs = vpIDs;
               interventionIDs = getInterventionIDs(myWorksheet);
               mnSDRows = nan(nEntries,1);
               binRows = nan(nEntries,1);
               distRows = nan(nEntries,1);
			   distRows2D = cell(nEntries,2);
			   corRows = cell(nEntries,2);								 

               for rowCounter = 1 : nEntries
                    interventionID = simData.rowInfo{rowCounter,interventionIDIndex};
                    elementID = simData.rowInfo{rowCounter,elementIDIndex};
                    elementType = simData.rowInfo{rowCounter,elementTypeIndex};
                    wshInterventionIndex = find(ismember(interventionIDs,interventionID));
                    expTime = simData.rowInfo{rowCounter,expTimeIndex};
                    simFilterValues = double(myRECISTFilter{wshInterventionIndex}.filterMatrix);
                    simFilterTimes = myRECISTFilter{wshInterventionIndex}.time;
                    simFilterValuesInf = simFilterValues;
                    simFilterValuesInf(find((simFilterValues<1)))=-1;
                    
                    % To avoid re-searching for the right rows on every
                    % iteration mapel, we provide the indices here 
                    if mnsdDataFlag
                        temp = find((ismember(myMnSdData{:,'interventionID'},interventionID)) & ((myMnSdData{:,'time'})==expTime) & (ismember(myMnSdData{:,'elementID'},elementID)) & (ismember(myMnSdData{:,'elementType'},elementType)));
                        if ~isempty(temp)
                            mnSDRows(rowCounter) = temp;
                        end
                    end
                    if binDataFlag
                        temp = find((ismember(myBinTable{:,'interventionID'},interventionID)) & ((myBinTable{:,'time'})==expTime) & (ismember(myBinTable{:,'elementID'},elementID)) & (ismember(myBinTable{:,'elementType'},elementType)));
                        if ~isempty(temp)
                            binRows(rowCounter) = temp;
                        end
                    end
                    if distDataFlag
                        temp = find((ismember(myDistTable{:,'interventionID'},interventionID)) & ((myDistTable{:,'time'})==expTime) & (ismember(myDistTable{:,'elementID'},elementID)) & (ismember(myDistTable{:,'elementType'},elementType)));
                        if ~isempty(temp)
                            distRows(rowCounter) = temp;
                        end
                    end
                    if distData2DFlag
                        % One row of source data may be involved in
                        % multiple 2D distributions
                        temp = find((ismember(myDistTable2D{:,'interventionID1'},interventionID)) & ((myDistTable2D{:,'time1'})==expTime) & (ismember(myDistTable2D{:,'elementID1'},elementID)) & (ismember(myDistTable2D{:,'elementType1'},elementType)));
                        if ~isempty(temp)
                            distRows2D{rowCounter,1} = temp;
                        end
                        temp = find((ismember(myDistTable2D{:,'interventionID2'},interventionID)) & ((myDistTable2D{:,'time2'})==expTime) & (ismember(myDistTable2D{:,'elementID2'},elementID)) & (ismember(myDistTable2D{:,'elementType2'},elementType)));
                        if ~isempty(temp)
                            distRows2D{rowCounter,2} = temp;
                        end						
                    end
                    if corDataFlag
                        % One row of source data may be involved in
                        % multiple 2D distributions
                        temp = find((ismember(myCorTable{:,'interventionID1'},interventionID)) & ((myCorTable{:,'time1'})==expTime) & (ismember(myCorTable{:,'elementID1'},elementID)) & (ismember(myCorTable{:,'elementType1'},elementType)));
                        if ~isempty(temp)
                            corRows{rowCounter,1} = temp;
                        end
                        temp = find((ismember(myCorTable{:,'interventionID2'},interventionID)) & ((myCorTable{:,'time2'})==expTime) & (ismember(myCorTable{:,'elementID2'},elementID)) & (ismember(myCorTable{:,'elementType2'},elementType)));
                        if ~isempty(temp)
                            corRows{rowCounter,2} = temp;
                        end						
                    end 
                    % I don't see how to avoid looping this, given the way
                    % the data is structured.  Luckily we just get the data
                    % once
                    for vpCounter = 1 : nVPs
                        % We need to verify the desired result for the VP is
                        % present, otherwise we report the simData result as
                        % nan
                        if length(myWorksheet.results) > 0
                            curResult = myWorksheet.results{wshInterventionIndex, vpCounter};
                            % Results should be stored in a structure, we 
                            % assume if a structure is provided then it is a 
                            % valid result
                            if strcmp(class(curResult),'struct')
                                curResult = myWorksheet.results{wshInterventionIndex, vpCounter};
                                curTimeIndex = find(ismember(curResult.Names,'time'));
                                curVarIndex = find(ismember(curResult.Names,elementID));
                                curTime = curResult.Data(:,curTimeIndex);
                                curVar = curResult.Data(:,curVarIndex);
                                % Interpolate to get the time exactly right
                                interpolateValue = interp1(curTime,curVar,expTime,'linear');
                                % Account for VPs dropping off of therapy
                                filterIndex = find(expTime <= (simFilterTimes .* simFilterValuesInf(:,vpCounter)));
                                
                                if length(filterIndex)>0
                                    filterIndex = filterIndex(1);
                                    simData.Data(rowCounter, vpCounter) = interpolateValue;
                                else
                                    simData.Data(rowCounter, vpCounter) = nan;
                                end
                            else
                                simData.Data(rowCounter, vpCounter) = nan;
                            end
                        else
                            simData.Data(rowCounter, vpCounter) = nan;
                        end
                    end
               end
               
               % BR results for each VP for each row
               myDataSource = myBRTableRECIST(:,rowInfoNames);
               myDataSource = sortrows(myDataSource,{'interventionID','elementID','time','elementType'},{'ascend','ascend','ascend','ascend'});
               
               brRowInfo = table2cell(myDataSource);
               
               [nEntries, ~] = size(myDataSource);
               dataValues = nan(nEntries, length(vpIDs));
               interventionIDIndex = find(ismember(rowInfoNames,'interventionID'));
               expTimeIndex = find(ismember(rowInfoNames,'time'));
               rData.Data = dataValues;
               
               interventionIDs = getInterventionIDs(myWorksheet);
               brRows = nan(nEntries,1);
               
               for rowCounter = 1 : nEntries
                   interventionID = brRowInfo{rowCounter,interventionIDIndex};
                   wshInterventionIndex = find(ismember(interventionIDs,interventionID));
                   expTime = brRowInfo{rowCounter,expTimeIndex};
                   temp = find((ismember(myBRTableRECIST{:,'interventionID'},interventionID)) & ((myBRTableRECIST{:,'time'})==expTime) );
                   brRows(rowCounter) = temp;
                   % To avoid re-searching for the right rows on every
                   % iteration mapel, we provide the indices here
                   
                   % I don't see how to avoid looping this, given the way
                   % the data is structured.  Luckily we just get the data
                   % once
                   for vpCounter = 1 : nVPs
                       % We need to verify the desired result for the VP is
                       % present, otherwise we report the simData result as
                       % nan
                       if length(myWorksheet.results) > 0
                           curResult = myRECISTFilter{wshInterventionIndex}.bestResp(:,vpCounter);
                           curTime = myRECISTFilter{wshInterventionIndex}.time;
                           
                           % Interpolate to get the time exactly right
                           % Might want to make this "last"
                           interpolateValue = interp1(curTime,curResult,expTime,'previous');
                           brData.Data(rowCounter, vpCounter) = interpolateValue;
                       end
                   end
               end
               
               % R results for each VP for each row
               myDataSource = myRTableRECIST(:,rowInfoNames);
               myDataSource = sortrows(myDataSource,{'interventionID','elementID','time','elementType'},{'ascend','ascend','ascend','ascend'});
               
               rRowInfo = table2cell(myDataSource);
               
               [nEntries, ~] = size(myDataSource);
               dataValues = nan(nEntries, length(vpIDs));
               interventionIDIndex = find(ismember(rowInfoNames,'interventionID'));
               expTimeIndex = find(ismember(rowInfoNames,'time'));
               rData.Data = dataValues;
               
               interventionIDs = getInterventionIDs(myWorksheet);
               rRows = nan(nEntries,1);
               
               for rowCounter = 1 : nEntries
                   interventionID = brRowInfo{rowCounter,interventionIDIndex};
                   wshInterventionIndex = find(ismember(interventionIDs,interventionID));
                   expTime = brRowInfo{rowCounter,expTimeIndex};
                   temp = find((ismember(myRTableRECIST{:,'interventionID'},interventionID)) & ((myRTableRECIST{:,'time'})==expTime) );
                   rRows(rowCounter) = temp;
                   % To avoid re-searching for the right rows on every
                   % iteration mapel, we provide the indices here
                   
                   % I don't see how to avoid looping this, given the way
                   % the data is structured.  Luckily we just get the data
                   % once
                   for vpCounter = 1 : nVPs
                       % We need to verify the desired result for the VP is
                       % present, otherwise we report the simData result as
                       % nan
                       if length(myWorksheet.results) > 0
                           curResult = myRECISTFilter{wshInterventionIndex}.curResp(:,vpCounter);
                           curTime = myRECISTFilter{wshInterventionIndex}.time;
                           
                           % Interpolate to get the time exactly right
                           % Might want to make this "last"
                           interpolateValue = interp1(curTime,curResult,expTime,'previous');
                           rData.Data(rowCounter, vpCounter) = interpolateValue;
                       end
                   end
               end               
               
               
               simData.binRows = binRows;
               simData.mnSDRows = mnSDRows;
               simData.distRows = distRows;	
               simData.distRows2D = distRows2D;
			   simData.corRows = corRows;
               simData.brRows = brRows;
               simData.brData = brData.Data;
               simData.brRowInfo = brRowInfo; 
               simData.rRows = rRows;
               simData.rData = rData.Data;
               simData.rRowInfo = rRowInfo;                
               obj.simData = simData;
           end
      end

      function obj = addDistTableSimVals(obj)   
          % Here we simply get sim values that are
          % fixed during optimization and add them to the 
          % dist table.  This is done at initialization to speed
          % execution.
          %
          % ARGUMENTS
          %  (self):      Note that the following properties should be
          %               initialized (experimental and simulation data) 
          %               before calling this method:
          %                distTable
          %                simData
          %
          % RETURNS
          %  (self):      The VPop object is returned, but with updated
          %               properties:
          %                distTable	
          

          myDistTable = obj.distTable;          		  
          
          mySimData = obj.simData.Data;
          
          mySimRowInfo = obj.simData.rowInfo;
          
          mySimColNames = obj.simData.rowInfoNames;
          distRowsTarget = obj.simData.distRows;
          distRowsSource = find(distRowsTarget>0); 		
          
          simInterventionIDCol = find(ismember(mySimColNames, 'interventionID'));
          simTimeCol = find(ismember(mySimColNames, 'time'));
          simElementIDCol = find(ismember(mySimColNames, 'elementID'));
          simElementTypeCol = find(ismember(mySimColNames, 'elementType'));                        
		  
          if ~isempty(distRowsSource)
              distRowsTarget = distRowsTarget(distRowsSource);
          
              % 2 step assignment to speed execution
              % first to matrix, then convert back to table.
              [nDistRows, nDistCols] = size(myDistTable);          
				
              curSimValues = nan(nDistRows,  size(mySimData,2));
              curSimValues(distRowsTarget,:) = (mySimData(distRowsSource, :));
              
              [curSimValues, I] = sort(curSimValues, 2, 'ascend');
              for rowCounter = 1 : nDistRows
                  
                  keepIndices = find(~isnan(curSimValues(rowCounter,:)));
                  curVals = curSimValues(rowCounter,keepIndices);                  
                  myDistTable.('predSample'){rowCounter} = curVals(keepIndices);
				  myDistTable.('predIndices'){rowCounter} = I(rowCounter,keepIndices);
                  % Also get the indices to align the exp and sim samples
                  sample1 = myDistTable.('expSample'){rowCounter};
                  [sample1Ind, sample2Ind, SC] = alignSamples(sample1, curVals);
                  myDistTable.('expCombinedIndices'){rowCounter} = sample1Ind;
                  myDistTable.('simCombinedIndices'){rowCounter} = sample2Ind;
                  myDistTable.('combinedPoints'){rowCounter} = SC;
              end
              obj.distTable = myDistTable;												
          end   

      end

      function obj = addDistTable2DSimVals(obj)   
          % Here we simply get sim values that are
          % fixed during optimization and add them to the 
          % dist table.  This is done at initialization to speed
          % execution.
          %
          % ARGUMENTS
          %  (self):      Note that the following properties should be
          %               initialized (experimental and simulation data) 
          %               before calling this method:
          %                distTable2D
          %                simData
          %
          % RETURNS
          %  (self):      The VPop object is returned, but with updated
          %               properties:
          %                distTable2D	

          myDistTable = obj.distTable2D;          		  
          
          mySimData = obj.simData.Data;
          
          mySimRowInfo = obj.simData.rowInfo;
          
          mySimColNames = obj.simData.rowInfoNames;
          distRowsTarget1 = obj.simData.distRows2D(:,1);
		  distRowsTarget2 = obj.simData.distRows2D(:,2);
          distRowsSource1 = find(~cellfun(@isempty,distRowsTarget1));
          distRowsSource2 = find(~cellfun(@isempty,distRowsTarget2));		  
          
          simInterventionIDCol = find(ismember(mySimColNames, 'interventionID'));
          simTimeCol = find(ismember(mySimColNames, 'time'));
          simElementIDCol = find(ismember(mySimColNames, 'elementID'));
          simElementTypeCol = find(ismember(mySimColNames, 'elementType'));                        
		  
          if ~isempty(distRowsSource1)
              distRowsTarget1 = distRowsTarget1(distRowsSource1);
			  distRowsTarget2 = distRowsTarget2(distRowsSource2);
              % 2 step assignment to speed execution
              % first to matrix, then convert back to table.
              [nDistRows, nDistCols] = size(myDistTable);
              curSimValues1 = nan(nDistRows,  size(mySimData,2));
              curSimValues2 = nan(nDistRows,  size(mySimData,2));
              % We need a loop for the assignment
              for target1counter = 1 :length(distRowsTarget1)
                  targetRows = distRowsTarget1{target1counter};
                  for target1repCounter = 1 :length(targetRows)
                    curSimValues1(targetRows(target1repCounter),:) = (mySimData(distRowsSource1(target1counter), :));
                  end
              end
              for target2counter = 1 :length(distRowsTarget2)
                  targetRows = distRowsTarget2{target2counter};
                  for target2repCounter = 1 :length(targetRows)
                    curSimValues2(targetRows(target2repCounter),:) = (mySimData(distRowsSource2(target2counter), :));
                  end                  
              end              		  
			  % Unlike the 1D case we won't pre-sort
              %[curSimValues1, I] = sort(curSimValues1, 2, 'ascend');
			  %curSimValues2 = curSimValues2(:,I);
              for rowCounter = 1 : nDistRows
                  keepIndices = find(~isnan(curSimValues1(rowCounter,:)) & ~isnan(curSimValues2(rowCounter,:)));
                  curVals = [curSimValues1(rowCounter,keepIndices); curSimValues2(rowCounter,keepIndices)];          
                  myDistTable.('predSample'){rowCounter} = curVals;
				  myDistTable.('predIndices'){rowCounter} = keepIndices;
                  % Also get the quadrant measures
                  sample1 = myDistTable.('expSample'){rowCounter};
				  quadrantCounts1c = quadrantCount(sample1, curVals);
				  quadrantCounts1s = quadrantCount(sample1, sample1);
				  quadrantCounts2c = quadrantCount(curVals, sample1);
				  quadrantCounts2s = quadrantCount(curVals, curVals);
                  %myDistTable.('expCombinedIndices'){rowCounter} = sample1Ind;
                  %myDistTable.('simCombinedIndices'){rowCounter} = sample2Ind;
                  myDistTable.('combinedQuadrants')(rowCounter,:) = {quadrantCounts1c,quadrantCounts1s,quadrantCounts2c,quadrantCounts2s};
              end
              obj.distTable2D = myDistTable;												
          end   

      end	  

      function obj = addCorTableSimVals(obj)   
          % Here we simply get sim values that are
          % fixed during optimization and add them to the 
          % correlation table.  This is done at initialization to speed
          % execution.
          %
          % ARGUMENTS
          %  (self):      Note that the following properties should be
          %               initialized (experimental and simulation data) 
          %               before calling this method:
          %                corTable
          %                simData
          %
          % RETURNS
          %  (self):      The VPop object is returned, but with updated
          %               properties:
          %                corTable	

          myCorTable = obj.corTable;          		  
          
          mySimData = obj.simData.Data;
          
          mySimRowInfo = obj.simData.rowInfo;
          
          mySimColNames = obj.simData.rowInfoNames;
          corRowsTarget1 = obj.simData.corRows(:,1);
		  corRowsTarget2 = obj.simData.corRows(:,2);
          corRowsSource1 = find(~cellfun(@isempty,corRowsTarget1));
          corRowsSource2 = find(~cellfun(@isempty,corRowsTarget2));		  
          
          simInterventionIDCol = find(ismember(mySimColNames, 'interventionID'));
          simTimeCol = find(ismember(mySimColNames, 'time'));
          simElementIDCol = find(ismember(mySimColNames, 'elementID'));
          simElementTypeCol = find(ismember(mySimColNames, 'elementType'));                        
		  
          if ~isempty(corRowsSource1)
              corRowsTarget1 = corRowsTarget1(corRowsSource1);
			  corRowsTarget2 = corRowsTarget2(corRowsSource2);
              % 2 step assignment to speed execution
              % first to matrix, then convert back to table.
              [nCorRows, nCorCols] = size(myCorTable);
              curSimValues1 = nan(nCorRows,  size(mySimData,2));
              curSimValues2 = nan(nCorRows,  size(mySimData,2));
              % We need a loop for the assignment
              for target1counter = 1 :length(corRowsTarget1)
                  targetRows = corRowsTarget1{target1counter};
                  for target1repCounter = 1 :length(targetRows)
                    curSimValues1(targetRows(target1repCounter),:) = (mySimData(corRowsSource1(target1counter), :));
                  end
              end
              for target2counter = 1 :length(corRowsTarget2)
                  targetRows = corRowsTarget2{target2counter};
                  for target2repCounter = 1 :length(targetRows)
                    curSimValues2(targetRows(target2repCounter),:) = (mySimData(corRowsSource2(target2counter), :));
                  end                  
              end              		  
			  % Unlike the 1D case we won't pre-sort
              %[curSimValues1, I] = sort(curSimValues1, 2, 'ascend');
			  %curSimValues2 = curSimValues2(:,I);
              for rowCounter = 1 : nCorRows
                  keepIndices = find(~isnan(curSimValues1(rowCounter,:)) & ~isnan(curSimValues2(rowCounter,:)));
                  curVals = [curSimValues1(rowCounter,keepIndices); curSimValues2(rowCounter,keepIndices)];          
                  myCorTable.('predSample'){rowCounter} = curVals;
				  myCorTable.('predIndices'){rowCounter} = keepIndices;
              end
              obj.corTable = myCorTable;												
          end   

      end	    			  
      
          function obj = addPredTableVals(obj)
          % Once we have read the simData and generated a pw vector,
          % We need to add the predicted equivalents to the 
          % tables used for stat calculations.
          %
          % ARGUMENTS
          %  (self):      Note that the following properties should be
          %               initialized (experimental and simulation data) 
          %               before calling this method:
          %                mnSDTable
          %                binTable
          %                distTable
          %                distTable2D
          %                corTable		  
		  %				   brTableRECIST		  
		  %				   rTableRECIST		
          %                simData
          %
          % RETURNS
          %  (self):      The VPop object is returned, but with updated
          %               properties:
          %                mnSDTable
          %                binTable
          %                distTable
          %                distTable2D
          %                corTable	
		  %				   brTableRECIST		  
		  %				   rTableRECIST
          myMnSdData = obj.mnSDTable;
          myBinTable = obj.binTable;
          myDistTable = obj.distTable; 
          myDistTable2D = obj.distTable2D; 		  
		  myCorTable = obj.corTable;
          myBRTableRECIST = obj.brTableRECIST;
          myRTableRECIST = obj.rTableRECIST;
          
          mySimData = obj.simData.Data;
          myBRData = obj.simData.brData;
          myRData = obj.simData.rData;
          
          mySimRowInfo = obj.simData.rowInfo;
          myBRRowInfo = obj.simData.brRowInfo;
          myBRowInfo = obj.simData.rRowInfo;
          
          myPWs = obj.pws;
          mySimColNames = obj.simData.rowInfoNames;
          mnSDRowsTarget = obj.simData.mnSDRows;
          mnSDRowsSource = find(mnSDRowsTarget>0);												
          binRowsTarget = obj.simData.binRows;
          binRowsSource = find(binRowsTarget>0);
          distRowsTarget = obj.simData.distRows;
          distRowsSource = find(distRowsTarget>0);		  
          
          brRowsTarget = obj.simData.brRows;
          brRowsSource = find(brRowsTarget>0);  
          rRowsTarget = obj.simData.rRows;
          rRowsSource = find(rRowsTarget>0);            
          
          simInterventionIDCol = find(ismember(mySimColNames, 'interventionID'));
          simTimeCol = find(ismember(mySimColNames, 'time'));
          simElementIDCol = find(ismember(mySimColNames, 'elementID'));
          simElementTypeCol = find(ismember(mySimColNames, 'elementType'));                    
          brInterventionIDCol = find(ismember(mySimColNames, 'interventionID'));
          brTimeCol = find(ismember(mySimColNames, 'time'));   
          rInterventionIDCol = find(ismember(mySimColNames, 'interventionID'));
          rTimeCol = find(ismember(mySimColNames, 'time'));          

          if ~isempty(mnSDRowsSource)
              mnSDRowsTarget = mnSDRowsTarget(mnSDRowsSource);
          
              mnSDRows = obj.simData.mnSDRows;
              mnSDRows = mnSDRows(find(mnSDRows>0));

              %  myExpMnSdData has colnames
              % {'time', 'interventionID', 'elementID', 'elementType', 'expDataID', 'expTimeVarID', 'expVarID', 'weight', 'expN', 'expMean', 'expSD', 'simN', 'simMean', 'simSD'}
              % We want to match to keep:
              [nMnSdRows, nMnSdCols] = size(myMnSdData);          

              % 2 step assignment to speed execution
              % first to matrix, then convert back to table.
              curMean = nan(nMnSdRows,1);
              curPredN = nan(nMnSdRows,1);
              curSD = nan(nMnSdRows,1);
              curSimValues = nan(nMnSdRows,length(myPWs));
              curSimValues(mnSDRowsTarget,:) = (mySimData(mnSDRowsSource, :));

              for rowCounter = 1 : nMnSdRows
                   % Note that if any simulation result values are NaN, then 
                   % these summary stats will be nan as well.  So, the default 
                   % behavior is that failed simulation runs will preclude 
                   % calculation of population statistics, which seems a 
                   % desirable failure mode.
                       
                   keepIndices = find(~isnan(curSimValues(rowCounter,:)));
                   curPWs = myPWs(keepIndices) / sum(myPWs(keepIndices));

                   if obj.useEffN
                           curN = 1/sum(curPWs.^2);
                   else
                          % We could use the PW cutoff here, but it seems this  
                          % encourages the optimizer to try to push the weight onto a 
                          % few VPs to decrease N.  Instead, let's use the number of 
                          % VPs for the purpose of statistical comparison, especially
                          % during optimization.
                          % curN = sum(myPWs >= obj.pwCutoff);
                          curN = length(myPWs);
                   end                          
                   curPredN(rowCounter) = curN;   
                   curMean(rowCounter) = wtdMean(curSimValues(rowCounter,keepIndices),curPWs);
                   curSD(rowCounter) = wtdStd(curSimValues(rowCounter,keepIndices),curPWs);
              end

              myMnSdData.('predN') = (curPredN);
              myMnSdData.('predMean') = (curMean);
              myMnSdData.('predSD') = (curSD);     
              obj.mnSDTable = myMnSdData;
          end
          if ~isempty(binRowsSource)
              binRowsTarget = binRowsTarget(binRowsSource);
          
              % 2 step assignment to speed execution
              % first to matrix, then convert back to table.
              [nBinRows, nBinCols] = size(myBinTable);          
              curProbs = cell(nBinRows,1);     
              curPredN = nan(nBinRows,1);
              curSimValues = nan(nBinRows,length(myPWs));
              curSimValues(binRowsTarget,:) = (mySimData(binRowsSource, :));
              binEdgeValues = myBinTable{:,'binEdges'};          

              for rowCounter = 1 : nBinRows

                   
                   keepIndices = find(~isnan(curSimValues(rowCounter,:)));
                   curPWs = myPWs(keepIndices) / sum(myPWs(keepIndices)); 
                   if obj.useEffN
                       curN = 1/sum(curPWs.^2);
                   else
                       % We could use the PW cutoff here, but it seems this
                       % encourages the optimizer to try to push the weight onto a
                       % few VPs to decrease N.  Also allow the number of
                       % VPs for the purpose of statistical comparison, especially
                       % during optimization.
                       % curN = sum(myPWs >= obj.pwCutoff);
                       curN = length(myPWs);
                   end
                   
                   curPredN(rowCounter) = curN;
                   curProbs{rowCounter} = wtdBinProb(curSimValues(rowCounter,keepIndices), curPWs, binEdgeValues{rowCounter});
              end
              myBinTable.('predN') = (curPredN);
              myBinTable.('predBins') = curProbs;    
              obj.binTable = myBinTable;
          end
		  
         if ~isempty(distRowsSource)
          
			  [nDistRows, nDistCols] = size(myDistTable); 
			  assignPWs = cell(nDistRows,1);
              assignN = nan(nDistRows,1);
			  keepIndices = myDistTable.('predIndices');
              for rowCounter = 1 : nDistRows
                   % We have already found these
                   curPWs = myPWs(keepIndices{rowCounter}) / sum(myPWs(keepIndices{rowCounter})); 
                   if obj.useEffN
                       curN = 1/sum(curPWs.^2);
                   else
                       % We could use the PW cutoff here, but it seems this
                       % encourages the optimizer to try to push the weight onto a
                       % few VPs to decrease N.  Also allow the number of
                       % VPs for the purpose of statistical comparison, especially
                       % during optimization.
                       % curN = sum(myPWs >= obj.pwCutoff);
                       curN = length(myPWs);
                   end

                  % Since we assign in multiple values per row for the
                  % distribution, it looks like we have to loop this
                  % All variables in the column must be same size
                  assignN(rowCounter) = curN;
				  assignPWs{rowCounter} = curPWs;
              end
              myDistTable.('predN') = (assignN);
			  myDistTable.('predProbs') = (assignPWs);
              obj.distTable = myDistTable;														
         end 		

         if ~isempty(myDistTable2D)
			  distRowsTarget1 = obj.simData.distRows2D(:,1);
			  distRowsTarget2 = obj.simData.distRows2D(:,2);
			  distRowsSource1 = find(~cellfun(@isempty,distRowsTarget1));
			  distRowsSource2 = find(~cellfun(@isempty,distRowsTarget2));		 
          
			  [nDistRows, nDistCols] = size(myDistTable2D); 
			  assignPWs = cell(nDistRows,1);
              assignN = nan(nDistRows,1);
			  keepIndices = myDistTable2D.('predIndices');
              for rowCounter = 1 : nDistRows
                   % We have already found these
                   curPWs = myPWs(keepIndices{rowCounter}) / sum(myPWs(keepIndices{rowCounter})); 
                   if obj.useEffN
                       curN = 1/sum(curPWs.^2);
                   else
                       % We could use the PW cutoff here, but it seems this
                       % encourages the optimizer to try to push the weight onto a
                       % few VPs to decrease N.  Also allow the number of
                       % VPs for the purpose of statistical comparison, especially
                       % during optimization.
                       % curN = sum(myPWs >= obj.pwCutoff);
                       curN = length(myPWs);
                   end

                  % Since we assign in multiple values per row for the
                  % distribution, it looks like we have to loop this
                  % All variables in the column must be same size
                  assignN(rowCounter) = curN;
				  assignPWs{rowCounter} = curPWs;
              end
              myDistTable2D.('predN') = (assignN);
			  myDistTable2D.('predProbs') = (assignPWs);
              obj.distTable2D = myDistTable2D;														
         end 		  		  
         if ~isempty(myCorTable)
			  corRowsTarget1 = obj.simData.corRows(:,1);
			  corRowsTarget2 = obj.simData.corRows(:,2);
			  corRowsSource1 = find(~cellfun(@isempty,corRowsTarget1));
			  corRowsSource2 = find(~cellfun(@isempty,corRowsTarget2));		 
			  [nCorRows, nCorCols] = size(myCorTable); 
			  assignPWs = cell(nCorRows,1);
              assignN = nan(nCorRows,1);
			  curCor = nan(nCorRows,1);
			  keepIndices = myCorTable.('predIndices');
              for rowCounter = 1 : nCorRows
                   % We have already found these
                   curPWs = myPWs(keepIndices{rowCounter}) / sum(myPWs(keepIndices{rowCounter})); 
                   if obj.useEffN
                       curN = 1/sum(curPWs.^2);
                   else
                       % We could use the PW cutoff here, but it seems this
                       % encourages the optimizer to try to push the weight onto a
                       % few VPs to decrease N.  Also allow the number of
                       % VPs for the purpose of statistical comparison, especially
                       % during optimization.
                       % curN = sum(myPWs >= obj.pwCutoff);
                       curN = length(myPWs);
                   end

                  % Since we assign in multiple values per row for the
                  % distribution, it looks like we have to loop this
                  % All variables in the column must be same size
                  assignN(rowCounter) = curN;
				  assignPWs{rowCounter} = curPWs;
				  curwtdcorr = weightedcorrs(myCorTable.('predSample'){rowCounter}', curPWs');
				  curCor(rowCounter) = curwtdcorr(1,2);
              end
              myCorTable.('predN') = (assignN);
			  myCorTable.('predProbs') = (assignPWs);
			  myCorTable.('predCor') = (curCor);
              obj.corTable = myCorTable;								
			  
          end 			   
		  
          if ~isempty(brRowsSource)
              brRowsTarget = brRowsTarget(brRowsSource);      
              [nBRRows, nBinCols] = size(myBRTableRECIST);          
              curProbs = nan(nBRRows,4);     
              curPredN = nan(nBRRows,1);
              curSimValues = nan(nBRRows,length(myPWs));
              curSimValues(brRowsTarget,:) = (myBRData(brRowsSource, :));
              binEdgeValues = [.9,1.9,2.9];     
              
              if obj.useEffN
                  curN = 1/sum(myPWs.^2);
              else
                  % We could use the PW cutoff here, but it seems this
                  % encourages the optimizer to try to push the weight onto a
                  % few VPs to decrease N.  Instead, let's use the number of
                  % VPs for the purpose of statistical comparison, especially
                  % during optimization.
                  % curN = sum(myPWs >= obj.pwCutoff);
                  curN = length(myPWs);
              end

              for rowCounter = 1 : nBRRows
                   % 'binEdge1','binEdge2','binEdge3'
                   curPredN(rowCounter) = curN;
                   curProbs(rowCounter,:) = wtdBinProb(curSimValues(rowCounter,:), myPWs, binEdgeValues);
              end
              myBRTableRECIST.('predN') = (curPredN);
              myBRTableRECIST.('predCR') = (curProbs(:,1));
              myBRTableRECIST.('predPR') = (curProbs(:,2));
              myBRTableRECIST.('predSD') = (curProbs(:,3));
              myBRTableRECIST.('predPD') = (curProbs(:,4));     
              obj.brTableRECIST = myBRTableRECIST;
          end          

          if ~isempty(rRowsSource)
              rRowsTarget = rRowsTarget(rRowsSource);      
              [nRRows, nBinCols] = size(myRTableRECIST);          
              curProbs = nan(nRRows,4);     
              curPredN = nan(nRRows,1);
              curSimValues = nan(nRRows,length(myPWs));
              curSimValues(rRowsTarget,:) = (myRData(rRowsSource, :));
              binEdgeValues = [.9,1.9,2.9];     
                            
              if obj.useEffN
                  curN = 1/sum(myPWs.^2);
              else
                  % We could use the PW cutoff here, but it seems this
                  % encourages the optimizer to try to push the weight onto a
                  % few VPs to decrease N.  Instead, let's use the number of
                  % VPs for the purpose of statistical comparison, especially
                  % during optimization.
                  % curN = sum(myPWs >= obj.pwCutoff);
                  curN = length(myPWs);
              end
              
              for rowCounter = 1 : nRRows
                  
                   curPredN(rowCounter) = curN;
                   %curProbs(rowCounter,:) = wtdBinProb(curSimValues(rowCounter,keepIndices), curPWs, binEdgeValues);
                   curProbs(rowCounter,:) = wtdBinProb(curSimValues(rowCounter,:), myPWs, binEdgeValues);
              end
              myRTableRECIST.('predN') = (curPredN);
              myRTableRECIST.('predCR') = (curProbs(:,1));
              myRTableRECIST.('predPR') = (curProbs(:,2));
              myRTableRECIST.('predSD') = (curProbs(:,3));
              myRTableRECIST.('predPD') = (curProbs(:,4));     
              obj.rTableRECIST = myRTableRECIST;
          end              
          
      end

      function obj = VPopRECISTnoBin()
          % This is the constructor method for an instance of a VPop
          % (virtual population) object.
          obj.coeffsTable=[];
		  obj.coeffsDist=[];		  
          obj.pws = [];
          obj.mnSDTable = [];
          obj.binTable = [];
          obj.distTable = [];  
          obj.distTable2D = [];		  
		  obj.corTable = [];					  
          obj.brTableRECIST = [];
          obj.rTableRECIST = [];          
          obj.expData = [];          
          obj.simData = [];
          obj.gofMn = [];
          obj.gofSD = [];
          obj.gofBin = [];
          obj.gofDist = [];      
		  obj.gofDist2D = [];   
		  obj.gofCor = [];					
          obj.gofBR = [];  
          obj.gofR = [];                              
          obj.gof = [];          
          obj.spreadOut = 0;
          obj.minIndPVal = 0;	
          obj.useEffN = false;
          obj.exactFlag = true; 		  
          obj.optimizeTimeLimit = 10*60;
          obj.optimizeType = 'pso';    
          obj.optimizePopSize = 1000;   
		  obj.objectiveLimit = -Inf;
          obj.intSeed = -1;
          obj.tol = 1E-3;
          obj.nIters = 10000;
          obj.minEffN = 0;
          obj.relSLDvar = [];
          obj.absALDVar = [];
          obj.crCutoff = nan;            
          obj.recistSimFilter = [];           
      end

end

end
```
