## Supplementary File 3 for "A surrogate workflow for virtual population development: designing an amenable proxy for a hypothesis-test inspired nonlinear goodness-of-fit": updateBinTableFormat.m

```
function myUpdatedObject = updateBinTableFormat(myInputObject)
% This is a function to update the bin tables from the old
% format to the new one with allowed variable bin numbers.
% The transition was made around in-house rev1043.
%
% ARGUMENTS: 
%  myInputObject:     A VPop, VPopRECIST, VPopRECISTnoBin, 
%                      mapelOptions, mapelOptionsRECIST, or 
%                      mapelOptionsRECISTnoBin object to convert.
%
% RETURNS:
%  myUpdatedObject    A VPop, VPopRECIST, VPopRECISTnoBin, 
%                      mapelOptions, mapelOptionsRECIST, or 
%                      mapelOptionsRECISTnoBin object with
%                      the new binTable.


continueFlag = true;
if nargin > 1
    continueFlag = false;
    warning(['Too many input arguments for ',mfilename,'. Should provide: a VPop, VPopRECIST, VPopRECISTnoBin, mapelOptions, mapelOptionsRECIST, or mapelOptionsRECISTnoBin.'])
    continueFlag = false;	
elseif nargin > 0
    continueFlag = true;
else
    warning(['Insufficient input arguments for ',mfilename,'. Should provide: a VPop, VPopRECIST, VPopRECISTnoBin, mapelOptions, mapelOptionsRECIST, or mapelOptionsRECISTnoBin.'])
    continueFlag = false;
end

if ~(isa(myInputObject,'VPop') || isa(myInputObject,'VPopRECIST') || isa(myInputObject,'VPopRECISTnoBin') || isa(myInputObject,'mapelOptions') || isa(myInputObject,'mapelOptionsRECIST') || isa(myInputObject,'mapelOptionsRECISTnoBin'))
	warning(['Wrong input arguments for ',mfilename,'. Should provide: a VPop, VPopRECIST, VPopRECISTnoBin, mapelOptions, mapelOptionsRECIST, or mapelOptionsRECISTnoBin.'])
	continueFlag = false;
end

if continueFlag
	myUpdatedObject = myInputObject;
	if ~isempty(myInputObject.binTable)
		[nRows, nCols] = size(myInputObject.binTable);
		if (isa(myInputObject,'VPop') || isa(myInputObject,'mapelOptions'))
			referenceCol = 9;
		else
			referenceCol = 12;
		end
		myUpdateTable = myInputObject.binTable(:,1:referenceCol);
		binEdges = cell(nRows,1);
		expBins = cell(nRows,1);
		predBins = cell(nRows,1);
		for rowCounter = 1 : nRows
			curEdges = [myInputObject.binTable{rowCounter,'binEdge1'},myInputObject.binTable{rowCounter,'binEdge2'},myInputObject.binTable{rowCounter,'binEdge3'}];
			curExpBins = [myInputObject.binTable{rowCounter,'expBin1'}, myInputObject.binTable{rowCounter,'expBin2'}, myInputObject.binTable{rowCounter,'expBin3'}, myInputObject.binTable{rowCounter,'expBin4'}];
            curPredBins = [myInputObject.binTable{rowCounter,'predBin1'}, myInputObject.binTable{rowCounter,'predBin2'}, myInputObject.binTable{rowCounter,'predBin3'}, myInputObject.binTable{rowCounter,'predBin4'}];
		    binEdges{rowCounter} = curEdges;
		    expBins{rowCounter} = curExpBins;			
		    predBins{rowCounter} = curPredBins;			
		end
		myUpdateTable = [myInputObject.binTable(:,1:referenceCol), table(binEdges), myInputObject.binTable(:,'expN'), table(expBins), myInputObject.binTable(:,'predN'), table(predBins)];
		myUpdatedObject.binTable = myUpdateTable;
    end
end
end
```
