## Supplementary File 3 for "A surrogate workflow for virtual population development: designing an amenable proxy for a hypothesis-test inspired nonlinear goodness-of-fit": updateTablesWithSubpopformat.m

```
function myUpdatedObject = updateTablesWithSubpopformat(myInputObject)
% This is a function to update the tables attached to mapelOptions
% and VPop objects to convert to a new format of subpoptable.  
% The transition was made around in-house rev1228???.
%
% ARGUMENTS: 
%  myInputObject:     A VPop, VPopRECIST, VPopRECISTnoBin, 
%                      mapelOptions, mapelOptionsRECIST, or 
%                      mapelOptionsRECISTnoBin object to convert.
%
% RETURNS:
%  myUpdatedObject    A VPop, VPopRECIST, VPopRECISTnoBin, 
%                      mapelOptions, mapelOptionsRECIST, or 
%                      mapelOptionsRECISTnoBin object with
%                      the tables.

continueFlag = true;
if nargin > 1
    continueFlag = false;
    warning(['Too many input arguments for ',mfilename,'. Should provide: a VPop, VPopRECIST, mapelOptions, or mapelOptionsRECIST.'])
    continueFlag = false;	
elseif nargin > 0
    continueFlag = true;
else
    warning(['Insufficient input arguments for ',mfilename,'. Should provide: a VPop, VPopRECIST, mapelOptions, or mapelOptionsRECIST.'])
    continueFlag = false;
end

if ~(isa(myInputObject,'VPop') || isa(myInputObject,'VPopRECIST') || isa(myInputObject,'mapelOptions') || isa(myInputObject,'mapelOptionsRECIST') )
	warning(['Wrong input arguments for ',mfilename,'. Should provide: a VPop, VPopRECIST, mapelOptions, mapelOptionsRECIST.'])
	continueFlag = false;
end

myUpdatedObject = myInputObject;

if continueFlag
    updateFlag = false;
%     if isa(myInputObject, 'VPopRECIST') || isa(myInputObject, 'mapelOptionsRECIST')
%         updateTables = {'expData','mnSDTable','binTable','distTable','distTable2D','corTable','brTableRECIST','rTableRECIST'};
%     elseif isa(myInputObject, 'VPop') || isa(myInputObject, 'mapelOptions')
%         updateTables = {'expData','mnSDTable','binTable','distTable','distTable2D','corTable'};
%     end
    updateTables = myInputObject.subpopTable;
    [nRow, nCol] = size(updateTables);
    colnames = updateTables.Properties.VariableNames;
    curComparator = updateTables.comparator; %% this is to check if the format spells comparators out
    nComparator = size(curComparator,2);
    
    updateComparator = cell(nRow,1);
   for rowi = 2:nRow
       curRowcurCol = curComparator(rowi,:);
       for coli = 1:nComparator
            if(strcmp(' ',curRowcurCol{coli}))
                 curRowcurCol(:,coli) = [];
            end
       end
       updateComparator{rowi} = curRowcurCol;
   end
                        
    for colCounter = 1:length(colnames)
        if ismember(colnames{colCounter},{'time','interventionID','elementID','elementType','comparator','value'})
            curCol = updateTables.(colnames{colCounter}); 
            updateCol = cell(nRow,1);
            ncurCol = size(curCol,2);
            if isequal(colnames{colCounter},'time') || isequal(colnames{colCounter},'value')
                if isnumeric(curCol)
                    updateCol{1} = num2cell(curCol(1));
                    updateFlag = true;
                elseif iscell(curCol) & ncurCol>1
                    updateCol{1} = curCol(1,1);
                    updateFlag = true;
                end        
            end
            if nRow>1 
                if isequal(colnames{colCounter},'time') || isequal(colnames{colCounter},'value')
                    if nComparator==1
                        for rowi = 2:nRow
                            updateCol{rowi} = num2cell(curCol(rowi));
                        end
                    elseif nComparator>1  %% this is to make it compatible with Lu's local subpopluation format
                            for rowi = 2:nRow
                                curRowcurCol = curCol(rowi,:);
                                for coli = 1:ncurCol
                                    if(isnan(curRowcurCol{coli}) || strcmp(' ',curRowcurCol{coli}))
                                        curRowcurCol(:,coli) = [];
                                    end
                                end
                                updateCol{rowi} = curRowcurCol;
                            end
                    end
                else
                    if nComparator==1
                        for rowi = 2:nRow
                            updateCol(rowi) = num2cell(curCol(rowi));
                        end     
                    elseif nComparator>1  %% this is to make it compatible with Lu's local subpopluation format
                        if isequal(colnames{colCounter},'interventionID')
                            for rowi = 2:nRow
                                curRowcurCol = curCol(rowi,:);
                                if(length(curRowcurCol)<length(updateComparator{rowi}))
                                    curRowcurCol = [curRowcurCol,repmat(curRowcurCol(:,end),1,length(updateComparator{rowi})-length(curRowcurCol))];
                                end
                                updateCol{rowi} = curRowcurCol;
                            end                            
                        else
                            for rowi = 2:nRow
                                curRowcurCol = curCol(rowi,:);
                                for coli = 1:ncurCol
                                    if(strcmp(' ',curRowcurCol{coli}))
                                        curRowcurCol(:,coli) = [];
                                    end
                                end
                                updateCol{rowi} = curRowcurCol;
                            end  
                        end
                    end
                end
            end
            if updateFlag == true
                updateTables.(colnames{colCounter})=updateCol;  
            end
        end
    end
    myUpdatedObject.subpopTable =  updateTables;
    if updateFlag
        warning(['Detected old subpop format and updated ',class(myInputObject),' subpop tables to the new format.  Note that you will have to manually add a subpopTable to the object, which can be created by calling createSubpopTable with the appropriate worksheet.'])
    end
else
    warning(['Unable to run ',mfilename,'.  Returning input object.'])
end
```
