## Supplementary File 3 for "A surrogate workflow for virtual population development: designing an amenable proxy for a hypothesis-test inspired nonlinear goodness-of-fit": updateTablesWithSubpopReference.m

```
function myUpdatedObject = updateTablesWithSubpopReference(myInputObject)
% This is a function to update the tables attached to mapelOptions
% and VPop objects to include a row for the subpopulation index.  
% The transition was made around in-house rev1100.
%
% ARGUMENTS: 
%  myInputObject:     A VPop, VPopRECIST, VPopRECISTnoBin, 
%                      mapelOptions, mapelOptionsRECIST, or 
%                      mapelOptionsRECISTnoBin object to convert.
%
% RETURNS:
%  myUpdatedObject    A VPop, VPopRECIST, VPopRECISTnoBin, 
%                      mapelOptions, mapelOptionsRECIST, or 
%                      mapelOptionsRECISTnoBin object with
%                      the tables.

continueFlag = true;
if nargin > 1
    continueFlag = false;
    warning(['Too many input arguments for ',mfilename,'. Should provide: a VPop, VPopRECIST, mapelOptions, or mapelOptionsRECIST.'])
    continueFlag = false;	
elseif nargin > 0
    continueFlag = true;
else
    warning(['Insufficient input arguments for ',mfilename,'. Should provide: a VPop, VPopRECIST, mapelOptions, or mapelOptionsRECIST.'])
    continueFlag = false;
end

if ~(isa(myInputObject,'VPop') || isa(myInputObject,'VPopRECIST') || isa(myInputObject,'mapelOptions') || isa(myInputObject,'mapelOptionsRECIST') )
	warning(['Wrong input arguments for ',mfilename,'. Should provide: a VPop, VPopRECIST, mapelOptions, mapelOptionsRECIST.'])
	continueFlag = false;
end

myUpdatedObject = myInputObject;

if continueFlag
    updateFlag = false;
    if isa(myInputObject, 'VPopRECIST') || isa(myInputObject, 'mapelOptionsRECIST')
        updateTables = {'expData','mnSDTable','binTable','distTable','distTable2D','corTable','brTableRECIST','rTableRECIST'};
    elseif isa(myInputObject, 'VPop') || isa(myInputObject, 'mapelOptions')
        updateTables = {'expData','mnSDTable','binTable','distTable','distTable2D','corTable'};
    end
    for tableCounter = 1:length(updateTables)
        curTable = myInputObject.(updateTables{tableCounter});
        [nRows, nCols] = size(curTable);
        if nRows > 0
            curNames = curTable.Properties.VariableNames;
            if ~isequal(curNames{1},'subpopNo')
                newCol = num2cell(ones(nRows,1));
                newCol = cell2table(newCol);
                newCol.Properties.VariableNames = {'subpopNo'};
                curTable = [newCol,curTable];
                updateFlag = true;
                myUpdatedObject.(updateTables{tableCounter}) = curTable;
            end
        end
    end
    if updateFlag
        warning(['Detected missing columns and added subpopNo columns to update the ',class(myInputObject),' tables to the new format.  Note that you will have to manually add a subpopTable to the object, which can be created by calling createSubpopTable with the appropriate worksheet.'])
    end
else
    warning(['Unable to run ',mfilename,'.  Returning input object.'])
end
```
