## Supplementary File 3 for "A surrogate workflow for virtual population development: designing an amenable proxy for a hypothesis-test inspired nonlinear goodness-of-fit": Cheng_2017_AAPSJ.pdf

*The AAPS Journal, Vol. 19, No. 4, July 2017 (© 2017)*
DOI: 10.1208/s12248-017-0100-x

*Tutorial*

*Theme: Pharmaceutical Sciences in the Era of Big Data Computing*
*Guest Editor: Xiang-Qun Xie*

### QSP Toolbox: Computational Implementation of Integrated Workflow Components for Deploying Multi-Scale Mechanistic Models

**Yougan Cheng,[1] Craig J. Thalhauser,[1] Shepard Smithline,[1] Jyotsna Pagidala,[1] Marko Miladinov,[1] Heather E. Vezina,[1] Manish Gupta,[1] Tarek A. Leil,[1] and Brian J. Schmidt[1,2]**

*Received 5 January 2017; accepted 8 May 2017; published online 24 May 2017*

***Abstract.*** Quantitative systems pharmacology (QSP) modeling has become increasingly important in pharmaceutical research and development, and is a powerful tool to gain mechanistic insights into the complex dynamics of biological systems in response to drug treatment. However, even once a suitable mathematical framework to describe the pathophysiology and mechanisms of interest is established, final model calibration and the exploration of variability can be challenging and time consuming. QSP models are often formulated as multi-scale, multi-compartment nonlinear systems of ordinary differential equations. Commonly accepted modeling strategies, workflows, and tools have promise to greatly improve the efficiency of QSP methods and improve productivity. In this paper, we present the QSP Toolbox, a set of functions, structure array conventions, and class definitions that computationally implement critical elements of QSP workflows including data integration, model calibration, and variability exploration. We present the application of the toolbox to an ordinary differential equations-based model for antibody drug conjugates. As opposed to a single stepwise reference model calibration, the toolbox also facilitates simultaneous parameter optimization and variation across multiple *in vitro*, *in vivo*, and clinical assays to more comprehensively generate alternate mechanistic hypotheses that are in quantitative agreement with available data. The toolbox also includes scripts for developing and applying virtual populations to mechanistic exploration of biomarkers and efficacy. We anticipate that the QSP Toolbox will be a useful resource that will facilitate implementation, evaluation, and sharing of new methodologies in a common framework that will greatly benefit the community.

**KEY WORDS:** quantitative systems pharmacology; ordinary differential equations; optimization; virtual patient; virtual population.

#### INTRODUCTION

QSP has been characterized as a "quantitative analysis of the dynamic interactions between drug(s) and a biological system that aims to understand the behavior of the system as a whole (1)." There are various existing QSP approaches and applications, and one common feature of QSP models is that they strive to incorporate key biological pathways from the systems of interest and the pharmacology of therapeutic interventions, aiming not only a better holistic understanding of the biology but also "optimal and translatable pharmacological pathway interventions (2)." QSP models are often multi-scale in that they characterize processes that occur at multiple scales of space and time (e.g., ligand binding vs. disease progression) and mechanistic meaning that fundamental biological processes are represented with mechanistic fidelity. This "systems" approach can better inform target selection and the decision process for advancing compounds through preclinical and clinical research (3); as such, it is becoming increasingly important in pharmaceutical research and development as a potential means of reducing attrition and improving productivity (4–7). QSP models often are developed to impact drug discovery and development, and often enable the investigation of relationships between

**Electronic supplementary material** The online version of this article (doi:10.1208/s12248-017-0100-x) contains supplementary material, which is available to authorized users.

[1] Bristol-Myers Squibb, PO Box 4000, Princeton, New Jersey 08543-4000, USA.

1550-7416/17/0400-1002/0 © 2017 The Author(s). This article is an open access publication biological pathways and observed biomarkers, efficacious dose projections, and population variability (1).

QSP modeling approaches have been categorized into statistical data-driven, logic-based, differential equations, cellular automata and agent-based, and hybrid and integrated models (8). Ordinary differential equation (ODE) modeling frameworks are commonly, but not exclusively, employed in QSP models. ODE models may be broadly applied to describe tissue, cellular, and molecular and biochemical systems, with inherent strengths and limitations that must be evaluated for a given application (8,9).

The similarities in intended applications for many QSP models suggest common conceptual workflows for how to develop and apply QSP models. Furthermore, given mathematical similarities in QSP models, computational implementations of QSP workflows are also generalizable for many applications. Common workflows and their computational implementation facilitate: (i) standardization of modeling approaches within the community; (ii) increased efficiency of model development and application; (iii) greater sharing of models between groups; and (iv) providing guidance to modelers on best practices (10,11).

Conceptual QSP workflows spanning model development to application have been proposed. They have involved an assessment of pathways to include (which may be assisted by data (12, 13)), simulation of physiologic phenotypes, modeling of populations, and evaluation of predictions (13). Rather than a workflow, Friedrich proposed a model qualification method intended to guide staged QSP modeling projects (14). Critical elements include assessing: (i) whether a model scope is appropriate for a research question and if appropriate pathways are included; (ii) whether both qualitative uncertainty, such as the impact of knowledge gaps, and quantitative uncertainty, such as parameter values, are assessed; (iii) whether the model captures both known variability in mechanisms as well as variability in potential clinical outcomes; and (iv) whether the model results are both qualitatively consistent with relevant data and matches selected quantitative test data. Recently, Gadkar *et al.* proposed a comprehensive conceptual workflow for QSP modeling, which was described in six stages as: (i) "project needs and goals;" (ii) "reviewing the biology: determining the project scope;" (iii) "representing the biology: developing the model structure;" (iv) "capturing behaviors & building confidence: calibrating 'reference' subjects;" (v) "exploring knowledge gaps and variability: alternate parameterizations;" and (vi) "supporting experimental and clinical design: refining knowledge (8)."

While prior work has been presented in the context of components of potential conceptual workflows, arguably QSP is not at a point yet where generally accepted and optimized methods for accomplishing tasks frequently required on QSP projects have been enumerated. Furthermore, such best practices even once established are not static, and workflows should continue to evolve (15). Another shortcoming is the limited availability of computational implementations of proposed QSP workflows, which would help to standardize methods and enable QSP modelers to more efficiently exchange knowledge and compare techniques. For example, in sub-specialties of systems biology and also in pharmacometrics, standardized tools and add-ons exist to facilitate conceptual workflows exactly as applied in the literature (for a few tools available for systems biology and pharmacometrics, see (16–18)). Although tools exist to facilitate QSP workflows (19), readily available packages that directly handle integration of multiple stages in proposed conceptual workflows, especially model calibration, exploration of uncertainties and variability, and application for study design would broadly facilitate efforts in QSP model application for many researchers.

We provide the QSP Toolbox, a set of functions, structure arrays, and class definitions that computationally implement critical elements of QSP workflows, and demonstrate how to use it by method of example. In the current version, the QSP Toolbox reads ODE QSP models built in MATLAB® SimBiology®. We describe the implemented computational QSP workflow, the toolbox features and organization, and demonstrate various aspects of the toolbox utilities with a QSP model of the pharmacodynamics of antibody drug conjugates (ADCs) as well as a smaller test model for demonstrating some of the most computationally demanding algorithms. The examples and models are included with the toolbox. We anticipate that this tutorial to get started using the QSP Toolbox and implement steps in QSP workflows will be broadly beneficial for many modelers that may not have utilized these approaches, and will be a useful reference resource for QSP modelers that have implemented similar methods. We also anticipate releasing the QSP Toolbox as an additional tool will help to further strengthen the communication and efficiency within the QSP community (20).

*Availability and Requirements.* The QSP Toolbox is provided as MATLAB® files, and the following MATLAB® toolboxes are required: SimBiology, Optimization, Global Optimization, Parallel Computing, and Statistics and Machine Learning. A 12-core Haswell machine (2× E5-2620V3) with 64 GB RAM or better is recommended to run the tutorial examples. A working familiarity with MATLAB® and SimBiology® is also assumed.

#### QSP Workflow

As previously discussed, stages iv–vi described by Gadkar *et al.* are essential elements in the execution of the workflow after an initial mechanistic model is built (8). Since the QSP Toolbox addresses these elements, they are elaborated upon below.

##### *Calibration*

To ensure a QSP model can capture system behaviors with sufficient fidelity, it is critical to calibrate the model to observed data under various interventions or experimental conditions. QSP models often contain large numbers of parameters and it may be computationally demanding to perform optimization of all simultaneously. Sensitivity analysis, subsystem/modular calibration, and model reduction can be used to enable model calibration and avoid simultaneous large-scale parameter estimation (8). As noted previously, which of these methods are best suited for a project, as well as their order, may vary depending on project goals and model formulation (8).

Analyses of the sensitivity of model outputs to different parameters in the model can determine which parameters should be the focus of optimization. This reduces the dimensions of the optimization problem and the associated computational demand. In addition to specifying the parameters to explore, their potential ranges should also be specified (8), which requires a systematic evaluation of available data. We refer to the combination of a parameter and the allowed range as a mechanistic axis (see Table I for a more complete description). Global sensitivity analysis methods are generally useful during model calibration as they can help establish whether it is possible to calibrate model outcomes to within desired ranges, or if model revisions are needed (8). Additionally, sensitivity analysis can help to identify key parameters to focus on during model calibration (8), and even guide the evaluation of the potential impact of structural uncertainties in model equations (14).

Subsystem/modular calibration refers to initial calibration of simple model subsystems from specified experimental data and literature. Some parameters can be directly calculated based on physiological mechanisms and literature, and some parameters can be estimated utilizing specified experimental datasets. This subsystem/modular calibration can reduce the parameter space to estimate in the integrated model considerably. For example, Cheng and Othmer modularized a signal transduction network and the submodule parameters were estimated using a combination of experimental data and steady state analysis. The integrated model was able to capture various signal transduction characteristics (21). Subsystem/submodule calibration is capable of justifying the model topology and capturing corresponding datasets, which makes subsequent integrated model calibration manageable.

Model reduction takes a different approach by simplifying the model network topology to reduce the number of model parameters. By identification of relationships among model states, the system of differential equations is transformed into one of lower order which still retains the key dynamic information. Different model reduction techniques have been proposed (8,22), including lumped methods, sensitivity analysis-based techniques, and time-scale-based approaches. The number of parameters can be reduced substantially following these techniques, therefore making their identification based on experimental data more feasible. In addition to reducing the number of parameters that need to be considered during calibration, time-scale-based methods

**Table I.** Terminology used the QSP Toolbox and this tutorial

| Term | Description |
|---|---|
| Virtual patient | A single model parameterization. Here, a virtual patient may equivalently be called a virtual subject or, in the case of the included ADC model examples, a virtual xenograft |
| Virtual patient cohort | An ensemble of unique virtual patients |
| Mechanistic axis | A set of model parameters as well as upper and lower bounds for each. A mechanistic axis generally constitutes a single model parameter and bounds, but may include multiple parameters and bounds combined and scaled together. A linear or logarithmic scale may be applied. |
| Mechanistic axis coefficients | A numerical value that indicates how far along a mechanistic axis a parameter (or parameter set) lies for a particular virtual patient. A value of zero indicates the parameter value(s) are set at the lower bound, and a value of 1 indicates the parameter value(s) are set at the higher bound. |
| Response type element | A mapping of a virtual patient characteristic to desired values and a method for evaluating the degree to which they agree. For example, this may include data in response to an intervention at a given time point and a specification of the objective function that should be used to assess agreement. A response type element would generally include a mapping of simulated outcomes to target values. However, one might also employ this strategy to help to develop VPs biased towards mechanistic characteristics (parameter values). |
| Response type | A set of response type elements and a method for combining their respective objective function evaluations. |
| Plausible virtual patient | A virtual patient that meets constraints imposed on response type element objective functions and/or a combined objective function evaluation for the response type |
| Reference virtual patient | A plausible virtual patient that has been designated as a "reference" due to meeting additional physiologic criteria, for example biomarker trends characteristic of responders to a given therapy or a mean of population characteristics |
| Prevalence weight | A weight assigned to a plausible virtual patient in order to optimize the agreement of virtual patient cohort outcomes with observed data statistics. The prevalence weights for all included plausible virtual patients sum to one. |
| Virtual population | A cohort of plausible virtual patients and one associated set of prevalence weights. |
| Intervention | A simulated experiment or trial. In the QSP Toolbox, an intervention may include a pharmacological intervention or overwriting of some virtual patient parameters in order to properly recreate the effect of the experimental or clinical condition. |
| Worksheet | A data structure in the QSP Toolbox that includes VP definition, intervention definition, data, and response types to help develop and assess virtual patients. |
| Variant type | Each variant type contains a consistent grouping of parameters that are used to help define virtual patients and interventions. A variant type can essentially be thought of as a parameter set. The type name is specified by the description before the delimiter in the variant name, "___" by default. |
| Type value set | A particular set of values for all parameters in a variant type. A type value set could also be referred to as a value set. The value set name is specified after the delimiter in the variant name, "___" by default. |

can also improve model simulation performance by replacing "fast" processes with suitable quasi-steady state approximations. As one example, Schmidt *et al.* illustrate development of a quasi-steady state approximation to reduce a model of bone remodeling (23).

Model calibration may also require an appropriate optimization scheme that can match results of different experiments (for example, different cellular assays) or multiple clinical outcomes (for example, the response to alternate therapeutic interventions). Optimization strategies may focus on producing an exact match of the simulation with observed trajectories, or on producing responses that fall within bounds, which are often defined as a time series of numerical bounds of observed responses. In this latter case, it may be practical to identify many degenerate solutions with optimal objective function values, since alternate parameterizations may equivalently fall within target bounds. In either case, establishing agreement with datasets from different groups of individuals with different interventions may also be necessary.

##### *Variability*

Due to biological complexities, model parameters may be poorly constrained by available datasets (24). Therefore, it is important to evaluate the impacts of known variability and uncertainty, where QSP models often use ensembles of alternative parameterizations that appear to be plausibly in agreement with observed data (8,25,26). As one concrete example, it is not uncommon to find quantitatively different *in vitro* measures of the same process reported by two different labs. QSP models enable testing the impact of mechanistic differences that would be inferred from inter-lab variability on endpoints of interest *in silico*, and therefore help triage impactful uncertainty that must be resolved or explained from uncertainty that is not impactful and is therefore not worth investing experimental or clinical resources to resolve. Alternate parameterizations are referred to as virtual subjects (8), or virtual patients (VPs) (26,27), and the ensembles are referred to as a virtual cohort or cohort of virtual patients (see Table I). Note that verification of plausibility generally may or may not be assumed in the description of a virtual patient depending on the workflow and algorithm (25), and here these are specifically referred to as "plausible virtual patients" to avoid ambiguity.

Reference virtual patients (8), or any set of plausible virtual patients, can be used as a basis to develop a larger cohort of VPs by adding stochastic noise to their parameterizations. Moreover, one can generate a cohort of plausible VPs by repeated application of a sampling-acceptance/rejection algorithm. It is essential to make sure the cohort spans all typical phenotypes observed in experiments, or to identify a mechanistic rationale if this does not appear to be possible. Therefore, cohort generation can be a time-consuming iterative process of building a cohort, identifying missing phenotypes, and refining the cohort.

After iterative refinement of a cohort of VPs, they can be used to explore a broad range of responses following a variety of simulated interventions. However, population-level distributions for the cohort may not match experimentally and clinically observed response statistics (for example, mean, standard deviation, clinical response bins, etc., see Fig. 1). The cohort can provide a sense of mechanistic factors responsible for possible outcomes, but it is difficult to interpret whether the cohort resembles a clinical trial population or the probability of observing certain outcomes (25). To overcome this obstacle, strategies have been proposed to add another layer of optimization to improve the statistical agreement between simulations and observed data (25–29). Several algorithms have been proposed to assign a statistical weight, often called a prevalence weight, to each VP in the cohort so calculated statistics for the cohort better match observations (26–29). A VP cohort that has been calibrated to better match clinical data is called a virtual population (VPop, see Table I). Due to the availability of published source code for detailed review and also to the ability of the algorithm to integrate outcomes from alternate interventions based on marginal distribution data, the previously described Mechanistic Axes Population Ensemble Linkage (MAPEL) algorithm for assigning prevalence weights was implemented as an initial method for developing VPops (26). In MAPEL, a prevalence weight is assigned to each VP with the goal of optimizing VPop agreement with experimental or trial data distributions or summary statistics (see Fig. 1). A composite goodness-of-fit (GOF) statistic is calculated, enabling assessment of agreement to multiple endpoints across multiple interventions. The resulting VPop can then be used for prediction of responses to new interventions and to analyze prevalence-weighted correlations among parameters or simulated outcomes.

##### *Application*

One can explore different mechanistic hypotheses using a cohort of plausible VPs. Clustering algorithms, such as partitioning around medoids, can also be used to reduce cohort size while maintaining phenotypic and parametric variability. A weighted VPop, on the other hand, can be used to predict population-level statistics under new interventions, like mean, standard deviation, and clinical response fractions. One important feature of QSP models is that they are often used to generate hypotheses to identify/rule out mechanisms related to the questions of interests; therefore, they are exploratory in nature (30). It is often informative about underlying biology when there are discrepancies between a QSP model prediction and observed data.

#### THE QSP TOOLBOX: OVERVIEW

The QSP Toolbox is designed to support QSP workflows including calibrating the model using experimental data, defining and generating VPs, exploring model variability by the unweighted cohort of VPs and weighted VPops, and predicting new experimental/trial outcomes. In this section, we will introduce some key features of the toolbox and its organization.

##### Toolbox Features

To build an initial cohort of plausible VPs, one needs to develop a number of alternative parameterizations to capture observed variability. Meanwhile, model calibration often

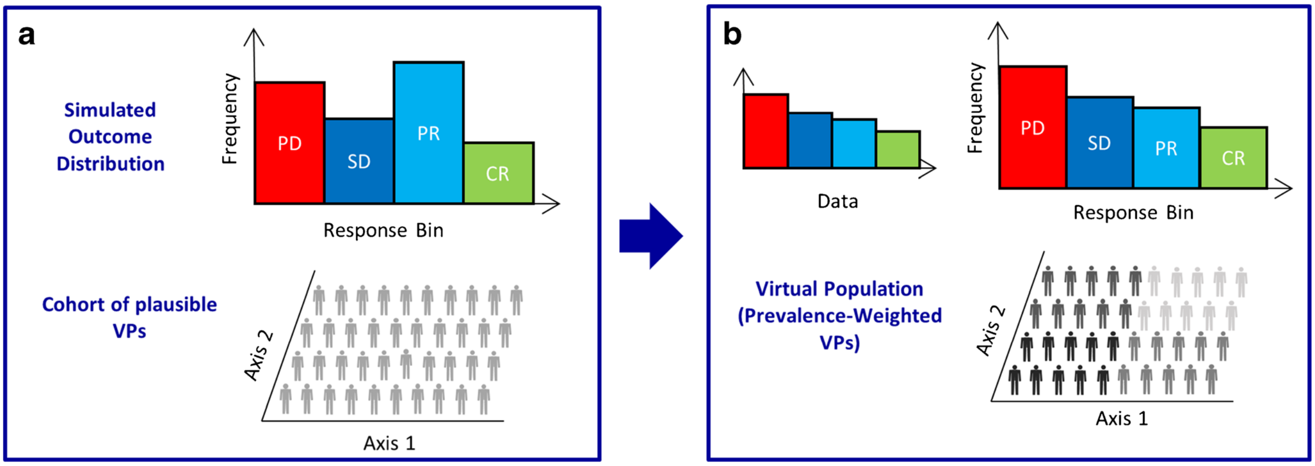


**Fig. 1.** Description of MAPEL algorithm. **a** Statistics for the simulated outcomes in the cohort may not match observed clinical or experimental data. *PD* progressive disease; *SD* stable disease; *PR* partial response; *CR* complete response (as an example, although other marginal distribution bins, mean, or standard deviation may be used in the included version of MAPEL). **b** The cohort of plausible VPs is prevalence-weighted to the calculated statistics match trial or experiment data. Each VP is assigned a prevalence weight, and the weights are optimized using MAPEL. Portions adapted from (26)

involves integration of various datasets from different conditions, described as different "interventions" in the QSP Toolbox terminology (Table I). For example, calibration may use cell culture or xenograft experiments from preclinical settings as well as different indications or dosing scenarios in clinical settings. To coordinate the calibration complexity and variability exploration in QSP workflow, the QSP Toolbox employs a "worksheet" structure array to manage VPs and interventions. To be more specific, the creation of a worksheet starts by setting relevant characteristics that defines a VP into parameter groups, which we define as variant types to maintain consistency with the terminology of variants in the underlying SimBiology® model. Parameters could include molecular, cellular level, or tissue level properties which can be determined subjectively or algorithmically. As the toolbox is implemented, if variants are specified in the SimBiology® project, the naming must follow a convention. Every variant must have a variant type (see Table I), and each VP must use exactly one set of values for each variant type in the project, denoted as a type value set. One can specify the characteristics that define a particular intervention with variants and doses. In SimBiology®, "doses" are specified separately from variants, but they are also in the intervention definition. Mechanistic axes can also be defined. The combination of the VP definition by type value sets and mechanistic axes as well as intervention by type value sets and doses defines a unique VP on an intervention for simulation. A scheme of the worksheet data structure is shown in Fig. 2. The worksheet data structure provides a convenient way of organizing VPs and interventions, enables quick access to manipulate optimization simultaneously across multiple parameters and interventions, and enables functions like clustering and reducing VPs.

Another important feature of the QSP Toolbox is the processing and integration of various experimental data. The QSP Toolbox maps experimental data to model variables and includes appropriate datasets in objective function evaluations when optimization is necessary. Various optimizations, samplings, and plotting scripts are also included in the toolbox to support statistical calibration to measured endpoints to develop a VPop.

Development of a cohort of plausible VPs and weighted VPops could be computationally expensive and time-consuming. The QSP Toolbox has the feature to compile the model to run in an accelerated manner and parallelize the computation by distributing simulations over available cores.

#### Toolbox Organization

Similar to various script-based toolboxes that have been developed to facilitate systems research (for a couple examples, see (17, 31)), the QSP Toolbox is provided in MATLAB® and can be run from the MATLAB® command line. It reads QSP models deployed in SimBiology®, a VP definition table file, an intervention definition table file, and various available experimental dataset files. By providing the functionality necessary for a more comprehensive QSP workflow, the toolbox helps to develop a cohort of VPs, weighted VPops, and miscellaneous visualizations. Predictions for new interventions using developed VPs and VPops can be easily achieved by adding new interventions to a worksheet. Utilization of the toolbox is demonstrated in detail in the examples, which are also discussed in the "Examples" section.

There are eight main folders in the package: "docs," "io," "external," "plotting," "worksheet," "cohort," "population," "examples," and "bakery." The "docs" and "examples" folders are helpful for getting started with the QSP Toolbox. The "docs" folder includes two posters demonstrating the development of one QSP model and application of the toolbox (32,33), and the "examples" folder provides script files of toolbox application that can be followed to learn how to use critical toolbox functions. Additional help is available in the header for the individual functions and object definitions, and can be accessed by typing "help [mFileName]" at the command line once the QSP Toolbox is initialized. Key elements of the QSP workflow are built in the "io," "external," "plotting," "worksheet," "cohort," and "population" folders. Various scripts for reading and writing information are included in "io," "external" includes several

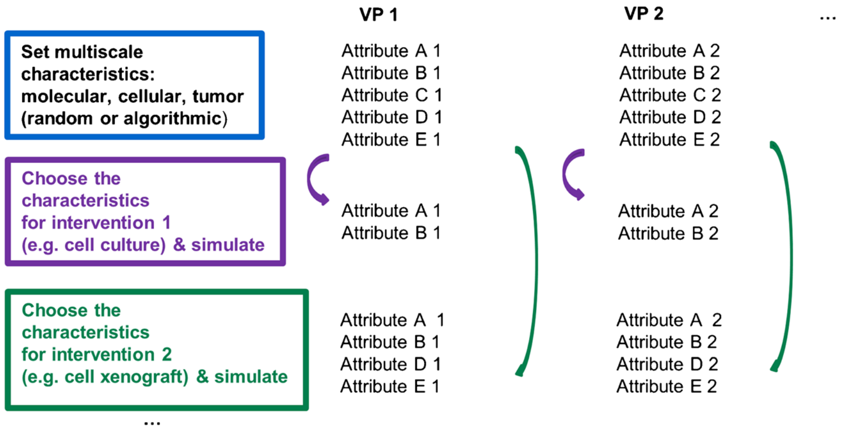


**Fig. 2.** A simplified illustration of the worksheet data structure and how parameters are selected for individual VP-intervention simulations from a worksheet. For the purpose of clarity, it is assumed attributes A–E define a VP and different VPs have different attribute values. VP 1 is defined by Attribute A 1-Attribute E 1, while VP 2 is defined by different values of these attributes denoted as Attribute A 2-Attribute E 2. When the cell culture "intervention" is simulated, the attributes that characterize VPs for this intervention are identified, which would be A and B in this case. Then we perform simulations, updating with the appropriate attribute values. Similarly, if we are simulating the cell xenograft "intervention," define simulations appropriate attributes. For example, a VP attribute such as vascular volume fraction would impact the xenograft simulation but not the cell culture simulation. Note that this simplified illustration captures critical aspects of the effective implementation of simulation parameterization. In the computational workflow, it is a multi-step process of taking the model's base parameterization, overwriting with VP-specific parameters from the variant types (if included), overwriting with VP-specific parameters from the mechanistic axes (if included), overwriting with parameters specified by variant types in the intervention definition (if included), and finally identifying specified doses for the simulation

third party files used by functions in the toolbox, and "plotting" includes various scripts to visualize the cohort of the VPs and VPops. Scripts in the "worksheet" folder build and manipulate the worksheet, scripts in the "cohort" folder help to establish a cohort of unweighted VPs and perform sensitivity analyses via sets of VPs, and scripts in the "population" are critical for developing virtual populations. Some key scripts are presented in Fig. 3. The "bakery" folder contains provisional scripts that are useful but are not systematically integrated yet with other functions.

Several experimental datasets using consistent *in vitro* cell culture and animal models (for example, N87 cell lines and N87 xenografts in mice) were generously provided by colleagues. They are included in the "examples" folder. These data were also used to guide model and toolbox development on a project as presented in the results in the "docs" folder (32,33). These data and results have been recently updated, and many are demonstrated computationally here. Animal experiments were conducted in full compliance with local, national, ethical, and regulatory principles and local licensing regulations, per the spirit of Association for Assessment and Accreditation of Laboratory Animal Care (AAALAC) International's expectations for animal care and use/ethics committees.

#### ANTIBODY DRUG CONJUGATE (ADC) PLATFORM

In the following sections, we will demonstrate the utility of the QSP Toolbox through examples where the QSP Toolbox is applied to a mechanistic ODE model of Antibody Drug Conjugate (ADC) efficacy. Before demonstrating the utility of the toolbox with this model, a brief description of ADCs and the structure of the model is required.

ADCs are a therapeutic modality that combines the affinity and specificity of antibodies with a cytotoxic drug, often referred to as payload. Unlike conventional chemotherapies that damage healthy tissues and cause severe side effects, ADCs have the potential to deliver cytotoxic agents to the tumor via cancer specific, over-expressed cell surface antigens (34,35). Exposure to the cytotoxic agent is thus potentially low in the normal tissues which have low expression of these cell surface antigens.

Structurally, there exist three components in an ADC (see Fig. 4a for illustration): (i) a targeting moiety, which could be antibody or other modality that confers affinity that permits tumor-specific localization of the ADC; (ii) linker, which is chemically optimized for preferentially releasing the payload inside the cancer cell; and (iii) payload, a cytotoxic molecule with high potency (36). The mechanism of the ADC's tumor-specific action starts with the affinity component of an ADC selectively binding a cell surface tumor antigen following transport into the tumor microenvironment. The ADC-antigen complex is internalized through receptor-mediated endocytosis. The ADC-antigen complex then traffics to lysosomal compartments and is degraded, releasing the cytotoxic payload inside the cell leading to cell death, often by binding microtubules or causing DNA damage (37). These processes are illustrated in Fig. 4b. Each component of an ADC must be optimized to fully realize the goal of a targeted therapy with improved efficacy and tolerability.

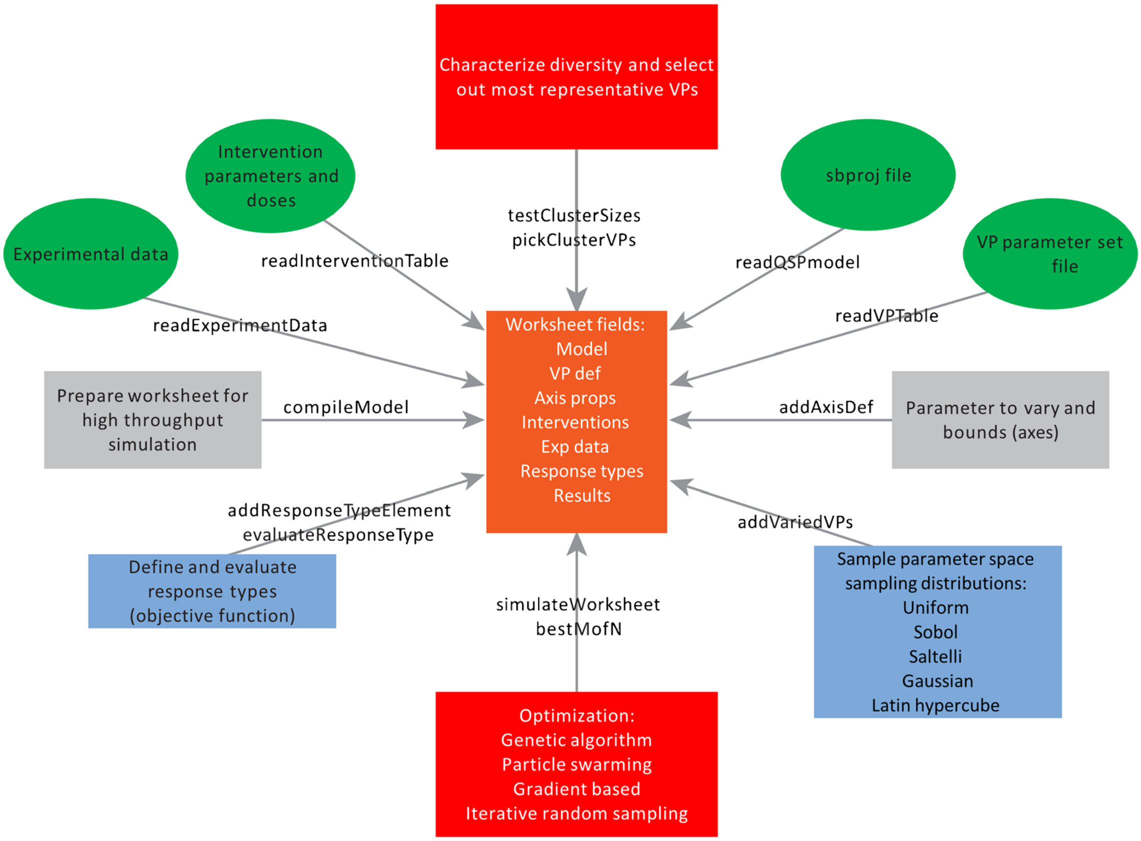


**Fig. 3.** Key scripts for VP cohort development. The worksheet data structure plays a central role during virtual patient cohort development, which begins from processing user-provided files and continues to store refinements though the iterations of cohort development. The annotations on the *arrows* represent the function names that relate the worksheet data structures and different processes. The *green ovals* represent how the user-provided files are read into the worksheet data structure. For example, readQSPmodel.m reads the SimBiology model from a sbproj file, readVPTable.m reads VP definitions from file and so on. The *gray rectangles* represent some important integration and preparation steps for cohort generation. compileModel.m updates a list of model parameters, initial conditions, and compartment values, collectively referred to as elements in the commenting, and also accelerates the model to prepare for high-throughput simulation. addAxisDef.m adds mechanistic axes and the corresponding bounds to define the parameter space to explore. The *blue rectangles* represent two key functions for sampling and rejection/acceptance. addVariedVPs.m provides different sampling strategies for the mechanistic axes. addResponseTypeElement.m and evaluateResponseType.m specify the experimental data used to calibrate the model and evaluate the performance of the sampled VPs. The *top red rectangle* provides functions to cluster and select representative VPs and the *bottom red rectangle* represent various optimization tools built in the toolbox

There are several processes complicating the investigation of ADC efficacy and tolerability. First, random conjugation of the cytotoxic payload to the targeting molecule yields heterogeneous mixtures of ADCs that often can include 0–8 payload species per molecule (37). With antibodies as a targeting modality, this results in transport of ADCs with

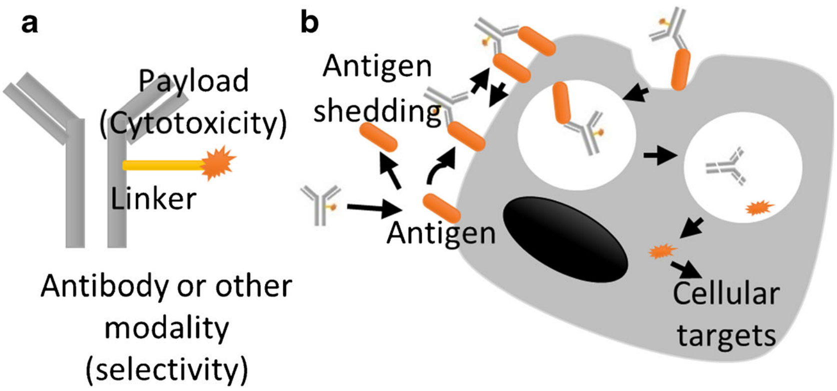


**Fig. 4.** ADC structure and function. **a** Structure of an ADC. **b** Delivery of cytotoxic payload to cancer cells by an ADC

different drug-to-antibody ratios (DARs) among different compartments. Second, selective tumor targeting could be a complicated process requiring consideration of both systemic clearance and tumor-specific factors that impact delivery into the microenvironment, such as vascularization and capillary permeability (38). Third, many tumor-specific surface antigens are actively shed from cancer cells and shedding could potentially influence not only the tissue distribution of ADCs, with not only implications for safety (39–41), but also potentially the ability to permeate from tumor vessels by nature of the impact of the size of the complex (38). Shed antigen might also impact the ability of ADCs to bind their cellular targets in tumors (42,43). A QSP model can be used to integrate many of these complex, nonlinear, and unintuitive processes involved in the cellular and physiological disposition of ADCs and their components. The model thus becomes a tool for investigation of the key determinants of ADC pharmacodynamics, can be used to optimize the chemical/physical properties of the ADC to maximize efficacy and tolerability, can be used to develop clinical dose projections based on preclinical data (44), and furthermore can be used to explore the clinical implications of mechanistic variability.

Several models have been proposed to mechanistically model ADC efficacy (43–52), many of which have been previously reviewed (41). Mechanistic models have often been applied to investigate antibodies as a modality, although modeling studies focused on immunotoxins have also been reported (43,52). It has also been suggested that ADCs are a special case of "affinity" drug conjugates that can include alternate modalities to confer specificity (32,33), such as adnectins (53). Mechanistic models of ADC efficacy often include three important components: PK of the ADC in the plasma, transport into the tumor microenvironment, and disposition of the ADC in a target cell.

We have developed a QSP model of ADC tumor delivery and efficacy, and a diagram view of the ADC platform is shown in Fig. 5. The ODE model is similar in many aspects to the approach taken by Shah *et al.* (44), but there are differences that impact the potential platform applications in the species that are represented, how the platform is initialized, and different sections of the model that can be activated. The platform includes a compartmental tumor model (38,54), and also a basic framework for the simulation of individual DAR (50), shedding, and bivalency (55). In support of extending the model to other modalities conferring specificity, valency for antigen (bivalency vs. monovalency) is controlled via a parameter that behaves as a switch. In addition, the lesion capillary permeability for each soluble species is calculated based on vessel properties and molecular weight (38).

Note that for the purposes of illustrating a QSP workflow, we have focused primarily on mechanistic tumor rather than pharmacokinetic variability. We have also not provided an analysis of cellular payload disposition in the examples, which would require additional data. We have developed the model providing equations with consistent clearance rates and DAR-proportional deconjugation and metabolism rates, and these equation forms may be refined with additional data to better support parameterizing DAR-dependent rates (56–58).

#### Examples

We demonstrate various aspects of the QSP Toolbox, primarily by applying the toolbox to the ADC QSP model with several datasets from cell culture and interventions with a naked antibody without attached payload. Seven examples are used to cover different features of the toolbox, including cross referencing VPs and interventions, generation of cohorts of VPs, and calibration of VPops. We give summaries of the examples, and one can find detailed comments in the example scripts included in the toolbox. Since N87 cell line and N87 xenograft data were used, we refer to the VPs as virtual xenografts (VXs). Examples 1–3 walk through toolbox functions with the ADC platform using a smaller set of parameters to explore and endpoints for calibration. Examples 4–6 illustrate iterative cohort refinement and VPops with a slightly expanded set of parameters and endpoints for calibration. Example 7 gives an example of a global sensitivity analysis.

It is generally recommended to run the examples by opening them and following the extensive comments, executing their steps either by copying and pasting commands at the prompt or by using the "Evaluate Selection" feature in the debugger. Note that in order to use the toolbox, the command "initQSPToolbox" must be run from the MATLAB® command line, and the root path to the toolbox should be added to the path manually. Subdirectories will be added at initialization. The following examples were run with MATLAB® Release 2016a.

##### *Example 1: Load, Simulate, and Plot Results from a Worksheet by Cross-Referencing VX and Interventions*

In this example, a SimBiology® model, VX definitions (the columns of a worksheet), and intervention definitions (the rows of a worksheet) are imported and integrated together into a worksheet. Parameter values from the model variants are used to define one VX. Four different interventions are applied: an N87 cell culture simulation, an N87 xenograft with buffer injection, an N87 xenograft with naked antibody injection, and an N87 xenograft with an $^{89}$Zr-labeled antibody injected. The simulated time courses of the observed biomarkers are plotted.

##### *Example 2: Vary Mechanistic Parameters with the Toolbox*

In example 2, mechanistic axes are specified to illustrate how to create alternate VXs. Five parameters are added, each with their own mechanistic axis with predefined bounds. These parameters are antigen shedding rate, ADC internalization rate, large vessel porosity, shed antigen clearance rate, and positron emission tomography (PET) label export rate. Additional parameterizations are sampled from the five dimensional hypercube defined by the bounds, the resulting 101 VXs are simulated each with the same four interventions as in example 1, and the simulated time courses of observed biomarkers are plotted and compared to experimental data for net internalization in culture, xenograft shed antigen concentration, and xenograft $^{89}$Zr PET label accumulation. The randomly generated VXs are compared to the experimental data. Results for these randomly generated VXs are shown in Fig. 6 a, b, c, d. Most VXs do not match the experimental internalization, shed antigen, or label accumulation data very well. Fig. 6e illustrates axis coefficients in the VXs. We also save the worksheet in example 2 to serve as a starting point for example 3.

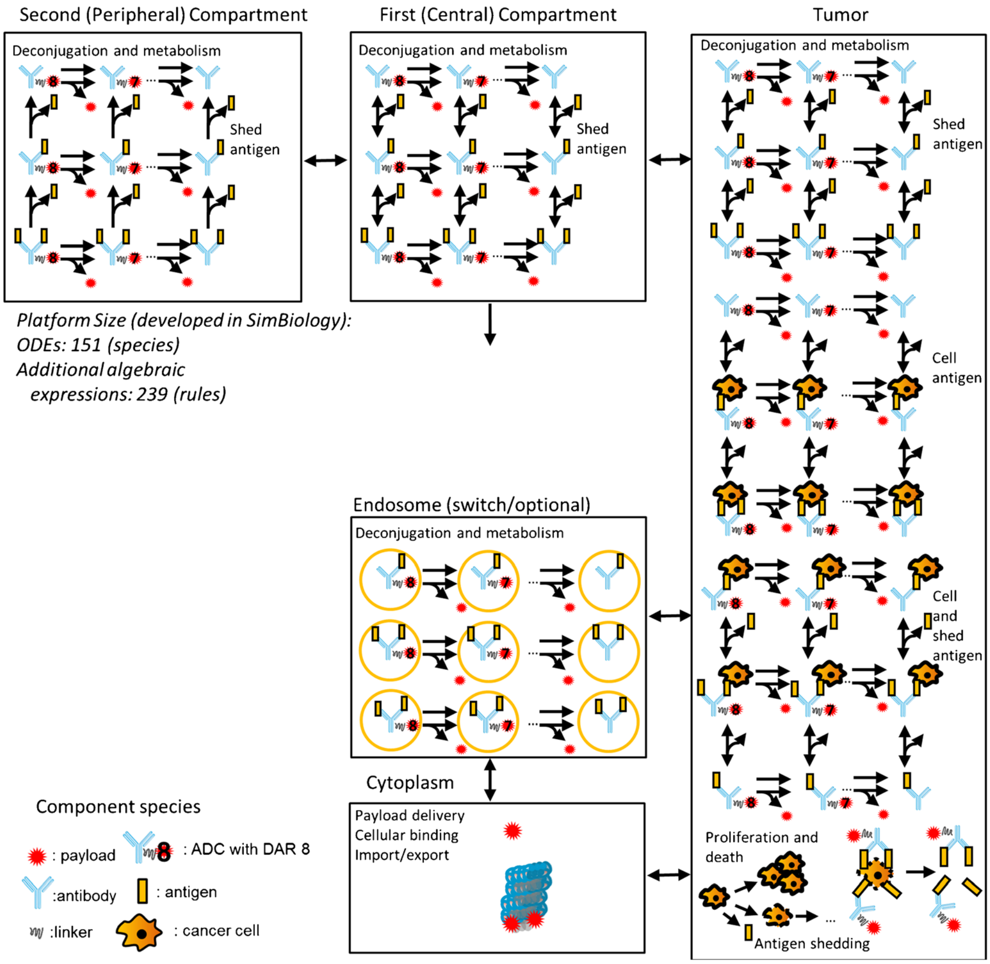


**Fig. 5.** A diagram of the ADC QSP model. In the platform, we currently implement an antibody drug conjugate, but the molecular weight, valency, and pharmacokinetics can be reparameterized in the platform. Two compartments are included for ADC and payload pharmacokinetics; there is a tumor compartment, and there is also an endosomal compartment that is not used in the examples but may be enabled to support additional data on cellular disposition or to enable exploratory simulations of cellular disposition. The biomolecular species on the platform that can form as a result of payload deconjugation and metabolism, shed antigen binding, cellular antigen, and intracellular processes binding are shown

By establishing an objective function, one can also quantitatively evaluate the agreement of VXs with data and keep the best. The objective function evaluation implemented here is detailed in one of the posters included in the documents folder (33). Individual response type elements map simulated model outcomes to experimental data and specify an objective function form (see Table I). The response type elements are each evaluated with their individual objective functions. The individual objective function evaluations are all summed together with equal weight in the evaluation of the response type. Fig. 6 f, g, i show simulations of the selected 10 best VX out of 300 randomly generated VXs.

*Example 3: Optimization of VXs*

In example 2, the randomly sampled VXs usually do not match the experimental measurements, although we can find VXs that match better by sampling larger numbers of VX. In example 3, the worksheet created in example 2 is loaded and 2 VXs from the 101 VXs are included as initial points for an optimization to best match experimental data. There are per-VX approaches where we run optimization once for each VX (That means we aim to identify two valid VXs). Alternatively, there are "cohort" approaches where we run optimization informed with all existing VXs in the initial pool, and the size of the returned cohort is also specified. We use a particle swarm and genetic algorithm in each approach, respectively. Simulation results of the optimized VXs are presented in Fig. 7 a, b, c, d and e, f, g, h, respectively. The optimization algorithms, either per-VX approach or cohort approach, can effectively create VXs in better agreement with the data. Available local optimization methods may be particularly attractive for certain applications with the per-VX approach.

*Example 4: An Iterative Workflow for Finding Additional Cohort VXs Given an Initial Set of Plausible VXs*

One can get an initial set of plausible VXs through sampling the hypervolume defined by the mechanistic axes bounds and accepting/rejecting VXs based on their response

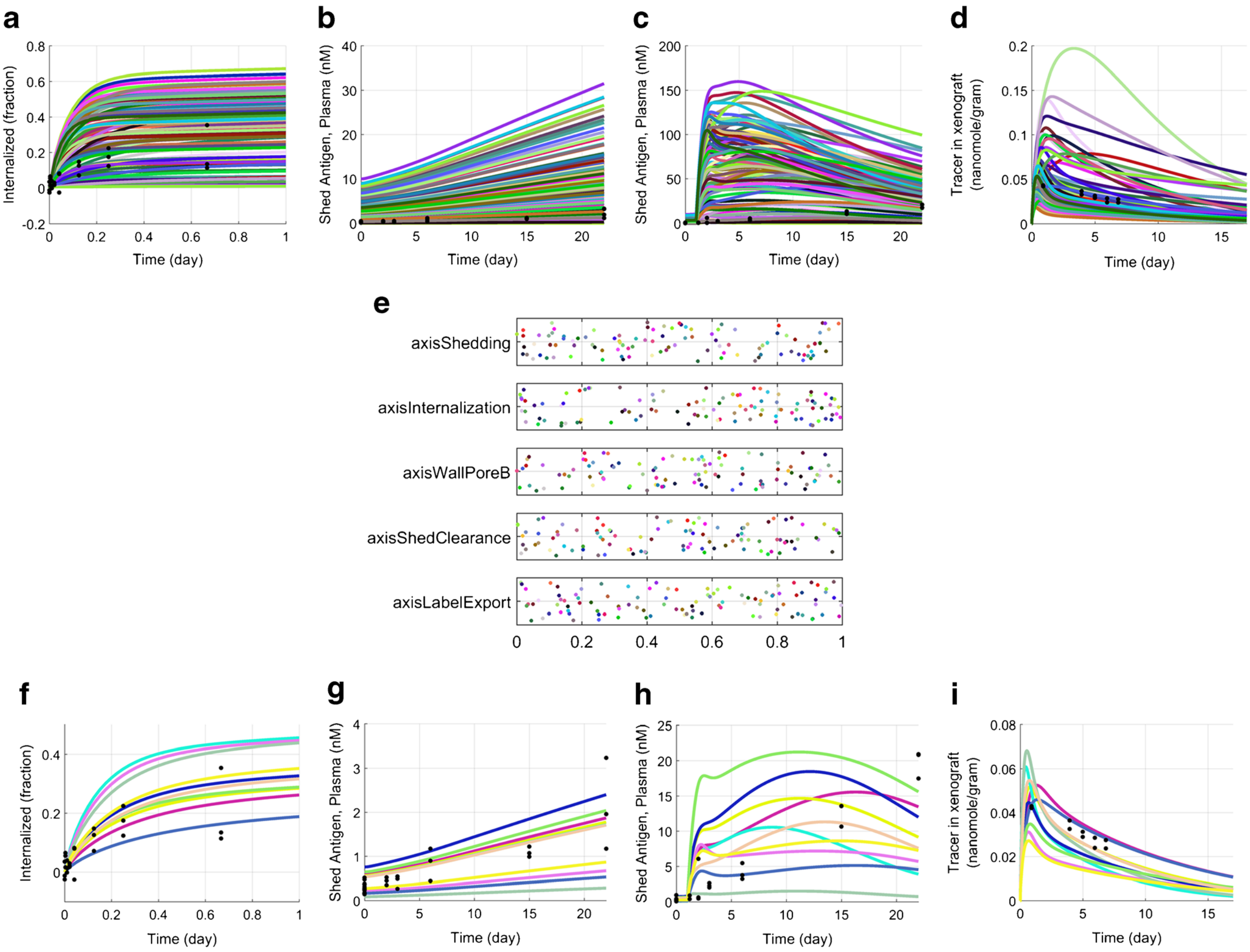


**Fig. 6.** Time courses of observed biomarker of VXs under four different interventions. **a**–**d** Simulated time courses of observed biomarkers of 100 randomly sampled and one initial VX. The *black dots* are the experimental data. **e** Sampled parameter distribution of the 101 VXs scaled between 0 and 1. **f**–**i** Selected simulated time courses of observed biomarkers of the best 10 VXs out of 300 VXs based on the quantitative evaluations of VX agreement with data

type elements (that is, individual simulation outcomes mapped to data) or an overall response type (combined evaluation) objective function value. If missing phenotypes are identified, an iterative and potentially time-consuming process is needed to refine the initial cohort. In this example, a process of generating additional plausible cohort VXs given an initial set of plausible VXs is illustrated. The initial cohort size is 1000 and we want to target a new cohort size of 1000. In order to generate new VXs, initial VXs are selected by using a partition around medoids clustering strategy to select 300 VXs. In iterative simulations, 7 new VXs are added for each existing VX, resulting 2100 additional VXs in total in the first iteration. These new VXs are generated by adding Gaussian noise to the axes. Note that other sampling distributions for worksheet VPs are available (type “help varyAxesOptions”). To enable more rapid exploration of the allowed solution space with each iteration, one mechanistic axis is selected for each VX and a new coefficient is also sampled from a uniform distribution. In practice, we may also set up a biased sampling to try to more efficiently generate specific phenotypes. From these 2100 VXs, we pick out the plausible VXs that satisfy our constraints on the objective function. If the number of accepted VXs exceeds the target, VXs are again selected based on clustering to bring the worksheet back down to target size before backing up progress by writing to file. In this example, 13 out of 2100 new VXs were selected as plausible VXs in the first iteration, and 5 new VXs were selected in the second. Note the rate of successful plausible VX generation, as well as generation of diversity and desired phenotypes, will vary substantially with the sampling strategy, applied response types and objective functions, and plausibility criteria.

##### *Example 5: Sample VPop Object*

Rather than first illustrating the VPop development workflow, example 6 presents a worksheet, virtual xenograft cohort, and VPop object from such an analysis. Note that the included virtual xenograft cohort also meets plausibility constraint cutoffs imposed on the response type elements, as described in the included poster (33). Also, when saving this worksheet, the simulated results were set as empty (type “[myWorksheet].results = {}”). The results may consume substantial disk space and MATLAB® file IO can be slow

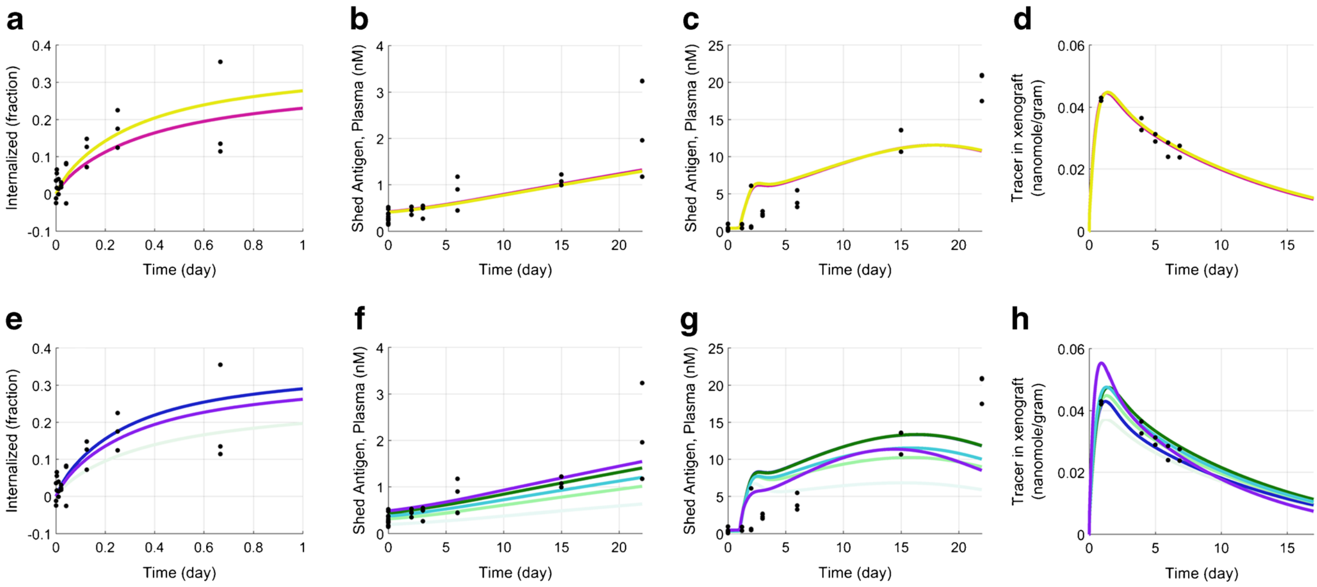


**Fig. 7.** Time courses of observed biomarkers of optimized VXs under four different interventions. **a**–**d** Simulated time courses of observed biomarkers of two valid VXs using per-VX approach. The *black dots* are the experimental data. **e**–**h** Simulated time courses of observed biomarkers of ten valid VXs using "cohort approach." The cohort size was set as 100 and VXs with a total objective function evaluation within a factor of 2 of the best are chosen. The optimization is set to terminate after 30 iterations

with files greater than about 2 GB. One can always regenerate results by simulating the worksheet.

The VPop and its prediction intervals are illustrated in Fig. 8. Note that here we include 6 measured outcomes on 4 interventions in the response type and variability in 13 mechanistic axes. The rationale for the included axes is available in the included poster (33). In the VPop object, there are simulation results from 1000 plausible VXs and each VX is assigned an optimized prevalence weight. The results suggest that valid VXs cover many of the parameter ranges, with a few exceptions (most notably shedding rate, see Fig. 8m). This suggests differences in the study cell line from bounds derived from the literature.

##### *Example 6: Workflow for Creating VPops Using MAPEL*

We demonstrated a previously developed VPop in example 5. Example 6 illustrates a workflow for creating VPops from a cohort of unweighted VXs using MAPEL (26), subject to an alteration of MAPEL's objective function as described in one of the included posters (33). Summary statistics for the VPop to match are calculated from experimental data. In the published MAPEL version, the goal is to match mean, standard deviation, and bin frequencies. It should also be noted that QSP models often may be calibrated with time-series data with various experimental and clinical assays, and one can often identify several time/experimental data points that would be difficult to match due to issues in the data. In some cases, one may be comfortable allowing some transient model behaviors to quantitatively deviate from observed data. Data for select time points may be omitted during optimization with the provided MAPEL implementation by setting their weight to zero. A diagnostic plotting function is provided to help assess how VXs in the worksheet span the data. Additional diagnostic plotting functions are available to help identify endpoints that may be problematic, as well as the distribution of prevalence weights in the VPop.

##### *Example 7: Sobol Global Sensitivity Analysis*

Sobol sensitivity analysis has been used to characterize systems pharmacology models (59,60). Sensitivity analysis has also been presented as an important method to support QSP workflows (8), and here we illustrate how to run a Sobol global sensitivity analysis using a test function that can be solved quickly, the six-parameter Sobol g-function (61). The general form of $k$ parameter Sobol g-function is given by:

$$g(x_1, x_2, \cdots, x_k) = \prod_{i=1}^{k} g_i(x_i) \tag{1}$$

With

$$g_i(x_i) = \frac{|4x_i - 2| + a_i}{1 + a_i} \tag{2}$$

In this example, the Sobol g-function is implemented as a SimBiology model and all six parameters are mechanistic axes with bounds between 0 and 1. Sampling for the first-order sensitivity and total-order sensitivity indices are determined using an efficient model evaluation strategy (61,62), and computational formulas for the evaluation of the indices were selected based on previous proposals (61–63). The first-order sensitivity indices and total-order sensitivity indices are compared to the respective analytical solutions in Fig. 9. Note that with sufficient time and a suitable computing environment with sufficient computing cores and memory, larger sampling runs may be applied to QSP models to calculate the sensitivity indices.

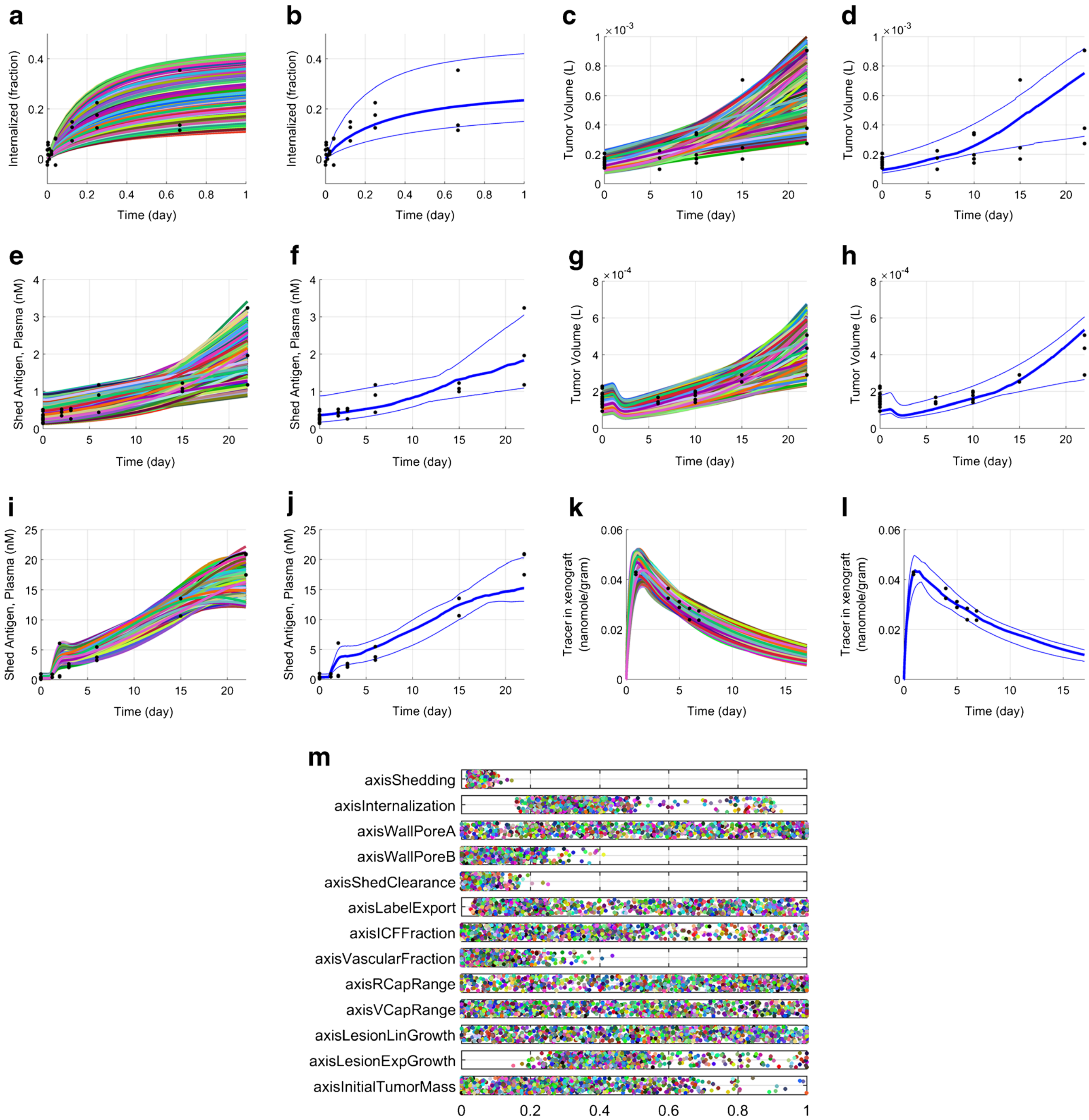


**Fig. 8.** VPop demonstration. *Plots* show indicated simulated outcomes for each VX in the cohort for: **a** cell culture; **c**, **e** buffer injection; **g**, **i**: naked antibody injection; **k**: $^{89}$Zr-labeled antibody injection. **b**, **d**, **f**, **h**, **j**, **l**: 5/50/95 prevalence-weighted VPop percentiles. That is, these lines are the prediction interval using the VPop. **m** Scaled mechanistic axis coefficients of 1000 VXs. Here, each axis maps to one model parameter, each dot maps to one virtual xenograft, and each parameter is scaled within each axis range

#### Advanced Concepts

The examples included in the tutorial walk through setting up a worksheet, developing and evaluating alternate VPs, establishing a cohort of VPs, and calibrating a VPop. QSP modelers will not necessarily want to replicate these steps exactly for their projects, but will be interested in applying variations on these steps with their own models and datasets. This will likely entail variations or expansion on the basic steps presented here. Capabilities to support both variations on these steps and additional workflows are provided. As one example, time-series data were available here, and response type elements based on datapoints (type "help responseTypeElementPoints" at the command line for additional information) were implemented to assess the agreement of simulated virtual xenograft outcomes with the experimental data. Additional response type element objects were also developed for the QSP Toolbox, and agreement with bounds over defined intervals can be assessed with other

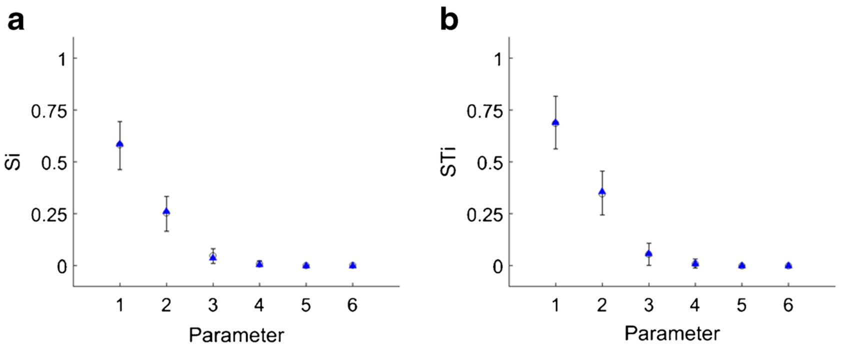


**Fig. 9.** Sobol global sensitivity analysis demonstrated by Sobol's g-function. The *lower error bar*, *circle*, and *upper error bar* indicate the 2.5, 50, and 97.5 percentile from bootstrapping, and the *triangle* indicates the analytical solution. **a** First-order sensitivity indices. **b** Total-order sensitivity indices

included classes (type "help responseTypeElementBounds" at the command line). This approach will be essential to generate VPs when it is desired to check that simulated outcomes fall within ranges that can be established from available literature. A response type element class is also available to help create VPs biased toward mechanistic parameter values (type "help responseTypeElementAxis"). An example demonstrating an analysis of correlation in a weighted virtual population is not provided. However, it is possible to use the examples as a starting point, and functions are provided to support the analysis (type "help evaluateCorrelations" (64). Functions to support a basic control coefficient analysis of sensitivity, similar to that applied to a model of immunogenicity (65), are provided (type "help runControlCoefficientsSimulations" and "help calculateControlCoefficients"). We also do not demonstrate implementation of pharmacokinetic (PK) variability according to a defined population model here, but for other projects we have implemented such variability by including the PK parameters as mechanistic axes and performing multivariate sampling for their coefficients from an established population PK model to set the axis coefficients. While the interventions generally implement a fixed dose, a provisional framework for simulating VP-customized dosing is provided in the "bakery" folder (type "help simulateWorksheetIndDoses"). Other workflows of interest to QSP modelers include the creation of alternate virtual populations. For example, it may be necessary to characterize how an enrollment criteria impacts a trial outcome or how much variation there might be in observed biomarkers for a given efficacy signal. If alternate enrollment criteria are based on a different class of patients with a separate set of published clinical trials, as needed the cohort VPs can be selected based on additional restrictions, and the new set of trial statistics can be used to guide the development of new sets of prevalence weights for the cohort. Similarly, multiple alternate virtual populations with acceptable composite goodness-of-fit scores can be developed to explore population variability in biomarkers.

In developing VPops that statistically match observations, we have taken the approach described previously of developing virtual cohorts of feasible virtual patients and then optimizing prevalence weights (26–29). Note that a related strategy of optimizing prevalence and then sampling VPs to define the VPop has also been implemented (25). Example 4 illustrates how to use existing plausible VPs as a basis to look for additional plausible VPs, and similar methods may be used to create VPs with missing phenotypes. One exciting area for future development that has also been suggested by Allen *et al.* is the development of methods to better direct the VP sampling step to more efficiently create VPs and missing or poorly represented phenotypes (25).

Virtual patient generation involves sampling a high-dimensional space and extensive model evaluations, which could be computationally expensive. Note that these calculations can be easily parallelized since the essence is evaluation and scoring of alternative parameterizations of a QSP model. The QSP Toolbox has built in a simple parallelization feature employing the parallel simulation of the VP-intervention combinations in MATLAB®. As there is a demand to simulate more virtual patients or run models that require longer wall clock simulation times, there are options to increase resources for QSP Toolbox deployment. Options for larger worker pools may be available with the MATLAB® Distributed Computing Server, and Amazon currently offers X1 instances with 64 physical cores and 2 TB of memory. Emakov *et al.* previously presented a framework that could utilize submission of compiled QSP model executables directly to a distributed cloud environment (66). We are currently in the process of developing a cloud deployment for the QSP Toolbox that writes directly to database tables to store simulated worksheet results.

The QSP Toolbox is dependent on interfacing with MATLAB® and SimBiology®. Advantages to this strategy include the capability to import models developed elsewhere that are available in Systems Biology Markup Language (SBML), good support with multiple solvers for ODE models, a library of relevant functions and optimization algorithms from which to draw, demonstrated capability to run models with more than hundreds of state variables, and parallel computing capabilities. Additional benefits from SimBiology® include support of additional considerations in QSP modeling such as drug doses and discontinuities, the graphical user interface for organizing large models, the availability of variants to help organize parameters, and general interest from the developer in continuing to improve SimBiology® as a tool for the QSP community. Although the QSP Toolbox is freely available, other associated software costs or preference for another scripting language may be disadvantages for some users. It is also worth noting that with a sufficient development community and dedicated effort, the QSP Toolbox or package to offer similar or additional QSP efficiencies could also be developed with a free and open source solution, as has been done with other tools in systems biology (for example, see (17, 67)).

#### CONCLUDING REMARKS

QSP models provide valuable mechanistic insights about a biological system and the effect of drug treatment on system behavior. Workflows to develop reasonable parameterizations, comprehensively integrate different experimental data, and investigate parameter uncertainties/variabilities are very important for QSP model deployment. In this paper, we introduced an implementation of a computational workflow, the QSP Toolbox, to systematically deploy multi-scale mechanistic models and demonstrated its capabilities in data integration, model calibration, and variability exploration using an ADC QSP model. It is anticipated that the QSP Toolbox will accelerate model application. Although the tutorial uses an ADC model as an example, the QSP Toolbox was developed to process and analyze ODE models deployed in MATLAB® SimBiology® regardless of therapeutic agent, disease area, or indication. Furthermore, as with computational tools in other fields, we anticipate releasing the QSP Toolbox will help to strengthen the efficiency of the QSP community (20). Here, we directly communicate and demonstrate many methods and provide a framework for others to contribute improved algorithms directly to. With more QSP modelers using and potentially contributing to the toolbox, we can help standardize and enhance the computational implementation of QSP workflow, which is essential to better compare modeling practice, communicate QSP models with a matrix team, and promote QSP modeling. Furthermore, the provided detailed examples and code base may also be useful for others to adapt and expand on workflows detailed here.

#### ACKNOWLEDGEMENTS

We gratefully acknowledge suggestions from Mr. Ricardo Paxson and Dr. Arthur Goldsipe of MathWorks® on improving aspects of the QSP Toolbox.

**Open Access** This article is distributed under the terms of the Creative Commons Attribution 4.0 International License (http://creativecommons.org/licenses/by/4.0/), which permits unrestricted use, distribution, and reproduction in any medium, provided you give appropriate credit to the original author(s) and the source, provide a link to the Creative Commons license, and indicate if changes were made.
