## Supplementary File 3 for "A surrogate workflow for virtual population development: designing an amenable proxy for a hypothesis-test inspired nonlinear goodness-of-fit": example01.m

```
% Here is an example to load, simulate, and plot results from a
% worksheet using the QSP toolbox

% We either need to start with the MATLAB active directory in 
% The QSP root directory or we need to add this before calling
% initQSPtoolbox.  For example:
% toolboxPath = 'DRIVE:/DIRs/QSPToolbox';
% addpath(toolboxPath);
% Toolbox initialization
initQSPToolbox;

% Read in the model
myWorksheet = readQSPModel('adc_platform_public_v0p15p1.sbproj');
% Read in the VP (column) definitions
myWorksheet = readVPTable(myWorksheet, 'example_vp_file.txt');
% Read in the intervention (row) definitions
myWorksheet = readInterventionTable(myWorksheet, 'example_intervention_file.txt');
% Take a look at the fields in the worksheet.
myWorksheet
% Note that there are a few conventions in how SimBiology's variants are
% implemented in the model, these will need to be followed when building
% your own VPs.
myWorksheet.vpDef{1}.variants
% Each variant has a unique prefix, followed by  a delimiter ("___"),
% followed by a suffix.
myWorksheet.variantProps
% The prefixes are called variantTypes
myWorksheet.variantProps.variantTypes
% Each variantType contains a consistent set of parameters.  The suffixes
% correspond to a different set of values defined by that prefix, here
% we have designated these as typeValueSets
myWorksheet.variantProps.typeValueSets
% That is, there is a 1 : many mapping of prefixes to suffixes.
% Also note that the VP definition includes 1 of each of the variantTypes.
% This is generally enforced.
% Interventions may be defined by both doses and variants.
myWorksheet.interventions{3}.definition
% Note that there may be variable variantTypes for each intervention.
% When we go to simulate, the parameters will be combined at the level of
% the individual VP and intervention. Parameters will overwrite eachother
% in a specific order, and this convention enables simulation of the same
% VP over different interventions like cell culture where we may want to
% set exchange with an allowable plasma compartment to zero, in contrast
% with a xenograft simulation.  The order of assignment is:
% 1. Base model parameter values
% 2. Parameters from VP definition in variants, which are assigned
%    in the order of appearance.
% 3. Parameters from VP parameter axes, which we will introduce in a later
%    example
% 4. Parameters set from intervenions.
% Note that we can take advantage of this order to overwrite parameters
% that we know will be fix by a given intervention.

% Set some of the simulation properties - how often to save a result,
% how long to simulate for.
myWorksheet.simProps.sampleTimes = [0:.025:35]';

% Simulate the worksheet.  Note that simulateWorksheet
% will invoke a check to see if
% the model is accelerated or not.  If not, it will and will re-do 
% the gathering of model elements and export but will also add the 
% acceleration.
% Also note that simulateWorksheet will prepare the jobs to run
% in parallel on all available cores.
myWorksheet = simulateWorksheet(myWorksheet);

% Plot results of interest
% Start with cell culture internalization result
myPlotOptions = plotOptions;
myPlotOptions.xLabelPretty = 'Time (day)';
myPlotOptions.yLabelPretty = 'Internalized antibody (fraction)';
myPlotOptions.xLim = [0 0.8];
myPlotOptions.xShiftSim = 0;
myPlotOptions.flagLegend = false;
myPlotOptions.varName = 'parrule_payload_fraction_of_initial_bound_intracellular';
myPlotOptions.interventionID = 'culture_internalization';
plotHandle = plotAcrossIntervention(myWorksheet, myPlotOptions);
%
% Check the shed antigen prediction for the untreated
myPlotOptions = plotOptions;
myPlotOptions.xLabelPretty = 'Time (day)';
myPlotOptions.yLabelPretty = {'Shed antigen plasma','concentration (nM)'};
myPlotOptions.xLim = [0 22];
myPlotOptions.xShiftSim = 10;
myPlotOptions.flagLegend = false;
myPlotOptions.varName = 'parrule_shed_total_plasma_concentration';
myPlotOptions.interventionID = 'buffer_injection_1200147_shed_assay';
plotHandle = plotAcrossIntervention(myWorksheet, myPlotOptions);
%
% Check the shed antigen predictions for the antibody-treated
myPlotOptions = plotOptions;
myPlotOptions.xLabelPretty = 'Time (day)';
myPlotOptions.yLabelPretty = {'Shed antigen plasma','concentration (nM)'};
myPlotOptions.xLim = [0 22];
myPlotOptions.xShiftSim = 10;
myPlotOptions.flagLegend = false;
myPlotOptions.varName = 'parrule_shed_total_plasma_concentration';
myPlotOptions.interventionID = 'antibody_injection_1200147_shed_assay';
plotHandle = plotAcrossIntervention(myWorksheet, myPlotOptions);
%
% Check the predictions for N87 PET xenografts
myPlotOptions = plotOptions;
myPlotOptions.xLabelPretty = 'Time (day)';
myPlotOptions.yLabelPretty = {'Tracer in xenograft','(nanomole/gram)'};
myPlotOptions.xLim = [0 17];
myPlotOptions.xShiftSim = 10;
myPlotOptions.flagLegend = false;
myPlotOptions.varName = 'parrule_payload_total_apparent_pet_tumor_nanomole_per_gram';
myPlotOptions.interventionID = 'agxab1pet_injection';
plotHandle = plotAcrossIntervention(myWorksheet, myPlotOptions);
```
