## Supplementary File 3 for "A surrogate workflow for virtual population development: designing an amenable proxy for a hypothesis-test inspired nonlinear goodness-of-fit": example02.m

```
% A modification from example01 to demonstrate how to add
% mechanistic parameters that can be easily varied in the toolbox,
% here these are called mechanistic axes.
%
% Toolbox initialization
% toolboxPath = 'DRIVE:/DIRs/QSPToolbox';
% addpath(toolboxPath);
initQSPToolbox;
% Read in the model
myWorksheet = readQSPModel('adc_platform_public_v0p15p1.sbproj');
myWorksheet = readVPTable(myWorksheet, 'example_vp_file.txt');
myWorksheet = readInterventionTable(myWorksheet, 'example_intervention_file.txt');
%
% Add mechanistic axes definitions to the worksheet.
% Note that adding axes will trigger a model export and getting the model
% "elements" (parameters, species, compartment values)
myWorksheet = addAxisDef(myWorksheet,'axisShedding',{'par_ag_shed_rate_t','parameter', [0, 10]});
myWorksheet = addAxisDef(myWorksheet,'axisInternalization',{'par_ag_endocytosis','parameter', [0, 10]});
myWorksheet = addAxisDef(myWorksheet,'axisWallPoreB',{'par_fraction_capillary_pore_area_per_unit_thickness_b','parameter', [0.2, 0.8]});
myWorksheet = addAxisDef(myWorksheet,'axisShedClearance',{'par_shed_antigen_clearance_p1','parameter', [20, 80]});
myWorksheet = addAxisDef(myWorksheet,'axisLabelExport',{'par_payload_secretion_rate','parameter', [0, 24]});

% We can also adjust tolerances as needed
myWorksheet.simProps.relativeTolerance = 1E-6;
% Even more strict tolerances can be used, commented out here
% myWorksheet.simProps.absoluteTolerance = 1.0000e-20;
% myWorksheet.simProps.relativeTolerance = 1.0000e-12;
% myWorksheet.simProps.absoluteToleranceScaling = false;

% Pick a VP to vary - we only have one in the worksheet
myVaryAxesOptions = varyAxesOptions();
% You can check the help
help myVaryAxesOptions
myVaryAxesOptions.newPerOld = 100;
myVaryAxesOptions.varyAxisIDs = getAxisDefIDs(myWorksheet);
myVaryAxesOptions.baseVPIDs = {'n87_xenograft_scenario_0006'};
myVaryAxesOptions.intSeed=1;
newWorksheet = varyAxes(myWorksheet, myVaryAxesOptions);
%
% Simulate the worksheet
newWorksheet.simProps.sampleTimes = [0:.025:35]';
newWorksheet = simulateWorksheet(newWorksheet);

% Add experimental datasets to the worksheet, so we can include
% them on the plot and include them in response types.
newWorksheet = readExperimentData(newWorksheet, 'fluorescence_quench_internalization_data_n87_ab1.txt');
newWorksheet = readExperimentData(newWorksheet, 'xenograft_data_1200115_n87_buffer_only.txt');
newWorksheet = readExperimentData(newWorksheet, 'xenograft_data_1200147_n87_buffer_only.txt');
newWorksheet = readExperimentData(newWorksheet, 'xenograft_data_1200147_n87_ab1_only.txt');
newWorksheet = readExperimentData(newWorksheet, 'xenograft_data_95305073_n87.txt');
% You can see the experimental data in the worksheet, it is read as tables.
% This does create some name length restrictions for the experimental
% variables
newWorksheet.expData{:}

%
% Plot results of interest
% Check the predictions for culture internalization
myPlotOptions = plotOptions;
myPlotOptions.flagSave = false;
myPlotOptions.xLabelPretty = 'Time (day)';
myPlotOptions.yLabelPretty = 'Internalized (fraction)';
myPlotOptions.xLim = [0 1];
myPlotOptions.yLim = [];
myPlotOptions.xShiftSim = 0;
myPlotOptions.flagLegend = false;
myPlotOptions.varName = 'parrule_payload_fraction_of_initial_bound_intracellular';
myPlotOptions.interventionID = 'culture_internalization';
myPlotOptions.expDataID = 'fluorescence_quench_internalization_data_n87_ab1';
myPlotOptions.expDataTimeVar = 'time_day';
myPlotOptions.expDataYVar = 'internalized_fr';
plotHandle = plotAcrossIntervention(newWorksheet, myPlotOptions);
%
% Shed antigen concentrations
myPlotOptions = plotOptions;
myPlotOptions.flagSave = false;
myPlotOptions.xLabelPretty = 'Time (day)';
myPlotOptions.yLabelPretty = 'Shed Antigen, Plasma (nM)';
myPlotOptions.xLim = [0 22];
myPlotOptions.yLim = [];
myPlotOptions.xShiftSim = 10;
myPlotOptions.flagLegend = false;
myPlotOptions.varName = 'parrule_shed_total_plasma_concentration';
myPlotOptions.interventionID = 'buffer_injection_1200147_shed_assay';
myPlotOptions.expDataID = 'xenograft_data_1200147_n87_buffer_only';
myPlotOptions.expDataTimeVar = 'TIME_PLUS_ONE_DAY';
myPlotOptions.expDataYVar = 'SOLAG_NANOMOLEPL';
plotHandle = plotAcrossIntervention(newWorksheet, myPlotOptions);
%
% Shed antigen concentrations
myPlotOptions = plotOptions;
myPlotOptions.flagSave = false;
myPlotOptions.xLabelPretty = 'Time (day)';
myPlotOptions.yLabelPretty = 'Shed Antigen, Plasma (nM)';
myPlotOptions.xLim = [0 22];
myPlotOptions.yLim = [];
myPlotOptions.xShiftSim = 10;
myPlotOptions.flagLegend = false;
myPlotOptions.varName = 'parrule_shed_total_plasma_concentration';
myPlotOptions.interventionID = 'antibody_injection_1200147_shed_assay'; 
myPlotOptions.expDataID = 'xenograft_data_1200147_n87_ab1_only';
myPlotOptions.expDataTimeVar = 'TIME_PLUS_ONE_DAY';
myPlotOptions.expDataYVar = 'SOLAG_NANOMOLEPL';
plotHandle = plotAcrossIntervention(newWorksheet, myPlotOptions);
%
% Check the predictions for N87 PET xenografts with the variants in place
myPlotOptions = plotOptions;
myPlotOptions.flagSave = false;
myPlotOptions.xLabelPretty = 'Time (day)';
myPlotOptions.yLabelPretty = {'Tracer in xenograft','(nanomole/gram)'};
myPlotOptions.xLim = [0 17];
myPlotOptions.yLim = [];
myPlotOptions.xShiftSim = 10;
myPlotOptions.flagLegend = false;
myPlotOptions.varName = 'parrule_payload_total_apparent_pet_tumor_nanomole_per_gram';
myPlotOptions.interventionID = 'agxab1pet_injection';
myPlotOptions.expDataID = 'xenograft_data_95305073_n87';
myPlotOptions.expDataTimeVar = 'time_day';
myPlotOptions.expDataYVar = 'accumulation_N87_nmole_per_gram_tumor';
plotHandle = plotAcrossIntervention(newWorksheet, myPlotOptions);
%
% We can also plot the sampled axes coefficients
myPlotCoefficientsOptions = plotCoefficientsOptions;
myPlotCoefficientsOptions.flagSave = true;
plotCoefficients(newWorksheet,myPlotCoefficientsOptions);
% We can also get the VP coefficient matrix
vpCoeffs = getVPCoeffs(newWorksheet)

% Response types serve as a mechanism to score how well we agree with data.
% We add one here.
newWorksheet = createResponseType(newWorksheet, 'N87_agx');
% Now add response type elements to the response type.
% Full definition of response type element ID (RTEID) requires:
%                               id
%                               responseTypeID
%                               modelYVar
%                               modelYVarType
%                               interventionID
%                               expDataID
%                               expDataTimeVar
%                               expDataYVar
responseTypeElementDef = {'culture_internalized', 'N87_agx', 'parrule_payload_fraction_of_initial_bound_intracellular', 'parameter', 'culture_internalization', 'fluorescence_quench_internalization_data_n87_ab1', 'time_day', 'internalized_fr'};
newWorksheet = addResponseTypeElement('points', newWorksheet, responseTypeElementDef);
responseTypeElementDef = {'buffer_injection_1200147_shed_assay', 'N87_agx', 'parrule_shed_total_plasma_concentration', 'parameter', 'buffer_injection_1200147_shed_assay', 'xenograft_data_1200147_n87_buffer_only', 'TIME_PLUS_11_DAY', 'SOLAG_NANOMOLEPL'};
newWorksheet = addResponseTypeElement('points', newWorksheet, responseTypeElementDef);
responseTypeElementDef = {'antibody_injection_1200147_shed_assay', 'N87_agx', 'parrule_shed_total_plasma_concentration', 'parameter', 'antibody_injection_1200147_shed_assay', 'xenograft_data_1200147_n87_ab1_only', 'TIME_PLUS_11_DAY', 'SOLAG_NANOMOLEPL'};
newWorksheet = addResponseTypeElement('points', newWorksheet, responseTypeElementDef);
responseTypeElementDef = {'agxab1pet_injection', 'N87_agx', 'parrule_payload_total_apparent_pet_tumor_nanomole_per_gram', 'parameter', 'agxab1pet_injection', 'xenograft_data_95305073_n87', 'time_10delayed_day', 'accumulation_N87_nmole_per_gram_tumor'};
newWorksheet = addResponseTypeElement('points', newWorksheet, responseTypeElementDef);
% We can also check the response type to make sure we have paired
% everything correctly
myPassCheck = verifyResponseType(newWorksheet, 'N87_agx')
% You can view the response types
newWorksheet.responseTypes
newWorksheet.responseTypes{1}.elements{:}

% We can also copy the worksheet as a backup in case we want to change
% anything later.
% Also note copyWorksheet gives the option of picking out a subset of VPs 
% if the IDs are specified
bakWorksheet = copyWorksheet(newWorksheet);

% We can also save the worksheet to file for later use.  We can pick up here
% in example03, where we will demonstrate application of optimization 
% methods.
% Also note that worksheets can become rather large in memory,
% depending on how many variables we have in the model results, how
% many VPs, how many interventions, how often we sample the simulations,
% etc...
% We can also speed disk IO if we just want to keep the VPs and
% other things in the worksheet but not the results,
% especially if we can re-simulate quickly.
% We can remove results from the worksheet by:
% bakWorksheet.results = {};
saveWorksheet(bakWorksheet, 'example02Worksheet')

% We can also make sure that we set everything up correctly
verifyResponseType(newWorksheet, 'N87_agx')
% We score how well we match the response type in myResponseTypeResult
myResponseTypeResult = evaluateResponseType(newWorksheet, 'N87_agx');

% Check the VP scores/ objective function values
myResponseTypeResult.vpValues

% In addition to the randomization and scoring, we can just iteratively
% sample VPs.  bestNofM randomly tries VPs and keeps the best ones
help bestNofM
help bestNofMOptions

% Now try testing 300 and keeping the best 10
myBestNofMOptions = bestNofMOptions
myBestNofMOptions.baseVPID = 'n87_xenograft_scenario_0006';
myBestNofMOptions.responseTypeID = 'N87_agx';
myBestNofMOptions.keepN = 10;
myBestNofMOptions.tryM = 300;
myBestNofMOptions.intSeed=1;

tic
newWorksheet2 = bestNofM(newWorksheet, myBestNofMOptions)
toc
% We can also review the responsetype results
myResponseTypeResult2 = evaluateResponseType(newWorksheet2, 'N87_agx');


% Now re-plot
myPlotOptions = plotOptions;
myPlotOptions.flagSave = false;
myPlotOptions.xLabelPretty = 'Time (day)';
myPlotOptions.yLabelPretty = 'Internalized (fraction)';
myPlotOptions.xLim = [0 1];
myPlotOptions.xShiftSim = 0;
myPlotOptions.yLim = [];
myPlotOptions.flagLegend = false;
myPlotOptions.varName = 'parrule_payload_fraction_of_initial_bound_intracellular';
myPlotOptions.interventionID = 'culture_internalization'; 
myPlotOptions.expDataID = 'fluorescence_quench_internalization_data_n87_ab1';
myPlotOptions.expDataTimeVar = 'time_day';
myPlotOptions.expDataYVar = 'internalized_fr';
plotHandle = plotAcrossIntervention(newWorksheet2, myPlotOptions);
%
% Shed antigen concentrations
myPlotOptions = plotOptions;
myPlotOptions.flagSave = false;
myPlotOptions.xLabelPretty = 'Time (day)';
myPlotOptions.yLabelPretty = 'Shed Antigen, Plasma (nM)';
myPlotOptions.xLim = [0 22];
myPlotOptions.yLim = [];
myPlotOptions.xShiftSim = 10;
myPlotOptions.flagLegend = false;
myPlotOptions.varName = 'parrule_shed_total_plasma_concentration';
myPlotOptions.interventionID = 'buffer_injection_1200147_shed_assay'; 
myPlotOptions.expDataID = 'xenograft_data_1200147_n87_buffer_only';
myPlotOptions.expDataTimeVar = 'TIME_PLUS_ONE_DAY';
myPlotOptions.expDataYVar = 'SOLAG_NANOMOLEPL';
plotHandle = plotAcrossIntervention(newWorksheet2, myPlotOptions);
%
% Shed antigen concentrations
myPlotOptions = plotOptions;
myPlotOptions.flagSave = false;
myPlotOptions.xLabelPretty = 'Time (day)';
myPlotOptions.yLabelPretty = 'Shed Antigen, Plasma (nM)';
myPlotOptions.xLim = [0 22];
myPlotOptions.yLim = [];
myPlotOptions.xShiftSim = 10;
myPlotOptions.flagLegend = false;
myPlotOptions.varName = 'parrule_shed_total_plasma_concentration';
myPlotOptions.interventionID = 'antibody_injection_1200147_shed_assay'; 
myPlotOptions.expDataID = 'xenograft_data_1200147_n87_ab1_only';
myPlotOptions.expDataTimeVar = 'TIME_PLUS_ONE_DAY';
myPlotOptions.expDataYVar = 'SOLAG_NANOMOLEPL';
plotHandle = plotAcrossIntervention(newWorksheet2, myPlotOptions);
%
% Check the predictions for N87 PET xenografts with the variants in place
myPlotOptions = plotOptions;
myPlotOptions.flagSave = false;
myPlotOptions.xLabelPretty = 'Time (day)';
myPlotOptions.yLabelPretty = {'Tracer in xenograft','(nanomole/gram)'};
myPlotOptions.xLim = [0 17];
myPlotOptions.yLim = [0 0.08];
myPlotOptions.xShiftSim = 10;
myPlotOptions.flagLegend = false;
myPlotOptions.varName = 'parrule_payload_total_apparent_pet_tumor_nanomole_per_gram';
myPlotOptions.interventionID = 'agxab1pet_injection'; 
myPlotOptions.expDataID = 'xenograft_data_95305073_n87';
myPlotOptions.expDataTimeVar = 'time_day';
myPlotOptions.expDataYVar = 'accumulation_N87_nmole_per_gram_tumor';
plotHandle = plotAcrossIntervention(newWorksheet2, myPlotOptions);
%
```
