## Supplementary File 3 for "A surrogate workflow for virtual population development: designing an amenable proxy for a hypothesis-test inspired nonlinear goodness-of-fit": example03.m

```
% A slight modification from example02 to demonstrate
% using the optimization algorithms and response types.
% Note that after updating MATLAB from 2016a to newer versions, the
% output of the optimizations no longer exactly matched those
% presented in 
% Cheng Y, Thalhauser CJ, Smithline S, et al. QSP Toolbox: 
% Computational Implementation of Integrated Workflow Components for 
% Deploying Multi-Scale Mechanistic Models. The AAPS journal. 
% 2017;19(4):1002-1016.
%
% In addition to running in 2017a, the provided 
% SimBiology model format was updated to 2017a. 

%
% Toolbox initialization
% toolboxPath = 'DRIVE:/DIRs/QSPToolbox';
% addpath(toolboxPath);
initQSPToolbox;
% We start with the model from example02
myWorksheet = loadWorksheet('example02Worksheet')


% We keep 2 VPs from this worksheet for illustration of some of the
% optimization algorithms.
myVPIDs = getVPIDs(myWorksheet);
newWorksheet = copyWorksheet(myWorksheet, myVPIDs(1:2));


% Try optimizing each individual VP base and comparing to data
mySimulateOptions = simulateOptions;
mySimulateOptions.rerunExisting = true;
mySimulateOptions.responseTypeID = 'N87_agx';
mySimulateOptions.optimizeAxisIDs = getAxisDefIDs(newWorksheet);
% There are "cohort" approaches and per-VP approaches to the optimization.
% here, we try a pso-based per-VP approach where we run optimization once 
% for each VP, and we include each worksheet VP as a seed in each
% individual optimization run.
mySimulateOptions.optimizeType = 'pso';
mySimulateOptions.optimizePopSize = 100;
% It took about 2 minutes for about 30 iterations with 2xE5-2698v3 CPUs (Win7 64)
% Also using MinGW64 Compiler for acceleration steps.
% mySimulateOptions.optimizeTimeLimit = 60*2;
mySimulateOptions.optimizeMaxIter = 30;
mySimulateOptions.intSeed=1;
tic
newWorksheet = simulateWorksheet(newWorksheet, mySimulateOptions)
toc

% Evaluate the response type
newResponseTypeResult = evaluateResponseType(newWorksheet, 'N87_agx');

% Now plot.  Since the VP definition variants are the same for all VPs in
% this worksheet, it is likely they converge to similar values with a
% global optimization technique.
myPlotOptions = plotOptions;
myPlotOptions.flagSave = false;
myPlotOptions.xLabelPretty = 'Time (day)';
myPlotOptions.yLabelPretty = 'Internalized (fraction)';
myPlotOptions.xLim = [0 1];
myPlotOptions.yLim = [];
myPlotOptions.xShiftSim = 0;
myPlotOptions.flagLegend = false;
myPlotOptions.varName = 'parrule_payload_fraction_of_initial_bound_intracellular';
myPlotOptions.interventionID = 'culture_internalization'; 
myPlotOptions.expDataID = 'fluorescence_quench_internalization_data_n87_ab1';
myPlotOptions.expDataTimeVar = 'time_day';
myPlotOptions.expDataYVar = 'internalized_fr';
plotHandle = plotAcrossIntervention(newWorksheet, myPlotOptions);
%
% Shed antigen concentrations
myPlotOptions = plotOptions;
myPlotOptions.flagSave = false;
myPlotOptions.xLabelPretty = 'Time (day)';
myPlotOptions.yLabelPretty = 'Shed Antigen, Plasma (nM)';
myPlotOptions.xLim = [0 22];
myPlotOptions.yLim = [];
myPlotOptions.xShiftSim = 10;
myPlotOptions.flagLegend = false;
myPlotOptions.varName = 'parrule_shed_total_plasma_concentration';
myPlotOptions.interventionID = 'buffer_injection_1200147_shed_assay'; 
myPlotOptions.expDataID = 'xenograft_data_1200147_n87_buffer_only';
myPlotOptions.expDataTimeVar = 'TIME_PLUS_ONE_DAY';
myPlotOptions.expDataYVar = 'SOLAG_NANOMOLEPL';
plotHandle = plotAcrossIntervention(newWorksheet, myPlotOptions);
%
% Shed antigen concentrations
myPlotOptions = plotOptions;
myPlotOptions.flagSave = false;
myPlotOptions.xLabelPretty = 'Time (day)';
myPlotOptions.yLabelPretty = 'Shed Antigen, Plasma (nM)';
myPlotOptions.xLim = [0 22];
myPlotOptions.yLim = [];
myPlotOptions.xShiftSim = 10;
myPlotOptions.flagLegend = false;
myPlotOptions.varName = 'parrule_shed_total_plasma_concentration';
myPlotOptions.interventionID = 'antibody_injection_1200147_shed_assay'; 
myPlotOptions.expDataID = 'xenograft_data_1200147_n87_ab1_only';
myPlotOptions.expDataTimeVar = 'TIME_PLUS_ONE_DAY';
myPlotOptions.expDataYVar = 'SOLAG_NANOMOLEPL';
plotHandle = plotAcrossIntervention(newWorksheet, myPlotOptions);
%
% Check the predictions for N87 PET xenografts with the variants in place
myPlotOptions = plotOptions;
myPlotOptions.flagSave = false;
myPlotOptions.xLabelPretty = 'Time (day)';
myPlotOptions.yLabelPretty = {'Tracer in xenograft','(nanomole/gram)'};
myPlotOptions.xLim = [0 17];
myPlotOptions.yLim = [0 0.06];
myPlotOptions.xShiftSim = 10;
myPlotOptions.flagLegend = false;
myPlotOptions.varName = 'parrule_payload_total_apparent_pet_tumor_nanomole_per_gram';
myPlotOptions.interventionID = 'agxab1pet_injection'; 
myPlotOptions.expDataID = 'xenograft_data_95305073_n87';
myPlotOptions.expDataTimeVar = 'time_day';
myPlotOptions.expDataYVar = 'accumulation_N87_nmole_per_gram_tumor';
plotHandle = plotAcrossIntervention(newWorksheet, myPlotOptions);

% Note if we want to return multiple solutions by pooling existing VPs 
% as basis, if we use the cohort optimization like 'gacohort' optimization 
% type, but in this case all base VPs need to have the same variants in the
% definition and they can only vary with respect to the axis coefficients.
myVPIDs = getVPIDs(myWorksheet);
newWorksheet = copyWorksheet(myWorksheet, myVPIDs(1:2));
mySimulateOptions = simulateOptions;
mySimulateOptions.rerunExisting = true;
mySimulateOptions.responseTypeID = 'N87_agx';
mySimulateOptions.optimizeAxisIDs = getAxisDefIDs(newWorksheet);
mySimulateOptions.optimizeType = 'gacohort';
mySimulateOptions.optimizePopSize = 100;
% It took about 2 minutes for about 30 iterations on a 2xE5-2698v3 CPU (Win7 64)
% mySimulateOptions.optimizeTimeLimit = 60*2;
mySimulateOptions.optimizeMaxIter = 30;
mySimulateOptions.intSeed=1;

tic
newWorksheet = simulateWorksheet(newWorksheet, mySimulateOptions)
toc

% Note that some algorithms, like the genetic algorithm, do not select
% out duplicate solutions in their results.  We can filter this down a
% little more 
newWorksheet = removeDuplicateVPs(newWorksheet);
newResponseTypeResult = evaluateResponseType(newWorksheet, 'N87_agx');
% Note that the entire last generation is returned in the worksheet.
% We can pick the best VPs.
myVPvalues = newResponseTypeResult.vpValues;
myVPvalues = sort(myVPvalues);
% Let's take all VPs that ge to within a factor of 2 of the best
% objective function value
myIndices = find(newResponseTypeResult.vpValues <= myVPvalues(1)*2);
allVPIDs = getVPIDs(newWorksheet);
newWorksheet = copyWorksheet(newWorksheet,allVPIDs(myIndices));


% Now plot
myPlotOptions = plotOptions;
myPlotOptions.flagSave = false;
myPlotOptions.xLabelPretty = 'Time (day)';
myPlotOptions.yLabelPretty = 'Internalized (fraction)';
myPlotOptions.xLim = [0 1];
myPlotOptions.yLim = [];
myPlotOptions.xShiftSim = 0;
myPlotOptions.flagLegend = false;
myPlotOptions.varName = 'parrule_payload_fraction_of_initial_bound_intracellular';
myPlotOptions.interventionID = 'culture_internalization'; 
myPlotOptions.expDataID = 'fluorescence_quench_internalization_data_n87_ab1';
myPlotOptions.expDataTimeVar = 'time_day';
myPlotOptions.expDataYVar = 'internalized_fr';
plotHandle = plotAcrossIntervention(newWorksheet, myPlotOptions);
%
% Shed antigen concentrations
myPlotOptions = plotOptions;
myPlotOptions.flagSave = false;
myPlotOptions.xLabelPretty = 'Time (day)';
myPlotOptions.yLabelPretty = 'Shed Antigen, Plasma (nM)';
myPlotOptions.xLim = [0 22];
myPlotOptions.yLim = [];
myPlotOptions.xShiftSim = 10;
myPlotOptions.flagLegend = false;
myPlotOptions.varName = 'parrule_shed_total_plasma_concentration';
myPlotOptions.interventionID = 'buffer_injection_1200147_shed_assay'; 
myPlotOptions.expDataID = 'xenograft_data_1200147_n87_buffer_only';
myPlotOptions.expDataTimeVar = 'TIME_PLUS_ONE_DAY';
myPlotOptions.expDataYVar = 'SOLAG_NANOMOLEPL';
plotHandle = plotAcrossIntervention(newWorksheet, myPlotOptions);
%
% Shed antigen concentrations
myPlotOptions = plotOptions;
myPlotOptions.flagSave = false;
myPlotOptions.xLabelPretty = 'Time (day)';
myPlotOptions.yLabelPretty = 'Shed Antigen, Plasma (nM)';
myPlotOptions.xLim = [0 22];
myPlotOptions.yLim = [];
myPlotOptions.xShiftSim = 10;
myPlotOptions.flagLegend = false;
myPlotOptions.varName = 'parrule_shed_total_plasma_concentration';
myPlotOptions.interventionID = 'antibody_injection_1200147_shed_assay'; 
myPlotOptions.expDataID = 'xenograft_data_1200147_n87_ab1_only';
myPlotOptions.expDataTimeVar = 'TIME_PLUS_ONE_DAY';
myPlotOptions.expDataYVar = 'SOLAG_NANOMOLEPL';
plotHandle = plotAcrossIntervention(newWorksheet, myPlotOptions);
%
% Check the predictions for N87 PET xenografts with the variants in place
myPlotOptions = plotOptions;
myPlotOptions.flagSave = false;
myPlotOptions.xLabelPretty = 'Time (day)';
myPlotOptions.yLabelPretty = {'Tracer in xenograft','(nanomole/gram)'};
myPlotOptions.xLim = [0 17];
myPlotOptions.yLim = [0 0.06];
myPlotOptions.xShiftSim = 10;
myPlotOptions.flagLegend = false;
myPlotOptions.varName = 'parrule_payload_total_apparent_pet_tumor_nanomole_per_gram';
myPlotOptions.interventionID = 'agxab1pet_injection'; 
myPlotOptions.expDataID = 'xenograft_data_95305073_n87';
myPlotOptions.expDataTimeVar = 'time_day';
myPlotOptions.expDataYVar = 'accumulation_N87_nmole_per_gram_tumor';
plotHandle = plotAcrossIntervention(newWorksheet, myPlotOptions);

% Plot the axes coefficients
plotCoefficients(newWorksheet);
```
