## Supplementary File 3 for "A surrogate workflow for virtual population development: designing an amenable proxy for a hypothesis-test inspired nonlinear goodness-of-fit": example04.m

```
% Here we illustrate an iterative workflow for finding additional cohort
% VPs given an initial set of plausible VPs.
% Examples originally run with 2xE5-2698v3 CPUs (Win7 64)
% Also using MinGW64 Compiler for acceleration steps.
%
% Note that after updating the model from 2016a and also adjusting the tolerances,
% although they may be similar,
% the results may not exactly match 2016a results, especially following
% random sampling, such as those discussed in
% Cheng Y, Vezina HE, Gupta M, Pan C, Leil TA, Schmidt BJ. 
% Development of a Quantitative Systems Pharmacology (QSP)
% Toolbox and Virtual Population (VPop) for Affinity-Drug Conjugate (ADC) Research.
% Journal of pharmacokinetics and pharmacodynamics. 2016;43(1):S47.
% and
% Cheng Y, Thalhauser CJ, Smithline S, et al. QSP Toolbox: 
% Computational Implementation of Integrated Workflow Components for 
% Deploying Multi-Scale Mechanistic Models. The AAPS journal. 
% 2017;19(4):1002-1016.
%
% Toolbox initialization
% toolboxPath = 'DRIVE:/DIRs/QSPToolbox';
% addpath(toolboxPath);
initQSPToolbox;

% When we save large worksheets locally, often we set
% myWorksheet.results = {}, since the results
% may be quite large and MATLAB has issues with file IO
% > 2 GB
myWorksheet = loadWorksheet('example_cohort_worksheet1');
% First we repopulate the results.
myWorksheet = simulateWorksheet(myWorksheet);
% Now check to make sure all of the VPs make the plausibility cutoffs
myResponseSummaryTable = createResponseSummaryTable(myWorksheet, 'N87_agx');
myIndex1 = find(ismember(myResponseSummaryTable.rowNames,'culture_internalized'));
myIndices1 = find(myResponseSummaryTable.values(myIndex1,:)<=1.0);
myIndex2 = find(ismember(myResponseSummaryTable.rowNames,'buffer_injection_1200147_growth_assay'));
myIndices2 = find(myResponseSummaryTable.values(myIndex2,:)<=1.0);
myIndex3 = find(ismember(myResponseSummaryTable.rowNames,'buffer_injection_1200147_shed_assay'));
myIndices3 = find(myResponseSummaryTable.values(myIndex3,:)<=0.75);
myIndex4 = find(ismember(myResponseSummaryTable.rowNames,'antibody_injection_1200147_growth_assay'));
myIndices4 = find(myResponseSummaryTable.values(myIndex4,:)<=1.0);
myIndex5 = find(ismember(myResponseSummaryTable.rowNames,'antibody_injection_1200147_shed_assay'));
myIndices5 = find(myResponseSummaryTable.values(myIndex5,:)<=0.75);
myIndex6 = find(ismember(myResponseSummaryTable.rowNames,'agxab1pet_injection'));
myIndices6 = find(myResponseSummaryTable.values(myIndex6,:)<=1.0);
myIndices = intersect(intersect(intersect(intersect(intersect(myIndices1,myIndices2),myIndices3),myIndices4),myIndices5),myIndices6);
% How many VPs fulfill all?  This should be all 1,000, although
% it is possible a few fail due to differences that could be caused by
% issues like alterations to the solver or tolerance.  This changed
% a little in the update from 2016a to later releases.
length(myIndices)

% Check how many VPs total.
allVPIDs = getVPIDs(myWorksheet);
length(allVPIDs)
failVPIDs = setdiff(allVPIDs, allVPIDs(myIndices));
length(failVPIDs)
% Just in case a few fail the test and you wish to exclude, 
% you can use the following code
% to pick out the right ones.
% You can manually trigger recompilation with
% myWorksheet = compileModel(myWorksheet,true);
% It may also be triggered for othe reasons.
% You can also check the solver settings in:
% myWorksheet.simProps
% This is an example of code you can use to pick out the VPs that fail
% on the responseTypeElement cutoffs.
% failWorksheet = copyWorksheet(myWorksheet,failVPIDs);
% myWorksheet = copyWorksheet(myWorksheet,allVPIDs(myIndices));
% failResponseSummaryTable = createResponseSummaryTable(failWorksheet, 'N87_agx');
% % Doublecheck how many VPs total pass.
% allVPIDs = getVPIDs(myWorksheet);
% length(allVPIDs)

% Here are settings to update the cohort through the randomizations
% You can keep running for however long you would like to keep trying
% to generate diversity.
nRandomizations = 2;
% The worksheet will cap out at targetCohortSize.  Then we will cluster
% the VPs and start randomizing from the slsected VPs.
targetCohortSize = 1000;
% We'll limit how much the worksheet can tempoararily expand to in 
% iterations to save on memory.
maxWshVPs = 2500;
myVaryAxesOptions = varyAxesOptions;
% We'll vary most of the new VP axes with a gaussian, but also
% pick one axis to draw from a uniform distribution to speed up the
% sampling
myVaryAxesOptions.varyMethod = 'gaussian';
myVaryAxesOptions.gaussianStd = 0.05;
myVaryAxesOptions.varyAxisIDs = getAxisDefIDs(myWorksheet);


tic;
% We'll also backup the worksheet to file every so many randomizations.
lastBackCounter = -10;
for rCounter = 1 : nRandomizations
    % Reseed the rng with the counter before randomizing here
    rng(rCounter);
    [nAxis, nVPs] = size(getVPCoeffs(myWorksheet)); 
    % We can select VPs by clustering if there are too many
    % so we can generate new ones.  Note if we reach targetCohortSize,
    % it is expected a save would be tiggered in the previous run-through.
    % If we have trouble generating specific phenotypes, we could set  
    % up a biased sampling here as well but that isn't demonstrated here.
    if nVPs >= targetCohortSize
        myClusterPickOptions = clusterPickOptions;
        myClusterPickOptions = myClusterPickOptions.setDefaultFromWorksheet(myWorksheet);
        myClusterPickOptions.normalizeType = 'min-max';
        myClusterPickOptions.nClusters = round(targetCohortSize*.3);
        myClusterPickOptions.edgeVPFlag = true;
        myClusterPickOptions.intSeed = -1;
        myMedoidResult = pickClusterVPs(myWorksheet,myClusterPickOptions);
        myVPIDs = myMedoidResult.('pickedVPIDs');
        myWorksheet = copyWorksheet(myWorksheet, myVPIDs);
    end   
    
    myVaryAxesOptions.additionalIDString = num2str(rCounter);
    [nAxis, nPreviousVPs] = size(getVPCoeffs(myWorksheet));
    previousVPIDs = getVPIDs(myWorksheet); 
    myVaryAxesOptions.baseVPIDs = previousVPIDs;
    myVaryAxesOptions.newPerOld = max(floor(maxWshVPs-nPreviousVPs)/length(myVaryAxesOptions.baseVPIDs),1);
    % First we add the VPIDs
    myWorksheet = addVariedVPs(myWorksheet, myVaryAxesOptions);    
    allVPIDs = getVPIDs(myWorksheet);
    [nAxis, nVPs] = size(getVPCoeffs(myWorksheet));     
    for vpCounter = 1 : nVPs
        curVPID = allVPIDs{vpCounter};
        % We want to avoid overwriting coefficients for
        % the old best VPs in each iteration
        if sum(ismember(previousVPIDs,curVPID)) < 1
            % We'll randomly pick an axis to draw from a uniform normal
            % to speed the sampling
            axisIndex = randsample([1:nAxis],1);
            myWorksheet.axisProps.axisVP.coefficients(axisIndex,vpCounter) = rand(1);
        end
    end    
       
    % Now we run the simulations for the altered variables
    mySimulateOptions = simulateOptions;
    mySimulateOptions.rerunExisting = false;
    mySimulateOptions.optimizeType = 'none';
    myWorksheet = simulateWorksheet(myWorksheet);
    
    % Now we will pick out the VPs that satisfy our constraints on the
    % response type / response type elements objective function ("feasible" VPs)
    myResponseSummaryTable = createResponseSummaryTable(myWorksheet, 'N87_agx');
    myIndex1 = find(ismember(myResponseSummaryTable.rowNames,'culture_internalized'));
    myIndices1 = find(myResponseSummaryTable.values(myIndex1,:)<=1.0);
    myIndex2 = find(ismember(myResponseSummaryTable.rowNames,'buffer_injection_1200147_growth_assay'));
    myIndices2 = find(myResponseSummaryTable.values(myIndex2,:)<=1.0);
    myIndex3 = find(ismember(myResponseSummaryTable.rowNames,'buffer_injection_1200147_shed_assay'));
    myIndices3 = find(myResponseSummaryTable.values(myIndex3,:)<=0.75);
    myIndex4 = find(ismember(myResponseSummaryTable.rowNames,'antibody_injection_1200147_growth_assay'));
    myIndices4 = find(myResponseSummaryTable.values(myIndex4,:)<=1.0);
    myIndex5 = find(ismember(myResponseSummaryTable.rowNames,'antibody_injection_1200147_shed_assay'));
    myIndices5 = find(myResponseSummaryTable.values(myIndex5,:)<=0.75);
    myIndex6 = find(ismember(myResponseSummaryTable.rowNames,'agxab1pet_injection'));
    myIndices6 = find(myResponseSummaryTable.values(myIndex6,:)<=1.0);
    myIndices = intersect(intersect(intersect(intersect(intersect(myIndices1,myIndices2),myIndices3),myIndices4),myIndices5),myIndices6);
    length(myIndices)
    allVPIDs = getVPIDs(myWorksheet);
    length(allVPIDs)
    
    myVPIDs = allVPIDs(myIndices);
    myWorksheet = copyWorksheet(myWorksheet, myVPIDs);
    length(myVPIDs)
    
    % If the worksheet grows bigger than the target size, we select VPs
    % based on clustering to bring the worksheet back down to
    % target size before writing to file
    if length(myVPIDs) > targetCohortSize
        myClusterPickOptions = clusterPickOptions;
        myClusterPickOptions = myClusterPickOptions.setDefaultFromWorksheet(myWorksheet);
        myClusterPickOptions.normalizeType = 'min-max';
        myClusterPickOptions.nClusters = targetCohortSize;
        myClusterPickOptions.edgeVPFlag = true;
        myClusterPickOptions.intSeed = -1;
        rng(rCounter);
        myMedoidResult = pickClusterVPs(myWorksheet,myClusterPickOptions);
        myVPIDs = myMedoidResult.('pickedVPIDs');
        myWorksheet = copyWorksheet(myWorksheet, myVPIDs);
    end   
    rCounter
    toc/60
    if (((rCounter - lastBackCounter) >= 1) && (length(myVPIDs) == targetCohortSize))
        lastBackCounter = rCounter;
        tempWorksheet = myWorksheet;
        tempWorksheet.results = {};
        saveWorksheet(tempWorksheet, 'example04_test');
    end
end

myPlotOptions = plotOptions;
myPlotOptions.flagSave = false;
myPlotOptions.xLabelPretty = 'Time (day)';
myPlotOptions.yLabelPretty = 'Internalized (fraction)';
myPlotOptions.xLim = [0 1];
myPlotOptions.xShiftSim = 0;
myPlotOptions.flagLegend = false;
myPlotOptions.varName = 'parrule_payload_fraction_of_initial_bound_intracellular';
myPlotOptions.interventionID = 'culture_internalization'; 
myPlotOptions.expDataID = 'fluorescence_quench_internalization_data_n87_ab1';
myPlotOptions.expDataTimeVar = 'time_day';
myPlotOptions.expDataYVar = 'internalized_fr';
plotHandle = plotAcrossIntervention(myWorksheet, myPlotOptions);

% lesion growth
myPlotOptions = plotOptions;
myPlotOptions.flagSave = false;
myPlotOptions.xLabelPretty = 'Time (day)';
myPlotOptions.yLabelPretty = 'Tumor Volume (L)';
myPlotOptions.xLim = [0 22];
myPlotOptions.xShiftSim = 10;
myPlotOptions.flagLegend = false; 
myPlotOptions.varName = 'parrule_tumor_volume';
myPlotOptions.interventionID = 'buffer_injection_1200147_shed_assay';
myPlotOptions.expDataID = 'xenograft_data_1200147_n87_buffer_only';
myPlotOptions.expDataTimeVar = 'TIME_PLUS_ONE_DAY';
myPlotOptions.expDataYVar = 'TUMOR_VOLUME_L';
plotHandle = plotAcrossIntervention(myWorksheet, myPlotOptions);

%
% Shed antigen concentrations
myPlotOptions = plotOptions;
myPlotOptions.flagSave = false;
myPlotOptions.xLabelPretty = 'Time (day)';
myPlotOptions.yLabelPretty = 'Shed Antigen, Plasma (nM)';
myPlotOptions.xLim = [0 22];
myPlotOptions.xShiftSim = 10;
myPlotOptions.flagLegend = false;
myPlotOptions.varName = 'parrule_shed_total_plasma_concentration';
myPlotOptions.interventionID = 'buffer_injection_1200147_shed_assay'; 
myPlotOptions.expDataID = 'xenograft_data_1200147_n87_buffer_only';
myPlotOptions.expDataTimeVar = 'TIME_PLUS_ONE_DAY';
myPlotOptions.expDataYVar = 'SOLAG_NANOMOLEPL';
plotHandle = plotAcrossIntervention(myWorksheet, myPlotOptions);

%
myPlotOptions = plotOptions;
myPlotOptions.flagSave = false;
myPlotOptions.xLabelPretty = 'Time (day)';
myPlotOptions.yLabelPretty = 'Tumor Volume (L)';
myPlotOptions.xLim = [0 22];
myPlotOptions.xShiftSim = 10;
myPlotOptions.flagLegend = false; 
myPlotOptions.varName = 'parrule_tumor_volume';
myPlotOptions.interventionID = 'antibody_injection_1200147_shed_assay';
myPlotOptions.expDataID = 'xenograft_data_1200147_n87_ab1_only';
myPlotOptions.expDataTimeVar = 'TIME_PLUS_ONE_DAY';
myPlotOptions.expDataYVar = 'TUMOR_VOLUME_L';
plotHandle = plotAcrossIntervention(myWorksheet, myPlotOptions);


myPlotOptions = plotOptions;
myPlotOptions.flagSave = false;
myPlotOptions.xLabelPretty = 'Time (day)';
myPlotOptions.yLabelPretty = 'Shed Antigen, Plasma (nM)';
myPlotOptions.xLim = [0 22];
myPlotOptions.xShiftSim = 10;
myPlotOptions.flagLegend = false;
myPlotOptions.varName = 'parrule_shed_total_plasma_concentration';
myPlotOptions.interventionID = 'antibody_injection_1200147_shed_assay'; 
myPlotOptions.expDataID = 'xenograft_data_1200147_n87_ab1_only';
myPlotOptions.expDataTimeVar = 'TIME_PLUS_ONE_DAY';
myPlotOptions.expDataYVar = 'SOLAG_NANOMOLEPL';
plotHandle = plotAcrossIntervention(myWorksheet, myPlotOptions);
%
myPlotOptions.xLabelPretty = 'Time (day)';
myPlotOptions.flagSave = false;
myPlotOptions.yLabelPretty = {'Tracer in xenograft','(nanomole/gram)'};
myPlotOptions.xLim = [0 17];
myPlotOptions.xShiftSim = 10;
myPlotOptions.flagLegend = false;
myPlotOptions.varName = 'parrule_payload_total_apparent_pet_tumor_nanomole_per_gram';
myPlotOptions.interventionID = 'agxab1pet_injection'; 
myPlotOptions.expDataID = 'xenograft_data_95305073_n87';
myPlotOptions.expDataTimeVar = 'time_day';
myPlotOptions.expDataYVar = 'accumulation_N87_nmole_per_gram_tumor';
plotHandle = plotAcrossIntervention(myWorksheet, myPlotOptions);

myPlotCoefficientsOptions = plotCoefficientsOptions;
myPlotCoefficientsOptions.fontSize = 12;
plotCoefficients(myWorksheet,myPlotCoefficientsOptions);


% Since the 2017 AAPS paper, resampling steps were automized more in
% the expandVPopEffN function.  However, the resampling is guided by 
% calibration targets.  This requires first making a valid mapelOptions
% object, which is explained in later example06.  
```
