## Supplementary File 3 for "A surrogate workflow for virtual population development: designing an amenable proxy for a hypothesis-test inspired nonlinear goodness-of-fit": example06.m

```
% Here, we illustrate workflow for creating VPops using MAPEL

% Toolbox initialization
% toolboxPath = 'DRIVE:/DIRs/QSPToolbox';
% addpath(toolboxPath);
initQSPToolbox;

%% load in VP cohort: existing worksheet.
% Load in the worksheet and regenerate results.
myWorksheetFileName = 'example_cohort_worksheet1'; % check this RT?
myWorksheet = loadWorksheet(myWorksheetFileName);
mySimulateOptions = simulateOptions();
mySimulateOptions.rerunExisting = true; 

% add in target time point to ensure we store the simulation results at the correct time points
myMapelOptions = mapelOptions;
myMapelOptions.expData = convertResponseTypeToExpDataTable(myWorksheet, 'N87_agx', 'USUBJID');
myWorksheet.simProps.sampleTimes = sort(unique([0:.025:35,myMapelOptions.expData.time'])','ascend');

% record only target biomarkers to save storage space
myWorksheet.simProps.saveElementResultIDs={'parrule_payload_fraction_of_initial_bound_intracellular'
    'parrule_tumor_volume'
    'parrule_shed_total_plasma_concentration'
    'parrule_payload_total_apparent_pet_tumor_nanomole_per_gram'};
myWorksheet = simulateWorksheet(myWorksheet, mySimulateOptions);

saveWorksheetAutoSplit(myWorksheet,'example_cohort_worksheet_SIM');

%% step 2: set up mapelOption, run one mapel, which is prevelance weighting
% Now we create a mapelOptions structure
help mapelOptions
myMapelOptions = mapelOptions;

% First we need to calculate the summary statistics.  In the published
% MAPEL version, we matched mean, standard deviation, and binned data.
myMapelOptions.expData = convertResponseTypeToExpDataTable(myWorksheet, 'N87_agx', 'USUBJID');
myMapelOptions.mnSDTable = convertExpDataToMnSDTable(myMapelOptions);
% We create a default subpopTable for 'all', since there are noise
% subpopulations of special interest for this calibration.
myMapelOptions.subpopTable=createSubpopTable(myWorksheet);
% Note that this will automatically generate the bin table for now.
% Sometimes, we may want to create our own bin tables rather than using the
% default bin edges.
% Note the bin functions have changes since Cheng et al. AAPS J 2017,
% so bin edges may vary slightly from the available historical
% VPOP used later in this example.
myMapelOptions.pwStrategy = 'bin';
myMapelOptions.binTable = convertExpDataToBinTable(myMapelOptions);
[nAxis,nVP] = size(myWorksheet.axisProps.axisVP);
% nVPs is the largest total number of bins we should create.
% We need at least 1 VP per bin
% or we risk assigning weight to areas
% not represented by VPs.  This sets the number of bins on a
% per-axis basis, so there are a total 4 * 13 total population bins
myMapelOptions.nBins = 4;
% We set additional parameters for the optimization
myMapelOptions.optimizeType = 'pso';
% We usually don't need very small tolerances to find good solutions, which
% we can assess by the GOF statistic and effN.  The objective is on a
% logarithmic scale.  The optimization may time out before convergence,
% we need to check the GOF to see if the solution is acceptable in terms
% of agreement with data.  Multiple VPop solutions may be acceptable.
myMapelOptions.tol = 1E-2;
myMapelOptions.randomStart = true;
myMapelOptions.spreadOut = 1;
myMapelOptions.minEffN = 30;
myMapelOptions.optimizePopSize = 1000;

% We may want to run a short optimization to create a VPop object
% just to see how well our experimental and simulated data ranges agree
myMapelOptions.optimizeTimeLimit = 10;
myVPop = mapel(myWorksheet, myMapelOptions);

%% diagnostic plot to identify problematic data points . could the tumor bl be unweighted here?
% This diagnostic plot for the population doesn't require a VPop solution, 
% but it does require
% the VPop have both the experimental and simulation data populated.
% ExpData is usually populated manually, but simData is usually populated 
% by the getSimData method of the VPop in the call to mapel.
help plotSimExpDataVPop
plotSimExpDataVPop(myVPop)
% Outputs/interventions/times where the red circles do not cover the
% ranges of the blue triangles well suggest there is not good model/cohort
% coverage and the population fitting have issues.  You need to assess
% whether this means the cohort need to be expanded, whether this is an
% issue with the data (for example, if there is non-mechanistic measurement
% noise in the data that we don't capture), or if there could be an issue
% with the model that we need to fix, discuss with experimenters, or
% accept.  If we decide we want to accept an issue and we do not want it to
% influence fits, we need to set the corresponding weight to 0.


% We had identified several time/experiment datapoints that would be 
% difficult to match due to issues in the experimental data or model 
% behaviors.  We can set the influence of these on the optimization to 0.
myMapelOptions.mnSDTable{1,'weightSD'}=0;
myMapelOptions.mnSDTable{1,'weightMean'}=0;
myMapelOptions.binTable{1,'weight'}=0;

myMapelOptions.mnSDTable{2,'weightSD'}=0;
myMapelOptions.mnSDTable{2,'weightMean'}=0;
myMapelOptions.binTable{2,'weight'}=0;

myMapelOptions.mnSDTable{3,'weightSD'}=0;
myMapelOptions.mnSDTable{3,'weightMean'}=0;
myMapelOptions.binTable{3,'weight'}=0;

myMapelOptions.mnSDTable{26,'weightSD'}=0;
myMapelOptions.mnSDTable{26,'weightMean'}=0;
myMapelOptions.binTable{26,'weight'}=0;

myMapelOptions.mnSDTable{27,'weightSD'}=0;
myMapelOptions.mnSDTable{27,'weightMean'}=0;
myMapelOptions.binTable{27,'weight'}=0;

myMapelOptions.mnSDTable{28,'weightSD'}=0;
myMapelOptions.mnSDTable{28,'weightMean'}=0;
myMapelOptions.binTable{28,'weight'}=0;

%% re-optimization - do mapel  a few times (in case of random seed)
% Convergence times may vary between systems
myMapelOptions.optimizeTimeLimit = 2*60*60;
% We can iterate through a few times in case we have a bad seed
% population in the fitting.
for myTestCounter = 1 : 1
        mySaveName = ['example06_',num2str(myTestCounter)];
        myMapelOptions.intSeed = myTestCounter;
        myVPop = mapel(myWorksheet, myMapelOptions); 
        saveVPop(myVPop,mySaveName);
        % We can also display the summary statistic with useEffN
        myVPop.useEffN = true;
        myVPop = evaluateGOF(myVPop);        
        myVPop.mnSDTable{1,'predN'}    
        myVPop.gof
end

%% additional diagnostic plot of expData and Vpop simulation to identify problematic data points. could the tumor bl be unweighted here?
% This is an additional diagnostic table for the places where we have
% experimental and simulation data.  Note that these can become
% rather crowded when we have a lot of distinct
% outputs/interventions/times and don't exclude non-weighted times, but
% can be generally useful to get a quick sense of issues with the
% population fitting.
myPlotOptions = plotOptions;
% We want to include the mn/sd plots
myPlotOptions.flagPlotUnweighted = true;
plotMnSDVPop(myVPop, myPlotOptions);

% There is also a quick plot to visualize how the prevalence weights
% are distributed in the VPop
plotPWHist(myVPop)

%% restartMapel from a preexisting VPop, add random noise
% Note that we may find a good VPop and may want to use this as a seed for
% additional VPops, or we may want to pick up with a VPop fitting
% run that was stopped early.
% For example, we can start from a previous VPop as an initial guess,
% and optionally add noise to the initial probabilities
myVPop = loadVPop('example_vpop');
% Set this to 0 if we want to use the exact same starting point
myRandomStart = 0.01;
for myTestCounter = 1 : 1
    myVPop.intSeed = myTestCounter;
    myTestCounter
    newVPop = restartMapel(myVPop, myRandomStart);
    mySaveName = ['example06_rerun_',num2str(myTestCounter)];
    saveVPop(newVPop,mySaveName);
    newVPop.useEffN = true;
    newVPop = evaluateGOF(newVPop);
    newVPop.mnSDTable{1,'predN'}
    newVPop.gof
end

%% above ones are all run on 'bin' mapel strategy ...
%% from this one on it is 'direct'

%% restartMapel 'direct' on the existing vpop
% These additional sections were not in the AAPS 2017 publication but
% highlight additional algorithms implemented since then.  For the first
% example, we run "MAPEL" without imposing the binned axis
% distributions and allowing the prevalence weights to vary
myVPop = loadVPop('example_vpop');
myVPop.pwStrategy = 'direct';
myVPop.optimizeType = 'gapso';
newVPop = restartMapel(myVPop);
newVPop.useEffN = true;
newVPop = evaluateGOF(newVPop);
newVPop.mnSDTable{1,'predN'}
newVPop.gof


%% linearCalibration module
% This is a sample run with the linearCalibration module
% Set up the optimization options, which will automatically spit back a set
% of default values

myOptimOptions = LinearCalibrationOptions();
% Define the probabilities of the cumulative
% distribution functions (CDF) to fit. The default is to fit all points on
% the CDF. We will only fit certain probabilities
% along the CDF in order to make the fit faster. These probabilities are
% defined here:
myOptimOptions.cdfProbsToFit = 0.05:0.05:0.95;
myOptimOptions.pdf2DProbsToFitN = 5;
myOptimOptions.responseValTransformation='none';
% multiple algorithms are implemented, we chose 'quadproEffN' for demonstration
% this is also the algorithm we use in the iterative vpop development workflow
myOptimOptions.optimizationAlgorithm = "quadprogEffN";				
myOptimOptions.priorPrevalenceWeightAssumption = 'specified';
myOptimOptions.oldVPop = myVPop;  

% Initialize a 'LinearCalibration' object:
linearCalibrationObject = LinearCalibration(myVPop,'optimOptions',myOptimOptions);
% Run the optimization:
linearCalibrationObject = linearCalibrationObject.run();
% Update VPop:
myInitialPWs = linearCalibrationObject.OptimizationResults.optimalPrevalenceWeightsNormalized';
myVPop.pws = myInitialPWs(1,:);

% Update table values.  Not strictly necessary but a nice
% step to diagnose and does have much computational cost            
newVPop = myVPop.addPredTableVals();  
% Next update the individual GOF statistics
newVPop.useEffN = true;
newVPop = evaluateGOF(newVPop);
newVPop.mnSDTable{1,'predN'}
newVPop.gof


%% Here, we demonstrate our original VPop developement workflow with expandVPopEffN with particle swarm optimization
% A automatic VPop resampling and reweighting workflow to generate 
% a well calibrated VPop (p-value>0.9) using wrapper function expandVPopEffN
myVPop = loadVPop('example_vpop');

% We had identified several time/experiment datapoints that would be 
% difficult to match and set the influence of these on the optimization to 0.
myMapelOptions.mnSDTable{9,'weightSD'}=0;
myMapelOptions.mnSDTable{9,'weightMean'}=0;
myMapelOptions.binTable{9,'weight'}=0;

myMapelOptions.mnSDTable{20,'weightSD'}=0;
myMapelOptions.mnSDTable{20,'weightMean'}=0;
myMapelOptions.binTable{20,'weight'}=0;

% update the VPop, add expDataID to check for the calibration target, for use in linearCalibration and expandVPopEffN
myVPop=getSimData(myVPop,myWorksheet)
myVPop=myVPop.addTableSimVals
myVPop=myVPop.addPredTableVals

myVPop.pwStrategy = 'direct';
myVPop.optimizeType = 'gapso';
myMapelOptions = initializeVPopPropertiesToOption(myVPop);
myExpandVPopEffNOptions = expandVPopEffNOptions;
% An identifying suffix
myExpandVPopEffNOptions.suffix = 'example06OriginalWf';
myExpandVPopEffNOptions.wsIterCounter = 0;
% Normally target 150 and often pause at 50
% in initial run.  We'll stop here too in order to
% adjust expansion sized
myExpandVPopEffNOptions.targetEffN = 40;
myMapelOptions.minEffN = 0;
myExpandVPopEffNOptions.maxNewPerIter= 10;
myExpandVPopEffNOptions.expandCohortSize = 5000;
myExpandVPopEffNOptions.effNDelta = 2;
myExpandVPopEffNOptions.minPVal = 0.9;
myExpandVPopEffNOptions.nTries = 1;
myExpandVPopEffNOptions.nRetries = 1;
myExpandVPopEffNOptions.verbose = true;
myExpandVPopEffNOptions.restartPVal = 1E-4;
myExpandVPopEffNOptions.expandRandomStart = 0;
myExpandVPopEffNOptions.varyMethod = 'gaussian';
myExpandVPopEffNOptions.resampleStd = 0.05;
myExpandVPopEffNOptions.expandEdgeVPs=true;
myExpandVPopEffNOptions.maxNewPerOld = 3;
% Here, we enforce any checks on sampled VPs,
% before simulation and adjust VPs if needed.
myExpandVPopEffNOptions.screenFunctionName = '';
myExpandVPopEffNOptions.nCluster = 400;

% To run the original workflow: make sure set linearExpandFlag=false
myExpandVPopEffNOptions.linearExpandFlag = false;
myExpandVPopEffNOptions.minEffNlinearflag = false;


% This takes a while to run.  Every iteration will
% write to file when a better VPop is found
% and a new worksheet will write to file
% when new VPs are added.
% You can delete intermediate iterations between your start file and
% stopping point.  They are included to allow you to more easily
% track progress.
[newWorksheet, newVPop] = expandVPopEffN(myWorksheet,myExpandVPopEffNOptions,myMapelOptions,myVPop);

%% We also demonstrate our new surrogate VPop developement workflow with expandVPopEffN with linearCalibration
% A automatic VPop resampling and reweighting workflow to generate 
% a well calibrated VPop (p-value>0.9) using wrapper function expandVPopEffN
myVPop = loadVPop('example_vpop');

% We had identified several time/experiment datapoints that would be 
% difficult to match and set the influence of these on the optimization to 0.
myMapelOptions.mnSDTable{9,'weightSD'}=0;
myMapelOptions.mnSDTable{9,'weightMean'}=0;
myMapelOptions.binTable{9,'weight'}=0;

myMapelOptions.mnSDTable{20,'weightSD'}=0;
myMapelOptions.mnSDTable{20,'weightMean'}=0;
myMapelOptions.binTable{20,'weight'}=0;

% update the VPop, add expDataID to check for the calibration target, for use in linearCalibration and expandVPopEffN
myVPop=getSimData(myVPop,myWorksheet)
myVPop=myVPop.addTableSimVals
myVPop=myVPop.addPredTableVals

myVPop.pwStrategy = 'direct';
myVPop.optimizeType = 'gapso';
myMapelOptions = initializeVPopPropertiesToOption(myVPop);
myExpandVPopEffNOptions = expandVPopEffNOptions;
% An identifying suffix
myExpandVPopEffNOptions.suffix = 'example06SurrogateWf';
myExpandVPopEffNOptions.wsIterCounter = 0;
% Normally target 150 and often pause at 50
% in initial run.  We'll stop here too in order to
% adjust expansion sized
myExpandVPopEffNOptions.targetEffN = 40;
myMapelOptions.minEffN = 0;
myExpandVPopEffNOptions.maxNewPerIter= 10;
myExpandVPopEffNOptions.expandCohortSize = 5000;
myExpandVPopEffNOptions.effNDelta = 2;
myExpandVPopEffNOptions.minPVal = 0.9;
myExpandVPopEffNOptions.nTries = 1;
myExpandVPopEffNOptions.nRetries = 1;
myExpandVPopEffNOptions.verbose = true;
myExpandVPopEffNOptions.restartPVal = 1E-4;
myExpandVPopEffNOptions.expandRandomStart = 0;
myExpandVPopEffNOptions.varyMethod = 'gaussian';
myExpandVPopEffNOptions.resampleStd = 0.05;
myExpandVPopEffNOptions.expandEdgeVPs=true;
myExpandVPopEffNOptions.maxNewPerOld = 3;
% Here, we enforce any checks on sampled VPs,
% before simulation and adjust VPs if needed.
myExpandVPopEffNOptions.screenFunctionName = '';
myExpandVPopEffNOptions.nCluster = 400;

% all above seetings are the same as the original workflow 
% Different settings to run the surrogate workflow: make sure set linearExpandFlag=true
myExpandVPopEffNOptions.linearExpandFlag = true;
myExpandVPopEffNOptions.minEffNlinearflag = true;
% also set cohortSize and maxIter if running this workflow
myExpandVPopEffNOptions.linearExpandCohortSize = 5000;
myExpandVPopEffNOptions.maxIterlinearExpand = 20;
myExpandVPopEffNOptions.nVPMax = 1000;
myExpandVPopEffNOptions.minPVallinear = 0.1;


% This takes a while to run.  Every iteration will
% write to file when a better VPop is found
% and a new worksheet will write to file
% when new VPs are added.
% You can delete intermediate iterations between your start file and
% stopping point.  They are included to allow you to more easily
% track progress.
[newWorksheet, newVPop] = expandVPopEffN(myWorksheet,myExpandVPopEffNOptions,myMapelOptions,myVPop);
```
