## Supplementary File 3 for "A surrogate workflow for virtual population development: designing an amenable proxy for a hypothesis-test inspired nonlinear goodness-of-fit": example07.m

```
% We illustrate how to run a Sobol global sensitivity 
% analysis.  Sobol's g function is used as an example here.
% Sobol's g function test was also used in 
% Saltelli, A., Making best use of model evaluations to compute
% sensitivity indices. Computer Physics Communications, 2002.
% See Fig 2 a,b for Saltelli's results.
%
% Toolbox initialization
% toolboxPath = 'DRIVE:/DIRs/QSPToolbox';
% addpath(toolboxPath);
initQSPToolbox;
% Read in the model
myWorksheet = readQSPModel('gfunction_test.sbproj');
% Create the VP (column) definitions
myWorksheet = createVPs(myWorksheet, {'gVP'},{{'ai_values___default';'xi_values___center'}},'');
myWorksheet = createIntervention(myWorksheet, 'calculate_g',cell(0,2));

% Add mechanistic axes definitions to the worksheet
myWorksheet = addAxisDef(myWorksheet,'axisX1',{'par_x1','parameter', [0, 1]});
myWorksheet = addAxisDef(myWorksheet,'axisX2',{'par_x2','parameter', [0, 1]});
myWorksheet = addAxisDef(myWorksheet,'axisX3',{'par_x3','parameter', [0, 1]});
myWorksheet = addAxisDef(myWorksheet,'axisX4',{'par_x4','parameter', [0, 1]});
myWorksheet = addAxisDef(myWorksheet,'axisX5',{'par_x5','parameter', [0, 1]});
myWorksheet = addAxisDef(myWorksheet,'axisX6',{'par_x6','parameter', [0, 1]});

% This is just a test of the simulation.
myWorksheet.simProps.sampleInterval = .1;
myWorksheet.simProps.simTime = 1;
myWorksheet = simulateWorksheet(myWorksheet);

% Sobol's g function is not an ODE and the variables
% are not necessarily time,
% but just to demonstrate the function in
% SimBiology and with toolbox worksheets, we
% have to implement this as an output of a dynamical model

mySobolSampleOptions = sobolSampleOptions;
allVPIDs = getVPIDs(myWorksheet);
allInterventionIDs = getInterventionIDs(myWorksheet);
mySobolSampleOptions.baseVPID = allVPIDs{1};
mySobolSampleOptions.interventionID = 'calculate_g';
mySobolSampleOptions.varyAxisIDs = getAxisDefIDs(myWorksheet);
mySobolSampleOptions.nRandomizationsPerSample = 1024;
mySobolSampleOptions.maxBatchSimulateN = 10000*10;
mySobolSampleOptions.saveElementResultIDs = {'parrule_gtotal'};
mySobolSampleOptions.saveFileName = '';
mySobolSampleOptions.intSeed = 0;
mySampledWorksheet = runSobolSample(myWorksheet,mySobolSampleOptions);

mySobolSensitivityOptions = sobolSensitivityOptions;
mySobolSensitivityOptions.nBootstraps = 10000;
mySobolSensitivityOptions.interventionID = mySobolSampleOptions.interventionID;
mySobolSensitivityOptions.analyzeElementResultIDs = {'parrule_gtotal'};
mySobolSensitivityOptions.subSampleSplitType = 'base2';
%mySobolSensitivityOptions.subSampleSplitType = 'none'
mySobolSensitivityOptions.analyzeTime = 1;
mySobolSensitivityOptions.intSeed=1;
mySobolResults = runSobolSensitivity(mySampledWorksheet, mySobolSensitivityOptions);

% Quick visual predictive check of the accuracy of the Si values.
% These are the ai values used in Saltelli 2002.
[analyticalSi,analyticalSTi] = calculateGFunctionSi([0;.5;3;9;99;99]);

nParams = length(mySobolResults.axisDefIDs);
nSplits = length(mySobolResults.sampleSize);

% Plot the sum of the first order sensitivity indices results as a
% function of the subsample split size
figure;
plotHandle = errorbar(mySobolResults.sampleSize,mySobolResults.('parrule_gtotal').firstOrderMedian(:,nParams+1),mySobolResults.('parrule_gtotal').firstOrderUpperCI(:,nParams+1)-mySobolResults.('parrule_gtotal').firstOrderMedian(:,nParams+1),mySobolResults.('parrule_gtotal').firstOrderMedian(:,nParams+1)-mySobolResults.('parrule_gtotal').firstOrderLowerCI(:,nParams+1),'bo');
%set(plotHandle,'xscale','log');
set(gca,'fontsize', 18);
xlabel('Sample size (N)')
ylabel('Sum of Si, 6-par G-function')
xlim([-1, 1050])

% Plot the sum of the total order sensitivity indices results as a
% function of the subsample split size
figure;
plotHandle = errorbar(mySobolResults.sampleSize,mySobolResults.('parrule_gtotal').totalMedian(:,nParams+1),mySobolResults.('parrule_gtotal').totalUpperCI(:,nParams+1)-mySobolResults.('parrule_gtotal').totalMedian(:,nParams+1),mySobolResults.('parrule_gtotal').totalMedian(:,nParams+1)-mySobolResults.('parrule_gtotal').totalLowerCI(:,nParams+1),'bo');
%set(plotHandle,'xscale','log');
set(gca,'fontsize', 18);
xlabel('Sample size (N)')
ylabel('Sum of STi, 6-par G-function')
xlim([-1, 1050])


% Plot the first order sensitivity indices results along with the
% analytical solutions
figure;
hold on
errorbar([1;2;3;4;5;6],mySobolResults.('parrule_gtotal').firstOrderMedian(nSplits,1:6),mySobolResults.('parrule_gtotal').firstOrderUpperCI(nSplits,1:6)-mySobolResults.('parrule_gtotal').firstOrderMedian(nSplits,1:6),mySobolResults.('parrule_gtotal').firstOrderMedian(nSplits,1:6)-mySobolResults.('parrule_gtotal').firstOrderLowerCI(nSplits,1:6),'ko');
%set(gca,'fontsize', 18);
xlabel('Parameter')
ylabel('Si')
set(gca,'XTick',[1,2,3,4,5,6])
set(gca,'YTick',[0,0.25,0.5,0.75,1])
xlim([0 7])
ylim([-.1 1.1])
hold on
set(gca,'fontsize', 18);
plot([1;2;3;4;5;6],analyticalSi,'b^','MarkerSize',6,'MarkerFaceColor','blue');

% Plot the total order sensitivity indices results along with the
% analytical solutions
figure;
hold on
errorbar([1;2;3;4;5;6],mySobolResults.('parrule_gtotal').totalMedian(nSplits,1:6),mySobolResults.('parrule_gtotal').totalUpperCI(nSplits,1:6)-mySobolResults.('parrule_gtotal').totalMedian(nSplits,1:6),mySobolResults.('parrule_gtotal').totalMedian(nSplits,1:6)-mySobolResults.('parrule_gtotal').totalLowerCI(nSplits,1:6),'ko');
%set(gca,'fontsize', 18);
xlabel('Parameter')
ylabel('STi')
set(gca,'XTick',[1,2,3,4,5,6])
set(gca,'YTick',[0,0.25,0.5,0.75,1])
xlim([0 7])
ylim([-.1 1.1])
hold on
set(gca,'fontsize', 18);
plot([1;2;3;4;5;6],analyticalSTi,'b^','MarkerSize',6,'MarkerFaceColor','blue');
```
