## Supplementary File 3 for "A surrogate workflow for virtual population development: designing an amenable proxy for a hypothesis-test inspired nonlinear goodness-of-fit": calculateGFunctionSi.m

```
function [mySi,mySTi] = calculateGFunctionSi(aimatrix)
% A simple function to calculate the analytical solutions for
% first order (Si) and total (STi) sensitivity indices in
% Sobol's G function,
% given an nAi X 1 matrix of ai values 
% See:
% Saltelli, A., et al. (2009). "Global Sensitivity Analysis: The Primer."
% p. 123-125 for the analytical sensitivity index calculation.
%
% ARGUMENTS:
% aimatrix
%
% RETURNS:
% mySi
% mySTi

nAi = length(aimatrix);
% vi is the variance in term
vi = zeros(nAi,1);
for curAiCounter = 1 : nAi
    vi(curAiCounter) = 1/(3*(1+aimatrix(curAiCounter))^2);
end

% We also calculat the total variances
vti = zeros(nAi,1);
for curAiCounter = 1 : nAi
    vti(curAiCounter) = vi(curAiCounter);
    for hotCounter = 2 : nAi
        curCombs = combnk([1:nAi],hotCounter);
        combFinder = (curCombs == curAiCounter);
        combFinder = find(sum(combFinder,2)>0);
        curCombs = curCombs(combFinder,:);
        [nRows, nCols] = size(curCombs);
        curTerm = 0;
        for curRow = 1 : nRows
            tempTerm = 1;
            for curCol = 1 : nCols
                tempTerm = tempTerm * vi(curCombs(curRow,curCol));
            end
            curTerm = curTerm + tempTerm;
        end
        vti(curAiCounter) = vti(curAiCounter) + curTerm;
    end
end

vProd = 1;
for i = 1 : nAi
    vProd = vProd * (1 + vi(i));
end

vTot = -1 + vProd;

mySi = vi / vTot;
mySTi = vti / vTot;

end
```
