## Supplementary File 3 for "A surrogate workflow for virtual population development: designing an amenable proxy for a hypothesis-test inspired nonlinear goodness-of-fit": compare_correlation_coefficients.m

```
% This function compare if two correlation coefficients are significantly
% different.
% The correlation coefficients were tansfered to z scores using fisher's r
% to z transformation. 
% ref: http://core.ecu.edu/psyc/wuenschk/docs30/CompareCorrCoeff.pdf
%--------------------------------------------------------------------------
% Inputs: (1) r1: correlation coefficient of the first correlation (2) r2:
% correlation coefficient of the second correlation (3) n1: number of
% samples used to compute the first correlation (4) n2: number of samples
% used to compute the second correlation
%--------------------------------------------------------------------------
% Output: (1) p: p value, the probability that H0 (the correlation
% coefficiets are not different) is correct
%--------------------------------------------------------------------------
% Example :
% x = rand(20,1); 
% y1= x+rand(20,1)*0.05;
% y2= x+rand(20,1)*0.5;
% r1=corr(x,y1);
% r1=corr(x,y2);
% p = compare_correlation_coefficients(r1,r2,length(x),length(x));
%--------------------------------------------------------------------------
function p = compare_correlation_coefficients(r1,r2,n1,n2)
t_r1 = 0.5*log((1+r1)/(1-r1));
t_r2 = 0.5*log((1+r2)/(1-r2));
z = (t_r1-t_r2)/sqrt(1/(n1-3)+1/(n2-3));
p = (1-normcdf(abs(z),0,1))*2;
end
```
