## Supplementary File 3 for "A surrogate workflow for virtual population development: designing an amenable proxy for a hypothesis-test inspired nonlinear goodness-of-fit": genpath_exclude.m

```
% pathStr = genpath_exclude(basePath,ignoreDirs)
%
% Extension of Matlab's "genpath" function, except this will exclude
% directories (and their sub-tree) given by "ignoreDirs". 
%
%
%
% Inputs:
%    basePath: string.  The base path for which to generate path string.
%
%    excludeDirs: cell-array of strings. all directory names to ignore. Note,
%                 these strings are passed into regexp surrounded by
%                 '^'   and '$'.  If your directory name contains special
%                 characters to regexp, they must be escaped.  For example,
%                 use '\.svn' to ignore ".svn" directories.  You may also
%                 use regular expressions to ignore certian patterns. For
%                 example, use '*._ert_rtw' to ignore all directories ending
%                 with "_ert_rtw".
%
% Outputs:
%    pathStr: string. semicolon delimited string of paths. (see genpath)
% 
% See also genpath
%
% ---CVS Keywords----
% $Author: jhopkin $
% $Date: 2009/10/27 19:06:19 $
% $Name:  $
% $Revision: 1.5 $

% $Log: genpath_exclude.m,v $
% Revision 1.6  2020/11/26 19:06:19  bjs
% Exclude MATLAB "package" directories, "+"
%
% Revision 1.5  2009/10/27 19:06:19  jhopkin
% fixed regexp handling.  added more help comments
%
% Revision 1.4  2008/11/25 19:04:29  jhopkin
% minor cleanup.  Made input more robust so that if user enters a string as 'excudeDir' rather than a cell array of strings this function will still work.  (did this by moving the '^' and '$' to surround the entire regexp string, rather than wrapping them around each "excludeDir")
%
% Revision 1.3  2008/11/25 18:43:10  jhopkin
% added help comments
%
% Revision 1.1  2008/11/22 00:23:01  jhopkin
% *** empty log message ***
%

function p = genpath_exclude(d,excludeDirs)
	% if the input is a string, then use it as the searchstr
	if ischar(excludeDirs)
		excludeStr = excludeDirs;
	else
		excludeStr = '';
		if ~iscellstr(excludeDirs)
			error('excludeDirs input must be a cell-array of strings');
		end
		
		for i = 1:length(excludeDirs)
			excludeStr = [excludeStr '|^' excludeDirs{i} '$'];
		end
	end

	
	% Generate path based on given root directory
	files = dir(d);
	if isempty(files)
	  return
	end

	% Add d to the path even if it is empty.
	p = [d pathsep];

	% set logical vector for subdirectory entries in d
	isdir = logical(cat(1,files.isdir));
	%
	% Recursively descend through directories which are neither
	% private nor "class" directories.
	%
	dirs = files(isdir); % select only directory entries from the current listing

	for i=1:length(dirs)
		dirname = dirs(i).name;
		%NOTE: regexp ignores '.', '..', '@.*', and 'private' directories by default. 
		if ~any(regexp(dirname,['^\.$|^\.\.$|^\@.*|^\+.*|^private$|^resources$|' excludeStr ],'start'))
		  p = [p genpath_exclude(fullfile(d,dirname),excludeStr)]; % recursive calling of this function.
		end
	end
end
```
