## Supplementary File 3 for "A surrogate workflow for virtual population development: designing an amenable proxy for a hypothesis-test inspired nonlinear goodness-of-fit": kstest_2s_2d.m

```
function [H, pValue, KSstatistic] = kstest_2s_2d(x1, x2, alpha)

% kstest_2s_2d - FUNCTION Two-sample Two-dimensional Kolmogorov-Smirnov Test
%
% Usage:[H, pValue, KSstatistic] = kstest_2s_2d(x1, x2 <, alpha>)
%
% The paired-sample Kolmogorov-Smirnov test is a statistical test used to
% determine whether two sets of data arise from the same or different
% distributions.  The null hypothesis is that both data sets were drawn
% from the same continuous distribution.
% 
% The algorithm in this function is taken from Peacock [1].
% 
% 'x1' is an [Nx2] matrix, each row containing a two-dimensional sample.
% 'x2' is an [Mx2] matrix, each row likewise containing a two-dimensional
% sample. The optional argument 'alpha' is used to set the desired
% significance level for rejecting the null hypothesis.
% 
% 'H' is a logical value: true indicates that the null hypothesis should be
% rejected.  'pValue' is an estimate for the P value of the test statistic.
% 'KSstatistic' is the raw value for the test statistic ('D' in [1]).
% 
% In contrast to kstest2, this function can only perform a two-tailed test.
% This is because Peacock does not provide a method for estimating P in the
% one-tailed case [1].  Suggestions for a one-tailed test are welcome.
%
% References: [1] J. A. Peacock, "Two-dimensional goodness-of-fit testing
%  in astronomy", Monthly Notices Royal Astronomy Society 202 (1983)
%  615-627.
%    Available from: http://articles.adsabs.harvard.edu/cgi-bin/nph-iarticle_query?1983MNRAS.202..615P&defaultprint=YES&filetype=.pdf
%

% Author: Dylan Muir (From kstest_2s_2d by Qiuyan Peng @ ECE/HKUST) Date:
% 13th October, 2012

%%

if nargin < 2
   error('stats:kstest2:TooFewInputs','At least 2 inputs are required.');
end


%%
%
% x1,x2 are both 2-column matrices
%

if ((size(x1,2)~=2)||(size(x2,2)~=2))
   error('stats:kstest2:TwoColumnMatrixRequired','The samples X1 and X2 must be two-column matrices.');
end
n1 = size(x1,1);
n2 = size(x2,1);


%%
%
% Ensure the significance level, ALPHA, is a scalar
% between 0 and 1 and set default if necessary.
%

if (nargin >= 3) && ~isempty(alpha)
   if ~isscalar(alpha) || (alpha <= 0 || alpha >= 1)
      error('stats:kstest2:BadAlpha',...
         'Significance level ALPHA must be a scalar between 0 and 1.');
   end
else
   alpha  =  0.05;
end


%%
%
% Calculate F1(x) and F2(x), the empirical (i.e., sample) CDFs.
%

% - A function handle to perform comparisons in all possible directions
fhCounts = @(x, edge)([(x(:, 1) > edge(1)) & (x(:, 2) > edge(2))...
   (x(:, 1) <= edge(1)) & (x(:, 2) > edge(2))...
   (x(:, 1) <= edge(1)) & (x(:, 2) <= edge(2))...
   (x(:, 1) > edge(1)) & (x(:, 2) <= edge(2))]);

KSstatistic = -inf;

fprintf('Measuring [%6.2f%%]...', 0);

for iX = 1:(n1+n2)
   % - Choose a starting point
   if (iX<=n1)
      edge = x1(iX,:);
   else
      edge = x2(iX-n1,:);
   end
   
   % - Estimate the CDFs for both distributions around this point
   vfCDF1 = sum(fhCounts(x1, edge)) ./ n1;
   vfCDF2 = sum(fhCounts(x2, edge)) ./ n2;
   
   % - Two-tailed test statistic
   vfThisKSTS = abs(vfCDF1 - vfCDF2);
   fKSTS = max(vfThisKSTS);
   
   % - Final test statistic is the maximum absolute difference in CDFs
   if (fKSTS > KSstatistic)
      KSstatistic = fKSTS;
   end
   
   if (mod(iX, 1000) == 0)
      fprintf('\b\b\b\b\b\b\b\b\b\b\b%6.2f%%]...', iX / (n1+n2)*100);
   end
end

fprintf('\n');


%% Peacock Z calculation and P estimation

n      =  n1 * n2 /(n1 + n2);
Zn = sqrt(n) * KSstatistic;
Zinf = Zn / (1 - 0.53 * n^(-0.9));
pValue = 2 * exp(-2 * (Zinf - 0.5).^2);

% Clip invalid values for P
if (pValue > 1)
   pValue = 1;
end

H = (pValue <= alpha);
```
