## Supplementary File 3 for "A surrogate workflow for virtual population development: designing an amenable proxy for a hypothesis-test inspired nonlinear goodness-of-fit": quadrantCount.m

```
function quadrantChecks = quadrantCount(x1, x2)
% This function is roughly a 2D equivalent to creating the comparison grid for the
% 1D KS test.  Here, all of the quadrant comparisons are run for each point in
% x1 as a reference against x2
%
%  Arguments
%   x1:              2xN1 matrix of reference points
%   x2:              2xN2 matrix of reference points
%
%  Returns
%   quadrantChecks   N1x(4 quadrants * N2) 
%                    matrix of binary comparisons

[~,nsample1] = size(x1);
[~,nsample2] = size(x2);

% - A function handle to perform comparisons in all possible directions
pointCompare = @(x, point)([((x(1,:) > point(1)) & (x(2,:) > point(2))),...
   ((x(1, :) <= point(1)) & (x(2, :) > point(2))),...
   ((x(1, :) <= point(1)) & (x(2, :) <= point(2))),...
   ((x(1, :) > point(1)) & (x(2, :) <= point(2)))]);

quadrantChecks = false(nsample1,4*nsample2);
for j = 1:(nsample1)
	point = x1(:,j);
	quadrantChecks(j,:) = pointCompare(x2, point);
end
```
