## Supplementary File 3 for "A surrogate workflow for virtual population development: designing an amenable proxy for a hypothesis-test inspired nonlinear goodness-of-fit": weightedKS2D.m

```
function myPVal = weightedKS2D(sample1, sample2, quadrantCounts1cross, quadrantCounts1self, quadrantCounts2cross, quadrantCounts2self, W1, W2, selectN1, selectN2)
% This function performs a two-sided 2D KS test and returns a p-value.
% It is assumed quadrant counts are performed prior to running this test.
% Note these tests currently can require a substantial computational
% overhead and a shortcut is included to speed the calculation by ignoring
% data with low weight.
%
% References: 
%  Peacock, JA. Two-dimensional goodness-of-fit testing
%  in astronomy, Mon Not R astr Soc 202 (1983)
%  615-627.
%
%  Fasano, G., Franceschini, A., 1987. A 
%  multidimensional version of the Kolmogorov-Smirnov 
%  test. Mon Not R ast Soc 225, 155–170.
%
%  Press et al., 1993. Numerical recipes in C: 
%  The art of scientific computing, second edition.
%
% original written by:
% Dylan Muir (From kstest_2s_2d by Qiuyan Peng @ ECE/HKUST) Date:
% 13th October, 2012
% This code substantially modified to account for prevalence weighting
% in accordance with license from a MATLAB implementation,
% Modification by Brian Schmidt, August 2018
% The original code also did not appear to take into consideration
% the comparison for all the pooled (N1+N2) x (N1+N2) points
% as proposed by Peacock. The original code
% was closer to the method by Fasano & Franceschini.  This code has
% ben updated to better correspond to the F&F method
% by also using the weighted correlations for the calculation
% of the test statistic in addition to the 
% observed points.
%
% ARGUMENTS
%  sample1                a 2xlength(sample1) matrix of observed values
%  sample2                a 2xlength(sample2) matrix of observed values
%  quadrantCounts1cross   quadrant counts with each value in sample 1 as reference against sample 2
%  quadrantCounts1self
%  quadrantCounts2cross   quadrant counts with each value in sample 2 as reference against sample 1
%  quadrantCounts2self
%  W1                     a 1xlength(sample1) vector of weights for sample 1
%  W2                     a 1xlength(sample2) vector of weights for sample 2
%  selectN1               number of datapoints to use from sample1, the
%                          most highly weighted are selected.  This is done
%                          to speed calculation and avoid spending time
%                          on unweighted portions of the sample space.
%                          Set to 0 to use all.
%  selectN2               number of datapoints to use from sample2, the
%                          most highly weighted are selected.  This is done
%                          to speed calculation and avoid spending time
%                          on unweighted portions of the sample space.
%                          Set to 0 to use all.
%
% RETURNS
%  myPVal                 the p-value resulting from the two-tailed test

nTotal1 = length(W1);
nTotal2 = length(W2);

if (selectN1 == 0) ||  (selectN1 == nTotal1)
    nSamples1 = nTotal1;
else
    nSamples1 = selectN1;
    [~, indices1] = sort(W1, 'descend');
    indices1 = indices1(1:nSamples1);
    quadrantCounts1cross = quadrantCounts1cross(indices1,:);
    quadrantCounts2cross = quadrantCounts2cross(:,[indices1,nTotal1+indices1,2*nTotal1+indices1,3*nTotal1+indices1]);
    quadrantCounts1self = quadrantCounts1self(indices1,[indices1,nTotal1+indices1,2*nTotal1+indices1,3*nTotal1+indices1]);
    sample1 = sample1(:,indices1);
    W1 = W1(indices1);
    W1 = W1/sum(W1);
end
if (selectN2 == 0) ||  (selectN2 == length(W2))
    nSamples2 = nTotal2;
else        
    nSamples2 = selectN2;    
    [~, indices2] = sort(W2, 'descend');
    indices2 = indices2(1:nSamples2);
    quadrantCounts1cross = quadrantCounts1cross(:,[indices2,nTotal2+indices2,2*nTotal2+indices2,3*nTotal2+indices2]);
    quadrantCounts2cross = quadrantCounts2cross(indices2,:);
    quadrantCounts2self = quadrantCounts2self(indices2,[indices2,nTotal2+indices2,2*nTotal2+indices2,3*nTotal2+indices2]);
    sample2 = sample2(:,indices2);
    W2 = W2(indices2);
    W2 = W2/sum(W2);    
end

b11 = nSamples1;
b12 = nSamples1*2;
b13 = nSamples1*3;
b21 = nSamples2;
b22 = nSamples2*2;
b23 = nSamples2*3;
KSstatistic = ones(nSamples1+nSamples2,1);
WQuadC = zeros(nSamples1+nSamples2,4);
WQuadS = zeros(nSamples1+nSamples2,4);

WQuadC(1:nSamples1,:) = [sum(quadrantCounts1cross(1:nSamples1,1:b21).*(ones(nSamples1,1).*W2),2),sum(quadrantCounts1cross(1:nSamples1,b21+1:b22).*(ones(nSamples1,1).*W2),2),...
	  sum(quadrantCounts1cross(1:nSamples1,b22+1:b23).*(ones(nSamples1,1).*W2),2),sum(quadrantCounts1cross(1:nSamples1,b23+1:end).*(ones(nSamples1,1).*W2),2)];	  
WQuadC(nSamples1+1:nSamples1+nSamples2,:) = [sum(quadrantCounts2cross(1:nSamples2,1:b11).*(ones(nSamples2,1).*W1),2),sum(quadrantCounts2cross(1:nSamples2,b11+1:b12).*(ones(nSamples2,1).*W1),2),...
	  sum(quadrantCounts2cross(1:nSamples2,b12+1:b13).*(ones(nSamples2,1).*W1),2),sum(quadrantCounts2cross(1:nSamples2,b13+1:end).*(ones(nSamples2,1).*W1),2)];
WQuadS(1:nSamples1,:) = [sum(quadrantCounts1self(1:nSamples1,1:b11).*(ones(nSamples1,1).*W1),2),sum(quadrantCounts1self(1:nSamples1,b11+1:b12).*(ones(nSamples1,1).*W1),2),...
	  sum(quadrantCounts1self(1:nSamples1,b12+1:b13).*(ones(nSamples1,1).*W1),2),sum(quadrantCounts1self(1:nSamples1,b13+1:end).*(ones(nSamples1,1).*W1),2)];	  
WQuadS(nSamples1+1:nSamples1+nSamples2,:) = [sum(quadrantCounts2self(1:nSamples2,1:b21).*(ones(nSamples2,1).*W2),2),sum(quadrantCounts2self(1:nSamples2,b21+1:b22).*(ones(nSamples2,1).*W2),2),...
	  sum(quadrantCounts2self(1:nSamples2,b22+1:b23).*(ones(nSamples2,1).*W2),2),sum(quadrantCounts2self(1:nSamples2,b23+1:end).*(ones(nSamples2,1).*W2),2)];	 	  
KSstatistic = max(abs(WQuadC - WQuadS),[],2);
% See page 649 in Press et al.
% For the two-sample FF test they suggest
% taking the average of the
% comparisons against sample 1 and sample2
% as opposed to the max.
% KSstatistic = max(KSstatistic);
KSstatistic = (max(KSstatistic(1:nSamples1))+max(KSstatistic(nSamples1+1:end)))/2;

% Fasano & Franceschini also include the correlation
corr1 = weightedcorrs(sample1', W1'); 
corr1 = corr1(1,2);
corr2 = weightedcorrs(sample2', W2'); 
corr2 = corr2(1,2);
corrr = sqrt(1 - 0.5*(corr1^2 + corr2^2));

N1 = 1/sum(W1.^2);
N2 = 1/sum(W2.^2);

% See page 649 in Numerical Recipes
% This is an infinite series.  Here we assume 102 terms is enough
% TODO: may want to add a check to validate desired precision
% for the series
nPool =  N1 * N2 /(N1 + N2);
iVals =  (1:102)';
zVal = (sqrt(nPool)*KSstatistic) / (1 + corrr*(.25 - .75/sqrt(nPool)));
myPVal = 2 * sum((-1).^(iVals-1).*exp(-2*zVal^2*iVals.^2));
myPVal = min(max(myPVal,0),1);


end
```
