## Supplementary File 3 for "A surrogate workflow for virtual population development: designing an amenable proxy for a hypothesis-test inspired nonlinear goodness-of-fit": myfisher22.m

```
function Pvalue=myfisher22(x,varargin)
%Fisher's exact test of 2x2 contingency tables permits calculation of
%precise probabilities in situation where, as a consequence of small cell
%frequencies, the much more rapid normal approximation and chi-square
%calculations are liable to be inaccurate. The Fisher's exact test involves
%the computations of several factorials to obtain the probability of the
%observed and each of the more extreme tables. Factorials growth quickly,
%so it's necessary use logarithms of factorials. This computations is very
%easy in Matlab because x!=gamma(x+1) and log(x!)=gammaln(x+1). This
%function is fully vectorized to speed up the computation.
%The routine coumputes the Power and, if necessary, the sample sizes needed
%to achieve a power=0.80 using a modified asymptotic normal method with
%continuity correction as described by Hardeo Sahai and Anwer Khurshid in
%Statistics in Medicine, 1996, Vol. 15, Issue 1: 1-21.
%
% Syntax: 	myfisher22(x,alpha,plts)
%      
%     Inputs:
%           X - 2x2 data matrix 
%           ALPHA - significance level (default = 0.05).
%           PLTS - Flag to set if you don't want (0) or want (1) view the plot
%           of Wald Statistics distribution (default=0)
%     Outputs:
%           - Three p-values
%           - Power and sample sizes
%
%   Example:
%
%                                    Vaccine
%                               Yes           No
%                            ---------------------
%                    Yes         7            12
% Infectious status                 
%                     No         8            3
%                            ---------------------
%
%   Calling on Matlab the function: 
%             myfisher22([7 12; 8 2])
%
%   Answer is:
%
% --------------------------------------------------------------------------------
% 2x2 matrix Fisher's exact test
% --------------------------------------------------------------------------------
%     Tables    left_tail    right_tail    two_tails    Mid_p_correction
%     ______    _________    __________    _________    ________________
% 
%     11        0.032884     0.99635       0.050175     0.035557        
% 
% Power Computation (Asymptotic normal method)
% --------------------------------------------------------------------------------
% alpha = 0.0500  Sum of row 1 (n1) = 10  Sum of row 2 (n2) = 19
% --------------------------------------------------------------------------------
%              one_tail    two_tails
%              ________    _________
% 
%     Z1_b     0.65726     0.29123  
%     Power    0.74449     0.61456  
% 
%  
% To achieve a recommended Power=0.80
%           one_tail    two_tails
%           ________    _________
% 
%     n1    15          29       
%     n2    18          34       
% 
% --------------------------------------------------------------------------------
%
%           Created by Giuseppe Cardillo
%          
%
% To cite this file, this would be an appropriate format:
% Cardillo G. (2007) MyFisher22: a very compact routine for Fisher's exact
% test on 2x2 matrix
% http://www.mathworks.com/matlabcentral/fileexchange/15434

%Input error handling
p = inputParser;
addRequired(p,'x',@(x) validateattributes(x,{'numeric'},{'real','finite','integer','nonnegative','nonnan','size',[2 2]}));
addOptional(p,'alpha',0.05, @(x) validateattributes(x,{'numeric'},{'scalar','real','finite','nonnan','>',0,'<',1}));
addOptional(p,'plts',0, @(x) isnumeric(x) && isreal(x) && isfinite(x) && isscalar(x) && (x==0 || x==1));
parse(p,x,varargin{:});
alpha=p.Results.alpha; plts=p.Results.plts;
clear p

Rs=sum(x,2); %rows sum
Cs=sum(x); %columns sum
N=sum(Rs); %sum of all elements

%Rearrange the matrix if necessary.
flip=0; %flip flag
if ~issorted(Rs)
    x=flipud(x);
    Rs=sort(Rs);
    flip=flip+1;
end
if ~issorted(Cs)
    x=fliplr(x);
    Cs=sort(Cs);
    flip=flip+1;
end

%           X - 2x2 data matrix composed like this: 
%          ___________
%         |  A  |  B  |Rs1
%         |_____|_____|
%         |  C  |  D  |Rs2
%         |_____|_____|____
%           Cs1  Cs2    N
%recall that Fisher's P=[Rs(1)!Rs(2)!Cs(1)!Cs(2)!]/[N!A!B!C!D!]
%Log(A*B)=Log(A)+Log(B) and Log(A/B)=Log(A)-Log(B)

%A 2x2 matrix has only 1 degree of freedom...
A=0:1:min(Rs(1),Cs(1)); %all possible values for the first cell
z=[A;Rs(2)-Cs(1)+A;Rs(1)-A;Cs(1)-A;]; %all possible matrices

%          The first table will be: 
%          _____________
%         |  0  | Rs1   |Rs1
%         |_____|_______|
%         | Cs1 |Cs2-Rs1|Rs2
%         |_____|_______|____
%           Cs1  Cs2      N
%
%P0=[Rs(1)!Rs(2)!Cs(1)!Cs(2)!]/[N!0!Rs(1)!Cs(1)!D!]=[Rs(2)!Cs(2)!]/[N!D!]
np=zeros(1,length(A)); %p-values vector preallocation
lz=log(z);
%LOG(X!)=GAMMALN(x+1)
np(1)=sum(gammaln([Rs(2)+1 Cs(2)+1])-gammaln([N+1 z(2)+1]));
%remember that 
%np(i+1)=np(i)*[B(i)*C(i)]/(A(i+1)*D(i+1)]=np(i)*f(i) 
%This formula is vectorizable and is recursive!
%using Logarithm to improve computation
%log(np(i+1))=log(np(i))+[log(B(i))+log(C(i))]-[log(A(i+1))+log(D(i+1))]=log(np(i))+f(i)
%J>1
%log(np(J))=log(np(1))+sum(f(1:J-1))=log(np(1))+cumsum(f)
f=sum(lz(3:4,1:end-1))-sum(lz(1:2,2:end));
np(2:end)=np(1)+cumsum(f);
np=exp(np);

%now compute the 1-tailed p-values and the 2-tailed p-value
W=x(1)+1;
if flip~=1 %choose direction
    P=[sum(np(1:W)) sum(np(W:end)) sum(np(np<=np(W)))];
else
    P=[sum(np(W:end)) sum(np(1:W)) sum(np(np<=np(W)))];
end

%display results
tr=repmat('-',1,80);
disp(tr)
disp('2x2 matrix Fisher''s exact test')
disp(tr)
disp(array2table([size(z,2),P,0.5*np(W)+sum(np(np<np(W)))],'VariableNames',{'Tables','left_tail','right_tail','two_tails','Mid_p_correction'}));

%power (Asymptotic normal method)
Za=-realsqrt(2).*erfcinv(2.*[1-alpha 1-alpha/2]);
Zb=-realsqrt(2)*erfcinv(1.6); %1-Beta=0.8
p=x(:,1)./Rs;
d=abs(diff(p));
k=Rs(2)/Rs(1);
q=1-p;
pm=(p(1)+k*p(2))/(k+1);
qm=1-pm;
Z1_b=abs((realsqrt(Rs(1)*d^2)-Za.*realsqrt((1+1/k)*pm*qm))/realsqrt(p(1)*q(1)+p(2)*q(2)/k));
pwr=0.5.*erfc(-Z1_b./realsqrt(2));
fprintf('Power Computation (Asymptotic normal method)\n')
disp(tr)
fprintf('alpha = %0.4f  Sum of row 1 (n1) = %d  Sum of row 2 (n2) = %d\n',alpha,Rs)
disp(tr)
disp(array2table([Z1_b;pwr],'VariableNames',{'one_tail','two_tails'},'RowNames',{'Z1_b','Power'}))
if any(pwr<0.8)
    %sample size (Modified Asymptotic normal method with continuity correction)
    nstar=(Za.*realsqrt(pm*qm*(1+1/k))+Zb.*realsqrt(p(1)*q(1)+p(2)*q(2)/k)).^2./d^2;
    n1=round(nstar./4.*(1+realsqrt(1+2*(k+1)./(k*d.*nstar))).^2);
    n2=round(k.*n1);
    disp(' ')
    disp('To achieve a recommended Power=0.80')
    disp(array2table([n1' n2'],'VariableNames',{'one_tail','two_tails'},'RowNames',{'n1','n2'}))
end
disp(tr)

if plts
    %Display plot
    D=sum(z([1 3],:))./N;
    TX=(z(1,:)./Cs(1)-z(3,:)./Cs(2))./realsqrt(D.*(1-D).*sum(1./Cs));
    
    %The Matlab BAR function doesn't work properly. So I set-up a my proper BAR
    %function using the FILL function.
    hold on
    Wh=2*abs(TX(2)-TX(1))/5; %BarWidth established on the x-ticks
    leTX=TX(np<=np(W)); lenp=np(np<=np(W)); %less or equal than observed
    MX1=repmat(leTX,4,1); MY1=repmat(lenp,4,1);
    MX1([1 2],:)=MX1([1 2],:)-Wh; MX1([3 4],:)=MX1([3 4],:)+Wh; MY1([1 4],:)=0;
    H1=fill(MX1,MY1,'b');
    H1Group = hggroup; %Group all this bar
    set(H1,'Parent',H1Group)
    set(get(get(H1Group,'Annotation'),'LegendInformation'),'IconDisplayStyle','on'); % Include this hggroup in the legend
    
    eTX=TX(W); enp=np(W); %Observed table
    MX2=repmat(eTX,4,1); MY2=repmat(enp,4,1);
    MX2([1 2],:)=MX2([1 2],:)-Wh; MX2([3 4],:)=MX2([3 4],:)+Wh; MY2([1 4],:)=0;
    fill(MX2,MY2,'g');

    if any(np>np(W))
        gTX=TX(np>np(W)); gnp=np(np>np(W)); %greater than observed
        MX3=repmat(gTX,4,1); MY3=repmat(gnp,4,1);
        MX3([1 2],:)=MX3([1 2],:)-Wh; MX3([3 4],:)=MX3([3 4],:)+Wh; MY3([1 4],:)=0;
        H3=fill(MX3,MY3,'r');
        H3Group = hggroup; %Group all this bar
        set(H3,'Parent',H3Group)
        set(get(get(H3Group,'Annotation'),'LegendInformation'),'IconDisplayStyle','on'); % Include this hggroup in the legend
        legend('P<=P_o_b_s_e_r_v_e_d _t_a_b_l_e','P_o_b_s_e_r_v_e_d _t_a_b_l_e','P>P_o_b_s_e_r_v_e_d _t_a_b_l_e');
    else
        legend('P<=P_o_b_s_e_r_v_e_d _t_a_b_l_e','P_o_b_s_e_r_v_e_d _t_a_b_l_e')
    end
    hold off
    
    axis square
    title('Distribution of Wald Statistic for Fisher''s Exact test','FontName','Arial','FontSize',12,'FontWeight','Bold');
    xlabel('T(X)','FontName','Arial','FontSize',12,'FontWeight','Bold');
    txt=['P[T(X)|X€Gamma(' num2str(Rs(1)) ')]'];
    ylabel(txt,'FontSize',12,'FontWeight','Bold');
end

if nargout
    Pvalue=P;
end
```
