## Supplementary File 3 for "A surrogate workflow for virtual population development: designing an amenable proxy for a hypothesis-test inspired nonlinear goodness-of-fit": myfisher23.m

```
function Pvalue=myfisher23(x)
%P=MYFISHER23(X)- Fisher's Exact Probability Test on 2x3 matrix.
%Fisher's exact test of 2x3 contingency tables permits calculation of
%precise probabilities in situation where, as a consequence of small cell
%frequencies, the much more rapid normal approximation and chi-square
%calculations are liable to be inaccurate. The Fisher's exact test involves
%the computations of several factorials to obtain the probability of the
%observed and each of the more extreme tables. Factorials growth quickly,
%so it's necessary use logarithms of factorials. This computations is very
%easy in Matlab because x!=gamma(x+1) and log(x!)=gammaln(x+1). This
%function is fully vectorized to speed up the computation.
%
% Syntax: 	myfisher23(x)
%      
%     Inputs:
%           X - 2x3 data matrix 
%     Outputs:
%           - Three p-values
%
%   Example:
%
%                A   B   C
%           -------------------
%      X         0   3   2
%           -------------------    
%      Y         6   5   1
%           -------------------
%
%   Calling on Matlab the function: 
%             myfisher23([0 3 2; 6 5 1])
%
%   Answer is:
%
% --------------------------------------------------------------------------------
% 2x3 matrix Fisher's exact test
% --------------------------------------------------------------------------------
%     Tables    two_tails_p_value    Mid_p_correction
%     ______    _________________    ________________
% 
%     18        0.088235             0.074661  
%
%           Created by Giuseppe Cardillo
%          
%
% To cite this file, this would be an appropriate format:
% Cardillo G. (2007) MyFisher23: a very compact routine for Fisher's exact
% test on 2x3 matrix
% http://www.mathworks.com/matlabcentral/fileexchange/15399


%Input error handling
p = inputParser;
addRequired(p,'x',@(x) validateattributes(x,{'numeric'},{'real','finite','integer','nonnegative','nonnan','size',[2 3]}));
parse(p,x);
clear p

Rs=sum(x,2); %rows sum
Cs=sum(x); %columns sum
N=sum(Rs); %Total observations

%If necessed, rearrange matrix
if ~issorted(Cs)
    [Cs,ind]=sort(Cs);
    x=x(:,ind);
    clear ind
end
if ~issorted(Rs)
    [Rs,ind]=sort(Rs);
    x=x(ind,:);
    clear ind
end

%recall that Fisher's P=[Prod(Rs!)*Prod(Cs!)]/[N!*prod(X(i,j)!)]
%Log(A*B)=Log(A)+Log(B) and Log(A/B)=Log(A)-Log(B)
%Construct all possible tables
%A 2x3 matrix has 2 degrees of freedom...
A=0:1:min(Rs(1),Cs(1)); %all possible values of X(1,1)
B=min(Cs(2),Rs(1)-A); %max value of X(1,2) given X(1,1)
C=max(Rs(1)-A-Cs(3),0); % min value of X(1,2) given X(1,1)
et=sum(B-C+ones(size(B))); %tables to evaluate
Tables=zeros(et,6); %Matrix preallocation
%compute the index
stop=cumsum(B-C+1);
start=[1 stop(1:end-1)+1];
%In the first round of the for cycle, Column 1 assignment should be skipped
%because it is already zero. So, modify the cycle...
Tables(start(1):stop(1),2)=C(1):1:B(1); %Put in the Column2 all the possible values of X(1,2) given X(1,1)
for I=2:length(A)
    Tables(start(I):stop(I),1)=A(I); %replicate the A(I) value for B(I)-C(I)+1 times
    %Put in the Column2 all the possible values of X(1,2) given X(1,1)
    Tables(start(I):stop(I),2)=C(I):1:B(I); 
end
clear A B start stop
%The degrees of freedom are finished, so complete the table...
%...Put all the possible values of X(1,3) given X(1,1) and X(1,2)
Tables(:,3)=Rs(1)-sum(Tables(:,1:2),2);
%Complete the second row given the first row
Tables(:,4:6)=repmat(Cs,et,1)-Tables(:,1:3);

%Compute log(x!) using the gammaln function
zf=gammaln(Tables+1); %compute log(x!)
K=sum(gammaln([Rs' Cs]+1))-gammaln(N+1); %The costant factor K=log(prod(Rs!)*prod(Cs!)/N!)
np=exp(K-sum(zf,2)); %compute the p-value of each possible matrix
[~,obt]=ismember(x(1,:),Tables(:,1:3),'rows'); %Find the observed table
clear zf K tf
%Finally compute the probability for 2-tailed test
P=sum(np(np<=np(obt)));

%display results
tr=repmat('-',1,80);
disp(tr)
disp('2x3 matrix Fisher''s exact test')
disp(tr)
disp(array2table([et,P,0.5*np(obt)+sum(np(np<np(obt)))],'VariableNames',{'Tables','two_tails_p_value','Mid_p_correction'}));

if nargout
    Pvalue=P;
end
```
