## Supplementary File 3 for "A surrogate workflow for virtual population development: designing an amenable proxy for a hypothesis-test inspired nonlinear goodness-of-fit": myfisher24.m

```
function Pvalue=myfisher24(x)
%P=MYFISHER24(X)- Fisher's Exact Probability Test on 2x4 matrix.
% Fisher's exact test of 2x4 contingency tables permits calculation of
% precise probabilities in situation where, as a consequence of small cell
% frequencies, the much more rapid normal approximation and chi-square
% calculations are liable to be inaccurate. The Fisher's exact test involves
% the computations of several factorials to obtain the probability of the
% observed and each of the more extreme tables. Factorials growth quickly,
% so it's necessary use logarithms of factorials. This computations is very
% easy in Matlab because x!=gamma(x+1) and log(x!)=gammaln(x+1). This
% function is now fully vectorized to speed up the computation.
%
% Syntax: 	myfisher24(x)
%      
%     Inputs:
%           X - 2x4 data matrix 
%     Outputs:
%           - Three p-values
%
%   Example:
%
%                A   B   C   D
%           -------------------
%      X         2   3   0   1
%           -------------------    
%      Y         4   0   2   6
%           -------------------
%                                       
%
%   x=[2 3 0 1; 4 0 2 6];
%
%   Calling on Matlab the function: 
%             myfisher24(x)
%
%   Answer is:
%
% 2x4 matrix Fisher's exact test: 54 tables were evaluated
% -----------------------------------------------------------------
% 		 p-value (2-tails): 0.0523594053
% -----------------------------------------------------------------
%
%           Created by Giuseppe Cardillo
%          
%
% To cite this file, this would be an appropriate format:
% Cardillo G. (2007) MyFisher24: a very compact routine for Fisher's exact
% test on 2x4 matrix
% http://www.mathworks.com/matlabcentral/fileexchange/19842

% Input Error handling
%Input Error handling
if ~isequal(size(x),[2 4])
    if isequal(size(x),[4 2])
        x=x';
    else
        error('Input matrix must be a 2x4 matrix')
    end
end
if ~all(isfinite(x(:))) || ~all(isnumeric(x(:)))
    error('Warning: all X values must be numeric and finite')
end
if ~isequal(x(:),round(x(:)))
    error('Warning: X data matrix values must be whole numbers')
end

Rs=sum(x,2); %rows sum
Cs=sum(x); %columns sum
N=sum(Rs); %total observations

%If necessed, rearrange matrix
if ~issorted(Cs)
    [Cs,ind]=sort(Cs);
    x=x(:,ind);
end
if ~issorted(Rs)
    [Rs,ind]=sort(Rs);
    x=x(ind,:);
end
%recall that Fisher's P=[Prod(Rs!)*Prod(Cs!)]/[N!*prod(X(i,j)!)]
%Log(A*B)=Log(A)+Log(B) and Log(A/B)=Log(A)-Log(B)

%Costruct all possible tables
%A 2x4 matrix has 3 degrees of freedom...
A=0:1:min(Rs(1),Cs(1)); %all possible values of X(1,1)
B=min(Cs(2),Rs(1)-A); %max value of X(1,2) given X(1,1)
et=0; BigM=cell(length(A),2); %preallocation
for I=1:length(A);
    %set of max values of X(1,3) given X(1,1) and X(1,2)
    BigM{I,1}=min(Cs(3),Rs(1)-(A(I)+(0:1:B(I)))); 
    %how many tables are generated for each value of X(1,1)?
    BigM{I,2}=sum(BigM{I,1}+ones(size(BigM{I,1})),2);
    et=et+BigM{I,2}; %tables to evaluate
end
Tables=zeros(et,8); %Matrix preallocation
%Fill the Columns
%IdxA are vectors with the same length of A: they indicates how many rows 
%of the first column must be filled with the A(I) value 
idxAstop=cumsum(cell2mat(BigM(:,2)));
idxAstart=[1; idxAstop(1:end-1)+1];

for I=1:length(idxAstart)
    %In the first round of the for cycle skip the Column 1 assignment
    %because it is already zero.
    if I>1
        Tables(idxAstart(I):idxAstop(I),1)=A(I);
        prevA=idxAstop(I-1);
    else
        prevA=0;
    end
%IdxB vectors indicates how many rows of the second column must be filled 
%with the Jth value 0<=J<=B(I) and how many rows of column 3 must be filled
%with the X(1,3) set.
    Ct=BigM{I,1};
    idxBstop=prevA+cumsum(Ct+1);
    idxBstart=[prevA+1 idxBstop(1:end-1)+1];
    for J=1:B(I)+1
        %In the first round of the for cycle skip the Column 2 assignment
        %because it is already zero.
        if J>1
            Tables(idxBstart(J):idxBstop(J),2)=J-1;
        end
        Tables(idxBstart(J):idxBstop(J),3)=0:1:Ct(J);
    end
end

clear A B BigM Ct idxAstart idxAstop idxBstart idxBstop I J prevA
%The degrees of freedom are finished, so complete the table...
%...Put all the possible values of X(1,4) given X(1,1:3)
Tables(:,4)=Rs(1)-sum(Tables(:,1:3),2);

% Modified here from Giuseppe Cardillo's implementation.
% http://www.mathworks.com/matlabcentral/fileexchange/19842-myfisher24
% 23 Dec 2009	v 1.5
% We need to separately enforce the constraint that
% Tables(:,4) <= Cs(4) for some cases. 
% For example, check the evaluation of
% x = [0, 4, 9, 2; 0, 4, 2, 11] in v 1.5.
Tables=Tables(find(Tables(:,4) <= Cs(4)),:);
[et, ~] = size(Tables);

%Complete the second row given the first row
Tables(:,5:8)=repmat(Cs,et,1)-Tables(:,1:4);

%Compute log(x!) using the gammaln function
zf=gammaln(Tables+1); %compute log(x!)
K=sum(gammaln([Rs' Cs]+1))-gammaln(N+1); %The costant factor K=log(prod(Rs!)*prod(Cs!)/N!)
np=exp(K-sum(zf,2)); %compute the p-value of each possible matrix

[tf,obt]=ismember(x(1,:),Tables(:,1:4),'rows'); %Find the observed table

%Finally compute the probabilities for 2-tailed test
P=sum(np(np<=np(obt)));

%display results
tr=repmat('-',1,65); %Set up the divisor
disp(' ')
fprintf('2x4 matrix Fisher''s exact test: %0.0f tables were evaluated\n',et)
disp(tr)
fprintf('\t\t p-value (2-tails): %0.10f\n',P); 
disp(tr)
fprintf('Mid-p correction: %0.10f\n',0.5*np(obt)+sum(np(np<np(obt)))); 
disp(tr)
disp(' ')

if nargout
    Pvalue=P;
end
```
