## Supplementary File 3 for "A surrogate workflow for virtual population development: designing an amenable proxy for a hypothesis-test inspired nonlinear goodness-of-fit": myfisher.m

```
function Pout=myfisher(x,varargin)
%P=MYFISHER(X)- Fisher's Exact Probability Test for a RxC matrix.
% Fisher's exact test permits calculation of precise probabilities in situation 
% where, as a consequence of small cell frequencies, the much more rapid normal 
% approximation and chi-square calculations are liable to be inaccurate. 
% The Fisher's exact test involves the computations of several factorials 
% to obtain the probability of the observed and each of the more extreme tables. 
% Factorials growth quickly, so it's necessary use logarithms of factorials. 
% This computations is very easy in Matlab because:
% x!=gamma(x+1) and log(x!)=gammaln(x+1). 
% Moreover, when the matrix has many Rows and Columns, the computation of all the
% set of possible matrices is very time expensive.
% This function uses this strategy:
% 1) if the input is a 2x2, 2x3, 2x4 or 3x3 matrix it uses (or download) ad hoc,
% previously written by me, function;
% 2) else it uses a Monte Carlo approach.
% Finally, this function uses the Peter J. Acklam rldecode function, and so I
% want to acknowledge him.
%
% Syntax: 	p=myfisher(x)
%      
%     Inputs:
%           X - data matrix 
%           delta - If Monte Carlo simulation is needed, the simulation
%           size to ensure that p-value is within delta units of the true
%           one with (1-alpha)*100% confidence. Psycometrika 1979; Vol.44:75-83.
%           By default delta=0.01;
%           alpha - Significance level. By default alpha=0.05;
%     Outputs:
%           P - 2-tailed p-value
%
%
%           Created by Giuseppe Cardillo
%          
%
% To cite this file, this would be an appropriate format:
% Cardillo G. (2010) MyFisher: the definitive function for the Fisher's exact
% and conditional test for any RxC matrix
% http://www.mathworks.com/matlabcentral/fileexchange/26883


%Input error handling
p = inputParser;
addRequired(p,'x',@(x) validateattributes(x,{'numeric'},{'real','finite','integer','nonnegative','nonnan','2d'}));
addOptional(p,'delta',0.01, @(x) validateattributes(x,{'numeric'},{'scalar','real','finite','nonnan','>',0,'<',1}));
addOptional(p,'alpha',0.05, @(x) validateattributes(x,{'numeric'},{'scalar','real','finite','nonnan','>',0,'<',1}));
parse(p,x,varargin{:});
alpha=p.Results.alpha; delta=p.Results.delta;
clear p

%chech if you can use a previously written function of mine
[rows,columns]=size(x);
% Had to modify this if statement to use the function to assess 2xN inputs.  
% Making a note in concordance with GPU3.  Date: 9/6/19
if (rows==2) && (columns < 5)
% was: if rows==2     
    if columns==2
        assert(exist('myfisher22.m','file')~=0,'You must download myfisher22.m function from https://it.mathworks.com/matlabcentral/fileexchange/15434-myfisher22')
        myfisher22(x,alpha)
    elseif columns==3
        assert(exist('myfisher23.m','file')~=0,'You must download myfisher23.m function from https://it.mathworks.com/matlabcentral/fileexchange/15399-myfisher23')
        myfisher23(x,alpha)
    elseif columns==4
        assert(exist('myfisher24.m','file')~=0,'You must download myfisher24.m function from https://it.mathworks.com/matlabcentral/fileexchange/19842-myfisher24')        
        myfisher24(x,alpha)
    end
elseif rows==3 && columns==3
    assert(exist('myfisher33.m','file')~=0,'You must download myfisher33.m function from https://it.mathworks.com/matlabcentral/fileexchange/15482-myfisher33')
    myfisher33(x,alpha)
else
    clear rows columns

    C=sum(x); %columns sums
    R=sum(x,2); %rows sums
    N=sum(x(:)); %sum of all cells
    Kf=sum(gammaln([R' C]+1))-gammaln(N+1); %The costant factor K=log(prod(R!)*prod(C!)/N!)
    zf=gammaln(x+1); %compute log(x!)
    op=exp(Kf-sum(zf(:))); %compute the p-value of the observed matrix

    %Each matrix can be transformed into a Nx2 matrix:
    % Example:
    %                                       Sex
    %                                Male         Female  R
    %                              ---------------------
    %                    Recovered |   3      |     6   | 9
    %              Response        |----------|---------|
    %                    Deceased  |   8      |     2   | 10
    %                              ---------------------
    %                            C    11           8      19 N
    %
    % In the first cell (R=1 C=1) there are 3 elements;
    % In the second cell (R=1 C=2) there are 6 elements; and so on
    % We can construct this 19x2 matrix:
    % table =
    % 
    %      1     1
    %      1     1
    %      1     1
    %      1     2
    %      1     2
    %      1     2
    %      1     2
    %      1     2
    %      1     2
    %      2     1
    %      2     1
    %      2     1
    %      2     1
    %      2     1
    %      2     1
    %      2     1
    %      2     1
    %      2     2
    %      2     2

    r=1:1:length(R); %create the base array for the first column of the Nx2 matrix
    %(using the example r = 1 2)
    c=repmat(1:1:length(C),1,size(x,1)); %create the base array for the second column of the Nx2 matrix
    %(using the example c = 1 2 1 2)
    table=zeros(N,2); %Nx2 matrix preallocation
    
    table(:,1)=rldecode(R',r); %expand the r array and put it in the first column
    %(using the example the rows sums R = 9 10 and r = 1 2; so we must expand r in
    %this way: 1 must be expanded 9 times and 2 must be expanded 10 times).
    
    tmp=reshape(x',1,[]); %create an array concatenating elements by rows (thanks Jos!)
    %(using the example tmp = 3 6 8 2)
    table(:,2)=rldecode(tmp,c); %expand the c array and put it in the second column
    %(using the example the rows sums c = 1 2 1 2 and tmp=3 6 8 2; so we must expand c in
    %this way: 1 must be expanded 3 times, 2 must be expanded 6 times, 1 must be expanded 8 times and 2 must be expanded 2 times).
    clear R r C c tmp %clear the useless variables

    %Now the Monte Carlo algotithm starts: shuffling the second column we will
    %obtain a new x matrix with the same rows and columns sums of the original.

    %tbs=simulation size to ensure that p-value is within delta units of the true
    %one with (1-alpha)*100% confidence. Psycometrika 1979; Vol.44:75-83.
    tbs=round(((-realsqrt(2)*erfcinv(2-alpha))/(2*delta))^2);
    MCC=0; %Monte Carlo counter
    for I=1:tbs
        %shuffle the second column of table using the Fisher-Yates shuffle Sattolo's
        %version. This is faster than Matlab RANDPERM: to be clearer: Fisher-Yates
        %is O(n) while Randperm is O(nlog(n))
        for J=N:-1:2
            s=ceil((J-1).*rand);
            tmp=table(s,2); table(s,2)=table(J,2); table(J,2)=tmp;
        end
        g=zeros(size(x)); %Construct a new table
        %This cycle is faster than Matlab ACCUMARRAY.
        for J=1:N
            g(table(J,1),table(J,2))=g(table(J,1),table(J,2))+1; %add one to the cell
        end
        zf=gammaln(g+1); %compute log(x!)
        gpv=exp(Kf-sum(zf(:))); %compute the p-value of the new matrix
        if gpv<=op %if the current p-value is less or equal than the observed p-value...
            MCC=MCC+1; %update the counter
        end
    end
    P=MCC/tbs; %Monte Carlo p-value
    tr=repmat('-',1,80);
    disp(tr)
    disp('Fisher''s test - Conventional Monte Carlo Method')
    disp(tr)
    disp(array2table([tbs P],'VariableNames',{'Tables','p_value'}))
    fprintf('p-value is within %0.4f units of the true one with %0.4f%% confidence\n',delta,(1-alpha)*100)
    disp(tr)
end
if nargout
    Pout=P;
end
end

function y = rldecode(len, val)
%RLDECODE Run-length decoding of run-length encode data.
%
%   X = RLDECODE(LEN, VAL) returns a vector XLEN with the length of each run
%   and a vector VAL with the corresponding values.  LEN and VAL must have the
%   same lengths.
%
%   Example: rldecode([ 2 3 1 2 4 ], [ 6 4 5 8 7 ]) will return
%
%      x = [ 6 6 4 4 4 5 8 8 7 7 7 7 ];
%
%   See also RLENCODE.

%   Author:      Peter J. Acklam
%   Time-stamp:  2002-03-03 13:50:38 +0100
%  
%   URL:         http://home.online.no/~pjacklam

% keep only runs whose length is positive
KK = len > 0;
len = len(KK);
val = val(KK);

% now perform the actual run-length decoding
II = cumsum(len);             % LENGTH(LEN) flops
JJ = zeros(1, II(end));
JJ(II(1:end-1)+1) = 1;         % LENGTH(LEN) flops
JJ(1) = 1;
y = val(cumsum(JJ));          % SUM(LEN) flops
end
```
