## Supplementary File 3 for "A surrogate workflow for virtual population development: designing an amenable proxy for a hypothesis-test inspired nonlinear goodness-of-fit": nnls.m

```
function [x,w,info]=nnls(C,d,opts)
% nnls  Non negative least squares Cx=d x>=0 w=C'(d-Cx)<=0
%  2012-08-21  Matlab8  W.Whiten
%  2013-02-17  Line 52 added
%  Copyright (C) 2012, W.Whiten (personal) BSD license
%  (http://opensource.org/licenses/BSD-3-Clause)
%
% [x,w,info]=nnls(C,d,opts)
%  C    Coefficient matrix
%  d    Rhs vector
%  opts Struct containing options: (optional)
%        .Accy  0 fast version, 1 refines final value (default), 
%                 2 uses accurate steps but very slow on large cases, 
%                 faster on small cases, result usually identical to 1
%        .Order True or [], or order to initially include positive terms
%                 if included will supply info.Order, if x0 available use 
%                 find(x0>0), but best saved from previous run of nnls
%        .Tol   Tolerance test value, default zero, use multiple of eps
%        .Iter  Maximum number of iterations, should not be needed.
%
%  x    Positive solution vector x>=0
%  w    Lagrange multiplier vector w(x==0)<= approx zero
%  info Struct with extra information: 
%        .iter  Number of iterations used
%        .wsc0  Estimated size of errors in w
%        .wsc   Maximum of test values for w
%        .Order Order variables used, use to restart nnls with opts.Order
%
% Exits with x>=0 and w<= zero or slightly above 0 due to
%  rounding and to ensure for convergence
% Using faster matrix operations then refines answer as default (Accy 1).
% Accy 0 is more robust in singular cases.
%
% Follows Lawson & Hanson, Solving Least Squares Problems, Ch 23.

[~,n]=size(C);
maxiter=4*n;

% inital values
P=false(n,1);
x=zeros(n,1);
z=x;

w=C'*d;

% wsc_ are scales for errors
wsc0=sqrt(sum(w.^2));
wsc=zeros(n,1);
tol=3*eps;
accy=1;
pn1=0;
pn2=0;
pn=zeros(1,n);

% see if option values have been given
ind=true;
if(nargin>2)
    if(isfield(opts,'Tol'))
        tol=opts.Tol;
        wsc(:)=wsc0*tol;
    end
    if(isfield(opts,'Accy'))
        accy=opts.Accy;
    end
    if(isfield(opts,'Iter'))
        maxiter=opts.Iter;
    end
end

% test if to use normal matrix for speed
if(accy<2)
    A=C'*C;
    b=C'*d;
    %L=zeros(n,n);
    LL=zeros(0,0);
    lowtri=struct('LT',true);
    uptri=struct('UT',true);
end

% test if initial information given
if(nargin>2)
    if(isfield(opts,'Order') && ~islogical(opts.Order))
        pn1=length(opts.Order);
        pn(1:pn1)=opts.Order;
        P(pn(1:pn1))=true;
        ind=false;
    end
    if(~ind && accy<2)
        %L(1:pn1,1:pn1)=chol(A(pn(1:pn1),pn(1:pn1)),'lower');
        UU(1:pn1,1:pn1)=chol(A(pn(1:pn1),pn(1:pn1)));
        LL=UU';
    end
    pn2=pn1;
end

% loop until all positive variables added
iter=0;
while(true)
    
    % Check if no more terms to be added
    if(ind && (all(P==true) || all(w(~P)<=wsc(~P))))
        if(accy~=1)
            break
        end
        accy=2;
        ind=false;
    end

    % skip if first time and initial Order given
    if(ind)
        % select best term to add
        ind1=find(~P);
        [~,ind2]=max(w(ind1)-wsc(ind1));
        ind1=ind1(ind2);
        P(ind1)=true;
        pn2=pn1+1;
        pn(pn2)=ind1;
    end
    
    % loop until all negative terms are removed
    while(true)
        
        % check for divergence
        iter=iter+1;
        if(iter>=2*n)
            if(iter>maxiter)
                error(['nnls Failed to converge in ' num2str(iter)  ...
                    ' iterations'])
                %warning(['nnls Failed to converge in ' num2str(iter)  ...
                %    ' iterations'])
                %return
            elseif(mod(iter,n)==0)
                wsc=(wsc+wsc0*tol)*2;
            end
        end
        
        % solve using suspected positive terms
        z(:)=0;
        if(accy>=2)
            z(P)=C(:,P)\d;
        else
            % add row to the lower triangular factor
            for i=pn1+1:pn2
                i1=i-1;
                %LL=L(1:i1,1:i1);
                %LL=LL(1:i1,1:i1);
                t=linsolve(LL,A(pn(1:i1),pn(i)),lowtri);
                %t=LL\A(pn(1:i1),pn(i));
                %L(i,1:i1)=t;
                %LL(i,1:i1)=t;
                AA=A(pn(i),pn(i));
                tt=AA-t'*t;
                if(tt<=AA*tol)
                    tt=1e300;
                else
                    tt=sqrt(tt);
                end
                %L(i,i)=sqrt(tt);
                %LL(i,i)=sqrt(tt);
                LL(i,1:i)=[t',tt];
                UU(1:i,i)=[t;tt];
            end
            
            % solve using lower triangular factor
            %LL=L(1:pn2,1:pn2);
            t=linsolve(LL,b(pn(1:pn2)),lowtri);
            %t=LL\b(pn(1:pn2));
            %UU=LL';
            %z(pn(1:pn2))=linsolve(UU,t,uptri);
            z(pn(1:pn2))=linsolve(UU,t,uptri);
            %z(pn(1:pn2))=LL'\t;
            % or could use this to solve without updating factors
            %z(pn(1:pn2))=A(pn(1:pn2),pn(1:pn2))\b(pn(1:pn2));
        end
        pn1=pn2;
        
        % check terms are positive
        if(all(z(P)>=0))
            x=z;
            if(accy<2)
                w=b-A*x;
            else
                w=C'*(d-C*x);
            end
            wsc(P)=max(wsc(P),2*abs(w(P)));
            ind=true;
            break
        end
        
        % select and remove worst negative term
        ind1=find(z<0);
        [alpha,ind2]=min(x(ind1)./(x(ind1)-z(ind1)+realmin));
        ind1=ind1(ind2);
        
        % test if removing last added, increase wsc to avoid loop
        if(x(ind1)==0 && ind)
            w=C'*(d-C*z);
            wsc(ind1)=(abs(w(ind1))+wsc(ind1))*2;
        end
        P(ind1)=false;
        x=x-alpha*(x-z);
        pn1=find(pn==ind1);
        pn(pn1:end)=[pn(pn1+1:end),0];
        pn1=pn1-1;
        pn2=pn2-1;
        if(accy<2)
            LL=LL(1:pn1,1:pn1);
            UU=UU(1:pn1,1:pn1);
        end
        ind=true;
    end
end

% info result required
if(nargout>2)
    info.iter=iter;
    info.wsc0=wsc0*eps;
    info.wsc=max(wsc);
    if(nargin>2 && isfield(opts,'Order'))
        info.Order=pn(1:pn1);
    end
end

return
end
```
