## Supplementary File 3 for "A surrogate workflow for virtual population development: designing an amenable proxy for a hypothesis-test inspired nonlinear goodness-of-fit": nnlstest1.m

```
% Test cases for nnls
%  2012-08-22  Matlab8  W.Whiten


disp('C=randn(20,10);')
C=randn(20,10);
disp('C=randn(20,10);')
d=randn(20,1);

x0=nnls(C,d);

%#ok<*NASGU>
disp('tic;[x0,w0,info]=nnls(C,d,struct(''Accy'',0));toc')
tic;[x0,w0,info]=nnls(C,d,struct('Accy',0));toc
disp(['Min x  ', num2str(min(x0)), '    Max w  ', num2str(max(w0(x0==0)))])
disp(' ')
disp('tic;[x1,w1,info]=nnls(C,d,struct(''Accy'',1));toc')
tic;[x1,w1,info]=nnls(C,d,struct('Accy',1));toc
disp(['Min x  ', num2str(min(x1)), '    Max w  ', num2str(max(w1(x1==0)))])
disp(' ')
disp('tic;[x2,w2,info]=nnls(C,d,struct(''Accy'',2));toc')
tic;[x2,w2,info]=nnls(C,d,struct('Accy',2));toc
disp(['Min x  ', num2str(min(x2)), '    Max w  ', num2str(max(w2(x2==0)))])
disp(' ')
disp(' ')

disp('tic;[x0,w0,info]=nnls(C,d,struct(''Accy'',0,''Order'',1:10));toc')
tic;[x0,w0,info]=nnls(C,d,struct('Accy',0,'Order',1:10));toc
disp(['Min x  ', num2str(min(x0)), '    Max w  ', num2str(max(w0(x0==0)))])
disp(' ')
disp('tic;[x1,w1,info]=nnls(C,d,struct(''Accy'',1,''Order'',1:10));toc')
tic;[x1,w1,info]=nnls(C,d,struct('Accy',1,'Order',1:10));toc
disp(['Min x  ', num2str(min(x1)), '    Max w  ', num2str(max(w1(x1==0)))])
disp(' ')
disp('tic;[x2,w2,info]=nnls(C,d,struct(''Accy'',2,''Order'',1:10));toc')
tic;[x2,w2,info]=nnls(C,d,struct('Accy',2,'Order',1:10));toc
disp(['Min x  ', num2str(min(x2)), '    Max w  ', num2str(max(w2(x2==0)))])
disp(' ')
disp(' ')


disp('C=randn(5000,1000);')
C=randn(5000,1000);
disp('d=randn(5000,1);')
d=randn(5000,1);
disp('tic;[x0,w0,info]=nnls(C,d,struct(''Accy'',0,''Order'',[]));toc')
tic;[x0,w0,info]=nnls(C,d,struct('Accy',0,'Order',[]));toc
disp(['Min x  ', num2str(min(x0)), '    Max w  ', num2str(max(w0(x0==0)))])
disp(' ')
disp('tic;[x1,w1,info]=nnls(C,d,struct(''Accy'',1,''Order'',[]));toc')
tic;[x1,w1,info]=nnls(C,d,struct('Accy',1,'Order',[]));toc
disp(['Min x  ', num2str(min(x0)), '    Max w  ', num2str(max(w0(x0==0)))])
disp(' ')
disp('About 3 mintues (about 100 times longer then last)')
disp('tic;[x2,w2,info]=nnls(C,d,struct(''Accy'',2,''Order'',[]));toc')
tic;[x2,w2,info]=nnls(C,d,struct('Accy',2,'Order',[]));toc %#ok<ASGLU>
disp(['Min x  ', num2str(min(x0)), '    Max w  ', num2str(max(w0(x0==0)))])
disp(' ')
disp('tic;[x0,w0,info]=nnls(C,d,struct(''Accy'',0,''Order'',info.Order));toc')
tic;[x0,w0,info]=nnls(C,d,struct('Accy',0,'Order',info.Order));toc
disp(['Min x  ', num2str(min(x0)), '    Max w  ', num2str(max(w0(x0==0)))])
disp(' ')
disp('tic;[x2,w2,info]=nnls(C,d,struct(''Accy'',2,''Order'',info.Order));toc')
tic;[x2,w2,info]=nnls(C,d,struct('Accy',2,'Order',info.Order));toc
disp(['Min x  ', num2str(min(x2)), '    Max w  ', num2str(max(w2(x2==0)))])
disp(' ')

disp('lsqnoneg is also slow on this case')
disp('tic;x=lsqnonneg(C,d);tic')
tic;x=lsqnonneg(C,d);toc
disp(['Min x  ', num2str(min(x)), '    Difference x-x1  ', num2str(max(abs(x1-x)))])
disp(' ')


% nnlstest1
% C=randn(20,10);
% C=randn(20,10);
% Undefined function or variable 'D'.
% Error in nnlstest1 (line 10)
% x0=nnls(D,c); 
% nnlstest1
% C=randn(20,10);
% C=randn(20,10);
% tic;[x0,w0,info]=nnls(C,d,struct('Accy',0));toc
% Elapsed time is 0.001989 seconds.
% Min x  0    Max w  -2.5133
%  
% tic;[x1,w1,info]=nnls(C,d,struct('Accy',1));toc
% Elapsed time is 0.002882 seconds.
% Min x  0    Max w  -2.5133
%  
% tic;[x2,w2,info]=nnls(C,d,struct('Accy',2));toc
% Elapsed time is 0.001377 seconds.
% Min x  0    Max w  -2.5133
%  
%  
% tic;[x0,w0,info]=nnls(C,d,struct('Accy',0,'Order',1:10));toc
% Elapsed time is 0.002009 seconds.
% Min x  0    Max w  -2.5133
%  
% tic;[x1,w1,info]=nnls(C,d,struct('Accy',1,'Order',1:10));toc
% Elapsed time is 0.001952 seconds.
% Min x  0    Max w  -2.5133
%  
% tic;[x2,w2,info]=nnls(C,d,struct('Accy',2,'Order',1:10));toc
% Elapsed time is 0.001060 seconds.
% Min x  0    Max w  -2.5133
%  
%  
% C=randn(5000,1000);
% d=randn(5000,1);
% tic;[x0,w0,info]=nnls(C,d,struct('Accy',0,'Order',[]));toc
% Elapsed time is 1.987874 seconds.
% Min x  0    Max w  -0.026687
%  
% tic;[x1,w1,info]=nnls(C,d,struct('Accy',1,'Order',[]));toc
% Elapsed time is 3.206198 seconds.
% Min x  0    Max w  -0.026687
%  
% About 3 mintues (about 100 times longer then last)
% tic;[x2,w2,info]=nnls(C,d,struct('Accy',2,'Order',[]));toc
% Elapsed time is 229.973385 seconds.
% Min x  0    Max w  -0.026687
%  
% tic;[x0,w0,info]=nnls(C,d,struct('Accy',0,'Order',info.Order));toc
% Elapsed time is 0.419892 seconds.
% Min x  0    Max w  -0.026687
%  
% tic;[x2,w2,info]=nnls(C,d,struct('Accy',2,'Order',info.Order));toc
% Elapsed time is 1.279451 seconds.
% Min x  0    Max w  -0.026687
%  
% lsqnoneg is also slow on this case
% tic;x=lsqnonneg(C,d);tic
% Elapsed time is 230.101080 seconds.
% Min x  0    Difference x-x1  0
```
