## Supplementary File 3 for "A surrogate workflow for virtual population development: designing an amenable proxy for a hypothesis-test inspired nonlinear goodness-of-fit": Readme.txt

nnls - Alternative to lsqnonneg: can be faster on large problems,
 improved convergence control, optional restart vector

Solves non negative least squares:
    min wrt x: (d-Cx)'*(d-Cx) subject to: x>=0

This version of nnls aims to solve convergance problems that can occur 
with the 2011-2012 version of lsqnonneg, and provides a fast solution of 
large problems. Includes an option to give initial positive terms for x 
for faster solution of iterative problems using nnls. 

For some large problems nnls can be faster than lsqnonneg, 
see test file (nnlstest.m).


Simple usage:  x=nnls(C,d)

[x,w,info]=nnls(C,d,opts)
 C    Coefficient matrix
 d    Rhs vector
 opts Struct containing options: (optional)
       .Accy  0 fast version, 1 refines final value (default), 
                2 uses accurate steps but  very slow on large cases, 
                faster on small cases, result usually identical to 1
       .Order True or [], or order to initially include positive terms
                if included will supply info.Order, if x0 available use 
                find(x0>0), but best saved from previous run of nnls
       .Tol   Tolerance test value, default zero, use multiple of eps
       .Iter  Maximum number of iterations, should not be needed.

 x    Positive solution vector x>=0
 w    Lagrange multiplier vector w(x==0)<= approx zero
 info Struct with extra information: 
       .iter  Number of iterations used
       .wsc0  Estimated size of errors in w
       .wsc   Maximum of test values for w
       .Order Order variables used, use to restart nnls with opts.Order

Examples in nnlstest.m
