## Supplementary File 3 for "A surrogate workflow for virtual population development: designing an amenable proxy for a hypothesis-test inspired nonlinear goodness-of-fit": weightedcorrs.m

```
function R = weightedcorrs(Y, w)
%
%   WEIGHTEDCORRS returns a symmetric matrix R of weighted correlation
%   coefficients calculated from an input T-by-N matrix Y whose rows are
%   observations and whose columns are variables and an input T-by-1 vector
%   w of weights for the observations. This function may be a valid
%   alternative to CORRCOEF if observations are not all equally relevant
%   and need to be weighted according to some theoretical hypothesis or
%   knowledge.
%
%   R = WEIGHTEDCORRS(Y, w) returns a positive semidefinite matrix R,
%   i.e. all its eigenvalues are non-negative (see Example 1).
%
%   WEIGHTEDCORRS is such that
%          WEIGHTEDCORRS(Y, w) == WEIGHTEDCORRS(a * Y + b, w)
%   where a and b are two real numbers (see Example 1).
%
%   Furthermore, the result provided by the function doesn't change if the
%   unit system of each column of Y is changed through an arbitrary affine
%   transformation y = a * x + b, where a and b are two real numbers, with
%   a > 0 (see Example 2).
%
%   If w = ones(size(Y, 1), 1), no difference exists between
%   WEIGHTEDCORRS(Y, w) and CORRCOEF(Y) (see Example 4).
%
%
%   REFERENCE: the mathematical formulas in matrix notation, together with
%   the code, is also available in
%   F. Pozzi, T. Di Matteo, T. Aste,
%   "Exponential smoothing weighted correlations",
%   The European Physical Journal B, Volume 85, Issue 6, 2012.
%   DOI:10.1140/epjb/e2012-20697-x. 
%
% % ======================================================================
% % EXAMPLE 0: some common shapes in 2D.
% % ======================================================================
%
%   T = 1000;                                                                     % number of observations
%   % CHOOSE WEIGHTS
%   alpha = 2 / T;
%   w0 = 1 / sum(exp(((1:T) - T) * alpha));
%   w = w0 * exp(((1:T) - T) * alpha);                                            % weights: exponential decay
%
% %%%%%%%%%%%%%%%%%%%%%%%%%%%%%%%%%%%%%%%%%%%%%%%%%%%%%%%%%%%%%%%%%%%%%%%%%
% % a) A line is always a line!
%   Y(:, 1) = 1:T;
%   Y(:, 2) = rand * Y(:, 1) + rand;                                              % Linear relation
%   r1 = corrcoef(Y);
%   r1 = r1(2)                                                                    % Traditional Correlation
%   r2 = weightedcorrs(Y, w); 
%   r2 = r2(2)                                                                    % Weighted Correlation
% %%%%%%%%%%%%%%%%%%%%%%%%%%%%%%%%%%%%%%%%%%%%%%%%%%%%%%%%%%%%%%%%%%%%%%%%%
% % b) An horizontal line has a correlation equal to 0
%   Y(1:T, 1) = rand;
%   Y(:, 2) = 1:T;                                                                % Linear relation
%   r1 = corrcoef(Y);
%   r1 = r1(2)                                                                    % Traditional Correlation
%   r2 = weightedcorrs(Y, w); 
%   r2 = r2(2)                                                                    % Weighted Correlation
% %%%%%%%%%%%%%%%%%%%%%%%%%%%%%%%%%%%%%%%%%%%%%%%%%%%%%%%%%%%%%%%%%%%%%%%%%
% % c) A vertical line has a correlation equal to 0
%   Y(:, 1) = 1:T;
%   Y(:, 2) = rand;                                                               % Linear relation
%   r1 = corrcoef(Y);
%   r1 = r1(2)                                                                    % Traditional Correlation
%   r2 = weightedcorrs(Y, w); 
%   r2 = r2(2)                                                                    % Weighted Correlation
% %%%%%%%%%%%%%%%%%%%%%%%%%%%%%%%%%%%%%%%%%%%%%%%%%%%%%%%%%%%%%%%%%%%%%%%%%
% % d) A symmetric parabolic shape ... makes a huge difference!
%   Y(:, 1) = (1:T) - T / 2 - 0.5;
%   Y(:, 2) = rand * Y(:, 1) .^ 2 + rand;                                         % Parabolic relation (symmetric)
%   r1 = corrcoef(Y);
%   r1 = r1(2)                                                                    % Traditional Correlation
%   r2 = weightedcorrs(Y, w); 
%   r2 = r2(2)                                                                    % Weighted Correlation
% %%%%%%%%%%%%%%%%%%%%%%%%%%%%%%%%%%%%%%%%%%%%%%%%%%%%%%%%%%%%%%%%%%%%%%%%%
% % e) A circle
%   angle = (2 * rand(T, 1) - 1) * pi;
%   rho = rand(T, 1);
%   Y(:, 1) = rho .* cos(angle);
%   Y(:, 2) = rho .* sin(angle);                                                  % Circle
%   r1 = corrcoef(Y);
%   r1 = r1(2)                                                                    % Traditional Correlation
%   r2 = weightedcorrs(Y, w); 
%   r2 = r2(2)                                                                    % Weighted Correlation
% %%%%%%%%%%%%%%%%%%%%%%%%%%%%%%%%%%%%%%%%%%%%%%%%%%%%%%%%%%%%%%%%%%%%%%%%%
% % f) An exponential curve
%   Y(:, 1) = 1:T;
%   Y(:, 2) = exp(3 * (1:T) / T);                                                 % Exponential relation
%   r1 = corrcoef(Y);
%   r1 = r1(2)                                                                    % Traditional Correlation
%   r2 = weightedcorrs(Y, w); 
%   r2 = r2(2)                                                                    % Weighted Correlation
% %%%%%%%%%%%%%%%%%%%%%%%%%%%%%%%%%%%%%%%%%%%%%%%%%%%%%%%%%%%%%%%%%%%%%%%%%
% % g) A logarithm
%   Y(:, 1) = 1:T;
%   Y(:, 2) = log(1:T);                                                           % Logarithmic relation
%   r1 = corrcoef(Y);
%   r1 = r1(2)                                                                    % Traditional Correlation
%   r2 = weightedcorrs(Y, w); 
%   r2 = r2(2)                                                                    % Weighted Correlation
% %%%%%%%%%%%%%%%%%%%%%%%%%%%%%%%%%%%%%%%%%%%%%%%%%%%%%%%%%%%%%%%%%%%%%%%%%
%
%
% % ======================================================================
% % EXAMPLE 1: verify some of the properties for weighted correlations.
% % ======================================================================
%
% % GENERATE CORRELATED STOCHASTIC PROCESSES
%   N = 500;                                                                      % number of variables
%   T = 1000;                                                                     % number of observations
%   Y = randn(T, N);                                                              % shocks from standardized normal distribution
%   lambda = 2;
%   Y = Y + repmat(poissrnd(lambda, T, 1) .* (2 * rand(T, 1) - 1), 1, N);         % shocks plus common factor
  % Y = cumsum(Y);                                                                % correlated stochastic processes
%
% % CHOOSE WEIGHTS
%   alpha = 2 / T;
%   w0 = 1 / sum(exp(((1:T) - T) * alpha));
%   w = w0 * exp(((1:T) - T) * alpha);                                            % weights: exponential decay
%
% % COMPUTE CORRELATION MATRIX
%   r1 = weightedcorrs(Y, w);                                                     % Weighted Correlation matrix for Y
%
% % COMPUTE CORRELATION MATRIX FOR MODIFIED DATA
%   a = 1000 * (2 * rand - 1);                                                    % Multiplicative coefficient
%   b = 1000 * (2 * rand - 1);                                                    % Shift
%   r2 = weightedcorrs(a * Y + b, w);                                             % Weighted Correlation matrix for modified Y
%
% % FIND RELEVANT INDEXES AND PLOT A SCATTER DIAGRAM
%   I = ones(N);
%   indexes = find(tril(I, -1));                                                  % Indexes of lower triangular square matrix
%
%   figure('Position', [50   50   950   630]);
%   plot([-1:1], [-1:1], 'r', 'LineWidth', 10);                                   % 45� Line
%   hold on;
%   plot(r1(indexes), r2(indexes), '*', 'MarkerSize', 6);                         % Comparison between the coefficients of the two Matrices
%   title('Identical correlations for Y and a * Y + b', ...                       % title label for the plot
%       'FontSize', 20, 'FontWeight', 'Bold');                                    % ... font properties
%   xlabel('Correlations for Y', 'FontSize', 16, 'FontWeight', 'Bold');
%   ylabel('Correlations for modified Y = a * Y + b', ...
%       'FontSize', 16, 'FontWeight', 'Bold');
%   set(gca, 'FontSize', 14, 'FontWeight', 'Bold');
%   grid on;
%
% % PLOT HISTOGRAM OF DIFFERENCES BETWEEN COEFFICIENTS
% % OF THE TWO CORRELATION MATRICES
%   figure('Position', [50   50   950   630]);
%   hist(r1(indexes) - r2(indexes), 100);                                         % Histogram of the differences (which are always negligible)
%   title('Identical correlations for Y and a * Y + b', ...                       % title label for the plot ...
%       'FontSize', 20, 'FontWeight', 'Bold');                                    % ... font properties
%   xlabel('Negligible differences', 'FontSize', 16, 'FontWeight', 'Bold');
%   ylabel('Absolute Frequencies', 'FontSize', 16, 'FontWeight', 'Bold');
%   set(gca, 'FontSize', 14, 'FontWeight', 'Bold');
%   grid on;
%
% % VERIFY SYMMETRY
%   all(all(r1 == r1'))
%   all(all(r2 == r2'))
%
% % VERIFY POSITIVENESS FOR ALL EIGENVALUES
%   all(eig(r1) >= 0)
%   all(eig(r2) >= 0)
%
%
% % ======================================================================
% % EXAMPLE 2: the unit system of each column of Y is changed through
% % arbitrary affine transformations.
% % ======================================================================
%
% % GENERATE CORRELATED STOCHASTIC PROCESSES
%   N = 500;                                                                      % number of variables
%   T = 1000;                                                                     % number of observations
%   Y = randn(T, N);                                                              % shocks from standardized normal distribution
%   lambda = 2;
%   Y = Y + repmat(poissrnd(lambda, T, 1) .* (2 * rand(T, 1) - 1), 1, N);         % shocks plus common factor
%   Y = cumsum(Y);                                                                % correlated stochastic processes
%
% % CHOOSE WEIGHTS
%   alpha = 2 / T;
%   w0 = 1 / sum(exp(((1:T) - T) * alpha));
%   w = w0 * exp(((1:T) - T) * alpha);                                            % weights: exponential decay
%
% % COMPUTE CORRELATION MATRIX
%   r1 = weightedcorrs(Y, w);                                                     % Weighted Correlation matrix for Y
%
% % COMPUTE CORRELATION MATRIX FOR MODIFIED DATA
%   a = 1000 * rand(1, N);                                                        % Multiplicative coefficients (positive)
%   b = 1000 * (2 * rand(1, N) - 1);                                              % Shifts
%   r2 = weightedcorrs(repmat(a, T, 1) .* Y + repmat(b, T, 1), w);                % Weighted Correlation matrix for modified Y
%
% % FIND RELEVANT INDEXES AND PLOT A SCATTER DIAGRAM
%   I = ones(N);
%   indexes = find(tril(I, -1));                                                  % Indexes of lower triangular square matrix
%
%   figure('Position', [50   50   950   630]);
%   plot([-1:1], [-1:1], 'r', 'LineWidth', 10);                                   % 45� Line
%   hold on;
%   plot(r1(indexes), r2(indexes), '*', 'MarkerSize', 6);                         % Comparison between the coefficients of the two Matrices
%   str = 'Identical correlations after arbitrary affine transformations';
%   title(str, 'FontSize', 20, 'FontWeight', 'Bold');                             % title label for the plot
%   xlabel('Correlations for Y', 'FontSize', 16, 'FontWeight', 'Bold');
%   ylabel('Correlations for modified Y', ...
%       'FontSize', 16, 'FontWeight', 'Bold');
%   set(gca, 'FontSize', 14, 'FontWeight', 'Bold');
%   grid on;
%
% % PLOT HISTOGRAM OF DIFFERENCES BETWEEN COEFFICIENTS
% % OF THE TWO CORRELATION MATRICES
%   figure('Position', [50   50   950   630]);
%   hist(r1(indexes) - r2(indexes), 100);                                         % Histogram of the differences (which are always negligible)
%   title('Identical correlations for Y and modified Y', ...                      % title label for the plot
%       'FontSize', 20, 'FontWeight', 'Bold');                                    % ... font properties
%   xlabel('Negligible differences', 'FontSize', 16, 'FontWeight', 'Bold');
%   ylabel('Absolute Frequencies', 'FontSize', 16, 'FontWeight', 'Bold');
%   set(gca, 'FontSize', 14, 'FontWeight', 'Bold');
%   grid on;
%
%
% % ======================================================================
% % Example 3: differences with respect to traditional correlations.
% % ======================================================================
%
% % GENERATE CORRELATED STOCHASTIC PROCESSES
%   N = 500;                                                                      % number of variables
%   T = 1000;                                                                     % number of observations
%   Y = randn(T, N);                                                              % shocks from standardized normal distribution
%   lambda = 2;
%   Y = Y + repmat(poissrnd(lambda, T, 1) .* (2 * rand(T, 1) - 1), 1, N);         % shocks plus common factor
%   Y = cumsum(Y);                                                                % correlated stochastic processes
%
% % CHOOSE WEIGHTS AND PLOT THEM
%   alpha = 2 / T;
%   w0 = 1 / sum(exp(((1:T) - T) * alpha));
%   w = w0 * exp(((1:T) - T) * alpha);                                            % weights: exponential decay
%
%   figure('Position', [50   50   950   630]);
%   plot([floor(-T / 2):ceil(T + T / 2)], ...                                     % x values ...
%       w0 * exp(((floor(-T / 2):ceil(T + T / 2)) - T) * alpha), ...              % ... y values ...
%       '.r', 'MarkerSize', 7);                                                   % ... plot properties
%   hold on;
%   plot(w, '.', 'MarkerSize', 30);
%   title('Exponential weights assigned to observations', ...                     % title label for the plot
%       'FontSize', 20, 'FontWeight', 'Bold');                                    % ... font properties
%   xlabel('Observations', 'FontSize', 16, 'FontWeight', 'Bold');
%   ylabel('Weights', 'FontSize', 16, 'FontWeight', 'Bold');
%   set(gca, 'FontSize', 14, 'FontWeight', 'Bold');
%   grid on;
%
% % COMPUTE CORRELATION MATRICES
%   r1 = weightedcorrs(Y, w);                                                     % Weighted Correlation matrix
%   r2 = corrcoef(Y);                                                             % Traditional Correlation matrix
%
% % FIND RELEVANT INDEXES AND PLOT A SCATTER DIAGRAM
%   I = ones(N);
%   indexes = find(tril(I, -1));                                                  % Indexes of lower triangular square matrix
%
%   figure('Position', [50   50   950   630]);
%   plot(r1(indexes), r2(indexes), '.');                                          % Comparison with the Traditional Correlation matrix
%   hold on;
%   plot([-1:1], [-1:1], 'r', 'LineWidth', 5);                                    % 45� Line
%   title('Scatter Diagram for Traditional and Weighted Correlations', ...        % title label for the plot ...
%       'FontSize', 20, 'FontWeight', 'Bold');                                    % ... font properties
%   xlabel('Weighted Correlation coefficients', ...                               % x label ...
%       'FontSize', 16, 'FontWeight', 'Bold');                                    % ... font properties
%   ylabel('Traditional Correlation coefficients', ...                            % y label ...
%       'FontSize', 16, 'FontWeight', 'Bold');                                    % ... font properties
%   set(gca, 'FontSize', 14, 'FontWeight', 'Bold');
%   grid on;
%
% % PLOT HISTOGRAM OF DIFFERENCES BETWEEN COEFFICIENTS
% % OF THE TWO CORRELATION MATRICES
%   figure('Position', [50   50   950   630]);
%   hist(r2(indexes) - r1(indexes), 100);                                         % Differences between the two correlation matrices
%   title('Differences between Traditional and Weighted Correlations', ...        % title label for the plot ...
%       'FontSize', 20, 'FontWeight', 'Bold');                                    % ... font properties
%   ylabel('Absolute Frequencies', 'FontSize', 16, 'FontWeight', 'Bold');
%   xlabel('Differences between Traditional and Weighted Correlations', ...       % x label ...
%       'FontSize', 16, 'FontWeight', 'Bold');                                    % ... font properties
%   set(gca, 'FontSize', 14, 'FontWeight', 'Bold');
%   grid on;
%
%
% % ======================================================================
% % Example 4: If weights are uniform, then the traditional correlation
% % matrix is obtained.
% % ======================================================================
%
% % GENERATE CORRELATED STOCHASTIC PROCESSES
%   N = 500;                                                                      % number of variables
%   T = 1000;                                                                     % number of observations
%   Y = randn(T, N);                                                              % shocks from standardized normal distribution
%   lambda = 2;
%   Y = Y + repmat(poissrnd(lambda, T, 1) .* (2 * rand(T, 1) - 1), 1, N);         % shocks plus common factor
%   Y = cumsum(Y);                                                                % correlated stochastic processes
%
% % CHOOSE WEIGHTS AND PLOT THEM
%   w = ones(T, 1) / T;                                                           % weights: uniform
%
%   figure('Position', [50   50   950   630]);
%   plot(w, '*', 'MarkerSize', 7);
%   title('Uniform Weights assigned to observations', ...                         % title label for the plot ...
%       'FontSize', 20, 'FontWeight', 'Bold');                                    % ... font properties
%   xlabel('Observations', 'FontSize', 16, 'FontWeight', 'Bold');
%   ylabel('Weights', 'FontSize', 16, 'FontWeight', 'Bold');
%   axis([0 T 0 2 * w(1)]);
%   set(gca, 'FontSize', 14, 'FontWeight', 'Bold');
%   grid on;
%
% % COMPUTE CORRELATION MATRICES
%   r1 = weightedcorrs(Y, w);                                                     % Weighted Correlation matrix
%   r2 = corrcoef(Y);                                                             % Traditional Correlation matrix
%
% % FIND RELEVANT INDEXES AND PLOT A SCATTER DIAGRAM
%   I = ones(N);
%   indexes = find(tril(I, -1));                                                  % Indexes of lower triangular square matrix
%
%   figure('Position', [50   50   950   630]);
%   plot([-1:1], [-1:1], 'r', 'LineWidth', 10);                                   % 45� Line
%   hold on;
%   plot(r1(indexes), r2(indexes), '*', 'MarkerSize', 6);                         % Comparison with the Traditional Correlation matrix
%   title('Scatter Diagram for Traditional and Weighted Correlations', ...        % title label for the plot ...
%       'FontSize', 20, 'FontWeight', 'Bold');                                    % ... font properties
%   xlabel('Weighted Correlation coefficients (uniform weights!!!)', ...          % x label ...
%       'FontSize', 16, 'FontWeight', 'Bold');                                    % ... font properties
%   ylabel('Traditional Correlation coefficients', ...                            % y label ...
%       'FontSize', 16, 'FontWeight', 'Bold');                                    % ... font properties
%   set(gca, 'FontSize', 14, 'FontWeight', 'Bold');
%   grid on;
%
% % PLOT HISTOGRAM OF DIFFERENCES BETWEEN COEFFICIENTS
% % OF THE TWO CORRELATION MATRICES
%   figure('Position', [50   50   950   630]);
%   hist(r2(indexes) - r1(indexes), 100);                                         % Difference between the two correlation matrices
%   temp = 'Differences between Traditional and Weighted ';
%   temp(2, :) = 'Correlation coefficients (uniform weights!!!)';
%   title(temp, 'FontSize', 20, 'FontWeight', 'Bold');
%   ylabel('Absolute Frequencies', 'FontSize', 16, 'FontWeight', 'Bold');
%   xlabel(temp, 'FontSize', 16, 'FontWeight', 'Bold');
%   set(gca, 'FontSize', 14, 'FontWeight', 'Bold');
%   grid on;
%
%
% % ======================================================================
% % Example 5: more reliable dynamic correlations (it may take some mins):
% % sensitive with respect to recent shocks, not to remote ones!
% % ======================================================================
%
% % GENERATE CORRELATED STOCHASTIC PROCESSES
%   N = 500;                                                                      % number of variables
%   T = 1000;                                                                     % number of observations
%   Y = randn(T, N);                                                              % shocks from standardized normal distribution
%   lambda = 2;
%   Y = Y + repmat(poissrnd(lambda, T, 1) .* (2 * rand(T, 1) - 1), 1, N);         % shocks plus common factor
%   Y = cumsum(Y);                                                                % correlated stochastic processes
%
% % CHOOSE WEIGHTS
%   delta = 100;                                                                  % Running window
%   alpha = 2 / delta;
%   w0 = 1 / sum(exp(((1:delta) - delta) * alpha));
%   w = w0 * exp(((1:delta) - delta) * alpha);                                    % weights: exponential decay
%
% % FIND RELEVANT INDEXES AND PLOT A SCATTER DIAGRAM
%   I = ones(N);
%   indexes = find(tril(I, -1));                                                  % indexes of lower triangular square matrix
%
% % COMPUTE DYNAMIC CORRELATION MATRICES
%   for i = 1:(T - delta + 1)
%     temp = weightedcorrs(Y(i:(delta + i - 1), :), w);                           % Dynamic Weighted Correlation matrix
%     r1(i) = mean(temp(indexes));                                                % consider only the lower matrix, in vectorial form, compute the mean
%     temp = corrcoef(Y(i:(delta + i - 1), :));                                   % Dynamic Traditional Correlation matrix
%     r2(i) = mean(temp(indexes));                                                % consider only the lower matrix, in vectorial form, compute the mean
%   end
%
% % PLOT THE AVERAGE CORRELATIONS, BOTH WEIGHTED AND TRADITIONAL
%   figure('Position', [50   50   950   630]);
%   plot(r2, '.g', 'MarkerSize', 24);                                             % the red curve is less sensitive with respect
%   hold on;                                                                      % to remote shocks at time (t - delta + 1) and more
%   plot(r1, '.r', 'MarkerSize', 18);                                             % sensitive with respect to present shocks at time t
%   title('Moving Average Correlations', ...                                      % title label for the plot ...
%       'FontSize', 20, 'FontWeight', 'Bold');                                    % ... font properties
%   xlabel('Time', 'FontSize', 16, 'FontWeight', 'Bold');
%   temp = '       Moving Average Correlations       ';
%   temp(2, :) = '(green --> traditional; red --> weighted)';
%   ylabel(temp, 'FontSize', 16, 'FontWeight', 'Bold');
%   set(gca, 'FontSize', 14, 'FontWeight', 'Bold');
%   grid on;
%
%
% % ======================================================================
% % Example 6: Bell-shaped weights centered on single observations
% % ======================================================================
%
% % GENERATE CORRELATED STOCHASTIC PROCESSES
%   N = 2;                                                                        % number of variables
%   T = 1000;                                                                     % number of observations
%   Y = randn(T, N);                                                              % shocks from standardized normal distribution
%   lambda = 2;
%   Y = Y + repmat(poissrnd(lambda, T, 1) .* (2 * rand(T, 1) - 1), 1, N);         % shocks plus common factor
%   Y = cumsum(Y);                                                                % correlated stochastic processes
%   alpha = 16 / T / T;
%
% % PLOT BELL-SHAPED WEIGHTS IN THREE CASES: AT THE BEGINNING,
% % IN THE MIDDLE AND AT THE END OF THE PERIOD
%   figure('Position', [50   50   950   630]);
%   i = 1;
%   w = exp(-alpha * ((1:T) - i) .^ 2);
%   w = w / sum(w);
%   plot(w, 'g', 'LineWidth', 7);
%   hold on;
%   i = floor(T / 2);
%   w = exp(-alpha * ((1:T) - i) .^ 2);
%   w = w / sum(w);
%   plot(w, 'r', 'LineWidth', 7);
%   i = T;
%   w = exp(-alpha * ((1:T) - i) .^ 2);
%   w = w / sum(w);
%   plot(w, 'b', 'LineWidth', 7);
%   title('Weights assigned to observations', ...                                 % title label for the plot ...
%       'FontSize', 20, 'FontWeight', 'Bold');                                    % ... font properties
%   xlabel('Observations', 'FontSize', 16, 'FontWeight', 'Bold');
%   ylabel('Weights', 'FontSize', 16, 'FontWeight', 'Bold');
%   set(gca, 'FontSize', 14, 'FontWeight', 'Bold');
%   grid on;
%   legend('Weights centered on t = 1', ...                                       % Legend: first item ...
%       sprintf('Weights centered on t = %d', floor(T / 2)), ...                  % ... second item ...
%       sprintf('Weights centered on t = %d', T), ...                             % ... third item ...
%       'Location', 'North');                                                     % Legend Location
%
% % COMPUTE CORRELATIONS FOR EACH POSSIBLE CENTRAL OBSERVATION
%   for i = 1:T
%      w = exp(-alpha * ((1:T) - i) .^ 2);                                        % weights: exponential decay
%      w = w / sum(w);
%      temp = weightedcorrs(Y, w);                                                % Weighted Correlation matrix
%      r(i) = temp(2);
%   end
%   temp = corrcoef(Y);
%   rr = temp(2);
%
% % PLOT CENTERED CORRELATION FOR EACH POSSIBLE CENTRAL OBSERVATION
%   figure('Position', [50   50   950   630]);
%   plot([1, T], [rr, rr], 'b', 'LineWidth', 10);
%   hold on;
%   plot(r, '.r', 'MarkerSize', 24);
%   title('Correlations with bell-shaped weights', ...                            % title label for the plot ...
%       'FontSize', 20, 'FontWeight', 'Bold');                                    % ... font properties
%   xlabel('Central Observation', 'FontSize', 16, 'FontWeight', 'Bold');
%   ylabel(['   Correlations with bell-shaped weights  '; ...                     % y label, first line ...
%       'centered on different Central Observations'], ...                        % y label, second line ...
%       'FontSize', 16, 'FontWeight', 'Bold');                                    % ... font properties
%   set(gca, 'FontSize', 14, 'FontWeight', 'Bold');
%   grid on;
%
% % PLOT HISTOGRAM OF CENTERED CORRELATIONS COMPUTED
% % FOR EACH POSSIBLE CENTRAL OBSERVATION
%   figure('Position', [50   50   950   630]);
%   hist(r, T / 20);
%   title(['     Histogram of the correlation computed with      '; ...           % title label, first line ...
%       'bell-shaped weights on different Central Observations'], ...             % title label, second line ...
%       'FontSize', 20, 'FontWeight', 'Bold');                                    % ... font properties
%   ylabel('Absolute Frequencies', 'FontSize', 16, 'FontWeight', 'Bold');
%   xlabel('Correlation', 'FontSize', 16, 'FontWeight', 'Bold');
%   set(gca, 'FontSize', 14, 'FontWeight', 'Bold');
%   grid on;
%
%
% % ======================================================================
%
%   See also CORRCOEF, COV, STD, MEAN.
%
% % ======================================================================
%
%-*-* -*-* -*-* -*-* -*-* -*-* -*-* -*-* -*-* -*-* -*-* -*-* -*-* -*-* -*-* -*-*%
%                                                                                               %
%            Author: Liber Eleutherios                                             %
%            E-Mail:                             %
%            Date: 23 July 2008                                                      %
%            Updated: 6 June 2012                                                 %
%                                                                                               %
%-*-* -*-* -*-* -*-* -*-* -*-* -*-* -*-* -*-* -*-* -*-* -*-* -*-* -*-* -*-* -*-*%
%
% % ======================================================================
%

% Check input
% Modifed to allow for zeros
ctrl = isvector(w) & isreal(w) & ~any(isnan(w)) & ~any(isinf(w)) & all(w >= 0);
if ctrl
  w = w(:) / sum(w);                                                          % w is column vector
else
  error('Check w: it needs be a vector of real non-negative numbers with no infinite or nan values!')
end
ctrl = isreal(Y) & ~any(isnan(Y)) & ~any(isinf(Y)) & (size(size(Y), 2) == 2);
if ~ctrl
  error('Check Y: it needs be a 2D matrix of real numbers with no infinite or nan values!')
end
ctrl = length(w) == size(Y, 1);
if ~ctrl
  error('size(Y, 1) has to be equal to length(w)!')
end

[T, N] = size(Y);                                                             % T: number of observations; N: number of variables
temp = Y - repmat(w' * Y, T, 1);                                              % Remove mean (which is, also, weighted)
temp = temp' * (temp .* repmat(w, 1, N));                                     % Covariance Matrix (which is weighted)
temp = 0.5 * (temp + temp');                                                  % Must be exactly symmetric
R = diag(temp);                                                               % Variances
R = temp ./ sqrt(R * R');                                                     % Matrix of Weighted Correlation Coefficients
```
