## Supplementary File 3 for "A surrogate workflow for virtual population development: designing an amenable proxy for a hypothesis-test inspired nonlinear goodness-of-fit": initQSPToolbox.m

```
function initQSPtoolbox()
% Initialize Quantiative Systems Pharmacology Toolbox.  This script should
% be run from the command line when you want to start using the toolbox.
% ARGUMENTS
% None
%
% RETURNS
% None
%

% Get the toolbox directory
rootPath=which('initQSPToolbox.m');
global QSPTDIR
QSPTDIR = rootPath(1:end-(length('initQSPToolbox.m')+1));

% Add QSP toolbox paths
path(path,[QSPTDIR, filesep, 'external',filesep,'genpath_exclude']);
myPaths = genpath_exclude(QSPTDIR,{'docs','\.svn'});
if (~isdeployed)
    addpath(myPaths);
end
```
