## Supplementary File 3 for "A surrogate workflow for virtual population development: designing an amenable proxy for a hypothesis-test inspired nonlinear goodness-of-fit": loadMapelOptions.m

```
function myMapelOptions = loadMapelOptions(myFileName, myPath, myFormat)
% Load a mapelOptions object from file.
%
% ARGUMENTS
% fileName: a filename, suffix will be appended based on format
% path:     (optional) save file path
% format:   (optional)save file format, currently only support 'mat'
%
% RETURNS
% myMapelOptions
%

% Perform initial checks on the provided arguments
flagContinue = true;
if nargin > 3
    warning(['Too many arguments provided to ',mfilename,', require: fileName, and optionally path, format.'])
    flagContinue = false;
elseif nargin > 2
    flagContinue = true;
elseif nargin > 1
    myFormat = 'mat';
    flagContinue = true;
elseif nargin > 0
    myPath = '';
    flagContinue = true;  
    myFormat = 'mat';
else
    warning(['Insufficient arguments provided to ',mfilename,', require: fileName, and optionally path, format.'])
    flagContinue = false;
end
myMapelOptions = mapelOptions;

if flagContinue
    if ~(sum(ismember({'mat'},lower(myFormat))) == 1)
        warning(['Unsupported file format specified in ',mfilename,'. Support: "mat".'])
        flagContinue = false;
    else
        myFormat = lower(myFormat);
        % This doesn't need to be specified for load     
        % saveVersion = '-v7.3';
    end
end

if flagContinue
    if strcmp('mat', myFormat)
        fullFileName = [myPath,myFileName,'.',myFormat];
        myMapelOptions = load(fullFileName, '-mat');
        myMapelOptions = myMapelOptions.myMapelOptions;
		myMapelOptions = checkUpdateObjectVersion(myMapelOptions);
    end
else
    warning('Unable to load in ',mfilename,'. Returning a blank mapelOptions.')
end
end
```
