## Supplementary File 3 for "A surrogate workflow for virtual population development: designing an amenable proxy for a hypothesis-test inspired nonlinear goodness-of-fit": loadWorksheet.m

```
function myWorksheet = loadWorksheet(myFileName, myPath, myFormat, opts)
% Load a worksheet to file.
%
% ARGUMENTS 
%   fileName:    (optional) a filename, suffix will be appended
%   based on format 
%   path:        (optional) save file path 
%   format:      (optional) save file format, currently only support 'mat' 
%   fullPath:    (optional, keyword) full path to the worksheet file, including file
%   extension.
%
% RETURNS
% myWorksheet
%

arguments
    myFileName (1,:) char='';
    myPath (1,:) char='';
    myFormat (1,:) char='mat';
    opts.fullPath (1,:) {mustBeFileOrEmpty}='';
end

% Perform initial checks on the provided arguments
flagContinue = true;
if isempty(myFileName) && isempty(opts.fullPath)
    warning("File not specified. Either the first positional argument " + ...
            "is specified with" + ...
            "the name of the worksheet file (without extension) or " + ...
            "the optional keyword argument " + ...
            "fullPath " + ...
            "must be specified " + ...
            "with the full path to the worksheet file (inclding extension).");
    flagContinue = false;
end

myWorksheet = createWorksheet();

if flagContinue
    if ~(sum(ismember({'mat'},lower(myFormat))) == 1)
        warning(['Unsupported file format specified in ',mfilename,'. Support: "mat".'])
        flagContinue = false;
    else
        myFormat = lower(myFormat);
        % saveVersion doesn't need to be specified for load     
    end
end

if flagContinue
    if strcmp('mat', myFormat)
        if ~isempty(myFileName)
            fullFileName = [myPath,myFileName,'.',myFormat];
        else
            fullFileName = opts.fullPath;
        end
        myWorksheet = load(fullFileName, '-mat');
        % We allow for loading from the simple save or the more 
        % disk space and MATLAB IO conscious save here
        myFields = fields(myWorksheet);
        if ((length(myFields) == 1) && (sum((ismember(myFields,'myWorksheet')))==1))
            myWorksheet = myWorksheet.myWorksheet;
        elseif ((length(myFields) == 11) && (sum((ismember(myFields,{'myWorksheet','myVPCoeffs','myVPIDs','pointBaseVPIndices','baseVPVariantSets', 'myAxisIDs','myAxisElementNames','myAxisElementTypes','myAxisBounds','myAxisScale','myResponseTypes',})))==11))
            myVPIDs = myWorksheet.myVPIDs;
            nVPs = length(myVPIDs);
            pointBaseVPIndices = myWorksheet.pointBaseVPIndices;
            myVPCoeffs = myWorksheet.myVPCoeffs;
            baseVPVariantSets = myWorksheet.baseVPVariantSets;
            myAxisIDs = myWorksheet.myAxisIDs;
            myAxisElementNames = myWorksheet.myAxisElementNames;
            myAxisElementTypes = myWorksheet.myAxisElementTypes;
            myAxisBounds = myWorksheet.myAxisBounds;
            myAxisScale = myWorksheet.myAxisScale;
            myResponseTypes = myWorksheet.myResponseTypes;
            myWorksheet = myWorksheet.myWorksheet;           
            [nAxis,~] = size(myVPCoeffs);
            % Rather than calling createVPs, it will likely be faster to
            % run this step similar to addVariedVPs.
            % So we implement this a little differently here.
            myWorksheet.vpDef=cell(1,nVPs);
            templateVP = struct();
            templateVP.ID = myVPIDs{1};
            templateVP.variants = baseVPVariantSets{1};
            for vpCounter = 1 : nVPs
                templateVP.ID = myVPIDs{vpCounter};
                templateVP.variants = baseVPVariantSets{pointBaseVPIndices(1,vpCounter)};
                myWorksheet.vpDef{vpCounter} = templateVP;
            end
            % Add the axis definitions back
            myWorksheet.axisProps.axisDef = cell(nAxis,1);
            for axisCounter = 1 : nAxis
                myWorksheet.axisProps.axisDef{axisCounter,1} = axisDef({myAxisIDs{axisCounter}, myAxisElementNames{axisCounter}, myAxisBounds{axisCounter}, myAxisElementTypes{axisCounter}, myAxisScale{axisCounter}});
            end   
            myWorksheet.axisProps.axisVP = axisVP(myVPCoeffs);
            % Add the response types back
            for responseTypeCounter = 1 : length(myResponseTypes)
                nElements = length(myResponseTypes{responseTypeCounter}.elements);
                for elementCounter = 1 : nElements;
                    rteStruct = myResponseTypes{responseTypeCounter}.elements{elementCounter};
                    curClass = rteStruct.class;
                    curFields = fields(rteStruct);
                    curFields = setdiff(curFields,'class');
                    curRTE = feval(curClass,{});
                    for propertyCounter = 1 : length(curFields)
                        curField = curFields{propertyCounter};
                        curRTE = set(curRTE, curField, rteStruct.(curField));
                    end
                    myResponseTypes{responseTypeCounter}.elements{elementCounter} = curRTE;
                end
            end     
            myWorksheet.responseTypes = myResponseTypes;
        else
            warning(['Unable to identify loaded mat file as an anticipated worksheet file in ',mfilename,'. Returning an empty worksheet.'])
            myWorksheet = createWorksheet();
        end
    end
else
    warning(['Unable to load in ',mfilename,'. Returning an empty worksheet.'])
end
end

function mustBeFileOrEmpty(str)
    if isempty(str)
        return
    end

    mustBeFile(str)
end
```
