## Supplementary File 3 for "A surrogate workflow for virtual population development: designing an amenable proxy for a hypothesis-test inspired nonlinear goodness-of-fit": loadWorksheetAutoMerge.m

```
function myWorksheet = loadWorksheetAutoMerge(myFileName, myPath, myFormat)
% Load a worksheet to file.
%
% ARGUMENTS
% fileName:    a filename, suffix will be appended based on format
% path:        (optional) save file path
% format:      (optional)save file format, currently only support 'mat'
%
% RETURNS
% myWorksheet
%

% Perform initial checks on the provided arguments
flagContinue = true;
if nargin > 3
    warning(['Too many arguments provided to ',mfilename,', require: fileName, and optionally path, format.'])
    flagContinue = false;
elseif nargin > 2
    flagContinue = true;
elseif nargin > 1
    myFormat = 'mat';
    flagContinue = true;
elseif nargin > 0
    myPath = '';
    flagContinue = true;  
    myFormat = 'mat';
else
    warning(['Insufficient arguments provided to ',mfilename,', require: fileName, and optionally path, format.'])
    flagContinue = false;
end
%myWorksheet = createWorksheet();

if flagContinue
    if ~(sum(ismember({'mat'},lower(myFormat))) == 1)
        warning(['Unsupported file format specified in ',mfilename,'. Support: "mat".'])
        flagContinue = false;
    else
        myFormat = lower(myFormat);
        % saveVersion doesn't need to be specified for load     
    end
end

if flagContinue
    if strcmp('mat', myFormat)
        checkFiles = 0;
        checkFileName = [myPath,[myFileName,'___',num2str(checkFiles+1)],'.',myFormat];
        while exist(checkFileName,'file') == 2
            checkFiles = checkFiles + 1;
            checkFileName = [myPath,[myFileName,'___',num2str(checkFiles+1)],'.',myFormat];
        end
        if checkFiles>0
            myWorksheet = loadWorksheet([myFileName,'___',num2str(1)], myPath, myFormat);
            for splitCounter=2:(checkFiles)
                curWorksheet = loadWorksheet([myFileName,'___',num2str(splitCounter)], myPath, myFormat);
                myWorksheet=mergeWorksheets(myWorksheet,curWorksheet);
            end
        else
            myWorksheet = loadWorksheet(myFileName, myPath, myFormat);
        end
    end
else
    warning(['Unable to load in ',mfilename,'. Returning an empty worksheet.'])
end
end
```
