## Supplementary File 3 for "A surrogate workflow for virtual population development: designing an amenable proxy for a hypothesis-test inspired nonlinear goodness-of-fit": readExperimentData.m

```
function myWorksheet = readExperimentData(myWorksheet, fileName)
% Read experiment result data from file
% ARGUMENTS
% myWorksheet: a worksheet, required
% fileName:    tab-delimited text file to read data from.
%
% RETURNS
% worksheet: an updated worksheet with the VPs.
%
% Note that we don't map subject or group information, or to 
% simulation variables here.  For now, we'll keep this free until
% it is actually needed at optimization.  If it seems this is consistent,
% we can implement specification here in the future.
% We also won't specify the time variable here, since we
% may include alternate initialization times in the simulation
%
continueFlag = true;
%
switch nargin
    case 2
        tokens = regexp(fileName,'(.*)\.(\w*)','tokens');
        if length(tokens) > 0
            noPathName = strsplit(tokens{1}{1},filesep);
            filePath = strjoin(noPathName(1:length(noPathName)-1),filesep);
            noPathName = noPathName{length(noPathName)};
            fileTypeExt = tokens{1}{2};
        else
            warning(['Cannot load file in ',mfilename,', file should have an extension.'])
            continueFlag = false;
        end
    case 1
        % Open a dialog to select file
        % Only support SimBiology models for now
        [fileNameFull,filePath] = uigetfile({'*.sbproj'});
        if (fileNameFull)
            tokens = regexp(fileName,'(.*)\.(\w*)','tokens');
            if length(tokens) > 0
                noPathName = tokens{1}{1};
                fileTypeExt = tokens{1}{2};
            else
                warning(['Cannot load file in ',mfilename,', file should have an extension.'])
                continueFlag = false; 
            end
        else
            warning(['Cannot load file in ',mfilename,'.'])
            continueFlag = false;
        end
    case 0
        warning(['Require a worksheet and filename in ',mfilename,'.'])     
        continueFlag = false;        
end

if continueFlag
    switch fileTypeExt
        case 'txt'
            fileType = 'TXT';
        otherwise
            warning(['Cannot process this file type in ',mfilename,'.'])
            continueFlag = false;
    end      
end


if continueFlag
    if length(filePath) < 1
        filePath=which([noPathName,'.',fileTypeExt]);
        filePath=filePath(1:end-(length([noPathName,'.',fileTypeExt])+1));
    end
end

if continueFlag
    if exist([filePath,filesep,noPathName,'.',fileTypeExt], 'file') ~= 2
        warning(['Cannot locate file in ',mfilename,'.'])
        continueFlag = false;
    end
end

if continueFlag
    existingExpDataIDs = getExpDataIDs(myWorksheet);
    if sum(ismember(existingExpDataIDs,noPathName)) > 0
        warning(['Specified expDataID',noPathName,' already present in myWorksheet while running ',mfilename,'.'])
        continueFlag = false;
    end
end

if continueFlag
    switch fileType
        case 'TXT'
            % Note: there appears to be an issue with long
            % strings in tdfread(), where variant lengths may exceed 
            % MATLAB's namelengthmax.  The way these files are formatted,
            % the variant names should be entries rather than variable names 
            % with the MATLAB tblread command.  This appears to be an issue
            % with tdfread and does not impact the file reading in 2015b.
            % For reading experimental data, readtable is being used.
            % We may need to switch if the variable names are too long,
            % but there haven't been any issues so far.
            warning('off','all');
            %unformatTable = tdfread([fileName,'.txt'],'\t');
            unformatTable = readtable([filePath,filesep,noPathName,'.',fileTypeExt],'Delimiter','\t','Format','auto');
            warning('on','all');
            if ~(strcmp(class(unformatTable),'table'))
                warning(['Cannot read specified file in ',mfilename,'.'])
                continueFlag = false;
            else
                %tableNames = fields(unformatTable);
                %tableVars = properties(unformatTable);
                tableVars = unformatTable.Properties.VariableNames;
                for variableCounter = 1 : length(tableVars)
                    curName = tableVars{variableCounter};
                    curData = unformatTable{:, curName};
                    % readtable doesn't know what to do with NaN's, and the whole
                    % column gets converted to text.  Fix that here.
                    if strcmp(class(curData),'cell')
                        for rowCounter = 1 : length(curData)
                            curEntry = curData{rowCounter};
                            if sum(ismember({'.','NaN','NAN'},curEntry))>0
                                curEntry = nan;
                            else
                                [num, status] = str2num(curEntry);
                                if status == 1
                                    curEntry = num;
                                end
                            end
                            newData{rowCounter,1} = curEntry;
                        end
                        if sum(~(cellfun(@isnumeric,newData))) == 0
                            unformatTable.(curName) = cell2mat(newData);
                        else
                            unformatTable.(curName) = newData;
                        end
                    end
                end
                % We will overwrite datasets that have the same name
                expDataIDs = getExpDataIDs(myWorksheet);
                if sum(ismember(expDataIDs, noPathName)) > 1
                    warning(['Degenerate experimental data names in ',mfilename,'.'])
                    continueFlag = false;
                elseif sum(ismember(expDataIDs, noPathName)) < 1
                    [nDataSets, dummy] = size(myWorksheet.expData);
                    theData = struct();
                    theData.ID = noPathName;
                    theData.data = unformatTable;
                    myWorksheet.expData{nDataSets+1,1} = theData;
                else
                    theIndex = find(ismember(expDataIDs, noPathName));
                    theData = struct();
                    theData.ID = noPathName;
                    theData.data = unformatTable;
                    myWorksheet.expData{theIndex,1} = theData;
                end
            end
    end
else
    warning(['Unable to read file in ',mfilename,'.  Returning input worksheet.'])
end

end
```
