## Supplementary File 3 for "A surrogate workflow for virtual population development: designing an amenable proxy for a hypothesis-test inspired nonlinear goodness-of-fit": readInterventionTable.m

```
function myWorksheet = readInterventionTable(myWorksheet, fileName)
% Read an intervention definition from a file.  Note that if there are
% issues with any of the interventions in the file, the default behavior is
% to ignore the addition of all of the interventions in the file.
% If the worksheet already has interventions, the ones from file are
% appended.
%
% ARGUMENTS
% worksheet: a worksheet, required
% fileName: the name of tab-delimited text file to read intervention 
%       definitions from, note that the full filename including extension
%       should be specified.
%       1st column serves as headers and may only contain the following
%       keywords:
%       ID: name to be assigned to intervention
%       VARIANT:  1st column, indicates rest of row contains
%                 variant names to use for intervention.  These will
%                 take priority over VP-level variants when simulating the
%                 intervention.  Unlike VP definition files, these do not
%                 need to be "square", and may be left blank
%       DOSE:     additional parameters to modify to apply to the VPs
%
% RETURNS
% worksheet: an updated worksheet with the interventions.
%


continueFlag = true;
switch nargin
    case 2
        tokens = regexp(fileName,'(.*)\.(\w*)','tokens');
        if length(tokens) > 0
            noPathName = strsplit(tokens{1}{1},filesep);
            filePath = strjoin(noPathName(1:length(noPathName)-1),filesep);
            noPathName = noPathName{length(noPathName)};
            fileTypeExt = tokens{1}{2};
        else
            warning(['Cannot load file in ',mfilename,', file should have an extension.'])
            continueFlag = false;
        end
    case 1
        % Open a dialog to select file
        [fileNameFull,filePath] = uigetfile({'*.txt'});
        if (fileNameFull)
            tokens = regexp(fileName,'(.*)\.(\w*)','tokens');
            if length(tokens) > 0
                noPathName = tokens{1}{1};
                fileTypeExt = tokens{1}{2};
            else
                warning(['Cannot load file in ',mfilename,', file should have an extension.'])
                continueFlag = false;
            end                
        else
            warning(['Cannot load file in ',mfilename,'.'])
            continueFlag = false;
        end
    case 0
        warning(['Require a worksheet and filename in ',mfilename,'.'])     
        continueFlag = false;
end

if continueFlag    
    switch fileTypeExt
        case 'txt'
            fileType = 'TXT';
        otherwise
            warning(['Cannot process this file type in ',mfilename,'.'])
            continueFlag = false;
    end  
end   

if continueFlag
    if length(filePath) < 1
        filePath=which([noPathName,'.',fileTypeExt]);
        filePath=filePath(1:end-(length([noPathName,'.',fileTypeExt])+1));
    end
end

if continueFlag
    if exist([filePath,filesep,noPathName,'.',fileTypeExt], 'file') ~= 2
        warning(['Cannot locate file in ',mfilename,'.'])
        continueFlag = false;
    end
end

if continueFlag
    switch fileType
        case 'TXT'
            % Note: there appears to be an issue with long
            % strings here, where variant lengths may exceed 
            % MATLAB's namelengthmax.  The way these files are formatted,
            % the variant names should be entries rather than variable names 
            % with the MATLAB tblread command.  This appears to be an issue
            % with tdfread and does not impact the file reading in 2015b.
            % wanted to note this erroneous warning and 
            % in case we write our own table reading script that
            % doesn't have a similar issue.
            warning('off','all');
            unformatInterventionTable = tdfread([filePath,filesep,noPathName,'.',fileTypeExt],'\t');
            warning('on','all');
            if ~(strcmp(class(unformatInterventionTable),'struct'))
                warning(['Cannot read specified file in ', mfilename, '.'])
                continueFlag = false;
            else
            
                interventionNames = fields(unformatInterventionTable);
                id_pos = find(ismember(interventionNames,'ID'));
                if length(id_pos) ~= 1
                    warning(['Exactly one row headed with ID is required in ', mfilename,'.'])
                    continueFlag = false;
                else
                    interventionNames{1} = {};
                    interventionNames = interventionNames(~cellfun('isempty', interventionNames));
                    rowLabels = unformatInterventionTable.('ID');
                    rowLabels = cellstr(rowLabels);
                    % Check all row labels are acceptable
                    passcheck = [find(ismember(rowLabels,'VARIANT'));find(ismember(rowLabels,'DOSE'))];
                    if length(passcheck) ~= length(rowLabels)
                        warning(['Only acceptable row headers are "DOSE" and "VARIANT" in ',mfilename,'.'])
                        continueFlag = false;
                    end
                end

            end
    end
end

if continueFlag
    % Now check to make sure all of the variants and doses are
    % allowed and in the
    % SimBiology file
    passInterventions = cell(0,1);
    failedInterventions = cell(0,1);
    variantDelimiter = myWorksheet.variantProps.delimiter;
    variantTypes = myWorksheet.variantProps.variantTypes;
    variantNames = myWorksheet.variantProps.typeValueSets;
    doseNames = getDoseNames(myWorksheet);
    previousInterventionIDs = getInterventionIDs(myWorksheet); 
    [nInterventionResults, nVPResults] = size(myWorksheet.results);

    for interventionCounter = 1 : length(interventionNames)
        currentPass = true;
        interventionID = interventionNames{interventionCounter};
        currentVariantsDoses = unformatInterventionTable.(interventionID);
        % Fully blank interventions, like what might happen with a control,
        % get read in as NaN's
        if isnumeric(currentVariantsDoses)
            currentVariantsDoses = num2cell(currentVariantsDoses);
            currentVariantsDoses(cellfun(@isnan,currentVariantsDoses))={''};
        end
        currentVariantsDoses = cellstr(currentVariantsDoses);
        % We allow blank entries, all of the interventions
        % in the file don't need the same # variants/doses
        blankLocations = find(ismember(currentVariantsDoses,''));
        for blankCounter = 1 : length(blankLocations)
            blankIndex = blankLocations(blankCounter);
            currentVariantsDoses{blankIndex} = {};
        end
        %currentVariantsDoses = currentVariantsDoses(~cellfun('isempty',currentVariantsDoses));
        arrangedCurrentVariants = cell(length(variantTypes),1);
        arrangedCurrentDoses = cell(length(doseNames),1);

        for row_counter = 1 : length(rowLabels)
            row_type = rowLabels{row_counter};
            if ~(isempty(currentVariantsDoses{row_counter}))
                if strcmp(row_type, 'VARIANT')
                    variantName = currentVariantsDoses{row_counter};
                    split_result = strsplit(variantName, variantDelimiter);
                    variantType = variantTypes{find(ismember(variantTypes, split_result{1}))};
                    if length(find(ismember(variantTypes,variantType))) == 1
                        curIndices = find(ismember(variantTypes,variantType));
                        arrangedCurrentVariants{curIndices} = variantName;
                        if length(find(ismember(variantName,variantNames))) < 1
                            disp(['Unable to find VARIANTNAME ',variantName,' from intervention file in model.'])
                        end
                    else
                        currentPass = false;
                    end
                elseif strcmp(row_type, 'DOSE')
                    doseName = currentVariantsDoses{row_counter};
                    if find(ismember(doseNames, doseName))
                        arrangedCurrentDoses{find(ismember(doseNames,doseName))} = doseName;
                    else
                        currentPass = false;
						warning(['Unable to find DOSE ',doseName,' from intervention file in model.'])
                    end
                end
            end
        end
        
        if sum(ismember(previousInterventionIDs,interventionID)) > 0
            currentPass = false;
        end        

        arrangedCurrentVariants = arrangedCurrentVariants(~cellfun('isempty',arrangedCurrentVariants));
        arrangedCurrentDoses = arrangedCurrentDoses(~cellfun('isempty',arrangedCurrentDoses));
        nVariants = length(arrangedCurrentVariants);
        nDoses = length(arrangedCurrentDoses);
        variantLabel = cell(nVariants,1);
        doseLabel = cell(nDoses,1);
        if (((nDoses + nVariants) > 0) && (currentPass == true))
            if length(variantLabel) > 0
                variantLabel(:) = {'VARIANT'};
            end
            if length(doseLabel) > 0
                doseLabel(:) = {'DOSE'};
            end
            passInterventions{length(passInterventions) +1,1} = [variantLabel, arrangedCurrentVariants; doseLabel, arrangedCurrentDoses];
        elseif (currentPass == true)
            %failedInterventions{length(failedInterventions)+1} = interventionID;
            warning(['No model variants or doses for intervention ',interventionID,' in ',mfilename,'.'])
            passInterventions{length(passInterventions) +1,1} = [variantLabel, arrangedCurrentVariants; doseLabel, arrangedCurrentDoses];
        else
            failedInterventions{length(failedInterventions)+1} = interventionID;
        end
    end

    if length(failedInterventions) == 0
        [nPassInterventions, dummy] = size(passInterventions);
        for newInterventionCounter = 1 : nPassInterventions
            newIntervention = passInterventions{newInterventionCounter};
            interventionID = interventionNames{newInterventionCounter};
            arrangedIntervention = passInterventions{newInterventionCounter};
            myWorksheet = createIntervention(myWorksheet, interventionID, arrangedIntervention);
        end    
        if nInterventionResults > 0
            % We pad with empty results if there are results for prior
            % interventions in the worksheet
            myWorksheet.results = [myWorksheet.results;cell(nPassInterventions,nVPResults)];
        end
    else
        failedInterventions = [failedInterventions',[repmat({','},numel(failedInterventions)-1,1);{[]}]]';
        failedInterventions = [failedInterventions{:}];
        warning(['Failed importing interventions ',failedInterventions,' in ',mfilename,'.'])   
    end
else
    warning(['Unable to read intervention file in ',mfilename,'. Exiting and returning input worksheet.'])
end
```
