## Supplementary File 3 for "A surrogate workflow for virtual population development: designing an amenable proxy for a hypothesis-test inspired nonlinear goodness-of-fit": readQSPModel.m

```
function worksheet = readQSPModel(fileName, modelname)
% Read a model from file, and set up a worksheet structure
% 
% ARGUMENTS
% fileName:  name of the file to read.  A dialogue box will prompt if not
%            specified.  Supported file extensions (must be included):
%             .sbproj: SimBiology model
% modelname: a simbiology project may have multiple associated models.
%            specify the right one here, or leave blank to just
%            use the first model
%
% RETURNS
% worksheet
%
% Initialize the worksheet structure
worksheet = createWorksheet();
continueFlag = true;
switch nargin
    case 2
        tokens = regexp(fileName,'(.*)\.(\w*)','tokens');
        if length(tokens) > 0
            noPathName = strsplit(tokens{1}{1},filesep);
            filePath = strjoin(noPathName(1:length(noPathName)-1),filesep);
            noPathName = noPathName{length(noPathName)};
            fileTypeExt = tokens{1}{2};
        else
            warning(['Cannot load file in ',mfilename,', file should have an extension.'])
            continueFlag = false;
        end
    case 1
        tokens = regexp(fileName,'(.*)\.(\w*)','tokens');
        if length(tokens) > 0
            noPathName = strsplit(tokens{1}{1},filesep);
            filePath = strjoin(noPathName(1:length(noPathName)-1),filesep);
            noPathName = noPathName{length(noPathName)};
            fileTypeExt = tokens{1}{2};
            modelname = '';
        else
            warning(['Cannot load file in ',mfilename,', file should have an extension.'])
            continueFlag = false;
        end        
    case 0
        % Open a dialog to select file
        % Only support SimBiology models for now
        [fileNameFull,filePath] = uigetfile({'*.sbproj'});
        if (fileNameFull)
            tokens = regexp(fileNameFull,'(.*)\.(\w*)','tokens');
            if length(tokens) > 0
                noPathName = tokens{1}{1};
                fileTypeExt = tokens{1}{2};
            else
                warning(['Cannot load file in ',mfilename,', file should have an extension.'])
                continueFlag = false; 
            end                

        else
            warning(['Cannot load file in ',mfilename,'.'])
            continueFlag = false;
        end
        modelname = '';
end

if continueFlag
    switch fileTypeExt
        case 'sbproj'
            fileType = 'SB';
        otherwise
            warning(['Cannot process this file type in ',mfilename,'.'])
            continueFlag = false;
    end        
end

if continueFlag
    if length(filePath) < 1
        filePath=which([noPathName,'.',fileTypeExt]);
        filePath=filePath(1:end-(length([noPathName,'.',fileTypeExt])+1));
    end
end

if continueFlag
    if exist([filePath,filesep,noPathName,'.',fileTypeExt], 'file') ~= 2
        warning(['Cannot locate file in ',mfilename,'.'])
        continueFlag = false;
    end
end

if continueFlag
    switch fileType
        case 'SB'
             temp = sbioloadproject([filePath,filesep,noPathName,'.',fileTypeExt]);
             nModels = length(fields(temp));
             modelNames = cell(1,nModels);
             dummyNames = fields(temp);
             for modelCounter = 1 : nModels
                 modelNames{modelCounter} = temp.(dummyNames{modelCounter}).Name;
             end
             modelPos = find(ismember(modelNames,modelname));
             if length(modelPos) < 1
                 % By default we take the first model
                 worksheet.model = temp.(dummyNames{1});
                 if length(modelname) > 0
                     warning(['Cannot find model with specified name, using the first: ',modelNames{1},'.'])
                 elseif length(modelNames) > 1
                     disp(['Model to use not specified, using the first: ',modelNames{1},'.'])
                 end
             else
                 worksheet.model = temp.(dummyNames{modelPos(1)});
             end
    end

    worksheet.project.fileName = [noPathName,'.',fileTypeExt];
    worksheet.project.filePath = filePath;

    % We assume a convention where variant names are constructed
    % according to the pattern: xxxxxxxxxxxxxxx___yyyyyyyyyy
    worksheet.variantProps.variantTypes = getVariantTypes(worksheet);
    worksheet.variantProps.typeValueSets = getVariantNames(worksheet);
else
    warning(['Unable to read file in ',mfilename,'.  Returning an empty worksheet.'])
end
```
