## Supplementary File 3 for "A surrogate workflow for virtual population development: designing an amenable proxy for a hypothesis-test inspired nonlinear goodness-of-fit": readVPTable.m

```
function myWorksheet = readVPTable(myWorksheet, fileName)
% Read VP definitions from a file
% ARGUMENTS
% worksheet: a worksheet, required
% fileName: tab-delimited text file to read vp definitions from.
%       1st column must follow this format:
%       ID: name to be assigned to VP
%       VARIANTS: 1st column, indicates rest of row contains
%                 variant names to use for VP.  Must include
%                 exactly one entry to for each variantType
%                 in worksheet.variantProps.variantTypes
%       parameter_name: additional parameters to modify to apply to the VPs
%
% RETURNS
% worksheet: an updated worksheet with the VPs.
%

continueFlag = true;
switch nargin     
    case 2
        tokens = regexp(fileName,'(.*)\.(\w*)','tokens');
        if length(tokens) > 0
            noPathName = strsplit(tokens{1}{1},filesep);
            filePath = strjoin(noPathName(1:length(noPathName)-1),filesep);
            noPathName = noPathName{length(noPathName)};
            fileTypeExt = tokens{1}{2};
        else
            warning(['Cannot load file in ',mfilename,', file should have an extension.'])
            continueFlag = false;
        end
    case 1
        % Open a dialog to select file
        [fileNameFull,filePath] = uigetfile({'*.txt'});
        if (fileNameFull)
            tokens = regexp(fileName,'(.*)\.(\w*)','tokens');
            if length(tokens) > 0
                noPathName = tokens{1}{1};
                fileTypeExt = tokens{1}{2};
            else
                warning(['Cannot load file in ',mfilename,', file should have an extension.'])
                continueFlag = false; 
            end                
        else
            warning(['Cannot load file in ',mfilename,'.'])
            continueFlag = false;
        end
    case 0
        warning(['Require a worksheet and filename in ',mfilename,'.'])     
        continueFlag = false;
end


if continueFlag    
    switch fileTypeExt
        case 'txt'
            fileType = 'TXT';
        otherwise
            warning(['Cannot process this file type in ',mfilename,'.'])
            continueFlag = false;
    end  
end

if continueFlag
    if length(filePath) < 1
        filePath=which([noPathName,'.',fileTypeExt]);
        filePath=filePath(1:end-(length([noPathName,'.',fileTypeExt])+1));
    end
end

if continueFlag
    if exist([filePath,filesep,noPathName,'.',fileTypeExt], 'file') ~= 2
        warning(['Cannot locate file in ',mfilename,'.'])
        continueFlag = false;
    end
end

if continueFlag
    switch fileType
        case 'TXT'
            % Note: there appears to be an issue with long
            % strings here, where variant lengths may exceed 
            % MATLAB's namelengthmax.  The way these files are formatted,
            % the variant names should be entries rather than variable names 
            % with the MATLAB tblread command.  This appears to be an issue
            % with tdfread and does not impact the file reading in 2015b.
            % wanted to note this erroneous warning and 
            % in case we write our own table reading script that
            % doesn't have a similar issue.
            warning('off','all');
            unformatVPTable = tdfread([filePath,filesep,noPathName,'.',fileTypeExt],'\t');
            warning('on','all');
            if ~(strcmp(class(unformatVPTable),'struct'))
                error('Cannot read specified file.')
            end
            vpNames = fields(unformatVPTable);
            id_pos = find(ismember(vpNames,'ID'));
            if length(id_pos) ~= 1
                warning(['Exactly one row headed with ID is required in ',mfilename,'.'])
                continueFlag = false;
            else
                vpNames{1} = {};
                vpNames = vpNames(~cellfun('isempty',vpNames));
                rowLabels = unformatVPTable.('ID');
                rowLabels = cellstr(rowLabels);
            end
        otherwise
            warning(['Cannot process this file type in ',mfilename,'.'])
            continueFlag = false;
    end
end

if continueFlag
    % Now apply checks to make sure VPs and variants are read
    % correctly
    passVPs = cell(1,0);
    failedVPs = cell(1,0);
    failedRowVPs = cell(1,0);
    failedVariantNames = cell(1,0);
    failedVariantTypes = cell(1,0);
    failedVPVariantPresent = cell(1,0);
    failedVPPresent = cell(1,0);    
    variantDelimiter = myWorksheet.variantProps.delimiter;
    variantTypes = myWorksheet.variantProps.variantTypes;
    variantNames = myWorksheet.variantProps.typeValueSets;
    missingVariantTypes = cell(1,0);

    previousVPIDs = getVPIDs(myWorksheet); 
    for vpCounter = 1 : length(vpNames)
        currentPass = true;
        failrowtypecheck = false;
        failvppresentcheck=false;
        failvarianttypecheck = false;      
        failvariantnamecheck = false;
        failvariantpresentcheck = false;
        % We might want to add a check on the VP ID
        % here for uniqueness
        vpID = vpNames{vpCounter};
        currentVariants = unformatVPTable.(vpID);
        if isnumeric(currentVariants)
            currentVariants = num2cell(currentVariants);
            currentVariants(cellfun(@isnan,currentVariants))={''};
        end        
        currentVariants = cellstr(currentVariants);
        % We allow blank entries, all of the interventions
        % in the file don't need the same # variants/doses
        blankLocations = find(ismember(currentVariants,''));
        for blankCounter = 1 : length(blankLocations)
            blankIndex = blankLocations(blankCounter);
            currentVariants{blankIndex} = {};
        end        
        arrangedCurrentVariants = cell(length(variantTypes),1);
        arrangedCurrentVariantTypes = cell(length(variantTypes),1);

        for row_counter = 1 : length(rowLabels)
            row_type = rowLabels{row_counter};
            if ~(isempty(currentVariants{row_counter}))
                if row_type == 'VARIANT'
                    variantName = currentVariants{row_counter};
                    split_result = strsplit(variantName, variantDelimiter);
                    variantType = variantTypes{find(ismember(variantTypes, split_result{1}))};
                    if length(find(ismember(variantTypes,variantType))) == 1
                        arrangedCurrentVariants{find(ismember(variantTypes,variantType))} = variantName;
                        arrangedCurrentVariantTypes{find(ismember(variantTypes,variantType))} = variantType;
                    else
                        % Fail if trying to add a variant type we don't
                        % have in the model
                        currentPass = false;
                        failvarianttypecheck = true;
                        failedVariantTypes = [failedVariantTypes, variantType]
                    end
                    % Also check if not just the variant type is in the model 
                    % but also whether the variant itself is available in the model
                    if sum(ismember(variantNames, variantName)) == 0
                        currentPass = false;
                        failvariantnamecheck = true;
                        failedVariantNames = [failedVariantNames, variantName]                
                    end
                else
                    % only 'VARIANT' supported
                    % Individual parameters not yet supported
                    currentPass = false;
                    failrowtypecheck = true;
                end
            end
        end
        % Final check: Make sure we got all of the Variant Types needed for
        % VP specification
        if sum(cellfun('isempty',arrangedCurrentVariants)) > 0
            currentPass = false;
            failvariantpresentcheck = true;
            arrangedCurrentVariantTypes = arrangedCurrentVariantTypes(~cellfun('isempty',arrangedCurrentVariantTypes));
            curMissingVariantTypes = setdiff(variantTypes, arrangedCurrentVariantTypes);
            for vCounter = 1 : length(curMissingVariantTypes)
                curMissingVariant = curMissingVariantTypes{vCounter};
                if sum(ismember(missingVariantTypes,curMissingVariant)) < 1
                    missingVariantTypes{1, length(missingVariantTypes)+1} = curMissingVariant;
                end
            end
        end
        % We also check for uniqueness
        if sum(ismember(previousVPIDs,vpID)) > 0
            currentPass = false;
            failvppresentcheck = true;
        end
        if currentPass
            passVPs{1, length(passVPs)+1} = {vpID, arrangedCurrentVariants};
        else
            failedVPs{1, length(failedVPs)+1} = vpID;
            if failrowtypecheck
                failedRowVPs{1, length(failedRowVPs)+1} = vpID;
            end
            if failvariantpresentcheck
                failedVPVariantPresent{1,length(failedVPVariantPresent)+1} = vpID;
            end
            if failvppresentcheck
                failedVPPresent{1,length(failedVPPresent)+1} = vpID;
            end            
        end
    end

    if length(failedVPs) > 0
        warning(['Failed VPs, ignoring: ',strjoin(failedVPs)])
        if length(failedRowVPs) > 0
            warning(['Note, failed thse VPs due to unrecognized row specification: ',strjoin(failedRowVPs),' in ',mfilename,'.'])
        end
        if length(failedVPVariantPresent) > 0
            warning(['Note, failed the following VPs due to missing variant types: ',strjoin(failedVPVariantPresent),' in ',mfilename,'.'])
            warning(['Variant types with issues: ',strjoin(missingVariantTypes),' in ',mfilename,'.'])
        end
        if length(failedVariantNames) > 0
            warning(['Note, the following variants were specified in the file but could not be identified in the model: ',strjoin(failedVariantNames),' in ',mfilename,'.'])
        end    
        if length(failedVPPresent) > 0
            warning(['Note, the following VP IDs were specified in the file but were already present in the worksheet: ',strjoin(failedVPPresent),' in ',mfilename,'.'])
        end
    end
    
    if length(passVPs) > 0

        newVPIDs = cell(1,length(passVPs));
        newVPVariants = cell(1,length(passVPs));

        for newVPCounter = 1 : length(passVPs)
            newVPIDs{newVPCounter} = passVPs{newVPCounter}{1};
            newVPVariants{newVPCounter} = passVPs{newVPCounter}{2};
        end
        myWorksheet = createVPs(myWorksheet, newVPIDs, newVPVariants);
    end
        
    
else
    warning(['Unable to read VP file in ',mfilename,'. Exiting and returning input worksheet.'])
end  

end
```
