## Supplementary File 3 for "A surrogate workflow for virtual population development: designing an amenable proxy for a hypothesis-test inspired nonlinear goodness-of-fit": saveVPop.m

```
function saveVPop(myVPop,myFileName, myFormat, myPath)
% Save a VPop object to file.
%
% ARGUMENTS
% myVPop:   a VPop
% fileName: a filename, suffix will be appended based on format
% format:   (optional)save file format, currently only support 'mat'
% path:     (optional) save file path
%
% RETURNS
% nothing
%

% Perform initial checks on the provided arguments
flagContinue = true;
if nargin > 4
    warning(['Too many arguments provided to ',mfilename,', require: myVPop, fileName, and optionally format, path.'])
    flagContinue = false;
elseif nargin > 3
    flagContinue = true;
elseif nargin > 2
    myPath = '';
    flagContinue = true;
elseif nargin > 1
    myPath = '';
    flagContinue = true;  
    myFormat = 'mat';
else
    warning(['Insufficient arguments provided to ',mfilename,', require: myVPop, fileName, and optionally format, path.'])
    flagContinue = false;
end

% Create the new worksheet and verify the IDs
if flagContinue
    if ~(sum(ismember({'mat'},lower(myFormat))) == 1)
        warning(['Unsupported file format specified in ',mfilename,'. Support: "mat".'])
        flagContinue = false;
    else
        myFormat = lower(myFormat);
        % Force v7.3, which is needed to save files > 2GB. 
        % However, the HDF5 format used by v7.3 may be slow.
        % I'm still exploring whether any optimization is possible or
        % needed to improve read/write times.     
        saveVersion = '-v7.3';
    end
end

if flagContinue
    if strcmp('mat', myFormat)
        fullFileName = [myPath,myFileName,'.',myFormat];
        % This is fast to recalculate and can take up space on disk
        myVPop.coeffsDist='';
        save(fullFileName, 'myVPop', saveVersion);
    end
else
    warning('Unable to save in ',mfilename,'. Exiting without save...')
end
end
```
