## Supplementary File 3 for "A surrogate workflow for virtual population development: designing an amenable proxy for a hypothesis-test inspired nonlinear goodness-of-fit": saveWorksheet.m

```
function saveWorksheet(myWorksheet, myFileName, oldSaveFlag, myPath, myFormat)
% Save a worksheet to file. Note the default is to save to MATLAB v7 if
% this is possible.  For worksheets that take up more disk space,
% expecially with results, you may need to save to v7.3
%
% ARGUMENTS
% myWorksheet:     a worksheet
% myFileName:      a filename, suffix will be appended based on format
% oldSaveFlag:     (optional) whether to use an older save format, which
%                  is simpler to read/write but potnetial takes more space
%                  on disk and is slower for IO.  Default is false.
%                  If true, the variation on save worksheet that
%                  takes advantage of some likely duplication in worksheet 
%                  structure to reduce the memory footprint and time to
%                  write to file, so it will be useful for larger
%                  worksheets.
% myPath:          (optional) save file path, if not the current MATLAB path.
%                  leave as '' to keep current
% myFormat:        (optional)save file format. Currently support:
%                   'v7':   *.mat file with v7 formatting.  Generally gives
%                           good results.  Default.
%                   'v7.3'  *.mat file with v7.3 formatting.  Generally
%                           not as compact and quick as saving worksheets
%                           v7 but may be needed in some cases.
%
% RETURNS
% nothing
%

% Perform initial checks on the provided arguments
mflagContinue = true;
if nargin > 5
    warning(['Too many arguments provided to ',mfilename,', require: myWorksheet, fileName, and optionally format, path.'])
    flagContinue = false;
elseif nargin > 4
    flagContinue = true;
elseif nargin > 3
    myFormat = 'v7';    
    flagContinue = true;
elseif nargin > 2
    myFormat = 'v7';  
    myPath = '';
    flagContinue = true;
elseif nargin > 1
    myFormat = 'v7';  
    myPath = '';    
    oldSaveFlag = false;    
    flagContinue = true;  
else
    warning(['Insufficient arguments provided to ',mfilename,', require: myWorksheet, fileName, and optionally format, path.'])
    flagContinue = false;
end

if flagContinue
    allowedFormats = {'v7','v7.3'};
    matFormats = allowedFormats;
    if ~(sum(ismember(allowedFormats,lower(myFormat))) == 1)
        warning(['Unsupported file format specified in ',mfilename,'. Support: "v7", "v7.3".'])
        flagContinue = false;
    else
        myFormat = lower(myFormat);
        if (sum(ismember(matFormats,lower(myFormat))) == 1)
            % Benchmarking the worksheet saves found better performance for
            % -v7 than -v7.3.
            % Also a useful reference:
            % http://undocumentedmatlab.com/blog/improving-save-performance
            myExtension = 'mat';
            
        end
    end
    % We won't check the path in advance
    if ~islogical(oldSaveFlag)
        warning(['Invalid oldSaveFlag specified for ',mfilename,', a boolean (true/false) should be specified.'])
    end    
    
end

if flagContinue
    fullFileName = [myPath,myFileName,'.',myExtension];
    if ~oldSaveFlag
        myVPCoeffs = getVPCoeffs(myWorksheet);
        [nAxis, nVPs] = size(myVPCoeffs);
        [pointBaseVPIndices, baseVPVariantSets] = getUniqueBaseVPVariantSets(myWorksheet, ones(1,nVPs));
        myVPIDs = getVPIDs(myWorksheet);
        myAxisID = cell(nAxis,1);
        myAxisElementNames = cell(nAxis,1);
        myAxisElementTypes = cell(nAxis,1);
        myAxisBounds = cell(nAxis,1);
        myAxisScale = cell(nAxis,1);
        for axisDefCounter = 1 : nAxis
            myAxisIDs{axisDefCounter} = myWorksheet.axisProps.axisDef{axisDefCounter}.id;
            myAxisElementNames{axisDefCounter} = myWorksheet.axisProps.axisDef{axisDefCounter}.elementNames;
            myAxisElementTypes{axisDefCounter} = myWorksheet.axisProps.axisDef{axisDefCounter}.elementTypes;
            myAxisBounds{axisDefCounter} = myWorksheet.axisProps.axisDef{axisDefCounter}.bounds;
            myAxisScale{axisDefCounter} = myWorksheet.axisProps.axisDef{axisDefCounter}.scale;            
        end
        myResponseTypes = myWorksheet.responseTypes;
        for responseTypeCounter = 1 : length(myResponseTypes)
            nElements = length(myResponseTypes{responseTypeCounter}.elements);
            for elementCounter = 1 : nElements;
                curRTE = myResponseTypes{responseTypeCounter}.elements{elementCounter};
                rteStruct = struct();
                rteStruct.class = class(curRTE);
                curProperties = properties(curRTE);
                for propCounter = 1 : length(curProperties)
                    rteStruct.(curProperties{propCounter}) = get(curRTE, curProperties{propCounter});
                end
                myResponseTypes{responseTypeCounter}.elements{elementCounter} = rteStruct;
            end
        end
        
        % We eliminate custom objects from what will be written 
        % to file. This should make it easier to update worksheet as needed
        % in the future with new object definitions. These are 
        % re-instantiated and re-populated with loadWorksheet.
        myWorksheet.axisProps.axisDef = cell(0,1);
        myWorksheet.axisProps.axisVP = cell(0,0);
        myWorksheet.vpDef = cell(1,0);  
        myWorksheet.responseTypes = cell(0,1);
        % At some point we may also want to explore   
        % optimal ways of writing the results structure to
        % file.
    end
    if ~oldSaveFlag
        if strcmp('mat', myExtension)
            save(fullFileName, 'myWorksheet', 'myVPIDs', 'myVPCoeffs', 'pointBaseVPIndices', 'baseVPVariantSets', 'myAxisIDs','myAxisElementNames','myAxisElementTypes','myAxisBounds','myAxisScale','myResponseTypes', ['-',myFormat]);
        end
    else
        if strcmp('mat', myExtension)
            save(fullFileName, 'myWorksheet', ['-',myFormat]);
        end
    end
else
    warning('Unable to save in ',mfilename,'. Exiting without save...')
end
end
```
