## Supplementary File 3 for "A surrogate workflow for virtual population development: designing an amenable proxy for a hypothesis-test inspired nonlinear goodness-of-fit": saveWorksheetAutoSplit.m

```
function saveWorksheetAutoSplit(myWorksheet, myFileName, oldSaveFlag, myPath, myFormat)
% Save a worksheet to file. Note the default is to save to MATLAB v7 if
% this is possible. This save command will try to see if the worksheet
% exceeds 2GB and split the worksheet into multiple subworksheet, and 
% identifier for subworksheet is ___
%
% ARGUMENTS
% myWorksheet:     a worksheet
% myFileName:      a filename, suffix will be appended based on format
% oldSaveFlag:     (optional) whether to use an older save format, which
%                  is simpler to read/write but potnetial takes more space
%                  on disk and is slower for IO.  Default is false.
%                  If true, the variation on save worksheet that
%                  takes advantage of some likely duplication in worksheet 
%                  structure to reduce the memory footprint and time to
%                  write to file, so it will be useful for larger
%                  worksheets.
% myPath:          (optional) save file path, if not the current MATLAB path.
%                  leave as '' to keep current
% myFormat:        (optional)save file format. Currently support:
%                   'v7':   *.mat file with v7 formatting.  Generally gives
%                           good results.  Default.
%                   'v7.3'  *.mat file with v7.3 formatting.  Generally
%                           not as compact and quick as saving worksheets
%                           v7 but may be needed in some cases.
% mySplit: 
%
% RETURNS
% nothing
%

% Perform initial checks on the provided arguments
flagContinue = true;
if nargin > 5
    warning(['Too many arguments provided to ',mfilename,', require: myWorksheet, fileName, and optionally format, path.'])
    flagContinue = false;
elseif nargin > 4
    flagContinue = true;
elseif nargin > 3
    myFormat = 'v7';    
    flagContinue = true;
elseif nargin > 2
    myFormat = 'v7';  
    myPath = '';
    flagContinue = true;
elseif nargin > 1
    myFormat = 'v7';  
    myPath = '';    
    oldSaveFlag = false;    
    flagContinue = true;  
else
    warning(['Insufficient arguments provided to ',mfilename,', require: myWorksheet, fileName, and optionally format, path.'])
    flagContinue = false;
end

if flagContinue
    allowedFormats = {'v7','v7.3'};
    matFormats = allowedFormats;
    if ~(sum(ismember(allowedFormats,lower(myFormat))) == 1)
        warning(['Unsupported file format specified in ',mfilename,'. Support: "v7", "v7.3".'])
        flagContinue = false;
    else
        myFormat = lower(myFormat);
        if (sum(ismember(matFormats,lower(myFormat))) == 1)
            % Benchmarking the worksheet saves found better performance for
            % -v7 than -v7.3.
            % Also a useful reference:
            % http://undocumentedmatlab.com/blog/improving-save-performance
            myExtension = 'mat';
            
        end
    end
    % We won't check the path in advance
    if ~islogical(oldSaveFlag)
        warning(['Invalid oldSaveFlag specified for ',mfilename,', a boolean (true/false) should be specified.'])
    end    
    
end

if flagContinue
    lastwarn('');
    warning('off','MATLAB:save:sizeTooBigForMATFile')
    saveWorksheet(myWorksheet, myFileName, oldSaveFlag, myPath, myFormat);
    [warnMsg, warnId] = lastwarn;
    if ~isempty(warnMsg) && (strcmp(warnId,'MATLAB:save:sizeTooBigForMATFile'))
        warning(['MATLAB:save:sizeTooBigForMATFile detected in ',mfilename,'. Will automatically split the worksheet into subworsheets to save into files.'])
        nSplit=1;
        successSave = false;
        allVPIDs = getVPIDs(myWorksheet);
        nVPs = length(allVPIDs);
        while ~successSave
            nSuccess = 0;
            nSplit = nSplit+1;
            saveVPsize = ceil(nVPs/nSplit);
            for splitCounter=1:nSplit
                if (nSuccess + 1) >= splitCounter
                    vpStartIndex = (splitCounter-1)*saveVPsize + 1;
                    vpEndIndex = min(vpStartIndex + saveVPsize - 1, nVPs);
                    curWorksheet = copyWorksheet(myWorksheet, allVPIDs(vpStartIndex:vpEndIndex));
                    lastwarn('');
                    saveWorksheet(curWorksheet,[myFileName,'___',num2str(splitCounter)],oldSaveFlag, myPath, myFormat);
                    [warnMsg, warnId] = lastwarn;
                    if ~(~isempty(warnMsg) && (strcmp(warnId,'MATLAB:save:sizeTooBigForMATFile')))
                        nSuccess = nSuccess+1;
                    end
                end
            end
            if nSuccess >= nSplit
                successSave = true;
            end
        end
    end
    warning('on','MATLAB:save:sizeTooBigForMATFile')
end
end
```
