## Supplementary File 3 for "A surrogate workflow for virtual population development: designing an amenable proxy for a hypothesis-test inspired nonlinear goodness-of-fit": writeResultStruct.m

```
function myFID = writeResultStruct(myResult, myFileName)
% This function takes a results structure and writes it to a tab-delimited
% text file.  Note that you need to include the file extension in
% myFileName and should be in the desired directory.
%
% ARGUMENTS
%  myResult:                            A struct with the desired data;
%                                       essentially a result struct.
%  myFileName:                          The name of the file to write to.
%
% RETURNS
%  myFID:                               A file identifier for MATLAB.
%

% Perform initial checks on the provided arguments
flagContinue = true;

myFID = 0;

if nargin > 2
    warning(['Too many input arguments to ',mfilename,'. Require: mySimStruct, myFileName.'])
    flagContinue = false;
elseif nargin < 2 
    warning(['Insufficient input arguments to ',mfilename,'. Require: mySimStruct, myFileName.'])
    flagContinue = false;   
end

if flagContinue
    if ~strcmp(class(myResult),'struct')
        warning(['Expecting a result structure with "Name" and "Data" fields in call to ',mfilename,'.  Exiting.'])
        flagContinue = false; 
    end
    if ~strcmp(class(myFileName),'char')
        warning(['Expecting a string for a filename in call to ',mfilename,'.  Exiting.'])
        flagContinue = false; 
    end    
end

if flagContinue
    if ~((sum(ismember(fields(myResult),'Names')) == 1) & (sum(ismember(fields(myResult),'Data')) == 1))
        warning(['Expecting a result structure with "Name" and "Data" fields in call to ',mfilename,'.  Exiting.'])
        flagContinue = false; 
    end
end

if flagContinue
    myFID=fopen(myFileName,'wt');
    nCols = length(myResult.Names);
    for colCounter = 1:nCols-1
        fprintf(myFID,[myResult.Names{colCounter},'\t']);
    end
    fprintf(myFID, [myResult.Names{nCols},'\n']);
    [nRows,nCols] = size(myResult.Data);
    for rowCounter = 1:nRows
        fprintf(myFID,'%e\t',myResult.Data(rowCounter,1:(nCols-1)));
        fprintf(myFID,'%e\n',myResult.Data(rowCounter,nCols));
    end    
    fclose(myFID);
else
    warning(['Unable to complete ', mfilename,', exiting.'])
end
end
```
