## Supplementary File 3 for "A surrogate workflow for virtual population development: designing an amenable proxy for a hypothesis-test inspired nonlinear goodness-of-fit": oncology_trial_data_integration.R

```
# oncology_trial_data_integration.R
#
# Developed by Quantitative Systems Pharmacology group at Bristol-Myers Squibb
# to piece together heterogeneous oncology datasets and guide QSP modeling
#
#
# Version	Date		Author/Description
# --------- ----------- --------------------------------------------------------------------
# 0.48     08/09/2023  LH/ fixed minor bugs in check_and_convert_numeric, read.delim in get_lesion_type to read in non-English characters correctly
#                          and get_patient_time_series_subdata default arguments
# 0.47     10/31/2022  LH/ fixed a bug in add_CBOR_recist function, to read only lesion scan rows
# 0.46     08/07/2022  LH/ added a add_CBOR_recist function, 
#                          edited create_distribution_tables_recist function to add CBRSCORE, CBRSCORENEW columns to the integrated_patient_data table
# 0.45     07/29/2022  LH/ fixed a bug in get_lesion_type, allow correct lymphe node denotation of target lesions, remove the empty denotation, updated default lymphnode_marker in get_lesion_type, get_nontarget_lesions, get_nontarget_progression_time_lnsize_filter
# 0.44     06/04/2022  LH/ fixed a bug in get_demographic_data, not output the whole table when reading data frame as input
# 0.43     06/01/2022  LH/ fixed a issue with missing data when OFFTRTIME is earlier than lesion scan time due to the alignment, by:
#                          1) edited calculate_lesion_summary to read OFFTRTIME and assign it as a time series attribute "ONTRT" onto the trial_list
#                          2) edited create_distribution_tables_recist function to get offtreatment time as maximum of OFFTRTIME and aligned "ONTRT=1" lesion scan date, also assign intermediate time points with correct ONTRT
# 0.42     04/19/2022  RS/ added lymphnode_marker argument to get_nontarget_progression_time_lnsize_filter (was hard-coded before)
# 0.41     11/18/2021  LH/ fixed a bug in get_patient_time_series_subdata, only read time series entry when date info exist
#                          edited in get_demographic_data, so it can read a data frame as filename, if one already prepared the data_table
# 0.40     10/11/2021  LH/ fixed a bug in convert_to_date function, so it is able to automatically convert dates in both format: yyyy-mm-dd and yyyy/mm/dd
# 0.39     08/23/2021  LH/ added a new input in get_patient_time_series_subdata: output_var_name. So the subclass_id can be mapped to the corresponding output name; by default, the output name is same as subclass_id
# 0.38     04/30/2021  EL/ Fixed a date format bug. Some dates were being recognized as character instead of double. A line is added at 276. 
# 0.37     03/03/2021  YC/ renamed LH's script to 0.37, fix a bug when values are not numeric in calculate_ratios
# 0.36     11/23/2020  LH/ fixed a bug in get_patient_time_series_subdata for: e.g. reading timepoint response: If the value is character, keep it, do not assign it as NA; 
#                                                                                                               if duplicate time points exit, take the latest one.
#                          added a new input in get_lesion_type: lymphnode_marker. This allows more than typical lymph node denotations.    
#                          added three new inputs in get_nontarget_lesions:lymphnode_marker
#                                                                          filter_var and filter_value. This allows filter information based on information in 'filter_var' column, 
#                                                                                                            e.g. if the nontarget denotation includes new lesion
#                          added three new inputs in get_demographic_data: filter_var and filter_value: This allows filter information based on information in 'filter_var' column.
#                                                                          order_var: previously when there is multiple rows corresponding to a patient, is the latest row but not latest date being read. 
#                                                                                          This allows reorder the data_table by 'order_var' column, e.g. select the latest time point
#                          fixed bugs in create_distribution_tables_recist: previous output matrix lack the last row. output_time was varied.line 3114
#                                                                           fix the response categorization after off-treatment time. all others should be 'OS', not 'PD2'
#                                     
#           09/30/2020  LH/ fixed a bug in the initial checking part in align_time series_data function
#
# 0.35      07/31/2020  YC/ added new OS input when assigning tumor responses at line 2872 based on ds_sp_200714 off treatment reason
#                           The new added reasons are "COMPLETED TREATMENT AS PER PROTOCOL","FOLLOWUP NO LONGER REQUIRED PER PROTOCOL"
# 0.34      03/04/2020  YC/ fixed calculation that gives Inf
# 0.33      02/21/2020    YC/ added new OS input when assigning tumor responses at line 2720 based on ds_sp200108 ca 224020 off treatment reason
#                             also changed line 2356 where cur_lesion_measure is NA but is.numeric(cur_lesion_measure) is still true
# 0.32      01/25/2020    BJS/ updated read.delim calls to better recognize anticipated missing 
#                              value strings.
# 0.31      12/16/2019    BJS/ added get_nontarget_progression_time_lnsize_filter, which filters nontarget LNs from
#                              triggering PD in accordance with RECIST 1.1
# 0.30      12/14/2019    BJS/ added new OS input when assigning tumor responses at line 2720 based ib ca 224020 off treatment reason
# 0.29      09/30/2019    YC/ added codes between 1467-1483 to fix an issue where index lesion numbers at pretreatment are assigned NA
# 0.28      09/27/2019    YC/ added new OS input when assigning tumor responses at line 2700 based ib ca 184022 off treatment reason
# 0.27_yc   03/12/2019    YC/ added new OS input when assigning tumor responses at line 2699 based on ca184169 off treatment reason
# 0.26_yc   xxxxxxxx    YC/ bug fix and added baseline volume and offtreatment time integration
# 0.24   09/29/18    BJS/Update to get_patient_time_series_subdata to handle strings that
#                        are entered.  Update calculate_demographic_time to handle missing entries.
# 0.23	 08/16/18    BJS/Add PD2 classification for nontarget progression.
# 0.22	 11/22/17    BJS/Add additional checks into get_patient_time_series_subdata for numeric data.
# 0.21	 10/30/17    BJS/Minor update for adding index lesions indentified by numeric codes.
# 0.20	 10/11/17    BJS/Fixed a bug in 5 mm over nadir increase, where regrowing lesions couldn't
#                        reclassify based on thresholds other than PD.
# 0.19	 10/10/17    BJS/Added 5 mm over nadir increase.
# 0.18	 10/10/17    BJS/Fixed issues in create_distribution_tables_recist: removed the grouping of
#                        patients by treatmenr group.  Also fixed a bug in calculation of change
#                        from nadir for RECIST classification.
# 0.17	 9/24/17     BJS/Add additional processing to account for special considerations with LN index lesions.
# 0.16	 9/23/17     BJS/Add functions for non-target lesions and LN for deriving response status.
# 0.15	 8/11/17     BJS/Add a function to read in an integrated patient dataset from file
#                         and create calibration target files for PW algorithms for lesion
#                         responses.
# 0.14	 8/4/17     BJS/Add a function for adjusting time-series data to a common
#                         timepoint.  Updated lesion calculations and where there are stored
#                         to streamline writing outputs to file.
# 0.13	 7/24/17     BJS/Add a function to calculate off treatment times based on dates
# 0.12	 6/30/17	 BJS/Add calculation of fraction change in SLD to calculate_lesion_summary
#                          Also corrected a bug in get_patient_time_series_subdata(), force conversion to
#                          a numeric data type.
# 0.11       7/17/15     BJS/Added functionality for getting baseline lesion characteristics
# 0.10       4/16/15     BJS/Added function for calculating summary (mean, median, max)
#                          for different patient groups
# 0.09	 3/21/15     BJS/Additional updates to facilitate reading new files
# 0.08       3/19/15     BJS/Fixed a bug that was switching lesion site and lesion type
#                          identifiers when reading in lesion information.  Additional
#                          miscellaneous changes made to accomodate "less clean" datasets
# 0.07       3/12/15     BJS/Changes to enable reading in long gene expression tables
#                          as time series data
# 0.06       3/12/15     BJS/Small bugfix where script was variable to null with
#                          unchanged names
# 0.05       3/11/15     BJS/Small bugfixes to handle single lesion studies
# 0.04       2/25/15     BJS/Changes to lesion derived measures to 
#                        - Handle RECIST studies where only 1 tumor measure is made
#                          TODO: not tested yet
#                        - Provide best calculated response
#                        - Provide ratios of measures to baseline
# 0.03       2/19/15    BJS/Implement techniques based on GBS protocol for 209038 for
#                         calculating lesion measures and handling missing data 
# 0.02       2/8/15     BJS/Re-do lesion level measures so they are {MEASURE{TIME:VALUE}} rather
#                         than {MEASURE{TIME:[VALUE1, VALUE2, ...]}}
#				  Added functions for calculating lesion summaries at the patient level
#                         Added functions for updating variable names
# 0.01	 12/5/14	BJS/initial trial release, probably still an alpha 
#                       version
# TODO: Calculation of TAPD - time after previous dose - for outputs
#       Add function to calculate outputs relative to baseline (ratios)
#
#
# *************** README ***************
# File contains scripts to facilitate the integration of
# PK, outcome, and biomarker datasets.
#
# Dependencies:
# Note, at BMS, to avoid intereference from the firewall:
# setInternet2()
# chooseCRANmirror()
# install.packages('my_package')
#
# Dependencies:
# install.packages('gtools')
#
# Not currently needed, later may want to convert 
# lists to hash tables/dictionaries: install.packages('hash')
#
# The goal is to form an integrated dataset
# for a trial, a key/value pair hierarchy
# This has been implemented in R as lists
# rather than hash tables
#
# Once we have the data in this format, we can create "flat" dataframes
# and export to table files for import into
# other applications such as SimBiology
#

create_patient_data_structure = function() 
  # Inputs:
  #  None
  # Output:
  #  patient_list: a list that has been formatted
  #   appropriately to organize patient information
{
  patient_list = list()
  patient_list[['first_dose_date']] = NULL
  patient_list[['all_dose_dates']] = c()
  patient_list[['demographics']] = list()
  patient_list[['index_lesions']] = list()
  patient_list[['time_series']] = list()
  # Genomics data not yet implemented separated, 
  # could probably pair at lesion or
  # patient level
  # patient_list[['genomics']] = list()
  
  patient_list
}


create_trial_data_structure = function(filename, patient_id_header = "USUBJID", trial_list = '')
  # This function looks in a file with name filename,
  # and assigns the patients in trial_list to
  # the appropriate treatment group
  # Inputs:
  #  filename: the name of a file to read
  #  patient_id_header: the header name in the file for the patient identifier data 
  #  trial_list: a dictionary with trial data.  If not provided,
  #   one will be generated.
  # Output:
  #  trial_list: dictionary with patients
{
  data_table = read.delim(filename, na.strings = c("#N/A","."), check.names = FALSE)
  patient_ids = unique(data_table[,patient_id_header])
  if (class(trial_list) != "list")
  {
    trial_list = list()
  }
  for (the_patient_id in patient_ids)
  {
    if (!(the_patient_id %in% names(trial_list)))
    {
      patient_list = create_patient_data_structure()
      trial_list[[the_patient_id]] = patient_list
    }
  }
  trial_list
}


get_demographic_data = function(trial_list, filename, demographic_header_vector = c("TRTGRP","WT","OFFTRTPH","I_STAGE"), patient_id_header = "USUBJID", filter_var='', filter_value='', order_var='')
  # This function looks in a file with name filename,
  # and adds the demographic annotation to the patient
  # Note this isn't necessarily just demographic data,
  # but could be any data that can be captured
  # as a single key: value pair without time
  # series considerations
  # Inputs:
  #  trial_list
  #  filename: the name of a file to read
  #  demographic_header_vector: a list of demographic headers
  #  patient_id_header: the header name in the file for the patient identifier data 
#  treatment_key: the headername with the information on treatment group
# Output:
#  trial_list: list with patient data read in
{
  continue_flag = TRUE
  options(stringsAsFactors = FALSE)
  if (class(filename) == "character")
  {
    data_table = read.delim(filename, na.strings = c("#N/A","."), check.names = FALSE)
  } else if (class(filename) == "data.frame") {
    data_table = filename
  }
  
  if(nchar(filter_var)>0){
    data_table = data_table[which(data_table[,filter_var]!=filter_value),]
  }
  
  patient_ids = unique(data_table[,patient_id_header])
  row_names = rownames(data_table)
  if (!(length(row_names) == length(patient_ids)))
  {
    if (class(filename) == "character"){
      print_string = paste('Warning, ',filename,' does not contain 1 patient ID per row.  Only the latest rows occurring in the file for each patient will be retained.',sep = "")
    }else{
      print_string = paste('Warning, input data frame does not contain 1 patient ID per row.  Only the latest rows occurring in the file for each patient will be retained.',sep = "")
    }
    print(print_string)
    # This is just a warning, can proceed
    # With the caveat that we overwrite values
    # and just use the most recent
    
    #
    if(nchar(order_var)>0){
      print("reorder by the order_var ...")
      data_table = data_table[order(data_table[,order_var]),]	
    }
    continue_flag = TRUE
  } else {
    rownames(data_table) = data_table[,patient_id_header]
  }
  patient_ids = intersect(patient_ids, names(trial_list))
  the_rownames = rownames(data_table[which(data_table[, patient_id_header] %in% patient_ids),])
  
  if (continue_flag)
  {
    # for (the_patient_id in patient_ids)
    for (the_row_name in the_rownames)
    {
      the_patient_id = data_table[the_row_name, patient_id_header]
      # Now read in the 'demographics'
      for (the_demo_key in demographic_header_vector)
      {
        the_datum = check_and_convert_numeric(data_table[the_row_name, the_demo_key])
        the_datum_date = convert_to_date(the_datum)
        if (class(the_datum_date) == "Date")
        {
          the_datum = the_datum_date
        }
        trial_list[[the_patient_id]][['demographics']][[the_demo_key]] = the_datum
      }
    }
  }
  trial_list
}


get_initial_dose_date = function(trial_list, filename, dose_date_header = "DOSESD", patient_id_header = "USUBJID", drop_if_missing = FALSE)
  # This function looks in a file with name filename,
  # and pulls the treatment start date
  # Inputs:
  #  trial_list:
  #  filename:          the name of a file to read
  #  dose_date_header:  the header name with the information on dose dates
  #  patient_id_header: the header name in the file for the patient identifier data 
  #  drop_if_missing:   (boolean) whether to drop missing variables 
  # Output:
  #  trial_list: dictionary with patient dose date data
{
  continue_flag = TRUE
  options(stringsAsFactors = FALSE)
  data_table = read.delim(filename, na.strings = c("#N/A","."), check.names = FALSE)
  patient_ids = unique(data_table[,patient_id_header])
  patient_ids = intersect(patient_ids, names(trial_list))
  row_names = rownames(data_table)
  
  if (continue_flag)
  {	
    for (the_patient_id in patient_ids)
    {
      # Add patients in case we find new ones
      # if (!(the_patient_id %in% names(trial_list)))
      # {
      # 	patient_list = create_patient_data_structure()
      # 	trial_list[[the_patient_id]] = patient_list
      # }
      # Now make a list of [dose_dates]
      # so we can then add the 1st ones
      # into the trial_list
      check_rows = which(data_table[,patient_id_header]==the_patient_id)			
      date_list = data_table[check_rows,dose_date_header]
      new_date_list = vector()
      class(new_date_list) = "Date"
      for (the_date in date_list)
      {
        the_date = convert_to_date(the_date)
        the_date = as.Date(the_date)
        new_date_list = c(new_date_list, the_date)
      }
      new_date_list = sort(new_date_list)
      trial_list[[the_patient_id]][['first_dose_date']] = new_date_list[1]
      if (drop_if_missing)
      {
        if (length(new_date_list) == 0)
        {
          trial_list = trial_list[names(trial_list) != the_patient_id]
        }
      }
    }
  }
  trial_list
}


get_lesion_type = function(trial_list, filename, lesion_id_header = "DZTU", lesion_type_header = "LSTYL", disease_site_header = "DZSITE", patient_id_header = "USUBJID", index_lesion_values = c("index","INDEX","TARGET","target",1),lymphnode_marker=c(" node"," node ", "node ", "lymphnode", "lymph node","lymph","lynph", " ln","ln ", " ln "))
  # This function looks in a file with name filename,
  # and pulls the lesion information, but not measures
  # Inputs:
  #  trial_list:          dict with all the trial information that has been compiled so far
  #  filename:            the name of a file to read
  #  lesion_id_header:    the header name in the file for the tumor lesion information 
  #  lesion_type_header:  the headername with the information on lesion types, only target/index lesions will be retained
  #  disease_site_header: the headername with the information on where the lesion comes from
  #  patient_id_header:   the header name in the file for the patient identifier data
  #  index_lesion_values: values for index lesions in lesion_type_header
  # Output:
#  trial_list: dictionary updated with lesion data
{
  continue_flag = TRUE
  options(stringsAsFactors = FALSE)
  data_table = read.delim(filename, na.strings = c("#N/A","."), check.names = FALSE, fileEncoding="latin1") 
  # Lu added this fileEncoding command on 20230717, because there is error in reading e.g. cd184169 files: Error in tolower("sd\xdeL") : invalid multibyte string 1
  # fileEncoding = "UTF-8") this will result in losing data
  patient_ids = unique(data_table[,patient_id_header])
  patient_ids = intersect(patient_ids, names(trial_list))
  row_names = rownames(data_table)
  
  if (continue_flag)
  {	
    for (the_patient_id in patient_ids)
    {
      # Add patients in case we find new ones
      if (!(the_patient_id %in% names(trial_list)))
      {
        trial_list[[the_patient_id]] = create_patient_data_structure()
      }
      # Now make a list of [lesion_id_header]
      # so we can then add the 1st ones
      # into the trial_list
      check_rows = which(data_table[,patient_id_header]==the_patient_id)
      check_table = data_table[check_rows,]
      keep_lesion_ids = vector()
      for (the_row in rownames(check_table))
      {
        if (check_table[the_row,lesion_type_header] %in% index_lesion_values)
        {
          keep_lesion_ids = c(keep_lesion_ids,check_table[the_row,lesion_id_header])
        }
      }
      keep_lesion_ids = unique(keep_lesion_ids)
      for (the_lesion_id in keep_lesion_ids)
      {
        check_rows = which(check_table[,lesion_id_header] == the_lesion_id)
        lesion_information = check_table[check_rows,disease_site_header]
        ulesion_information = unique(lesion_information)
        # filter out the empty information site information
        ulesion_information = setdiff(ulesion_information,"")
        if (!(the_lesion_id %in% names(trial_list[[the_patient_id]][['index_lesions']])))
        {
          trial_list[[the_patient_id]][['index_lesions']][[the_lesion_id]] = list()
          filtered_lesion_information = ulesion_information[which.max(tabulate(match(lesion_information, ulesion_information)))]
          trial_list[[the_patient_id]][['index_lesions']][[the_lesion_id]][[disease_site_header]] = filtered_lesion_information
          # Also add a flag to indicate whether this is a lymph node, which is treated differently in evaluating complete response
          #if ((grepl(" node", tolower(filtered_lesion_information), fixed = TRUE)) | (grepl(" node ", tolower(filtered_lesion_information), fixed = TRUE)) | (grepl("lymphnode", tolower(filtered_lesion_information), fixed = TRUE)) | (grepl("LN", tolower(filtered_lesion_information), fixed = TRUE)))
          if (grepl(paste(lymphnode_marker,collapse="|"), tolower(filtered_lesion_information), fixed = F))
            #if (grepl(paste(lymphnode_marker,collapse="|"), toupper(filtered_lesion_information), fixed = F))
          {
            trial_list[[the_patient_id]][['index_lesions']][[the_lesion_id]][["NODE_FLAG"]] = 1
          } else {
            trial_list[[the_patient_id]][['index_lesions']][[the_lesion_id]][["NODE_FLAG"]] = 0
          }
        }
        
      }
    }
  }
  trial_list
}


get_lesion_level_measures = function(trial_list, filename, lesion_id_header = "DZTU", lesion_eval_date_header = "DZEVD", lesion_measure_names = c("MEASURE"), patient_id_header = "USUBJID")
  # This function looks in a file with name filename,
  # and pulls the lesion measure information, combining them in under tumor_measure_name
  # Inputs:
  #  trial_list: list structure with all the trial information that has been compiled so far
  #  filename: the name of a file to read
  #  lesion_id_header: the header name in the file for the tumor lesion information 
  #  lesion_eval_date_header: date the lesion sizes were evaluated, in a format that will be recognized by R
  #  lesion_measure_names: keys for lesions size, or other measure, entered as a vector since there may be multiple, e.g. perpendicular diameter measures
  #  patient_id_header: the header name in the file for the patient identifier data
  # Output:
  #  trial_list: dictionary updated with lesion data
{
  continue_flag = TRUE
  options(stringsAsFactors = FALSE)
  data_table = read.delim(filename, na.strings = c("#N/A","."), check.names = FALSE)
  patient_ids = unique(data_table[,patient_id_header])
  row_names = rownames(data_table)
  # We won't add new patients at this point, since we reference to
  # the first dose date, which may not be here.
  patient_ids = intersect(patient_ids, names(trial_list))
  
  if (continue_flag)
  {	
    for (the_patient_id in patient_ids)
    {
      # Now make a list of [lesion_id_header]
      # so we can then add the 1st ones
      # into the trial_list
      check_rows = which(data_table[,patient_id_header]==the_patient_id)
      check_table = data_table[check_rows,]
      # We only add the lesion data if we have
      # evidence that this has been recognized as an index lesion
      keep_lesion_ids = vector()
      reference_date = trial_list[[the_patient_id]][['first_dose_date']]
      for (the_row in rownames(check_table))
      {
        if (check_table[the_row,lesion_id_header] %in% names(trial_list[[the_patient_id]][['index_lesions']]))
        {
          keep_lesion_ids = c(keep_lesion_ids,check_table[[the_row,lesion_id_header]])
        }
      }
      keep_lesion_ids = unique(keep_lesion_ids)
      for (the_lesion_id in keep_lesion_ids)
      {
        for (lesion_measure_name in lesion_measure_names)
        {
          trial_list[[the_patient_id]][['index_lesions']][[the_lesion_id]][[lesion_measure_name]] = list()
          check_rows = which(check_table[,lesion_id_header] == the_lesion_id)
          for (the_row in check_rows)
          {
            the_date = convert_to_date(check_table[the_row,lesion_eval_date_header])
            if (!(is.null(the_date)))
            {
              days_post_dose = the_date - reference_date
              current_datum = check_and_convert_numeric(check_table[the_row,lesion_measure_name])
              # Force numeric list name to character, avoids issues in R
              trial_list[[the_patient_id]][['index_lesions']][[the_lesion_id]][[lesion_measure_name]][[as.character(as.numeric(days_post_dose))]] = current_datum
            }
          }
        }
      }
    }
  }
  trial_list
}


get_patient_time_series_data = function(trial_list, filename, date_header = "", atafd_header = "", measure_header_vector = c(), patient_id_header = "USUBJID", transpose = FALSE)
  # This function looks in a file with name filename,
  # and pulls the patient measures that have been recorded
  # in a time-series format, e.g. general biomarkers or PK data
  # Inputs:
  #  trial_list: dict with all the trial information that has been compiled so far
  #  filename: the name of a file to read.  This may also be a data frame if
  #            one has already been prepared to read to the patient data structure.
  #  date_header: date for the time-series measure
  #   atafd_header: actual time after first dose (days) header.  optional, date header will be used if provided,
  #   but at least one of these must be provided
  #  measure_header_vector: an R vector (e.g. c("header1","header2",...)) of
#   header names
#  patient_id_header: the header name in the file for the patient identifier data
#  transpose: TRUE/FALSE, whether to transpose the table, only used for reading from file
#
# Output:
#  trial_list: dictionary updated with lesion data
#
{
  continue_flag = TRUE
  options(stringsAsFactors = FALSE)
  if (class(filename) == "character")
  {
    # We set check.names to FALSE so we can have hyphens
    if (transpose == FALSE)
    {
      data_table = read.delim(filename, na.strings = c("#N/A","."), check.names = FALSE)
    }
    if (transpose == TRUE)
    {
      data_table = read.tdelim(filename, na.strings = c("#N/A","."), check.names = FALSE)
    }
  } else if (class(filename) == "data.frame") {
    data_table = filename
  } else {
    print("A recognized file or dataframe must be provided.  Exiting...")
    continue_flag = FALSE
  }
  
  # Enforce uppercase to avoid case mismatch issues
  measure_header_vector = toupper(measure_header_vector)
  patient_id_header = toupper(patient_id_header)
  rownames(data_table) = toupper(rownames(data_table))
  colnames(data_table) = toupper(colnames(data_table))
  data_table[,patient_id_header] = toupper(data_table[,patient_id_header])
  patient_ids = (unique(data_table[,patient_id_header]))
  row_names = toupper(rownames(data_table))
  
  patient_ids = unique(data_table[,patient_id_header])
  row_names = rownames(data_table)
  # We won't add new patients at this point, since we reference to
  # the first dose date, which may not be here.
  patient_ids = intersect(patient_ids, names(trial_list))
  if (length(patient_ids) == 0 && class(filename) == "character")
  {
    cat("No patient IDs from the trial_list were found in the file: ",filename,".\n",sep='')
  }
  
  atafd_header = toupper(atafd_header)
  date_header = toupper(date_header)
  
  if (nchar(date_header) == 0)
  {
    if (nchar(atafd_header) == 0)
    {
      continue_flag = FALSE	
    } else {
      time_header = atafd_header
    }
  } else {
    time_header = date_header
  }
  
  if (continue_flag)
  {
    for (the_patient_id in patient_ids)
    {
      check_rows = which(data_table[,patient_id_header]==the_patient_id)
      reference_date = trial_list[[the_patient_id]][['first_dose_date']]
      for (measure_header in measure_header_vector)
      {
        temp_list = list()
        for (the_row in check_rows)
        {
          
          the_value = check_and_convert_numeric(data_table[the_row,measure_header])
          # We loosen the restriction that time series data must be numeric to be of interest
          # if (is.numeric(the_value))
          # {
          if (time_header == date_header)
          {
            the_date = convert_to_date(data_table[the_row,time_header])
            temp_list[[as.character(as.numeric(the_date-reference_date))]] = the_value
          } else {
            # Otherwise we assume this is already corrected ATAFD
            the_date = check_and_convert_numeric(data_table[the_row,time_header])
            temp_list[[as.character(the_date)]] = the_value
          }
          # }
        }
        
        
        # Force numeric list name to character, avoids issues in R
        # Also nice since this avoids potential roundoff/floating point issues
        if (length(temp_list) > 0)
        {
          trial_list[[the_patient_id]][['time_series']][[measure_header]] = temp_list
        }
      }
    }
  }
  trial_list
}


get_patient_dataframe = function(trial_list)
  # This function takes patient data
  # and converts it to a flat data table
{
  patient_list = names(trial_list)
  # You can change these defaults, this was just consistent with my datasets
  column_names_1 = c("USUBJID","first_dose_date")
  column_names_demo = c()
  column_names_time_series = c()
  for (the_patient_id in patient_list)
  {
    column_names_demo = union(column_names_demo, names(trial_list[[the_patient_id]][["demographics"]]))
  }
  # Now we scan through the time_series data for headers
  for (the_patient_id in patient_list)
  {
    column_names_time_series = union(column_names_time_series, names(trial_list[[the_patient_id]][["time_series"]]))
  }
  column_names_time_series = c("TIME","TAPD",column_names_time_series)
  column_names = c(column_names_1, column_names_demo, column_names_time_series, recursive = TRUE)
  patient_data_frame = data.frame(matrix(nrow = 0,ncol=length(column_names)))
  colnames(patient_data_frame) = column_names
  # Now we start building the data frame.  We start at the time series data.
  for (the_patient_id in patient_list)
  {
    #print(the_patient_id)
    # First we scan through and get the needed time points
    time_point_vector = c()
    for (the_measurement in column_names_time_series)
    {
      time_point_vector = union(time_point_vector,(names(trial_list[[the_patient_id]][["time_series"]][[the_measurement]])))
    }
    # Force these to numeric, they should be characters in R so far
    time_point_vector = as.numeric(time_point_vector)
    time_point_vector = sort(time_point_vector)
    one_patient_data_frame = data.frame(matrix(NA, nrow = length(time_point_vector), ncol = length(column_names)))
    names(one_patient_data_frame) = column_names
    row_counter = 0
    for (the_time_point in time_point_vector)
    {
      row_counter = row_counter + 1
      the_time_point_char = as.character(the_time_point)
      #print(column_names_time_series)
      for (the_column_name in column_names_time_series)
      {
        # column_index = which(the_column_name, column_names)
        if (the_column_name == "TIME")
        {
          one_patient_data_frame[row_counter,the_column_name] = the_time_point
        } else if (the_column_name == "TAPD")
        {
          if (length(one_patient_data_frame[["all_dose_dates"]])>0)
          {
            the_date = the_time_point + one_patient_data_frame[["first_dose_date"]]
            last_date = find(the_date > one_patient_data_frame[["all_dose_dates"]])
            last_date = one_patient_data_frame[["all_dose_dates"]][last_date[1]]
            one_patient_data_frame[row_counter,the_column_name] = as.numeric(the_date - last_date)
          } else {
            one_patient_data_frame[row_counter,the_column_name] = NA
          }
          
        } else if (the_column_name %in% names(trial_list[[the_patient_id]][["time_series"]])) {
          if(the_time_point_char %in% names(trial_list[[the_patient_id]][["time_series"]][[the_column_name]])) 
          {
            one_patient_data_frame[row_counter,the_column_name] = trial_list[[the_patient_id]][["time_series"]][[the_column_name]][[the_time_point_char]]
          }
        }
      }
      #print(column_names_demo)
      for (the_column_name in column_names_demo)
      {
        if (the_column_name %in% names(trial_list[[the_patient_id]][["demographics"]])) 
        {
          if (class(trial_list[[the_patient_id]][["demographics"]][[the_column_name]]) == "Date")
          {
            one_patient_data_frame[row_counter,the_column_name] = as.character(trial_list[[the_patient_id]][["demographics"]][[the_column_name]])
          } else {
            #print(the_column_name)
            #browser()
            one_patient_data_frame[row_counter,the_column_name] = trial_list[[the_patient_id]][["demographics"]][[the_column_name]]
          }
        }
      }
      #print(column_names_1)
      for (the_column_name in column_names_1)
      {
        if (the_column_name == "USUBJID")
        {
          one_patient_data_frame[row_counter,the_column_name] = the_patient_id
        } else if (the_column_name == "first_dose_date") {
          if ("first_dose_date" %in% names(trial_list[[the_patient_id]])) 
          {
            one_patient_data_frame[row_counter,the_column_name] = as.character(trial_list[[the_patient_id]][[the_column_name]])
          } else {
            one_patient_data_frame[row_counter,the_column_name] = NA
            print(paste("Note: no first dose date information available for ",the_patient_id,"!",sep = ""))
          }
        }
      }
    }
    patient_data_frame = rbind(patient_data_frame, one_patient_data_frame)
  }
  patient_data_frame
}


get_lesion_dataframe = function(trial_list)
  # This function takes lesion data
  # and converts it to a flat data table
{
  patient_list = names(trial_list)
  # You can change these defaults, this was just consistent with my datasets
  column_names_1 = c("USUBJID","DZTU")
  column_names_demo = c()
  column_names_time_series = c("TIME")
  measure_names_time_series = list()
  
  # First scan through the time series measures and determine how many elements are needed for each
  measure_lengths_time_series = list()
  for (the_patient_id in patient_list)
  {
    for (the_lesion_id in names(trial_list[[the_patient_id]][['index_lesions']]))
    {
      for (the_variable in names(trial_list[[the_patient_id]][['index_lesions']][[the_lesion_id]]))
      {
        if (class(trial_list[[the_patient_id]][['index_lesions']][[the_lesion_id]][[the_variable]]) == "list")
        {
          if (!(the_variable %in% names(measure_lengths_time_series)))
          {
            measure_lengths_time_series[[the_variable]] = 0
          }
          for (the_time_point in names(trial_list[[the_patient_id]][['index_lesions']][[the_lesion_id]][[the_variable]]))
          {
            the_sub_counter = 0
            for (the_sub_measure in trial_list[[the_patient_id]][['index_lesions']][[the_lesion_id]][[the_variable]][[the_time_point]])
            {
              the_sub_counter = the_sub_counter + 1
            }
            measure_lengths_time_series[[the_variable]] = max(measure_lengths_time_series[[the_variable]], the_sub_counter)
          }
        } else {
          # Otherwise, we can go ahead and specify the column name
          column_names_demo = union(column_names_demo, c(the_variable))
        }				
      }
    }
  }
  
  # Now we create the variable names to flatten repeated lesion measures at each time point
  measure_names_time_series = list()
  for (the_measure_name in names(measure_lengths_time_series))
  {
    if (measure_lengths_time_series[[the_measure_name]] == 1)
    {
      column_names_time_series = union(column_names_time_series, the_measure_name)
      measure_names_time_series[[the_measure_name]] = the_measure_name
    } else {
      for (the_sub_counter in seq(1, measure_lengths_time_series[[the_variable]]))
      {
        column_names_time_series = union(column_names_time_series, c(paste(the_measure_name,the_sub_counter,sep="_")))
        measure_names_time_series[[the_variable]] = c(measure_names_time_series[[the_variable]], paste(the_measure_name,the_sub_counter,sep = "_"))				
      }
    }
  }
  column_names = c(column_names_1, column_names_demo, column_names_time_series, recursive = TRUE)
  lesion_data_frame = data.frame(matrix(nrow = 0,ncol=length(column_names)))
  colnames(lesion_data_frame) = column_names
  
  for (the_patient_id in patient_list)
  {
    for (the_lesion_id in names(trial_list[[the_patient_id]][['index_lesions']]))
    {
      # First we scan through and get the needed time points for the lesion
      time_point_vector = c()
      for (the_measurement in names(measure_names_time_series))
      {
        if (the_measurement %in% names(trial_list[[the_patient_id]][['index_lesions']][[the_lesion_id]]))
        {
          time_point_vector = union(time_point_vector,names(trial_list[[the_patient_id]][['index_lesions']][[the_lesion_id]][[the_measurement]]))
        }
      }
      
      # Define a new data frame here for the lesion
      time_point_vector = as.numeric(time_point_vector)
      time_point_vector = sort(time_point_vector)
      one_lesion_data_frame = data.frame(matrix(NA, nrow = length(time_point_vector), ncol = length(column_names)))
      names(one_lesion_data_frame) = column_names
      row_counter = 0
      
      for (the_time_point in time_point_vector)
      {
        row_counter = row_counter + 1
        the_time_point_char = as.character(the_time_point)
        one_lesion_data_frame[row_counter,"TIME"] = the_time_point
        for (the_measure_id in names(measure_names_time_series))
        {
          if (the_measure_id %in% names(trial_list[[the_patient_id]][['index_lesions']][[the_lesion_id]]))
          {				
            the_sub_measure_id_vector = measure_names_time_series[[the_measure_id]]
            the_sub_measure_length = min(length(the_sub_measure_id_vector),length(trial_list[[the_patient_id]][['index_lesions']][[the_lesion_id]][[the_measure_id]][[the_time_point_char]]))
            if (the_sub_measure_length > 0)
            {
              for (the_sub_measure_index in seq(1, the_sub_measure_length))
              {
                the_sub_measure_id = the_sub_measure_id_vector[the_sub_measure_index]
                one_lesion_data_frame[row_counter,the_sub_measure_id] = trial_list[[the_patient_id]][['index_lesions']][[the_lesion_id]][[the_measure_id]][[the_time_point_char]][the_sub_measure_index]
              }
            }
          }
        }
        
        one_lesion_data_frame[row_counter,"USUBJID"] = the_patient_id
        one_lesion_data_frame[row_counter,"DZTU"] = the_lesion_id
        for (the_column_name in column_names_demo)
        {
          if (the_column_name %in% names(trial_list[[the_patient_id]][['index_lesions']][[the_lesion_id]])) 
          {
            one_lesion_data_frame[row_counter,the_column_name] = trial_list[[the_patient_id]][['index_lesions']][[the_lesion_id]][[the_column_name]]
          }
        }
        
      }
      lesion_data_frame = rbind(lesion_data_frame, one_lesion_data_frame)			
    }
  }
  lesion_data_frame
}			


get_lesion_baseline_dataframe = function(trial_list)
  # This function takes lesion data
  # and converts it to a flat data table
{
  patient_list = names(trial_list)
  column_names = c("USUBJID", "DZTU", "LESIONSIZE1_MM", "LESIONSIZE2_MM")
  lesion_data_frame = data.frame(matrix(nrow = 0,ncol=length(column_names)))
  for (the_patient_id in patient_list)
  {
    if ("baseline_lesions" %in% names(trial_list[[the_patient_id]]))
    {
      for (the_lesion_id in names(trial_list[[the_patient_id]][["baseline_lesions"]]))
      {
        measure_1 = trial_list[[the_patient_id]][["baseline_lesions"]][[the_lesion_id]][["LESIONSIZE1_MM"]]
        measure_2 = trial_list[[the_patient_id]][["baseline_lesions"]][[the_lesion_id]][["LESIONSIZE2_MM"]]
        the_new_row = data.frame(matrix(nrow = 1,ncol=length(column_names)))
        the_new_row[1,1] = the_patient_id
        the_new_row[1,2] = the_lesion_id
        # Only keep lesions with at least 1 non-null baseline measure
        if (!is.null(measure_1))
        {
          the_new_row[1,3] = measure_1
          
          if (!is.null(measure_2))
          {
            the_new_row[1,4] = measure_2 
          }
          
          lesion_data_frame = rbind(lesion_data_frame, the_new_row)
        }
      }
    }
  }
  colnames(lesion_data_frame) = column_names
  lesion_data_frame
}


get_patient_time_series_subdata = function(trial_list, filename, date_header="", atafd_header="", measure_header="", subclass_ids=c(), output_var_name = subclass_ids, data_header="", patient_id_header = "USUBJID")
  # This function looks in a file with name filename,
  # and pulls the patient measures that have been recorded
  # in a time-series format but broken into subfields, e.g. as is
  # generally done for lab data
  # Inputs:
  #  trial_list: dict with all the trial information that has been compiled so far
  #  filename: the name of a file to read
  #  date_header: date for the time-series measure
  #   atafd_header: actual time after first dose (days) header.  optional, date header will be used if provided,
  #   but at least one of these must be provided
  #  measure_header: the header to scan for subclass IDs
#  sublass_ids: a vector of subclass IDs to include, e.g. these are picked out of the measure_header column
#               as being the measures of interest
# output_var_name: a vector corresponding to subclass_ids, as the name we want the output to be
#  data_header: column that includes the values
#  patient_id_header: the header name in the file for the patient identifier data
#
# Output:
#  trial_list: list updated with patient data
{
  
  continue_flag = TRUE
  options(stringsAsFactors = FALSE)
  original_data_table = read.delim(filename, na.strings = c("#N/A","."), check.names = FALSE)
  
  if (length(output_var_name)>0){
    require(plyr)
    original_data_table[,measure_header] <- mapvalues(original_data_table[,measure_header], 
                                                      from=subclass_ids, 
                                                      to=output_var_name)
    subclass_ids =  output_var_name
  }
  
  # We enforce uppercase as much as possible to avoid issues in variable names and identifiers
  ## I came across issues in consistency in the test name when importing data
  patient_id_header = toupper(patient_id_header)
  measure_header = toupper(measure_header)
  subclass_ids = toupper(subclass_ids)
  rownames(original_data_table) = toupper(rownames(original_data_table))
  colnames(original_data_table) = toupper(colnames(original_data_table))
  # browser()
  original_data_table[,patient_id_header] = toupper(original_data_table[,patient_id_header])
  patient_ids = (unique(original_data_table[,patient_id_header]))
  row_names = (rownames(original_data_table))
  # We won't add new patients at this point, since we reference to
  # the first dose date, which may not be here.
  #browser()
  patient_ids = intersect(patient_ids, names(trial_list))
  original_data_table[,measure_header] = toupper(original_data_table[,measure_header])	
  if (length(subclass_ids) == 0)
  {
    # We just take all the subclasses if none are specified
    subclass_ids = (unique(original_data_table[,measure_header]))
  }
  data_header = toupper(data_header)
  atafd_header = toupper(atafd_header)
  date_header = toupper(date_header)	
  if (nchar(date_header) == 0)
  {
    if (nchar(atafd_header) == 0)
    {
      continue_flag = FALSE	
    } else {
      time_header = atafd_header
    }
  } else {
    time_header = date_header
  }
  
  # browser()
  # We don't force the date_header or measure_header to an upper
  new_column_names = (c(patient_id_header, time_header, subclass_ids, recursive = TRUE))
  new_data_frame = data.frame(matrix(nrow = 0,ncol=length(new_column_names)))
  colnames(new_data_frame) = new_column_names
  if (continue_flag)
  {
    # Now we start building the new data frame.
    for (the_patient_id in patient_ids)
    {
      all_rows_to_combine = c()
      for (the_subclass_id in subclass_ids)
      {
        # Get the rows with the measure that we want to work with
        the_rows = intersect(which(original_data_table[,patient_id_header]==the_patient_id),which(original_data_table[,measure_header]==the_subclass_id))
        
        if (length(the_rows) > 0)
        {
          all_rows_to_combine = c(all_rows_to_combine, the_rows)
        }
      }
      
      # Now scan through and check the time points to "collapse"
      the_date_vector = c()
      for (the_row_number in all_rows_to_combine)
      {
        the_date = original_data_table[the_row_number,time_header]
        if (!is.null(the_date) & the_date!=""){
          the_date_vector = c(the_date_vector, c(the_date))
        }
      }
      the_date_vector = unique(the_date_vector)
      
      if (length(the_date_vector) > 0)
      {
        # browser()
        one_patient_data_frame = data.frame(matrix(NA, nrow = length(the_date_vector), ncol = length(new_column_names)))
        colnames(one_patient_data_frame) = new_column_names
        for (the_one_patient_row in seq(1, length(the_date_vector)))
        {
          the_date = the_date_vector[the_one_patient_row]
          one_patient_data_frame[the_one_patient_row,patient_id_header] = the_patient_id
          one_patient_data_frame[the_one_patient_row,time_header] = the_date
          for (the_subclass_id in subclass_ids)
          {
            
            the_original_row_number = intersect(which(original_data_table[,patient_id_header]==the_patient_id),intersect(which(original_data_table[,measure_header]==the_subclass_id),which(original_data_table[,time_header]==the_date)))
            # It is possible to end up with more than 1 measure, if we are tracking
            # measure by the date and we take multiple measures on the same date,
            # e.g. sampling at 1 or 4 hr intervals.  In this case try to use atafd header
            # with good specificity for the time points to
            # separate them.  Here, we just take the median of
            # all datapoints from the same day.  This seems safer
            # than the mean, since concentration data may not
            # be normally distributed.
            if (length(the_original_row_number) >= 1)
            {
              the_value = original_data_table[the_original_row_number,data_header]
              the_value = check_and_convert_numeric(the_value)
              
              if (length(the_value) >= 1)
              {
                if (is.numeric(the_value))
                {
                  if ((sum(is.na(the_value)) > 0) & (length(the_value) > 1))
                  {
                    keep_indices = which(!(is.na(the_value)))
                    the_value = the_value[keep_indices]
                  }
                  the_value = median(the_value)
                }else if (is.character(the_value)) {
                  # If this is a character string we set
                  # the value to the latest one
                  the_value = the_value[length(the_value)]
                }
                # changed here. if it is a character, we keep output as character, take the latest one. e.g. timepoint response
              } 
              one_patient_data_frame[the_one_patient_row,the_subclass_id] = the_value
            }
          }
        }
        new_data_frame = rbind(new_data_frame, one_patient_data_frame)
      }
    }
    
    
    # Now that we have reformatted the data we can just read it
    # to the trial/patient data structure
    # But first we have to apply a small tweak to the dates in case we read in date data
    # from file as absolute time (days) after first dose rather than dates
    
    if (time_header == atafd_header)
    {
      
      for (the_rowname in rownames(new_data_frame))
      {
        the_patient_id = new_data_frame[the_rowname,patient_id_header]
        first_dose_date = trial_list[[the_patient_id]][["first_dose_date"]]
        the_time = as.numeric(new_data_frame[the_rowname,time_header])
        new_data_frame[the_rowname,time_header] = as.character(first_dose_date + the_time)
      }
    }
    trial_list = get_patient_time_series_data(trial_list, new_data_frame, date_header = time_header, measure_header_vector = subclass_ids, patient_id_header = patient_id_header)
  }
  trial_list
}


check_and_convert_numeric = function(the_datum)
  # This function takes an input and
  # tries to convert it to a number.
{
  # We suppress warnings here, because we
  # know trying to convert a string will result in a
  # warning.
  options(warn=-1)
  if (!is.null(the_datum))
  {
    if (!all(is.na(as.numeric(the_datum))))
    {
      the_datum = as.numeric(the_datum)
      # Here, we default nissing values to NA.  Could use NULL as well.
    } else if (all(the_datum %in% c('.',''))) { # it might have c("","")  
      the_datum = NA
    }
  }
  the_datum
}


convert_to_date = function(the_date)
  # This function supplement
  # R's as.Date function by identifying
  # date formats commonly used in BMS clinical 
  # datasets and converting them to an R dates
{
  #if (is.null(the_date))
  #{
  #	# Pass-through...
  #	the_date = the_date
  #
  #} else if ((class(the_date) == "Date") | (is.na(the_date)))
  if ((class(the_date) == "Date") | (is.na(the_date)))
  {
    # Pass-through...
    the_date = the_date
  } else {
    the_date = as.character(the_date)
    # TODO can expand this to be more exhaustive of potential
    # date formats
    if (nchar(the_date) == 9)
    {
      # the_date = tolower(the_date)
      # We could be more explicit here for the allowed ranges
      pattern_1 = "[[:digit:]][[:digit:]][[:digit:]][[:digit:]][[:alpha:]][[:alpha:]][[:alpha:]][[:digit:]][[:digit:]]"
      pattern_2 = "[[:digit:]][[:digit:]][[:alpha:]][[:alpha:]][[:alpha:]][[:digit:]][[:digit:]][[:digit:]][[:digit:]]"
      if (grepl(pattern_1,the_date))
      {
        the_date = as.Date(the_date,format="%Y%b%d")
      } else if (grepl(pattern_2,the_date))
      {
        the_date = as.Date(the_date,format="%d%b%Y")
      } else 
      {
        the_date = NULL
      }
    } else if (nchar(the_date) == 8)
    {
      pattern_3 = "[[:digit:]][[:digit:]][[:digit:]][[:digit:]][[:digit:]][[:digit:]][[:digit:]][[:digit:]]"
      # We could add pattern_4 to deal with cases like "7-Mar-11" but be careful!  We're coercing
      # an ambiguous assumption about the date formatting.
      # This introduces an additional issue with the century: e.g. 19XX or 20XX,
      # so this needs to be done in the file and not made automatic.
      # pattern_4= "[[:digit:]][[:punct:]][[:alpha:]][[:alpha:]][[:alpha:]][[:punct:]][[:digit:]][[:digit:]]"
      if (grepl(pattern_3,the_date))
      {
        the_date = as.Date(the_date,format="%Y%m%d")
        # } else if (grepl(pattern_4,the_date))
        # {
        # 	the_date = as.Date(the_date,format="%d%m%Y")
      } else
      {
        the_date = NULL
      }
    } else if (nchar(the_date) == 10) 
    {
      pattern_5 = "[[:digit:]][[:digit:]][[:digit:]][[:digit:]][[:punct:]][[:digit:]][[:digit:]][[:punct:]][[:digit:]][[:digit:]]"
      # Although not totally unambiguous, we assume "YYYY-MM-DD" format
      if (grepl(pattern_5,the_date))
      {
        if (grepl("/",the_date)){
          the_date = as.Date(the_date,format="%Y/%m/%d")
        }else{
          the_date = as.Date(the_date,format="%Y-%m-%d")
        }
      } else 
      {
        the_date = NULL
      }			
    } else {
      the_date = NULL
    }
  }
  the_date
}


update_variable_names = function(trial_list, filename, old_name_header, new_name_header)
  # This function replaces variable names.  Uppercase is also coerced here tO avoId
  # nAMIng aND rETRIevAl isSUes.
  # Arguments:
  #  trial_list
  #  filename
  #  old_name_header
  #  new_name_header
  # Output:
  #  trial_list: input list with new variable names
{
  
  continue_flag = TRUE
  options(stringsAsFactors = FALSE)
  name_change_table = read.delim(filename, na.strings = c("#N/A","."), check.names = FALSE)
  
  # We enforce uppercase as much as possible to avoid issues in 
  # variable names and identifiers.  This seems to come up often in
  # data entry
  
  name_change_table[,new_name_header] = toupper(name_change_table[,new_name_header])
  name_change_table[,old_name_header] = toupper(name_change_table[,old_name_header])
  
  old_names_in_trial_list = list()
  old_names_in_trial_list[["demographics"]] = c()
  old_names_in_trial_list[["index_lesions"]] = c()
  old_names_in_trial_list[["time_series"]] = c()
  # old_names_in_trial_list[["genomics"]] = c()
  # one_deep_list = c("demographics", "time_series", "genomics")
  one_deep_list = c("demographics", "time_series")
  
  
  for (the_patient_id in names(trial_list))
  {
    for (the_one_deep_id in one_deep_list)
    {
      # print(the_one_deep_id)	
      for (the_name in names(trial_list[[the_patient_id]][[the_one_deep_id]]))
      {
        # print(the_name)
        the_name_upper = toupper(the_name)
        if (the_name_upper %in% name_change_table[, old_name_header])
        {
          if (!(the_name %in% old_names_in_trial_list[[the_one_deep_id]]))
          {
            old_names_in_trial_list[[the_one_deep_id]] = c(old_names_in_trial_list[[the_one_deep_id]], the_name, recursive = TRUE)
          }
        }
      }
    }
    the_two_deep_id = "index_lesions"
    for (the_lesion_id in names(trial_list[[the_patient_id]][[the_two_deep_id]]))
    {
      for (the_name in names(trial_list[[the_patient_id]][[the_two_deep_id]][[the_lesion_id]]))
      {
        the_name_upper = toupper(the_name)
        if (the_name_upper %in% name_change_table[, old_name_header])
        {
          if (!(the_name %in% old_names_in_trial_list[[the_two_deep_id]]))
          {
            old_names_in_trial_list[[the_two_deep_id]] = c(old_names_in_trial_list[[the_two_deep_id]], the_name, recursive = TRUE)
          }
        }				
      }
    }
  }
  # browser()
  # Make a key:value type of list to simplify replacement
  old_name_key_new_name_value = list()
  for (the_row_name in rownames(name_change_table))
  {
    the_old_name = name_change_table[the_row_name, old_name_header]
    the_new_name = name_change_table[the_row_name, new_name_header]
    old_name_key_new_name_value[[the_old_name]] = the_new_name
  }
  # Now rename
  for (the_one_deep_id in one_deep_list)
  {
    for (the_old_name in (old_names_in_trial_list[[the_one_deep_id]]))
    {
      the_old_name_upper = toupper(the_old_name)
      for (the_patient_id in names(trial_list))
      {
        # browser()
        the_old_names = (names(trial_list[[the_patient_id]][[the_one_deep_id]]))
        if (the_old_name %in% the_old_names)
        {
          the_new_name = old_name_key_new_name_value[[the_old_name_upper]]
          # browser()
          trial_list[[the_patient_id]][[the_one_deep_id]][[the_new_name]] = trial_list[[the_patient_id]][[the_one_deep_id]][[the_old_name]]
          if (the_new_name != the_old_name)
          {
            trial_list[[the_patient_id]][[the_one_deep_id]][[the_old_name]] = NULL
          }
        }
      }
    }
  }
  the_two_deep_id = "index_lesions"
  for (the_old_name in (old_names_in_trial_list[[the_two_deep_id]]))
  {
    the_old_name_upper = toupper(the_old_name)
    for (the_patient_id in names(trial_list))
    {
      for (the_lesion_id in names(trial_list[[the_patient_id]][[the_two_deep_id]]))
      {
        the_old_names = (names(trial_list[[the_patient_id]][[the_two_deep_id]][[the_lesion_id]]))
        #print(the_old_name_upper %in% the_old_names_uppercase)
        if (the_old_name %in% the_old_names)
        {
          the_new_name = old_name_key_new_name_value[[the_old_name_upper]]
          trial_list[[the_patient_id]][[the_two_deep_id]][[the_lesion_id]][[the_new_name]] = trial_list[[the_patient_id]][[the_two_deep_id]][[the_lesion_id]][[the_old_name]]
          trial_list[[the_patient_id]][[the_two_deep_id]][[the_lesion_id]][[the_old_name]] = NULL
          #print(the_new_name)
        }
      }
    }
    
  }	
  trial_list
}


calculate_lesion_summary = function(trial_list,lesion_measure_1,lesion_measure_2 = NA)
  # This is a function to calculate a summary of lesion data and add
  # it to the VP-level readout.  
  #
  # Arguments:
  #  trial_list
  #  lesion_measure_1: name of lesion measure 1, a diameter in mm
  #  lesion_measure_2: name of lesion measure 2, a diameter in mm.  
  #   If left as NA, it will be assumed that we have 1 measure per lesion.
  # 
  # Outputs:
  #  INDEX_LESION_N: number of index lesions at a time point
#  INDEX_LESION_BSLD: sum of longest diameters prior to the first dosing
#  INDEX_LESION_SLD: sum of longest diameters at a time point
#  INDEX_LESION_AVG_LD: sum of longest diameters at a time point / N
#  INDEX_LESION_MED_LD: median of longest diameters at a time point
#  INDEX_LESION_AVG_VOLUME: total volume of index lesion /N
#  Conditional:
#  INDEX_LESION_BSPD: sum of product of perpendicular diameters prior to the first dosing
#  INDEX_LESION_SPD: sum of product of perpendicular diameters at a time point
#  INDEX_LESION_AVG_PD: sum of product of perpendicular diameters at a time point / N
#  INDEX_LESION_MED_PD: median of product of perpendicular diameters at a time point 

# Note on calculations: lesion sum method based on 
#  Global Biometric Sciences
#  SAS Dataset Specification
#  Project/Study: CA209038
#  Dataset: LETM, LEEF
#  Document Revision: 1.0
#
#  BSXD = sum of lesion measurements prior to the first dosing. 
#  If same tumor lesion measured multiple times, the one with the 
#  latest date prior to or equal to first dosing is used
#
# If there is at least one target lesion reported at a visit 
# (even with missing tumor measurement), which is not found at baseline 
#  SXD= sum(TMRTOT)- TMRTOT from tumor lesions not exist at baseline;
# If there is no missing value of tumor measurement and 
# all baseline lesions are reported:
#  SXD = sum(TMRTOT)
# Else
#  SXDTMP = sum(non missing TMRTOT)
#  If SXDTMP >= 1.2 * previous NSXD  and SXDTMP >= previous NSXD+5
#   SXD=SXDTMP
#  Else
#   SXD=missing
#  End
# End
# If BSXD is not missing, NSXD is the smallest non-missing SPD from all previous visits,
# Otherwise NSXD is missing
#
#  Note effectively that lesions will full (no missing) data only will be considered in
#  the calculations at the patient level here.  At some point, we may want to add in 
#  "lesion split" events as these are included in the GBS CA209038 specification
#  
{
  library('gtools')
  # If these fields already exist for a given patient, we 
  # will simply update/overwrite them.
  for (the_patient_id in names(trial_list))
  {
    # first we compile the lesion data to get a matrix for each time [lesion, m1, m2]
    compiled_lesion_data = reorganize_lesion_data_by_time(trial_list[[the_patient_id]], lesion_measure_1, lesion_measure_2)
    
    # Now reorganize this so we fill in any missing values for constant-dimension 
    # matrices at each time point
    compiled_lesion_data = fill_measure_matrices(compiled_lesion_data)
    
    
    trial_list[[the_patient_id]][["time_series"]][["ONTRT"]]=list()
    trial_list[[the_patient_id]][["time_series"]][["INDEX_LESION_N"]]=list()
    if (!(is.na(lesion_measure_2)))
    {
      trial_list[[the_patient_id]][["time_series"]][["INDEX_LESION_SPD"]]=list()
      trial_list[[the_patient_id]][["time_series"]][["INDEX_LESION_AVG_PD"]]=list()
      trial_list[[the_patient_id]][["time_series"]][["INDEX_LESION_MED_PD"]]=list()
    }
    trial_list[[the_patient_id]][["time_series"]][["INDEX_LESION_SLD"]]=list()
    trial_list[[the_patient_id]][["time_series"]][["INDEX_LESION_AVG_LD"]]=list()
    trial_list[[the_patient_id]][["time_series"]][["INDEX_LESION_MED_LD"]]=list()
    trial_list[[the_patient_id]][["time_series"]][["INDEX_LESION_AVG_VOLUME"]]=list()
    the_times = names(compiled_lesion_data)
    if (!(is.null(the_times)))
    {
      the_times = mixedsort(the_times)
      
      the_lesion_ids = compiled_lesion_data[[the_times[1]]][,1]
      n_lesions = length(the_lesion_ids)
      the_baseline_time_strings = c()
      for (the_time_string in the_times)
      {
        if (check_and_convert_numeric(the_time_string) <= 0)
        {
          the_baseline_time_strings = c(the_baseline_time_strings, the_time_string)
        }
      }
      the_baseline_time_strings = mixedsort(the_baseline_time_strings)
      baseline_measures = data.frame(matrix(data = NA, n_lesions, 3))
      baseline_measures[,1] = the_lesion_ids
      
      for (the_row_number in seq_along(baseline_measures[,1]))
      {
        the_value_1 = NA
        the_value_2 = NA
        # First we identify the baseline measure
        # and enter those that are not missing important values
        for (the_time_string in the_baseline_time_strings)
        {
          the_value_1 = compiled_lesion_data[[the_time_string]][the_row_number,2]
          if (!(is.na(lesion_measure_2)))
          {
            the_value_2 = compiled_lesion_data[[the_time_string]][the_row_number,3]
            # We add a check, if in this study we are looking for 2 measures but only 1 is valid
            # we will invalidate/ignore this lesion measure
            if (!(is.numeric(the_value_2)) || is.na(the_value_2))
            {
              the_value_1 = NA
              the_value_2 = NA
            }
          }
          if ( ((!(is.na(the_value_1))) && (is.na(lesion_measure_2))) || ((!(is.na(the_value_1))) && (!(is.na(the_value_2)))) )
          {
            baseline_measures[the_row_number,2] = the_value_1
            baseline_measures[the_row_number,3] = the_value_2
          }
          #print(the_patient_id)
          #print(baseline_measures)
        }
      }
      
      lesion_measures_missing_at_baseline = c()
      for (the_row_number in seq_along(baseline_measures[,1])) 
      {
        value_1 = baseline_measures[the_row_number,2]
        value_2 = baseline_measures[the_row_number,3]
        if (is.na(value_1) && is.na(value_2))
        {
          lesion_measures_missing_at_baseline = c(lesion_measures_missing_at_baseline, baseline_measures[the_row_number,1])
        }
      }
      lesion_id_present_at_baseline = setdiff(the_lesion_ids,lesion_measures_missing_at_baseline)
      lesion_row_numbers_present_at_baseline = c()
      for (the_row_number in seq_along(baseline_measures[,1]))
      {
        the_lesion_id = baseline_measures[the_row_number, 1]
        if (the_lesion_id %in% lesion_id_present_at_baseline)
        {
          lesion_row_numbers_present_at_baseline = c(lesion_row_numbers_present_at_baseline, the_row_number)
        }
      }
      
      # Add the baseline measures for the patient
      trial_list[[the_patient_id]][["baseline_lesions"]] = list()
      for (the_row_number in seq_along(baseline_measures[,1]))
      {
        the_lesion_id = baseline_measures[the_row_number, 1]
        trial_list[[the_patient_id]][["baseline_lesions"]][[the_lesion_id]] = list()
        trial_list[[the_patient_id]][["baseline_lesions"]][[the_lesion_id]][["LESIONSIZE1_MM"]] = baseline_measures[the_row_number, 2]
        
        
        if (!(is.na(lesion_measure_2)))
        {
          trial_list[[the_patient_id]][["baseline_lesions"]][[the_lesion_id]][["LESIONSIZE2_MM"]] = baseline_measures[the_row_number, 3]
        }
      }
      
      # We only consider lesions that are not missing baseline measures.
      # Split lesions are different, but we ignore this for now.
      # Will handle in a different update
      nonmissing_baseline_matrix = baseline_measures[lesion_row_numbers_present_at_baseline, ]
      bnlesions = length(nonmissing_baseline_matrix[,2])
      bdia_prod = nonmissing_baseline_matrix[,2] * nonmissing_baseline_matrix[,3]
      
      if (!(is.na(lesion_measure_2)))
      {
        bl_dia = apply(nonmissing_baseline_matrix[,2:3], 1, max)
      } else {
        bl_dia = nonmissing_baseline_matrix[,2]
      }
      baseline_sld = sum(bl_dia)
      sld_reference = c(baseline_sld)
      
      # Now we have baseline measures.  Now we can evaluate on-therapy measures.
      cur_sld = c() 
      cur_bsld = c()
      cur_asld = c()
      cur_spd = c()
      cur_bspd = c()
      cur_aspd = c()
      
      # Also create a matrix of lymph node flags
      lesion_ids_to_check = nonmissing_baseline_matrix[,1]
      node_flags = matrix(0,length(lesion_ids_to_check),1)
      if (length(lesion_ids_to_check)>0)
      {
        
        for (node_counter in seq(1,length(node_flags)))
        {
          node_flags[node_counter] = trial_list[[the_patient_id]][["index_lesions"]][[lesion_ids_to_check[node_counter]]][["NODE_FLAG"]]
        }
      }
      
      
      for (the_time_string in the_times)
      {
        current_lesion_matrix = compiled_lesion_data[[the_time_string]][lesion_row_numbers_present_at_baseline, ]
        nlesions = NA
        dia_prod = NA
        l_dia = NA
        max_node_l_dia = NA
        nonnode_l_dia = NA
        
        # First we filter the time data and set to NA non-numeric values
        # or, on older trials, if we had 2 measures but both are not present
        ## SCAN THROUGH ROWS, SET NAs ##
        for (the_row_index in seq_along(current_lesion_matrix[,1]))
        {
          the_value_1 = current_lesion_matrix[the_row_index,2]
          the_value_2 = current_lesion_matrix[the_row_index,3]
          
          if (!(is.na(lesion_measure_2)))
          {
            if ( (!(is.numeric(the_value_1))) || (!(is.numeric(the_value_2))) || is.na(the_value_1) || is.na(the_value_2))
            {
              current_lesion_matrix[the_row_index,2] = NA
              current_lesion_matrix[the_row_index,3] = NA
            }
          } else {
            # If lesion_measure_2 is NA, then we just need to check the first value
            if(!(is.numeric(the_value_1)) || is.na(the_value_1)) 	
            {
              current_lesion_matrix[the_row_index,2] = NA
            }					
          }
        }
        
        # Get the summary stats for the current time point
        if (sum(is.na(current_lesion_matrix[,2])) == 0)
        {
          nlesions = nrow(current_lesion_matrix)
          dia_prod = current_lesion_matrix[,2] * current_lesion_matrix[,3]
          
          
          if (!(is.na(lesion_measure_2)))
          {
            l_dia = apply(current_lesion_matrix[,2:3], 1, max)
          } else {
            l_dia = current_lesion_matrix[,2]
          }
          if (length(l_dia) > 0)
          {
            max_node_l_dia = max(node_flags*l_dia)
            nonnode_l_dia = ((1-node_flags)*l_dia)
          }
        } else 
        {
          # Get the rows with no missing values
          no_missing = which( !(is.na(current_lesion_matrix[,2])) )
          no_missing_lesion_current_matrix = current_lesion_matrix[no_missing,]
          nlesions_tmp = nrow(no_missing_lesion_current_matrix)
          dia_prod_tmp = no_missing_lesion_current_matrix[,2] * no_missing_lesion_current_matrix[,3]
          if (!(is.na(lesion_measure_2)))
          {
            l_dia_tmp = apply(no_missing_lesion_current_matrix[,2:3], 1, max)
          } else {
            l_dia_tmp = no_missing_lesion_current_matrix[,2]
          }
          
          
          
          NSLD = min(sld_reference)
          sld_temp = sum(l_dia_tmp)
          if (!(is.na(NSLD)))
          {
            # These are the GBS rules from 209038 (THIS IF IS COMMENTED BY YOUGAN FOR 184022)
            # Note this may cause issues.
            # Note this result in measures dropping out
            # if extra measures are taken prior to baseline
            
            #browser()
            #print(NSLD)
            #print(the_patient_id)
            #print(l_dia)
            #print(l_dia_tmp)
            #print(sld_temp)
            #print(lesion_measure_2)
            #print(current_lesion_matrix)
            #print(no_missing_lesion_current_matrix)
            if ((sld_temp >= (1.2 * NSLD)) && (sld_temp > (NSLD + 5)))
            {
              nlesions = nlesions_tmp
              dia_prod = dia_prod_tmp
              l_dia = l_dia_tmp
              nonnode_l_dia = ((1-node_flags[no_missing])*l_dia_tmp)
            }
            
          }	
          
        }
        if(!(is.na(sum(l_dia))))
        {
          sld_reference = c(sld_reference, sum(l_dia)) 
        }
        
        trial_list[[the_patient_id]][["time_series"]][["INDEX_LESION_N"]][[the_time_string]] = nlesions
        
        if (!(is.na(lesion_measure_2)))
        {
          
          trial_list[[the_patient_id]][["time_series"]][["INDEX_LESION_SPD"]][[the_time_string]] = sum(dia_prod)
          trial_list[[the_patient_id]][["time_series"]][["INDEX_LESION_AVG_PD"]][[the_time_string]] = sum(dia_prod)/nlesions
          trial_list[[the_patient_id]][["time_series"]][["INDEX_LESION_MED_PD"]][[the_time_string]] = median(dia_prod)
        }		
        
        # Yougan: Note that if the pretreatment lesions are measured at seperate time points, this will result in NA assignment
        # for l_dia, nlesions etc. This might be on purpuse, but I am changing it. For example if a patient got index lesion A B
        # measured at day -20, and C, D measured at day -1, the algorithm still get the baseline measures fine. But, here it calculates
        # the specific time point behavior, and at day -20 and -1 l_dia, nlesions will be assigned as NA
        
        if (as.numeric(the_time_string) <= 0)
        {
          trial_list[[the_patient_id]][["time_series"]][["INDEX_LESION_N"]][[the_time_string]] = bnlesions
          nlesions = bnlesions
          l_dia = bl_dia
          if (length(l_dia) > 0)
          {
            max_node_l_dia = max(node_flags*l_dia)
            nonnode_l_dia = ((1-node_flags)*l_dia)
          }
        }
        
        
        trial_list[[the_patient_id]][["time_series"]][["INDEX_LESION_SLD"]][[the_time_string]] = sum(l_dia)
        trial_list[[the_patient_id]][["time_series"]][["INDEX_LESION_AVG_LD"]][[the_time_string]] = sum(l_dia)/nlesions
        trial_list[[the_patient_id]][["time_series"]][["INDEX_LESION_AVG_VOLUME"]][[the_time_string]] = (0.2291*sum((l_dia/2)^3))/nlesions
        trial_list[[the_patient_id]][["time_series"]][["INDEX_LESION_MED_LD"]][[the_time_string]] = median(l_dia)
        trial_list[[the_patient_id]][["time_series"]][["INDEX_LESION_MAXLN"]][[the_time_string]] = max_node_l_dia
        trial_list[[the_patient_id]][["time_series"]][["INDEX_LESION_NLNSLD"]][[the_time_string]] = sum(nonnode_l_dia)
        
        if (as.numeric(the_time_string) > 0)
        {
          cur_sld = c(cur_sld, sum(l_dia)) 
          cur_bsld = c(cur_bsld, sum(bl_dia))
          cur_spd = c(cur_spd, sum(dia_prod))
          cur_bspd = c(cur_bspd, sum(bdia_prod))
        }
        cur_asld = c(cur_asld, sum(l_dia))
        cur_aspd = c(cur_aspd, sum(dia_prod))
        
        OFFTIME = trial_list[[the_patient_id]][["demographics"]][["OFFTRTIME"]]
        if (!is.na(OFFTIME)){
          if (as.numeric(the_time_string) <= as.numeric(OFFTIME))
          {
            trial_list[[the_patient_id]][["time_series"]][["ONTRT"]][[the_time_string]] = 1
          }else
          {
            trial_list[[the_patient_id]][["time_series"]][["ONTRT"]][[the_time_string]] = 0
          }
        }else{
          trial_list[[the_patient_id]][["time_series"]][["ONTRT"]][[the_time_string]] = NA
        }
        
        
      }
      
      sld_ratio = (cur_sld)/cur_bsld
      spd_ratio = (cur_spd)/cur_bspd
      asld_ratio = (cur_asld)/cur_bsld
      aspd_ratio = (cur_aspd)/cur_bspd			
      if (length(na.omit(sld_ratio)) > 0)
      {
        b_sld_ratio = min(na.omit(sld_ratio))-1
      } else {
        b_sld_ratio = NA
        #sld_ratio = NA
      }
      if (length(na.omit(spd_ratio)) > 0)
      {
        b_spd_ratio = min(na.omit(spd_ratio))-1
      } else {
        b_spd_ratio = NA
        #spd_ratio = NA
      }
      
      
      # trial_list[[the_patient_id]][["demographics"]][["INDEX_LESION_SLD_BOR"]][[the_time_string]] = sld_score
      # trial_list[[the_patient_id]][["demographics"]][["INDEX_LESION_SPD_BOR"]][[the_time_string]] = spd_score
      for (the_i in seq_along(the_times))
      {
        the_time_string = the_times[the_i]
        trial_list[[the_patient_id]][["time_series"]][["INDEX_LESION_SLD_RELCH"]][[the_time_string]] = asld_ratio[the_i]-1
        if (!(is.na(lesion_measure_2)))
        {
          trial_list[[the_patient_id]][["time_series"]][["INDEX_LESION_SPD_RELCH"]][[the_time_string]] = aspd_ratio[the_i]-1
        }
      }
      trial_list[[the_patient_id]][["demographics"]][["INDEX_LESION_BN"]] = bnlesions
      trial_list[[the_patient_id]][["demographics"]][["INDEX_LESION_BSLD"]] = sum(bl_dia)		
      trial_list[[the_patient_id]][["demographics"]][["INDEX_LESION_BAVG_LD"]] = sum(bl_dia)/bnlesions
      trial_list[[the_patient_id]][["demographics"]][["INDEX_LESION_BMED_LD"]] = median(bl_dia)
      trial_list[[the_patient_id]][["demographics"]][["INDEX_LESION_SLD_RELCHMIN"]] = b_sld_ratio
      if (!(is.na(lesion_measure_2)))
      {	
        trial_list[[the_patient_id]][["demographics"]][["INDEX_LESION_BSPD"]] = sum(bdia_prod)
        trial_list[[the_patient_id]][["demographics"]][["INDEX_LESION_BAVG_PD"]] = sum(bdia_prod)/bnlesions
        trial_list[[the_patient_id]][["demographics"]][["INDEX_LESION_BMED_PD"]] = median(bdia_prod)
        trial_list[[the_patient_id]][["demographics"]][["INDEX_LESION_SPD_RELCHMIN"]] = b_spd_ratio	
      }
      
    }
  }
  trial_list
}


reorganize_lesion_data_by_time = function(trial_list_one_patient, lesion_measure_1, lesion_measure_2)
  # This function regroups lesion measure data so time is the higher level key
  # and organizes the measures in a data frame
  # to facilitate subsequent calculation of derived measures at each time point.
  #
  # Arguments
  #  trial_list_one_patient
  #  lesion_measure_1
  #  lesion_measure_2: may be NA if only 1 measure was taken
  #
  # Output
  #  compiled_lesion_data
#
{
  lesion_source_data = trial_list_one_patient[['index_lesions']]
  compiled_lesion_data = list()
  for (the_index_lesion_id in names(lesion_source_data))
  {
    # First we make sure we have both measures for a lesion at each time point
    # Didn't see a shorter way to code this in the base R package
    vector_a = names(lesion_source_data[[the_index_lesion_id]][[lesion_measure_1]])
    if (!(is.na(lesion_measure_2)))
    {
      vector_b = names(lesion_source_data[[the_index_lesion_id]][[lesion_measure_2]])
    } else {
      vector_b = vector_a
    }
    vector_i = intersect(vector_a,vector_b)
    if ((length(vector_i) == length(vector_a)) && (length(vector_i) == length(vector_b)))
    {
      the_time_names = names(lesion_source_data[[the_index_lesion_id]][[lesion_measure_1]])
      if (NA %in% the_time_names)
      {
        # It's possible the time value is missing, in which case we ignore that time point
        the_time_names = setdiff(the_time_names, c(NA))
      }
      
      # We need to perform checks to make sure we either 
      # have values or missing values entered for a lesion.
      # E.g., if you tell me a lesion measures 5, that's OK, 
      # or if you tell me it measures NA that's OK too,
      # but if you tell me it measures "red" I need to ignore this.
      ignore_lesion = FALSE
      if (length(the_time_names) < 1)
      {
        ignore_lesion = TRUE
      }
      
      if (!(ignore_lesion))
      {
        for (the_time in the_time_names)
        {
          value_1 = lesion_source_data[[the_index_lesion_id]][[lesion_measure_1]][[the_time]]
          
          if (!(is.na(lesion_measure_2)))
          {
            value_2 = lesion_source_data[[the_index_lesion_id]][[lesion_measure_2]][[the_time]]
          } else {
            value_2 = NA
          }
          if (((is.numeric(value_1) || is.na(value_1)) && (is.numeric(value_2) || is.na(value_2))))
          {
            if (!(the_time %in% names(compiled_lesion_data)))
            {
              compiled_lesion_data[[the_time]] = data.frame(matrix(nrow = 0,ncol=3))
            }
            row_to_add = data.frame(the_index_lesion_id,value_1,value_2)
            compiled_lesion_data[[the_time]] = rbind(compiled_lesion_data[[the_time]], row_to_add)
          }
        }
      }
    }
  }
  compiled_lesion_data
}


fill_measure_matrices = function(compiled_lesion_data)
  # This function fills out lesion measure data frames across
  # all time points so that it is easier to reference which measures
  # are not taken at a given time point.  For the purposes of
  # calculation, these need to be noted and indicated as missing
  # values.
  #
  # Arguments:
  #  compiled_lesion_data
  #
  # Outputs: 
  #  reorganized_lesion_data
#
# Now reorganize this so we have for each time [lesion, m1, m2] with all lesions in the header
{
  lesion_id_vector = c()
  reorganized_lesion_data = list()
  for (the_time in names(compiled_lesion_data))
  {
    the_matrix = compiled_lesion_data[[the_time]]
    current_lesions = the_matrix[,1]
    lesion_id_vector = union(lesion_id_vector, current_lesions)
  }
  lesion_id_vector = sort(lesion_id_vector)
  number_lesions = length(lesion_id_vector)
  for (the_time in names(compiled_lesion_data))
  {
    the_old_matrix = compiled_lesion_data[[the_time]]
    the_new_matrix = data.frame(matrix(data = NA, nrow = number_lesions, ncol=3))
    the_new_matrix[,1] = lesion_id_vector
    for (the_old_row in seq_along(the_old_matrix[,1]))
    {
      the_lesion_id = the_old_matrix[the_old_row,1]
      the_new_row = which(lesion_id_vector==the_lesion_id)
      the_new_row = the_new_row[1]
      the_new_matrix[the_new_row,] = the_old_matrix[the_old_row,]
    }
    
    reorganized_lesion_data[[the_time]] = the_new_matrix
  }
  reorganized_lesion_data
}


get_time_series_matrix = function(one_measure_time_series)
  # This function gets data for 1 measure and
  # returns a dataframe organized by time point.
  #
  # Arguments
  #  one_measure_time_series
  #
  # Output
  #  one_time_series_measure_dataframe
  #
{
  library('gtools')
  the_times = names(one_measure_time_series)
  the_times = mixedsort(the_times)
  one_time_series_measure_dataframe = data.frame(matrix(data = NA, nrow = length(the_times),ncol = 3))
  i = 1
  for (the_time_name in the_times)
  {
    one_time_series_measure_dataframe[i,1] = the_time_name
    one_time_series_measure_dataframe[i,2] = as.numeric(the_time_name)
    one_time_series_measure_dataframe[i,3] = as.numeric(one_measure_time_series[[the_time_name]])
    i = i + 1
  }
  one_time_series_measure_dataframe
}


calculate_ratios = function(trial_data, measure_names = c(), transform = "BRATIO")
  # This function identifies the last measure prior to the initiation 
  # of therapy and then calculates the ratio for each time point
  #
  # Arguments:
  #  trial_data
  #  measure_names: a vector of variable names in "time_series" to get
  #                 baseline ratios for
  #  transform: "BRATIO" "BRATIOOFLOG2" "LOG2BRATIOOFEXP2" "BRATIOOFEXP2"  "DIFFERENCE" (DIFFERENCE added by Lu)
{
  for (the_patient_id in names(trial_list))
  {
    #print(the_patient_id)
    # We need to scan through each measure
    for (the_measure_name in measure_names)
    {
      #print(the_measure_name)
      # We can use the lesion measure function for this purpose
      if (the_measure_name %in% names(trial_list[[the_patient_id]][["time_series"]]))
      {
        the_measure_list = (trial_list[[the_patient_id]][["time_series"]][[the_measure_name]])
        compiled_measure_data = get_time_series_matrix(the_measure_list)		
        
        # Get the index to the baseline measure
        the_baseline_row_index = max(which(compiled_measure_data[,2]<=0))
        the_baseline_value = compiled_measure_data[the_baseline_row_index,3]
        #if (!is.na(the_baseline_value))
        #{
        # Apply any transforms
        if (transform == "BRATIOOFLOG2") 
        {
          compiled_measure_data[,3] = log2(compiled_measure_data[,3])/log2(the_baseline_value)
        } else if(transform == "BRATIOOFEXP2") 
        {
          compiled_measure_data[,3] = 2^(compiled_measure_data[,3])/2^(the_baseline_value)
        } else if(transform == "LOG2BRATIOOFEXP2") 
        {
          compiled_measure_data[,3] = log2(2^(compiled_measure_data[,3])/2^(the_baseline_value))
        } else if(transform == "BRATIO")
        {
          compiled_measure_data[,3] = compiled_measure_data[,3]/the_baseline_value
        } else
        {
          compiled_measure_data[,3] = compiled_measure_data[,3] - the_baseline_value
        }
        if (nrow(compiled_measure_data)>0){
          compiled_measure_data[(compiled_measure_data[,3]==Inf) & (!is.na(compiled_measure_data[,3])),3] = NA
        }
        the_new_ratio_name = paste(the_measure_name, transform, sep = "_")
        trial_list[[the_patient_id]][["time_series"]][[the_new_ratio_name]] = list()
        for (the_index in seq_along(compiled_measure_data[,1]))
        {
          the_time_point = compiled_measure_data[the_index,1]
          the_value = compiled_measure_data[the_index,3]
          trial_list[[the_patient_id]][["time_series"]][[the_new_ratio_name]][[the_time_point]] = the_value
        }
        #}
      }
    }
  }
  trial_list
}

calculate_demographic_time = function(trial_list, newtime_name = "OFFTRTIME", date_name = "DATE_DISCONTINUE")
  # This function calculates a time output given a date variable, assumed to be in the
  # demographic patient level data as opposed to time series measures, and adds that output
  # as an additional variable.  This is intended to facilitate making important
  # comparisons within the data such interpreting lesion sizes relative to
  # when a patient stops therapy.
  #
  # Arguments:
  #  trial_list
  #  newtime_name:    the time variable the results of the calculations will be stored to.
  #  date_name:       the name of the later of the date variables, for example the
  #                   off treatment date
#
# Returns:
#  trial_list
#
{
  firstdate_name = "first_dose_date"	
  for (the_patient_id in names(trial_list))
  {
    pass1_flag = FALSE
    pass2_flag = FALSE
    if (firstdate_name %in% names(trial_list[[the_patient_id]]))
    {
      if (class(trial_list[[the_patient_id]][[firstdate_name]]) == "Date")
      {
        pass1_flag = TRUE
      }
    }
    if (date_name %in% names(trial_list[[the_patient_id]][["demographics"]]))
    {
      if (length(trial_list[[the_patient_id]][["demographics"]][[date_name]]) > 0)
      {
        if (!(is.na(trial_list[[the_patient_id]][["demographics"]][[date_name]])))
        {
          date2 = convert_to_date(trial_list[[the_patient_id]][["demographics"]][[date_name]])
          if (class(date2) == "Date")
          {
            pass2_flag = TRUE
          }
        }
      } else {
        pass2_flag = FALSE
        trial_list[[the_patient_id]][["demographics"]][[date_name]] = NA
      }
    }
    if (pass1_flag && pass2_flag)
    {
      # Since this is a demographic variable rather than a reserved
      # time variable it is just kept as numeric
      trial_list[[the_patient_id]][["demographics"]][[newtime_name]] = date2-trial_list[[the_patient_id]][[firstdate_name]]
    }
    else
    {
      trial_list[[the_patient_id]][["demographics"]][[newtime_name]] = NA
    }
  }
  trial_list
}

align_timeseries_data = function(trial_list, var_name_vector, target_time_vector, bin_edge_matrix, stop_time_name = NULL, interpolate_option="NEAREST", drop_missing = FALSE)
  # This is a function to take time series variables and align them to a desired
  # set of output times.  This is intended to help develop marginal distributions
  #
  # Arguments
  #  trial_list
  #  var_name_vector:    A vector of variable names to be aligned, should be time_series variables
  #  target_time_vector: A vector of the target times for the output
  #  bin_edge_matrix:    A vector of the in edges to use when assigning output values for the target times.
  #                      time values >= the lower bin edge (n) and < the upper bin edge (n+1) are considered
  #                      in the evaluation of the value assigned for the (n) spot in the target_time_vector
  #  stop_time_name:     Variable with time to stop the measurements at, for example the off treatment time.
#                      measures after this time are ignored.
#                      We will look under "demographic" variables.
#  interpolate_option: Method for assigning the data to the target time point.  supported options include:
#                      "NEAREST": the value from the nearest valid time point in the bin is used, whether it
#                                 occurs before or after
#                      "LAST": the value from the last time point is used
#  drop_missing:       (boolean) Whether to drop target times with values that are missing
#  
# Returns
#  trial_list 
{
  # Perform initial checks
  continue_flag = TRUE
  #marker_counter = 0
  marker_with_at_least_one = 0
  n_patients = length(names(trial_list)) # Lu: remove one 'names'
  for (marker_index in seq_along(var_name_vector)) # Lu: changed from 'marker_counter' to 'marker_index'
  {
    marker_counter = 0 # Lu: moved to here
    the_marker_name = var_name_vector[marker_index] # Lu: changed from 'marker_counter' to 'marker_index'
    for (the_patient_id in names(trial_list))
    {
      if (the_marker_name %in% union(names(trial_list[[the_patient_id]][["time_series"]]),names(trial_list[[the_patient_id]][["demographics"]]))) # Lu: add demographic names
      {
        marker_counter = marker_counter + 1
      }
    }
    if (marker_counter < n_patients)
    {
      print(paste("Only found data for ",as.character(marker_counter)," of ",as.character(n_patients)," patients for ",the_marker_name,".", sep = ""))
    }
    if (marker_counter > 0)
    {
      marker_with_at_least_one = marker_with_at_least_one+1
    }
  }
  if (marker_with_at_least_one<1)
  {
    print(paste("No marker data, exiting..."))
    continue_flag = FALSE
  }
  
  stop_counter = 0	
  if (!(is.null(stop_time_name)))
  {
    for (the_patient_id in names(trial_list))
    {
      if (stop_time_name %in% names(trial_list[[the_patient_id]][["demographics"]]))
      {
        stop_counter = stop_counter + 1
      }
    }
    if (stop_counter < n_patients)
    {
      print(paste("Only found stop data for ",as.character(stop_counter)," of ",as.character(n_patients)," patients.", sep = ""))
    }
    if (stop_counter < 1)
    {
      print("No stop data as specified, exiting...")
      continue_flag = FALSE
    }		
  }	
  
  interpolate_option = check_arguments(interpolate_option, c("NEAREST","LAST"), force = FALSE)
  if (is.null(interpolate_option)) 
  {
    continue_flag = FALSE
    print("Please specify a valid interpolate_option argument, exiting...")
  }	
  
  
  n_time_points = length(target_time_vector)
  bindim = dim(bin_edge_matrix)
  n_bin_edges = bindim[1]
  fail_t_points = 0
  if (n_time_points != (n_bin_edges))
  {
    continue_flag = FALSE
    print("The bin_edge_matrix should have same number of rows as the target_time_vector has entries, exiting...")
  } else {
    for (time_counter in seq_along(target_time_vector))
    {
      if ((target_time_vector[time_counter]< bin_edge_matrix[time_counter,1]) | (target_time_vector[time_counter]>= bin_edge_matrix[time_counter,2]))
      {
        continue_flag = FALSE
        fail_t_points = fail_t_points + 1
      }
    }
  }
  if 	(fail_t_points > 0)
  {
    print("Each element of the target_time_vector should be greater than or equal to the corresponding first row entry in the bin_edge_matrix or and less than the second entry, exiting...")
  }
  
  
  if (continue_flag)
  {
    for (the_patient_id in names(trial_list))
    {
      for (the_marker_name in var_name_vector)
      {
        if (the_marker_name %in% names(trial_list[[the_patient_id]][["time_series"]]))
          # if (the_marker_name %in% union(names(trial_list[[the_patient_id]][["time_series"]]),names(trial_list[[the_patient_id]][["demographics"]])))
        {
          #print(the_patient_id)
          #print(the_marker_name)
          #trial_list[[the_patient_id]][["time_series"]][[the_marker_name]]
          cur_data = get_time_series_matrix(trial_list[[the_patient_id]][["time_series"]][[the_marker_name]])
          # Filter out data not within the cutoff time
          if (!(is.null(stop_time_name)))
          {
            if (stop_time_name %in% names(trial_list[[the_patient_id]][["demographics"]]))
            {
              stop_time = trial_list[[the_patient_id]][["demographics"]][[stop_time_name]]
              if (is.numeric(stop_time))
              {
                cur_data=cur_data[which(cur_data[,2]<=stop_time),]
              }
            }
          }
          interpolated_values = list()
          for (target_time_counter in seq_along(target_time_vector))
          {
            t0 = bin_edge_matrix[target_time_counter,1]
            t1 = bin_edge_matrix[target_time_counter,2]
            tc = target_time_vector[target_time_counter]
            point_data = cur_data[which((cur_data[,2]>=t0)&(cur_data[,2]<t1)),]
            if (interpolate_option=="LAST")
            {
              cur_indices = which(point_data[,2]<=tc)
              if (length(cur_indices) > 0)
              {
                cur_value = point_data[cur_indices[length(cur_indices)],3]
              } else {
                cur_value = NA
              }
            } else
            {
              cur_index = which.min(abs(point_data[,2]-tc))
              if (length(cur_index) > 0)
              {
                cur_value = point_data[cur_index,3]
              } else {
                cur_value = NA
              }
            }
            if ((!(is.na(cur_value))) | (!(drop_missing)))
            {
              interpolated_values[[as.character(tc)]] = cur_value
            }
            
          }
          
          # We could add in some function here to filter the results
          # That are not full.  But keep all results for now.
          # Now overwrite the data with the interpolation results
          
          trial_list[[the_patient_id]][["time_series"]][[the_marker_name]] = interpolated_values
        }
        
      }
    }
  }	
  trial_list
}


read.tdelim = function(filename, na.strings = c("#N/A","."), check.names = FALSE) 
  # This is just essentially a wrapper for read.delim
  # but is needed to read transposed data
{ 
  x <- read.table(filename, na.strings = na.strings, check.names = check.names, sep = "\t", header = FALSE)
  x <- as.data.frame(t(x))
  colnames(x) = x[1,]
  x = x[-1,]
  x
}


calculate_summary = function(data_frame, condition_column, data_column_vector = c(), comparison_vector = c(), time_column = "TIME")
  # STILL IN DEVELOPMENT
  # Take a data_frame and get summary statistics from patients (rows)
  # that fall within pre-defined criteria at each time point
  # Arguments:
  #  data_frame: a data.frame object
  #  condition_column: the column name that contains the
  #                    conditions that it is desired to 
  #                    extract information from.
  #  data_column_vector: a vector of columns to calculate 
  #                      summary data for  
  #  comparison_vector: a vector of reference values
#         we look for the row value in
#         the condition_column to equal one of the elements
#         If we want to seach for values
#         that fall within a range we need to pre-screen
#         and assign to a bucket before calling
#         this function
#  type: valid values are "EQ", "BET"
#  time_column: colname for the time column in the data_frame  
#
# TODO: think about binning time rather than requiring equality
{
  
  # Run checks on the specified variable, make a graceful exit if
  # it looks like there are issues.
  continue_flag = TRUE
  condition_column = check_arguments(condition_column, colnames(data_frame), force = FALSE)
  if (is.null(condition_column)) 
  {
    continue_flag = FALSE
    print("Please specify a condition_column argument, exiting...")
  } else if(!(condition_column %in% colnames(data_frame))) {
    continue_flag = FALSE
    print("Please specify a condition column that is in the column names of the data frame.  Exiting...")
    # If something is specified, we will test it to make sure it makes sense.
    # If not, we use all of what is available
  } else if (length(comparison_vector) > 0) {
    comparison_vector = intersect(comparison_vector,unique(data_frame[,condition_column]))
    if (length(comparison_vector) == 0) 
    {
      print("Please specify comparison conditions in comparison_vector that are elements of condition_column, exiting...")
      continue_flag = FALSE
    }
  } else {
    comparison_vector = unique(data_frame[,condition_column])
  }
  
  
  time_column = check_arguments(time_column, colnames(data_frame), force = FALSE)
  if (is.null(time_column)) 
  {
    stop("Please specify a time_column argument that is in the colnames of the data frame.  Existing...")
    continue_flag = FALSE
  }  
  
  
  data_frame_return = data_frame
  shape = dim(data_frame)
  nrows = shape[1]
  ncols = shape[2]
  
  # We will calculate median, min, max, mean, SD for each set of
  # data at a time point.
  if (continue_flag)
  {
    data_column_vector = intersect(data_column_vector,colnames(data_frame))
    data_column_vector = setdiff(data_column_vector, c(time_column,condition_column))
    # Check remaining arguments
    if (length(data_column_vector) < 1)
    {
      # This is not implemented because we may be carrying additional non-numeric
      # data like the patient ID
      continue_flag = FALSE
      print("Please specify column names ")
    } else {
      # As an additional check:
      data_column_vector = setdiff(data_column_vector, c(time_column,condition_column))
      data_column_vector = intersect(data_column_vector, colnames(data_frame))
      if (length(data_column_vector) < 1) {continue_flag = FALSE}
    }
  }
  
  if (continue_flag)
  {
    # For every var to summarize, we have 5 new columns:
    # median, 0.25 quartile, 0.75 quartile, mean, sd, n
    column_increment = 6;
    # plus condition, time, and N
    ncols_new = length(data_column_vector)*column_increment + 2
    data_frame_return = data.frame(matrix(NA,0,ncols_new))
    all_new_names = c(condition_column,time_column)
    final_var_name_list = list()
    for (the_root in data_column_vector)
    {
      median_name = paste(the_root,"MED",sep = "_")
      qr1_name = paste(the_root,"25QR",sep = "_")
      qr2_name = paste(the_root,"75QR",sep = "_")
      mean_name = paste(the_root,"MEAN",sep = "_")
      sd_name = paste(the_root,"SD",sep = "_")
      n_name = paste(the_root,"N",sep = "_")
      final_var_name_list[[the_root]] = list()
      final_var_name_list[[the_root]][["MED"]] = median_name
      final_var_name_list[[the_root]][["25QR"]] = qr1_name
      final_var_name_list[[the_root]][["75QR"]] = qr2_name
      final_var_name_list[[the_root]][["MEAN"]] = mean_name
      final_var_name_list[[the_root]][["SD"]] = sd_name
      final_var_name_list[[the_root]][["N"]] = n_name
      all_new_names = c(all_new_names, median_name, qr1_name, qr2_name, mean_name, sd_name, n_name, recursive = TRUE)
    } 
    colnames(data_frame_return) = all_new_names
    # The number of rows will vary based on the time points
    # We are grouping by time point, so scan first
    # within each condition group then by data at
    # a time point
    for (current_condition in comparison_vector)
    {
      # May want to add some binning here
      the_time_points = unique(data_frame[which(data_frame[, condition_column] == current_condition), time_column])
      the_time_points = sort(the_time_points)
      for (the_time_point in the_time_points)
      {
        #data_frame[which(data_frame[, condition_column] == current_condition), time_column])
        the_sub_data_frame = data_frame[intersect(which(data_frame[, condition_column]== current_condition), which(data_frame[, time_column] == the_time_point)),]
        # CONTINUE HERE
        the_new_row = data.frame(matrix(NA, 1, ncols_new))
        colnames(the_new_row) = all_new_names
        for (the_var_name in names(final_var_name_list))
        {
          # Check for at least 1 numeric element
          the_values = the_sub_data_frame[,the_var_name]
          median_name = final_var_name_list[[the_var_name]][["MED"]]
          qr1_name = final_var_name_list[[the_var_name]][["25QR"]]
          qr2_name = final_var_name_list[[the_var_name]][["75QR"]]
          mean_name = final_var_name_list[[the_var_name]][["MEAN"]]
          sd_name = final_var_name_list[[the_var_name]][["SD"]]
          n_name = final_var_name_list[[the_var_name]][["N"]]
          cur_n = sum(is.numeric(the_values))
          the_new_row[1,n_name] = cur_n
          
          if (sum(is.numeric(the_values)) > 0)
          {
            # We know we are dealing with potentially unclean data and 
            # missing values, so go ahead and ignore NA's
            the_new_row[1, median_name] = median(the_values,na.rm=TRUE)
            the_new_row[1, qr1_name] = quantile(the_values,probs = c(0.25),na.rm=TRUE)
            the_new_row[1, qr2_name] = quantile(the_values,probs = c(0.75),na.rm=TRUE)
            the_new_row[1, mean_name] = mean(the_values,na.rm=TRUE)
            the_new_row[1, sd_name] = sd(the_values,na.rm=TRUE)
          }
        }
        if (sum(!(is.na(the_new_row[1,])))>0)
        {
          the_new_row[1, time_column] = the_time_point
          the_new_row[1, condition_column] = current_condition
          data_frame_return = rbind(data_frame_return, the_new_row)
        }
      }
    }
  }
  data_frame_return
}


check_arguments = function(value, allowed_values, force = FALSE)
  # A little helper function to check if values are allowed
  # mainly intended to check if input arguments are OK
  # Arguments:
  #  value: the value to test
  #  allowed_values: a vector of allowed values
  #  force: if TRUE, if value is not in allowed_values,
  #         value will be coerced to the first element 
  #         of allowed_values
{
  pass_check_flag = FALSE
  if (value %in% allowed_values)
  {
    value_to_return = value
  } else if (force == TRUE) {
    if (length(allowed_values) > 0)
    {
      value_to_return = allowed_values[1]
    } else {
      value_to_return = NULL
    }
  } else {
    value_to_return = NULL
  } 
  value_to_return
}

get_nontarget_progression_time = function(trial_list, lesion_data_filename = "le.txt", file_var = "LEEVAL", file_value = 15, date_var = "LEDN", output_var_name = "PD_UNEQNONTARGET", patient_id_var = "USUBJID")
  # A function to scan lesion information file for nontarget progression, for example either unequivocal or 
  # new target lesions.
  # This will be added as a 0/1 that switches to 1 one once there is is unequivocal progression
  # Time series can be aligned similar to other variables 
  # Arguments:
  #  trial_list:            
  #  lesion_data_filename:  file to open, should be tab-delimited text with top row as headers
  #  file_var:              file variable to check
  #  file_var_value:        file variable value to check for in order to set the flag to true    
  #  date_var:              variable with dates to check.  Note we will try to coerce to date
  #                         and data without a date is ignored.  Date format YYYYMMMDD is preferred.
#  output_var_name:		  what to assign the new time series output to.  Preferred "standard" variables are:
#                            "PD_UNEQNONTARGET": unequivocal progression of nontarget lesions
#                            "PD_NEW": progression due to appearance of new target lesions
#  patient_id_var
#
# Returns
#  trial_list:            The input list structure with the additional data/flag
#                         in "time_series" for each patient and value for each timepoint
#   
#
{
  options(stringsAsFactors = FALSE)
  
  data_table = read.delim(lesion_data_filename, na.strings = c("#N/A","."), check.names = FALSE)
  my_size = dim(data_table)
  temp = as.Date("0001-01-01") + 1:my_size[1]
  for (row_counter in seq(1,my_size[1]))
  {
    the_date = (data_table[row_counter,date_var])
    if ((is.character(the_date)))
      if (nchar(the_date) > 0)
      {
        temp[row_counter] = convert_to_date(the_date)
      } else {
        temp[row_counter] = NA
        #data_table[row_counter,date_var] = NA
      } else {
        temp[row_counter] = NA
        #data_table[row_counter,date_var] = NA
      }
  }
  data_table[,date_var] = temp
  data_table = data_table[which((!(is.na(data_table[,date_var])))),]
  data_table = data_table[order(data_table[,date_var]),]
  
  patient_ids = names(trial_list)
  for (cur_patient_id in patient_ids)
  {
    cur_rows = which(data_table[,patient_id_var]==cur_patient_id) 
    cur_data = data_table[cur_rows,]
    cur_rows = which(cur_data[,file_var]==file_value) 
    all_dates = unique(cur_data[,date_var])
    all_dates = all_dates[order(all_dates)]
    flag_vector = matrix(0,length(all_dates),1)
    if (length(all_dates) > 0)
    {
      prog_dates = cur_data[cur_rows,date_var]
      #prog_dates = prog_dates[which((!(is.na(prog_dates))))]
      prog_dates = unique(prog_dates)
      prog_dates = prog_dates[order(prog_dates)]
      prog_date = prog_dates[1]
      flag_indices = which(all_dates>=prog_date)
      flag_vector[flag_indices] = 1
    }
    all_dates = all_dates - trial_list[[cur_patient_id]][["first_dose_date"]]
    trial_list[[cur_patient_id]][["time_series"]][[output_var_name]] = list()
    for (date_counter in seq(1, length(all_dates)))
    {
      trial_list[[cur_patient_id]][["time_series"]][[output_var_name]][[as.character(all_dates[date_counter])]] = flag_vector[date_counter]
    }
    
  }
  
  trial_list
  
}


get_nontarget_progression_time_lnsize_filter = function(trial_list, lesion_data_filename = "le.txt", file_var = "LEEVAL", file_value = 15, date_var = "LEDN", output_var_name = "PD_UNEQNONTARGET", patient_id_var = "USUBJID", lesion_description_var = "LESITE", lesion_measure_var = "LED1ME", lymphnode_marker=c(" node"," node ", "node ", "lymphnode", "lymph node","lymph","lynph", " ln","ln ", " ln "))
  # A function to scan lesion information file for nontarget progression, for example either unequivocal or 
  # new target lesions.  This function also checks whether the nontarget lesions are lymph nodes
  # which are only considered pathological if their short axis measure is greater than or equal to 10 mm by RECIST 1.1
  # This will be added as a 0/1 that switches to 1 one once there is is unequivocal progression
  # Time series can be aligned similar to other variables 
  # Arguments:
  #  trial_list:            
  #  lesion_data_filename:  file to open, should be tab-delimited text with top row as headers
  #  file_var:              file variable to check
  #  file_var_value:        file variable value to check for in order to set the flag to true    
  #  date_var:              variable with dates to check.  Note we will try to coerce to date
#                         and data without a date is ignored.  Date format YYYYMMMDD is preferred.
#  output_var_name:		  what to assign the new time series output to.  Preferred "standard" variables are:
#                            "PD_UNEQNONTARGET": unequivocal progression of nontarget lesions
#                            "PD_NEW": progression due to appearance of new target lesions
#  patient_id_var
#  lesion_description_var
#  lesion_measure_var
#  lymphnode_marker       vector of strings that are used to label lymph nodes in data set
#
# Returns
#  trial_list:            The input list structure with the additional data/flag
#                         in "time_series" for each patient and value for each timepoint
#   
#
{
  options(stringsAsFactors = FALSE)
  path_ln_min = 10
  data_table = read.delim(lesion_data_filename, na.strings = c("#N/A","."), check.names = FALSE)
  my_size = dim(data_table)
  temp = as.Date("0001-01-01") + 1:my_size[1]
  for (row_counter in seq(1,my_size[1]))
  {
    the_date = (data_table[row_counter,lesion_eval_date_header])
    if ((is.character(the_date)))
      if (nchar(the_date) > 0)
      {
        temp[row_counter] = convert_to_date(the_date)
      } else {
        temp[row_counter] = NA
        #data_table[row_counter,lesion_eval_date_header] = NA
      } else {
        temp[row_counter] = NA
        #data_table[row_counter,lesion_eval_date_header] = NA
      }
  }
  data_table[,date_var] = temp
  data_table = data_table[which((!(is.na(data_table[,lesion_eval_date_header])))),]
  data_table = data_table[order(data_table[,lesion_eval_date_header]),]
  
  patient_ids = names(trial_list)
  for (cur_patient_id in patient_ids)
  {
    cur_rows = which(data_table[,patient_id_var]==cur_patient_id)
    cur_data = data_table[cur_rows,]
    cur_rows = which(cur_data[,file_var]==file_value)
    pass_check = cur_rows
    # Also check the rows, if they are a LN we will check to see if we can exclude based on size
    for (row_counter in seq(1, length(cur_rows)))
    {
      ulesion_information = cur_data[cur_rows[row_counter],lesion_description_var]
      if (length(ulesion_information) > 0)
      {
        #if ((grepl(" node", tolower(ulesion_information), fixed = TRUE)) | (grepl(" node ", tolower(ulesion_information), fixed = TRUE)) | (grepl("lymphnode", tolower(ulesion_information), fixed = TRUE)))
        if (grepl(paste(lymphnode_marker,collapse="|"), tolower(ulesion_information), fixed = F))
        {
          {
            cur_lesion_measure = check_and_convert_numeric(cur_data[cur_rows[row_counter],lesion_measure_var])
            #browser()
            #  CONVERT TO NUMERIC  cur_lesion_measure
            if (is.numeric(cur_lesion_measure) && (!is.na(cur_lesion_measure)))
            {
              # lymph nodes only considered pathologic if greater than 10 mm
              # we only remove the lesion if there is evidence to
              # support removal, otherwise we assume the
              # pathologist included it to indicated off target
              # progression
              if (cur_lesion_measure < path_ln_min)
              {
                pass_check[row_counter] = NA
              }
            }
          }				
        }
      }
    }
    #cur_rows = cur_rows[!is.na(cur_rows)]
    cur_rows = cur_rows[which(!is.na(cur_rows) & !is.na(pass_check))]
    #cur_rows = cur_rows[cur_check]
    #cur_data = cur_data[cur_rows,]
    
    all_dates = unique(cur_data[,lesion_eval_date_header])
    all_dates = all_dates[order(all_dates)]
    flag_vector = matrix(0,length(all_dates),1)
    if (length(all_dates) > 0)
    {
      
      prog_dates = cur_data[cur_rows,lesion_eval_date_header]
      #prog_dates = prog_dates[which((!(is.na(prog_dates))))]
      prog_dates = unique(prog_dates)
      prog_dates = prog_dates[order(prog_dates)]
      prog_date = prog_dates[1]
      flag_indices = which(all_dates>=prog_date)
      flag_vector[flag_indices] = 1
    }
    all_dates = all_dates - trial_list[[cur_patient_id]][["first_dose_date"]]
    trial_list[[cur_patient_id]][["time_series"]][[output_var_name]] = list()
    for (date_counter in seq(1, length(all_dates)))
    {
      trial_list[[cur_patient_id]][["time_series"]][[output_var_name]][[as.character(all_dates[date_counter])]] = flag_vector[date_counter]
    }
    
  }
  
  trial_list
  
}


## modified by Lu 
get_nontarget_lesions = function(trial_list, lesion_data_filename = "le.txt", lesion_type_var = "LETYPE", target_value = 1, lesion_id_var = "LETUCD", lesion_measure_var = "LED1ME", lesion_description_var = "LESITE", date_var = "LEDN", eval_var = "LEEVAL", exclude_codes = c(2), patient_id_var = "USUBJID",lymphnode_marker=c(" node"," node ", "node ", "lymphnode", "lymph node","lymph","lynph", " ln","ln ", " ln "),filter_var='', filter_value='')
  # A simple function to get the number of reported nontarget lesions for LN and non-LN non-target lesions at each time point 
  # Arguments:
  #  trial_list:            
  #  lesion_data_filename:   file to open, should be tab-delimited text with top row as headers
  #  lesion_type_var:        file variable to check for lesion type
  #  target_value:            value for index lesions: these will be ignored
  #  lesion_id_var
  #  lesion_measure_var
  #  lesion_description_var: variable with the text description, this will be checked for lymph nodes       
  #  date_var:               variable with dates to check.  Note we will try to coerce to date
  #                          and data without a date is ignored.  Date format YYYYMMMDD is preferred.
#  eval_var:               variable with lesion evaluations
#  exclude_codes:          a vector of codes for lesions to exclude - for example "absent"
#  patient_id_var
#
# Returns
#  trial_list:             updated data structure.  Number of non-target lesions evaluated will be noted in:
#                           NONTARGETNLN_N, NONTARGETLN_N
#                             
#   
#
{
  options(stringsAsFactors = FALSE)
  data_table = read.delim(lesion_data_filename, na.strings = c("#N/A","."), check.names = FALSE)
  # First filter on non-target lesions
  if(nchar(filter_var)>0){
    data_table = data_table[which(((data_table[,lesion_type_var]!=target_value)) & (data_table[,filter_var]!=filter_value)),]
  }else{
    data_table = data_table[which(((data_table[,lesion_type_var]!=target_value))),]
  }
  
  # Now don't filter on lesions with measures - we won't require measures to be present
  # data_table = data_table[which((!(is.na(data_table[,lesion_measure_var])))),]
  data_table = data_table[which((!(is.na(data_table[,eval_var])))),]
  # We can also filter out lesions that don't fulfill our consideration criteria
  data_table = data_table[which(!(is.element(data_table[,eval_var],exclude_codes))),]
  
  my_size = dim(data_table)
  temp = as.Date("0001-01-01") + 1:my_size[1]
  for (row_counter in seq(1,my_size[1]))
  {
    the_date = (data_table[row_counter,lesion_eval_date_header])
    if ((is.character(the_date)))
      if (nchar(the_date) > 0)
      {
        temp[row_counter] = convert_to_date(the_date)
      } else {
        temp[row_counter] = NA
        #data_table[row_counter,lesion_eval_date_header] = NA
      } else {
        temp[row_counter] = NA
        #data_table[row_counter,lesion_eval_date_header] = NA
      }
  }
  data_table[,date_var] = temp
  data_table = data_table[which((!(is.na(data_table[,lesion_eval_date_header])))),]
  data_table = data_table[order(data_table[,lesion_eval_date_header]),]
  
  
  patient_ids = names(trial_list)
  for (cur_patient_id in patient_ids)
  {
    cur_patient_rows = which(data_table[,patient_id_var]==cur_patient_id) 
    cur_patient_data = data_table[cur_patient_rows,]
    all_dates = unique(cur_patient_data[,lesion_eval_date_header])
    all_dates = all_dates[order(all_dates)]
    
    # Should get times from index lesions too, so we don't miss non-index lesions if they were
    # measured at odd times.
    my_times = c()
    my_index_lesions = names(trial_list[[cur_patient_id]][["index_lesions"]])
    for (my_lesion in my_index_lesions)
    {
      my_vals = names(trial_list[[cur_patient_id]][["index_lesions"]][[my_lesion]][[lesion_measure_var]])
      if (length(my_vals) > 0)
      {
        my_times = c(my_times,(as.numeric((my_vals))))
      }
    }
    my_times = unique(my_times)
    if (length(my_times > 0))
    {
      my_times = my_times + trial_list[[cur_patient_id]][["first_dose_date"]]
      all_dates = c(all_dates,my_times)
      all_dates = unique(all_dates)
      all_dates = all_dates[order(all_dates)]
    }
    
    # We want 2 lesion-related variables at each time point
    NONTARGETNLN_N_vector = matrix(0,length(all_dates),1)
    NONTARGETLN_N_vector = matrix(0,length(all_dates),1)
    trial_list[[cur_patient_id]][["time_series"]][["NONTARGETLN_N"]] = list()
    trial_list[[cur_patient_id]][["time_series"]][["NONTARGETNLN_N"]] = list()	
    lesions_to_test = unique(cur_patient_data[,lesion_id_var])
    lesions_to_test = lesions_to_test[which(!(is.na(lesions_to_test)))]
    
    
    
    if (length(all_dates) > 0)
    {
      for (date_counter in seq(1, length(all_dates)))
      {
        nnln_lesions = 0
        nln_lesions = 0
        # There is more efficient code to do this but I am in a hurry...
        for (lesion_id in lesions_to_test)
        {
          cur_rows = which(cur_patient_data[,lesion_id_var]==lesion_id)
          cur_data = cur_patient_data[cur_rows,]
          if (all_dates[date_counter] >= min(cur_data[,lesion_eval_date_header]))
          {
            if (all_dates[date_counter] <= max(cur_data[,lesion_eval_date_header]))
            {
              ulesion_information = unique(cur_data[,lesion_description_var])
              ulesion_information = ulesion_information[ulesion_information==max(ulesion_information)]
              ulesion_information = ulesion_information[1]
              #ln_rows = ((grepl(" node", tolower(ulesion_information), fixed = TRUE)) | (grepl(" node ", tolower(ulesion_information), fixed = TRUE)) | (grepl("lymphnode", tolower(ulesion_information), fixed = TRUE)))
              ln_rows = (grepl(paste(lymphnode_marker,collapse="|"), tolower(ulesion_information), fixed = F))
              #ln_rows = (grepl(paste(lymphnode_marker,collapse="|"), toupper(ulesion_information), fixed = F))
              if (ln_rows)
              {
                nln_lesions = nln_lesions+1
              } else {
                nnln_lesions = nnln_lesions +1
              }
            }	
          }	
        }
        NONTARGETLN_N_vector[date_counter] = nln_lesions
        NONTARGETNLN_N_vector[date_counter] = nnln_lesions					
      }
      
      all_dates = all_dates - trial_list[[cur_patient_id]][["first_dose_date"]]
      for (date_counter in seq(1, length(all_dates)))
      {
        trial_list[[cur_patient_id]][["time_series"]][["NONTARGETLN_N"]][[as.character(all_dates[date_counter])]] = NONTARGETLN_N_vector[date_counter]
        trial_list[[cur_patient_id]][["time_series"]][["NONTARGETNLN_N"]][[as.character(all_dates[date_counter])]] = NONTARGETNLN_N_vector[date_counter]
      }
    }
  }
  
  trial_list
  
}


create_distribution_tables_recist = function(patient_data_filename, response_varname = "INDEX_LESION_SLD_RELCH", offtrtime_varname = "OFFTRTIME", ontrt_varname = "ONTRT", offtrreason_varname = "OFFTRTREASON", patient_id_varname="USUBJID",time_varname="TIME", sld_varname = "INDEX_LESION_SLD", maxln_varname = "INDEX_LESION_MAXLN",nonlnsld_varname = "INDEX_LESION_NLNSLD",nontarget_pd_varnames=c("PD_UNEQNONTARGET","PD_NEW"),nontarget_varnames=c("NONTARGETNLN_N","NONTARGETLN_N"))
  # A function to create tables of patient data arranged so each row shows
  # the individual responses at a given timepoint.
  # This also uses RECIST1.1 criteria for creating and defining
  # response bins 
  # Arguments:
  #  patient_data_filename: a file with the integrated patient data
  #  response_varname:      the variable in the file with the lesion response of interest
  #  offtrtime_varname:     the number of quantitative output bins
  #  offtrreason_varname:   the variable with the reason for going off treatment
  #  patient_id_varname:    the variable with the patient ID
  #  time_varname:          the variable with the time
#  sld_varname:           SLD for the index lesions
#  maxln_varname:         a variable with the largest target LN measure
#  nonlnsld_varname:      a variable with the SLD measure not including lymph node
#                         lesions        				  
#  nontarget_pd_varnames: a vector of variable names with nontarget time series 
#                         pd measures where a 1 will trigger a rating of PD
#  nontarget_varnames:	  a vector of variable names with indications of the number of nontarget lesions remaining
#
# Returns
#  A list with 4 data frames
#   output_matrix:           output lesion measure changes
#   output_matrix_score:     response score for patient 
#   output_matrix_bor_score: BOR score for patients
#   on_therapy_matrix:       boolean matrix for whether a patient is on therapy 
#   integrate_patient_data:  the table read in from patient_data_filename, updated with RSCORE,BRSCORE, RSCORENEW,BRSCORENEW, CBRSCORE, CBRSCORENEW
{
  
  my_table = read.table(patient_data_filename, header = TRUE, check.names = FALSE, sep = '\t',na.strings = ".", stringsAsFactors = FALSE)
  output_times = sort(unique(my_table[,time_varname]))
  #for (treatment_counter in seq(1,n_treatments))
  #{
  #cur_treatment = treatment_types[treatment_counter]
  # We can reduce the number of columns right away
  temp = my_table[,c(patient_id_varname,offtrtime_varname,ontrt_varname,offtrreason_varname,time_varname,response_varname,sld_varname,maxln_varname,nonlnsld_varname,nontarget_pd_varnames,nontarget_varnames)]
  
  #if (is.na(cur_treatment))
  #{
  #	temp = temp[is.na(temp[,treatmengp_varname]),]
  #} else {
  #	temp = temp[which(temp[,treatmengp_varname] == cur_treatment),]
  #}
  
  # Now walk through the patient IDs and output the data
  my_patient_ids = sort(unique(temp[,patient_id_varname]))
  n_patients = length(my_patient_ids)
  
  n_time_points = length(output_times)
  
  output_matrix_1 = matrix(NA, n_time_points, n_patients)
  sld_matrix_1 = matrix(NA, n_time_points, n_patients)
  nontargetpd_matrix_1 = matrix(NA, n_time_points, n_patients)
  maxln_matrix_1 = matrix(NA, n_time_points, n_patients)
  nonlnsld_matrix_1 = matrix(NA, n_time_points, n_patients)
  nontarget_matrix_1 = matrix(NA, n_time_points, n_patients)
  output_matrix_score_1 = matrix(NA, n_time_points, n_patients)
  output_matrix_bor_score_1 = matrix(NA, n_time_points, n_patients)
  on_therapy_matrix_1 = matrix(NA, n_time_points, n_patients)
  ## Yougan addition starts: add the rscore, brscore, and ontherapy alternatives
  ## where we don't track off treatment
  output_matrix_score_2 = matrix(NA, n_time_points, n_patients)
  output_matrix_bor_score_2 = matrix(NA, n_time_points, n_patients)
  on_therapy_matrix_2 = matrix(NA, n_time_points, n_patients)
  colnames(output_matrix_score_2) = my_patient_ids
  colnames(output_matrix_bor_score_2) = my_patient_ids
  colnames(on_therapy_matrix_2) = my_patient_ids
  rownames(output_matrix_score_2) = output_times
  rownames(output_matrix_bor_score_2) = output_times
  rownames(on_therapy_matrix_2) = output_times
  ## Yougan addition ends
  
  
  
  colnames(output_matrix_1) = my_patient_ids
  colnames(sld_matrix_1) = my_patient_ids
  colnames(output_matrix_score_1) = my_patient_ids
  colnames(output_matrix_bor_score_1) = my_patient_ids
  colnames(on_therapy_matrix_1) = my_patient_ids
  
  rownames(output_matrix_1) = output_times
  rownames(sld_matrix_1) = output_times
  rownames(output_matrix_score_1) = output_times
  rownames(output_matrix_bor_score_1) = output_times
  rownames(on_therapy_matrix_1) = output_times
  
  row_matrix = matrix(NA, n_time_points, n_patients)
  
  
  
  # Right now we just export the lesion data
  for (time_point_counter in seq_along(output_times))
  {
    for (patient_counter in seq_along(my_patient_ids))
    {
      cur_rows = which(temp[,patient_id_varname]==my_patient_ids[patient_counter])
      cur_row  = which(temp[cur_rows,time_varname]==output_times[time_point_counter])
      
      if (length(cur_row)>0)
      {
        output_matrix_1[time_point_counter,patient_counter] = temp[cur_rows[cur_row],response_varname]
        sld_matrix_1[time_point_counter,patient_counter] = temp[cur_rows[cur_row],sld_varname]
        nontargetpd_matrix_1[time_point_counter,patient_counter] = max(temp[cur_rows[cur_row],nontarget_pd_varnames])
        maxln_matrix_1[time_point_counter,patient_counter] = temp[cur_rows[cur_row],maxln_varname]
        nonlnsld_matrix_1[time_point_counter,patient_counter] = temp[cur_rows[cur_row],nonlnsld_varname]
        nontarget_matrix_1[time_point_counter,patient_counter] = sum(temp[cur_rows[cur_row],nontarget_varnames])
        # If we have no measures at a time point for nontarget lesions, we have to assume they are 0 to move forward
        if (is.na(nontarget_matrix_1[time_point_counter,patient_counter]))
        {
          nontarget_matrix_1[time_point_counter,patient_counter] = 0
        }
        row_matrix[time_point_counter,patient_counter] = cur_rows[cur_row]
      } else {
        output_matrix_1[time_point_counter,patient_counter] = NA
        sld_matrix_1[time_point_counter,patient_counter] = NA
        row_matrix[time_point_counter,patient_counter] = NA
        nontargetpd_matrix_1[time_point_counter,patient_counter] = NA#max(temp[cur_rows[cur_row],nontarget_pd_varnames])
        maxln_matrix_1[time_point_counter,patient_counter] = NA#temp[cur_rows[cur_row],maxln_varname]
        nonlnsld_matrix_1[time_point_counter,patient_counter] = NA#temp[cur_rows[cur_row],nonlnsld_varname]	
        nontarget_matrix_1[time_point_counter,patient_counter] = NA
      }
    }
  }
  
  
  # Now we want to step through each VP and create a matrix
  # of PD, SD, PR, CR at each time point.  We also add "PD2" for
  # PD we can't model as lesion size changes, i.e. PD in non-target
  # lesions
  # Also include off treatment for other reasons like death (DT) or toxicity (RX)
  # get baseline value, get minimum
  remove_columns = c()
  for (patient_counter in seq_along(my_patient_ids))
  {
    # Assume the first measure is baseline
    # Some patients will not have any lesion measures.
    if (!(is.na(output_matrix_1[1,patient_counter])))
    {
      
      min_rel_sld = output_matrix_1[1,patient_counter]
      min_sld = sld_matrix_1[1,patient_counter]
      cur_row = row_matrix[1,patient_counter]
      
      # off_treatment_time = temp[cur_row,offtrtime_varname]
      ## Lu modified this, take the maximum of offtrttime and the maximum aligned ONTRT time (lesion scan time) as the offtreatment time
      ## it is tricky, because we also need to populate ONTRT=1 between the lesion scan times for biomarker data collection
      pt_rows = row_matrix[,patient_counter]
      pt_rows = pt_rows[!is.na(pt_rows)]
      pt_temp = temp[pt_rows,]
      
      offtrtime = pt_temp[1,offtrtime_varname]
      ontrt = pt_temp[,ontrt_varname]
      ontrttime = pt_temp[which(ontrt==1),time_varname]
      off_treatment_time = max(c(offtrtime,ontrttime))
      
      if (is.na(off_treatment_time))
      {
        off_treatment_time = Inf
      }
      off_treatment_reason = temp[cur_row,offtrreason_varname]
      continue_flag = TRUE
      # We will keep patients that just have NA recorded for now
      # and rely on the PW algorithms to filter them appropriately.
      last_treatment_counter = 1
      
      for (time_point_counter in seq_along(output_times))
      {
        if (continue_flag)
        {	
          cur_time = output_times[time_point_counter]
          cur_rel_sld = output_matrix_1[time_point_counter,patient_counter]
          cur_sld = sld_matrix_1[time_point_counter,patient_counter]
          cur_nontarget_pd = nontargetpd_matrix_1[time_point_counter,patient_counter]
          cur_maxln = maxln_matrix_1[time_point_counter,patient_counter]
          cur_nonlnsld = nonlnsld_matrix_1[time_point_counter,patient_counter]
          cur_nontarget = nontarget_matrix_1[time_point_counter,patient_counter]					
          
          if (is.na(cur_rel_sld))
          {
            cur_pd_check_1 = NA
            cur_pd_check_2 = NA
          } else {
            if (cur_rel_sld < min_rel_sld)
            {
              min_rel_sld = cur_rel_sld
              min_sld = cur_sld
            }
            
            # RECIST measures also need to track relative to nadir
            # to trigger PD classification appropriately
            if (min_rel_sld != 0)
            {
              if (min_rel_sld > -1)
              {
                cur_pd_check_1 = (cur_rel_sld - min_rel_sld)/(min_rel_sld + 1)
                cur_pd_check_2 = cur_sld - min_sld
              } else {
                # We check to trigger PD classification based on increase if we have
                # no target lesion left
                cur_pd_check_1 = .21
                cur_pd_check_2 = cur_sld - min_sld
              }
            } else {
              cur_pd_check_1 = 0
              cur_pd_check_2 = 0
            }
          }
          
          if (cur_time <= off_treatment_time)
            
          {
            last_treatment_counter = time_point_counter
            if (!(is.na(cur_rel_sld)))
            {
              if ((((cur_pd_check_1 >= 0.2) & (cur_pd_check_2 >= 5)) | (cur_rel_sld >= 0.2)) ) {
                output_matrix_score_1[time_point_counter,patient_counter] = "PD"
                
              } else if (cur_nontarget_pd>0) {
                # Nontarget PD will be flagged as different, decide how to best handle with PW algorithm
                output_matrix_score_1[time_point_counter,patient_counter] = "PD2"
                
                #} else if ((cur_pd_check_1 < 0.2) & ((cur_rel_sld <= -1) | ((cur_nonlnsld==0)&(cur_maxln<10))) & (cur_nontarget_pd<=0) & (cur_nontarget<=0))
              } else if (((cur_rel_sld <= -1) | ((cur_nonlnsld==0)&(cur_maxln<10))) & (cur_nontarget_pd<=0) & (cur_nontarget<=0))
              {
                if (cur_rel_sld <= -1)
                {
                  # This is a CR with disapearance of all lesions
                  output_matrix_score_1[time_point_counter,patient_counter] = "CR"
                }
                else 
                {
                  # This is a CR with some residual target lymph node vlume.  So the 
                  # relative SLD will not be -1, but we will have a complete response.
                  # We could try to refine this as "CR2" for modeling lesion dynamics,
                  # but it is not a common case and has not been problematic yet.
                  output_matrix_score_1[time_point_counter,patient_counter] = "CR"
                }
                
                #} else if ((cur_pd_check_1 < 0.2) & (cur_rel_sld <= -.3)) {
              } else if (cur_rel_sld <= -.3) {
                output_matrix_score_1[time_point_counter,patient_counter] = "PR"
                
                
                #} else if ((cur_pd_check_1 < 0.2) & (cur_rel_sld < 0.2)) {
              } else if (cur_rel_sld < 0.2) {
                output_matrix_score_1[time_point_counter,patient_counter] = "SD"
                
                #} else if (((cur_pd_check_1 >= 0.2) & (cur_pd_check_2 < 5))) {
                #	# In this case the increase does not merit a classification
                #	# as PD
                #	if (((cur_rel_sld <= -1) | ((cur_nonlnsld==0)&(cur_maxln<10))) & (cur_nontarget_pd<=0) & (cur_nontarget<=0))
                #	{
                #		output_matrix_score_1[time_point_counter,patient_counter] = "CR"
                #	} else if ((cur_rel_sld <= -.3)) {
                #		output_matrix_score_1[time_point_counter,patient_counter] = "PR"
                #	} else if ((cur_rel_sld < 0.2)) {
                #		output_matrix_score_1[time_point_counter,patient_counter] = "SD"	
                #	#}
                #	#	output_matrix_score_1[time_point_counter,patient_counter] = output_matrix_score_1[time_point_counter-1,patient_counter]
                #	} else {
                #		# print(my_patient_ids[patient_counter])
                #		output_matrix_score_1[time_point_counter,patient_counter] = NA
                #	}
              } else {
                print(paste("Warning: no valid measure for patient ",my_patient_ids[patient_counter]," at time ",toString(cur_time),".", sep = ""))
                output_matrix_score_1[time_point_counter,patient_counter] = NA
                
              }
            } else if (!(is.na(cur_nontarget_pd))) {
              if (cur_nontarget_pd > 0)
              {
                # We still need to check for other sources of PD even if index measures were not taken
                # Nontarget PD will be flagged as different, decide how to best handle with PW algorithm
                output_matrix_score_1[time_point_counter,patient_counter] = "PD2"
                
              } else {
                output_matrix_score_1[time_point_counter,patient_counter] = NA
                
              }
            } else {
              output_matrix_score_1[time_point_counter,patient_counter] = NA
              
            }
            
            on_therapy_matrix_1[time_point_counter,patient_counter] = 1
            
            # Only look at BOR after the baseline measure, the baseline is not a response
            if (cur_time > 0)
            {
              # This should probably always be 2, but be safe here
              initial_index=(which(output_times>0))[1]
              if ("CR" %in% output_matrix_score_1[initial_index:time_point_counter,patient_counter])
              {
                output_matrix_bor_score_1[time_point_counter,patient_counter] = "CR"
                # We would add a CR2 check here if we did implement such a classification.
              } else if ("PR" %in% output_matrix_score_1[initial_index:time_point_counter,patient_counter]) {
                output_matrix_bor_score_1[time_point_counter,patient_counter] = "PR"
              } else if ("SD" %in% output_matrix_score_1[initial_index:time_point_counter,patient_counter]) {
                output_matrix_bor_score_1[time_point_counter,patient_counter] = "SD"
              } else if ("PD" %in% output_matrix_score_1[initial_index:time_point_counter,patient_counter]) {
                output_matrix_bor_score_1[time_point_counter,patient_counter] = "PD"
              } else if ("PD2" %in% output_matrix_score_1[initial_index:time_point_counter,patient_counter]) {
                output_matrix_bor_score_1[time_point_counter,patient_counter] = "PD2"									
              } else {
                output_matrix_bor_score_1[time_point_counter,patient_counter] = NA
              }	
            }
            
            
            # this ends the before drop-off time point processing
          } else {
            # We may want to add a scenario for complete response here.... though there may be some issues 
            # in using an investigator assessed CR.  
            # Since we include more considerations in computing response, we will not use investigator assessment here.  Let's see how
            # this algorithm does
            # if ((off_treatment_reason %in% c("MAXIMUM CLINICAL BENEFIT","COMPLETE RESPONSE")) & ("CR" %in% output_matrix_score[last_treatment_counter,patient_counter]))
            if (("CR" %in% output_matrix_score_1[last_treatment_counter,patient_counter]))
            {
              output_matrix_score_1[time_point_counter,patient_counter] = "CR"
              # We would add a CR2 check here if we did implement CR2 to handle residual target
              # lymph node size handling.
              # We will keep similar strict criteria for PD, this means the investigator may score some patients as PD that we do not
              # } else if ((off_treatment_reason %in% c("DISEASE PROGRESSION")) & ("PD" %in% output_matrix_score[last_treatment_counter,patient_counter]))
            } else if (("PD" %in% output_matrix_score_1[last_treatment_counter,patient_counter]))
            {
              output_matrix_score_1[time_point_counter,patient_counter] = "PD"
            } else if (("PD2" %in% output_matrix_score_1[last_treatment_counter,patient_counter]))
            {
              output_matrix_score_1[time_point_counter,patient_counter] = "PD2"
              # We will keep similar strict criteria for PD, this means the investigator may score some patients as PD that we do not
            } else if (off_treatment_reason %in% union(unique(temp[,offtrreason_varname]),c("MAXIMUM CLINICAL BENEFIT","STUDY DRUG TOXICITY","COMPLETED TREATMENT AS PER PROTOCOL","FOLLOWUP NO LONGER REQUIRED PER PROTOCOL","SUBJECT WITHDREW CONSENT","SUBJECT NO LONGER MEETS STUDY CRITERIA","SUBJ REQUEST TO DISCONTINUE STUDY TRT","ADVERSE EVENT UNRELATED TO STUDY DRUG","SUBJECT REQUEST TO DISCONTINUE STUDY TREATMENT","NOT REPORTED","OTHER","DT","ADVERSE EVENT","DEATH","LOST TO FOLLOW-UP","NA","NA.","COMPLETION OF MAXIMUM CYCLES","DISEASE PROGRESSION","COMPLETE RESPONSE","MAXIMUM CLINICAL BENEFIT","POOR/NON-COMPLIANCE","COMPLETED","PROGRESSIVE DISEASE","WITHDRAWAL BY SUBJECT","FAILURE TO MEET STUDY CRITERIA","DOCUMENTED DISEASE PROGRESSION","ADMINISTRATIVE REASON BY SPONSOR","DETERIORATION W/O DOCUMENTED PROGRESSION","NEVER TREATED WITH STUDY MEDICATION","PHYSICIAN DECISION","SUBJECT REQUEST","AE/SAE MYELITIS","CREATININE INCREASED GRADE 3","ENTERIC COLITIS","DRUG INDUCED LIVER INJURY","EROSIVE GASTRIC DRUG RELATED","LONGLASTING DIARRHEA","HEPATITIS","AUTOIMMUNE COLITIS WITH FEVER","COLITIS","PATIENT HAS OPTED TO WITHDRAW CONSENT FROM TREATMENT & ALL RELATED PROCEDURES DUE TO RISKS/BENEFITS & THE GEOGRAPHICAL CHALLENGE OF THE STUDY. PATIENT IS OKAY TO BE CONTACTED FOR SURVIVAL ONLY.","THE PATIENT DECIDED TO BE NOT FOLLOWED FOR CLINICAL AND SURVIVAL FOLLOW UP NEITHER","ISCHEMIC ICTUS","SAE HEPATITIS","HEPATITIS AND CHOLANGITIS","PANKREATITIS","THE PATIENT HAS TWO CANCER DIAGNOSIS","PATIENT WILL RETURN TO HOME COUNTRY AND WILL NOT BE FOLLOWED UP","LACK OF CLINICAL EFFICIENCY","ABDOMINAL ADENOPATHY BIOPSY SHOWED FOLLICULAR LYMPHOMA","IMMUNE RELATED COLITIS","SELECTIVE SURGERY TO REMOVE  L HEMIABDOMINAL MASS.IT WAS THE ONLY TARGET LESION FOR RECIST","PATIENT REQUESTED TO PURSUE SURGICAL TREATMENT OF THE R ORBIT LESION","BUT DATE OF DEATH CAN BE REPORTED","SUBJECT WOULD LIKE TO WITHDRAW CONSENT FROM ALL STUDY-RELATED PROCEDURES, INCLUDING RECONSENTING, SAFETY & SURVIVAL FOLLOW UPS. PATIENT IS INTERESTED IN JOINING A DIFFERENT CLINICAL TRIAL ELSEWHERE.","PATIENT HAS REQUESTED TO WITHDRAW FROM STUDY TX DUE TO FINANCIAL BURDEN.",NA)))  {
              output_matrix_score_1[time_point_counter,patient_counter] = "OS"
            } else {
              print(paste("Warning: off treatment without recognized rationale for ",my_patient_ids[patient_counter],"!  It is: ",off_treatment_reason,".  Inputting PD2.", sep=""))
              output_matrix_score_1[time_point_counter,patient_counter] = "PD2"
            }
            on_therapy_matrix_1[time_point_counter,patient_counter] = 0
            # ("OS" %in% output_matrix_bor_score[1:time_point_counter,patient_counter])
            # {
            # 		output_matrix_bor_score[time_point_counter,patient_counter] = "OS"
            # } else if 
            initial_index=(which(output_times>0))[1]
            if ("CR" %in% output_matrix_score_1[initial_index:time_point_counter,patient_counter]) 
            {
              output_matrix_bor_score_1[time_point_counter,patient_counter] = "CR"
              # We would add a CR2 check here if we did implement CR2 to handle residual target
              # lymph node size handling.									
            } else if ("PR" %in% output_matrix_score_1[initial_index:time_point_counter,patient_counter]) {
              output_matrix_bor_score_1[time_point_counter,patient_counter] = "PR"
            } else if ("SD" %in% output_matrix_score_1[initial_index:time_point_counter,patient_counter]) {
              output_matrix_bor_score_1[time_point_counter,patient_counter] = "SD"
            } else if ("PD" %in% output_matrix_score_1[initial_index:time_point_counter,patient_counter]) {
              output_matrix_bor_score_1[time_point_counter,patient_counter] = "PD"
            } else if ("PD2" %in% output_matrix_score_1[initial_index:time_point_counter,patient_counter]) {
              output_matrix_bor_score_1[time_point_counter,patient_counter] = "PD2"									
            } else {
              output_matrix_bor_score_1[time_point_counter,patient_counter] = NA
            }
          } # this should be the end of the old dropoff algorithm
        }
        
        
        
        #Yougan addition starts: we won't distinguish time_point_counter is before or after drop-off time and calculated rscore and brscore as it is
        
        
        if (!(is.na(cur_rel_sld)))
        {
          if ((((cur_pd_check_1 >= 0.2) & (cur_pd_check_2 >= 5)) | (cur_rel_sld >= 0.2)) ) {
            output_matrix_score_2[time_point_counter,patient_counter] = "PD"
            
          } else if (cur_nontarget_pd>0) {
            # Nontarget PD will be flagged as different, decide how to best handle with PW algorithm
            output_matrix_score_2[time_point_counter,patient_counter] = "PD2"
            
            #} else if ((cur_pd_check_1 < 0.2) & ((cur_rel_sld <= -1) | ((cur_nonlnsld==0)&(cur_maxln<10))) & (cur_nontarget_pd<=0) & (cur_nontarget<=0))
          } else if (((cur_rel_sld <= -1) | ((cur_nonlnsld==0)&(cur_maxln<10))) & (cur_nontarget_pd<=0) & (cur_nontarget<=0))
          {
            output_matrix_score_2[time_point_counter,patient_counter] = "CR"
            
            #} else if ((cur_pd_check_1 < 0.2) & (cur_rel_sld <= -.3)) {
          } else if (cur_rel_sld <= -.3) {
            output_matrix_score_2[time_point_counter,patient_counter] = "PR"
            
            
            #} else if ((cur_pd_check_1 < 0.2) & (cur_rel_sld < 0.2)) {
          } else if (cur_rel_sld < 0.2) {
            output_matrix_score_2[time_point_counter,patient_counter] = "SD"
            
            #} else if (((cur_pd_check_1 >= 0.2) & (cur_pd_check_2 < 5))) {
            #	# In this case the increase does not merit a classification
            #	# as PD
            #	if (((cur_rel_sld <= -1) | ((cur_nonlnsld==0)&(cur_maxln<10))) & (cur_nontarget_pd<=0) & (cur_nontarget<=0))
            #	{
            #		output_matrix_score_2[time_point_counter,patient_counter] = "CR"
            #	} else if ((cur_rel_sld <= -.3)) {
            #		output_matrix_score_2[time_point_counter,patient_counter] = "PR"
            #	} else if ((cur_rel_sld < 0.2)) {
            #		output_matrix_score_2[time_point_counter,patient_counter] = "SD"	
            #	#}
            #	#	output_matrix_score_2[time_point_counter,patient_counter] = output_matrix_score_2[time_point_counter-1,patient_counter]
            #	} else {
            #		# print(my_patient_ids[patient_counter])
            #		output_matrix_score_2[time_point_counter,patient_counter] = NA
            #	}
          } else {
            print(paste("Warning: no valid measure for patient ",my_patient_ids[patient_counter]," at time ",toString(cur_time),".", sep = ""))
            output_matrix_score_2[time_point_counter,patient_counter] = NA
            
          }
        } else if (!(is.na(cur_nontarget_pd))) {
          if (cur_nontarget_pd > 0)
          {
            # We still need to check for other sources of PD even if index measures were not taken
            # Nontarget PD will be flagged as different, decide how to best handle with PW algorithm
            output_matrix_score_2[time_point_counter,patient_counter] = "PD2"
            
          } else {
            output_matrix_score_2[time_point_counter,patient_counter] = NA
            
          }
        } else {
          output_matrix_score_2[time_point_counter,patient_counter] = NA
          
        }
        
        on_therapy_matrix_2[time_point_counter,patient_counter] = 1
        
        # Only look at BOR after the baseline measure, the baseline is not a response
        if (cur_time > 0)
        {
          # This should probably always be 2, but be safe here
          initial_index=(which(output_times>0))[1]
          if ("CR" %in% output_matrix_score_2[initial_index:time_point_counter,patient_counter])
          {
            output_matrix_bor_score_2[time_point_counter,patient_counter] = "CR"
          } else if ("PR" %in% output_matrix_score_2[initial_index:time_point_counter,patient_counter]) {
            output_matrix_bor_score_2[time_point_counter,patient_counter] = "PR"
          } else if ("SD" %in% output_matrix_score_2[initial_index:time_point_counter,patient_counter]) {
            output_matrix_bor_score_2[time_point_counter,patient_counter] = "SD"
          } else if ("PD" %in% output_matrix_score_2[initial_index:time_point_counter,patient_counter]) {
            output_matrix_bor_score_2[time_point_counter,patient_counter] = "PD"
          } else if ("PD2" %in% output_matrix_score_2[initial_index:time_point_counter,patient_counter]) {
            output_matrix_bor_score_2[time_point_counter,patient_counter] = "PD2"									
          } else {
            output_matrix_bor_score_2[time_point_counter,patient_counter] = NA
          }	
        }
        
        
        
        
        #Yougan addition ends
        
        # the following } ends the time_point_counter for loop
      }
    } else {
      output_matrix_score_1[1,patient_counter] = "OS"
      output_matrix_score_2[1,patient_counter] = "OS"   #yougan comment: this is to take care cases patients don't have any lesion measurements, not even at baseline
      
    }
  }
  temp2 = matrix(NA, (length(output_times)), 1)
  temp2 = data.frame(temp2, check.names = FALSE)
  # temp2[,1] = cur_treatment
  # temp2[,2] = response_varname
  temp2[,1] = output_times
  colnames(temp2) = c(time_varname)
  # "INTERVENTION_ID","EXP_VAR_ID",
  output_matrix = cbind(temp2,data.frame(output_matrix_1, check.names = FALSE))
  output_matrix_score = cbind(temp2,data.frame(output_matrix_score_1, check.names = FALSE))
  output_matrix_bor_score = cbind(temp2, data.frame(output_matrix_bor_score_1, check.names = FALSE))
  on_therapy_matrix = cbind(temp2, data.frame(on_therapy_matrix_1, check.names = FALSE))
  
  
  # yougan addition starts: add data frames for the new columns to be added
  output_matrix_score_new = cbind(temp2,data.frame(output_matrix_score_2, check.names = FALSE))
  output_matrix_bor_score_new = cbind(temp2, data.frame(output_matrix_bor_score_2, check.names = FALSE))
  on_therapy_matrix_new = cbind(temp2, data.frame(on_therapy_matrix_2, check.names = FALSE))
  # yougan addition ends 
  
  
  #}	
  
  #output_times = sort(unique(my_table[,time_varname]))
  my_patient_ids = sort(unique(my_table[,patient_id_varname]))
  
  # Now add the on therapy variables to the input matrix
  if (!("ONTRT" %in% colnames(my_table)))
  {
    my_table[,"ONTRT"]=NA
  }
  for (patient_counter in seq_along(my_patient_ids))
  {
    for (time_point_counter in seq_along(output_times))
    {
      cur_row = intersect(which(my_table[,patient_id_varname] == my_patient_ids[patient_counter]),which(my_table[,time_varname] == output_times[time_point_counter]))
      my_table[cur_row,"ONTRT"] = on_therapy_matrix[time_point_counter,my_patient_ids[patient_counter]]
    }
  }
  
  
  # yougan addition starts here
  # now add new on therapy variables to the input matrix
  
  if (!("ONTRTNEW" %in% colnames(my_table)))
  {
    my_table[,"ONTRTNEW"]=NA
  }
  for (patient_counter in seq_along(my_patient_ids))
  {
    for (time_point_counter in seq_along(output_times))
    {
      cur_row = intersect(which(my_table[,patient_id_varname] == my_patient_ids[patient_counter]),which(my_table[,time_varname] == output_times[time_point_counter]))
      my_table[cur_row,"ONTRTNEW"] = on_therapy_matrix_new[time_point_counter,my_patient_ids[patient_counter]]
    }
  }
  # yougan addition ends here
  
  
  
  
  
  
  # Now we drop the first rows before treatment where it is not informative for response:
  initial_index=(which(output_times>0))[1]
  #	output_times = output_times[initial_index:length(output_times)]
  output_matrix = output_matrix[initial_index:length(output_times),]
  output_matrix_score = output_matrix_score[initial_index:length(output_times),]
  output_matrix_bor_score = output_matrix_bor_score[initial_index:length(output_times),]
  on_therapy_matrix = on_therapy_matrix[initial_index:length(output_times),]
  
  output_matrix_score_new = output_matrix_score_new[initial_index:length(output_times),]
  output_matrix_bor_score_new = output_matrix_bor_score_new[initial_index:length(output_times),]
  on_therapy_matrix_new = on_therapy_matrix_new[initial_index:length(output_times),]
  
  output_times = output_times[initial_index:length(output_times)] # Lu: moved to here, because output_times will reduce 1 length after this command line
  
  # Now add the response variables to the input matrix
  if (!("RSCORE" %in% colnames(my_table)))
  {
    my_table[,"RSCORE"]=""
    my_table[,"RSCORE"]=NA
  }
  if (!("BRSCORE" %in% colnames(my_table)))
  {
    my_table[,"BRSCORE"]=""
    my_table[,"BRSCORE"]=NA
  }
  for (patient_counter in seq_along(my_patient_ids))
  {
    for (time_point_counter in seq_along(output_times))
    {
      cur_row = intersect(which(my_table[,patient_id_varname] == my_patient_ids[patient_counter]),which(my_table[,time_varname] == output_times[time_point_counter]))
      my_table[cur_row,"RSCORE"] = output_matrix_score[time_point_counter,my_patient_ids[patient_counter]]
      my_table[cur_row,"BRSCORE"] = output_matrix_bor_score[time_point_counter,my_patient_ids[patient_counter]]
    }
  }
  
  
  # Yougan addition starts
  # now add the new response variables to the input matrix
  if (!("RSCORENEW" %in% colnames(my_table)))
  {
    my_table[,"RSCORENEW"]=""
    my_table[,"RSCORENEW"]=NA
  }
  if (!("BRSCORENEW" %in% colnames(my_table)))
  {
    my_table[,"BRSCORENEW"]=""
    my_table[,"BRSCORENEW"]=NA
  }
  for (patient_counter in seq_along(my_patient_ids))
  {
    for (time_point_counter in seq_along(output_times))
    {
      cur_row = intersect(which(my_table[,patient_id_varname] == my_patient_ids[patient_counter]),which(my_table[,time_varname] == output_times[time_point_counter]))
      my_table[cur_row,"RSCORENEW"] = output_matrix_score_new[time_point_counter,my_patient_ids[patient_counter]]
      my_table[cur_row,"BRSCORENEW"] = output_matrix_bor_score_new[time_point_counter,my_patient_ids[patient_counter]]
    }
  }
  
  # Yougan additional ends
  
  # Yougan comment: I don't know what are these, but will copy the same codes to the new variables
  
  # Lu add cbor/cbornew columns to the integrated table
  my_table = add_CBOR_recist(my_table, ontrt_varname = "ONTRT", patient_id_varname,time_varname,
                             response_varname, nontarget_pd_varnames="PD_UNEQNONTARGET", rscore_varname="RSCORE", cbrscore_varname="CBRSCORE", output_times = output_times)
  my_table = add_CBOR_recist(my_table, ontrt_varname = "ONTRTNEW", patient_id_varname,time_varname,
                             response_varname, nontarget_pd_varnames="PD_UNEQNONTARGET", rscore_varname="RSCORENEW", cbrscore_varname="CBRSCORENEW", output_times = output_times)
  
  my_return_tables = list()
  my_return_tables[["output_matrix"]] = output_matrix
  my_return_tables[["output_matrix_score"]] = output_matrix_score
  my_return_tables[["output_matrix_bor_score"]] = output_matrix_bor_score
  my_return_tables[["on_therapy_matrix"]] = on_therapy_matrix	
  
  my_return_tables[["output_matrix_score_new"]] = output_matrix_score_new
  my_return_tables[["output_matrix_bor_score_new"]] = output_matrix_bor_score_new
  my_return_tables[["on_therapy_matrix_new"]] = on_therapy_matrix_new
  
  my_return_tables[["integrated_patient_data"]] = my_table
  my_return_tables
}


add_CBOR_recist = function(my_table, ontrt_varname = "ONTRT", patient_id_varname="USUBJID",time_varname="TIME", response_varname = "INDEX_LESION_SLD_RELCH", nontarget_pd_varnames="PD_UNEQNONTARGET", rscore_varname="RSCORE", cbrscore_varname = "CBRSCORE", output_times = output_times)
  # A function to add confirmed BRSCOREs to the integrated_patient_data table, based on RECIST 1.1 guideline. 
  # reference: RECIST 1.1 guideline, Table 3 - Best overall response when confirmation of CR and PR required.
  # response bins 
  # Arguments:
  #  my_table:   input talbe, e.g. integrated_patient_data table with BRSCORE
  #  ontrt_varname
  #  patient_id_varname:    the variable with the patient ID
  #  time_varname:          the variable with the time
  #  response_varname:      the variable in the file with the lesion response of interest
  #  nontarget_pd_varnames: the variable for nontarget pd, used to filter lesion scan time points
  #  rscore_varname:       the unconfimred rscores: RSCORE or RSCORENEW
  #  cbrscore_varname:      the name for confirmed brscores

# Returns
#  add_CBOR_recist:  updated my_table with the new confirmed brscore column added in

{
  require(plyr)
  my_table[,cbrscore_varname] = my_table[,rscore_varname]
  my_table_sld = my_table[which(!is.na(my_table[,response_varname])),]
  TConfirmation = 28 # 4 weeks to confirm CR, PR
  #  alltimepoints = sort(unique(my_table_sld[,time_varname]))
  #  minSDduration =  alltimepoints[3]-alltimepoints[2]  # take this as minimum criterion for SD duration
  minSDduration = 42 # 6 weeks to confirm SD # based on general RECIST guideline:6-8weeks. Ca099003: 35 days, Ca209568: 6 weeks
  
  # Now walk through the patient IDs and output the data
  my_patient_ids = sort(unique(my_table[,patient_id_varname]))
  n_patients = length(my_patient_ids)
  for (i in 1:n_patients){
    patient_id = my_patient_ids[i]
    dat_patient = my_table[which(my_table[,patient_id_varname] == patient_id),]
    rowindices = which(my_table[,patient_id_varname] == patient_id)
    sldrowindices = which(!is.na(dat_patient[,nontarget_pd_varnames])) # used to filter lesion scan time points # sometimes there is no index lesion measurements recorded at the scan date, so cannot use response_varname to filter
    dat_patient_sld = dat_patient[sldrowindices,]
    
    # dat_patient_sld = dat_patient[which(!is.na(dat_patient[,response_varname])),]
    # output_times = unique(c(0,output_times)) # put time 0 back
    # sldrowindices = which(dat_patient[,time_varname] %in% output_times)
    # dat_patient_sld = dat_patient[which(dat_patient[,time_varname] %in% output_times),]
    if (nrow(dat_patient_sld)>=2){
      timepoints = dat_patient_sld[,time_varname]
      for (j in 2:length(timepoints)){  # start from the second timepoint
        if (!is.na(dat_patient_sld[j,rscore_varname]) && dat_patient_sld[j,rscore_varname]=="CR"){
          if ((j+1) <= length(timepoints)){
            if (!is.na(dat_patient_sld[j+1,rscore_varname]) && dat_patient_sld[j+1,rscore_varname] == "PR"){
              if (dat_patient_sld[j,time_varname] >= minSDduration){
                dat_patient_sld[j,cbrscore_varname] = "SD"
              }else{
                dat_patient_sld[j,cbrscore_varname] = "PD"  # assuming through algorithm, we always get true CR
              }
            }else if (!is.na(dat_patient_sld[j+1,rscore_varname]) && dat_patient_sld[j+1,rscore_varname] %in% c("SD","PD")){
              if (dat_patient_sld[j,time_varname] >= minSDduration){
                dat_patient_sld[j,cbrscore_varname] = "SD"
              }else{
                dat_patient_sld[j,cbrscore_varname] = "PD"
              }
            }else if (!is.na(dat_patient_sld[j+1,rscore_varname]) && dat_patient_sld[j+1,rscore_varname] %in% c("PD2")){
              if (dat_patient_sld[j,time_varname] >= minSDduration){
                dat_patient_sld[j,cbrscore_varname] = "SD"
              }else{
                dat_patient_sld[j,cbrscore_varname] = "PD2"
              }
            }else if (is.na(dat_patient_sld[j+1,rscore_varname]) | dat_patient_sld[j+1,rscore_varname] %in% c("OS","NE")){
              if (dat_patient_sld[j,time_varname] >= minSDduration){
                dat_patient_sld[j,cbrscore_varname] = "SD"
              }else{
                dat_patient_sld[j,cbrscore_varname] = NA
              }
            }else if (!is.na(dat_patient_sld[j+1,rscore_varname]) && dat_patient_sld[j+1,rscore_varname] == "CR"){
              if (timepoints[j+1]-timepoints[j] >= TConfirmation){
                dat_patient_sld[j,cbrscore_varname] = "CR"
              }else if ((j+2) <= length(timepoints)){
                if (!is.na(dat_patient_sld[j+2,rscore_varname]) && dat_patient_sld[j+2,rscore_varname] == "PR"){
                  if (dat_patient_sld[j,time_varname] >= minSDduration){
                    dat_patient_sld[j,cbrscore_varname] = "SD"
                  }else{
                    dat_patient_sld[j,cbrscore_varname] = "PD"  # assuming through algorithm, we always get true CR
                  }
                }else if (!is.na(dat_patient_sld[j+2,rscore_varname]) && dat_patient_sld[j+2,rscore_varname] %in% c("SD","PD")){
                  if (dat_patient_sld[j,time_varname] >= minSDduration){
                    dat_patient_sld[j,cbrscore_varname] = "SD"
                  }else{
                    dat_patient_sld[j,cbrscore_varname] = "PD"
                  }
                }else if (!is.na(dat_patient_sld[j+2,rscore_varname]) && dat_patient_sld[j+2,rscore_varname] %in% c("PD2")){
                  if (dat_patient_sld[j,time_varname] >= minSDduration){
                    dat_patient_sld[j,cbrscore_varname] = "SD"
                  }else{
                    dat_patient_sld[j,cbrscore_varname] = "PD2"
                  }
                }else if (is.na(dat_patient_sld[j+2,rscore_varname]) | dat_patient_sld[j+2,rscore_varname] %in% c("OS","NE")){
                  if (dat_patient_sld[j,time_varname] >= minSDduration){
                    dat_patient_sld[j,cbrscore_varname] = "SD"
                  }else{
                    dat_patient_sld[j,cbrscore_varname] = NA
                  }
                }else if (!is.na(dat_patient_sld[j+2,rscore_varname]) && dat_patient_sld[j+2,rscore_varname] == "CR"){
                  if (timepoints[j+2]-timepoints[j] >= TConfirmation){
                    dat_patient_sld[j,cbrscore_varname] = "CR"
                  }else { # simpify ... up to 2 subsequent time point scan checked
                    if (dat_patient_sld[j,time_varname] >= minSDduration){
                      dat_patient_sld[j,cbrscore_varname] = "SD"
                    }else{
                      dat_patient_sld[j,cbrscore_varname] = NA
                    }
                  }
                }
              }
            }
          }else{ # when there is no subsequent timepoint assessment, just take it as the next timepoint is NE
            if (!is.na(dat_patient_sld[j,rscore_varname]) && dat_patient_sld[j,time_varname] >= minSDduration){
              dat_patient_sld[j,cbrscore_varname] = "SD"
            }else{
              dat_patient_sld[j,cbrscore_varname] = NA
            }
          }
        }else if (!is.na(dat_patient_sld[j,rscore_varname]) && dat_patient_sld[j,rscore_varname]=="PR"){
          if ((j+1) <= length(timepoints)){
            if (!is.na(dat_patient_sld[j+1,rscore_varname]) && dat_patient_sld[j+1,rscore_varname] == "SD"){
              dat_patient_sld[j,cbrscore_varname] = "SD"
            }else if (!is.na(dat_patient_sld[j+1,rscore_varname]) && dat_patient_sld[j+1,rscore_varname] %in% c("PD")){
              if (dat_patient_sld[j,time_varname] >= minSDduration){
                dat_patient_sld[j,cbrscore_varname] = "SD"
              }else{
                dat_patient_sld[j,cbrscore_varname] = "PD"
              }
            }else if (!is.na(dat_patient_sld[j+1,rscore_varname]) && dat_patient_sld[j+1,rscore_varname] %in% c("PD2")){
              if (dat_patient_sld[j,time_varname] >= minSDduration){
                dat_patient_sld[j,cbrscore_varname] = "SD"
              }else{
                dat_patient_sld[j,cbrscore_varname] = "PD2"
              }
            }else if (is.na(dat_patient_sld[j+1,rscore_varname]) | dat_patient_sld[j+1,rscore_varname] %in% c("OS","NE")){
              if ((j+2) <= length(timepoints) && !is.na(dat_patient_sld[j+2,rscore_varname]) && dat_patient_sld[j+2,rscore_varname]=="PR"){
                dat_patient_sld[j,cbrscore_varname] = "PR"  # PR-NE-PR is considered as confirmed PR (recist 1.1)
              }else if (dat_patient_sld[j,time_varname] >= minSDduration){
                dat_patient_sld[j,cbrscore_varname] = "SD"
              }else{
                dat_patient_sld[j,cbrscore_varname] = NA
              }
            }else if (!is.na(dat_patient_sld[j+1,rscore_varname]) && dat_patient_sld[j+1,rscore_varname] %in% c("CR","PR")){
              if (timepoints[j+1]-timepoints[j] >= TConfirmation){
                dat_patient_sld[j,cbrscore_varname] = "PR"
              }else if ((j+2) <= length(timepoints)){
                if (!is.na(dat_patient_sld[j+2,rscore_varname]) && dat_patient_sld[j+2,rscore_varname] == "SD"){
                  dat_patient_sld[j,cbrscore_varname] = "SD"
                }else if (!is.na(dat_patient_sld[j+2,rscore_varname]) && dat_patient_sld[j+2,rscore_varname] %in% c("PD")){
                  if (dat_patient_sld[j,time_varname] >= minSDduration){
                    dat_patient_sld[j,cbrscore_varname] = "SD"
                  }else{
                    dat_patient_sld[j,cbrscore_varname] = "PD"
                  }
                }else if (!is.na(dat_patient_sld[j+2,rscore_varname]) && dat_patient_sld[j+2,rscore_varname] %in% c("PD2")){
                  if (dat_patient_sld[j,time_varname] >= minSDduration){
                    dat_patient_sld[j,cbrscore_varname] = "SD"
                  }else{
                    dat_patient_sld[j,cbrscore_varname] = "PD2"
                  }
                }else if (is.na(dat_patient_sld[j+2,rscore_varname]) | dat_patient_sld[j+2,rscore_varname] %in% c("OS","NE")){
                  if ((j+2) <= length(timepoints) && !is.na(dat_patient_sld[j+2,rscore_varname]) && dat_patient_sld[j+2,rscore_varname]=="PR"){
                    dat_patient_sld[j,cbrscore_varname] = "PR"  # PR-NE-PR is considered as confirmed PR (recist 1.1)
                  }else if (dat_patient_sld[j,time_varname] >= minSDduration){
                    dat_patient_sld[j,cbrscore_varname] = "SD"
                  }else{
                    dat_patient_sld[j,cbrscore_varname] = NA
                  }
                }else if (!is.na(dat_patient_sld[j+2,rscore_varname]) && dat_patient_sld[j+2,rscore_varname] %in% c("CR","PR")){
                  if (timepoints[j+2]-timepoints[j] >= TConfirmation){
                    dat_patient_sld[j,cbrscore_varname] = "PR"
                  }else{  # simpify ... up to 2 subsequent time point scan checked
                    if (!is.na(dat_patient_sld[j,rscore_varname]) && dat_patient_sld[j,time_varname] >= minSDduration){
                      dat_patient_sld[j,cbrscore_varname] = "SD"
                    }else{
                      dat_patient_sld[j,cbrscore_varname] = NA
                    }
                  }
                }
              }else{ # when there is no subsequent timepoint assessment, just take it as the next timepoint is NE
                if (!is.na(dat_patient_sld[j,rscore_varname]) && dat_patient_sld[j,time_varname] >= minSDduration){
                  dat_patient_sld[j,cbrscore_varname] = "SD"
                }else{
                  dat_patient_sld[j,cbrscore_varname] = NA
                }
              }
            }
          }else{ # when there is no subsequent timepoint assessment, just take it as the next timepoint is NE
            if (!is.na(dat_patient_sld[j,rscore_varname]) && dat_patient_sld[j,time_varname] >= minSDduration){
              dat_patient_sld[j,cbrscore_varname] = "SD"
            }else{
              dat_patient_sld[j,cbrscore_varname] = NA
            }
          }
        }else if (!is.na(dat_patient_sld[j,rscore_varname]) && dat_patient_sld[j,rscore_varname]=="SD"){
          if (dat_patient_sld[j,time_varname] >= minSDduration){
            dat_patient_sld[j,cbrscore_varname] = "SD"
          }else{
            dat_patient_sld[j,cbrscore_varname] = NA
          }
        }else if (is.na(dat_patient_sld[j,rscore_varname])){
          dat_patient_sld[j,cbrscore_varname] = NA
        }
      }
    }
    
    if (nrow(dat_patient_sld)>=2){
      dat_patient_sld[,cbrscore_varname] <- mapvalues(dat_patient_sld[,cbrscore_varname],
                                                      from=c("CR","PR","SD","PD","PD2","OS",NA,"."),
                                                      to=c(0,1,2,3,4,5,6,7), warn_missing = FALSE)
      dat_patient_sld[,cbrscore_varname] = cummin(as.numeric(dat_patient_sld[,cbrscore_varname]))
      
      dat_patient_sld[,cbrscore_varname] <- mapvalues(dat_patient_sld[,cbrscore_varname],
                                                      from=c(0,1,2,3,4,5,6,7),
                                                      to=c("CR","PR","SD","PD","PD2","OS",NA,"."),
                                                      warn_missing = FALSE)
    }
    
    my_table[rowindices[sldrowindices],cbrscore_varname] = dat_patient_sld[,cbrscore_varname]
  }
  my_table[which(my_table[,cbrscore_varname]=="."),cbrscore_varname] = NA
  my_table[which(my_table[,cbrscore_varname]=="OS"),cbrscore_varname] = NA
  add_CBOR_recist = my_table
}
```
