## Supplementary File 3 for "A surrogate workflow for virtual population development: designing an amenable proxy for a hypothesis-test inspired nonlinear goodness-of-fit": plotAcrossVPOptions.m

```
classdef plotAcrossVPOptions
% Here, we define the plotAcrossVPOptions class 
%
%  Properties
%      flagSave:        Boolean (true/false) indicating whether to save the
%                       figure.  Default is false.
%      fileName:        String indicating the save filename, '.tif' will be
%                       appended.  Leave as '' for a default name if saving.
%      flagLegend:      Boolean (true/false) indicating whether to add a
%                       legend. Default is false.
%      scale:           String indicating X/Y scale, valid strings are:
%                       'xlinylin' (default)
%                       'xlinylog'
%                       'xlogylog'
%                       'xlogylin'
%      xLim:            1x2 numeric matrix of X plotting limits.  Leave as
%                       [] for MATLAB's default.
%      yLim:            1x2 numeric matrix of Y plotting limits.  Leave as
%                       [] for MATLAB's default.
%      xShiftSim:       Double, x-value (time) to shift 
%                       simulation results by. (Optional), leave as 0
%                       so as not to shift.
%      xShiftData:      Double, x-value (time) to shift 
%                       data by. (Optional), leave as 0
%                       so as not to shift.
%      xLabelPretty:    String, will be applied to xlabel. Optional,
%                       if left as '' a default label will be used.
%      yLabelPretty:    String, will be applied to ylabel. Optional,
%                       if left as '' a default label will be used.
%      interventionsIDs:           Cell array of strings, interventions IDs
%                       from the worksheet to
%                       include.  (Optional) If {} or plotting a population
%                       all VPs are included.
%      varName:         String, variable name from the results to plot.  
%      vpID:            String, vpID from the worksheet to plot.
%   
   properties
      flagSave
      fileName
      flagLegend
      scale
      xLim
      yLim
      xShiftSim
      xShiftData
      xLabelPretty
      yLabelPretty
      interventionIDs
      varName
      vpID 
   end
   methods
      function obj = set.flagSave(obj,flagvalue)
          if (flagvalue == true) || (flagvalue == false)
              obj.flagSave = flagvalue;
          else
            error('Invalid flagSave value, expecting true/false')
          end
      end
      function obj = set.fileName(obj,myFileName)
          if (ischar(myFileName) == true) 
              obj.fileName = myFileName;
          else
              error('Invalid fileName value, expecting a string')
          end
      end           
      function obj = set.flagLegend(obj,flagvalue)
          if (flagvalue == true) || (flagvalue == false)
              obj.flagLegend = flagvalue;
          else
            error('Invalid flagLegend value, expecting true/false')
          end
      end         
      function obj = set.scale(obj,scalevalue)
          if (strcmpi(scalevalue,'xlinylin') ||...
               strcmpi(scalevalue,'xlinylog') ||...
               strcmpi(scalevalue,'xlogylog') || ...
               strcmpi(scalevalue,'xlogylin'))
            obj.scale = scalevalue;
          else
            error('Invalid scale setting, allowed setting: "xlinylin," "xlinylog," "xlogylog," "xlogylin."')
          end
      end       
      function obj = set.xLim(obj,xLimVal)
          if ((isnumeric(xLimVal) == true) &&...
                  ((length(xLimVal) == 0) || (length(xLimVal) == 2)))
              obj.xLim = xLimVal;
          else
            error('Invalid xLim value, expecting a 1x2 numeric matrix')
          end
      end 
      function obj = set.yLim(obj,ylim)
          if ((isnumeric(ylim) == true) &&...
                  ((length(ylim) == 0) || (length(ylim) == 2)))
              obj.yLim = ylim;
          else
            error('Invalid yLim value, expecting a 1x2 numeric matrix')
          end
      end 
      function obj = set.xShiftSim(obj,xShiftSimVal)
          if (isnumeric(xShiftSimVal) == true) 
              obj.xShiftSim = xShiftSimVal;
          else
            error('Invalid xShiftSim value, expecting a numeric value')
          end
      end 
      function obj = set.xShiftData(obj,xShiftDataVal)
          if (isnumeric(xShiftDataVal) == true) 
              obj.xShiftData = xShiftDataVal;
          else
            error('Invalid xShiftData value, expecting a numeric value')
          end
      end 
      function obj = set.xLabelPretty(obj,xLabelPrettyVal)
          if (ischar(xLabelPrettyVal) == true) 
              obj.xLabelPretty = xLabelPrettyVal;
          else
            error('Invalid xLabelPretty value, expecting a string')
          end
      end       
      function obj = set.yLabelPretty(obj,yLabelPrettyVal)
          if (ischar(yLabelPrettyVal) == true) || (iscell(yLabelPrettyVal) == true)
              obj.yLabelPretty = yLabelPrettyVal;
          else
            error('Invalid yLabelPretty value, expecting a string (or cell array of strings for multiple lines).')
          end
      end     
      function obj = set.interventionIDs(obj,myInterventionIDs)
          if (iscell(myInterventionIDs) == true) 
              obj.interventionIDs = myInterventionIDs;
          else
            error('Invalid myInterventionIDs value, expecting a cell array')
          end
      end     
      function obj = set.varName(obj,myVarName)
          if (ischar(myVarName) == true) 
              obj.varName = myVarName;
          else
            error('Invalid myVarName value, expecting a string')
          end
      end 
      function obj = set.vpID(obj,myVPID)
          if (ischar(myVPID) == true) 
              obj.vpID = myVPID;
          else
            error('Invalid myVPID value, expecting a string')
          end
      end       
      
      function passCheck = verify(obj,myWorksheet)
          passCheck = true;
          passCheck = obj.verifyVPID(myWorksheet);
          passInterventions = obj.verifyInterventionIDs(myWorksheet);
          passCheck = passCheck && passInterventions;
          [nInterventionResults, nVPResults] = size(myWorksheet.results);
          myVPIDs = getVPIDs(myWorksheet);
          myInterventionIDs = obj.interventionIDs
          if passInterventions
              if nInterventionResults > 0
                  myIndex = find(ismember(myVPIDs,obj.vpID));
                  myResultClasses = cellfun(@class,myWorksheet.results(:,myIndex), 'UniformOutput', false);
                  nSimResults = sum(strcmp(myResultClasses,'struct'));
                  if nSimResults == nInterventionResults;
                      passCheck = passCheck && true;
                  else
                      warning(['Simulation results not fully populated for specified VP in worksheet provided to ',mfilename,'.'])
                      passCheck = false;
                  end
              else
                  warning(['Simulation results not fully populated for specified VP in worksheet provided to ',mfilename,'.'])
                  passCheck = false;
              end
          end
      end
          
      function passCheck = verifyInterventionIDs(obj,myWorksheet)  
          passCheck = true;
          interventionIDs = obj.interventionIDs;
          % Check whether the desired Interventions exists
          allInterventionIDs = getInterventionIDs(myWorksheet);
          keepIndices = find(ismember(allInterventionIDs,interventionIDs));
          interventionIDs = allInterventionIDs(keepIndices);
          if length(keepIndices) < 1
              warning(['Specified interventionIDs not in worksheet provided to ',mfilename,'.'])
              passCheck = false;
          end
      end
          
      function passCheck = verifyVPID(obj,myWorksheet)  
          passCheck = true;      
          vpID = obj.vpID;
          vpIDs = getVPIDs(myWorksheet);
          if ischar(vpID)
              if sum(ismember(vpIDs,vpID)) == 0
                  passCheck = false;
                  warning(['Specified vpID not in worksheet provided to ',mfilename,'.'])
              elseif sum(ismember(vpIDs,vpID)) > 1
                  passCheck = false;
                  warning(['Specified vpID is degenerate in worksheet provided to ',mfilename,'.'])
              end
          else
              passCheck = false;
              warning(['No vpID in options provided to ',mfilename,'.'])
          end
      end
          
      
      function value = get(obj,propName)
          switch propName
              case 'flagSave'
                  value = obj.flagSave;
              case 'fileName'
                  value = obj.fileName;                  
              case 'flagLegend'
                  value = obj.flagLegend;
              case 'scale'
                  value = obj.scale;                  
              %case 'axesCor'
              %    value = obj.axesCor;
              case 'xLim'
                  value = obj.xLim;
              case 'yLim'
                  value = obj.yLim;
              case 'xShiftSim'
                  value = obj.xShiftSim;
              case 'xLabelPretty'
                  value = obj.xLabelPretty;
              case 'yLabelPretty'
                  value = obj.yLabelPretty;
              case 'interventionIDs'
                  value = obj.interventionIDs;
              case 'varName'
                  value = obj.varName;  
              case 'vpID'
                  value = obj.vpID;                   
              otherwise
                  error(['Error: ',propName ,' is not a valid ',mfilename,' property.'])
          end
      end      
      
      % The constructor method must have the same name as the class
      function obj = plotOptions()
            obj.flagSave = false;
            obj.fileName = '';            
            obj.flagLegend = false;
            obj.scale = 'xlinylin';
            obj.xLim = [];
            obj.yLim = [];
            obj.xShiftSim = 0;
            obj.xShiftData = 0;
            obj.xLabelPretty = '';
            obj.yLabelPretty = '';   
            obj.interventionIDs = {};
            obj.varName = '';
            obj.vpID = '';
      end
   end
end
```
