## Supplementary File 3 for "A surrogate workflow for virtual population development: designing an amenable proxy for a hypothesis-test inspired nonlinear goodness-of-fit": plotCoefficientsOptions.m

```
classdef plotCoefficientsOptions
% Here, we define the plotCoefficientsOptions class to help quickly make
% plots of worksheet coefficients
%
% Properties:
%      flagSave      boolean (true/false), whether to save the plot.
%                    Default is false.
%      fileName      String indicating the save filename, '.tif' will be
%                    appended.  Leave as '' for a default name if saving.
%      leftPosition  a value (generally between 0 and 1)
%                    to specify how far to
%                    move the left margin of the figure
%      width         a value (generally between zero and 1)
%                    to specify the relative
%                    width of the figure
%      xLim          a 1x2 array of upper/lower coefficient limits for all
%                    axes on the plot
%      fontSize      Text, tick label size.  Default is 14.
%      flagSort      Boolean, whether to sort axes by covered range.
%                    Default is false.
%
   properties
      flagSave
      fileName
      leftPosition
      width
      xLim
      fontSize
      flagSort
   end
   methods
      function obj = set.flagSave(obj,flagvalue)
          if (flagvalue == true) || (flagvalue == false)
              obj.flagSave = flagvalue;
          else
            error('Invalid flagSave value, expecting true/false')
          end
      end
      function obj = set.fileName(obj,myFileName)
          if (ischar(myFileName) == true) 
              obj.fileName = myFileName;
          else
              error('Invalid fileName value, expecting a string')
          end
      end
      
      function obj = set.leftPosition(obj, myLeftPosition)
          if (isnumeric(myLeftPosition) == true)
              obj.leftPosition = myLeftPosition;
          else
            error('Invalid leftPosition value')
          end
      end
      
      function obj = set.width(obj, myWidth)
          if ((isnumeric(myWidth) == true) && (myWidth >= 0))
              obj.width = myWidth;
          else
            warning(['Invalid width value in ',mfilename,'.'])
          end
      end      
      function obj = set.flagSort(obj,flagvalue)
          if (flagvalue == true) || (flagvalue == false)
              obj.flagSort = flagvalue;
          else
            error('Invalid flagSort value, expecting true/false')
          end
      end      
      
      function obj = set.xLim(obj, myXLim)
            if isnumeric(myXLim)
                if (isequal(size(myXLim), [1 2])) && isequal(~isnan(myXLim), [1 1])
                    if ~isequal(max(myXLim),min(myXLim))
                        obj.xLim =  [min(myXLim), max(myXLim)];
                    else
                        error(['Unable to assign ',myXLim,' to xLim property in ',mfilename,'.  Expecting a 1x2 numeric matrix with unique, non-NaN values.'])    
                    end
                else
                    error(['Unable to assign ',myXLim,' to xLim property in ',mfilename,'.  Expecting a 1x2 numeric matrix with unique, non-NaN values.'])
                end
            else
                error(['Unable to assign ',myXLim,' to xLim property in ',mfilename,'.  Expecting a 1x2 numeric matrix with unique, non-NaN values.'])
            end
        end       
      
      function obj = set.fontSize(obj,myFontSize)
          if ((isnumeric(myFontSize) == true) && (myFontSize >= 0))
              obj.fontSize = myFontSize;
          else
            error(['Invalid fontSize value in ',mfilename,'.'])
          end
      end        

      
      function value = get(obj,propName)
          switch propName
              case 'flagSave'
                  value = obj.flagSave;
              case 'fileName'
                  value = obj.fileName;                  
              case 'leftPosition'
                  value = obj.leftPosition;
              case 'width'
                  value = obj.width;                  
              case 'xLim'
                  value = obj.xLim;
              case 'fontSize'
                  value = obj.fontSize;
              case 'flagSort'
                  value = obj.flagSort;                  
          end
      end      
      
      % The constructor method must have the same name as the class
      function obj = plotCoefficientsOptions()
        obj.flagSave = false;
        obj.fileName = '';
        obj.leftPosition = 0.3;
        obj.width = 0.65;
        obj.xLim = [0 1];
        obj.fontSize = 14;
        obj.flagSort = false;
      end
   end
end
```
