## Supplementary File 3 for "A surrogate workflow for virtual population development: designing an amenable proxy for a hypothesis-test inspired nonlinear goodness-of-fit": plotOptions.m

```
classdef plotOptions
% Here, we define the plotOptions class to help quickly make acceptable
% plots of results
%
%  Properties
%      flagPlotUnweighted Boolean (true/false) indicating whether to show 
%                       unweighted.  Default is false.
%      flagSave:        Boolean (true/false) indicating whether to save the
%                       figure.  Default is false.
%      fileName:        String indicating the save filename, '.tif' will be
%                       appended.  Leave as '' for a default name if saving.
%      flagLegend:      Boolean (true/false) indicating whether to add a
%                       legend. Default is false.
%      flagPD2:         Boolean (true/false) indicating whether to include
%                       PD2 at first lesion scan (upto last time point) in plotBRVPop. PD2 estimated from regression model.
%      flagPD2Exp:  Boolean (true/false) indicating whether to include
%                       PD2 at first lesion scan (upto last time point) in plotBRVPop. PD2 directly based on expPD2
%      scale:           String indicating X/Y scale, valid strings are:
%                       'xlinylin' (default)
%                       'xlinylog'
%                       'xlogylog'
%                       'xlogylin'
%      xLim:            1x2 numeric matrix of X plotting limits.  Leave as
%                       [] for MATLAB's default.
%      yLim:            1x2 numeric matrix of Y plotting limits.  Leave as
%                       [] for MATLAB's default.
%      xShiftSim:       Double, x-value (time) to shift 
%                       simulation results by. (Optional), leave as 0
%                       so as not to shift.
%      xShiftData:      Double, x-value (time) to shift 
%                       data by. (Optional), leave as 0
%                       so as not to shift.
%      xScale:          Double, x-value (time) to scale 
%                       by. (Optional), leave as 1
%                       so as not to scale.
%      yScale:          Double, y-value (time) to scale 
%                       by. (Optional), leave as 1
%                       so as not to scale.
%      xLabelPretty:    String, will be applied to xlabel. Optional,
%                       if left as '' a default label will be used.
%      yLabelPretty:    String, will be applied to ylabel. Optional,
%                       if left as '' a default label will be used.
%      expDataID:       String, experimental dataset ID to plot from the
%                       worksheet.  (Optional) Leave as '' so as not 
%                       to include experimental data.
%      expDataTimeVar:  String for the time variable in the experimental 
%                       dataset, if data is included. 
%      expDataYVar:     String for the Y variable in the experimental 
%                       dataset, if data is included.
%      vpIDs:           Cell array of strings, VP IDs from the worksheet to
%                       include.  (Optional) If {} or plotting a population
%                       all VPs are included.
%      varName:         String, variable name from the results to plot.  
%      interventionID:  String, intervention ID from the worksheet to plot.
%   
   properties
      flagPlotUnweighted
      flagSave
      fileName
      flagLegend
      flagPD2
      flagPD2Exp
      scale
      xLim
      yLim
      xShiftSim
      xShiftData
      xScale  
      yScale
      xLabelPretty
      yLabelPretty
      expDataID
      expDataTimeVar
      expDataYVar
      vpIDs
      varName
      interventionID 
   end
   methods
      function obj = set.flagPlotUnweighted(obj,flagvalue)
          if (flagvalue == true) || (flagvalue == false)
              obj.flagPlotUnweighted = flagvalue;
          else
            error('Invalid flagPlotUnweighted value, expecting true/false')
          end
      end       
      function obj = set.flagSave(obj,flagvalue)
          if (flagvalue == true) || (flagvalue == false)
              obj.flagSave = flagvalue;
          else
            error('Invalid flagSave value, expecting true/false')
          end
      end
      function obj = set.fileName(obj,myFileName)
          if (ischar(myFileName) == true) 
              obj.fileName = myFileName;
          else
              error('Invalid fileName value, expecting a string')
          end
      end           
      function obj = set.flagLegend(obj,flagvalue)
          if (flagvalue == true) || (flagvalue == false)
              obj.flagLegend = flagvalue;
          else
            error('Invalid flagLegend value, expecting true/false')
          end
      end     
      function obj = set.flagPD2(obj,flagvalue)
          if (flagvalue == true) || (flagvalue == false)
              obj.flagPD2 = flagvalue;
          else
            error('Invalid flagPD2 value, expecting true/false')
          end
      end
      function obj = set.flagPD2Exp(obj,flagvalue)
          if (flagvalue == true) || (flagvalue == false)
              obj.flagPD2Exp = flagvalue;
          else
            error('Invalid flagPD2Exp value, expecting true/false')
          end
      end
      function obj = set.scale(obj,scalevalue)
          if (strcmpi(scalevalue,'xlinylin') ||...
               strcmpi(scalevalue,'xlinylog') ||...
               strcmpi(scalevalue,'xlogylog') || ...
               strcmpi(scalevalue,'xlogylin'))
            obj.scale = scalevalue;
          else
            error('Invalid scale setting, allowed setting: "xlinylin," "xlinylog," "xlogylog," "xlogylin."')
          end
      end       
      function obj = set.xLim(obj,xLimVal)
          if ((isnumeric(xLimVal) == true) &&...
                  ((length(xLimVal) == 0) || (length(xLimVal) == 2)))
              obj.xLim = xLimVal;
          else
            error('Invalid xLim value, expecting a 1x2 numeric matrix')
          end
      end 
      function obj = set.yLim(obj,ylim)
          if ((isnumeric(ylim) == true) &&...
                  ((length(ylim) == 0) || (length(ylim) == 2)))
              obj.yLim = ylim;
          else
            error('Invalid yLim value, expecting a 1x2 numeric matrix')
          end
      end 
      function obj = set.xShiftSim(obj,xShiftSimVal)
          if (isnumeric(xShiftSimVal) == true) 
              obj.xShiftSim = xShiftSimVal;
          else
            error('Invalid xShiftSim value, expecting a numeric value')
          end
      end 
      function obj = set.xShiftData(obj,xShiftDataVal)
          if (isnumeric(xShiftDataVal) == true) 
              obj.xShiftData = xShiftDataVal;
          else
            error('Invalid xShiftData value, expecting a numeric value')
          end
      end 
      function obj = set.xScale(obj,xScaleVal)
          if (isnumeric(xScaleVal) == true) 
              obj.xScale = xScaleVal;
          else
            error('Invalid xScaleVal value, expecting a numeric value')
          end
      end  
      function obj = set.yScale(obj,yScaleVal)
          if (isnumeric(yScaleVal) == true) 
              obj.yScale = yScaleVal;
          else
            error('Invalid yScaleVal value, expecting a numeric value')
          end
      end 
      function obj = set.xLabelPretty(obj,xLabelPrettyVal)
          if (ischar(xLabelPrettyVal) == true) 
              obj.xLabelPretty = xLabelPrettyVal;
          else
            error('Invalid xLabelPretty value, expecting a string')
          end
      end       
      function obj = set.yLabelPretty(obj,yLabelPrettyVal)
          if (ischar(yLabelPrettyVal) == true) || (iscell(yLabelPrettyVal) == true)
              obj.yLabelPretty = yLabelPrettyVal;
          else
            error('Invalid yLabelPretty value, expecting a string (or cell array of strings for multiple lines).')
          end
      end
      function obj = set.expDataID(obj,myExpDataID)
          if (ischar(myExpDataID) == true) 
              obj.expDataID = myExpDataID;
          else
            error('Invalid myExpDataID value, expecting a string')
          end
      end      
      function obj = set.expDataTimeVar(obj,myExpDataTimeVar)
          if (ischar(myExpDataTimeVar) == true) 
              obj.expDataTimeVar = myExpDataTimeVar;
          else
            error('Invalid myExpDataTimeVar value, expecting a string')
          end
      end         
      function obj = set.expDataYVar(obj,myExpDataYVar)
          if (ischar(myExpDataYVar) == true) 
              obj.expDataYVar = myExpDataYVar;
          else
            error('Invalid myExpDataTimeVar value, expecting a string')
          end
      end          
      function obj = set.vpIDs(obj,myVPIDs)
          if (iscell(myVPIDs) == true) 
              obj.vpIDs = myVPIDs;
          else
            error('Invalid myVPIDs value, expecting a cell array')
          end
      end     
      function obj = set.varName(obj,myVarName)
          if (ischar(myVarName) == true) 
              obj.varName = myVarName;
          else
            error('Invalid myVarName value, expecting a string')
          end
      end 
      function obj = set.interventionID(obj,myInterventionName)
          if (ischar(myInterventionName) == true) 
              obj.interventionID = myInterventionName;
          else
            error('Invalid myInterventionName value, expecting a string')
          end
      end       
      
      function passCheck = verify(obj,myWorksheet)
          passCheck = true;
          passCheck = obj.verifyVPIDs(myWorksheet);
          passInterventions = obj.verifyInterventionIDs(myWorksheet);
          passCheck = passCheck && passInterventions;
          flagExpData = false;
          if ((length(obj.expDataID) > 0) || (length(obj.expDataTimeVar) > 0) || (length(obj.expDataYVar) > 0))
              flagExpData = true;
          end
          if flagExpData
              passCheck = passCheck && obj.verifyExpData(myWorksheet);
          end
          [nInterventionResults, nVPResults] = size(myWorksheet.results);
          myInterventionIDs = getInterventionIDs(myWorksheet);
          if passInterventions
              if nInterventionResults == length(myInterventionIDs)
                  myIndex = find(ismember(myInterventionIDs,obj.interventionID));
                  myResultClasses = cellfun(@class,myWorksheet.results(myIndex,:), 'UniformOutput', false);
                  nSimResults = sum(strcmp(myResultClasses,'struct'));
                  if nSimResults == nVPResults;
                      passCheck = passCheck && true;
                  else
                      warning(['Simulation results not fully populated for specified intervention in worksheet provided to ',mfilename,'.'])
                      passCheck = false;
                  end
              else
                  warning(['Simulation results not fully populated for specified intervention in worksheet provided to ',mfilename,'.'])
                  passCheck = false;
              end
          end
      end
          
      function passCheck = verifyVPIDs(obj,myWorksheet)  
          passCheck = true;
          vpIDs = obj.vpIDs;
          % Check whether the desired VPs exists
          allVPIDs = getVPIDs(myWorksheet);
          keepIndices = find(ismember(allVPIDs,vpIDs));
          vpIDs = allVPIDs(keepIndices);
          if length(keepIndices) < 1
              warning(['Specified vpIDs not in worksheet provided to ',mfilename,'.'])
              passCheck = false;
          end
      end
          
      function passCheck = verifyInterventionIDs(obj,myWorksheet)  
          passCheck = true;      
          interventionID = obj.interventionID;
          interventionIDs = getInterventionIDs(myWorksheet);
          if sum(ismember(interventionIDs,interventionID)) == 0
              passCheck = false;
              warning(['Specified interventionID not in worksheet provided to ',mfilename,'.'])
          elseif sum(ismember(interventionIDs,interventionID)) > 1
              passCheck = false;
              warning(['Specified interventionID is degenerate in worksheet provided to ',mfilename,'.'])
          end
      end
          
      function passCheck = verifyExpData(obj,myWorksheet)
          passCheck = true;
          expDataID = obj.expDataID;
          allExpIDs = getExpDataIDs(myWorksheet);
          if sum(ismember(allExpIDs, expDataID)) < 1
              passCheck = false;
              warning(['Specified experimental data set ID not present in worksheet, ignoring'])
          elseif sum(ismember(allExpIDs, expDataID)) > 1
              passCheck = false;
              warning(['Specified experimental data set ID degenerate in worksheet, ignoring'])
          end
          if (passCheck)
              expData = getExpData(expDataID, myWorksheet);
              expDataTimeVar = obj.expDataTimeVar;
              if sum(ismember(expData.Properties.VariableNames,expDataTimeVar)) < 1
                  warning(['Specified time variable name not present in data set, ignoring experimental data'])
                  passCheck = false;
              elseif sum(ismember(expData.Properties.VariableNames,expDataTimeVar)) > 1
                  passCheck = false;
                  warning(['Specified time variable name degenerate in worksheet, ignoring experimental data'])
              end
              expDataYVar = obj.expDataYVar;
              if sum(ismember(expData.Properties.VariableNames,expDataYVar)) < 1
                  warning(['Specified dependent variable name not present in data set, ignoring experimental data'])
                  passCheck = false;
              elseif sum(ismember(allExpIDs, expDataYVar)) > 1
                  passCheck = false;
                  warning(['Specified dependent variable name degenerate in worksheet, ignoring experimental data'])
              end
          end
      end
      
      function value = get(obj,propName)
          switch propName
              case 'flagPlotUnweighted'
                  value = obj.flagPlotUnweighted;              
              case 'flagSave'
                  value = obj.flagSave;
              case 'fileName'
                  value = obj.fileName;                  
              case 'flagLegend'
                  value = obj.flagLegend;
              case 'flagPD2'
                  value = obj.flagPD2;    
              case 'flagPD2Exp'
                  value = obj.flagPD2Exp;  
              case 'scale'
                  value = obj.scale;                  
              %case 'axesCor'
              %    value = obj.axesCor;
              case 'xLim'
                  value = obj.xLim;
              case 'yLim'
                  value = obj.yLim;
              case 'xShiftSim'
                  value = obj.xShiftSim;
              case 'xScale'
                  value = obj.xScale;      
              case 'yScale'
                  value = obj.yScale;  
              case 'xLabelPretty'
                  value = obj.xLabelPretty;
              case 'yLabelPretty'
                  value = obj.yLabelPretty;
              case 'expDataID'
                  value = obj.expDataID;
              case 'expDataTimeVar'
                  value = obj.expDataTimeVar;
              case 'expDataYVar'
                  value = obj.expDataYVar;
              case 'vpIDs'
                  value = obj.vpIDs;
              case 'varName'
                  value = obj.varName;  
              case 'interventionID'
                  value = obj.interventionID;                   
              otherwise
                  error(['Error: ',propName ,' is not a valid ',mfilename,' property.'])
          end
      end      
      
      % The constructor method must have the same name as the class
      function obj = plotOptions()
            obj.flagPlotUnweighted = false;          
            obj.flagSave = false;
            obj.fileName = '';            
            obj.flagLegend = false;
            obj.flagPD2 = false;    
            obj.flagPD2Exp = false; 
            obj.scale = 'xlinylin';
            obj.xLim = [];
            obj.yLim = [];
            obj.xShiftSim = 0;
            obj.xShiftData = 0;
            obj.xScale = 1;   
            obj.yScale = 1;  
            obj.xLabelPretty = '';
            obj.yLabelPretty = '';   
            obj.expDataID = '';
            obj.expDataTimeVar = '';
            obj.expDataYVar = '';
            obj.vpIDs = {};
            obj.varName = '';
            obj.interventionID = '';
      end
   end
end
```
