## Supplementary File 3 for "A surrogate workflow for virtual population development: designing an amenable proxy for a hypothesis-test inspired nonlinear goodness-of-fit": plotAcrossIntervention.m

```
function plotHandleVector = plotAcrossIntervention(myWorksheet, myPlotOptions)
% This function plots results for a single variable stored in a worksheet.
% One or more VPs are plotted for a single intervention.
%
% ARGUMENTS
% myWorksheet:      (required) Worksheet with simulation results
% myPlotOptions:    (required) A plotOptions structure to adjust the plotting
%
% RETURNS
% plotHandle
%

% Check number of inputs.
% We define fail behavior as warn rather than error here
% This just seems a more reasonable way to fail if we can generate plots
% rather than interrupting higher level execution with an error.
plotHandleVector = [];
flagExpData = false;

if nargin > 2
    flagContinue = false;
    warning(['Too many input arguments for ',mfilename,', require: myWorksheet, myPlotOptions. Exiting.'])
elseif nargin > 1
    flagContinue = true;
else
    flagContinue = false;
    warning(['Insufficient input arguments for ',mfilename,', require: myWorksheet, myPlotOptions. Exiting.']) 
end

if flagContinue
    if ~strcmp(class(myPlotOptions),'plotOptions')
        flagContinue = false;
        warning(['Invalid plotOptions object for ',mfilename,', exiting.'])
    end           
end

if flagContinue
    if length(myPlotOptions.vpIDs) < 1
        myPlotOptions.vpIDs = getVPIDs(myWorksheet);
    end
end

if flagContinue
    flagContinue = myPlotOptions.verify(myWorksheet);
    if ~flagContinue
        warning(['The plotOptions failed verification for ',mfilename,', exiting.'])
    end
end

if flagContinue
    if ((length(myPlotOptions.expDataID) > 0) || (length(myPlotOptions.expDataTimeVar) > 0) || (length(myPlotOptions.expDataYVar) > 0))
        flagExpData = true;
    end
end

if flagContinue
    vpIDs = myPlotOptions.vpIDs;
    % Check whether the desired VPs exists
    allVPIDs = getVPIDs(myWorksheet);
    keepIndices = find(ismember(allVPIDs,vpIDs));
    interventionID = myPlotOptions.interventionID;    
    varName = myPlotOptions.varName;
end

if flagExpData
    % Filter out nan's, this will cause issues 
    expDataID = myPlotOptions.expDataID;
    expData = getExpData(expDataID, myWorksheet);
    expDataTimeVar = myPlotOptions.expDataTimeVar;
    expDataYVar = myPlotOptions.expDataYVar;
    expDataTime = expData.(expDataTimeVar);
    expDataYVar = expData.(expDataYVar);
    the_indices = find((~isnan(expDataTime)) & (~isnan(expDataYVar)));
    expDataTime = expDataTime(the_indices);
    expDataYVar = expDataYVar(the_indices);
end

% TO CONSIDER:
% May want to scan VPs for the simulation lag parameter and
% update the plot options
xVarName = 'time';   
legendEntries = cell(1,0);
if flagContinue
    % Now we can finally generate the plots!
    figure;
    nVP = length(keepIndices);
    % Reseed the random number generator to get consistent if randomly sequenced plot colors
    rng('default');
    randColors = rand(nVP,3);
    interventionIDs = getInterventionIDs(myWorksheet);
    interventionIndex = find(ismember(interventionIDs,interventionID));
    for vpCounter = 1 : nVP
        vpIndex = keepIndices(vpCounter);
        legendEntries{1,vpCounter} = myWorksheet.vpDef{vpIndex}.('ID');       
        varNames = myWorksheet.results{interventionIndex,vpIndex}.Names;
        timeIndex = find(ismember(varNames,xVarName));
        varIndex = find(ismember(varNames,varName));
        timeVal = myWorksheet.results{interventionIndex,vpIndex}.Data(:,timeIndex);
        varVal = myWorksheet.results{interventionIndex,vpIndex}.Data(:,varIndex);
        hold on
        plotHandleVector(vpCounter) = plot((timeVal-myPlotOptions.xShiftSim)/myPlotOptions.xScale, varVal/myPlotOptions.yScale, 'color', randColors(vpCounter,:), 'LineWidth',4); 
        if (strcmp(myPlotOptions.scale,'xlogylin') || strcmp(myPlotOptions.scale,'xlogylog'))
            set(gca,'xscale','log'); % gca
        end
        if (strcmp(myPlotOptions.scale,'xlogylog') || strcmp(myPlotOptions.scale,'xlinylog'))
            set(gca,'yscale','log'); % gca
        end        
    end
    if flagExpData
        plotHandleVector(nVP+1) = plot((expDataTime-myPlotOptions.xShiftData)/myPlotOptions.xScale, expDataYVar/myPlotOptions.yScale, 'k.','MarkerSize',20); 
        if (strcmp(myPlotOptions.scale,'xlogylin') || strcmp(myPlotOptions.scale,'xlogylog'))
            set(gca,'xscale','log'); % gca
        end
        if (strcmp(myPlotOptions.scale,'xlogylog') || strcmp(myPlotOptions.scale,'xlinylog'))
            set(gca,'yscale','log'); % gca
        end    
        set(get(get(plotHandleVector(nVP+1),'Annotation'),'LegendInformation'),'IconDisplayStyle','off');
    end
    if length(myPlotOptions.yLim) > 0
        ylim(myPlotOptions.yLim);
    end 
    if length(myPlotOptions.xLabelPretty) > 0
        xlabel(myPlotOptions.xLabelPretty,'interpreter','none');
    else
        xlabel(xVarName,'interpreter','none');
    end
    if length(myPlotOptions.yLabelPretty) > 0
        ylabel(myPlotOptions.yLabelPretty,'interpreter','none');
    else
        ylabel(varName,'interpreter','none');
    end    
    if length(myPlotOptions.xLim) > 0
        xlim(myPlotOptions.xLim);
    end    
    if length(myPlotOptions.yLim) > 0
        ylim(myPlotOptions.yLim);
    end     
    grid on
    set(gca,'fontsize', 18);
    if myPlotOptions.flagLegend
        legend(legendEntries,'interpreter','none');
    end
    if myPlotOptions.flagSave
        if length(myPlotOptions.fileName) > 0
            print([myPlotOptions.fileName,'.tif'],'-dtiff','-r300');
        else
            theDate = date;
            formatOut = 'yymmdd';
            theDate = datestr(theDate,formatOut);            
            print([interventionID,'_',varName,'_',theDate,'.tif'],'-dtiff','-r300');
        end
    end
end

end
```
