## Supplementary File 3 for "A surrogate workflow for virtual population development: designing an amenable proxy for a hypothesis-test inspired nonlinear goodness-of-fit": plotAcrossVP.m

```
function plotHandleVector = plotAcrossVP(myWorksheet, myPlotAcrossVPOptions)
% This function plots results for a single variable stored in a worksheet.
% Results for one or more interventions for a single VP are plotted.
%
% ARGUMENTS
% myWorksheet:            (required) Worksheet with simulation results
% myPlotAcrossVPOptions:  (required) A plotAcrossVPOptions structure to adjust the plotting
%
% RETURNS
% plotHandle
%

% Check number of inputs.
% We define fail behavior as warn rather than error here
% This just seems a more reasonable way to fail if we can generate plots
% rather than interrupting higher level execution with an error.
plotHandleVector = [];
flagExpData = false;

if nargin > 2
    flagContinue = false;
    warning(['Too many input arguments for ',mfilename,', require: myWorksheet, myPlotAcrossVPOptions. Exiting.'])
elseif nargin > 1
    flagContinue = true;
else
    flagContinue = false;
    warning(['Insufficient input arguments for ',mfilename,', require: myWorksheet, myPlotAcrossVPOptions. Exiting.']) 
end

if flagContinue
    if ~strcmp(class(myPlotAcrossVPOptions),'plotAcrossVPOptions')
        flagContinue = false;
        warning(['Invalid plotAcrossVPOptions object for ',mfilename,', exiting.'])
    end           
end

if flagContinue
    if length(myPlotAcrossVPOptions.interventionIDs) < 1
        myPlotAcrossVPOptions.interventionIDs = getInterventionIDs(myWorksheet);
    end
end

if flagContinue
    flagContinue = myPlotAcrossVPOptions.verify(myWorksheet);
    if ~flagContinue
        warning(['The plotAcrossVPOptions failed verification for ',mfilename,', exiting.'])
    end
end


if flagContinue
    interventionIDs = myPlotAcrossVPOptions.interventionIDs;
    % Check whether the desired VPs exists
    allInterventionIDs = getInterventionIDs(myWorksheet);
    keepIndices = find(ismember(allInterventionIDs,interventionIDs));
    vpID = myPlotAcrossVPOptions.vpID;    
    varName = myPlotAcrossVPOptions.varName;
    allVPIDs = getVPIDs(myWorksheet);
end


% TO CONSIDER:
% May want to scan VPs for the simulation lag parameter and
% update the plot options
xVarName = 'time';   

if flagContinue
    % Now we can finally generate the plots!
    figure;
    nIntervention = length(keepIndices);
    % Reseed the random number generator to get consistent if randomly sequenced plot colors
    rng('default');
    randColors = rand(nIntervention,3);
    vpIndex = find(ismember(allVPIDs,vpID));
    legendEntries = cell(1,nIntervention);
    for interventionCounter = 1 : nIntervention
        interventionIndex = keepIndices(interventionCounter);
        legendEntries{1,interventionCounter} = myWorksheet.interventions{interventionIndex}.('ID');       
        varNames = myWorksheet.results{interventionIndex,vpIndex}.Names;
        timeIndex = find(ismember(varNames,xVarName));
        varIndex = find(ismember(varNames,varName));
        timeVal = myWorksheet.results{interventionIndex,vpIndex}.Data(:,timeIndex);
        varVal = myWorksheet.results{interventionIndex,vpIndex}.Data(:,varIndex);
        hold on
        plotHandleVector(interventionCounter) = plot(timeVal-myPlotAcrossVPOptions.xShiftSim, varVal, 'color', randColors(interventionCounter,:), 'LineWidth',4); 
        if (strcmp(myPlotAcrossVPOptions.scale,'xlogylin') || strcmp(myPlotAcrossVPOptions.scale,'xlogylog'))
            set(gca,'xscale','log'); % gca
        end
        if (strcmp(myPlotAcrossVPOptions.scale,'xlogylog') || strcmp(myPlotAcrossVPOptions.scale,'xlinylog'))
            set(gca,'yscale','log'); % gca
        end        
    end
    if flagExpData
        plotHandleVector(nVP+1) = plot(expDataTime, expDataYVar, 'k.','MarkerSize',20); 
        if (strcmp(myPlotAcrossVPOptions.scale,'xlogylin') || strcmp(myPlotAcrossVPOptions.scale,'xlogylog'))
            set(gca,'xscale','log'); % gca
        end
        if (strcmp(myPlotAcrossVPOptions.scale,'xlogylog') || strcmp(myPlotAcrossVPOptions.scale,'xlinylog'))
            set(gca,'yscale','log'); % gca
        end    
        set(get(get(plotHandleVector(nVP+1),'Annotation'),'LegendInformation'),'IconDisplayStyle','off');
    end
    if length(myPlotAcrossVPOptions.yLim) > 0
        ylim(myPlotAcrossVPOptions.yLim);
    end 
    if length(myPlotAcrossVPOptions.xLabelPretty) > 0
        xlabel(myPlotAcrossVPOptions.xLabelPretty);
    else
        xlabel(xVarName,'interpreter','none');
    end
    if length(myPlotAcrossVPOptions.yLabelPretty) > 0
        ylabel(myPlotAcrossVPOptions.yLabelPretty);
    else
        ylabel(varName,'interpreter','none');
    end    
    if length(myPlotAcrossVPOptions.xLim) > 0
        xlim(myPlotAcrossVPOptions.xLim);
    end    
    if length(myPlotAcrossVPOptions.yLim) > 0
        ylim(myPlotAcrossVPOptions.yLim);
    end     
    grid on
    set(gca,'fontsize', 18);
    if myPlotAcrossVPOptions.flagLegend
        legend(legendEntries,'interpreter','none');
    end
    if myPlotAcrossVPOptions.flagSave
        if length(myPlotAcrossVPOptions.fileName) > 0
            %saveas(gca,[myPlotAcrossVPOptions.fileName,'.tif']);
            print([myPlotAcrossVPOptions.fileName,'.tif'],'-dtiff','-r300');
        else
            theDate = date;
            formatOut = 'yymmdd';
            theDate = datestr(theDate,formatOut);            
            print([interventionID,'_',varName,'_',theDate,'.tif'],'-dtiff','-r300');
        end
    end
end

end
```
