## Supplementary File 3 for "A surrogate workflow for virtual population development: designing an amenable proxy for a hypothesis-test inspired nonlinear goodness-of-fit": plotBinVPop.m

```
function plotHandle = plotBinVPop(myVPop, myPlotOptions)
% This function takes the bin data in a VPop distribution table
% to plot the experimental data side-by-side with the population
% to facilitate diagnosing fitting during
% the population calibration/prevalence weighting process.
%
% ARGUMENTS
% myVPop:           (required) A VPop, VPopRECIST, or VPopRECISTnoBin.  Note  
%                   that both the experimental and simulation result fields should  
%                   be populated
% myPlotOptions:    (optional) A plotOptions structure.
%                   Note that not all arguments are used.
%
% RETURNS
% plotHandle
%
plotHandle = [];
flagContinue = false;

if nargin > 2
    flagContinue = false;
    warning(['Too many input arguments to ',mfilename,'. Required: myVPop.'])
elseif nargin > 1
    flagContinue = true;    
elseif nargin > 0
    myPlotOptions = plotOptions;
    
    flagContinue = true;
else
    flagContinue = false;
    warning(['Insufficient input arguments to ',mfilename,'. Required: myVPop.'])
end

if flagContinue
    if ~(ismember(class(myVPop), {'VPop','VPopRECIST','VPopRECISTnoBin'}))
        flagContinue = false;
        warning(['Invalid VPop for ',mfilename,'.'])
    end  
    if ~strcmp(class(myPlotOptions),'plotOptions')
        flagContinue = false;
        warning(['Invalid plotOptions for ',mfilename,'.'])
    end        
end

if flagContinue
    if ~(strcmp(class(myVPop.binTable), 'table'))
        flagContinue = false;
        warning(['Invalid VPop.binTable for ',mfilename,'.'])
    end  
end

if (flagContinue)
	myTable = myVPop.binTable;
    [myNPlots, ~] = size(myTable);
    if ~(myPlotOptions.flagPlotUnweighted) && (myNPlots > 0)    
        myTable(myTable.weight==0, :) =[];
        [myNPlots, ~] = size(myTable);
    end
    if myNPlots < 1
        flagContinue = false;
        warning(['Invalid VPop.binTable for ',mfilename,'.'])        
    end
end


if (flagContinue)
    
    
    for rowCounter = 1 : myNPlots
        plotNames{rowCounter} = {myTable{rowCounter,'interventionID'}{1},myTable{rowCounter,'elementID'}{1},num2str(myTable{rowCounter,'time'})};
    end
    
end

if (flagContinue)
    nPlotsVer = floor(sqrt(myNPlots));
    nPlotsHor = ceil(myNPlots / nPlotsVer);
    plotHandle = figure;
    for plotCounter = 1 : myNPlots
        subplot(nPlotsVer, nPlotsHor, plotCounter);
        expBins = myTable{plotCounter,'expBins'};
        predBins = myTable{plotCounter,'predBins'};
        binEdges = myTable{plotCounter,'binEdges'};
        binEdges = binEdges{1};
        expBins = expBins{1};
        predBins = predBins{1};
        binLabels = cell(1, length(binEdges)+1);
        binLabels{1} = ['< ',num2str(binEdges(1))];
        binLabels{length(binEdges)+1} = ['>= ',num2str(binEdges(length(binEdges)))];
        for binCounter = 1 : (length(binEdges)-1)
            binLabels{binCounter+1} = [num2str(binEdges(binCounter)),' - ',num2str(binEdges(binCounter+1))];
        end
        barplot = bar([expBins;predBins]');
        set(gca,'XTickLabel',binLabels);
        xtickangle(45);
        set(barplot(1),'facecolor','w') 
        set(barplot(2),'facecolor',[.2,.2,.2]) 
        legend({'Trials','VPop'})
        title(gca,plotNames{plotCounter},'interpreter','none','FontWeight','Normal');
        set(gca,'box','on');
        set(gca,'fontsize', 10);
    end

    if myPlotOptions.flagSave
        theDate = date;
        formatOut = 'yymmdd';
        theDate = datestr(theDate,formatOut);
        print(['VPopBin_',theDate,'.tif'],'-dtiff','-r300');
    end
end

end
```
