## Supplementary File 3 for "A surrogate workflow for virtual population development: designing an amenable proxy for a hypothesis-test inspired nonlinear goodness-of-fit": plotBiomarker.m

```
function plotHandle = plotBiomarker(myWorksheet, myVPop, Biomarker, Patientgroup)
% This function takes a Worksheet, a VPop and a cell array of Biomarker
% definitions, and returns a plot handle to Whisker plots of the biomarkers
% along with available experimental data. If exactly 2 Biomarkers are
% provided and their ElementIDs are different Biomarker 1 is plotted vs.
% Biomarker 2 along with experimental data if available.
%
% If Patientgroup is provided as a field to Biomarker the 1d Whisker plots 
% are restricted to the respective patient subgroups (CR, PR, SD, PD).
% If Patientgroup is provided as a 4th argument the 2d biomarker plot 
% is restricted to the respective patient subgroups (CR, PR, SD, PD).
% If Patientgroup.Time is left empty ('') Patientgroup data is pulled from
% the last available time point in simData and expData.
%
% ARGUMENTS
% myWorksheet:      (required) A worksheet with results field populated
% myVPop:           (required) A VPop, VPopRECIST, or VPopRECISTnoBin.
%                    myVPop should correspond to myWorksheet.
% Biomarker:        (required) cell array of biomarker definitions where
%                    each entry is a struct with the following fields
%                       .ExpDataID (if empty ('') no data is plotted)
%                       .TrtGrp (defined in myWorksheet.expData)
%                       .ExpVarID (defined in myWorksheet.expData)
%                       .Time (make sure Time exists in expData and simData)
%                       .InterventionID (defined in myWorksheet.interventions)
%                       .ElementID (defined in myWorksheet.simProps.saveElementIDs)
%                       .Percentile ([alpha 1-alpha], only applies to Whisker plots)
%                       .Patientgroup.CR (true or false)
%                       .Patientgroup.PR (true or false)
%                       .Patientgroup.SD (true or false)
%                       .Patientgroup.PD (true or false)
%                       .Patientgroup.Time (day at which RECIST criteria are evaluated)
%
% Patientgroup:     (optional) struct with following fields:
%                   .CR (true or false)
%                   .PR (true or false)
%                   .SD (true or false)
%                   .PD (true or false)
%                   .Time (day at which RECIST criteria are evaluated)
%
% RETURNS
% plotHandle
%

nBiomarkers = length(Biomarker);

PatientgroupFlag = false;
plot1dExpFlag = false(nBiomarkers,1);
plot2dFlag = false;
plot2dExpFlag = false;

if nBiomarkers == 2 && ~strcmp(Biomarker{1}.ElementID, Biomarker{2}.ElementID) ...
        && strcmp(Biomarker{1}.InterventionID, Biomarker{2}.InterventionID)
    % # of biomarkers == 2 AND ElementIDs are different AND InterventionIDs
    % are the same
    plot2dFlag = true;
    
    if nargin > 3
        PatientgroupFlag = true;
        Patientgroup.InterventionID = Biomarker{1}.InterventionID;
        [curWorksheet, curVPop] = getPatientgroup(myWorksheet, Patientgroup, myVPop);
    else
        curWorksheet = myWorksheet;
        curVPop = myVPop;
        
    end
    
    if ~isempty(Biomarker{1}.ExpDataID) && ~isempty(Biomarker{2}.ExpDataID)
        % get experimental results for Biomarker 1 and Biomarker 2
        plot2dExpFlag = true;
        myExpDataIDs = getExpDataIDs(curWorksheet);
        expDataIDBM1 = Biomarker{1}.ExpDataID;
        expDataIDBM2 = Biomarker{2}.ExpDataID;
        expDataBM1Ind = find(ismember(myExpDataIDs, expDataIDBM1));
        expDataBM2Ind = find(ismember(myExpDataIDs, expDataIDBM2));
        
        if ~isempty(Biomarker{1}.TrtGrp)
            candidateRows1 = find((curWorksheet.expData{expDataBM1Ind}.data{:,'TIME'} == Biomarker{1}.Time) ...
            & ismember(curWorksheet.expData{expDataBM1Ind}.data{:,'TRTGRP'},Biomarker{1}.TrtGrp) ...
            & ~isnan(curWorksheet.expData{expDataBM1Ind}.data{:,Biomarker{1}.ExpVarID}));
        else
            candidateRows1 = find((curWorksheet.expData{expDataBM1Ind}.data{:,'TIME'} == Biomarker{1}.Time) ...
            & ~isnan(curWorksheet.expData{expDataBM1Ind}.data{:,Biomarker{1}.ExpVarID}));
        end
        
        % get USUBJIDs for candidateRows1
        candidateUSUBJID1 = ((curWorksheet.expData{expDataBM1Ind}.data{candidateRows1,'USUBJID'}));
        
        % restrict candidateRows1 to requested Patientgroup
        if PatientgroupFlag
            
            if isempty(Patientgroup.Time)
                % pull Patientgroup data from last available Time in
                % BRSCORE Column of expDataTable
                curBRSCORE = cell(length(candidateUSUBJID1),1);
                relSLDRows = zeros(length(candidateUSUBJID1),1);
                relSLDRowInd = false(length(candidateUSUBJID1),1);
                
                for k=1:length(candidateUSUBJID1)
                    
                    if ~isempty(Biomarker{1}.TrtGrp)
                        curSLDRows = find(ismember(curWorksheet.expData{expDataBM1Ind}.data{:,'USUBJID'}, candidateUSUBJID1(k)) ...
                        & ~isnan(curWorksheet.expData{expDataBM1Ind}.data{:,'TIME'}) ...
                        & ismember(curWorksheet.expData{expDataBM1Ind}.data{:,'TRTGRP'},Biomarker{1}.TrtGrp) ...
                        & ~isnan(curWorksheet.expData{expDataBM1Ind}.data{:,'INDEX_LESION_SLD_RELCH'}) ...
                        & cellfun(@ischar,curWorksheet.expData{expDataBM1Ind}.data{:,'BRSCORENEW'}));
                    else
                        curSLDRows = find(ismember(curWorksheet.expData{expDataBM1Ind}.data{:,'USUBJID'}, candidateUSUBJID1(k)) ...
                        & ~isnan(curWorksheet.expData{expDataBM1Ind}.data{:,'TIME'}) ...
                        & ~isnan(curWorksheet.expData{expDataBM1Ind}.data{:,'INDEX_LESION_SLD_RELCH'}) ...
                        & cellfun(@ischar,curWorksheet.expData{expDataBM1Ind}.data{:,'BRSCORENEW'}));
                    end
                    
                    if ~isempty(curSLDRows)
                        curBRSCORE{k,1} = curWorksheet.expData{expDataBM1Ind}.data{curSLDRows,'BRSCORENEW'}{end};
                        relSLDRows(k,1) = curSLDRows(end);
                        relSLDRowInd(k,1) = true;
                    end
                end
                % get rid of empty rows
                curBRSCORE = curBRSCORE(relSLDRowInd);
                relSLDRows = relSLDRows(relSLDRowInd);
                
            else
                % pull Patientgroup data from BRSCORE Column of
                % expDataTable at Time specified in Patientgroup.Time
                
                if ~isempty(Biomarker{1}.TrtGrp)
                    relSLDRows = find(ismember(curWorksheet.expData{expDataBM1Ind}.data{:,'USUBJID'}, candidateUSUBJID1) ...
                    & (curWorksheet.expData{expDataBM1Ind}.data{:,'TIME'} == Patientgroup.Time) ...
                    & ismember(curWorksheet.expData{expDataBM1Ind}.data{:,'TRTGRP'},Biomarker{1}.TrtGrp) ...
                    & ~isnan(curWorksheet.expData{expDataBM1Ind}.data{:,'INDEX_LESION_SLD_RELCH'}) ...
                    & cellfun(@ischar,curWorksheet.expData{expDataBM1Ind}.data{:,'BRSCORENEW'}));
                else
                    relSLDRows = find(ismember(curWorksheet.expData{expDataBM1Ind}.data{:,'USUBJID'}, candidateUSUBJID1) ...
                    & (curWorksheet.expData{expDataBM1Ind}.data{:,'TIME'} == Patientgroup.Time) ...
                    & ~isnan(curWorksheet.expData{expDataBM1Ind}.data{:,'INDEX_LESION_SLD_RELCH'}) ...
                    & cellfun(@ischar,curWorksheet.expData{expDataBM1Ind}.data{:,'BRSCORENEW'}));
                end
                
                curBRSCORE = curWorksheet.expData{expDataBM1Ind}.data{relSLDRows,'BRSCORENEW'};
                
            end
            
            CRInd = false(length(curBRSCORE),1);
            PRInd = false(length(curBRSCORE),1);
            SDInd = false(length(curBRSCORE),1);
            PDInd = false(length(curBRSCORE),1);
            
            if Patientgroup.CR
                CRInd = ismember(curBRSCORE,'CR');
            end
            
            if Patientgroup.PR
                PRInd = ismember(curBRSCORE,'PR');
            end
            
            if Patientgroup.SD
                SDInd = ismember(curBRSCORE,'SD');
            end
            
            if Patientgroup.PD
                PDInd = ismember(curBRSCORE,'PD');
            end
            
            BRInd = logical(CRInd + PRInd + SDInd + PDInd);
            
            % get USUBJIDs for rows in relSLDRows corresponding to the
            % desired Patientgroups
            BRscoreUSUBJID1 = curWorksheet.expData{expDataBM1Ind}.data{relSLDRows(BRInd),'USUBJID'};
            
            if ~isempty(Biomarker{1}.TrtGrp)
                candidateRows1 = find(ismember(curWorksheet.expData{expDataBM1Ind}.data{:,'USUBJID'}, BRscoreUSUBJID1) ...
                & (curWorksheet.expData{expDataBM1Ind}.data{:,'TIME'} == Biomarker{1}.Time) ...
                & ismember(curWorksheet.expData{expDataBM1Ind}.data{:,'TRTGRP'},Biomarker{1}.TrtGrp) ...
                & ~isnan(curWorksheet.expData{expDataBM1Ind}.data{:,Biomarker{1}.ExpVarID}));
            else
                candidateRows1 = find(ismember(curWorksheet.expData{expDataBM1Ind}.data{:,'USUBJID'}, BRscoreUSUBJID1) ...
                & (curWorksheet.expData{expDataBM1Ind}.data{:,'TIME'} == Biomarker{1}.Time) ...
                & ~isnan(curWorksheet.expData{expDataBM1Ind}.data{:,Biomarker{1}.ExpVarID}));
            end
            
        end
        
        % get USUBJIDs for candidateRows1
        candidateUSUBJID1 = ((curWorksheet.expData{expDataBM1Ind}.data{candidateRows1,'USUBJID'}));
        
        % make sure to pick experimental results for the same patient IDs
        if ~isempty(Biomarker{2}.TrtGrp)
            candidateRows2 = find(ismember(curWorksheet.expData{expDataBM2Ind}.data{:,'USUBJID'},candidateUSUBJID1) ...
            & (curWorksheet.expData{expDataBM2Ind}.data{:,'TIME'} == Biomarker{2}.Time) ...
            & ismember(curWorksheet.expData{expDataBM2Ind}.data{:,'TRTGRP'},Biomarker{2}.TrtGrp) ...
            & ~isnan(curWorksheet.expData{expDataBM2Ind}.data{:,Biomarker{2}.ExpVarID}));
        else
            candidateRows2 = find(ismember(curWorksheet.expData{expDataBM2Ind}.data{:,'USUBJID'},candidateUSUBJID1) ...
            & (curWorksheet.expData{expDataBM2Ind}.data{:,'TIME'} == Biomarker{2}.Time) ...
            & ~isnan(curWorksheet.expData{expDataBM2Ind}.data{:,Biomarker{2}.ExpVarID}));
        end
        candidateUSUBJID2 = ((curWorksheet.expData{expDataBM2Ind}.data{candidateRows2,'USUBJID'}));
        
        candidateUSUBJID = intersect(candidateUSUBJID1, candidateUSUBJID2);
        
        % find rows in candidateRows that match common candidateUSUBJID
        candidateRows1sub = find(ismember(curWorksheet.expData{expDataBM1Ind}.data{candidateRows1,'USUBJID'},candidateUSUBJID));
        candidateRows2sub = find(ismember(curWorksheet.expData{expDataBM2Ind}.data{candidateRows2,'USUBJID'},candidateUSUBJID));
        
    else
        warning('No experimental data available.')
    end
    
    % set initial dose day and time at which to pull Biomarker simData
    initDoseInd = ismember('nodose',myVPop.mnSDTable.interventionID);
    if ~isempty(initDoseInd)
        initDoseDay = myVPop.mnSDTable.time(initDoseInd);
        
    else
        initDoseDay = 2000;
    end
    
    % get simulation results for Biomarker 1 and Biomarker2
    data1 = getResultOutputforIntervention(curWorksheet,Biomarker{1}.InterventionID, Biomarker{1}.ElementID);
    myRow1 = find(data1.Data(:,1 )== (initDoseDay + Biomarker{1}.Time));
    data2 = getResultOutputforIntervention(curWorksheet,Biomarker{2}.InterventionID, Biomarker{2}.ElementID);
    myRow2 = find(data2.Data(:,1 )== (initDoseDay + Biomarker{2}.Time));
    
else % create 1d Boxplots for all Biomarkers
    
    BiomarkerExpData = cell(nBiomarkers,1);
    nBiomarkerExpData = zeros(nBiomarkers,1);
    BiomarkerData = [];
    BiomarkerDataPrcntl = cell(2*nBiomarkers,1);
    allInterventionIDs = getInterventionIDs(myWorksheet);
    allElementIDNames = myWorksheet.results{1}.Names;
    myExpDataIDs = getExpDataIDs(myWorksheet);
    plotIDX = 2.*[1:nBiomarkers]-1;
    
    for i=1:nBiomarkers
        % gather biomarker data if available
        
        expDataIDBM = Biomarker{i}.ExpDataID;
        expDataBMInd = find(ismember(myExpDataIDs, expDataIDBM));
        
        
        if isfield(Biomarker{i},'Patientgroup')
            PatientgroupFlag = true;
            PatientGroup = Biomarker{i}.Patientgroup;
            if Biomarker{i}.InterventionID == "nodose"
                if isfield(Biomarker{i}.Patientgroup,'InterventionID')
                    PatientGroup.InterventionID = Biomarker{i}.Patientgroup.InterventionID;
                else
                    warning(['Please set InterventionID in Biomarker.Patientgroup',newline, ...
                        'to something else than >>nodose<<'])
                end
            else
                PatientGroup.InterventionID = Biomarker{i}.InterventionID;
            end
            [curWorksheet, curVPop] = getPatientgroup(myWorksheet, PatientGroup, myVPop);
            
        else
            curWorksheet = myWorksheet;
            curVPop = myVPop;
        end
        
        % find rows in ExpDataTable with matching TIME, TRTGRP and EXPVarID
        if ~isempty(Biomarker{i}.TrtGrp)
            candidateRows = find((curWorksheet.expData{expDataBMInd}.data{:,'TIME'} == Biomarker{i}.Time) ...
            & ismember(curWorksheet.expData{expDataBMInd}.data{:,'TRTGRP'},Biomarker{i}.TrtGrp) ...
            & ~isnan(curWorksheet.expData{expDataBMInd}.data{:,Biomarker{i}.ExpVarID}));
        else
            candidateRows = find((curWorksheet.expData{expDataBMInd}.data{:,'TIME'} == Biomarker{i}.Time) ...
            & ~isnan(curWorksheet.expData{expDataBMInd}.data{:,Biomarker{i}.ExpVarID}));
        end
        
        % restrict candidateRows to respective Patientgroup
        if PatientgroupFlag
            
            % store candidateUSUBJIDs for which Biomarker data is available
            candidateUSUBJID = ((curWorksheet.expData{expDataBMInd}.data{candidateRows,'USUBJID'}));
            
            if isempty(Biomarker{i}.Patientgroup.Time)
                % pull Patientgroup data from last available Time in
                % BRSCORE Column of expDataTable
                curBRSCORE = cell(length(candidateUSUBJID),1);
                relSLDRows = zeros(length(candidateUSUBJID),1);
                relSLDRowInd = false(length(candidateUSUBJID),1);
                
                for k=1:length(candidateUSUBJID)
                    
                    if ~isempty(Biomarker{i}.TrtGrp)
                        curSLDRows = find(ismember(curWorksheet.expData{expDataBMInd}.data{:,'USUBJID'}, candidateUSUBJID(k)) ...
                        & ~isnan(curWorksheet.expData{expDataBMInd}.data{:,'TIME'}) ...
                        & ismember(curWorksheet.expData{expDataBMInd}.data{:,'TRTGRP'},Biomarker{i}.TrtGrp) ...
                        & ~isnan(curWorksheet.expData{expDataBMInd}.data{:,'INDEX_LESION_SLD_RELCH'}) ...
                        & cellfun(@ischar,curWorksheet.expData{expDataBMInd}.data{:,'BRSCORENEW'}));
                    else
                        curSLDRows = find(ismember(curWorksheet.expData{expDataBMInd}.data{:,'USUBJID'}, candidateUSUBJID(k)) ...
                        & ~isnan(curWorksheet.expData{expDataBMInd}.data{:,'TIME'}) ...
                        & ~isnan(curWorksheet.expData{expDataBMInd}.data{:,'INDEX_LESION_SLD_RELCH'}) ...
                        & cellfun(@ischar,curWorksheet.expData{expDataBMInd}.data{:,'BRSCORENEW'}));
                    end
                    
                    if ~isempty(curSLDRows)
                        curBRSCORE{k,1} = curWorksheet.expData{expDataBMInd}.data{curSLDRows,'BRSCORENEW'}{end};
                        relSLDRows(k,1) = curSLDRows(end);
                        relSLDRowInd(k,1) = true;
                    end
                end
                % get rid of empty rows
                curBRSCORE = curBRSCORE(relSLDRowInd);
                relSLDRows = relSLDRows(relSLDRowInd);
            else
                % pull Patientgroup data from BRSCORE Column of
                % expDataTable at Time specified in Patientgroup.Time
                
                if ~isempty(Biomarker{i}.TrtGrp)
                    relSLDRows = find(ismember(curWorksheet.expData{expDataBMInd}.data{:,'USUBJID'}, candidateUSUBJID) ...
                    & (curWorksheet.expData{expDataBMInd}.data{:,'TIME'} == Biomarker{i}.Patientgroup.Time) ...
                    & ismember(curWorksheet.expData{expDataBMInd}.data{:,'TRTGRP'},Biomarker{i}.TrtGrp) ...
                    & ~isnan(curWorksheet.expData{expDataBMInd}.data{:,'INDEX_LESION_SLD_RELCH'}) ...
                    & cellfun(@ischar,curWorksheet.expData{expDataBMInd}.data{:,'BRSCORENEW'}));
                else
                    relSLDRows = find(ismember(curWorksheet.expData{expDataBMInd}.data{:,'USUBJID'}, candidateUSUBJID) ...
                    & (curWorksheet.expData{expDataBMInd}.data{:,'TIME'} == Biomarker{i}.Patientgroup.Time) ...
                    & ~isnan(curWorksheet.expData{expDataBMInd}.data{:,'INDEX_LESION_SLD_RELCH'}) ...
                    & cellfun(@ischar,curWorksheet.expData{expDataBMInd}.data{:,'BRSCORENEW'}));
                end
                
                curBRSCORE = curWorksheet.expData{expDataBMInd}.data{relSLDRows,'BRSCORENEW'};
                
            end
            
            CRInd = false(length(curBRSCORE),1);
            PRInd = false(length(curBRSCORE),1);
            SDInd = false(length(curBRSCORE),1);
            PDInd = false(length(curBRSCORE),1);
            
            if Biomarker{i}.Patientgroup.CR
                CRInd = ismember(curBRSCORE,'CR');
            end
            
            if Biomarker{i}.Patientgroup.PR
                PRInd = ismember(curBRSCORE,'PR');
            end
            
            if Biomarker{i}.Patientgroup.SD
                SDInd = ismember(curBRSCORE,'SD');
            end
            
            if Biomarker{i}.Patientgroup.PD
                PDInd = ismember(curBRSCORE,'PD');
            end
            
            BRInd = logical(CRInd + PRInd + SDInd + PDInd);
            nBiomarkerExpData(i,1) = sum(BRInd); % variable not yet used!
            
            % get USUBJIDs for rows in relSLDRows corresponding to the
            % desired Patientgroups
            BRscoreUSUBJID = curWorksheet.expData{expDataBMInd}.data{relSLDRows(BRInd),'USUBJID'};
            
            if ~isempty(Biomarker{i}.TrtGrp)
                candidateRows = find(ismember(curWorksheet.expData{expDataBMInd}.data{:,'USUBJID'}, BRscoreUSUBJID) ...
                & (curWorksheet.expData{expDataBMInd}.data{:,'TIME'} == Biomarker{i}.Time) ...
                & ismember(curWorksheet.expData{expDataBMInd}.data{:,'TRTGRP'},Biomarker{i}.TrtGrp) ...
                & ~isnan(curWorksheet.expData{expDataBMInd}.data{:,Biomarker{i}.ExpVarID}));
            
            else
                candidateRows = find(ismember(curWorksheet.expData{expDataBMInd}.data{:,'USUBJID'}, BRscoreUSUBJID) ...
                & (curWorksheet.expData{expDataBMInd}.data{:,'TIME'} == Biomarker{i}.Time) ...
                & ~isnan(curWorksheet.expData{expDataBMInd}.data{:,Biomarker{i}.ExpVarID}));
            end
        end
        
        if ~isempty(candidateRows)
            plot1dExpFlag(i) = true;
            BiomarkerExpData{i} = curWorksheet.expData{expDataBMInd}.data{candidateRows,Biomarker{i}.ExpVarID};
            
            % find and eliminate '0's in ExpData
            ZeroIDX = find(BiomarkerExpData{i} ==0);
            BiomarkerExpData{i}(ZeroIDX) =[];
        else
            warning(['No data available for biomarker: ',Biomarker{i}.ElementID], newline)
        end
        

        %         BRSCORERows = find(ismember(curWorksheet.expData{expDataBMInd}.data{:,'USUBJID'}, candidateUSUBJID{14}) ...
        %             & ismember(curWorksheet.expData{expDataBMInd}.data{:,'TRTGRP'}, Biomarker{i}.TrtGrp) ...
        %             & ~isnan(curWorksheet.expData{expDataBMInd}.data{:,'TIME'}));
        %         table(curWorksheet.expData{expDataBMInd}.data{BRSCORERows,'TIME'}, ...
        %             curWorksheet.expData{expDataBMInd}.data{BRSCORERows,'BRSCORE'})
        
        
        
        % sample biomarker according to prevalence weights
        allVPIDs = getVPIDs(curWorksheet);
        nVPs = length(allVPIDs);
        
        [pws_sorted, pws_Ind] = sort(curVPop.pws);
        pws_cum = NaN(nVPs,1);
        
        for j=1:nVPs
            pws_cum(j) = sum(pws_sorted(1:j));
        end
        
        % sample size for resampling from VPop
        nSamples = 2*nVPs;
        curBiomarkerSample = NaN(1, nSamples);
        curInterventionID = Biomarker{i}.InterventionID;
        curElementID = Biomarker{i}.ElementID;
        % set initial dose day and time at which to pull Biomarker simData
        initDoseInd = ismember('nodose',myVPop.mnSDTable.interventionID);
        if ~isempty(initDoseInd)
            initDoseDay = myVPop.mnSDTable.time(initDoseInd);
            curTime = initDoseDay + Biomarker{i}.Time;
        else
            curTime = 2000 + Biomarker{i}.Time;
        end
        
        % find indices for intervention and time
        [~, curInterventionInd] = ismember(curInterventionID, allInterventionIDs);
        curTimeInd = find(curWorksheet.results{curInterventionInd,1}.Data(:,1) == curTime);
        [~, curElementInd] = ismember(curElementID, allElementIDNames);
        
        for k=1:nSamples
            vp_rand = rand(1);
            ind_rand = find(vp_rand < pws_cum,1);
            curBiomarkerSample(1,k) = curWorksheet.results{curInterventionInd, pws_Ind(ind_rand)}.Data(curTimeInd, curElementInd);
        end
        BiomarkerDataPrcntl{2*i-1} = quantile(curBiomarkerSample, Biomarker{i}.Percentile);
        BiomarkerData = [BiomarkerData; curBiomarkerSample', (2*i-1).*ones(length(curBiomarkerSample),1)];
        if length(BiomarkerExpData{i}) >= 5
            BiomarkerDataPrcntl{2*i} = quantile(BiomarkerExpData{i}, Biomarker{i}.Percentile);
            BiomarkerData = [BiomarkerData; BiomarkerExpData{i}, 2*i.*ones(length(BiomarkerExpData{i}),1)];
        else
            plotIDX(i+1:end) = plotIDX(i+1:end)-1;
        end
        
    end
    
    figure
    plotHandle = boxplot(BiomarkerData(:,1),BiomarkerData(:,2));
    set(plotHandle(7,:),'Visible','off')
    hold on
    
    % adjust whiskers to percentile defined in the Percentile field
    for i=1:nBiomarkers
        upWhisker = get(plotHandle(1,plotIDX(i)),'YData');
        set(plotHandle(1,plotIDX(i)), 'YData', [upWhisker(1) BiomarkerDataPrcntl{2*i-1}(2)]);
        set(plotHandle(3,plotIDX(i)), 'YData', [BiomarkerDataPrcntl{2*i-1}(2) BiomarkerDataPrcntl{2*i-1}(2)]);
        dnWhisker = get(plotHandle(2,plotIDX(i)),'YData');
        set(plotHandle(2,plotIDX(i)), 'YData', [BiomarkerDataPrcntl{2*i-1}(1) dnWhisker(2)]);
        set(plotHandle(4,plotIDX(i)), 'YData', [BiomarkerDataPrcntl{2*i-1}(1) BiomarkerDataPrcntl{2*i-1}(1)]);
        
        if length(BiomarkerExpData{i}) >= 5
            
            upWhisker = get(plotHandle(1,plotIDX(i) + 1),'YData');
            set(plotHandle(1,plotIDX(i) + 1), 'YData', [upWhisker(1) BiomarkerDataPrcntl{2*i}(2)]);
            set(plotHandle(3,plotIDX(i) + 1), 'YData', [BiomarkerDataPrcntl{2*i}(2) BiomarkerDataPrcntl{2*i}(2)]);
            dnWhisker = get(plotHandle(2,plotIDX(i) + 1),'YData');
            set(plotHandle(2,plotIDX(i) + 1), 'YData', [BiomarkerDataPrcntl{2*i}(1) dnWhisker(2)]);
            set(plotHandle(4,plotIDX(i) + 1), 'YData', [BiomarkerDataPrcntl{2*i}(1) BiomarkerDataPrcntl{2*i}(1)]);
            
            plot(plotIDX(i) + 1,BiomarkerExpData{i},'ko');
        else
            if plot1dExpFlag(i) == true
               plot(plotIDX(i),BiomarkerExpData{i},'ko');
            end
        end
    end
end

if plot2dFlag
    figure
    plotHandle = scatter(data1.Data(myRow1,2:end),data2.Data(myRow2,2:end),5000*curVPop.pws);
    if plot2dExpFlag
        hold on
        scatter(myWorksheet.expData{expDataBM1Ind}.data{candidateRows1(candidateRows1sub),Biomarker{1}.ExpVarID}, ...
            myWorksheet.expData{expDataBM2Ind}.data{candidateRows2(candidateRows2sub),Biomarker{2}.ExpVarID},'filled')
    end
end
end
```
