## Supplementary File 3 for "A surrogate workflow for virtual population development: designing an amenable proxy for a hypothesis-test inspired nonlinear goodness-of-fit": plotBRVPop.m

```
function plotHandle = plotBRVPop(myVPop, myPlotOptions)
% This function takes the data in a VPop BR table
% to plot the experimental data side-by-side with the population
% to facilitate diagnosing fitting during
% the population calibration/prevalence weighting process.
%
% ARGUMENTS
% myVPop:           (required) A VPopRECIST or VPopRECISTnoBin object.
% myPlotOptions:    (optional) A plotOptions structure.
%                   Note that not all arguments are used.
%
% RETURNS
% plotHandle
%
plotHandle = [];
flagContinue = false;

if nargin > 2
    flagContinue = false;
    warning(['Too many input arguments to ',mfilename,'. Required: myVPop.'])
elseif nargin > 1
    flagContinue = true;    
elseif nargin > 0
    myPlotOptions = plotOptions;
    
    flagContinue = true;
else
    flagContinue = false;
    warning(['Insufficient input arguments to ',mfilename,'. Required: myVPop.'])
end

if flagContinue
    if ~(ismember(class(myVPop), {'VPopRECIST','VPopRECISTnoBin'}))
        flagContinue = false;
        warning(['Invalid VPopRECIST for ',mfilename,'.'])
    end  
    if ~strcmp(class(myPlotOptions),'plotOptions')
        flagContinue = false;
        warning(['Invalid plotOptions for ',mfilename,'.'])
    end        
end

if flagContinue
    if ~(strcmp(class(myVPop.brTableRECIST), 'table'))
        flagContinue = false;
        warning(['Invalid VPop.brTableRECIST for ',mfilename,'.'])
    end  
end

if (flagContinue)
    if myPlotOptions.flagPD2    
        % Make sure this is updated
        myVPop = predNPDTFLS(myVPop);
    end
   % moved these arguments to below. we want to read from the updated brTableRECIST
        myTable = myVPop.brTableRECIST;
        [myNPlots, ~] = size(myTable);
        if ~(myPlotOptions.flagPlotUnweighted) && (myNPlots > 0)
            myTable(myTable.weight==0, :) =[];
            [myNPlots, ~] = size(myTable);
        end    
        if myNPlots < 1
            flagContinue = false;
            warning(['Invalid VPop.brTableRECIST for ',mfilename,'.'])        
        end

        for rowCounter = 1 : myNPlots
            plotNames{rowCounter} = {myTable{rowCounter,'interventionID'}{1},myTable{rowCounter,'elementID'}{1},num2str(myTable{rowCounter,'time'})};
        end

    nPlotsVer = floor(sqrt(myNPlots));
    nPlotsHor = ceil(myNPlots / nPlotsVer);
    plotHandle = figure;
    for plotCounter = 1 : myNPlots
        subplot(nPlotsVer, nPlotsHor, plotCounter);
        if ~myPlotOptions.flagPD2 && ~myPlotOptions.flagPD2Exp
            expBins = [myTable{plotCounter,'expCR'},myTable{plotCounter,'expPR'},myTable{plotCounter,'expSD'},myTable{plotCounter,'expPD'}];
            predBins = [myTable{plotCounter,'predCR'},myTable{plotCounter,'predPR'},myTable{plotCounter,'predSD'},myTable{plotCounter,'predPD'}];
        elseif myPlotOptions.flagPD2 && ~myPlotOptions.flagPD2Exp
            expNPD21LS = myTable{plotCounter,'expNPD21LS'};
            predNPD21LS = myTable{plotCounter,'predNPD21LS'};
            if isnan(expNPD21LS)
                expNPD21LS = 0;
            end
            if isnan(predNPD21LS)
                predNPD21LS = 0;
            end            
            predN = myTable{plotCounter,'predN'};
            expN = myTable{plotCounter,'expN'};
            expPD = (expNPD21LS + myTable{plotCounter,'expPD'}*expN)/(expN+expNPD21LS);
            expSD = (myTable{plotCounter,'expSD'}*expN)/(expN+expNPD21LS);
            expPR = (myTable{plotCounter,'expPR'}*expN)/(expN+expNPD21LS);
            expCR = (myTable{plotCounter,'expCR'}*expN)/(expN+expNPD21LS);
            predPD = (predNPD21LS + myTable{plotCounter,'predPD'}*predN)/(predN+predNPD21LS);
            predSD = (myTable{plotCounter,'predSD'}*predN)/(predN+predNPD21LS);
            predPR = (myTable{plotCounter,'predPR'}*predN)/(predN+predNPD21LS);
            predCR = (myTable{plotCounter,'predCR'}*predN)/(predN+predNPD21LS);
            expBins = [expCR,expPR,expSD,expPD];
            predBins = [predCR,predPR,predSD,predPD];
        elseif ~myPlotOptions.flagPD2 && myPlotOptions.flagPD2Exp
            expNPD21LS = myTable{plotCounter,'expNPD21LS'};
            predNPD21LS = myTable{plotCounter,'predNPD21LS'};
            if isnan(expNPD21LS)
                expNPD21LS = 0;
            end
            if isnan(predNPD21LS)
                predNPD21LS = 0;
            end            
            predN = myTable{plotCounter,'predN'};
            expN = myTable{plotCounter,'expN'};
            expPD = (expNPD21LS + myTable{plotCounter,'expPD'}*expN)/(expN+expNPD21LS);
            expSD = (myTable{plotCounter,'expSD'}*expN)/(expN+expNPD21LS);
            expPR = (myTable{plotCounter,'expPR'}*expN)/(expN+expNPD21LS);
            expCR = (myTable{plotCounter,'expCR'}*expN)/(expN+expNPD21LS);
            predPD = (expNPD21LS*predN/expN + myTable{plotCounter,'predPD'}*predN)/(predN+expNPD21LS*predN/expN);
            predSD = (myTable{plotCounter,'predSD'}*predN)/(predN+expNPD21LS*predN/expN);
            predPR = (myTable{plotCounter,'predPR'}*predN)/(predN+expNPD21LS*predN/expN);
            predCR = (myTable{plotCounter,'predCR'}*predN)/(predN+expNPD21LS*predN/expN);
            expBins = [expCR,expPR,expSD,expPD];
            predBins = [predCR,predPR,predSD,predPD];
        elseif myPlotOptions.flagPD2 && myPlotOptions.flagPD2Exp
               warning('Both flagPD2 and flagPD2Ex are true, need to choose one to estimate PD2 rates and plot on ...');       
        end            
        binLabels = {'CR','PR','SD','PD'};
        barplot = bar([expBins;predBins]');
        set(gca,'XTickLabel',binLabels);
        xtickangle(45);
        set(barplot(1),'facecolor','w') 
        set(barplot(2),'facecolor',[.2,.2,.2]) 
        if plotCounter==1
            legend({'Trials','VPop'})
        end
        title(gca,plotNames{plotCounter},'interpreter','none','FontWeight','Normal');
        set(gca,'box','on');
        set(gca,'fontsize', 10);
    end

    if myPlotOptions.flagSave
        theDate = date;
        formatOut = 'yymmdd';
        theDate = datestr(theDate,formatOut);
        print(['VPopBR_',theDate,'.tif'],'-dtiff','-r300');
    end
end

end
```
