## Supplementary File 3 for "A surrogate workflow for virtual population development: designing an amenable proxy for a hypothesis-test inspired nonlinear goodness-of-fit": plotCoefficients.m

```
function plotHandle = plotCoefficients(myWorksheet, myPlotCoefficientsOptions)
% This function plots the axis coefficients for all of the VPs in a
% worksheet.
%
% ARGUMENTS
% myWorksheet:               (required) Worksheet with coefficients
% myPlotCoefficientsOptions: (optional) Optional structure to adjust the plotting
%
% RETURNS
% plotHandle
%
plotHandle = [];
flagExpData = false;

if nargin > 2
    flagContinue = false;
    warning(['Too many input arguments to ',mfilename,'. Required: myWorksheet; optional: myPlotCoefficientsOptions.'])
elseif nargin > 1
    flagContinue = true;
elseif nargin > 0
    flagContinue = true;
    myPlotCoefficientsOptions = plotCoefficientsOptions;
end

if flagContinue
    if ~strcmp(class(myPlotCoefficientsOptions),'plotCoefficientsOptions')
        flagContinue = false;
        warning('Invalid plotOptions for ',mfilename,'.')
    end  
    allCoefficients = (getVPCoeffs(myWorksheet));
    [nAxis, nVP] = size(allCoefficients);
    if nAxis < 1
        flagContinue = false;
        warning('Worksheet contains no axes to plot for call to ',mfilename,'.')
    end            
end

if (flagContinue)
    allCoefficients = transpose(allCoefficients);
    allAxisIDs = getAxisDefIDs(myWorksheet);
    nAxis = length(allAxisIDs);
    rng('default');
    randColors = rand(nVP,3);    
    jitterVals = 0.5*rand(nVP,1)-.25; 
    plotHandle = figure;
    plotOrder = 1:nAxis;
    if myPlotCoefficientsOptions.flagSort
        coefficientRange = max(allCoefficients,[],1) - min(allCoefficients,[],1);
        [~, plotOrder] = sort(coefficientRange,'ascend');
    end
    for plotCounter = 1 : nAxis
        subplot(nAxis, 1, plotCounter);
        yPos = (nAxis - plotCounter)/(nAxis+1) + 0.5/(nAxis+1);
        pos = get(gca, 'Position');
        pos(1) = myPlotCoefficientsOptions.leftPosition;
        pos(3) = myPlotCoefficientsOptions.width;
        set(gca, 'Position', pos);
        [ksdens, u, bb]=ksdensity(allCoefficients(:,plotOrder(plotCounter)));
        ksdens=ksdens/max(ksdens)*0.3;
        median=nanmedian(allCoefficients(:,plotOrder(plotCounter)));
        scatter(allCoefficients(:,plotOrder(plotCounter)),jitterVals,10,randColors,'filled');
        ylim([-0.3 0.3]);
        xlim(myPlotCoefficientsOptions.xLim);
        set(gca, 'YTick', [0]);
        set(gca, 'YTickLabel', {allAxisIDs{plotOrder(plotCounter)}},'TickLabelInterpreter','none');
        set(gca,'box','on');
        grid on;
        set(gca,'fontsize', myPlotCoefficientsOptions.fontSize);
        if plotCounter < nAxis
            set(gca,'XTickLabel', '');
        end
    end
    if myPlotCoefficientsOptions.flagSave
        if length(myPlotCoefficientsOptions.fileName) > 0
            print([myPlotCoefficientsOptions.fileName,'.tif'],'-dtiff','-r300');
        else        
            theDate = date;
            formatOut = 'yymmdd';
            theDate = datestr(theDate,formatOut);
            print(['worksheetAxisCoefficients_',theDate,'.tif'],'-dtiff','-r300');
        end
    end
end

end
```
