## Supplementary File 3 for "A surrogate workflow for virtual population development: designing an amenable proxy for a hypothesis-test inspired nonlinear goodness-of-fit": plotDist2DVPop.m

```
function plotHandle = plotDist2DVPop(myVPop, myPlotOptions)
% This function takes the 2D distribution data in a VPop distribution table
% to plot the experimental data side-by-side with the population
% to facilitate diagnosing fitting during
% the population calibration/prevalence weighting process.
%
% ARGUMENTS
% myVPop:           (required) A VPop.  Note there should be a distTable2D.
% myPlotOptions:    (optional) A plotOptions structure.
%                   Note that not all arguments are used.
%
% RETURNS
% plotHandle
%
plotHandle = [];
flagContinue = false;

if nargin > 2
    flagContinue = false;
    warning(['Too many input arguments to ',mfilename,'. Required: myVPop.'])
elseif nargin > 1
    flagContinue = true;    
elseif nargin > 0
    myPlotOptions = plotOptions;
    
    flagContinue = true;
else
    flagContinue = false;
    warning(['Insufficient input arguments to ',mfilename,'. Required: myVPop.'])
end

if flagContinue
    if ~(ismember(class(myVPop), {'VPop','VPopRECIST','VPopRECISTnoBin'}))
        flagContinue = false;
        warning(['Invalid VPop for ',mfilename,'.'])
    end  
    if ~strcmp(class(myPlotOptions),'plotOptions')
        flagContinue = false;
        warning(['Invalid plotOptions for ',mfilename,'.'])
    end        
end

if flagContinue
    if ~(strcmp(class(myVPop.distTable), 'table'))
        flagContinue = false;
        warning(['Invalid VPop.distTable for ',mfilename,'.'])
    end  
end

if (flagContinue)
    myTable = myVPop.distTable2D;
    [myNPlots, ~] = size(myTable);
    if ~(myPlotOptions.flagPlotUnweighted) && (myNPlots > 0)
        myTable(myTable.weight==0, :) =[];
        [myNPlots, ~] = size(myTable);
    end        
    if myNPlots < 1
        flagContinue = false;
        warning(['Invalid VPop.distTable for ',mfilename,'.'])        
    end
end


if (flagContinue)
    plotNames1 = cell(1, myNPlots);
    plotNames2 = cell(1, myNPlots);
    for rowCounter = 1 : myNPlots
        plotNames1{rowCounter} = {myTable{rowCounter,'interventionID1'}{1},myTable{rowCounter,'elementID1'}{1},num2str(myTable{rowCounter,'time1'})};
		plotNames2{rowCounter} = {myTable{rowCounter,'interventionID2'}{1},myTable{rowCounter,'elementID2'}{1},num2str(myTable{rowCounter,'time2'})};
    end
    pwScale = true;
end

if (flagContinue)
    nPlotsVer = floor(sqrt(myNPlots));
    nPlotsHor = ceil(myNPlots / nPlotsVer);
    plotHandle = figure;
    for plotCounter = 1 : myNPlots
        subplot(nPlotsVer, nPlotsHor, plotCounter);
        expN = myTable{plotCounter,'expN'};
        predN = myTable{plotCounter,'predN'};
		if pwScale
			expW = 1./(expN*ones(1,expN));
			predW = myTable{plotCounter,'predProbs'}{1};
			if min(predW) <= 0
				minPredW = min(predW) + 1E-6;
				predW = predW + minPredW;
				predW = predW./sum(predW);
			end
		else
			expW = 0.05*ones(1,expN);
			[~, nPWs] = dim(predW);
			predW = 0.05*ones(1,nPWs);
		end
        expSample = myTable{plotCounter,'expSample'}{1};
        predSample = myTable{plotCounter,'predSample'}{1};
		
		hold on
		scatter(predSample(1,:),predSample(2,:),500*predW,'bo');
		scatter(expSample(1,:),expSample(2,:),500*expW,'ro','filled');
		hold off
		
        grid on
        xlabel(plotNames1{plotCounter},'interpreter','none','FontWeight','Normal');
		ylabel(plotNames2{plotCounter},'interpreter','none','FontWeight','Normal');
        set(gca,'box','on');
        set(gca,'fontsize', 10);
        
    end

    if myPlotOptions.flagSave
        theDate = date;
        formatOut = 'yymmdd';
        theDate = datestr(theDate,formatOut);
        print(['VPopDistCDF2D_',theDate,'.tif'],'-dtiff','-r300');
    end
end

end
```
