## Supplementary File 3 for "A surrogate workflow for virtual population development: designing an amenable proxy for a hypothesis-test inspired nonlinear goodness-of-fit": plotInterventionVPop.m

```
function plotHandleVector = plotInterventionVPop(myWorksheet, myVPop, myPlotOptions, existingFigure)
% This function plots results for a single variable stored in a worksheet.
% The returned plots show the 5th, 50th, and 95th percentile.
%
% ARGUMENTS
% myWorksheet:      (required) Worksheet with simulation results
% myVPop:           (required) Virtual population with prevalence weight
%                   solution.  VPopRECIST also allowed, in which
%                   case only on-therapy VPs will be plotted.
% myPlotOptions:    Options structure to adjust the plotting
% existingFigure:   'true' if you want to plot into an existing figure
%                   (such as a subplot), 'false' creates a new figure
%                   DEFAULT is false
%
% RETURNS
% plotHandle
%

% Check number of inputs.
% We define fail behavior as warn rather than error here
% This just seems a more reasonable way to fail if we can generate plots
% rather than interrupting higher level execution with an error.
plotHandleVector = [];
flagExpData = false;
flagSubPlot = false;

if nargin > 4
    flagContinue = false;
    warning(['Too many input arguments for ',mfilename,', require: myWorksheet, myVPop, myPlotOptions; optionally: existingFigure. Exiting.'])
elseif nargin > 2
    flagContinue = true;
else
    flagContinue = false;
    warning(['Insufficient input arguments for ',mfilename,', require: myWorksheet, myVPop, myPlotOptions. Exiting.'])  
end

if flagContinue
    if ~strcmp(class(myPlotOptions),'plotOptions')
        flagContinue = false;
        warning(['Invalid plotOptions object for ',mfilename,', exiting.'])
    end        
    if ~ismember(class(myVPop), {'VPop','VPopRECIST','VPopRECISTnoBin'})
        flagContinue = false;
        warning(['Invalid VPop for ',mfilename,'.'])
    end             
end

if flagContinue    
    if length(myVPop.pws) < 1
        flagContinue = false;
        warning(['VPop contains no prevalence weight results for ',mfilename,', exiting.'])
    end            
end


if flagContinue
    if length(myPlotOptions.vpIDs) < 1
        myPlotOptions.vpIDs = getVPIDs(myWorksheet);
    end
end

if flagContinue
    vpIDs = myPlotOptions.vpIDs;
    allVPIDs = getVPIDs(myWorksheet);
    if (sum(ismember(allVPIDs,vpIDs))<length(allVPIDs))
        warning(['All VPs are considered for the purpose of plotting the results of the VPop in ',mfilename,'. Cannot proceed with a subset, exiting.'])
        flagContinue = false;
    end
end

if flagContinue
    if ((length(myPlotOptions.expDataID) > 0) || (length(myPlotOptions.expDataTimeVar) > 0) || (length(myPlotOptions.expDataYVar) > 0))
        flagExpData = true;
    end
end

if flagContinue
    flagContinue = myPlotOptions.verify(myWorksheet);
    if ~flagContinue
        warning(['The plotOptions failed verification for ',mfilename,', exiting.'])
    end
end

   
if flagContinue
    nVPs = length(allVPIDs);
    interventionID = myPlotOptions.interventionID;    
    varName = myPlotOptions.varName;
end

if flagContinue && nargin == 4
    if ~islogical(existingFigure)
        warning(['4th argument must be logical (true or false). Assuming FALSE by default.'])
    else
        flagSubPlot = existingFigure;
    end
end

if flagExpData && flagContinue
    % Filter out nan's, this will cause issues 
    expDataID = myPlotOptions.expDataID;
    expData = getExpData(expDataID, myWorksheet);
    expDataTimeVar = myPlotOptions.expDataTimeVar;
    expDataYVar = myPlotOptions.expDataYVar;
    expDataTime = expData.(expDataTimeVar);
    expDataYVar = expData.(expDataYVar);
    the_indices = find((~isnan(expDataTime)) & (~isnan(expDataYVar)));
    expDataTime = expDataTime(the_indices);
    expDataYVar = expDataYVar(the_indices);
end

if flagContinue
    % We reconstruct data here rather than getting from VPop in case
    % new variables will be plotted that weren't in the calibration
    interventionIDs = getInterventionIDs(myWorksheet);
    wshInterventionIndex = find(ismember(interventionIDs,interventionID));
    [nTimePoints, ~] = size(myWorksheet.results{wshInterventionIndex, 1}.Data);
    dataToPlot = nan(nTimePoints,nVPs);
    % Time should be constant
    for vpCounter = 1 : nVPs
        curResult = myWorksheet.results{wshInterventionIndex, vpCounter};
        curTimeIndex = find(ismember(curResult.Names,'time'));
        curVarIndex = find(ismember(curResult.Names,varName));
        curTime = curResult.Data(:,curTimeIndex);
        curData = curResult.Data(:,curVarIndex);
        % Interpolate to get the time exactly right
        % interpolateValue = interp1(curTime,curVar,expTime,'linear');
        dataToPlot(:, vpCounter) = curData/myPlotOptions.yScale;
    end
    
    % Plot 5/50/95 prediction interval
    intervalsToPlot = nan(nTimePoints,3);
    for timePointCounter = 1 : nTimePoints
        curData = dataToPlot(timePointCounter,:);
        curPWs = myVPop.pws;
        if isa(myVPop,'VPopRECIST')
            keepIndices = find(myVPop.recistSimFilter{wshInterventionIndex}.filterMatrix(timePointCounter,:));
            curData = curData(keepIndices);
            curPWs = curPWs(keepIndices)/sum(curPWs(keepIndices));
        end
        [curData, indices] = sort(curData);
        curPWs = curPWs(indices);
        curPWs = cumsum(curPWs);
        % PWs may be very small, and may be zero especially with rounding.
        % so we'll use both "first" and "last" occurrence of the PW comsum
        % and take the median of intervening values as the point for
        % to inform the percentile extrapolation
        [temp, indicesFirst] = unique(curPWs,'first');
        [curPWs, indicesLast] = unique(curPWs,'last');
        nUnique = length(indicesFirst);
        filteredData = nan(1, nUnique);
        for dataPointCounter = 1 : nUnique
            currentValues = curData(indicesFirst(dataPointCounter):indicesLast(dataPointCounter));
            filteredData(dataPointCounter) = median(currentValues);
        end

        lbPercentile = 0.05;
        ubPercentile = 0.95;
        if curPWs(1) > 0.05
            lbPercentile = curPWs(1);
            warning(['Lowest percentile in the VPop data is higher than 5, plotting ' num2str(round(lbPercentile*100)) 'th to 95th percentile ...']);
        end
% 
%         if curPWs(end) < 0.95
%             lbPercentile = curPWs(1);
%             warning(sprintf('Lowest percentile in the VPop data is higher than 5, plotting %fth to 95 percentile ...'));
%         end
        intervalsToPlot(timePointCounter,:) = interp1(curPWs,filteredData,[lbPercentile,0.50,ubPercentile],'linear');

%         intervalsToPlot(timePointCounter,:) = interp1(curPWs,filteredData,[lbPercentile,0.50,0.95],'linear','extrap');
    end
    % Carry last observation forward to patch NaN.  This will fail if first
    % value is nan
    for iCounter = 1: 3
        curIndices = (~isnan(intervalsToPlot(:,iCounter)));
        temp = intervalsToPlot(curIndices,iCounter);
        intervalsToPlot(:,iCounter) = temp(cumsum(curIndices));
    end
    
    if flagSubPlot
        % plot into existing figure
    else
        figure;
    end
    hold on
    plotHandleVector(1) = fill([(curTime-myPlotOptions.xShiftSim)/myPlotOptions.xScale;flipud((curTime-myPlotOptions.xShiftSim)/myPlotOptions.xScale)],[intervalsToPlot(:,1);flipud(intervalsToPlot(:,2))],'b','edgecolor','none');
    set(plotHandleVector(1),'FaceAlpha',0.5);
    plotHandleVector(3) = fill([(curTime-myPlotOptions.xShiftSim)/myPlotOptions.xScale;flipud((curTime-myPlotOptions.xShiftSim)/myPlotOptions.xScale)],[intervalsToPlot(:,2);flipud(intervalsToPlot(:,3))],'b','edgecolor','none');
    set(plotHandleVector(3),'FaceAlpha',0.5); 
    plotHandleVector(1:3) = plot((curTime-myPlotOptions.xShiftSim)/myPlotOptions.xScale, intervalsToPlot(:,1), 'b-', (curTime-myPlotOptions.xShiftSim)/myPlotOptions.xScale, intervalsToPlot(:,2),'b-', (curTime-myPlotOptions.xShiftSim)/myPlotOptions.xScale, intervalsToPlot(:,3),'b-', 'LineWidth',4);
    set(plotHandleVector(1),'linewidth',1)
    set(plotHandleVector(3),'linewidth',1)
    if (strcmp(myPlotOptions.scale,'xlogylin') || strcmp(myPlotOptions.scale,'xlogylog'))
        set(gca,'xscale','log'); % gca
    end
    if (strcmp(myPlotOptions.scale,'xlogylog') || strcmp(myPlotOptions.scale,'xlinylog'))
        set(gca,'yscale','log'); % gca
    end
    if flagExpData
        plotHandleVector(4) = plot((expDataTime-myPlotOptions.xShiftData)/myPlotOptions.xScale, expDataYVar/myPlotOptions.yScale, 'k.','MarkerSize',20); 
        if (strcmp(myPlotOptions.scale,'xlogylin') || strcmp(myPlotOptions.scale,'xlogylog'))
            set(gca,'xscale','log'); % gca
        end
        if (strcmp(myPlotOptions.scale,'xlogylog') || strcmp(myPlotOptions.scale,'xlinylog'))
            set(gca,'yscale','log'); % gca
        end    
        %set(get(get(plotHandleVector(nVP+1),'Annotation'),'LegendInformation'),'IconDisplayStyle','off');
    end
    if length(myPlotOptions.yLim) > 0
        ylim(myPlotOptions.yLim);
    end 
    if length(myPlotOptions.xLabelPretty) > 0
        xlabel(myPlotOptions.xLabelPretty,'interpreter','none');
    else
        xlabel('time','interpreter','none');
    end
    if length(myPlotOptions.yLabelPretty) > 0
        ylabel(myPlotOptions.yLabelPretty,'interpreter','none');
    else
        ylabel(varName,'interpreter','none');
    end    
    if length(myPlotOptions.xLim) > 0
        xlim(myPlotOptions.xLim);
    end    
    if length(myPlotOptions.yLim) > 0
        ylim(myPlotOptions.yLim);
    end     
    grid on
    set(gca,'fontsize', 18);
    if myPlotOptions.flagSave
        if length(myPlotOptions.fileName) > 0
            % saveas(gca,[myPlotOptions.fileName,'.tif']);
            print([myPlotOptions.fileName,'.tif'],'-dtiff','-r300');
        else
            theDate = date;
            formatOut = 'yymmdd';
            theDate = datestr(theDate,formatOut);            
            print([interventionID,'_',varName,'_vpop_',theDate,'.tif'],'-dtiff','-r300');
        end
    end
end
end
```
