## Supplementary File 3 for "A surrogate workflow for virtual population development: designing an amenable proxy for a hypothesis-test inspired nonlinear goodness-of-fit": plotMnSDVPop.m

```
function plotHandle = plotMnSDVPop(myVPop, myPlotOptions)
% This function takes the mean & SD matching data in a VPop MnSD table
% to plot the experimental data side-by-side with the population
% to facilitate diagnosing fitting during
% the population calibration/prevalence weighting process.
%
% ARGUMENTS
% myVPop:           (required) A VPop.  Note that both the 
%                   experimental and simulation result fields should be 
%                   populated
% myPlotOptions:    (optional) A plotOptions structure.
%                   Note that not all arguments are used.
%
% RETURNS
% plotHandle
%
plotHandle = [];
flagContinue = false;

if nargin > 2
    flagContinue = false;
    warning(['Too many input arguments to ',mfilename,'. Required: myVPop.'])
elseif nargin > 1
    flagContinue = true;    
elseif nargin > 0
    myPlotOptions = plotOptions;    
    flagContinue = true;
else
    flagContinue = false;
    warning(['Insufficient input arguments to ',mfilename,'. Required: myVPop.'])
end

if flagContinue
    if ~(ismember(class(myVPop), {'VPop','VPopRECIST','VPopRECISTnoBin'}))
        flagContinue = false;
        warning(['Invalid VPop for ',mfilename,'.'])
    end  
    if ~strcmp(class(myPlotOptions),'plotOptions')
        flagContinue = false;
        warning(['Invalid plotOptions for ',mfilename,'.'])
    end        
end

if flagContinue
    if ~(strcmp(class(myVPop.mnSDTable), 'table'))
        flagContinue = false;
        warning(['Invalid VPop.mnSDTable for ',mfilename,'.'])
    end  
end

if (flagContinue)
	myTable = myVPop.mnSDTable;
    [myNPlots, ~] = size(myTable);
    if ~(myPlotOptions.flagPlotUnweighted) && (myNPlots > 0)
        myTable(myTable.weightMean==0 | myTable.weightSD==0, :) =[];
        [myNPlots, ~] = size(myTable);
    end
    [myNPlots, ~] = size(myTable);
    if myNPlots < 1
        flagContinue = false;
        warning(['Invalid VPop.mnSDTable for ',mfilename,'.'])        
    end
end

if (flagContinue)
    myMnValExp = myTable{:,'expMean'};
    myUBValExp = myTable{:,'expSD'};
    myLBValExp = myTable{:,'expSD'};
    
    myMnValVPop = myTable{:,'predMean'};
    myUBValVPop = myTable{:,'predSD'};
    myLBValVPop = myTable{:,'predSD'};
	
	logNVPop = myTable{:,'logN'};
	
	
	expMn = myTable{:,'expMean'};
    expSD = myTable{:,'expSD'};
    predSD = myTable{:,'predSD'};
	predMn = myTable{:,'predMean'};
	logN = myTable{:,'logN'};
		
	% Convert lognormal summary data
	if sum(logN) > 0
		expSDBak = expSD;
		expMnBak = expMn;
		predSDBak = predSD;
		predMnBak = predMn;
		predMn(logN) = log(predMnBak(logN)./sqrt(1+(predSDBak(logN).^2)./(predMnBak(logN).^2)));
		expMn(logN) = log(expMnBak(logN)./sqrt(1+(expSDBak(logN).^2)./(expMnBak(logN).^2)));
		predSD(logN) = sqrt(log(1+(predSDBak(logN).^2)./(predMnBak(logN).^2)));
		expSD(logN) = sqrt(log(1+(expSDBak(logN).^2)./(expMnBak(logN).^2)));
	end	
	
    for rowCounter = 1 : myNPlots
        plotNames{rowCounter} = {myTable{rowCounter,'interventionID'}{1},myTable{rowCounter,'elementID'}{1},num2str(myTable{rowCounter,'time'})};
    end
    
end

if (flagContinue)
    nPlotsVer = floor(sqrt(myNPlots));
    nPlotsHor = ceil(myNPlots / nPlotsVer);
    plotHandle = figure;
    for plotCounter = 1 : myNPlots
		if logN(plotCounter)
			curExpUV = exp(expMn(plotCounter)+expSD(plotCounter));
			curPredUV = exp(predMn(plotCounter)+predSD(plotCounter));
			curExpLV = exp(expMn(plotCounter)-expSD(plotCounter));
			curPredLV = exp(predMn(plotCounter)-predSD(plotCounter));	
			curExpMn = exp(expMn(plotCounter));
			curPredMn = exp(predMn(plotCounter));
		else
			curExpUV = (expMn(plotCounter)+expSD(plotCounter));
			curPredUV = (predMn(plotCounter)+predSD(plotCounter));
			curExpLV = (expMn(plotCounter)-expSD(plotCounter));
			curPredLV = (predMn(plotCounter)-predSD(plotCounter));	
			curExpMn = (expMn(plotCounter));
			curPredMn = (predMn(plotCounter));			
		end
	
        subplot(nPlotsVer, nPlotsHor, plotCounter);
        fill([-2,2,2,-2],[curExpUV,curExpUV,curExpLV,curExpLV], [0.75 0.75 0.75])
        hold on
        errorbar(-1,curExpMn,curExpMn-curExpLV,curExpUV-curExpMn,'ko-','MarkerFaceColor','k','LineWidth',3);
        hold on
        errorbar(1,curPredMn,curPredMn-curPredLV,curPredUV-curPredMn,'ro-','MarkerFaceColor','r','LineWidth',3);
        hold on
        xlim([-1.25 1.25]);
        
        set(gca,'TickLabelInterpreter', 'none');
        set(gca, 'XTick', [-1 1]);
        set(gca, 'XTickLabel', {'Exp','VPop'});
        set(gca,'TickLabelInterpreter', 'none');
        grid on
        title(gca,plotNames{plotCounter},'interpreter','none','FontWeight','Normal');
        
        set(gca,'box','on');
        set(gca,'fontsize', 10);
        
    end

    if myPlotOptions.flagSave
        theDate = date;
        formatOut = 'yymmdd';
        theDate = datestr(theDate,formatOut);
        print(['VPopMnSD_',theDate,'.tif'],'-dtiff','-r300');
    end
end

end
```
