## Supplementary File 3 for "A surrogate workflow for virtual population development: designing an amenable proxy for a hypothesis-test inspired nonlinear goodness-of-fit": plotPWHist.m

```
function plotHandleVector = plotPWHist(myVPop, myPlotOptions)
% This function plots a histogram for non-zero prevalence weights for a
% VPop
%
% ARGUMENTS
% myVPop:           (required) Virtual population with prevalence weight
%                   solution
% myPlotOptions:    (optional) Options structure to adjust the plotting.
%                   Note that not all arguments are used.
%
% RETURNS
% plotHandle
%

plotHandleVector = [];

if nargin > 2
    flagContinue = false;
    warning('Too many input arguments for ',mfilename,', require: myVPop, optionally myPlotOptions. Exiting.')
elseif nargin > 1
    flagContinue = true;
elseif nargin > 0
    flagContinue = true;
    myPlotOptions = plotOptions();
    myPlotOptions.xLabelPretty = 'log10(Prevalence Weight)';
    myPlotOptions.yLabelPretty = 'Count';    
    myPlotOptions.scale = 'xlogylin';
else
    flagContinue = false;
    warning('Insufficient input arguments for ',mfilename,', require: myVPop, myPlotOptions. Exiting.') 
end

if flagContinue
    if ~strcmp(class(myPlotOptions),'plotOptions')
        flagContinue = false;
        warning('Invalid plotOptions for ',mfilename,'.')
    end    
    if ~isa(myVPop,'VPop') && ~isa(myVPop, 'VPopRECIST')
        flagContinue = false;
        warning('Invalid VPop for ',mfilename,'.')
    end 
end

if flagContinue    
    if length(myVPop.pws) < 1
        flagContinue = false;
        warning('VPop contains no prevalence weight results for ',mfilename,'.')
    end            
end


if flagContinue
    nbins = 20;
    allPWs = myVPop.pws;
    maxPW = max(allPWs);
    minPW = min(allPWs);
    allPWNoZero = find(allPWs > 0);
    allPWNoZero = allPWs(allPWNoZero);
    minPWNoZero = min(allPWNoZero);
    if strcmp(myPlotOptions.scale,'xlogylin') || strcmp(myPlotOptions.scale,'xlogylog')
        minLog = log10(minPWNoZero);
        maxLog = log10(maxPW);
        interval = (maxLog - minLog) / (nbins);
        binEdges = [minLog : interval: maxLog];
        binMid = binEdges(1:(nbins)) + interval/2;
        myCounts = histc(log10(allPWNoZero),binEdges);
        myCounts(nbins) = myCounts(nbins) + myCounts(nbins+1);
        myCounts(nbins+1) = [];
    else
        interval = (maxPW - minPWNoZero) / (nbins);
        binEdges = [minPWNoZero : interval : maxPW];
        binMid = binEdges(1:(nbins)) + interval/2;
        myCounts = histc((allPWNoZero),binEdges);
        myCounts(nbins) = myCounts(nbins) + myCounts(nbins+1);
        myCounts(nbins+1) = [];
    end
    cumCounts = cumsum(myCounts);
    figure;
    hold on
    plotHandleVector(1:2) = plot(binMid,myCounts,'.k-',binMid,cumCounts,'.k:', 'LineWidth',4);
    if (strcmp(myPlotOptions.scale,'xlogylog') || strcmp(myPlotOptions.scale,'xlinylog'))
        set(gca,'yscale','log'); % gca
    end
    set(gca,'XTick',binEdges);
    set(gca,'XTickLabel',binEdges);
    set(gca, 'XTickLabelRotation', 90);
    if length(myPlotOptions.yLim) > 0
        ylim(myPlotOptions.yLim);
    end 
    if length(myPlotOptions.xLabelPretty) > 0
        xlabel(myPlotOptions.xLabelPretty);
    end
    if length(myPlotOptions.yLabelPretty) > 0
        ylabel(myPlotOptions.yLabelPretty);
    else
        ylabel(myPlotOptions.varName,'interpreter','none');
    end    
    if length(myPlotOptions.xLim) > 0
        xlim(myPlotOptions.xLim);
    end    
    if length(myPlotOptions.yLim) > 0
        ylim(myPlotOptions.yLim);
    end     
    grid on
    set(gca,'fontsize', 18);
    if myPlotOptions.flagSave
        theDate = date;
        formatOut = 'yymmdd';
        theDate = datestr(theDate,formatOut);
        print(['VPop_',theDate,'.tif'],'-dtiff','-r300');
    end
end
```
