## Supplementary File 3 for "A surrogate workflow for virtual population development: designing an amenable proxy for a hypothesis-test inspired nonlinear goodness-of-fit": plotSimExpDataVPop.m

```
function plotHandle = plotSimExpDataVPop(myVPop, myPlotOptions)
% This function plots the simData stored in a population with the corresponding
% experimental data side-by-side to facilitate diagnosing fitting issues.
%
% ARGUMENTS
% myVPop:           (required) A VPop.  Note that both the experimental and
%                   simulation result fields should be populated.
% myPlotOptions:    (optional) A plotOptions structure.
%                   Note that not all arguments are used.
%
% RETURNS
% plotHandle
%
plotHandle = [];
flagContinue = false;

if nargin > 2
    flagContinue = false;
    warning(['Too many input arguments to ',mfilename,'. Required: myVPop.'])
elseif nargin > 1
    flagContinue = true;    
elseif nargin > 0
    myPlotOptions = plotOptions;    
    flagContinue = true;
else
    flagContinue = false;
    warning(['Insufficient input arguments to ',mfilename,'. Required: myVPop.'])
end

if flagContinue
    if ~(strcmp(class(myVPop),'VPop'))
        flagContinue = false;
        warning('Invalid VPop for ',mfilename,'.')
    end
    if ~strcmp(class(myPlotOptions),'plotOptions')
        flagContinue = false;
        warning('Invalid plotOptions for ',mfilename,'.')
    end  
end

if flagContinue
    if ~(strcmp(class(myVPop.expData), 'table'))
        flagContinue = false;
        warning('Invalid VPop.expData for ',mfilename,'.')
    end  
    if ~(strcmp(class(myVPop.simData),'struct'))
        flagContinue = false;
        warning('Invalid VPop.simData for ',mfilename,'.')
    end      
end

if (flagContinue)
    matchedNamesFixed = {'time', 'interventionID', 'elementID', 'elementType', 'expVarID'};
    nVPs = length(myVPop.simData.vpIDs);
    expDataNameIndices = nan(1,5);
    simDataNameIndices = nan(1,5);
    for fieldCounter = 1 : length(matchedNamesFixed)
        simDataNameIndices(fieldCounter) = find(ismember(myVPop.simData.rowInfoNames, matchedNamesFixed{fieldCounter}));
    end
    expRowInfo = (myVPop.expData(:,matchedNamesFixed));
    simRowInfo = cell2table(myVPop.simData.rowInfo(:,simDataNameIndices));
    simRowInfo.Properties.VariableNames = matchedNamesFixed;
    [nRowExp, ~] = size(expRowInfo);
    [nRowSim, ~] = size(simRowInfo);
    % We want to pair rows
    rowMap = nan(min(nRowExp,nRowSim),2);
    if nRowExp < nRowSim
        for rowCounter = 1 : nRowSim
            testRow = simRowInfo(rowCounter,:);
            expIndex = find(ismember(expRowInfo,testRow,'rows'));
            if length(expIndex) == 1
                rowMap(rowCounter,:) = [rowCounter, expIndex];
            end
        end
    else
        for rowCounter = 1 : nRowExp
            testRow = expRowInfo(rowCounter,:);
            simIndex = find(ismember(simRowInfo,testRow,'rows'));
            if length(simIndex) == 1
                rowMap(rowCounter,:) = [simIndex, rowCounter];
            end
        end        
    end
    keepRows = find(~isnan(rowMap(:,1)) & ~isnan(rowMap(:,2)));
    rowMap = rowMap(keepRows,:);
    [nSubPlot,~] = size(rowMap);
    simToPlot = nan(nSubPlot,nVPs);
    expToPlot = cell(nSubPlot,0);
    plotNames = cell(nSubPlot,0);

    commonNames = loadCommonNames();
    if isa(myVPop,'VPop')
        tableVariableNames = commonNames.VPOPTABLEVARNAMESFIXED;
		nDataHeaderCols = length(commonNames.VPOPTABLEVARNAMESFIXED);
    else
        tableVariableNames = commonNames.VPOPRECISTTABLEVARNAMESFIXED;
		nDataHeaderCols = length(commonNames.VPOPRECISTTABLEVARNAMESFIXED);
    end

    for rowCounter = 1 : length(keepRows)
        simRow = rowMap(rowCounter,1);
        expRow = rowMap(rowCounter,2);
        curVPData = myVPop.simData.Data(simRow, :);
        curExpData = myVPop.expData{expRow, (nDataHeaderCols+1):end};
        curExpData = curExpData(~isnan(curExpData));
        simToPlot(rowCounter,:) = curVPData;
        expToPlot{rowCounter} = curExpData;
        plotNames{rowCounter} = [myVPop.expData{expRow,'interventionID'}{1},'_',myVPop.expData{expRow,'elementID'}{1},'_',num2str(myVPop.expData{expRow,'time'})];
    end
    
    rng('default');
    %randColors = rand(nVP,3);    
    jitterVals = rand(nVPs,1); 
    plotHandle = figure;
    for plotCounter = 1 : nSubPlot
        subplot(nSubPlot, 1, plotCounter);
        yPos = (nSubPlot - plotCounter)/(nSubPlot+1) + 0.5/(nSubPlot+1);
        pos = get(gca, 'Position');
        pos(1) = 0.3;
        pos(3) = 0.65;
        set(gca, 'Position', pos);
        plot(simToPlot(plotCounter,:),jitterVals,'ro',expToPlot{plotCounter},jitterVals(1:length(expToPlot{plotCounter}))-1,'b^');
        set(gca,'TickLabelInterpreter', 'none');
        set(gca, 'YTick', [0]);
        set(gca, 'YTickLabel', {plotNames{plotCounter}});
        ylim([-1 1]);        
        set(gca,'XTickLabel', '');
        set(gca,'box','on');
        
        grid on;
        set(gca,'fontsize', 10);
%         if plotCounter < nAxis
%             set(gca,'XTickLabel', '');
%         end
    end        
    if myPlotOptions.flagSave
        theDate = date;
        formatOut = 'yymmdd';
        theDate = datestr(theDate,formatOut);
        print(['simDataVPop_',theDate,'.tif'],'-dtiff','-r300');
    end
end

end
```
