## Supplementary File 3 for "A surrogate workflow for virtual population development: designing an amenable proxy for a hypothesis-test inspired nonlinear goodness-of-fit": adjustLambda.m

```
function [myVPop, linearCalibrationObject] = adjustLambda(myVPop,targetEffN)
% using Bisect method to adjust lambda to allow VPop to reach targetEffN (+/- 10 is enough). used in the beginning of 
%
% ARGUMENTS
% myVPop:         An instance of a VPop, VPopRECIST object.  Populating the tables
%
% RETURNS
% myVpop:         A VPop is returned, with the fields populated:
%                  lambda
continueFlag = false;
if nargin > 2
    warning(['Too many input arguments provided to,',mfilename,'.  Requires: myVPop.'])
    continueFlag = false;
elseif nargin > 0
    continueFlag = true;
else
    continueFlag = false;
    warning(['Insufficient input arguments provided to,',mfilename,'.  Requires: myVPop.'])
end

if continueFlag
         myVPop=evaluateMSE(myVPop);
         vpPrevalenceWeights=myVPop.pws';
         SubgroupSumWeights = myVPop.LinearProblemMatrices.vpIsInSubgroup*vpPrevalenceWeights;
         myVPop.LinearProblemMatricesSubgroupSumWeights = SubgroupSumWeights; % calculate the subgroup weights 
         myOptimOptions = LinearCalibrationOptions();
         myOptimOptions.cdfProbsToFit = 0.05:0.05:0.95;
         myOptimOptions.pdf2DProbsToFitN = 5;
         myOptimOptions.responseValTransformation='none';
         myOptimOptions.optimizationAlgorithm = "quadprogEffN";				
         myOptimOptions.priorPrevalenceWeightAssumption = 'specified';
         myOptimOptions.method = "bestFit";
         myOptimOptions.oldVPop = myVPop; 

         linearCalibrationObject = LinearCalibration(myVPop,'optimOptions',myOptimOptions);   
         try
            linearCalibrationObject = linearCalibrationObject.run('closeParallelPoolWhenFinished',false);
            myVPop = linearCalibrationObject.OptimizedVPop;
         catch
            myVPop.lambda = 0; 
         end
         curVPopEffN = 1/sum(myVPop.pws.^2);
         
  %      tic;
    if (myVPop.lambda>0 && abs(curVPopEffN-targetEffN)>10) || (myVPop.lambda==0 && curVPopEffN < targetEffN && abs(curVPopEffN-targetEffN)>10)       
         % if current EffN > targetEffN, reduce it until it finds lower bound
         if curVPopEffN > targetEffN
            Blambda = myVPop.lambda;
            BEffN = curVPopEffN;
            Alambda = Blambda;
            AEffN = BEffN;
            while AEffN > targetEffN
                Alambda = Alambda/10;
                myVPop.lambda = Alambda;
                
                myOptimOptions.oldVPop = myVPop; 
                linearCalibrationObject = LinearCalibration(myVPop,'optimOptions',myOptimOptions); 
                linearCalibrationObject.LinearProblemMatrices = myVPop.LinearProblemMatrices;
                try
                    linearCalibrationObject = linearCalibrationObject.run('closeParallelPoolWhenFinished',false);
                    myVPop = linearCalibrationObject.OptimizedVPop;
                    myVPop.lambda = linearCalibrationObject.lambda;         
                catch
                    myVPop.lambda = 0;
                end
                AEffN = 1/sum(myVPop.pws.^2);
            end
         else
            Alambda = myVPop.lambda;
            AEffN = curVPopEffN;
            Blambda = Alambda;
            BEffN = AEffN;
            while BEffN < targetEffN
                Blambda = max(Blambda*10,Blambda+10);  % in case Blambda starts with 0
                myVPop.lambda = Blambda;
                
                myOptimOptions.oldVPop = myVPop; 
                linearCalibrationObject = LinearCalibration(myVPop,'optimOptions',myOptimOptions);  
                linearCalibrationObject.LinearProblemMatrices = myVPop.LinearProblemMatrices;
                try
                    linearCalibrationObject = linearCalibrationObject.run('closeParallelPoolWhenFinished',false);
                    myVPop = linearCalibrationObject.OptimizedVPop;
                    myVPop.lambda = linearCalibrationObject.lambda;
                catch
                    myVPop.lambda = 0;
                end
                BEffN = 1/sum(myVPop.pws.^2);
            end
         end
  %       toc;
        
        % Bisect to find the Clabmda that gives EffN= targetEffN
            iter = 1;
            while iter <= 20 
            %    iter
                Clambda = 0.5*(Alambda+Blambda); % take mean
                myVPop.lambda = Clambda;
                
                myOptimOptions.oldVPop = myVPop; 
                linearCalibrationObject = LinearCalibration(myVPop,'optimOptions',myOptimOptions);  
                linearCalibrationObject.LinearProblemMatrices = myVPop.LinearProblemMatrices;
                try
                    linearCalibrationObject = linearCalibrationObject.run('closeParallelPoolWhenFinished',false);
                    myVPop = linearCalibrationObject.OptimizedVPop;
                    myVPop.lambda = linearCalibrationObject.lambda;
                catch
                    myVPop.lambda = 0;
                end
                CEffN = 1/sum(myVPop.pws.^2);
                if abs(CEffN - targetEffN) < 10 || abs(Blambda-Alambda)<1e-3
                        break;
                end
                iter = iter+1;
                
                if CEffN>targetEffN
                    Blambda = Clambda;
                else
                    Alambda = Clambda;
                end  
            end
    end
end
end
```
