## Supplementary File 3 for "A surrogate workflow for virtual population development: designing an amenable proxy for a hypothesis-test inspired nonlinear goodness-of-fit": addPredTableVals.m

```
function vpop = addPredTableVals(vpop)
    % Once we have read the simData and generated a pw vector,
    % We need to add the predicted equivalents to the
    % tables used for statistical comparison.
    %
    % ARGUMENTS
    %  (self):      Note that the following properties should be
    %               initialized (experimental and simulation data)
    %               before calling this method:
    %                mnSDTable
    %                binTable
    %                distTable
    %                distTable2D
    %                corTable
    %                subpopTable
    %                simData
    %
    % RETURNS
    %  (self):      The VPop object is returned, but with updated
    %               properties:
    %                mnSDTable
    %                binTable
    %                distTable
    %                distTable2D
    %                corTable
    %                subpopTable

    myMnSdData = vpop.mnSDTable;
    myBinTable = vpop.binTable;
    myDistTable = vpop.distTable;
    myDistTable2D = vpop.distTable2D;
    myCorTable = vpop.corTable;
    mySubpopTable = vpop.subpopTable;

    vpIndicesSubpop = mySubpopTable.('vpIndices');

    mySimData = vpop.simData.Data;    
    myPWs = vpop.pws;
    mySimColNames = vpop.simData.rowInfoNames;

    mnSDRowsTarget = vpop.simData.mnSDRows;
    mnSDRowsSource = find(mnSDRowsTarget>0);

    binRowsTarget = vpop.simData.binRows;
    binRowsSource = find(binRowsTarget>0);

    distRowsTarget = vpop.simData.distRows;
    distRowsSource = find(distRowsTarget>0);

    simInterventionIDCol = find(ismember(mySimColNames, 'interventionID'));
    simTimeCol = find(ismember(mySimColNames, 'time'));
    simElementIDCol = find(ismember(mySimColNames, 'elementID'));
    simElementTypeCol = find(ismember(mySimColNames, 'elementType'));


    if ~isempty(mnSDRowsSource)
        mnSDRowsTarget = mnSDRowsTarget(mnSDRowsSource);

        mnSDRows = vpop.simData.mnSDRows;
        mnSDRows = mnSDRows(find(mnSDRows>0));
        mySubpopNo = myMnSdData.('subpopNo');

        [nMnSdRows, nMnSdCols] = size(myMnSdData);

        % 2 step assignment to speed execution
        % first to matrix, then convert back to table.
        curMean = nan(nMnSdRows,1);
        curPredN = nan(nMnSdRows,1);
        curSD = nan(nMnSdRows,1);
        curSimValues = nan(nMnSdRows,length(myPWs));
        curSimValues(mnSDRowsTarget,:) = (mySimData(mnSDRowsSource, :));
        keepIndices = myMnSdData.('predIndices');
        for rowCounter = 1 : nMnSdRows
            curPWs = myPWs(keepIndices{rowCounter}) / sum(myPWs(keepIndices{rowCounter}));
            if vpop.useEffN
                curN = 1/sum(curPWs.^2);
            else
                % We could use the PW cutoff here, but it seems this
                % encourages the optimizer to try to push the weight onto a
                % few VPs to decrease N.  Instead, let's use the number of
                % VPs for the purpose of statistical comparison, especially
                % during optimization.
                % curN = sum(myPWs >= obj.pwCutoff);
                curN = length(myPWs);
            end
            curPredN(rowCounter) = curN;
            curMean(rowCounter) = wtdMean(curSimValues(rowCounter,keepIndices{rowCounter}),curPWs);
            curSD(rowCounter) = wtdStd(curSimValues(rowCounter,keepIndices{rowCounter}),curPWs);
        end

        myMnSdData.('predN') = (curPredN);
        myMnSdData.('predMean') = (curMean);
        myMnSdData.('predSD') = (curSD);
        vpop.mnSDTable = myMnSdData;
    end

    if ~isempty(binRowsSource)
        binRowsTarget = binRowsTarget(binRowsSource);

        % 2 step assignment to speed execution
        % first to matrix, then convert back to table.
        [nBinRows, nBinCols] = size(myBinTable);
        curProbs = cell(nBinRows,1);
        curPredN = nan(nBinRows,1);
        curSimValues = nan(nBinRows,length(myPWs));
        curSimValues(binRowsTarget,:) = (mySimData(binRowsSource, :));
        binEdgeValues = myBinTable{:,'binEdges'};

        mySubpopNo = myBinTable.('subpopNo');
        keepIndices = myBinTable.('predIndices');
        for rowCounter = 1 : nBinRows
            curPWs = myPWs(keepIndices{rowCounter}) / sum(myPWs(keepIndices{rowCounter}));
            if vpop.useEffN
                curN = 1/sum(curPWs.^2);
            else
                % We could use the PW cutoff here, but it seems this
                % encourages the optimizer to try to push the weight onto a
                % few VPs to decrease N.  Also allow the number of
                % VPs for the purpose of statistical comparison, especially
                % during optimization.
                % curN = sum(myPWs >= obj.pwCutoff);
                curN = length(myPWs);
            end
            curPredN(rowCounter) = curN;
            curProbs{rowCounter} = wtdBinProb(curSimValues(rowCounter,keepIndices{rowCounter}), curPWs, binEdgeValues{rowCounter});
        end
        myBinTable.('predN') = (curPredN);
        myBinTable.('predBins') = curProbs;
        vpop.binTable = myBinTable;
    end

    if ~isempty(distRowsSource)
        [nDistRows, nDistCols] = size(myDistTable);
        assignPWs = cell(nDistRows,1);
        assignN = nan(nDistRows,1);
        keepIndices = myDistTable.('predIndices');


        for rowCounter = 1 : nDistRows
            % We have already found these
            curPWs = myPWs(keepIndices{rowCounter}) / sum(myPWs(keepIndices{rowCounter}));


            if vpop.useEffN
                curN = 1/sum(curPWs.^2);
            else
                % We could use the PW cutoff here, but it seems this
                % encourages the optimizer to try to push the weight onto a
                % few VPs to decrease N.  Also allow the number of
                % VPs for the purpose of statistical comparison, especially
                % during optimization.
                % curN = sum(myPWs >= obj.pwCutoff);
                curN = length(myPWs);
            end

            % Since we assign in multiple values per row for the
            % distribution, it looks like we have to loop this
            % All variables in the column must be same size
            assignN(rowCounter) = curN;
            assignPWs{rowCounter} = curPWs;
        end
        myDistTable.('predN') = (assignN);
        myDistTable.('predProbs') = (assignPWs);
        vpop.distTable = myDistTable;
    end

    if ~isempty(myDistTable2D)
        distRowsTarget1 = vpop.simData.distRows2D(:,1);
        distRowsTarget2 = vpop.simData.distRows2D(:,2);
        distRowsSource1 = find(~cellfun(@isempty,distRowsTarget1));
        distRowsSource2 = find(~cellfun(@isempty,distRowsTarget2));

        [nDistRows, nDistCols] = size(myDistTable2D);
        assignPWs = cell(nDistRows,1);
        assignN = nan(nDistRows,1);
        keepIndices = myDistTable2D.('predIndices');


        for rowCounter = 1 : nDistRows
            % We have already found these
            curPWs = myPWs(keepIndices{rowCounter}) / sum(myPWs(keepIndices{rowCounter}));


            if vpop.useEffN
                curN = 1/sum(curPWs.^2);
            else
                % We could use the PW cutoff here, but it seems this
                % encourages the optimizer to try to push the weight onto a
                % few VPs to decrease N.  Also allow the number of
                % VPs for the purpose of statistical comparison, especially
                % during optimization.
                % curN = sum(myPWs >= obj.pwCutoff);
                curN = length(myPWs);
            end

            % Since we assign in multiple values per row for the
            % distribution, it looks like we have to loop this
            % All variables in the column must be same size
            assignN(rowCounter) = curN;
            assignPWs{rowCounter} = curPWs;
        end
        myDistTable2D.('predN') = (assignN);
        myDistTable2D.('predProbs') = (assignPWs);
        vpop.distTable2D = myDistTable2D;
    end

    if ~isempty(myCorTable)
        corRowsTarget1 = vpop.simData.corRows(:,1);
        corRowsTarget2 = vpop.simData.corRows(:,2);
        corRowsSource1 = find(~cellfun(@isempty,corRowsTarget1));
        corRowsSource2 = find(~cellfun(@isempty,corRowsTarget2));
        [nCorRows, nCorCols] = size(myCorTable);
        assignPWs = cell(nCorRows,1);
        assignN = nan(nCorRows,1);
        curCor = nan(nCorRows,1);
        keepIndices = myCorTable.('predIndices');
        for rowCounter = 1 : nCorRows
            % We have already found these
            curPWs = myPWs(keepIndices{rowCounter}) / sum(myPWs(keepIndices{rowCounter}));

            if vpop.useEffN
                curN = 1/sum(curPWs.^2);
            else
                % We could use the PW cutoff here, but it seems this
                % encourages the optimizer to try to push the weight onto a
                % few VPs to decrease N.  Also allow the number of
                % VPs for the purpose of statistical comparison, especially
                % during optimization.
                % curN = sum(myPWs >= obj.pwCutoff);
                curN = length(myPWs);
            end

            % Since we assign in multiple values per row for the
            % distribution, it looks like we have to loop this
            % All variables in the column must be same size
            assignN(rowCounter) = curN;
            assignPWs{rowCounter} = curPWs;
            curwtdcorr = weightedcorrs(myCorTable.('predSample'){rowCounter}', curPWs');
            curCor(rowCounter) = curwtdcorr(1,2);
        end
        myCorTable.('predN') = (assignN);
        myCorTable.('predProbs') = (assignPWs);
        myCorTable.('predCor') = (curCor);
        vpop.corTable = myCorTable;
    end

    [nSubpopRows, nSubpopCols] = size(mySubpopTable);
    if nSubpopRows > 1
        curProbs = ones(nSubpopRows,1);
        for rowCounter = 2 : nSubpopRows
            curIndices = vpIndicesSubpop{rowCounter};
            curPWs = myPWs(curIndices);
            curProbs(rowCounter) = sum(curPWs);
        end
        mySubpopTable.('predW') = (curProbs);
        vpop.subpopTable = mySubpopTable;
    end
end
```
