## Supplementary File 3 for "A surrogate workflow for virtual population development: designing an amenable proxy for a hypothesis-test inspired nonlinear goodness-of-fit": addTableSimVals.m

```
function obj = addTableSimVals(obj)
    % Here we simply get sim values that are
    % fixed during optimization and add them to the
    % tables.  This is done at initialization to speed
    % execution.
    %
    % ARGUMENTS
    %  (self):      Note that the following properties should be
    %               initialized (experimental and simulation data)
    %               before calling this method:
    %                mnSDTable
    %                binTable
    %                distTable
    %                distTable2D
    %                simData
    %                corTable
    %                subpopTable
    %
    % RETURNS
    %  (self):      The VPop object is returned, but with updated
    %               properties:
    %                mnSDTable
    %                binTable
    %                distTable
    %                distTable2D
    %                simData
    %                corTable
    %                subpopTable


    mySubpopTable = obj.subpopTable;
    mySimData = obj.simData.Data;
    mySimRowInfo = obj.simData.rowInfo;
    mySimColNames = obj.simData.rowInfoNames;
    simInterventionIDCol = find(ismember(mySimColNames, 'interventionID'));
    simTimeCol = find(ismember(mySimColNames, 'time'));
    simElementIDCol = find(ismember(mySimColNames, 'elementID'));
    simElementTypeCol = find(ismember(mySimColNames, 'elementType'));


    myTable = obj.mnSDTable;
    rowsTarget = obj.simData.mnSDRows;
    rowsSource = find(rowsTarget>0);
    if ~isempty(rowsSource)
        mySubpopNo = myTable.('subpopNo');
        vpIndicesSubpop = mySubpopTable.('vpIndices');
        rowsTarget = rowsTarget(rowsSource);
        % 2 step assignment to speed execution
        % first to matrix, then convert back to table.
        [nRows, ~] = size(myTable);
        curSimValues = nan(nRows,  size(mySimData,2));
        curSimValues(rowsTarget,:) = (mySimData(rowsSource, :));
        for rowCounter = 1 : nRows
            subpopIndices = vpIndicesSubpop{mySubpopNo(rowCounter)};
            rowSimValues = curSimValues(rowCounter,:);
            keepIndices = intersect(find(~isnan(rowSimValues)),subpopIndices,'stable');
            [curVals, sortIndices] = sort(rowSimValues, 'ascend');
            keepIndices = intersect(sortIndices,keepIndices,'stable');
            %curVals = curSimValues(rowCounter,keepIndices);
            myTable.('predSample'){rowCounter} = rowSimValues(keepIndices);
            myTable.('predIndices'){rowCounter} = keepIndices;
        end
        obj.mnSDTable = myTable;
    end

    myTable = obj.binTable;
    rowsTarget = obj.simData.binRows;
    rowsSource = find(rowsTarget>0);
    if ~isempty(rowsSource)
        mySubpopNo = myTable.('subpopNo');
        vpIndicesSubpop = mySubpopTable.('vpIndices');
        rowsTarget = rowsTarget(rowsSource);
        % 2 step assignment to speed execution
        % first to matrix, then convert back to table.
        [nRows, ~] = size(myTable);
        curSimValues = nan(nRows,  size(mySimData,2));
        curSimValues(rowsTarget,:) = (mySimData(rowsSource, :));
        for rowCounter = 1 : nRows
            subpopIndices = vpIndicesSubpop{mySubpopNo(rowCounter)};
            rowSimValues = curSimValues(rowCounter,:);
            keepIndices = intersect(find(~isnan(rowSimValues)),subpopIndices,'stable');
            [curVals, sortIndices] = sort(rowSimValues, 'ascend');
            keepIndices = intersect(sortIndices,keepIndices,'stable');
            %curVals = curSimValues(rowCounter,keepIndices);
            myTable.('predSample'){rowCounter} = rowSimValues(keepIndices);
            myTable.('predIndices'){rowCounter} = keepIndices;
        end
        obj.binTable = myTable;
    end

    myTable = obj.distTable;
    rowsTarget = obj.simData.distRows;
    rowsSource = find(rowsTarget>0);
    if ~isempty(rowsSource)
        mySubpopNo = myTable.('subpopNo');
        vpIndicesSubpop = mySubpopTable.('vpIndices');
        rowsTarget = rowsTarget(rowsSource);

        % 2 step assignment to speed execution
        % first to matrix, then convert back to table.
        [nRows, nDistCols] = size(myTable);

        curSimValues = nan(nRows,  size(mySimData,2));
        curSimValues(rowsTarget,:) = (mySimData(rowsSource, :));
        curSimValues = nan(nRows,  size(mySimData,2));
        curSimValues(rowsTarget,:) = (mySimData(rowsSource, :));
        for rowCounter = 1 : nRows

            subpopIndices = vpIndicesSubpop{mySubpopNo(rowCounter)};
            % Should account for subpops
            keepIndices = find(~isnan(curSimValues(rowCounter,:)));

            subpopIndices = vpIndicesSubpop{mySubpopNo(rowCounter)};
            rowSimValues = curSimValues(rowCounter,:);
            keepIndices = intersect(find(~isnan(rowSimValues)),subpopIndices,'stable');
            [curVals, sortIndices] = sort(rowSimValues, 'ascend');
            keepIndices = intersect(sortIndices,keepIndices,'stable');
            curVals = rowSimValues(keepIndices);
            myTable.('predSample'){rowCounter} = curVals;
            myTable.('predIndices'){rowCounter} = keepIndices;

            % Also get the indices to align the exp and sim samples
            sample1 = myTable.('expSample'){rowCounter};
            [sample1Ind, sample2Ind, SC] = alignSamples(sample1, curVals);
            myTable.('expCombinedIndices'){rowCounter} = sample1Ind;
            myTable.('simCombinedIndices'){rowCounter} = sample2Ind;
            myTable.('combinedPoints'){rowCounter} = SC;

        end
        obj.distTable = myTable;
    end

    myTable = obj.distTable2D;
    rowsTarget1 = obj.simData.distRows2D(:,1);
    rowsTarget2 = obj.simData.distRows2D(:,2);
    rowsSource1 = find(~cellfun(@isempty,rowsTarget1));
    rowsSource2 = find(~cellfun(@isempty,rowsTarget2));
    if ~isempty(rowsSource1)
        mySubpopNo = myTable.('subpopNo');
        vpIndicesSubpop = mySubpopTable.('vpIndices');
        rowsTarget1 = rowsTarget1(rowsSource1);
        rowsTarget2 = rowsTarget2(rowsSource2);
        % 2 step assignment to speed execution
        % first to matrix, then convert back to table.
        [nDistRows, nDistCols] = size(myTable);
        curSimValues1 = nan(nDistRows,  size(mySimData,2));
        curSimValues2 = nan(nDistRows,  size(mySimData,2));
        % We need a loop for the assignment
        for target1counter = 1 :length(rowsTarget1)
            targetRows = rowsTarget1{target1counter};
            for target1repCounter = 1 :length(targetRows)
                curSimValues1(targetRows(target1repCounter),:) = (mySimData(rowsSource1(target1counter), :));
            end
        end
        for target2counter = 1 :length(rowsTarget2)
            targetRows = rowsTarget2{target2counter};
            for target2repCounter = 1 :length(targetRows)
                curSimValues2(targetRows(target2repCounter),:) = (mySimData(rowsSource2(target2counter), :));
            end
        end
        % Unlike the 1D case we won't sort
        for rowCounter = 1 : nDistRows
            keepIndices = find(~isnan(curSimValues1(rowCounter,:)) & ~isnan(curSimValues2(rowCounter,:)));
            subpopIndices = vpIndicesSubpop{mySubpopNo(rowCounter)};
            % Should account for subpops
            keepIndices = intersect(keepIndices,subpopIndices);

            curVals = [curSimValues1(rowCounter,keepIndices); curSimValues2(rowCounter,keepIndices)];
            myTable.('predSample'){rowCounter} = curVals;
            myTable.('predIndices'){rowCounter} = keepIndices;
        end
        obj.distTable2D = myTable;
    end

    myTable = obj.corTable;
    rowsTarget1 = obj.simData.corRows(:,1);
    rowsTarget2 = obj.simData.corRows(:,2);
    rowsSource1 = find(~cellfun(@isempty,rowsTarget1));
    rowsSource2 = find(~cellfun(@isempty,rowsTarget2));
    if ~isempty(rowsSource1)
        mySubpopNo = myTable.('subpopNo');
        vpIndicesSubpop = mySubpopTable.('vpIndices');
        rowsTarget1 = rowsTarget1(rowsSource1);
        rowsTarget2 = rowsTarget2(rowsSource2);
        % 2 step assignment to speed execution
        % first to matrix, then convert back to table.
        [nCorRows, nCorCols] = size(myTable);
        curSimValues1 = nan(nCorRows,  size(mySimData,2));
        curSimValues2 = nan(nCorRows,  size(mySimData,2));
        % We need a loop for the assignment
        for target1counter = 1 :length(rowsTarget1)
            targetRows = rowsTarget1{target1counter};
            for target1repCounter = 1 :length(targetRows)
                curSimValues1(targetRows(target1repCounter),:) = (mySimData(rowsSource1(target1counter), :));
            end
        end
        for target2counter = 1 :length(rowsTarget2)
            targetRows = rowsTarget2{target2counter};
            for target2repCounter = 1 :length(targetRows)
                curSimValues2(targetRows(target2repCounter),:) = (mySimData(rowsSource2(target2counter), :));
            end
        end
        % Unlike the 1D case we won't sort
        for rowCounter = 1 : nCorRows
            keepIndices = find(~isnan(curSimValues1(rowCounter,:)) & ~isnan(curSimValues2(rowCounter,:)));
            subpopIndices = vpIndicesSubpop{mySubpopNo(rowCounter)};
            % Should account for subpops
            keepIndices = intersect(keepIndices,subpopIndices);

            curVals = [curSimValues1(rowCounter,keepIndices); curSimValues2(rowCounter,keepIndices)];
            myTable.('predSample'){rowCounter} = curVals;
            myTable.('predIndices'){rowCounter} = keepIndices;

        end
        obj.corTable = myTable;
    end

end
```
