## Supplementary File 3 for "A surrogate workflow for virtual population development: designing an amenable proxy for a hypothesis-test inspired nonlinear goodness-of-fit": assignCoeffs.m

```
function obj = assignCoeffs(obj, myWorksheet)
% Copy the VP coefficients from a worksheet to the `coeffsTable` property of the
% VPop object. 
% 
% EXAMPLE
% >> myVPop.assignCoeffs(myWorksheet)
%
% SEE ALSO
% VPop.coeffsTable
    mycoeffsTable = getVPCoeffs(myWorksheet);
    obj.coeffsTable = mycoeffsTable;
end
```
