## Supplementary File 3 for "A surrogate workflow for virtual population development: designing an amenable proxy for a hypothesis-test inspired nonlinear goodness-of-fit": assignIndices.m

```
function obj = assignIndices(obj, myWorksheet, myMapelOptions)
    % This method takes a worksheet and mapelOptions structure and
    % creates a bin table and other poperties needed for the virtual
    % population and MAPEL algorithm.
    %
    % ARGUMENT
    %  (self)
    %  myWorksheet:    a worksheet structure, with VPs, axes, and
    %                  coefficients
    %  myMapelOptions: an instace of mapelOptions, with
    %                  properties
    %                   nBins
    %                   equalBinBreaks
    %
    % RETURNS
    %  (self): properties of the virtual population are updated:
    %           indexTable
    %           binEdges
    %           binMidPoints
    %
    % SEE ALSO
    % VPop.indexTable, VPop.binEdges, VPop.binMidPoints
    % mapelOptions

    myVPCoeffs = getVPCoeffs(myWorksheet);
    [nAxis, nVP] = size(myVPCoeffs);
    myIndexTable = ones(nAxis, nVP);
    nBins = myMapelOptions.nBins;
    myBinEdges = nan(nAxis, (nBins+1));
    myBinMidPoints = nan(nAxis, nBins);
    equalBinBreaks = myMapelOptions.equalBinBreaks;
    for axisCounter = 1 : nAxis
        % We currently only implement continuous variables from
        % the paper/R MAPEL
        if equalBinBreaks
            % Rather than scaling based on the sampled min and
            % max as in the R MAPEL algorithm, we adjust according
            % to the allowed,
            % and this range is [0, 1] by the axis definition.
            myBinEdges(axisCounter,:) =  (0:1/nBins:1);
        else
            myPercentiles =  (0:1/nBins:1)';
            myBinEdges(axisCounter,:) = (quantile(myVPCoeffs(axisCounter,:),myPercentiles));
        end
        % We'll use the cdf convention that FX(x) = P(X <= x), so
        % check if a value is <= the bin upper cutoff in order to
        % assign it.
        % Note that effectively we want to ignore the first bin of 0
        % as a cutoff and lump this in with the first bin.
        for binCounter = 1 : nBins
            myBinMidPoints(axisCounter,binCounter) = (myBinEdges(axisCounter,binCounter) + myBinEdges(axisCounter,binCounter+1))/2;
            % First bins are assigned an index of 0 by this method.
            myIndexTable(axisCounter, :) = myIndexTable(axisCounter, :) + (myVPCoeffs(axisCounter, :) > myBinEdges(axisCounter,binCounter+1));
        end
    end
    obj.indexTable = myIndexTable;
    obj.binEdges = myBinEdges;
    obj.binMidPoints = myBinMidPoints;
end
```
