## Supplementary File 3 for "A surrogate workflow for virtual population development: designing an amenable proxy for a hypothesis-test inspired nonlinear goodness-of-fit": assignPWs.m

```
function obj = assignPWs(obj)
    % Update prevalence weight assignment for the virtual population.
    % Note that existing predicted population results will be set
    % to NaN in:
    %  mnSDTable
    %  binTable
    %  distTable
    %  distTable2D
    %  corTable
    %
    % And also the following properties will be reset:
    %  gofMn
    %  gofSD
    %  gofBin
    %  gofDist
    %  gofDist2D
    %  gofCor
    %  gof
    %
    % Only relevant for obj.pwStrategy = 'axis'
    % ARGUMENTS:
    %  (self): No additional arguments, but the VPop object must have
    %          previously assigned the following properties:
    %           indexTable
    %           binProbs
    %
    % RETURNS:
    %  (self): Returns the VPop with the updated PW properties.
    %
    myIndexTable = obj.indexTable;
    [myNAxis, myNVP] = size(myIndexTable);
    myPWs = zeros(1,myNVP);
    myBinProbs = obj.binProbs;
    %pwCutoff = obj.pwCutoff;
    for axisCounter = 1 : myNAxis
        % Note this will result in -Inf if any bins have
        % prob of zero.  MATLAB's exponential at the end corrects
        % this, but there is a danger of getting back all
        % -inf if all of the bin probabilities are small.
        myPWs = myPWs + log(myBinProbs(axisCounter,myIndexTable(axisCounter,:)));
    end
    % Carried over from paper MAPEL: force
    % biggest value to ~1 to avoid underflow
    myPWs = myPWs - max(myPWs);
    myPWs = exp(myPWs); %/ sum(exp(myPWs));
    %myPWs = myPWs.*(myPWs >= pwCutoff);
    myPWs = myPWs / sum(myPWs);
    obj.pws = myPWs;

    % If we re-calculate PWs, eliminate any
    % previous PW-dependent properties - i.e. those
    % that start with "pred" except predIndices
    myMnSDTable = obj.mnSDTable;
    [nRows, nCols] = size(myMnSDTable);
    if nRows > 0
        % myString = 'pred';
        % varNames = myMnSDTable.Properties.VariableNames;
        % myPos = find(strncmpi(myString,varNames,length(myString)));
        % nanifyVars = varNames(myPos);
        nanifyVars = {'predN','predMean','predSD'};
        for varCounter = 1 : length(nanifyVars)
            myMnSDTable.(nanifyVars{varCounter}) = nan(nRows,1);
        end
    end
    obj.mnSDTable = myMnSDTable;

    myBinTable = obj.binTable;
    [nRows, nCols] = size(myBinTable);
    if nRows > 0
        % myString = 'pred';
        % varNames = myBinTable.Properties.VariableNames;
        % myPos = find(strncmpi(myString,varNames,length(myString)));
        % nanifyVars = varNames(myPos);
        nanifyVars = {'predN','predBins'};
        for varCounter = 1 : length(nanifyVars)
            myBinTable.(nanifyVars{varCounter}) = nan(nRows,1);
        end
    end
    obj.binTable = myBinTable;

    myDistTable = obj.distTable;
    [nRows, nCols] = size(myDistTable);
    if nRows > 0
        myNanStrings = {'predN'};
        varNames = myDistTable.Properties.VariableNames;
        myPos = find(ismember(varNames,myNanStrings));
        nanifyVars = varNames(myPos);
        for varCounter = 1 : length(nanifyVars)
            myDistTable.(nanifyVars{varCounter}) = nan(nRows,1);
        end
        myNanStrings = {'predProbs'};
        varNames = myDistTable.Properties.VariableNames;
        myPos = find(ismember(varNames,myNanStrings));
        nanifyVars = varNames(myPos);
        myDistTable{1 : nRows,nanifyVars} = {nan};
    end
    obj.distTable = myDistTable;

    myTable = obj.distTable2D;
    if ~isempty(myTable)
        [nRows, nCols] = size(myTable);
        if nRows > 0
            myNanStrings = {'predN'};
            varNames = myTable.Properties.VariableNames;
            myPos = find(ismember(varNames,myNanStrings));
            nanifyVars = varNames(myPos);
            for varCounter = 1 : length(nanifyVars)
                myTable.(nanifyVars{varCounter}) = nan(nRows,1);
            end
            myNanStrings = {'predProbs'};
            varNames = myTable.Properties.VariableNames;
            myPos = find(ismember(varNames,myNanStrings));
            nanifyVars = varNames(myPos);
            myTable{1 : nRows,nanifyVars} = {nan};
        end
        obj.distTable2D = myTable;
    end

    myTable = obj.corTable;
    if ~isempty(myTable)
        [nRows, nCols] = size(myTable);
        if nRows > 0
            myNanStrings = {'predN','predCor'};
            varNames = myTable.Properties.VariableNames;
            myPos = find(ismember(varNames,myNanStrings));
            nanifyVars = varNames(myPos);
            for varCounter = 1 : length(nanifyVars)
                myTable.(nanifyVars{varCounter}) = nan(nRows,1);
            end
            myNanStrings = {'predProbs'};
            varNames = myTable.Properties.VariableNames;
            myPos = find(ismember(varNames,myNanStrings));
            nanifyVars = varNames(myPos);
            myTable{1 : nRows,nanifyVars} = {nan};
        end
        obj.corTable = myTable;
    end

    myTable = obj.subpopTable;
    [nRows, nCols] = size(myTable);
    if nRows > 1
        nanifyVars = 'predW';
        myTable.(nanifyVars{varCounter})(2:end) = nan(nRows-1,1);
    end
    obj.subpopTable = myTable;

    obj.gofMn = [];
    obj.gofSD = [];
    obj.gofBin = [];
    obj.gofDist = [];
    obj.gofDist2D = [];
    obj.gofCor = [];
    obj.gof = [];
end
```
