## Supplementary File 3 for "A surrogate workflow for virtual population development: designing an amenable proxy for a hypothesis-test inspired nonlinear goodness-of-fit": get.m

```
function value = get(obj, propName)
    % value = get(vpop, propName) get the value of property `propName` from the virtual
    % population object.
    switch propName
        case 'coeffsTable'
            value = obj.coeffsTable;
        case 'coeffsDist'
            value = obj.coeffsDist;
        case 'pwStrategy'
            value = obj.pwStrategy;
        case 'indexTable'
            value = obj.indexTable;
        case 'binEdges'
            value = obj.binEdges;
        case 'binMidPoints'
            value = obj.binMidPoints;
        case 'binProbs'
            value = obj.binProbs;
        case 'pws'
            value = obj.pws;
        case 'mnSDTable'
            value = obj.mnSDTable;
        case 'binTable'
            value = obj.binTable;
        case 'distTable'
            value = obj.distTable;
        case 'distTable2D'
            value = obj.distTable2D;
        case 'corTable'
            value = obj.corTable;
        case 'subpopTable'
            value = obj.subpopTable;
        case 'expData'
            value = obj.expData;
        case 'simData'
            value = obj.simData;
        case 'gofMn'
            value = obj.gofMn;
        case 'gofSD'
            value = obj.gofSD;
        case 'gofBin'
            value = obj.gofBin;
        case 'gofDist'
            value = obj.gofDist;
        case 'gofDist2D'
            value = obj.gofDist2D;
        case 'gofCor'
            value = obj.gofCor;
        case 'gof'
            value = obj.gof;
        case 'spreadOut'
            value = obj.spreadOut;
        case 'minIndPVal'
            value = obj.minIndPVal;
        case 'useEffN'
            value = obj.useEffN;
        case 'exactFlag'
            value = obj.exactFlag;
        case 'optimizeTimeLimit'
            value = obj.optimizeTimeLimit;
        case 'optimizeType'
            value = obj.optimizeType;
        case 'optimizePopSize'
            value = obj.optimizePopSize;
        case 'objectiveLimit'
            value = obj.objectiveLimit;
        case 'poolRestart'
            value = obj.poolRestart;
        case 'poolClose'
            value = obj.poolClose;
        case 'intSeed'
            value = obj.intSeed;
        case 'tol'
            value = obj.tol;
        case 'nIters'
            value = obj.nIters;
        case 'minEffN'
            value = obj.minEffN;
        otherwise
            error(['Error: ',propName ,' is not a valid ',mfilename,' property.'])
    end
end
```
