## Supplementary File 3 for "A surrogate workflow for virtual population development: designing an amenable proxy for a hypothesis-test inspired nonlinear goodness-of-fit": getSimData.m

```
function obj = getSimData(obj, myWorksheet)
    % Get all of the simulation datapoints for the VPs that will
    % be needed for calculating population statistics
    %
    % ARGUMENTS
    %  (self):      Note that the following properties should be
    %               initialized (experimental data) before calling this
    %               method:
    %                mnSDTable
    %                binTable
    %                distTable
    %                distTable2D
    %				corTable
    %  myWorksheet: A worksheet with the simulation results.
    %
    % RETURNS
    %  (self):      The VPop object is returned, but with an updated
    %               simData property.
    %
    %
    continueFlag = true;
    mnsdDataFlag = false;
    binDataFlag = false;
    distDataFlag = false;
    distData2DFlag = false;
    corDataFlag = false;
    myMnSdData = obj.mnSDTable;
    myBinTable = obj.binTable;
    myDistTable = obj.distTable;
    myDistTable2D = obj.distTable2D;
    myCorTable = obj.corTable;
    % We want the {expVarID, element, elementType, time} sets
    % from the experimental summaries that will be paired
    % with real data.
    rowInfoNames = {'expVarID','interventionID','elementID','elementType','time','expDataID'};
    rowInfoNames2D = {'expVarID1','expVarID2','interventionID1','interventionID2','elementID1','elementID2','elementType1','elementType2','time1','time2','expDataID1','expDataID2'};
    rowInfoNames2D1 = {'expVarID1','interventionID1','elementID1','elementType1','time1','expDataID1'};
    rowInfoNames2D2 = {'expVarID2','interventionID2','elementID2','elementType2','time2','expDataID2'};
    brInfoNames = {'expVarID','interventionID','elementID','elementType','time'};
    rInfoNames = brInfoNames;
    myDataSource = cell2table(cell(0,6), 'VariableNames', rowInfoNames);

    %            rowInfoNames = {'expVarID','interventionID','elementID','elementType','time'};
    % 		   rowInfoNames2D = {'expVarID1','expVarID2','interventionID1','interventionID2','elementID1','elementID2','elementType1','elementType2','time1','time2'};
    %            rowInfoNames2D1 = {'expVarID1','interventionID1','elementID1','elementType1','time1'};
    %            rowInfoNames2D2 = {'expVarID2','interventionID2','elementID2','elementType2','time2'};
    %            myDataSource = cell2table(cell(0,5), 'VariableNames', rowInfoNames);
    if ~isempty(myMnSdData)
        myDataSource = [myDataSource; myMnSdData(:,rowInfoNames)];
        mnsdDataFlag = true;
    end
    if ~isempty(myBinTable)
        myDataSource = [myDataSource; myBinTable(:,rowInfoNames)];
        binDataFlag = true;
    end
    if ~isempty(myDistTable)
        myDataSource = [myDataSource; myDistTable(:,rowInfoNames)];
        distDataFlag = true;
    end
    if ~isempty(myDistTable2D)
        distData2DFlag = true;
        myDataSource1 = myDistTable2D(:,rowInfoNames2D1);
        myDataSource2 = myDistTable2D(:,rowInfoNames2D2);
        myDataSource1.Properties.VariableNames = rowInfoNames;
        myDataSource2.Properties.VariableNames = rowInfoNames;
        myDataSource = [myDataSource; myDataSource1];
        myDataSource = [myDataSource; myDataSource2];
    end
    if ~isempty(myCorTable)
        corDataFlag = true;
        myDataSource1 = myCorTable(:,rowInfoNames2D1);
        myDataSource2 = myCorTable(:,rowInfoNames2D2);
        myDataSource1.Properties.VariableNames = rowInfoNames;
        myDataSource2.Properties.VariableNames = rowInfoNames;
        myDataSource = [myDataSource; myDataSource1];
        myDataSource = [myDataSource; myDataSource2];
    end
    [nrows,ncols] = size(myDataSource);
    if nrows < 1
        warning(['No mnSDTable, binTable, distTable, distTable2D, or corTable assigned before calling getSimData in ',mfilename,'. The data to get is not known. Exiting.'])
        continueFlag = false;
    end
    if continueFlag
        % Eliminate duplicate rows
        myDataSource = unique(myDataSource, 'rows');
        myDataSource = sortrows(myDataSource, ...
            {'interventionID','elementID','time','elementType','expDataID'},...
            {'ascend','ascend','ascend','ascend','ascend'});
    end

    if continueFlag
        myCheckVars = myDataSource.('elementID');
        % If the length is 0, all variables should be written.
        if length(myWorksheet.simProps.saveElementResultIDs) > 0
            myMissingVars = ~ismember(myCheckVars,myWorksheet.simProps.saveElementResultIDs);
            myMissingVars = myCheckVars(myMissingVars);
            if length(myMissingVars) > 0
                disp(['Missing needed variables for ',mfilename,' in myWorksheet.simProps.saveElementResultIDs.  They are: ',strjoin(myMissingVars,', '),'.  Attempting to continue...'])
            end
        end
    end

    if continueFlag
        [nEntries, ~] = size(myDataSource);

        % Info on each row of simvalues
        rowInfo = table2cell(myDataSource);
        % We will need to use a structure, like we did for results,
        % since some of the VP names will certainly be longer
        % than what is permitted by variable name lengths
        % Also, use the values
        % Row indices for the data are also added to speed subsequent calculations.

        vpIDs = getVPIDs(myWorksheet);
        nVPs = length(vpIDs);
        % Simulation results for each VP for each row
        dataValues = nan(nEntries, length(vpIDs));
        interventionIDIndex = find(ismember(rowInfoNames,'interventionID'));
        elementIDIndex = find(ismember(rowInfoNames,'elementID'));
        elementTypeIndex = find(ismember(rowInfoNames,'elementType'));
        expTimeIndex = find(ismember(rowInfoNames,'time'));
        expDataIDIndex = find(ismember(rowInfoNames,'expDataID'));
        simData.Data = dataValues;
        simData.rowInfoNames = rowInfoNames;
        simData.rowInfo = rowInfo;
        simData.vpIDs = vpIDs;
        interventionIDs = getInterventionIDs(myWorksheet);
        mnSDRows = nan(nEntries,1);
        binRows = nan(nEntries,1);
        distRows = nan(nEntries,1);
        distRows2D = cell(nEntries,2);
        corRows = cell(nEntries,2);

        for rowCounter = 1 : nEntries
            interventionID = simData.rowInfo{rowCounter,interventionIDIndex};
            elementID = simData.rowInfo{rowCounter,elementIDIndex};
            elementType = simData.rowInfo{rowCounter,elementTypeIndex};
            wshInterventionIndex = find(ismember(interventionIDs,interventionID));
            expTime = simData.rowInfo{rowCounter,expTimeIndex};
            expDataID = simData.rowInfo{rowCounter,expDataIDIndex};
            % To avoid re-searching for the right rows on every
            % iteration mapel, we provide the indices here
            if mnsdDataFlag
                temp = find((ismember(myMnSdData{:,'interventionID'},interventionID)) ...
                    & ((myMnSdData{:,'time'})==expTime) ...
                    & (ismember(myMnSdData{:,'elementID'},elementID)) ...
                    & (ismember(myMnSdData{:,'elementType'},elementType) ...
                    & (ismember(myMnSdData{:,'expDataID'},expDataID))));
                if ~isempty(temp)
                    mnSDRows(rowCounter) = temp;
                end
            end
            if binDataFlag
                temp = find((ismember(myBinTable{:,'interventionID'},interventionID)) ...
                    & ((myBinTable{:,'time'})==expTime) ...
                    & (ismember(myBinTable{:,'elementID'},elementID)) ...
                    & (ismember(myBinTable{:,'elementType'},elementType)) ...
                    & (ismember(myBinTable{:,'expDataID'},expDataID)));
                if ~isempty(temp)
                    binRows(rowCounter) = temp;
                end
            end
            if distDataFlag
                temp = find((ismember(myDistTable{:,'interventionID'},interventionID)) ...
                    & ((myDistTable{:,'time'})==expTime) ...
                    & (ismember(myDistTable{:,'elementID'},elementID)) ...
                    & (ismember(myDistTable{:,'elementType'},elementType)) ...
                    & (ismember(myDistTable{:,'expDataID'},expDataID)));
                if ~isempty(temp)
                    distRows(rowCounter) = temp;
                end
            end
            if distData2DFlag
                % One row of source data may be involved in
                % multiple 2D distributions
                temp = find((ismember(myDistTable2D{:,'interventionID1'},interventionID)) & ((myDistTable2D{:,'time1'})==expTime) & (ismember(myDistTable2D{:,'elementID1'},elementID)) & (ismember(myDistTable2D{:,'elementType1'},elementType)));
                if ~isempty(temp)
                    distRows2D{rowCounter,1} = temp;
                end
                temp = find((ismember(myDistTable2D{:,'interventionID2'},interventionID)) & ((myDistTable2D{:,'time2'})==expTime) & (ismember(myDistTable2D{:,'elementID2'},elementID)) & (ismember(myDistTable2D{:,'elementType2'},elementType)));
                if ~isempty(temp)
                    distRows2D{rowCounter,2} = temp;
                end
            end
            if corDataFlag
                % One row of source data may be involved in
                % multiple 2D distributions
                temp = find((ismember(myCorTable{:,'interventionID1'},interventionID)) & ((myCorTable{:,'time1'})==expTime) & (ismember(myCorTable{:,'elementID1'},elementID)) & (ismember(myCorTable{:,'elementType1'},elementType)));
                if ~isempty(temp)
                    corRows{rowCounter,1} = temp;
                end
                temp = find((ismember(myCorTable{:,'interventionID2'},interventionID)) & ((myCorTable{:,'time2'})==expTime) & (ismember(myCorTable{:,'elementID2'},elementID)) & (ismember(myCorTable{:,'elementType2'},elementType)));
                if ~isempty(temp)
                    corRows{rowCounter,2} = temp;
                end
            end
            % I don't see how to avoid looping this, given the way
            % the data are structured.  Luckily we just get the data
            % once
            rowIndex = nan;
            for vpCounter = 1 : nVPs
                % We need to verify the desired result for the VP is
                % present, otherwise we report the simData result as
                % nan
                if length(myWorksheet.results) > 0
                    curResult = myWorksheet.results{wshInterventionIndex, vpCounter};
                    % Results should be stored in a structure, we
                    % assume if a structure is provided then it is a
                    % valid result
                    if strcmp(class(curResult),'struct')
                        curResult = myWorksheet.results{wshInterventionIndex, vpCounter};
                        curTimeIndex = find(ismember(curResult.Names,'time'));
                        curVarIndex = find(ismember(curResult.Names,elementID));
                        curTime = curResult.Data(:,curTimeIndex);
                        curVar = curResult.Data(:,curVarIndex);
                        % First check to see if the right row is
                        % already known.  We assume the result
                        % size is consistent across VPs.
                        if isnan(rowIndex)
                            if sum(curTime == expTime) == 1
                                rowIndex = find(curTime == expTime);
                                interpolateValue = curVar(rowIndex);
                            else
                                interpolateValue = interp1(curTime,curVar,expTime,'linear');
                                warning([num2str(expTime),' not found in the worksheet result, interpolation implemented ...']);
                            end
                        else
                            interpolateValue = curVar(rowIndex);
                        end
                        simData.Data(rowCounter, vpCounter) = interpolateValue;
                    else
                        simData.Data(rowCounter, vpCounter) = nan;
                    end
                else
                    simData.Data(rowCounter, vpCounter) = nan;
                end
            end
        end
        simData.binRows = binRows;
        simData.mnSDRows = mnSDRows;
        simData.distRows = distRows;
        simData.distRows2D = distRows2D;
        simData.corRows = corRows;
        obj.simData = simData;
    end
end
```
