## Supplementary File 3 for "A surrogate workflow for virtual population development: designing an amenable proxy for a hypothesis-test inspired nonlinear goodness-of-fit": startProbs.m

```
function obj = startProbs(obj, myRandomStart)
    % This method initializes bin probabilities, and is used
    % prior to the optization.
    % Only relevant for obj.pwStrategy = 'bin'
    %
    % ARGUMENTS
    %  (self)
    %  myRandomStart:  A boolean variable (true/false), if true
    %                  the bin probabilities will be set
    %                  randomly and if false they will be uniform
    if nargin < 1
        myRandomStart = false;
    end
    myBinMidPoints = obj.binMidPoints;
    [nAxes, nBins] = size(myBinMidPoints);
    if ~myRandomStart
        myUniformStartProbs = ones(nAxes, nBins) ./ nBins;
    else
        myUniformStartProbs = rand([nAxes, nBins]);
        for axisCounter = 1 : nAxes
            myUniformStartProbs(axisCounter,:) = myUniformStartProbs(axisCounter,:)/sum(myUniformStartProbs(axisCounter,:));
        end
    end
    obj.binProbs = myUniformStartProbs;
end
```
