## Supplementary File 3 for "A surrogate workflow for virtual population development: designing an amenable proxy for a hypothesis-test inspired nonlinear goodness-of-fit": VPop.m

```
classdef VPop
    % This is a virtual population class.  This contains the results of a call
    % to MAPEL. There should be too much here to that needs to be changed
    % directly, with the biggest possible exception being adjusting useEffN
    % before re-evaluating the GOF.
    %
    % BRIEF DESCRIPTIONS OF PROPERTIES
    %  (Type `help VPop.<propertyName>` for detailed descriptions)
    %
    %  coeffsTable:  (Don't manipulate) An nAxis x nVP matrix of VP coefficients.
    %
    %  coeffsDist:   (Don't manipulate) A nVP X vVP matrix with VP the
    %                 distance for the axis coefficients
    %
    %  pwStrategy:	 Strategy for the optimization.  
    %
    %  indexTable:   (Don't manipulate) A nAxis X nVP table that indicates in which 
    %                 axis bin each VP falls into.  
    %
    %  binEdges:     (Don't manipulate) Edges for the axis bins. 
    %
    %  binMidPoints: (Don't manipulate) Mid points for the axis bins, populated
    %                 by the assignIndices method.  ONLY NEEDED FOR BIN PWSTRATEGY
    %
    %  binProbs:     (Don't manipulate) Probabilities for each axis bin.  ONLY NEEDED FOR BIN PWSTRATEGY
    %
    %  pws:          (Don't manipulate) A 1xnVP vector of prevalence weights,
    %                which are calculated based on the assignPWs.
    %
    %  coefficients: a nAxisxnVP matrix of VP parameter coefficients.
    %
    %  expData:      (Don't manipulate) A table of experimental data used to
    %                 guide optimization.  
    %
    %  simData:      (Don't manipulate) A table of simulation results that 
    %                 pairs to the experimental data.  
    %
    %  mnSDTable:    (Don't manipulate) Summary table of experimental and 
    %                simulated (weighted) mean and standard deviation.  
    %
    %  binTable:     (Don't manipulate) Summary table of experimental and  
    %                simulated binned (weighted) data for a virtual population
    %                
    %  distTable:    (Don't manipulate) Summary table of experimental and  
    %                simulated distribution data for a virtual population
    %                				
    %  distTable2D:	 (Don't manipulate) A table to enable calibrating 2D distributions
    %
    %  corTable:	 (Don't manipulate) A table to enable calibrating pairwise correlations
    %
    %  subpopTable:  (Don't manipulate) Contains criteria to create subpopulations
    %                 from simulated VPs.				  
    %
    %  gofMn:        (Don't manipulate) Goodness of fit statistics for 
    %                individual endpoint/time mean comparisons between data
    %                and virtual population.  
    %
    %  gofSD:        (Don't manipulate) Goodness of fit statistics for 
    %                individual endpoint/time standard deviation comparisons  
    %                between data and virtual population. 
    %
    %  gofBin:       (Don't manipulate) Goodness of fit statistics for 
    %                individual endpoint/time bin distribution comparisons 
    %                between data and virtual population.  
    %
    %  gofDist:      (Don't manipulate) Goodness of fit statistics for 
    %                empirical distribution comparisons 
    %                between data and virtual population.
    %
    %  gofCor:		 (Don't manipulate) Goodness of fit statistics for comparisons between
    %                correlations estimated from data and from virtual population
    %                simulation.
    %
    %  gofDist2D:     (Don't manipulate) Goodness of fit statistics for
    %                 comparisons between bivariate joint distributions
    %                 estimated from data and from virtual population
    %                 simulation.
    %
    %  gof:          (Don't manipulate) Composite goodness-of-fit result.    
    %
    %  useEffN:      Whether or not to use the effN for optimization and
    %                evaluation of the GOF. 
    %
    %  exactFlag:    Whether to check to apply Fisher's exact test instead
    %                of the chi-square approximation for binned distribution
    %                test. 
    %
    %  spreadOut:    Used for optimization, if nonzero, will  
    %                apply a penalty proportional to spreadOut to increase how
    %                the prevalence weight is distributed.
    %
    %  minIndPVal:   Used for optimization, if nonzero, will  
    %                apply a large penalty if any of the individual pvalues.
    %                fall below the target.
    %
    %  optimizeTimeLimit:   Time limit for optimizing the VPop in s.
    %
    %  optimizeType:     Type of optimization algorithm to employ.  Default is
    %                    "pso". Other options: 'ga', 'pso', 'papso', 'gapso',
    %                    'gapapso', 'simplex', 'surrogate'.
    %
    %  optimizePopSize:     Number of solutions to try in each optimization
    %                       generation. Default is 1000.
    %
    %  objectiveLimit:		stopping condition for optimization.
    %
    %  poolClose:           Whether to close the pool at the end of the optimization.
    %
    %  poolRestart:         Whether to restart the pool at the beginning of the optimization.
    %
    %  intSeed:             A non-negative integer seed to initialize the 
    %                       random number generator.  Set to -1 to avoid
    %                       changing the state of the random number generator.
    %
    %  tol:                 Numerical tolerance for optimization.
    %
    %  nIters:             (Optional) Maximum number of outer-loop iterations in optimization. 
    %                       Default is 10,000.
    %
    %  minEffN:             Minimum effective N. The default is 0.
    %
    %  MSE:                  mean sum of residuals from linear calibration
    %  (subpopulation/dropouts weighted).
    %
    %  LinearProblemMatrices:  the linearMatrix made from oldVPop linearCalibrationObject, actually this will be the reduced matrix for the rescaled oldVPop with only nVPmax columns.
    %
    %  LinearProblemMatricesSubgroupSumWeights: the subweight directly recorded
    %                           fom oldVPop linearCalibrationObject.
    %
    %  LinearProblemMatricesobservationDescriptions: the subweightDescription
    %                            directly fom oldVPop linearCalibrationObject,
    %                            to keep track of biomarkers during cohort
    %                            expansion.
    %
    %
    % SEE ALSO
    % mapel

    properties (SetAccess = public, GetAccess = public)
        % (Don't manipulate) Table of VP coefficients. A nAxis X nVP matrix with
        % VP the coefficients used for prevalence weights with the bin pw
        % strategy.
        coeffsTable

        % (Don't manipulate) A nVP X vVP matrix with VP the distance for the
        % axis coefficients
        coeffsDist

        % Strategy for the optimization.  
        % Allowed values:
        %  'direct' (default value) If this is set then PWs are optimized directly. 
        %  'bin'     specified in a strategy similar to the original MAPEL
        %  algorithm.
        pwStrategy

        % (Don't manipulate) A nAxis X nVP table that indicates in which
        % axis bin each VP falls into.  This is usually populated at the
        % beginning of the MAPEL algorithm by the assignIndices method.
        % ONLY NEEDED FOR BIN PWSTRATEGY
        indexTable

        % (Don't manipulate) Edges for the axis bins. These are also
        % populated by the assignIndices method.
        binEdges 

        % (Don't manipulate) Mid points for the axis bins, populated by the
        % assignIndices method.  ONLY NEEDED FOR BIN PWSTRATEGY
        binMidPoints

        %(Don't manipulate) Probabilities for each axis bin.  These are directly
        %varied in the optimization, and the results are used to calculate the
        %individual prevalence weights.  ONLY NEEDED FOR BIN PWSTRATEGY
        binProbs

        % (Don't manipulate) A 1xnVP vector of prevalence weights, which are
        % calculated based on the assignPWs.
        pws

        %(Don't manipulate) A table of experimental data used to guide
        %optimization.  It is usually taken from the mapelOptions.
        expData

        % (Don't manipulate) A table of simulation results that pairs to the
        % experimental data.  Usually assigned in MAPEL.
        simData

        % (Don't manipulate) Summary table of experimental and simulated
        % (weighted) mean and standard deviation.  Usually copied from the
        % mapelOptions and updated with the weighted simulation results from the
        % VPop fitting. If you want to adjust a fit and restart, you may want to
        % adjust some of the weights here.
        mnSDTable

        % (Don't manipulate) Summary table of experimental and simulated binned
        % (weighted) data for a virtual population Usually copied from the
        % mapelOptions and updated with the weighted simulation results from the
        % VPop fitting. If you want to adjust a fit and restart, you may want to
        % adjust some of the weights here.
        binTable

        % (Don't manipulate) Summary table of experimental and simulated
        % distribution data for a virtual population Usually copied from the
        % mapelOptions and updated with the weighted simulation results from the
        % VPop fitting. If you want to adjust a fit and restart, you may want to
        % adjust some of the weights here.
        distTable

        % (Don't manipulate) A table to enable calibrating 2D distributions
        distTable2D
        
        % (Don't manipulate) A table to enable calibrating pairwise correlations
        corTable

        % (Don't manipulate) Contains criteria to create subpopulations from
        % simulated VPs
        subpopTable

        % (Don't manipulate) Goodness of fit statistics for individual
        % endpoint/time mean comparisons between data and virtual population.
        % Usually calcualated by the evaluateGOF function.
        gofMn

        % (Don't manipulate) Goodness of fit statistics for individual
        % endpoint/time standard deviation comparisons between data and virtual
        % population.  Usually calculated by the evaluateGOF function.
        gofSD

        % (Don't manipulate) Goodness of fit statistics for individual
        % endpoint/time bin distribution comparisons between data and virtual
        % population.  Usually calculated by the evaluateGOF function.
        gofBin

        % (Don't manipulate) Goodness of fit statistics for empirical
        % distribution comparisons between data and virtual population.  Usually
        % calculated by the evaluateGOF function.
        gofDist

        % (Don't manipulate) Goodness of fit statistics for comparisons between
        % bivariate joint distributions estimated from data and from virtual
        % population simulation.
        gofDist2D

        % (Don't manipulate) Goodness of fit statistics for comparisons between
        % correlations estimated from data and from virtual population
        % simulation.
        gofCor

        % (Don't manipulate) Composite goodness-of-fit result. Usually updated
        % by a call to the evaluateGOF.
        gof
    end
    properties (SetAccess = public)
        % Whether or not to use the effN for optimization and evaluation of the
        % GOF.  It is generally recommended to useEffN for evaluation of the
        % final fit but not during optimization.
        useEffN

        % Whether to check to apply Fisher's exact test instead
        % of the chi-square approximation for binned distribution test.  This
        % may be strictly correct but can slow calculations, often with small
        % impacts on GOF calculations.
        exactFlag

        % Used for optimization, if nonzero, will apply a penalty proportional
        % to spreadOut to increase how the prevalence weight is distributed.
        spreadOut

        % Used for optimization, if nonzero, will apply a large penalty if any
        % of the individual pvalues. fall below the target.
        minIndPVal

        % Time limit for optimizing the VPop in second
        optimizeTimeLimit

        % Type of optimization algorithm to employ.  Default is "pso". Allowed
        % options:
        %		"ga" - MATLAB's GA
        %		"pso" - MATLAB's PSO
        %		"papso" - MATLAB's PAPSO
        %		"gapso" - MATLAB's GA, polished by MATLAB's PSO
        %		"gapapso" - MATLAB's GA, polished by MATLAB's PAPSO
        %		"simplex" - MATLAB's simplex
        %		"surrogate" - a short run of MATLAB's surrogate,
        %                     polished by MATLAB's PSO
        optimizeType

        % Number of solutions to try in each optimization generation.  Most
        % directly impacts GA and PSO steps.  This is the population size to use
        % in the optimization runs. Default is 1000.
        optimizePopSize

        % Stopping condition for optimization
        objectiveLimit

        % Whether to close the pool at the end of optimization
        poolClose

        % Whether to restart the parallel pool at the start of optimization
        poolRestart

        % A non-negative integer seed to initialize the random number generator.
        % Set to -1 to avoid changing the state of the random number generator.
        intSeed

        % Numerical tolerance for optimization
        tol

        % Maximum number of outer-loop iterations to use in optimization.
        % If optimizeType is 'simplex', it's the number of iterations. If
        % optimizeType is 'ga', 'gapso', it is the number of generations. For
        % other optimization methods, this property is not used
        nIters

        % Minimum effective N. A large penalty is applied if effN drops below
        % this number during optimization, better ensuring solutions that weight
        % multiple VPs. The minEffN setting operates independently of useEffN,
        % and often works better than modifying spreadOut. Default: 0
        minEffN

        % Mean sum of residuals from linear calibration (subpopulation/dropouts weighted)
        MSE

        % The linearMatrix made from oldVPop linearCalibrationObject, actually
        % this will be the reduced matrix for the rescaled oldVPop with only
        % nVPmax columns. This is for the next iteration MSE minimization
        LinearProblemMatrices

        % The subweight directly recorded fom oldVPop linearCalibrationObject
        LinearProblemMatricesSubgroupSumWeights

        % The subweightDescription directly fom oldVPop linearCalibrationObject,
        % to keep track of biomarkers during cohort expansion
        LinearProblemMatricesobservationDescriptions

        % Lagrange multiplier for constrained optimization when using linear
        % calibration
        lambda
    end

    methods (Access = public)
        % Get a property of the VPop
        value = get(obj, propName)

        % Copy VP coefficients from a Worksheet to VPop
        obj = assignCoeffs(obj, myWorksheet)

        % Assign binning indices to VPs using information from worksheet and MAPEL
        % options
        obj = assignIndices(obj, myWorksheet, myMapelOptions)

        % Initialize bin probabilities. Only relevant for obj.pwStrategy = 'bin'
        obj = startProbs(obj, myRandomStart)

        % Update prevalance weight assignment for the virtual population
        obj = assignPWs(obj)

        % Initialize prevalence weights prior to the optimization
        obj = startPWs(obj, myWorksheet, myRandomStart)

        % Get all of the simulation datapoints for the VPs that will be needed
        % for calculating population statistics
        obj = getSimData(obj, myWorksheet)

        % Get simulation values that are fixed during optimization and add them
        % to the tables
        obj = addTableSimVals(obj)

        % Once we have read the simData and generated a prevalence weight
        % vector, we need to add the predicted equivalents to the tables used
        % for statistical comparison
        obj = addPredTableVals(obj)        
    end

    % Constructors
    methods
        function obj = VPop()
            % Default constructor for VPop object.
            obj.coeffsTable=[];
    		  obj.coeffsDist=[];
              obj.pwStrategy = 'direct';
              obj.indexTable = [];
              obj.binEdges = [];
              obj.binMidPoints = [];
              obj.binProbs = [];
              obj.pws = [];
              obj.mnSDTable = [];
              obj.binTable = [];
              obj.distTable = [];
              obj.distTable2D = [];
              obj.corTable = [];
              obj.subpopTable = [];
              obj.expData = [];
              obj.simData = [];
              obj.gofMn = [];
              obj.gofSD = [];
              obj.gofBin = [];
              obj.gofDist = [];
              obj.gofDist2D = [];
              obj.gofCor = [];
              obj.gof = [];
              obj.spreadOut = 0;
              obj.minIndPVal = 0;
              obj.spreadOut = 0;
              obj.minIndPVal = 0;
              obj.useEffN = false;
              obj.exactFlag = true;
              obj.optimizeTimeLimit = 10*60;
              obj.optimizeType = 'pso';
              obj.optimizePopSize = 1000;
              obj.objectiveLimit = -Inf;
              obj.poolClose = true;
              obj.poolRestart = true;
              obj.intSeed = -1;
              obj.tol = 1E-3;
              obj.nIters = 10000;
              obj.minEffN = 0;
              obj.MSE = [];
              obj.LinearProblemMatrices = [];
              obj.LinearProblemMatricesSubgroupSumWeights = [];
              obj.LinearProblemMatricesobservationDescriptions = [];
              obj.lambda = [];
        end
    end

    %% This block of methods are internal set methods. They must be defined in the
    % same file as the class definition. They cannot be called directly in user's
    % code but is called by MATLAB whenever the user assigns a new value to VPop's
    % properties with the "=" syntax. For example, when the user executes >>
    % vpop.coeffsTable = myCoeffsTable MATLAB will call the method
    % `set.coeffsTable(vpop, myCoeffsTable)` under the hood.
    methods        
        function obj = set.coeffsTable(obj,myCoeffsTable)
            obj.coeffsTable = myCoeffsTable;
        end
        function obj = set.coeffsDist(obj,myCoeffsDist)
            obj.coeffsDist = myCoeffsDist;
        end
        function obj = set.pwStrategy(obj,myPWStrategy)
            if sum(ismember({'direct','bin'},lower(myPWStrategy))) == 1
                obj.pwStrategy = lower(myPWStrategy);
            else
                error(['Property pwStrategy in ',mfilename,' must be "direct" or "bin"'])
            end
        end
        function obj = set.indexTable(obj,myIndexTable)
            obj.indexTable = myIndexTable;
        end

        function obj = set.binEdges(obj,myBins)
            obj.binEdges = myBins;
        end

        function obj = set.binMidPoints(obj,myBinMidPoints)
            obj.binMidPoints = myBinMidPoints;
        end

        function obj = set.binProbs(obj,myBinProbs)
            % We enforce constraints for binProbs:
            % a minimum value is 1E-14 rows must sum to 1
            % This helps avoid issues with the solver as well
            [nrows,ncols] = size(myBinProbs);
            if ((nrows > 0) && (ncols > 0))
                myBinProbs = myBinProbs ./ (repmat(sum(myBinProbs,2),1,ncols));
                adjustFlag = myBinProbs < 1E-14;
                myNadjust = sum(adjustFlag,2);
                myValDecrease = 1E-14 ./ myNadjust;
                myValDecrease(isinf(myValDecrease)) = 0 ;
                myBinProbs = (myBinProbs + adjustFlag * 1E-14) - ((1- adjustFlag) .* (repmat(myValDecrease,1,ncols)));
                myBinProbs = myBinProbs ./ (repmat(sum(myBinProbs,2),1,ncols));
            end
            obj.binProbs = myBinProbs;
        end

        function obj = set.pws(obj,myPWs)
            obj.pws = myPWs;
        end

        function obj = set.mnSDTable(obj,myMnSDTable)
            obj.mnSDTable = myMnSDTable;
        end

        function obj = set.binTable(obj,myBinTable)
            obj.binTable = myBinTable;
        end

        function obj = set.distTable(obj,myDistTable)
            obj.distTable = myDistTable;
        end
        function obj = set.distTable2D(obj,myDistTable)
            obj.distTable2D = myDistTable;
        end
        function obj = set.corTable(obj,myCorTable)
            obj.corTable = myCorTable;
        end

        function obj = set.subpopTable(obj,mySubpopTable)
            obj.subpopTable = mySubpopTable;
        end

        function obj = set.expData(obj,myExpData)
            obj.expData = myExpData;
        end

        function obj = set.simData(obj,mySimData)
            obj.simData = mySimData;
        end

        function obj = set.gofMn(obj,myGOF)
            obj.gofMn = myGOF;
        end

        function obj = set.gofSD(obj,myGOF)
            obj.gofSD = myGOF;
        end

        function obj = set.gofBin(obj,myGOF)
            obj.gofBin = myGOF;
        end

        function obj = set.gofDist(obj,myGOF)
            obj.gofDist = myGOF;
        end

        function obj = set.gofDist2D(obj,myGOF)
            obj.gofDist2D = myGOF;
        end

        function obj = set.gofCor(obj,myGOF)
            obj.gofCor = myGOF;
        end

        function obj = set.gof(obj,myGOF)
            obj.gof = myGOF;
        end

        function obj = set.useEffN(obj,myUseEffN)
            if islogical(myUseEffN)
                obj.useEffN = myUseEffN;
            else
                error(['Property useEffN in ',mfilename,' must be logical.'])
            end
        end

        function obj = set.exactFlag(obj,myFlag)
            obj.exactFlag = myFlag;
        end

        function obj = set.spreadOut(obj,mySpreadOut)
            if (mySpreadOut >= 0)
                obj.spreadOut = mySpreadOut;
            else
                error(['Property spreadOut in ',mfilename,' must be >= 0.'])
            end
        end

        function obj = set.minIndPVal(obj,myMinIndPVal)
            if (myMinIndPVal >= 0) & (myMinIndPVal <= 1)
                obj.minIndPVal = myMinIndPVal;
            else
                error(['Property minIndPVal in ',mfilename,' must be between 0 and 1.'])
            end
        end

        function obj = set.optimizeTimeLimit(obj,myOptimizeTimeLimit)
            obj.optimizeTimeLimit = myOptimizeTimeLimit;
        end

        function obj = set.optimizeType(obj,myOptimizeType)
            if sum(ismember({'pso','papso','ga','gapso','gapapso','simplex','surrogate'},lower(myOptimizeType))) == 1
                if sum(ismember({'papso','gapapso'},lower(myOptimizeType))) == 1
                    opts = optimoptions('particleswarm');
                    if ~isprop(opts,'UseAsync')
                        if isequal('papso',lower(myOptimizeType))
                            disp(['Parallel asynchronous mode not available for ',lower(myOptimizeType),' in ',mfilename,'.  Using pso.'])
                            myOptimizeType = 'pso';
                        else
                            disp(['Parallel asynchronous mode not available for ',lower(myOptimizeType),' in ',mfilename,'.  Using gapso.'])
                            myOptimizeType = 'gapso';
                        end
                    end
                end
    			  obj.optimizeType = lower(myOptimizeType);
            else
                error(['Property optimizeType in ',mfilename,' must be "ga," "pso," "gapso," "simplex," or "surrogate.  Parallel asynchronous options may also be available, "papso" and "gapapso."'])
            end
        end

        function obj = set.optimizePopSize(obj,myPopulationSize)
            if ((isnumeric(myPopulationSize) == true) && (myPopulationSize >= 0))
                obj.optimizePopSize = myPopulationSize;
            else
                error('Invalid optimizePopSize value')
            end
        end

        function obj = set.objectiveLimit(obj,myObjectiveLimit)
            if ((isnumeric(myObjectiveLimit) == true))
                obj.objectiveLimit = myObjectiveLimit;
            else
                error('Invalid objectiveLimit value')
            end
        end

        function obj = set.poolClose(obj,myInput)
            if islogical(myInput)
                obj.poolClose = myInput;
            else
                error(['Property poolClose in ',mfilename,' must be logical.'])
            end
        end

        function obj = set.poolRestart(obj,myInput)
            if islogical(myInput)
                obj.poolRestart = myInput;
            else
                error(['Property poolRestart in ',mfilename,' must be logical.'])
            end
        end

        function obj = set.intSeed(obj,myIntSeed)
            if ((mod(myIntSeed,1) == 0) && (myIntSeed>=-1))
                obj.intSeed = (myIntSeed);
            else
                error(['Invalid intSeed specified for ',mfilename,', a non-negative integer should be specified, or -1 to ignore.'])
            end
        end

        function obj = set.tol(obj,myTol)
            if (myTol > 0)
                obj.tol = myTol;
            else
                error(['Property tol in ',mfilename,' must be > 0.'])
            end
        end

        function obj = set.nIters(obj,myNIters)
            if (myNIters > 0) && (mod(myNIters,1) == 0)
                obj.nIters = myNIters;
            else
                error(['Property nIters in ',mfilename,' must be a positive integer.'])
            end
        end

        function obj = set.minEffN(obj,myMinEffN)
            if ((isnumeric(myMinEffN) == true) && (myMinEffN >= 0))
                obj.minEffN = myMinEffN;
            else
                error('Invalid minEffN value')
            end
        end
    end

    %% Short methods that can be kept in the class def file
    methods 
        function effN = computeEffN(obj)
            % Compute the VPop's effective sample size (a.k.a. EffN).
            arguments
                obj (1,1) VPop; 
            end
            effN = 1.0/sum(obj.pws.^2);
        end
    end
end
```
