## Supplementary File 3 for "A surrogate workflow for virtual population development: designing an amenable proxy for a hypothesis-test inspired nonlinear goodness-of-fit": correlationOptions.m

```
classdef correlationOptions
% Options object for checking for correlations in a prevalence-weighted
% cohort.
%
% PROPERTIES
% interventionID:          (Required) Worksheet intervention to run
%                          analysis on.
% pws:                     (Optional) a 1xnVP vector of prevalence weights
%                          for VPs in the worksheet.  If not provided,
%                          1/nVP (unweighted) will be assumed for each
% analyzeElementResultIDs: (Optional) a 1xNvar cell array of strings of
%                          variables to include in the analysis.  If not
%                          provided, all variables that exhibit variability
%                          in the results will be considered
% analyzeTimes:            (Optional) a 1xNvar vector of
%                          time values to include in the analysis.  If not
%                          provided, the last simulated time will be used.
% minAbsValFilter:         (Optional) If a value >0 is provided,
%                          variables that do not achieve a value greater
%                          than this in at least 1 VP the simulations will
%                          be filteredfrom the analysis.  Intended to
%                          help with noise.
% addAxis:                (Optional) Whether to include mechanistic axes
%                         coefficients in the analysis.  If true, they will
%                         be added.  Default is false.
    
   properties
      interventionID
      pws
      analyzeElementResultIDs
      analyzeTimes
      minAbsValFilter
      addAxis
   end
   
   methods

      function obj = set.interventionID(obj,myInterventionID)
          if ischar(myInterventionID)
              obj.interventionID = myInterventionID;
          else
              error(['Invalid interventionID specified for ',mfilename,', an intervention ID string should be specified.'])
          end
      end     
      
      function obj = set.pws(obj,myPWs)
          if isnumeric(myPWs)
              obj.pws = myPWs;
          else
              error(['Invalid PWs specified for ',mfilename,', a numeric vector should be specified, ideally a nVPx1 vector that sums to 1.'])
          end
      end      
      
      function obj = set.analyzeElementResultIDs(obj,myAnalyzeElementResultIDs)
          if iscell(myAnalyzeElementResultIDs)
              obj.analyzeElementResultIDs = (myAnalyzeElementResultIDs);
          else
              error(['Invalid analyzeElementResultIDs specified for ',mfilename,', a 1xnVar cell array of variable ID strings should be specified.'])
          end
      end     
      
      function obj = set.addAxis(obj,myAddAxis)
          if islogical(myAddAxis)
               obj.addAxis = myAddAxis;
          else
               error(['Property addAxis in ',milename,' must be true or false.'])
          end
      end      
      
      function obj = set.analyzeTimes(obj,myAnalyzeTimes)
          failFlag = false;
          if isnumeric(myAnalyzeTimes)
            obj.analyzeTimes = (myAnalyzeTimes);
          else
            failFlag = true;
          end
          if failFlag
              error(['Invalid myAnalyzeTimes specified for ',mfilename,'. A 1xnVar cell array of numbers should be specified.'])
          end
      end      
      
      function obj = set.minAbsValFilter(obj,myMinAbsValFilter)
          failFlag = false;
          if isnumeric(myMinAbsValFilter)
              if myMinAbsValFilter >= 0
                obj.minAbsValFilter = myMinAbsValFilter;
              else
                  failFlag = true;
              end
          else
              failFlag = true;
          end
          if failFlag
              error(['Invalid minAbsValFilter specified for ',mfilename,', a nonnegative number should be specified.'])
          end
      end            
      
      function value = get(obj,propName)
          switch propName                 
              case 'interventionID'
                  value = obj.interventionID;   
              case 'pws'
                  value = obj.pws;  
              case 'analyzeElementResultIDs'
                  value = obj.analyzeElementResultIDs;                     
              case 'analyzeTimes'
                  value = obj.analyzeTimes;  
              case 'analyzeTimes'
                  value = obj.analyzeTimes;   
              case 'minAbsValFilter'
                  value = obj.minAbsValFilter;
              case 'addAxis'
                  value = obj.addAxis;                  
          end
      end     
      
      function passCheck = verify(obj,myWorksheet)
          passCheck = true;
          allInterventionIDs = getInterventionIDs(myWorksheet);
          allVPIDs = getVPIDs(myWorksheet);
          [~, nVPResults]= size(myWorksheet.results);
          if sum(ismember(allInterventionIDs,  obj.interventionID)) < 1
              warning(['Specified interventionID should be selected from interventionIDs in the worksheet in ',mfilename,'.'])
              passCheck = false;
          end
          if nVPResults ~= length(allVPIDs)
              warning(['Results for all VPs not available from worksheet in ',mfilename,'.'])
              passCheck = false;
          end          
          if passCheck
              myInterventionIndex = find(ismember(allInterventionIDs,  obj.interventionID));
              flagResultCheck = strcmp(cellfun(@class,myWorksheet.results(myInterventionIndex,:), 'UniformOutput', false),'struct');
              if sum(flagResultCheck) < length(allVPIDs)
                  warning(['Results for all VPs for intervention ',obj.interventionID,' not available from worksheet in ',mfilename,'.'])
                  passCheck = false;
              end
          end
          if passCheck
              
              % Otherwise, we need to make sure any analyzeResultsIDs
              % specified are available in the results
              if length(obj.analyzeElementResultIDs) > 0
                  testResult = myWorksheet.results{myInterventionIndex,1};
                  testNames = testResult.Names;
                  myTimeIndex=find(ismember(testResult.Names,'time'));
                  testNames(myTimeIndex)=[];
                  if sum(ismember(obj.analyzeElementResultIDs,testNames)) < length(obj.analyzeElementResultIDs)
                      warning(['Specified analyzeElementResultIDs not all available from worksheet in ',mfilename,'.'])
                      passCheck = false;
                  end
              end
              
              timesToCheck = unique(obj.analyzeTimes);
              % We'll use the first result as a proxy, they should all be
              % identical
              % Note we don't include a check for the uniqueness and
              % numerical accuracy of the simulation times here but
              % these should be fixed in the worksheet simProps
              testResult=myWorksheet.results{myInterventionIndex,1};
              myTimeIndex=find(ismember(testResult.Names,'time'));
              if sum(ismember(timesToCheck,testResult.Data(:,myTimeIndex))) < length(timesToCheck);
                  warning(['Results for all requested times for intervention ',obj.interventionID,' not available in worksheet results in ',mfilename,'.'])
                  passCheck = false;                  
              end
          end
          [~,myNpws] = size(obj.pws);
          if myNpws ~= length(allVPIDs)
              warning(['Prevalence weights should be a nVPx1 numeric vector in ',mfilename,'.'])
              passCheck = false;
          else
              % We'll also enforce that the PWs sum to 1 here
              myPWs = obj.pws - max(obj.pws);
              myPWs = exp(myPWs);
              myPWs = myPWs / sum(myPWs);
              obj.pws = myPWs;
          end
          if ~isequal(size(obj.analyzeElementResultIDs),size(obj.analyzeTimes))
              warning(['A 1xnVar cell array should be specified for the analyzeElementResultIDs, and a 1xnVar matrix should be specified for the analyzeTimes in ',mfilename,'.'])
              passCheck = false;
          end 
      end
          
      function obj = correlationOptions() 
          obj.interventionID='';
          obj.pws=ones(0,1);
          obj.analyzeElementResultIDs=cell(1,0);
          obj.analyzeTimes=zeros(1,0);   
          obj.minAbsValFilter = 0;
          obj.addAxis = false;
      end
   end
end
```
