## Supplementary File 3 for "A surrogate workflow for virtual population development: designing an amenable proxy for a hypothesis-test inspired nonlinear goodness-of-fit": expandVPopEffNOptions.m

```
classdef expandVPopEffNOptions
% Options and data required to to run expandVPopEffN
%
%  suffix:             (Required) a text descriptor string that will be included in what 
%                      will be written to file.  This is also used
%                      in setting VP identities.
%                       Default = '' and should be specified  
%  wsIterCounter:      (Optional) tracks the iterations through the algorithm.  WARNING: Keep
%                      incrementing to avoid issues with repeated VPIDs, and do not repeat
%                      a previous value.
%                       Default = 0  
%  targetEffN:         (Optional) algorithm stops when it is achieves a valid VPop
%                      with this effN
%                       Default = 100  
%  maxNewPerIter:      (Optional) Maximum new VPs we can add per iteration.  Set to 
%                      -1 to use the VPop effN
%                       Default = 50  
%  expandCohortSize:   (Optional) size of the cohort to generate for testing when
%                      making new VPs
%                       Default = 2500                       
%  effNDelta:          (Optional) How much to increment the minEffN each iterationv
%                       Default = 2
%  useMapelIntSeed:    (Optional) Whether to use the integer seed in the mapel options to
%                       start the random number generator (true) or to use seeds fixed
%                       by the algorithm for repeatability (false).
%                       Default = false
%  minPVal:            (Optional) minimum p-value for a good VPop
%                       Default = 0.9
%  nCluster:           (Optional) Number of VPs to keep following a cluster
%                       step.  If set to nan, expandVPopEffN will make
%                       an intelligent guess.
%                       Default = nan;
%  nTries:             (Optional) Number of times to try a solution before triggering expansion.
%                       Default = 5
%  nVPMax:             (Optional) Maximum number of VPs in the cohort.
%                       Default = Inf    
%  nRetries:           (Optional) Number of times to try a solution that is not quite at acceptance.
%                       Default = 30
%  restartPVal:        (Optional) p-value that will trigger a restart of the algorithm.
%                       Default = 1E-4
%  nSamplePastCompletion (Optional) if > 0, will continue to sample this many
%                        VPs and reweight to develop alternate VPops once
%                        targetEffN is achieved.
%                       Default = 0
%  expandRandomStart:  (Optional) If 0, the prevalence 
%                      weights are not perturbed from initial guesses on the first iteration or
%                      following a successful solution for the population and expansion.
%                      Otherwise the transformed initial bin
%                      probabilities are perturbed 
%                      with random (normally distributed) noise that has
%                      normalized standard deviation of the specified value. 
%					   Set to Inf for a fully random initial start. 
%                       Default = 0; 
%  varyMethod :        Method to use to guide resampling.  See varyAxesOptions,
%                       currently allowed values are 'gaussian' and 'localPCA'
%  resampleStd:        Degree of normalized variability to add to children of
%                       highest weighted VPs in each iteration when resampling. 
%                       Default = 0.05;
%                       If the varyMethod is gaussian, this is the std of
%                       the resampling in the normalized units of the axis,
%                       and values < 1 are appropriate.
%                       If the varyMethod is localPCA, this is the standard
%                       deviation relative to the PCA result, and values
%                       greater than 1 should help to increase the
%                       variability in the VPs.
%  maxNewPerOld:       Maximum number of VPs to keep per old parent VP for each iteration.
%                       This number can be exceeded if allowed by maxNewPerIter and the number of
%                       parent VPs, but in the case when there aren't enough maxNewPerIter at least
%                       this many will be kept.
%                       Default = 2;
%  expandEdgeVPs:      Whether to compare the simulation ranges to the
%                       ranges in the data and try to use any "edge" VPs
%                       as seeds every iteration if the range is not   
%                       covered well.  Default = false;
%  plausibleResponseTypeIDs A cell array of responseTypeIDs that will be used
%                       as a post-simulation check to make sure VPs are
%                       plasuble.  If left blank, all worksheet
%                       responseTypes will be used.  Note VPs will be
%                       screened for plausibility relative to the starting
%                       worksheet.
%  screenFunctionName  A string with a name for a VP pre-simulation
%                       screening function.
%                       The function will be called every iteration when
%                       VPs are expanded.  The function should be in a file
%                       in the MATLAB path with the same filename, do 
%                       not include '.m'.  The function should accept
%                       a worksheet and a list of VPIDs to screen as input,
%                       and should return a screened worksheet as output.
%  selectByParent:	   Whether to select children VPs each iteration based on
%                       their parent, in which case maxNewPerOld is important to
%                       select VPs, or to pool all the children together and
%                       pick them based on their scores.
%  verbose:            (Optional) report progress to screen
%                       Default = true;
%  linearExpandFlag:   (Optional) Whether or not expand through linear calibration workflow first. Default = true
%  minPVallinear:      (Optional) Minimum p-value to terminate linear expansion phase 
%                       and enter pso optimization phase. Default = 0.5
%  maxIterlinearExpand: (Optional) Maximum iteration number to terminate linear expansion phase 
%                       and enter pso optimization phase. Default = 200
%  minEffNlinearflag:   (Optional) Whether or not start with effN determined from a linear calibration first. Default = true
%  linearExpandCohortSize: (Optional) expandCohortSize during linearExpansion phase. Default = 1500

   properties
		suffix
		wsIterCounter
		targetEffN
		maxNewPerIter
		expandCohortSize
		effNDelta
		useMapelIntSeed
		minPVal
        nCluster
		nTries
        nVPMax
		nRetries
		restartPVal
		expandRandomStart
		varyMethod
		resampleStd
        nSamplePastCompletion
		maxNewPerOld
		expandEdgeVPs
        plausibleResponseTypeIDs
        screenFunctionName
		selectByParent
		verbose
        linearExpandFlag 
        minPVallinear
        maxIterlinearExpand
        minEffNlinearflag
        linearExpandCohortSize
   end

   methods     

      function obj = set.suffix(obj,mySuffix)
          if ischar(mySuffix)
              if contains(mySuffix,'_')
                  disp(['Underscore "_" characters not supported in suffix in ',mfilename,'.  Replacing with hyphens "-".'])
                  mySuffix = strrep(mySuffix,'_','-');
              end
              obj.suffix = (mySuffix);
          else
              error(['Invalid suffix specified in ',mfilename,', a string should be specified.'])
          end
      end         
   
      function obj = set.wsIterCounter(obj,myWsIterCounter)
          failFlag = false;
          if isnumeric(myWsIterCounter) 
              if isequal(size(myWsIterCounter),[1 1])
                  if round(myWsIterCounter) >= 0
                      obj.wsIterCounter = round(myWsIterCounter);
                  else
                      failFlag = true;
                  end
              else
                  failFlag = true;
              end
          else
              failFlag = true;
          end
          if failFlag
              error(['Invalid wsIterCounter specified for ',mfilename,'. A nonnegative number should be specified.'])
          end
      end   
   
      function obj = set.targetEffN(obj,myTargetEffN)
          failFlag = false;
          if isnumeric(myTargetEffN) 
              if isequal(size(myTargetEffN),[1 1])
			  	  if mod(myTargetEffN,1) ~= 0
					  warning(['A whole number is required for targetEffN in ',mfilename,'. The provided value will be rounded.']) 
				  end
                  if round(myTargetEffN) >= 0
                      obj.targetEffN = round(myTargetEffN);
                  else
                      failFlag = true;
                  end
              else
                  failFlag = true;
              end
          else
              failFlag = true;
          end
          if failFlag
              error(['Invalid targetEffN specified for ',mfilename,'. A whole number should be specified.'])
          end
      end      
   
      function obj = set.maxNewPerIter(obj,myMaxNewPerIter)
          failFlag = false;
          if isnumeric(myMaxNewPerIter) 
              if isequal(size(myMaxNewPerIter),[1 1])
                  if round(myMaxNewPerIter) > 0
                      obj.maxNewPerIter = round(myMaxNewPerIter);
                  else
                      failFlag = true;
                  end
              else
                  failFlag = true;
              end
          else
              failFlag = true;
          end
          if failFlag
              error(['Invalid maxNewPerIter specified for ',mfilename,'. A positive number should be specified.'])
          end
      end     
 
      function obj = set.expandCohortSize(obj,myExpandCohortSize)
          failFlag = false;
          if isnumeric(myExpandCohortSize) 
              if isequal(size(myExpandCohortSize),[1 1])
                  if round(myExpandCohortSize) >= 0
                      obj.expandCohortSize = round(myExpandCohortSize);
                  else
                      failFlag = true;
                  end
              else
                  failFlag = true;
              end
          else
              failFlag = true;
          end
          if failFlag
              error(['Invalid expandCohortSize specified for ',mfilename,'. A nonnegative number should be specified.'])
          end
      end    

      function obj = set.effNDelta(obj,myEffNDelta)
          failFlag = false;
          if isnumeric(myEffNDelta) 
              if isequal(size(myEffNDelta),[1 1])
				  if mod(myEffNDelta,1)~=0
					  warning(['A whole number is required for effNDelta in ',mfilename,'. The provided value will be rounded.']) 
				  end
                  if round(myEffNDelta) >= 0.5
                      obj.effNDelta = round(myEffNDelta);
                  elseif myEffNDelta >= 0
                      obj.effNDelta = 0;
					  warning(['A zero effNDelta is specified in ',mfilename,'. Note the calling function may never terminate if an increased effN is required.']) 
                  else
                      failFlag = true;					  
                  end
              else
                  failFlag = true;
              end
          else
              failFlag = true;
          end
          if failFlag
              error(['Invalid effNDelta specified for ',mfilename,'. A positive number should be specified.'])
          end
      end  	  
		
	  function obj = set.useMapelIntSeed(obj,myUseMapelIntSeed)
          if islogical(myUseMapelIntSeed)
               obj.useMapelIntSeed = myUseMapelIntSeed;
          else
               error(['Property useMapelIntSeed in ',milename,' must be true or false.'])
          end
      end   			
		
      function obj = set.minPVal(obj,myMinPVal)
          failFlag = false;
          if isnumeric(myMinPVal) 
              if isequal(size(myMinPVal),[1 1])
                  if ((myMinPVal >= 0) && (myMinPVal <= 1))
                      obj.minPVal = myMinPVal;
                  else
                      failFlag = true;
                  end
              else
                  failFlag = true;
              end
          else
              failFlag = true;
          end
          if failFlag
              error(['Invalid minPVal specified for ',mfilename,'. A number between 0 and 1 should be specified.'])
          end
      end  
      
      function obj = set.nCluster(obj,myVal)
          failFlag = false;
          if isnumeric(myVal) 
              if isequal(size(myVal),[1 1])
                  if round(myVal) > 0
                      obj.nCluster = round(myVal);
                  elseif isnan(myVal) 
                      obj.nCluster = myVal;
                  else
                      failFlag = true;
                  end
              else
                  failFlag = true;
              end
          else
              failFlag = true;
          end
          if failFlag
              error(['Invalid nCluster specified for ',mfilename,'. A positive number, or NaN, should be specified.'])
          end
      end        
	
      function obj = set.nTries(obj,myNTries)
          failFlag = false;
          if isnumeric(myNTries) 
              if isequal(size(myNTries),[1 1])
                  if round(myNTries) > 0
                      obj.nTries = round(myNTries);
                  else
                      failFlag = true;
                  end
              else
                  failFlag = true;
              end
          else
              failFlag = true;
          end
          if failFlag
              error(['Invalid nTries specified for ',mfilename,'. A positive number should be specified.'])
          end
      end  
      
      function obj = set.nVPMax(obj,myNVPMax)
          failFlag = false;
          if isnumeric(myNVPMax) 
              if isequal(size(myNVPMax),[1 1])
                  if round(myNVPMax) > 0
                      obj.nVPMax = round(myNVPMax);
                  else
                      failFlag = true;
                  end
              else
                  failFlag = true;
              end
          else
              failFlag = true;
          end
          if failFlag
              error(['Invalid nVPMax specified for ',mfilename,'. A positive number should be specified.'])
          end
      end       

      function obj = set.nRetries(obj,myNRetries)
          failFlag = false;
          if isnumeric(myNRetries) 
              if isequal(size(myNRetries),[1 1])
                  if round(myNRetries) > 0
                      obj.nRetries = round(myNRetries);
                  else
                      failFlag = true;
                  end
              else
                  failFlag = true;
              end
          else
              failFlag = true;
          end
          if failFlag
              error(['Invalid nRetries specified for ',mfilename,'. A positive number should be specified.'])
          end
      end  		  
	  
      function obj = set.restartPVal(obj,myRestartPVal)
          failFlag = false;
          if isnumeric(myRestartPVal) 
              if isequal(size(myRestartPVal),[1 1])
                  if ((myRestartPVal >= 0) && (myRestartPVal <= 1))
                      obj.restartPVal = myRestartPVal;
                  else
                      failFlag = true;
                  end
              else
                  failFlag = true;
              end
          else
              failFlag = true;
          end
          if failFlag
              error(['Invalid restartPVal specified for ',mfilename,'. A number between 0 and 1 should be specified.'])
          end
      end  	  
	  
      function obj = set.expandRandomStart(obj,myExpandRandomStart)
          failFlag = false;
          if isnumeric(myExpandRandomStart) 
              if isequal(size(myExpandRandomStart),[1 1])
                  if (myExpandRandomStart >= 0)
                      obj.expandRandomStart = myExpandRandomStart;
                  else
                      failFlag = true;
                  end
              else
                  failFlag = true;
              end
          else
              failFlag = true;
          end
          if failFlag
              error(['Invalid expandRandomStart specified for ',mfilename,'. A number >= 0 should be specified.'])
          end
      end  	  

      function obj = set.varyMethod(obj,myVaryMethod)
          allowedSettings = {'gaussian', 'localpca'};
          if sum(ismember(allowedSettings, lower(myVaryMethod)))>0
              obj.varyMethod = lower(myVaryMethod);
          else
              error('Invalid varyMethod specified for ',mfilename,', should specify one of: ',strjoin(allowedSettings,', '),'.')
          end
      end 
	  
      function obj = set.resampleStd(obj,myValue)
          failFlag = false;
          if isnumeric(myValue) 
              if isequal(size(myValue),[1 1])
                  if (myValue >= 0)
                      obj.resampleStd = myValue;
                  else
                      failFlag = true;
                  end
              else
                  failFlag = true;
              end
          else
              failFlag = true;
          end
          if failFlag
              error(['Invalid resampleStd specified for ',mfilename,'. A number >= 0 should be specified.'])
          end
      end     	
      
      function obj = set.nSamplePastCompletion(obj,nSamplePastCompletion)
          failFlag = false;
          if isnumeric(nSamplePastCompletion) 
              if isequal(size(nSamplePastCompletion),[1 1])
                  if round(nSamplePastCompletion) >= 0
                      obj.nSamplePastCompletion = round(nSamplePastCompletion);
                  else
                      failFlag = true;
                  end
              else
                  failFlag = true;
              end
          else
              failFlag = true;
          end
          if failFlag
              error(['Invalid nSamplePastCompletion specified for ',mfilename,'. A nonnegative number should be specified.'])
          end
      end         

      function obj = set.maxNewPerOld(obj,myMaxNewPerOld)
          failFlag = false;
          if isnumeric(myMaxNewPerOld) 
              if isequal(size(myMaxNewPerOld),[1 1])
                  if (myMaxNewPerOld >= 1)
                      obj.maxNewPerOld = round(myMaxNewPerOld);
                  else
                      failFlag = true;
                  end
              else
                  failFlag = true;
              end
          else
              failFlag = true;
          end
          if failFlag
              error(['Invalid maxNewPerOld specified for ',mfilename,'. A number >= 1 should be specified.'])
          end
      end  	  

	  function obj = set.expandEdgeVPs(obj,myInput)
          if islogical(myInput)
               obj.expandEdgeVPs = myInput;
          else
               error(['Property expandEdgeVPs in ',milename,' must be true or false.'])
          end
      end

	  function obj = set.plausibleResponseTypeIDs(obj,myValue)
          if iscell(myValue)
               obj.plausibleResponseTypeIDs = myValue;
          else
               error(['Property plausibleResponseTypeIDs in ',milename,' must be a cell.'])
          end
      end        
      
	  function obj = set.screenFunctionName(obj,myScreenFunctionName)
          if ischar(myScreenFunctionName)
               obj.screenFunctionName = myScreenFunctionName;
          else
               error(['Property screenFunctionName in ',milename,' must be a character array.'])
          end
      end       
	  
	  function obj = set.selectByParent(obj,mySelectByParent)
          if islogical(mySelectByParent)
               obj.selectByParent = mySelectByParent;
          else
               error(['Property selectByParent in ',milename,' must be true or false.'])
          end
      end   	  
	  
	  function obj = set.verbose(obj,myVerbose)
          if islogical(myVerbose)
               obj.verbose = myVerbose;
          else
               error(['Property verbose in ',milename,' must be true or false.'])
          end
      end         
 
      
      function value = get(obj,propName)
          switch propName
              case 'suffix'
                  value = obj.suffix;
              case 'wsIterCounter'
                  value = obj.wsIterCounter;
              case 'targetEffN'
                  value = obj.targetEffN;  
              case 'maxNewPerIter'
                  value = obj.maxNewPerIter;                   
              case 'expandCohortSize'
                  value = obj.expandCohortSize;				  
              case 'effNDelta'
                  value = obj.effNDelta;
			  case 'useMapelIntSeed'
				  value = obj.useMapelIntSeed;
              case 'minPVal'
                  value = obj.minPVal;
              case 'nCluster'
                  value = obj.nCluster;                  
              case 'nTries'
                  value = obj.nTries;
              case 'nVPMax'
                  value = obj.nVPMax;                  
              case 'nRetries'
                  value = obj.nRetries;
              case 'restartPVal'
                  value = obj.restartPVal;	
              case 'expandRandomStart'
                  value = obj.expandRandomStart;
              case 'varyMethod'
                  value = obj.varyMethod;				  
              case 'resampleStd'
                  value = obj.resampleStd;
              case 'nSamplePastCompletion'
                  value = obj.nSamplePastCompletion;                  
              case 'maxNewPerOld'
                  value = obj.maxNewPerOld;	
              case 'expandEdgeVPs'
                  value = obj.expandEdgeVPs;
              case 'plausibleResponseTypeIDs'
                  value = obj.plausibleResponseTypeIDs;                  
              case 'screenFunctionName'
                  value = obj.screenFunctionName;
              case 'selectByParent'
                  value = obj.selectByParent;
              case 'verbose'
                  value = obj.verbose;
              otherwise
                  error(['Error: ',propName ,' is not a valid ',mfilename,' property.'])
          end
      end

      % TODO: ADD METHODS TO VERIFY AGAINST WORKSHEET
            
      function obj = expandVPopEffNOptions()
          % This is the constructor method for instances of 
          % expandVPopEffNOptions objects.
          obj.suffix = '';
          obj.wsIterCounter = 0;
          obj.targetEffN = 100;
          obj.maxNewPerIter = 50;
		  obj.expandCohortSize = 2500;
          obj.effNDelta = 2;
		  obj.useMapelIntSeed = false;
          obj.minPVal = 0.9;
          obj.nCluster = nan;           
          obj.nTries = 5;
          obj.nVPMax = Inf;          
          obj.nRetries = 30;
		  obj.restartPVal = 1E-4;
		  obj.expandRandomStart = 0;
		  obj.varyMethod = 'gaussian';
		  obj.resampleStd = 0.05;
          obj.nSamplePastCompletion = 0;
		  obj.maxNewPerOld = 2;
		  obj.expandEdgeVPs = false;
          obj.plausibleResponseTypeIDs = {};
          obj.screenFunctionName = '';
		  obj.selectByParent = true;
		  obj.verbose = true;	
          obj.linearExpandFlag = true;
          obj.minPVallinear = 0.5;
          obj.maxIterlinearExpand = 200;
          obj.minEffNlinearflag = true;
          obj.linearExpandCohortSize = 1500;
      end
   end
end
```
