## Supplementary File 3 for "A surrogate workflow for virtual population development: designing an amenable proxy for a hypothesis-test inspired nonlinear goodness-of-fit": mapelOptions.m

```
classdef mapelOptions
% Options and data required to to run MAPEL
%
% expData:           (Required) Contains experimental data.  Must be
%                     populated before calling MAPEL.
% mnSDTable:         (Optional) Contains mean/SD data to match.  Must be
%                     populated with experimental data before calling MAPEL,
%                     or leave unassigned if no mean/SD targets.
% binTable:          (Optional) Containing data on a binned pdf function to
%                     match.  Must be 
%                     populated with experimental data before calling MAPEL,
%                     or leave unassigned if no bin targets.
% distTable:         (Optional) Contains data on a cdf function to
%                     match.  Must be
%                     populated with experimental data before calling MAPEL,
%                     or leave unassigned if no distribution targets.
% distTable2D:       (Optional) Contains data on a 2D pdf function to
%                     match.  Must be
%                     populated with experimental data before calling MAPEL,
%                     or leave unassigned if no 2D distribution targets.
% corTable:	         (Optional) A table to enable calibrating pairwise correlations.
%                     Leave unassigned if there are not correlation targets to match.
% subpopTable:       Contains criteria to create subpopulations
%                     from simulated VPs.												   
% pwStrategy:	     Strategy for the optimization.  Allowed values:
%                     'direct' (default value) If this is set then PWs are optimized directly
%                     'bin'     specified in a strategy similar to teh original MAPEL algorithm 
% nBins:             (Optional) Number of bins per continuous axis.  The
%                    default is 2, which may be too constrictive in many
%                    situations.
% initialProbs:      (Optional) A NAxis X Nbins matrix of initial 
%                    probabilities. If set to be [], probabilities are
%                    initialized to be uniform.  The default is [].
%                    ONLY USED IN BIN PWSTRATEGY
% initialPWs:        (Optional) 
%                     - A 1 x nVP vector of initial pws. It is also 
%                       possible to provide multiple initial guesses as an
%                       m x nVP vector. OR
%                     - (default) If set to be [], probabilities are
%                       initialized to be uniform.  
%                       The default is []. OR
%                     - if set to -1, we will try to find a near optimum starting
%                       point based on a linearized problem formulation.
%                       ONLY USED IN DIRECT PWSTRATEGY	  
% randomStart:       (Optional) 
%					 - If 0 (default), the initial 
%                      parameters are not perturbed in anyway.  
%                    - If greater than 0, the 
%                      transformed initial probabilities are perturbed  
%                      with random (normally distributed) noise that has
%                      normalized standard deviation of the specified value.  
%                      Suggest a value less than 1.
%                    - If initialProbs/initialPWs are not specified, a value of zero 
%					   results in uniform initial probabilities
%                    - If initialProbs/initialPWs are not specified, a value > 0 
%                      results in uniformly sampled initial probabilities,
%                      renormalized 
% nIters:            (Optional) default is 10,000.
%                    simplex: number of iterations
%                    ga,gapso: maximum number of generations
%                    ignored for other optimizeType options.
% equalBinBreaks     (Optional) Boolean (true/false). Whether to adjust bin 
%                    edges so they cover an equal numeric interval (true),
%                    or a ~ equal number of VPs are
%                    included in each bin (false).  Default is false.
%                    ONLY USED IN BIN PWSTRATEGY												
% tol:               (Optional) Stopping tolerance for the solvers.
%                    Default is 1E-3.
% spreadOut:         (Optional) Extra argument for "findFit" function that 
%                    tries to force more VPs to be used in the solution.
%                    This is also addressed with minEffN.  Default is 0.
% minIndPVal:        Used for optimization, if nonzero, will  
%                    apply a large penalty if any of the individual pvalues.
%                    fall below the target.
% useEffN:           (Optional) Boolean (true/false). Whether to use the  
%                    total number of PWs above or effective N measure for 
%                    optimization and calculating GOF statistics.  It is 
%                    strongly recommended to set this to false during
%                    optimization.  It is set to false by
%                    default.
%  exactFlag:        Whether to check to apply Fisher's exact test instead
%                    of the chi-square approximation for binned distribution
%                    test.  This may be strictly correct but can slow
%                    calculations, often with small impacts on GOF
%                    calculations.
% optimizeTimeLimit: (Optional) Time limit to stop the solver, does not 
%                    apply to the simplex method.  Default is 10*60 s,
%                    which will not be sufficient in many cases.
% optimizeType:      Type of optimization algorithm to employ.  Default is
%                     "pso".
%						"ga" - MATLAB's GA
%						"pso" - MATLAB's PSO
%						"papso" - MATLAB's PAPSO
%						"gapso" - MATLAB's GA, polished by MATLAB's PSO
%						"gapapso" - MATLAB's GA, polished by MATLAB's PAPSO
%						"simplex" - MATLAB's simplex
%						"surrogate" - a short run of MATLAB's surrogate,
%                                     polished by MATLAB's PSO
% optimizePopSize:   (Optional) most directly impacts GA and PSO steps.  This is
%                    the population size to use in the optimization runs.
%                    Default is 1000.
% objectiveLimit:    stopping condition for optimization
% poolClose:         Whether to close the pool at the end of the optimization  
% poolRestart:       Whether to restart the pool at the beginning of the optimization 
% intSeed:           (Optional) Random seed, set to -1 to avoid re-seeding
%                    the random number generator.  Default is -1.
% minEffN:           (Optional) Distinct from useEffN, this is an additional
%                    constraint placed on optimization.  Solutions that
%                    do not meet this EffN criteria are heavily penalized.
%                    The default is 0.

   properties
      expData
      mnSDTable
      binTable
      distTable      
	  distTable2D		
	  corTable
	  subpopTable	  
      pwStrategy
      nBins
      initialProbs
      initialPWs
      randomStart
      nIters
      equalBinBreaks
      tol
      spreadOut
	  minIndPVal	  
      useEffN
      exactFlag
      optimizeTimeLimit
      optimizeType
      optimizePopSize 
	  objectiveLimit
	  poolClose
	  poolRestart
      intSeed
      minEffN
   end

   methods     

      function obj = set.expData(obj,myExpData)
          obj.expData = myExpData;
      end       
       
      function obj = set.mnSDTable(obj,myExpMnSdData)
          obj.mnSDTable = myExpMnSdData;
      end

      function obj = set.binTable(obj,myExpBinData)
          obj.binTable = myExpBinData;
      end

      function obj = set.distTable(obj,myExpDistData)
          obj.distTable = myExpDistData;
      end      

      function obj = set.distTable2D(obj,myExpDistData2D)
          obj.distTable2D = myExpDistData2D;
      end 
	  
      function obj = set.corTable(obj,myCorTable)
          obj.corTable = myCorTable;
      end 	  
	  
      function obj = set.subpopTable(obj,mySubPopTable)
          obj.subpopTable = mySubPopTable;
      end	  

      function obj = set.pwStrategy(obj,myPWStrategy)
          if sum(ismember({'direct','bin'},lower(myPWStrategy))) == 1
              obj.pwStrategy = lower(myPWStrategy);
          else
              error(['Property pwStrategy in ',mfilename,' must be "direct" or "bin"'])
          end
      end 
	  
      function obj = set.nBins(obj,myNBins)
          if mod(myNBins,1) == 0
              obj.nBins = myNBins;
          else
              error(['Invalid nBins in ',mfilename,'.'])
          end
      end      

      function obj = set.initialProbs(obj,myInitialProbs)
          obj.initialProbs = myInitialProbs;
      end      
	  
      function obj = set.initialPWs(obj,myInitialPWs)
          obj.initialPWs = myInitialPWs;
      end    	
      
      function obj = set.randomStart(obj,myRandomStart)
          if (myRandomStart >= 0)
               obj.randomStart = myRandomStart;
          else
               error(['Property randomStart in ',mfilename,' must be > 0.'])
          end
      end  
      
      function obj = set.nIters(obj,myNIters)
          if (myNIters > 0) && (mod(myNIters,1) == 0)
               obj.nIters = myNIters;
          else
               error(['Property nIters in ',mfilename,' must be a positive integer.'])
          end
      end            

      function obj = set.equalBinBreaks(obj,myEqualBinBreaks)
          if islogical(myEqualBinBreaks)
               obj.equalBinBreaks = myEqualBinBreaks;
          else
               error(['Property equalBinBreaks in ',mfilename,' must be true or false.'])
          end
      end  
      
      function obj = set.tol(obj,myTol)
          if (myTol > 0)
               obj.tol = myTol;
          else
               error(['Property tol in ',mfilename,' must be > 0.'])
          end
      end      
       
      function obj = set.spreadOut(obj,mySpreadOut)
          if (mySpreadOut >= 0)
               obj.spreadOut = mySpreadOut;
          else
               error(['Property spreadOut in ',mfilename,' must be >= 0.'])
          end
      end  
	  
      function obj = set.minIndPVal(obj,myMinIndPVal)
          if (myMinIndPVal >= 0) & (myMinIndPVal <= 1)
               obj.minIndPVal = myMinIndPVal;
          else
               error(['Property minIndPVal in ',mfilename,' must be between 0 and 1.'])
          end
      end 			  

      function obj = set.useEffN(obj,myUseEffN)
          if islogical(myUseEffN)
               obj.useEffN = myUseEffN;
          else
               error(['Property useEffN in ',mfilename,' must be logical.'])
          end
      end  

      function obj = set.exactFlag(obj,myFlag)
          if islogical(myFlag)
               obj.exactFlag = myFlag;
          else
               error(['Property exactFlag in ',mfilename,' must be logical.'])
          end
      end     
      
      function obj = set.optimizeTimeLimit(obj,myOptimizeTimeLimit)
          obj.optimizeTimeLimit = myOptimizeTimeLimit;
      end  
      
      function obj = set.optimizeType(obj,myOptimizeType)
          if sum(ismember({'pso','papso','ga','gapso','gapapso','simplex','surrogate'},lower(myOptimizeType))) == 1
              if sum(ismember({'papso','gapapso'},lower(myOptimizeType))) == 1
				opts = optimoptions('particleswarm');
				if ~isprop(opts,'UseAsync')
					if isequal('papso',lower(myOptimizeType))
						disp(['Parallel asynchronous mode not available for ',lower(myOptimizeType),' in ',mfilename,'.  Using pso.'])
						myOptimizeType = 'pso';
					else
						disp(['Parallel asynchronous mode not available for ',lower(myOptimizeType),' in ',mfilename,'.  Using gapso.'])
						myOptimizeType = 'gapso';
					end
				end
              end
			  obj.optimizeType = lower(myOptimizeType);
          else
              error(['Property optimizeType in ',mfilename,' must be "ga," "pso," "gapso," "simplex," or "surrogate.  Parallel asynchronous options may also be available, "papso" and "gapapso."'])
          end
      end     
      
      function obj = set.optimizePopSize(obj,myPopulationSize)
          if ((isnumeric(myPopulationSize) == true) && (myPopulationSize >= 0))
              obj.optimizePopSize = myPopulationSize;
          else
            error('Invalid optimizePopSize value')
          end
      end     
	  
      function obj = set.objectiveLimit(obj,myObjectiveLimit)
          if ((isnumeric(myObjectiveLimit) == true))
              obj.objectiveLimit = myObjectiveLimit;
          else
            error('Invalid objectiveLimit value')
          end
      end 	  	  
	  
      function obj = set.poolClose(obj,myInput)
          if islogical(myInput)
               obj.poolClose = myInput;
          else
               error(['Property poolClose in ',mfilename,' must be logical.'])
          end
      end  	  
	  
      function obj = set.poolRestart(obj,myInput)
          if islogical(myInput)
               obj.poolRestart = myInput;
          else
               error(['Property poolRestart in ',mfilename,' must be logical.'])
          end
      end  			  
      
      function obj = set.intSeed(obj,myIntSeed)
          if ((mod(myIntSeed,1) == 0) && (myIntSeed>=-1))
              obj.intSeed = (myIntSeed);
          else
              error(['Invalid intSeed specified for ',mfilename,', a non-negative integer should be specified, or -1 to ignore.'])
          end
      end
      
      function obj = set.minEffN(obj,myMinEffN)
          if ((isnumeric(myMinEffN) == true) && (myMinEffN >= 0))
              obj.minEffN = myMinEffN;
          else
              error('Invalid minEffN value')
          end
      end         
      
      function value = get(obj,propName)
          switch propName
              case 'expData'
                  value = obj.expData;
              case 'mnSDTable'
                  value = obj.mnSDTable;
              case 'binTable'
                  value = obj.binTable;  
              case 'distTable'
                  value = obj.distTable;                   
              case 'distTable2D'
                  value = obj.distTable2D;
              case 'corTable'
                  value = obj.corTable;	
              case 'subpopTable'
                  value = obj.subpopTable;					  
              case 'pwStrategy'
                  value = obj.pwStrategy;
              case 'nBins'
                  value = obj.nBins;
              case 'initialProbs'
                  value = obj.initialProbs;
              case 'randomStart'
                  value = obj.randomStart;
              case 'nIters'
                  value = obj.nIters;
              case 'equalBinBreaks'
                  value = obj.equalBinBreaks;
              case 'tol'
                  value = obj.tol;
              case 'spreadOut'
                  value = obj.spreadOut;
              case 'minIndPVal'
                  value = obj.minIndPVal;				  
              case 'useEffN'
                  value = obj.useEffN;
              case 'exactFlag'
                  value = obj.exactFlag;                  
              case 'optimizeTimeLimit'
                  value = obj.optimizeTimeLimit;
              case 'optimizeType'
                  value = obj.optimizeType;  
              case 'optimizePopSize'
                  value = obj.optimizePopSize;
              case 'objectiveLimit'
                  value = obj.objectiveLimit;	
              case 'poolRestart'
                  value = obj.poolRestart;
              case 'poolClose'
                  value = obj.poolClose;					  
              case 'intSeed'
                  value = obj.intSeed;   
              case 'minEffN'
                  value = obj.minEffN; 
              otherwise
                  error(['Error: ',propName ,' is not a valid ',mfilename,' property.'])
          end
      end

      % TODO: ADD METHODS TO VERIFY AGAINST WORKSHEET
            
      function obj = mapelOptions()
          % This is the constructor method for instances of mapelOptions
          % objects.
          %
          obj.expData = [];
          obj.mnSDTable = [];
          obj.binTable = [];
          obj.distTable = [];          
		  obj.distTable2D = []; 
		  obj.corTable = []; 
          obj.subpopTable = []; 			  
		  obj.pwStrategy = 'direct';								  
          obj.nBins = 2;
          obj.initialProbs = [];
          obj.initialPWs = [];	
          obj.randomStart = 0;
          obj.nIters = 10000;
          obj.equalBinBreaks = false;
          obj.tol = 1E-3;
          obj.spreadOut = 0;
          obj.minIndPVal = 0;			  		  		  
          obj.useEffN = false;
          obj.exactFlag = true;
          obj.optimizeTimeLimit = 10*60;
          obj.optimizeType = 'pso';
          obj.optimizePopSize = 1000;
		  obj.objectiveLimit = -Inf;	
          obj.poolClose = true;  
		  obj.poolRestart = true;			  
          obj.intSeed = -1;   
          obj.minEffN = 0;          
      end
   end
end
```
