## Supplementary File 3 for "A surrogate workflow for virtual population development: designing an amenable proxy for a hypothesis-test inspired nonlinear goodness-of-fit": VPopRECIST.m

```
classdef VPopRECIST
% This is a virtual population class.  This contains the results if a call
% to MAPEL.  There should be too much here to that needs to
% be changed directly, with the biggest possible exception being adjusting 
% useEffN before re-evaluating the GOF.  However, there is a substantial
% amount of customization possible by resuming fits with an existing
% VPop so you may find reasons to modify the properties.
%
% This object uses RECIST-based classifications for VP responses.  This
% impacts population observations.  For example, once a VP has observed "PD"
% biomarkers for later time points will not be included in calculations since
% this VP would generally be off trial.
%
% PROPERTIES:
%  coeffsTable:  (Don't manipulate) A nAxis X nVP matrix with VP the
%                 coefficients used for prevalence weights with the bin pw strategy
%  coeffsDist:   (Don't manipulate) A nVP X vVP matrix with VP the
%                 distance for the axis coefficients
%  pwStrategy:	 Strategy for the optimization.  Allowed values:
%                 'direct' (default value) If this is set then PWs are optimized directly
%                 'bin'     specified in a strategy similar to teh original MAPEL algorithm 
%  indexTable:   (Don't manipulate) A nAxis X nVP table that indicates in which 
%                 axis bin each VP falls into.  This is usually populated at 
%                 the beginning of the MAPEL algorithm by the assignIndices
%                 method.  ONLY NEEDED FOR BIN PWSTRATEGY
%  binEdges:     (Don't manipulate) Edges for the axis bins. These are also
%                 populated by the assignIndices method.
%  binMidPoints: (Don't manipulate) Mid points for the axis bins, populated
%                 by the assignIndices method.  ONLY NEEDED FOR BIN PWSTRATEGY
%  binProbs:     (Don't manipulate) Probabilities for each axis bin.  These are directly
%                 varied in the optimization, and the results are used
%                to calculate the individual prevalence weights.  ONLY NEEDED FOR BIN PWSTRATEGY
%  pws:          (Don't manipulate) A 1xnVP vector of prevalence weights,
%                which are calculated based on the assignPWs.
%  coefficients: a nAxisxnVP matrix of VP parameter coefficients.
%  expData:      (Don't manipulate) A table of experimental data used to
%                 guide optimization.  It is usually taken from the
%                 mapelOptions.
%  simData:      (Don't manipulate) A table of simulation results that 
%                 pairs to the experimental data.  Usually assigned in
%                 MAPEL.
%  mnSDTable:    (Don't manipulate) Summary table of experimental and 
%                simulated (weighted) mean and standard deviation.  Usually
%                copied from the mapelOptions and updated with
%                the weighted simulation results from the VPop fitting.
%                If you want to adjust a fit and restart, you may want to 
%                adjust some of the weights here.
%  binTable:     (Don't manipulate) Summary table of experimental and  
%                simulated binned (weighted) data for a virtual population
%                Usually copied from the mapelOptions and updated with
%                the weighted simulation results from the VPop fitting.
%                If you want to adjust a fit and restart, you may want to 
%                adjust some of the weights here.
%  distTable:    (Don't manipulate) Summary table of experimental and  
%                simulated distribution data for a virtual population
%                Usually copied from the mapelOptions and updated with
%                the weighted simulation results from the VPop fitting.
%                If you want to adjust a fit and restart, you may want to 
%                adjust some of the weights here. 						
%  distTable2D:	 (Don't manipulate) A table to enable calibrating 2D distributions
%  corTable:	 (Don't manipulate) A table to enable calibrating pairwise correlations
%  brTableRECIST: (Don't manipulate) A table with best RECIST responses
%  rTableRECIST: (Don't manipulate) A table with RECIST responses to better 
%  subpopTable:  (Don't manipulate) Contains criteria to create subpopulations
%                 from simulated VPs.
%                calibrate the distributions at each time step
%  gofMn:        (Don't manipulate) Goodness of fit statistics for 
%                individual endpoint/time mean comparisons between data
%                and virtual population.  Usually calcualated by the
%                evaluateGOF function.
%  gofSD:        (Don't manipulate) Goodness of fit statistics for 
%                individual endpoint/time standard deviation comparisons  
%                between data and virtual population.  Usually calculated 
%                by the evaluateGOF function.
%  gofBin:       (Don't manipulate) Goodness of fit statistics for 
%                individual endpoint/time bin distribution comparisons 
%                between data and virtual population.  Usually calculated 
%                by the evaluateGOF function.
%  gofDist:      (Don't manipulate) Goodness of fit statistics for 
%                empirical distribution comparisons 
%                between data and virtual population.  Usually calculated 
%                by the evaluateGOF function.	
%  gofCor:			
%  gofBR:
%  gofR:
%  gof:          (Don't manipulate) Composite goodness-of-fit result.
%                Usually updated by a call to the evaluateGOF.
%  useEffN:      Whether or not to use the effN for optimization and
%                evaluation of the GOF.  It is generally recommended to
%                useEffN for evaluation of the final fit but not
%                during optimization.
%  exactFlag:    Whether to check to apply Fisher's exact test instead
%                of the chi-square approximation for binned distribution
%                test.  This may be strictly correct but can slow
%                calculations, often with small impacts on GOF
%                calculations.																	  	  
%  spreadOut:    Used for optimization, if nonzero, will  
%                apply a penalty proportional to spreadOut to increase how
%                the prevalence weight is distributed.
%  minIndPVal:   Used for optimization, if nonzero, will  
%                apply a large penalty if any of the individual pvalues.
%                fall below the target.
%  optimizeTimeLimit:   Time limit for optimizing the VPop in s.
%  optimizeType:     	Type of optimization algorithm to employ.  Default is
%                    	"pso".  Options:
%							"ga" - MATLAB's GA
%							"pso" - MATLAB's PSO
%							"papso" - MATLAB's PAPSO
%							"gapso" - MATLAB's GA, polished by MATLAB's PSO
%							"gapapso" - MATLAB's GA, polished by MATLAB's PAPSO
%							"simplex" - MATLAB's simplex
%							"surrogate" - a short run of MATLAB's surrogate,
%                                     polished by MATLAB's PSO
%  optimizePopSize:     Number of solutions to try in each optimization
%                       generation.  Most directly impacts GA and PSO steps.  This 
%                       is the population size to use in the optimization 
%                       runs. Default is 1000.
%  objectiveLimit:		stopping condition for optimization
%  poolClose:           Whether to close the pool at the end of the optimization  
%  poolRestart:         Whether to restart the pool at the beginning of the optimization 
%  intSeed:             A non-negative integer seed to initialize the 
%                       random number generator.  Set to -1 to avoid
%                       changing the state of the random number generator.
%  tol:                 Numerical tolerance for optimization.
%  nIters:             (Optional) default is 10,000.
%                      simplex: number of iterations
%                      ga,gapso: maximum number of generations
%                      ignored for other optimizeType options.
%  minEffN:             Minimum effective N.  A large penalty is applied
%                       if the effN drops below this N during optimization,
%                       better ensuring solutions that weight multiple VPs.
%                       The minEffN setting operates independently of 
%                       useEffN, and often works better than modifying 
%                       spreadOut.  The default is 0.
%  relSLDvar:         a variable name that will be used for RECIST classification
%  absALDVar:         a variable to indicate lesion size, used for CR cutoff
%  crCutoff:          numeric value for diameter to use for CR cutoff
%  recistSimFilter:   A structure with data on which simulated patients are
%                    on therapy
%  MSE:                  mean sum of residuals from linear calibration (subpopulation/dropouts weighted)
%  LinearProblemMatrices:  the linearMatrix made from oldVPop linearCalibrationObject, actually this will be the reduced matrix for the rescaled oldVPop with only nVPmax columns. This is for the next iteration MSE minimization
%  LinearProblemMatricesSubgroupSumWeights: the subweight directly recorded fom oldVPop linearCalibrationObject
%  LinearProblemMatricesobservationDescriptions: the subweightDescription directly fom oldVPop linearCalibrationObject, to keep track of biomarkers during cohort expansion
%  lambda:            the inequality constraint from linearCalibration "fmincon" or the regularization parameter used during linearCalibration "quadprogEffN" 

properties
	coeffsTable
	coeffsDist
	pwStrategy
    indexTable
    binEdges
    binMidPoints
    binProbs
    pws
    expData
    simData      
    mnSDTable
    binTable
    distTable
	distTable2D
	corTable
    brTableRECIST
    rTableRECIST  
	subpopTable	
    gofMn
    gofSD
    gofBin
    gofDist	
	gofDist2D
	gofCor
    gofBR    
    gofR    
    gof
    useEffN  
    exactFlag			 
    spreadOut     
	minIndPVal	
    optimizeTimeLimit
    optimizeType
    optimizePopSize
	objectiveLimit
	poolClose
	poolRestart
    intSeed  
    tol
    nIters
    minEffN
    relSLDvar 
    absALDVar
    crCutoff    
    recistSimFilter    
    MSE
    LinearProblemMatrices
    LinearProblemMatricesSubgroupSumWeights
    LinearProblemMatricesobservationDescriptions
    lambda
end

methods
      function obj = set.coeffsTable(obj,myCoeffsTable)
          obj.coeffsTable = myCoeffsTable;
      end
      function obj = set.coeffsDist(obj,myCoeffsDist)
          obj.coeffsDist = myCoeffsDist;
      end	  	  
      function obj = set.pwStrategy(obj,myPWStrategy)
          if sum(ismember({'direct','bin'},lower(myPWStrategy))) == 1
              obj.pwStrategy = lower(myPWStrategy);
          else
              error(['Property pwStrategy in ',mfilename,' must be "direct" or "bin"'])
          end
      end   	  
      function obj = set.indexTable(obj,myIndexTable)
          obj.indexTable = myIndexTable;
      end

      function obj = set.binEdges(obj,myBins)
          obj.binEdges = myBins;
      end

      function obj = set.binMidPoints(obj,myBinMidPoints)
          obj.binMidPoints = myBinMidPoints;
      end

      function obj = set.binProbs(obj,myBinProbs)
          % We enforce constraints for binProbs:
          % a minimum value is 1E-14 rows must sum to 1
          % This helps avoid issues with the solver as well
          [nrows,ncols] = size(myBinProbs);
          if ((nrows > 0) && (ncols > 0))
            myBinProbs = myBinProbs ./ (repmat(sum(myBinProbs,2),1,ncols));
            adjustFlag = myBinProbs < 1E-14;
            myNadjust = sum(adjustFlag,2);
            myValDecrease = 1E-14 ./ myNadjust; 
            myValDecrease(isinf(myValDecrease)) = 0 ;
            myBinProbs = (myBinProbs + adjustFlag * 1E-14) - ((1- adjustFlag) .* (repmat(myValDecrease,1,ncols)));
            myBinProbs = myBinProbs ./ (repmat(sum(myBinProbs,2),1,ncols));
          end
          obj.binProbs = myBinProbs;
      end
	  
      function obj = set.pws(obj,myPWs)
          obj.pws = myPWs;
      end
	  
      function obj = set.mnSDTable(obj,myMnSDTable)
          obj.mnSDTable = myMnSDTable;
      end

      function obj = set.binTable(obj,myBinTable)
          obj.binTable = myBinTable;
      end

      function obj = set.distTable(obj,myDistTable)
          obj.distTable = myDistTable;
      end    				    
      function obj = set.distTable2D(obj,myDistTable)
          obj.distTable2D = myDistTable;
      end     													 						
      function obj = set.corTable(obj,myCorTable)
          obj.corTable = myCorTable;
      end       
      function obj = set.brTableRECIST(obj,myBRTableRECIST)
          obj.brTableRECIST = myBRTableRECIST;
      end 
      
      function obj = set.rTableRECIST(obj,myRTableRECIST)
          obj.rTableRECIST = myRTableRECIST;
      end 

      function obj = set.subpopTable(obj,mySubpopTable)
          obj.subpopTable = mySubpopTable;
      end	  	  
      
      function obj = set.expData(obj,myExpData)
          obj.expData = myExpData;
      end

      function obj = set.simData(obj,mySimData)
          obj.simData = mySimData;
      end

      function obj = set.gofMn(obj,myGOF)
          obj.gofMn = myGOF;
      end

      function obj = set.gofSD(obj,myGOF)
          obj.gofSD = myGOF;
      end
      
      function obj = set.gofBin(obj,myGOF)
          obj.gofBin = myGOF;
      end     

      function obj = set.gofDist(obj,myGOF)
          obj.gofDist = myGOF;
      end   
      
      function obj = set.gofDist2D(obj,myGOF)
          obj.gofDist2D = myGOF;
      end 

      function obj = set.gofCor(obj,myGOF)
          obj.gofCor = myGOF;
      end 
	  
      function obj = set.gofBR(obj,myGOF)
          obj.gofBR = myGOF;
      end      
      
      function obj = set.gofR(obj,myGOF)
          obj.gofR = myGOF;
      end           
      
      function obj = set.gof(obj,myGOF)
          obj.gof = myGOF;
      end          
      
      function obj = set.useEffN(obj,myUseEffN)
          if islogical(myUseEffN)
               obj.useEffN = myUseEffN;
          else
               error(['Property useEffN in ',mfilename,' must be logical.'])
          end
      end  
      
      function obj = set.exactFlag(obj,myFlag)
          obj.exactFlag = myFlag;
      end         
      
      function obj = set.spreadOut(obj,mySpreadOut)
          if (mySpreadOut >= 0)
               obj.spreadOut = mySpreadOut;
          else
               error(['Property spreadOut in ',mfilename,' must be >= 0.'])
          end
      end    
	  
      function obj = set.minIndPVal(obj,myMinIndPVal)
          if (myMinIndPVal >= 0) & (myMinIndPVal <= 1)
               obj.minIndPVal = myMinIndPVal;
          else
               error(['Property minIndPVal in ',mfilename,' must be between 0 and 1.'])
          end
      end 	 	  
      
      function obj = set.optimizeTimeLimit(obj,myOptimizeTimeLimit)
          obj.optimizeTimeLimit = myOptimizeTimeLimit;
      end  
      
      function obj = set.optimizeType(obj,myOptimizeType)
          if sum(ismember({'pso','papso','ga','gapso','gapapso','simplex','surrogate'},lower(myOptimizeType))) == 1
              if sum(ismember({'papso','gapapso'},lower(myOptimizeType))) == 1
				opts = optimoptions('particleswarm');
				if ~isprop(opts,'UseAsync')
					if isequal('papso',lower(myOptimizeType))
						disp(['Parallel asynchronous mode not available for ',lower(myOptimizeType),' in ',mfilename,'.  Using pso.'])
						myOptimizeType = 'pso';
					else
						disp(['Parallel asynchronous mode not available for ',lower(myOptimizeType),' in ',mfilename,'.  Using gapso.'])
						myOptimizeType = 'gapso';
					end
				end
              end
			  obj.optimizeType = lower(myOptimizeType);
          else
              error(['Property optimizeType in ',mfilename,' must be "ga," "pso," "gapso," "simplex," or "surrogate.  Parallel asynchronous options may also be available, "papso" and "gapapso."'])
          end
      end
      
      function obj = set.optimizePopSize(obj,myPopulationSize)
          if ((isnumeric(myPopulationSize) == true) && (myPopulationSize >= 0))
              obj.optimizePopSize = myPopulationSize;
          else
            error('Invalid optimizePopSize value')
          end
      end      

      function obj = set.objectiveLimit(obj,myObjectiveLimit)
          if ((isnumeric(myObjectiveLimit) == true))
              obj.objectiveLimit = myObjectiveLimit;
          else
            error('Invalid objectiveLimit value')
          end
      end 	       
	  
      function obj = set.poolClose(obj,myInput)
          if islogical(myInput)
               obj.poolClose = myInput;
          else
               error(['Property poolClose in ',mfilename,' must be logical.'])
          end
      end  	  
	  
      function obj = set.poolRestart(obj,myInput)
          if islogical(myInput)
               obj.poolRestart = myInput;
          else
               error(['Property poolRestart in ',mfilename,' must be logical.'])
          end
      end  		 	  
      
      function obj = set.intSeed(obj,myIntSeed)
          if ((mod(myIntSeed,1) == 0) && (myIntSeed>=-1))
              obj.intSeed = (myIntSeed);
          else
              error(['Invalid intSeed specified for ',mfilename,', a non-negative integer should be specified, or -1 to ignore.'])
          end
      end      
      
      function obj = set.tol(obj,myTol)
          if (myTol > 0)
               obj.tol = myTol;
          else
               error(['Property tol in ',mfilename,' must be > 0.'])
          end
      end       
      
      function obj = set.nIters(obj,myNIters)
          if (myNIters > 0) && (mod(myNIters,1) == 0)
               obj.nIters = myNIters;
          else
               error(['Property nIters in ',mfilename,' must be a positive integer.'])
          end
      end         
      
      function obj = set.minEffN(obj,myMinEffN)
          if ((isnumeric(myMinEffN) == true) && (myMinEffN >= 0))
              obj.minEffN = myMinEffN;
          else
            error('Invalid minEffN value')
          end
      end       
      
      function obj = set.relSLDvar(obj,myRelSLDvar)
          obj.relSLDvar = myRelSLDvar;
      end       
      
      function obj = set.absALDVar(obj,myAbsALDVar)
          obj.absALDVar = myAbsALDVar;
      end     
      
      function obj = set.crCutoff(obj,myCRCutoff)
          obj.crCutoff = myCRCutoff;
      end       
      
      function obj = set.recistSimFilter(obj,myRECISTSimFilter)
          obj.recistSimFilter = myRECISTSimFilter;
      end
      
      function value = get(obj,propName)
          switch propName
              case 'coeffsTable'
                  value = obj.coeffsTable;		
			  case 'coeffsDist'
                  value = obj.coeffsDist;	
              case 'pwStrategy'
                  value = obj.pwStrategy;				  
              case 'indexTable'
                  value = obj.indexTable;
              case 'binEdges'
                  value = obj.binEdges;
              case 'binMidPoints'
                  value = obj.binMidPoints;
              case 'binProbs'
                  value = obj.binProbs;   
              case 'pws'
                  value = obj.pws; 				  			  
              case 'mnSDTable'
                  value = obj.mnSDTable; 
              case 'binTable'
                  value = obj.binTable; 
              case 'distTable'
                  value = obj.distTable;   							  
              case 'distTable2D'
                  value = obj.distTable2D;
              case 'corTable'
                  value = obj.corTable;				  
              case 'brTableRECIST'
                  value = obj.brTableRECIST;   
              case 'rTableRECIST'
                  value = obj.rTableRECIST; 
              case 'subpopTable'
                  value = obj.subpopTable; 				  
              case 'expData'
                  value = obj.expData;                  
              case 'simData'
                  value = obj.simData;            
              case 'gofMn'
                  value = obj.gofMn;      
              case 'gofSD'
                  value = obj.gofSD;   
              case 'gofBin'
                  value = obj.gofBin;   
              case 'gofDist'
                  value = obj.gofDist;   
              case 'gofDist2D'
                  value = obj.gofDist2D;   
              case 'gofCor'
                  value = obj.gofCor;   
              case 'gofBR'
                  value = obj.gofBR;   
              case 'gofR'
                  value = obj.gofR;                     
              case 'gof'
                  value = obj.gof;                     
              case 'spreadOut'
                  value = obj.spreadOut;
              case 'minIndPVal'
                  value = obj.minIndPVal;				  
              case 'useEffN'
                  value = obj.useEffN;
              case 'exactFlag'
                  value = obj.exactFlag;     
              case 'optimizeTimeLimit'
                  value = obj.optimizeTimeLimit;
              case 'optimizeType'
                  value = obj.optimizeType;  
              case 'optimizePopSize'
                  value = obj.optimizePopSize;
              case 'objectiveLimit'
                  value = obj.objectiveLimit;	
              case 'poolRestart'
                  value = obj.poolRestart;
              case 'poolClose'
                  value = obj.poolClose;				  
              case 'intSeed'
                  value = obj.intSeed;             
              case 'tol'
                  value = obj.tol;
              case 'nIters'
                  value = obj.nIters;       
              case 'minEffN'
                  value = obj.minEffN;  
              case 'relSLDvar'
                  value = obj.relSLDvar; 
              case 'absALDVar'
                  value = obj.absALDVar; 
              case 'crCutoff'
                  value = obj.crCutoff; 
              case 'recistSimFilter'
                  value = obj.recistSimFilter;                  
              otherwise
                  error(['Error: ',propName ,' is not a valid ',mfilename,' property.'])
          end
      end
	  
      function obj = assignCoeffs(obj,myWorksheet)
          mycoeffsTable = getVPCoeffs(myWorksheet);
          obj.coeffsTable = mycoeffsTable;
      end	  

      function obj = assignIndices(obj, myWorksheet, myMapelOptions)
          % This method takes a worksheet and mapelOptions structure and
          % creates a bin table and other poperties needed for the virtual
          % population and MAPEL algorithm.
          %
          % ARGUMENT
          %  (self)
          %  myWorksheet:    a worksheet structure, with VPs, axes, and
          %                  coefficients
          %  myMapelOptions: an MAPEL options object instance, with
          %                  properties
          %                   nBins
          %                   equalBinBreaks
          %
          % RETURNS
          %  (self): properties of the virtual population are updated:
          %           indexTable
          %           binEdges
          %           binMidPoints
          %  
          
          myVPCoeffs = getVPCoeffs(myWorksheet);
          [nAxis, nVP] = size(myVPCoeffs);
          myIndexTable = ones(nAxis, nVP);
          nBins = myMapelOptions.nBins;
          myBinEdges = nan(nAxis, (nBins+1));
          myBinMidPoints = nan(nAxis, nBins);
          equalBinBreaks = myMapelOptions.equalBinBreaks;
          for axisCounter = 1 : nAxis
              % We currently only implement continuous variables from
              % the paper/R MAPEL   
              if equalBinBreaks
                   % Rather than scaling based on the sampled min and
                   % max as in the R MAPEL algorithm, we adjust according
                   % to the allowed,
                   % and this range is [0, 1] by the axis definition.
                   myBinEdges(axisCounter,:) =  (0:1/nBins:1);
              else
                   myPercentiles =  (0:1/nBins:1)';
                   myBinEdges(axisCounter,:) = (quantile(myVPCoeffs(axisCounter,:),myPercentiles));
              end
              % We'll use the cdf convention that FX(x) = P(X <= x), so 
              % check if a value is <= the bin upper cutoff in order to 
              % assign it.
              % Note that effectively we want to ignore the first bin of 0
              % as a cutoff and lump this in with the first bin.
              for binCounter = 1 : nBins
                   myBinMidPoints(axisCounter,binCounter) = (myBinEdges(axisCounter,binCounter) + myBinEdges(axisCounter,binCounter+1))/2;
                   % First bins are assigned an index of 0 by this method.
                   myIndexTable(axisCounter, :) = myIndexTable(axisCounter, :) + (myVPCoeffs(axisCounter, :) > myBinEdges(axisCounter,binCounter+1));
              end     
          end
          obj.indexTable = myIndexTable;
          obj.binEdges = myBinEdges;
          obj.binMidPoints = myBinMidPoints;
      end

      function obj = startProbs(obj, myRandomStart)
          % This method initializes bin probabilities, and is used
          % prior to the optization.
		  % Only relevant for obj.pwStrategy = 'bin'
          %
          % ARGUMENTS
          %  (self)
          %  myRandomStart:  A boolean variable (true/false), if true
          %                  the bin probabilities will be set
          %                  randomly and if false they will be uniform
           if nargin < 1
                myRandomStart = false;
           end
           myBinMidPoints = obj.binMidPoints;
           [nAxes, nBins] = size(myBinMidPoints);
           if ~myRandomStart
                myUniformStartProbs = ones(nAxes, nBins) ./ nBins;
           else
                myUniformStartProbs = rand([nAxes, nBins]);
                for axisCounter = 1 : nAxes
                    myUniformStartProbs(axisCounter,:) = myUniformStartProbs(axisCounter,:)/sum(myUniformStartProbs(axisCounter,:));
                end
           end
           obj.binProbs = myUniformStartProbs;
      end

      function obj = assignPWs(obj)
          % Update prevalence weight assignment for the virtual population.
          % Note that existing predicted population results will be set
          % to NaN in:
          %  mnSDTable
          %  binTable
		  %  distTable
		  %  distTable2D
		  %  corTable		  
          %  brTableRECIST
          %  rTableRECIST          
          %
          % And also the following properties will be reset:
          %  gofMn
          %  gofSD
          %  gofBin
		  %  gofDist
		  %  gofDist2D
		  %  gofCor		  
          %  gofBR
          %  gofR          
          %  gof
          %
		  % Only relevant for obj.pwStrategy = 'axis'
          % ARGUMENTS:
          %  (self): No additional arguments, but the VPop object must have
          %          previously assigned the following properties:
          %           indexTable
          %           binProbs
          %
          % RETURNS:
          %  (self): Returns the VPop with the updated PW properties.
          %
           myIndexTable = obj.indexTable;
           [myNAxis, myNVP] = size(myIndexTable);
           myPWs = zeros(1,myNVP);
           myBinProbs = obj.binProbs;
           %pwCutoff = obj.pwCutoff;
           for axisCounter = 1 : myNAxis
               % Note this will result in -Inf if any bins have
               % prob of zero.  MATLAB's exponential at the end corrects
               % this, but there is a danger of getting back all
               % -inf if all of the bin probabilities are small.
                myPWs = myPWs + log(myBinProbs(axisCounter,myIndexTable(axisCounter,:)));
           end
           % Carried over from paper MAPEL: force
           % biggest value to ~1 to avoid underflow
           myPWs = myPWs - max(myPWs);
           myPWs = exp(myPWs); %/ sum(exp(myPWs));
           %myPWs = myPWs.*(myPWs >= pwCutoff);
           myPWs = myPWs / sum(myPWs);
           obj.pws = myPWs;
           
           % If we re-calculate PWs, eliminate any
           % previous PW-dependent properties - i.e. those
           % that start with "pred" except predIndices
           myMnSDTable = obj.mnSDTable;
           [nRows, nCols] = size(myMnSDTable);
           if nRows > 0
                % myString = 'pred';
                % varNames = myMnSDTable.Properties.VariableNames;
                % myPos = find(strncmpi(myString,varNames,length(myString)));
                % nanifyVars = varNames(myPos);
                nanifyVars = {'predN','predMean','predSD'};
                for varCounter = 1 : length(nanifyVars)
                    myMnSDTable.(nanifyVars{varCounter}) = nan(nRows,1);
                end
           end
           obj.mnSDTable = myMnSDTable;

           myBinTable = obj.binTable;
           [nRows, nCols] = size(myBinTable);
           if nRows > 0
                % myString = 'pred';
                % varNames = myBinTable.Properties.VariableNames;
                % myPos = find(strncmpi(myString,varNames,length(myString)));
                % nanifyVars = varNames(myPos);
                nanifyVars = {'predN','predBins'};             
                for varCounter = 1 : length(nanifyVars)
                    myBinTable.(nanifyVars{varCounter}) = nan(nRows,1);
                end
           end
           obj.binTable = myBinTable;

           myDistTable = obj.distTable;
           [nRows, nCols] = size(myDistTable);
           if nRows > 0
                myNanStrings = {'predN'};
                varNames = myDistTable.Properties.VariableNames;
                myPos = find(ismember(varNames,myNanStrings));
                nanifyVars = varNames(myPos);
                for varCounter = 1 : length(nanifyVars)
                    myDistTable.(nanifyVars{varCounter}) = nan(nRows,1);
                end
                myNanStrings = {'predProbs'};
                varNames = myDistTable.Properties.VariableNames;
                myPos = find(ismember(varNames,myNanStrings));
                nanifyVars = varNames(myPos);
				myDistTable{1 : nRows,nanifyVars} = {nan};               
           end
           obj.distTable = myDistTable;   
		   
           myTable = obj.distTable2D;
		   if ~isempty(myTable)
			   [nRows, nCols] = size(myTable);
			   if nRows > 0
					myNanStrings = {'predN'};
					varNames = myTable.Properties.VariableNames;
					myPos = find(ismember(varNames,myNanStrings));
					nanifyVars = varNames(myPos);
					for varCounter = 1 : length(nanifyVars)
						myTable.(nanifyVars{varCounter}) = nan(nRows,1);
					end
					myNanStrings = {'predProbs'};
					varNames = myTable.Properties.VariableNames;
					myPos = find(ismember(varNames,myNanStrings));
					nanifyVars = varNames(myPos);
					myTable{1 : nRows,nanifyVars} = {nan};               
			   end
			   obj.distTable2D = myTable; 
		   end	

           myTable = obj.corTable;
		   if ~isempty(myTable)
			   [nRows, nCols] = size(myTable);
			   if nRows > 0
					myNanStrings = {'predN','predCor'};
					varNames = myTable.Properties.VariableNames;
					myPos = find(ismember(varNames,myNanStrings));
					nanifyVars = varNames(myPos);
					for varCounter = 1 : length(nanifyVars)
						myTable.(nanifyVars{varCounter}) = nan(nRows,1);
					end
					myNanStrings = {'predProbs'};
					varNames = myTable.Properties.VariableNames;
					myPos = find(ismember(varNames,myNanStrings));
					nanifyVars = varNames(myPos);
					myTable{1 : nRows,nanifyVars} = {nan};               
			   end
			   obj.corTable = myTable; 
		   end			   
           
           myBRTable = obj.brTableRECIST;
           [nRows, nCols] = size(myBRTable);
           if nRows > 0
                myString = 'pred';
                varNames = myBRTable.Properties.VariableNames;
                myPos = find(strncmpi(myString,varNames,length(myString)));
                nanifyVars = varNames(myPos);
                for varCounter = 1 : length(nanifyVars)
                    myBRTable.(nanifyVars{varCounter}) = nan(nRows,1);
                end	   
           end
           obj.brTableRECIST = myBRTable;  
           
           myRTable = obj.rTableRECIST;
           [nRows, nCols] = size(myRTable);
           if nRows > 0
                myString = 'pred';
                varNames = myRTable.Properties.VariableNames;
                myPos = find(strncmpi(myString,varNames,length(myString)));
                nanifyVars = varNames(myPos);
                for varCounter = 1 : length(nanifyVars)
                    myRTable.(nanifyVars{varCounter}) = nan(nRows,1);
                end	   
           end
           obj.rTableRECIST = myRTable;   
          
           myTable = obj.subpopTable;
           [nRows, nCols] = size(myTable);
           if nRows > 1
                nanifyVars = 'predW';
                myTable.(nanifyVars{varCounter})(2:end) = nan(nRows-1,1);
           end
           obj.subpopTable = myTable; 
           
           obj.gofMn = [];
           obj.gofSD = [];
           obj.gofBin = [];
		   obj.gofDist = [];	
		   obj.gofDist2D = [];		   
		   obj.gofCor = [];		   
		   obj.gofBR = [];				
		   obj.gofR = [];				           
           obj.gof = [];
      end

      function obj = startPWs(obj, myWorksheet, myRandomStart)
          % This method initializes pws, and is used
          % prior to the optization.
		  % Only relevant for obj.pwStrategy = 'direct'
          %
          % ARGUMENTS
          %  (self)
          %  myRandomStart:  A boolean variable (true/false), if true
          %                  prevalence weights will be set
          %                  randomly and if false they will be uniform
           if nargin < 1
                myRandomStart = false;
           end
          mycoeffsTable = getVPCoeffs(myWorksheet);
          [nAxis, nVP] = size(mycoeffsTable);
           if ~myRandomStart
                myUniformStartProbs = ones(1,nVP) ./ nVP;
           else
                myUniformStartProbs = rand([1, nVP]);
				myUniformStartProbsSum=sum(myUniformStartProbs);
                for axisCounter = 1 : nVP
                    myUniformStartProbs(1,axisCounter) = myUniformStartProbs(1,axisCounter)/myUniformStartProbsSum;
                end
           end
           obj.pws = myUniformStartProbs;
      end	  
	  
      function obj = getSimData(obj, myWorksheet)
           % Get all of the simulation datapoints for the VPs that will
           % be needed for calculating population statistics
           %
           % ARGUMENTS
           %  (self):      Note that the following properties should be
           %               initialized (experimental data) before calling this
           %               method:
           %                mnSDTable
           %                binTable
           %                distTable
		   %                distTable2D
		   %				corTable
		   %                brTableRECIST
		   %                rTableRECIST           
		   %                recistSimFilter;
           %  myWorksheet: A worksheet with the simulation results.
           %
           % RETURNS
           %  (self):      The VPop object is returned, but with an updated
           %               simData property.
           %
           %
           continueFlag = true;
           mnsdDataFlag = false;
           binDataFlag = false;
           distDataFlag = false;
           distData2DFlag = false;	
		   corDataFlag = false;
           brDataFlag = false;	
           rDataFlag = false;		              
           myMnSdData = obj.mnSDTable;
           myBinTable = obj.binTable;
		   myDistTable = obj.distTable;
		   myDistTable2D = obj.distTable2D;	  
		   myCorTable = obj.corTable;
           myBRTableRECIST = obj.brTableRECIST;
           myRTableRECIST = obj.rTableRECIST;           
           myRECISTFilter = obj.recistSimFilter;
           
           % We want the {expVarID, element, elementType, time} sets
           % from the experimental summaries that will be paired
           % with real data.
           rowInfoNames = {'expVarID','interventionID','elementID','elementType','time','expDataID'};
		   rowInfoNames2D = {'expVarID1','expVarID2','interventionID1','interventionID2','elementID1','elementID2','elementType1','elementType2','time1','time2','expDataID1','expDataID2'};	
           rowInfoNames2D1 = {'expVarID1','interventionID1','elementID1','elementType1','time1','expDataID1'};
           rowInfoNames2D2 = {'expVarID2','interventionID2','elementID2','elementType2','time2','expDataID2'};
           brInfoNames = {'expVarID','interventionID','elementID','elementType','time'};
           rInfoNames = brInfoNames;
           myDataSource = cell2table(cell(0,6), 'VariableNames', rowInfoNames);
           if ~isempty(myMnSdData)
               myDataSource = [myDataSource; myMnSdData(:,rowInfoNames)];
               mnsdDataFlag = true;
           end
           if ~isempty(myBinTable)
               myDataSource = [myDataSource; myBinTable(:,rowInfoNames)];
               binDataFlag = true;
           end           
           if ~isempty(myDistTable)
               myDataSource = [myDataSource; myDistTable(:,rowInfoNames)];
               distDataFlag = true;
           end          
           if ~isempty(myDistTable2D)
               distData2DFlag = true;
               myDataSource1 = myDistTable2D(:,rowInfoNames2D1);
               myDataSource2 = myDistTable2D(:,rowInfoNames2D2);
               myDataSource1.Properties.VariableNames = rowInfoNames;
               myDataSource2.Properties.VariableNames = rowInfoNames;
               myDataSource = [myDataSource; myDataSource1];
               myDataSource = [myDataSource; myDataSource2];
		   end
           if ~isempty(myCorTable)
               corDataFlag = true;		
               myDataSource1 = myCorTable(:,rowInfoNames2D1);
               myDataSource2 = myCorTable(:,rowInfoNames2D2);
               myDataSource1.Properties.VariableNames = rowInfoNames;
               myDataSource2.Properties.VariableNames = rowInfoNames;
               myDataSource = [myDataSource; myDataSource1];
               myDataSource = [myDataSource; myDataSource2];               
           end
           if ~isempty(myBRTableRECIST)
               brDataFlag = true;
           end     
           if ~isempty(myRTableRECIST)
               rDataFlag = true;
           end                
		   
           [nrows,ncols] = size(myDataSource);
           if (nrows < 1) && ~brDataFlag && ~rDataFlag
               warning(['No mnSDTable, binTable, distTable, distTable2D, corTable, brTableRECIST, or rTableRECIST assigned before calling getSimData in ',mfilename,'. The data to get is not known. Exiting.'])
               continueFlag = false;
           end
           
           % TODO: verify check for RECIST data "Source"
           
           if continueFlag
               % Eliminate duplicate rows
               myDataSource = unique(myDataSource, 'rows');
               myDataSource = sortrows(myDataSource, ...
                   {'interventionID','elementID','time','elementType','expDataID'},...
                   {'ascend','ascend','ascend','ascend','ascend'});
           end
           
           if continueFlag
               if ~isempty(myDataSource)
                   myCheckVars = myDataSource.('elementID');
                   % If the length is 0, all variables should be written.
                   if length(myWorksheet.simProps.saveElementResultIDs) > 0
                       myMissingVars = ~ismember(myCheckVars,myWorksheet.simProps.saveElementResultIDs);
                       myMissingVars = myCheckVars(myMissingVars);
                       if length(myMissingVars) > 0
                           disp(['Missing needed variables for ',mfilename,' in myWorksheet.simProps.saveElementResultIDs.  They are: ',strjoin(myMissingVars,', '),'.  Attempting to continue...'])
                       end
                   end
               end
           end
               
           if continueFlag    
               
               [nEntries, ~] = size(myDataSource);

               % Info on each row of simvalues
               rowInfo = table2cell(myDataSource);
               % We will need to use a structure, like we did for results,
               % since some of the VP names will certainly be longer
               % than what is permitted by variable name lengths
               % Also, use the values
               % Row indices for the data are also added to speed subsequent calculations.

               vpIDs = getVPIDs(myWorksheet);
               nVPs = length(vpIDs);
               % Simulation results for each VP for each row
               dataValues = nan(nEntries, length(vpIDs));
               interventionIDIndex = find(ismember(rowInfoNames,'interventionID'));
               elementIDIndex = find(ismember(rowInfoNames,'elementID'));
               elementTypeIndex = find(ismember(rowInfoNames,'elementType'));
               expTimeIndex = find(ismember(rowInfoNames,'time'));
               expDataIDIndex = find(ismember(rowInfoNames,'expDataID'));
               simData.Data = dataValues;
               simData.rowInfoNames = rowInfoNames;
               simData.rowInfo = rowInfo;
               simData.vpIDs = vpIDs;
               interventionIDs = getInterventionIDs(myWorksheet);
               mnSDRows = nan(nEntries,1);
               binRows = nan(nEntries,1);
               distRows = nan(nEntries,1);
			   distRows2D = cell(nEntries,2);
			   corRows = cell(nEntries,2);

               for rowCounter = 1 : nEntries
                    interventionID = simData.rowInfo{rowCounter,interventionIDIndex};
                    elementID = simData.rowInfo{rowCounter,elementIDIndex};
                    elementType = simData.rowInfo{rowCounter,elementTypeIndex};
                    wshInterventionIndex = find(ismember(interventionIDs,interventionID));
                    expTime = simData.rowInfo{rowCounter,expTimeIndex};
                    expDataID = simData.rowInfo{rowCounter,expDataIDIndex};
                    simFilterValues = double(myRECISTFilter{wshInterventionIndex}.filterMatrix);
                    simFilterTimes = myRECISTFilter{wshInterventionIndex}.time;
                    simFilterValuesInf = simFilterValues;
                    simFilterValuesInf(find((simFilterValues<1)))=-1;
                    
                    % To avoid re-searching for the right rows on every
                    % iteration mapel, we provide the indices here 
                    if mnsdDataFlag
                        temp = find((ismember(myMnSdData{:,'interventionID'},interventionID)) ...
                                 & ((myMnSdData{:,'time'})==expTime) ...
                                 & (ismember(myMnSdData{:,'elementID'},elementID)) ...
                                 & (ismember(myMnSdData{:,'elementType'},elementType) ...
                                 & (ismember(myMnSdData{:,'expDataID'},expDataID))));
						if ~isempty(temp)
                            mnSDRows(rowCounter) = temp;
                        end
                    end
                    if binDataFlag
                       temp = find((ismember(myBinTable{:,'interventionID'},interventionID)) ...
                                 & ((myBinTable{:,'time'})==expTime) ...
                                 & (ismember(myBinTable{:,'elementID'},elementID)) ...
                                 & (ismember(myBinTable{:,'elementType'},elementType)) ...
                                 & (ismember(myBinTable{:,'expDataID'},expDataID)));
                        if ~isempty(temp)
                            binRows(rowCounter) = temp;
                        end
                    end
                    if distDataFlag
                       temp = find((ismember(myDistTable{:,'interventionID'},interventionID)) ...
                                 & ((myDistTable{:,'time'})==expTime) ...
                                 & (ismember(myDistTable{:,'elementID'},elementID)) ...
                                 & (ismember(myDistTable{:,'elementType'},elementType)) ...
                                 & (ismember(myDistTable{:,'expDataID'},expDataID)));
                        if ~isempty(temp)
                            distRows(rowCounter) = temp;
                        end
                    end
                    if distData2DFlag
                        % One row of source data may be involved in
                        % multiple 2D distributions
                        temp = find((ismember(myDistTable2D{:,'interventionID1'},interventionID)) ...
                                 & ((myDistTable2D{:,'time1'})==expTime) ...
                                 & (ismember(myDistTable2D{:,'elementID1'},elementID)) ...
                                 & (ismember(myDistTable2D{:,'elementType1'},elementType)));
                        if ~isempty(temp)
                            distRows2D{rowCounter,1} = temp;
                        end
                        temp = find((ismember(myDistTable2D{:,'interventionID2'},interventionID)) ...
                                 & ((myDistTable2D{:,'time2'})==expTime) ...
                                 & (ismember(myDistTable2D{:,'elementID2'},elementID)) ...
                                 & (ismember(myDistTable2D{:,'elementType2'},elementType)));
                        if ~isempty(temp)
                            distRows2D{rowCounter,2} = temp;
                        end						
                    end
                    if corDataFlag
                        % One row of source data may be involved in
                        % multiple 2D distributions
                        temp = find((ismember(myCorTable{:,'interventionID1'},interventionID)) ...
                                 & ((myCorTable{:,'time1'})==expTime) ...
                                 & (ismember(myCorTable{:,'elementID1'},elementID)) ...
                                 & (ismember(myCorTable{:,'elementType1'},elementType)));
                        if ~isempty(temp)
                            corRows{rowCounter,1} = temp;
                        end
                        temp = find((ismember(myCorTable{:,'interventionID2'},interventionID)) ...
                                 & ((myCorTable{:,'time2'})==expTime) ...
                                 & (ismember(myCorTable{:,'elementID2'},elementID)) ...
                                 & (ismember(myCorTable{:,'elementType2'},elementType)));
                        if ~isempty(temp)
                            corRows{rowCounter,2} = temp;
                        end						
                    end 					
                    % I don't see how to avoid looping this, given the way
                    % the data are structured.  Luckily we just get the data
                    % once 
                    rowIndex = nan;
                    for vpCounter = 1 : nVPs
                        % We need to verify the desired result for the VP is
                        % present, otherwise we report the simData result as
                        % nan
                        if length(myWorksheet.results) > 0
                            curResult = myWorksheet.results{wshInterventionIndex, vpCounter};
                            % Results should be stored in a structure, we 
                            % assume if a structure is provided then it is a 
                            % valid result
                            if strcmp(class(curResult),'struct')
                                curResult = myWorksheet.results{wshInterventionIndex, vpCounter};
                                curTimeIndex = find(ismember(curResult.Names,'time'));
                                curVarIndex = find(ismember(curResult.Names,elementID));
                                curTime = curResult.Data(:,curTimeIndex);
                                curVar = curResult.Data(:,curVarIndex);
                                % First check to see if the right row is
                                % already known.  We assume the result
                                % size is consistent across VPs.
                                if isnan(rowIndex)
                                    if sum(curTime == expTime) == 1
                                        rowIndex = find(curTime == expTime);
                                        interpolateValue = curVar(rowIndex);
                                    else
                                        interpolateValue = interp1(curTime,curVar,expTime,'linear');
                                        warning([num2str(expTime),' not found in the worksheet result, interpolation implemented ...']);
                                    end
                                else
                                    interpolateValue = curVar(rowIndex);
                                end
                                % Account for VPs dropping off of therapy
                                filterIndex = find(expTime <= (simFilterTimes .* simFilterValuesInf(:,vpCounter)));
                                
                                if length(filterIndex)>0
                                    filterIndex = filterIndex(1);
                                    simData.Data(rowCounter, vpCounter) = interpolateValue;
                                else
                                    simData.Data(rowCounter, vpCounter) = nan;
                                end
                            else
                                simData.Data(rowCounter, vpCounter) = nan;
                            end
                        else
                            simData.Data(rowCounter, vpCounter) = nan;
                        end
                    end
               end
               
               % BR results for each VP for each row
               if ~isempty(myBRTableRECIST)
                   myDataSource = myBRTableRECIST(:,rowInfoNames);
                   myDataSource = sortrows(myDataSource, ...
                       {'interventionID','elementID','time','elementType','expDataID'}, ...
                       {'ascend','ascend','ascend','ascend','ascend'});
               
                   brRowInfo = table2cell(myDataSource);

                   [nEntries, ~] = size(myDataSource);
                   dataValues = nan(nEntries, length(vpIDs));
                   interventionIDIndex = find(ismember(rowInfoNames,'interventionID'));
                   expTimeIndex = find(ismember(rowInfoNames,'time'));
                   expDataIDIndex = find(ismember(rowInfoNames,'expDataID'));
                   rData.Data = dataValues;

                   interventionIDs = getInterventionIDs(myWorksheet);
                   brRows = nan(nEntries,1);

                   for rowCounter = 1 : nEntries
                       interventionID = brRowInfo{rowCounter,interventionIDIndex};
                       wshInterventionIndex = find(ismember(interventionIDs,interventionID));
                       expTime = brRowInfo{rowCounter,expTimeIndex};
                       expDataID = brRowInfo{rowCounter,expDataIDIndex};
                       temp = find((ismember(myBRTableRECIST{:,'interventionID'},interventionID)) ...
                                & ((myBRTableRECIST{:,'time'})==expTime) ... 
                                & ismember(myBRTableRECIST{:,'expDataID'},expDataID));
                       brRows(rowCounter) = temp;
                       % To avoid re-searching for the right rows on every
                       % iteration mapel, we provide the indices here
                       if length(myWorksheet.results) > 0
                           curStruct = myRECISTFilter{wshInterventionIndex};
                           curTime = curStruct.time;                   
                           % I don't see how to avoid looping this, given the way
                           % the data are structured.  Luckily we just get the data
                           % once
                           rowIndex = nan;                       
                           for vpCounter = 1 : nVPs
                               % We need to verify the desired result for the VP is
                               % present, otherwise we report the simData result as
                               % nan
                               curResult = curStruct.bestResp(:,vpCounter);

                               % First check to see if the right row is
                               % already known.  We assume the result
                               % size is consistent across VPs.
                               if isnan(rowIndex)
                                   if sum(curTime == expTime) == 1
                                       rowIndex = find(curTime == expTime);
                                       interpolateValue = single(curResult(rowIndex));
                                   else
                                       % Interpolate to get the time exactly right
                                       % Might want to make this "last"
                                       % Interp1 demands double or single for input                                   
                                       interpolateValue = interp1(curTime,single(curResult),expTime,'previous');
                                   end
                               else
                                   interpolateValue = single(curResult(rowIndex));
                               end
                               % Convert back
                               brData.Data(rowCounter, vpCounter) = int8(interpolateValue);
                           end
                       end
                   end
               else
                   brRows = [];
                   brData.Data = [];
                   brRowInfo = []; 
               end
               
               % R results for each VP for each row
               if ~isempty(myRTableRECIST)
                   myDataSource = myRTableRECIST(:,rowInfoNames);
                   myDataSource = sortrows(myDataSource, ...
                       {'interventionID','elementID','time','elementType','expDataID'}, ...
                       {'ascend','ascend','ascend','ascend','ascend'});

                   rRowInfo = table2cell(myDataSource);

                   [nEntries, ~] = size(myDataSource);
                   dataValues = nan(nEntries, length(vpIDs));
                   interventionIDIndex = find(ismember(rowInfoNames,'interventionID'));
                   expTimeIndex = find(ismember(rowInfoNames,'time'));
                   expDataIDIndex = find(ismember(rowInfoNames,'expDataID'));
                   rData.Data = dataValues;

                   interventionIDs = getInterventionIDs(myWorksheet);
                   rRows = nan(nEntries,1);

                   for rowCounter = 1 : nEntries
                       interventionID = rRowInfo{rowCounter,interventionIDIndex};
                       wshInterventionIndex = find(ismember(interventionIDs,interventionID));
                       expTime = rRowInfo{rowCounter,expTimeIndex};
                       expDataID = rRowInfo{rowCounter,expDataIDIndex};
                       temp = find((ismember(myRTableRECIST{:,'interventionID'},interventionID)) ... 
                                & ((myRTableRECIST{:,'time'})==expTime) ...
                                & ismember(myRTableRECIST{:,'expDataID'},expDataID));
                       rRows(rowCounter) = temp;
                       % To avoid re-searching for the right rows on every
                       % iteration mapel, we provide the indices here

                       if length(myWorksheet.results) > 0
                           curStruct = myRECISTFilter{wshInterventionIndex};
                           curTime = curStruct.time;   
                           % I don't see how to avoid looping this, given the way
                           % the data are structured.  Luckily we just get the data
                           % once
                           rowIndex = nan;                       
                           for vpCounter = 1 : nVPs
                               % We need to verify the desired result for the VP is
                               % present, otherwise we report the simData result as
                               % nan
                               curResult = curStruct.curResp(:,vpCounter);

                               % First check to see if the right row is
                               % already known.  We assume the result
                               % size is consistent across VPs.
                               if isnan(rowIndex)
                                   if sum(curTime == expTime) == 1
                                       rowIndex = find(curTime == expTime);
                                       interpolateValue = single(curResult(rowIndex));
                                   else
                                       % Interpolate to get the time exactly right
                                       % Might want to make this "last"
                                       % Interp1 demands double or single for input                                   
                                       interpolateValue = interp1(curTime,single(curResult),expTime,'previous');
                                   end
                               else
                                   interpolateValue = single(curResult(rowIndex));
                               end
                               % Convert back
                               rData.Data(rowCounter, vpCounter) = int8(interpolateValue);
                           end
                       end
                   end
               else
                   rRows = [];
                   rData.Data = [];
                   rRowInfo = [];     
               end
               
               
               simData.binRows = binRows;
               simData.mnSDRows = mnSDRows;
               simData.distRows = distRows;
               simData.distRows2D = distRows2D;
			   simData.corRows = corRows;
               simData.brRows = brRows;
               simData.brData = brData.Data;
               simData.brRowInfo = brRowInfo; 
               simData.rRows = rRows;
               simData.rData = rData.Data;
               simData.rRowInfo = rRowInfo;                
               obj.simData = simData;
           end
      end

      function obj = addTableSimVals(obj)   
          % Here we simply get sim values that are
          % fixed during optimization and add them to the 
          % tables.  This is done at initialization to speed
          % execution.
          %
          % ARGUMENTS
          %  (self):      Note that the following properties should be
          %               initialized (experimental and simulation data) 
          %               before calling this method:
          %                mnSDTable
          %                binTable
          %                distTable
          %                distTable2D
          %                simData
          %                corTable	
          %                subpopTable
          %
          % RETURNS
          %  (self):      The VPop object is returned, but with updated
          %               properties:
          %                mnSDTable
          %                binTable
          %                distTable
          %                distTable2D
          %                simData
          %                corTable	
          %                subpopTable
          

         
		  mySubpopTable = obj.subpopTable;		
          mySimData = obj.simData.Data;
          mySimRowInfo = obj.simData.rowInfo;
          mySimColNames = obj.simData.rowInfoNames;
          simInterventionIDCol = find(ismember(mySimColNames, 'interventionID'));
          simTimeCol = find(ismember(mySimColNames, 'time'));
          simElementIDCol = find(ismember(mySimColNames, 'elementID'));
          simElementTypeCol = find(ismember(mySimColNames, 'elementType'));                        
		  

          myTable = obj.mnSDTable;
          rowsTarget = obj.simData.mnSDRows;
          rowsSource = find(rowsTarget>0); 	
          if ~isempty(rowsSource)
			   mySubpopNo = myTable.('subpopNo');          
               vpIndicesSubpop = mySubpopTable.('vpIndices');
               rowsTarget = rowsTarget(rowsSource);
              % 2 step assignment to speed execution
              % first to matrix, then convert back to table.
              [nRows, ~] = size(myTable);          
              curSimValues = nan(nRows,  size(mySimData,2));
              curSimValues(rowsTarget,:) = (mySimData(rowsSource, :));
              for rowCounter = 1 : nRows
                  subpopIndices = vpIndicesSubpop{mySubpopNo(rowCounter)};		
                  rowSimValues = curSimValues(rowCounter,:);
                  keepIndices = intersect(find(~isnan(rowSimValues)),subpopIndices,'stable');
                  [curVals, sortIndices] = sort(rowSimValues, 'ascend');
                  keepIndices = intersect(sortIndices,keepIndices,'stable');
                  %curVals = curSimValues(rowCounter,keepIndices);                 
                  myTable.('predSample'){rowCounter} = rowSimValues(keepIndices);
				  myTable.('predIndices'){rowCounter} = keepIndices;
              end
              obj.mnSDTable = myTable;												
          end  

          myTable = obj.binTable;
          rowsTarget = obj.simData.binRows;
          rowsSource = find(rowsTarget>0); 	
          if ~isempty(rowsSource)
			   mySubpopNo = myTable.('subpopNo');          
               vpIndicesSubpop = mySubpopTable.('vpIndices');
               rowsTarget = rowsTarget(rowsSource);
              % 2 step assignment to speed execution
              % first to matrix, then convert back to table.
              [nRows, ~] = size(myTable);          
              curSimValues = nan(nRows,  size(mySimData,2));
              curSimValues(rowsTarget,:) = (mySimData(rowsSource, :));
              for rowCounter = 1 : nRows
                  subpopIndices = vpIndicesSubpop{mySubpopNo(rowCounter)};		
                  rowSimValues = curSimValues(rowCounter,:);
                  keepIndices = intersect(find(~isnan(rowSimValues)),subpopIndices,'stable');
                  [curVals, sortIndices] = sort(rowSimValues, 'ascend');
                  keepIndices = intersect(sortIndices,keepIndices,'stable');
                  %curVals = curSimValues(rowCounter,keepIndices);                 
                  myTable.('predSample'){rowCounter} = rowSimValues(keepIndices);
				  myTable.('predIndices'){rowCounter} = keepIndices;
              end
              obj.binTable = myTable;												
          end  

          myTable = obj.distTable;
          rowsTarget = obj.simData.distRows;
          rowsSource = find(rowsTarget>0); 	
          if ~isempty(rowsSource)
			   mySubpopNo = myTable.('subpopNo');          
               vpIndicesSubpop = mySubpopTable.('vpIndices');
               rowsTarget = rowsTarget(rowsSource);
          
              % 2 step assignment to speed execution
              % first to matrix, then convert back to table.
              [nRows, nDistCols] = size(myTable);          
				
              curSimValues = nan(nRows,  size(mySimData,2));
              curSimValues(rowsTarget,:) = (mySimData(rowsSource, :));
              curSimValues = nan(nRows,  size(mySimData,2));
              curSimValues(rowsTarget,:) = (mySimData(rowsSource, :));              
              for rowCounter = 1 : nRows


				  
                  subpopIndices = vpIndicesSubpop{mySubpopNo(rowCounter)};		
                  rowSimValues = curSimValues(rowCounter,:);
                  keepIndices = intersect(find(~isnan(rowSimValues)),subpopIndices,'stable');
                  [curVals, sortIndices] = sort(rowSimValues, 'ascend');
                  keepIndices = intersect(sortIndices,keepIndices,'stable');
                  curVals = rowSimValues(keepIndices);                 
                  myTable.('predSample'){rowCounter} = curVals;
				  myTable.('predIndices'){rowCounter} = keepIndices;				  
				  
                  % Also get the indices to align the exp and sim samples
                  sample1 = myTable.('expSample'){rowCounter};
                  if length(sample1 > 0) && length(curVals > 0)
                      [sample1Ind, sample2Ind, SC] = alignSamples(sample1, curVals);
                      myTable.('expCombinedIndices'){rowCounter} = sample1Ind;
                      myTable.('simCombinedIndices'){rowCounter} = sample2Ind;
                      myTable.('combinedPoints'){rowCounter} = SC;
                  else
                      myTable.('expCombinedIndices'){rowCounter} = [];
                      myTable.('simCombinedIndices'){rowCounter} = [];
                      myTable.('combinedPoints'){rowCounter} = [];
                  end

              end
              obj.distTable = myTable;												
          end   

          myTable = obj.distTable2D;
          rowsTarget1 = obj.simData.distRows2D(:,1);
		  rowsTarget2 = obj.simData.distRows2D(:,2);
          rowsSource1 = find(~cellfun(@isempty,rowsTarget1));
          rowsSource2 = find(~cellfun(@isempty,rowsTarget2));
          if ~isempty(rowsSource1)
              mySubpopNo = myTable.('subpopNo');
              vpIndicesSubpop = mySubpopTable.('vpIndices');
              rowsTarget1 = rowsTarget1(rowsSource1);
			  rowsTarget2 = rowsTarget2(rowsSource2);
              % 2 step assignment to speed execution
              % first to matrix, then convert back to table.
              [nDistRows, nDistCols] = size(myTable);
              curSimValues1 = nan(nDistRows,  size(mySimData,2));
              curSimValues2 = nan(nDistRows,  size(mySimData,2));
              % We need a loop for the assignment
              for target1counter = 1 :length(rowsTarget1)
                  targetRows = rowsTarget1{target1counter};
                  for target1repCounter = 1 :length(targetRows)
                    curSimValues1(targetRows(target1repCounter),:) = (mySimData(rowsSource1(target1counter), :));
                  end
              end
              for target2counter = 1 :length(rowsTarget2)
                  targetRows = rowsTarget2{target2counter};
                  for target2repCounter = 1 :length(targetRows)
                    curSimValues2(targetRows(target2repCounter),:) = (mySimData(rowsSource2(target2counter), :));
                  end                  
              end              		  
			  % Unlike the 1D case we won't sort
              for rowCounter = 1 : nDistRows
                  keepIndices = find(~isnan(curSimValues1(rowCounter,:)) & ~isnan(curSimValues2(rowCounter,:)));
				  subpopIndices = vpIndicesSubpop{mySubpopNo(rowCounter)};			
                  % Should account for subpops
                  keepIndices = intersect(keepIndices,subpopIndices);                  
                  
                  curVals = [curSimValues1(rowCounter,keepIndices); curSimValues2(rowCounter,keepIndices)];          
                  myTable.('predSample'){rowCounter} = curVals;
				  myTable.('predIndices'){rowCounter} = keepIndices;
              end
              obj.distTable2D = myTable;												
          end               

          myTable = obj.corTable;
          rowsTarget1 = obj.simData.corRows(:,1);
		  rowsTarget2 = obj.simData.corRows(:,2);
          rowsSource1 = find(~cellfun(@isempty,rowsTarget1));
          rowsSource2 = find(~cellfun(@isempty,rowsTarget2));	
          if ~isempty(rowsSource1)
              mySubpopNo = myTable.('subpopNo');
              vpIndicesSubpop = mySubpopTable.('vpIndices'); 
              rowsTarget1 = rowsTarget1(rowsSource1);
			  rowsTarget2 = rowsTarget2(rowsSource2);
              % 2 step assignment to speed execution
              % first to matrix, then convert back to table.
              [nCorRows, nCorCols] = size(myTable);
              curSimValues1 = nan(nCorRows,  size(mySimData,2));
              curSimValues2 = nan(nCorRows,  size(mySimData,2));
              % We need a loop for the assignment
              for target1counter = 1 :length(rowsTarget1)
                  targetRows = rowsTarget1{target1counter};
                  for target1repCounter = 1 :length(targetRows)
                    curSimValues1(targetRows(target1repCounter),:) = (mySimData(rowsSource1(target1counter), :));
                  end
              end
              for target2counter = 1 :length(rowsTarget2)
                  targetRows = rowsTarget2{target2counter};
                  for target2repCounter = 1 :length(targetRows)
                    curSimValues2(targetRows(target2repCounter),:) = (mySimData(rowsSource2(target2counter), :));
                  end                  
              end              	
			  % Unlike the 1D case we won't sort			  
              for rowCounter = 1 : nCorRows
                  keepIndices = find(~isnan(curSimValues1(rowCounter,:)) & ~isnan(curSimValues2(rowCounter,:)));
				  subpopIndices = vpIndicesSubpop{mySubpopNo(rowCounter)};			
                  % Should account for subpops
                  keepIndices = intersect(keepIndices,subpopIndices);                  
                  
                  curVals = [curSimValues1(rowCounter,keepIndices); curSimValues2(rowCounter,keepIndices)];          
                  myTable.('predSample'){rowCounter} = curVals;
				  myTable.('predIndices'){rowCounter} = keepIndices;

              end
              obj.corTable = myTable;												
          end             

      end

      
      function obj = addPredTableVals(obj)
          % Once we have read the simData and generated a pw vector,
          % We need to add the predicted equivalents to the 
          % tables used for stat calculations.
          %
          % ARGUMENTS
          %  (self):      Note that the following properties should be
          %               initialized (experimental and simulation data) 
          %               before calling this method:
          %                mnSDTable
          %                binTable
          %                distTable
          %                distTable2D
          %                corTable		  
		  %				   brTableRECIST
		  %				   rTableRECIST	
		  %                subpopTable
          %                simData
          %
          % RETURNS
          %  (self):      The VPop object is returned, but with updated
          %               properties:
          %                mnSDTable
          %                binTable
          %                distTable
          %                distTable2D
          %                corTable	
		  %				   brTableRECIST	 % add 8 columns:expCRPD2,expPRPD2, ...	  
		  %				   rTableRECIST
		  %                subpopTable			  
          
          myMnSdData = obj.mnSDTable;
          myBinTable = obj.binTable;
          myDistTable = obj.distTable;          		  
          myDistTable2D = obj.distTable2D;
		  myCorTable = obj.corTable;
          myBRTableRECIST = obj.brTableRECIST;
          myRTableRECIST = obj.rTableRECIST;
		  mySubpopTable = obj.subpopTable;
		  
		  vpIndicesSubpop = mySubpopTable.('vpIndices');
          
          mySimData = obj.simData.Data;
          myBRData = obj.simData.brData;
          myRData = obj.simData.rData;
          
          mySimRowInfo = obj.simData.rowInfo;
          myBRRowInfo = obj.simData.brRowInfo;
          myBRowInfo = obj.simData.rRowInfo;
          
          
          myPWs = obj.pws;
          mySimColNames = obj.simData.rowInfoNames;
          mnSDRowsTarget = obj.simData.mnSDRows;
          mnSDRowsSource = find(mnSDRowsTarget>0);												
          binRowsTarget = obj.simData.binRows;
          binRowsSource = find(binRowsTarget>0);
          distRowsTarget = obj.simData.distRows;
          distRowsSource = find(distRowsTarget>0); 																															 															 
          
          brRowsTarget = obj.simData.brRows;
          brRowsSource = find(brRowsTarget>0);  
          rRowsTarget = obj.simData.rRows;
          rRowsSource = find(rRowsTarget>0);            
          
          simInterventionIDCol = find(ismember(mySimColNames, 'interventionID'));
          simTimeCol = find(ismember(mySimColNames, 'time'));
          simElementIDCol = find(ismember(mySimColNames, 'elementID'));
          simElementTypeCol = find(ismember(mySimColNames, 'elementType'));                    
          brInterventionIDCol = find(ismember(mySimColNames, 'interventionID'));
          brTimeCol = find(ismember(mySimColNames, 'time'));   
          rInterventionIDCol = find(ismember(mySimColNames, 'interventionID'));
          rTimeCol = find(ismember(mySimColNames, 'time'));          

          if ~isempty(myMnSdData) % (mnSDRowsSource)
              mnSDRowsTarget = mnSDRowsTarget(mnSDRowsSource);
          
              mnSDRows = obj.simData.mnSDRows;
              mnSDRows = mnSDRows(find(mnSDRows>0));
			  mySubpopNo = myMnSdData.('subpopNo');

              [nMnSdRows, nMnSdCols] = size(myMnSdData);          

              % 2 step assignment to speed execution
              % first to matrix, then convert back to table.
              curMean = nan(nMnSdRows,1);
              curPredN = nan(nMnSdRows,1);
              curSD = nan(nMnSdRows,1);						 
              curSimValues = nan(nMnSdRows,length(myPWs));
              curSimValues(mnSDRowsTarget,:) = (mySimData(mnSDRowsSource, :));
              keepIndices = myMnSdData.('predIndices');
              for rowCounter = 1 : nMnSdRows
                  if length(keepIndices{rowCounter}) > 1
                      curPWs = myPWs(keepIndices{rowCounter}) / sum(myPWs(keepIndices{rowCounter}));
                      if obj.useEffN
                          curN = 1/sum(curPWs.^2);
                      else
                          % We could use the PW cutoff here, but it seems this
                          % encourages the optimizer to try to push the weight onto a
                          % few VPs to decrease N.  Instead, let's use the number of
                          % VPs for the purpose of statistical comparison, especially
                          % during optimization.
                          % curN = sum(myPWs >= obj.pwCutoff);
                          curN = length(myPWs);
                      end
                      curPredN(rowCounter) = curN;
                      curMean(rowCounter) = wtdMean(curSimValues(rowCounter,keepIndices{rowCounter}),curPWs);
                      curSD(rowCounter) = wtdStd(curSimValues(rowCounter,keepIndices{rowCounter}),curPWs);
                  end
              end

              myMnSdData.('predN') = (curPredN);
              myMnSdData.('predMean') = (curMean);
              myMnSdData.('predSD') = (curSD);     
              obj.mnSDTable = myMnSdData;
          end
		  
          if ~isempty(myBinTable) %Lu changed from (binRowsSource)
              binRowsTarget = binRowsTarget(binRowsSource);
          
              % 2 step assignment to speed execution
              % first to matrix, then convert back to table.
              [nBinRows, nBinCols] = size(myBinTable);          
              curProbs = cell(nBinRows,1);     
              curPredN = nan(nBinRows,1);
              curSimValues = nan(nBinRows,length(myPWs));
              curSimValues(binRowsTarget,:) = (mySimData(binRowsSource, :));
              binEdgeValues = myBinTable{:,'binEdges'};
			  
			  mySubpopNo = myBinTable.('subpopNo');
              keepIndices = myBinTable.('predIndices');
              for rowCounter = 1 : nBinRows
                   curPWs = myPWs(keepIndices{rowCounter}) / sum(myPWs(keepIndices{rowCounter}));
                   if obj.useEffN
                       curN = 1/sum(curPWs.^2);
                   else
                       % We could use the PW cutoff here, but it seems this
                       % encourages the optimizer to try to push the weight onto a
                       % few VPs to decrease N.  Also allow the number of
                       % VPs for the purpose of statistical comparison, especially
                       % during optimization.
                       % curN = sum(myPWs >= obj.pwCutoff);
                       curN = length(myPWs);
                   end
                   curPredN(rowCounter) = curN;
                   curProbs{rowCounter} = wtdBinProb(curSimValues(rowCounter,keepIndices{rowCounter}), curPWs, binEdgeValues{rowCounter});
              end
              myBinTable.('predN') = (curPredN);
              myBinTable.('predBins') = curProbs;     
              obj.binTable = myBinTable;
          end
		  
         if ~isempty(myDistTable)
			  [nDistRows, nDistCols] = size(myDistTable); 
			  assignPWs = cell(nDistRows,1);
              assignN = nan(nDistRows,1);
			  keepIndices = myDistTable.('predIndices');
              for rowCounter = 1 : nDistRows
                   % We have already found these
                   curPWs = myPWs(keepIndices{rowCounter}) / sum(myPWs(keepIndices{rowCounter})); 
                   if obj.useEffN
                       curN = 1/sum(curPWs.^2);
                   else
                       % We could use the PW cutoff here, but it seems this
                       % encourages the optimizer to try to push the weight onto a
                       % few VPs to decrease N.  Also allow the number of
                       % VPs for the purpose of statistical comparison, especially
                       % during optimization.
                       % curN = sum(myPWs >= obj.pwCutoff);
                       curN = length(myPWs);
                   end

                  % Since we assign in multiple values per row for the
                  % distribution, it looks like we have to loop this
                  % All variables in the column must be same size
                  assignN(rowCounter) = curN;
				  assignPWs{rowCounter} = curPWs;
              end
              myDistTable.('predN') = (assignN);
			  myDistTable.('predProbs') = (assignPWs);
              obj.distTable = myDistTable;														
         end
		 
         if ~isempty(myDistTable2D)
			  distRowsTarget1 = obj.simData.distRows2D(:,1);
			  distRowsTarget2 = obj.simData.distRows2D(:,2);
			  distRowsSource1 = find(~cellfun(@isempty,distRowsTarget1));
			  distRowsSource2 = find(~cellfun(@isempty,distRowsTarget2));		 
          
			  [nDistRows, nDistCols] = size(myDistTable2D); 
			  assignPWs = cell(nDistRows,1);
              assignN = nan(nDistRows,1);
			  keepIndices = myDistTable2D.('predIndices');
              for rowCounter = 1 : nDistRows
                   % We have already found these
                   curPWs = myPWs(keepIndices{rowCounter}) / sum(myPWs(keepIndices{rowCounter})); 
                   if obj.useEffN
                       curN = 1/sum(curPWs.^2);
                   else
                       % We could use the PW cutoff here, but it seems this
                       % encourages the optimizer to try to push the weight onto a
                       % few VPs to decrease N.  Also allow the number of
                       % VPs for the purpose of statistical comparison, especially
                       % during optimization.
                       % curN = sum(myPWs >= obj.pwCutoff);
                       curN = length(myPWs);
                   end

                  % Since we assign in multiple values per row for the
                  % distribution, it looks like we have to loop this
                  % All variables in the column must be same size
                  assignN(rowCounter) = curN;
				  assignPWs{rowCounter} = curPWs;
              end
              myDistTable2D.('predN') = (assignN);
			  myDistTable2D.('predProbs') = (assignPWs);
              obj.distTable2D = myDistTable2D;														
         end 		
		 
         if ~isempty(myCorTable)
			  corRowsTarget1 = obj.simData.corRows(:,1);
			  corRowsTarget2 = obj.simData.corRows(:,2);
			  corRowsSource1 = find(~cellfun(@isempty,corRowsTarget1));
			  corRowsSource2 = find(~cellfun(@isempty,corRowsTarget2));		 
			  [nCorRows, nCorCols] = size(myCorTable); 
			  assignPWs = cell(nCorRows,1);
              assignN = nan(nCorRows,1);
			  curCor = nan(nCorRows,1);
			  keepIndices = myCorTable.('predIndices');
              for rowCounter = 1 : nCorRows
                   % We have already found these
                   curPWs = myPWs(keepIndices{rowCounter}) / sum(myPWs(keepIndices{rowCounter}));  
                   if obj.useEffN
                       curN = 1/sum(curPWs.^2);
                   else
                       % We could use the PW cutoff here, but it seems this
                       % encourages the optimizer to try to push the weight onto a
                       % few VPs to decrease N.  Also allow the number of
                       % VPs for the purpose of statistical comparison, especially
                       % during optimization.
                       % curN = sum(myPWs >= obj.pwCutoff);
                       curN = length(myPWs);
                   end

                  % Since we assign in multiple values per row for the
                  % distribution, it looks like we have to loop this
                  % All variables in the column must be same size
                  assignN(rowCounter) = curN;
				  assignPWs{rowCounter} = curPWs;
                  curwtdcorr = weightedcorrs(myCorTable.('predSample'){rowCounter}', curPWs');
                  curCor(rowCounter) = curwtdcorr(1,2);	  
              end
              myCorTable.('predN') = (assignN);
			  myCorTable.('predProbs') = (assignPWs);
			  myCorTable.('predCor') = (curCor);
              obj.corTable = myCorTable;								
          end 			  
		  
          if ~isempty(myBRTableRECIST)
              brRowsTarget = brRowsTarget(brRowsSource);      
              [nBRRows, nBinCols] = size(myBRTableRECIST);          
              curProbs = nan(nBRRows,4);     
              curPredN = nan(nBRRows,1);
              curSimValues = nan(nBRRows,length(myPWs));
              curSimValues(brRowsTarget,:) = (myBRData(brRowsSource, :));
              binEdgeValues = [.9,1.9,2.9];     
			  
			  mySubpopNo = myBRTableRECIST.('subpopNo');

              for rowCounter = 1 : nBRRows
				   curIndices = vpIndicesSubpop{mySubpopNo(rowCounter)};
                   % 'binEdge1','binEdge2','binEdge3'
				   curPWs = myPWs(curIndices);
				   curPWs = curPWs./sum(curPWs);
				   if obj.useEffN
						curPredN(rowCounter) = 1/sum(curPWs.^2);
				   else
						curPredN(rowCounter) = length(myPWs);
                   end
                   if ~isempty(curPWs) % catch edge cases when some rows contain no VPs
                        curProbs(rowCounter,:) = wtdBinProb(curSimValues(rowCounter,curIndices), curPWs, binEdgeValues);
                   end
              end
              myBRTableRECIST.('predN') = (curPredN);
              myBRTableRECIST.('predCR') = (curProbs(:,1));
              myBRTableRECIST.('predPR') = (curProbs(:,2));
              myBRTableRECIST.('predSD') = (curProbs(:,3));
              myBRTableRECIST.('predPD') = (curProbs(:,4));
              obj.brTableRECIST = myBRTableRECIST;
          end          

          if ~isempty(myRTableRECIST) %(rRowsSource)
              rRowsTarget = rRowsTarget(rRowsSource);      
              [nRRows, nBinCols] = size(myRTableRECIST);          
              curProbs = nan(nRRows,4);     
              curPredN = nan(nRRows,1);
              curSimValues = nan(nRRows,length(myPWs));
              curSimValues(rRowsTarget,:) = (myRData(rRowsSource, :));
              binEdgeValues = [.9,1.9,2.9];   

			  mySubpopNo = myRTableRECIST.('subpopNo');
                            
              for rowCounter = 1 : nRRows
				   curIndices = vpIndicesSubpop{mySubpopNo(rowCounter)};			  
                   % 'binEdge1','binEdge2','binEdge3'
				   curPWs = myPWs(curIndices);
				   curPWs = curPWs./sum(curPWs);
				   if obj.useEffN
						curPredN(rowCounter) = 1/sum(curPWs.^2);
				   else
						curPredN(rowCounter) = length(myPWs);
                   end
                   if ~isempty(curPWs) % catch edge cases when some rows contain no VPs
                        curProbs(rowCounter,:) = wtdBinProb(curSimValues(rowCounter,curIndices), curPWs, binEdgeValues);
                   end
              end
              myRTableRECIST.('predN') = (curPredN);
              myRTableRECIST.('predCR') = (curProbs(:,1));
              myRTableRECIST.('predPR') = (curProbs(:,2));
              myRTableRECIST.('predSD') = (curProbs(:,3));
              myRTableRECIST.('predPD') = (curProbs(:,4));     
              obj.rTableRECIST = myRTableRECIST;
          end              
		  
          [nSubpopRows, nSubpopCols] = size(mySubpopTable);		
		  if nSubpopRows > 1
            curProbs = ones(nSubpopRows,1);
			for rowCounter = 2 : nSubpopRows
				curIndices = vpIndicesSubpop{rowCounter};
				curPWs = myPWs(curIndices);
				curProbs(rowCounter) = sum(curPWs);
			end
            mySubpopTable.('predW') = (curProbs);
			obj.subpopTable = mySubpopTable;
		  end
      end

      function obj = VPopRECIST()
          % This is the constructor method for an instance of a VPop
          % (virtual population) object.
          obj.coeffsTable=[];
		  obj.coeffsDist=[];
		  obj.pwStrategy = 'direct';
          obj.indexTable = [];
          obj.binEdges = [];
          obj.binMidPoints = [];
          obj.binProbs = [];
          % obj.vpIDs = cell(1,0);		  
          obj.pws = [];
          % obj.subpopTable = [];		  
          obj.mnSDTable = [];
          obj.binTable = [];
          obj.distTable = [];         
          obj.distTable2D = [];	
		  obj.corTable = [];
          obj.brTableRECIST = [];
          obj.rTableRECIST = [];   
          obj.subpopTable = [];
          obj.expData = [];          
          obj.simData = [];
          obj.gofMn = [];
          obj.gofSD = [];
          obj.gofBin = [];
          obj.gofDist = [];          		  
		  obj.gofDist2D = [];  
		  obj.gofCor = [];
          obj.gofBR = [];  
          obj.gofR = [];                              
          obj.gof = [];          
          obj.spreadOut = 0; 
		  obj.minIndPVal = 0;	
          obj.useEffN = false;
          obj.exactFlag = true; 		  
          obj.optimizeTimeLimit = 10*60;
          obj.optimizeType = 'pso';    
          obj.optimizePopSize = 1000;  
		  obj.objectiveLimit = -Inf;
          obj.poolClose = true;  
		  obj.poolRestart = true;		  
          obj.intSeed = -1;
          obj.tol = 1E-3;
          obj.nIters = 10000;
          obj.minEffN = 0;
          obj.relSLDvar = [];
          obj.absALDVar = [];
          obj.crCutoff = nan;            
          obj.recistSimFilter = [];
          obj.MSE = [];
          obj.LinearProblemMatrices = [];
          obj.LinearProblemMatricesSubgroupSumWeights = [];
          obj.LinearProblemMatricesobservationDescriptions = [];
          obj.lambda = [];
      end

end

end
```
