## Supplementary File 3 for "A surrogate workflow for virtual population development: designing an amenable proxy for a hypothesis-test inspired nonlinear goodness-of-fit": createVPRangeTable.m

```
function vpRangeTable = createVPRangeTable(arg1, arg2)
% This function checks the data in the tables to be calibrated,
% finds their simulation data, and, if the experiemental source data exists
% it then checks the range in the simulated data to see if it covered the
% range in the experimental data.
%
% ARGUMENTS
%  arg1:            A virtual population OR a worksheet.
%  arg2:            (optional) If a worksheet is provided as arg1,
%                     a mapelOptions must be given.  This option is given
%                     so we can check the worksheet as soon as we have
%                     simulation results and the population targets set up.
% 
% RETURNS
% vpRangeTable:      A table with the comparison of simulated and observed
%                    ranges, with information on "edge" VPs that may
%                    be of interest.
%

% First check input arguments
if nargin > 2
    warning(['Too many input arguments to',mfilename,'. Requires: myVPop OR myWorksheet, myMapelOptions.'])
    flagContinue = false;
elseif nargin > 1
    myWorksheet = arg1;
    myMapelOptions = arg2;
    myVPop = [];
    flagContinue = true;     
elseif nargin > 0
    myWorksheet = [];
    myMapelOptions = [];
    myVPop = arg1;    
    flagContinue = true;     
else
    warning(['Insufficient input arguments to ',mfilename,'. Requires: myVPop.'])
    flagContinue = false;    
end

vpRangeTable = [];

if (isempty(myVPop) && flagContinue)
    % First copy all of the needed properties over
    myVPop = initializeOptionPropertiesToVPop(myMapelOptions);
    if isa(myVPop,'VPopRECIST')
        myVPop.recistSimFilter = createRECISTSimFilter(myWorksheet, myVPop);
    end
    myVPop = myVPop.getSimData(myWorksheet);
    myVPop.subpopTable = updateSubpopTableVPs(myVPop.subpopTable, myWorksheet);
end

if flagContinue
    myCheckElement = cell(0,6);
    if ~isempty(myVPop.mnSDTable)
        % We will only consider mn/sd data for ranges if the variance is
        % included, we will not try to match if we are only calibrating
        % the mean
        keepRows = find(myVPop.mnSDTable{:, 'weightSD'} > 0);
        if length(keepRows) > 0
            myVPop.mnSDTable(:, 'weight') = table(max(myVPop.mnSDTable{:, 'weightMean'},myVPop.mnSDTable{:, 'weightSD'}));
            myCheckElement = [myCheckElement; table2cell(myVPop.mnSDTable(keepRows,{'elementID','elementType','interventionID','time','weight','subpopNo'}))];
        end
    end
    if ~isempty(myVPop.distTable)
        keepRows = find(myVPop.distTable{:, 'weight'} > 0);
        if length(keepRows) > 0        
            myCheckElement = [myCheckElement; table2cell(myVPop.distTable(keepRows,{'elementID','elementType','interventionID','time','weight','subpopNo'}))];
        end
    end
    if ~isempty(myVPop.binTable)
        keepRows = find(myVPop.binTable{:, 'weight'} > 0);
        if length(keepRows) > 0           
            myCheckElement = [myCheckElement; table2cell(myVPop.binTable(keepRows,{'elementID','elementType','interventionID','time','weight','subpopNo'}))];
        end
    end    
    if ~isempty(myVPop.distTable2D)
        keepRows = find(myVPop.distTable2D{:, 'weight'} > 0);
        if length(keepRows) > 0            
            myCheckElement = [myCheckElement; table2cell(myVPop.distTable2D(keepRows,{'elementID1','elementType1','interventionID1','time1','weight','subpopNo'}))];
            myCheckElement = [myCheckElement; table2cell(myVPop.distTable2D(keepRows,{'elementID2','elementType2','interventionID2','time2','weight','subpopNo'}))];
        end
    end  
    if ~isempty(myVPop.corTable)
        keepRows = find(myVPop.corTable{:, 'weight'} > 0);
        if length(keepRows) > 0         
            myCheckElement = [myCheckElement; table2cell(myVPop.corTable(keepRows,{'elementID1','elementType1','interventionID1','time1','weight','subpopNo'}))];
            myCheckElement = [myCheckElement; table2cell(myVPop.corTable(keepRows,{'elementID2','elementType2','interventionID2','time2','weight','subpopNo'}))];
        end
    end    
    [~,idx]=unique(cell2table(myCheckElement(:,[1:4,6])),'rows'); % not include weight to the unique row selection
    myCheckElement = myCheckElement(idx,:);
    myCheckElement = myCheckElement(cell2mat(myCheckElement(:,5))>0,:);
    myCheckElement = myCheckElement(:,[1:4,6]);
    [nCheckElements, ~] = size(myCheckElement);

     % Find column in ExpDataTable where data starts
     commonNames = loadCommonNames();
     myExpDataTable = myVPop.expData;
     if isa(myVPop, 'VPopRECIST')
         firstDataIndex = find(ismember(myExpDataTable.Properties.VariableNames, commonNames.VPOPRECISTTABLEVARNAMESFIXED{end}))+1;
     else
         firstDataIndex = find(ismember(myExpDataTable.Properties.VariableNames, commonNames.VPOPTABLEVARNAMESFIXED{end}))+1;
     end
    
     expDataTableRows = size(myExpDataTable,1); % number of experimental data
     mySimData = myVPop.simData.Data;
     myVPIDs = myVPop.simData.vpIDs;
    
    % define pointers into columns of rowInfoTable
    simTimeCol = find(ismember(myVPop.simData.rowInfoNames,'time'));
    simInterventionIDCol = find(ismember(myVPop.simData.rowInfoNames,'interventionID'));
    simElementIDCol = find(ismember(myVPop.simData.rowInfoNames,'elementID'));
    simExpVarIDCol = find(ismember(myVPop.simData.rowInfoNames,'expVarID'));
    
    %% TODO: later might want to change myVPop.simData.rowInfo, add subpopNo in. for now use the expRowFinder's
    rowInfoTable = cell2table(myVPop.simData.rowInfo,'VariableNames',myVPop.simData.rowInfoNames);
    expRowFinder = [myExpDataTable{:,'elementID'},myExpDataTable{:,'elementType'},myExpDataTable{:,'interventionID'},num2cell(myExpDataTable{:,'time'}),num2cell(myExpDataTable{:,'subpopNo'})];
    simRowFinder = [rowInfoTable{:,'elementID'},rowInfoTable{:,'elementType'},rowInfoTable{:,'interventionID'},num2cell(rowInfoTable{:,'time'})];
    
     if nCheckElements > 0

         nFoundElements = 0;
         expMin = nan(nCheckElements,1);
         expMax = nan(nCheckElements,1);
         % In the future, we might also want to
         % add checks on the 10/90 percentiles
         % in the data.
         exp10 = nan(nCheckElements,1);
         exp90 = nan(nCheckElements,1);         
         simMin = nan(nCheckElements,1);
         simMax = nan(nCheckElements,1);
         expN = nan(nCheckElements,1);
         vpMinInd = cell(nCheckElements,1);
         vpMaxInd = cell(nCheckElements,1);
         vpIDsMin = cell(nCheckElements,1);
         vpIDsMax = cell(nCheckElements,1);
         rowInfo = cell(nCheckElements,5);
         for checkCounter = 1 : nCheckElements
             curCheckElements = myCheckElement(checkCounter, :);
             
             match  = strcmp(expRowFinder(:, 1), curCheckElements{1}) & ...
                strcmp(expRowFinder(:, 2), curCheckElements{2}) & ...
                strcmp(expRowFinder(:, 3), curCheckElements{3}) & ...
                cat(1, expRowFinder{:, 4}) == curCheckElements{4} & ...
                cat(1, expRowFinder{:, 5}) == curCheckElements{5};
             expRow  = find(match);
             
             match  = strcmp(simRowFinder(:, 1), curCheckElements{1}) & ...
                strcmp(simRowFinder(:, 2), curCheckElements{2}) & ...
                strcmp(simRowFinder(:, 3), curCheckElements{3}) & ...
                cat(1, simRowFinder{:, 4}) == curCheckElements{4};
             simRow  = find(match);             
             
             % expRow = find(ismember(expRowFinder,curCheckElements,'rows'));
             % simRow = find(ismember(expRowFinder,curCheckElements,'rows'));
             
             if (length(expRow) == 1) && (length(simRow) == 1)
                nFoundElements = nFoundElements + 1;
                expN(nFoundElements) = sum(~isnan(myExpDataTable{expRow,firstDataIndex:end}));
                expMin(nFoundElements) = min(myExpDataTable{expRow,firstDataIndex:end});
                expMax(nFoundElements) = max(myExpDataTable{expRow,firstDataIndex:end});
                temp = prctile(myExpDataTable{expRow,firstDataIndex:end},[10 90]);
                exp10(nFoundElements) = temp(1);
                exp90(nFoundElements) = temp(2);
                subpopIdx = curCheckElements{5};
                vpIndices = myVPop.subpopTable.vpIndices{subpopIdx};
                simMin(nFoundElements) = min((mySimData(simRow,vpIndices)));
                simMax(nFoundElements) = max((mySimData(simRow,vpIndices)));
                vpMinInd{nFoundElements} = find(mySimData(simRow,:) == simMin(nFoundElements));
                vpMaxInd{nFoundElements} = find(mySimData(simRow,:) == simMax(nFoundElements));
                vpIDsMin{nFoundElements} = myVPIDs(vpMinInd{nFoundElements});
                vpIDsMax{nFoundElements} = myVPIDs(vpMaxInd{nFoundElements}); 
                rowInfo(nFoundElements,:) = [myExpDataTable{expRow,'elementID'},myExpDataTable{expRow,'elementType'},myExpDataTable{expRow,'interventionID'},myExpDataTable{expRow,'time'},myExpDataTable{expRow,'subpopNo'}];
             end
         end
         if nFoundElements > 0
             expN = expN(1:nFoundElements);
             expMin = expMin(1:nFoundElements);
             expMax = expMax(1:nFoundElements);
             exp10 = exp10(1:nFoundElements);
             exp90 = exp90(1:nFoundElements);             
             simMin = simMin(1:nFoundElements);
             simMax = simMax(1:nFoundElements);
             vpMinInd = vpMinInd(1:nFoundElements);
             vpMaxInd = vpMaxInd(1:nFoundElements);
             vpIDsMin = vpIDsMin(1:nFoundElements);   
             vpIDsMax = vpIDsMax(1:nFoundElements); 
             rowInfo = rowInfo(1:nFoundElements,:);
			 
             rangeCover = (min(simMax,expMax) - max(simMin,expMin)) ./ (expMax - expMin);
             minRangeMissing = max((simMin - expMin),zeros(nFoundElements,1))./(expMax - expMin);
             maxRangeMissing = max((expMax - simMax),zeros(nFoundElements,1))./(expMax - expMin);             
             %percentileCover80 = (min(simMax,exp90) - max(simMin,exp10)) ./ (exp90 - exp10);
             % maxMissing = max((expMax - simMax),zeros(nFoundElements,1))./(expMax - expMin);
             min10PercentileMissing = max((simMin - exp10),zeros(nFoundElements,1))./(expMax - exp10);   
             max90PercentileMissing = max((exp90 - simMax),zeros(nFoundElements,1))./(exp90 - expMin);
             elementID = rowInfo(:,1);
             elementType = rowInfo(:,2);
             interventionID = rowInfo(:,3);
             time = rowInfo(:,4);
             subpopNo = rowInfo(:,5);
             
             [~, sortIndices] = sort(rangeCover,'ascend');
             
             vpRangeTable = table(subpopNo,elementID,elementType,interventionID,time,expN,vpIDsMin,vpIDsMax,simMin,simMax,expMin,expMax,rangeCover,minRangeMissing,maxRangeMissing,min10PercentileMissing,max90PercentileMissing);
             vpRangeTable = vpRangeTable(sortIndices,:);
             
         end
     end
else
    warning(['Unable to proceed in ',mfilename,'.'])
end

end
```
