## Supplementary File 3 for "A surrogate workflow for virtual population development: designing an amenable proxy for a hypothesis-test inspired nonlinear goodness-of-fit": convertExpDataTo2DDistTable.m

```
function myDistTable = convertExpDataTo2DDistTable(myVPop,myExpVars1,myExpVars2,mySimTimepoints1,mySimTimepoints2,myInterventions1,myInterventions2,myExpDataIDs1,myExpDataIDs2)
% This function takes experimental data from a newer 2D table and
% converts it to a 2D dist table format for use in MAPEL
%
% ARGUMENTS:
% myVPop:                 A VPop object with a populated expData field.  A
%                          mapelOptions structure is also OK.
% myExpVars1,myExpVars2:  Two paired cell arrays of length N 
%                          (number of 2D distributions to calibrate to)
%                          with the experimental/data variable names
% mySimTimepoints1,mySimTimepoints2:  Two paired cell arrays of length N 
%                                     (number of 2D distributions to calibrate to)
%                                     with the simulation timepoints
% myInterventions1,myInterventions2:  Two paired cell arrays of length N 
%                                     (number of 2D distributions to calibrate to)
%                                      with the interventions
%  myExpDataIDs1,myExpDataIDs2:       Optional: Two paired cell arrays of length N 
%                                      (number of 2D distributions to calibrate to)
%                                      with the ExpDataIDs.
%
%
% RETURNS
% myDistTable
%


Length_2D_calib = size(myExpVars1,2);
% myExpDataIDs1 and myExpDataIDs2 are optional
% If they do not exist, we default them to a value
if ~ismember('myExpDataIDs1',who)
    myExpDataIDs1 = cell(1,Length_2D_calib);
end
if ~ismember('myExpDataIDs2',who)
    myExpDataIDs2 = cell(1,Length_2D_calib);
end


continueFlag = true;

if nargin > 9
    warning(['Too many input arguments for ',mfilename,'. Should provide: myVPop (or mapelOptions),myExpVars1,myExpVars2,mySimTimepoints1,mySimTimepoints2,myInterventions1,myInterventions2 with myExpDataIDs1 and myExpDataIDs2 optional.'])
    continueFlag = false;
elseif nargin < 7
    warning(['Insufficient input arguments for ',mfilename,'. Should provide: myVPop (or mapelOptions),myExpVars1,myExpVars2,mySimTimepoints1,mySimTimepoints2,myInterventions1,myInterventions2 with myExpDataIDs1 and myExpDataIDs2 optional.'])
    continueFlag = false;
else
    continueFlag = true;
end


if continueFlag
    if sum(ismember({'VPop','VPopRECIST','VPopRECISTnoBin','mapelOptions','mapelOptionsRECIST','mapelOptionsRECISTnoBin'},class(myVPop))) < 1
        warning(['Wrong input arguments for ',mfilename,'. Should provide a RECIST type myVPop (or mapelOptions).'])
        continueFlag = false;
    end
end
        
if continueFlag
    if sum(ismember({'table'},class(myVPop.expData))) < 1
        warning(['Wrong input arguments for ',mfilename,'. Should provide: myVPop (or mapelOptions) with a populated expData property.'])
        continueFlag = false;
    end
end

if continueFlag
    commonNames = loadCommonNames();
    nRows = length(myExpVars1);

        
    for rowCounter = 1 : nRows
        myExpVars1Cur = myExpVars1{rowCounter};
        myExpVars2Cur = myExpVars2{rowCounter};
        mySimTimepoints1Cur = mySimTimepoints1(rowCounter);
        mySimTimepoints2Cur = mySimTimepoints2(rowCounter);
        myInterventions1Cur = myInterventions1{rowCounter};
        myInterventions2Cur = myInterventions2{rowCounter};
        myExpDataIDs1Cur = myExpDataIDs1{rowCounter};
        myExpDataIDs2Cur = myExpDataIDs2{rowCounter};

        if rowCounter == 1
            % The new headers will be a little different since we are
            % 2 measures on each row
            % This is not yest supported as we will have yet to add
            % PatientIDVar for non-RECIST
            if isa(myVPop,'VPop') || isa(myVPop,'mapelOptions')
                tableVariableNames = commonNames.VPOP2DTABLEVARNAMESFIXED;
                nDataHeaderCols = length(tableVariableNames);
                oldTableVariableNames = commonNames.VPOPEXPTABLEVARNAMESFIXED;
                nOldDataHeaderCols = length(oldTableVariableNames);
            else
                tableVariableNames = commonNames.VPOPRECIST2DTABLEVARNAMESFIXED;
                nDataHeaderCols = length(tableVariableNames);
                oldTableVariableNames = commonNames.VPOPRECISTEXPTABLEVARNAMESFIXED;
                nOldDataHeaderCols = length(oldTableVariableNames);                
            end
            tableVariableNames = [tableVariableNames,{'weight','expN', 'expSample', 'predN', 'predIndices', 'predSample', 'predProbs'}];
            myDistTable = cell2table(cell(0,length(tableVariableNames)));
            myDistTable.Properties.VariableNames = tableVariableNames;
     
        end
        
        if isempty(myExpDataIDs1{1})           
            sourceRow1 = find(ismember(myVPop.expData.('expVarID'),myExpVars1Cur) ...
                            & ismember(myVPop.expData.('time'),mySimTimepoints1Cur) ...
                            & ismember(myVPop.expData.('interventionID'),myInterventions1Cur));
                    
            sourceRow2 = find(ismember(myVPop.expData.('expVarID'),myExpVars2Cur) ... 
                            & ismember(myVPop.expData.('time'),mySimTimepoints2Cur) ...
                            & ismember(myVPop.expData.('interventionID'),myInterventions2Cur));
        else
            
            sourceRow1 = find(ismember(myVPop.expData.('expVarID'),myExpVars1Cur) ...
                            & ismember(myVPop.expData.('time'),mySimTimepoints1Cur) ...
                            & ismember(myVPop.expData.('interventionID'),myInterventions1Cur) ...
                            & ismember(myVPop.expData.('expDataID'),myExpDataIDs1Cur));
                    
            sourceRow2 = find(ismember(myVPop.expData.('expVarID'),myExpVars2Cur) ... 
                            & ismember(myVPop.expData.('time'),mySimTimepoints2Cur) ...
                            & ismember(myVPop.expData.('interventionID'),myInterventions2Cur) ...
                            & ismember(myVPop.expData.('expDataID'),myExpDataIDs2Cur));
        end
                
        curData = myVPop.expData{[sourceRow1,sourceRow2],nOldDataHeaderCols+1:end};
        curData = curData(:,find(min(~isnan(curData),[],1)));
        %[~,idx] = sort(curData(1,:),'ascend');
        %curData=curData(:,idx);
        [~,nEntries] = size(curData);
        if ~isequal(myVPop.expData{sourceRow1,'subpopNo'},myVPop.expData{sourceRow2,'subpopNo'})
            disp(['Attempting to correlate different subpopulations in ',mfilename,' for rows ',num2str(sourceRow1),' and ',num2str(sourceRow2),'.  Assuming the first of the pair.'])
        end
        if isa(myVPop,'VPop') || isa(myVPop,'mapelOptions')
            curRow = {myVPop.expData{sourceRow1,'subpopNo'},myVPop.expData{sourceRow1,'time'},myVPop.expData{sourceRow2,'time'},myVPop.expData{sourceRow1,'interventionID'},myVPop.expData{sourceRow2,'interventionID'},myVPop.expData{sourceRow1,'elementID'},myVPop.expData{sourceRow2,'elementID'},myVPop.expData{sourceRow1,'elementType'},myVPop.expData{sourceRow2,'elementType'},myVPop.expData{sourceRow1,'expDataID'},myVPop.expData{sourceRow2,'expDataID'},myVPop.expData{sourceRow1,'expTimeVarID'},myVPop.expData{sourceRow2,'expTimeVarID'},myVPop.expData{sourceRow1,'expVarID'},myVPop.expData{sourceRow2,'expVarID'},myVPop.expData{sourceRow1,'PatientIDVar'},myVPop.expData{sourceRow2,'PatientIDVar'}};
        else
            curRow = {myVPop.expData{sourceRow1,'subpopNo'},myVPop.expData{sourceRow1,'time'},myVPop.expData{sourceRow2,'time'},myVPop.expData{sourceRow1,'interventionID'},myVPop.expData{sourceRow2,'interventionID'},myVPop.expData{sourceRow1,'elementID'},myVPop.expData{sourceRow2,'elementID'},myVPop.expData{sourceRow1,'elementType'},myVPop.expData{sourceRow2,'elementType'},myVPop.expData{sourceRow1,'expDataID'},myVPop.expData{sourceRow2,'expDataID'},myVPop.expData{sourceRow1,'expTimeVarID'},myVPop.expData{sourceRow2,'expTimeVarID'},myVPop.expData{sourceRow1,'expVarID'},myVPop.expData{sourceRow2,'expVarID'},myVPop.expData{sourceRow1,'PatientIDVar'},myVPop.expData{sourceRow2,'PatientIDVar'},myVPop.expData{sourceRow1,'TRTVar'},myVPop.expData{sourceRow2,'TRTVar'},myVPop.expData{sourceRow1,'BRSCOREVar'},myVPop.expData{sourceRow2,'BRSCOREVar'},myVPop.expData{sourceRow1,'RSCOREVar'},myVPop.expData{sourceRow2,'RSCOREVar'}};
        end 
        curRow = [curRow,{1, nEntries, {curData}, nan, {nan}, {nan}, {nan}}];
        curRow = cell2table(curRow);
        curRow.Properties.VariableNames = myDistTable.Properties.VariableNames; 
        
        myDistTable = [myDistTable; curRow];   
    end
        
else
    warning(['Unable to complete ',mfilename,', exiting.'])
	myDistTable = [];
end            

end
```
