## Supplementary File 3 for "A surrogate workflow for virtual population development: designing an amenable proxy for a hypothesis-test inspired nonlinear goodness-of-fit": convertExpDataToBinTable.m

```
function myBinTable = convertExpDataToBinTable(myVPop, nBins)
% This function takes experimental data and
% converts it to a bin table format for use in MAPEL
%
% ARGUMENTS:
% myVPop:           A VPop object with a populated expData field.  A
%                    mapelOptions structure is also OK.
% nBins:            Number of bins.  If not given, 4 are assumed.
%                    The same number of bins are used for all datasets.
%
% RETURNS
% myBinTable
%

continueFlag = true;
if nargin > 2
    continueFlag = false;
    warning(['Too many input arguments for ',mfilename,'. Should provide: myVPop and optionally nBins.'])
    continueFlag = false;
elseif nargin > 1
    continueFlag = true;	
elseif nargin > 0
	nBins = 4;
    continueFlag = true;
else
    warning(['Insufficient input arguments for ',mfilename,'. Should provide: myVPop.'])
    continueFlag = false;
end

if continueFlag
    if sum(ismember({'VPop','VPopRECIST','VPopRECISTnoBin','mapelOptions','mapelOptionsRECIST','mapelOptionsRECISTnoBin'},class(myVPop))) < 1
        warning(['Wrong input arguments for ',mfilename,'. Should provide: myVPop (or mapelOptions).'])
        continueFlag = false;
    end
end
        
if continueFlag
    if sum(ismember({'table'},class(myVPop.expData))) < 1
        warning(['Wrong input arguments for ',mfilename,'. Should provide: myVPop (or mapelOptions) with a populated expData property.'])
        continueFlag = false;
    end
end

if continueFlag
    commonNames = loadCommonNames();
    [nRows, ~] = size(myVPop.expData);
    if isa(myVPop,'VPop') || isa(myVPop,'mapelOptions')
        tableVariableNames = commonNames.VPOPTABLEVARNAMESFIXED;
		nDataHeaderCols = length(commonNames.VPOPTABLEVARNAMESFIXED);
    else
        tableVariableNames = commonNames.VPOPRECISTTABLEVARNAMESFIXED;
		nDataHeaderCols = length(commonNames.VPOPRECISTTABLEVARNAMESFIXED);
    end
    uniqueExpDataTime = unique(myVPop.expData{:,'time'});
	% Bins will be set to capture
	% a similar number of experimental datapoints
	% in each bin by default		
	binPercentiles = 100/nBins;
	binPercentiles = (1 : 1: (nBins-1))*binPercentiles;	
    for rowCounter = 1 : nRows
        if rowCounter == 1
            %tableVariableNames = [tableVariableNames,{'weight','binEdge1','binEdge2','binEdge3','expN','expBin1','expBin2','expBin3','expBin4','predN','predBin1','predBin2','predBin3','predBin4'}];
			tableVariableNames = [tableVariableNames,{'weight','binEdges','expN','expBins','predN','predIndices','predBins'}];
            myBinTable = cell2table(cell(0,length(tableVariableNames)));
            myBinTable.Properties.VariableNames = tableVariableNames;
        end
        curData = myVPop.expData{rowCounter,nDataHeaderCols+1:end};
        curData = curData(~isnan(curData));
        curRow = table2cell(myVPop.expData(rowCounter,1:nDataHeaderCols));
        myElementID = myVPop.expData{rowCounter,'elementID'};
        allCurVarRows = find(ismember(table2cell(myVPop.expData(:, 'elementID')),myElementID));
        allCurVarValues = myVPop.expData{allCurVarRows, nDataHeaderCols+1:end};
        allCurVarValues = allCurVarValues(~isnan(allCurVarValues));
		myBinEdgeValues = prctile(allCurVarValues, binPercentiles);
        curProbs = wtdBinProb(curData, ones(1, length(curData))/length(curData), myBinEdgeValues);
        expN = length(curData);
        curRow = [curRow,{1,{myBinEdgeValues}, expN, {curProbs}, nan, {nan},{nan(1,nBins)}}];        
        curRow = cell2table(curRow);
        curRow.Properties.VariableNames = myBinTable.Properties.VariableNames; 
        myBinTable = [myBinTable; curRow];   
    end
else
    warning(['Unable to complete ',mfilename,', exiting.'])
	myBinTable = [];
end            

end
```
