## Supplementary File 3 for "A surrogate workflow for virtual population development: designing an amenable proxy for a hypothesis-test inspired nonlinear goodness-of-fit": convertMapelOptionsToExpDataTable.m

```
function myExpDataTable = convertMapelOptionsToExpDataTable(myWorksheet, myMapelOptions)
% This function takes experimental data embedded in a worksheet and
% converts it to a data table format for use in MAPEL, guided by the
% data tables in the mapelOptions. 
%
% ARGUMENTS:
% myWorksheet:           a worksheet containing the data
% myMapelOptions:        a mapelOptions object ('mapelOptions','mapelOptionsRECIST','mapelOptionsRECISTnoBin')
%
% RETURNS
% myExpDataTable

continueFlag = true;
if nargin > 2
    continueFlag = false;
    warning(['Too many input arguments for ',mfilename,'. Should provide: myWorksheet, myMapelOptions.'])
    continueFlag = false;
elseif nargin < 2
    warning(['Insufficient input arguments for ',mfilename,'. Should provide: myWorksheet, myMapelOptions.'])
    continueFlag = false;   
end

if continueFlag
    if sum(ismember(class(myMapelOptions),{'mapelOptions','mapelOptionsRECIST','mapelOptionsRECISTnoBin'}))<0;
        warning(['Should provide: myWorksheet, myMapelOptions for ',mfilename,'.'])
        continueFlag = false;
    elseif sum(ismember(class(myMapelOptions),{'mapelOptions'}))<0
        warning(['The provided myMapelOptions for ',mfilename,' is not RECIST-based.  This function does not yet support 2D distributions with non-RECIST workflows.'])
    end
end

if continueFlag
    if isa(myMapelOptions, 'mapelOptions')
		lookupRows  = cell2table(cell(0,7), 'VariableNames', {'interventionID', 'elementID', 'elementType', 'expDataID', 'expTimeVarID', 'expVarID', 'PatientIDVar'});
    else
        lookupRows  = cell2table(cell(0,10), 'VariableNames', {'interventionID', 'elementID', 'elementType', 'expDataID', 'expTimeVarID', 'expVarID', 'PatientIDVar', 'TRTVar', 'BRSCOREVar', 'RSCOREVar'});
    end
    tablesToCheck = {'mnSDTable','binTable','distTable'};
	tablesToCheck2 = {'distTable2D','corTable'};
	subpopNo = nan(0,1);
    for tableCounter = 1: length(tablesToCheck)
        curTable = get(myMapelOptions,tablesToCheck{tableCounter});
		[nRows, nCols] = size(curTable);
		if nRows > 0
			if isa(myMapelOptions, 'mapelOptions')
				lookupRows = [lookupRows;curTable(:,{'interventionID', 'elementID', 'elementType', 'expDataID', 'expTimeVarID', 'expVarID', 'PatientIDVar'})];
				subpopNo = [subpopNo; curTable{:,'subpopNo'}];
			else
				lookupRows = [lookupRows;curTable(:,{'interventionID', 'elementID', 'elementType', 'expDataID', 'expTimeVarID', 'expVarID', 'PatientIDVar', 'TRTVar', 'BRSCOREVar', 'RSCOREVar'})];
				subpopNo = [subpopNo; curTable{:,'subpopNo'}];
			end
		end
    end
    for tableCounter = 1: length(tablesToCheck2)
        curTable = get(myMapelOptions,tablesToCheck2{tableCounter});
		[nRows, nCols] = size(curTable);
		if nRows > 0
			if isa(myMapelOptions, 'mapelOptions')
				lookupRows = [lookupRows;curTable(:,{'interventionID1', 'elementID1', 'elementType1', 'expDataID1', 'expTimeVarID1', 'expVarID1', 'PatientIDVar1'})];
				subpopNo = [subpopNo; curTable{:,'subpopNo'}];
				lookupRows = [lookupRows;curTable(:,{'interventionID2', 'elementID2', 'elementType2', 'expDataID2', 'expTimeVarID2', 'expVarID2', 'PatientIDVar2'})];
				subpopNo = [subpopNo; curTable{:,'subpopNo'}];
			else
				lookupRows = [lookupRows;curTable(:,{'interventionID1', 'elementID1', 'elementType1', 'expDataID1', 'expTimeVarID1', 'expVarID1', 'PatientIDVar1', 'TRTVar1', 'BRSCOREVar1', 'RSCOREVar1'})];
				subpopNo = [subpopNo; curTable{:,'subpopNo'}];
				lookupRows = [lookupRows;curTable(:,{'interventionID2', 'elementID2', 'elementType2', 'expDataID2', 'expTimeVarID2', 'expVarID2', 'PatientIDVar2', 'TRTVar2', 'BRSCOREVar2', 'RSCOREVar2'})];
				subpopNo = [subpopNo; curTable{:,'subpopNo'}];
			end
		end
    end
    [lookupRows2, IA, IC] = unique(lookupRows,'rows','first');
    % IC provides a useful reference for the replicated blocks
    % in lookupRows.  We run a quick check to make sure these
    % map with the subpopulations, and provide a quick note of caution
    % if they do not...
    uniqueBlocks = unique(IC,'rows','first');
    for checkCounter = 1 : length(IA)
        rowsToCheck = find(IC == uniqueBlocks(checkCounter));
        if length(unique(subpopNo(rowsToCheck))) > 1
            disp(['Found sets of {intevention,elementID,elementType,expDataID,expTimeVarID,expVarID,...} that map to more than one subpulation in ',mfilename,'.  It is not expected that different subpopulations would be calibrated to the same data.  The combination that is causing the conflict is:'])
            lookupRows(rowsToCheck(1),:)
            disp(['Proceeding by assigning subpopNo ',num2str(subpopNo(rowsToCheck(1))),' to the expDataTable row.'])
        end
    end
    lookupRows = lookupRows2;
	% We don't double check the subpop data for conflicts
	% where the uniqueness is not consistent with the
	% other data, which could happen if there is a mistake
	% when entering the tables and updating the subpop information
	subpopNo = subpopNo(IA);
    lookupRows = lookupRows2;
    [nRows, nCols] = size(lookupRows);
    for rowCounter = 1 : nRows
        if rowCounter == 1
            initialRows = 0;
            if isa(myMapelOptions, 'mapelOptions')
				myExpDataTable = createExpDataTable(myWorksheet, lookupRows{rowCounter,'interventionID'}{1}, lookupRows{rowCounter,'elementID'}{1}, lookupRows{rowCounter,'elementType'}{1}, lookupRows{rowCounter,'expDataID'}{1}, lookupRows{rowCounter,'expTimeVarID'}{1}, lookupRows{rowCounter,'expVarID'}{1}, lookupRows{rowCounter,'PatientIDVar'}{1});
            else
                myExpDataTable = createExpDataTableRECIST(myWorksheet, lookupRows{rowCounter,'interventionID'}{1}, lookupRows{rowCounter,'elementID'}{1}, lookupRows{rowCounter,'elementType'}{1}, lookupRows{rowCounter,'expDataID'}{1}, lookupRows{rowCounter,'expTimeVarID'}{1}, lookupRows{rowCounter,'expVarID'}{1}, lookupRows{rowCounter,'PatientIDVar'}{1}, lookupRows{rowCounter,'TRTVar'}, lookupRows{rowCounter,'BRSCOREVar'}{1}, lookupRows{rowCounter,'RSCOREVar'}{1});
            end
        else
            [initialRows,~] = size(myExpDataTable);
            if isa(myMapelOptions, 'mapelOptions')
				myExpDataTable = createExpDataTable(myWorksheet, lookupRows{rowCounter,'interventionID'}{1}, lookupRows{rowCounter,'elementID'}{1}, lookupRows{rowCounter,'elementType'}{1}, lookupRows{rowCounter,'expDataID'}{1}, lookupRows{rowCounter,'expTimeVarID'}{1}, lookupRows{rowCounter,'expVarID'}{1}, lookupRows{rowCounter,'PatientIDVar'}{1}, myExpDataTable);
            else
                myExpDataTable = createExpDataTableRECIST(myWorksheet, lookupRows{rowCounter,'interventionID'}{1}, lookupRows{rowCounter,'elementID'}{1}, lookupRows{rowCounter,'elementType'}{1}, lookupRows{rowCounter,'expDataID'}{1}, lookupRows{rowCounter,'expTimeVarID'}{1}, lookupRows{rowCounter,'expVarID'}{1}, lookupRows{rowCounter,'PatientIDVar'}{1}, lookupRows{rowCounter,'TRTVar'}, lookupRows{rowCounter,'BRSCOREVar'}{1}, lookupRows{rowCounter,'RSCOREVar'}{1}, myExpDataTable);
            end
        end
        [finalRows,~] = size(myExpDataTable);
        if finalRows - initialRows > 0
            % Also add the known subpop information
            myExpDataTable{initialRows+1:finalRows,'subpopNo'} = subpopNo(rowCounter); 
        end
    end

else
    warning(['Unable to complete ',mfilename,', exiting.'])
end    
```
