## Supplementary File 3 for "A surrogate workflow for virtual population development: designing an amenable proxy for a hypothesis-test inspired nonlinear goodness-of-fit": convertResponseTypeToExpDataTable.m

```
function myExpDataTable = convertMapelOptionsToExpDataTable(myWorksheet, myResponseTypeID, patientIDVar)
% This function takes experimental data embedded in a worksheet and
% converts it to a data table format for use in MAPEL, guided by a
% response type
%
% ARGUMENTS:
%  myWorksheet:       A worksheet with the experimental data and response
%                      type attached
%  myResponseTypeID:  The response type to gather experimental data for
%  patientIDVar:      The patient ID variable.  This must be specified,
%                      and enables calibrating multivariate relationships
%                      later.  Note that in the current version, the same
%                      ID variable name must be used in all experimental
%                      datasets referenced.
%
% RETURNS
% myExpDataTable

continueFlag = true;
if nargin > 3
    continueFlag = false;
    warning(['Too many input arguments for ',mfilename,'. Should provide: myWorksheet, myResponseTypeID, patientIDVar.'])
    continueFlag = false;
elseif nargin > 2
    continueFlag = true;
else
    warning(['Insufficient input arguments for ',mfilename,'. Should provide: myWorksheet, myResponseTypeID.'])
    continueFlag = false;
end

if continueFlag
    allResponseTypeIDs = getResponseTypeIDs(myWorksheet);
    
    if sum(ismember(allResponseTypeIDs, myResponseTypeID)) < 1
        warning(['Specified responseTypeID not found in the provided worksheet in call to ',mfilename,'.'])
        continueFlag = false;
    
    else
        myResponseType = getResponseType(myWorksheet, myResponseTypeID);
        [nRTE,~] = size(myResponseType.elements);
        for responseTypeElementCounter = 1 : nRTE
            myRTE = myResponseType.elements{responseTypeElementCounter};
            if ~(strcmp(class(myRTE), 'responseTypeElementPoints'))
                warning(['Included responseTypeElementID ,', myRTE.id,', not of class responseTypeElementPoints, this case is not currently handled in ',myResponseTypeID,'.'])
                continueFlag = false;                
            end
        end
    end
end

if continueFlag
    [nRTE, dummy] = size(myResponseType.elements);
    for responseTypeElementCounter = 1 : nRTE
        % We may want to add a filter for the first data point,
        % in case initial conditions are specified,
        % or this can be done post-processing.
        myRTE = myResponseType.elements{responseTypeElementCounter};
        myInterventionID = myRTE.interventionID;
        myElementID = myRTE.modelYVar;
        myElementType = myRTE.modelYVarType;
        myExpDataID = myRTE.expDataID;
        myExpTimeVarID = myRTE.expDataTimeVar;
        myExpVarID = myRTE.expDataYVar;
        if responseTypeElementCounter == 1
            myExpDataTable = createExpDataTable(myWorksheet, myInterventionID, myElementID, myElementType, myExpDataID, myExpTimeVarID, myExpVarID, patientIDVar);
        else
            myExpDataTable = createExpDataTable(myWorksheet, myInterventionID, myElementID, myElementType, myExpDataID, myExpTimeVarID, myExpVarID, patientIDVar, myExpDataTable);
        end
    end
else
    warning(['Unable to complete ',mfilename,', exiting.'])
end    
```
