## Supplementary File 3 for "A surrogate workflow for virtual population development: designing an amenable proxy for a hypothesis-test inspired nonlinear goodness-of-fit": createBinTable.m

```
function myBinTable = createBinTable(myWorksheet,myExpDataIDs,PatientIDVar,myVars,myBinEdges,mySimVars,mySimVarTypes,timeVar,startTime,myInterventionIDs, nBins)
% This function takes experimental data embedded in a worksheet and
% converts it to a bin table for nonRECIST mapelOptions, bypassing reading from a mapelOptions
% "experimental data table" and directly taking data from the worksheet
%
% Note the cell arrays below are based on calibration variable x interventionID
% combinations.  That is, if multiple time points are available they will automatically be extracted.
% You can go back and remove rows you do not want or set weights to 0.
%
% ARGUMENTS:
%  myWorksheet:       A worksheet with the experimental data attached
%  myExpDataIDs:      A cell array, of length N intervention/var combinations
%                      with the experimental dataset IDs
%  PatientIDVar:      ID for the variable with the patient IDs
%  myVars:            A cell array, of length N intervention/var combinations
%                      variables in the dataset to get mean/SD summary data for.
%  mySimVars:         Simulation variable names, of length N intervention/var
%                      combinations, map to myVars.
%  myBinEdges         A cell array, of length N intervention/var combinations
%  mySimVars:         simulation variable names, 1 x n calibration criteria, map to myVars
%  mySimVarTypes:     A cell array, of length N intervention/var combinations
%                      with values to indicate the type of simulation
%                      variables, i.e.
%                      'parameter', 'compartment', 'species', 'observable'
%  timeVar:           ID for the variable with time in the experimental dataset
%                     NOTE: it is assumed start of therapy is t=startTime.
%  startTime		  Time to start therapy.
%  myInterventionIDs: A cell array, of length N intervention/var combinations
%                      with all of the intervention names from the simulations
%
% RETURNS
%  myBinTable

% TODO: Add proofing of inputs

continueFlag = true;
commonNames = loadCommonNames();
tableVariableNames = [commonNames.VPOPTABLEVARNAMESFIXED,{'weight','binEdges','expN','expBins','predN', 'predIndices','predBins'}];
myBinTable = cell2table(cell(0,length(tableVariableNames)));
myBinTable.Properties.VariableNames = tableVariableNames;


% Proofing of the input
% variables starts here
if continueFlag
	testN = length(myExpDataIDs);
	if sum([length(myExpDataIDs),length(myVars),length(mySimVars),length(mySimVarTypes),length(myInterventionIDs)] ~= testN) > 0
		continueFlag = false;
		warning(['Not all input cell array arguments are of consistent length in ',mfilename,'.  Exiting.'])
	end
end

if continueFlag
    
    allExpDataIDs = getExpDataIDs(myWorksheet);
    allInterventionIDs = getInterventionIDs(myWorksheet);
    
    for varCounter = 1 : testN
    
        curVar = myVars{varCounter};
        myExpDataID = myExpDataIDs{varCounter};
        interventionID = myInterventionIDs{varCounter};
  
        curIndex = find(ismember(allExpDataIDs,myExpDataID));
        
        if isempty(curIndex)
            warning(['Not able to find experimental dataset ',myExpDataID,' in ',mfilename,'.'])
            continueFlag = false;
        else        
            curData = myWorksheet.expData{curIndex}.data;   % nonRECIST: no off-treatment info, take all data
            [nRows,nCols] = size(curData);
            if nRows < 1
                warning(['Not able to find on treatment data for experimental dataset ',myExpDataID,' in ',mfilename,'.'])
                continueFlag = false;
            else
                if sum(ismember(curData.Properties.VariableNames,myVars{varCounter})) < 1
                    warning(['Not able to find experimental variable ',myVars{varCounter},' in dataset ',myExpDataID,' in ',mfilename,'.'])
                    continueFlag = false;
                end
                if sum(ismember(allInterventionIDs,interventionID)) < 1
                    warning(['Not able to find intervention ID ',interventionID,' associated with variable ',myVars{varCounter},' in dataset ',myExpDataID,' in ',mfilename,'.'])
                    continueFlag = false;                
                end
            end
        end
    end
	
	% Also proof whether the intervention/model variable inputs are unique.
	% Non-uniqueness would suggest mapping the same model/intervention
	% combination onto multiple outputs, which should not be allowed	
	mySimVars = reshape(mySimVars,testN,1);
	mySimVarTypes = reshape(mySimVarTypes,testN,1);
	myInterventionIDs = reshape(myInterventionIDs,testN,1);
    myExpDataIDs = reshape(myExpDataIDs,testN,1);
	combineRows = [mySimVars,mySimVarTypes,myInterventionIDs,myExpDataIDs];
	combineRows = cell2table(combineRows);
	[nRows,~] = size(unique(combineRows,'rows'));
	if nRows ~= testN
		warning(['Not all model variables, variable types, intervention sets are unique in call to ',mfilename,'.  Exiting.'])
		continueFlag = false;    
	end
end


if continueFlag

    for varCounter = 1 : testN

        curVar = myVars{varCounter};
        myExpDataID = myExpDataIDs{varCounter};
        interventionID = myInterventionIDs{varCounter};        

        curIndex = find(ismember(allExpDataIDs,myExpDataID));
        curData = myWorksheet.expData{curIndex}.data;

        selectData = curData;    % no filter based on RECIST, take all data
        curRows = find(~isnan(selectData{:,curVar}));        
        if length(curRows) >= 2
            % First get the bin edges from all variable data points
            allCurVarValues = selectData{curRows, curVar};
            myBinEdgeValues = myBinEdges{varCounter};
			nBins = length(myBinEdgeValues)+1;
            % Now we need to step through and evaluate for the times
            curTimes = unique(selectData{curRows,timeVar});
            for curTimeCounter = 1: length(curTimes)
                curDataRows = find(selectData{curRows,timeVar} == curTimes(curTimeCounter));
                if length(curDataRows) >= 2
                    curData = selectData{curRows(curDataRows),curVar};
                    % We only keep add a row if the largest value is bigger
                    % than the smallest.  Sometimes when normailizing to
                    % baseline variables with zero variability are created
                    % at T=0 and it makes sense to filter these.
                    if max(curData) > min(curData)
                        curProbs = wtdBinProb(curData', ones(1, length(curData))/length(curData), myBinEdgeValues);
                        expN = length(curData);
						% When initially transferring the data over,
						% assume it is from the "all" subpop.						
                        curRow = {1, curTimes(curTimeCounter), interventionID, mySimVars{varCounter}, mySimVarTypes{varCounter}, myExpDataID, timeVar, curVar, PatientIDVar};
						curRow = [curRow,{1,{myBinEdgeValues}, expN, {curProbs}, nan, {nan}, {nan(1,nBins)}}];
                        curRow = cell2table(curRow);
                        curRow.Properties.VariableNames = myBinTable.Properties.VariableNames; 
                        myBinTable = [myBinTable; curRow];
                    end
                end
            end
        end
    end
end
        
end
```
