## Supplementary File 3 for "A surrogate workflow for virtual population development: designing an amenable proxy for a hypothesis-test inspired nonlinear goodness-of-fit": createExpDataTable.m

```
function myExpDataTable = createExpDataTable(myWorksheet, myInterventionID, elementID, elementType, expDataID, expTimeVarID, expVarID, PatientIDVar, previousExpDataTable)
% Convert experimental data attached to a worksheet
% into a format that can
% can be used in the population weighting algorithms.
%
% Note myExpDataTable currently assumes the data is intended the
% all VPs.  If you wish to map to a different subopulation,
% you can edit the subpopNo reference for each row entry after
% the table is made.
%
% ARGUMENTS
% myWorksheet:           a worksheet containing the data
% myInterventionID:      the ID of the worksheet intervention to map the 
% elementID:             data to the variable ID in the simulation results
% elementType:           type of variable to map to: parameter, 
%                        compartment, species
% expDataID:             dataset ID for the experimental data
% expTimeVarID:          name of the time variable in the experimental 
%                        dataset to map onto the simulation time.  Note if 
%                        the simulated intervention uses a delay or time 
%                        shift, there may need to be a shift in the 
%                        experimental data here to align the two.  Time 
%                        units should be consistent.
% expVarID:              Column ID for the experimental data to map
% PatientIDVar:          variable with the patient IDs
% previousExpDataTable:  a previous output of the function, if provided the 
%                        additional results will be appended to the end. 
%
% RETURNS
% myExpDataTable:       A MATLAB table with the data and mappings.  This will
%                   have headings:
%                   'time', 'interventionID', 'elementID', 'elementType', 'expDataID', 'expTimeVarID', 'expVarID', '', 'expVal1', 'expVal2', ... , 'expValN'
%
continueFlag = true;
myExpDataTable = table();
if nargin > 9
    continueFlag = false;
    warning(['Too many input arguments for ',mfilename,'. Should provide: myWorksheet, myInterventionID, elementID, elementType, expDataID, expTimeVarID, expVarID, PatientIDVar; optionally: previousExpDataTable.'])
    continueFlag = false;
elseif nargin > 8
    continueFlag = true;
elseif nargin > 7
    previousExpDataTable = table();
else
    warning(['Insufficient input arguments for ',mfilename,'. Should provide: myWorksheet, myInterventionID, elementID, elementType, expDataID, expTimeVarID, expVarID, PatientIDVar; optionally: previousExpDataTable.'])
    continueFlag = false;
end

if continueFlag
    worksheetInterventionIDs = getInterventionIDs(myWorksheet);
    if sum(ismember(worksheetInterventionIDs,myInterventionID)) < 1
        continueFlag = false;
        warning(['Missing specified InterventionID in worksheet supplied to ',mfilename,'.'])
    end
    if sum(ismember(fields(myWorksheet),'compiled')) < 1
        continueFlag = false;
        warning(['Worksheet supplied to ',mfilename,' should have a compiled model.'])
    else
        elementType = lower(elementType);
        myWorksheetElements = myWorksheet.compiled.elements;
        if sum(ismember(myWorksheetElements(:,2),elementType)) < 1
            continueFlag = false;
            warning(['The elementType supplied to ',mfilename,' should be species, compartment, or parameter.'])
        else
            if sum(ismember(myWorksheetElements(:,1),elementID)) < 1
                continueFlag = false;
                warning(['The elementID supplied to ',mfilename,' was not identified in the compiled model.'])
            end
        end
    end
    worksheetExpDataIDs = getExpDataIDs(myWorksheet);
    if sum(ismember(worksheetExpDataIDs,expDataID)) < 1
        continueFlag = false;
        warning(['The expDataID supplied to ',mfilename,' was not identified in the supplied worksheet.']);       
    else
        myExpTable = getExpData(expDataID, myWorksheet);
        if sum(ismember(myExpTable.Properties.VariableNames, expTimeVarID)) < 1
            continueFlag = false;
            warning(['The expTimeVarID supplied to ',mfilename,' was not identified in the specified dataset.'])
        end
        if sum(ismember(myExpTable.Properties.VariableNames, expVarID)) < 1
            continueFlag = false;
            warning(['The expVarID supplied to ',mfilename,' was not identified in the specified dataset.'])
        end
        if sum(ismember(myExpTable.Properties.VariableNames, PatientIDVar)) < 1
            continueFlag = false;
            warning(['The PatientIDVar supplied to ',mfilename,' was not identified in the specified dataset.'])
        end
            
    end
end
  
if continueFlag
    commonNames = loadCommonNames();
    tableVariableNamesFixed = commonNames.VPOPEXPTABLEVARNAMESFIXED;
    [nPreviousRows, nPreviousColumns] = size(previousExpDataTable);
    if nPreviousRows >= 1
        previousVariableNames = previousExpDataTable.Properties.VariableNames;
        previousPatientIDs = setdiff(previousVariableNames,tableVariableNamesFixed);        
        if sum(ismember(tableVariableNamesFixed, previousVariableNames)) ~= length(tableVariableNamesFixed)
            continueFlag = false;
            warning(['When providing a previousExpDataTable to ',mfilename,', the VariableNames must include: ',strjoin(tableVariableNamesFixed,', '),'.'])
        %elseif sum(strncmpi('expVal',diffNames,6)) ~= length(diffNames)
        %    continueFlag = false;
        %    warning(['When providing a previousExpDataTable to ',mfilename,', the VariableNames must include only: ',strjoin(tableVariableNamesFixed,', '),', and "expValXX".'])       
        else
            existingN = length(previousPatientIDs);
            myExpDataTable = previousExpDataTable;
        end
    else
        existingN = 0;
		previousPatientIDs = cell(1,0);
        myExpDataTable = cell2table(cell(0,length(tableVariableNamesFixed)));
        myExpDataTable.Properties.VariableNames = tableVariableNamesFixed;
        myExpDataTable.Properties.VariableDescriptions = tableVariableNamesFixed;
    end
end

if continueFlag
    expDataTime = myExpTable.(expTimeVarID);
    expDataVar = myExpTable.(expVarID);
	expDataPatientID = myExpTable.(PatientIDVar);
    the_indices = find((~isnan(expDataTime)) & (~isnan(expDataVar)));
    expDataTime = expDataTime(the_indices);
    expDataVar = expDataVar(the_indices);
	expDataPatientID = expDataPatientID(the_indices);
    uniqueExpDataTime = unique(expDataTime);
    for timePointCounter = 1 : length(uniqueExpDataTime)
        curTime = uniqueExpDataTime(timePointCounter);
        originalRowIndices = find(expDataTime == curTime);
        allValues = (expDataVar(originalRowIndices));
		allIDs = expDataPatientID(originalRowIndices);
        % Enforce converting any numeric IDs to strings.
        if isnumeric(allIDs)
            [nIDs, ~] = size(allIDs);
            allIDsold = allIDs;
            allIDs=cell(nIDs,1);
            for idCounter = 1 : nIDs
                allIDs{idCounter} = num2str(allIDsold(idCounter));
            end
        end      
        curN = length(originalRowIndices);
		
		% Find the indices for pre-existing Patient IDs in the current row
		% TODO: expect this is slow, try to speed up
		initialCells = {1, curTime, myInterventionID, elementID, elementType, expDataID, expTimeVarID, expVarID, PatientIDVar};
		previousVariableNames = myExpDataTable.Properties.VariableDescriptions;
       
		foundMate = [];
		curRow = nan(1,length(previousVariableNames)-length(initialCells));
        arrangedNames = cell(1,length(previousVariableNames)-length(initialCells));
		for i = 1 : curN
			curTest = allIDs(i);
			curLogicals = ismember(previousVariableNames,curTest);
			if sum(curLogicals) > 0
				foundMate = [foundMate,i];
				curRow(find(curLogicals)-length(initialCells)) = allValues(i);
                arrangedNames(find(curLogicals)-length(initialCells)) = allIDs(i);
			end
		end
		allValues(foundMate) = [];
        foundIDs = allIDs(foundMate);
		allIDs(foundMate) = [];
		curRow = [cell2table(initialCells),array2table(curRow)];
        %curRow = cell2table(curRow);
        curRow.Properties.VariableNames = myExpDataTable.Properties.VariableNames;
        curRow.Properties.VariableDescriptions = myExpDataTable.Properties.VariableDescriptions; 
        myExpDataTable = [myExpDataTable; curRow];
		if length(allValues) > 0
			[nRows, ~] = size(myExpDataTable);
			curVals = nan(nRows,length(allValues));
			curVals(nRows,:) = allValues;
			curVals = array2table(curVals);%,'VariableNames',allIDs);
			%curVals.VariableNames = allIDs;
            curVals.Properties.VariableNames = genvarname(allIDs);
            curVals.Properties.VariableDescriptions = allIDs;
			myExpDataTable = [myExpDataTable, curVals];
		end		
		
    end
else
    warning(['Unable to complete ',mfilename,', exiting.'])
end

end
```
