## Supplementary File 3 for "A surrogate workflow for virtual population development: designing an amenable proxy for a hypothesis-test inspired nonlinear goodness-of-fit": createMnSDTable.m

```
function myMnSDTable = createMnSDTable(myWorksheet,myExpDataIDs,PatientIDVar,myVars,mySimVars,mySimVarTypes,timeVar,startTime,myInterventionIDs)
% This function takes experimental data embedded in a worksheet and
% converts it to a mean/sd table for nonRECIST mapelOptions, bypassing reading from a mapelOptions
% "experimental data table" and directly taking data from the worksheet,
%
% Note the cell arrays below are based on calibration variable x interventionID
% combinations.  That is, if multiple time points are available they will automatically be extracted.
% You can go back and remove rows you do not want or set weights to 0.
%
% ARGUMENTS:
%  myWorksheet:       A worksheet with the experimental data attached
%  myExpDataIDs:      A cell array, of length N intervention/var combinations
%                      with the experimental dataset IDs
%  PatientIDVar:      ID for the variable with the patient IDs
%  myVars:            A cell array, of length N intervention/var combinations
%                      variables in the dataset to get mean/SD summary data for.
%  mySimVars:         Simulation variable names, of length N intervention/var
%                      combinations, map to myVars.
%  mySimVarTypes:     A cell array, of length N intervention/var combinations
%                      with values to indicate the type of simulation
%                      variables, i.e.
%                      'parameter', 'compartment', 'species', 'observable'
%  timeVar:           ID for the variable with time in the experimental dataset
%                     NOTE: it is assumed start of therapy is t=startTime.
%  startTime		  Time to start therapy.
%  myInterventionIDs: A cell array, of length N intervention/var combinations
%                      with all of the intervention names from the simulations
%
% RETURNS
%  myMnSDTable

continueFlag = true;

commonNames = loadCommonNames();
tableVariableNames = [commonNames.VPOPTABLEVARNAMESFIXED,{'logN','weightMean', 'weightSD', 'expN', 'expMean', 'expSD', 'predN', 'predIndices', 'predSample', 'predMean', 'predSD'}];
myMnSDTable = cell2table(cell(0,length(tableVariableNames)));
myMnSDTable.Properties.VariableNames = tableVariableNames;

% Proofing of the input
% variables starts here
if continueFlag
	testN = length(myExpDataIDs);
	if sum([length(myExpDataIDs),length(myVars),length(mySimVars),length(mySimVarTypes),length(myInterventionIDs)] ~= testN) > 0
		continueFlag = false;
		warning(['Not all input cell array arguments are of consistent length in ',mfilename,'.  Exiting.'])
	end
end

if continueFlag
    
    allExpDataIDs = getExpDataIDs(myWorksheet);
    allInterventionIDs = getInterventionIDs(myWorksheet);
    
    for varCounter = 1 : testN
    
        curVar = myVars{varCounter};
        myExpDataID = myExpDataIDs{varCounter};
        interventionID = myInterventionIDs{varCounter};
  
        curIndex = find(ismember(allExpDataIDs,myExpDataID));
        
        if isempty(curIndex)
            warning(['Not able to find experimental dataset ',myExpDataID,' in ',mfilename,'.'])
            continueFlag = false;
        else        
            curData = myWorksheet.expData{curIndex}.data; % nonRECIST: no off-treatment info, take all data
            [nRows,nCols] = size(curData);
            if nRows < 1
                warning(['Not able to find on treatment data for experimental dataset ',myExpDataID,' in ',mfilename,'.'])
                continueFlag = false;
            else
                if sum(ismember(curData.Properties.VariableNames,myVars{varCounter})) < 1
                    warning(['Not able to find experimental variable ',myVars{varCounter},' in dataset ',myExpDataID,' in ',mfilename,'.'])
                    continueFlag = false;
                end
                if sum(ismember(allInterventionIDs,interventionID)) < 1
                    warning(['Not able to find intervention ID ',interventionID,' associated with variable ',myVars{varCounter},' in dataset ',myExpDataID,' in ',mfilename,'.'])
                    continueFlag = false;                
                end
            end
        end
    end
	
	% Also proof whether the intervention/model variable inputs are unique.
	% Non-uniqueness would suggest mapping the same model/intervention
	% combination onto multiple outputs, which should not be allowed	
	mySimVars = reshape(mySimVars,testN,1);
	mySimVarTypes = reshape(mySimVarTypes,testN,1);
	myInterventionIDs = reshape(myInterventionIDs,testN,1);
    myExpDataIDs = reshape(myExpDataIDs,testN,1);
	combineRows = [mySimVars,mySimVarTypes,myInterventionIDs,myExpDataIDs];
	combineRows = cell2table(combineRows);
	[nRows,~] = size(unique(combineRows,'rows'));
	if nRows ~= testN
		warning(['Not all model variables, variable types, intervention sets are unique in call to ',mfilename,'.  Exiting.'])
		continueFlag = false;    
	end
end


if continueFlag
    expN = nan(0,1);
    expMean = nan(0,1);
    expSD = nan(0,1);
    expTime = nan(0,1);
	subpopNo = nan(0,1);
    interverventionIDCell = cell(0,1);
    elementIDCell = cell(0,1);
    elementTypeCell = cell(0,1);
    expDataIDCell = cell(0,1);
    expTimeVarIDCell = cell(0,1);
    expVarIDCell = cell(0,1);  
    PatientIDCell = cell(0,1);
   
    for varCounter = 1 : testN
        curVar = myVars{varCounter};
        myExpDataID = myExpDataIDs{varCounter};
        interventionID = myInterventionIDs{varCounter};
  
        curIndex = find(ismember(allExpDataIDs,myExpDataID));
        curData = myWorksheet.expData{curIndex}.data;

        selectData = curData;  % no filter based on RECIST, take all data
        curRows = find(~isnan(selectData{:,curVar}));
        if length(curRows) >= 2
            curTimes = unique(selectData{curRows,timeVar});
            for curTimeCounter = 1: length(curTimes)
                curDataRows = find(selectData{curRows,timeVar} == curTimes(curTimeCounter));
                curData = selectData{curRows(curDataRows),curVar};
                if length(curDataRows) >= 2
					% When initially transferring the data over,
					% assume it is from the "all" subpop.
					subpopNo = [subpopNo; 1];
                    expN = [expN; length(curDataRows)];
                    expMean = [expMean; mean(curData)];
                    expSD = [expSD; std(curData)];
                    expTime = [expTime; curTimes(curTimeCounter)];
                    interverventionIDCell = [interverventionIDCell; interventionID];
                    elementIDCell = [elementIDCell; mySimVars{varCounter}];
                    elementTypeCell = [elementTypeCell; mySimVarTypes{varCounter}];
                    expDataIDCell = [expDataIDCell; myExpDataID];
                    expTimeVarIDCell = [expTimeVarIDCell; timeVar];
                    expVarIDCell = [expVarIDCell; curVar];
                    PatientIDCell = [PatientIDCell; PatientIDVar];                 
                end
            end
        end
    end
    
    myMnSDTable = table(subpopNo, expTime,interverventionIDCell,elementIDCell,elementTypeCell,expDataIDCell,expTimeVarIDCell,expVarIDCell,PatientIDCell,false(length(expTime),1),ones(length(expTime),1),ones(length(expTime),1),expN,expMean,expSD,nan(length(expTime),1),repmat({{nan}},length(expTime),1),repmat({nan},length(expTime),1),nan(length(expTime),1),nan(length(expTime),1));
    % We also filter where SD > 0.  SD = 0 can show up when normalizing
    % to a baseline value, and it doesn't make sense to calibrate
    % the baseline value to zero.
    myMnSDTable.Properties.VariableNames = tableVariableNames;
    myMnSDTable = myMnSDTable(find(myMnSDTable{:,'expSD'}>0),:);
else
    warning(['Unable to complete ',mfilename,', exiting.'])    
end    
```
