## Supplementary File 3 for "A surrogate workflow for virtual population development: designing an amenable proxy for a hypothesis-test inspired nonlinear goodness-of-fit": createResponseTablesRECIST.m

```
function [myCBRTableRECIST, myBRTableRECIST, myRTableRECIST] = createResponseTablesRECIST(myWorksheet,myExpDataIDs,PatientIDVar,TRTVar,CBRSCOREVar,BRSCOREVar,RSCOREVar,timeVar,startTime,myInterventionIDs, prefixStopPairs)
% This function takes experimental data embedded in a worksheet and
% converts it to bin tables with BRSCORE and RSCORE data
%
% ARGUMENTS:
%  myWorksheet:       A worksheet with the experimental data attached
%  myExpDataIDs:      A cell array, of length N intervention combinations
%                      with the experimental dataset IDs
%  PatientIDVar:      ID for the variable with the patient IDs
%  TRTVar:            ID for the binary variable indicating whether 
%                      a patient is on treatment
%  CBRSCOREVar:        ID for the confirmed best overall response to date variable.
%  BRSCOREVar:        ID for the best overall response to date variable.
%  RSCOREVar:         ID for the current response variable.
%  RSCOREVar:         variable with current response.
%  timeVar:           variable with time information
%  startTime:         time in the data when treatment starts
%  myInterventionIDs: A cell array, of length N intervention/var combinations
%                      with all of the intervention names from the simulations
%  prefixStopPairs:   (optional) A cell array of cell arrays, i.e.:
%                      {{prefix1, time1},{prefix2, time2} ...}
%                      containing VP prefix and trial stop time.
%                      This is included to correct for trials that
%                      end earlier and we want to avoid biasing results by
%                      treating a large number of patients as toxicity-related
%                      drop outs.  With USUBJID conventions, the prefix
%                      generally matches a trial, i.e. CAXXXYYYY.
%  
% RETURNS
%  myCBRTableRECIST   Will be empty if there is no CBRSCORE information in input data file
%  myBRTableRECIST
%  myRTableRECIST
%

% TODO: might want to add more proofing for prefixStopPairs

continueFlag = false;
if nargin > 11
    warning(['Too many input arguments to ',mfilename, '. Arguments should be: myWorksheet,myExpDataIDs,PatientIDVar,TRTVar,BRSCOREVar,RSCOREVar,timeVar,startTime,myInterventionIDs, optionally prefixStopPairs.'])
    continueFlag = false;
elseif nargin > 10
    continueFlag = true;
elseif nargin > 9
    prefixStopPairs = cell(1,0);
    continueFlag = true; 
else
    warning(['Too few input arguments to ',mfilename, '. Arguments should be: myWorksheet,myExpDataIDs,PatientIDVar,TRTVar,BRSCOREVar,RSCOREVar,timeVar,startTime,myInterventionIDs, optionally prefixStopPairs.'])
    continueFlag = false;
end

% Proofing of the input
% variables starts here
if continueFlag
	testN = length(myExpDataIDs);
	if sum([length(myExpDataIDs),length(myInterventionIDs)] ~= testN) > 0
		continueFlag = false;
		warning(['myExpDataIDs and myInterventionIDs are not of consistent length in ',mfilename,'.  Exiting.'])
	end
end

if continueFlag
    
    allExpDataIDs = getExpDataIDs(myWorksheet);
    allInterventionIDs = getInterventionIDs(myWorksheet);
    
    for varCounter = 1 : testN
    
        myExpDataID = myExpDataIDs{varCounter};
        interventionID = myInterventionIDs{varCounter};
  
        curIndex = find(ismember(allExpDataIDs,myExpDataID));
        
        if isempty(curIndex)
            warning(['Not able to find experimental dataset ',myExpDataID,' in ',mfilename,'.'])
            continueFlag = false;
        else        
            curData = myWorksheet.expData{curIndex}.data;
            % First remove off treatment rows
            curData = curData(find(curData{:,TRTVar}==1),:);
            [nRows,nCols] = size(curData);
            if nRows < 1
                warning(['Not able to find on treatment data for experimental dataset ',myExpDataID,' in ',mfilename,'.'])
                continueFlag = false;
            else
                if sum(ismember(allInterventionIDs,interventionID)) < 1
                    warning(['Not able to find intervention ID ',interventionID,' associated with dataset ',myExpDataID,' in ',mfilename,'.'])
                    continueFlag = false;                
                end
            end
        end
    end
	
	% Also proof whether the intervention/dataset inputs are unique.
	% Non-uniqueness would suggest a redundancy in specification
	% which could be a mistake
	myInterventionIDs = reshape(myInterventionIDs,testN,1);
	myExpDataIDs = reshape(myExpDataIDs,testN,1);
	combineRows = [myInterventionIDs,myExpDataIDs];
	combineRows = cell2table(combineRows);
	[nRows,~] = size(unique(combineRows,'rows'));
	if nRows ~= testN
		warning(['The experimental datasets and intervention sets are not uniquely paired in call to ',mfilename,'.  Exiting.'])
		continueFlag = false;    
	end
end

nStopTrials = length(prefixStopPairs);
prefixes = cell(1,nStopTrials);
stopTimes = nan(1,nStopTrials);
if nStopTrials > 0
    for prefixCounter = 1 : nStopTrials
        prefixes{prefixCounter} = prefixStopPairs{prefixCounter}{1};
        stopTimes(prefixCounter) = prefixStopPairs{prefixCounter}{2};
    end
end

commonNames = loadCommonNames();
rTableVariableNames = [commonNames.VPOPRECISTRESPONSETABLEVARNAMESFIXED,{'weight','expN','expCR','expPR','expSD','expPD','predN','predCR','predPR','predSD','predPD'}];
brTableVariableNames = [commonNames.VPOPRECISTRESPONSETABLEVARNAMESFIXED,{'weight','expN','expCR','expPR','expSD','expPD','predN','predCR','predPR','predSD','predPD','expNPD21LS','predNPD21LS'}]; % ,'expNMS1LS'
myBRTableRECIST = cell2table(cell(0,length(brTableVariableNames)));
myBRTableRECIST.Properties.VariableNames = brTableVariableNames;
myRTableRECIST = cell2table(cell(0,length(rTableVariableNames)));
myRTableRECIST.Properties.VariableNames = rTableVariableNames;
myCBRTableRECIST = myBRTableRECIST;

if continueFlag

    for varCounter = 1 : testN
        
        myExpDataID = myExpDataIDs{varCounter};
        interventionID = myInterventionIDs{varCounter};            
        
        curIndex = find(ismember(allExpDataIDs,myExpDataID));
        curData = myWorksheet.expData{curIndex}.data;
        
        % First remove off treatment rows
        curData = curData(find(curData{:,TRTVar}==1),:);
        % Need to make all table vars into string for this to work
        new_variable = arrayfun(@(value) cast(value{1}, 'char'), curData.(BRSCOREVar), 'uniform', 0);
        curData.(BRSCOREVar) = new_variable;
        new_variable = arrayfun(@(value) cast(value{1}, 'char'), curData.(RSCOREVar), 'uniform', 0);
        curData.(RSCOREVar) = new_variable; 
        if ismember(CBRSCOREVar,curData.Properties.VariableNames)
            new_variable = arrayfun(@(value) cast(value{1}, 'char'), curData.(CBRSCOREVar), 'uniform', 0);
            curData.(CBRSCOREVar) = new_variable;
            % We use CBR to filter
            curData = curData(find(ismember(curData{:,CBRSCOREVar},{'PD','SD','PR','CR','PD2'})),:); 
        else
            % We are just looking for BR, so this filter makes sense
            curData = curData(find(ismember(curData{:,BRSCOREVar},{'PD','SD','PR','CR','PD2'})),:); 
        end
                
        % Now also filter once a patient hits CR, PD
        curPatientIDs = unique(curData{:,PatientIDVar},'stable');
        nPatients = length(curPatientIDs);
        allRows = nan(0,1);
        PD2onlyRows = nan(0,1);
        for patientCounter = 1 : nPatients
            curRows = find(ismember(curData{:,PatientIDVar},curPatientIDs(patientCounter)));
            %lastRows = find(ismember(curData{curRows,BRSCOREVar},{'PD','CR'}) | ismember(curData{curRows,RSCOREVar},{'PD','CR'}));
            % First filter out PD2 and subsequent time points
			% We will exclude all data with PD2 for now
            if ismember(CBRSCOREVar,curData.Properties.VariableNames)
                lastRows = find(ismember(curData{curRows,CBRSCOREVar},{'PD2'}) | ismember(curData{curRows,BRSCOREVar},{'PD2'}) | ismember(curData{curRows,RSCOREVar},{'PD2'}));
            else
                lastRows = find(ismember(curData{curRows,BRSCOREVar},{'PD2'}) | ismember(curData{curRows,RSCOREVar},{'PD2'}));
            end
			if ~isempty(lastRows)
                if ismember(1,lastRows) % corrected from lastRows == 1  % look for PD2 patients (at least PD2 at first lesion scan)
                    PD2onlyRows = [PD2onlyRows;curRows(1:lastRows)];
                    lastRows = lastRows(1)-1;
                else
                    lastRows = lastRows(1)-1;
                end
                if lastRows > 0
                    curRows = curRows(1:lastRows);
                else
                    curRows = [];
                end
            end
            if length(curRows) > 0  
                lastRows = find(ismember(curData{curRows,RSCOREVar},{'PD','CR'}));
                if (length(lastRows) > 1)
                    curRows = curRows(1:lastRows(1));
                end
                allRows = [allRows; curRows];
            end
        end
        pd2Data = curData(PD2onlyRows,:);
        selectData = curData(allRows,:);
        new_variable = arrayfun(@(value) cast(value{1}, 'char'), selectData.(BRSCOREVar), 'uniform', 0);
        selectData.(BRSCOREVar) = new_variable;  
        new_variable = arrayfun(@(value) cast(value{1}, 'char'), selectData.(RSCOREVar), 'uniform', 0);
        selectData.(RSCOREVar) = new_variable;   
        if ismember(CBRSCOREVar,selectData.Properties.VariableNames)
            new_variable = arrayfun(@(value) cast(value{1}, 'char'), selectData.(CBRSCOREVar), 'uniform', 0);
            selectData.(CBRSCOREVar) = new_variable;   
        end

        % We want a full matrix with nTimePoint x nVP with the BRSCORE.
        % We also want to remove times that are missing a RSCORE measure
        % Sometimes missing values are entered as blanks
        selectData = selectData(find(~ismember(selectData{:,RSCOREVar},{' ',char(0)})),:);
        
        allTimes = unique(selectData{:,timeVar});
        allTimes = sort(allTimes,'ascend');

        % This may be redundant.  That is, we don't expect a response
        % unless we are beyond startTime anyway.
        allTimes = allTimes(find(allTimes>startTime));
        fullCell = cell(length(allTimes),nPatients);
        cumPD2 = zeros(length(allTimes),1);
        for patientCounter = 1 :nPatients
            patientID = curPatientIDs{patientCounter};
            curRows = find(ismember(selectData{:,PatientIDVar},patientID));
            curData = selectData(curRows,:);
            lastBRSCORE = '.';
            for timeCounter = 1 : length(allTimes)
                curTime = allTimes(timeCounter);
				% See if we can update lastBRSCORE, if not we will propagate
				% the last value
                if sum(ismember(curData{:,timeVar},curTime)) > 0
                    curRow = find((ismember(curData{:,timeVar},curTime)));
                    lastBRSCORE = curData{curRow,BRSCOREVar};
                    lastBRSCORE = lastBRSCORE{1};
                end
                % We need to add a check in case the trial has stopped
                passCheck = true;
                if length(prefixes) > 0
                    for prefixCounter = 1:length(prefixes)
                        if startsWith(patientID,prefixes{prefixCounter},'IgnoreCase',true) && (curTime > stopTimes(prefixCounter))
                            passCheck = false;
                        end
                    end
                end
                if ~passCheck
                    lastBRSCORE = '.';
                end
                fullCell{timeCounter,patientCounter} = lastBRSCORE;
                
            end
            
        end
        
        for timeCounter = 1 : length(allTimes)
            curTime = allTimes(timeCounter);
            cumPD2(timeCounter) = sum(pd2Data{:,timeVar}<=curTime);
        end
		
		% Now we have fullCell as nTimePoints x nVP
		% Table of best responses.  Now get the summary
		% information to guide calibration.
        
        % Now we want to count...
        for timeCounter = 1 : length(allTimes)
            curData = fullCell(timeCounter,:);
            nCR = sum(ismember(curData,'CR'));
            nPR = sum(ismember(curData,'PR'));
            nSD = sum(ismember(curData,'SD'));
            nPD = sum(ismember(curData,'PD'));
            expN = nCR+nPR+nSD+nPD;
            curProbs = [nCR, nPR, nSD, nPD] / expN;
            curRow = {1, allTimes(timeCounter), 'BRSCORE', interventionID, 'BRSCORE','derived',myExpDataID, timeVar,PatientIDVar,TRTVar,BRSCOREVar,RSCOREVar};
            curRow = [curRow,{1, expN, curProbs(1), curProbs(2), curProbs(3), curProbs(4), nan, nan, nan, nan, nan,cumPD2(timeCounter),nan}];
            curRow = cell2table(curRow);
            curRow.Properties.VariableNames = myBRTableRECIST.Properties.VariableNames; 
            myBRTableRECIST = [myBRTableRECIST; curRow];    
        end
        
        % %% CBRTableRECIST
        % We want a full matrix with nTimePoint x nVP with the BRSCORE.
        % We also want to remove times that are missing a RSCORE measure
        % Sometimes missing values are entered as blanks
        if ismember(CBRSCOREVar,selectData.Properties.VariableNames)
            allTimes = unique(selectData{:,timeVar});
            allTimes = sort(allTimes,'ascend');

            % This may be redundant.  That is, we don't expect a response
            % unless we are beyond startTime anyway.
            allTimes = allTimes(find(allTimes>startTime));
            fullCell = cell(length(allTimes),nPatients);
            cumPD2 = zeros(length(allTimes),1);
            for patientCounter = 1 :nPatients
                patientID = curPatientIDs{patientCounter};
                curRows = find(ismember(selectData{:,PatientIDVar},patientID));
                curData = selectData(curRows,:);
                lastCBRSCORE = '.';
                for timeCounter = 1 : length(allTimes)
                    curTime = allTimes(timeCounter);
                    % See if we can update lastCBRSCORE, if not we will propagate
                    % the last value
                    if sum(ismember(curData{:,timeVar},curTime)) > 0
                        curRow = find((ismember(curData{:,timeVar},curTime)));
                        lastCBRSCORE = curData{curRow,CBRSCOREVar};
                        lastCBRSCORE = lastCBRSCORE{1};
                    end
                    % We need to add a check in case the trial has stopped
                    passCheck = true;
                    if length(prefixes) > 0
                        for prefixCounter = 1:length(prefixes)
                            if startsWith(patientID,prefixes{prefixCounter},'IgnoreCase',true) && (curTime > stopTimes(prefixCounter))
                                passCheck = false;
                            end
                        end
                    end
                    if ~passCheck
                        lastCBRSCORE = '.';
                    end
                    fullCell{timeCounter,patientCounter} = lastCBRSCORE;

                end

            end

            for timeCounter = 1 : length(allTimes)
                curTime = allTimes(timeCounter);
                cumPD2(timeCounter) = sum(pd2Data{:,timeVar}<=curTime);
            end

            % Now we have fullCell as nTimePoints x nVP
            % Table of best responses.  Now get the summary
            % information to guide calibration.

            % Now we want to count...
            for timeCounter = 1 : length(allTimes)
                curData = fullCell(timeCounter,:);
                nCR = sum(ismember(curData,'CR'));
                nPR = sum(ismember(curData,'PR'));
                nSD = sum(ismember(curData,'SD'));
                nPD = sum(ismember(curData,'PD'));
                expN = nCR+nPR+nSD+nPD;
                curProbs = [nCR, nPR, nSD, nPD] / expN;
                curRow = {1, allTimes(timeCounter), 'CBRSCORE', interventionID, 'CBRSCORE','derived',myExpDataID, timeVar,PatientIDVar,TRTVar,CBRSCOREVar,RSCOREVar};
                curRow = [curRow,{1, expN, curProbs(1), curProbs(2), curProbs(3), curProbs(4), nan, nan, nan, nan, nan,cumPD2(timeCounter),nan}];
                curRow = cell2table(curRow);
                curRow.Properties.VariableNames = myCBRTableRECIST.Properties.VariableNames; 
                myCBRTableRECIST = [myCBRTableRECIST; curRow];    
            end      
        end
        
        % Repeat for the RSCORE
        % We want a full matrix with nTimePoint x nVP with the RSCORE.
        % We want to keep the last value once a patient hits PD or CR
        % but also allow missing values if a patients stops therapy before
        % PD or CR.
        % If we do not have data at a time point but the patient has
        % not hit PD/CR yet, we impute from the last observation
        allTimes = unique(selectData{:,timeVar});
        allTimes = sort(allTimes,'ascend');
        % This may be redundant.  That is, we don't expect a response
        % unless we are beyond startTime anyway.
        allTimes = allTimes(find(allTimes>startTime));
        fullCell = cell(length(allTimes),nPatients);
        for patientCounter = 1 :nPatients
            patientID = curPatientIDs{patientCounter};
            curRows = find(ismember(selectData{:,PatientIDVar},patientID));
            curData = selectData(curRows,:);
            lastRSCORE = '';
            % Last time we have for the current VP
            lastTime = max(curData{:,timeVar});
            for timeCounter = 1 : length(allTimes)
                curTime = allTimes(timeCounter);
                % Update the last RSCORE if we have
                % a measure for the current time for the current VP
                if sum(ismember(curData{:,timeVar},curTime)) > 0
                    curRow = find((ismember(curData{:,timeVar},curTime)));
                    lastRSCORE = curData{curRow,RSCOREVar};
                    lastRSCORE = lastRSCORE{1};
                end
                if curTime <= lastTime
                    fullCell{timeCounter,patientCounter} = lastRSCORE;
                else
                    % Check to see if the last event should be carried
                    % forward.
                    if ismember(lastRSCORE,{'CR','PD'})
                        % In this case we carry CR, PD forward unless
                        % we've specified the trial has ended via the prefix.
                        passCheck = true;
                        if length(prefixes) > 0
                            for prefixCounter = 1:length(prefixes)
                                if startsWith(patientID,prefixes{prefixCounter},'IgnoreCase',true) && (curTime > stopTimes(prefixCounter))
                                    passCheck = false;
                                end
                            end
                        end
                        if passCheck
                            fullCell{timeCounter,patientCounter} = lastRSCORE;
                        else
                            fullCell{timeCounter,patientCounter} = '.';
                        end
                    else
                        % In the case there was a dropout without hitting
                        % CR/PD treat subsequent values as missing
                        fullCell{timeCounter,patientCounter} = '.';
                    end
                end
            end
        end
        % Now we want to count...
        for timeCounter = 1 : length(allTimes)
            curData = fullCell(timeCounter,:);
            nCR = sum(ismember(curData,'CR'));
            nPR = sum(ismember(curData,'PR'));
            nSD = sum(ismember(curData,'SD'));
            nPD = sum(ismember(curData,'PD'));
            expN = nCR+nPR+nSD+nPD;
            curProbs = [nCR, nPR, nSD, nPD] / expN;
            curRow = {1, allTimes(timeCounter), 'RSCORE', interventionID, 'RSCORE','derived',myExpDataID, timeVar,PatientIDVar,TRTVar,BRSCOREVar,RSCOREVar};
            curRow = [curRow,{1, expN, curProbs(1), curProbs(2), curProbs(3), curProbs(4), nan, nan, nan, nan, nan}];
            curRow = cell2table(curRow);
            curRow.Properties.VariableNames = myRTableRECIST.Properties.VariableNames; 
            myRTableRECIST = [myRTableRECIST; curRow];    
        end   
          
    end
        
end
end
```
