## Supplementary File 3 for "A surrogate workflow for virtual population development: designing an amenable proxy for a hypothesis-test inspired nonlinear goodness-of-fit": createSubpopTable.m

```
function mySubpopTable = createSubpopTable(myWorksheet,mySubpopIDs,myTimePoints,myInterventionIDs,myElement1Names,myElement1Types,myComparators,myValues)
% This function creates a table with subpopulation information
% to be used in VPop calibration.  Note if only "myWorksheet" is provided,
% as an input argument, a subpopulation table with a single row
% for all VPs is provided.  This "all" subpopulation is included as
% the first row in all subpopulations.
%
% ARGUMENTS:
%  myWorksheet:       A worksheet with the experimental data attached
%  mySubpopIDs:       A cell array, of length N subpopulations
%                      with names for the subpopulations.  Note a subpopulation for 
%                      "all" will automatically be prefixed.  If none given,
%                      a table with just 'all' wil be generated.
%  myTimePoints:      A cell array of time points for the comparison.  For axis, 
%                      and all, the numeric value is ignored
%                      but a dummy value like nan should be entered.
%  myInterventionIDs: A cell array of cell arrays of interventionIDs, or axisIDs, 
%                      to use for each comparison.  For all, ' ' is used. 
%  myElement1Names:   A cell array of N strings with names for the
%                      variables to be used to set the subpopulations.
%                      For subpop 'all' this will be set to ' '.
%  myElement1Types:   A cell array of N strings with the types of variables to check.
%                      Should be one of:
%                      'result'  A dynamic variable to look up in the results
%                      'axis'    An axis coefficient to look up
%                      For subpop 'all' this will be set to ' '.
%  myComparators:     A cell array of of cell arrays of logical comparators entered as strings.
%                      Each string should be one of:
%                       '>' , '>=', '==', '<=', '<'. Note ' ' is used for all.
%  myValues:          A cell array of numeric values to compare against in the
%                      right hand side of the comparison.
%                     Note fixed reference values currently must be specified rather than 
%                      doing a comparison between two model variables.
%
% RETURNS
%  mySubpopTable
%


if nargin == 1
    mySubpopIDs = {};
    myTimePoints = [];
    myInterventionIDs = {};
    myElement1Names = {};
    myElement1Types = {};
    myComparators = {};
    myValues = [];
end

continueFlag = true;
commonNames = loadCommonNames();
tableVariableNamesFixed = commonNames.SUBPOPTABLEVARNAMESFIXED;
tableVariableNames = [tableVariableNamesFixed,{'vpIDs','vpIndices','predW'}];
mySubpopTable = cell2table(cell(0,length(tableVariableNames)));
mySubpopTable.Properties.VariableNames = tableVariableNames;

% Proofing of the input
% variables starts here
if continueFlag
	testN = length(mySubpopIDs);
	if sum([length(mySubpopIDs),length(myInterventionIDs),length(myElement1Names),length(myElement1Types),length(myComparators),length(myValues),length(myTimePoints)] ~= testN) > 0
		continueFlag = false;
		warning(['Not all input cell array arguments are of consistent length in ',mfilename,'.  Exiting.'])
	end
end

if continueFlag
	allInterventionIDs = getInterventionIDs(myWorksheet);
    if sum(cellfun(@(x) sum(ismember(x,allInterventionIDs)),myInterventionIDs)) < sum(cellfun(@(x) length(x),myInterventionIDs))       
		warning(['Not all interventionIDs found in worksheet in ',mfilename,'.  Exiting.'])
		continueFlag = false;
    end
	myResultIDs = myWorksheet.simProps.saveElementResultIDs;
	myAxisIDs = getAxisDefIDs(myWorksheet);
    if sum(cellfun(@(x) sum(ismember(x,[myResultIDs,myAxisIDs])),myElement1Names)) < sum(cellfun(@(x) length(x),myElement1Names))
		warning(['Not all saveElementResultIDs/axisDefIDs found in worksheet in ',mfilename,'.  Exiting.'])
		continueFlag = false;
    end		
end


if continueFlag
    %% reformat the first row
	allVPIDs = getVPIDs(myWorksheet);
    curRow = ['all',{{{nan}}},{{{}}},{{{}}}, {{{}}},{{{}}},{{{nan}}},{{allVPIDs}},{{[1:length(allVPIDs)]}},1];
	
    curRow = cell2table(curRow);
    curRow.Properties.VariableNames = tableVariableNames;
	mySubpopTable = [mySubpopTable; curRow]; 
	for rowCounter = 1 : testN     
        curRow = [mySubpopIDs{rowCounter},{myTimePoints(rowCounter)},{myInterventionIDs(rowCounter)},{myElement1Names(rowCounter)}, {myElement1Types(rowCounter)},{myComparators(rowCounter)},{myValues(rowCounter)},{{}},{{[]}},nan];
		curRow = cell2table(curRow);
        curRow.Properties.VariableNames = tableVariableNames;
		mySubpopTable = [mySubpopTable; curRow];     
	end
		
    allExpDataIDs = getExpDataIDs(myWorksheet);
    allInterventionIDs = getInterventionIDs(myWorksheet);

else
    warning(['Unable to complete ',mfilename,', exiting.'])    
	mySubpopTable = [];
end    
```
