## Supplementary File 3 for "A surrogate workflow for virtual population development: designing an amenable proxy for a hypothesis-test inspired nonlinear goodness-of-fit": evaluateCorrelations.m

```
function myCorrelationResults = evaluateCorrelations(myWorksheet,myCorrelationOptions)
% This function takes a worksheet and evaluates correlations in the output
% variables.
%
% ARGUMENTS
% myWorksheet:               an instance of a worksheet
% myCorrelationOptions:      an instance of a correlationOptions
%                            object
%
% RETURNS
% myCorrelationResults:       a structure with the fields:
%                              analyzeElementResultIDs: a 1xnVar cell array 
%                              of strings of variables to include in the
%                              analysis.
%                              analyzeTimes: a 1xnVar matrix of time values
%                              data: nVar x nVar matrix of correlations
%
myCorrelationResults = struct();
myCorrelationResults.analyzeElementResultIDs = cell(1,0);
myCorrelationResults.analyzeTimes = nan(1,0);
myCorrelationResults.data = nan(0,0);
myCorrelationResults.correlation = nan(0,0);
myCorrelationResults.pws = nan(0,1);


continueFlag = false;
if nargin > 2
    warning(['Too many input arguments to ',mfilename, '. Arguments should be: myWorksheet and myCorrelationOptions.'])
    continueFlag = false;
elseif nargin > 1
    continueFlag = true;
else
    warning(['Insufficient input arguments to ',mfilename, '. Arguments should be: myWorksheet and myCorrelationOptions.'])
    continueFlag = false;
end

if continueFlag
    allVPIDs = getVPIDs(myWorksheet);
    nVPs = length(allVPIDs);
    % If no prevalence weights are provided we assume equal weights
    % when testing for correlations
    if length(myCorrelationOptions.pws) < 1
        myCorrelationOptions.pws = ones(1,nVPs)/nVPs;
    end
    continueFlag = myCorrelationOptions.verify(myWorksheet);
end

flagAutomatedAnalyzeResultIDs = false;
flagAutomatedAnalyzeResultTimes = false;
if continueFlag
    allInterventionIDs = getInterventionIDs(myWorksheet);
    myInterventionIndex = find(ismember(allInterventionIDs,  myCorrelationOptions.interventionID));
    if length(myCorrelationOptions.analyzeElementResultIDs) < 1
        flagAutomatedAnalyzeResultIDs = true;
        testResult = myWorksheet.results{myInterventionIndex,1};
        flagAutomatedAnalyzeResultIDs = true;
        myCorrelationOptions.analyzeElementResultIDs = testResult.Names;
        timeIndex = find(ismember(myCorrelationOptions.analyzeElementResultIDs,'time'));
        myCorrelationOptions.analyzeElementResultIDs(timeIndex) = [];
    end
    if length(myCorrelationOptions.analyzeTimes) < 1
        myCorrelationOptions.analyzeTimes = ones(1,length(myCorrelationOptions.analyzeElementResultIDs))*max(myWorksheet.simProps.sampleTimes);
        flagAutomatedAnalyzeResultTimes = true;
    end
end

% if we auotmated either of these double-check the options
if flagAutomatedAnalyzeResultIDs || flagAutomatedAnalyzeResultTimes
    continueFlag = myCorrelationOptions.verify(myWorksheet);
end

if continueFlag
    % Preallocate the matrix
    nVars = length(myCorrelationOptions.analyzeElementResultIDs);
    dataMatrix = nan(nVPs,nVars);
    % We'll have to create a data matrix for each unique resultID
    uniqueAnalyzeResultIDs = unique(myCorrelationOptions.analyzeElementResultIDs);
    nGrabLoops = length(uniqueAnalyzeResultIDs);
    for grabLoopCounter = 1 : nGrabLoops
        curVarID = uniqueAnalyzeResultIDs{grabLoopCounter};
        % It may be possible to speed this up by re-writing some of this
        % without including the checks at the beginning of the call...
        % just run like this for now.
        curResult = getResultOutputforIntervention(myWorksheet, myCorrelationOptions.interventionID, curVarID);
        % We maintain the VP order in this call
        outputVarIndices = find(ismember(myCorrelationOptions.analyzeElementResultIDs,curVarID));
        for outputVarCounter = 1 : length(outputVarIndices)
            outputIndex = outputVarIndices(outputVarCounter);
            % Time is always appended to the first element
            timeIndex = find(curResult.Data(:,1) == myCorrelationOptions.analyzeTimes(outputIndex));
            dataMatrix(:,outputIndex) = (curResult.Data(timeIndex,2:end))';
        end
    end
    myCorrelationResults.analyzeElementResultIDs = myCorrelationOptions.analyzeElementResultIDs;
    myCorrelationResults.analyzeTimes = myCorrelationOptions.analyzeTimes;
    myCorrelationResults.data = dataMatrix;    
    if myCorrelationOptions.addAxis
        myCorrelationResults.data = [transpose(getVPCoeffs(myWorksheet)),myCorrelationResults.data];
        myAxisDefIDs=getAxisDefIDs(myWorksheet);
        myCorrelationResults.analyzeTimes = [zeros(1,length(myAxisDefIDs)),myCorrelationResults.analyzeTimes];
        myCorrelationResults.analyzeElementResultIDs = [myAxisDefIDs,myCorrelationResults.analyzeElementResultIDs];
    end
    myCorrelationResults.pws = myCorrelationOptions.pws;
    % Remove vars with NaNs
    nanVarIndices = find(sum(isnan(myCorrelationResults.data),1)>0);
    myCorrelationResults.data(:,nanVarIndices)=[];
    myCorrelationResults.analyzeElementResultIDs(nanVarIndices)=[];
    myCorrelationResults.analyzeTimes(nanVarIndices)=[];
    % Remove constant vars
    constantVarIndices = find(max(myCorrelationResults.data,[],1)<=min(myCorrelationResults.data,[],1));
    myCorrelationResults.data(:,constantVarIndices)=[];
    myCorrelationResults.analyzeElementResultIDs(constantVarIndices)=[];
    myCorrelationResults.analyzeTimes(constantVarIndices)=[];    
    % Remove vars that are too small (and may reflect solver issues)
    if myCorrelationOptions.minAbsValFilter > 0
        smallVarIndices = find(max(abs(myCorrelationResults.data),[],1) < myCorrelationOptions.minAbsValFilter);
        myCorrelationResults.data(:,smallVarIndices)=[];
        myCorrelationResults.analyzeElementResultIDs(smallVarIndices)=[];
        myCorrelationResults.analyzeTimes(smallVarIndices)=[];          
    end
    % Now we have pre-processed the data and can look at correlations
    myCorrelationResults.correlation = weightedcorrs(myCorrelationResults.data, myCorrelationResults.pws');
else
    warning(['Unable to run ',mfilename, '. Exiting.'])
end
end
```
