## Supplementary File 3 for "A surrogate workflow for virtual population development: designing an amenable proxy for a hypothesis-test inspired nonlinear goodness-of-fit": evaluateGOF.m

```
function myVPop = evaluateGOF(myVPop)
% Evaluate the goodness-of-fit for a given
% axis weight solution (VPop).  This has been implemented
% as a stand-alone function rather than a method.
%
% ARGUMENTS
% myVPop:         An instance of a VPop, VPopRECIST object.  Populating the tables
%                 with experiment and simulated data should be performed
%                 prior to calling this function:
%                  mnSDTable
%                  binTable
%                  distTable
%                  distTable2D
%                  corTable
%                  brTableRECIST (only if a VPopRECIST object)	
%                  rTableRECIST (only if a VPopRECIST object)									   
%
% RETURNS
% myVpop:         A VPop is returned, with the fields populated:
%                  gofMn
%                  gofSD
%                  gofBin
%                  gofDist
%                  gofDist2D
%                  gofCor
%                  gofBR (only if a VPopRECIST object)
%                  gofR (only if a VPopRECIST object)
%                  gof
%

continueFlag = false;
if nargin > 1
    warning(['Too many input arguments provided to,',mfilename,'.  Requires: myVPop.'])
    continueFlag = false;
elseif nargin > 0
    continueFlag = true;
else
    continueFlag = false;
    warning(['Insufficient input arguments provided to,',mfilename,'.  Requires: myVPop.'])
end

if continueFlag
    if ~ismember(class(myVPop),{'VPop','VPopRECIST','VPopRECISTnoBin'})
        continueFlag = false;
        warning(['Input VPop not recognized in call to ',mfilename,'.'])
    end       
end

if continueFlag
    % First, update the predicted values
    myVPop = myVPop.addPredTableVals();

    myMnSDTable = myVPop.mnSDTable;
    myBinTable = myVPop.binTable;
    myDistTable = myVPop.distTable;
	myDistTable2D = myVPop.distTable2D;
	myCorTable = myVPop.corTable;	
	if isa(myVPop,'VPopRECIST') || isa(myVPop,'VPopRECISTnoBin')
		myBRTableRECIST = myVPop.brTableRECIST;
		myRTableRECIST = myVPop.rTableRECIST;        
	end

    if ~isempty(myMnSDTable)
        % Evaluate mn sd first
        % meanResids = myMnSDTable{:,'expMean'} - myMnSDTable{:,'predMean'};
        expN = myMnSDTable{:,'expN'};
        predN = myMnSDTable{:,'predN'};
		expMn = myMnSDTable{:,'expMean'};
        expSD = myMnSDTable{:,'expSD'};
        predSD = myMnSDTable{:,'predSD'};
		predMn = myMnSDTable{:,'predMean'};
		logN = myMnSDTable{:,'logN'};
		
		% Convert lognormal summary data
		if sum(logN) > 0
			expSDBak = expSD;
			expMnBak = expMn;
			predSDBak = predSD;
			predMnBak = predMn;
			predMn(logN) = log(predMnBak(logN)./sqrt(1+(predSDBak(logN).^2)./(predMnBak(logN).^2)));
			expMn(logN) = log(expMnBak(logN)./sqrt(1+(expSDBak(logN).^2)./(expMnBak(logN).^2)));
			predSD(logN) = sqrt(log(1+(predSDBak(logN).^2)./(predMnBak(logN).^2)));
			expSD(logN) = sqrt(log(1+(expSDBak(logN).^2)./(expMnBak(logN).^2)));
		end

        % First we apply the t-test
        denominator = ( ((predN-1).*predSD.^2 + (expN-1).*expSD.^2)./(predN+expN-2) ).^.5;
        denominator = denominator.*((predN + expN)./(predN.*expN)).^.5;
        deltaMean = predMn - expMn;
        tStat = deltaMean./denominator;
        % We want the two-sided test
        meanPvals = tcdf(-1*abs(tStat),predN+expN-2);
        meanPvals = 2 * meanPvals;

        % Next we apply the F-test
        sdPvals = fcdf((predSD./expSD).^2, predN-1, expN-1);

        % We want the two-sided test result
        sdPvals = 2 * min([sdPvals, 1-sdPvals],[],2);
		
		% This can fail, for example
		% if the standard deviation is
		% not defined.
		% In that case, we set the pvalue to zero
		repPos = find(isnan(meanPvals));
		meanPvals(repPos) = 0;
		repPos = find(isnan(sdPvals));
		sdPvals(repPos) = 0;		
		
        myVPop.gofMn = meanPvals;
        myVPop.gofSD = sdPvals;
    else
        myVPop.gofMn = [];
        myVPop.gofSD = [];
    end
    if ~isempty(myBinTable)
        expN = myBinTable{:,'expN'};
        predN = myBinTable{:,'predN'};
        expBins = myBinTable{:, 'expBins'};
        predBins = myBinTable{:, 'predBins'};		
        %expBinCounts = round(repmat(expN,1,4) .* [myBinTable{:, 'expBin1'}, myBinTable{:, 'expBin2'}, myBinTable{:, 'expBin3'}, myBinTable{:, 'expBin4'}]);
        %predBinCounts = round(repmat(predN,1,4) .* [myBinTable{:, 'predBin1'}, myBinTable{:, 'predBin2'}, myBinTable{:, 'predBin3'}, myBinTable{:, 'predBin4'}]);
        [nStatRows, ~] = size(expBins);
        binPvals = nan(nStatRows,1);
        for rowCounter = 1 : nStatRows 
			curExpBins = expBins{rowCounter};
			curPredBins = predBins{rowCounter};
			[~, nCurExpBins] = size(curExpBins);
			[~, nCurPredBins] = size(curPredBins);
			expBinCounts = round(repmat(expN(rowCounter),1,nCurExpBins) .* curExpBins);
			predBinCounts = round(repmat(predN(rowCounter),1,nCurPredBins) .* curPredBins);
            % The desired test is related to a 2xN contigency table.  
            binPval = contingency2N(expBinCounts, predBinCounts,myVPop.exactFlag);
            % 3rd party "myfisher" series of tests are used if the sample size is small.
            binPvals(rowCounter) = binPval;
        end
        % Write these pvals to the VPop before returning it.
        myVPop.gofBin = binPvals;
    else
        myVPop.gofBin = [];
    end
    if ~isempty(myDistTable)
        [nStatRows, ~] = size(myDistTable);
        ksPvals = nan(nStatRows,1);
		predW = myDistTable.('predProbs');
		predInd = myDistTable.('simCombinedIndices');
		expInd = myDistTable.('expCombinedIndices');
        expN = myDistTable.('expN');
        predN = myDistTable.('predN');	
		SC = myDistTable.('combinedPoints');
        for rowCounter = 1 : nStatRows 
            % Custom KS test for weighted samples
            expW = 1./(expN(rowCounter)*ones(1,expN(rowCounter)));
            if length(SC{rowCounter} > 0) 
                ksPval = weightedKSPreGrid(SC{rowCounter}, expInd{rowCounter}, predInd{rowCounter}, expW, predW{rowCounter}, expN(rowCounter), predN(rowCounter));
                ksPvals(rowCounter) = ksPval;
            else
                ksPvals(rowCounter) = nan;
            end
        end	
        % set Nans to zero
        repPos = find(isnan(ksPvals));
		ksPvals(repPos) = 0;	
        
        % Write these pvals to the VPop before returning it.
        myVPop.gofDist = ksPvals;
    else
        myVPop.gofDist = [];
	end
    if ~isempty(myDistTable2D)
        [nStatRows, ~] = size(myDistTable2D);
        ksPvals2D = nan(nStatRows,1);
		predW = myDistTable2D.('predProbs');
		predInd = myDistTable2D.('predIndices');
        expN = myDistTable2D.('expN');
        predN = myDistTable2D.('predN');
		expSample = myDistTable2D.('expSample');
		predSample = myDistTable2D.('predSample');
        for rowCounter = 1 : nStatRows 
			expW = 1./(expN(rowCounter)*ones(1,expN(rowCounter)));
            curPredW = predW{rowCounter};
            % Custom 2DKS test for weighted samples

            sample1 = expSample{rowCounter};
            curVals = predSample{rowCounter};

            sample1KeepN = expN(rowCounter);
            [~,sample2KeepN] = size(predSample{rowCounter});            
            if ~myVPop.exactFlag
                % We'll reduce the size to speed up calculations in this
                % case.  When trying different parameters, 10 x the effN
                % gave very good estimates of P (within 10%) of the full matrix
                % value.  But 2 x was substantially faster if a 50% error in the 
                % returned p value is acceptable.

                sample2KeepN = min(max(min(sample2KeepN,2*ceil(predN(rowCounter))),3),sample2KeepN);

                [~, indices2] = sort(curPredW, 'descend');
                indices2 = indices2(1:sample2KeepN);
                curVals = curVals(:,indices2);
                curPredW = curPredW(indices2);
                curPredW = curPredW/sum(curPredW);
            end

            % These can take a while to generate but are also large
            % to store on file.  So they are generated as needed.
            % Fortunately, if most of the calculations are done
            % without the exactFlag, the comparison grid can
            % be kept smaller and these run faster.
            quadrantCounts1c = quadrantCount(sample1, curVals);
            quadrantCounts1s = quadrantCount(sample1, sample1);
            quadrantCounts2c = quadrantCount(curVals, sample1);
            quadrantCounts2s = quadrantCount(curVals, curVals); 
            if length(curVals) > 0
                ksPval = weightedKS2D(sample1, curVals, quadrantCounts1c, quadrantCounts1s, quadrantCounts2c, quadrantCounts2s, expW, curPredW, sample1KeepN, sample2KeepN);
                ksPvals2D(rowCounter) = ksPval;
            else
                ksPvals2D(rowCounter) = nan;
            end
            % set Nans to zero
            repPos = find(isnan(ksPvals2D));
            ksPvals2D(repPos) = 0;	
        
        end		
        % Write these pvals to the VPop before returning it.
        myVPop.gofDist2D = ksPvals2D;
    else
        myVPop.gofDist2D = [];
	end	
    if ~isempty(myCorTable)
        [nStatRows, ~] = size(myCorTable);
        corPvals2D = nan(nStatRows,1);
        expN = myCorTable.('expN');
        predN = myCorTable.('predN');
		expCor = myCorTable.('expCor');
		predCor = myCorTable.('predCor');
        for rowCounter = 1 : nStatRows 
			corPvals2D(rowCounter) = compare_correlation_coefficients(expCor(rowCounter),predCor(rowCounter),expN(rowCounter),predN(rowCounter));
        end		
        % Write these pvals to the VPop before returning it.
        myVPop.gofCor = corPvals2D;
    else
        myVPop.gofCor = [];
	end	
	if isa(myVPop,'VPopRECIST')
		if ~isempty(myBRTableRECIST)
			expN = myBRTableRECIST{:,'expN'};
			predN = myBRTableRECIST{:,'predN'};
			expBinCounts = round(repmat(expN,1,4) .* [myBRTableRECIST{:, 'expCR'}, myBRTableRECIST{:, 'expPR'}, myBRTableRECIST{:, 'expSD'}, myBRTableRECIST{:, 'expPD'}]);
			predBinCounts = round(repmat(predN,1,4) .* [myBRTableRECIST{:, 'predCR'}, myBRTableRECIST{:, 'predPR'}, myBRTableRECIST{:, 'predSD'}, myBRTableRECIST{:, 'predPD'}]);
			[nStatRows, nBins] = size(expBinCounts);
			binPvals = nan(nStatRows,1);
			for rowCounter = 1 : nStatRows 
				% The desired test is a 2x4 contigency table.  
				chiPval = contingency2N(expBinCounts(rowCounter,:), predBinCounts(rowCounter,:),myVPop.exactFlag);
				% A 3rd party myfisher24() is used for the small sample size table.
				binPvals(rowCounter) = chiPval;
			end
			% Write these pvals to the VPop before returning it.
			myVPop.gofBR = binPvals;
		else
			myVPop.gofBR = [];
        end
		if ~isempty(myRTableRECIST)
			expN = myRTableRECIST{:,'expN'};
			predN = myRTableRECIST{:,'predN'};
			expBinCounts = round(repmat(expN,1,4) .* [myRTableRECIST{:, 'expCR'}, myRTableRECIST{:, 'expPR'}, myRTableRECIST{:, 'expSD'}, myRTableRECIST{:, 'expPD'}]);
			predBinCounts = round(repmat(predN,1,4) .* [myRTableRECIST{:, 'predCR'}, myRTableRECIST{:, 'predPR'}, myRTableRECIST{:, 'predSD'}, myRTableRECIST{:, 'predPD'}]);
			[nStatRows, nBins] = size(expBinCounts);
			binPvals = nan(nStatRows,1);
			for rowCounter = 1 : nStatRows 
				% The desired test is a 2x4 contigency table.  
				chiPval = contingency2N(expBinCounts(rowCounter,:), predBinCounts(rowCounter,:),myVPop.exactFlag);
				% A 3rd party myfisher24() is used for the small sample size table.
				binPvals(rowCounter) = chiPval;
			end
			% Write these pvals to the VPop before returning it.
			myVPop.gofR = binPvals;
		else
			myVPop.gofR = [];
		end         
	end			   
    if isa(myVPop,'VPopRECIST')
		myVPop = compositeGOFRECIST(myVPop);
	else
		myVPop = compositeGOF(myVPop);
	end
else
    warning(['Unable to run ',mfilename,'.  Returning input.'])
end    

end
```
