## Supplementary File 3 for "A surrogate workflow for virtual population development: designing an amenable proxy for a hypothesis-test inspired nonlinear goodness-of-fit": evaluateMSE.m

```
    function myVPop = evaluateMSE(myVPop)
        % count the subpopulation and dropouts in:
        vpPrevalenceWeights=myVPop.pws';
        if isempty(myVPop.LinearProblemMatrices)
             myOptimOptions = LinearCalibrationOptions();
             myOptimOptions.cdfProbsToFit = 0.05:0.05:0.95;
             myOptimOptions.pdf2DProbsToFitN = 5;
             myOptimOptions.responseValTransformation='none';
             myOptimOptions.optimizationAlgorithm = "quadprogEffN";				
             myOptimOptions.priorPrevalenceWeightAssumption = 'specified';
             myOptimOptions.oldVPop = myVPop;              
%             myOptimOptions = LinearCalibrationOptions();
%             myOptimOptions.cdfProbsToFit = 0.05:0.05:0.95;
%             myOptimOptions.pdf2DProbsToFitN = 5;
%             myOptimOptions.responseValTransformation='none';
%             myOptimOptions.optimizationAlgorithm = "fmincon";				
%             myOptimOptions.priorPrevalenceWeightAssumption = 'specified';
%             myOptimOptions.targetEffNConstraint = 1; % just put any non-zero effN constraint to get the matrix
%             myOptimOptions.minSubWeightConstraint = 0;
                 
            Objlinear = LinearCalibration(myVPop,myVPop,'optimOptions',myOptimOptions);
            Objlinear = Objlinear.constructLinearProblemMatrices();
            myVPop.LinearProblemMatrices = Objlinear.LinearProblemMatrices;             
        end
        C = myVPop.LinearProblemMatrices.independentVarValsWeighted; 
        d = myVPop.LinearProblemMatrices.observationValsWeighted;
        SubgroupSumWeights = myVPop.LinearProblemMatrices.vpIsInSubgroup*vpPrevalenceWeights;
     %   myVPop.LinearProblemMatricesSubgroupSumWeights = SubgroupSumWeights; % don't recalculate the subgroupweight from the rescaled weights. keep it the same as oldVPop
        Cactual = (1./SubgroupSumWeights).*(C.*myVPop.LinearProblemMatrices.vpIsInSubgroup); 
        residuals = Cactual*vpPrevalenceWeights-d;
        residuals(isnan(residuals)) = 0; % added this in case there are one or less VPs in the row, mn and std will be NaNs
        N = length(d); % just fix it as the number of rows, so it is comparable between iterations
        myVPop.MSE = sum(residuals.^2)/N;
    end

 
```
