## Supplementary File 3 for "A surrogate workflow for virtual population development: designing an amenable proxy for a hypothesis-test inspired nonlinear goodness-of-fit": expandVPopEffN.m

```
function [myWorksheet, newVPop] = expandVPopEffN(myWorksheet,myExpandVPopEffNOptions,myMapelOptions, hotStartVPop)
% This function takes a worksheet, options object
% mapelOptions object, as well as a reference 
% worksheet for response type bounds,
% tries to fit a VPop. Once successful it adds
% VPs to the worksheet, then tries to fit a VPop with higher effN,
% expanding the effective N in MAPEL in through consecutive iterations.  
% Essentially, it will run MAPEL iteratively, generating VPops until it 
% finds one with an acceptable GOF, generate new VPs based on the most 
% weighted VPs, and select VPs from each most weighted "seed" VP to add.
% Worksheets and VPops are written to file as the algorithm progresses.
% After each iteration, acceptable VPop results and the new
% worksheet are written to file, before the minEffN is increased in MAPEL.
% The algorithm will keep running until it finds a VPop that satisfies the 
% target effN.
% 
% ARGUMENTS:
%  myWorksheet:             A worksheet to expand
%  myExpandVPopEffNOptions: An expandVPopEffNOptions object
%  myMapelOptions:          A mapelOptions object
%  hotStartVPop:            A VPop object, will try to hot
%                            start from this solution if provided.
%                            Leave empty or '' if no hot start desired.
%
% RETURNS:
%  myWorksheet
%  newVPop
%
%  NOTES: 
%   The algorithm
%   may encounter problems if an old suffix and
%   wsIterCounter combination are re-used, if existing 
%   VPs in the worksheet have names identical to those 
%   selected by the algorithm.
%

continueFlag = false;
if nargin > 4
    warning(['Too many input arguments to ',mfilename, '. Arguments should be: myWorksheet, myExpandVPopEffNOptions, and myMapelOptions. Optionally: hotStartVPop.'])
    continueFlag = false;
    hotStartVPop = '';    
elseif nargin > 3
    continueFlag = true;
else 
    warning(['Insufficient input arguments to ',mfilename, '. Arguments should be: myWorksheet, myExpandVPopEffNOptions, and myMapelOptions. Optionally: hotStartVPop.'])
    continueFlag = false;
end

if continueFlag
    if ~strcmp(class(myExpandVPopEffNOptions),'expandVPopEffNOptions')
        warning(['Argument myExpandVPopEffNOptions to ',mfilename, ' should be of class expandVPopEffNOptions.'])
        continueFlag = false;
    end
    if ~ismember(class(myMapelOptions),{'mapelOptions','mapelOptionsRECIST','mapelOptionsRECISTnoBin'})
        warning(['Argument myMapelOptions to ',mfilename, ' should be of class mapelOptions or mapelOptionsRECIST or mapelOptionsRECISTnobin.'])
        continueFlag = false;
    end      
end 
    
if continueFlag
    allResponseTypeIDs = getResponseTypeIDs(myWorksheet);
    if length(myExpandVPopEffNOptions.plausibleResponseTypeIDs) == 0
        myExpandVPopEffNOptions.plausibleResponseTypeIDs = allResponseTypeIDs;
        disp(['No plausibleResponseTypeIDs defined by myExpandVPopEffNOptions in call to ',mfilename, '.  Using all worksheet responseTypeIDs.'])
        continueFlag = true;
    elseif sum(ismember(myExpandVPopEffNOptions.plausibleResponseTypeIDs,allResponseTypeIDs)) < length(myExpandVPopEffNOptions.plausibleResponseTypeIDs)
        warning(['The response types for myExpandVPopEffNOptions were not all found in the worksheet provided to ',mfilename, '.'])
        continueFlag = false;
    end   
end

if continueFlag    
    
    % INITIALIZATION STARTS
    suffix = myExpandVPopEffNOptions.suffix;
    wsIterCounter = myExpandVPopEffNOptions.wsIterCounter;
    targetEffN = myExpandVPopEffNOptions.targetEffN;
    maxNewPerIter = myExpandVPopEffNOptions.maxNewPerIter;
    effNDelta = myExpandVPopEffNOptions.effNDelta;
    minPVal = myExpandVPopEffNOptions.minPVal;
    nTries = myExpandVPopEffNOptions.nTries;
    nRetries = myExpandVPopEffNOptions.nRetries;
    verbose = myExpandVPopEffNOptions.verbose;
    expandCohortSize = myExpandVPopEffNOptions.expandCohortSize;
    restartPVal = myExpandVPopEffNOptions.restartPVal;
    nVPMax = myExpandVPopEffNOptions.nVPMax;
    samplePastCompletion = myExpandVPopEffNOptions.nSamplePastCompletion;
    nClusterSpec = myExpandVPopEffNOptions.nCluster;    
    linearExpandFlag = myExpandVPopEffNOptions.linearExpandFlag;
    minPVallinear = myExpandVPopEffNOptions.minPVallinear;
    maxIterlinearExpand = myExpandVPopEffNOptions.maxIterlinearExpand;
    minEffNlinearflag = myExpandVPopEffNOptions.minEffNlinearflag;
    linearExpandCohortSize = myExpandVPopEffNOptions.linearExpandCohortSize;

    % Maximal optimizePopSize.  We'll try to use smaller to save time.
    maxOptimizePopSize = myMapelOptions.optimizePopSize;

    if linearExpandFlag
        % while in linear calibration workflow. set exactFlag=true;
        myMapelOptions.exactFlag = true;
    else
        % while in pso workflow, set these for fast optimization 
        myMapelOptions.exactFlag = false;
    end
    % Enforce using effN, since
    % we enforce using it for the VPops
    myMapelOptions.useEffN = true;
    % Also minimize pool restarts
    myMapelOptions.poolRestart=false;
    myMapelOptions.poolClose=false;
    
    % We will manually refresh the pool once at the start.
    if ~isempty(gcp('nocreate'))
        delete(gcp);
    end
    if isempty(gcp('nocreate'))
        % We will use default pool settings
        mySimulateOptions = simulateOptions;
        mySimulateOptions = checkNWorkers(mySimulateOptions);		
        myPool = parpool(mySimulateOptions.clusterID,mySimulateOptions.nWorkers,'SpmdEnabled',false);
        % addAttachedFiles(myPool,{'evaluateObjective.m','evaluateObjectiveNoBin.m'});
    end	    

    % Make sure results are present.  Here, we don't re-simulate VPs if 
    % results are present.
    mySimulateOptions.rerunExisting=false;
	mySimulateOptions.poolRestart=false;
	mySimulateOptions.poolClose=false; 
    myWorksheet = simulateWorksheet(myWorksheet,mySimulateOptions);

	% Set up a reference worksheet size and effN
	% to help with clustering decisions
    startNVPs = length(getVPIDs(myWorksheet));
    startEffN = myMapelOptions.minEffN;	
    % Basis and nVPs for clustering
	basisEffN = 0;    
	nSteps = floor(((startEffN - basisEffN) / effNDelta)); 
    
    % Also check the number of VPs from the axes and data targets
    myClusterPickOptions = clusterPickOptions;       
    myClusterPickOptions = myClusterPickOptions.setDefaultFromWorksheet(myWorksheet);
    myClusterPickOptions = myClusterPickOptions.setClusterElementFromOptions(myMapelOptions);  
    nClusterAxis = length(myClusterPickOptions.clusterAxisIDs);
    [nClusterOutput, ~] = size(myClusterPickOptions.clusterElement);
    nFromCluster = ceil(1.05*(2*(nClusterOutput+nClusterAxis)));
    
    if ~isnan(nClusterSpec)
        % In this case we try to give preference to nClusterSpec
        if nClusterSpec < nFromCluster
            if verbose
                disp(['Specified number of VPs ',num2str(nClusterSpec),' as a basis for clustering steps too small, increasing to ',num2str(nFromCluster),'.'])
            end
            basisNVPs = nFromCluster;
        else
            basisNVPs = nClusterSpec;
        end
    else
        basisNVPs = min(max(floor(startNVPs - nSteps * maxNewPerIter),nFromCluster),nVPMax-samplePastCompletion);
        if verbose
            disp(['Setting ',num2str(basisNVPs),' VPs as a basis for clustering steps.'])
        end           
    end
	curEffN = startEffN;
    
    startedSamplePastCompletionFlag = false;

    % Create a screen table
    % so we can enforce new VPs will need pass within bounds
    % set by the starting worksheet.
    myScreenTable = createScreenTable(myWorksheet, myExpandVPopEffNOptions.plausibleResponseTypeIDs, true);
    
    % Note intSeed will be incremented +1 soon after.
    if (~ismember(class(hotStartVPop),{'VPop','VPopRECIST'}))
        nVPopsFound = 0;
        if (myExpandVPopEffNOptions.useMapelIntSeed) && (myMapelOptions.intSeed > -1)
            intSeedTarget = myMapelOptions.intSeed-1;
        else
            intSeedTarget = -1;
        end
    else
        nVPopsFound = 1;
        oldVPop = hotStartVPop;
        oldVPop.useEffN = true;
        oldVPop.exactFlag = true;
        oldVPop = evaluateGOF(oldVPop);	
        if (myExpandVPopEffNOptions.useMapelIntSeed) && (myMapelOptions.intSeed > -1)    
            intSeedTarget = myMapelOptions.intSeed-1;
        else
            intSeedTarget = oldVPop.intSeed-1;
        end
    end
    % INITIALIZATION ENDS
    
    % START ITERATIONS
    % OUTER LOOP
    % Set the initial best p value to a negative
		% number to force the algorithm to record
		% and write out the first VPop calibration
		% attempt, especially to aid troubleshooting
		% if calibrations fail.
        if nVPopsFound > 0
            bestPVal = oldVPop.gof;
            curVPopEffN = 1/sum(oldVPop.pws.^2);
        else
            bestPVal = -1;
            curVPopEffN = 0;
        end
        
        wsIterCounter0 = wsIterCounter;
        maxIterlinearExpand = maxIterlinearExpand+wsIterCounter0;

    if linearExpandFlag
        disp('Start linear expansion ...');
        while ~((bestPVal >= minPVallinear && curVPopEffN >= targetEffN) || wsIterCounter >= maxIterlinearExpand)
                myTestCounter = 0;		
                curEffN = targetEffN;
                myMapelOptions.minEffN = curEffN;
            % NOW FOR A GIVEN EFFN ITERATE UNTIL WE FIND A GOOD VPOP
            % AND EXPAND THE WORKSHEET WHEN WE DO
                myTestCounter = myTestCounter + 1;
                % We check whether to reseed with myTestCounter
                % for repeatability of the sequence
                
                % to initialize the seed properly at the starting iteration
                if wsIterCounter==wsIterCounter0
                    intSeedTarget = intSeedTarget+1;
                    myMapelOptions.intSeed=intSeedTarget;
                end

                % read oldVPop or worksheet
                if (myTestCounter == 1) && (nVPopsFound > 0)                    
                    newVPop = oldVPop;
                else
                    % For consistency, on the first iteration
                    % if a VPop isn't provided we will use the
                    % myExpandVPopEffNOptions.expandRandomStart
                    % value to decide randomization status.
                    curMapelOptions = myMapelOptions;
                    if (myTestCounter == 1) 
                        curMapelOptions.randomStart = myExpandVPopEffNOptions.expandRandomStart;
                    end
                    newVPop = mapelLinearExpand(myWorksheet, curMapelOptions, []);
                    
                    curVPopEffN = 1/sum(newVPop.pws.^2);
                    curMSE = newVPop.MSE;
%                     allVPIDs = getVPIDs(myWorksheet);
%                     nVPs = length(allVPIDs);  
                  %   if nVPs <= nVPMax
                    mySaveName = ['vpop_',suffix,'_iter',num2str(wsIterCounter),'_effN',num2str(round(curVPopEffN,0)),'_pvalue',num2str(round(newVPop.gof,4)),'_MSE',num2str(round(curMSE,4)),'_curBest'];
                    saveVPop(newVPop, mySaveName); % TODO: save iter0, size is different from nVPmax. modify in the future
                  %   end
                end

                oldVPop = newVPop;

                % We will sample for more VPs if we hove not finished
                % expanding or we want to resample for more past
                % completion of the expansion
                if linearExpandCohortSize > 0

                    % Get ready to expand VPs
                    wsIterCounter = wsIterCounter+1;
                    % There should be no -1 intseeds.
                    rng(oldVPop.intSeed, 'twister');
                    lastWorksheet = myWorksheet;

                    allVPIDs = getVPIDs(myWorksheet);
                    nVPs = length(allVPIDs);

                    pwExpandCutoff = -1; % to select all VPs
                    curMaxNewPerIter = linearExpandCohortSize; % maxNewPerIter;
                    curEdgeVPs = myExpandVPopEffNOptions.expandEdgeVPs;  
                    curMaxNewPerOld = myExpandVPopEffNOptions.maxNewPerOld;
                    useScoresForNew = true;
                    [myWorksheet, newPassNames] = expandWorksheetVPsFromVPop(myWorksheet, oldVPop, myMapelOptions, suffix,wsIterCounter, curMaxNewPerIter, myScreenTable, linearExpandCohortSize, myExpandVPopEffNOptions.varyMethod, myExpandVPopEffNOptions.resampleStd, curMaxNewPerOld, curEdgeVPs, myExpandVPopEffNOptions.selectByParent, useScoresForNew, pwExpandCutoff, myExpandVPopEffNOptions.screenFunctionName, linearExpandFlag);
                    % Limit the worksheet size if desired
                    allVPIDs = getVPIDs(myWorksheet);
                    nVPs = length(allVPIDs);    
                end

              %  clear lastWorksheet;                
                % if nVP <= nVPmax, save worksheet here.
                if nVPs <= nVPMax
                    saveWorksheetAutoSplit(myWorksheet,['myWorksheet_',suffix,'_iter',num2str(wsIterCounter)]);
                end
             
                intSeedTarget = intSeedTarget+1;
                myMapelOptions.intSeed=intSeedTarget;

                % now do a linear MSE optimization, then remove extra VPs and renormalize the pws to sum=1 again
                myExtraPWs = [];                
                % This is the first iteration with the current effN
                % and we have already found VPops before
                newVPop = initializeOptionPropertiesToVPop(myMapelOptions);
                if isa(myMapelOptions,'mapelOptionsRECIST')                   
                     newVPop.recistSimFilter = createRECISTSimFilter(myWorksheet, newVPop, false);                    
                end		
                newVPop = newVPop.getSimData(myWorksheet);
                newVPop.subpopTable = updateSubpopTableVPs(newVPop.subpopTable,myWorksheet);
                if strcmpi(newVPop.pwStrategy,'bin') %ismember(newVPop.pwStrategy,'bin')
                        newVPop = newVPop.assignIndices(myWorksheet, myMapelOptions);
                        % We assume the old bin probs are compatible
                        % with the new indices assignments.  That is,
                        % there are an equal number of bins and axes.
                        newVPop.binProbs = oldVPop.binProbs;
                else
                        % Otherwise we are weighting on VPs directly.
                        % We need to assign pws for added vps, also
                        % the CoeffsTable
                        newVPop=newVPop.assignCoeffs(myWorksheet);
                        [~,nVP_nobin]=size(newVPop.coeffsTable);
                        nVP_diff=nVP_nobin-length(oldVPop.pws);
                        if nVP_diff > 0
                            % If we have more VPs to calibrate in this
                            % VPop than the last, we can provide an
                            % intelligent guess on spreading the PWs
                            % to the new VPs.
                            newVPop.pws = [oldVPop.pws,zeros(1,nVP_diff)];
                            newVPop.pws = transferPWtoChildren(newVPop,nVP_diff,0);
                            myExtraPWs = transferPWtoChildren(newVPop,nVP_diff,2);
                        else
                            % If there are the same number of VPs
                            % just use the old PWs
                            newVPop.pws = oldVPop.pws;
                        end
                end
                % TEST: IF WE ARE LOADING IN A FULLY SUCCESSFUL
                % VPOP SO WE CAN SKIP MAPEL               
                newVPop.useEffN = true;
                newVPop.exactFlag = true;
                newVPop=addTableSimVals(newVPop);                
                newVPop = evaluateGOF(newVPop);
               % newVPop.exactFlag = myMapelOptions.exactFlag;
               % curVPopEffN = 1/sum(newVPop.pws.^2);
                % We randomize here with magnitude myExpandVPopEffNOptions.expandRandomStart 
                % If the magnitude is set to infinity, we just fully randomize before "restarting"
                % If set to zero, no randomization will be applied

                if isinf(myExpandVPopEffNOptions.expandRandomStart)
                      if ismember(newVPop.pwStrategy,'direct')
                          newVPop = newVPop.startPWs(myWorksheet,true);
                      else
                          newVPop = newVPop.startProbs(true);
                      end	
                      newVPop = restartMapelLinearExpand(newVPop, 0, myExtraPWs, oldVPop);
                 else
                      newVPop = restartMapelLinearExpand(newVPop, myExpandVPopEffNOptions.expandRandomStart, myExtraPWs, oldVPop);
                 end
                % Now check the results for the current fresh start
                % iteration
                % already done this check in mapel function.
                oldVPopEffN = 1/sum(oldVPop.pws.^2);
              %  curMSE = oldVPop.MSE;
                curVPopEffN = 1/sum(newVPop.pws.^2);
                oldMSE = newVPop.MSE;

                %% remove the lowest weighted VPs from this new fit if exceed nVPMax:
                nVPs = length(newVPop.pws);  
                curEdgeVPs = myExpandVPopEffNOptions.expandEdgeVPs;  % determine whether to cluster at each step to keep edge VPs
                if nVPs > nVPMax 
                    if ~curEdgeVPs
                        disp(['Number of VPs, ',num2str(nVPs),', exceeds nVPMax, ',num2str(nVPMax),', allowed for a worksheet in ',mfilename,'.  Reducing to the max allowed based on pws.'])
                        
                        [sortpw, sortindex] = sort(newVPop.pws,'descend');
                        keepindex = sortindex(1:nVPMax);
                        myVPIDs = getVPIDs(myWorksheet);
                        keepVPIDs = myVPIDs(keepindex);
                        [myWorksheet, newVPop] = subsetWorksheetVPop(myWorksheet, newVPop, keepVPIDs, false);                            
                        saveWorksheetAutoSplit(myWorksheet,['myWorksheet_',suffix,'_iter',num2str(wsIterCounter)]);

                      %% Evaluate MSE again after removing VPover
                        newVPop.useEffN = true;
                        newVPop.exactFlag = true;
                        newVPop=addTableSimVals(newVPop);
                        newVPop = evaluateGOF(newVPop);
                        newVPop = evaluateMSE(newVPop);
                        curVPopEffN = 1/sum(newVPop.pws.^2);
                        curMSE = newVPop.MSE;

                        if verbose
                            disp(['Previous best pvalue ',num2str(max(0,bestPVal)),' with effN ',num2str(round(oldVPopEffN,0)), ' with lambda ', num2str(oldVPop.lambda),'.']) % before expansion
%                             disp(['--------------- after removing VPs to keep nVPmax cohort size -----------'])
                            disp(['Current solution effN ', num2str(curVPopEffN), ' with pvalue ',num2str(newVPop.gof),' with MSE ',num2str(newVPop.MSE), ' in ',mfilename,'.'])
                            if curMSE > oldMSE*1.01
%                                 disp(['oldMSE = ' num2str(round(oldMSE,4))]);
%                                 disp(['after removing VPs to keep VPmax cohort size, new MSE = ' num2str(round(curMSE,4))]);
                                warning(['MSE goes up more than 1%, maybe good VPs are being thrown away. could check']) % could consider to throw away VPs adaptively in the future
                            end
                        end  
                    else
                        disp(['Number of VPs, ',num2str(nVPs),', exceeds nVPMax, ',num2str(nVPMax),', allowed for a worksheet in ',mfilename,'.  Reducing to the max allowed based on clustering and pws.'])
                        lastVPIDs = getVPIDs(lastWorksheet);
                        allVPIDs = getVPIDs(myWorksheet); % Lu ADD on 4/21/2023
                        nVPover = nVPs - nVPMax;
                        % Remove the old, low weighted ones that would not
                        % be kept by clustering a reasonable minimum set.
                        % If some were parents we won't spread their weight, 
                        % in the next iteration, but that should be OK
                        % Decide now far to "reduce" the worksheet
                        myClusterPickOptions = clusterPickOptions;
                        myClusterPickOptions = myClusterPickOptions.setDefaultFromWorksheet(myWorksheet);
                        myClusterPickOptions = myClusterPickOptions.setClusterElementFromOptions(myMapelOptions);
                        myClusterPickOptions.intSeed = intSeedTarget;
                        myClusterPickOptions.edgeVPFlag = true;
                        myClusterPickOptions.nClusters = basisNVPs;
                        myClusterPickOptions.algorithm = 'auto';
                        myClusterPickOptions.distance = 'correlation';
                        myClusterPickOptions.nWorkers = mySimulateOptions.nWorkers;
                        myClusterPickOptions.poolClose = mySimulateOptions.poolClose;
                        myClusterPickOptions.poolRestart = mySimulateOptions.poolRestart;
                        myClusterPickOptions.clusterID = mySimulateOptions.clusterID;
                        myClusterPickOptions.verbose = false;
                        myMedoidResult = pickClusterVPs(myWorksheet,myClusterPickOptions);
                        clusterVPIDs = myMedoidResult.('pickedVPIDs');                        
                        
                        % keep the top VPs with highest weight from each subpopulation
                        vpIsInSubgroup=newVPop.LinearProblemMatrices.vpIsInSubgroup;
                        [uniqueSubpop,ia,ic]=unique(vpIsInSubgroup,'rows');
                        keepsubpopVPIDs = [];
                        for i = 1:size(uniqueSubpop,1)
                             SubpopVPIndices = find(uniqueSubpop(i,:)==1); % find the VPs belongs to each Subpop
                             SubpopVPIDs = allVPIDs(SubpopVPIndices);
                            [~,idx] = sort(newVPop.pws(SubpopVPIndices),'descend');
                            if ~isempty(idx)
                                keepsubpopVPIDs = [keepsubpopVPIDs, SubpopVPIDs(idx(1:min(1,length(idx))))];
                            end
                        end

                        ClusterSubpopIDs=unique([clusterVPIDs,keepsubpopVPIDs]);
                        nonClusterIdx = find(~ismember(allVPIDs,ClusterSubpopIDs));                       
%                         nonClusterIdx = find(~ismember(lastVPIDs,clusterVPIDs));

                        [~,idx] = sort(newVPop.pws(nonClusterIdx),'ascend');
                        nonClusterDiscardIDs = allVPIDs(nonClusterIdx(idx(1:nVPover)));
                        keepVPIDs = allVPIDs(~ismember(allVPIDs,nonClusterDiscardIDs));

                         [myWorksheet, newVPop] = subsetWorksheetVPop(myWorksheet, newVPop, keepVPIDs, false);
                        saveWorksheetAutoSplit(myWorksheet,['myWorksheet_',suffix,'_iter',num2str(wsIterCounter)]);
                      %% Evaluate MSE again after removing VPover
                        newVPop.useEffN = true;
                        newVPop.exactFlag = true;
                        newVPop=addTableSimVals(newVPop);
                        newVPop = evaluateGOF(newVPop);
                        newVPop = evaluateMSE(newVPop);
                        curVPopEffN = 1/sum(newVPop.pws.^2);
                        curMSE = newVPop.MSE;

                        if verbose
                            disp(['Previous best pvalue ',num2str(max(0,bestPVal)),' with effN ',num2str(round(oldVPopEffN,0)), ' with lambda ', num2str(oldVPop.lambda),'.']) % before expansion
%                             disp(['--------------- after removing VPs to keep nVPmax cohort size -----------'])
                            disp(['Current solution effN ', num2str(curVPopEffN), ' with pvalue ',num2str(newVPop.gof),' with MSE ',num2str(newVPop.MSE), ' in ',mfilename,'.'])
                            if curMSE > oldMSE*1.01
                                warning(['MSE goes up more than 1%, maybe good VPs are being thrown away. could check']) % could consider to throw away VPs adaptively in the future
                            end
                        end  
                    end
                else
                    curMSE = oldMSE;
                end
                mySaveName = ['vpop_',suffix,'_iter',num2str(wsIterCounter),'_effN',num2str(round(curVPopEffN,0)),'_pvalue',num2str(round(newVPop.gof,4)),'_MSE',num2str(round(curMSE,4)),'_curBest'];
                saveVPop(newVPop, mySaveName);
                bestPVal = newVPop.gof;
                nVPopsFound = nVPopsFound+1;
                oldVPop = newVPop;
                if (bestPVal >= minPVallinear && curVPopEffN >= targetEffN) || wsIterCounter >= maxIterlinearExpand
                    disp(['minPVallinear or maxIterlinearExpand criteria reached, Exiting linear expansion phase ...']);
                    break;
                end
        end
    end
    %% do one time pso optimization with this linear expanded VPop as initial cohort and pw guess
    if nVPopsFound > 0
        bestPVal = oldVPop.gof;
    end
    if linearExpandFlag && ((bestPVal >= minPVallinear && curVPopEffN >= targetEffN) || wsIterCounter >= maxIterlinearExpand)
        disp(['Start p-value gof pso optimization using the linear expanded VPop as the initial VP cohort ...']); 
        curEffN = targetEffN; 
        myMapelOptions.minEffN = curEffN;        
        myMapelOptions.exactFlag = false; % for pso optimization to speed up a bit
        bestPVal = oldVPop.gof;  % the VPop after removing to nVPMax and LC optimized, reached critiera and returned  
        % to initialize the seed properly at the starting iteration
        if wsIterCounter==wsIterCounter0 % only if we start from the final linear expanded VPop, we need to increase the intSeed
            intSeedTarget = intSeedTarget+1;
            myMapelOptions.intSeed=intSeedTarget;
        end
        
        nVPs = length(oldVPop.pws);  
        if nVPs > nVPMax
            %% first remove VPs to nVPMax (if different) for pso speed, and check GOF. will be used if we allow different nVPMax for linear and pso expansion phase
            if verbose
                disp(['Remove lowest weighted VPs to keep nVP = ' num2str(nVPMax)])
            end                         
             [sortpw, sortindex] = sort(oldVPop.pws,'descend');
             keepindex = sortindex(1:nVPMax);
             myVPIDs = getVPIDs(myWorksheet);
             keepVPIDs = myVPIDs(keepindex);
             [myWorksheet, newVPop] = subsetWorksheetVPop(myWorksheet, oldVPop, keepVPIDs, false);  
             %% Evaluate GOF again after removing VPover
             oldVPopEffN = 1/sum(oldVPop.pws.^2);
             newVPop.useEffN = true;
             newVPop.exactFlag = true;
             newVPop=addTableSimVals(newVPop);
             newVPop = evaluateGOF(newVPop);
             newVPop = evaluateMSE(newVPop);
             curVPopEffN = 1/sum(newVPop.pws.^2);
           %  curMSE = newVPop.MSE;
             if verbose
%                      disp(['--------------- after removing VPs to keep nVPmax cohort size -----------'])
%                      disp(['Previous best p value ',num2str(max(0,bestPVal)),' with effN ',num2str(oldVPopEffN),'.'])
                     disp(['Current solution effN ', num2str(curVPopEffN), ' with pvalue ',num2str(newVPop.gof),' with MSE ',num2str(newVPop.MSE), ' in ',mfilename,'.'])
                     if newVPop.gof < minPVallinear
                            warning(['GOF less than minPVallinear, maybe good VPs are being thrown away. could check']) % could consider to throw away VPs adaptively in the future
                     end
             end  
        else
             newVPop = oldVPop;
        end
        wsIterCounter = wsIterCounter+1;
        saveWorksheetAutoSplit(myWorksheet,['myWorksheet_',suffix,'_iter',num2str(wsIterCounter)]); % if no VP reduction, we still save again the same cohort, to ensure wsIterCounter consistency
         
%         disp(['Final adjustment of labmda to give targetEffN ...']);
%         tic;
%         newVPop = adjustLambda(newVPop,targetEffN);
%         toc;
%         curVPopEffN = 1/sum(newVPop.pws.^2);
%         curMSE=newVPop.MSE;
%         mySaveName = ['vpop_',suffix,'_iter',num2str(wsIterCounter),'_lambda',num2str(round(newVPop.lambda,2)),'_effN',num2str(round(curVPopEffN,0)), '_pvalue ',num2str(round(newVPop.gof,4)), '_MSE ',num2str(round(curMSE,4)),'_curBest'];
%         saveVPop(newVPop, mySaveName);
         
         % for reproducibility
         newVPop.useEffN = myMapelOptions.useEffN;
         intSeedTarget = intSeedTarget+1;
         myMapelOptions.intSeed = intSeedTarget;
         newVPop.intSeed = myMapelOptions.intSeed;

         newVPop.exactFlag = myMapelOptions.exactFlag; % false; % for pso to speed up
         myExtraPWs = []; 
         newVPop.minEffN = myMapelOptions.minEffN;
         if isinf(myExpandVPopEffNOptions.expandRandomStart)
              if ismember(newVPop.pwStrategy,'direct')
                     newVPop = newVPop.startPWs(myWorksheet,true);
              else
                     newVPop = newVPop.startProbs(true);
              end
              newVPop = restartMapel(newVPop, 0, myExtraPWs);
         else
              newVPop = restartMapel(newVPop, myExpandVPopEffNOptions.expandRandomStart, myExtraPWs); % linearExpandFlag=0, do pso
         end   
        % Now check the results for the current fresh start
        % iteration
        % Force the exact flag for this check.
        oldVPopEffN = 1/sum(oldVPop.pws.^2);
        newVPop.useEffN = true;
        newVPop.exactFlag = true;
        newVPop=addTableSimVals(newVPop);
        newVPop = evaluateGOF(newVPop);
        newVPop = evaluateMSE(newVPop);
        curVPopEffN = 1/sum(newVPop.pws.^2);
        curMSE=newVPop.MSE; 
        if verbose
               disp(['Previous best pvalue ',num2str(max(0,bestPVal)),' with effN ',num2str(oldVPopEffN),'.'])
               disp(['after p-value gof pso optimization on the linear expanded- VP cohort, current solution effN ', num2str(curVPopEffN), ' with pvalue ',num2str(newVPop.gof), ' with MSE ',num2str(curMSE),' in ',mfilename,'.'])
        end       
        % If the results didn't look too bad, we will restart
        if ((newVPop.gof < minPVal) && (newVPop.gof >= restartPVal) && (curVPopEffN >= curEffN))
            disp(['This pvalue not quite at acceptance, restarting MAPEL with this as initial guess.  Using myMapelOptions.randomStart value to set the magnitude of randomization.'])
            myRandomStart = myMapelOptions.randomStart;
            restartCounter = 1;
            if (newVPop.gof > bestPVal)
                % Save here in case this is the best
                mySaveName = ['vpop_',suffix,'_iter',num2str(wsIterCounter),'_effN',num2str(curEffN), '_pvalue ',num2str(round(newVPop.gof,4)), '_MSE ',num2str(round(curMSE,4)),'_curBest'];
                saveVPop(newVPop, mySaveName);
                bestPVal = newVPop.gof;   
            end								
            while (((restartCounter <= nRetries) && ~(newVPop.gof >= minPVal)) || ((newVPop.gof > (0.96 * minPVal)) && (restartCounter <= max(2*nRetries,2)) && ~(newVPop.gof >= minPVal)))
                   % We will keep trying a restart until we get an OK value or we run out of retries.  If we're really close we will let the algorithm keep iterating.
                   restartVPop = newVPop;
                   restartVPop.useEffN = myMapelOptions.useEffN;
                   restartVPop.exactFlag = false;
                   intSeedTarget = intSeedTarget+1;
                   restartVPop.intSeed = intSeedTarget;
                   % Decrease the tolerance stringency for restarting
                   restartVPop.tol = newVPop.tol*0.1;
                   %restartVPop = restartMapel(restartVPop, myRandomStart);
                   myExtraPWs = [];
                   oldVPop = newVPop;
                   restartVPop = restartMapel(restartVPop, myRandomStart, myExtraPWs);                        
                   restartVPop.tol = newVPop.tol; % why set .tol back?
                   restartVPop.useEffN = true;
                   restartVPop.exactFlag = true;
                   restartVPop = evaluateGOF(restartVPop);				
                   restartVPopEffN = 1/sum(restartVPop.pws.^2);
                   if verbose
                        disp(['Previous best pvalue ',num2str(max(0,bestPVal)),' with effN ',num2str(curEffN),'.'])
                        disp(['Current restart solution effN ', num2str(restartVPopEffN), ' with pvalue ',num2str(restartVPop.gof),' in ',mfilename,'.'])
                        disp(['Current restart iteration ', num2str(restartCounter),'.'])
                   end
                   restartCounter = restartCounter + 1;
                   if ((restartVPop.gof >= newVPop.gof) && (restartVPopEffN >= curEffN))
                            newVPop = restartVPop;
                            curVPopEffN = restartVPopEffN;
                            newVPop = evaluateMSE(newVPop);
                            curMSE=newVPop.MSE; 
                            if (newVPop.gof > bestPVal)
                                % We also do this during the restart loop so we can track progress
                                mySaveName = ['vpop_',suffix,'_iter',num2str(wsIterCounter),'_effN',num2str(curEffN), '_pvalue ',num2str(round(newVPop.gof,4)), '_MSE ',num2str(round(curMSE,4)),'_curBest'];
                                saveVPop(newVPop, mySaveName);
                                bestPVal = newVPop.gof;   
                            end
                   end
             end
        end
        if ((newVPop.gof > minPVal) && (curVPopEffN >= curEffN))
            disp(['After one p-value gof pso optimization, target VPop achieved ...']);
            % save this VPop
            % This handles the case where we found a better
            % VPop than we had but it didn't trigger a restart
            mySaveName = ['vpop_',suffix,'_iter',num2str(wsIterCounter),'_effN',num2str(curEffN), '_pvalue ',num2str(round(newVPop.gof,4)), '_MSE ',num2str(round(curMSE,4)),'_curBest'];
            saveVPop(newVPop, mySaveName);
            bestPVal = newVPop.gof;
            linearExpandFlag = 0; % directly go to SPC at curEffN; TODO: SPC would be more efficient to run on ~400 workers, but we keep it the same for now
            minEffNlinearflag = 0; 
            startEffN = curEffN; 
        else
            disp(['After one p-value gof pso optimization, target VPop not achieved ... Continue with pso expansion workflow ...']);
            linearExpandFlag = 0;
            mySaveName = ['vpop_',suffix,'_iter',num2str(wsIterCounter),'_effN',num2str(curEffN), '_pvalue ',num2str(round(newVPop.gof,4)), '_MSE ',num2str(round(curMSE,4)),'_curBest'];
            saveVPop(newVPop, mySaveName);
        end        
        oldVPop = newVPop;
    end
    
    %% pso expansion phase: could either from an oldVPop (loaded, or from linear-pso part of workflow); or from mapel starting vpop    
    if linearExpandFlag==0 % && ~((bestPVal > minPVal) && (curVPopEffN >= targetEffN))
        if minEffNlinearflag % flag: estimate minEffN from MSE fit of current VP cohort
            %% if minEffN is not given, start with a MSE with no effN constraint to determine the minEffN.
            disp(['Start pso expansion from a MSE fit determined minEffN...']);
            % if pvalue didn't reach threshold or if no linearexpansion was selected
            % do one round of MSE fitting with relaxed effN constraint first. 
            myTestCounter = 0;
           % curEffN = 1; % give a minimal non-zero constraint, so fmincon will work
            wsIterCounter = wsIterCounter+1;
            saveWorksheetAutoSplit(myWorksheet,['myWorksheet_',suffix,'_iter',num2str(wsIterCounter)]); % it is the same cohort, but we save it again, to keep wsIterCounter consistency
            
           % myMapelOptions.minEffN = curEffN;
            intSeedTarget = intSeedTarget+1;
            myMapelOptions.intSeed = intSeedTarget;
            myTestCounter = myTestCounter+1;            
            
            if (nVPopsFound > 0)
                 myExtraPWs = []; 
                 newVPop = oldVPop;
                 newVPop.minEffN = myMapelOptions.minEffN;
                 newVPop.intSeed = myMapelOptions.intSeed;
                 if isinf(myExpandVPopEffNOptions.expandRandomStart)
                      if ismember(newVPop.pwStrategy,'direct')
                             newVPop = newVPop.startPWs(myWorksheet,true);
                      else
                             newVPop = newVPop.startProbs(true);
                      end
                      newVPop = restartMapelLinearExpand(newVPop, 0, myExtraPWs, []);
                 else
                      newVPop.lambda = 0; % relax effN constraint
                      newVPop = restartMapelLinearExpand(newVPop, myExpandVPopEffNOptions.expandRandomStart, myExtraPWs, newVPop);
                 end
            else
                % For consistency, on the first iteration
                % if a VPop isn't provided we will use the
                % myExpandVPopEffNOptions.expandRandomStart
                % value to decide randomization status.
                curMapelOptions = myMapelOptions;
                if (myTestCounter == 1) 
                    curMapelOptions.randomStart = myExpandVPopEffNOptions.expandRandomStart;
                end
                newVPop = mapelLinearExpand(myWorksheet, curMapelOptions,[]);
            end
            % Now check the results for the current fresh start
            % iteration
            % Force the exact flag for this check.
            newVPop.useEffN = true;
            newVPop.exactFlag = true;
            newVPop = evaluateGOF(newVPop);
            curVPopEffN = 1/sum(newVPop.pws.^2);
            curMSE=newVPop.MSE; % is this able to derive MSE?
            if verbose
                    disp(['Linear calibration with relaxed effN constraint...']);
                    disp(['Current solution effN ', num2str(curVPopEffN), ' with pvalue ',num2str(newVPop.gof), ' with MSE ',num2str(curMSE),' in ',mfilename,'.'])
                    disp(['Current iteration ', num2str(myTestCounter),'.']);
            end
            % save this VPop
            % This handles the case where we found a better
            % VPop than we had but it didn't trigger a restart
            mySaveName = ['vpop_',suffix,'_iter',num2str(wsIterCounter),'_effN',num2str(round(curVPopEffN,0)), '_pvalue ',num2str(round(newVPop.gof,4)), '_MSE ',num2str(round(newVPop.MSE,4)),'_curBest'];
            saveVPop(newVPop, mySaveName);
            bestPVal = newVPop.gof;

            %% then use this effN as minEffN during pso fitting
         %   linearExpandFlag = 0;
            curEffN = round(curVPopEffN); 
           % myMapelOptions.minEffN = curEffN;
            nVPopsFound = nVPopsFound+1;
            oldVPop = newVPop;
            basisEffN = 0; % curEffN; % go back to effN = 0 when recluster.
        else            
            if startEffN >= 1
                curEffN = startEffN; % this will be the defined myMapelOptions.minEffN from setup script
            else
                curEffN = 1;  % effN will always be > 1. this can ensure fmincon to run
                disp(['minEffN too small, increasing it to ', num2str(curEffN), '... ']);
            end
            basisEffN = 0;
        end
        
        disp(['Start p-value gof pso expansion with minEffN = ', num2str(curEffN), ', basisEffN =', num2str(basisEffN), '... ']);        
        myMapelOptions.exactFlag = false;
        % START ITERATIONS
        % OUTER LOOP
        while (curEffN < (targetEffN + effNDelta)) || (curEffN > (targetEffN + effNDelta) && bestPVal < minPVal)  % MIGHTNEEDFIX: added to catch switch from middle
            % Set the initial best p value to a negative
            % number to force the algorithm to record
            % and write out the first VPop calibration
            % attempt, especially to aid troubleshooting
            % if calibrations fail.
            bestPVal = -1;
            myMapelOptions.minEffN = curEffN; 
            myTestCounter = 0;		

            % NOW FOR A GIVEN EFFN ITERATE UNTIL WE FIND A GOOD VPOP
            % AND EXPAND THE WORKSHEET WHEN WE DO
            while bestPVal < minPVal
                myTestCounter = myTestCounter + 1;
                % We check whether to reseed with myTestCounter
                % for repeatability of the sequence
                % use myExpandVPopEffNOptions.linearExpandFlag to identify if the starting VPop is from linear workflow or pso workflow
                if (~(myExpandVPopEffNOptions.linearExpandFlag)) || (myExpandVPopEffNOptions.linearExpandFlag && wsIterCounter>(maxIterlinearExpand+1))
                    intSeedTarget = intSeedTarget+1;
                    myMapelOptions.intSeed=intSeedTarget;
                end

                % We will try to restart with initial probabilities, if
                % available, and refresh worksheet data if this is the the first
                % iteration.  We'll also enforce the MAPEL options in
                % case they are tweaked.  Note we don't just directly restart
                % since there may be new simData in the worksheet.

                if (myTestCounter == 1) && (nVPopsFound > 0)
                    myExtraPWs = [];                
                    % This is the first iteration with the current effN
                    % and we have already found VPops before
                    newVPop = initializeOptionPropertiesToVPop(myMapelOptions);
                    if isa(myMapelOptions,'mapelOptionsRECIST')                   
                        newVPop.recistSimFilter = createRECISTSimFilter(myWorksheet, newVPop, false);                    
                    end		
                    newVPop = newVPop.getSimData(myWorksheet);
                    newVPop.subpopTable = updateSubpopTableVPs(newVPop.subpopTable,myWorksheet);
                    if ismember(newVPop.pwStrategy,'bin')
                        newVPop = newVPop.assignIndices(myWorksheet, myMapelOptions);
                        % We assume the old bin probs are compatible
                        % with the new indices assignments.  That is,
                        % there are an equal number of bins and axes.
                        newVPop.binProbs = oldVPop.binProbs;
                    else
                        % Otherwise we are weighting on VPs directly.
                        % We need to assign pws for added vps, also
                        % the CoeffsTable
                        newVPop=newVPop.assignCoeffs(myWorksheet);
                        [~,nVP_nobin]=size(newVPop.coeffsTable);
                        nVP_diff=nVP_nobin-length(oldVPop.pws);
                        if nVP_diff > 0
                            % If we have more VPs to calibrate in this
                            % VPop than the last, we can provide an
                            % intelligent guess on spreading the PWs
                            % to the new VPs.
                            newVPop.pws = [oldVPop.pws,zeros(1,nVP_diff)];
                            newVPop.pws = transferPWtoChildren(newVPop,nVP_diff,0);
                            myExtraPWs = transferPWtoChildren(newVPop,nVP_diff,2);
                        else
                            % If there are the same number of VPs
                            % just use the old PWs
                            % nVP_diff should not be < 0
                            % pws_addition=sum(ones(1,nVP_diff)*1/nVP_nobin);
                            % newVPop.pws=[oldVPop.pws*(1-pws_addition),
                            % ones(1,nVP_diff)*1/nVP_nobin]; 
                            newVPop.pws = oldVPop.pws;
                        end
                    end


                    % TEST: IF WE ARE LOADING IN A FULLY SUCCESSFUL
                    % VPOP SO WE CAN SKIP MAPEL               
                    newVPop.useEffN = true;
                    newVPop.exactFlag = true;
                    newVPop=addTableSimVals(newVPop);                
                    newVPop = evaluateGOF(newVPop);
                    newVPop.exactFlag = myMapelOptions.exactFlag; % set to false for fast calculation
                    curVPopEffN = 1/sum(newVPop.pws.^2);
                    if ~(~(newVPop.gof < minPVal) && (curVPopEffN >= curEffN))
                        % We randomize here with magnitude 
                        % myExpandVPopEffNOptions.expandRandomStart
                        % If the magnitude is set to infinity, we just
                        % fully randomize before "restarting"
                        % If set to zero, no randomization will be applied
                        if isinf(myExpandVPopEffNOptions.expandRandomStart)
                            if ismember(newVPop.pwStrategy,'direct')
                                newVPop = newVPop.startPWs(myWorksheet,true);
                            else
                                newVPop = newVPop.startProbs(true);
                            end	
                            newVPop = restartMapel(newVPop, 0, myExtraPWs);
                        else
                            newVPop = restartMapel(newVPop, myExpandVPopEffNOptions.expandRandomStart, myExtraPWs);
                        end
                    end
                else
                    % For consistency, on the first iteration
                    % if a VPop isn't provided we will use the
                    % myExpandVPopEffNOptions.expandRandomStart
                    % value to decide randomization status.
                    curMapelOptions = myMapelOptions;
                    if (myTestCounter == 1) 
                        curMapelOptions.randomStart = myExpandVPopEffNOptions.expandRandomStart;
                    end
                    newVPop = mapel(myWorksheet, curMapelOptions);
                end

                % Now check the results for the current fresh start
                % iteration
                % Force the exact flag for this check.
                newVPop.useEffN = true;
                newVPop.exactFlag = true;
                newVPop = evaluateGOF(newVPop);
                newVPop = evaluateMSE(newVPop);
                curVPopEffN = 1/sum(newVPop.pws.^2);

                if verbose
                    disp(['Previous best pvalue ',num2str(max(0,bestPVal)),' with effN ',num2str(curEffN),'.'])
                    disp(['Current solution effN ', num2str(curVPopEffN), ' with pvalue ',num2str(newVPop.gof), ' in ',mfilename,'.']) % not output MSE info
                    disp(['Current iteration ', num2str(myTestCounter),'.'])
                end

                % If the results didn't look too bad, we will restart
                if ((newVPop.gof < minPVal) && (newVPop.gof >= restartPVal) && (curVPopEffN >= curEffN))
                    disp(['This pvalue not quite at acceptance, restarting MAPEL with this as initial guess.  Using myMapelOptions.randomStart value to set the magnitude of randomization.'])
                    myRandomStart = myMapelOptions.randomStart;
                    restartCounter = 1;
                    if (newVPop.gof > bestPVal)
                        % Save here in case this is the best
                        if (curEffN==1)
                            curEffN = curEffN-1;
                        end
                        mySaveName = ['vpop_',suffix,'_iter',num2str(wsIterCounter),'_effN',num2str(curEffN),'_curBest'];
                        saveVPop(newVPop, mySaveName);
                        bestPVal = newVPop.gof;   
                    end								
                    while (((restartCounter <= nRetries) && ~(newVPop.gof ...
                          >= minPVal)) || ((newVPop.gof > (0.96 * minPVal)) ...
                          && (restartCounter <= max(2*nRetries,2)) && ~(newVPop.gof >= minPVal)))
                        % We will keep trying a restart until we get an
                        % OK value or we run out of retries.  If we're
                        % really close we will let the algorithm
                        % keep iterating.
                        restartVPop = newVPop;
                        restartVPop.useEffN = myMapelOptions.useEffN;
                        restartVPop.exactFlag = myMapelOptions.exactFlag;
                        intSeedTarget = intSeedTarget+1;
                        restartVPop.intSeed = intSeedTarget;
                        % Decrease the tolerance stringency for restarting
                        restartVPop.tol = newVPop.tol*0.1;
                        %restartVPop = restartMapel(restartVPop, myRandomStart);
                        myExtraPWs = [];
                        oldVPop = newVPop;
                        restartVPop = restartMapel(restartVPop, myRandomStart, myExtraPWs);                        
                        restartVPop.tol = newVPop.tol;
                        restartVPop.useEffN = true;
                        restartVPop.exactFlag = true;
                        restartVPop = evaluateGOF(restartVPop);				
                        restartVPopEffN = 1/sum(restartVPop.pws.^2);
                        if verbose
                            disp(['Previous best pvalue ',num2str(max(0,bestPVal)),' with effN ',num2str(curEffN),'.'])
                            disp(['Current restart solution effN ', num2str(restartVPopEffN), ' with pvalue ',num2str(restartVPop.gof),' in ',mfilename,'.'])
                            disp(['Current restart iteration ', num2str(restartCounter),'.'])
                        end
                        restartCounter = restartCounter + 1;
                        if ((restartVPop.gof >= newVPop.gof) && (restartVPopEffN >= curEffN))
                            newVPop = restartVPop;
                            curVPopEffN = restartVPopEffN;
                            if (newVPop.gof > bestPVal)
                                % We also do this during the restart loop so we can
                                % track progress
                                if (curEffN==1)
                                    curEffN = curEffN-1;
                                end
                                mySaveName = ['vpop_',suffix,'_iter',num2str(wsIterCounter),'_effN',num2str(curEffN),'_curBest'];
                                saveVPop(newVPop, mySaveName);
                                bestPVal = newVPop.gof;   
                            end
                        end
                    end
                end
                if ((newVPop.gof > bestPVal) && (curVPopEffN >= ...
                                                 curEffN))
                    % This handles the case where we found a better
                    % VPop than we had but it didn't trigger a restart
                    if (curEffN==1)
                        curEffN = curEffN-1;
                    end
                    mySaveName = ['vpop_',suffix,'_iter',num2str(wsIterCounter),'_effN',num2str(curEffN),'_curBest'];
                    saveVPop(newVPop, mySaveName);
                    bestPVal = newVPop.gof;
                end
                % If we have surpassed nTries VPop iterations without success 
                % we will try to adjust the VPs

                if (mod(myTestCounter,nTries) == 0) && (nVPopsFound > 0) && ~((newVPop.gof > minPVal) && (1/sum(newVPop.pws.^2) >= curEffN))
                    % We will try adding VPs based on the last, best
                    % VPop if the effN is not much bigger than the basis
                    if curEffN <= (basisEffN + 2*effNDelta)
                        if ((expandCohortSize > 0) && (maxNewPerIter > 0))
                            % There should be no -1 intseeds.
                            useScoresForNew = true;
                            rng(newVPop.intSeed, 'twister');
                            oldVPop = newVPop;
                            [myWorksheet, newPassNames] = expandWorksheetVPsFromVPop(myWorksheet,oldVPop, myMapelOptions,suffix,wsIterCounter, maxNewPerIter, myScreenTable, expandCohortSize, myExpandVPopEffNOptions.varyMethod, myExpandVPopEffNOptions.resampleStd, myExpandVPopEffNOptions.maxNewPerOld, myExpandVPopEffNOptions.expandEdgeVPs, myExpandVPopEffNOptions.selectByParent,useScoresForNew, 0.01, myExpandVPopEffNOptions.screenFunctionName, linearExpandFlag);
                            % We force myTestCounter to 0
                            % so we can generate a new
                            % initial VPop spreading
                            % PWs from parents to children.
                            myTestCounter = 0;
                            bestPVal = -1;
                            wsIterCounter = wsIterCounter + 1;                       
                            if verbose
                                disp(['Unable to find an acceptable VPop with initial worksheet in ',num2str(nTries),' VPop fit restarts, added ', num2str(length(newPassNames)), ' VPs to the worksheet in ',mfilename,' to start worksheet iteration ',num2str(wsIterCounter),'.'])
                            end       
                            saveWorksheetAutoSplit(myWorksheet,['myWorksheet_',suffix,'_iter',num2str(wsIterCounter)]);                         
                        end    
                    else
                        % Otherwise, if we've increased the effN a bit,
                        % we will try clustering the worksheet
                        % and continuing from a lower effN
                        if ((expandCohortSize > 0) && (maxNewPerIter > 0))
                            % Decide now far to "reduce" the worksheet
                            reduceFactor = 0;
                            % This calculation is just left here for reference
                            % as reduceFactor = 0                        
                            nSteps = floor(((curEffN - basisEffN) / effNDelta)*reduceFactor);
                            curEffN = basisEffN + nSteps * effNDelta; 
                            nVPTarget = basisNVPs + nSteps * maxNewPerIter;
                            if nVPTarget <= 0
                                nVPTarget = startNVPs;
                                curEffN = startEffN;
                            end
            
                            % Cluster on worksheet axes, data being calibrated
                            % and keeping the edge VPs
                            myClusterPickOptions = clusterPickOptions;       
                            myClusterPickOptions = myClusterPickOptions.setDefaultFromWorksheet(myWorksheet);
                            myClusterPickOptions = myClusterPickOptions.setClusterElementFromOptions(myMapelOptions);  
                            myClusterPickOptions.intSeed = intSeedTarget;
                            myClusterPickOptions.edgeVPFlag = true;
                            myClusterPickOptions.nClusters = nVPTarget;
                            myClusterPickOptions.algorithm = 'auto';
                            myClusterPickOptions.distance = 'correlation';
                            myClusterPickOptions.nWorkers = mySimulateOptions.nWorkers;
                            myClusterPickOptions.poolClose = mySimulateOptions.poolClose;
                            myClusterPickOptions.poolRestart = mySimulateOptions.poolRestart;
                            myClusterPickOptions.clusterID = mySimulateOptions.clusterID;
                            myClusterPickOptions.verbose = verbose;
                            myMedoidResult = pickClusterVPs(myWorksheet,myClusterPickOptions);
                            myVPIDs = myMedoidResult.('pickedVPIDs');
                            [myWorksheet, oldVPop] = ...
                                subsetWorksheetVPop(myWorksheet, newVPop, myVPIDs, false);
                            % We also need to update the  current
                            % effN, pValue
                            bestPVal = -1;
                            
                            if curEffN < 1 
                                curEffN = 1;  % effN will always be > 1. and this can ensure fmincon to run
                                disp(['Rescluster, basisEffN too small, increasing it to ', num2str(curEffN), '... ']);
                            end
                            
                            myMapelOptions.minEffN = curEffN;
                            wsIterCounter = wsIterCounter + 1;
                            % We reset the testCounter so
                            % we can force trying to use the current
                            % VPop PW guess from the subset
                            myTestCounter = 0;
                            if verbose
                                disp(['Unable to find an acceptable VPop with initial worksheet in ',num2str(nTries),' VPop fit restarts, clustered to ', num2str(nVPTarget), ' VPs to the worksheet in ',mfilename,' to start worksheet iteration ',num2str(wsIterCounter),' with effN ',num2str(curEffN),'.'])
                            end                            
                            saveWorksheetAutoSplit(myWorksheet,['myWorksheet_',suffix,'_iter',num2str(wsIterCounter)]);                         
                        end
                    end
                end
            end
            nVPopsFound = nVPopsFound+1;
            oldVPop = newVPop;

            % We will sample for more VPs if we hove not finished
            % expanding or we want to resample for more past
            % completion of the expansion
            if (curEffN < targetEffN) || ((curEffN >= targetEffN) && (samplePastCompletion > 0))
                if expandCohortSize > 0
                    % Get ready to expand VPs
                    wsIterCounter = wsIterCounter+1;
                    % There should be no -1 intseeds.
                    rng(newVPop.intSeed, 'twister');
                    lastWorksheet = myWorksheet;
                    useScoresForNew = true;
                    if (curEffN >= targetEffN) && (samplePastCompletion > 0)
                        % If we're continuing to resample past achieving the target effN
                        pwExpandCutoff = 1/(targetEffN*2);
                        curMaxNewPerIter = samplePastCompletion;
                        curEdgeVPs = false;
                        nHigh = sum(oldVPop.pws>=pwExpandCutoff);
                        curMaxNewPerOld = ceil(curMaxNewPerIter/nHigh);
                        useScoresForNew = false;
                        % We'll also fix the target effN to the current size
                        % on the first iteration
                        if ~startedSamplePastCompletionFlag
                            allVPIDs = getVPIDs(myWorksheet);
                            nVPs = length(allVPIDs);
                            nVPMax = min(nVPs,nVPMax);                        
                            startedSamplePastCompletionFlag = true;
                            if verbose
                                disp(['Fixing maximum number of cohort VPs for sampling past completion to ',num2str(nVPMax),'.'])
                            end
                        end
                    else
                        pwExpandCutoff = 0.01;
                        curMaxNewPerIter = maxNewPerIter;
                        curEdgeVPs = myExpandVPopEffNOptions.expandEdgeVPs;  
                        curMaxNewPerOld = myExpandVPopEffNOptions.maxNewPerOld;
                    end
                    [myWorksheet, newPassNames] = expandWorksheetVPsFromVPop(myWorksheet, oldVPop, myMapelOptions, suffix,wsIterCounter, curMaxNewPerIter, myScreenTable, expandCohortSize, myExpandVPopEffNOptions.varyMethod, myExpandVPopEffNOptions.resampleStd, curMaxNewPerOld, curEdgeVPs, myExpandVPopEffNOptions.selectByParent,useScoresForNew, pwExpandCutoff, myExpandVPopEffNOptions.screenFunctionName, linearExpandFlag);
                    % Limit the worksheet size if desired
                    allVPIDs = getVPIDs(myWorksheet);
                    nVPs = length(allVPIDs);    
                    if nVPs > nVPMax
                        if verbose
                            disp(['Number of VPs, ',num2str(nVPs),', exceeds nVPMax, ',num2str(nVPMax),', allowed for a worksheet in ',mfilename,'.  Reducing to the max allowed.'])
                        end
                        lastVPIDs = getVPIDs(lastWorksheet);
                        nVPover = nVPs - nVPMax;
                        % Remove the old, low weighted ones that would not
                        % be kept by clustering a reasonable minimum set.
                        % If some were parents we won't spread their weight, 
                        % in the next iteration, but that should be OK
                        % Decide now far to "reduce" the worksheet
                        reduceFactor = 0;
                        % +1 is needed since we increment the curEffN
                        % This calculation is just left here for reference
                        % as reduceFactor = 0
                        nSteps = floor((((curEffN - basisEffN) / effNDelta)+1)*reduceFactor);
                        nVPTarget = basisNVPs + nSteps * maxNewPerIter;
                        if nVPTarget <= 0
                            nVPTarget = startNVPs;
                        end
                        myClusterPickOptions = clusterPickOptions;
                        myClusterPickOptions = myClusterPickOptions.setDefaultFromWorksheet(lastWorksheet);
                        myClusterPickOptions = myClusterPickOptions.setClusterElementFromOptions(myMapelOptions);
                        myClusterPickOptions.intSeed = intSeedTarget;
                        myClusterPickOptions.edgeVPFlag = true;
                        myClusterPickOptions.nClusters = nVPTarget;
                        myClusterPickOptions.algorithm = 'auto';
                        myClusterPickOptions.distance = 'correlation';
                        myClusterPickOptions.nWorkers = mySimulateOptions.nWorkers;
                        myClusterPickOptions.poolClose = mySimulateOptions.poolClose;
                        myClusterPickOptions.poolRestart = mySimulateOptions.poolRestart;
                        myClusterPickOptions.clusterID = mySimulateOptions.clusterID;
                        myClusterPickOptions.verbose = false;
                        myMedoidResult = pickClusterVPs(lastWorksheet,myClusterPickOptions);
                        clusterVPIDs = myMedoidResult.('pickedVPIDs');
                        nonClusterIdx = find(~ismember(lastVPIDs,clusterVPIDs));

                        [~,idx] = sort(oldVPop.pws(nonClusterIdx),'ascend');
                        nonClusterDiscardIDs = lastVPIDs(nonClusterIdx(idx(1:nVPover)));
                        keepVPIDs = lastVPIDs(~ismember(lastVPIDs,nonClusterDiscardIDs));

                        [~, oldVPop] = subsetWorksheetVPop(lastWorksheet, oldVPop, keepVPIDs, false);
                        keepVPIDs = [keepVPIDs, allVPIDs(length(lastVPIDs) + 1 : nVPs)];
                        myWorksheet = copyWorksheet(myWorksheet, keepVPIDs);
                    end
                    clear lastWorksheet;
                    % Save the new worksheet that will be used.
                    saveWorksheetAutoSplit(myWorksheet,['myWorksheet_',suffix,'_iter',num2str(wsIterCounter)]);   
                    if verbose
                        disp(['Added ', num2str(length(newPassNames)), ' VPs to the worksheet in ',mfilename,' to start worksheet iteration ',num2str(wsIterCounter),'.'])
                    end
                end
            end

            % Increment target if needed
            % we definitely increment if we are not
            % at the target yet
            if (curEffN < targetEffN)
                curEffN = curEffN + effNDelta;
            % We will increment at target just to trigger exiting the
            % loop, only if we are not supposed to just keep sampling
            % past completion
            elseif (samplePastCompletion <= 0)
                if curEffN ==1
                    curEffN = curEffN-1;
                end
                curEffN = curEffN + effNDelta;              
            end

            % This is just here to avoid the case where effNDelta
            % could bypass the target without hitting it exactly
            % and ending the loop early
            if (samplePastCompletion > 0) && (curEffN > targetEffN)
                curEffN = targetEffN;
            end

            myMapelOptions.minEffN = curEffN;        
        end
    end
            
	% Close the pool before exiting.
	if ~isempty(gcp('nocreate'))
		delete(gcp);
    end
else
    warning(['Exiting ',mfilename, '. Returning input worksheet and hot start VPop.'])    
    newVPop = hotStartVPop;
end
end
```
