## Supplementary File 3 for "A surrogate workflow for virtual population development: designing an amenable proxy for a hypothesis-test inspired nonlinear goodness-of-fit": LinearCalibration.m

```
% Major change in LinearCalibration functions:
% matrix construction: VP mask matrix; SD and Correlation rewritten; 2Ddist matrix rewritten
% constrained least square optimization (fmincon): 2-step, 2-arm. implement in parallel. objective function rewritten
% changed in 'bestfit','bestInitials','bagging'

classdef LinearCalibration
% Performs fast linear least-squares optimization of prevalence weights for
% data in a QSPToolbox, using the 'lsqlin' or 'lsqnonneg' function. Can
% also perform bootstrapping to calculate confidence intervals, and bagging
% to reduce variability in the calibration.
%
% INPUT ARGUMENTS
% -myVPop: (VPop object) Doesn't need to include the prevalence weights.
%
% PROPERTIES
% -OptimOptions: (LinearCalibrationObtions object) Specifies optimization parameters. 
% Refer to the documentation in LinearCalibrationObtions.m
% -InputVPop: (VPop object) Equal to the 'myVPop' supplied by the user
% -LinearProblemMatrices: (struct) Contains matrices and information
% related to the weighted linear least-squares fitting problem w^(1/2)Ax=w^(1/2)b, where w is
% weights, A is the values of the independent variables, x is the
% prevalence weights we are optimizing, and b is the observation values.
% Contains the following fields:
% --independentVarVals: (matrix) Corresponds to the matrix 'A' in the linear
% problem. nObservations x nVPs
% --observationVals: (vector) Corresponds to the vector 'b' in the linear
% problem. nObservations x 1
% --observationWeights: (vector) Corresponds to the weights 'w' in the linear
% problem. nObservations x 1
% --dataGroup: (vector) The data group index of each observation (a "data
% group" is a set of non-independent observations). nObservations x 1
% --observationDescriptions: (cell array) String descriptions of the 
% different observations -- useful for debugging. nObservations x 1
% --independentVarValsWeighted: (matrix) Corresponds to the product 'w^(1/2)A' in the linear
% problem. nObservations x nVPs
% --observationValsWeighted: (vector) Corresponds to the product 'w^(1/2)b' in the linear
% problem. nObservations x 1
% -OptimalPrevalenceWeightsNormalized: (vector) Prevalence weights returned
% by the optimization, normalized so they sum to 1.
% -OptimalPrevalenceWeightsNormalizedConfidenceIntervals (matrix): 95%
% confidence intervals on 'OptimalPrevalenceWeightsNormalized' 
% -OptimizedVPop: (VPop object) 'InputVPop' with 'InputVPop.pws' set to the optimal
% prevalence weights renormalized so that they sum to 1, and with the tables
% and goodness-of-fits recalculated.
% -IgnoredObservationsDescriptions: (cell array) Descriptions of
% observations which were ignored in the optimization, typically because
% the observation is equal to zero and the fit is performed on a relative
% basis, thus the observation must be ignored so as not to divide by zero.
% -InputVPopStatistics: (struct) Residuals and root-mean-squared error
% based on the prevalence weights of the input VPop.
% -RMSE: (scalar) Root-mean-squared error from the fitting (i.e., on the
% training data set)
% -FractionRunsConverged: (scalar) Fraction of runs which converged
% !!! should make first character lower case
properties
    OptimOptions
    InputVPop
    LinearProblemMatrices
    OptimalPrevalenceWeightsNormalized
    OptimalPrevalenceWeightsNormalizedConfidenceIntervals
    OptimizedVPop
    IgnoredObservationsDescriptions
    InputVPopStatistics
    RMSE % Old definition of RMSE, might want to delete it or replace with new definition! keep it for now to maintain backward compatibility.
    FractionRunsConverged
    oldVPop % the last VPop (from last iteration), so that the linear calibration step1-subweight fitting could be fixed based on last VPop.
    MSE % actual mean squared error count all observations in, and with updated subpopulation weights
    lambda % Output from fmincon linear calibration fit, record for tracking purpose only
end

% Hidden properties:
properties (Hidden = true)
    NumVPs % number of VPs
    TimeElapsedMinutes
    LinearProblemMatricesParticular % Modified LinearProblemMatrices in cases where we want to select particular data groups, etc. This is the matrices which is actually being fitted.
    OptimizationResults % Optimization results
    CrossValidationResults
end

% Private properties:
properties (Access = private)

end


%% PUBLIC METHODS

methods
    function Obj = LinearCalibration(myVPop,varargin)
    % Constructor for initializing a LinearCalibration object.
    %
    % INPUT ARGUMENTS
    % -myVPop: (VPop object) Used for extracting experimental and
    % simulation data. The prevalence weights do not need to be
    % specified.
    % -optimOptions: (LinearCalibrationOptions object) Optional argument
    % for customizing the optimization. Specified as a name-value pair.
    % 
    
        % Parse the optional function arguments:
        p = inputParser;
        addOptional(p,'optimOptions',LinearCalibrationOptions)
        parse(p,varargin{:});
        Obj.OptimOptions = p.Results.optimOptions;
        
        % Save the user-supplied VPop:
        Obj.InputVPop = myVPop;
        
        Obj.oldVPop = Obj.OptimOptions.oldVPop; % might be empty if start the first iteration or run mapel pvalue-GOF optimization.
        if (~isempty(Obj.OptimOptions.oldVPop)) & (~isempty(Obj.OptimOptions.oldVPop.lambda))
            Obj.lambda = Obj.OptimOptions.oldVPop.lambda; % might be empty if start the first iteration or run mapel pvalue-GOF optimization.
        else
            Obj.lambda = 1; % initiate with lambda = 1, will adjust adaptively at later iterations
        end
        
        % Number of VPs:
        Obj.NumVPs = size(Obj.InputVPop.simData.Data,2);
        
        % add path to nnls function
        % Change the current folder to the folder of this m-file.
        % https://www.mathworks.com/matlabcentral/answers/72309-to-change-current-folder-to-the-folder-of-m-file#answer_82461
        dirMem = pwd();
        if(~isdeployed)
          cd(fileparts(which(mfilename)));
        end
        nnlsDir = 'nnls';
        exportFigDir = 'altmany-export_fig-b1a7288';
        if exist(nnlsDir,'dir')
            addpath(genpath(nnlsDir));
        end
        if exist(exportFigDir,'dir')
            addpath(genpath(exportFigDir));
        end
        if(~isdeployed)
        	cd(dirMem);
        end
    end
    
    function Obj = run(Obj,varargin) % Edited here in 'bestFit','bestInitials','bagging', included subpopulation and dropouts: 2 step, 2 arm optimization of weights.
    % Constructs the linear matrices, runs optimization, and constructs a
    % new VPop post-optimization.
    %    timerTotal = tic();

        % Parse the optional arguments:
        p = inputParser;
        addOptional(p,'closeParallelPoolWhenFinished',true,@islogical);
        parse(p,varargin{:});
        
        % reset the optimization results
        Obj.OptimizationResults = [];
        
        % Construct matrices:
        if isempty(Obj.LinearProblemMatrices)
            timerMatrixConstruction = tic();
            Obj = Obj.constructLinearProblemMatrices();
            Obj.TimeElapsedMinutes.matrixConstruction = toc(timerMatrixConstruction)/60;
        end
        Obj0 = Obj; % record the matrix
        % Run optimization:
        % Start timer:
     %   timerOptimization = tic();
        % Array of data group indices which we can sample from:
        dataGroups = (unique(Obj.LinearProblemMatrices.dataGroup))';
        % Number of data groups:
        nDataGroups = length(dataGroups);
                
        % Run optimization based on the specified method:
        if strcmpi(Obj.OptimOptions.method,"bestFit")
             % Run a single best-fit optimization: 
             % step1: estimate subpopulation weights from fullvpop; step2: estimate all PWs, fixing subweights based on step1 results
             % Two arms running in parallel. 1) step1 fitting from uniform initial PWs; 2) step1 from oldVPop subweights.            
            % C = Obj.LinearProblemMatrices.independentVarValsWeighted;
             % d = Obj.LinearProblemMatrices.observationValsWeighted;
             siCount = 2;
             nVPs=Obj.NumVPs;                        
             MSEs = zeros(1,siCount);
             pw2_eachsi = zeros(nVPs,siCount);
             lambdas=MSEs;         
             
        %     for si=1:siCount 
                 % parfor si=1:siCount % parfor took too long time on java.util.concurrent.LinkedBlockingQueue. 
                 % It might be the data transfer from the client to the workers. https://www.mathworks.com/matlabcentral/answers/41353-parfor-takes-a-lot-of-time-to-run
           %      if si==1
           si=1;
                     
                     Obj = Obj0;
                     vpPrevalenceWeights = Obj.InputVPop.pws';
                    
                     Obj.LinearProblemMatrices.SubgroupSumWeights = Obj.LinearProblemMatrices.vpIsInSubgroup*vpPrevalenceWeights;
                     if ~isempty(find(abs(Obj.LinearProblemMatrices.SubgroupSumWeights-1)>1e-6, 1))  % check if exists subpopulation
                        % only when there is subpopulation exists, implement 2 step optimization
                        % arm 1, step 1: fit with only fullvpop dataset, to estimate subpopulation weights for step2
                     	Obj.LinearProblemMatricesParticular = [];
                        Obj = Obj.runOptimizationfullvpop(); 
                        vpPrevalenceWeights = Obj.OptimizationResults.optimalPrevalenceWeightsNormalized;
                          
                        % arm 1, step 2: fit with full dataset: including subpopulation and dropout rows 
                        % calculate the subgroup weights based on the vpPrevalenceWeights from step 1
                        Obj.LinearProblemMatrices.SubgroupSumWeights = Obj.LinearProblemMatrices.vpIsInSubgroup*vpPrevalenceWeights;
                        Obj.LinearProblemMatricesParticular = [];
                        Obj.InputVPop.pws = vpPrevalenceWeights'; % update inputvpop.pws as initial x0 for step2 optimization
                        Obj = Obj.runOptimization();   
                        vpPrevalenceWeights=Obj.OptimizationResults.optimalPrevalenceWeightsNormalized;
                     else  
                         Obj.LinearProblemMatricesParticular = [];
                         Obj = Obj.runOptimizationfullvpop(); 
                         vpPrevalenceWeights = Obj.OptimizationResults.optimalPrevalenceWeightsNormalized;
                     end
                        % Note this MSE is from updated weights, not the same as MSEfval output from runOptimization function
                        C = Obj.LinearProblemMatricesParticular.independentVarValsWeighted; 
                        d = Obj.LinearProblemMatricesParticular.observationValsWeighted;
                        SubgroupSumWeights = Obj.LinearProblemMatrices.vpIsInSubgroup*vpPrevalenceWeights;
                        Cactual = (1./SubgroupSumWeights).*(C.*Obj.LinearProblemMatricesParticular.vpIsInSubgroup); 
                        Resnew = Cactual*vpPrevalenceWeights-d;
                        Resnew(isnan(Resnew)) = 0; % this is added in case there is one or less VP in some rows, then there will be NaNs for mean and std rows
                        pw2_eachsi(:,si) = vpPrevalenceWeights;
                        N = length(d); % just fix it as the number of rows, so it is comparable between iterations
                        Obj.MSE = sum(Resnew.^2)/N;
                        MSEs(si) = Obj.MSE;
                        lambdas(si) = Obj.lambda;
                     
           %      end
           %      if si==2
           si=2;
                     Obj = Obj0;
                     if ~isempty(Obj.oldVPop) && ~isempty(Obj.oldVPop.LinearProblemMatricesSubgroupSumWeights)
                             Obj.LinearProblemMatrices.SubgroupSumWeights = Obj.oldVPop.LinearProblemMatricesSubgroupSumWeights;
                             Obj.LinearProblemMatricesParticular = [];      

                             % in case during the vpop expansion iterations, some VPs being excluded, and there will be biomarkers (rows) with no VPs  
                             if ~isempty(Obj.oldVPop.LinearProblemMatricesobservationDescriptions)
                                 for i=1:size(Obj.oldVPop.LinearProblemMatrices.observationDescriptions,1)
                                       if ~sum(ismember(Obj.LinearProblemMatrices.observationDescriptions,Obj.oldVPop.LinearProblemMatricesobservationDescriptions{i}))
                                            Obj.LinearProblemMatrices.SubgroupSumWeights(i) = NaN;                             
                                       end
                                 end
                             end
                             Obj.LinearProblemMatrices.SubgroupSumWeights(isnan(Obj.LinearProblemMatrices.SubgroupSumWeights))=[];
                             Obj = Obj.runOptimization();   
                             vpPrevalenceWeights=Obj.OptimizationResults.optimalPrevalenceWeightsNormalized;
                             C = Obj.LinearProblemMatricesParticular.independentVarValsWeighted; 
                             d = Obj.LinearProblemMatricesParticular.observationValsWeighted;
                             SubgroupSumWeights = Obj.LinearProblemMatrices.vpIsInSubgroup*vpPrevalenceWeights;
                             Cactual = (1./SubgroupSumWeights).*(C.*Obj.LinearProblemMatricesParticular.vpIsInSubgroup); 
                             Resnew = Cactual*vpPrevalenceWeights-d;
                             Resnew(isnan(Resnew)) = 0; % this is added in case there is one or less VP in some rows, then there will be NaNs for some mean and std rows
                             pw2_eachsi(:,si) = vpPrevalenceWeights;
                             N = length(d); % just fix it as the number of rows, so it is comparable between iterations
                             Obj.MSE = sum(Resnew.^2)/N;
                             MSEs(si) = Obj.MSE;
                             lambdas(si) = Obj.lambda;
                     end
            %     end
       %     end
         
            ind = find(MSEs>0);
            if ~isempty(ind)
                MSEs = MSEs(ind);
                pw2_eachsi = pw2_eachsi(:,ind);
                index = find(MSEs==min(MSEs));
    %             MSEs % output to check which arm is selected. best if alternating arms are selected
    %             index
                Obj.MSE = MSEs(index(1));
                Obj.lambda = lambdas(index(1));
                Obj.OptimizationResults.optimalPrevalenceWeightsNormalized = pw2_eachsi(:,index(1));
                Obj.OptimizationResults.optimalPrevalenceWeightsPriorToNormalization = pw2_eachsi(:,index(1));
                Obj.LinearProblemMatrices.SubgroupSumWeights = Obj.LinearProblemMatrices.vpIsInSubgroup*Obj.OptimizationResults.optimalPrevalenceWeightsNormalized; % added on 220612
                Obj.FractionRunsConverged = 0;
            else
                index = 1;
                pw2_eachsi = pw2_eachsi(:,1);
                Obj.MSE = [];
                Obj.lambda = [];
                Obj.OptimizationResults.optimalPrevalenceWeightsNormalized = pw2_eachsi(:,index(1));
                Obj.OptimizationResults.optimalPrevalenceWeightsPriorToNormalization = pw2_eachsi(:,index(1));
                Obj.LinearProblemMatrices.SubgroupSumWeights = Obj.LinearProblemMatrices.vpIsInSubgroup*Obj.OptimizationResults.optimalPrevalenceWeightsNormalized; % added on 220612
                Obj.FractionRunsConverged = 0;
            end
            if ~any(isnan(Obj.OptimizationResults.optimalPrevalenceWeightsNormalized))
                   Obj.FractionRunsConverged = 1;
                   Obj.OptimizationResults.exitFlag = 1;
            else
                Obj.OptimizationResults.exitFlag = -1;
            end
        elseif strcmpi(Obj.OptimOptions.method,"bestFitInitials")
            % scan nInitials with effN constraint ranging around curEffN, as initial guess for pso 
            % Run a single best-fit optimization: 
             % for fast calculation, only do one step based on input pws (equal)   
             C = Obj.LinearProblemMatrices.independentVarValsWeighted;
             % d = Obj.LinearProblemMatrices.observationValsWeighted;
             nInitials = 1;
             nVPs=size(C,2);                        
             pw2_eachsi = zeros(nVPs,nInitials);
             % for fast calculation, only do one step based on input pws (equal)   
             Obj = Obj0;
                     
             vpPrevalenceWeights = Obj.InputVPop.pws';
             Obj.LinearProblemMatrices.SubgroupSumWeights = Obj.LinearProblemMatrices.vpIsInSubgroup*vpPrevalenceWeights;
             Obj.LinearProblemMatricesParticular = [];
             Obj.InputVPop.pws = vpPrevalenceWeights';
             Obj = Obj.runOptimization();   
             vpPrevalenceWeights=Obj.OptimizationResults.optimalPrevalenceWeightsNormalized;
             % Note this MSE is from updated weights, not the same as MSEfval output from runOptimization function
             C = Obj.LinearProblemMatricesParticular.independentVarValsWeighted; 
             d = Obj.LinearProblemMatricesParticular.observationValsWeighted;
             SubgroupSumWeights = Obj.LinearProblemMatrices.vpIsInSubgroup*vpPrevalenceWeights;
             Cactual = (1./SubgroupSumWeights).*(C.*Obj.LinearProblemMatricesParticular.vpIsInSubgroup); 
             Resnew = Cactual*vpPrevalenceWeights-d;
             Resnew(isnan(Resnew)) = 0;
             pw2_eachsi(:,1) = vpPrevalenceWeights;
             N = length(d); % just fix it as the number of rows, so it is comparable between iterations
             Obj.MSE = sum(Resnew.^2)/N;
             Obj.OptimizationResults.optimalPrevalenceWeightsNormalized = pw2_eachsi(:,1);
             Obj.OptimizationResults.optimalPrevalenceWeightsPriorToNormalization = pw2_eachsi(:,1);
             Obj.FractionRunsConverged = 0;
             if ~any(isnan(Obj.OptimizationResults.optimalPrevalenceWeightsNormalized))
                   Obj.FractionRunsConverged = 1;
                   Obj.OptimizationResults.exitFlag = 1;
             end
        elseif strcmpi(Obj.OptimOptions.method,"bootstrap")
        % Run a single best-fit optimization, and then calculate the
        % confidence intervals on the prevalence weights using
        % bootstrapping
            % First, run the single best-fit optimization:
            Obj = Obj.runOptimization();            
            % Specify the options for bootstrapping:
            bootstrapOptions = statset();
            bootstrapOptions.UseParallel = true;
            bootstapFunctionHandle = @(includedDataGroupsParticular) bootstrapFunction(Obj,includedDataGroupsParticular);
            
            % Calculate the confidence intervals using bootstrapping:
            Obj.OptimalPrevalenceWeightsNormalizedConfidenceIntervals = bootci(Obj.OptimOptions.nBootstrapIterations,{bootstapFunctionHandle,dataGroups},'Options',bootstrapOptions);

        elseif strcmpi(Obj.OptimOptions.method,"bagging") % need to revise
        % Use bagging to reduce the variability in the estimation of the
        % optimal prevalence weights
            
            % Create a vector of VP indices that 
            vpIndices = (1:Obj.NumVPs)';

            % Convert the fraction of VPs per iteration into the number of VPs
            nVPsPerBaggingIteration = floor(Obj.OptimOptions.fractionVPsPerBaggingIteration*Obj.NumVPs);

            % Perform the bagging
            % First, initialize a cell array to store the VP indices
            % included in each iteration:
            includedVPIndices = cell(1,Obj.OptimOptions.nBootstrapIterations);
            % Also initialize a cell array to store the optimal
            % prevalence weights calculated in each iteration:
            optimizationPrevalenceWeightsOrderedAccordingToSampledVPIndices = cell(1,Obj.OptimOptions.nBootstrapIterations);
            runConverged = zeros(1,Obj.OptimOptions.nBootstrapIterations);
            parfor iBagging = 1:Obj.OptimOptions.nBootstrapIterations
                                % In case we want to be verbose:
                % tictoc = tic();
                % disp(['** BAGGING: starting Iteration ' num2str(iBootstrap) ' of ' num2str(Obj.OptimOptions.nBootstrapIterations) ' **']);
                
                % Sample the data groups for this iteration. Like bootstrapping, the sampling is
                % performed with replacement, so that some data groups may
                % be represented more than once (in which case they would
                % be weighted more highly in the fit).
                includedDataGroups = datasample(dataGroups,nDataGroups);
                
                % Sample the VPs for this iteration, without replacement
                % (since having a VP represented more than once can create
                % issues in the fit).
                includedVPIndices{iBagging} = datasample(vpIndices,nVPsPerBaggingIteration,'Replace',false);
                
                % Rebuild the matrices based on the sampled VPs and data
                % groups:
                ObjParticular = Obj.selectFromLinearProblemMatrices('includedDataGroups',includedDataGroups,'includedVPIndices',includedVPIndices{iBagging});
                
                % Run the optimization:
                ObjParticular = ObjParticular.runOptimization();
                
                % Save the optimal prevalence weights for this iteration.
                % Note that the results for each iteration is ordered
                % according to the ordering of the sampled VP indices
                % (hence why above we also saved the ordering of the VP indices)
                optimizationPrevalenceWeightsOrderedAccordingToSampledVPIndices{iBagging} = ObjParticular.OptimizationResults.optimalPrevalenceWeightsNormalized;
                
                if ~any(isnan(ObjParticular.OptimizationResults.optimalPrevalenceWeightsNormalized))
                    runConverged(iBagging) = 1;
                end
                
                % In case we want to be verbose:
                % disp(['Time elapsed for Bagging Iteration ' num2str(iBootstrap) ' [seconds]: ' num2str(toc(tictoc))]);
            end

            Obj.FractionRunsConverged = sum(runConverged)/Obj.OptimOptions.nBootstrapIterations;
            
            % Consolidate the optimal prevalence weights into a matrix.
            % The matrix is initialized with zeros so that any VP not
            % included in an iteration is automatically assigned a
            % prevalence weight of zero for that iteration (this is what 
            % we want, so we'll keep it that way). 
            optimalPrevalenceWeightsByIteration = zeros(Obj.NumVPs,Obj.OptimOptions.nBootstrapIterations);
            for iBagging = 1:Obj.OptimOptions.nBootstrapIterations
                % The indexing of the left-hand side of the following line
                % is performed to account for the fact that the ordering of
                % the VPs is different in different iterations of the
                % bagging:
                optimalPrevalenceWeightsByIteration(includedVPIndices{iBagging},iBagging) = optimizationPrevalenceWeightsOrderedAccordingToSampledVPIndices{iBagging};
            end

            % Theory suggests that the mean of the predictions of each
            % iteration is the same as the prediction from the mean
            % prevalence weight. Thus, we can save only the mean
            % prevalence weight:
            Obj.OptimizationResults.optimalPrevalenceWeightsPriorToNormalization = nanmean(optimalPrevalenceWeightsByIteration,2);
        
            % Normalize the optimal prevalence weights:
            Obj.OptimizationResults.optimalPrevalenceWeightsNormalized = ...
                Obj.OptimizationResults.optimalPrevalenceWeightsPriorToNormalization./sum(Obj.OptimizationResults.optimalPrevalenceWeightsPriorToNormalization);
        else
            error('Unrecognized optimization method.');
        end
        % Post-optimization analysis:
        Obj.OptimalPrevalenceWeightsNormalized = Obj.OptimizationResults.optimalPrevalenceWeightsNormalized;
        %  Obj = Obj.performPostOptimizationAnalysis();
        %  Obj.RMSE = Obj.OptimizationResults.rmseConsideringOptimalPrevalenceWeightsNormalized;
        
        % delete the parallel pool unless specified not to do so
        if p.Results.closeParallelPoolWhenFinished
            poolobj = gcp('nocreate');
            delete(poolobj);
        end
        
%         Obj.TimeElapsedMinutes.optimization = toc(timerOptimization)/60;  % commented this to save time
%         
%         % Construct new VPop according to the optimal prevalence weights
%         timerVPopUpdate = tic();
          if sum(~isnan(Obj.OptimalPrevalenceWeightsNormalized))
                Obj = Obj.constructNewVPopPostOptimization();
          end
%         Obj.TimeElapsedMinutes.vPopUpdate = toc(timerVPopUpdate)/60;
%         
%         Obj.TimeElapsedMinutes.totalTime = toc(timerTotal)/60;
        
        % Nested function used in the 'bootstrap' method:
        function bestFitPrevalenceWeights = bootstrapFunction(Obj,includedDataGroupsParticular)
            % Rebuild the matrices based on the sampled data groups:
            ObjParticular = Obj.selectFromLinearProblemMatrices('includedDataGroups',includedDataGroupsParticular);
            % Run optimization:
            ObjParticular = ObjParticular.runOptimization();
            % Output the best-fit prevalence weights:
            bestFitPrevalenceWeights = ObjParticular.OptimizationResults.optimalPrevalenceWeightsNormalized;
        end
    end
    
    function Obj = runCrossValidation(Obj)
    % Runs leave-one-out cross-validation. Each iteration of the
    % cross-validation leaves out the observations comprising one data
    % group, fits the remainder of the observations, and tests the fit
    % against the observations in the data group which was left out. Thus,
    % there will be as many iterations as there are data groups.
    %
    
        % Array of data group indices which we can sample from:
        dataGroupIndices = unique(Obj.LinearProblemMatrices.dataGroup)';
        
        % Number of data groups:
        nDataGroups = length(dataGroupIndices);
        
        % The total number of observations:
        nObservations = length(Obj.LinearProblemMatrices.observationVals);
        
        % Initialize a matrix that will store the training residuals from all
        % 'nDataGroups' iterations:
        trainingResidualsSquared = nan(nObservations,nDataGroups);
        
        % Initialize a vector that will store the test residual for each
        % observation (each observation will be tested in exactly one
        % iteration -- when the corresponding data group is the one being
        % left out).
        testResidualsSquared = nan(nObservations,1);
        
        % Loop through iterations and perform cross-validation.
        for iDataGroup = 1:nDataGroups
            dataGroup = dataGroupIndices(iDataGroup);
            tictoc = tic();
            disp(['CROSS-VALIDATION: Testing Data Group Index ' num2str(iDataGroup) ' of ' num2str(nDataGroups)]);
            
            % Masking index for selecting observations in the data group:
            iMaskDataGroup = Obj.LinearProblemMatrices.dataGroup == dataGroup;
            
            % The data groups to include in the fit for this iteration:
            dataGroupsToInclude = dataGroupIndices;
            dataGroupsToInclude(iDataGroup) = [];
            
            % Isolate the observations corresponding to
            % 'dataGroupsToInclude':
            ObjParticular = Obj.selectFromLinearProblemMatrices('includedDataGroups',dataGroupsToInclude);
            ObjParticular.LinearProblemMatrices = ObjParticular.LinearProblemMatricesParticular;
            
            % Run the fit:
            ObjParticular = ObjParticular.run('closeParallelPoolWhenFinished',false);
            
            % Calculate the training residuals:
            [trainingResiduals,~,~] = ObjParticular.calculateResidualsAndRootMeanSquaredErrorForSpecifiedPWs();
            trainingResidualsSquared(~iMaskDataGroup,iDataGroup) = trainingResiduals.^2;
            
            % Calculate the test residuals:
            ObjParticular.LinearProblemMatrices = Obj.LinearProblemMatrices;
            [testResiduals,~,~] = ObjParticular.evaluateDataGroupResponse(dataGroup);
            testResidualsSquared(iMaskDataGroup) = testResiduals.^2;
            
            disp(['Time elapsed for Cross-Validating Data Group Index ' num2str(iDataGroup) ' [seconds]: ' num2str(toc(tictoc))]);
        end
        % Delete fields which don't make sense for cross-validation:
        Obj.OptimizedVPop = [];
        Obj.OptimalPrevalenceWeightsNormalized = [];
        Obj.RMSE = [];
        % Calculate the training and test RMSEs, using weighted means (note that the mean is weighted since the squared residuals are weighted with weights which sum to 1):
        % For the training residuals, we should weight each of them by
        % nDataGroups-1 since this is how many times they are represented.
        % NaN values are expected where the observation was left out in the
        % test set.
        trainingMSE = nansum(trainingResidualsSquared(:)/(nDataGroups-1));
        testMSE = sum(testResidualsSquared);
        Obj.CrossValidationResults.trainingRMSE = sqrt(trainingMSE);
        Obj.CrossValidationResults.testRMSE = sqrt(testMSE);
        
        % close the parallel pool
        poolobj = gcp('nocreate');
        delete(poolobj);
        
        % for debugging:
        % detect which data groups have smaller test residuals compared to
        % training residuals
%         trainingResidualsSquaredWeighted = trainingWeights.*trainingResidualsSquared./sum(trainingWeights(:));
%         testResidualsSquaredWeighted = testWeights.*testResidualsSquared./sum(testWeights);
%         trainingResidualsSquaredWeightedConsolidatedPerObservation = sum(trainingResidualsSquaredWeighted,2);
%         testMinusTrainingResidualsSquaredWeighted = testResidualsSquaredWeighted-trainingResidualsSquaredWeighted;
%         crossValidationResidualsTable = table((1:nObservations)',Obj.LinearProblemMatrices.observationDescriptions,Obj.LinearProblemMatrices.dataGroup,trainingResidualsSquaredWeightedConsolidatedPerObservation,testResidualsSquaredWeighted,testMinusTrainingResidualsSquaredWeighted,...
%             'VariableNames',{'observationIndex','observationDescriptions','dataGroup','trainingResidualsSquaredWeighted','testResidualsSquaredWeighted','testMinusTrainingResidualsSquaredWeighted'});
%         crossValidationResidualsSortedTable = sortrows(crossValidationResidualsTable,'testMinusTrainingResidualsSquaredWeighted');
    end  

    function resultsOut = useCrossValidationToOptimizeBagging(Obj,outputDir,varargin)
    % Guides the optimization of the fraction of VPs to be selected per
    % bagging iteration. This function will generate plots of
    % root-mean-squared error (RMSE), effective N, and run time; as a
    % function of the fraction, and will save these plots in 'outputDir'.
    %
    % INPUT ARGUMENTS
    % -outputDir: (character array) Directory where to store the results.
    % Must end with '/'.
    % -vPopToPredictAgainst: (VPop object) If an external prediction is
    % desired, supply the data to predict against using this argument.
    % Specified as a name-value pair. Default is [], in which case no
    % external prediction is performed.
    % -fractionVPsPerBaggingIterationToAnalyze: (numeric vector)Indicates
    % which fractions to analyze. This argument is optional, and is
    % specified as a name-value pair. The default is '0.05:0.05:1'.
    % -calculateGOF: (scalar boolean) Indicates whether the function should
    % calculare and return the composite goodness-of-fit calculated using
    % the QSP Toolbox. This argument is optional, and is
    % specified as a name-value pair. The default is 'false'.
    %
    % OUTPUTS
    % -resultsOut: (struct) Contains the following fields:
    % --optimizationResultsTable (table): Contains the cross-validation
    % results for each of the fractions
    % --inputVPop: (struct) Contains results from analysis of
    % 'Obj.InputVPop', in case we'd like to use them as a comparison in
    % order to judge the optimization
    % -resultsTable: (table) Results of analysis.
    %
        
        tictocMain = tic();
        
        % Parse the optional arguments:
        p = inputParser;
        addOptional(p,'vPopToPredictAgainst',[]);
        addOptional(p,'fractionVPsPerBaggingIterationToAnalyze',[0.001 0.005 0.01 0.05 0.1 0.25 0.5 0.75 1],@isnumeric);
        % addOptional(p,'fractionVPsPerBaggingIterationToAnalyze',[0.01:0.01:0.04, 0.05:0.05:1],@isnumeric);
        addOptional(p,'calculateGOF',false);
        parse(p,varargin{:});
        
        % Create the output directory if it doesn't already exist
        if ~exist(outputDir,'dir')
            mkdir(outputDir);
        end

        % Specify to use bagging:
        Obj.OptimOptions.method = "bagging";
        
        % Start a log file for this run:
        fid = fopen([outputDir 'runLog.txt'],'w');

        % The number of fractions to test:
        nFractions = length(p.Results.fractionVPsPerBaggingIterationToAnalyze);
        
        % Let's do a simple best-fit (with no bagging) run, so that we can
        % compare against it:
        ObjNoBagging = Obj;
        ObjNoBagging.OptimOptions.method = "bestFit";
        outputDirParticular = [outputDir 'noBagging/'];
        [results.noBaggingRMSE,results.noBaggingEffN,results.noBaggingGOF] = runCrossValidationWithComparisons(ObjNoBagging,outputDirParticular,p);
        
        % If adequate information is speified in the InputVPop, calculate
        % some statistics for comparison:
        if ~isempty(Obj.InputVPop.pws)
            % Effective N:
            results.inputVPopEffN.effNWholeFit = Obj.calculateEffN(Obj.InputVPop.pws);
            % RMSE:
            Obj = Obj.calculateInputVPopStatistics('vPopToPredictAgainst',p.Results.vPopToPredictAgainst,'saveFigDir',[outputDir '/inputVPopPredictions/'],'closeFigsAutomatically',true);
            results.inputVPopRMSE.rmseWholeFit = Obj.InputVPopStatistics.rootMeanSquaredError;
            if ~isempty(p.Results.vPopToPredictAgainst)
                results.inputVPopRMSE.rmsePrediction = Obj.InputVPopStatistics.predictionRootMeanSquaredError;
            end
            % Goodness of fit:
            if p.Results.calculateGOF && ~isempty(Obj.InputVPop.gof)
                results.inputVPopRMSE.gofWholeFit = Obj.InputVPop.gof;
            end
        else
            results.inputVPop = [];
        end
        
        % Loop through the specified bagging fractions, and perform
        % cross-validation for each of the freactions:
        for iFraction = 1:nFractions
            tictocParticularFraction = tic();
            
            disp(['------------- STARTING CROSS-VALIDATION FOR FRACTION INDEX ' num2str(iFraction) ' of ' num2str(nFractions) ' -------------']);
            
            % Specify the fraction VPs parameter for this iteration:
            Obj.OptimOptions.fractionVPsPerBaggingIteration = p.Results.fractionVPsPerBaggingIterationToAnalyze(iFraction);
            
            % Make a directory specific to this fraction:
            fractionVPsStr = strrep(num2str(p.Results.fractionVPsPerBaggingIterationToAnalyze(iFraction)),'.','p');
            outputDirParticular = [outputDir 'fractionIndex_' num2str(iFraction) '__fraction_' fractionVPsStr '/'];

            % run the cross-validation:
            [results.baggingRMSE(iFraction),results.baggingEffN(iFraction),results.baggingGOF(iFraction),results.baggingRunTime(iFraction)] = runCrossValidationWithComparisons(Obj,outputDirParticular,p);
            

            
            disp(['Time elapsed [minutes] for Cross-Validation Fraction Index ' num2str(iFraction) ': ' num2str(toc(tictocParticularFraction)/60)]);
        end
        for iFraction = 1:nFractions
            results.baggingRMSE(iFraction).fractionVPsPerBaggingIteration = p.Results.fractionVPsPerBaggingIterationToAnalyze(iFraction);
            results.baggingEffN(iFraction).fractionVPsPerBaggingIteration = p.Results.fractionVPsPerBaggingIterationToAnalyze(iFraction);
            results.baggingGOF(iFraction).fractionVPsPerBaggingIteration = p.Results.fractionVPsPerBaggingIterationToAnalyze(iFraction);
            results.baggingRunTime(iFraction).fractionVPsPerBaggingIteration = p.Results.fractionVPsPerBaggingIterationToAnalyze(iFraction);
        end

        % Close the parallel pool
        poolobj = gcp('nocreate');
        delete(poolobj);
        
        % Put the results into a table
        resultsFieldNamesTable = fields(results);
        for iFieldTable = 1:length(resultsFieldNamesTable)
            resultsFieldNamesCols = fields(results.(resultsFieldNamesTable{iFieldTable}));
            resultsOut.(resultsFieldNamesTable{iFieldTable}) = table();
            for iFieldCols = 1:length(resultsFieldNamesCols)
                resultsTableParticular = ...
                    table([results.(resultsFieldNamesTable{iFieldTable}).(resultsFieldNamesCols{iFieldCols})]','VariableNames',resultsFieldNamesCols(iFieldCols));
                resultsOut.(resultsFieldNamesTable{iFieldTable}) = [resultsOut.(resultsFieldNamesTable{iFieldTable}) resultsTableParticular];
            end
        end

        % Save the results to the output directory
        save([outputDir 'results.mat'],'resultsOut');
        
        % Generate the plots
        generateAndSaveCrossValidationPlots(outputDir,resultsOut,p);
        
        totalTimeElapsedHours = toc(tictocMain)/3600;
        disp(['Total time [hours] elapsed for the bagging optimization: ' num2str(totalTimeElapsedHours)]);
        fprintf(fid,'Total time [hours] elapsed for the bagging optimization: %f.\n\n', totalTimeElapsedHours);

        % internal function for running cross-validation:
        function [resultsRMSE,resultsEffN,resultsGOF,resultsRunTime] = runCrossValidationWithComparisons(Obj,outputDir,p)
            
            if ~exist(outputDir,'dir')
                mkdir(outputDir);
            end
            
            if isempty(gcp('nocreate'))
                parpool();
            end
            
            % First, run a fit on all of the data
            tictocRun = tic();
            Obj = Obj.run('closeParallelPoolWhenFinished',false);
            resultsRunTime.calibrationTimeForSingleNonCVRunSeconds = toc(tictocRun);
            resultsRMSE.rmseWholeFit = Obj.RMSE;
            resultsEffN.effNWholeFit = Obj.calculateEffN(Obj.OptimizedVPop.pws);
            resultsGOF.gofWholeFit = Obj.OptimizedVPop.gof;
            
            % Run the cross-validation:
            linearCalibrationObjectCrossValidation = Obj.runCrossValidation();
            resultsRMSE.rmseTraining = linearCalibrationObjectCrossValidation.CrossValidationResults.trainingRMSE;
            resultsRMSE.rmseTest = linearCalibrationObjectCrossValidation.CrossValidationResults.testRMSE;
            
            % for debugging:
%             if results.rmse.test < results.rmse.training
%                 nan;
%             end
            
            % Predict against the combo, using the whole-fit calibration
            if ~isempty(p.Results.vPopToPredictAgainst)
                outputDirPrediction = outputDir;
                resultsRMSE.rmsePrediction = Obj.predictAgainstSpecifiedVPop(p.Results.vPopToPredictAgainst,'saveFigDir',outputDirPrediction,'closeFigsAutomatically',true,'figFileNamePrefix','predictionVPop');
            end

            % Save the objects:
            save([outputDir 'linearCalibrationObject.mat'],'Obj');
            save([outputDir 'linearCalibrationObjectCrossValidation.mat'],'linearCalibrationObjectCrossValidation');

            % Generate plots of the fits for this fraction:
            Obj.plotInputVPopFits('saveFigDir',outputDir,'closeFigsAutomatically',true);
            Obj.plotOptimizedVPopFits('saveFigDir',outputDir,'closeFigsAutomatically',true);
            Obj.plotScatterOfOptimVsInputPrevalenceWeights('saveFigDir',outputDir,'closeFigsAutomatically',true);
            
        end
    
    end
    
    function rmse = predictAgainstSpecifiedVPop(Obj,myVPop,varargin)
    % Calculates the root-mean-squared error (RMSE) for the model's prediction of
    % the data specified in a VPop. This function does not perform any fit;
    % it only assesses its prediction according to the optimal prevalence
    % weights. In addition to calculating the RMSE, this function will also
    % generate and save figures of the prediction.
    %
    % INPUT ARGUMENTS
    % -myVPop (VPop object): Contains the data against which the
    % predictions will be tested.
    % -saveFigDir: (string, optional) Directory where to save figures. Does not
    % include the filename (the filename is generated automatically). But
    % it must include a slash at the end of the string. Default is '',
    % which generates the figure in the working directory. Specified as a
    % name-value pair.
    % -closeFigsAutomatically: (boolean, optional) Specifies whether the figures
    % should be automatically closed after they are generated. Default is
    % 'false'. Specified as a name-value pair.
    % -figFileNamePrefix: (character vector) prefix of saved figure
    % filename. Default is 'optimizedVPop'.
    % 
    
        % Our approach will be to create a new object ('ObjNewVPop') corresponding to
        % 'myVPop' and construct the matrices. Intead of fitting the
        % constructed matrices, we set the prevalence weights equal to the
        % ones in 'Obj'. Then, we calculate the RMSE and produce figures of
        % the "fit" of the prediction.
        
        % Instantiate an object corresponding to the specified VPop:
        ObjNewVPop = LinearCalibration(myVPop,'optimOptions',Obj.OptimOptions);
        
        % Construct the matrices:
        ObjNewVPop = ObjNewVPop.constructLinearProblemMatrices();
        
        % Bypass the optimization:
        ObjNewVPop.OptimalPrevalenceWeightsNormalized = Obj.OptimalPrevalenceWeightsNormalized;
        
        % Calculate the RMSE:
        [~,rmse,~] = ObjNewVPop.calculateResidualsAndRootMeanSquaredErrorForSpecifiedPWs();
        
        % Below, we use function that were initially developped to be run
        % post-optimization, but we are using them here to assess the
        % prediction (since we bypassed the optimization above):
        ObjNewVPop = ObjNewVPop.constructNewVPopPostOptimization();
        ObjNewVPop.plotOptimizedVPopFits(varargin{:});
    end
    
    function [residuals,rmse,weights] = evaluateDataGroupResponse(Obj,dataGroup)
    % Calculates the residuals and RMSE for the specified dataGroup
    %
    % INPUT ARGUMENTS
    % -dataGroup: (scalar)
    %
        
        % Isolate the specified data group from the linear problem
        % matrices, and then calculate the statistics:
        ObjParticular = Obj.selectFromLinearProblemMatrices('includedDataGroups',dataGroup);
        [residuals,rmse,weights] = ...
            ObjParticular.calculateResidualsAndRootMeanSquaredErrorForSpecifiedPWs();
    end
    
    
    function Obj = calculateInputVPopStatistics(Obj,varargin)
    % Computes the residuals and root-mean-squared error for the input VPop
    % prevalence weights
    
        % Parse the optional arguments:
        p = inputParser;
        addOptional(p,'vPopToPredictAgainst',[]);
        addOptional(p,'saveFigDir','inputVPopPredictions/');
        addOptional(p,'closeFigsAutomatically',false);
        parse(p,varargin{:});
    
        % Instantiate an object corresponding to the Input VPop:
        ObjInputVPop = LinearCalibration(Obj.InputVPop,'optimOptions',Obj.OptimOptions);
        
        ObjInputVPop.OptimalPrevalenceWeightsNormalized = Obj.InputVPop.pws;
        
        % Construct the matrices:
        if ~isempty(Obj.LinearProblemMatrices)
            ObjInputVPop.LinearProblemMatrices = Obj.LinearProblemMatrices;
        else
            ObjInputVPop = ObjInputVPop.constructLinearProblemMatrices();
        end
        
        % calculate residuals and RMSE:
        [Obj.InputVPopStatistics.residuals,Obj.InputVPopStatistics.rootMeanSquaredError,~] = ...
            ObjInputVPop.calculateResidualsAndRootMeanSquaredErrorForSpecifiedPWs();
        
        % If indicated, perform prediction against the specified VPop:
        if ~isempty(p.Results.vPopToPredictAgainst)
            Obj.InputVPopStatistics.predictionRootMeanSquaredError = ...
                ObjInputVPop.predictAgainstSpecifiedVPop(p.Results.vPopToPredictAgainst,'saveFigDir',p.Results.saveFigDir,'closeFigsAutomatically',p.Results.closeFigsAutomatically,'figFileNamePrefix','predictionVPop');
        end
    end
    
    function plotInputVPopFits(Obj,varargin)
    % Figures of input VPop fits
    %
    % INPUT ARGUMENTS
    % -saveFigDir: (string, optional) Directory where to save figures. Does not
    % include the filename (the filename is generated automatically). But
    % it must include a slash at the end of the string. Default is '',
    % which generates the figure in the working directory. Specified as a
    % name-value pair.
    % -closeFigsAutomatically: (boolean, optional) Specifies whether the figures
    % should be automatically closed after they are generated. Default is
    % 'false'. Specified as a name-value pair.
    % -figFileNamePrefix: (character vector) prefix of saved figure
    % filename. Default is 'inputVPop'.
    %
    
        % Parse the optional arguments:
        p = inputParser;
        addOptional(p,'saveFigDir','');
        addOptional(p,'closeFigsAutomatically',false);
        addOptional(p,'figFileNamePrefix','inputVPop');
        parse(p,varargin{:});

        figFilepathPrefix = [p.Results.saveFigDir p.Results.figFileNamePrefix];
        if ~isempty(p.Results.saveFigDir) && ~exist(p.Results.saveFigDir,'dir')
            mkdir(p.Results.saveFigDir);
        end
        createVPopPlots(Obj.InputVPop,p.Results.figFileNamePrefix,figFilepathPrefix,p.Results.closeFigsAutomatically)
    end
    
    function plotOptimizedVPopFits(Obj,varargin)
    % Figures of new (optimized) VPop fits
    %
    % INPUT ARGUMENTS
    % -saveFigDir: (string, optional) Directory where to save figures. Does not
    % include the filename (the filename is generated automatically). But
    % it must include a slash at the end of the string. Default is '',
    % which generates the figure in the working directory. Specified as a
    % name-value pair.
    % -closeFigsAutomatically: (boolean, optional) Specifies whether the figures
    % should be automatically closed after they are generated. Default is
    % 'false'. Specified as a name-value pair.
    % -figFileNamePrefix: (character vector) prefix of saved figure
    % filename. Default is 'optimizedVPop'.
    %
    
        % Parse the optional arguments:
        p = inputParser;
        addOptional(p,'saveFigDir','');
        addOptional(p,'closeFigsAutomatically',false);
        addOptional(p,'figFileNamePrefix','optimizedVPop');
        parse(p,varargin{:});
    
        figFilepathPrefix = [p.Results.saveFigDir p.Results.figFileNamePrefix];
        if ~isempty(p.Results.saveFigDir) && ~exist(p.Results.saveFigDir,'dir')
            mkdir(p.Results.saveFigDir);
        end
        createVPopPlots(Obj.OptimizedVPop,p.Results.figFileNamePrefix,figFilepathPrefix,p.Results.closeFigsAutomatically)
    end
    
    function plotScatterOfOptimVsInputPrevalenceWeights(Obj,varargin)
	% Scatter plot of new vs old prevalence weights
    %
    % INPUT ARGUMENTS
    % -saveFigDir: (string, optional) Directory where to save figures. Does not
    % include the filename (the filename is generated automatically). But
    % it must include a slash at the end of the string. Default is '',
    % which generates the figure in the working directory. Specified as a
    % name-value pair.
    % -closeFigsAutomatically: (boolean, optional) Specifies whether the figures
    % should be automatically closed after they are generated. Default is
    % 'false'. Specified as a name-value pair.
    % -figFileNamePrefix: (character vector) prefix of saved figure
    % filename. Default is 'prevalenceWeightsNewVsOldScatterPlot'.
    %
        
        % Parse the optional arguments:
        p = inputParser;
        addOptional(p,'saveFigDir','');
        addOptional(p,'closeFigsAutomatically',false);
        addOptional(p,'figFileNamePrefix','prevalenceWeightsNewVsOldScatterPlot');
        parse(p,varargin{:});
    
        if ~isempty(p.Results.saveFigDir) && ~exist(p.Results.saveFigDir,'dir')
            mkdir(p.Results.saveFigDir);
        end
        figHandle = figure('Name','prevalence weight optimal vs input');
        scatter(Obj.InputVPop.pws,Obj.OptimizedVPop.pws,'LineWidth',1.5);
        xlabel('prevalence weights in input VPop');
        ylabel('optimal prevalence weights');
        ax = gca();
        ax.FontSize = 20;
        ax.XGrid = 'on';
        ax.YGrid = 'on';
        linearCalibrationSaveFig(figHandle,[p.Results.saveFigDir p.Results.figFileNamePrefix]);
        if p.Results.closeFigsAutomatically
            close(figHandle);
        end
    end
    
    function plotPrevalenceWeightConfidenceIntervals(Obj,varargin)
    % Plots the confidence intervals of the prevalence weights.
    % 
    % INPUT ARGUMENTS
    % -saveFigDir: (string, optional) Directory where to save figures. Does not
    % include the filename (the filename is generated automatically). But
    % it must include a slash at the end of the string. Default is '',
    % which generates the figure in the working directory. Specified as a
    % name-value pair.
    % -closeFigsAutomatically: (boolean, optional) Specifies whether the figures
    % should be automatically closed after they are generated. Default is
    % 'false'. Specified as a name-value pair.
    
        % Parse the optional arguments:
        p = inputParser;
        addOptional(p,'saveFigDir','');
        addOptional(p,'closeFigsAutomatically',false);
        parse(p,varargin{:});
    
        % Make plot of confidence intervals:
        if ~isempty(p.Results.saveFigDir) && ~exist(p.Results.saveFigDir,'dir')
            mkdir(p.Results.saveFigDir);
        end
        vpIndices = 1:Obj.NumVPs;
        figHandle = figure;
        lineObjects = plot([vpIndices;vpIndices],Obj.OptimalPrevalenceWeightsNormalizedConfidenceIntervals);
        lineColors = [lineObjects(:).Color];
        lineColors = (reshape(lineColors,[3 Obj.NumVPs]))';
        hold on;
        scatter(vpIndices,Obj.OptimalPrevalenceWeightsNormalized,[],lineColors);
        ax = gca();
        ax.FontSize = 20;
%         ax.YScale = 'log';
        ax.YGrid = 'on';
        xlabel('VP index');
        ylabel('optimal prevalence weight and 95% CI');
        xlimPadding = 40;
        xlim([-xlimPadding Obj.NumVPs+xlimPadding]);
        linearCalibrationSaveFig(figHandle,[p.Results.saveFigDir 'prevalenceWeightConfidenceIntervals']);
        if p.Results.closeFigsAutomatically
            close(figHandle);
        end
        
        figHandle = figure;
        ciWidth = Obj.OptimalPrevalenceWeightsNormalizedConfidenceIntervals(2,:) - Obj.OptimalPrevalenceWeightsNormalizedConfidenceIntervals(1,:);
        scatter(Obj.OptimalPrevalenceWeightsNormalized',ciWidth);
        ax = gca();
        ax.FontSize = 20;
        ax.XScale = 'log';
%         ax.YScale = 'log';
        ax.YGrid = 'on';
        xlabel('optimal prevalence weight');
        ylabel('width of 95% CI');
        linearCalibrationSaveFig(figHandle,[p.Results.saveFigDir 'prevalenceWeightConfidenceIntervalWidthsVsEstimate']);
        if p.Results.closeFigsAutomatically
            close(figHandle);
        end
    end

end


%% STATIC METHODS

methods (Static)
    function effN = calculateEffN(prevalenceWeights)
        if abs(sum(prevalenceWeights)-1) > 1e-8 % 1e-12
            % OptimalityTolerance:	Termination tolerance on the first-order optimality, a positive scalar. The default is 1e-8
            error('Prevalence weights must sum to 1.');
        end
     %   effN = 2^(-nansum(prevalenceWeights.*log2(prevalenceWeights)));
         effN = 1/sum(prevalenceWeights.^2); % use the effN defined from importance weighting. consistent with GOF method
    end
end

%% HIDDEN METHODS

methods (Hidden = true)
    
   %% updated, dropoff, subpopulation counted, need to revise objective function
function Obj = constructLinearProblemMatrices(Obj)
%     % Constructs the matrices for the linearized fitting problem.        
        % The prevalence weights are required to renormalize VP prevalence weights
        % in cases where some VP simulation values are NaN since they dropped off
        % therapy. With VPs missing, the prevalence weights need to be renormalized
        % before calculating the weighted mean and standard deviation of a
        % simulation variable across the virtual population.
%         ignoreNaN = false;
%         if strcmpi(Obj.OptimOptions.priorPrevalenceWeightAssumption,"uniform")
%             pws = (1/Obj.NumVPs)*ones(1,Obj.NumVPs);
%         elseif strcmpi(Obj.OptimOptions.priorPrevalenceWeightAssumption,"specified")
%             pws = Obj.InputVPop.pws;
%         elseif strcmpi(Obj.OptimOptions.priorPrevalenceWeightAssumption,"ignoreDropout")
%             pws = NaN;
%             ignoreNaN = true;
%         else
%             error('The specified optimOptions.priorPrevalenceWeightAssumption is not supported.');
%         end
        
        % 0 for not adjusted based on the number of rows a data group takes up 
        % 1 for 1/sqrt(nRows)
        % 2 for 1/(nRows)
        % 3 for 1/(nRows)^2
        % Original behavior is 0, but in spot checks 3 looked like
        % it started closer to some of the targets from hypothesis testing
        groupWeightMethod = 3;

        % Convert the VPop simData into a table, for easier indexing later
        % on:
        mySimData = Obj.InputVPop.simData.Data;
          
        % Define a struct array of length 'nDataGroups', where a data group
        % contains all of the datapoints that are dependent (not
        % independent) on each other (e.g., all of the datapoints
        % corresponding to a CDF). Each element of this array is a 
        dataConsolidated = [];       % consider how to preallocate dataConsolidated in future release

        % Extract data for 'binTable'
        myTable = Obj.InputVPop.binTable;
        if istable(myTable)
            for iBinTableRow = 1:height(myTable)
            % Loop through rows in table

%                 % Determine the index of the row in the simData table that corresponds
%                 % to this particular 'binTable' row:
                % extract bin edges
         %       binTableVariableNames = myTable.Properties.VariableNames;
				binEdges = [-Inf, myTable{iBinTableRow,'binEdges'}{1}, Inf];

                % Extract the simulation values, and bin them according to the binEdges
                % determined above
                keepIndices = myTable{iBinTableRow,'predIndices'}{1}; % this is indices of pts on therapy & in subpopulation   
                simValspts = myTable{iBinTableRow,'predSample'}{1};
                simVals = NaN(1,size(mySimData,2));
                simVals(keepIndices) = simValspts;
                simValsBinned=discretize(simVals,binEdges); % fixed original bugs in reading NaN values in simVals
                                
                nObservations = length(binEdges)-1;
                dataParticular = initDataParticular(Obj.NumVPs,nObservations,Obj.OptimOptions.binTableGroupWeight);
                % Loop through bins and define an observation row for each bin
                for iBin = 1:nObservations
                    % The observed response is taken to be the experimentally observed
                    % probability of this bin
					observationValParticularBin = myTable{iBinTableRow,'expBins'}{1};
					observationValParticularBin = observationValParticularBin(iBin);
                    % Description of this observation:
                    descriptionParticular = ['binTable; Row ' num2str(iBinTableRow) '; Bin ' num2str(iBin)];
                    % The values for the independent variables will be set to 1 for
                    % VPs in the bin, and 0 for those outside the bin:
                    independentVarValsParticularBin = simValsBinned;
                    
                    nanValsIndexMask = isnan(simValsBinned);   
                    independentVarValsParticularBin(independentVarValsParticularBin~=iBin & (~nanValsIndexMask)) = 0; 
                    independentVarValsParticularBin(independentVarValsParticularBin==iBin & (~nanValsIndexMask)) = 1; 
                    dataParticular.independentVarVals(iBin,:) = independentVarValsParticularBin;
                    dataParticular.vpIsInSubgroup(iBin,:) = [~nanValsIndexMask]; 
                    dataParticular.independentVarVals(iBin,nanValsIndexMask) = 0;                   
                    
                    dataParticular.observationVals(iBin) = observationValParticularBin;
                    dataParticular.observationWeights(iBin) = myTable.weight(iBinTableRow);
                    dataParticular.observationDescriptions{iBin} = descriptionParticular;
                    if (nargin(Obj.OptimOptions.expWeightFuncHandle)==1) % first round and the last booststrapping pw points
                       STDbin = sqrt(dataParticular.observationVals(iBin)*(1-dataParticular.observationVals(iBin)));
                       
                       % if there are less than 1% or more than 99% vpIsInSubgroup for a bin, remove it from matrix
                       vpfractionSim = sum(dataParticular.independentVarVals(iBin,:))/sum(dataParticular.vpIsInSubgroup(iBin,:));
                       if vpfractionSim <= 0.01 || vpfractionSim >=0.99
                            dataParticular.expWeight(iBin) = 0;
                       else
                           if STDbin <= 0.1   % cutoff to avoid overweighting super low value bins: std ~= 0.1, when there is 1% response.
                              STDbin = 0.1;
                           end
                           dataParticular.expWeight(iBin) = sqrt(myTable.expN(iBinTableRow))/STDbin;
                       end
                    else % bootstrapping getting initial pws
                        dataParticular.expWeight(iBin) = Obj.OptimOptions.expWeightFuncHandle(myTable.expN(iBinTableRow),nan,descriptionParticular);
                    end
                    dataParticular.expWeight(iBin) = dataParticular.expWeight(iBin)/nObservations;
                end

                if groupWeightMethod == 0
                    dataParticular.dataGroupWeight = 1;
                elseif groupWeightMethod == 1
                    dataParticular.dataGroupWeight = 1/sqrt(length(dataParticular.observationVals));
                elseif groupWeightMethod == 2
                    dataParticular.dataGroupWeight = 1/(length(dataParticular.observationVals));
                elseif groupWeightMethod == 3
                    dataParticular.dataGroupWeight = 1/(length(dataParticular.observationVals))^2;                    
                end
                dataConsolidated = [dataConsolidated dataParticular];
            end
        end

        % Extract data from distTable
        myTable = Obj.InputVPop.distTable;
        if istable(myTable)
            nRows = height(myTable);
            for iDistTableRow = 1:nRows
            % loop through table rows
                % Some of the code below is adapted from the QSP Toolbox function
                % 'plotDistCDFVPop.m':
                expN = myTable{iDistTableRow,'expN'};
            %     predN = myTable{iDistTableRow,'predN'};
                expW = 1./(expN*ones(1,expN));
                predW = myTable{iDistTableRow,'predProbs'}{1};
            %     expSample = myTable{iDistTableRow,'expSample'}{1};
            %     predSample = myTable{iDistTableRow,'predSample'}{1};
                SC = myTable{iDistTableRow,'combinedPoints'}{1};
                expInd = myTable{iDistTableRow,'expCombinedIndices'}{1};
                predInd = myTable{iDistTableRow,'simCombinedIndices'}{1};		
                [CDFexp, ~] = alignCDFsPreGrid(SC, expInd, predInd, expW, predW);
                
                    % If the user hasn't specified to fit only certain probabilities on the
                    % CDF curve, then fit all of the probabilities. This may take a very
                    % long time if the CDF contains many points.
                    if isstring(Obj.OptimOptions.cdfProbsToFit) && strcmpi(Obj.OptimOptions.cdfProbsToFit,"all")
                        cdfProbsToFit = CDFexp;
                    elseif isnumeric(Obj.OptimOptions.cdfProbsToFit)
                        cdfProbsToFit = Obj.OptimOptions.cdfProbsToFit;
                    else
                        error('Unsupported specification for Obj.OptimOptions.cdfProbsToFit');
                    end
                
                if ~isempty(CDFexp)
                    nObservations = length(cdfProbsToFit); % or maybe should also try +1
                    dataParticular = initDataParticular(Obj.NumVPs,nObservations,Obj.OptimOptions.distTableGroupWeight);
                    for iIncludedProbs = 1:nObservations
                    % loop through the probabilities to fit
                        targetProb = cdfProbsToFit(iIncludedProbs);
                        % find the index that corresponds to the closest probability to the
                        % target probability:
                        % Lu: to do: maybe this could be simplified by a percentile function in matlab
                        closestIndicesToTargetProb = find(abs(CDFexp-targetProb) == min(abs(CDFexp-targetProb)));
                        % In case there are multiple closest indices, take the middle of
                        % the indices:
                        middleClosestIndexToTargetProb = round(closestIndicesToTargetProb(1) + (closestIndicesToTargetProb(end)-closestIndicesToTargetProb(1))/2);
                        % The response value is the experimentally observed CDF
                        % probability:
                        observationValParticular = CDFexp(middleClosestIndexToTargetProb);
                        descriptionParticular = ['distTable; Row ' num2str(iDistTableRow) '; Included Prob ' num2str(iIncludedProbs)];
                        % The values of the independent variables will be set to 1 for
                        % VPs that contribute to the particular point on the CDF, and
                        % to 0 for VPs that don't contribute to that point:
                        if isnan(Obj.InputVPop.distTable.simCombinedIndices{iDistTableRow}(middleClosestIndexToTargetProb))
                            vpIndicesInBin = [];
                        else
                            vpIndicesInBin = Obj.InputVPop.distTable.predIndices{iDistTableRow}(1:Obj.InputVPop.distTable.simCombinedIndices{iDistTableRow}(middleClosestIndexToTargetProb));
                        end
                        independentVarValParticular = NaN(1,Obj.NumVPs); % so dropout patients are NaN
                        independentVarValParticular(Obj.InputVPop.distTable.predIndices{iDistTableRow}) = 0; % this counts in on therapy & subpopulation
                        independentVarValParticular(vpIndicesInBin) = 1;
                        dataParticular.independentVarVals(iIncludedProbs,:) = independentVarValParticular;
                        nanValsIndexMask = isnan(independentVarValParticular);                 
                        dataParticular.vpIsInSubgroup(iIncludedProbs,:) = [~nanValsIndexMask]; 
                        dataParticular.independentVarVals(iIncludedProbs,nanValsIndexMask) = 0;
                        dataParticular.observationVals(iIncludedProbs) = observationValParticular;
                        dataParticular.observationWeights(iIncludedProbs) = Obj.InputVPop.distTable.weight(iDistTableRow);
                        dataParticular.observationDescriptions{iIncludedProbs} = descriptionParticular;
                        if (nargin(Obj.OptimOptions.expWeightFuncHandle)==1)
                               STDdist = sqrt(dataParticular.observationVals(iIncludedProbs)*(1-dataParticular.observationVals(iIncludedProbs)));
                               
                               vpfractionSim = sum(dataParticular.independentVarVals(iIncludedProbs,:))/sum(dataParticular.vpIsInSubgroup(iIncludedProbs,:));
                               if vpfractionSim <= 0.01 || vpfractionSim >=0.99
                                    dataParticular.expWeight(iIncludedProbs) = 0;
                               else
                                   if STDdist <= 0.1 % cutoff to avoid overweighting super low value bins: std ~= 0.1, when there is 1% response.
                                       STDdist = 0.1;
                                   end
                                   dataParticular.expWeight(iIncludedProbs) = sqrt(Obj.InputVPop.distTable.expN(iDistTableRow))/STDdist;
                               end
                        else
                            dataParticular.expWeight(iIncludedProbs) = Obj.OptimOptions.expWeightFuncHandle(Obj.InputVPop.distTable.expN(iDistTableRow),nan,descriptionParticular);
                        end
                        dataParticular.expWeight(iIncludedProbs) = dataParticular.expWeight(iIncludedProbs)/(nObservations); 
                    end
                else % when there is no VP simulation in a certain 'subpop', assign everything to be 0 (To maintain the dimension of matrix. )
                    nObservations = length(cdfProbsToFit); % or maybe should also try +1
                    dataParticular = initDataParticular(Obj.NumVPs,nObservations,Obj.OptimOptions.distTableGroupWeight);
                    for iIncludedProbs = 1:nObservations
                    % loop through the probabilities to fit
                        targetProb = cdfProbsToFit(iIncludedProbs);
                        % find the index that corresponds to the closest probability to the
                        % target probability:
                        % Lu: to do: maybe this could be simplified by a percentile function in matlab
%                         closestIndicesToTargetProb = find(abs(CDFexp-targetProb) == min(abs(CDFexp-targetProb)));
%                         % In case there are multiple closest indices, take the middle of
%                         % the indices:
%                         middleClosestIndexToTargetProb = round(closestIndicesToTargetProb(1) + (closestIndicesToTargetProb(end)-closestIndicesToTargetProb(1))/2);
%                         % The response value is the experimentally observed CDF
%                         % probability:
%                         observationValParticular = CDFexp(middleClosestIndexToTargetProb);
                        descriptionParticular = ['distTable; Row ' num2str(iDistTableRow) '; Included Prob ' num2str(iIncludedProbs)];
                        % The values of the independent variables will be set to 1 for
                        % VPs that contribute to the particular point on the CDF, and
                        % to 0 for VPs that don't contribute to that point:
                        vpIndicesInBin = Obj.InputVPop.distTable.predIndices{iDistTableRow};
                        
                        independentVarValParticular = NaN(1,Obj.NumVPs); % so dropout patients are NaN
                        independentVarValParticular(Obj.InputVPop.distTable.predIndices{iDistTableRow}) = 0; % this counts in on therapy & subpopulation
                        independentVarValParticular(vpIndicesInBin) = 1;
                        dataParticular.independentVarVals(iIncludedProbs,:) = independentVarValParticular;
                        nanValsIndexMask = isnan(independentVarValParticular);                 
                        dataParticular.vpIsInSubgroup(iIncludedProbs,:) = [~nanValsIndexMask]; 
                        dataParticular.independentVarVals(iIncludedProbs,nanValsIndexMask) = 0;
                        dataParticular.observationVals(iIncludedProbs) = observationValParticular;
                        dataParticular.observationWeights(iIncludedProbs) = Obj.InputVPop.distTable.weight(iDistTableRow);
                        dataParticular.observationDescriptions{iIncludedProbs} = descriptionParticular;
                        if (nargin(Obj.OptimOptions.expWeightFuncHandle)==1)
                               STDdist = sqrt(dataParticular.observationVals(iIncludedProbs)*(1-dataParticular.observationVals(iIncludedProbs)));
                               vpfractionSim = sum(dataParticular.independentVarVals(iIncludedProbs,:))/sum(dataParticular.vpIsInSubgroup(iIncludedProbs,:));
                               if vpfractionSim <= 0.01 || vpfractionSim >=0.99
                                    dataParticular.expWeight(iIncludedProbs) = 0;
                               else
                                   if STDdist <= 0.1 % cutoff to avoid overweighting super low value bins: std ~= 0.1, when there is 1% response.
                                       STDdist = 0.1;
                                   end
                                   dataParticular.expWeight(iIncludedProbs) = sqrt(Obj.InputVPop.distTable.expN(iDistTableRow))/STDdist;
                               end
                        else
                            dataParticular.expWeight(iIncludedProbs) = Obj.OptimOptions.expWeightFuncHandle(Obj.InputVPop.distTable.expN(iDistTableRow),nan,descriptionParticular);
                        end
                        dataParticular.expWeight(iIncludedProbs) = dataParticular.expWeight(iIncludedProbs)/(nObservations); 
                    end
                    
                end

                    if groupWeightMethod == 0
                        dataParticular.dataGroupWeight = 1;
                    elseif groupWeightMethod == 1
                        dataParticular.dataGroupWeight = 1/sqrt(length(dataParticular.observationVals));
                    elseif groupWeightMethod == 2
                        dataParticular.dataGroupWeight = 1/(length(dataParticular.observationVals));
                    elseif groupWeightMethod == 3
                        dataParticular.dataGroupWeight = 1/(length(dataParticular.observationVals))^2;
                    end                
                    dataConsolidated = [dataConsolidated dataParticular];
            end
        end

        % Extract data related to the 'brTable'
        % RECIST Class definitions
        % 0 = CR, 1 = PR, 2 = SD, 3 = PD
        recistClassStr = {'CR','PR','SD','PD'};
        resistClassNum = [0 1 2 3];
        if isa(Obj.InputVPop, 'VPopRECIST')
            if istable(Obj.InputVPop.brTableRECIST)
                for iBRTblRow = 1:height(Obj.InputVPop.brTableRECIST)  %% will not have NaN or dropout situation. only need to consider Subpopulation
                % Loop through table rows
                    % Extract the simulation data:
                    iMaskSimDataRow = Obj.InputVPop.simData.brRows == iBRTblRow;
                    brDataParticular = Obj.InputVPop.simData.brData(iMaskSimDataRow,:);
                    nObservations = length(resistClassNum);
                    dataParticular = initDataParticular(Obj.NumVPs,nObservations,Obj.OptimOptions.brTableRECISTGroupWeight);
                    for iRECISTClass = 1:nObservations
                    % loop through RECIST classes
                        % The response value is the experimentally observed probability for
                        % this RECIST class:
                        observationValParticular = Obj.InputVPop.brTableRECIST.(['exp' recistClassStr{iRECISTClass}])(iBRTblRow);
                        descriptionParticular = ['brTableRECIST; Row ' num2str(iBRTblRow) '; RECIST Class ' num2str(iRECISTClass)];
                        % The values of the independent variables will be set to 1 for
                        % VPs that are in the RECIST class, and 0 for those that aren't
                        dataParticular.vpIsInSubgroup(iRECISTClass,:) = 0; % only vps in subpop are 1
                        subpopNo = Obj.InputVPop.brTableRECIST.subpopNo(iBRTblRow);
                        dataParticular.vpIsInSubgroup(iRECISTClass,Obj.InputVPop.subpopTable.vpIndices{subpopNo,1}) = 1;
                        vpsInClass = brDataParticular == resistClassNum(iRECISTClass);
                        dataParticular.independentVarVals(iRECISTClass,:) = vpsInClass;
                        dataParticular.observationVals(iRECISTClass) = observationValParticular;
                        dataParticular.observationWeights(iRECISTClass) = Obj.InputVPop.brTableRECIST.weight(iBRTblRow);
                        dataParticular.observationDescriptions{iRECISTClass} = descriptionParticular;
                       % dataParticular.expWeight(iRECISTClass) = Obj.OptimOptions.expWeightFuncHandle(Obj.InputVPop.brTableRECIST.expN(iBRTblRow),nan,descriptionParticular);
                       if (nargin(Obj.OptimOptions.expWeightFuncHandle)==1)
                               STDbr = sqrt(dataParticular.observationVals(iRECISTClass)*(1-dataParticular.observationVals(iRECISTClass)));
                               % if there are less than 1% or more than 99% vpIsInSubgroup for a bin, remove it from matrix
                               vpfractionSim = sum(dataParticular.independentVarVals(iRECISTClass,:))/sum(dataParticular.vpIsInSubgroup(iRECISTClass,:));
                               if vpfractionSim <= 0.01 || vpfractionSim >=0.99
                                    dataParticular.expWeight(iRECISTClass) = 0;
                               else
                                   if STDbr <= 0.1
                                      STDbr = 0.1;
                                   end
                                   dataParticular.expWeight(iRECISTClass) = sqrt(Obj.InputVPop.brTableRECIST.expN(iBRTblRow))/STDbr;
                               end
                       else
                           dataParticular.expWeight(iRECISTClass) = Obj.OptimOptions.expWeightFuncHandle(Obj.InputVPop.brTableRECIST.expN(iBRTblRow),nan,descriptionParticular);
                       end
                       dataParticular.expWeight(iRECISTClass) = dataParticular.expWeight(iRECISTClass)/nObservations;
                    end
                    if groupWeightMethod == 0
                        dataParticular.dataGroupWeight = 1;
                    elseif groupWeightMethod == 1
                        dataParticular.dataGroupWeight = 1/sqrt(length(dataParticular.observationVals));
                    elseif groupWeightMethod == 2
                        dataParticular.dataGroupWeight = 1/(length(dataParticular.observationVals));
                    elseif groupWeightMethod == 3
                        dataParticular.dataGroupWeight = 1/(length(dataParticular.observationVals))^2;                    
                    end  
                    dataConsolidated = [dataConsolidated dataParticular];
                end
            end
        end
        
        if isa(Obj.InputVPop, 'VPopRECIST')       
            if istable(Obj.InputVPop.rTableRECIST)
                for iRTblRow = 1:height(Obj.InputVPop.rTableRECIST)
                    iMaskSimDataRow = Obj.InputVPop.simData.rRows == iRTblRow;
                    rDataParticular = Obj.InputVPop.simData.rData(iMaskSimDataRow,:);
                    nObservations = length(resistClassNum);
                    dataParticular = initDataParticular(Obj.NumVPs,nObservations,Obj.OptimOptions.rTableRECISTGroupWeight);
                    for iRECISTClass = 1:nObservations
                    % loop through RECIST classes
                        % The response value is the experimentally observed probability for
                        % this RECIST class:
                        observationValParticular = Obj.InputVPop.rTableRECIST.(['exp' recistClassStr{iRECISTClass}])(iRTblRow);
                        descriptionParticular = ['rTableRECIST; Row ' num2str(iRTblRow) '; RECIST Class ' num2str(iRECISTClass)];
                        % The values of the independent variables will be set to 1 for
                        % VPs that are in the RECIST class, and 0 for those that aren't
                        dataParticular.vpIsInSubgroup(iRECISTClass,:) = 0; % only vps in subpop are 1
                        subpopNo = Obj.InputVPop.rTableRECIST.subpopNo(iRTblRow);
                        dataParticular.vpIsInSubgroup(iRECISTClass,Obj.InputVPop.subpopTable.vpIndices{subpopNo,1}) = 1;                                               
                        vpsInClass = rDataParticular == resistClassNum(iRECISTClass);
                        dataParticular.independentVarVals(iRECISTClass,:) = vpsInClass;
                        dataParticular.observationVals(iRECISTClass) = observationValParticular;
                        dataParticular.observationWeights(iRECISTClass) = Obj.InputVPop.rTableRECIST.weight(iRTblRow);
                        dataParticular.observationDescriptions{iRECISTClass} = descriptionParticular;
                       % dataParticular.expWeight(iRECISTClass) = Obj.OptimOptions.expWeightFuncHandle(Obj.InputVPop.rTableRECIST.expN(iRTblRow),nan,descriptionParticular);
                       if (nargin(Obj.OptimOptions.expWeightFuncHandle)==1)
                            STDr = sqrt(dataParticular.observationVals(iRECISTClass)*(1-dataParticular.observationVals(iRECISTClass)));
                            % if there are less than 1% or more than 99% vpIsInSubgroup for a bin, remove it from matrix
                               vpfractionSim = sum(dataParticular.independentVarVals(iRECISTClass,:))/sum(dataParticular.vpIsInSubgroup(iRECISTClass,:));
                               if vpfractionSim <= 0.01 || vpfractionSim >=0.99
                                    dataParticular.expWeight(iRECISTClass) = 0;
                               else
                                   if STDr <= 0.1
                                        STDr = 0.1; 
                                   end
                                   dataParticular.expWeight(iRECISTClass) = sqrt(Obj.InputVPop.rTableRECIST.expN(iRTblRow))/STDr;
                               end
                       else
                           dataParticular.expWeight(iRECISTClass) = Obj.OptimOptions.expWeightFuncHandle(Obj.InputVPop.rTableRECIST.expN(iRTblRow),nan,descriptionParticular);      
                       end
                       dataParticular.expWeight(iRECISTClass) = dataParticular.expWeight(iRECISTClass)/nObservations;
                    end
                    if groupWeightMethod == 0
                        dataParticular.dataGroupWeight = 1;
                    elseif groupWeightMethod == 1
                        dataParticular.dataGroupWeight = 1/sqrt(length(dataParticular.observationVals));
                    elseif groupWeightMethod == 2
                        dataParticular.dataGroupWeight = 1/(length(dataParticular.observationVals));
                    elseif groupWeightMethod == 3
                        dataParticular.dataGroupWeight = 1/(length(dataParticular.observationVals))^2;                    
                    end  
                    dataConsolidated = [dataConsolidated dataParticular];
                end
            end
        end

        % Extract data for mean and standard deviation
        % Rewrote the variance comparison by fitting to E(X2), no longer need approximation
        myTable = Obj.InputVPop.mnSDTable;
        if istable(myTable)
            for iSumStatRow = 1:height(myTable)
                % Loop through table rows
                nObservations = 2;
                dataParticular = initDataParticular(Obj.NumVPs,nObservations,Obj.OptimOptions.mnSDTableGroupWeight);

                % Extract the appropriate simulation data:               
                keepIndices = myTable{iSumStatRow,'predIndices'}{1}; % this is indices of pts on therapy & in subpopulation   
                simValspts = myTable{iSumStatRow,'predSample'}{1};
                simVals = NaN(1,size(mySimData,2));
                simVals(keepIndices) = simValspts;

                % Some simVals will be NaN for VPs that dropped off of therapy. Thus,
                % we need to scale the prevalence weights so that the sum of the
                % weights for the VPs still on therapy adds to 1.
                nanValsIndexMask = isnan(simVals);
                dataParticular.vpIsInSubgroup(1:2,:) = [~nanValsIndexMask;~nanValsIndexMask];

                % Fit mean                           
                % The response value is the experimental mean:
                dataParticular.observationVals(1) = Obj.InputVPop.mnSDTable.expMean(iSumStatRow);
                dataParticular.observationDescriptions{1} = ['mnSDTable; Row ' num2str(iSumStatRow) '; mean'];
                % The values for the independent variables are simply the
                % simulation values
                dataParticular.independentVarVals(1,:) = simVals;                              
                dataParticular.independentVarVals(1,nanValsIndexMask) = 0;
                dataParticular.observationWeights(1) = Obj.InputVPop.mnSDTable.weightMean(iSumStatRow);
               % dataParticular.expWeight(1) = Obj.OptimOptions.expWeightFuncHandle(Obj.InputVPop.mnSDTable.expN(iSumStatRow),Obj.InputVPop.mnSDTable.expSD(iSumStatRow),dataParticular.observationDescriptions{1});
               if (nargin(Obj.OptimOptions.expWeightFuncHandle)==1) 
                    STDmean = Obj.InputVPop.mnSDTable.expSD(iSumStatRow);
                    if STDmean == 0
                        STDmean = 1e-6;
                    end
                    % we don't top variance for mn and sd, because this is not Bernoulli distribution, some fracton values are ~1e-6. 
                    dataParticular.expWeight(1) = sqrt(Obj.InputVPop.mnSDTable.expN(iSumStatRow))/STDmean;
               else
                    dataParticular.expWeight(1) = Obj.OptimOptions.expWeightFuncHandle(Obj.InputVPop.mnSDTable.expN(iSumStatRow),Obj.InputVPop.mnSDTable.expSD(iSumStatRow),dataParticular.observationDescriptions{1});
               end

               % Fit SD (actually, variance)
               simX2 = simVals.^2;
               % E(Xsquare), after drop out. match gof comparison better
              if ~isempty(Obj.InputVPop.expData)
                iSample = (Obj.InputVPop.expData.time == Obj.InputVPop.mnSDTable.time(iSumStatRow)) & ...
                    strcmp(Obj.InputVPop.expData.interventionID,Obj.InputVPop.mnSDTable.interventionID{iSumStatRow}) & ...
                    strcmp(Obj.InputVPop.expData.elementID,Obj.InputVPop.mnSDTable.elementID{iSumStatRow}) & ...
                    strcmp(Obj.InputVPop.expData.elementType,Obj.InputVPop.mnSDTable.elementType{iSumStatRow}) & ...
                    strcmp(Obj.InputVPop.expData.expVarID,Obj.InputVPop.mnSDTable.expVarID{iSumStatRow}) & ...
                    strcmp(Obj.InputVPop.expData.expDataID,Obj.InputVPop.mnSDTable.expDataID{iSumStatRow});
              else
                  iSample = [];
              end
              if (find(iSample))
                    numericIndices = cellfun(@isnumeric,table2cell(Obj.InputVPop.expData(iSample,:)));
                    rvIndices = ismember(Obj.InputVPop.expData.Properties.VariableNames,{'subpopNo','time'});
                    expVals = Obj.InputVPop.expData{iSample,numericIndices & (~rvIndices)}; % for VPop non-RECIST and RECIST, the num column indices are different. e.g. example 06
                    expVals = expVals(~isnan(expVals));
                    if length(expVals)>1
                        expMeanX2 = mean(expVals.^2);
                        expSTDX2 = std(expVals.^2);
                    else
                        % Sometimes there is only invalid one patient in the expData or pseudo data (need to think how to flag?), then we treat them as we only have summary statistics (mean, std from literature), we cannot get expVals, 
                        % thus, we just get expMeanX2 = expVar + (expMeanX)^2
                        expMeanX = Obj.InputVPop.mnSDTable.expMean(iSumStatRow);
                        expSTDX = Obj.InputVPop.mnSDTable.expSD(iSumStatRow);
                        expMeanX2 = expSTDX^2 + (expMeanX)^2;
                        expMeanX4 = expMeanX^4 + 6*expMeanX^2*expSTDX^2 + 3*expSTDX^4;     % assume normal distribution: E(x^4)=u^4+6*u^2*sigma^2+3*sigma^4 . checked with known data, this is more correct!
                        expVarX2 = expMeanX4 - (expMeanX2)^2; % do this only for summary data: Var(X^2)=E(X^4)-E(X^2)^2;
                        expSTDX2 = sqrt(expVarX2);
                    end
              else                                   
                    % Sometimes we only have summary statistics (mean, std from literature), we cannot get expVals, 
                    % thus, we just get expMeanX2 = expVar + (expMeanX)^2
                    expMeanX = Obj.InputVPop.mnSDTable.expMean(iSumStatRow);
                    expSTDX = Obj.InputVPop.mnSDTable.expSD(iSumStatRow);
                    expMeanX2 = expSTDX^2 + (expMeanX)^2;
                    expMeanX4 = expMeanX^4 + 6*expMeanX^2*expSTDX^2 + 3*expSTDX^4;     % assume normal distribution: E(x^4)=u^4+6*u^2*sigma^2+3*sigma^4 . checked with known data, this is more correct!
                    expVarX2 = expMeanX4 - (expMeanX2)^2; % do this only for summary data: Var(X^2)=E(X^4)-E(X^2)^2;
                    expSTDX2 = sqrt(expVarX2);
              end
                
              % The response value will be the variance:
              %dataParticular.observationVals(2) = Obj.InputVPop.mnSDTable.expSD(iSumStatRow)^2;
              dataParticular.observationVals(2) = expMeanX2;
              dataParticular.observationDescriptions{2} = ['mnSDTable; Row ' num2str(iSumStatRow) '; variance'];
              expMean = Obj.InputVPop.mnSDTable.expMean(iSumStatRow);
              % residualsSquared = (simVals-expMean).^2;
              % dataParticular.independentVarVals(2,:) = residualsSquared;
              dataParticular.independentVarVals(2,:) = simX2;
              dataParticular.independentVarVals(2,nanValsIndexMask) = 0;
              dataParticular.observationWeights(2) = Obj.InputVPop.mnSDTable.weightSD(iSumStatRow);
              % dataParticular.expWeight(2) = Obj.OptimOptions.expWeightFuncHandle(Obj.InputVPop.mnSDTable.expN(iSumStatRow),Obj.InputVPop.mnSDTable.expSD(iSumStatRow),dataParticular.observationDescriptions{2});
              if (nargin(Obj.OptimOptions.expWeightFuncHandle)==1) 
                    if expSTDX2 ==0
                        expSTDX2 = 1e-6;
                    end
                    % we don't top variance for mn and sd, because this is not bernouli distribution, some fracton values are ~1e-6. 
                    dataParticular.expWeight(2) = sqrt(Obj.InputVPop.mnSDTable.expN(iSumStatRow))/expSTDX2;
              else
                    dataParticular.expWeight(2) = Obj.OptimOptions.expWeightFuncHandle(Obj.InputVPop.mnSDTable.expN(iSumStatRow),Obj.InputVPop.mnSDTable.expSD(iSumStatRow),dataParticular.observationDescriptions{2});
              end
              dataConsolidated = [dataConsolidated dataParticular];
            end
        end
        
        %%%%%%%%%%%%%%%%%%%%%%%%%%%%%%%%%%%%%%%%%%%%%%%%%%%%%%%%%%%%%%%%%%%%%%%%%%%%%%%%%%%%%%%%%%%%%%%%%%%%%%%%%%%%%%%%%%%%%%%%%%%%%%%%%%%%%%%%%%%%%%%%%                
        % Rewrote the 2D correlation comparison by fitting to E(X*Y), no need approximation
        % Extract data for 2D correlation
        myTable = Obj.InputVPop.corTable;
        if istable(myTable)
            for iSumStatRow = 1:height(myTable)
                % Loop through table rows
                nObservations = 1;
                dataParticular = initDataParticular(Obj.NumVPs,nObservations,Obj.OptimOptions.corTableGroupWeight);
                % Previous version extract data from Obj.InputVPop.mnSDTable; It is better to extract the experimental data from myVPop.expData table, in case sometimes we don't have all data categories. more robust
                iMaskSample1Row = (Obj.InputVPop.expData.time== Obj.InputVPop.corTable.time1(iSumStatRow)) & ...
                    strcmp(Obj.InputVPop.expData.interventionID,Obj.InputVPop.corTable.interventionID1{iSumStatRow}) & ...
                    strcmp(Obj.InputVPop.expData.elementID,Obj.InputVPop.corTable.elementID1{iSumStatRow}) & ...
                    strcmp(Obj.InputVPop.expData.elementType,Obj.InputVPop.corTable.elementType1{iSumStatRow}) & ...
                    strcmp(Obj.InputVPop.expData.expVarID,Obj.InputVPop.corTable.expVarID1{iSumStatRow}) & ...
                    strcmp(Obj.InputVPop.expData.expDataID,Obj.InputVPop.corTable.expDataID1{iSumStatRow});
              %  iSample1=Obj.InputVPop.expData{iMaskSample1Row,13:end};
                numericIndices = cellfun(@isnumeric,table2cell(Obj.InputVPop.expData(iMaskSample1Row,:)));
                rvIndices = ismember(Obj.InputVPop.expData.Properties.VariableNames,{'subpopNo','time'});
                iSample1 = Obj.InputVPop.expData{iMaskSample1Row,numericIndices & (~rvIndices)}; % for VPop non-RECIST and RECIST, the num column indices are different. e.g. example 06
                                    
                iMaskSample2Row = (Obj.InputVPop.expData.time== Obj.InputVPop.corTable.time2(iSumStatRow)) & ...
                    strcmp(Obj.InputVPop.expData.interventionID,Obj.InputVPop.corTable.interventionID2{iSumStatRow}) & ...
                    strcmp(Obj.InputVPop.expData.elementID,Obj.InputVPop.corTable.elementID2{iSumStatRow}) & ...
                    strcmp(Obj.InputVPop.expData.elementType,Obj.InputVPop.corTable.elementType2{iSumStatRow}) & ...
                    strcmp(Obj.InputVPop.expData.expVarID,Obj.InputVPop.corTable.expVarID2{iSumStatRow}) & ...
                    strcmp(Obj.InputVPop.expData.expDataID,Obj.InputVPop.corTable.expDataID2{iSumStatRow});
              %  iSample2=Obj.InputVPop.expData{iMaskSample2Row,13:end};
                numericIndices = cellfun(@isnumeric,table2cell(Obj.InputVPop.expData(iMaskSample2Row,:)));
                rvIndices = ismember(Obj.InputVPop.expData.Properties.VariableNames,{'subpopNo','time'});
                iSample2 = Obj.InputVPop.expData{iMaskSample2Row,numericIndices & (~rvIndices)}; % for VPop non-RECIST and RECIST, the num column indices are different. e.g. example 06
                         
                iSample12 =  ~isnan(iSample1) & ~isnan(iSample2);
                expVals1 = iSample1(iSample12);
                expVals2 = iSample2(iSample12);
                expMeanXY = mean(expVals1.*expVals2);
                expSTDXY = std(expVals1.*expVals2);

                keepIndices = myTable{iSumStatRow,'predIndices'}{1}; % this is indices of pts on therapy & in subpopulation   
                simValspts = myTable{iSumStatRow,'predSample'}{1};
                simVals = NaN(2,size(mySimData,2));
                simVals(:,keepIndices) = simValspts;
                simVals1 = simVals(1,:);
                simVals2 = simVals(2,:);
                % simValscorr = ((simVals1-sampleexpmean(1)).*(simVals2-sampleexpmean(2)))/(sampleexpsd(1)*(sampleexpsd(2)));

                % Some simVals will be NaN for VPs that dropped off of therapy. Thus,
                % we need to scale the prevalence weights so that the sum of the
                % weights for the VPs still on therapy adds to 1.
                nanValsIndexMask = isnan(simVals1) | isnan(simVals2);
                
                % Fit the correlations
                % The response value is the experimental correlation:
                %dataParticular.observationVals(1) = Obj.InputVPop.corTable.expCor(iSumStatRow);
                dataParticular.observationVals(1) = expMeanXY;
                dataParticular.observationDescriptions{1} = ['corTable; Row ' num2str(iSumStatRow) '; correlation'];
                % The values for the independent variables are simply the
                % simulation values
                % dataParticular.independentVarVals(1,:) = simValscorr;   
                simXY =  simVals1.*simVals2;
                dataParticular.independentVarVals(1,:) = simXY;             
                dataParticular.vpIsInSubgroup(1,:) = [~nanValsIndexMask];
                dataParticular.independentVarVals(1,nanValsIndexMask) = 0;
                % mask dropouts and subpopulations
                dataParticular.independentVarVals(1,nanValsIndexMask) = 0;
                dataParticular.observationWeights(1) = Obj.InputVPop.corTable.weight(iSumStatRow);
                % dataParticular.expWeight(1) = Obj.OptimOptions.expWeightFuncHandle(Obj.InputVPop.corTable.expN(iSumStatRow),nan,dataParticular.observationDescriptions{1});
                if (nargin(Obj.OptimOptions.expWeightFuncHandle)==1) 
                    if expSTDXY == 0
                        expSTDXY = 1e-6;
                    end
                    % we don't top variance for mn and sd, because this is not bernouli distribution, some fracton values are ~1e-6. 
                    dataParticular.expWeight(1) = sqrt(Obj.InputVPop.corTable.expN(iSumStatRow))/expSTDXY;
                else
                    dataParticular.expWeight(1) = Obj.OptimOptions.expWeightFuncHandle(Obj.InputVPop.corTable.expN(iSumStatRow),nan,dataParticular.observationDescriptions{1});
                end
                dataConsolidated = [dataConsolidated dataParticular];
            end
        end
        %%%%%%%%%%%%%%%%%%%%%%%%%%%%%%%%%%%%%%%%%%%%%%%%%%%%%%%%%%%%%%%%%%%%%%%%%%%%%%%%%%%%%%%%%%%%%%%%%%%%%%%%%%%%%%%%%%%%%%%%%%%%%%%%%%%%%%%%%%%%%%%%%

		 % Now let's try to formulate the linear problem for 2D PDF
        % ***************************************************************************************************
        % Extract data from distTable
        if istable(Obj.InputVPop.distTable2D)
            nRows = height(Obj.InputVPop.distTable2D);
            myTable = Obj.InputVPop.distTable2D;
            for iDistTableRow = 1:nRows
                % loop through table rows                
                % split each dimension to bins by euqal quantile
                % count the number of data points in each grid; divide by the total data points
                expSample = myTable{iDistTableRow,'expSample'}{1};
                expVals1 = expSample(1,:);
                expVals2 = expSample(2,:);

                % read the mesh information
                if isnumeric(Obj.OptimOptions.pdf2DProbsToFitN)
                    pdf2DProbsToFitN = Obj.OptimOptions.pdf2DProbsToFitN;
                else
                    error('Unsupported specification for Obj.OptimOptions.pdf2DProbsToFitN. Must be a integer.');
                end
%                 xmesh = [min(expVals1),quantile(expVals1,(pdf2DProbsToFitN-1)),max(expVals1)];
%                 ymesh = [min(expVals2),quantile(expVals2,(pdf2DProbsToFitN-1)),max(expVals2)];
                xmesh = [-Inf,quantile(expVals1,(pdf2DProbsToFitN-1)),Inf];
                ymesh = [-Inf,quantile(expVals2,(pdf2DProbsToFitN-1)),Inf];
                % create the mesh coordinates
                [Xmesh,Ymesh] = meshgrid(xmesh,ymesh);
                % build the bin edges
                %[nnodey,nnodex]=size(Xmesh);
                Xbinloweredges = Xmesh(1:(end-1),1:(end-1));
                Xbinupperedges = Xmesh(2:end,2:end);
                Ybinloweredges = Ymesh(1:(end-1),1:(end-1));
                Ybinupperedges = Ymesh(2:end,2:end);
                
                % vectorize bin edges
                Xbinloweredgesv = Xbinloweredges(:);
                Xbinupperedgesv = Xbinupperedges(:);
                Ybinloweredgesv = Ybinloweredges(:);
                Ybinupperedgesv = Ybinupperedges(:);
                
                % now read the simulation data
                keepIndices = myTable{iDistTableRow,'predIndices'}{1};  
                simSample = myTable{iDistTableRow,'predSample'}{1};
                simVals = NaN(2,size(mySimData,2));
                simVals(:,keepIndices) = simSample;
                simVals1 = simVals(1,:);
                simVals2 = simVals(2,:);
                
                nanValsIndexMask = isnan(simVals1) | isnan(simVals2);
                
                % the length of vectors should be the same so we can choose
                % any to get number of Observations
                nObservations = length(Ybinupperedgesv);
                dataParticular = initDataParticular(Obj.NumVPs,nObservations,Obj.OptimOptions.distTable2DGroupWeight);
                
                for iIncludedProbs = 1:nObservations
                    % first calculate the target bin probability
                    observationValParticular = NaN(1,size(expSample,2)); % so otherwise dropout patients are NaN
                    observationValParticular(:) = 0; 
                    observationValParticular((expVals1 >= Xbinloweredgesv(iIncludedProbs)) & (expVals1 <= Xbinupperedgesv(iIncludedProbs)) ...
                        & (expVals2 >= Ybinloweredgesv(iIncludedProbs)) & (expVals2 <= Ybinupperedgesv(iIncludedProbs)))=1;              
                    targetProb = sum(observationValParticular)/size(expSample,2);
                    observationValParticular = targetProb;

                    descriptionParticular = ['distTable2D; Row ' num2str(iDistTableRow) '; Included 2D bin ' num2str(iIncludedProbs)];
                    % The values of the independent variables will be set to 1 for
                    % VPs that contribute to the particular point on the 2D PDF (within the bin), and
                    % to 0 for VPs that don't contribute to that point:
                    independentVarValParticular = NaN(1,Obj.NumVPs); % so otherwise dropout patients are NaN
                    independentVarValParticular(~nanValsIndexMask) = 0; 

                    independentVarValParticular((simVals1 >= Xbinloweredgesv(iIncludedProbs)) & ...
                        (simVals1 <= Xbinupperedgesv(iIncludedProbs)) ...
                        & (simVals2 >= Ybinloweredgesv(iIncludedProbs)) ...
                        & (simVals2 <= Ybinupperedgesv(iIncludedProbs)))=1;
                    
                    dataParticular.independentVarVals(iIncludedProbs,:) = independentVarValParticular;
                    dataParticular.vpIsInSubgroup(iIncludedProbs,:) = [~nanValsIndexMask];
                    dataParticular.independentVarVals(iIncludedProbs,nanValsIndexMask) = 0;
                    dataParticular.observationVals(iIncludedProbs) = observationValParticular;
                    dataParticular.observationWeights(iIncludedProbs) = Obj.InputVPop.distTable2D.weight(iDistTableRow);
                    dataParticular.observationDescriptions{iIncludedProbs} = descriptionParticular;
                     if (nargin(Obj.OptimOptions.expWeightFuncHandle)==1)                        
                        STDdist2D = sqrt(dataParticular.observationVals(iIncludedProbs)*(1-dataParticular.observationVals(iIncludedProbs)));
                        % if there are less than 1% or more than 99% vpIsInSubgroup for a bin, remove it from matrix
                        vpfractionSim = sum(dataParticular.independentVarVals(iIncludedProbs,:))/sum(dataParticular.vpIsInSubgroup(iIncludedProbs,:));
                        if vpfractionSim <= 0.01 || vpfractionSim >=0.99
                             dataParticular.expWeight(iIncludedProbs) = 0;
                        else
                               if STDdist2D <= 0.1
                                   STDdist2D = 0.1;   
                               end
                               dataParticular.expWeight(iIncludedProbs) = sqrt(Obj.InputVPop.distTable2D.expN(iDistTableRow))/STDdist2D;
                        end
                     else
                        dataParticular.expWeight(iIncludedProbs) = Obj.OptimOptions.expWeightFuncHandle(Obj.InputVPop.distTable2D.expN(iDistTableRow),nan,descriptionParticular);
                     end
                    dataParticular.expWeight(iIncludedProbs) = dataParticular.expWeight(iIncludedProbs)/(nObservations); 
                end

                if groupWeightMethod == 0
                    dataParticular.dataGroupWeight = 1;
                elseif groupWeightMethod == 1
                    dataParticular.dataGroupWeight = 1/sqrt(length(dataParticular.observationVals));
                elseif groupWeightMethod == 2
                    dataParticular.dataGroupWeight = 1/(length(dataParticular.observationVals));
                elseif groupWeightMethod == 3
                    dataParticular.dataGroupWeight = 1/(length(dataParticular.observationVals))^2;                    
                end  
                dataConsolidated = [dataConsolidated dataParticular];
            end
            warning('on','MATLAB:integral2:maxFunEvalsPass');
            warning('on','MATLAB:integral2:maxFunEvalsFail');
        end
        
        % ***************************************************************************************************
        % Loop through the data groups and perform the following
        % operations:
        % 1. Ensure that there are no missing values.
        % 2. Delete observations which are zero if the objective is
        % calculated relative to the observation value (since we don't want
        % to divide by zero)
        % 3. Calculate the number of observation in the data group.
        Obj.IgnoredObservationsDescriptions = {};
        numObservationsInEachDataGroup = nan(1,length(dataConsolidated));
        for iDataCons = 1:length(dataConsolidated)
            % As a sanity check, ensure that none of the fields in
            % 'dataConsolidated have any missing values:
            fieldNames = fields(dataConsolidated(iDataCons));
            for iField = 1:length(fieldNames)
                if ( iscell(dataConsolidated(iDataCons).(fieldNames{iField})) && any(cellfun(@isempty,dataConsolidated(iDataCons).(fieldNames{iField})),'all') ) || ...
                   ( isnumeric(dataConsolidated(iDataCons).(fieldNames{iField})) && any(isnan(dataConsolidated(iDataCons).(fieldNames{iField})),'all') )
                    error('Missing data values were produced when constructing the linear problem matrices.');
                end
            end

            % delete observations = 0 if fit is relative. % might preassign dataConsolidated for better performance
            if strcmpi(Obj.OptimOptions.responseValTransformation,"relative")
                iMaskZeroObs = dataConsolidated(iDataCons).observationVals == 0;
                Obj.IgnoredObservationsDescriptions = ...
                    [Obj.IgnoredObservationsDescriptions; dataConsolidated(iDataCons).observationDescriptions(iMaskZeroObs)];
                dataConsolidated(iDataCons).independentVarVals(iMaskZeroObs,:) = []; 
                dataConsolidated(iDataCons).observationVals(iMaskZeroObs) = [];
                dataConsolidated(iDataCons).observationWeights(iMaskZeroObs) = [];
                dataConsolidated(iDataCons).observationDescriptions(iMaskZeroObs) = [];
                dataConsolidated(iDataCons).expWeight(iMaskZeroObs) = [];
                dataConsolidated(iDataCons).vpIsInSubgroup(iMaskZeroObs,:) = []; 
            end
            numObservationsInEachDataGroup(iDataCons) = length(dataConsolidated(iDataCons).observationVals);
        end

        % In 'dataConsolidated', the data is separated according to data
        % groups. We still need to consolidate this data into large
        % matrices in order to fit the optimization problem:
        
        % The cumulative number of observations across the data groups:
        cumulativeNumObservations = cumsum(numObservationsInEachDataGroup);
        cumulativeNumObservations = [0 cumulativeNumObservations];
        
        % Initialize 'A' in Ax=b
        % Will eventually be matrix of size 'nObservations' x 'Obj.NumVPs'
        Obj.LinearProblemMatrices.independentVarVals = nan(cumulativeNumObservations(end),Obj.NumVPs);
        
        % Initialize 'b' in Ax=b
        % Will eventually be vector of size 'nObservations' x 1
        Obj.LinearProblemMatrices.observationVals = nan(cumulativeNumObservations(end),1);
        
        % Each observation has associated a weight
        % Will eventually be vector of size 'nObservations' x 1
        Obj.LinearProblemMatrices.observationWeights = nan(cumulativeNumObservations(end),1);
        Obj.LinearProblemMatrices.dataGroupWeights = nan(cumulativeNumObservations(end),1);
        
        % Keep track of the data group corresponding to each row:
        Obj.LinearProblemMatrices.dataGroup = nan(cumulativeNumObservations(end),1);
        
        % String descriptions of the different observations -- useful for
        % debugging.
        % Will eventually be vector of size 'nObservations' x 1
        Obj.LinearProblemMatrices.observationDescriptions = cell(cumulativeNumObservations(end),1);
        
        Obj.LinearProblemMatrices.numObservationsInDataGroup = nan(cumulativeNumObservations(end),1);
        
        Obj.LinearProblemMatrices.expWeight = nan(cumulativeNumObservations(end),1);
        
        Obj.LinearProblemMatrices.vpIsInSubgroup = zeros(size(Obj.LinearProblemMatrices.independentVarVals)); % Mask matrix, same dimension as independentVarVals
        % Now, construct the large matrices, consolidating all of the data groups:
        for iDataCons = 1:length(dataConsolidated)
            startRowIndex = cumulativeNumObservations(iDataCons)+1;
            endRowIndex = cumulativeNumObservations(iDataCons+1);
            Obj.LinearProblemMatrices.independentVarVals(startRowIndex:endRowIndex,:) = dataConsolidated(iDataCons).independentVarVals;
            Obj.LinearProblemMatrices.observationVals(startRowIndex:endRowIndex,:) = dataConsolidated(iDataCons).observationVals;
            Obj.LinearProblemMatrices.observationWeights(startRowIndex:endRowIndex,:) = dataConsolidated(iDataCons).observationWeights;
            Obj.LinearProblemMatrices.dataGroupWeights(startRowIndex:endRowIndex,:) = dataConsolidated(iDataCons).dataGroupWeight;
            Obj.LinearProblemMatrices.dataGroup(startRowIndex:endRowIndex,:) = iDataCons;
            Obj.LinearProblemMatrices.observationDescriptions(startRowIndex:endRowIndex,:) = dataConsolidated(iDataCons).observationDescriptions;
            Obj.LinearProblemMatrices.numObservationsInDataGroup(startRowIndex:endRowIndex,:) = numObservationsInEachDataGroup(iDataCons);
            Obj.LinearProblemMatrices.expWeight(startRowIndex:endRowIndex,:) = dataConsolidated(iDataCons).expWeight;
            Obj.LinearProblemMatrices.vpIsInSubgroup(startRowIndex:endRowIndex,:) = dataConsolidated(iDataCons).vpIsInSubgroup;
        end

        % Calculate and incorporate the observation weights:
        Obj.LinearProblemMatrices = incorporateLinearProblemMatricesWeights(Obj.LinearProblemMatrices,Obj.OptimOptions); % this is not necessary step now, but kept for now.
        
        % Nested function to initialize a 'dataParticular' struct:
        function dataParticular = initDataParticular(nVPs,nObservations,dataGroupWeight)
            dataParticular.independentVarVals = nan(nObservations,nVPs); % 'A' in Ax=b
            dataParticular.vpIsInSubgroup = nan(nObservations,nVPs); % maskMatrix: if the VP is not included, drop off =0, others 1. J.*A
            dataParticular.observationVals = nan(nObservations,1); % 'b' in Ax=b
            dataParticular.dataGroupWeight = dataGroupWeight; % weight for the data group
            dataParticular.observationWeights = nan(nObservations,1); % weight for the observation (in addition to the data group)
            dataParticular.observationDescriptions = cell(nObservations,1); % description of the observation
            dataParticular.expWeight = nan(nObservations,1); % experimental sample size
        end
 end
    
    function Obj = selectFromLinearProblemMatrices(Obj,varargin)
    % Selects only particular data groups and VPs from 'Obj.LinearProblemMatrices',
    % and stores the results in 'Obj.LinearProblemMatricesParticular'.
    % Optional input arguments are 'includedDataGroups' and
    % 'includedVPIndices'. Results are stored in
    % Obj.LinearProblemMatricesParticular.
    %
    
        % Parse optional arguments
        in = inputParser();
        in.addParameter('includedDataGroups',[],@isnumeric);
        in.addParameter('includedVPIndices',[],@isnumeric);
        in.parse(varargin{:});
    
        fieldNames = fields(Obj.LinearProblemMatrices);
        nFields = length(fieldNames);
        
        if isempty(in.Results.includedDataGroups)
            Obj.LinearProblemMatricesParticular = Obj.LinearProblemMatrices;
        else
            observationIndicesToInclude = [];
            % Determine which observations to include:
            for iDataGrp = 1:length(in.Results.includedDataGroups)
                observationIndicesToInclude = [observationIndicesToInclude; find(Obj.LinearProblemMatrices.dataGroup == in.Results.includedDataGroups(iDataGrp))];
            end
            % Extract those observations:
            for iField = 1:nFields
                Obj.LinearProblemMatricesParticular.(fieldNames{iField}) = Obj.LinearProblemMatrices.(fieldNames{iField})(observationIndicesToInclude,:); 
            end
        end
        
        if ~isempty(in.Results.includedVPIndices)
            for iField = 1:nFields
                if size(Obj.LinearProblemMatricesParticular.(fieldNames{iField}),2) > 1
                    Obj.LinearProblemMatricesParticular.(fieldNames{iField}) = Obj.LinearProblemMatricesParticular.(fieldNames{iField})(:,in.Results.includedVPIndices);
                end
            end
        end        
    end
    
    function Obj = runOptimization(Obj)
    % Run the optimization. 
        % Start timer
        tictoc = tic();
        
        if isempty(Obj.LinearProblemMatricesParticular)
            Obj.LinearProblemMatricesParticular = Obj.LinearProblemMatrices;
        end
        nVPs = Obj.NumVPs;
        
        % Run the optimization
        if strcmpi(Obj.OptimOptions.optimizationAlgorithm,"lsqnonneg")
            % Set optimization options:
            options = optimset('Display','none');
            if isstruct(Obj.OptimOptions.optimizationAlgorithmOptions)
                fieldNames = fields(Obj.OptimOptions.optimizationAlgorithmOptions);
            else
                fieldNames = {};
            end
            for iField = 1:length(fieldNames)
                options.(fieldNames{iField}) = Obj.OptimOptions.optimizationAlgorithmOptions.(fieldNames{iField});
            end
            % Run the optimization:
            [Obj.OptimizationResults.optimalPrevalenceWeightsPriorToNormalization,Obj.OptimizationResults.normOfresidualsSquared,Obj.OptimizationResults.residuals,Obj.OptimizationResults.exitFlag,Obj.OptimizationResults.output,Obj.OptimizationResults.lagrangeMultipliers] = ...
                lsqnonneg(Obj.LinearProblemMatricesParticular.independentVarValsWeighted,Obj.LinearProblemMatricesParticular.observationValsWeighted,options);
        elseif strcmpi(Obj.OptimOptions.optimizationAlgorithm,"nnls")
            % For poor problems, the function will throw a bunch of
            % warnings, despite still being able to converge. Let's turn
            % off these warnings.
             warning('off','MATLAB:nearlySingularMatrix');
             warning('off','MATLAB:rankDeficientMatrix');
%             % Run the optimization. Note that this optimization function
            % takes the options as a struct, so we can input them directly.
            Obj.OptimOptions.optimizationAlgorithmOptions.Iter = 30000;  % increased 
            [Obj.OptimizationResults.optimalPrevalenceWeightsPriorToNormalization,Obj.OptimizationResults.lagrangeMultipliers,~] = ...
                nnls(Obj.LinearProblemMatricesParticular.independentVarValsWeighted,Obj.LinearProblemMatricesParticular.observationValsWeighted,Obj.OptimOptions.optimizationAlgorithmOptions);
%             % turn the warnings back on:
            warning('on','MATLAB:nearlySingularMatrix');
            warning('on','MATLAB:rankDeficientMatrix');
             Obj.OptimizationResults.exitFlag = 1;
        elseif strcmpi(Obj.OptimOptions.optimizationAlgorithm,"lsqlin")
            % Specifying an 'x0' would result in a warning that x0 gets ignored in
            % 'lsqlin'
             x0 = (1/nVPs)*ones(nVPs,1);               %           x0 = [];
           if ~(isempty(Obj.OptimizationResults))
                if ~any(isnan(Obj.OptimizationResults.optimalPrevalenceWeightsNormalized))
                    x0 = Obj.OptimizationResults.optimalPrevalenceWeightsNormalized;
                end
           end
            lb = zeros(nVPs,1);
            ub = [];
            if ~isempty(Obj.OptimOptions.maxPrevalenceWeight) && isfinite(Obj.OptimOptions.maxPrevalenceWeight)
                A = spdiags(ones(nVPs,1),0,nVPs,nVPs);
                b = Obj.OptimOptions.maxPrevalenceWeight .* ones(1,nVPs);
            else
                A = [];
                b = [];
            end
            % Constraint that sum of prevalence weights should be 1:
            Aeq = ones(1,nVPs);
            beq = 1;
            % Set optimization options:
         %   lsqlinOptions = optimoptions('lsqlin','Display','none'); % default iter=200
            lsqlinOptions = optimoptions('lsqlin','Display','none','MaxIterations',20000);
            if isstruct(Obj.OptimOptions.optimizationAlgorithmOptions)
                fieldNames = fields(Obj.OptimOptions.optimizationAlgorithmOptions);
            else
                fieldNames = {};
            end
            for iField = 1:length(fieldNames)
                lsqlinOptions.(fieldNames{iField}) = Obj.OptimOptions.optimizationAlgorithmOptions.(fieldNames{iField});
            end
            % Run the optimization:
            [Obj.OptimizationResults.optimalPrevalenceWeightsPriorToNormalization,Obj.OptimizationResults.normOfresidualsSquared,Obj.OptimizationResults.residuals,Obj.OptimizationResults.exitFlag,Obj.OptimizationResults.output,Obj.OptimizationResults.lagrangeMultipliers] = ...
                lsqlin(Obj.LinearProblemMatricesParticular.independentVarValsWeighted,Obj.LinearProblemMatricesParticular.observationValsWeighted,A,b,Aeq,beq,lb,ub,x0,lsqlinOptions);
        elseif strcmpi(Obj.OptimOptions.optimizationAlgorithm,"quadprog")
            % Specifying an 'x0' would result in a warning that x0 gets ignored in
            % 'lsqlin'
             x0 = (1/nVPs)*ones(nVPs,1);   % x0 = [];
           if ~(isempty(Obj.OptimizationResults))
                if ~any(isnan(Obj.OptimizationResults.optimalPrevalenceWeightsNormalized))
                    x0 = Obj.OptimizationResults.optimalPrevalenceWeightsNormalized;
                end
           end
            lb = zeros(nVPs,1);
            ub = [];
            if ~isempty(Obj.OptimOptions.maxPrevalenceWeight) && isfinite(Obj.OptimOptions.maxPrevalenceWeight)
                A = spdiags(ones(nVPs,1),0,nVPs,nVPs);
                b = Obj.OptimOptions.maxPrevalenceWeight .* ones(1,nVPs);
            else
                A = [];
                b = [];
            end
            % Constraint that sum of prevalence weights should be 1:
            [subgroup ia ic]= unique(Obj.LinearProblemMatricesParticular.vpIsInSubgroup,'rows');
            subweights = Obj.LinearProblemMatricesParticular.SubgroupSumWeights;

            Aeq = subgroup;
            beq = subweights(ia);

            % Set optimization options:            
            C = Obj.LinearProblemMatricesParticular.independentVarValsWeighted;
            d = Obj.LinearProblemMatricesParticular.observationValsWeighted;
            F = (C-d).*Obj.LinearProblemMatricesParticular.vpIsInSubgroup;
            ind=find(vecnorm(F,2,2)<1e-6);
            F(ind,:)=[];
             subweights = Obj.LinearProblemMatricesParticular.SubgroupSumWeights;
             subweights(ind,:) = [];
            indexsubweight0=find(subweights<1e-6);
            if isempty(indexsubweight0)
                G = (1./subweights).*F;
            else
%                 subweights(indexsubweight0)=1;   % not to scale when subweight is 0, avoid 1/0 problem, i.e. when there is no VP in this subpop last iteration
                subweights(indexsubweight0)=NaN;
                subweights(indexsubweight0)=min(subweights);
                G = (1./subweights).*F;
            end
            % Set optimization options:
            options = optimoptions('quadprog','Display','none','MaxIterations',20000);
            H = G'*G;
           f = zeros(size(H,1),1);
           options.ConvexCheck = 'on'; 
           [Obj.OptimizationResults.optimalPrevalenceWeightsPriorToNormalization,fval,Obj.OptimizationResults.exitFlag] = quadprog(H,f,A,b,Aeq,beq,lb,ub,x0,options);
           
           if Obj.OptimizationResults.exitFlag == -6  % try turn the convexcheck off
                  options.ConvexCheck = 'off';
                  [Obj.OptimizationResults.optimalPrevalenceWeightsPriorToNormalization,fval,Obj.OptimizationResults.exitFlag] = quadprog(H,f,A,b,Aeq,beq,lb,ub,x0,options);
                  if abs(sum(Obj.OptimizationResults.optimalPrevalenceWeightsPriorToNormalization)-1)<=1e-12 & min(Obj.OptimizationResults.optimalPrevalenceWeightsPriorToNormalization)>0 % if returns a valid solution, keep it
                      Obj.OptimizationResults.exitFlag = 1;
                  else
                      Obj.OptimizationResults.exitFlag = -6;
                  end
           end
        elseif strcmpi(Obj.OptimOptions.optimizationAlgorithm,"quadprogMaxResi")
            % Specifying an 'x0' would result in a warning that x0 gets ignored in
            % 'lsqlin'
             x0 = (1/nVPs)*ones(nVPs,1);   % x0 = [];
             x0 = [x0;1];

            lb = zeros(nVPs+1,1);
            ub = [];
            % Constraint that sum of prevalence weights should be 1:
            [subgroup ia ic]= unique(Obj.LinearProblemMatricesParticular.vpIsInSubgroup,'rows');
            subweights = Obj.LinearProblemMatricesParticular.SubgroupSumWeights;
            Aeq = subgroup;
            beq = subweights(ia);
            Aeq = [Aeq,zeros(size(Aeq,1),1)];

            % Set optimization options:            
            C = Obj.LinearProblemMatricesParticular.independentVarValsWeighted;
            d = Obj.LinearProblemMatricesParticular.observationValsWeighted;
            F = (C-d).*Obj.LinearProblemMatricesParticular.vpIsInSubgroup;
            ind=find(vecnorm(F,2,2)<1e-6);
            F(ind,:)=[];
             subweights = Obj.LinearProblemMatricesParticular.SubgroupSumWeights;
             subweights(ind,:) = [];
            indexsubweight0=find(subweights<1e-6);
            if isempty(indexsubweight0)
                G = (1./subweights).*F;
            else
%                 subweights(indexsubweight0)=1;   % not to scale when subweight is 0, avoid 1/0 problem, i.e. when there is no VP in this subpop last iteration
                subweights(indexsubweight0)=NaN;
                subweights(indexsubweight0)=min(subweights);
                G = (1./subweights).*F;
            end
            % Set optimization options:
            options = optimoptions('quadprog','Display','none','MaxIterations',20000);
           %  H = G'*G;
           H = zeros(nVPs+1,nVPs+1);
           H(end,end) = 1;
           f = zeros(size(H,1),1);
           
           A = [[G;-G],(-1)*ones(2*size(G,1),1)];
           b = zeros(size(A,1),1);
           
           options.ConvexCheck = 'on'; 
     %      tic;
           [Obj.OptimizationResults.optimalPrevalenceWeightsPriorToNormalization,fval,Obj.OptimizationResults.exitFlag] = quadprog(H,f,A,b,Aeq,beq,lb,ub,x0,options);
     %      toc;
           y_maxResi = Obj.OptimizationResults.optimalPrevalenceWeightsPriorToNormalization(end);
                   Obj.OptimizationResults.optimalPrevalenceWeightsPriorToNormalization = Obj.OptimizationResults.optimalPrevalenceWeightsPriorToNormalization(1:end-1);
                resi_G=G*Obj.OptimizationResults.optimalPrevalenceWeightsPriorToNormalization;
                maxResi = max(abs(resi_G));
           if Obj.OptimizationResults.exitFlag == -6  % try turn the convexcheck off
                  options.ConvexCheck = 'off';
                  [Obj.OptimizationResults.optimalPrevalenceWeightsPriorToNormalization,fval,Obj.OptimizationResults.exitFlag] = quadprog(H,f,A,b,Aeq,beq,lb,ub,x0,options);
                             y_maxResi = Obj.OptimizationResults.optimalPrevalenceWeightsPriorToNormalization(end);
                          Obj.OptimizationResults.optimalPrevalenceWeightsPriorToNormalization = Obj.OptimizationResults.optimalPrevalenceWeightsPriorToNormalization(1:end-1);
                        resi_G=G*Obj.OptimizationResults.optimalPrevalenceWeightsPriorToNormalization;
                        maxResi = max(abs(resi_G));
                  if abs(sum(Obj.OptimizationResults.optimalPrevalenceWeightsPriorToNormalization)-1)<=1e-12 & min(Obj.OptimizationResults.optimalPrevalenceWeightsPriorToNormalization)>0 % if returns a valid solution, keep it
                      Obj.OptimizationResults.exitFlag = 1;
                  else
                      Obj.OptimizationResults.exitFlag = -6;
                  end
           end
       elseif strcmpi(Obj.OptimOptions.optimizationAlgorithm,"quadprogEffN")
            % Specifying an 'x0' would result in a warning that x0 gets ignored in
            % 'lsqlin'
             %x0 = (1/nVPs)*ones(nVPs,1);   % x0 = [];
             x0 = Obj.InputVPop.pws';

            lb = zeros(nVPs,1); % lb = zeros(nVPs,1)+eps;
            ub = ones(nVPs,1);
            if ~isempty(Obj.OptimOptions.maxPrevalenceWeight) && isfinite(Obj.OptimOptions.maxPrevalenceWeight)
                A = spdiags(ones(nVPs,1),0,nVPs,nVPs);
                b = Obj.OptimOptions.maxPrevalenceWeight .* ones(1,nVPs);
            else
                A = [];
                b = [];
            end
            % Constraint that sum of prevalence weights should be 1:
            [subgroup ia ic]= unique(Obj.LinearProblemMatricesParticular.vpIsInSubgroup,'rows');
            subweights = Obj.LinearProblemMatricesParticular.SubgroupSumWeights;
            if size(subweights,1) ~= size(Obj.LinearProblemMatricesParticular.vpIsInSubgroup,1) % added to ensure starting from a previous subsetVPop with sometimes different dimensions
                 subweights = Obj.LinearProblemMatricesParticular.vpIsInSubgroup*x0;
            end
            Aeq = subgroup;
            beq = subweights(ia);

            % Set optimization options:            
            C = Obj.LinearProblemMatricesParticular.independentVarValsWeighted;
            d = Obj.LinearProblemMatricesParticular.observationValsWeighted;
            F = (C-d).*Obj.LinearProblemMatricesParticular.vpIsInSubgroup;
            ind=find(vecnorm(F,2,2)<1e-6);
            F(ind,:)=[];
%              subweights = Obj.LinearProblemMatricesParticular.SubgroupSumWeights;
             subweights(ind,:) = [];
            indexsubweight0=find(subweights<1e-6);
            if isempty(indexsubweight0)
                G = (1./subweights).*F;
            else
%                 subweights(indexsubweight0)=1;   % not to scale when subweight is 0, avoid 1/0 problem, i.e. when there is no VP in this subpop last iteration
                subweights(indexsubweight0)=NaN;
                subweights(indexsubweight0)=min(subweights);
                G = (1./subweights).*F;
            end
            % Set optimization options:
            options = optimoptions('quadprog','Display','none','MaxIterations',20000);
            H = G'*G;
            H = H/size(G,1);
            
            if (~isempty(Obj.lambda))
                lambda = Obj.lambda;
            else
                lambda=0;
            end
            H = H + lambda*eye(size(H,1));
           f = zeros(size(H,1),1);
           options.ConvexCheck = 'on'; 
        %   tic;
           [Obj.OptimizationResults.optimalPrevalenceWeightsPriorToNormalization,fval,Obj.OptimizationResults.exitFlag] = quadprog(H,f,A,b,Aeq,beq,lb,ub,x0,options);
        %   toc;
           
           if Obj.OptimizationResults.exitFlag == -6  % quadprog sometimes cannot continue if it is not strictly convex, when the convexcheck fail, we try turn the convexcheck off and run it
                  options.ConvexCheck = 'off';
                  [Obj.OptimizationResults.optimalPrevalenceWeightsPriorToNormalization,fval,Obj.OptimizationResults.exitFlag] = quadprog(H,f,A,b,Aeq,beq,lb,ub,x0,options);
                  % set the negative but within tolerance pws to 0
                  Obj.OptimizationResults.optimalPrevalenceWeightsPriorToNormalization(Obj.OptimizationResults.optimalPrevalenceWeightsPriorToNormalization<0 & abs(Obj.OptimizationResults.optimalPrevalenceWeightsPriorToNormalization)<=options.ConstraintTolerance) = 0;
                  % if returns a valid solution, keep the solution
                  if abs(sum(Obj.OptimizationResults.optimalPrevalenceWeightsPriorToNormalization)-1)<=1e-3
                      Obj.OptimizationResults.exitFlag = 1;
                  else
                      Obj.OptimizationResults.exitFlag = -6;
                  end
           end            
     elseif strcmpi(Obj.OptimOptions.optimizationAlgorithm,"fmincon")
            % Specifying an 'x0' from last fit
            x0 = Obj.InputVPop.pws';
            %  x0 = (1/nVPs)*ones(nVPs,1);   % x0 = [];
            % Constraint that sum of prevalence weights should be 1:
            [subgroup ia ic]= unique(Obj.LinearProblemMatricesParticular.vpIsInSubgroup,'rows');
            subweights = Obj.LinearProblemMatricesParticular.SubgroupSumWeights;
            Aeq = subgroup;
            beq = subweights(ia);
            
            % add inequality constraint: pw > 0          
            A = -eye(nVPs);
            b = zeros(nVPs,1);

            % Set up matrix:            
            C = Obj.LinearProblemMatricesParticular.independentVarValsWeighted;
            d = Obj.LinearProblemMatricesParticular.observationValsWeighted;
            F = (C-d).*Obj.LinearProblemMatricesParticular.vpIsInSubgroup;
            ind=find(vecnorm(F,2,2)<1e-6); % remove 0 rows, 1 rows
            F(ind,:)=[];
            subweights = Obj.LinearProblemMatricesParticular.SubgroupSumWeights;
            subweights(ind,:) = [];
            indexsubweight0=find(subweights<1e-6);
            if isempty(indexsubweight0)
                G = (1./subweights).*F;
            else
%                 subweights(indexsubweight0)=1;   % not to scale when subweight is 0, avoid 1/0 problem, i.e. when there is no VP in this subpop last iteration
                subweights(indexsubweight0)=NaN;
                subweights(indexsubweight0)=min(subweights);
                G = (1./subweights).*F;
            end
            H = G'*G;
            H = H/size(G,1);
            f = zeros(size(H,1),1);
           
            % Set optimization options:
            options = optimoptions(@fmincon,'Algorithm','interior-point',...
                     'SpecifyObjectiveGradient',true,'SpecifyConstraintGradient',true,...
                     'HessianFcn',@(x,lambda)hessian(x,lambda,H),...
                     'Display','none','MaxIterations',2000); %,'OptimalityTolerance',1e-18); %, 'StepTolerance',1e-24);
%             options = optimoptions(@fmincon,'Algorithm','sqp',...
%                      'SpecifyObjectiveGradient',true,'SpecifyConstraintGradient',true,...
%                      'HessianFcn',@(x,lambda)hessian(x,lambda,H),...
%                      'Display','none','MaxIterations',2000);
            warning('off','MATLAB:nearlySingularMatrix');
            warning('off','MATLAB:rankDeficientMatrix');     
            objct = @(x)objective(x,H);
            effN = Obj.OptimOptions.targetEffNConstraint;
            constr = @(x)constraints(x,effN,A,b,Aeq,beq);
        %    tic;
            [Obj.OptimizationResults.optimalPrevalenceWeightsPriorToNormalization,fval,Obj.OptimizationResults.exitFlag,~,lambda] = fmincon(objct,x0,[],[],[],[],[],[],constr,options);
            Obj.lambda = lambda.ineqnonlin(1);  
        %     toc;
        %    Obj.MSEfval = fval;   
            warning('on','MATLAB:nearlySingularMatrix');
            warning('on','MATLAB:rankDeficientMatrix');
        else
            error('The specified optimization algorithm is not recognized.');
        end
        
        if strcmpi(Obj.OptimOptions.optimizationAlgorithm,"fmincon")
            % if the fit failed to converge, set the solution as NaN:
            if Obj.OptimizationResults.exitFlag < 1 && Obj.OptimizationResults.exitFlag > -3
                Obj.OptimizationResults.optimalPrevalenceWeightsPriorToNormalization = nan*Obj.OptimizationResults.optimalPrevalenceWeightsPriorToNormalization;
            elseif Obj.OptimizationResults.exitFlag >= 1 || Obj.OptimizationResults.exitFlag == -3
                % Enforce non-negativity for values below options.ConstraintTolerance. because negative values won't pass following checks in the algorithm, e.g. weightedcorrs. sometimes fmincon continue to assign a negaitve value which should be 0. 
                % Might try to optimize X^2 to get around it. https://comp.soft-sys.matlab.narkive.com/VAgw8dSq/preventing-negative-values-in-fmincon
                Obj.OptimizationResults.optimalPrevalenceWeightsPriorToNormalization(Obj.OptimizationResults.optimalPrevalenceWeightsPriorToNormalization<=0 & abs(Obj.OptimizationResults.optimalPrevalenceWeightsPriorToNormalization)<=options.ConstraintTolerance) = 1e-16;
            end
            % normalize the prevalence weights to sum to 1
            Obj.OptimizationResults.optimalPrevalenceWeightsNormalized = Obj.OptimizationResults.optimalPrevalenceWeightsPriorToNormalization;
        elseif strcmpi(Obj.OptimOptions.optimizationAlgorithm,"quadprogEffN")
            % if the fit failed to converge, set the solution as NaN:
            if Obj.OptimizationResults.exitFlag < 1 
                Obj.OptimizationResults.optimalPrevalenceWeightsPriorToNormalization = nan*Obj.OptimizationResults.optimalPrevalenceWeightsPriorToNormalization;
            elseif Obj.OptimizationResults.exitFlag >= 1 
                % Enforce non-negativity for values below options.ConstraintTolerance. because negative values won't pass following checks in the algorithm, e.g. weightedcorrs. sometimes fmincon continue to assign a negaitve value which should be 0. 
                % Might try to optimize X^2 to get around it. https://comp.soft-sys.matlab.narkive.com/VAgw8dSq/preventing-negative-values-in-fmincon
                Obj.OptimizationResults.optimalPrevalenceWeightsPriorToNormalization(Obj.OptimizationResults.optimalPrevalenceWeightsPriorToNormalization<=0 & abs(Obj.OptimizationResults.optimalPrevalenceWeightsPriorToNormalization)<=options.ConstraintTolerance) = 1e-16;
            end
            % normalize the prevalence weights to sum to 1
            Obj.OptimizationResults.optimalPrevalenceWeightsNormalized = Obj.OptimizationResults.optimalPrevalenceWeightsPriorToNormalization;
        else
            Obj.OptimizationResults.optimalPrevalenceWeightsNormalized = ...
                 Obj.OptimizationResults.optimalPrevalenceWeightsPriorToNormalization./sum(Obj.OptimizationResults.optimalPrevalenceWeightsPriorToNormalization);
        end
         Obj.OptimizationResults.timeElapsedMinutes = toc(tictoc)/60;
        
            % nested function for setting up constraints in fmincon
             function [y,yeq,grady,gradyeq] = constraints(x,effN,A,b,Aeq,beq)
                    % inequality constratints: the constraint on effN
                    %                          n=length(x) non-negativity constraints
                    % equality constraints: subpopulation weight constraints (including the sum-to-one constraint).
                    n = length(x);
                    y = [x'*x - 1/effN, (A*x - b)'];
                    yeq = (Aeq*x - beq)';
                    grady = [2*x, A'];
                    gradyeq = Aeq';
             end 
            % nested function for setting up hessian in fmincon     
             function hess = hessian(x,lambda,H)
                    n = length(x);
                    hess = 2*(H + lambda.ineqnonlin(1)*eye(n));
             end   
            % nested function for setting up objective function in fmincon  
             function [y,grady] = objective(x,H)
                    y = x'*H*x;               
                    if nargout>1
                        grady = 2*H*x;
                    end
             end  
        
    end
    
    function Obj = runOptimizationfullvpop(Obj)
        % Start timer
        tictoc = tic();

        if isempty(Obj.LinearProblemMatricesParticular)
            Obj.LinearProblemMatricesParticular = Obj.LinearProblemMatrices;
        end
        
        nVPs = size(Obj.LinearProblemMatricesParticular.independentVarValsWeighted,2);        
        % Run the optimization
        if strcmpi(Obj.OptimOptions.optimizationAlgorithm,"quadprog") || strcmpi(Obj.OptimOptions.optimizationAlgorithm,"quadprogMaxResi")
            % Specifying an 'x0' would result in a warning that x0 gets ignored
            x0 = (1/nVPs)*ones(nVPs,1);
            if ~(isempty(Obj.OptimizationResults))
                if ~any(isnan(Obj.OptimizationResults.optimalPrevalenceWeightsNormalized))
                    x0 = Obj.OptimizationResults.optimalPrevalenceWeightsNormalized;
                end
            end
            lb = zeros(nVPs,1);
            ub = [];
            if ~isempty(Obj.OptimOptions.maxPrevalenceWeight) && isfinite(Obj.OptimOptions.maxPrevalenceWeight)
                A = spdiags(ones(nVPs,1),0,nVPs,nVPs);
                b = Obj.OptimOptions.maxPrevalenceWeight .* ones(1,nVPs);
            else
                A = [];
                b = [];
            end
            Aeq = ones(1,nVPs);
            beq = 1;
            % Set optimization options:               
            C = Obj.LinearProblemMatricesParticular.independentVarValsWeighted;
            d = Obj.LinearProblemMatricesParticular.observationValsWeighted; 
            fullvpop = find(sum(Obj.LinearProblemMatricesParticular.vpIsInSubgroup,2)==size(C,2));        
            C = C(fullvpop,:);
            d = d(fullvpop,:);
            F = (C-d).*Obj.LinearProblemMatricesParticular.vpIsInSubgroup(fullvpop,:);
            ind=find(vecnorm(F,2,2)<1e-6);
            F(ind,:)=[];
            subweights = Obj.LinearProblemMatricesParticular.SubgroupSumWeights(fullvpop,:);
            subweights(ind,:) = [];
            indexsubweight0=find(subweights<1e-6);
            if isempty(indexsubweight0)
                G = (1./subweights).*F;
            else
%                 subweights(indexsubweight0)=1;   % not to scale when subweight is 0, avoid 1/0 problem, i.e. when there is no VP in this subpop last iteration
                subweights(indexsubweight0)=NaN;
                subweights(indexsubweight0)=min(subweights);
                G = (1./subweights).*F;
            end
            
            % Set optimization options:
            options = optimoptions('quadprog','Display','none','MaxIterations',20000);
           
            %% bagging to get one si: do 6 round of full vpop fitting
            vpopNo = ceil(size(C,2)/size(G,1));
            nVPsPerBaggingIteration = round(size(C,2)/vpopNo);
            removeVPIndices = [];
            includedVPIndices = [];
            pwbagging=zeros(nVPs,vpopNo);  
            for vpopi = 1:vpopNo
                 removeVPIndices = [removeVPIndices includedVPIndices];
                 RemainedSample = setdiff(1:size(C,2),removeVPIndices);
                 if length(RemainedSample)>=nVPsPerBaggingIteration
                        includedVPIndices = datasample(RemainedSample,nVPsPerBaggingIteration,'Replace',false);
                 else
                        includedVPIndices = RemainedSample;
                 end
                 Gi = G(:,includedVPIndices);
                 H = Gi'*Gi;
                 f = zeros(size(H,1),1);
                 Aeq = ones(1,length(includedVPIndices));
                 lb = zeros(length(includedVPIndices),1);
                 options.ConvexCheck = 'on'; 
                 [Obj.OptimizationResults.optimalPrevalenceWeightsPriorToNormalization,fval,Obj.OptimizationResults.exitFlag] = quadprog(H,f,A,b,Aeq,beq,lb,ub,x0,options);
                 if Obj.OptimizationResults.exitFlag == -6  % try turn the convexcheck off
                        options.ConvexCheck = 'off';
                        [Obj.OptimizationResults.optimalPrevalenceWeightsPriorToNormalization,fval,Obj.OptimizationResults.exitFlag] = quadprog(H,f,A,b,Aeq,beq,lb,ub,x0,options);
                        if abs(sum(Obj.OptimizationResults.optimalPrevalenceWeightsPriorToNormalization)-1)<=1e-12 & min(Obj.OptimizationResults.optimalPrevalenceWeightsPriorToNormalization)>0 % if returns a valid solution, keep it
                                 Obj.OptimizationResults.exitFlag = 1;
                        else
                                 Obj.OptimizationResults.exitFlag = -6;
                        end
                  end                      
                  % if the fit failed to converge, set the solution as NaN:
                  if Obj.OptimizationResults.exitFlag < 1
                         Obj.OptimizationResults.optimalPrevalenceWeightsPriorToNormalization = nan*Obj.OptimizationResults.optimalPrevalenceWeightsPriorToNormalization;
                  end
                  pwbagging(includedVPIndices,vpopi) = Obj.OptimizationResults.optimalPrevalenceWeightsPriorToNormalization;
            end
            Obj.OptimizationResults.optimalPrevalenceWeightsPriorToNormalization = nanmean(pwbagging,2); 
      elseif strcmpi(Obj.OptimOptions.optimizationAlgorithm,"quadprogEffN")
            % Specifying an 'x0' would result in a warning that x0 gets ignored
            % x0 = (1/nVPs)*ones(nVPs,1);
            x0 = Obj.InputVPop.pws';
            Aeq = ones(1,nVPs);
            beq = 1;
            
            if ~isempty(Obj.OptimOptions.maxPrevalenceWeight) && isfinite(Obj.OptimOptions.maxPrevalenceWeight)
                A = spdiags(ones(nVPs,1),0,nVPs,nVPs);
                b = Obj.OptimOptions.maxPrevalenceWeight .* ones(1,nVPs);
            else
                A = [];
                b = [];
            end
            
            lb = zeros(nVPs,1); % lb = zeros(nVPs,1)+eps;
            ub = ones(nVPs,1);

            % set up matrix:               
            C = Obj.LinearProblemMatricesParticular.independentVarValsWeighted;
            d = Obj.LinearProblemMatricesParticular.observationValsWeighted; 
            fullvpop = find(sum(Obj.LinearProblemMatricesParticular.vpIsInSubgroup,2)==size(C,2));        
            C = C(fullvpop,:);
            d = d(fullvpop,:);
            F = (C-d).*Obj.LinearProblemMatricesParticular.vpIsInSubgroup(fullvpop,:);
            ind=find(vecnorm(F,2,2)<1e-6);
            F(ind,:)=[];
            subweights = Obj.LinearProblemMatricesParticular.SubgroupSumWeights(fullvpop,:);
            subweights(ind,:) = [];
            indexsubweight0=find(subweights<1e-6);
            if isempty(indexsubweight0)
                G = (1./subweights).*F;
            else
%                 subweights(indexsubweight0)=1;   % not to scale when subweight is 0, avoid 1/0 problem, i.e. when there is no VP in this subpop last iteration
                subweights(indexsubweight0)=NaN;
                subweights(indexsubweight0)=min(subweights);
                G = (1./subweights).*F;
            end
            H = G'*G;
            H = H/size(G,1);
            % Set optimization options:
            options = optimoptions('quadprog','Display','none','MaxIterations',20000);
            if (~isempty(Obj.lambda))
                lambda = Obj.lambda;
            else
                lambda=0;
            end
            H = H + lambda*eye(size(H,1));
            f = zeros(size(H,1),1);
                 
           options.ConvexCheck = 'on'; 
             warning('off','optim:quadprog:NullHessian');
%              diary('linearOut');
%            tic;
           [Obj.OptimizationResults.optimalPrevalenceWeightsPriorToNormalization,fval,Obj.OptimizationResults.exitFlag] = quadprog(H,f,A,b,Aeq,beq,lb,ub,x0,options);
%            toc;
           
           if Obj.OptimizationResults.exitFlag == -6  
               % quadprog sometimes cannot continue if it is numerically not strictly convex, but actually often it can still reach an feasible solution. 
               % https://www.mathworks.com/matlabcentral/answers/420851-quadprog-new-feature
               % So when the convexcheck fail, we try turn the convexcheck off and run it again             
                  options.ConvexCheck = 'off';
                  [Obj.OptimizationResults.optimalPrevalenceWeightsPriorToNormalization,fval,Obj.OptimizationResults.exitFlag] = quadprog(H,f,A,b,Aeq,beq,lb,ub,x0,options);
                  % set the negative but within tolerance pws to 0
                  Obj.OptimizationResults.optimalPrevalenceWeightsPriorToNormalization(Obj.OptimizationResults.optimalPrevalenceWeightsPriorToNormalization<0 & abs(Obj.OptimizationResults.optimalPrevalenceWeightsPriorToNormalization)<=options.ConstraintTolerance) = 0;
                  % if returns a valid solution, keep the solution
                  if abs(sum(Obj.OptimizationResults.optimalPrevalenceWeightsPriorToNormalization)-1)<=1e-3
                      Obj.OptimizationResults.exitFlag = 1;
                  else
                      Obj.OptimizationResults.exitFlag = -6;
                  end
           end
           warning('on','optim:quadprog:NullHessian');
     elseif strcmpi(Obj.OptimOptions.optimizationAlgorithm,"fmincon") 
            % Specifying an 'x0' would result in a warning that x0 gets ignored
            % x0 = (1/nVPs)*ones(nVPs,1);
            x0 = Obj.InputVPop.pws';
            Aeq = ones(1,nVPs);
            beq = 1;
            
            % add inequality constraint: all subpop weights >= minSubWeightConstraint
            [subgroup , ~, ~]= unique(Obj.LinearProblemMatricesParticular.vpIsInSubgroup,'rows');
            minSubWeightConstraint = Obj.OptimOptions.minSubWeightConstraint;
            A = [-eye(nVPs);-subgroup];
            b = [zeros(nVPs,1);-minSubWeightConstraint*ones(size(subgroup,1),1)];

            % set up matrix:               
            C = Obj.LinearProblemMatricesParticular.independentVarValsWeighted;
            d = Obj.LinearProblemMatricesParticular.observationValsWeighted; 
            fullvpop = find(sum(Obj.LinearProblemMatricesParticular.vpIsInSubgroup,2)==size(C,2));        
            C = C(fullvpop,:);
            d = d(fullvpop,:);
            F = (C-d).*Obj.LinearProblemMatricesParticular.vpIsInSubgroup(fullvpop,:);
            ind=find(vecnorm(F,2,2)<1e-6);
            F(ind,:)=[];
            subweights = Obj.LinearProblemMatricesParticular.SubgroupSumWeights(fullvpop,:);
            subweights(ind,:) = [];
            indexsubweight0=find(subweights<1e-6);
            if isempty(indexsubweight0)
                G = (1./subweights).*F;
            else
%                 subweights(indexsubweight0)=1;   % not to scale when subweight is 0, avoid 1/0 problem, i.e. when there is no VP in this subpop last iteration
                subweights(indexsubweight0)=NaN;
                subweights(indexsubweight0)=min(subweights);
                G = (1./subweights).*F;
            end
            H = G'*G;
            H = H/size(G,1);
            f = zeros(size(H,1),1);
                 
            % Set optimization options:
            options = optimoptions(@fmincon,'Algorithm','interior-point',...
                     'SpecifyObjectiveGradient',true,'SpecifyConstraintGradient',true,...
                     'HessianFcn',@(x,lambda)hessian(x,lambda,H),...
                     'Display','none','MaxIterations',10000);
            warning('off','MATLAB:nearlySingularMatrix'); % can investigate later
            warning('off','MATLAB:rankDeficientMatrix');  
            objct = @(x)objective(x,H);
            effN = Obj.OptimOptions.targetEffNConstraint;
            constr = @(x)constraints(x,effN,A,b,Aeq,beq);
        %     tic;
            [Obj.OptimizationResults.optimalPrevalenceWeightsPriorToNormalization,fval,Obj.OptimizationResults.exitFlag,~,lambda] = fmincon(objct,x0,[],[],[],[],[],[],constr,options);
        %    toc; 
            Obj.lambda = lambda.ineqnonlin(1);   
%             Obj.MSEfval = fval;   
            warning('on','MATLAB:nearlySingularMatrix');
            warning('on','MATLAB:rankDeficientMatrix');  
        else
            error('The specified optimization algorithm is not recognized.');
        end
        
        if strcmpi(Obj.OptimOptions.optimizationAlgorithm,"fmincon")
            % if the fit failed to converge, set the solution as NaN:
            if Obj.OptimizationResults.exitFlag < 1 && Obj.OptimizationResults.exitFlag > -3
                Obj.OptimizationResults.optimalPrevalenceWeightsPriorToNormalization = nan*Obj.OptimizationResults.optimalPrevalenceWeightsPriorToNormalization;
            elseif Obj.OptimizationResults.exitFlag >= 1 || Obj.OptimizationResults.exitFlag == -3
                % Enforce non-negativity for values below options.ConstraintTolerance. because negative values won't pass following checks in the algorithm, e.g. weightedcorrs. sometimes fmincon continue to assign a negaitve value which should be 0. 
                % Might try to optimize X^2 to get around it. https://comp.soft-sys.matlab.narkive.com/VAgw8dSq/preventing-negative-values-in-fmincon
                Obj.OptimizationResults.optimalPrevalenceWeightsPriorToNormalization(Obj.OptimizationResults.optimalPrevalenceWeightsPriorToNormalization<=0 & abs(Obj.OptimizationResults.optimalPrevalenceWeightsPriorToNormalization)<=options.ConstraintTolerance) = 1e-16;
            end
            % normalize the prevalence weights to sum to 1
            Obj.OptimizationResults.optimalPrevalenceWeightsNormalized = Obj.OptimizationResults.optimalPrevalenceWeightsPriorToNormalization;
       elseif strcmpi(Obj.OptimOptions.optimizationAlgorithm,"quadprogEffN")
            % if the fit failed to converge, set the solution as NaN:
            if Obj.OptimizationResults.exitFlag < 1 
                Obj.OptimizationResults.optimalPrevalenceWeightsPriorToNormalization = nan*Obj.OptimizationResults.optimalPrevalenceWeightsPriorToNormalization;
            elseif Obj.OptimizationResults.exitFlag >= 1 
                % Enforce non-negativity for values below options.ConstraintTolerance. because negative values won't pass following checks in the algorithm, e.g. weightedcorrs. sometimes fmincon continue to assign a negaitve value which should be 0. 
                % Might try to optimize X^2 to get around it. https://comp.soft-sys.matlab.narkive.com/VAgw8dSq/preventing-negative-values-in-fmincon
                Obj.OptimizationResults.optimalPrevalenceWeightsPriorToNormalization(Obj.OptimizationResults.optimalPrevalenceWeightsPriorToNormalization<=0 & abs(Obj.OptimizationResults.optimalPrevalenceWeightsPriorToNormalization)<=options.ConstraintTolerance) = 1e-16;
            end
            % normalize the prevalence weights to sum to 1
            Obj.OptimizationResults.optimalPrevalenceWeightsNormalized = Obj.OptimizationResults.optimalPrevalenceWeightsPriorToNormalization;
       else
            Obj.OptimizationResults.optimalPrevalenceWeightsNormalized = ...
                 Obj.OptimizationResults.optimalPrevalenceWeightsPriorToNormalization./sum(Obj.OptimizationResults.optimalPrevalenceWeightsPriorToNormalization);
       end
        Obj.OptimizationResults.timeElapsedMinutes = toc(tictoc)/60;
        
            % nested function for setting up constraints in fmincon
             function [y,yeq,grady,gradyeq] = constraints(x,effN,A,b,Aeq,beq)
                    % inequality constratints: the constraint on effN
                    %                          n=length(x) non-negativity constraints
                    % equality constraints: subpopulation weight constraints (including the sum-to-one constraint).
                    n = length(x);
                    y = [x'*x - 1/effN, (A*x - b)'];
                    yeq = (Aeq*x - beq)';
                    grady = [2*x, A'];
                    gradyeq = Aeq';
             end 
        	% nested function for setting up hessian in fmincon     
             function hess = hessian(x,lambda,H)
                    n = length(x);
                    hess = 2*(H + lambda.ineqnonlin(1)*eye(n));
             end   
            % nested function for setting up objective function in fmincon  
             function [y,grady] = objective(x,H)
                    y = x'*H*x;               
                    if nargout>1
                        grady = 2*H*x;
                    end
             end  
    end
    
    function Obj = performPostOptimizationAnalysis(Obj)
    % Computes some statistics based on the optimal prevalence weight
        
        Obj.OptimizationResults.sumOfOptimalPrevalenceWeightsPriorToNormalization = sum(Obj.OptimizationResults.optimalPrevalenceWeightsPriorToNormalization);
        
        [Obj.OptimizationResults.residualsAfterRenormalizationOfPrevalenceWeights,Obj.OptimizationResults.rmseConsideringOptimalPrevalenceWeightsNormalized,~] = ...
            Obj.calculateResidualsAndRootMeanSquaredErrorForSpecifiedPWs();
        
%         % The following calculation follows from the calculations performed
%         % in 'lsqnonneg':
%         optimizationResults.lagrangeMultipliersAfterRenormalizationOfPrevalenceWeights = ...
%             Obj.LinearProblemMatrices.independentVarVals'*optimizationResults.residualsAfterRenormalizationOfPrevalenceWeights;
        
        Obj.OptimizationResults.effectiveN = Obj.calculateEffN(Obj.OptimizationResults.optimalPrevalenceWeightsNormalized);
    end
    
    function Obj = constructNewVPopPostOptimization(Obj)
    % Constructs a new VPop corresponding to the optimal prevalence weights
        
        % Sanity check that prevalence weights sum to one
        if abs(sum(Obj.OptimalPrevalenceWeightsNormalized)-1)  > 1e-3
            error('Prevalence weights do not sum to 1 after renormalization.')
        end
        
        % Copy the old VPop and replace the prevalence weights:
        Obj.OptimizedVPop = Obj.InputVPop;
        Obj.OptimizedVPop.pws = (Obj.OptimalPrevalenceWeightsNormalized(:))';

        % recalculate the tables and goodness of fit 
        Obj.OptimizedVPop=addPredTableVals(Obj.OptimizedVPop);
        Obj.OptimizedVPop=evaluateGOF(Obj.OptimizedVPop);
    end
    
    function [residuals,rmse,weights] = calculateResidualsAndRootMeanSquaredErrorForSpecifiedPWs(Obj)
    % Calculate residuals and RMSE corresponding to the specified
    % prevalence weights
        
        % The statistics are calculated according to the matrices in
        % 'Obj.LinearProblemMatricesParticular'. If this property is empty,
        % set it equal to 'Obj.LinearProblemMatrices'
        if isempty(Obj.LinearProblemMatricesParticular)
            Obj.LinearProblemMatricesParticular = Obj.LinearProblemMatrices;
        end
    
        weights = Obj.LinearProblemMatricesParticular.ultimateWeights;
        
        % The actual residuals and RMSE used in the fitting:
       % residuals = Obj.LinearProblemMatricesParticular.observationValsWeighted - Obj.LinearProblemMatricesParticular.independentVarValsWeighted*Obj.OptimalPrevalenceWeightsNormalized(:);
        % weighted mean (note that this is a weighted mean since the squared residuals are weighted with weights which sum to 1: (this will be NaN when the weights sum to zero):
        
        vpPrevalenceWeights=Obj.OptimizationResults.optimalPrevalenceWeightsNormalized;
        C = Obj.LinearProblemMatricesParticular.independentVarValsWeighted; 
        d = Obj.LinearProblemMatricesParticular.observationValsWeighted;
        SubgroupSumWeights = Obj.LinearProblemMatrices.vpIsInSubgroup*vpPrevalenceWeights;
        Cactual = (1./SubgroupSumWeights).*(C.*Obj.LinearProblemMatricesParticular.vpIsInSubgroup); 
        residuals = Cactual*vpPrevalenceWeights-d;
        residuals(find(isnan(residuals))) = 0; % added this in case there are one or less VPs in the row, mn and std will be NaNs
        mse = sum(residuals.^2);
        rmse = sqrt(mse);
        
        % for debugging
        % In the fitting, each residual is scaled according to the number
        % of observations in the data group, so that the influence of
        % a data group on the RMSE doesn't depend on the number of
        % observations in the data group. However, if we're analyzing a
        % particular data group's residuals after fitting, it would be more
        % informative to consider the unscaled version, which is calculated
        % below:
%         residualsCorrectedForN = residuals.*sqrt(Obj.LinearProblemMatricesParticular.numObservationsInDataGroup);
%         rmseCorrectedForN = sqrt(mean(residualsCorrectedForN.^2));
        
        % for debugging:
%         uniqueDataGroups = unique(Obj.LinearProblemMatricesParticular.dataGroup);
%         uniqueDataGroups = uniqueDataGroups(:);
%         nDataGroups = length(uniqueDataGroups);
%         rmseCorrectedForNByDataGroup = nan(nDataGroups,1);
%         dataGroupDescriptions = cell(nDataGroups,1);
%         for iGrp = 1:nDataGroups
%             iMaskCrp = Obj.LinearProblemMatricesParticular.dataGroup == iGrp;
%             residualsParticular = residualsCorrectedForN(iMaskCrp);
%             rmseCorrectedForNByDataGroup(iGrp) = sqrt(mean(residualsParticular.^2));
%             observationDescriptionsParticular = Obj.LinearProblemMatricesParticular.observationDescriptions(iMaskCrp);
%             dataGroupDescriptions{iGrp} = observationDescriptionsParticular{1};
%         end
%         rmseByDataGroupTable = table(uniqueDataGroups,dataGroupDescriptions,rmseCorrectedForNByDataGroup,'VariableNames',{'dataGroup','firstObservationDescription','rmseCorrectedForNForDataGroup'});
%         
%         residualsDiagnosticsTable = table(...
%             Obj.LinearProblemMatricesParticular.dataGroup,...
%             Obj.LinearProblemMatricesParticular.observationDescriptions,....
%             residualsCorrectedForN,...
%             Obj.LinearProblemMatricesParticular.numObservationsInDataGroup,...
%             Obj.LinearProblemMatricesParticular.observationVals,...
%             (Obj.LinearProblemMatricesParticular.independentVarVals*prevalenceWeights(:)),...
%             Obj.LinearProblemMatricesParticular.observationValsWeighted.*sqrt(Obj.LinearProblemMatricesParticular.numObservationsInDataGroup),...
%             (Obj.LinearProblemMatricesParticular.independentVarValsWeighted*prevalenceWeights(:)).*sqrt(Obj.LinearProblemMatricesParticular.numObservationsInDataGroup),...
%             'VariableNames',{'dataGroup','observationDescriptions','residualsCorrectedForN','numObservationsInDataGroup','observationVal','predVal','observationValWeighted_x_N','predValWeighted_x_N'});
%         
%         residualsDiagnosticsTable = outerjoin(residualsDiagnosticsTable,rmseByDataGroupTable,'Keys',{'dataGroup'},'RightVariables',{'rmseCorrectedForNForDataGroup'});
%         residualsDiagnosticsTable = sortrows(residualsDiagnosticsTable,'rmseCorrectedForNForDataGroup');
    end

end

end


%% INTERNAL HELPER FUNCTIONS

function createVPopPlots(vPop,figName,figFilepathPrefix,closeFigsAutomatically)
% Internal helper function for creating VPop plots

    plotVPopFunctionHandles = {@plotBinVPop,@plotBRVPop,@plotMnSDVPop,@plotDistCDFVPop,@plotDist2DVPop};
    for iFunc = 1:length(plotVPopFunctionHandles)
        try
            figHandle = plotVPopFunctionHandles{iFunc}(vPop);
        catch
            warning(['Unable to generate plots using ' func2str(plotVPopFunctionHandles{iFunc}) '. This could be because there is no data of this type to plot.']);
            continue;
        end
        if isempty(figHandle)
            warning(['Unable to generate plots using ' func2str(plotVPopFunctionHandles{iFunc}) '. This could be because there is no data of this type to plot.']);
            continue;
        end
        if ~isempty(figName)
            figHandle.Name = [figName ' - ' func2str(plotVPopFunctionHandles{iFunc})];
        end
        linearCalibrationSaveFig(figHandle,[figFilepathPrefix '_' func2str(plotVPopFunctionHandles{iFunc})]);
        if closeFigsAutomatically
            close(figHandle);
        end
    end
end

function linearCalibrationSaveFig(figHandle,figFilepathWithoutExtension)
% Internal helper function for saving figures
    figure(figHandle);
    set(figHandle, 'Position', [1 1 1680 1050]); % make the figure large before saving
    savefig(figHandle, figFilepathWithoutExtension);
    % export_fig figFilepathWithoutExtension -png -jpg -tif -transparent;
    print(figHandle, [figFilepathWithoutExtension '.tif'],'-dtiff','-r300');
end

function linearProblemMatrices = incorporateLinearProblemMatricesWeights(linearProblemMatrices,optimOptions)
% Internal helper function used during the construction of the linear
% problem matrices for calculating and incorporating the observation
% weights
        
    % Calculate the total weight for each observation:
    % The weights need to be square-rooted, since otherwise they will
    % get squared when the residuals are squared in the least
    % squares routine:
    % this is simplied for only accounting for the different observational weights
         linearProblemMatrices.ultimateWeights = linearProblemMatrices.expWeight.* linearProblemMatrices.observationWeights;
%          ...
%         sqrt( linearProblemMatrices.expWeight.* linearProblemMatrices.observationWeights );
%      linearProblemMatrices.ultimateWeights = ...
%         sqrt( linearProblemMatrices.expWeight.* linearProblemMatrices.observationWeights );

%     linearProblemMatrices.ultimateWeights = ...
%         sqrt( linearProblemMatrices.expWeight .* linearProblemMatrices.observationWeights .* linearProblemMatrices.dataGroupWeights ./ linearProblemMatrices.numObservationsInDataGroup );
% 
%     % normalize so that the squared weights sum to 1
%     normalizationDenominator = sqrt(sum(linearProblemMatrices.ultimateWeights.^2));
	% !!! the sqrt can be taken in the following line instead, to be more
	% efficient
  %  linearProblemMatrices.ultimateWeights = linearProblemMatrices.ultimateWeights/normalizationDenominator;
    
    % Relative transformation of response values, if performed, is performed after 
    % normalization of weights (since we don't want the value of one
    % observation to change the weight of another, and since it is not
    % correct to normalize based on the transformation)
%     if strcmpi(optimOptions.responseValTransformation,"relative")
%         % I don't think the following needs to be square rooted,
%         % since we are assuming that the residuals, not the squared
%         % residuals, are proportional to absolute values.
%         linearProblemMatrices.ultimateWeights = ...
%             abs(linearProblemMatrices.ultimateWeights ./ linearProblemMatrices.observationVals);
%     end
    
    % factor in weights
    ultimateWeightsMatrix = spdiags(linearProblemMatrices.ultimateWeights,0,length(linearProblemMatrices.ultimateWeights),length(linearProblemMatrices.ultimateWeights));
    linearProblemMatrices.independentVarValsWeighted = ultimateWeightsMatrix*linearProblemMatrices.independentVarVals;
    linearProblemMatrices.observationValsWeighted = ultimateWeightsMatrix*linearProblemMatrices.observationVals;
end

function generateAndSaveCrossValidationPlots(outputDir,resultsOut,p)
% Internal helper function for generating the cross-validation plots

    %% Define plot visual parameters:
    plotParameters.baggingRMSE.rmseWholeFit.displayName = 'whole-fit RMSE';
    plotParameters.baggingRMSE.rmseWholeFit.lineStyle = '-';
    plotParameters.baggingRMSE.rmseWholeFit.marker = 'o';
    plotParameters.baggingRMSE.rmseWholeFit.color = [0 0 255]/255;
    plotParameters.baggingRMSE.rmseTraining.displayName = 'CV training RMSE';
    plotParameters.baggingRMSE.rmseTraining.lineStyle = ':';
    plotParameters.baggingRMSE.rmseTraining.marker = '+';
    plotParameters.baggingRMSE.rmseTraining.color = [0 147 147]/255;
    plotParameters.baggingRMSE.rmseTest.displayName = 'CV test RMSE';
    plotParameters.baggingRMSE.rmseTest.lineStyle = '--';
    plotParameters.baggingRMSE.rmseTest.marker = 'square';
    plotParameters.baggingRMSE.rmseTest.color = [0 255 255]/255;
    plotParameters.baggingRMSE.rmsePrediction.displayName = 'prediction RMSE';
    plotParameters.baggingRMSE.rmsePrediction.lineStyle = '-.';
    plotParameters.baggingRMSE.rmsePrediction.marker = 'x';
    plotParameters.baggingRMSE.rmsePrediction.color = [round(255/2) 0 255]/255;
    plotParameters.baggingEffN.effNWholeFit.displayName = 'effective n';
    plotParameters.baggingEffN.effNWholeFit.lineStyle = '-';
    plotParameters.baggingEffN.effNWholeFit.marker = 'o';
    plotParameters.baggingEffN.effNWholeFit.color = [255 0 0]/255;
    plotParameters.baggingGOF.gofWholeFit.displayName = 'whole-fit GOF';
    plotParameters.baggingGOF.gofWholeFit.lineStyle = '-';
    plotParameters.baggingGOF.gofWholeFit.marker = 'o';
    plotParameters.baggingGOF.gofWholeFit.color = [0 255 255]/255;
    plotParameters.inputVPopRMSE = plotParameters.baggingRMSE;
    plotParameters.inputVPopEffN = plotParameters.baggingEffN;
    plotParameters.inputVPopGOF = plotParameters.baggingGOF;
    plotParameters.noBaggingRMSE = plotParameters.baggingRMSE;
    plotParameters.noBaggingEffN = plotParameters.baggingEffN;
    plotParameters.noBaggingGOF = plotParameters.baggingGOF;
    
    
    %% Plot the results RMSE and effective N:
    figfilename = 'baggingCrossValidationRMSEAndEffN';
    figHandle = figure;
    xlabel('fraction VPs per bagging sample')
    yyaxis left;
    hold on;
    makeLinePlots(plotParameters.baggingRMSE,resultsOut.baggingRMSE);
    ylabel('root-mean-squared error (RMSE)');
    % Place the bottom of the y-axis at the origin:
    yLimMem = ylim();
    ylim([0 yLimMem(2)]);
    yyaxis right;
    hold on;
    makeLinePlots(plotParameters.baggingEffN,resultsOut.baggingEffN);
    ylabel('effective num. virtual patients (n_{eff})');
    % Place the bottom of the y-axis at the origin:
    yLimMem = ylim();
    ylim([0 yLimMem(2)]);
    ax = gca();
    ax.XScale = 'log';
    ax.FontSize = 20;
    grid on;
    figFilepathWithoutExtension = [outputDir figfilename 'WithoutLegend'];
    linearCalibrationSaveFig(figHandle,figFilepathWithoutExtension);
    legend;
    figFilepathWithoutExtension = [outputDir figfilename 'WithLegend'];
    linearCalibrationSaveFig(figHandle,figFilepathWithoutExtension);

    %% Make RMSE/EffN bar plot
    
    % First, plot RMSE:
    
    figfilename = 'barRMSEandEffN';
    
    leftAxisDescription = 'baggingRMSE';
    rightAxisDescription = 'baggingEffN';
        
    subgroupTable = constructSubGroupTable(plotParameters,{leftAxisDescription,rightAxisDescription});
    [uniqueSubTypes,uniqueSubTypesIndices,~] = unique(subgroupTable.descriptionsSubType,'stable');
    subgroupLabels = (subgroupTable.displayNames(uniqueSubTypesIndices))';
    subgroupColors = subgroupTable.barColors(uniqueSubTypesIndices);
    
    groupLabels = {};
    barYLeft = [];

    barYLeft = [barYLeft ; constructBarYRow(resultsOut,subgroupTable,uniqueSubTypes,'inputVPopRMSE',1)];
    groupLabels = [groupLabels ; {'input VPop'}];
    
    barYLeft = [barYLeft ; constructBarYRow(resultsOut,subgroupTable,uniqueSubTypes,'noBaggingRMSE',1)];
    groupLabels = [groupLabels ; {'no bagging'}];
    
    for iFraction = 1:height(resultsOut.baggingRMSE)
        barYLeft = [barYLeft ; constructBarYRow(resultsOut,subgroupTable,uniqueSubTypes,'baggingRMSE',iFraction)];
        groupLabels = [groupLabels ; {['bagging with fraction: ' num2str(resultsOut.baggingRMSE.fractionVPsPerBaggingIteration(iFraction))]}];
    end
    
    figHandle = figure;
    yyaxis left;
    barHandleLeft = bar(barYLeft);
    for iBarLeft = 1:length(barHandleLeft)
        barHandleLeft(iBarLeft).FaceColor = subgroupColors{iBarLeft};
    end
    
    ylabel({'root-mean-squared'; 'error (RMSE)'});
    
    % Place the bottom of the y-axis at the origin:
    yLimMem = ylim();
    ylim([0 yLimMem(2)]);
    
    % Second, plot EffN:
    
    barYRight = [];
    
    barYRight = [barYRight ; constructBarYRow(resultsOut,subgroupTable,uniqueSubTypes,'inputVPopEffN',1)];
    
    barYRight = [barYRight ; constructBarYRow(resultsOut,subgroupTable,uniqueSubTypes,'noBaggingEffN',1)];
    
    for iFraction = 1:height(resultsOut.baggingEffN)
        barYRight = [barYRight ; constructBarYRow(resultsOut,subgroupTable,uniqueSubTypes,'baggingEffN',iFraction)];
    end
    
    yyaxis right;
    barHandleRight = bar(barYRight);
    for iBarRight = 1:length(barHandleRight)
        barHandleRight(iBarRight).FaceColor = subgroupColors{iBarRight};
    end
    
    ylabel({'effective num. virtual patients';'(n_{eff}) for the whole-fit case'});
    
    % Place the bottom of the y-axis at the origin:
    yLimMem = ylim();
    ylim([0 yLimMem(2)]);
    
    ax = gca();
    ax.XTick = 1:length(groupLabels);
    ax.XTickLabel = groupLabels';
    xtickangle(45);
    ax.FontSize = 20;
    ax.YGrid = 'on';
    figFilepathWithoutExtension = [outputDir figfilename 'WithoutLegend'];
    linearCalibrationSaveFig(figHandle,figFilepathWithoutExtension);
    legend(subgroupLabels);
    figFilepathWithoutExtension = [outputDir figfilename 'WithLegend'];
    linearCalibrationSaveFig(figHandle,figFilepathWithoutExtension);
    

    %% Make GOF/EffN Bar plot
    
    % First, plot RMSE:
    
    figfilename = 'barGOFandEffN';

    leftAxisDescription = 'baggingGOF';
    rightAxisDescription = 'baggingEffN';
    
    subgroupTable = constructSubGroupTable(plotParameters,{leftAxisDescription,rightAxisDescription});
    [uniqueSubTypes,uniqueSubTypesIndices,~] = unique(subgroupTable.descriptionsSubType,'stable');
    subgroupLabels = (subgroupTable.displayNames(uniqueSubTypesIndices))';
    subgroupColors = subgroupTable.barColors(uniqueSubTypesIndices);
    
    groupLabels = {};
    barYLeft = [];
    
    barYLeft = [barYLeft ; constructBarYRow(resultsOut,subgroupTable,uniqueSubTypes,'inputVPopGOF',1)];
    groupLabels = [groupLabels ; {'input VPop'}];
    
    barYLeft = [barYLeft ; constructBarYRow(resultsOut,subgroupTable,uniqueSubTypes,'noBaggingGOF',1)];
    groupLabels = [groupLabels ; {'no bagging'}];
    
    for iFraction = 1:height(resultsOut.baggingGOF)
        barYLeft = [barYLeft ; constructBarYRow(resultsOut,subgroupTable,uniqueSubTypes,'baggingGOF',iFraction)];
        groupLabels = [groupLabels ; {['bagging with fraction: ' num2str(resultsOut.baggingGOF.fractionVPsPerBaggingIteration(iFraction))]}];
    end
    
    figHandle = figure;
    yyaxis left;
    barHandleLeft = bar(barYLeft);
    for iBarLeft = 1:length(barHandleLeft)
        barHandleLeft(iBarLeft).FaceColor = subgroupColors{iBarLeft};
    end
    
    ylabel({'goodness-of-fit (GOF) p-value';'for the whole-fit case'});
    
    % Place the bottom of the y-axis at the origin:
    yLimMem = ylim();
    ylim([0 yLimMem(2)]);
    
    % Second, plot EffN:
    
    barYRight = [];
    
    barYRight = [barYRight ; constructBarYRow(resultsOut,subgroupTable,uniqueSubTypes,'inputVPopEffN',1)];
    
    barYRight = [barYRight ; constructBarYRow(resultsOut,subgroupTable,uniqueSubTypes,'noBaggingEffN',1)];
    
    for iFraction = 1:height(resultsOut.baggingEffN)
        barYRight = [barYRight ; constructBarYRow(resultsOut,subgroupTable,uniqueSubTypes,'baggingEffN',iFraction)];
    end
    
    yyaxis right;
    barHandleRight = bar(barYRight);
    for iBarRight = 1:length(barHandleRight)
        barHandleRight(iBarRight).FaceColor = subgroupColors{iBarRight};
    end
    
    ylabel({'effective num. virtual patients (n_{eff})';'for the whole-fit case'});
    
    % Place the bottom of the y-axis at the origin:
    yLimMem = ylim();
    ylim([0 yLimMem(2)]);
    
    ax = gca();
    ax.XTick = 1:length(groupLabels);
    ax.XTickLabel = groupLabels';
    xtickangle(45);
    ax.FontSize = 20;
    ax.YGrid = 'on';
    figFilepathWithoutExtension = [outputDir figfilename 'WithoutLegend'];
    linearCalibrationSaveFig(figHandle,figFilepathWithoutExtension);
    legend(subgroupLabels);
    figFilepathWithoutExtension = [outputDir figfilename 'WithLegend'];
    linearCalibrationSaveFig(figHandle,figFilepathWithoutExtension);
    
    
    %%  Plot the GOF
    if p.Results.calculateGOF
        figfilename = 'baggingGOFAndEffN';
        figHandle = figure;
        xlabel('fraction VPs per bagging sample')
        yyaxis left;
        hold on;
        makeLinePlots(plotParameters.baggingGOF,resultsOut.baggingGOF);
        ylabel({'goodness-of-fit (GOF) p-value';'for the whole-fit case'});
        % Place the bottom of the y-axis at the origin:
        yLimMem = ylim();
        ylim([0 yLimMem(2)]);
        yyaxis right;
        hold on;
        makeLinePlots(plotParameters.baggingEffN,resultsOut.baggingEffN);
        ylabel({'effective num. virtual patients';'(n_{eff}) for the whole-fit case'});
        % Place the bottom of the y-axis at the origin:
        yLimMem = ylim();
        ylim([0 yLimMem(2)]);
        ax = gca();
        ax.XScale = 'log';
        ax.FontSize = 20;
        grid on;
        figFilepathWithoutExtension = [outputDir figfilename];
        linearCalibrationSaveFig(figHandle,figFilepathWithoutExtension);
    end

    
    %%  Plot the time taken for a single bagging run:
    figfilename = 'timeForSingleBaggingRun';
    figHandle = figure;
    xlabel('fraction VPs per bagging sample')
    hold on;
    plot(resultsOut.baggingRunTime.fractionVPsPerBaggingIteration,resultsOut.baggingRunTime.calibrationTimeForSingleNonCVRunSeconds,'Marker','o','MarkerSize',12,'LineWidth',2,'DisplayName','run time');
    ylabel('run time for bagging [seconds]');
    % Place the bottom of the y-axis at the origin:
    yLimMem = ylim();
    ylim([0 yLimMem(2)]);
    ax = gca();
    ax.XScale = 'log';
    ax.FontSize = 20;
    grid on;
    figFilepathWithoutExtension = [outputDir figfilename];
    linearCalibrationSaveFig(figHandle,figFilepathWithoutExtension);

    
    %% Nested functions
    
    function makeLinePlots(plotParameters,resultsOutTable)
        fieldNamesInternal = fields(plotParameters);
        for iFieldInternal = 1:length(fieldNamesInternal)
            if any(strcmp(resultsOutTable.Properties.VariableNames,fieldNamesInternal{iFieldInternal}))
                plot(resultsOutTable.fractionVPsPerBaggingIteration,resultsOutTable.(fieldNamesInternal{iFieldInternal}),'LineStyle',plotParameters.(fieldNamesInternal{iFieldInternal}).lineStyle,'Marker',plotParameters.(fieldNamesInternal{iFieldInternal}).marker,'Color',plotParameters.(fieldNamesInternal{iFieldInternal}).color,'MarkerSize',12,'LineWidth',1.5,'DisplayName',plotParameters.(fieldNamesInternal{iFieldInternal}).displayName);
            end
        end
    end
    
    function subgroupTable = constructSubGroupTable(plotParameters,axisDescriptions)
        descriptionsMain = {};
        descriptionsSubType = {};
        displayNames = {};
        barColors = {};
        for iDescription = 1:length(axisDescriptions)
            fieldNamesInternal = fields(plotParameters.(axisDescriptions{iDescription}));
            for iFieldInternal = 1:length(fieldNamesInternal)
                descriptionsMain = [descriptionsMain ; axisDescriptions{iDescription}];
                descriptionsSubType =  [descriptionsSubType ; fieldNamesInternal{iFieldInternal}];
                displayNames = [displayNames ; plotParameters.(axisDescriptions{iDescription}).(fieldNamesInternal{iFieldInternal}).displayName];
                barColors = [barColors ; plotParameters.(axisDescriptions{iDescription}).(fieldNamesInternal{iFieldInternal}).color];
            end
        end
        subgroupTable = table(descriptionsMain,descriptionsSubType,displayNames,barColors);
    end
    
    function barYParticularRow = constructBarYRow(resultsOut,subgroupTable,uniqueSubTypes,descriptionMain,iResult)
        iMaskDescriptionMain = strcmp(subgroupTable.descriptionsMain,descriptionMain);
        descriptionSubTypes = subgroupTable.descriptionsSubType(iMaskDescriptionMain);
        barYParticularRow = nan(1,length(uniqueSubTypes));
        
        for iSubType = 1:length(uniqueSubTypes)
            if isfield(resultsOut,descriptionMain) && any(strcmp(resultsOut.(descriptionMain).Properties.VariableNames,uniqueSubTypes{iSubType}))
                barYParticularRow(iSubType) = resultsOut.(descriptionMain).(uniqueSubTypes{iSubType})(iResult);
            end
        end
    end
end
```
