## Supplementary File 3 for "A surrogate workflow for virtual population development: designing an amenable proxy for a hypothesis-test inspired nonlinear goodness-of-fit": LinearCalibrationOptions.m

```
classdef LinearCalibrationOptions
% Here, we define the LinearCalibrationOptions class to run a VPopOptimization
%
%  Properties%  
%  optimizationAlgorithm: 	(string) Specifies which Matlab optimization
%                           function to use. Options are "nnls", "lsqlin" and
%                           "lsqnonneg". Default is "nnls".
%  method:                  (string) Specifies whether to do a single,
%                           best-fit optimization, or to use use an
%                           iterative technique (either bootstrapping or
%                           bagging). Options are: "bestFit", "bootstrap", and "bagging".
%                           "bestFit" will run a single optimization and return the optimal
%                           prevalence weights. "bootstrap" will also do this, but will also run
%                           many iterations with perturbed data (sampled with replacement), in 
%                           order to calculate confidence intervals on the prevalence weight 
%                           estimates. "bagging" will also run many iterations with with 
%                           perturbed data (sampled with replacement), but will also select only
%                           a subset of VPs for each iteration, in order to reduce the
%                           variability in the model fit. Only "bootstrap" will calculate
%                           confidence intervals. Default is "bestFit". Specified as name-value
%                           pair.
%  cdfProbsToFit: 			(vector) Probabilities of the cumulative
% 							distribution functions (CDF) to fit. Default is to fit all points in the
% 							CDF, which may take a long time. For faster
% 							fitting, it's recommended to fit the CDFs more
% 							sparsely; e.g., 'cdfProbsToFit =
% 							0.05:0.05:0.95'. Default is "all", which will
% 							fit all points in the CDF.

%  pdf2DProbsToFitN: 	    (scalar) an integer to specify 2D meshgrid
%                            density. It would generate a
%                            pdf2DProbsToFitN*pdf2DProbsToFitN 2D mesh to
%                            generate a linear problem. Default is set to 2

%  binTableGroupWeight: 				(scalar) Weight for data in myVPop.binTable
%  distTableGroupWeight: 				(scalar) Weight for data in myVPop.distTable
%  brTableRECISTGroupWeight:			(scalar) Weight for data in myVPop.brTableRECIST
%  rTableRECISTGroupWeight: 			(scalar) Weight for data in myVPop.rTableRECIST
%  mnSDTableGroupWeight:				(scalar) Weight for data in myVPop.mnSDTable
%  corTableGroupWeight:                 (scalar) Weight for data in myVPop.corTable
%  distTable2DGroupWeight:              (scalar) Weight for data in
%                                        myVPop.distTable2D
%										GroupWeights Specify fitting weights for each of the
% 										different types of data. Default is to set all of the weights equal to 1,
% 										so that all of the different data groups are weighed equally.
%  priorPrevalenceWeightAssumption:		(string) A priori
% 										knowledge of prevalence weights is required to renormalize VP prevalence weights
% 										in cases where some VP simulation values are NaN since they dropped off
% 										therapy. With VPs missing, the prevalence weights need to be renormalized
% 										before calculating the weighted mean and standard deviation of a
% 										simulation variable across the virtual population. Three options are
% 										available: 
%										(1) "uniform": assume uniform prevalence weights (default) assumes that
% 										all VPs have the same prevalence weight -- this isn't expected to result
% 										in inaccurate fitting if the probability of dropping off therapy doesn't
% 										correlate with prevalence weight, since then the average prevalence
% 										weight of a large chunk of NaN VPs should be approximately the uniform
% 										prevalence weight, 1/nVP, and for a small chunk of NaN VPs, the
% 										prevalence weight renormalization correction factor should be negligible;
% 										(2) "specified": assume prevalence weights uses the prevalence weights
% 										specified in myVPop.pws as the prior prevalence weights; and 
%										(3)  "ignoreDropout": do not fit simulation data containing NaN VPs, ignores any data in which
% 										at least some of the VPs are NaN which can happen with modeled dropouts
%  responseValTransformation:	(string): Specifies whether the
% 								observations should be transformed  so that the residuals relative to the
% 								values of the observations are what are being fitting. Options:
% 								"relative" (default) and "none"
%  maxPrevalenceWeight:     (scalar): maximum prevalence weight allowed.
%                           Note: this probably shouldn't be set if using
%                           bootstrapping or bagging, since those
%                           methods tend to distribute weights by
%                           averaging; specifying a 'maxPrevalenceWeight'
%                           would impose a limit at each iteration of these
%                           algorithms, not on the average of the
%                           iterations. Default is 'Inf'.
%  nBootstrapIterations:    (scalar) Number of bootstrapping iterations to
%                           perform. This option is only used if 'method'
%                           is "bootstrap" or "bagging". Default is
%                           1000.
%  fractionVPsPerBaggingIteration:  (scalar) Fraction of VPs to sample
%                                   per bagging iteration. This option is
%                                   only used if 'method' is "bootstrap" or
%                                   "bagging". Default is 0.9.
%  optimizationAlgorithmOptions:    (struct) Changes the options of the
%                                   'optimizationAlgorithm' from their
%                                   default values to the values specified
%                                   here.
% expWeightFuncHandle:     (function handle). Specifies a weight for a
%                           group based on the experimental sample size and
%                           standard deviation. The function should take
%                           three arguments, in the following order: (1, scalar)
%                           experimental sample size; (2, scalar)
%                           experimental standard deviation; and (3, char
%                           vector) description of the data group.
%
% minSubWeightConstraint:   Specify minimum subpopulation weight constraint, default = 0
% targetEffNConstraint:     TargetEffN for linear calibration.
% oldVPop:                  oldVPop to ensure MSE reduction, might be empty for GOF optimization. Default = []
      
   properties
      optimizationAlgorithm = "nnls"
      method = "bestFit"
      cdfProbsToFit = "all"
      pdf2DProbsToFitN = 2
	  binTableGroupWeight = 1
	  distTableGroupWeight = 1
	  brTableRECISTGroupWeight = 1
	  rTableRECISTGroupWeight = 1
      mnSDTableGroupWeight = 1
      corTableGroupWeight = 1
      distTable2DGroupWeight = 1
	  priorPrevalenceWeightAssumption = "uniform"
	  responseValTransformation = "relative";
      maxPrevalenceWeight = Inf
      nBootstrapIterations = 1000
      fractionVPsPerBaggingIteration = 0.9
      optimizationAlgorithmOptions = []
      expWeightFuncHandle = @(expN) sqrt(expN) % original: @(expN,expSTD,expDataGrp) sqrt(expN) 
      minSubWeightConstraint = 0 
      targetEffNConstraint = 150 
      oldVPop = [] 
   end
   methods
      function obj = set.optimizationAlgorithm(obj,myValue)
          if (strcmpi(myValue,"lsqlin") ||...
               strcmpi(myValue,"lsqnonneg") ||...
               strcmpi(myValue,"nnls") ||...
               strcmpi(myValue,"quadprog") ||... 
               strcmpi(myValue,"quadprogMaxResi") ||... 
               strcmpi(myValue,"quadprogEffN") ||... 
               strcmpi(myValue,"fmincon")) 
            obj.optimizationAlgorithm = myValue;
          else
            error(['Invalid optimizationAlgorithm setting in ',mfilename,', allowed setting: "lsqlin," "lsqnonneg," "nnls," "quadprog," "quadprogMaxResi," "quadprogEffN," "fmincon."'])
          end
      end	
      function obj = set.method(obj,myValue)
          if (strcmpi(myValue,"bestFit") ||...
               strcmpi(myValue,"bootstrap") ||...
               strcmpi(myValue,"bagging") ||... 
               strcmpi(myValue,"bestFitInitials"))
            obj.method = myValue;
          else
            error(['Invalid method setting in ',mfilename,', allowed setting: "bestFit," "bootstrap," "bagging," "bestFitInitials."'])
          end
      end	
      function obj = set.cdfProbsToFit(obj,myValue)
          if isnumeric(myValue) || (isstring(myValue) && strcmpi(myValue,"all"))
              obj.cdfProbsToFit = myValue;
          else
            error(['Invalid cdfProbsToFit value in ',mfilename,', expecting a numeric vector or the string "all"'])
          end
      end 	 
      function obj = set.pdf2DProbsToFitN(obj,myValue)
          if isnumeric(myValue)
              obj.pdf2DProbsToFitN = myValue;
          else
            error(['Invalid pdf2DProbsToFitN value in ',mfilename,', expecting a numeric vector (integer)'])
          end
      end 
      function obj = set.binTableGroupWeight(obj,myValue)
          if isnumeric(myValue) 
              obj.binTableGroupWeight = myValue;
          else
            error(['Invalid binTableGroupWeight value in ',mfilename,', expecting a numeric value'])
          end
      end	
      function obj = set.distTableGroupWeight(obj,myValue)
          if isnumeric(myValue) 
              obj.distTableGroupWeight = myValue;
          else
            error(['Invalid distTableGroupWeight value in ',mfilename,', expecting a numeric value'])
          end
      end	  	  
      function obj = set.brTableRECISTGroupWeight(obj,myValue)
          if isnumeric(myValue) 
              obj.brTableRECISTGroupWeight = myValue;
          else
            error(['Invalid brTableRECISTGroupWeight value in ',mfilename,', expecting a numeric value'])
          end
      end	
      function obj = set.rTableRECISTGroupWeight(obj,myValue)
          if isnumeric(myValue) 
              obj.rTableRECISTGroupWeight = myValue;
          else
            error(['Invalid rTableRECISTGroupWeight value in ',mfilename,', expecting a numeric value'])
          end
      end	
      function obj = set.mnSDTableGroupWeight(obj,myValue)
          if isnumeric(myValue) 
              obj.mnSDTableGroupWeight = myValue;
          else
            error(['Invalid mnSDTableGroupWeight value in ',mfilename,', expecting a numeric value'])
          end
      end
      function obj = set.corTableGroupWeight(obj,myValue)
          if isnumeric(myValue) 
              obj.corTableGroupWeight = myValue;
          else
            error(['Invalid corTableGroupWeight value in ',mfilename,', expecting a numeric value'])
          end
      end
      function obj = set.distTable2DGroupWeight(obj,myValue)
          if isnumeric(myValue) 
              obj.distTable2DGroupWeight = myValue;
          else
            error(['Invalid distTable2DGroupWeight value in ',mfilename,', expecting a numeric value'])
          end
      end
      function obj = set.priorPrevalenceWeightAssumption(obj,myValue)
          if (strcmpi(myValue,"uniform") ||...
               strcmpi(myValue,"specified") ||...
               strcmpi(myValue,"ignoreDropout"))
            obj.priorPrevalenceWeightAssumption = myValue;
          else
            error(['Invalid priorPrevalenceWeightAssumption setting in ',mfilename,', allowed setting: "uniform," "specified," "ignoreDropout."'])
          end
      end
      function obj = set.responseValTransformation(obj,myValue)
%           obj.responseValTransformation = myValue;
          if (strcmpi(myValue,"relative") ||...
               strcmpi(myValue,"none"))
            obj.responseValTransformation = myValue;
          else
            error(['Invalid responseValTransformation setting in ',mfilename,', allowed setting: "relative," "none."'])
          end
      end  
      function obj = set.maxPrevalenceWeight(obj,myValue)
          if isnumeric(myValue) 
              obj.maxPrevalenceWeight = myValue;
          else
            error(['Invalid maxPrevalenceWeight value in ',mfilename,', expecting a numeric value'])
          end
      end	
      function obj = set.nBootstrapIterations(obj,myValue)
          if isnumeric(myValue) 
              obj.nBootstrapIterations = myValue;
          else
            error(['Invalid nBootstrapIterations value in ',mfilename,', expecting a numeric value'])
          end
      end	
      function obj = set.fractionVPsPerBaggingIteration(obj,myValue)
          if isnumeric(myValue) 
              obj.fractionVPsPerBaggingIteration = myValue;
          else
            error(['Invalid fractionVPsPerBaggingIteration value in ',mfilename,', expecting a numeric value'])
          end
      end
      function obj = set.optimizationAlgorithmOptions(obj,myValue)
          obj.optimizationAlgorithmOptions = myValue;
      end
      function obj = set.expWeightFuncHandle(obj,myValue)
          obj.expWeightFuncHandle = myValue;
      end
      function obj = set.minSubWeightConstraint(obj,myValue)
          obj.minSubWeightConstraint = myValue;
      end
      function obj = set.targetEffNConstraint(obj,myValue)
          obj.targetEffNConstraint = myValue;
      end
      
      function value = get(obj,propName)
          switch propName
              case 'verbose'
                  value = obj.verbose;
              case 'cdfProbsToFit'
                  value = obj.cdfProbsToFit;
              case 'pdf2DProbsToFitN'
                  value = obj.pdf2DProbsToFitN;
              case 'toleranceScalingFactor'
                  value = obj.toleranceScalingFactor;                  
              case 'binTableGroupWeight'
                  value = obj.binTableGroupWeight;
              case 'distTableGroupWeight'
                  value = obj.distTableGroupWeight;                  
              case 'brTableRECISTGroupWeight'
                  value = obj.brTableRECISTGroupWeight;
              case 'rTableRECISTGroupWeight'
                  value = obj.rTableRECISTGroupWeight;
              case 'mnSDTableGroupWeight'
                  value = obj.mnSDTableGroupWeight;
              case 'corTableGroupWeight'
                  value = obj.corTableGroupWeight;
              case 'distTable2DGroupWeight'
                  value = obj.distTable2DGroupWeight;
              case 'priorPrevalenceWeightAssumption'
                  value = obj.priorPrevalenceWeightAssumption;
              case 'responseValTransformation'
                  value = obj.responseValTransformation;		
              case 'maxPrevalenceWeight'
                  value = obj.maxPrevalenceWeight;	
              case 'nBootstrapIterations'
                  value = obj.maxPrevalenceWeight;	
              case 'fractionVPsPerBaggingIteration'
                  value = obj.maxPrevalenceWeight;	
              case 'optimizationAlgorithmOptions'
                  value = obj.optimizationAlgorithmOptions;	
              case 'expWeightFuncHandle'
                  value = obj.expWeightFuncHandle;	
              otherwise
                  error(['Error: ',propName ,' is not a valid ',mfilename,' property.'])
          end
      end      

      % The constructor method must have the same name as the class
      function obj = LinearCalibrationOptions()
			
      end
   end
end
```
