## Supplementary File 3 for "A surrogate workflow for virtual population development: designing an amenable proxy for a hypothesis-test inspired nonlinear goodness-of-fit": increaseEffNWithLinearCalibrate.m

```
function myVPop = increaseEffNWithLinearCalibrate(myVPop)
% Uses least-squares optimization and bagging to increase EffN, likely 
% sacrificing goodness-of-fit p-value.

tictoc = tic();

disp('Increasing EffN by using linear calibration with bagging...');

% Set up the optimization options, which will automatically spit back a set
% of default values
optimOptions = LinearCalibrationOptions();

% Define the probabilities of the cumulative
% distribution functions (CDF) to fit. The default is to fit all points on
% the CDF. We will only fit certain probabilities
% along the CDF in order to make the fit faster. These probabilities are
% defined here:
optimOptions.cdfProbsToFit = 0.05:0.05:0.95;

% Other options:
optimOptions.optimizationAlgorithm = "nnls";
optimOptions.optimizationAlgorithmOptions.Accy = 0;
optimOptions.method = "bagging";
optimOptions.fractionVPsPerBaggingIteration = 0.05;

% Initialize a 'LinearCalibration' object:
linearCalibrationObject = LinearCalibration(myVPop,'optimOptions',optimOptions);

% Run the optimization:
linearCalibrationObject = linearCalibrationObject.run();

% Update VPop:
myVPop = linearCalibrationObject.OptimizedVPop;

timeElapsedSec = toc(tictoc);
disp(['Finished increasing EffN. Time elapsed [minutes]: ' num2str(timeElapsedSec/60)]);

end
```
