## Supplementary File 3 for "A surrogate workflow for virtual population development: designing an amenable proxy for a hypothesis-test inspired nonlinear goodness-of-fit": mapelLinearExpand.m

```
function myVPop = mapelLinearExpand(myWorksheet, myMapelOptions, oldVPop)
% This is the main function to call MAPEL, which will develop a virtual
% population based on the simulation results in the worksheet and the
% settings in the mapelOptions.
%
% ARGUMENTS
%  myWorksheet:            (Required) a worksheet instance.  We allow a 
%                          special pass-through where a VPop object can also
%                          be given here to get a couple critical
%                          properties (simData, indexTable, binEdges
%                          binMidPoints) for mapel to be called from
%                          a restart, but in this case all other
%                          properties are taken from the options.
%  myMapelOptions:         (Required) an instance of a mapelOptions object.
%  oldVPop:                 the VPop from last iteration, might contain less Num of VPs. Default = []
%
% % Returns
%  VPop:                    an instance of a VPop
%

continueFlag = false;
if nargin > 3
    warning(['Too many input arguments provided to ',mfilename,'.  Requires: myWorksheet and a mapelOptions (or mapelOptionsRECIST) object'])
    continueFlag = false;
elseif nargin > 1
    continueFlag = true;
else
    continueFlag = false;
    warning(['Insufficient input arguments provided to ',mfilename,'.  Requires: myWorksheet and a mapelOptions (or mapelOptionsRECIST) object.'])
end

if continueFlag
    if ~ismember(class(myMapelOptions),{'mapelOptions','mapelOptionsRECIST'})
        continueFlag = false;
        warning(['Input mapelOptions not recognized in call to ',mfilename,'.  Requires: myWorksheet and a mapelOptions or mapelOptionsRECIST object.'])
    end       
end

if continueFlag
    % First copy all of the needed properties over
    myVPop = initializeOptionPropertiesToVPop(myMapelOptions);
    extraPWs = [];
    
    % Now add additional properties not in the mapelOptions
    if isa(myVPop,'VPopRECIST')      
        if ~isa(myWorksheet,'VPopRECIST')
            myVPop.recistSimFilter = createRECISTSimFilter(myWorksheet, myVPop, false);
        else   
            myVPop.recistSimFilter = myWorksheet.recistSimFilter;
        end
    end	    	 

    % Next, we create a table of nVPs x nBins to index each VP axis into a
    % bin for subsequent calculations.
    % We go to the simulation results in the worksheet and create
    % a table of the needed results to compare to the experimental data
    if strcmp(myVPop.pwStrategy, 'bin')
        if ~isa(myWorksheet,'VPop') && ~isa(myWorksheet,'VPopRECIST')
            myVPop = myVPop.assignIndices(myWorksheet, myMapelOptions);
            myVPop = myVPop.getSimData(myWorksheet);
			% myVPop.vpIDs = getVPIDs(myWorksheet);
			myVPop = myVPop.assignCoeffs(myWorksheet);
            if myVPop.spreadOut > 0
                myVPop.coeffsDist = single(pdist2(myVPop.coeffsTable',myVPop.coeffsTable'));
            end
            % Also enforce an update of the subpopulation table if a
            % worksheet is provided
            myVPop.subpopTable = updateSubpopTableVPs(myVPop.subpopTable, myWorksheet);            
        else
            myVPop.simData = myWorksheet.simData;
			% myVPop.vpIDs = myWorksheet.vpIDs;
            myVPop.indexTable = myWorksheet.indexTable;
            myVPop.binEdges = myWorksheet.binEdges;
            myVPop.binMidPoints = myWorksheet.binMidPoints;
			myVPop.coeffsTable = myWorksheet.coeffsTable;
            if myVPop.spreadOut > 0
                myVPop.coeffsDist = single(pdist2(myVPop.coeffsTable',myVPop.coeffsTable'));
            end
        end
    else %noBin
        if ~isa(myWorksheet,'VPop') && ~isa(myWorksheet,'VPopRECIST')
            myVPop = myVPop.getSimData(myWorksheet);
			% myVPop.vpIDs = getVPIDs(myWorksheet);
            myVPop = myVPop.assignCoeffs(myWorksheet);
            if myVPop.spreadOut > 0
                myVPop.coeffsDist = single(pdist2(myVPop.coeffsTable',myVPop.coeffsTable'));
            end
            % Also enforce an update of the subpopulation table if a
            % worksheet is provided
            myVPop.subpopTable = updateSubpopTableVPs(myVPop.subpopTable, myWorksheet);   
        else
            myVPop.simData = myWorksheet.simData;
			% myVPop.vpIDs = myWorksheet.vpIDs;
            % PWs will still be taken from the mapelOptions (later)
            myVPop.coeffsTable = myWorksheet.coeffsTable;
            if myVPop.spreadOut > 0
                myVPop.coeffsDist = single(pdist2(myVPop.coeffsTable',myVPop.coeffsTable'));
            end         
        end
    end
    												
    if myVPop.intSeed > -1
        rng(myVPop.intSeed, 'twister');
    end        
    % Also assign the now static sim data into the bin tables initially
    % rather than during execution to reduce table assignments.
    myVPop = myVPop.addTableSimVals();
	
	% Get the parallel pools ready
	optimizeType = myVPop.optimizeType;
	if myVPop.poolRestart
		if ~isempty(gcp('nocreate'))
			delete(gcp);
		end
	end

	% Create the pool early, there are a few processes that
	% may use it: both linearCalibration and the swarm optimization.
	% We don't start a pool for simplex
	% since simplex cannot use parallel processing
	% we may want to use the parallel pool to run multiple
	% MAPEL runs in parallel.	
	if sum(ismember({'simplex'},optimizeType)) < 1
		if isempty(gcp('nocreate'))
			% We will use default pool settings
			mySimulateOptions = simulateOptions;
			mySimulateOptions = checkNWorkers(mySimulateOptions);		
			myPool = parpool(mySimulateOptions.clusterID,mySimulateOptions.nWorkers,'SpmdEnabled',false);
		else
			myPool = [];
		end
	else
		myPool = [];
	end
    	
    if strcmp(myVPop.pwStrategy, 'bin')
        % We adopt the index table convention from the original MAPEL paper
        myIndexTable = myVPop.indexTable;
        [myNAxis, nVP] = size(myIndexTable);
        myBinEdges = myVPop.binEdges;
        myBinMidPoints = myVPop.binMidPoints;
        myNBins = myMapelOptions.nBins;

        % We can continue from a previous run if we have a valid
        % initial probability table.
        myInitialProbs = myMapelOptions.initialProbs;
        myRandomStart = myMapelOptions.randomStart;

        if isequal([myNAxis,myNBins], size(myInitialProbs))
            if myRandomStart > 0
                for axisCounter = 1 : myNAxis
                    myInitialProbsTrans = hyperTransform(myInitialProbs(axisCounter,:));
                    mySDs = myRandomStart*abs(myInitialProbsTrans);
                    myInitialProbsTrans = myInitialProbsTrans + randn(1,myNBins-1).*mySDs;
                    myInitialProbs(axisCounter,:) = invHyperTransform(myInitialProbsTrans); 
                end
            end
            myVPop.binProbs = myInitialProbs;
            myVPop = myVPop.assignPWs();
        else
            myVPop = myVPop.startProbs(myRandomStart>0);  
            myVPop = myVPop.assignPWs();
        end
    else 
		 [nAxis, nVP] = size(myVPop.coeffsTable);
        
		 myInitialPWs = myMapelOptions.initialPWs;
		 myRandomStart = myMapelOptions.randomStart;
		 curEffN = myMapelOptions.minEffN;
         
		% we do linear calibration at every iteration during linearExpand phase        
        if (isempty(oldVPop)) % if it is the first round, give a small lambda=1 to initiate
             myOptimOptions = LinearCalibrationOptions();
             myOptimOptions.cdfProbsToFit = 0.05:0.05:0.95;
             myOptimOptions.pdf2DProbsToFitN = 5;
             myOptimOptions.responseValTransformation='none';
             myOptimOptions.optimizationAlgorithm = "quadprogEffN";				
             myOptimOptions.priorPrevalenceWeightAssumption = 'specified';
             myOptimOptions.oldVPop = oldVPop;   
        else % continue linearExpand using quadprog with the specified lambda from last iteration oldVPop
             % if effN of oldVPop<=targetEffN, keep using this lambda; if larger, use lambda/2;
                oldVPopEffN = 1/sum(oldVPop.pws.^2);
                if oldVPopEffN >= curEffN*1.2
                    oldVPop.lambda = (oldVPop.lambda)/2;
                    disp('curEffN too big, reducing lambda to half ...');
                elseif oldVPopEffN <= curEffN*0.95
                    oldVPop.lambda = (oldVPop.lambda)*1.5;
                    disp('curEffN too small, increasing lambda to 1.5X ...');
                end
             myOptimOptions = LinearCalibrationOptions();
             myOptimOptions.cdfProbsToFit = 0.05:0.05:0.95;
             myOptimOptions.pdf2DProbsToFitN = 5;
             myOptimOptions.responseValTransformation='none';
             myOptimOptions.optimizationAlgorithm = "quadprogEffN";				
             myOptimOptions.priorPrevalenceWeightAssumption = 'specified';
             myOptimOptions.oldVPop = oldVPop;        
        end
         
         if isequal(1, length(myInitialPWs)) && (myInitialPWs < 0)
                % Need to add predTableVals to run.
                myVPop = myVPop.startPWs(myWorksheet,0);  % equal weights
         else
                myVPop.pws = myInitialPWs(1,:);
         end
         myVPop = myVPop.addPredTableVals(); 
         
         linearCalibrationObject = LinearCalibration(myVPop,'optimOptions',myOptimOptions);
         tic;
                linearCalibrationObject = linearCalibrationObject.run('closeParallelPoolWhenFinished',false);
         toc;

         if linearCalibrationObject.OptimizationResults.exitFlag >= 1 || linearCalibrationObject.OptimizationResults.exitFlag==-3
                 myInitialPWs = linearCalibrationObject.OptimizationResults.optimalPrevalenceWeightsNormalized'; %linearCalibrationObject.OptimizedVPop.pws;
         else
                 warning(['Unable to find optimal solution to linear problem in ',mfilename,'.  Proceeding with default options.'])
                 myInitialPWs = [];
         end
    end
    
    [~,nInitialPW] = size(myInitialPWs);	
    if isequal(nVP, nInitialPW)
			myVPop.pws = myInitialPWs(1,:);
    else
			myVPop = myVPop.startPWs(myWorksheet,myRandomStart>0);
    end  

    % Update table values.  Not strictly necessary but a nice
    % step to diagnose and does have much computational cost            
    myVPop = myVPop.addPredTableVals();  
    % Next update the individual GOF statistics
    myVPop = evaluateGOF(myVPop);

    if ~any(isnan(linearCalibrationObject.OptimizationResults.optimalPrevalenceWeightsNormalized))
        myVPop.LinearProblemMatrices = linearCalibrationObject.LinearProblemMatrices;
        myVPop.MSE = linearCalibrationObject.MSE; % will this change when I have findFit?
      %  myVPop.lambda = linearCalibrationObject.lambda; % output lambda from fmincon
        myVPop.LinearProblemMatricesSubgroupSumWeights = linearCalibrationObject.LinearProblemMatrices.vpIsInSubgroup*linearCalibrationObject.OptimizationResults.optimalPrevalenceWeightsNormalized;
        myVPop.LinearProblemMatricesobservationDescriptions = linearCalibrationObject.LinearProblemMatrices.observationDescriptions;
        
        minSubWeight = min(myVPop.LinearProblemMatricesSubgroupSumWeights);
        minSubWIndex = find(myVPop.LinearProblemMatricesSubgroupSumWeights==minSubWeight);
        disp(['minSubWeight =' num2str(round(minSubWeight,4)), ' at rowIndex = ' num2str(minSubWIndex(1))]);
    end
    myVPop.lambda = linearCalibrationObject.lambda;
	% Clean up the pool, if needed
	if myVPop.poolClose
		if ~isempty(gcp('nocreate'))
			delete(gcp);
		end
    end
else
    myVPop = VPop;
    warning(['Unable to run ',mfilename,'.  Returning an empty VPop.'])
end
end
```
