## Supplementary File 3 for "A surrogate workflow for virtual population development: designing an amenable proxy for a hypothesis-test inspired nonlinear goodness-of-fit": restartMapel.m

```
function myVPop = restartMapel(myVPop, myRandomStart, myExtraPWs)
% This is a function to start developing a VPop using data in an existing
% existing VPop as a starting point.
%
% ARGUMENTS
%  myVPop:           A VPop instance.  Note restarting will fail
%                    if there is not enough information in the VPop
%                    from a previous run.
%  myRandomStart:    (optional) A non-negative numeric value. If 0, the initial 
%                    parameters are not perturbed in anyway.  Otherwise the 
%                    transformed initial bin probabilities are perturbed  
%                    with random (normally distributed) noise that has
%                    normalized standard deviation of the specified value.  
%                    If no initialProbs are specified, a value of zero 
%                    results in uniform bin probabilities and a value >0 
%                    results in uniform normal initial bin probabilities  
%                    (renormalized so the margin is 1).  Default is 0.
%  myExtraPWs:       (optional) additional m x nVP pw matrix to include
%                     in optimization.  Only supported right now
%                     for 'direct.'
%
% Returns
%  VPop:             an instance of a VPop
%
continueFlag = false;
if nargin > 3
    warning(['Too many input arguments provided to ',mfilename,'.  Requires: myVpop and optionally myRandomStart, myExtraPWs.'])
    continueFlag = false;
elseif nargin > 2
    continueFlag = true;    
elseif nargin > 1
    myExtraPWs = [];
    continueFlag = true;
elseif nargin > 0
    myExtraPWs = [];
    myRandomStart = 0;
    continueFlag = true;
else
    continueFlag = false;
    warning(['Insufficient input arguments provided to ',mfilename,'.  Requires: myVpop and optionally myRandomStart, myExtraPWs.'])
end

if continueFlag
    if (~strcmp(class(myVPop),'VPop') & ~strcmp(class(myVPop),'VPopRECIST'))
        continueFlag = false;
        warning(['Input VPop not recognized in call to ',mfilename,'.  Requires: myVpop and optionally myRandomStart, myExtraPWs.'])
    end
    if ~(myRandomStart>=0)
        continueFlag = false;
        warning(['Input myRandomStart not recongized in call to ',mfilename,'.  myRandomStart must be a non-negative number.'])
    end        
end

if continueFlag
    % We'll just call MAPEL, so all we need to do is feed the input VPop
    % back to a mapelOptions object.
	if isa(myVPop,'VPopRECIST')
		myMapelOptions = mapelOptionsRECIST;
	elseif isa(myVPop,'VPop')
		myMapelOptions = mapelOptions;
	end
    myMapelOptions.expData = myVPop.expData;
    myMapelOptions.mnSDTable = myVPop.mnSDTable;
    myMapelOptions.binTable = myVPop.binTable;
    myMapelOptions.distTable = myVPop.distTable;
    myMapelOptions.distTable2D = myVPop.distTable2D;
	myMapelOptions.corTable = myVPop.corTable;
    myMapelOptions.subpopTable = myVPop.subpopTable;
	myMapelOptions.pwStrategy = myVPop.pwStrategy;
	if strcmp(myVPop.pwStrategy, 'bin')
		[nAxis, nBins] = size(myVPop.binProbs);
		myMapelOptions.nBins = nBins;
		myMapelOptions.initialProbs = myVPop.binProbs;
        % This property isn't kept for the VPop
        % Could check the binEdges of the VPop
		% myMapelOptions.equalBinBreaks = myVPop.equalBinBreaks;
    else
        if isempty(myExtraPWs)
            myMapelOptions.initialPWs = myVPop.pws;
        else
            myMapelOptions.initialPWs = [myVPop.pws;myExtraPWs];
        end
	end
    myMapelOptions.randomStart = myRandomStart;	
    myMapelOptions.nIters = myVPop.nIters;
    myMapelOptions.tol = myVPop.tol;
    myMapelOptions.spreadOut = myVPop.spreadOut;
    myMapelOptions.minIndPVal = myVPop.minIndPVal;	
	myMapelOptions.useEffN = myVPop.useEffN;
    myMapelOptions.exactFlag = myVPop.exactFlag;    
    myMapelOptions.optimizeTimeLimit = myVPop.optimizeTimeLimit;    
    myMapelOptions.optimizeType = myVPop.optimizeType;
    myMapelOptions.optimizePopSize = myVPop.optimizePopSize; 	
	myMapelOptions.objectiveLimit = myVPop.objectiveLimit; 
	myMapelOptions.poolClose = myVPop.poolClose; 
	myMapelOptions.poolRestart = myVPop.poolRestart; 
    myMapelOptions.intSeed = myVPop.intSeed;
    myMapelOptions.minEffN = myVPop.minEffN;
    myMapelOptions.nIters = myVPop.nIters; 
	if isa(myVPop,'VPopRECIST') 
		myMapelOptions.brTableRECIST = myVPop.brTableRECIST;
		myMapelOptions.rTableRECIST = myVPop.rTableRECIST;        
        myMapelOptions.relSLDvar = myVPop.relSLDvar;
        myMapelOptions.absALDVar = myVPop.absALDVar;
        myMapelOptions.crCutoff = myVPop.crCutoff;        
    end
	myVPop = mapel(myVPop, myMapelOptions);
else
    warning(['Unable to run ',mfilename,'.  Returning myVPop.'])
end
end
```
