## Supplementary File 3 for "A surrogate workflow for virtual population development: designing an amenable proxy for a hypothesis-test inspired nonlinear goodness-of-fit": addPD2Vals.m

```
      function updateVPopRECIST = addPD2Vals(updateVPopRECIST, refVPopRECIST, RegressionBasedOnBoth, flagPD2Exp)
          % Once we have model predicted PD and regression-based PD2,
          % we can add expPD2 and predPD2 into rate calibration
          % ARGUMENTS
          %  updateVPopRECIST (required):      Note that the following properties should be
          %               initialized (experimental and simulation data) 
          %               before calling this method:
          %                mnSDTable
          %                binTable
          %                distTable
          %                distTable2D
          %                corTable		  
		  %				   brTableRECIST
		  %				   rTableRECIST	
		  %                subpopTable
          %                simData
          %   refVPopRECIST (optional): the VPop where the brTableRECIST is 
          %                         used as reference to estimate PD2.
          %                         Default is empty.
          %   RegressionBasedOnBoth (optional): Boolean (true/false) 
          %                         indicating whether to estimate PD2 based on 
          %                         combined brTable (from refVPopRECIST and updateVPopRECIST) 
          %                         Default is false.
          %   flagPD2Exp    (optional): Boolean (true/false) 
          %                         indicating whether to use PD2 based on 
          %                         the PD2 rates from the Exp intervention, this can only be set to true when expPD2 is available.
          %                         Default is false.
          
          %
          % RETURNS
          %  updateVPopRECIST (updated): A VPop object is returned, but with updated properties:
          %                mnSDTable
          %                binTable
          %                distTable
          %                distTable2D
          %                corTable	
		  %				   brTableRECIST	 % with additional 8 columns:expCRPD2,expPRPD2, ..., predCRPD2,predPRPD2
		  %				   rTableRECIST
		  %                subpopTable	
          
        % Perform initial checks on the provided arguments
        flagContinue = true;
        if nargin > 4
            warning(['requires input arguments: updateVPopRECIST, optionally refVPopRECIST and RegressionBasedOnBoth.  Too many arguments provided.'])
            flagContinue = false;    
        elseif nargin > 3
            if RegressionBasedOnBoth && flagPD2Exp
                error('Both RegressionBasedOnBoth and flagPD2Exp are true, need to choose one to estimate PD2 rates and plot on ... ')
            else
                flagContinue = true;
            end
        elseif nargin > 2
            flagPD2Exp = false;
            flagContinue = true;
        elseif nargin < 2
            refVPopRECIST = [];
            RegressionBasedOnBoth = false;
            flagPD2Exp = false;
            flagContinue = true;
        elseif nargin < 3
            RegressionBasedOnBoth = false;
            flagPD2Exp = false;
            flagContinue = true;
        end

        if flagContinue
            if ~ismember(class(updateVPopRECIST),{'mapelOptionsRECIST','VPopRECIST'})
                flagContinue = false;
                warning(['Input not recognized in call to ',mfilename,'.  Requires: mapelOptionsRECIST or VPopRECIST object.'])
            elseif (~isempty(refVPopRECIST) & ~ismember(class(refVPopRECIST),{'mapelOptionsRECIST','VPopRECIST'}))
                flagContinue = false;
                warning(['Input not recognized in call to ',mfilename,'.  Requires: mapelOptionsRECIST or VPopRECIST object.'])                
            end       
        end
        
        % estimate PD2 based on brTableREF, and predict new response rates for myBRTableRECIST
        if flagContinue
            if (flagPD2Exp)
                disp(['Using the experimental PD2 rates from individual trials directly ...']);
            end
            
            if (isempty(refVPopRECIST))
                disp(['Estimating PD2 rates based on the provided updateVPopRECIST, return an updateVPopRECIST object with updated brTableRECIST ...']);
                brTableREF = updateVPopRECIST.brTableRECIST;
                myBRTableRECIST = updateVPopRECIST.brTableRECIST;
            elseif (~isempty(refVPopRECIST) & RegressionBasedOnBoth == false)
                disp(['Estimating PD2 rates based on the provided refVPopRECIST only, return an updateVPopRECIST object with updated brTableRECIST ...']);
                brTableREF = refVPopRECIST.brTableRECIST;
                myBRTableRECIST = updateVPopRECIST.brTableRECIST;
            elseif (~isempty(refVPopRECIST) & RegressionBasedOnBoth == true)
                disp(['Estimating PD2 rates based on both refVPopRECIST and updateVPopRECIST, return an updateVPopRECIST object with updated brTableRECIST ...']);
                brTableREF = [refVPopRECIST.brTableRECIST;updateVPopRECIST.brTableRECIST];
                myBRTableRECIST = updateVPopRECIST.brTableRECIST;
            end
            
            if ~flagPD2Exp
                % Only keep the last time point from each inidividual intervention (BOR), then train the regression model
                 testData = brTableREF(:,{'subpopNo','expDataID','interventionID'}); 
                 [C,IA,IC] = unique(testData,'rows','stable'); % unique(testData,'last','rows','stable');
                 LastRowindex=IA;
                 for j=1:length(IA)
                     indices=find(IC==j);
                     LastRowindex(j)=indices(end);
                 end            
                testRows = find(~isnan(brTableREF{:,{'expNPD21LS'}})); 
                regRows = intersect(testRows, LastRowindex);

                expN = table2array(brTableREF(regRows,'expN'));
                expPD=table2array(brTableREF(regRows,'expPD')).*expN;
                expSD=table2array(brTableREF(regRows,'expSD')).*expN;
                resp=table2array(brTableREF(regRows,'expNPD21LS'));
                fitTable = table(expPD,expSD,resp);
                lmResults = fitlm(fitTable,'resp ~ expPD + expSD','Intercept',false);

                % Lu: estimate a PD2 for each row: because for the same intervention different timepoints, the predicted PD patient number could be different   
                predRows = find(sum(isnan(myBRTableRECIST{:,{'predN','predCR','predPR','predSD','predPD'}}),2)==0);
                predN = myBRTableRECIST{predRows,{'predN'}};
                predPD = myBRTableRECIST{predRows,{'predPD'}}.*predN;
                predSD = myBRTableRECIST{predRows,{'predSD'}}.*predN;
                res = predPD.*lmResults.Coefficients{1,'Estimate'} + predSD.*lmResults.Coefficients{2,'Estimate'};
                myBRTableRECIST{predRows,'predNPD21LS'}=res;
            else
                expN = table2array(brTableREF(:,'expN'));
                predN = myBRTableRECIST{:,{'predN'}};
                expNPD21LS = myBRTableRECIST.('expNPD21LS');
                expNPD21LS(isnan(expNPD21LS)) = 0;
                myBRTableRECIST{:,'predNPD21LS'} = expNPD21LS.*predN./expN;
            end

            % only update the updateVPopRECIST.brTableRECIST
            brRowsTarget = updateVPopRECIST.simData.brRows;
            brRowsSource = find(brRowsTarget>0);        

          if ~isempty(brRowsSource)
              expNPD21LS = myBRTableRECIST.('expNPD21LS');
              predNPD21LS = myBRTableRECIST.('predNPD21LS');
              expNPD21LS(isnan(expNPD21LS)) = 0;
              predNPD21LS(isnan(predNPD21LS)) = 0;
            
              expN = myBRTableRECIST.('expN');
              expCR = myBRTableRECIST.('expCR');
              expPR = myBRTableRECIST.('expPR');
              expSD = myBRTableRECIST.('expSD');
              expPD = myBRTableRECIST.('expPD'); 
              expNPD2 = expN+expNPD21LS;
              myBRTableRECIST.('expCRPD2') = expCR.*expN./expNPD2;
              myBRTableRECIST.('expPRPD2') = expPR.*expN./expNPD2;
              myBRTableRECIST.('expSDPD2') = expSD.*expN./expNPD2;
              myBRTableRECIST.('expPDPD2') = (expPD.*expN+expNPD21LS)./expNPD2; 

              predN = myBRTableRECIST.('predN');
              predCR = myBRTableRECIST.('predCR');
              predPR = myBRTableRECIST.('predPR');
              predSD = myBRTableRECIST.('predSD');
              predPD = myBRTableRECIST.('predPD'); 
              predNPD2 = predN+predNPD21LS;
              myBRTableRECIST.('predCRPD2') = predCR.*predN./predNPD2;
              myBRTableRECIST.('predPRPD2') = predPR.*predN./predNPD2;
              myBRTableRECIST.('predSDPD2') = predSD.*predN./predNPD2;
              myBRTableRECIST.('predPDPD2') = (predPD.*predN+predNPD21LS)./predNPD2; 

              updateVPopRECIST.brTableRECIST = myBRTableRECIST;
          end          
        end
      end
      
      
      
```
