## Supplementary File 3 for "A surrogate workflow for virtual population development: designing an amenable proxy for a hypothesis-test inspired nonlinear goodness-of-fit": alignCDFsPreGrid.m

```
function [CDF1, CDF2] = alignCDFsPreGrid(SC, sample1ind, sample2ind, W1, W2)
% This function returns
% adjusted CDF vectors that can be compared element-wise
%
% ARGUMENTS
%  SC          the combined unique, sorted sample values
%  sample1ind  indices mapping values for sample1 to SC. 
%              Indices are assumed to point to the each unique, last value
%              following sorting.
%  sample2ind  indices mapping values for sample1 to SC. 
%              Indices are assumed to point to the each unique, last value
%              following sorting.   
%  W1          a 1xlength(sample1) vector of weights for sample 1
%  W2          a 1xlength(sample2) vector of weights for sample 2
%
% RETURNS
%  CDF1        a 1xlength unique sample values of increasing cumulative
%              probabilities in sample 1 corresponding to the unique 
%              combined sample values
%  CDF2        a 1xlength unique sample values of increasing cumulative
%              probabilities in sample 1 corresponding to the unique 
%              combined sample values
%
% First we need to interpolate the two samples so we can directly
% take the supremum function.
    W1 = W1 / sum(W1);
    W2 = W2 / sum(W2);
    W1 = cumsum(W1);
    W2 = cumsum(W2);
	% We need to re-calculate the cdf for sample1 and sample2
	% at all x-values in either
	% sample 1 and sample2 so we can directly calculate
	% the differences in the cdf at each point
    CDF1 = zeros(1, length(SC));
    CDF2 = zeros(1, length(SC));
    CDF1Indices = find(~isnan(sample1ind));
    CDF2Indices = find(~isnan(sample2ind));
    CDF1(CDF1Indices) = W1(sample1ind(CDF1Indices));
    CDF2(CDF2Indices) = W2(sample2ind(CDF2Indices));
    
end
```
