## Supplementary File 3 for "A surrogate workflow for virtual population development: designing an amenable proxy for a hypothesis-test inspired nonlinear goodness-of-fit": alignSamples.m

```
function [sample1Ind, sample2Ind, SC] = alignSamples(sample1, sample2)
% This function takes two samples, combines them, and returns
% indices vectors that can be used to map the
% original values in each sample onto the combined vector, prioritizing
% the last observation to facilitate generating empirical CDFs.
%
% ARGUMENTS
%  sample1     observed values for sample1. These are assumed to be sorted
%              in ascending order
%  sample2     observed values for sample2. These are assumed to be sorted
%              in ascending order    
%
% RETURNS
%  sample1Ind  a 1xlength unique sample indices that map elements in sample
%              1 to SC.
%  sample2Ind  a 1xlength unique sample indices that map elements in sample
%              2 to SC.
%  SC          combined sample values
%
	% We need to re-calculate the cdf for sample1 and sample2
	% at all x-values in either
	% sample 1 and sample2 so we can directly calculate
	% the differences in the cdf at each point
    [sample1,Inu1,~]=unique(sample1,'last');
    [sample2,Inu2,~]=unique(sample2,'last');
    SC = unique([sample1,sample2]);
    SC = sort(SC, 'ascend');
    % Get the indices of length SC where every entry in SC is found to be
    % >= the indicated index in sample1
    if length(sample1) > 1
        sample1Ind = interp1(sample1,[1:length(sample1)],SC,'previous');
    else
        % If there is just one unique value in sample1 
        sample1Ind = ones(1,length(SC));
        sample1Ind(sample1<SC) = nan;
    end        
    % Get the indices of length SC where every entry in SC is found to be
    % >= the indicated index in sample2    
    if length(sample2) > 1
        sample2Ind = interp1(sample2,[1:length(sample2)],SC,'previous');
    else
        sample2Ind = ones(1,length(SC));
        sample2Ind(sample2<SC) = nan;
    end
    % Right fill with last value, leave the left side as nan
    if sum(isnan(sample1Ind)) > 0
        t = 1:numel(sample1Ind);
        sample1Ind = interp1(t(~isnan(sample1Ind)),sample1Ind(~isnan(sample1Ind)),t,'previous','extrap');    
    end
    if sum(isnan(sample2Ind)) > 0 && sum(~isnan(sample2Ind)) > 1 %need > 2 points for interpolating
        t = 1:numel(sample2Ind);
        sample2Ind = interp1(t(~isnan(sample2Ind)),sample2Ind(~isnan(sample2Ind)),t,'previous','extrap');    
    end   
    % To update CDFs, we will need to map back to the non-unique but sorted
    % samples
    curInd = find(~isnan(sample1Ind));
    sample1Ind(curInd) = Inu1(sample1Ind(curInd));
    curInd = find(~isnan(sample2Ind));
    sample2Ind(curInd) = Inu2(sample2Ind(curInd));
end
```
