## Supplementary File 3 for "A surrogate workflow for virtual population development: designing an amenable proxy for a hypothesis-test inspired nonlinear goodness-of-fit": checkExpDataCorrelations.m

```
function [myCorrelationsCross,myCorrelationsAuto] = checkExpDataCorrelations(myMapelOptions)
% This is a function to check correlations in data loaded into
% a mapelOptions.
%
%  myMapelOptions:         (Required) an instance of a mapelOptions object.
%
% Returns
%  myCorrelationsCross:     a table with correlations between 
%                            biomarkers
%  myCorrelationsAuto:     a table with correlations within the same
%                            biomarkers but at, for example, different
%                            timepoints.  They may be
%                            of interest but will be much more numerous 
%                            so are separated here.

continueFlag = false;
if nargin > 1
    warning(['Too many input arguments provided to ',mfilename,'.  Requires: mapelOptions (or mapelOptionsRECIST) object'])
    continueFlag = false;
elseif nargin > 0
    continueFlag = true;
else
    continueFlag = false;
    warning(['Insufficient input arguments provided to ',mfilename,'.  Requires: myWorksheet and a mapelOptions (or mapelOptionsRECIST) object.'])
end

if continueFlag
    if ~ismember(class(myMapelOptions),{'mapelOptions','mapelOptionsRECIST'})
        continueFlag = false;
        warning(['Input mapelOptions not recognized in call to ',mfilename,'.  Requires: myWorksheet and a mapelOptions or mapelOptionsRECIST object.'])
    end       
end

rowNumber1 = [];
rowNumber2 = [];
interventionID1 = {};
interventionID2 = {};
time1 = [];
time2 = [];
expN = [];
elementID1 = {};
elementID2 = {};
expVarID1 = {};
expVarID2 = {};
expDataID1 = {};
expDataID2 = {};
data1 = {};
data2 = {};
corCoeff = [];
pVal = [];

if continueFlag
    expDataTable = myMapelOptions.expData;
    commonNames = loadCommonNames();
    if isa(myMapelOptions,'mapelOptions')    
        tableVariableNamesFixed = commonNames.VPOPEXPTABLEVARNAMESFIXED; 
    else
        tableVariableNamesFixed = commonNames.VPOPRECISTEXPTABLEVARNAMESFIXED;
    end

	
    [nRows,nCols] = size(expDataTable);
    nHeaderCols = length(tableVariableNamesFixed);
    
    dataRows = nHeaderCols+1 : (nCols-nHeaderCols);     
    
	outputRowCounter = 0;
    
    rowNumber1 = nan(nRows*(nRows-1),1);
    rowNumber2 = nan(nRows*(nRows-1),1);
    interventionID1 = cell(nRows*(nRows-1),1);
    interventionID2 = cell(nRows*(nRows-1),1);
    time1 = nan(nRows*(nRows-1),1);
    time2 = nan(nRows*(nRows-1),1);
    expN = nan(nRows*(nRows-1),1);
    elementID1 = cell(nRows*(nRows-1),1);
    elementID2 = cell(nRows*(nRows-1),1);
    expVarID1 = cell(nRows*(nRows-1),1);
    expVarID2 = cell(nRows*(nRows-1),1); 
    expDataID1 = cell(nRows*(nRows-1),1);
    expDataID2 = cell(nRows*(nRows-1),1);
    data1 = cell(nRows*(nRows-1),1);
    data2 = cell(nRows*(nRows-1),1);
    corCoeff = nan(nRows*(nRows-1),1);
    pVal = nan(nRows*(nRows-1),1);    
       
    for rowCounter1 = 1 : nRows
		for rowCounter2 = rowCounter1+1 : nRows
			if rowCounter1 ~= rowCounter2
				row1Data = expDataTable{rowCounter1,dataRows};
				row2Data = expDataTable{rowCounter2,dataRows};
				keepCols=find(~isnan(row1Data) & ~isnan(row2Data));
				if length(keepCols) > 2
					outputRowCounter = outputRowCounter+1;
					[R,P] = corrcoef(row1Data(keepCols)',row2Data(keepCols)');
					rowNumber1(outputRowCounter) = rowCounter1;
					rowNumber2(outputRowCounter) = rowCounter2;
					interventionID1{outputRowCounter} = expDataTable{rowCounter1,'interventionID'};
					interventionID2{outputRowCounter} = expDataTable{rowCounter2,'interventionID'};
					time1(outputRowCounter) = expDataTable{rowCounter1,'time'};
					time2(outputRowCounter) = expDataTable{rowCounter2,'time'};
                    expN(outputRowCounter) = length(keepCols);
					elementID1{outputRowCounter} = expDataTable{rowCounter1,'elementID'};
					elementID2{outputRowCounter} = expDataTable{rowCounter2,'elementID'};
					expVarID1{outputRowCounter} = expDataTable{rowCounter1,'expVarID'};
					expVarID2{outputRowCounter} = expDataTable{rowCounter2,'expVarID'};   
                    expDataID1{outputRowCounter} = expDataTable{rowCounter1,'expDataID'};
                    expDataID2{outputRowCounter} = expDataTable{rowCounter2,'expDataID'};
					corCoeff(outputRowCounter) = R(1,2);
					pVal(outputRowCounter) = P(1,2);
					data1{outputRowCounter} = row1Data(keepCols)';
					data2{outputRowCounter} = row2Data(keepCols)';
					
				end
			end
		end
	end
	
	if outputRowCounter > 0
		[sortP,sortIndices] = sort(pVal(1:outputRowCounter),'ascend');
        lastIndex = find(sortP<=0.05);
        if length(lastIndex) > 0
            sortIndices = sortIndices(1:length(lastIndex));
            myCorrelations = table(pVal(sortIndices),corCoeff(sortIndices),expN(sortIndices),rowNumber1(sortIndices),rowNumber2(sortIndices),time1(sortIndices),interventionID1(sortIndices),elementID1(sortIndices),expVarID1(sortIndices),expDataID1(sortIndices),time2(sortIndices),interventionID2(sortIndices),elementID2(sortIndices),expVarID2(sortIndices),expDataID2(sortIndices),data1(sortIndices),data2(sortIndices));
            myCorrelations.Properties.VariableNames = {'pVal','corCoeff','expN','rowNumber1','rowNumber2','time1','interventionID1','elementID1','expVarID1','expDataID1','time2','interventionID2','elementID2','expVarID2','expDataID2','data1','data2'};
            %myAutoRows = (myCorrelations{:,'elementID1'}==myCorrelations{:,'elementID2'});
            myAutoRows = cellfun(@isequal, myCorrelations{:,'elementID1'}, myCorrelations{:,'elementID2'});
            myCorrelationsAuto = myCorrelations(myAutoRows,:);
            myCorrelationsCross = myCorrelations(~myAutoRows,:);
        else
            rowNumber1 = [];
            rowNumber2 = [];
            interventionID1 = {};
            interventionID2 = {};
            time1 = [];
            time2 = [];
            expN = [];
            elementID1 = {};
            elementID2 = {};
            expVarID1 = {};
            expVarID2 = {};            
            data1 = {};
            data2 = {};
            corCoeff = [];
            pVal = [];
            myCorrelations = table(pVal,corCoeff,expN,rowNumber1,rowNumber2,time1,interventionID1,elementID1,expVarID1,expDataID1,time2,interventionID2,elementID2,expVarID2,expDataID2,data1,data2);
            myCorrelations.Properties.VariableNames = {'pVal','corCoeff','expN','rowNumber1','rowNumber2','time1','interventionID1','elementID1','expVarID1','expDataID1','time2','interventionID2','elementID2','expVarID2','expDataID2','data1','data2'};
            myCorrelationsCross = myCorrelations;
            myCorrelationsAuto = myCorrelations;
            warning(['Unable to find substantial correlations in ',mfilename,'.  Returning an empty myCorrelations.'])
        end
	else
        rowNumber1 = [];
        rowNumber2 = [];
        interventionID1 = {};
        interventionID2 = {};
        time1 = [];
        time2 = [];
        expN = [];
        elementID1 = {};
        elementID2 = {};
        expVarID1 = {};
        expVarID2 = {};        
        data1 = {};
        data2 = {};
        corCoeff = [];
        pVal = [];
        myCorrelations = table(pVal,corCoeff,expN,rowNumber1,rowNumber2,time1,interventionID1,elementID1,expVarID1,expDataID1,time2,interventionID2,elementID2,expVarID2,expDataID2,data1,data2);
        myCorrelations.Properties.VariableNames = {'pVal','corCoeff','expN','rowNumber1','rowNumber2','time1','interventionID1','elementID1','expVarID1','expDataID1','time2','interventionID2','elementID2','expVarID2','expDataID2','data1','data2'};
        myCorrelationsCross = myCorrelations;
        myCorrelationsAuto = myCorrelations;
        warning(['Unable to find substantial correlations in ',mfilename,'.  Returning empty outputs.'])
    end
	
else
    rowNumber1 = [];
    rowNumber2 = [];
    interventionID1 = {};
    interventionID2 = {};
    time1 = [];
    time2 = [];
    expN = [];
    elementID1 = {};
    elementID2 = {};
    expVarID1 = {};
    expVarID2 = {};
    data1 = {};
    data2 = {};
    corCoeff = [];
    pVal = [];
    myCorrelations = table(pVal,corCoeff,expN,rowNumber1,rowNumber2,time1,interventionID1,elementID1,expVarID1,expDataID1,time2,interventionID2,elementID2,expVarID2,expDataID2,data1,data2);
    myCorrelations.Properties.VariableNames = {'pVal','corCoeff','expN','rowNumber1','rowNumber2','time1','interventionID1','elementID1','expVarID1','expDataID1','time2','interventionID2','elementID2','expVarID2','expDataID2','data1','data2'};
    myCorrelationsCross = myCorrelations;
    myCorrelationsAuto = myCorrelations;

    warning(['Unable to run ',mfilename,'.  Returning empty outputs.'])
end
end
```
