## Supplementary File 3 for "A surrogate workflow for virtual population development: designing an amenable proxy for a hypothesis-test inspired nonlinear goodness-of-fit": checkLN.m

```
function [inputObj, myLNCheckTable] = checkLN(arg1, arg2)
% This function checks the assumptions for fitting the mean and standard
% deviation for a virtual population when we have the underlying data:
% i.e. are the data normally or lognormally distributed?  
% 
% ARGUMENTS:
%  arg1:               A virtual population OR a worksheet.
%  arg2:               (optional) If a worksheet is provided as arg1,
%                       a mapelOptions must be given.  This option is given
%                       so we can check the worksheet as soon as we have
%                       simulation results and the population targets set up.
%                       If a virtual population is given as arg1, then
%                       arg2 is not needed.
%
% RETURNS:
%  inputObj            An updated mapelOptions or VPop object, with 
%                       logN on mnSDTable updated to indicate the better 
%                       type of assumption.
%  myLNCheckTable:     A table of R-square slope and intercept from test probability
%                       plot relationships for normal and lognormal
%                       assumptions
%
%

% First check input arguments
if nargin > 2
    warning(['Too many input arguments to',mfilename,'. Requires: myVPop OR myWorksheet, myMapelOptions.'])
    flagContinue = false;
elseif nargin > 1
    myWorksheet = arg1;
    myMapelOptions = arg2;
    inputObj = myMapelOptions;
    myVPop = [];
    flagContinue = true;     
elseif nargin > 0
    myWorksheet = [];
    myMapelOptions = [];
    myVPop = arg1;    
    inputObj = myVPop;
    flagContinue = true;     
else
    warning(['Insufficient input arguments to ',mfilename,'. Requires: myVPop.'])
    flagContinue = false;    
end

newMnSDTable = [];
myLNCheckTable = [];

if (isempty(myVPop) && flagContinue)
    % First copy all of the needed properties over
    myVPop = initializeOptionPropertiesToVPop(myMapelOptions);
    if isa(myVPop,'VPopRECIST')
        myVPop.recistSimFilter = createRECISTSimFilter(myWorksheet, myVPop);
    end
    myVPop = myVPop.getSimData(myWorksheet);
    myVPop.subpopTable = updateSubpopTableVPs(myVPop.subpopTable, myWorksheet);
end

if flagContinue
    if isempty(myVPop.mnSDTable)
        flagContinue = false;
    else
        newMnSDTable = inputObj.mnSDTable;
    end
end

if flagContinue
    [nRows, ~] = size(newMnSDTable);
    myCheckElement = cell(0,5);
    if ~isempty(myVPop.mnSDTable)
        myCheckElement = [myCheckElement; table2cell(myVPop.mnSDTable(:,{'elementID','elementType','interventionID','time'}))];
    end
    [nCheckElements, ~] = size(myCheckElement);
    
     % Find column in ExpDataTable where data starts
     commonNames = loadCommonNames();
     myExpDataTable = myVPop.expData;
     if isa(myVPop, 'VPopRECIST')
         firstDataIndex = find(ismember(myExpDataTable.Properties.VariableNames, commonNames.VPOPRECISTTABLEVARNAMESFIXED{end}))+1;
     else
         firstDataIndex = find(ismember(myExpDataTable.Properties.VariableNames, commonNames.VPOPTABLEVARNAMESFIXED{end}))+1;
     end
     expDataTableRows = size(myExpDataTable,1); % number of experimental data
     
     rowInfoTable = cell2table(myVPop.simData.rowInfo,'VariableNames',myVPop.simData.rowInfoNames);
     expRowFinder = [myExpDataTable{:,'elementID'},myExpDataTable{:,'elementType'},myExpDataTable{:,'interventionID'},num2cell(myExpDataTable{:,'time'})];
    
     rsqN = nan(nCheckElements,1);
     rsqLN = nan(nCheckElements,1);
     slopeN = nan(nCheckElements,1);
     interceptN = nan(nCheckElements,1);
     slopeLN = nan(nCheckElements,1);
     interceptLN = nan(nCheckElements,1); 
     interceptLN = nan(nCheckElements,1); 
     logN = false(nCheckElements,1);
     
     for checkCounter = 1 : nCheckElements
         curCheckElements = myCheckElement(checkCounter, :);
         match  = strcmp(expRowFinder(:, 1), curCheckElements{1}) & ...
             strcmp(expRowFinder(:, 2), curCheckElements{2}) & ...
             strcmp(expRowFinder(:, 3), curCheckElements{3}) & ...
             cat(1, expRowFinder{:, 4}) == curCheckElements{4};
         expRow  = find(match);
         if (length(expRow) == 1)
             expData = myExpDataTable{expRow,firstDataIndex:end};
             expData = expData(~isnan(expData));
             expData = sort(expData, 'ascend');
             expN = length(expData);
             if expN >= 3
                 % From NIST handbook, see:
                 % https://www.itl.nist.gov/div898/handbook/eda/section3/normprpl.htm
                 uom = nan(1,expN);
                 uom(expN) = .5^(1/expN);
                 uom(1) = 1 - uom(expN);
                 for j = 2 : (expN-1)
                     uom(j) = (j - 0.3175)/(expN + 0.365);
                 end
             end
             myScores = zscore(uom);
             myScores = [ones(1,expN);myScores];
             % Get least-squares fit
             curFit = myScores'\expData';
             % Calculate residuals
             res1 = expData - curFit'*myScores;
             % Calculate R^2
             rsqN(checkCounter) = 1 - var(res1)/var(expData);
             slopeN(checkCounter) = curFit(2);
             interceptN(checkCounter) = curFit(1);
             % Repeat for lognormal
             if min(expData) > 0
                 expData = log(expData);
                 curFit = myScores'\expData';
                 % calculate residuals
                 res1 = expData - curFit'*myScores;
                 % calculate R^2
                 rsqLN(checkCounter) = 1 - var(res1)/var(expData);   
                 slopeLN(checkCounter) = curFit(2);
                 interceptLN(checkCounter) = curFit(1);   
             end
             if isnan(rsqLN(checkCounter))
                 logN(checkCounter) = false;
             elseif rsqLN(checkCounter) > rsqN(checkCounter)
                 logN(checkCounter) = true;
             end
             newMnSDTable{checkCounter,'logN'} = logN(checkCounter);
         end
     end
     inputObj.mnSDTable = newMnSDTable;
     myLNCheckTable = table(rsqN,rsqLN,slopeN,interceptN,slopeLN,interceptLN);
     myLNCheckTable = [myVPop.mnSDTable(:,{'elementID','elementType','interventionID','time'}),myLNCheckTable];
else
    warning(['Unable to proceed in ',mfilename,'.'])
end
         
         
         
    
```
