## Supplementary File 3 for "A surrogate workflow for virtual population development: designing an amenable proxy for a hypothesis-test inspired nonlinear goodness-of-fit": compositeGOF.m

```
function myVPop = compositeGOF(myVPop)
% Evaluate the composite goodness-of-fit (Fisher's combined test) for a 
% given axes weight solution
%
% ARGUMENTS
%  myVPop:     An instance of a VPop object. The fields should be
%              populated and individual p-values calculated before calling:
%               mnSDTable
%               binTable
%               gofMn
%               gofSD
%               gofBin
%               gofDist
%               gofDist2D
%               gofCor
%
% RETURNS
%  myVpop:     A VPop object with the composite GOF updated.
%

continueFlag = false;
if nargin > 1
    warning(['Too many input arguments provided to ',mfilename,'.  Requires: myVPop.'])
    continueFlag = false;
elseif nargin > 0
    continueFlag = true;
else
    continueFlag = false;
    warning(['Insufficient input arguments provided to ',mfilename,'.  Requires: myVPop.'])
end

if continueFlag
    if sum(ismember(class(myVPop),{'VPop','VPopRECIST'}))<1
        continueFlag = false;
        warning(['Input VPop not recognized in call to ',mfilename,'.'])
    end       
end

if continueFlag
    myMnSDTable = myVPop.mnSDTable;
    myBinTable = myVPop.binTable;
    myDistTable = myVPop.distTable;
    myDistTable2D = myVPop.distTable2D;	
	myCorTable = myVPop.corTable;    

    if ~isempty(myMnSDTable)
    % We will only include a term in the composite GOF if the corresponding
    % weight is > 0
        indicesMn = find(myMnSDTable.('weightMean') > 0);
        indicesSD = find(myMnSDTable.('weightSD') > 0);
        weightsMn = myMnSDTable.('weightMean')(indicesMn);
        weightsSD = myMnSDTable.('weightSD')(indicesSD);        
    else
        indicesMn = [];
        indicesSD = [];
        weightsMn = [];
        weightsSD = [];
    end
    if ~isempty(myBinTable)
        indicesBin = find(myBinTable.('weight') > 0);
        weightsBin = myBinTable.('weight')(indicesBin);
    else
        indicesBin = [];
        weightsBin = [];
    end
    if ~isempty(myDistTable)
        indicesDist = find(myDistTable.('weight') > 0);
        weightsDist = myDistTable.('weight')(indicesDist);
    else
        indicesDist = [];
        weightsDist = [];
    end    
    if ~isempty(myDistTable2D)
        indicesDist2D = find(myDistTable2D.('weight') > 0);
        weightsDist2D = myDistTable2D.('weight')(indicesDist2D);
    else
        indicesDist2D = [];
        weightsDist2D = [];
    end  	
    if ~isempty(myCorTable)
        indicesCor = find(myCorTable.('weight') > 0);
        weightsCor = myCorTable.('weight')(indicesCor);
    else
        indicesCor = [];
        weightsCor = [];
    end     
    allWeights = [weightsMn;weightsSD;weightsBin;weightsDist;weightsDist2D;weightsCor];
    pVals = [myVPop.gofMn(indicesMn);myVPop.gofSD(indicesSD);myVPop.gofBin(indicesBin);myVPop.gofDist(indicesDist);myVPop.gofDist2D(indicesDist2D);myVPop.gofCor(indicesCor)];
    if max(allWeights) == min(allWeights)
        dof = 2*length(pVals);
        fisherStat = -2*sum(log(pVals));
        myVPop.gof = chi2cdf(fisherStat,dof,'upper');
    else
        fisherStat = -2*sum(log(pVals).*allWeights);
%         myVPop.gof = woodF(allWeights,fisherStat);
        % hbe and woodP assume k df, not 2k.  replicating the entries
        % 2x corrects for this.
%         myVPop.gof = woodF(repmat(allWeights',1,2),fisherStat);        
        myVPop.gof = hbe(repmat(allWeights',1,2),fisherStat);
    end
        
else
    warning(['Unable to run ',mfilename,'.  Returning input.'])
end    
end
```
