## Supplementary File 3 for "A surrogate workflow for virtual population development: designing an amenable proxy for a hypothesis-test inspired nonlinear goodness-of-fit": contingency2N.m

```
function pVal = contingency2N(observations1, observations2, exactFlag)
% This function returns the p-value for a two-sided test of whether  
% the null hypothesis that two samples with four categories come from the
% same distribution.
% For small sample sizes, i.e. if any of the observations are < 5,
% an exact test is applied.  For sufficient sample sizes, the chisq  
% approximation is applied.
%
% ARGUMENTS
%  observations1: a 1xN vector of observations in each bin
%  observations2: a 1xN vector of observations in each bin
%  exactFlag:     boolean, whether to check if an exact test should be
%                 performed.  If false, the chi-square approximation is
%                 imposed.
%
% RETURNS
%  pVal
%
pVal = nan;
continueFlag = true;
[nRows1, nBins1] = size(observations1);
[nRows2, nBins2] = size(observations2);
if ((nBins1~=nBins2) || (nRows1~=1) || (nRows2~=1) || (nBins1<2))
    continueFlag = false;
    warning(['Unable to continue in ',mfilename,', two 1xN vectors of categorical observation counts should be supplied.'])
end

if continueFlag
	nBins = nBins1;
    no1t = sum(observations1);
    no2t = sum(observations2);
	epAll = nan(1,nBins);
	for epCounter = 1 : nBins
		epAll(epCounter) = (observations1(epCounter) + observations2(epCounter)) / (no1t + no2t);
	end
    % We use the rule of thumb:
    % mentioned in several places, including:
    % McHugh, ML, Biochem Med (Zagreb). 2013 Jun; 23(2): 143�149.
    % No more than 20% of the expected counts are less than 5 and 
    % all individual expected counts are 1 for the chisq approximation
    % to hold
    minFail = min(min([no1t, no2t]) * epAll) < 1;
    fiveFail = sum(sum(([no1t; no2t] * epAll) < 5)) > (0.2*(2*nBins));
    zeroFail = min(epAll) <= 0;
	if ((minFail || fiveFail) & (exactFlag || zeroFail))
		chisqFlag = false;
	else
		chisqFlag = true;
	end
end

if continueFlag
    if chisqFlag
		expected = [epAll * no1t, epAll * no2t];
        % We already apply our own check on the minimum number of observations
        % per bin.  Don't use the default 'emin' of 5.
        [h,pVal,stats] = chi2gof([1:(nBins*2)],'freq',[observations1, observations2],'expected',expected,'ctrs',[1:(nBins*2)],'nparams',nBins,'emin',0);
    else
        x = [observations1; observations2];
		% We use the 3rd party function for these 
		% calculations where chi-square approximation
		% is likely not good.
		if nBins < 3
			% Note evalc is used to supress the 
			% text displayed during function execution
			[T,pVal] = evalc('myfisher22(x);');		
			% Here, pVal is returned as a 1X3 p value matrix of [left tail, right tail, 2 tail]
			pVal = pVal(3);
		elseif nBins < 4
			% Note evalc is used to supress the 
			% text displayed during function execution
			[T,pVal] = evalc('myfisher23(x);');	
		elseif nBins < 5
			% There were some issues that we have debugged.
			% Note evalc is used to supress the 
			% text displayed during function execution
			[T,pVal] = evalc('myfisher24(x);');
		else
			% There were some issues that we have debugged.
			% Note evalc is used to supress the 
			% text displayed during function execution
			[T,pVal] = evalc('myfisher(x);');	
		end
    end
else
    warning(['Unable to continue in ',mfilename,', two 1xnBin vectors of categorical observation counts should be supplied.'])
end
end
```
