## Supplementary File 3 for "A surrogate workflow for virtual population development: designing an amenable proxy for a hypothesis-test inspired nonlinear goodness-of-fit": hbe.m

```
function myPval = hbe(coeff, x)
% This function applies Hall-Buckley-Eagleson method to give a p-value
% based on a positively-weighted sum of chi-squared
% random variables.
%
%
% References:
% Buckley, M.J., Eagleson, G.K.: An approximation to the distribution of
% quadratic forms in normal random variables. Aust. J. Stat. 30(1),
% 150–159 (1988)
%
% Hall, P.: Chi squared approximations to the distribution of a sum of
% independent random variables. Ann. Probab. 11(4), 1028–1036
% (1983)
%
% See also:
% Bodenham, D. A. & Adams, N. M. A comparison of efficient 
% approximations for a weighted sum of chi-squared random 
% variables. Stat Comput 26, 917–928 (2016).
%
% ARGUMENTS
%  coeff:   A vector of weighting coefficients
%  x:       The test statistic value
%
% RETURNS
%  myPval: 

c1 = sum(coeff);
c2 = 2*sum(coeff.^2);
c3 = 8*sum(coeff.^3);
nu = 8 * (c2^3) / (c3^2);
gk = nu/2;
gtheta = 2;
xchisqnu = sqrt(2 * nu / c2) * (x - c1) + nu;
myPval = gamcdf(xchisqnu, gk, gtheta, 'upper');

end
```
