## Supplementary File 3 for "A surrogate workflow for virtual population development: designing an amenable proxy for a hypothesis-test inspired nonlinear goodness-of-fit": predNPDTFLS.m

```
function myVPop = predNPDTFLS(myVPop)
% This function takes a VPop object and updates with 
% predictions for number of PD patients.
%
% ARGUMENTS
%  myVPop:                  (Required) a mapelOptionsRECIST or VPopRECIST
%                                     instance.
%
% Returns
%  myVPop:                  VPopRECIST with updated brTableRECIST.  This 
%                            gives predNPD21LS values.
%                            
%


continueFlag = true;
if nargin > 1
    warning(['Too many input arguments provided to ',mfilename,'.  Requires: VPopRECIST object.'])
    continueFlag = false;
elseif nargin < 1
    continueFlag = false;
    warning(['Insufficient input arguments provided to ',mfilename,'.  Requires: VPopRECIST object.'])
end

if continueFlag
    if ~ismember(class(myVPop),{'VPopRECIST'})
        continueFlag = false;
        warning(['Input myVPop not recognized in call to ',mfilename,'.  Requires: VPopRECIST object.'])
    end       
end


if continueFlag
    lmResults = regressPD2ScaleFactor(myVPop); % updated: use PD, SD as dependent variable, intercept=0
    % Update the values for the 
    % First find the blocks by the PD2 measures
    predRows = find(sum(isnan(myVPop.brTableRECIST{:,{'predN','predCR','predPR','predSD','predPD'}}),2)==0);
    predN = myVPop.brTableRECIST{predRows,{'predN'}};
    predPD = myVPop.brTableRECIST{predRows,{'predPD'}}.*predN;
    predSD = myVPop.brTableRECIST{predRows,{'predSD'}}.*predN;

    res = predPD.*lmResults.Coefficients{1,'Estimate'} + predSD.*lmResults.Coefficients{2,'Estimate'};
    myVPop.brTableRECIST{predRows,'predNPD21LS'}=res;
else
    lmResults = {};
    warning(['Unable to run ',mfilename,'.'])
end
end
```
