## Supplementary File 3 for "A surrogate workflow for virtual population development: designing an amenable proxy for a hypothesis-test inspired nonlinear goodness-of-fit": regressPD2ScaleFactor.m

```
function lmResults = regressPD2ScaleFactor(myMapelOptions)
% This function takes a mapelOptions or VPop object and returns 
% coefficients for a predicted PD2 scaling rate.
%
% ARGUMENTS
%  myMapelOptions:         (Required) a mapelOptionsRECIST or VPopRECIST
%                                     instance.
%
% Returns
%  lmResults:               Output of the regression.  This gives the
%                            a regression model for the observed PD normalized 
%                            to the number of patients recorded as nonPD2 
%                            (expN) at the first lesion scan.  Multiply by 
%                            expN to get the predicted expNPD21LS.  To
%                            given an expNPD21LS, one could use the
%                            predSD, predPD with the regression coefficients
%                            and scale by the the predN.
%                            
%


continueFlag = true;
if nargin > 1
    warning(['Too many input arguments provided to ',mfilename,'.  Requires: mapelOptionsRECIST or VPopRECIST object.'])
    continueFlag = false;
elseif nargin < 1
    continueFlag = false;
    warning(['Insufficient input arguments provided to ',mfilename,'.  Requires: mapelOptionsRECIST or VPopRECIST object.'])
end

if continueFlag
    if ~ismember(class(myMapelOptions),{'mapelOptionsRECIST','VPopRECIST'})
        continueFlag = false;
        warning(['Input not recognized in call to ',mfilename,'.  Requires: mapelOptionsRECIST or VPopRECIST object.'])
    end       
end


if continueFlag
    % First find the blocks by the PD2 measures
    % Only keep the last time point from each inidividual intervention (BOR), then train the regression model
     testData = myMapelOptions.brTableRECIST(:,{'subpopNo','expDataID','interventionID'}); 
     [C,IA,IC] = unique(testData,'rows','stable'); % unique(testData,'last','rows','stable');
     LastRowindex=IA;
     for j=1:length(IA)
         indices=find(IC==j);
         LastRowindex(j)=indices(end);
     end            
    testRows = find(~isnan(myMapelOptions.brTableRECIST{:,{'expNPD21LS'}})); 
    regRows = intersect(testRows, LastRowindex);
    expN = table2array(myMapelOptions.brTableRECIST(regRows,'expN'));
    expPD=table2array(myMapelOptions.brTableRECIST(regRows,'expPD')).*expN;
    expSD=table2array(myMapelOptions.brTableRECIST(regRows,'expSD')).*expN;
    resp=table2array(myMapelOptions.brTableRECIST(regRows,'expNPD21LS'));
    fitTable = table(expPD,expSD,resp);
    lmResults = fitlm(fitTable,'resp ~ expPD + expSD','Intercept',false);
else
    lmResults = {};
    warning(['Unable to run ',mfilename,'.  Returning an empty object.'])
end
end
```
