## Supplementary File 3 for "A surrogate workflow for virtual population development: designing an amenable proxy for a hypothesis-test inspired nonlinear goodness-of-fit": weightedKS.m

```
function myPVal = weightedKS(sample1, sample2, W1, W2, N1, N2)
% This function performs a KS test. Rather than assuming the
% percentiles are computed from the rank-order it 
% adjusts the percentiles for observations according to the
% provided weights.
%
% References:
% Press et al, Numerical Recipes - The Art of Scientific Computing
% (2007) P 736-740
% Stevens MA. Use of the Kolmogorov-Smirnov, Cramer-Von Mises and 
% Related Statistics Without Extensive Tables. 
% Journal of the Royal Statistical Society Series B (Methodological). 
% 1970;32(1):115-122.
%
%
% ARGUMENTS
%  sample1     observed values for sample1. These are assumed to be sorted
%              in ascending order
%  sample2     observed values for sample2. These are assumed to be sorted
%              in ascending order    
%  W1          a 1xlength(sample1) vector of weights for sample 1
%  W2          a 1xlength(sample2) vector of weights for sample 2
%  N1          number of observations underlying sample1
%  N2          number of observations underlying sample2
%
% RETURNS
%  myPVal      the p-value resulting from the two-tailed test

    % First calculate CDF along the sample sample axis
    [CDF1, CDF2, SC] = alignCDFs(sample1, sample2, W1, W2);
    % See Press et al, Numerical Recipes - The Art of Scientific Computing
    % (2007) P 736-740	
    D = max(abs(CDF2-CDF1));
    nPool =  N1 * N2 /(N1 + N2);
    zVal =  max((sqrt(nPool) + 0.12 + 0.11/sqrt(nPool)) * D , 0);
	% See page 334, special functions, in Numerical Recipes
	% This is an infinite series.  Here we assume 100 terms is enough
	% TODO: may want to add a check to validate desired precision
    % for the series
    iVals =  (1:100)';
    myPVal  =  2 * sum((-1).^(iVals-1).*exp(-2*zVal^2*iVals.^2));
    myPVal  =  min(max(myPVal, 0), 1);
end
    
```
