## Supplementary File 3 for "A surrogate workflow for virtual population development: designing an amenable proxy for a hypothesis-test inspired nonlinear goodness-of-fit": woodF.m

```
function myPval = woodF(coeff, x)
% This function applied Wood's F method to give a p-value
% based on a positively-weighted sum of chi-squared
% random variables.
%
% Note there is a fundamental issue where
% Wood's F method will be unable to match moments 
% for some cases. 
% A simple example is when the coefficient vector is of length 1.
% These cases are not common.  Hall-Buckley-Eagleson method is
% also available.
%
% Reference:
% Wood, A.T.A.: An F approximation to the distribution of a 
% linear combination of chi-squared variables. Commun. Stat. Simul. Comput.
% 18(4), 1439�1456 (1989)
%
% See also:
% Bodenham, D. A. & Adams, N. M. A comparison of efficient 
% approximations for a weighted sum of chi-squared random 
% variables. Stat Comput 26, 917�928 (2016).
%
% ARGUMENTS
%  coeff:   A vector of weighting coefficients
%  x:       The test statistic value
%
% RETURNS
%  myPval: 
%

c1 = sum(coeff);
c2 = 2*sum(coeff.^2);
c3 = 8*sum(coeff.^3);
	
r1 = 4 * c2^2 * c1  +  c3 * (c2 - c1^2);
r2 = c3 * c1 - 2* c2^2;
beta = r1/r2;
alpha1 = 2*c1 * (c3*c1  +  c1^2 * c2 - c2^2)/r1;
alpha2 = 3 + 2*c2*(c2 + c1^2)/r2;
x = x * alpha2/(alpha1*beta);
myPval = 1-fcdf(x, 2*alpha1, 2*alpha2);
end
```
