## Supplementary File 3 for "A surrogate workflow for virtual population development: designing an amenable proxy for a hypothesis-test inspired nonlinear goodness-of-fit": wtdBinProb.m

```
function wpp = wtdBinProb(measVals, pws, binEdgeValues)
% This function calculates a discrete/binned pdf based on observations and
% weights.  Here, we strictly enforce the first dimension is 1
% so there are no errors.
%
% ARGUMENTS
%  measVals:       A 1 X nVP vector of measurement values
%  pws:            A 1 X nVP vector of prevalence weights
%  binEdgeValues:  A 1 X Nbin-1 vector of bin dividing edge values.  Note
%                  that values smaller than the first bin edge are counted
%                  in the first bin and values >= than the upper bin edge
%                  are counted in the last returned bin.
%
% RETURNS
%  wpp:            A 1 X Nbin vector of weighted pdf in each bin
%
continueFlag = true;
wpp = nan;
numericTolerance = max(eps(1) * length(pws) * 10,1E-14);
% We just check if matrix/vector dimensions are OK here.
if nargin > 3
    continueFlag = false;
    warning(['A 1 x nVP vector of values and weights, and vector of inner bin edges, are all required for ',mfilename,'. Too many input arguments. Returning NaN.'])
elseif nargin > 2
    continueFlag = true;
else
    continueFlag = false;
    warning(['A 1 x nVP vector of values and weights, and vector of inner bin edges, are all required for ',mfilename,'. Insufficient input arguments. Returning NaN.'])
end

if continueFlag
    [size1r, size1c] = size(measVals);
    [size2r, size2c] = size(pws);
    [size3r, size3c] = size(binEdgeValues);
    if (size1c~= size2c)
        continueFlag = false;
    end
    if ((size1r ~= 1) || (size2r ~= 1) || (size3r ~= 1))
        continueFlag = false;
    end
    if (size3c < 1)
        continueFlag = false;
    end    
    if ~continueFlag
        warning(['A 1 x nVP vector of values and weights, and 1 X (nBin-1) vector of inner bin edges, are all required for ',mfilename,'. Returning NaN.'])
    end
end

if continueFlag
    if ((sum(pws) > (1 + numericTolerance)) || (sum(pws) < (1 - numericTolerance)))
        warning(['Supplied pws to ',mfilename,' should sum to 1. Returning NaN.'])
        continueFlag = false;
    end
end  

if continueFlag
    catMat = cat(1, measVals, pws);
    [sortVal, Indices] = sort(measVals);
    catMat = catMat(:,Indices);
    nBins = length(binEdgeValues) + 1;
    wpp = zeros(1,length(nBins));
    curIndices = catMat(1,:) < binEdgeValues(1);
    wpp(1,1) = sum(curIndices .* catMat(2,:));
    for binCounter = 2 : (nBins-1)
        curIndices = catMat(1,:) < binEdgeValues(binCounter);
        wpp(1,binCounter) = sum(curIndices .* catMat(2,:)) - sum(wpp(1:(binCounter-1)));
    end
    curIndices = catMat(1,:) >= binEdgeValues(nBins-1);
    wpp(1,nBins) = sum(curIndices .* catMat(2,:));
end

end
```
