## Supplementary File 3 for "A surrogate workflow for virtual population development: designing an amenable proxy for a hypothesis-test inspired nonlinear goodness-of-fit": wtdMean.m

```
function wmn = wtdMean(measVals,pws)
% Calculation of the weighted mean.
%
% ARGUMENTS
%  measVals: A 1 X nVP vector of measurement values
%  pws:      A 1 X nVP vector of prevalence weights
%
% RETURNS
%  wmn:      weighted mean
%

continueFlag = true;
wmn = nan;
% We just check if matrix/vector dimensions are OK here.
if nargin > 2
    continueFlag = false;
    warning(['A 1 x nVP (or nVP X 1) vector of values and weights are both required for ',mfilename,'. Too many input arguments.  Returning NaN.'])
elseif nargin > 1
    continueFlag = true;
else
    continueFlag = false;
    warning(['A 1 x nVP (or nVP X 1) vector of values and weights are both required for ',mfilename,'. Insufficient input arguments.  Returning NaN.'])
end

if continueFlag
    [size1r, size1c] = size(measVals);
    [size2r, size2c] = size(pws);	
    if ((size1r~= size2r) || (size1c~= size2c))
        continueFlag = false;
    end
    if (min(size1r, size1c) ~= 1) || (min(size2r, size2c) ~= 1)
        continueFlag = false;
    end
    if ~continueFlag
        warning(['A 1 x nVP (or nVP X 1) vector of values and weights are both required for ',mfilename,'. Returning NaN.'])
    end
end

if continueFlag
	% We allow an error of eps * the number of VPs * 10
	% We could in strictly enforce eps * the number of VPs but
	% this criterion is relaxed a little.
	% 10^-14 was used as a minimum in the bin probabilities.  The
	% bin probabilities should all be fixed and
	% PWs should all be corrected to sum to 1, but
	% here 10^-14 is set as a minimum allowed error in the check
	numericTolerance = max(eps(1) * length(pws) * 10,1E-14);
    if ((sum(pws) > (1 + numericTolerance)) || (sum(pws) < (1 - numericTolerance)))
        warning(['Supplied pws to ',mfilename,' should sum to 1. Returning NaN.'])
        continueFlag = false;
    end
end  

if continueFlag
    wmn = sum(pws .* measVals);
end

end
```
