## Supplementary File 3 for "A surrogate workflow for virtual population development: designing an amenable proxy for a hypothesis-test inspired nonlinear goodness-of-fit": testExpandVPopSettings.m

```
function myTestResultTable = testExpandVPopSettings(myWorksheet,myMapelOptions, myScreenFunctionName)
% This function takes an input worksheet and mapelOptions, identifies a VP
% to expand around, and then tests different expansion settings.  It reports
% back a table with the summary information fromt he expansion tests.
% This function is provided as a potential precursor to expandVPopEffN,
% to get an idea of anticipated performance with different settings.
%
% ARGUMENTS
%  myWorksheet:         A worksheet.  All responseTypes are used for screening.
%  myMapelOptions:      A mapelOptions object
%  myScreenFunctionName A pre-simulation screening function
%
% RETURNS
%  myTestResultTable:   A table with useful information for different expansion settings
%

continueFlag = false;
if nargin > 3
    warning(['Too many input arguments to ',mfilename, '. Arguments should be: myWorksheet, myMapelOptions, myScreenFunctionName.'])
    continueFlag = false;
    hotStartVPop = '';    
elseif nargin > 2
    continueFlag = true;
else 
    warning(['Too few input arguments to ',mfilename, '. Arguments should be: myWorksheet, myMapelOptions, myScreenFunctionName.'])
    continueFlag = false;
end


if continueFlag
	
    % Set these for fast optimization
    myMapelOptions.exactFlag = false;
	% Enforce using effN, since
	% we enforce using it for the VPops
    myMapelOptions.useEffN = true;
	% Also minimize pool restarts
	myMapelOptions.poolRestart=false;
	myMapelOptions.poolClose=false;
	% Be very loose with the required effN, just need
	% one reasonable VP for resampling
	myMapelOptions.minEffN = 0;
	
	allResponseTypeIDs = getResponseTypeIDs(myWorksheet);
	myScreenTable = createScreenTable(myWorksheet, allResponseTypeIDs, true);
	
	% We will manually refresh the pool once at the start.
	if ~isempty(gcp('nocreate'))
		delete(gcp);
	end
	if isempty(gcp('nocreate'))
		% We will use default pool settings
		mySimulateOptions = simulateOptions;
		mySimulateOptions = checkNWorkers(mySimulateOptions);		
		myPool = parpool(mySimulateOptions.clusterID,mySimulateOptions.nWorkers,'SpmdEnabled',false);
	end	    

    % Make sure results are present.  Here, we don't re-simulate VPs if 
    % results are present.
    mySimulateOptions.rerunExisting=false;
	mySimulateOptions.poolRestart=false;
	mySimulateOptions.poolClose=false; 
    myWorksheet = simulateWorksheet(myWorksheet,mySimulateOptions);
	
    % Make sure results are present.  Here, we don't re-simulate VPs if 
    % results are present.
    myWorksheet = simulateWorksheet(myWorksheet);
	
	% Seed the rng if requested
	if myMapelOptions.intSeed > -1
		rng(myMapelOptions.intSeed, 'twister');
	end  
	
	disp(['Runnning a VPop optimization to identify a test VP for expansion in ',mfilename,'.'])
    % Set this small just so it runs fast.
    myMapelOptions.optimizePopSize = 1000;
    myVPop.optimizeType = 'gapso';
	myVPop = mapel(myWorksheet, myMapelOptions);
    curVPopEffN = 1/sum(myVPop.pws.^2);
    disp(['VPop solution effN ', num2str(curVPopEffN), ' with pvalue ',num2str(myVPop.gof),' in ',mfilename,'.  Proceeding to expansion test.'])
	
	[~,baseVPIndices] = sort(myVPop.pws,'descend');
    baseVPIndices = baseVPIndices(1:4);
	originalVPIDs = getVPIDs(myWorksheet);
	baseVPIDs = originalVPIDs(baseVPIndices);
	
	% Now start a loop over the test conditions
	nPerVPTest = 50;
	resampleStd = [5; 4; 2; 1; 0.5; 0.25; 0.1; 0.3; 0.2; .1; .05; 0.01];
	varyMethod = {'localpca';'localpca';'localpca';'localpca';'localpca';'localpca';'localpca';'gaussian';'gaussian';'gaussian';'gaussian';'gaussian'};
	nTests = length(resampleStd);    
	passRate = nan(nTests, 1);
	vpDistance = nan(nTests, 1);
    allAxis = getAxisDefIDs(myWorksheet);
    nAxis = length(allAxis);
	
	for testCounter = 1 : nTests
		myVaryAxesOptions = varyAxesOptions;
		myVaryAxesOptions.varyMethod = varyMethod{testCounter};
		myVaryAxesOptions.gaussianStd = resampleStd(testCounter);
		myVaryAxesOptions.varyAxisIDs = allAxis;
		myVaryAxesOptions.intSeed = -1;
		myVaryAxesOptions.baseVPIDs = baseVPIDs;
		myVaryAxesOptions.newPerOld = nPerVPTest;
		myVaryAxesOptions.additionalIDString = 'resampleTest';
		jitteredWorksheet = addVariedVPs(myWorksheet, myVaryAxesOptions);
		curVPIDs = getVPIDs(jitteredWorksheet);
		newIndices = (find(~ismember(curVPIDs,originalVPIDs)));
		newVPIDs = curVPIDs(newIndices);   

		% We will also randomize a coefficient for the new VPs, like
		% in the resampling method
		for vpCounter = 1 : length(newVPIDs);
			curVPID = newVPIDs{vpCounter};
			% randomize one of the new VP axis coefficients
			axisIndex = randsample([1:nAxis],1);
			vpIndex = find(ismember(curVPIDs,curVPID));
			jitteredWorksheet.axisProps.axisVP.coefficients(axisIndex,vpIndex) = rand(1);
		end

		mySimulateOptions = simulateOptions;
		mySimulateOptions.rerunExisting = false;
		mySimulateOptions.optimizeType = 'none';
		% Inherit the pool properties
		mySimulateOptions.poolRestart = false;
		mySimulateOptions.poolClose = false;    
    
		% Also screen the worksheet if a function is provided
		if length(myScreenFunctionName) > 0
			jitteredWorksheet = eval([myScreenFunctionName,'(jitteredWorksheet,newVPIDs,mySimulateOptions)']);
		end
    
		jitteredWorksheet = simulateWorksheet(jitteredWorksheet,mySimulateOptions);
		curVPIDs = getVPIDs(jitteredWorksheet);
		originalIndices = (find(ismember(curVPIDs,originalVPIDs)));
		newIndices = (find(~ismember(curVPIDs,originalVPIDs)));
		newVPIDs = curVPIDs(newIndices); 
		nSimulated = length(newIndices);        

		% Identify the VPs that don't fulfill the worksheet response
		% as well as those in the initial worksheet in order to filter
		% them.
		disp(['Screening results from ',varyMethod{testCounter},' with variability setting ',num2str(resampleStd(testCounter)),' in ',mfilename,'.'])        
		jitteredWorksheet = screenWorksheetVPs(jitteredWorksheet, myScreenTable, true, newVPIDs);
		curVPIDs = getVPIDs(jitteredWorksheet);
		originalIndices = (find(ismember(curVPIDs,originalVPIDs)));
		newIndices = (find(~ismember(curVPIDs,originalVPIDs)));
		newVPIDs = curVPIDs(newIndices); 

		passRate(testCounter) = length(newIndices)/nSimulated;
		
        % Also get the distances but distinguish
        % VPs sampled around the same parent.
        allCoefficients = getVPCoeffs(jitteredWorksheet);
        curDistances = zeros(1,length(baseVPIDs));
        curN = zeros(1,length(baseVPIDs));
        for parentCounter = 1:length(baseVPIDs)
            curChildrenBaseIndices = cellfun(@isempty,strfind(curVPIDs,[baseVPIDs{parentCounter},'_resampleTest_']));
            curChildrenBaseIndices = [find(ismember(curVPIDs,baseVPIDs{parentCounter})),  find(~curChildrenBaseIndices)];
            % We can leave the parent in when calculating distances.
            if length(curChildrenBaseIndices) > 1
                curCoefficients = allCoefficients(:,curChildrenBaseIndices);
                P = pdist2(curCoefficients',curCoefficients').^2;
                curDistances(parentCounter) = mean(P(P>0));
                curN(parentCounter) = length(curChildrenBaseIndices);
            end
        end
        
        if sum(curN) > 0
            vpDistance(testCounter)=sum((curDistances(curN>0).*curN(curN>0))/sum(curN(curN>0)));
        end

	end
	
	myTestResultTable = table(varyMethod,resampleStd,passRate,vpDistance);
	%myTestResultTable.Properties.VariableNames = {'responseTypeID','responseTypeElementID','responseTypeElementType','interventionID','modelYVar','modelYVarType','weight','value'};

else
	warning(['Unable to proceed in ',mfilename,'.  Exiting.'])
	myTestResultTable = [];
end
end
```
