## Supplementary File 3 for "A surrogate workflow for virtual population development: designing an amenable proxy for a hypothesis-test inspired nonlinear goodness-of-fit": align2DPDFs.m

```
function [data1pdf, data2pdf, combinedPoints] = align2DPDFs(data1, data2, data2PWs)
% This function takes two samples and weights, and returns
% adjusted sample and CDF vectors that can be compared element-wise
%
% ARGUMENTS
%  data1       observed values for sample1. These need not be sorted,
%               but should be provided as a 2xN1 matrix
%  data2       observed values for sample2. These need not be sorted,
%               but should be provided as a 2xN2 matrix.
%  data2PWs    a 1xN2 vector of weights for sample 2.  It is assumed
%               all observations in sample1 are weighted equally.
%
% RETURNS
%  data1pdf:       a pdf for data in sample1 mapped onto the combinedPoints
%  data2pdf:       a pdf for data in sample2 mapped onto the combinedPoints
%  combinedPoints: a set of points that the pdfs in data1 and data2 are mapped onto.

% First combine points, to get ranges
combinedPoints = [(unique(data1','rows'))',(unique(data2','rows'))'];

% We will apply smoothing to the PDFs.  Calculate a bandwidth to use
% base on the range in the data
bw1 = (max(combinedPoints(1,:),[],2)-min(combinedPoints(1,:),[],2))/10;
bw2 = (max(combinedPoints(2,:),[],2)-min(combinedPoints(2,:),[],2))/10;

% Get the experimental density onto the model sampled points, with
% smoothing
data1pdf = ksdensity(data1',combinedPoints','Bandwidth',[bw1;bw2]);
f = scatteredInterpolant(combinedPoints(1,:)', combinedPoints(2,:)', data1pdf, 'nearest');

% Unfortunately, we have to integrate over discontinuous surfaces (see
% below)
% and it's very likely there will be some residual error.  We may want
% to revisit this in future MATLAB releases.
warning('off','MATLAB:integral2:maxFunEvalsPass');
% We may want to turn this back on with improved performance of
% the 2D integration
warning('off','MATLAB:integral2:maxFunEvalsFail');
% We use large tolerances.  int does not have to be very precise,
% and MATLAB's integral2 function can use much overhead if
% we require a small tolerance.
int = integral2(@(x,y) f(x,y), min(combinedPoints(1,:)), max(combinedPoints(1,:)), min(combinedPoints(2,:)), max(combinedPoints(2,:)),'method','tiled','AbsTol',1e-2,'RelTol',1e-1);

% Normalize the 2D PDF
data1pdf = data1pdf/int;

% We will resample from data2 before smoothing.  We can't apply
% PWs in the smoothing function directly
% Note: we may want to adjust the rng seed as an additional
% input to make this reproducible
% rng(0);
data2pdf = datasample(data2', 1E3, 'Weights',data2PWs');
data2pdf = ksdensity(data2pdf,combinedPoints','Bandwidth',[bw1;bw2]);
% We have to use 'nearest', which is discontinuous but
% the only practical option.  Otherwise, MATLAB may calculate a
% negative interpolant with the available options, which does not 
% make sense for a pdf surface
f = scatteredInterpolant(combinedPoints(1,:)', combinedPoints(2,:)', data2pdf, 'nearest');
int = integral2(@(x,y) f(x,y), min(combinedPoints(1,:)), max(combinedPoints(1,:)), min(combinedPoints(2,:)), max(combinedPoints(2,:)),'method','tiled','AbsTol',1e-2,'RelTol',1e-1);
data2pdf = data2pdf/int;
warning('on','MATLAB:integral2:maxFunEvalsPass');
warning('on','MATLAB:integral2:maxFunEvalsFail');

end
```
