## Supplementary File 3 for "A surrogate workflow for virtual population development: designing an amenable proxy for a hypothesis-test inspired nonlinear goodness-of-fit": calculateExpWeight.m

```
function w = calculateExpWeight(expN, expSTD, dataGroupDescription, simN)
% This is a utility function to find calculation of
% different weights for the different data group types
% in linear calibrate.
%
% ARGUMENTS
%  expN:                  An experimental N
%  expSTD:                An experimental standard deviation  
%  dataGroupDescription:  Data group description.  Should be one of:
%							binTable
%                           distTable
%                           brTableRECIST
%                           rTableRECIST
%                           mnSDTablemean
%                           mnSDTablevariance      
%  simN:                  Assumed N for weighting for the simulation results
% RETURNS
%  w:                     A weighting factor.
%

if strcmp(dataGroupDescription(1:length('binTable')), 'binTable')
                w = sqrt((expN*simN)/(expN+simN));
 
elseif strcmp(dataGroupDescription(1:length('distTable')), 'distTable')
                w = sqrt((expN*simN)/(expN+simN));
                
elseif strcmp(dataGroupDescription(1:length('distTable2D')), 'distTable2D')
                w = sqrt((expN*simN)/(expN+simN));
 
elseif strcmp(dataGroupDescription(1:length('brTableRECIST')), 'brTableRECIST')
                w = sqrt((expN*simN)/(expN+simN));
 
elseif strcmp(dataGroupDescription(1:length('rTableRECIST')), 'rTableRECIST')
                w = sqrt((expN*simN)/(expN+simN));
 
elseif strcmp(dataGroupDescription(1:length('mnSDTable')), 'mnSDTable') && strcmp(dataGroupDescription(end-length('mean')+1:end), 'mean')
                w = 1/(expSTD*sqrt(1/expN+1/simN));
 
elseif strcmp(dataGroupDescription(1:length('mnSDTable')), 'mnSDTable') && strcmp(dataGroupDescription(end-length('variance')+1:end), 'variance')
                w = 1/(expSTD*sqrt(1/expN+1/simN));
elseif strcmp(dataGroupDescription(1:length('corTable')), 'corTable')
    if expN>3 && simN>3
        w = 1/sqrt(1/(expN-3)+1/(simN-3));
    else
        warning(['Either expN or simN is less than or equal to 3, modify weight calculation in ' mfilename ' to avoid NaN'])
        w = 1/sqrt(1/(expN)+1/(simN));
    end
else
                error('data group not supported');
end
 
end
```
