## Supplementary File 3 for "A surrogate workflow for virtual population development: designing an amenable proxy for a hypothesis-test inspired nonlinear goodness-of-fit": calculateExpWeightFix.m

```
function w = calculateExpWeightFix(expN, expSTD, dataGroupDescription, simN)
% This is a utility function to fix the experimental
% weight to 1 for the different data group types
% in linear calibrate.
%
% ARGUMENTS
%  expN:                  An experimental N, these are ignored here
%  expSTD:                An experimental standard deviation  
%  dataGroupDescription:  Data group description.  Should be one of:
%							binTable
%                           distTable
%                           brTableRECIST
%                           rTableRECIST
%                           mnSDTablemean
%                           mnSDTablevariance      
%                           corTable
%  simN:                  Assumed N for weighting for the simulation results
% RETURNS
%  w:                     A weighting factor.
%

% We will also try supplementing the VP scores scaled into PWs.  This
% likely won't be very effective at finding optimal solutions but will add
% points where VPs in sparser regions of the distributions relative to the data 
% are more highly weighted which could help to weight to where it is needed
% without overly focusing on a few VPs
if strcmp(dataGroupDescription(1:length('binTable')), 'binTable')
                w = 1;
 
elseif strcmp(dataGroupDescription(1:length('distTable')), 'distTable')
                w = 1;
                
elseif strcmp(dataGroupDescription(1:length('distTable2D')), 'distTable2D')
                w = 1;
 
elseif strcmp(dataGroupDescription(1:length('brTableRECIST')), 'brTableRECIST')
                w = 1;
 
elseif strcmp(dataGroupDescription(1:length('rTableRECIST')), 'rTableRECIST')
                w = 1;
 
elseif strcmp(dataGroupDescription(1:length('mnSDTable')), 'mnSDTable') && strcmp(dataGroupDescription(end-length('mean')+1:end), 'mean')
                w = 1;
 
elseif strcmp(dataGroupDescription(1:length('mnSDTable')), 'mnSDTable') && strcmp(dataGroupDescription(end-length('variance')+1:end), 'variance')
                w = 1;
elseif strcmp(dataGroupDescription(1:length('corTable')), 'corTable')
                w = 1;
else
                error('data group not supported');
end
 
end
```
