## Supplementary File 3 for "A surrogate workflow for virtual population development: designing an amenable proxy for a hypothesis-test inspired nonlinear goodness-of-fit": calculateExpWeightFixSingle.m

```
function w = calculateExpWeightFixSingle(expN, expSTD, dataGroupDescription, simN, keepType)
% This is a utility function to fix the experimental
% weight to 1 for a single data group type
% in linear calibrate.
%
% ARGUMENTS
%  expN:                  An experimental N, these are ignored here
%  expSTD:                An experimental standard deviation  
%  dataGroupDescription:  Data group description.  Should be one of:
%							binTable
%                           distTable
%                           brTableRECIST
%                           rTableRECIST
%                           mnSDTablemean
%                           mnSDTablevariance      
%                           corTable
%  simN:                  Assumed N for weighting for the simulation results
%                           keepType: data group description to keep
%                           nonzero.  Note: use mnSDTable rather than 
%                           mnSDTablemean, mnSDTablevariance
% RETURNS
%  w:                     A weighting factor.
%

if strcmp(dataGroupDescription(1:length('binTable')), 'binTable')
    if strcmp(keepType,'binTable')
        w = 1;
    else
        w = 0;
    end
 
elseif strcmp(dataGroupDescription(1:length('distTable')), 'distTable')
    if strcmp(keepType,'distTable')
        w = 1;
    else
        w = 0;
    end
    
elseif strcmp(dataGroupDescription(1:length('distTable2D')), 'distTable2D')
    if strcmp(keepType,'distTable')
        w = 1;
    else
        w = 0;
    end
 
elseif strcmp(dataGroupDescription(1:length('brTableRECIST')), 'brTableRECIST')
    if strcmp(keepType,'distTable')
        w = 1;
    else
        w = 0;
    end
 
elseif strcmp(dataGroupDescription(1:length('rTableRECIST')), 'rTableRECIST')
    if strcmp(keepType,'rTableRECIST')
        w = 1;
    else
        w = 0;
    end
 
elseif strcmp(dataGroupDescription(1:length('mnSDTable')), 'mnSDTable') && strcmp(dataGroupDescription(end-length('mean')+1:end), 'mean')
    if strcmp(keepType,'mnSDTable')
        w = 1;
    else
        w = 0;
    end

 
elseif strcmp(dataGroupDescription(1:length('mnSDTable')), 'mnSDTable') && strcmp(dataGroupDescription(end-length('variance')+1:end), 'variance')
    if strcmp(keepType,'mnSDTable')
        w = 1;
    else
        w = 0;
    end
elseif strcmp(dataGroupDescription(1:length('corTable')), 'corTable')
    if strcmp(keepType,'corTable')
        w = 1;
    else
        w = 0;
    end
else
                error('data group not supported');
end
 
end
```
