## Supplementary File 3 for "A surrogate workflow for virtual population development: designing an amenable proxy for a hypothesis-test inspired nonlinear goodness-of-fit": combineBins.m

```
function myVPop = combineBins(myVPop)
% Take an existing population and combine the axis bins, such that N bins
% becomes N/2 bins.  This is an auxiliary function that may be helpful
% if testing using different numbers of axis bins for the VPop.
% Note that you need to start with an even number of bins per axis
% for this function.
%
% ARGUMENTS
% myVPop:         An instance of a VPop object. Properties should be
%                 populated:
%                  binEdges
%                  indexTable
%                  mnSDTable
%                  binTable
%                  simData
%
% RETURNS
% myVpop:         VPop with updated properties:
%                  binEdges
%                  binMidPoints
%                  binProbs
%                  gofMn
%                  gofSD
%                  gofBin
%                  gof
%

continueFlag = false;
if nargin > 1
    warning([mfilename,' requires input argument: myVPop.  Too many arguments provided.']);
elseif nargin >0
    continueFlag = true;
else
    warning([mfilename,' requires input argument: myVPop.  Insufficient arguments provided.'])
end

if continueFlag
    if class(myVPop) ~= 'VPop'
        warning(['An argument of class VPop is required in ',mfilename,'. Exiting.'])
        continueFlag = false;
    end
end

if continueFlag
    [myNAxis, myNBins] = size(myVPop.binMidPoints);
    if mod(myNBins,2) > 0
        warning(['Unable to combine bins in ',mfilename,'. An even number of bins is required. Exiting'])
        continueFlag = false;
    end
end

if continueFlag
    for axisCounter = 1 : myNAxis
        for binCounter = 1 : (myNBins/2)
            downAdjustBin = 2 * binCounter;
            myIndices = find(myVPop.indexTable(axisCounter,:) == downAdjustBin);
            indexTable(axisCounter,myIndices) = binCounter;
        end
    end
    myVPop.binEdges = myVPop.binEdges(:,[1:2:length(myVPop.binEdges(axisCounter,:))]);
    newMidPoints = nan(myNAxis,myNBins/2);
    newBinProbs = nan(myNAxis,myNBins/2);
    for binCounter = 1 : (myNBins/2)
        newMidPoints(:,binCounter) = 0.5*myVPop.binEdges(:,binCounter)+0.5*myVPop.binEdges(:,binCounter+1);
        newBinProbs(:,binCounter) = myVPop.binProbs(:,2*(binCounter-1)+1)+myVPop.binProbs(:,2*(binCounter));
    end
    myVPop.binMidPoints = newMidPoints;
    myVPop.binProbs = newBinProbs;
    myVPop = evaluateGOF(myVPop);
end
```
