## Supplementary File 3 for "A surrogate workflow for virtual population development: designing an amenable proxy for a hypothesis-test inspired nonlinear goodness-of-fit": createRECISTSimFilter.m

```
function myRecistSimFilter = createRECISTSimFilter(myWorksheet, myVPopRECIST, poolObsTimes, startTime)     
% Scan the simulation results and create a weight
% filter based on which VPs would still be on therapy
% assuming we remove CR and PD patients from study
%
% ARGUMENTS
%  myWorksheet:     a worksheet
%  myVPopRECIST:    a VPopRECIST object.  The properties should
%                        be populated:
%                    brTableRECIST:  used to define the lesion evaluation times, dynamics
%                     in between these time points are ignored.  Required
%                     fields are:
%                    relSLDvar          
%                    absALDVar   
%                    crCutoff  
%  poolObsTimes:    Whether to pool observation times to create a
%                    RECISTSimFilter for other interventions.
%                    Default is true.
%  startTime:       start time in simulation that corresponds to day0 of Intervention
%                   default startTime = 2000;  first 2000 days are for initializationb in IO model and workflow

%
% RETURNS
%  myRecistSimFilter: an nIntervention x 1 cell array of structures
%                    each with fields: 
%                    'curResp':     current response in 0-3 class
%                                   where generally 0 = "CR", 1 = "PR",
%                                               2 = "SD", 3 = "PD"
%                                               nan = VP is off study
%                    'bestResp':    best response. according to each row indicator: BRSCOREVar, it is calculated as CBOR or BOR, SimFilter criteria based on RECIST1.1
%                                                   
%                    'curRespNotCorrected':   RECIST classifications except without correction
%                                             (relative to baseline and
%                                             ignoring the clinical observation times.
%                                             This is in contrast to �curResp� and �bestResp�, which
%                                             are only updated at the clinical observation times, and
%                                             which are relative to the previous �bestResp�). 
%                    'curWeight:    whether simulated patient stays in
%                                   population for calibration at current time
%                                   point.  Achieving 0
%                                   or 3 results in removal.
%                    'time':        time vector
%

% Perform initial checks on the provided arguments
flagContinue = true;
myResponseSummaryTable = resultTable({});
binEdge = [-0.3;0.2];
if nargin > 4
    warning([mfilename,' requires input arguments: myWorksheet, myVPopRECIST, optionally poolObsTimes, startTime.  Too many arguments provided.'])
    flagContinue = false;    
elseif nargin < 2
    warning([mfilename,' requires input arguments: myWorksheet, myVPopRECIST, optionally poolObsTimes, startTime.  Too few arguments provided.'])
    flagContinue = false;
elseif nargin == 4
    flagContinue = true;
elseif nargin == 3
    flagContinue = true;
    startTime = 2000;
elseif nargin < 3
    flagContinue = true;
    poolObsTimes = true;
    startTime = 2000;
end

if flagContinue
    myBRTableRECIST = myVPopRECIST.brTableRECIST;
    myRTableRECIST = myVPopRECIST.rTableRECIST;    
    myRelSLDvar = myVPopRECIST.relSLDvar;  
    myAbsALDVar = myVPopRECIST.absALDVar; 
    crCutoff = myVPopRECIST.crCutoff;     
    myVPIDs = getVPIDs(myWorksheet);
    myInterventionIDs = getInterventionIDs(myWorksheet);
    allInterventionIDs = myInterventionIDs;
    [nInterventionResults, nVPresults] = size(myWorksheet.results);
    if((length(myVPIDs) ~= nVPresults)|(length(myInterventionIDs) ~= nInterventionResults))
        warning([mfilename,' requires fully populated results, exiting.'])
        flagContinue = false;        
    end
    if flagContinue
        if (sum(ismember(myWorksheet.results{1,1}.Names,myRelSLDvar)) < 1)
            flagContinue = false;  
        end
    end
end

myRecistSimFilter = cell(length(myInterventionIDs),1);
for interventionCounter = 1 :length(myInterventionIDs)
    curRecistFilter.interventionID = myInterventionIDs{interventionCounter};
    myRecistSimFilter{interventionCounter} = curRecistFilter;
end

if flagContinue  
    % Check the interventions that already have RECIST classification times
    % in the data
    if ~isempty(myBRTableRECIST)
        interventionsRECISTClass = unique(myBRTableRECIST{:,'interventionID'});
        if sum(~ismember(myInterventionIDs, interventionsRECISTClass)) > 0
            if poolObsTimes
                disp([mfilename,' found not all interventions to have RECIST classification times in myVPopRECIST.brTableRECIST for scoring.  Creating a simFilter for mechanistic dropouts using classification times for interventions where available, and pooling all observation times from other interventions where not.'])
            else
                disp([mfilename,' found not all interventions to have RECIST classification times in myVPopRECIST.brTableRECIST for scoring.  Creating a simFilter for mechanistic dropouts using classification times for interventions where available, and assuming no dropouts from other interventions where not.'])
            end
        end
        if sum(~ismember(interventionsRECISTClass,myInterventionIDs)) > 0
            disp([mfilename,' found not all RECIST classification interventions are present.  Proceeding, but be careful about this mismatch as there are interventions in the VPop that are not in the worksheet!'])
            flagContinue = true;
        end    
    else
        disp([mfilename,' found no interventions to have RECIST classification times in myVPopRECIST.brTableRECIST for scoring.  Creating a simFilter for mechanistic dropouts assuming no dropouts from interventions.'])
        poolObsTimes = false;
    end
end

if flagContinue
    binEdge = sort(binEdge,'ascend');
    for interventionIndex = 1 : length(myInterventionIDs)
        myInterventionID = myInterventionIDs{interventionIndex};
        interventionCounter = (find(ismember(allInterventionIDs,myInterventionID)));
        sampleData = myWorksheet.results{interventionCounter,1};

        [nTimePoints, nVars] = size(sampleData.Data);
        curData = nan(nTimePoints,nVPresults + 1);
        curSize = nan(nTimePoints,nVPresults + 1);
        varIndices = find(ismember(sampleData.Names,myRelSLDvar));
        varSizeIndices = find(ismember(sampleData.Names,myAbsALDVar));
        timeVector = sampleData.Data(:,1);
        curData(:,1) = timeVector;
        curSize(:,1) = timeVector;
        for vpCounter = 1 : length(myVPIDs)
            sampleData = myWorksheet.results{interventionCounter,vpCounter}.Data;
            % Get all of the relative SLD data for the VP
            curData(:,vpCounter + 1) = sampleData(:,varIndices);
            % Get all of the absolute lesion size data for the VP
            % (used for CR classification)
            curSize(:,vpCounter + 1) = sampleData(:,varSizeIndices);
        end
        curDataRecalc = curData;
        curSizeRecalc = curSize;

        % Get the observation times for the current intervention
        if ~isempty(myBRTableRECIST)
            obsTimes = find(ismember(myBRTableRECIST{:,'interventionID'},myInterventionID));
            if length(obsTimes) > 0
                obsTimes = unique(myBRTableRECIST{obsTimes,'time'});
                obsTimes = sort(obsTimes,'ascend');
            else
                % If there are no observed times for the intervention in the
                % data, for now we will simply take the pooled observation
                % times across studies where we do have measures.
                if poolObsTimes
                    obsTimes = unique(myBRTableRECIST{:,'time'});
                    obsTimes = sort(obsTimes,'ascend');
                else
                    obsTimes = max(timeVector);
                end
            end
        else
            obsTimes = max(timeVector);
        end

        % Recalculate values only at the observed times
        % Discretized observations so they are only updated at clinical
        % lesion scan times.
        obsTimes = [obsTimes;nan];
        for vpCounter = 1 : length(myVPIDs)
            % How many observation time points have passed
            pointsFound = 0;
            for timeCounter = 1 : nTimePoints
                % Prior to first observation, assign nan
                if timeVector(timeCounter) < obsTimes(1)
                    curDataRecalc(timeCounter,vpCounter + 1) = nan;
                    curSizeRecalc(timeCounter,vpCounter + 1) = nan;
                elseif timeVector(timeCounter) >= obsTimes(1+pointsFound)
                    pointsFound = pointsFound+1;
                    lastIndex = timeCounter;
                    curDataRecalc(timeCounter,vpCounter + 1) = curData(timeCounter,vpCounter + 1);
                    curSizeRecalc(timeCounter,vpCounter + 1) = curSize(timeCounter,vpCounter + 1);
                else
                    curDataRecalc(timeCounter,vpCounter + 1) = curData(lastIndex,vpCounter + 1);
                    curSizeRecalc(timeCounter,vpCounter + 1) = curSize(lastIndex,vpCounter + 1);
                end
            end
        end
        % Nominal PD VPs
        bin3 = (binEdge(2)<curDataRecalc(:,2:end)) & (curSizeRecalc(:,2:end)>=crCutoff);
        % Nominal SD VPs
        bin2 = (binEdge(1)<curDataRecalc(:,2:end)) & (curSizeRecalc(:,2:end)>=crCutoff);
        % Nominal PR VPs
        bin1 = (curDataRecalc(:,2:end)<=binEdge(1)) & (curSizeRecalc(:,2:end)>=crCutoff);
        curResp =  3 * (bin3 > 0) + 2 * (bin2 > 0) .* (bin3 < 1) + 1 * (bin1 > 0) .* (bin3 < 1) .* (bin2 < 1);
        
        % Also calculate the current response from baseline, ignoring
        % observation times:
        % Nominal PD VPs
        bin3Continuous = (binEdge(2)<curData(:,2:end)) & (curSize(:,2:end)>=crCutoff);
        % Nominal SD VPs
        bin2Continuous = (binEdge(1)<curData(:,2:end)) & (curSize(:,2:end)>=crCutoff);
        % Nominal PR VPs
        bin1Continuous = (curData(:,2:end)<=binEdge(1)) & (curSize(:,2:end)>=crCutoff);
        curRespNotCorrected =  3 * (bin3Continuous > 0) + 2 * (bin2Continuous > 0) .* (bin3Continuous < 1) + 1 * (bin1Continuous > 0) .* (bin3Continuous < 1) .* (bin2Continuous < 1);
        
        % Need to add cumulative change from min, too
        % increase of 20%, binEdge(2), relative to nadir
        % should trigger PD classification
        cumMin = cummin(curDataRecalc(:,2:end),1);
        cumMin = (curDataRecalc(:,2:end) - min(cumMin,0))./(1+min(cumMin,0));
        % We use cummin to trigger PD
        bin3 = (binEdge(2)<cumMin);
        curResp =  curResp .* (bin3 < 1) + ( (bin3 > 0)) .* (3*(bin3 > 0));
        
        curResp(find(isnan(curDataRecalc(:,2:end)))) = nan;
        curCResp = curResp(:,vpCounter); % Lu add 20220808. initialization % 20220825 move to here. after 
        curBestResp = curResp(:,vpCounter); % Lu add 20220808. initialization
        TConfirmation = 28; % based on RECIST1.1 criteria (4 weeks)
        minSDInterval = 42; % based on RECIST1.1 criteria (6 weeks)
        
        % Only keep counting for VPs until they are PD/CR
        % Note we don't consider what happens with tox here.
        % Note we want to count them up to and INCLUDING the first time they are
        % PD/CR and then to drop them for the NEXT time point.
        curWeight = ~(curResp<=0) & ~(curResp>=3);
         for vpCounter = 1 : length(myVPIDs)
            myIndices = find(curWeight(:,vpCounter) < 1);
            if length(myIndices) > 0
                if myIndices(1) < nTimePoints
                    curWeight(myIndices(1),vpCounter) = 1;
                    curWeight((myIndices(1)+1):end,vpCounter) = 0;
                    curResp((myIndices(1)+1):end,vpCounter) = curResp(myIndices(1),vpCounter); % response from last valid timepoint carried over for next few dropout timepoints
                end
            end
            curCResp(:,vpCounter) = curResp(:,vpCounter); % Lu add 20220808. initialization % 20220825 move to here. after 
            curBestResp(:,vpCounter) = curResp(:,vpCounter); % Lu add 20220808. initialization
               %            based on in brTable the row indicator for this intervention: expVarID, elementID, BRSCOREVar             
            interventionIndices = find(ismember(myBRTableRECIST{:,'interventionID'},myInterventionID));
            BRSCOREVar = unique(myBRTableRECIST{interventionIndices,'BRSCOREVar'});
            if length(BRSCOREVar) > 1
                error(['Intervention ' myInterventionID ' contains mixed BOR criteria ... please double check!']);  
                % TODO: we might need to add functionalities to address situations that a same intervention but two different subgroups, using different criteria. 
                % then will need to also make sure the SimFilter is applied correctly for the subpop+intervention for all other table calculations ...
            elseif length(BRSCOREVar) == 1
                BORflag = BRSCOREVar;
            elseif isempty(BRSCOREVar)
%                 disp(['Intervention ' myInterventionID ' is not in the brTable, proceed with default BRSCORE.']);
                BORflag = 'BRSCORE';
            end
            
            if strcmp(BORflag,'BRSCORE')
                curBestResp(:,vpCounter) = cummin(curResp(:,vpCounter));
            elseif strcmp(BORflag,'CBRSCORE')         
                % implement confirmed bor
                dat_patient = curResp(:,vpCounter);
                obsindices = find(ismember(timeVector,obsTimes));
    %           for vpCounter = 1 : length(myVPIDs)
                % How many observation time points have passed
                pointsFound = 0;
                for timeCounter = 1 : nTimePoints
                    % Prior to first observation, assign nan
                    if timeVector(timeCounter) < obsTimes(1)
                        curCResp(timeCounter,vpCounter) = nan;
                    elseif timeVector(timeCounter) >= obsTimes(1+pointsFound)
                        pointsFound = pointsFound+1;
                        lastIndex = timeCounter;
                        t1 = obsindices(pointsFound); % timeCounter
                        pointsRemain = length(obsindices)-pointsFound;
                             if dat_patient(t1) == 0
                                 if pointsRemain > 0 && obsindices(pointsFound+1) <= nTimePoints
                                     t2 = obsindices(pointsFound+1);
                                      if dat_patient(t2)==1
                                          curCResp(t1,vpCounter) = 2 * ((timeVector(t1)-startTime) >= minSDInterval) + 3 *((timeVector(t1)-startTime) < minSDInterval);
                                      elseif dat_patient(t2)>1
                                          curCResp(t1,vpCounter) = 2 * ((timeVector(t1)-startTime) >= minSDInterval) + 3 *((timeVector(t1)-startTime) < minSDInterval);
                                      elseif dat_patient(t2)==0
                                          if (timeVector(t2)-timeVector(t1)) >= TConfirmation
                                                curCResp(t1,vpCounter) = 0;
                                          elseif (pointsRemain-1) > 0 && obsindices(pointsFound+2) <= nTimePoints
                                                  t3 = obsindices(pointsFound+2);
                                                  if dat_patient(t3)==1
                                                      curCResp(t1,vpCounter) = 2 * ((timeVector(t1)-startTime) >= minSDInterval) + 3 *((timeVector(t1)-startTime) < minSDInterval);
                                                  elseif dat_patient(t3)>1
                                                      curCResp(t1,vpCounter) = 2 * ((timeVector(t1)-startTime) >= minSDInterval) + 3 *((timeVector(t1)-startTime) < minSDInterval);
                                                  elseif dat_patient(t3)==0
                                                      if (timeVector(t3)-timeVector(t1)) >= TConfirmation
                                                            curCResp(t1,vpCounter) = 0;
                                                      elseif (timeVector(t1)-startTime) >= minSDInterval  % simpify ... up to 2 subsequent time point scan checked
                                                          curCResp(t1,vpCounter) = 2;
                                                      else
                                                          curCResp(t1,vpCounter) = NaN;
                                                      end
                                                  end
                                          elseif (timeVector(t1)-startTime) >= minSDInterval
                                                  curCResp(t1,vpCounter) = 2;
                                          else
                                                  curCResp(t1,vpCounter) = NaN;
                                          end
                                      end
                                 elseif (timeVector(t1)-startTime) >= minSDInterval
                                      curCResp(t1,vpCounter) = 2;
                                 else
                                      curCResp(t1,vpCounter) = NaN;
                                 end
                             elseif dat_patient(t1) == 1
                                 if pointsRemain > 0 && obsindices(pointsFound+1) <= nTimePoints
                                     t2 = obsindices(pointsFound+1);
                                      if dat_patient(t2)==2
                                          curCResp(t1,vpCounter) = 2;
                                      elseif dat_patient(t2) > 2
                                          curCResp(t1,vpCounter) = 2 * ((timeVector(t1)-startTime) >= minSDInterval) + 3 *((timeVector(t1)-startTime) < minSDInterval);
                                      elseif dat_patient(t2) <= 1
                                          if (timeVector(t2)-timeVector(t1)) >= TConfirmation
                                                curCResp(t1,vpCounter) = 1;
                                          elseif (pointsRemain-1) > 0 && obsindices(pointsFound+2) <= nTimePoints
                                                  t3 = obsindices(pointsFound+2);
                                                  if dat_patient(t3)==2
                                                      curCResp(t1,vpCounter) = 2;
                                                  elseif dat_patient(t3) > 2
                                                      curCResp(t1,vpCounter) = 2 * ((timeVector(t1)-startTime) >= minSDInterval) + 3 *((timeVector(t1)-startTime) < minSDInterval);
                                                  elseif dat_patient(t3) <= 1
                                                      if (timeVector(t3)-timeVector(t1)) >= TConfirmation
                                                          curCResp(t1,vpCounter) = 1;
                                                      elseif (timeVector(t1)-startTime) >= minSDInterval  
                                                          curCResp(t1,vpCounter) = 2;
                                                      else
                                                          curCResp(t1,vpCounter) = NaN;
                                                      end
                                                  end

                                          elseif (timeVector(t1)-startTime) >= minSDInterval
                                                curCResp(t1,vpCounter) = 2;
                                          else
                                                curCResp(t1,vpCounter) = NaN;
                                          end
                                      end
                                 elseif (timeVector(t1)-startTime) >= minSDInterval
                                      curCResp(t1,vpCounter) = 2; 
                                 else
                                      curCResp(t1,vpCounter) = NaN;
                                 end
                             elseif dat_patient(t1) == 2
                                 if (timeVector(t1)-startTime) >= minSDInterval
                                     curCResp(t1,vpCounter) = 2;
                                 else
                                     curCResp(t1,vpCounter) = NaN;
                                 end
                             end 
                        curCResp(timeCounter,vpCounter) = curCResp(t1,vpCounter);
                    else
                        curCResp(timeCounter,vpCounter) = curCResp(lastIndex,vpCounter);
                    end
                end
                curBestResp(:,vpCounter) = cummin(curCResp(:,vpCounter));
             end
          end

        myRecistSimFilter{interventionCounter}.filterMatrix = curWeight;
        % To save memory, we will convert to int8.  First, any nans should 
        % be set to -1.
        curResp(isnan(curResp)) = -1;
        curRespNotCorrected(isnan(curRespNotCorrected)) = -1;
        curBestResp(isnan(curBestResp))= -1;
        myRecistSimFilter{interventionCounter}.curResp = int8(curResp);
        myRecistSimFilter{interventionCounter}.curRespNotCorrected = int8(curRespNotCorrected);
        myRecistSimFilter{interventionCounter}.bestResp = int8(curBestResp);
        myRecistSimFilter{interventionCounter}.time = curData(:,1);
 
	end
end
```
