## Supplementary File 3 for "A surrogate workflow for virtual population development: designing an amenable proxy for a hypothesis-test inspired nonlinear goodness-of-fit": evaluateObjective.m

```
function myObjective = evaluateObjective(myVPop, myProbVectTrans)
% This function evaluates an objective function, which is similar to
% the composite goodness-of-fit but with a few critical differences, and is
% generally called only during the optimization / VPop fit to data.
% This function should be called just for VPops with 'bin' pwStrategy
%
% ARGUMENTS
%  myVPop:              a VPop object instance, with properties already
%                       assigned:
%                        binProbs:  Should at least be initialized
%                        spreadOut
%                        minEffN
%                        mnSDTable: should at least be populated with
%                                   experiment data
%                        binTable:  Also need experiment data
%                        distTable
%                        distTable2D
%                        corTable
%                        brTableRECIST (if applicable)
%                        rTableRECIST (if applicable)
%  myProbVectTrans:     a 1 x (nBins-1 * nAxis) vector of transformed 
%                       probabilities
%
% RETURNS
%  myObjective:         the objective function values
%

% First update pws with new probVect
[nAxis, nBins] = size(myVPop.binProbs);
myProbTrans = transpose(reshape(myProbVectTrans, nBins-1, nAxis));
binProbs = nan(nAxis, nBins);
for axisCounter = 1 : nAxis
    binProbs(axisCounter,:) = invHyperTransform(myProbTrans(axisCounter,:));
end

myVPop.binProbs = binProbs;
myVPop = myVPop.assignPWs();

% Next update the individual GOF statistics
myVPop = evaluateGOF(myVPop);

myObjective = 0;
effN = 1/(sum((myVPop.pws).^2));

if myVPop.spreadOut > 0
	[nAxis, nPWs] = size(myVPop.coeffsTable);
	% myVPop.coeffsTable is nVP x nAxis
    % Store this value to speed
	% myD = pdist2(myVPop.coeffsTable',myVPop.coeffsTable');
    myD = myVPop.coeffsDist;
	% We can take the weighted sum along rows or columns
	% to get the weighted distances, renormalize each 
	% sum by (1-pwi).
	% Then we sum each weighted sum by pwi to get the weighted
	% average distance
    % Vectorized this following calculation.  The new version
    % is still slow, but a little better
    % 	myDSum = nan(1, nPWs);   
    % 	for vpCounter = 1: nPWs
    % 		myDSum(vpCounter) = sum(myD(vpCounter,:).*(myVPop.pws/sum(myVPop.pws([1:vpCounter-1,vpCounter+1:nPWs]))));
    % 	end
    % 	myDAvg = sum(myDSum.*myVPop.pws);
    repPWs = repmat(myVPop.pws,nPWs,1);	
    ieye = ~eye(nPWs);
	myDSum = sum(myD.*repPWs./repmat(sum(repPWs.*ieye,2),1,nPWs),2);
    myDAvg = sum(myDSum'.*myVPop.pws);	
    
	% We re-normalize the effective distance
	% by roughly what we would expect for nVP evenly spaced
	% partices in
	% a nAxis space
	dIdeal = (1/nPWs)^(1/nAxis);
    notUnif = myVPop.spreadOut*dIdeal/myDAvg;
    myObjective = myObjective + notUnif;
end

if myVPop.minEffN > 0
    % Add in a linear penalty term and a small discontinuity so 
    % numerically the effN should be at least a little larger in the
    % optimization.
    myObjective = myObjective + (effN < myVPop.minEffN) * 1E6 * (myVPop.minEffN - effN) + (effN <= myVPop.minEffN);
end

% Renormalize the weights in the objective function so they sum to 1.
% This will make it more straightforward to compare how well the
% optimization is performing for the same model even as we add more
% contraints.
myVPop = normalizeWeights(myVPop, false);

myMnSDTable = myVPop.mnSDTable;
myBinTable = myVPop.binTable;
myDistTable = myVPop.distTable;
myDistTable2D = myVPop.distTable2D;
myCorTable = myVPop.corTable;
if isa(myVPop, 'VPopRECIST')
	myBRTable = myVPop.brTableRECIST;
	myRTable = myVPop.rTableRECIST;
else
	myBRTable = [];
	myRTable = [];
end	

% We use the same epsilon from the MAPEL paper
%epsilon = myVPop.epsilon;
epsilon = 1E-16;
gofMean = myVPop.gofMn;
gofSD = myVPop.gofSD;
gofBin = myVPop.gofBin;
gofDist = myVPop.gofDist;
gofDist2D = myVPop.gofDist2D;
gofCor = myVPop.gofCor;
if isa(myVPop, 'VPopRECIST')
	gofBR = myVPop.gofBR;
	gofR = myVPop.gofR;
end	
minIndPVal = myVPop.minIndPVal;


% May want to add checks here for the Mean/SD/Bin evaluations and whether
% there are entries before adding the respective terms
if ~isempty(myMnSDTable)
    myObjective = myObjective - 2*(sum(myMnSDTable{:,'weightMean'} .* log10(gofMean+epsilon)) + sum(myMnSDTable{:,'weightSD'} .* log10(gofSD+epsilon)));
	if minIndPVal > 0
		myObjective = myObjective + (sum((myMnSDTable{:,'weightMean'} > 0) .* (gofMean < minIndPVal) .* 1E6 .* (minIndPVal - gofMean)) + sum((myMnSDTable{:,'weightSD'} > 0) .* (gofSD < minIndPVal) .* 1E6 .* (minIndPVal - gofSD)));
	end	
end
if ~isempty(myBinTable)
    myObjective = myObjective - 2*(sum(myBinTable{:,'weight'} .* log10(gofBin+epsilon)));
	if minIndPVal > 0
		myObjective = myObjective + sum((myBinTable{:,'weight'} > 0) .* (gofBin < minIndPVal) .* 1E6 .* (minIndPVal - gofBin));
	end		
end
if ~isempty(myDistTable)
    myObjective = myObjective - 2*(sum(myDistTable{:,'weight'} .* log10(gofDist+epsilon)));
	if minIndPVal > 0
		myObjective = myObjective + sum((myDistTable{:,'weight'} > 0) .* (gofDist < minIndPVal) .* 1E6 .* (minIndPVal - gofDist));
	end			
end
if ~isempty(myDistTable2D)
    myObjective = myObjective - 2*(sum(myDistTable2D{:,'weight'} .* log10(gofDist2D+epsilon)));
	if minIndPVal > 0
		myObjective = myObjective + sum((myDistTable2D{:,'weight'} > 0) .* (gofDist2D < minIndPVal) .* 1E6 .* (minIndPVal - gofDist2D));
	end			
end
if ~isempty(myCorTable)
    myObjective = myObjective - 2*(sum(myCorTable{:,'weight'} .* log10(gofCor+epsilon)));
	if minIndPVal > 0
		myObjective = myObjective + sum((myCorTable{:,'weight'} > 0) .* (gofCor < minIndPVal) .* 1E6 .* (minIndPVal - gofCor));
	end			
end
if ~isempty(myBRTable)
    myObjective = myObjective - 2*(sum(myBRTable{:,'weight'} .* log10(gofBR+epsilon)));
	if minIndPVal > 0
		myObjective = myObjective + sum((myBRTable{:,'weight'} > 0) .* (gofBR < minIndPVal) .* 1E6 .* (minIndPVal - gofBR));
	end		
end
if ~isempty(myRTable)
    myObjective = myObjective - 2*(sum(myRTable{:,'weight'} .* log10(gofR+epsilon)));
	if minIndPVal > 0
		myObjective = myObjective + sum((myRTable{:,'weight'} > 0) .* (gofR < minIndPVal) .* 1E6 .* (minIndPVal - gofR));
	end		
end
end
```
