## Supplementary File 3 for "A surrogate workflow for virtual population development: designing an amenable proxy for a hypothesis-test inspired nonlinear goodness-of-fit": expandWorksheetVPsFromVPop.m

```
function [myWorksheet, newPassNames] = expandWorksheetVPsFromVPop(myWorksheet,newVPop, myMapelOptions,suffix,wsIterCounter, maxNewPerIter, myScreenTable, expandCohortSize, varyMethod, gaussianStd, maxNewPerOld, unweightedParents, selectByParent, useScoresForNew, pwExpandCutoff, myScreenFunctionName, linearExpandFlag)
% This function expands a worksheet given a VPop.  It selected out VPs to expand around,
% samples for new VPs, scores the "children" based on available data, and adds
% the best children to the worksheet.
%
%  myWorksheet
%  newVPop
%  myMapelOptions
%  suffix:               a text descriptor string that will be included in what 
%                         will be written to file.  This is also used
%                         in setting VP identities.
%  wsIterCounter:        tracks the iterations through the algorithm.  Keep
%                         incrementing to avoid issues with repeated VPIDs.
%  maxNewPerIter:        maximum new VPs we can add per iteration.  Set to 
%                         -1 to use the VPop effN
%  myScreenTable:        a screen table to idenify VPs to keep
%  expandCohortSize:     size of the cohort to generate for testing
%  varyMethod:           method for resampling.  i.e. 'gaussian' or 'localPCA'
%  gaussianStd:        ` standard deviation for the re-sampled parameters.
%                         note this is applied across all axes in the transformed
%                         units (i.e. within bounds normalized 0-1).
%  maxNewPerOld:         maximum number of children per weighted parent
%  unweightedParents:    Whether to include unweighted VPs as seeds for 
%                         expansion if they look like they are useful.
%  selectByParent:       a boolean variable indicating whether children
%                         VPs are selected for inclusion each iteraction 
%                         based on their parent, or just pooled together
%  pwExpandCutoff:       a cutoff on PW for whether to use a highly weighted
%                         VP as a seed
%  myScreenFunctionName: a string indicating a function to use for screening
%                         VPs before simulation.  It should take two input
%                         arguments: a worksheet and a list of VPIDs to
%                         screen.  It should return a worksheet with
%                         identical number of names of VPs, possibly 
%                         modified after bing checked against some
%                         criteria. '' indicates no screening.
%  linearExpandFlag:     If linearExpandFlag is true, we sample children around all VPs proportional to their PWs, and add all past-screen VPs; 
%                        if linearExpandFlag is false, we use the previous way: select high PW VPs as parent VPs, sample equally maxNewPerOld VPs to add
%                      
%                      
% RETURNS:
%  myWorksheet
%  newPassNames
%
% TODO: this function could use more input proofing or an options object.  This
%       is also mainly intended to be called from other functions, so it hasn't
%       been done yet.

originalVPIDs = getVPIDs(myWorksheet); 
if length(originalVPIDs) > length(newVPop.pws)
    warning(['More VPs in worksheet than PWs in VPop in ',mfilename,'.  Proceeding assuming the PWs correspond to the first VPs in the worksheet.'])
    continueFlag = true;
elseif length(originalVPIDs) < length(newVPop.pws)
    warning(['Fewer VPs in worksheet than PWs in VPop in ',mfilename,'.  Exiting...'])
    continueFlag = false; 
else
    continueFlag = true;
end

    
if continueFlag   
    myCoeffs = getVPCoeffs(myWorksheet);
    [nAxis, nOriginalVPs] = size(myCoeffs);	
    
    myVPRangeTable = createVPRangeTable(newVPop);
    % Consider an entry if it is off of one of the edges by more than 10%
    myCutoff = 0.1;
    % To avoid issues with outliers, we also consider the range OK if we at
    % least have covered from the 10th percentile up to the 90th
    % percentile.

    disp('---------')
    disp(['Check of VP filling ranges in ',mfilename,'.'])
    myIndicesLow = find((myVPRangeTable{:,'minRangeMissing'}>myCutoff) & (myVPRangeTable{:,'min10PercentileMissing'}>0));
    myIndicesHigh = find((myVPRangeTable{:,'maxRangeMissing'}>myCutoff) & (myVPRangeTable{:,'max90PercentileMissing'}>0));
    myIndices = unique(sort([myIndicesLow;myIndicesHigh],'ascend'),'stable');
    % Print portions of the table to screen so we can track progress.
    myVPRangeTable(myIndices,{'elementID','interventionID','time','expN','rangeCover','minRangeMissing','maxRangeMissing','min10PercentileMissing','max90PercentileMissing'})

    % We will pre-rank parent VPs we might want to add
    % to resampling based on the observed data    
    % This step will only be impactful if
    % unweightedParents is not false
    if unweightedParents
        myVPRangeTable = myVPRangeTable(myIndices,:);
        [nRows,nCols] = size(myVPRangeTable);
        myIndicesLow = find((myVPRangeTable{:,'minRangeMissing'}>myCutoff) & (myVPRangeTable{:,'min10PercentileMissing'}>0));
        myIndicesHigh = find((myVPRangeTable{:,'maxRangeMissing'}>myCutoff) & (myVPRangeTable{:,'max90PercentileMissing'}>0));     
        myValuesLow = myVPRangeTable{myIndicesLow,'minRangeMissing'};
        myValuesHigh = myVPRangeTable{myIndicesHigh,'maxRangeMissing'};
        nLowValues = length(myValuesLow);
        nHighValues = length(myValuesHigh);
        lowTracker = 1*ones(nLowValues,1);
        highTracker = 2*ones(nHighValues,1);         
        myValuesCombine = [myValuesLow;myValuesHigh];
        myIndicesCombine = [myIndicesLow;myIndicesHigh];
        [myValuesCombineSort, myValuesCombineIndices] = sort(myValuesCombine,'descend');
        combineTracker = [lowTracker;highTracker];
        combineTracker = combineTracker(myValuesCombineIndices);
        myIndicesCombine = myIndicesCombine(myValuesCombineIndices);
        % We will manually sort through the VPs
        parentVPs = cell(1,nLowValues + nHighValues);
        if (nLowValues + nHighValues)>0
            for parentCounter = 1 : (nLowValues + nHighValues)
                if combineTracker(parentCounter) > 1.5
                    myIndex = myIndicesCombine(parentCounter);
                    parentVPs(parentCounter) = myVPRangeTable{myIndex,'vpIDsMax'};
                else
                    myIndex = myIndicesCombine(parentCounter);
                    parentVPs(parentCounter) = myVPRangeTable{myIndex,'vpIDsMin'};
                end
            end

            parentVPs(isempty(parentVPs)) = [];
        end
        edgeVPIDs = getMinimalEdgeSet(parentVPs);
    else
        edgeVPIDs = {};
    end
    % Get the indices for the edgeVPs in the original VPIDs.  Order
    % should be preserved.
    edgeVPIndices = nan(1, length(edgeVPIDs));
    for vpCounter = 1 : length(edgeVPIDs)
        edgeVPIndices(vpCounter) = find(ismember(originalVPIDs,edgeVPIDs{vpCounter}));
    end  

    allResponseTypeIDs = getResponseTypeIDs(myWorksheet);
    nResponseTypes = length(allResponseTypeIDs);
    myPWs = newVPop.pws;
    curEffN = round(1/sum(myPWs.^2));
    highVPIndices1 = find(myPWs>=pwExpandCutoff);
    [~, highVPIndices2] = sort(myPWs, 'descend');
    highVPIndices2 = highVPIndices2(1 : curEffN);    
    highVPindicesCat = [highVPIndices2,highVPIndices1];
    [highVPIndices,i,j] = unique(highVPindicesCat, 'first'); 
    highVPIndices = highVPindicesCat(sort(i));
    highVPIDs = originalVPIDs(highVPIndices);

    if linearExpandFlag % During linearExpansion phase, we select all VP as parent VPs and sample chidren number proportional to their weight. no need to go through the scoring function, just add all VPs that pass screen
        keepLowPwEdgeIndices = find(myPWs(edgeVPIndices)<0.01); 
        edgeVPIDs = originalVPIDs(edgeVPIndices(keepLowPwEdgeIndices));

        if ~isempty(newVPop.LinearProblemMatrices)
            vpIsInSubgroup=newVPop.LinearProblemMatrices.vpIsInSubgroup;
            [uniqueSubpop,ia,ic]=unique(vpIsInSubgroup,'rows');
            if (~isempty(newVPop.LinearProblemMatricesSubgroupSumWeights))
                lowsubpop = find(sum(uniqueSubpop,2)'<= 5 | newVPop.LinearProblemMatricesSubgroupSumWeights(ia)'<=1e-16); % max(0.001*length(newVPop.pws),2)) or subpopulation weights=0;
            else
                lowsubpop = find(sum(uniqueSubpop,2)'<= 5); 
            end
            %% need to allocate and sample inside each of these subpop, because they might all have very low weights
            subpopvpDat = sum(vpIsInSubgroup(ia(lowsubpop),:),1);
            subpopvpIndices = unique(find(subpopvpDat>0)); % these are the parents
            subpopVPIDs = originalVPIDs(subpopvpIndices);
            subpopVPIDs = setdiff(subpopVPIDs,edgeVPIDs);
            subpopvpIndices = nan(1, length(subpopVPIDs));
            for vpCounter = 1 : length(subpopVPIDs)
                subpopvpIndices(vpCounter) = find(ismember(originalVPIDs,subpopVPIDs{vpCounter}));
            end 
        else
            subpopVPIDs = {};
            subpopvpIndices = [];
        end
        
        % sample children VPs proportionally to the parentVP weight
        highVPIDsMono = setdiff(highVPIDs,union(edgeVPIDs,subpopVPIDs)); % can also consider taking out subpop VPs
        AddParentVPIDs = highVPIDsMono; % union(highVPIDs,edgeVPIDs);
        allVPIDs = getVPIDs(myWorksheet);
        AddParentVPIndices = [];
        for i=1:length(AddParentVPIDs)
            ind = find(ismember(allVPIDs,AddParentVPIDs{i}));
            AddParentVPIndices = [AddParentVPIndices, ind];
        end

        AddParentVPpws = myPWs(AddParentVPIndices)./sum(myPWs(AddParentVPIndices));
        if length(edgeVPIDs)>0
            expandedgeVPCohortSize = max(length(edgeVPIDs)*maxNewPerOld,expandCohortSize*0.1);  
        else
            expandedgeVPCohortSize = 0;
        end
         % allocate 10% to sample around low subpopulation VPs
        % since they are all low weight, maybe just sample equally here
        if length(subpopVPIDs)>0
            expandlowsubpopVPCohortSize = max(length(subpopVPIDs)*maxNewPerOld,expandCohortSize*0.1);  
        else
            expandlowsubpopVPCohortSize = 0;
        end
         %
        expandhighVPCohortSize = expandCohortSize-expandedgeVPCohortSize-expandlowsubpopVPCohortSize;
        CountUniqueParentVPs = round(AddParentVPpws.*expandhighVPCohortSize);

        myVaryAxesOptions = varyAxesOptions;
        myVaryAxesOptions.varyMethod = varyMethod;
        myVaryAxesOptions.gaussianStd = gaussianStd;
        myVaryAxesOptions.varyAxisIDs = getAxisDefIDs(myWorksheet);
        myVaryAxesOptions.intSeed = newVPop.intSeed;

        myVaryAxesOptions.additionalIDString = [suffix,num2str(wsIterCounter)];

        disp('---')  
        disp(['Adding ',num2str(length(AddParentVPIDs)),' VPs as expansion seeds from prevalence weight considerations in ',mfilename,'.'])
        disp(['Adding ',num2str(length(edgeVPIDs)),' VPs as expansion seeds from phenotype range considerations in ',mfilename,'.'])
        disp(['Adding ',num2str(length(subpopVPIDs)),' VPs as expansion seeds from low subpopulation considerations in ',mfilename,'.'])
        
        % sample children VPs proportionally to the parentVP weight
        jitteredWorksheet = myWorksheet;
        for addCounter = 1: length(AddParentVPIDs)
            if CountUniqueParentVPs(addCounter)>0
                myVaryAxesOptions.newPerOld = CountUniqueParentVPs(addCounter);
               % CountUniqueParentVPs(addCounter)
                myVaryAxesOptions.baseVPIDs = AddParentVPIDs(addCounter);
               % AddParentVPIDs(addCounter)
                jitteredWorksheet = addVariedVPs(jitteredWorksheet, myVaryAxesOptions);
            end
        end

        % edgeVPs
        myVaryAxesOptionsEdge = myVaryAxesOptions;
        myVaryAxesOptionsEdge.gaussianStd = 0.05;
        myVaryAxesOptionsEdge.varyMethod = 'gaussian';
        for addCounter = 1: length(edgeVPIDs)
                myVaryAxesOptionsEdge.newPerOld = round(expandedgeVPCohortSize/length(edgeVPIDs)); %maxNewPerOld;
                myVaryAxesOptionsEdge.baseVPIDs = edgeVPIDs(addCounter);
                jitteredWorksheet = addVariedVPs(jitteredWorksheet, myVaryAxesOptionsEdge);
        end

        % low subpopulation VPs
        myVaryAxesOptionsSubpop = myVaryAxesOptions;
        myVaryAxesOptionsSubpop.gaussianStd = 0.05;
        myVaryAxesOptionsSubpop.varyMethod = 'gaussian';
        for addCounter = 1: length(subpopVPIDs)
                myVaryAxesOptionsSubpop.newPerOld = round(expandlowsubpopVPCohortSize/length(subpopVPIDs)); %maxNewPerOld;
                myVaryAxesOptionsSubpop.baseVPIDs = subpopVPIDs(addCounter);
                jitteredWorksheet = addVariedVPs(jitteredWorksheet, myVaryAxesOptionsSubpop);
        end
        
            % Decide how many of the VPs from the seed expansion we can add.
        if maxNewPerIter < 0
            maxNewPerIterChecked = curEffN;
        else
            maxNewPerIterChecked = maxNewPerIter;
        end

        curVPIDs = getVPIDs(jitteredWorksheet);
        newIndices = (find(~ismember(curVPIDs,originalVPIDs)));
        newVPIDs = curVPIDs(newIndices);

        % We will also randomize a coefficient
        for vpCounter = 1 : length(newVPIDs);
            curVPID = newVPIDs{vpCounter};
            % randomize one of the new VP axis coefficients
            axisIndex = randsample([1:nAxis],1);
            vpIndex = find(ismember(curVPIDs,curVPID));
            jitteredWorksheet.axisProps.axisVP.coefficients(axisIndex,vpIndex) = rand(1);
        end

        mySimulateOptions = simulateOptions;
        mySimulateOptions.rerunExisting = false;
        mySimulateOptions.optimizeType = 'none';
        % Inherit the pool properties
        mySimulateOptions.poolRestart = myMapelOptions.poolRestart;
        mySimulateOptions.poolClose = myMapelOptions.poolClose;  

        s = rng;    
        % Also screen the worksheet if a function is provided
        if length(myScreenFunctionName) > 0
            jitteredWorksheet = eval([myScreenFunctionName,'(jitteredWorksheet,newVPIDs,mySimulateOptions)']);
        end
        rng(s);

        jitteredWorksheet = simulateWorksheet(jitteredWorksheet,mySimulateOptions);
        curVPIDs = getVPIDs(jitteredWorksheet);
        originalIndices = (find(ismember(curVPIDs,originalVPIDs)));
        newIndices = (find(~ismember(curVPIDs,originalVPIDs)));
        newVPIDs = curVPIDs(newIndices);     

        % Identify the VPs that don't fulfill the worksheet response
        % as well as those in the initial worksheet in order to filter
        % them.
        disp('---')
        disp(['Screening newly simulated VPs in ',mfilename,'.'])
        jitteredWorksheet = screenWorksheetVPs(jitteredWorksheet, myScreenTable, true, newVPIDs);
        disp('---------')

        myWorksheet = jitteredWorksheet;        
        curVPIDs = getVPIDs(jitteredWorksheet);
        newIndices = (find(~ismember(curVPIDs,originalVPIDs)));
        newVPIDs = curVPIDs(newIndices); 
        newPassNames = newVPIDs;
    else
        [edgeVPIDs,sortIndicesPick] = setdiff(edgeVPIDs,highVPIDs,'stable');
        edgeVPIndices = edgeVPIndices(sortIndicesPick);
        % Now combine considerations for heavily weighted VPs and
        % parents that look like they are useful.
        disp(['Adding ',num2str(length(highVPIDs)),' VPs as expansion seeds from prevalence weight considerations in ',mfilename,'.'])
        disp(['Adding ',num2str(length(edgeVPIDs)),' VPs as expansion seeds from phenotype range considerations in ',mfilename,'.'])
        highVPIDsMono = highVPIDs;
        highVPindicesMono = highVPIndices;
        nHighVPs = length(highVPIDsMono);
        nEdgeVPs = length(edgeVPIDs);
        highVPIDs = cell(1,nHighVPs + nEdgeVPs);
        highVPIndices = nan(1,nHighVPs + nEdgeVPs);
        nHighAdded=1;
        nEdgeAdded = 0;
        highVPIDs{1} = highVPIDsMono{1};
        highVPIndices(1) = highVPindicesMono(1);
        % Need an intelligent way of interlacing
        % the concerns of range coverage and
        % spreading out PWs.
        for addCounter = 2: length(highVPIDs)
            if ((nEdgeAdded < nHighAdded) && (nEdgeAdded < nEdgeVPs)) || (nHighAdded >= nHighVPs)
                nEdgeAdded = nEdgeAdded + 1;
                highVPIDs{addCounter} = edgeVPIDs{nEdgeAdded};
                highVPIndices(addCounter) = edgeVPIndices(nEdgeAdded);    
            else
                nHighAdded = nHighAdded + 1;
                highVPIDs{addCounter} = highVPIDsMono{nHighAdded};
                highVPIndices(addCounter) = highVPindicesMono(nHighAdded); 
            end
        end

        % Decide how many of the VPs from the seed expansion we can add.
        if maxNewPerIter < 0
            maxNewPerIterChecked = curEffN;
        else
            maxNewPerIterChecked = maxNewPerIter;
        end

        % We'll allow more than maxNewPerOld if it looks like we want
        % to allow many new VPs per seed.  We won't allow more
        % if we are selecting children based on parent and
        % we have many parents. If we are pooling all the children
        % together and picking the best, we will ignore this.
        maxNewPerOld = max(maxNewPerOld,ceil(maxNewPerIterChecked/length(highVPIndices)));
        % We allow 3x more child VP tries based on weight than edges
        simChildScale = (expandCohortSize/length(highVPIDs)*(nEdgeAdded+nHighAdded)/(1.5*nHighAdded+.5*nEdgeAdded));
        nChildSimulations = (ceil(1.5*nHighAdded*simChildScale)+ceil(.5*nEdgeAdded*simChildScale));

        myVaryAxesOptions = varyAxesOptions;
        myVaryAxesOptions.varyMethod = varyMethod;
        myVaryAxesOptions.gaussianStd = gaussianStd;
        myVaryAxesOptions.varyAxisIDs = getAxisDefIDs(myWorksheet);
        myVaryAxesOptions.intSeed = newVPop.intSeed;

        myVaryAxesOptions.additionalIDString = [suffix,num2str(wsIterCounter)];
        myVaryAxesOptions.baseVPIDs = highVPIDs;
        myVaryAxesOptions.newPerOld = ceil(expandCohortSize/length(highVPIDs));

        if selectByParent
            disp(['Note due to the maxNewPerIter and maxNewPerOld settings, it is possible only children from ',num2str(ceil(maxNewPerIterChecked/maxNewPerOld)),' VP parents will be selected in ',mfilename,'.'])    
        end
        disp('---')     

        % We will add more children based on weight than edges
        myVaryAxesOptions.newPerOld = ceil(simChildScale*1.5);
        myVaryAxesOptions.baseVPIDs = highVPIDs(1);
        jitteredWorksheet = addVariedVPs(myWorksheet, myVaryAxesOptions);
        % We enforce the Gaussian method with small 1% variance for edgeVPs
        myVaryAxesOptionsEdge = myVaryAxesOptions;
        myVaryAxesOptionsEdge.gaussianStd = 0.05;
        myVaryAxesOptionsEdge.varyMethod = 'gaussian';
        for addCounter = 2: length(highVPIDs)
            if sum(ismember(highVPIDs(addCounter),highVPIDsMono)) > 0
                myVaryAxesOptions.newPerOld = ceil(simChildScale*1.5);
                myVaryAxesOptions.baseVPIDs = highVPIDs(addCounter);
                jitteredWorksheet = addVariedVPs(jitteredWorksheet, myVaryAxesOptions);
            else
                myVaryAxesOptionsEdge.newPerOld = ceil(simChildScale*0.5);
                myVaryAxesOptionsEdge.baseVPIDs = highVPIDs(addCounter);
                jitteredWorksheet = addVariedVPs(jitteredWorksheet, myVaryAxesOptionsEdge);   
            end
        end

        curVPIDs = getVPIDs(jitteredWorksheet);
        newIndices = (find(~ismember(curVPIDs,originalVPIDs)));
        newVPIDs = curVPIDs(newIndices);

        % We will also randomize a coefficient
        for vpCounter = 1 : length(newVPIDs);
            curVPID = newVPIDs{vpCounter};
            % randomize one of the new VP axis coefficients
            axisIndex = randsample([1:nAxis],1);
            vpIndex = find(ismember(curVPIDs,curVPID));
            jitteredWorksheet.axisProps.axisVP.coefficients(axisIndex,vpIndex) = rand(1);
        end

        mySimulateOptions = simulateOptions;
        mySimulateOptions.rerunExisting = false;
        mySimulateOptions.optimizeType = 'none';
        % Inherit the pool properties
        mySimulateOptions.poolRestart = myMapelOptions.poolRestart;
        mySimulateOptions.poolClose = myMapelOptions.poolClose;    

         s = rng;    
        % Also screen the worksheet if a function is provided
        if length(myScreenFunctionName) > 0
            jitteredWorksheet = eval([myScreenFunctionName,'(jitteredWorksheet,newVPIDs,mySimulateOptions)']);
        end
        rng(s);

        jitteredWorksheet = simulateWorksheet(jitteredWorksheet,mySimulateOptions);
        curVPIDs = getVPIDs(jitteredWorksheet);
        originalIndices = (find(ismember(curVPIDs,originalVPIDs)));
        newIndices = (find(~ismember(curVPIDs,originalVPIDs)));
        newVPIDs = curVPIDs(newIndices);     

        % Identify the VPs that don't fulfill the worksheet response
        % as well as those in the initial worksheet in order to filter
        % them.
        disp('---')
        disp(['Screening newly simulated VPs in ',mfilename,'.'])
        jitteredWorksheet = screenWorksheetVPs(jitteredWorksheet, myScreenTable, true, newVPIDs);
        disp('---------')
        curVPIDs = getVPIDs(jitteredWorksheet);
        originalIndices = (find(ismember(curVPIDs,originalVPIDs)));
        newIndices = (find(~ismember(curVPIDs,originalVPIDs)));
        newVPIDs = curVPIDs(newIndices); 

        % We create a "dummy" vpop object to help score
        % the simulated VPs and see which look
        % like they will be more useful.
        if isa(newVPop,'VPop')
            testVPop = VPop;
        elseif isa(newVPop,'VPopRECIST')
            testVPop = VPopRECIST;
        elseif isa(newVPop,'VPopRECISTnoBin')
            testVPop = VPopRECISTnoBin;
        end
        testVPop.expData = newVPop.expData;
        testVPop.mnSDTable = newVPop.mnSDTable;
        testVPop.binTable = newVPop.binTable;
        testVPop.distTable = newVPop.distTable;
        testVPop.distTable2D = newVPop.distTable2D;   
        testVPop.corTable = newVPop.corTable;
        testVPop.subpopTable = newVPop.subpopTable;    
        if isa(newVPop,'VPopRECIST') || isa(newVPop,'VPopRECISTnoBin')
            testVPop.brTableRECIST = newVPop.brTableRECIST;
            testVPop.rTableRECIST = newVPop.rTableRECIST;        
            testVPop.relSLDvar = newVPop.relSLDvar;
            testVPop.absALDVar = newVPop.absALDVar;
            testVPop.crCutoff = newVPop.crCutoff;             
            testVPop.recistSimFilter = createRECISTSimFilter(jitteredWorksheet, testVPop, false);
        end
        if ~isa(newVPop,'VPopRECISTnoBin')
            testVPop = testVPop.assignIndices(jitteredWorksheet, myMapelOptions);
        end
        testVPop = testVPop.getSimData(jitteredWorksheet);
        testVPop.subpopTable = updateSubpopTableVPs(testVPop.subpopTable, jitteredWorksheet);
        testVPop = testVPop.addTableSimVals();  
        % For evaluation
        % coerce pws to be the same.
        testVPop.pws = ones(1,length(curVPIDs))./length(curVPIDs);
        testVPop = testVPop.addPredTableVals();    

        % Now get the score matrix for the new VPs
        % We can remove the LC scoring option for now
        % if isa(newVPop,'VPopRECIST')
        %     newVPScores = scoreWorksheetVPsLC(testVPop,originalIndices,newIndices);
        % else
            newVPScores = scoreWorksheetVPs(testVPop,originalIndices,newIndices);
        % end
        % Zero out scores if we are going to ignore them when selecting
        % which children to keep.
        if ~useScoresForNew
            newVPScores = newVPScores*0;
        end
        if unweightedParents
            % We can score all VPs but will focus on the edge VPs
            [nRows, ~] = size(myVPRangeTable);
            nVPs = length(curVPIDs);
            edgeVPScores = zeros(nRows,nVPs);
            allChildrenBaseIndices = zeros(1,length(curVPIDs));
            % Get the indices for the edge VP children
            for edgeVPCounter = 1 : length(edgeVPIDs)
                % We will apply different criteria for the VPs being
                % selected from the highly weighted ones and the ones being
                % selected as edgeVPs.
                parentID=edgeVPIDs(edgeVPCounter);
                childrenBase = strcat(parentID, ['_',suffix,num2str(wsIterCounter)]);
                curChildrenBaseIndices = cellfun(@isempty,strfind(curVPIDs,childrenBase));
                curChildrenBaseIndices=find(~curChildrenBaseIndices);
                curChildrenBaseIndices=intersect(curChildrenBaseIndices,newIndices); 
                allChildrenBaseIndices(curChildrenBaseIndices) = 1;
            end
            % We will step through each endpoint
            % Then get the simdata for the new VPs for the endpoint
            % then set the VPs that are non-edge children to zero
            % Then find the index of the minimum and set the corresponding
            % in the scores to 1.
            for rowCounter = 1 : nRows
                myElementID = myVPRangeTable{rowCounter, 'elementID'};
                myElementType = myVPRangeTable{rowCounter, 'elementType'};
                myInterventionID = myVPRangeTable{rowCounter, 'interventionID'};
                myTime = myVPRangeTable{rowCounter, 'time'};
                myElementIDCol = find(ismember(testVPop.simData.rowInfoNames,'elementID'));
                myElementTypeCol = find(ismember(testVPop.simData.rowInfoNames,'elementType'));
                myInterventionIDCol = find(ismember(testVPop.simData.rowInfoNames,'interventionID'));
                myTimeCol = find(ismember(testVPop.simData.rowInfoNames,'time'));
                simDataRow = find(ismember(testVPop.simData.rowInfo(:,myElementIDCol),myElementID) & ismember(testVPop.simData.rowInfo(:,myElementTypeCol),myElementType) & ismember(testVPop.simData.rowInfo(:,myInterventionIDCol),myInterventionID) & (cell2mat(testVPop.simData.rowInfo(:,myTimeCol)) == cell2mat(myTime)));
                curData = testVPop.simData.Data(simDataRow,:);
                if ((myVPRangeTable{rowCounter, 'minRangeMissing'} > myCutoff) && (myVPRangeTable{rowCounter,'min10PercentileMissing'}>0));
                    curVals = sort(curData(find(allChildrenBaseIndices)),'ascend');
                    curVals = unique(curVals,'stable');
                    % Any extreme edge is scored.
                    % Currently, number 2 is not scored though 
                    % this code is set up so we could add it back in.
                    if length(curVals) >= 1 
                        edgeVPScores(rowCounter,:) = 1 * (curData == curVals(1));
                        % 2nd place does not get a nonzero score
                        % + 0 * (curData == curVals(2));
                        % Enforce that the children also must expand the range
                        % to be scored
                        simMin = myVPRangeTable{rowCounter, 'simMin'};
                        edgeVPScores(rowCounter,:) = edgeVPScores(rowCounter,:) .* (curData < simMin);
                    end
                end
                if ((myVPRangeTable{rowCounter, 'maxRangeMissing'} > myCutoff) && (myVPRangeTable{rowCounter,'max90PercentileMissing'}>0))
                    curVals = sort(curData(find(allChildrenBaseIndices)),'descend');
                    curVals = unique(curVals,'stable');
                    if length(curVals) >= 1 
                        edgeVPScores(rowCounter,:) = 1 * (curData == curVals(1));
                        % + 0 * (curData == curVals(2));
                        % Enforce that the children also must expand the range
                        % to be scored
                        simMax = myVPRangeTable{rowCounter, 'simMax'};
                        edgeVPScores(rowCounter,:) = edgeVPScores(rowCounter,:) .* (curData > simMax);    
                    end
                end                
            end
            % We'll just use 1 row for the edge scores,
            % we won't separate the edges.
            edgeVPScores = sum(edgeVPScores(:,newIndices),1);

        end

        newPassNames = cell(1,0);
        if selectByParent
            % Select valid VPs from each higher weighted seed VP
            for highCounter = 1 : length(highVPIDs)
                % We will apply different criteria for the VPs being
                % selected from the highly weighted ones and the ones being
                % selected as edgeVPs.
                parentID=highVPIDs(highCounter);
                childrenBase = strcat(parentID, ['_',suffix,num2str(wsIterCounter)]);
                allChildrenBaseIndices = cellfun(@isempty,strfind(curVPIDs,childrenBase));
                allChildrenBaseIndices=find(~allChildrenBaseIndices);
                allChildrenBaseIndices=intersect(allChildrenBaseIndices,newIndices);
                childIDs = curVPIDs(allChildrenBaseIndices);
                childIDs = newVPIDs(find(ismember(newVPIDs,childIDs)));
                if sum(ismember(highVPIDsMono,parentID)) > 0
                    childScores = newVPScores(:,find(ismember(newVPIDs,childIDs)));
                else
                    % For VPs selected based on edges, apply a different
                    % selection strategy
                    % We'll only consider edge VPs with scores > 0 
                    childScores = edgeVPScores(:,find(ismember(newVPIDs,childIDs)));
                    keepChildren = find(childScores > 0);
                    childScores = childScores(keepChildren);
                    childIDs = childIDs(keepChildren);
                end
                % We will prioritize VPs that score well
                [nScoresPerVP, nCurChildren] = size(childScores);
                sortIndices = nan(nScoresPerVP, nCurChildren);
                for rowCounter = 1 : nScoresPerVP
                    nonZeroIndices = find(childScores(rowCounter,:)>0);
                    [curScores, curIndices] = sort(childScores(rowCounter,nonZeroIndices),'descend');
                    childScores(rowCounter,:) = nan(1,nCurChildren);
                    childScores(rowCounter,1:length(nonZeroIndices)) = curScores;
                    sortIndices(rowCounter,1:length(nonZeroIndices)) = nonZeroIndices(curIndices);
                end
                sortIndices=reshape(sortIndices,1,[]);
                sortIndices = sortIndices(find(~isnan(sortIndices)));
                sortIndices = unique(sortIndices,'stable');
                childIDs = childIDs(sortIndices);
                npass = length(childIDs);
                if (npass > 0)
                    newPassNames = [newPassNames,childIDs(1:min(npass,maxNewPerOld))];
                end
            end
            if maxNewPerIterChecked < 1
                newPassNames = cell(1,0);
            elseif length(newPassNames) > maxNewPerIterChecked
                newPassNames = newPassNames(1 : maxNewPerIterChecked);
            elseif length(newPassNames) < maxNewPerIterChecked
                addIndices = find(~ismember(newVPIDs,newPassNames));
                if length(addIndices) > 0
                    addVPIDs = newVPIDs(addIndices);
                    addScores = newVPScores(addIndices);
                    [addScores, addIndices] = sort(addScores, 'descend');
                    nToAdd = min(maxNewPerIterChecked-length(newPassNames),length(addScores));
                    addVPIDs = addVPIDs(addIndices(1:nToAdd));
                    newPassNames = [newPassNames, addVPIDs];
                end
            end		
        else
            % Otherwise, we pool the children from
            % all of our test VPs and just take the best
            % Note this does not yet support the separate edge scores.
            [nScoresPerVP, nCurChildren] = size(newVPScores);
            childScores = newVPScores;
            sortIndices = nan(nScoresPerVP, nCurChildren);
            for rowCounter = 1 : nScoresPerVP
                nonZeroIndices = find(childScores(rowCounter,:)>0);
                [curScores, curIndices] = sort(childScores(rowCounter,nonZeroIndices),'descend');
                childScores(rowCounter,:) = nan(1,nCurChildren);
                childScores(rowCounter,1:length(nonZeroIndices)) = curScores;
                sortIndices(rowCounter,1:length(nonZeroIndices)) = nonZeroIndices(curIndices);
            end
            sortIndices=reshape(sortIndices,1,[]);
            sortIndices = sortIndices(find(~isnan(sortIndices)));
            sortIndices = unique(sortIndices,'stable');        
            childIDs = newVPIDs(sortIndices);
            npass = length(childIDs);
            % We just allow up to maxNewPerIter
            % in the case where we are pooling children
            if (npass > 0)
                newPassNames = childIDs(1:min(npass,maxNewPerIter));
            end
        end

        % The repeats should already be screened, but just in case
        newPassNames = unique(newPassNames,'first');

        % Finalize the updated cohort
        mergeNames = [originalVPIDs,newPassNames];
        myWorksheet = copyWorksheet(jitteredWorksheet,mergeNames);
    end
    %         % Repopulate results.  Note that this is not necessary
    %         % and some time to each iteration, but help to avoid issues
    %         % if the original worksheet was carrying old, invalid results.
    %         % This is therefore removed, better to check the worksheet
    %         % VPs before calling expandVPopEffN.
    %         myWorksheet.results = {};
    %         myWorksheet = simulateWorksheet(myWorksheet);
else
    warning(['Unable to proceed in ',mfilename,'.  Returning input worksheet.'])
end
end
```
