## Supplementary File 3 for "A surrogate workflow for virtual population development: designing an amenable proxy for a hypothesis-test inspired nonlinear goodness-of-fit": extractLowPs.m

```
function [myVPopLowPs, lowPTables] = extractLowPs(myVPop, myWorksheet, thresh, outputFlag)
% Extract all low p value rows in input VPop tables
%
% ARGUMENTS
%  myVPop:    a VPop, VPopRECIST or VPopRECISTnoBin object
%  myWorksheet: a worksheet, with simulation results of the input VPop cohort
%  thresh:    Optional. the p value threshold 
%             Default is 0.01.
%  outputFlag: Boolean (true/false) indicating whether to write extracted information to screen
%              Default is true.
%  
% RETURNS
%  myVPopLowPs: a VPop or VPopRECIST object with low p value and nonzero weight part of the tables
%  lowPTables:  the table structure that contains all the extracted low p value information with the corresponding row indices and gof. for quick visual check.

flagContinue = true;
if nargin > 4
    warning([mfilename,' requires input arguments: myVPop, myWorksheet, optionally thresh, outputFlag.  Too many arguments provided.'])
    flagContinue = false;
elseif nargin < 2
    warning([mfilename,' requires input arguments: myVPop, myWorksheet, optionally thresh, outputFlag.  Too few arguments provided.'])
    flagContinue = false;
elseif nargin < 3
    flagContinue = true;
    thresh = 0.01;
    outputFlag = true;
elseif nargin < 4
    flagContinue = true;
    outputFlag = true;
end

if flagContinue
    if ~ismember(class(myVPop),{'VPop','VPopRECIST','VPopRECISTnoBin'})
        flagContinue = false;
        warning(['Input VPop not recognized in call to ',mfilename,'.  Requires: a VPop or VPopRECIST object and myWorksheet.'])
    end  
    % In-depth check of worksheet properties
    mySimulateOptions = simulateOptions;
    passCheck = mySimulateOptions.verify(myWorksheet);
    if ~passCheck
        warning(['Input Worksheet not recognized in call to ',mfilename,'.  Specified simulation options for ',mfilename,' not valid.'])
        flagContinue = false;
    end   
end


if flagContinue
    myMnSDTable = myVPop.mnSDTable;
    myBinTable = myVPop.binTable;
    myDistTable = myVPop.distTable;
    myDistTable2D = myVPop.distTable2D;
    myCorTable = myVPop.corTable;
    if isa(myVPop,'VPopRECIST') || isa(myVPop,'VPopRECISTnoBin')
        myBRTableRECIST = myVPop.brTableRECIST;
        myRTableRECIST = myVPop.rTableRECIST;
    end
    
    % initialize myVPopLowPs and lowPTables
    myVPopLowPs = myVPop;
    lowPTables.MnTable = [];
    lowPTables.SdTable = [];
    lowPTables.binTable = [];
    lowPTables.DistTable = [];
    lowPTables.Dist2DTable = [];
    lowPTables.corTable = [];
    lowPTables.corTable = [];
    if isa(myVPop,'VPopRECIST') || isa(myVPop,'VPopRECISTnoBin')
        lowPTables.brTableRECIST = [];
        lowPTables.rTableRECIST = [];
    end
    
    if ~isempty(myMnSDTable)
        curGOFMn = myVPop.gofMn<thresh;
        weighted = myVPop.mnSDTable{:,'weightMean'}>0;
        rowIndices = find(curGOFMn & weighted);
        gofMn = myVPop.gofMn(rowIndices);
        gofMn = table(gofMn);
        myVPopLowPs.mnSDTable = myVPop.mnSDTable(rowIndices,:);
        rowIndices = table(rowIndices);
        lowPTables.MnTable = [rowIndices, gofMn, myVPop.mnSDTable(curGOFMn & weighted,{'subpopNo','time','interventionID','elementID','expDataID','expTimeVarID','expVarID'})];
        
        curGOFSD = myVPop.gofSD<thresh;
        weighted = myVPop.mnSDTable{:,'weightSD'}>0;
        rowSdIndices = find(curGOFSD & weighted);
        rowIndices = setdiff(rowSdIndices, rowIndices{:,1}); % remove the Mnrows that has been extracted above
        gofSD = myVPop.gofSD(rowSdIndices);
        gofSD = table(gofSD);
        myVPopLowPs.mnSDTable = [myVPopLowPs.mnSDTable;myVPop.mnSDTable(rowIndices,:)]; % remove the Mnrows that has been extracted above 
        rowSdIndices = table(rowSdIndices);
        lowPTables.SdTable = [rowSdIndices, gofSD, myVPop.mnSDTable(rowSdIndices{:,1},{'subpopNo','time','interventionID','elementID','expDataID','expTimeVarID','expVarID'})]; % show all low sd pvalue rows, not removing the ones that been extracted to mnTable
    end
    
    if ~isempty(myBinTable)
        curGOFBIN = myVPop.gofBin<thresh;
        weighted = myVPop.binTable{:,'weight'}>0;
        rowIndices = find(curGOFBIN & weighted);
        gofBin = myVPop.gofBin(rowIndices);
        gofBin = table(gofBin);
        myVPopLowPs.binTable = myVPop.binTable(rowIndices,:);
        rowIndices = table(rowIndices);
        lowPTables.binTable = [rowIndices, gofBin, myVPop.binTable(curGOFBIN & weighted,{'subpopNo','time','interventionID','elementID','expDataID','expTimeVarID','expVarID'})];
    end
    
    if ~isempty(myDistTable)
        curGOFDIST = myVPop.gofDist<thresh;
        weighted = myVPop.distTable{:,'weight'}>0;
        rowIndices = find(curGOFDIST & weighted);
        gofDist = myVPop.gofDist(rowIndices);
        gofDist = table(gofDist);
        myVPopLowPs.distTable = myVPop.distTable(rowIndices,:);
        rowIndices = table(rowIndices);
        lowPTables.DistTable = [rowIndices, gofDist, myVPop.distTable(curGOFDIST & weighted,{'subpopNo','time','interventionID','elementID','expDataID','expTimeVarID','expVarID'})];
    end
    
    if ~isempty(myDistTable2D)
        curGOFDIST2D = myVPop.gofDist2D < thresh;
        weighted = myVPop.distTable2D{:,'weight'}>0;
        rowIndices = find(curGOFDIST2D & weighted);
        gofDist2D = myVPop.gofDist2D(rowIndices);
        gofDist2D = table(gofDist2D);
        myVPopLowPs.distTable2D = myVPop.distTable2D(rowIndices,:);
        rowIndices = table(rowIndices);
        lowPTables.Dist2DTable = [rowIndices, gofDist2D, myVPop.distTable2D(curGOFDIST2D & weighted,{'subpopNo','interventionID1','time1','elementID1','time2','elementID2','expDataID1','expVarID1', ...
            'expDataID2','expVarID2'})];
    end
    
    if ~isempty(myCorTable)
        curGOFCOR = myVPop.gofCor < thresh;
        weighted = myVPop.corTable{:,'weight'}>0;
        rowIndices = find(curGOFCOR & weighted);
        gofCor = myVPop.gofCor(rowIndices);
        gofCor = table(gofCor);
        myVPopLowPs.corTable = myVPop.corTable(rowIndices,:);
        rowIndices = table(rowIndices);
        lowPTables.corTable = [rowIndices, gofCor, myVPop.corTable(curGOFCOR & weighted,{'subpopNo','interventionID1','time1','elementID1','time2','elementID2','expDataID1','expVarID1', ...
            'expDataID2','expVarID2'})];
    end
    
    if isa(myVPop,'VPopRECIST')
        if ~isempty(myBRTableRECIST)
            curGOFBR = myVPop.gofBR < thresh;
            weighted = myVPop.brTableRECIST{:,'weight'}>0;
            rowIndices = find(curGOFBR & weighted);
            gofBR = myVPop.gofBR(rowIndices);
            gofBR = table(gofBR);
            myVPopLowPs.brTableRECIST = myVPop.brTableRECIST(rowIndices,:);
            rowIndices = table(rowIndices);
            lowPTables.brTableRECIST = [rowIndices, gofBR, myVPop.brTableRECIST(curGOFBR & weighted,{'subpopNo','time','interventionID','elementID','expDataID','expTimeVarID','expVarID'})];
        end
        
        if ~isempty(myRTableRECIST)
            curGOFR = myVPop.gofR < thresh;
            weighted = myVPop.rTableRECIST{:,'weight'}>0;
            rowIndices = find(curGOFR & weighted);
            gofR = myVPop.gofR(rowIndices);
            gofR = table(gofR);
            myVPopLowPs.rTableRECIST = myVPop.rTableRECIST(rowIndices,:);
            rowIndices = table(rowIndices);
            lowPTables.rTableRECIST = [rowIndices, gofR, myVPop.rTableRECIST(curGOFR & weighted,{'subpopNo','time','interventionID','elementID','expDataID','expTimeVarID','expVarID'})];
        end
    end
    
    myVPopLowPs = getSimData(myVPopLowPs, myWorksheet); % need to update simData first
    myVPopLowPs = evaluateGOF(myVPopLowPs);
    if outputFlag
        %% output to screen lowPTables
        disp('------- output all low p value rows ------');
        if (~isempty(lowPTables.MnTable))
            disp('lowPTables.MnTable = ');
            lowPTables.MnTable
        end
        if (~isempty(lowPTables.SdTable))
            disp('lowPTables.SdTable = ');
            lowPTables.SdTable
        end
        if (~isempty(lowPTables.binTable))
            disp('lowPTables.binTable = ');
            lowPTables.binTable
        end
        if (~isempty(lowPTables.DistTable))
            disp('lowPTables.DistTable = ');
            lowPTables.DistTable
        end
        if (~isempty(lowPTables.Dist2DTable))
            disp('lowPTables.Dist2DTable = ');
            lowPTables.Dist2DTable
        end
        if (~isempty(lowPTables.corTable))
            disp('lowPTables.corTable = ');
            lowPTables.corTable
        end
        if (isa(myVPop,'VPopRECIST') || isa(myVPop,'VPopRECISTnoBin')) && (~isempty(lowPTables.brTableRECIST))
            disp('lowPTables.brTableRECIST = ');
            lowPTables.brTableRECIST
        end
        if (isa(myVPop,'VPopRECIST') || isa(myVPop,'VPopRECISTnoBin')) && (~isempty(lowPTables.rTableRECIST))
            disp('lowPTables.rTableRECIST = ');
            lowPTables.rTableRECIST
        end
    end
else
    warning(['Unable to run ',mfilename,'.  Returning input.'])
end
end
      
      
      
      
      
      
```
